# Avoidance of pyroptosis accounts for the relatively high metastatic potential observed in early hybrid EMT states

**DOI:** 10.1101/2025.01.26.634904

**Authors:** Aakanksha Verma, Alessandro Genna, Yaron Vinik, Nishanth Belugali Nataraj, Maha Abedrabbo, Boobash Raj Selvadurai, Tithi Bhandari, Noa Aharoni, Palaniappan Ramesh, Boris Boguslavski, Diana Drago, Tomer-Meir Salame, Amir Prior, Yishai Levin, Eviatar Weizman, Rong Zhu, Carlos Caldas, Oscar M. Rueda, Sima Lev, Yosef Yarden

## Abstract

EMT converts epithelial (E) phenotypes to invasive mesenchymal (M) states. However, analyses of circulating tumor cells (CTCs) indicated that biphenotypic (E+M) CTCs better correlate with metastasis. Similarly, investigations of murine tumors undergoing EMT concluded that early E+M states posses the highest metastatic potential. To explore this, we selected in animals with breast cancer CTCs having progressively increasing intravasation abilities. This revealed that downregulation of arrestin Arrdc4 associates with CTC aggressiveness. In xenografts, depleting Arrdc4 accelerated tumor progression, whereas overexpression hindered progression in immunocompetent, but not in immunocompromised mice. Mechanistically, high Arrdc44 suppresses glucose uptake and enhances gasdermin E, triggering pyroptosis a type of pro-inflammatory cell death. Consistently, Arrdc4’s lowest levels characterize the most metastatic biphenotypic states. In patients, both epigenetic and chromosomal aberrations downregulate ARRDC4 and predict poor prognosis. In summary, the uncovered mechanism portrays pyroptosis of biphenotypic EMT cells as a rheostat of CTCs, which may resolve the controversy on the role played by EMT in metastasis.

## Introduction

Breast cancer (BC) is the leading cause of cancer-related deaths among women, with metastasis representing the most lethal phase of the disease progression ^1,2^. The process of metastasis follows a well-defined sequence, including tumor growth, intravasation into lymphatic and blood vessels, survival of cancer cells in the bloodstream and colonization of secondary sites ^3^. Prior research has shown that the presence of BC cells in the bloodstream correlates with lung metastasis, highlighting blood and lymphatic entry as critical steps in the metastatic process ^4^. Unlike non-metastatic cells, which tend to fragment when interacting with blood vessels, metastatic cells not only survive but also exhibit a stronger orientation towards the vessels. This behavior may be linked to the activation of specific gene expression programs, such as the WNT pathway, which inhibits anoikis, increases fibronectin levels, and enhances metastatic potential in pancreatic cancer models ^5^. Intravital imaging revealed that BC cells undergo re-programming during intravasation and that cancer stem cells are enriched in the circulation and at the metastatic site ^6^. Similarly, research on brain tumors suggested that stemness-related gene programs may aid in the survival of metastatic cells ^7^. For example, circulating tumor cells (CTCs) in glioblastoma patients exhibit a cancer stem cell-like phenotype with activation of SOX2, OCT4 and NANOG, along with WNT. In addition, the embryonic epithelial-to-mesenchymal transition (EMT) program can improve the metastatic capabilities of CTCs ^8^. However, rather than a complete EMT, an incomplete or “partial” EMT, characterized by the co-expression of epithelial (E) and mesenchymal (M) markers, may better support phenotypic plasticity in metastasis ^9,10^. Importantly, biphenotypic CTCs were detected in the blood of patients with lung, prostate, liver and breast cancer, and coexpression of E+M markers, rather than fully M or fully E phenotypes, have been assosciated with poor prognosis of patients with several cancer types, including BC ^11–13^.

Understanding the factors permitting circulating biphenotypic cells, and other CTCs, to survive in bloodstream and target parenchyma is critical. In general, patient’ CTCs have very short half-life, ranging from 1 to 2.4 hours ^14^. This rapid decline is attributable to oxidative stress, immune effector cells and physical parameters ^15^. Thus, although CTCs are infrequent in the blood of patients, they are frequently found in clusters having up to a 50-fold increase in metastatic potential ^16^. Clusters physically stabilized by surface adhesion molecules like E-cadherin and EpCAM may permit collective invasion. Indeed, it has been reported that E-cadherin-and vimentin-positive tumors have the worst prognosis among all BC cases ^17^. Further, these proteins are colocalized within the same tumor cells, suggesting the existence of an aggressive subpopulation in primary tumors. In addition, the existence of tumors with E-cadherin^high^/vimentin-positive expression might permit polarization into leader cells that guide follower cells at the cluster’s rear. A screen of proteins expressed on the cell surface of BC and SCC (squamous cell carcinoma) proposed that the molecular switch to the mesenchymal state initially comprises loss of EpCAM expression, which coincides emergence of vimentin expression ^18^. However, some EpCAM-negative subpopulations might continue coexpressing keratin 14 (K14), an epithelial marker and vimentin, a mesenchymal marker. These cells are likely the biphenotypic (or hybrid) tumor cells. Yet other populations that completely K14 expression correspond to full EMT tumor cells. It is interesting noting that internalization of adhesion molecules, rather than transcriptional regulation, might similarly control EMT in pancreatic cancer ^19^. It is also important noting that the early E+M subpopulation are relatively primed towards a hybrid phenotype, unlike the M state that cannot revert spontaneously to more epithelial phenotypes^18,20^.

Both the tumor microenvironment, such as inflammation and immune cells, and intrinsic factors within cancer cells, such as metabolic reprogramming, play crucial roles in EMT and the dissemination of tumor cells. For instance, transketolase levels were found to be higher in lymph node metastases compared to primary BC and inclusion of alpha-ketoglutarate increased levels of two tumor suppressor metabolites, succinate dehydrogenase and fumarate hydratase, leading to a reduction in two oncometabolites: succinate and fumarate ^21^. The role of immune surveillance in metastasis has been illustrated by case reports involving kidney transplants, where immunosuppressed recipients developed metastases from micrometastases that were previously present in the kidneys of melanoma patients who had served as organ donors ^22,23^. Notably, some transplanted kidneys were obtained from patients many years after the melanoma had been surgically removed. Similarly, relapses of dormant BC metastases can occur up to 15 years after the removal of the primary tumor ^24^, with cytotoxic immune cells potentially playing a role in BC dormancy ^25,26^. Employing a murine immunocompetent model, we uncovered two novel characteristics of biphenotypic states that may resolve the controversy on the role played by EMT in metastasis: (i) by downregulating a late EMT marker, E+M cells gain protection from a lytic cell death process, pyroptosis, triggered by the activation of inflammatory caspases, and (ii) by preventing endocytosis of glucose transporters, E+M cells sustain glucose uptake, which fuels their metabolism and proliferation.

## Results

### A genome-wide in vivo screen identifies Arrdc4 as a gene that is downregulated in murine CTCs, as well as in patients’ BC tissues

To identify genes regulating intravasation and survival of CTCs in the bloodstream, we established an in vivo screen, which is outlined in Figure 1A. Briefly, the screen made use of a GFP (green fluorescence protein) labeled derivative of the murine 4T1 cells, derived from a highly invasive line that can spontaneously metastasize from mammary gland tumors to multiple distant sites. The labeled cells were injected in the 4th mammary fat pad of 6 weeks old immunocompetent female Balb/c mice. Using cardiac puncture, we harvested CTCs five days after inoculation. Ficoll gradients and cytometry were employed to isolate individual GFP-positive CTCs (denoted: fresh G1-CTCs) directly into 96-well plates containing lysis buffer, for single cell RNA sequencing (scRNA-seq). Another fraction of harvested CTCs was cultured in vitro (G1’-CTCs) for 7-10 days, prior to reinjection of cells (pooled from 5 mice) in the mammary fat pad of naïve female Balb/c mice. This process was repeated to obtain G2, G3, G4 and G5 cells (‘fresh CTCs’), as well as G2’, G3’ and G4’ cells (‘cultured CTCs’). To examine the possibility that CTC-derived xenografts (CDXs) progressively gained increased aggressiveness, we determined the survival of mice bearing CTXs and noted that mice implanted with more advanced generations of CTCs showed progressively shorter survival (Fig. 1B; p<0.001). In a similar way, when cultured CTCs (2×10^4^ cells) were injected into the tail vein of naïve mice, the more advanced cells gave rise to significantly more lung metastases and the emerged metastases were relatively large (Figs. S1A and S1B). Taken together, these observations indicated that repeated cycles of CTC harvest-reinjection selected CTCs that were endowed with increasing competence to intravasate and form distant metastases.

**Figure 1:**
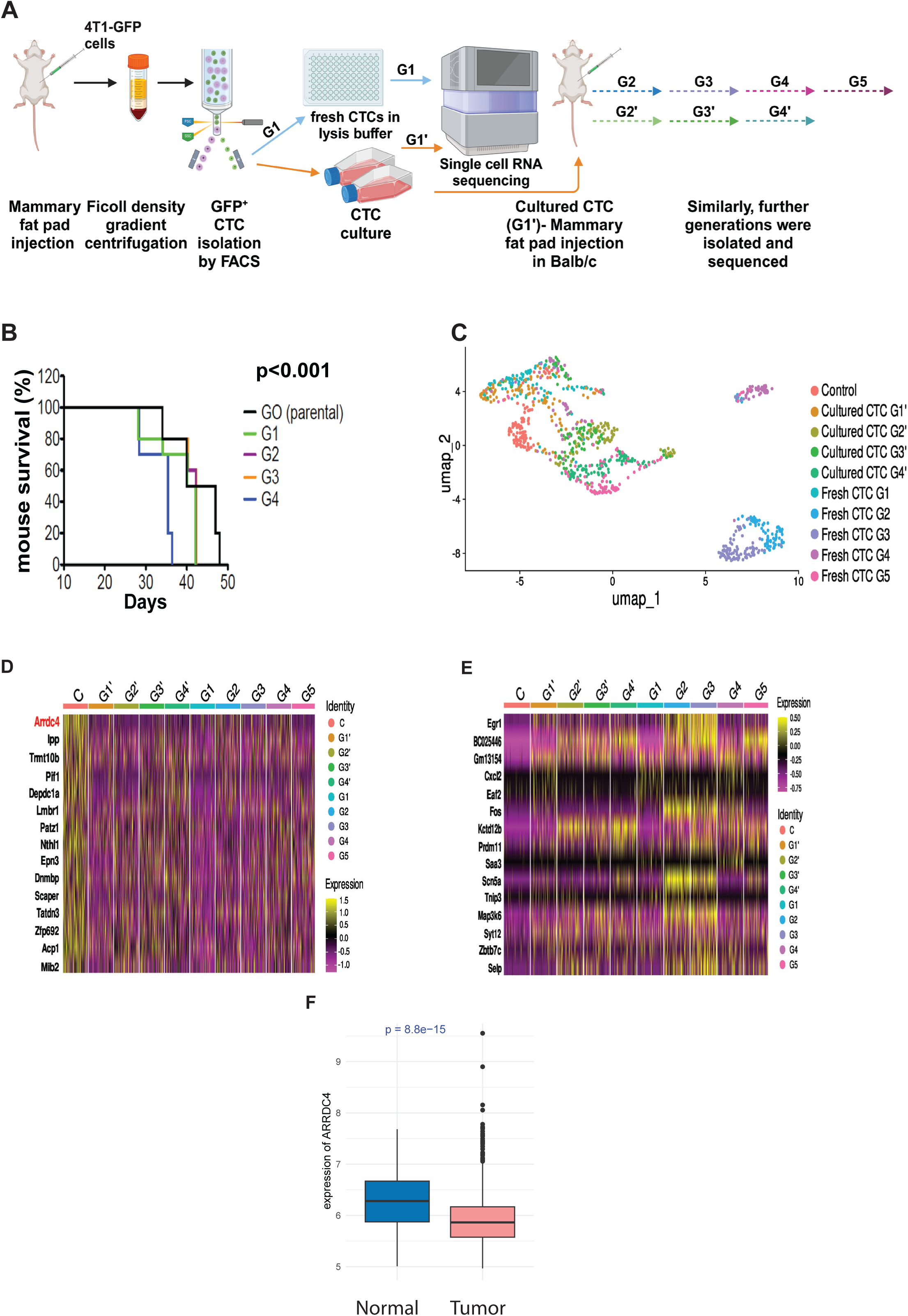
In vivo screens and single cell RNA sequencing uncover genes differentially expressed in murine CTCs. (**A**) Schematic representation of the experimental steps used for the isolation, sub-culturing, and implantation of CTCs in mice to generate CTC-derived xenografts (CTX). The scheme was prepared using BioRender.com. (**B**) Kaplan-Meier survival curves corresponding to groups of mice (n=6) that were implanted with the indicated generations of CTCs. Note that G0 corresponds to the parental (control) generation. The p-value relates to the difference between G4 and G0 (using the log-rank test). (**C**) Uniform Manifold Approximation and Projection (UMAP) visualization of all single cells, which were colored according to the respective populations of fresh CTCs (G1 through G5) and cultured CTCs (G1’ through G4’). (**D** and **E**) Heatmaps of single cell expression profiles of the top 15 differentially expressed genes in the control (parental) 4T1 cells versus the indicated CTC populations. Shown are the genes downregulated in CTCs (D) and the genes upregulated in CTCs. (**F**) Box plots comparing the expression of ARRDC4 in 1980 breast tumour samples and in 144 normal samples (METABRIC dataset; Wilcoxon rank-sum test).

A total of 874 single cell sequencing libraries were obtained from control, fresh CTCs and cultured CTCs. Their analysis made use of Seurat, an R package designed for exploration of scRNA-seq data. Uniform Manifold Approximation and Projection (UMAP) was used for non-linear dimension reduction. Figure 1C presents the different cell populations, indicating that more closely related populations of CTCs clustered together. For differential expression (DE) analysis we utilized edgeR and glmTreat ^27^, which selected genes displaying consistent trends in CTCs versus control cells. Next, we generated heatmaps of single cell expression of the top 15 genes upregulated in the control parental cells versus CTCs (Fig 1D) and, similarly, the top 15 genes downregulated in CTCs (Fig. 1E). The respective uppermost genes, EGR1 and ARRDC4, were consistently up-and downregulated in all CTC generations. Because previous studies established clearly positive effects of EGR1 in metastasis ^28,29^ but no previous study reported involvement of the alpha-arrestin called Arrdc4 in tumor progression, we focused on this gene. In line with clinical relevance, analysis of METABRIC, a database comprising information from ∼2000 BC samples and 140 normal breast tissue ^30–32^ revealed that ARRDC4 transcripts were significantly downregulated in the tumors relative to the surrounding normal mammary gland tissues (p=8.8e-15; Fig. 1F).

### ARRDC4 is downregulated in aggressive types of BC and it bears prognostic significance

Encouraged by the clinical input showing that downregulation of ARRDC4 is medically meaningful, we obtained an RNA expression plot (Fig. 2A) and immunoblots (Fig. 2B) demonstrating higher and more variable abundance of transcripts and proteins corresponding to Arrdc4 in the parental 4T1 cells, relative to the various CTC generations. These observations prompted analysis of the potential prognostic value of ARRDC4 in human BC. To this end, we utilized the longitudinal information available from METABRIC, which followed BC patients for >20 years since initial diagnosis. As illustrated by the Kaplan-Meier survival plots shown in Figure 2C, ARRDC4’s transcripts categorized into high and low expression levels correlated low ARRDC4 levels with poorer patient prognosis (p=0.00019 for disease free survival, DFS, and p<0.0001 for overall survival, OS). In line with this conclusion, low ARRDC4 statistically associated with high grade BC (Fig. S2A) and higher stage tumors, especially stages III and IV (Fig. 2D), which translates to faster-growing tumors that either already metastasized or they are likely to spread. Consistent with the aggressive features of ARRDC4-low BC, the respective tumors frequently belong to the triple negative BC group (Fig. 2E), which typically has adverse prognosis. In summary, downregulation of Arrdc4 is not limited to the murine tumor model—it extends to patients harboring relatively aggressive forms of BC

**Figure 2:**
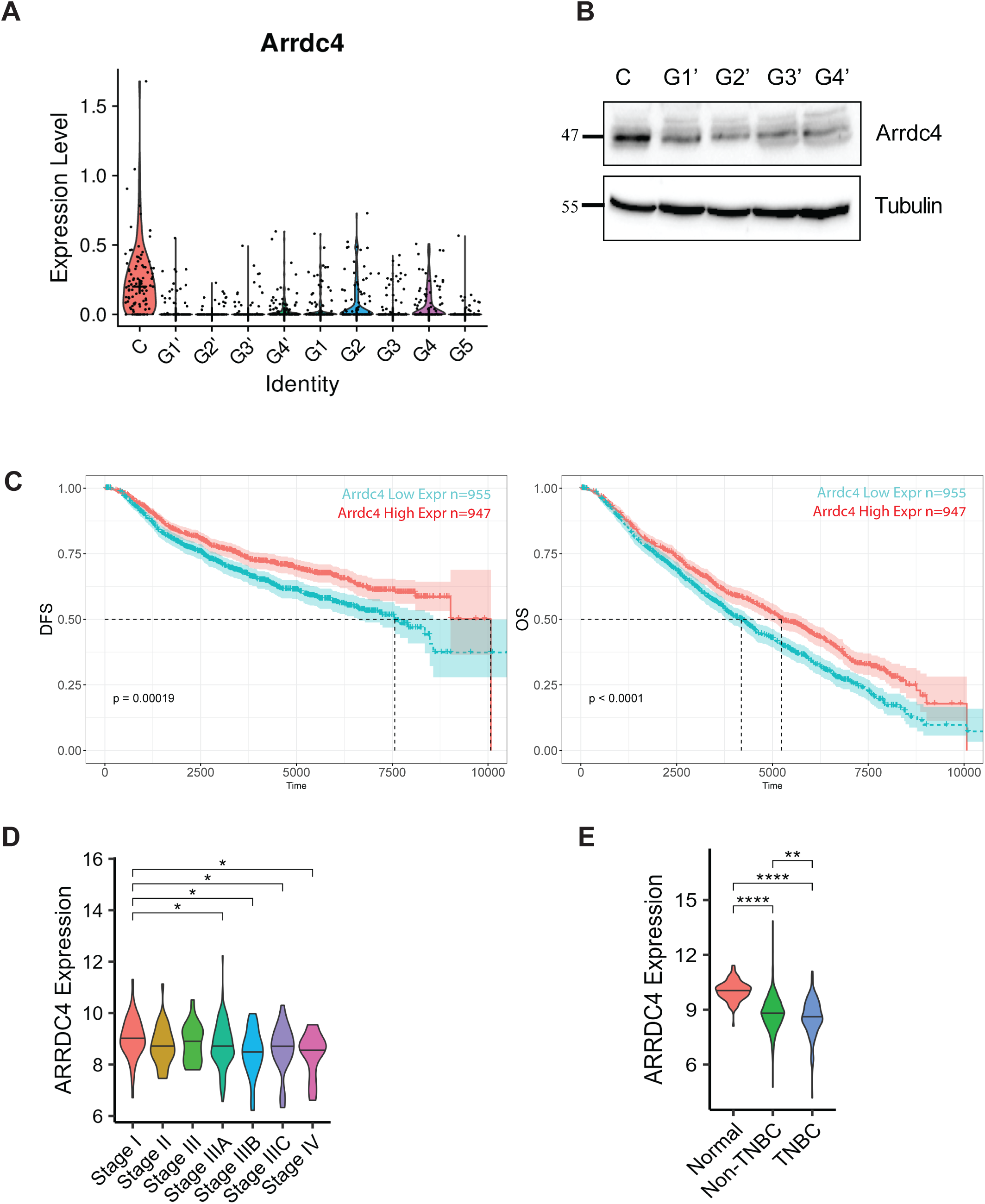
Arrdc4 is downregulated in murine CTCs and predicts shorter survival of patients with aggressive breast tumors. (**A**) The plot presents the relative expression of *Arrdc4* in parental 4T1 cells (control; C), fresh and cultured CTCs. Note that each dot represents RNA sequencing data from a single CTC. (**B**) Immunoblot showing Arrdc4 protein expression levels in the parental 4T1 cells and in the indicated generations of cultured CTCs. Tubulin was used as a gel loading control. (**C**) The full cohort of patients included in the METABRIC dataset was divided by their median into two nearly equal groups according to their ARRDC4’s mRNA expression abundance: low (blue) and high (red) expression. Breast cancer-related deaths, both disease-free survival (DFS; left) and overall survival (OS) are shown in the respective Kaplan-Meier curves. (**D**) Classification of breast cancer patients based on tumor stage and correlation with ARRDC4 expression levels (TCGA dataset; *, p<0.05). **(E)** Box plot showing classification of normal breast, triple negative BC (TNBC) and non-TNBC patients based on ARRDC4 expression (TCGA dataset: normal, n=114; non-TNBC, n= 879 and TNBC, n= 180 patients). The level of significance is shown as follows: **, p<0.01 and ****, p<0.0001.

### Both chromosomal aberrations and DNA methylation downregulate ARRDC4 in patients with breast cancer

While the functional in vivo screen, as well as the clinical data, supported the possibility that relatively low levels of ARRDC4 favor dissemination and survival of CTCs, the mechanisms underlying downregulation of ARRDC4 in patients remained unknown. Assuming that DNA methylation might contribute to ARRDC4 downregulation, we surveyed gene methylation in the TCGA dataset. This analysis concluded that *ARRDC4* methylation is associated with the absence of both the estrogen and the progesteron receptors (Fig. S2B; left panels), well characterized markers of luminal disease. Hence, we next analyzed the 10 integrated clusters (ICs) of BC, whose classification combines information on the genomic and transcriptomic data of tumors ^31^. This analysis uncovered an association between high *ARRDC4* methylation and the relatively aggressive, hormone receptor negative clusters IC5, IC9 and IC10 (Fig. S2B, right panel). These observations raised the possibility that methylation of *ARRDC4* might bear prognostic value. To examine this, we performed hazard ratio (HR) analysis that evaluated the risk of disease recurrence or patient death. For DFS, the analyzed dataset included 635 patients with no disease recurrence and 87 with disease recurrence (or death). Likewise, for OS (overall survival), the dataset included 678 survivors and 105 deaths. The results shown in Figure S2C imply that in both univariate and multivariate analyses the methylation of *ARRDC4* can serve as a poor prognosis factor.

Because large scale DNA sequencing and comparative genomic hybridization analyses revealed that BC is driven largely by genomic copy number aberrations (CNAs), rather than by point mutations or indels ^33^, we identified the TCN (total copy number) of *ARRDC4* in tumors and categorized it as either gain, neutral or loss, based on whether TCN was greater than, equal to, or less than the ploidy. Additionally, we considered ARRDC4’s mCN (minor copy number) to categorize it as loss of heterozygosity (LOH) or not LOH. The results of an initial survey of the TCGA dataset returned less significant output compared to those from METABRIC, likely due to the typically shorter TCGA’s patient follow-up time. Concentrating on METABRIC, the Kaplan-Meier plots for CNA and mCN data of ARRDC4 (Fig. S2D) indicated that the loss of the *ARRDC4* gene was a statistically significant poor prognostic factor. In summary, both epigenetic mechanisms and chromosomal aberrations permit BC to downregulate expression of ARRDC4, thereby gain relatively aggressive phenotypes resulting in shorter patient survival time, in line with the characteristics of the CTCs we analyzed in the murine model.

### Downregulation of murine Arrdc4 increases cell proliferation and migration in vitro, as well as enhances tumorigenesis and metastasis in animals

In light of the results of the in vivo screen and the surveys of clinical data, we predicted that reducing Arrdc4 expression would accelerate mammary tumor progression. Using a lentiviral knockdown approach, we generated three 4T1 clones exhibiting stable downregulation of Arrdc4. Each clone simultaneously expressed GFP and a small Arrdc4-specific small hairpin RNA (shRNA). A fourth clone expressing a scrambled shRNA (Scr) served as control. Stable downregulation was confirmed at the level of Arrdc4’s mRNA (Figs. 3A) and protein (Fig. 3B). Cell proliferation, as determined by means of the conversion of MTT (3-(4,5-dimethylthiazol-2-yl)-2,5-diphenyltetrazolium bromide) to insoluble formazan was reproducibly increased upon downregulation of Arrdc4 (Fig. S3A). Additionally, the abilities to form colonies (Figs. S3B and S3C) and migrate across a permeable membrane (in a modified Boyden chamber assay; Figs. S3D and S3E) were significantly enhanced in Arrdc4-depleted cells. Cell cycle analysis revealed yet another effect of Arrdc4 downregulation: knockdown of the transcript moderately increased the fractions of cells in the S (DNA synthesis) phase (Fig. 3C) and, as expected, no detectable apoptosis accompanied Arrdc4 downregulation (Fig. 3D).

**Figure 3:**
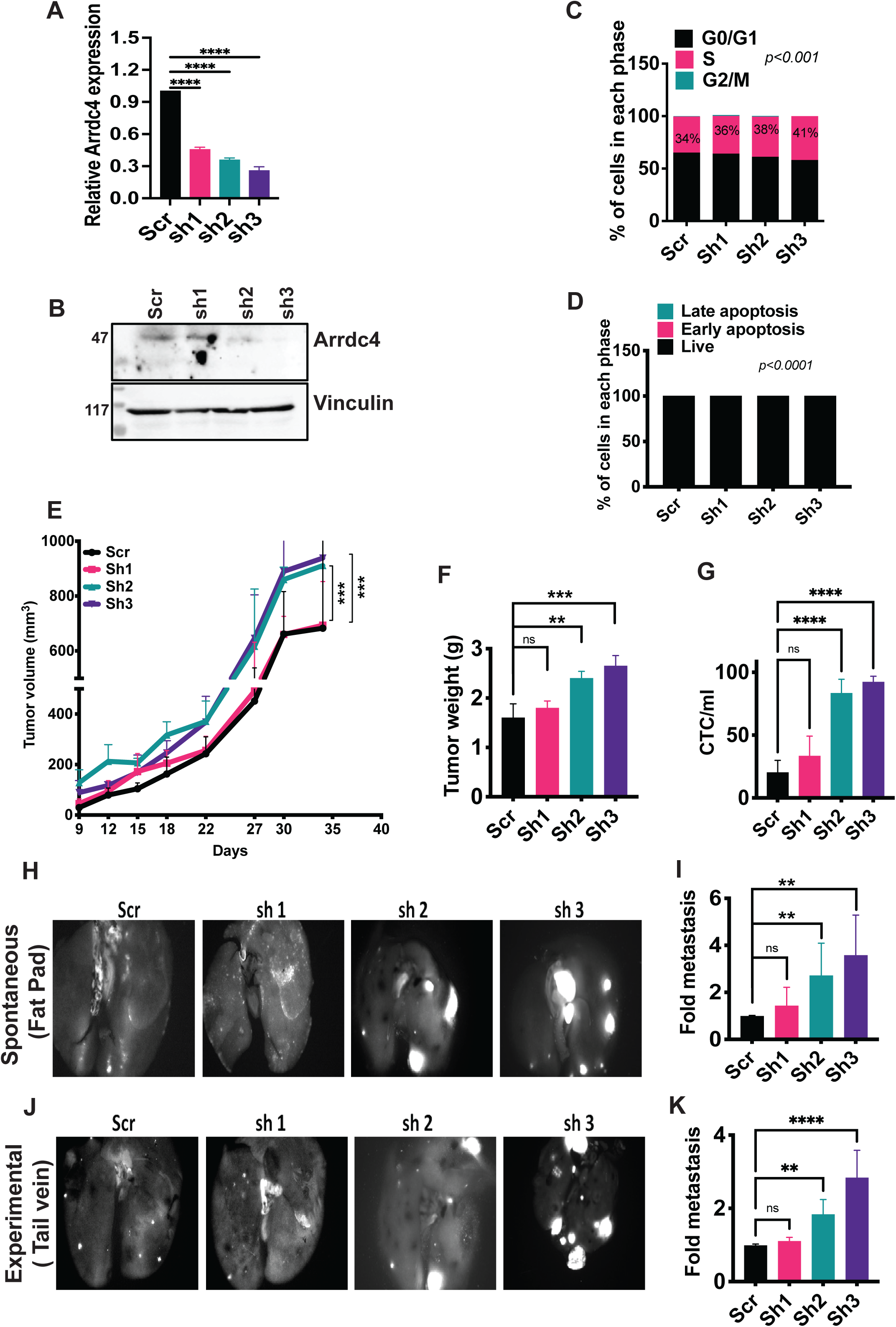
***Arrdc4* knockdown accelerates tumor growth and metastasis in immunocompetent animals.** (**A** and **B**) Three different shRNAs (sh1, sh2 and sh3), along with a control shRNA (scrambled; Scr), were separately used to transfect 4T1 cells. qRT-PCR (n=4; A) and immunoblotting (B) were used to quantify the respective levels. ****, p<0.0001. (**C** and **D**) The bromodeoxyuridine (BrdU) incorporation assay (C) and the annexin V staining assay (D) were respectively used to determine the cell cycle phase and the phase of apoptosis of the stable 4T1 clones from A. (**E**-**G**) Results showing the average tumor volume (±SD; panel E), tumor weight (F) and the number of CTCs (G) observed in 6 weeks old female Balb/c mice (n=6) that were injected in the mammary fad pads with 4T1 cells (1×10^6^). Either Scr or the indicated Arrcd4’s shRNA clones were examined. Note that CTCs were isolated from freshly collected blood using Ficoll-density gradient-based centrifugation and quantified using cytometry. (**H** and **I**) Representative images showing spontaneous lung metastases in mice that were injected in their mammary fat pads with 4T1 cells, either Scr or the indicated shArrdc4-expressing cells. The bar graph shows quantification of lung metastatic nodules. (**J** and **K**) Representative images obtained as in H, except that cells were injected in the tail veins of mice (experimental metastasis assay). The bar graph quantifies the corresponding lung metastases. The levels of significance in all panels are shown (ns, non-significant; **, p<0.01; ***, p<0.001 and ****, p<0.0001).

To evaluate the effects of Arrdc4 downregulation on tumorigenesis, female Balb/c mice were implanted with 4T1 cells (1×10^6^ per animal) that stably expressed shArrdc4 (sh1, sh2, or sh3) or a control shRNA (Scr). Tumor growth was notably accelerated in mice injected with sh2 and sh3 cells, as evidenced by larger tumor volumes and weights relative to the Scr control (Figs. 3E and 3F). Additionally, quantification of CTCs showed higher numbers in mice with sh2-and sh3-expressing tumors compared to those carrying Scr tumors (Fig. 3G). Notably, the CTC counts increased beyond the gains in tumor volume or weight. Consistent with these findings, spontaneous metastasis (i.e., spread of 4T1 cells from fat pads to lungs) was higher in mice injected with GFP-labeled sh2 or sh3 cells (1×10^6^ per animal; Figs. 3H and 3I). Similarly, shArrdc4 increased experimental metastasis (from vein to lungs) after injecting 2×10^4^ cells into the tail vein of mice (Figs. 3J and 3K). These results indicated that Arrdc4 downregulation enhances tumor growth and metastasis. Conceivably, this phenotype has been selected during the repeated CTC enrichment cycles we performed in animals, and it might explain why patients with tumors highly expressing ARRDC4 exhibit longer survival.

### Arrdc4 overexpression induces growth arrest, likely due to endocytosis of glucose transporters

We next investigated whether forced overexpression of Arrdc4 would reveal new functional properties, beyond those identified by using RNA interference. To explore this, we used lentiviral transduction and generated 4T1 clones stably co-expressing GFP and murine Arrdc4 (clones OX1, OX2 and OX3). An empty vector (EV) that drove only GFP expression served as control. We confirmed stable Arrdc4 overexpression at both transcript and protein levels (Figs. 4A and 4B). As anticipated, there was a significant decrease in cell proliferation with Arrdc4 overexpression (Fig. S4A), accompanied by a roughly two-fold reduction in the percentage of cells in the S-phase of the cell cycle (Fig. 4C), although no signs of apoptosis were observed (Fig. 4D). Consistent with this partial arrest, colony formation and transwell migration assays showed notable reductions in both proliferation (Figs. S4B and S4C) and migration (Figs. S4D and S4E). To examine the effects of Arrdc4 overexpression in animals, we implanted 4T1 cells (1.5×10^6^ per mouse) overexpressing either Arrdc4 (OX1, OX2 and OX3) or the empty vector (EV), into the mammary fat pads of immune-competent female Balb/c mice. Surprisingly, unlike the vigorous tumor growth seen in the EV cells, none of the three Arrdc4-overexpressing clones developed detectable tumors over a monitoring period of more than three months (Figs. 4E and 4F). Consistent with this finding, and unlike the EV group, no metastatic nodules were detected in the lungs after inoculation of OX cells into the tail vein or fat pads of Balb/c mice (Figs. S4F-S4I).

**Figure 4:**
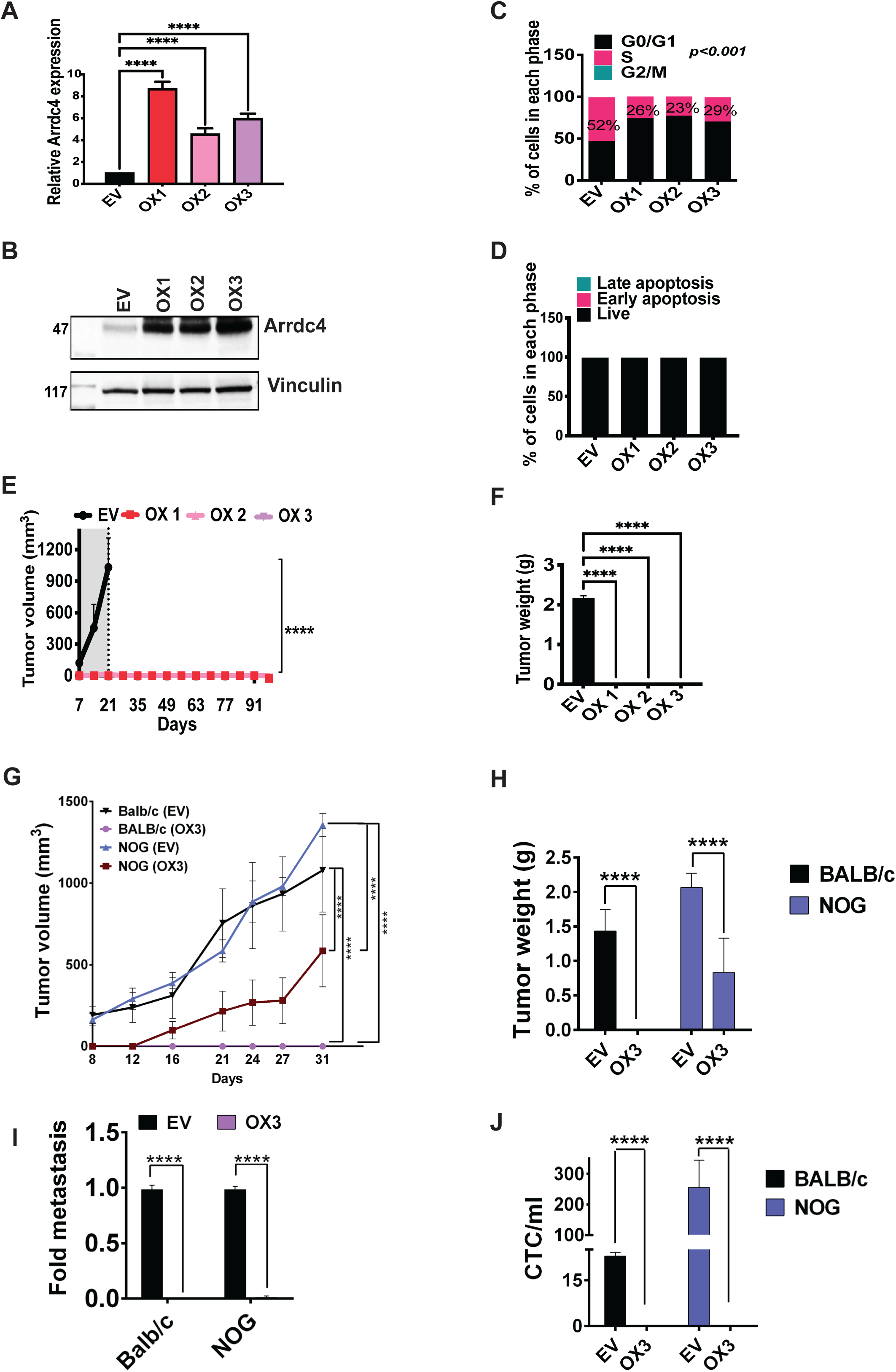
Arrdc4 overexpression in 4T1 cells prevents tumor progression in immunocompetent mice. (**A**) Arrdc4 was stably overexpressed in 4T1 cells and three independent stable clones, along with an empty vector (EV) control, were established. Shown are bar graphs representing the respective mRNA levels of Arrdc4 in each clone using qRT-PCR (n=4). (**B**) Immunoblots showing the protein levels of Arrdc4 in the empty vector control (EV), as well as in overexpressing (OX) clones. Vinculin was used as a gel loading control. (**C**) Bar graphs showing cell cycle analyses that employed BrDU (bromodeoxyuridine) incorporation followed by cytometry. (**D**) Cells were incubated with an anti-APC-Annexin V antibody and 7-aminoactinomycin D (7-AAD), and then subjected to cytometry that assayed early and late apoptosis. (**E**-**F**) Groups of Balb/c female mice (n=6) were injected in the fad pads with 1.5×10^6^ control (EV) 4T1 cells, or the derivative Arrdc4 OX clones. Tumor volumes were monitored for up to 95 days (E). Shown are average tumor volumes ±SD (in mm^3^). Tumor weights (in grams) are presented in F. **(G**-**J)** Groups of female Balb/c mice (n=6), or the immunocompromised strain NOG, were injected in the fad pad with Arrdc4 overexpressing cells (OX; 1.5×10^6^). Tumor volumes were monitored for 4 weeks (G) and both tumor weight (H) and spontaneous lung metastases (I) were determined in both strains of mice. Likewise, CTCs were isolated from freshly collected blood and quantified using cytometry (J). The levels of significance are presented as follows: ***, p<0.001 and ****, p<0.0001.

Based on these lines of evidence, we concluded that overexpression of Arrdc4 exhibits a strong inhibitory effect on 4T1 tumorigenic and metastatic growth, implying a tumor suppressive role. This likely mirrors the known function of alpha arrestins, which promote endosomal trafficking and lysosomal degradation of glucose transporters like Glut1 ^34^. To investigate whether Arrdc4 regulates Glut1 in 4T1 cells, we analyzed derivatives that overexpressed Arrdc4 (OX3) or expressed an Arrdc4-specific shRNA (sh3). To quantify Glut1 levels at the cell membrane, we adapted a previously established protocol ^35^. Four 4T1 derivatives, including those expressing scrambled shRNA and an empty plasmid, were seeded on glass coverslips and subsequently stained for Glut1. High-resolution images were obtained, and the amount of Glut1 in a small region across the cell edge (region of interest, ROI) was measured (Fig. S4J). As shown in Figure S4K, Arrdc4 downregulation significantly increased Glut1 abundance, in support of a model in which Arrdc4 normally inhibits glucose uptake, hence suppresses tumor growth and survival of CTCs.

### Arrdc4 induces immune-mediated tumor suppression

In addition to the ability of Arrdc4 to control intracellular trafficking of glucose transporters, the complete lack of both primary and secondary tumors of Arrdc4-overexpressing cells in immunocompetent mice suggested yet another function, namely engagement of one of the immune system’s suppressive mechanisms. To explore this hypothesis, we repeated the in vivo experiments using NOG (NOD/Shi-scid/IL-2Rγnull) mice, which exhibit severe immunodeficiency. In these experiments, we compared two derivatives of the 4T1 cell line: the OX3 clone, which overexpresses murine Arrdc4, and a control EV clone. Cells were implanted in the mammary fat pads of female mice (either NOG or Balb/c), and tumor growth was observed over a 30-day period. As expected, the EV clone rapidly formed tumors in both animal strains, while the OX3 clone did not develop tumors in Balb/c mice (Figs. 4G and 4H). In contrast, in the immunocompromised NOG mice the OX3 clone produced tumors but only after a latency period of 12 days (Fig. 4G). Interestingly, the OX3 clone did not produce any metastatic lung nodules (Fig. 4I), and no circulating tumor cells (CTCs) were detected in the blood of NOG mice bearing OX3 tumors (Fig. 4J). These results suggest that, in addition to its intrinsic role in inhibiting tumor cell proliferation, Arrdc4 also exerts extrinsic effects that depend on an intact immune system.

### Transcriptome and proteome analyses identify Gasdermin E as a potential mediator of the immune suppressive effects of Arrdc4

Arrdc4 belongs to the arrestin fold family, which controls the fate of transmembrane and other proteins ^36^. For example, Arrdc4 functions as an adaptor that physically interacts with the Itch/Nedd4 family of E3 ubiquitin ligases ^37^, to control trafficking of the transporters of glucose and divalent metal ions. To identify physically interacting partners of Arrdc4, we performed pull-down experiments using anti-GFP beads and extracts of 4T1 cells overexpressing a GFP-tagged Arrdc4. On-bead trypsin digestion was followed by reducion and alkylation, prior to chromatographic separation and mass spectrometry. Figure S5A shows the top 20 interacting partners, along with the respective numbers of unique peptides. As expected, this list included Itch, a member of the NEDD4 family. In addition, we detected Irs4, a member of the insulin receptor substrate family, which regulates glucose homeostasis. However, no direct immune effectors were uncovered by this approach.

As an alternative approach, we applied both RNA sequencing and whole-cell proteomic analysis on Arrdc4-overexpressing (OX) cells. RNA sequencing using the MARS-seq protocol identified 1,444 differentially expressed genes, whereas in-depth proteomic analysis detected 7,310 proteins with altered abundance (see Supplementary Excel File 1). Key differentially expressed genes (Fig. S5B) and proteins (Fig. S5C) are presented in volcano plots. To reduce complexity and eliminate noise signals, we present in Figure 5A a volcano plot showing genes and proteins that are differentially expressed and highly correlated in both RNA sequencing and deep proteomics analyses. The observed high positive correlation between idividual transcript levels and the respective protein abundance (R=0.75) ascribed high reliability to the transcriptome and proteome analyses we performed. The uppermost up-and down-regulated genes/proteins shown in Figure 5A in red and blue, respectively include a water channel (Aqp1), an adenylate kinase involved in energy metabolism (Ak1), a chaperone (peptidylprolyl isomerase C, Ppic), and an enzyme regulating cysteine proteases (Cstm1). Gratifyingly, genes associated with the epithelial-to-mesenchymal transition (EMT), particularly partial (biphenotypic) EMT, were identified in the integrated transcriptome/proteome list. For instance, we detected N-cadherin (Cdh2), which aids in cancer cell migration and metastasis, and Cadm1, a cell-to-cell adhesion molecule. Additionally, four epithelial keratins (K7, K8, K18, and K19) were downregulated in Arrdc4-overexpressing cells. In general, the downregulation of intermediate filament keratins, aside from vimentin, has been linked to partial or complete EMT ^38^. For instance, downregulation of K18 induces partial EMT and stemness through increasing EpCAM expression in BC ^39^. Equally important, we also observed increased abundance of Zeb1 and integrin beta-4. The latter, a component of the integrin α6β4 complex, is crucial for cell adhesion to the extracellular matrix (ECM), whereas Zeb1 is a master EMT transcription factor that represses the epithelial transcription factor TAp63α (tumor protein 63 isoform 1), promoting integrin beta-4 expression and identifying partially mesenchymal cancer stem cell-enriched populations of mammary carcinomas ^40^.

**Figure 5:**
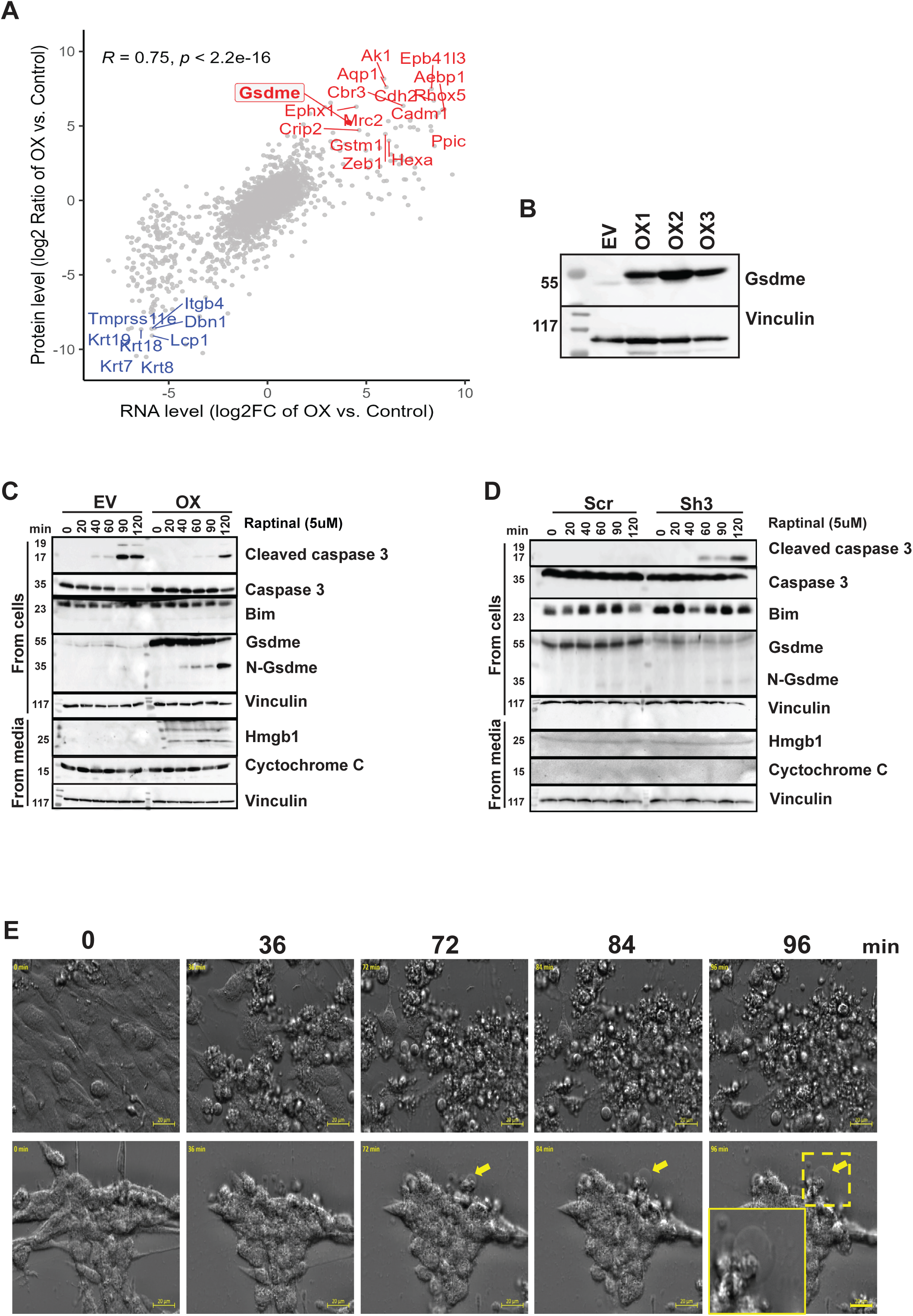
Arrdc4 promotes pyroptosis by means of means of engaging Gsdme. (**A**) A volcano plot showing genes and proteins that are differentially expressed and highly correlated in both RNA sequencing and deep proteomics analyses. The comparison was made between the ratio of protein expression in Arrdc4 OX (overexpressing) cells versus control cells (vertical axis) and the fold change in RNA expression level between Arrdc4 OX cells vs. control cells (horizontal axis). Pearson correlation was used to determine the correlation between the measurements. (**B**) An immunoblot showing protein levels of Gsdme in control (EV) and three Arrdc4-overexpressing 4T1 cells. (**C**) Control 4T1 (EV) and Arrdc4-overexpressing (OX) cells were treated with raptinal (5µM) for different time intervals, up to 120 minutes. The immunoblot examined activation of caspase 3 and Gsdme cleavage, along with release of the high mobility group box 1 (Hmgb1) protein. (**D**) 4T1 control (Scr) and Arrdc4-downregulated cells (sh3) were treated with raptinal as in C and both cell extracts and supernatants were analyzed as indicated. (**E**) Control (upper row) and Arrdc4-overexpressing cells (lower row) were cultured in glass chambered slides and treated with raptinal (5µM) for up to 96 minutes. Simultaneous live cell imaging that used the ZEISS Celldiscoverer 7 microscope (40X air objective) and an oblique frame was done from 0 to 96 minutes. Snaps captured during live imaging at the indicated time points are shown. Yellow arrows indicate cell ballooning characteristic to pyroptosis. The yellow frame shows the zoom-in image. Scale bar, 20 μm.

Along with Zeb1, high mRNA and protein corresponding to gasdermin E (Gsdme), and to a lesser extent also gasdermin D, were detected in Arrdc4-overexpressing cells. Gsdme acts as a tumor suppressor by regulating immune cell-mediated inflammatory cell death, known as pyroptosis ^41^. Caspase-3 cleavage of Gsdme generates a pore-forming amino-terminal fragment of Gsdme that induces cell death. Given that, in general, Gsdme’s tumor suppression is dependent on immune cells, these findings suggested that the elevation of Gsdme by Arrdc4 could explain the immune-mediated tumor suppression we observed when testing Arrdc4-overexpressing 4T1 clones in immunocompetent mice. Consistent with this, Gsdme was upregulated in all three independently isolated Arrdc4-overexpressing clones, relative to the control clone (Figs. 5B and S5D). Importantly, a previous report found that Zeb1 binds to and actiavtes Gsdme’s promoter ^42^, which might explain the co-expression we observed in derivatives of 4T1 cells.

To examine the putative role for Arrdc4 in pyroptosis, we treated Arrdc4-overexpressing cells (OX3) with raptinal, a caspase-3 activator, assuming that active caspase-3 would cleave Gsdme and generate a free N-terminus necessary for pore formation. Indeed, shortly after exposure to raptinal Arrdc4-overexpressing cells, unlike the control (EV) cells, showed a marked increase in Gsdme levels (Fig. 5C). Accordingly, only the Arrdc4-overexpressing cells displayed the pore-forming fragment of Gsdme (N-Gsdme). Additionally, the raptinal treated OX3 cells released HMGB1 (high mobility group box-1 protein), a proinflammatory cytokine and marker of pyroptosis. We then explored whether the same sequence of events, linking Arrdc4 to Gsdme cleavage, took place in Arrdc4-knockdown cells (clone sh3). In line with the low levels of Gsdme, sh3 cells did not release HMGB1 following raptinal treatment, despite the activation of caspase-3 (Fig. 5D). This observation suggests that a key difference between control and Arrdc4-overexpressing 4T1 cells is the activation of the pyroptotic pathway, which functions to suppress tumor growth and enhance immune inhibition.

Previous research has shown that granzymes and perforins released by NK cells, cytotoxic T lymphocytes, as well as macrophages, activate caspase 3, which then cleaves Gsdme ^41,43^. This cleavage leads to cell blebbing (or ballooning), a characteristic feature that distinguishes pyroptosis from other cell death mechanisms. To investigate this, we conducted live cell imaging following treatment with raptinal. Consistent with pyroptosis, cells overexpressing Arrdc4 exhibited the expected blebbing and ballooning (marked by yellow arrows in Fig. 5E, lower panels), whereas control (EV) cells did not. In summary, the data shown in Figures 5 and S5 indicated that Arrdc4 can facilitate partial EMT and elevate Gsdme levels, increasing the vulnerability of BC cells to pyroptosis. Consequently, by downregulating Arrdc4, disseminated BC cells may adjust their EMT process, ensure a steady glucose supply, and gain protection against inflammatory cell death.

### Arrdc4’s anti-tumor effect involves recruitment of perforin-and granzyme-positive immune cells to mammary tumors

To verify that Arrdc4’s tumor-suppressive effects are mediated by Gsdme, we used CRISPR-Cas9 to knock out the endogenous gasdermin E gene in Arrdc4-overexpressing 4T1 cells (OX3). As expected, immunoblotting confirmed the absence of the protein in the resulting cells (called: OX3 Gsdme-KO; Fig. S6A). These cells, along with the EV and OX3 cells, were implanted in the mammary fat pads of immunocompetent Balb/c mice (1.5×10^6^ cells per mouse), and tumor growth was monitored over 3-4 weeks (Fig. S6B). Whereas control cells (EV) exhibited robust tumor growth and OX3 cells did not form tumors, the Gsdme-deficient 4T1 derivatives initially displayed slow rate of tumor development, which sharply accelerated after three weeks. Considering our findings in immunocompromised mice, the biphasic growth curves of the Gsdme-deficient OX3 cells in immunocompetent mice validated that Gsdme is involved in Arrdc4-mediated immune suppression of tumor formation. These observations also implied that additional genes and yet-to-be-identified dynamic processes have roles to play in the ability of the Arrdc4-Gsdme pathway to regulate tumorigenesis.

The coordinated regulation of Arrdc4 and Gsdme suggested that upregulation of this pathway will result in the cleavage of Gsdme, with its cleavage product triggering the recruitment of cytotoxic immune cells prior to the onset of pyroptosis. To test this hypothesis in animals, despite the difficulty posed by the rapid elimination of Arrdc4-overexpressing cells in immunocompetent mice, we employed a tumor transplant strategy (refer to the scheme shown in Fig. 6A): two clones of Arrdc4-overexpressing 4T1 cells (OX1 and OX3), along with control EV cells, were first implanted in the fat pads of immunocompromised NOG mice. Once tumors reached a size of 150 mm³, they were surgically removed and transplanted in the fat pads of naïve female Balb/c mice. While the EV control tumors grew aggressively, none of the tumors from Arrdc4-overexpressing cells exhibited similar growth patterns (Fig. 6B). However, variability was observed among the recipient mice: the M1 Arrdc4-overexpressing tumor from animal number 1 remained stagnant, whereas the M2 tumor from another animal initially regressed but later resumed slow growth. Notably, despite these differences, resection and reimplantation of of M1’s and M2’s fragments in naïve Balb/c mice consistently led to tumor regression (Fig. S6C).

**Figure 6:**
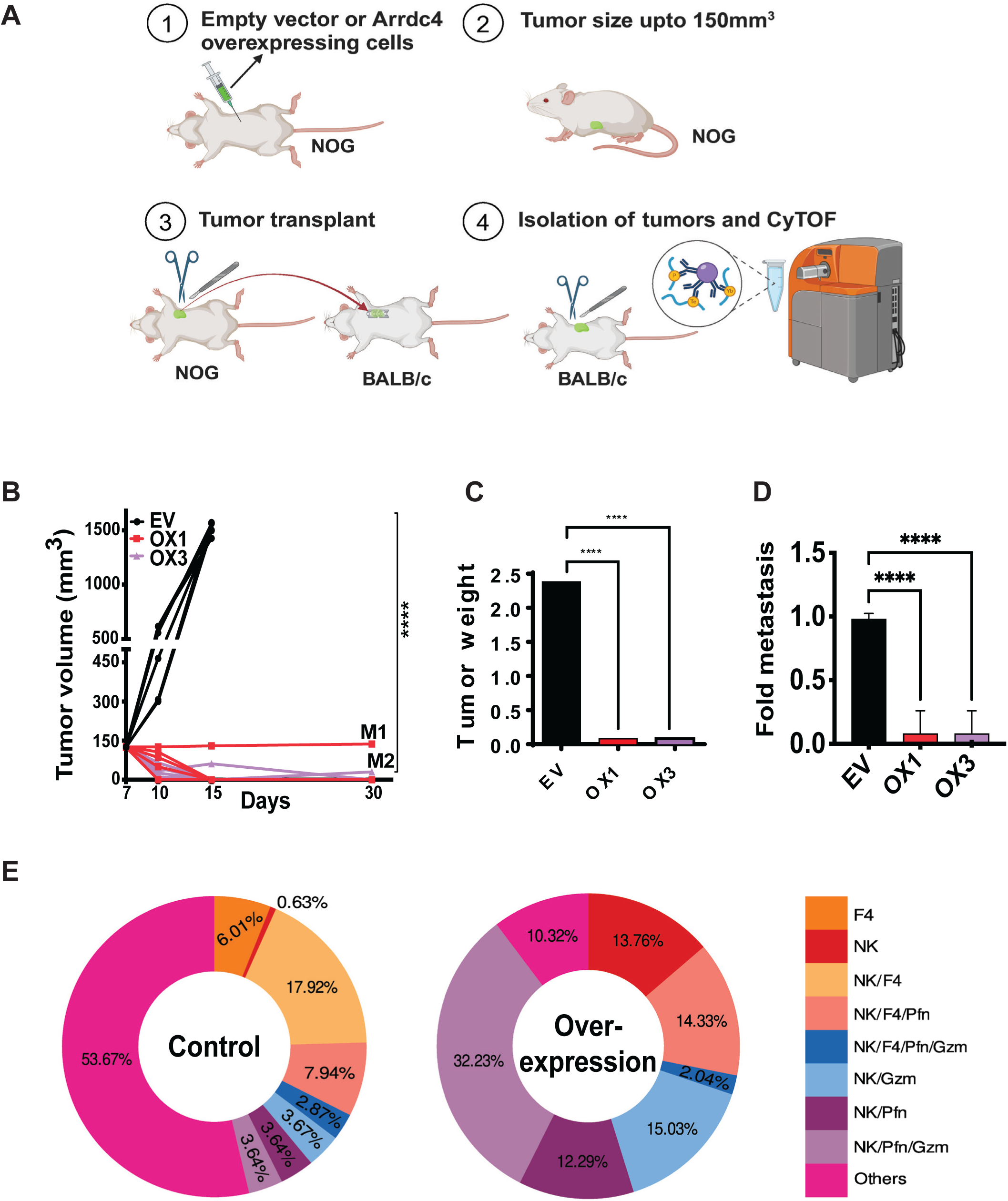
Suppression of Arrdc4-overexpressing tumors in animals involves recruitment of immune cells. (**A**) Schematic representation of the protocol used for tumor growth in immunocompromized (NOG) mice, followed by resection and transplantation of cancer cells in immunocompetent (Balb/c) mice. The figure was created using BioRender.com. (**B**-**D**) 4T1 control (EV) tumors and tumors overexpressing Arrdc4 (OX1 and OX3) were excised from NOG mice (see panel C). Next, 6 weeks old female Balb/c mice (5 per group) were implanted in their mammary fat pads with equal volume and weight of 4T1 empty vector (EV) and Arrdc4-overexpressing (OX1 and OX3) tumors originally isolated from NOG mice. The line graphs show the average tumor volumes. M1 and M2 refer to the recipient mice of the secondary Arrdc4-overexpressing cells. Also shown are bar graphs presenting the respective tumor weight (C) and events of spontaneous lung metastases (D). (**G**) Control 4T1 cells (EV) and Arrdc4-overexpressing tumors (OX) were isolated from tumor-bearing animals on day 10 after tumor cell injection into female Balb/c mice and used for CyTOF analysis. The assay made use of the following heavy metal-conjugated primary antibodies: anti-CD45 (a general immunological marker), CD8a (cytotoxic T cells), F4/80 (macrophages), Nkp46 (NK cells), granzyme B and perforins. Shown are the fractions of the respective immune cell types in EV tumors and in Arrdc4-overexpressing tumors.

As expected, we observed significant differences between the EV and OX groups of immunocompetent mice in terms of tumor weight and metastasis (Figs. 6C and 6D). To determine if these differences were accompanied by differential recuitment of immune cells, we applied cytometry by time-of-flight (CyTOF) on tumors collected on day 10 post inoculation in recipient mice. To achieve high-dimensional description of multiple markers at the single-cell level, CyTOF makes use of sets of antibodies conjugated to heavy metals. Practically, we utilized a pyroptosis centered panel, comprising antibodies targeting markers specific to CD8^+^, cytotoxic CD45-positive cells, NK cells and macrophages, as well as granzyme (Gzm)-specific and perforin (Pfn)-specific antibodies. Sample barcoding and three biological replicates were employed to address batch effects and also ensure reproducible results across multiple runs. The data from these replicates were aggregated, encompassing a total of 10,000 cells for both control and Arrdc4-overexpressing tumor samples. Data visualization used UMAP, which highlighted distinct cellular populations in control (Fig. S6D) and Arrdc4-overexpressing tumors (Fig. S6E). Importantly, the CyTOF analyses identified in the microenvironment of Arrdc4-OX tumors several key populations of cells that are likely associated with pyroptosis induction. Thus, cluster analysis showed that Arrdc4 overexpression led to increased infiltration by macrophages and NK cells positive for perforin-and/or granzyme (Fig. 6E), in line with pyroptosis initiation and progression. In contrast, control tumors displayed high proportion of undefined cells and fewer granzyme-and perforin-positive immune cells. Taken together, these studies supported the possibilty that Arrdc4 overexpression enhances tumor cell susceptibility to immune surveillance by means of engaging a cascade initiated by Gsdme cleavage and pore formation and culminating in the release of inflammatory agents and cell death.

### The Arrdc4-Gsdme module induces partial EMT

Although previous studies reported an association between gasdermins and EMT, no study directly linked an alpha-arrestin like Arrdc4 to loss of epithelial traits. For example, it has been reported that the ZEB family of mesenchymal transcription factors can activate the promoter of Gsdme ^42^, and TGF-beta, a highly potent inducer of ZEB proteins, can instigate cleavage of Gasdermin D ^44^. In line with the ability of Arrdc4 to increase expression of Zeb1 and downregulate epithelial keratins (Fig. 5A), Arrdc4-overexpressing cells displayed spindle-like morphologies resembling mesenchymal cells (Fig. 7A). In contrast, Arrdc4-depleted 4T1 cells (shArrdc4), as well as cells expressing a scrambled shRNA, displayed round morphologies. Next, we focused on the effect of Arrdc4 on TGF-beta induced EMT. Firstly, we starved for serum factors both the EV control and the Arrdc4-overexpressing (OX) cells, and then treated all cells with TGF-beta for increasing time intervals (Fig. 7B). Consistent with the possibility that both Arrdc4 overexpression and stimulation with TGF-beta induce EMT, Gsdme was detectable only in OX cells and these cells displayed loss of E-cadherinn, as well as up-regulation of vimentin, prior to TGF-beta treatment. Notably, TGF-beta did not downregulate E-cadherin in EV cells, in line with the reported paradoxical role for E-cadherin in 4T1 cells ^45,46^. We also observed alterations in the abundance and phosphorylation of H2AX (gamma-H2AX), a marker of DNA breaks ^47^ and a regulator of EMT ^48^. Collectively, these findings confirmed the ability of Arrdc4 to induce Gsdme and raised the possibility that this alpha-arrestin might strengthen the ability of TGF-beta to induce partial or complete EMT.

**Figure 7:**
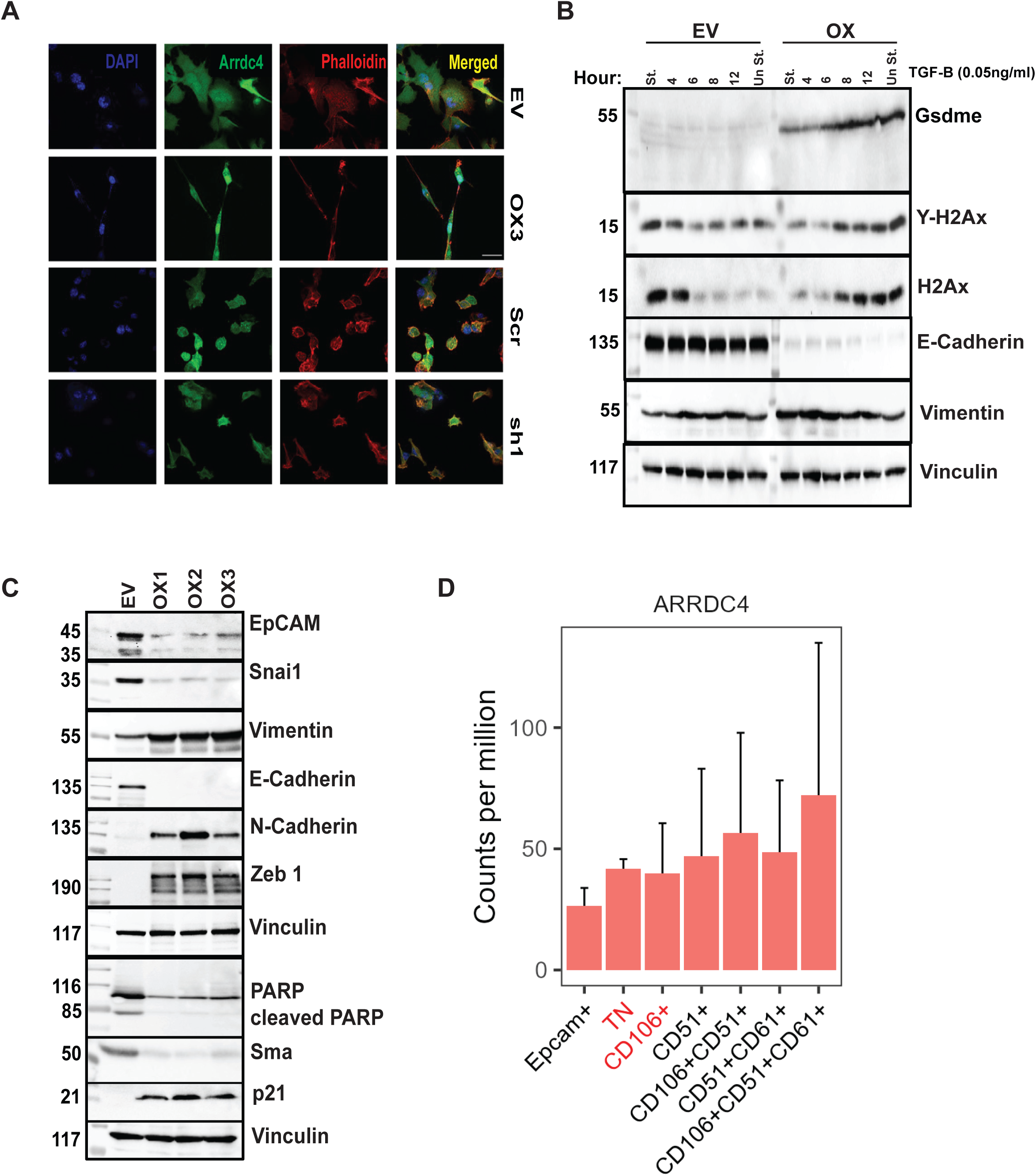
Arrdc4 elevates Gsdme and promotes partial EMT. (**A**) Immunofluorescence images comparing the morphology of control (EV) 4T1 cells, Arrdc4-overexpressing (OX) cells, along with Arrdc4 knockdown cells (sh1) and cells expressing scrambled shRNA (Scr). Phalloidin (red) was used to visualize actin filaments and DAPI (blue) was used to stain nuclei. Images were captured using a confocal microscope (100X magnification). Scale bar, 25 μm. (**B**) An immunoblot showing time course analyses of the induction of different proteins in 4T1 cells, either cells overexpressing Arrdc4 (OX) or control (EV) cells, which were treated with TGF-beta (50 pg/ml), two hours post serum starvation. Note that TGF-beta induces Gsdme and vimentin expression, as well as reduces E-cadherin expression, but these effects are largely limited to Arrdc4-overexpressing cells. (**C**) An immunoblot showing expression of both epithelial (EpCAM, Snai1 and E-cadherin) and mesenchymal markers (vimentin, N-cadherin and Zeb1), along with the expression of PARP, SMA (smooth muscle actin) and p21, in 4T1-Arrdc4 overexpressing (OX) cells as compared to 4T1 control (EV) cells. (**D**) Shown are levels of ARRDC4 expression in the long-lasting cellular states underlying EMT ^20^. TN refers to a triple negative state lacking expression of the epithelial cell adhesion molecule (EpCAM), as well as lacking expression of CD106 (Vascular Cell Adhesion Molecule 1; VCAM-1), CD51 (integrin alpha V; ITGAV) and CD61 (integrin beta-3; ITGB3). Note that TN and the next state, CD106+ (both shown in red), are considered the most metastatic transition states. The data were derived from GEO record GSE110587. Error bars refer to the standard error values (n=3).

To reinforce this conclusion, we extended the comparative analysis to all three Arrdc4-overexpressing (OX) clones, and also included additional EMT markers (Fig. 7C). Our assays confirmed, on the one hand, a positive effect of Arrdc4 on mesenchymal markers, such Zeb1, N-cadherin and vimentin and, on the other hand, a negative effect on epithelial markers like EpCAM and E-cadherin. Nevertheless, we also observed Arrdc4-induced downregulation of the mesenchymal markers smooth muscle actin (Sma) and Snail (Snai), implying that rather than switching from the epithelial (E) to the mesenchymal (M) state, Arrdc4 stabilizes a biphenotypic E+M state ^18,49^. Interestingly, poly (ADP-ribose) polymerase (PARP), which supports DNA repair, was downregulated in OX cells, wheraes p21, a marker of EMT-mediated cell cycle arrest, underwent upregulation. Taken together with other lines of evidence, it is plausible that 4T1-derived CTCs downregulate Arrdc4, as well as alter the abundance of a subset of mesenchymal markers, as a mechanism that confers a biphenotypic state characterized by low Gsdme expression, sustained glucose uptake and increased protection from pyroptosis.

### Loss of Arrdc4 typifies early biphenotypic EMT states known to be highly metastatic

Since downregulation of cytokeratins, especially K18 (Fig. 5A), induces partial EMT through regulation of EpCAM expression ^39^, and low abundance of Snail (SNAI1), another marker of Arrdc4-overexpressing cells, can contribute to the initial states of EMT ^50^, we suspected that low Arrdc4 levels would associate with early EMT hybrid states. To examine this prediction, we refered to a previous study that identified in primary tumors several subpopulations, from E to M through biphenotypic hybrid states differing in plasticity, invasiveness and metastatic potential ^20^. Utilizing the defining surface proteins, CD106 (Vcam1), CD51 (integrin alpha-V) and CD61 (integrin beta-3), we validated that Arrdc4’s levels were lowest in two early transition states: triple negative (TN) and the CD106-positive state (red labeled in Figure 7D). Importantly, it has been reported that these two states are endowed with the highest metastatic potential ^20^.

The unexpected link connecting ARRDC4 and Gasdermin E, as well as the concurrence of low ARRDC4, low GSDME and the most metastatic EMT states, motivated analyses of the other gasdermins and additional pyroptosis markers. As expected, ARRDC4 shared with mesenchymal markers a shallow trend of increasing abundance towards the M state (left box of Fig. S7A). In addition, analysis of a previously reported pyroptosis gene signature ^51^, which includes four pore-forming gasdermins (Fig. S7A, right box), similarly resolved two opposite trends: a group of pyroptosis markers, such as TNF and AIM2, that exhibited a descending gradient along the series of partial (hybrid) EMT states, and another group of markers that parallelly underwent up-regulation as the mesenchymal attributes gradually increased up the EMT ladder. Interestingly, GSDME and GSDMD, which are directly activated by caspases, belong to the latter group, whereas GSDMA and GSDMC, which have unique activation mechanisms, displayed a reciprocal trend (Fig. S7B). Although the significance of these expression gradients remains unknown, they suggest complex interactions among pyroptosis, EMT and metastasis. Relatedly, because biphenotypic states, rather than fully E or fully M states, have been associated with relatively poor clinical prognosis ^11,18^, the impact of the ARRDC4-GSDME module on metastasis might arbitrate immune tradeoffs. Stated in other words, while the more mesenchymal EMT states are better fitted for invasive growth, their high Arrdc4 levels are expected to confer susceptibility to pyroptosis. Inversely, due to their low Arrdc4 levels the more epithelial biphenotypic states, especially the most metastatic TN and CD106-positive states ^20,52^, might better seed metastases due to their ability to evade pyroptosis.

In summary, by selecting CTCs with gradually enhanced intravasation and survival in the bloodstream of tumor-bearing immunocompetent mice, we identified an alpha-arrestin protein, Arrdc4, as a metastasis suppressor. The best characterized function of Arrdc4 is to downregulate glucose transporters, suggesting that persistent supply of glucose is vital for effective dissemination of BC cells. Supporting this model, Arrdc4 overexpression reduced cell proliferation in vitro and slowed tumor growth in immunocompromised mice. Importantly, however, when Arrdc4-overexpressing cells were implanted in immunocompetent animals, they completely lost their tumor-forming capability and did not generate any CTCs. This surprising observation suggested that Arrdc4 induces potent immune suppression of CTCs. In line with this model, transcriptomic and proteomic studies revealed co-regulation between Arrdc4 and gasdermin E, a protein that triggers pyroptosis. Further research implied that the downregulation of Arrdc4 allows CTCs not only to sustain their glucose uptake but also evade inflammatory death by NK and other immune cells. Notably, reduced levels of the murine forms of both Arrdc4 and Gsdme are associated with partially epithelial biphenotypic EMT states, which have previously been linked to the most metastatic hybrid EMT states. Patient data support the clinical relevance of these findings: both ARRDC4 and GSDME are downregulated in mammary tumors compared to the adjacent normal tissue. Moreover, disabled mutants of human GSDME, which are oncogenic, have previously been reported ^41^, and we report herein that downregulation of ARRDC4 in patients with BC is driven by both epigenetic (DNA methylation) and genetic (copy number alterations) mechanisms. As expected, both aberrations can predict relatively short survival time of patients with BC.

## Discussion

CTCs are not only extremely rare in the bloodstream, with estimates of only one CTC per 1-10 million white blood cells, they also have very short half-life in circulation, estimated to be below 2.5 hours ^53,54^. The rapid clearance rates of CTCs have been attributed to the physical forces and shear stress in the bloodstream, which can damage CTCs. In addition, because CTCs often detach from the extracellular matrix, they may undergo anoikis, a form of programmed cell death, but some CTCs evolve resistance to anoikis ^55^. Another form of programmed cell death, pyroptosis, that is instigated by gasdermins and characterized by local inflammation, has been implicated in tumor suppression ^41^. However, no previous report has directly linked CTCs to pyroptosis. Unlike apoptosis, which is non-inflammatory, pyroptosis leads to the release of pro-inflammatory cytokines and danger-associated molecular patterns (DAMPs). Our in vivo screens led to the identification of a novel upstream regulator of gasdermins, ARRDC4, which inhibits another group of vital proteins, glucose transporters (Glut). Thus, by downregulating the abundance of ARRDC4, CTCs acquire on the one hand protection from pyroptosis and, on the other hand, they gain free access to glucose, thus enhancing their survival while en route. Notably, in addition to glucose transporters, ARRDC4 also regulates transporters of divalent metal ions ^56^, such as calcium, iron and zinc, which are known as regulators of cell adhesion, invasion and metastasis.

Our results and previous observations propose that granzymes and perforins released by NK cells, cytotoxic T lymphocytes and macrophages activate caspase 3, which then cleaves GSDME ^41,43^, leading to perforation of cancer cells that acquired mesenchymal traits. In contrast, other sets of blood cells might protect, rather than damage CTCs. For example, the circulating malignant cells might increase their chance to survive and reach distant niches by means of physical interactions with neutrophils ^57^. The latter can promote metastasis also in an indirect manner, by means of CTC shielding structures called neutrophil extracellular traps (NETs) ^58^, which promote EMT ^59^. Similarily, platelets can aggregate CTCs in patients’ bloodstream, accelerate their EMT and induce resistance to anoikis ^60^. Another protective cell type, macrophages, enhance CTC survival by means of a different mechanism, namely formation of macrophage-tumor cell hybrids in the blood of patients with multiple cancers ^61^. Thus, while in transit, distinct sets of blood-borne cells differentially determine the fate of CTCs, in part by controlling their mesenchymal phenotype.

Although both ARRDC4 and GSDME act as tumor suppressors that are lowly expressed in BC, compared to the surrounding tissue, their co-regulation is both novel and complex. Despite absence of DNA and RNA binding domains, ARRDC4 extensively modifies cell morphology, metabolism and EMT. These functions likely relate to ARRDC4-mediated regulation of different receptors, transporters and extracellular vesicles ^37,56^. One outcome of these alterations is the induction of EMT, including downregulation of E-cadherin, partial growth arrest and upregulation of ZEB1. As ZEB1 binds with and activates GSDME’s promoter ^42^, it is likely that the most mesenchymal states of the previously identified EMT hybrid states ^20^ are prone to GSDME-induced pyroptosis. Consistent with this possibility, the levels of both ARRDC4 and GSDME gradually increase along the series of biphenotypic EMT states and this pattern is shared by a few pyroptosis marker genes. In patients with BC, reduced GSDME levels are associated with decreased patient survival time, and multiple cancer-associated mutations affecting GSDME inhibit its function as a driver of pyroptosis ^41,62^. Likewise, we report herein that ARRDC4 expression is downregulated in BC and this is due to DNA methylation and copy number aberrations, changes that contribute to shorter patient survival time. Since the vast majority of cancer patients die as a consequence of their metastatic disease, the uncovered link between CTCs, EMT and pyroptosis might bear wider clinical relevance. In this context, it is wortwhile noting that the most metastatic EMT transition states are characterized by co-expression of both epithelial and mesenchymal markers ^52^. As we report herein these partial EMT states are also chracterized by low abundance of both ARRDC4 and GSDME. Hence, this corespondence might bottleneck metastasis by means of limiting glucose availability and elevating susceptibility to inflammatory cell death. Furthermore, since pro-inflammatory cytokines, such as IL-1-beta and IL-6, can induce EMT, a feed forward loop involving the ARRDC4-GSDME axis might restrict the contribtion of the mesenchymal states to metastasis. Future studies will likely uncover additional links connecting ARRDC4 to aerobic glycosylysis ^63^ and the occurence of CTC clusters having increased metastatic potential^16^.

## Materials and methods

### Cell culture

The mouse 4T1 (#CRL-2539), human HEK-293T (CRL-1573) and HeLa cell lines were purchased from the American Type Cell Culture collection (ATCC). 4T1 cells were grown in Roswell Park Memorial Institute 1640 (RPMI, Gibco, Cat: 21875034) supplemented with 10% (v/v) fetal bovine serum (FBS, Gibco, Cat:12657029) and 1% penicillin-streptomycin (i.e., 100 U/ml, Biological Industries, Israel; Cat: 030311B). HEK-293T and HeLa cells were grown in Dulbecco Modified Eagle Medium (DMEM, Gibco, Cat: 41965039) supplemented with 10% (v/v) FBS, 1% penicillin-streptomycin (i.e., 100 U/ml).

### Antibodies and Reagents

The following antibodies were used for immunoblotting: from Cell Signalling: ERK1/2 (#4695), pERK1/2 (T202/Y204; #9101), AKT1 (#2938), pAKT (S473; #4060), mTOR (#4517), caspase3 (#9622), cleaved caspase3 (#9661), BIM (#2933), γH2AX (#2577), cytochrome c (#4272), HMGB1 (#3935), p21 (#2947), SMA (#19245), PARP (#9542), snail (#3879), vimentin (#5741), N-cadherin (#13116), c-Cadherin (#3195), HA (#3724) and FLAG (#8146). From Millipore: GAPDH (#MAB374). From Thermo Scientific: tubulin (#PA5-58711), GLUT4 (#MA183191) and ARRDC4 antibody (#PA536518). From Abcam: GLUT1 (#ab32551), mCherry (#ab167453) and GSDME (#ab215191). From Santa Cruz Biotechnology: IRS4 (#sc-393207), Itgb4 (#sc-514426), Zeb1(#sc-25388), EpCAM (#sc-25308) and Ds Red antibody (sc-390909). From MBL: GFP antibody (598). From Sigma: vinculin (V9264), phalloidin (P1951). From Jackson ImmunoResearch Laboratories: Secondary peroxidase-conjugated anti-mouse IgG (715-035-151) and anti-rabbit IgG (#715-035-152). For CyTOF, we used the following heavy metal conjugated antibodies that were obtained from Fluidigm-CD45 (89Y; #3089003B), NKp46 (167Er; #3167008B), perforin (172Yb; #3172018B), F4/80 (159Tb; #3159009B) and granzyme B (171Yb; #3171002B). In addition, we used raptinal (Sigma Cat# SML-1745) and TGFbeta (Peprotech Cat# SML-1745) were used post serum starvation. Ficoll (Cytiva 17144002), FAST SYBR Green master mix (Thermofisher #4385612), Alexa Fluor 647 BrdU Kit (BioLegend #370706), Annexin A5 Apoptosis Detection Kit (BioLegend #640930), Dual-Luciferase Reporter Assay System (Promega #E1910), qScript cDNA sysnthesis kit (QuantaBio #95047), RNA isolation kit (Qiagen #217004)

### Plasmids and shRNAs

The following plasmids were kind gifts from Sharad Kumar (University of South Australia): pEGFPN1-Arrdc4 (WT), pEGFPN1-Arrdc4-PY and pEGFPN1-Arrdc4-F115L; Carlos E. Alvarez (Nationwide Children’s Hospital, Columbus, Ohio): pcDNA3.1/ARRDC4-mCherry and pcDNA3.1/ARRDC4_PPXY_mutantmCherry; Emad Alnemri (Thomas Jefferson University, Philadelphia): pEGFPN1-GSDME, from Addgene: Glut4 mCherry (Cat:64049), Glut1mCherry (Cat:105010), Gsdme (Cat:154876,154877), Irs4 (Cat:11363), HA-Ubiquitin-WT (Cat:17608) and HA-Ubiquitin-K48R (Cat:17604), and from Vector Builders: pEGFP-Bsd-mArrdc4 (Cat:VB230316-1026frs), pEGFP-puro-mArrdc4 (Cat:VB190704-1044bce) and pLVHygro-Tet3G (Cat:VB160122-10094). A set of 3 smart lentivirus shRNA pools were used for stable downregulation of Arrdc4 (Dharmacon, Cat: V3SM11244-01EG66412).

### Isolation of CTCs and single cell RNA sequencing (scRNA-seq)

A cell suspension containing 4T1-GFP cells (1×10^6^ in 0.1 ml) was injected into the mammary fat pad of 6- weeks old female BALB/c mice. After 5 days, blood was collected, using cardiac puncture, and mixed with heparin. Next, blood was diluted using saline and layered on Ficoll (Cytiva, #17144002). Red blood cells were removed using density gradient centrifugation and thereafter CTCs were purified using cytometry for GFP expression. Sorted cells were cultured in RPMI 1640 supplemented with fetal bovine serum (FBS; 10%) and puromycin, prior to scRNA-seq analysis. Single cells were sorted using FACSAria cell sorter and directly mixed with lysis buffer. Cells were sorted from 3 different types of samples: 4T1 cells, fresh CTCs collected from blood, and cultured CTCs. Sequencing libraries were prepared using Smart Seq2. A total of 874 single cells were sequenced and used for analysis that utilized the Seurat package (in R). Uniform Manifold Approximation and Projection (UMAP) was used for visualization and non-linear reduction. Differential gene expression analysis was performed using edgeR. The glmTreat function was used to determine significant changes in gene expression and the threshold was set at greater or equal to 2-fold change. Library preparation, sequencing and single cell analysis were performed at the core facilty of the Weizmann Institute of Science (G-INCPM).

### RNA extraction, qPCR and sequencing

RNA was isolated using RNeasy Mini Kit (Qiagen). RNA (1µg) was used for cDNA synthesis. cDNA was synthesized using qScript^TM^ cDNA synthesis Kit (Quanta BioSciences). Real Time quantitative PCR analyses were preformed using Fast SYBR Green Master Mix (Applied Biosystems). *GAPDH*, *Actin* and *B2M* were used as internal controls. We used the following forward and reverse primers: 5’GGGCTACTGTAACGGAGAA3’ and 5’TCCACTAGCCAAGTACGTCTG3’, respectively (corresponding to the murine Arrdc4). Similarly, we used the forward and reverse primers, 5’TGGTCACCTCGTTTACTG3’ and 5’AATGCTGGTGTGTTGACAATCG3’, respectively, for human ARRDC4. Libraries for RNA-seq were prepared following the bulk MARS-Seq protocol and sequenced using the NextSeq500 system. Quality checks, pre-processing, alignment, and differential expression analysis were performed using UMI-tools ^64^ and Salmon ^65^. Differential expression analysis was performed using the Limma and edgeR packages of R. Genes were considered to be differentially expressed if their adjusted p-value was smaller than 0.05 and their log Fold Change threshold was ±1.5.

### Cytometry by time of flight (CyTOF) and data analysis

Our protocol was based on a previously describe procedure ^66^. Up to 6 million cells were harvested, resuspended in MaxPar saline (Fluidigm) and centrifuged at 400g for 10 minutes with slow break (used throughout the protocol). Cells were labeled with 1.25µM Cell-ID Cisplatin (Fluidigm) according to the manufacturer’s instructions. Cells were washed with staining buffer (Fluidigm), then fixed using the Maxpar Nuclear Antigen Staining Buffer Set (Fluidigm) according to the manufacturer’s instructions. Up to 3 million cells per sample were barcoded and pooled using the Cell-ID 20-Plex Pd Barcoding Kit (Fluidigm) according to the manufacturer’s instructions. The pooled sample was blocked in 10% normal goat serum (NGS, Cell Signaling Technology) containing the cocktail of antibodies. Cells were then washed with staining buffer and fixed in 4% formaldehyde (Pierce, Thermo Fisher Scientific). On the acquisition day, the cells were stained using Cell-ID Intercalator-Iridium (Fluidigm) according to the manufacturer’s instructions. The cells were then washed in staining buffer and afterwards in Maxpar Cell Acquisition Solution Plus (Fluidigm). Cells were resuspended in cell acquisition solution containing 1:10 dilution of EQ Four Element Calibration Beads (Fluidigm), adjusted to attain a concentration of 300,000 cells/ml. Cells were filtered through a 35µm mesh cell strainer (Falcon) before acquiring data on a Fluidigm Helios CyTOF system. CyTOF data underwent the following pre-processing: Firstly, the CyTOF software was used for normalization and concatenation of the acquired data. Then, several gates were applied using the Cytobank platform (Beckman Coulter): First, the normalization beads were gated out using the 140Ce channel. Then, live single cells were gated using the cisplatin 195Pt, iridium DNA label in 193Ir, event length and the Gaussian parameters of width, center, offset and residual channels. CyTOF software was then used for sample de-barcoding.

### Liquid chromatography and mass spectrometry (LC/MS)

To identify Arrdc4’s binding partners, pull down experiments were performed using GFP beads that were (https://www.ptglab.com). Peptide samples were desalted using Oasis HLB μElution format (Waters, Milford, MA), vacuum dried and stored frozen. ULC/MS grade solvents were used and each sample was loaded using split-less nano-Ultra Performance Liquid Chromatography (10 kpsi nanoAcquity; Waters). The mobile phase was: A: water + 0.1% formic acid and B: acetonitrile + 0.1% formic acid. Desalting of the samples was performed online using a reversed-phase Symmetry C18 trapping column. The peptides were then separated using a T3 HSS nano-column (frpm Waters) at 0.35 µl/min. Peptides were eluted from the column into the mass spectrometer. Mass spectrometry used nanoUPLC coupled online through a nanoESI emitter (10 μm tip; New Objective; Woburn, MA, USA) to a quadrupole orbitrap mass spectrometer (Q Exactive HF) using a FlexIon nanospray apparatus (Proxeon). Data was acquired in a data dependent acquisition mode (DDA), using a Top-10 method. MS1 resolution was set to 120,000 (at m/z 200), mass range of m/z 375-1650, AGC of 3e6 and maximum injection time was set at 60 msec. MS2 resolution was set at 15,000, quadrupole isolation m/z 1.7, AGC of 1e5, dynamic exclusion of 20 sec and maximum injection time of 60 msec. Data processing and analysis used MaxQuant v1.6.6.0. The Andromeda search engine was used against the human proteome database. This was appended with common laboratory protein contaminants, and the following modifications: Carbamidomethylation of cysteine as a fixed modification and oxidation of methionine, deamidation of asparagine or glutamine and N-terminal acetylation as variables. The fold changes (ratios) between the corresponding samples were calculated on the basis of the LFQ intensities.

### Proteomic analyses

Cell pellets were lysed in 5% SDS in 50 mM Tris-HCl. Lysates were incubated at 96°C for 15 min, followed by ten cycles of 30s of sonication (Bioruptor Pico, Diagenode). Protein concentration was measured using the BCA assay (Thermo Scientific) and a total of 95 μg protein was reduced (with 5 mM dithiothreitol) and alkylated with 10 mM iodoacetamide (in the dark). Samples were loaded onto S-Trap mini-columns (Protifi). After loading, samples were washed with 90:10% methanol/50 mM triethylammonium bicarbonate, and then digested with trypsin for 1 h at 47°C. The digested peptides were eluted using 50 mM ammonium bicarbonate and trypsin was added to this fraction and incubated overnight at 37°C. Two more elutions were made using 0.2% formic acid and 0.2% formic acid in 50% acetonitrile. The three elutions were pooled together and vacuum-centrifuged. Each sample was fractionated using high pH reverse phase followed by low pH reverse phase separation. Digested proteins (0.1 mg) were loaded using high Performance Liquid Chromatography (Acquity H Class Bio, Waters). Peptides were separated on an XBridge C18 column (3×100mm, Waters) using the following gradient: 3% B (2 min), linear gradient to 40% B (50 min), 5 min to 95% B, maintained at 95% B for 5 minutes and then back to the initial conditions. Peptides were fractionated and each fraction was vacuum dried, then reconstituted in 50 µl 97:3 acetonitrile:water plus 0.1% formic acid, and pooled to six fractions. Each pooled fraction was loaded using split-less nano-Ultra Performance Liquid Chromatography (Ultimate 3000, Thermo Scientific). The mobile phase was: A: water + 0.1% formic acid, and B: acetonitrile + 0.1% formic acid. Desalting of the samples was performed online using a reverse-phase Symmetry C18 trapping column (PepMap, from Thermo Scientific). The peptides were then separated using a T3 HSS nano-column (from Waters). Peptides were eluted from the column into the mass spectrometer using the following gradient: 4% to 27% B (73 min), 27% to 90% B (7 min), maintained at 90% (6 min) and then back to initial conditions. NanoUPLC was used for mass spectrometry, coupled online through a nanoESI emitter (10 μm tip; New Objective; Woburn, MA) to a quadrupole orbitrap mass spectrometer (Exploris 480) using a FlexIon nanospray apparatus (Proxeon). Data was acquired in data dependent acquisition (DDA) mode, using a 2 sec cycle time. MS1 resolution was set at 120,000 (at m/z 200), mass range of m/z 380-1500, normalized AGC target of 200% and maximum injection time was set to 50 msec. MS2 resolution was set at 15,000, quadrupole isolation m/z 1.4, normalized AGC target of 75%, dynamic exclusion of 40 sec and auto mode for maximum injection time. Raw data was processed with MaxQuant v1.6.6.0, analyzed with the Andromeda search engine against the mouse proteome database (www.uniprot.com). Protein quantification was based on unique peptides only, the minimal peptide ratio count was set to 1 and match between runs was enabled. The ProteinGroups file was used for further calculations using Perseus version 1.6.2.3. The LFQ intensities were log transformed and only proteins that had at least 2 valid values in at least one experimental group (67% in at least one group) were kept. The remaining missing values were imputed. Student’s t-test was performed on basis of the LFQ intensities and the fold changes (ratios) between the corresponding samples were calculated.

### Knockout of gasdermin E

Gsdme knockout (KO) cells were generated using a lentivirus approach. For this, HEK293 cells were transfected with a plasmid encoding both Cas9 and the specific guide RNA (from Vector Builders). The plasmid also contained a hygromycin selection marker for mammalian cells. Viral particles were generated in HEK293 cells by co-transfection with two packaging plasmids, Pspax and Pmd2G. Viruses were used to infect 4T1 cells stably overexpressing Arrdc4. Following 24 hours of incubation with the virus, the cells were washed and the medium was changed to a hygromycin-containing medium, which permitted selection of Gsdme KO cells.

### Live microscopy imaging

Cells were seeded in 4 Well Glass Bottom slides (from IBIDI) and incubated overnight. On the next day, cells were analyzed using a CD7 Microscope preset to normal incubating conditions. Different regions of each slide were focused and defined. Then, cells were treated with 5µM raptinal, same points were readjusted, and live imaging was performed for nearly 3 hours using a 40X objective, a GFP channel and oblique frame. Data were analyzed using the Blue Zen software.

### Immunoprecipitation and immunoblotting

Cell lysates were collected in ice cold mild lysis buffer (50 mM HEPES, pH 7.5, 10% glycerol, 150 mM NaCl, 1% Triton X-100, 1 mM EDTA, 1 mM EGTA, 10 mM NaF and 30 mM β-glycerol phosphate). Proteins were immunoprecipitated using beads conjugated to an antibody. After 2 hours of incubation at 4°C on an end to end rotor, complexes were washed three times and bound proteins were eluted in 6X Laemmli sample buffer. Eluates were subjected to electrophoresis and immunoblotting. For immunoblotting, cleared cell lysates were resolved using electrophoresis, followed by electrophoretic transfer to a nitrocellulose membrane. Membranes were blocked in TBS-T solution (tris-buffered saline containing Tween-20) containing 1% low-fat milk, blotted overnight with a primary antibody, washed three times with TBS-T, incubated for 30 minutes with a secondary antibody linked to horseradish peroxidase, and washed once again with TBS-T. Immunoreactive bands were detected using ClarityTM and Clarity Max^TM^ Western-ECL blot substrates (Bio-Rad, Cat:1705061). Blots were imaged using Bio-Rad ChemiDoc^TM^ Imaging system. For pull down with GFP nano-trap Agarose (chromotek, #gta-20) or RFP nano-trap Agarose beads. Cells were transfected with the appropriate GFP/mCherry plasmids for 48 hours, cell lysates were collected in ice-cold lysis buffer (10mM Tris/Cl pH 7.5, 150 mM NaCl, 0.5mM EDTA, 0.5% NP-40) and incubated with GFP-trap or RFP-trap agarose beads overnight at 4°C. Next morning, beads were washed thrice using ice-cold wash buffer (10mM Tris-HCl pH 7.5, 150 mM NaCl, 0.5mM EDTA) and bound proteins were eluted in 6X Laemmli buffer. Eluates were subjected to electrophoresis and immunoblotting.

### Colony formation assays

For anchorage-dependent colony formation assays, 4T1 cells were plated in 60-mm plates at 1000 cells/plate and cultured for 5 days. Colonies were washed twice with saline, fixed in ice-cold methanol (20 min) and stained with 2% crystal violet (30 min). The wells were imaged using an Olympus microscope (Model-SZX2-ILLK) and the CellSens Entry software.

### Transwell migration assays

4T1 cells were trypsinized and suspended in serum-free medium. Approximately 20,000 cells were seeded in transwell inserts (8 µm pore size, Corning, Cat. #3422), that were placed into 24-well cell culture plates containing RPMI with 10% FBS. After incubation at 37°C for 16 hours, non-migrating cells on the inner membrane surface of the insert were removed with cotton swabs and those migrated at the base (outer membrane surface) of the inserts were fixed in 4% PFA (5 min), permeabilized with methanol (20 min) and stained with 2% crystal violet. Cells were imaged using Olympus optical Stereo zoom microscope (Model-SZX2-ILLK) using the CellSens Entry software.

### Cell proliferation assay

Cell proliferation was assessed using MTT (3-(4,5-dimethylthiazol-2-yl)-2,5-diphenyltetrazolium bromide). Cells (1000-3000 per well) were seeded in 96-well plates. Cells were cultured for 24, 48 or 72 hours and afterwards incubated for 3 hours at 37°C with the MTT solution (0.5 mg/ml). The formazan crystals formed by metabolically active cells were dissolved in DMSO and the absorbance was determined at 570 nm.

### Cell cycle and apoptosis assays

Cell cycle assays were performed using the Phase-Flow™ Alexa Fluor 647 BrdU Kit (from BioLegend, Cat. #370706) and analyzed using a BD FACSAria Fusion instrument controlled by BD FACS Diva software v8.0.1 (BD Biosciences). Apoptosis assays were performed using the APC Annexin V Apoptosis Detection Kit with 7-AAD (from BioLegend) and analyzed using a BD FACSAria Fusion instrument.

### Immunofluorescence and determination of membrane localizaion of Glut1

4T1 cells were seeded onto coverslips in 12-well plates. Approximately 20,000 cells/well were plated for immunofluorescence based analyses. For cell morphology experiments, GFP labelled cells were used and kept for 48 hours before fixation. The cells were fixed in 4% paraformaldehyde (PFA) for 20 minutes and washed thrice with saline. DAPI and Phalloidin red were added for 5 minutes and cover slips were again washed 3 times with saline and mounted as described. The quantification of Glut1’s abundance at cell membrane used an adaptation of a previously described protocol ^35^. In brief, 4T1 cells expressing GFP and either shArrdc4, an empty vector or an overexpressed Arrdc4 were seeded on glass coverslip and after 24 hours they where fixed and stained for Glut1 (Abcam ab32551). High-resolution images were taken using an Olympus X01 microscope and images were captured by Hamamtsu Orca v2 camera. Random fields were picked and the corresponding cells analyzed using FIJI ^67^. The cytosolic location of GFP guided the identification of cell edges, and a small area of 2mm around the cell edge was marked as a region of interest (ROI). A mask of the Glut 1 image was created using the built-in threshold function of imageJ and the ROI was superhimposed onto the mask of Glut1. The high intensity signals residing within the ROI were quantified.

### Animal experiments

All animal studies were approved by the Weizmann Institute’s Animal Care and Use Committee (IACUC). BALB/c (#000651) and NOG (#005557) mice were obtained from Jackson laboratory. Balb/c female mice (6 weeks old) were injected in the mammary fat pad with 1.5 million GFP labelled derivatives of 4T1 cells in a 0.1 ml suspension in saline. Alternatively, 1 million cells were injected in the mammary fat pad of 6 weeks old female NOG mice. Tumor volume was estimated using vernier calliper measurements of the longest axis, α/mm, and the perpendicular axis, b/mm. Tumor volume was calculated in accordance with the formula V = (4π/3) x (α/2)2 x (b/2). When tumor volumes reached approximately 1500 mm^3^, mice were euthanized. For tail vein metastasis experiments, 6-weeks old female Balb/c mice were injected with 20,000 cells directly to the tail vein and monitored for 1 month. After mice were sacrificed, lungs were extracted and analyzed for metastases using Olympus microscope for GFP fluorescence.

## Statistical and image analyses

Fiji (NIH doi:10.1038/nmeth.2019) and ZEN Microscopy Software (Zeiss RRID:SCR_013672) were used for image acquisition and analysis. GraphPad Prism v8.0.2 (GraphPad Software RRID:SCR_002798) was used for statistical analysis. Correlation between ARRDC4 expression levels and patient survival time was analyzed in clinical specimens using the Kaplan-Meier method. Data was reported as mean and the corresponding standard deviation (SD). Two-sample comparisons were made using two-sided Student T-test. For all animal experiments we used 2-way ANOVA with Sidak’s or the Dunnett’s multiple comparison test. All statistical analyses were performed using Graph Pad Prism version 6.04 (ns, non-significant; ∗, p ≤ 0.05; ∗∗, p ≤ 0.01; ∗∗∗, p ≤ 0.001; and ∗∗∗∗, p ≤ 0.0001).

## Data and resourse availability

All materials that are not commercially available will be made available to interested readers. Please contact the lead author, Dr. Yosef Yarden.

The data generated in this study are publicly available in Gene Expression Omnibus (GEO).

i. To review GEO accession GSE280819 (scRNAseq): go to https://www.ncbi.nlm.nih.gov/geo/query/acc.cgi?acc=GSE280819 Enter token mvuxyyqaznypjih into the box
ii. To review GEO accession GSE277374 (ARRDC4 RNAseq): go to https://www.ncbi.nlm.nih.gov/geo/query/acc.cgi?acc=GSE277374 Enter token idcpycaovfybbmh into the box

## Supporting information

Proteomic Table

## Acknowledgments

We thank Sharad Kumar (University of South Australia), Carlos E. Alvarez (Nationwide Children’s Hospital, Columbus, Ohio) and Emad Alnemri (Thomas Jefferson University, Philadelphia) for plasmids, and all our lab members for help and comments. This work was performed in the Marvin Tanner Laboratory for Research on Cancer. YY is the incumbent of the Harold and Zelda Goldenberg Professorial Chair in Molecular Cell Biology. Our studies were supported by the Israel Science Foundation, European Research Council (ERC), the Israel Cancer Research Fund (ICRF) and the Dr. Miriam and Sheldon G. Adelson Medical Research Foundation. O.M.R and R.Z. were supported by the NIHR Cambridge Biomedical Research Centre (BRC-1215-20014) and the Medical Research Council (UK; MC_UU_00002/16).

## Author contributions

AV, NBN, AG and YY designed the experiments. AV, SL and YY wrote the manuscript. AV, AG, NBN, MA, BRS, TB, PR, AP, T-MS and BB performed the experiments, YV, YL, EW, DD, RZ, CC, OR and YY analyzed the data.

