## Supplemental Figures for "Avoidance of pyroptosis accounts for the relatively high metastatic potential observed in early hybrid EMT states"

**Supplementary figures with titles and legends**

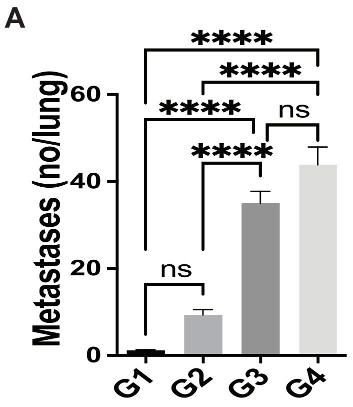


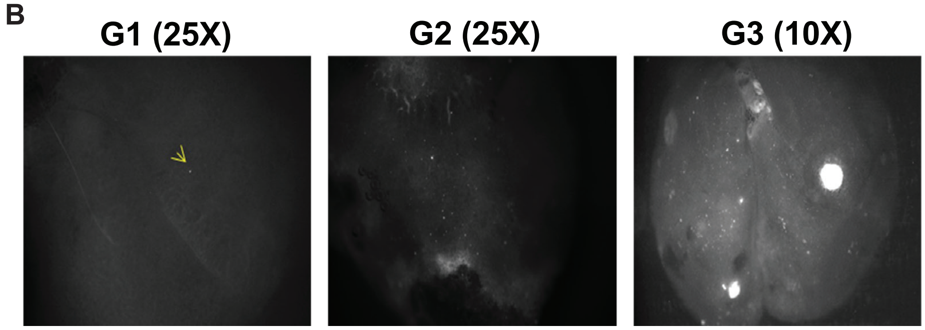


**Figure S1: Quantification of lung metastases post inoculation of isolated CTCs (related to Fig. 1).** (**A**) GFP-labeled 4T1 cells ($1x10^6$) were implanted in the fat pad of groups (n=6) of 6-weeks old female BALB/c mice and CTCs were harvested from blood 5 days later. CTCs isolated from several animals were pooled together and cells ($2x10^4$ per animal) were injected in the tail vein of naïve mice. This process was repeated to generate G1 through G4 (fresh CTCs). Shown are bar graphs depicting the numbers of resulting metastatic lung nodules 5 days later across the different generations of fresh CTCs (G1 through G4). ****, $p< 0.0001$; ns, non-significant. (**B**) Representative images of lung metastases that emerged post injection of G1, G2 and G3 cells ($2x10^4$) in the tail vein of 6-weeks old female BALB/c mice. The optic magnifications (either 10X or 25X) are indicated.

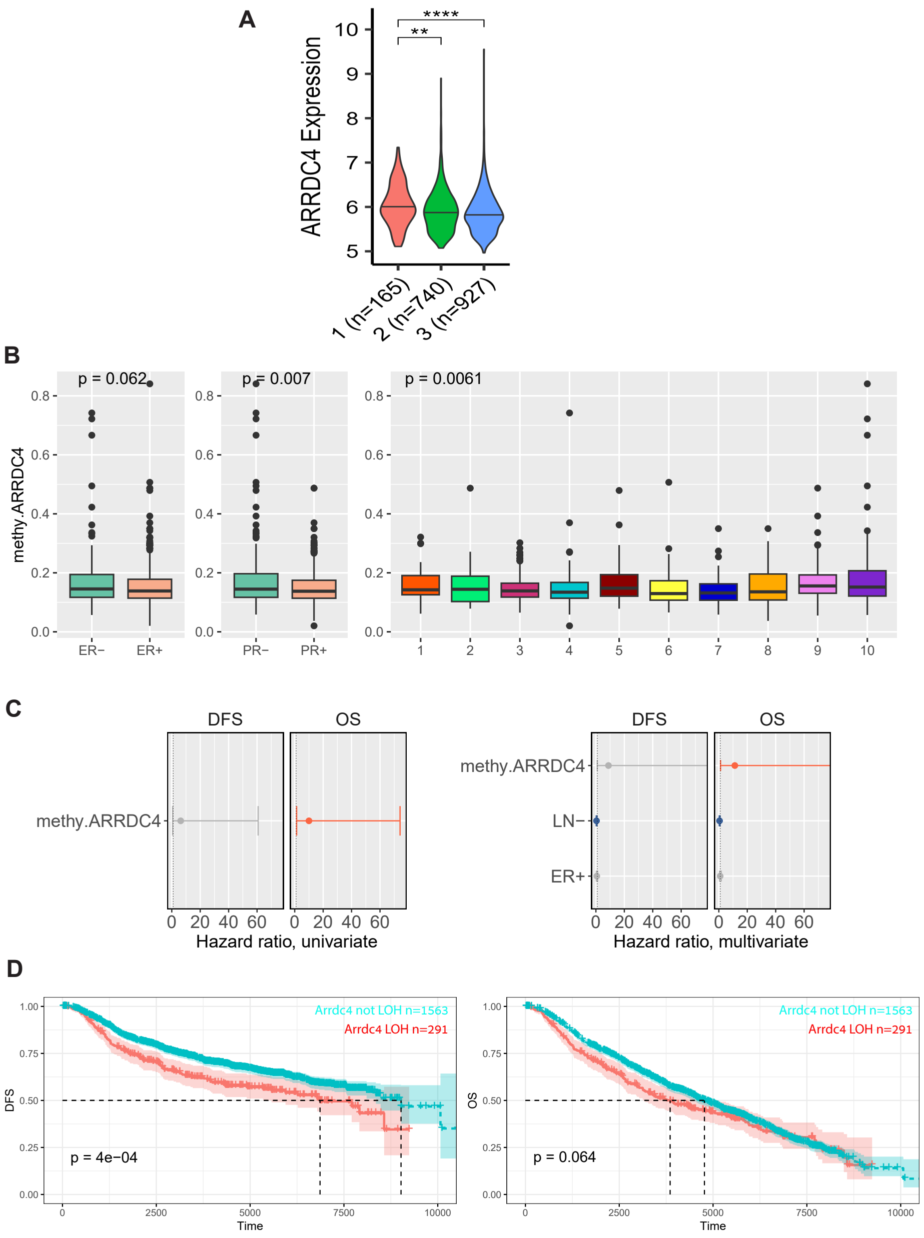


**Figure S2: Expression levels of ARRDC4 in breast cancer and prognostic values of the corresponding epigenetic and chromosomal aberrations (related to Fig. 2).** (**A**) Plot showing relevance of ARRDC4 expression to histological grade of breast tumors based on the METABRIC dataset. The level of significance is shown as follows: **, p<0.01 and ****, p<0.0001. (**B**) Box plots presenting the methylation level of *ARRDC4* according to ER and PR levels (left panels) and across the 10 subtypes of breast cancer (right panel; TCGA dataset). P-values are indicated. (**C**) Univariate and multivariate analyses of the hazard ratio of *ARRDC4*'s methylation (methy.ARRDC4; TCGA dataset). Note that according to both analyses, the methylation of *ARRDC4* can serve as a poor prognosis marker for patient survival time. OS, overall survival, DFS, disease free survival. (**D**) ARRDC4's mCN (minor copy number) was used to categorise chromosomal events as LOH or not LOH. The results of this analysis identified 291 cases, out of 1864 patients, as LOH. Disease free survival and overall survival of the two groups of patients are presented in Kaplan-Meier plots.

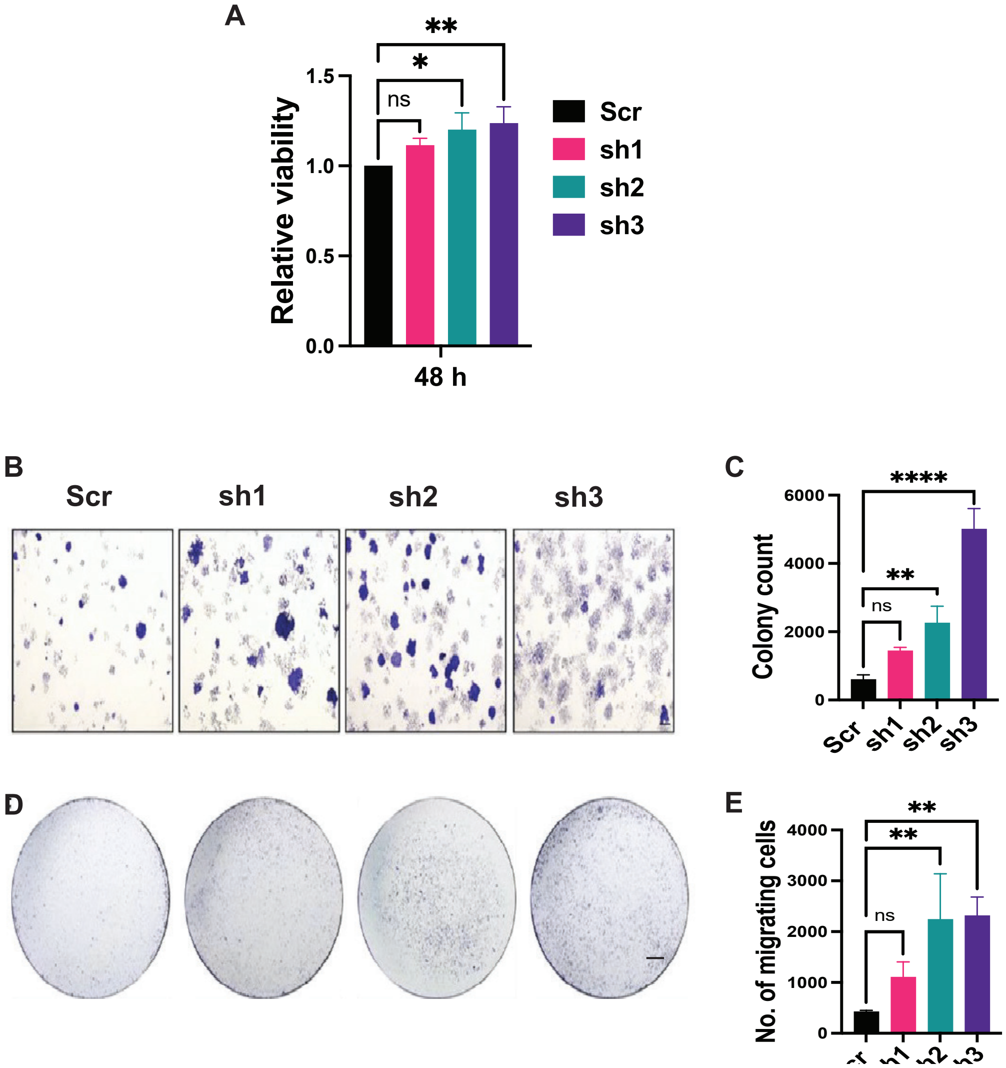


**Figure S3: Arrdc4 knockdown increases cell proliferation and migration (related to Fig. 3).** (**A**) Clones of 4T1 cells stably expressing the indicated ARRDC4-spedific shRNAs, or a control (Scr) shRNA, were seeded in 96-well plates and incubated for 48 hours, prior to a cell proliferation assay that used 3-(4,5-dimethylthiazol-2-yl)-2,5-diphenyltetrazolium bromide (MTT; n=4). (**B** and **C**) Representative images (B) showing colonies formed by control 4T1 cells (Scr) and by the indicated shRNA-expressing derivatives. Cells (1000 per well) were incubated for 5 days prior to photography and colony number quantification (n=4; panel C). Scale, 10 micrometers. (**D** and **E**) Representative images reflecting the migration potential of 4T1 derivative lines across transwell chambers containing media and serum in the lower compartment. Cells were placed in the upper compartment and allowed to migrate for 16 hours. Also shown are bar graphs (panel E) quantifying migration across the filter. The assay was performed in quadruplicates and the levels of significance are presented as follows: *, p<0.05; **, p<0.01; ***, p<0.001 and ****, p<0.0001, ns, non-significant. Scale bar, 500 micrometer.

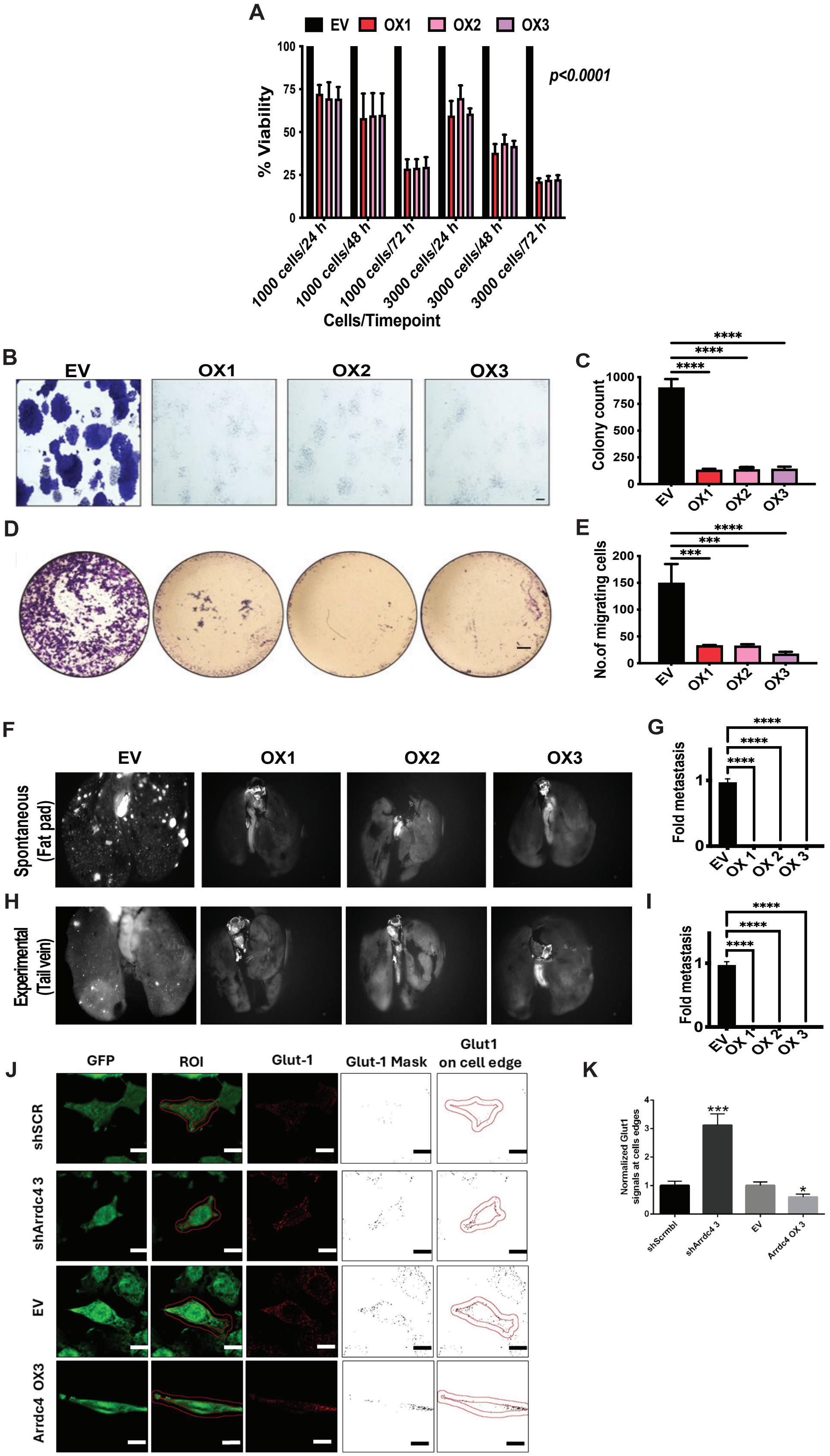
A
EV
OX1
OX2
OX3
% Viability
p<0.0001
1000 cells/24 h
1000 cells/48 h
1000 cells/72 h
3000 cells/24 h
3000 cells/48 h
3000 cells/72 h
Cells/Timepoint
B
C
Colony count
D
E
No.of migrating cells
F
G
Spontaneous (Fat pad)
Fold metastasis
H
I
Experimental (Tail vein)
J
GFP
ROI
Glut-1
Glut-1 Mask
Glut1 on cell edge
shSCR
shArrdc4 3
Arrdc4 OX3
K
Normalized Glut1 signals at cells edges
shScrmbl
shArrdc4 3
Arrdc4 OX 3

**Figure S4: Arrdc4 overexpression in 4T1 cells reduces proliferation in vitro, as well as inhibits metastasis in immunocompetent mice (related to Fig. 4).** (**A**) The indicated derivatives of 4T1 cells, including the empty vector (EV) control cells and the three overexpressing (OX1, OX2 and OX3) clones, were plated in 96-well plates at different densities (1000-3000 cells per well). Following incubation for different time intervals (24-72 hours), cell viability was determined in triplicates using MTT. (**B** and **C**) Colony forming potential of the indicated OX clones was analyzed by seeding 1000 cells per well and culturing the cells for 5 days. Shown are representative photographs (B) and a bar graph (C) presenting quantitation of colonies in quadruplicates (4 fields per well). Bar, 10 micrometers. (**D** and **E**) Control and OX clones were analyzed for migration using transwell chambers (n=4) containing media with serum in the lower compartments. Cells were incubated for 16 hours prior to photography (D). The bar graph presents quantification of cells that migrated across the separating filter. Scale bar, 500 micrometers. (**F** and **G**) Representative images (10X magnification) showing events of spontaneous lung metastases in mice pre-injected in the fat pads with control (EV) and Arrdc4-overexpressing (OX) 4T1 cells ($1.5x10^6$ per animal). Metastasis was monitored on day 95, photos were captured and metastases were quantified. (**H** and **I**) Representative images showing lung metastases (10X magnification) after 30 days in mice injected in the tail vein with EV or with the indicated OX cells ($2x10^4$ per mouse). The bar graph presents quantification of the corresponding lung metastases. (**J**) Shown are images of GFP-positive 4T1 cells stably expressing an ectopic Arrdc4 (or shArrdc4) along with control lines expressing an empty vector or a scrambled shRNA. Cells were immunostained for Glut1 and later the abundance of Glut1 at the cell surface was determined using an established protocol that focused on a specific region of interest (ROI; i.e., the cell surface). Scale bar 10 micrometers. (**K**) The bar graph represents the Glut1 positive signals in the ROI after normalization to the respective control. Twenty cells of each subline were analyzed. The levels of significance in all panels are shown as follows: *, p<0.05 and ***, p<0.001.

**A**

| S.No. | Protein name | Gene name | Peptides | Unique peptides |
|---|---|---|---|---|
| 1 | Insulin receptor substrate 4 | IRS4 | 47 | 47 |
| 2 | Golgin subfamily A member 4 | GOLGA4 | 42 | 42 |
| 3 | MICOS complex subunit MIC60 | IMMT | 36 | 36 |
| 4 | Sarcoplasmic/endoplasmic reticulum calcium ATPase 2 | ATP2A2 | 32 | 32 |
| 5 | Eukaryotic translation initiation factor 4 gamma 1 | EIF4G1 | 34 | 29 |
| 6 | Cytoskeleton-associated protein 4 | CKAP4 | 27 | 27 |
| 7 | Myb-binding protein 1A | MYBBP1A | 26 | 26 |
| 8 | Importin-5 | IPO5 | 26 | 24 |
| 9 | Zinc finger CCCH-type antiviral protein 1 | ZC3HAV1 | 22 | 22 |
| 10 | Golgin subfamily B member 1 | GOLGB1 | 22 | 21 |
| 11 | Kinase D-interacting substrate of 220 kDa | KIDINS220 | 20 | 20 |
| 12 | Large proline-rich protein BAG6 | BAG6 | 20 | 20 |
| 13 | E3 ubiquitin-protein ligase Itchy homolog | ITCH | 21 | 18 |
| 14 | Polyadenylate-binding protein 1;Polyadenylate-binding protein 3 | PABPC1;PABPC3 | 27 | 18 |
| 16 | DnaJ homolog subfamily A member 1 | DNAJA1 | 16 | 16 |
| 17 | Importin-4 | IPO4 | 16 | 16 |
| 18 | Inactive tyrosine-protein kinase 7 | PTK7 | 16 | 16 |
| 19 | Desmoglein-2 | DSG2 | 16 | 16 |
| 20 | Serine/threonine-protein kinase mTOR | MTOR | 15 | 15 |

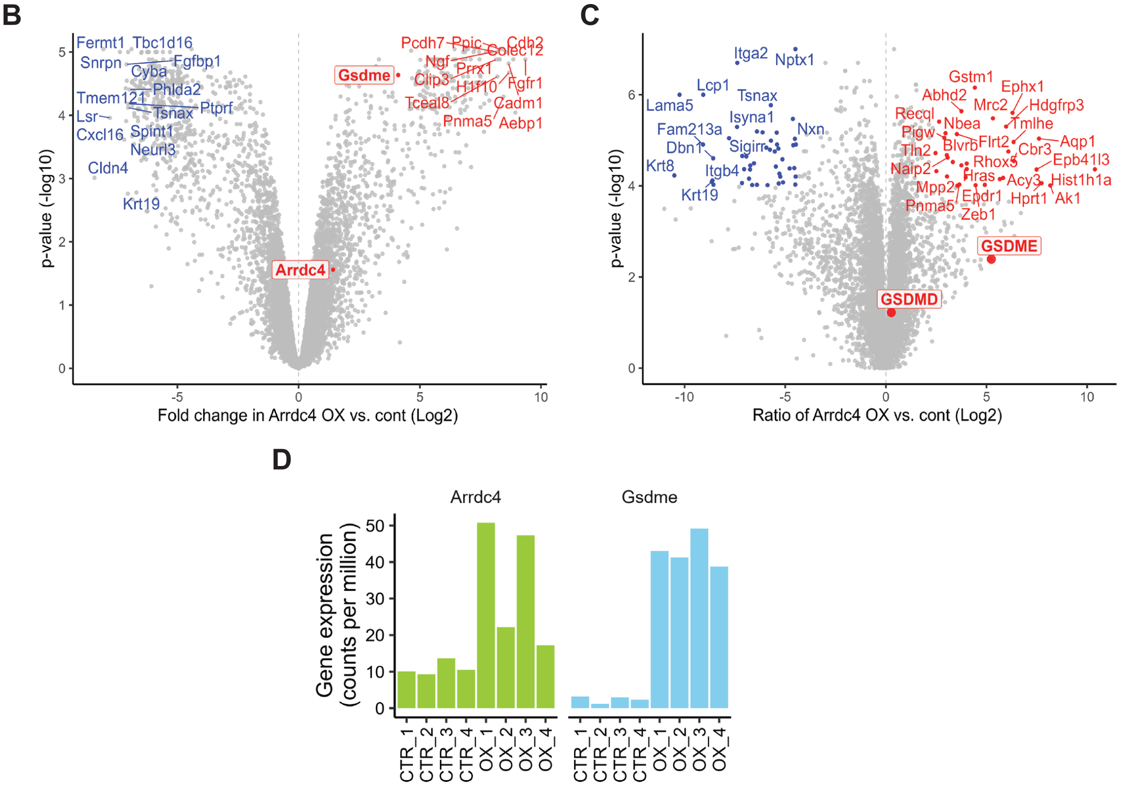


**Figure S5: RNA sequencing and deep proteomics analyses uncover candidates potentially involved in the phenotype associated with Arrdc4 overexpression (related to Fig. 5).** (**A**) Listed are the top 20 physically interacting partners of Arrdc4 based on the number of unique peptides determined following digestion of pulled-down interactors. Both protein names and gene names are indicated. (**B**) Shown is a volcano plot depicting the most upregulated (red) and downregulated (blue) transcripts identified using

RNA sequencing of 4T1 cells overexpressing Arrdc4 versus control cells (n=4 per group). Analysis was performed using Limma in R. (**C**) Volcano plot showing differentially expressed proteins (identified using deep proteomics analysis) comparing 4T1 cells overexpressing Arrdc4 versus control cells (n=3 per group). For each protein, we calculated the abundance ratio between the Arrdc4-overexpressing samples versus the control samples. Shown are the proteins with the highest (red) and lowest (blue) ratios. (**D**) Bar graphs demonstrating similarity between expression levels of Gsdme and Arrdc4 based on RNA-sequencing analyses of the indicated cell lines (CTR, empty vector control; OX, Arrdc4-overexpressing cells). Data are presented in counts per million and they were calculated using the edgeR package in R.

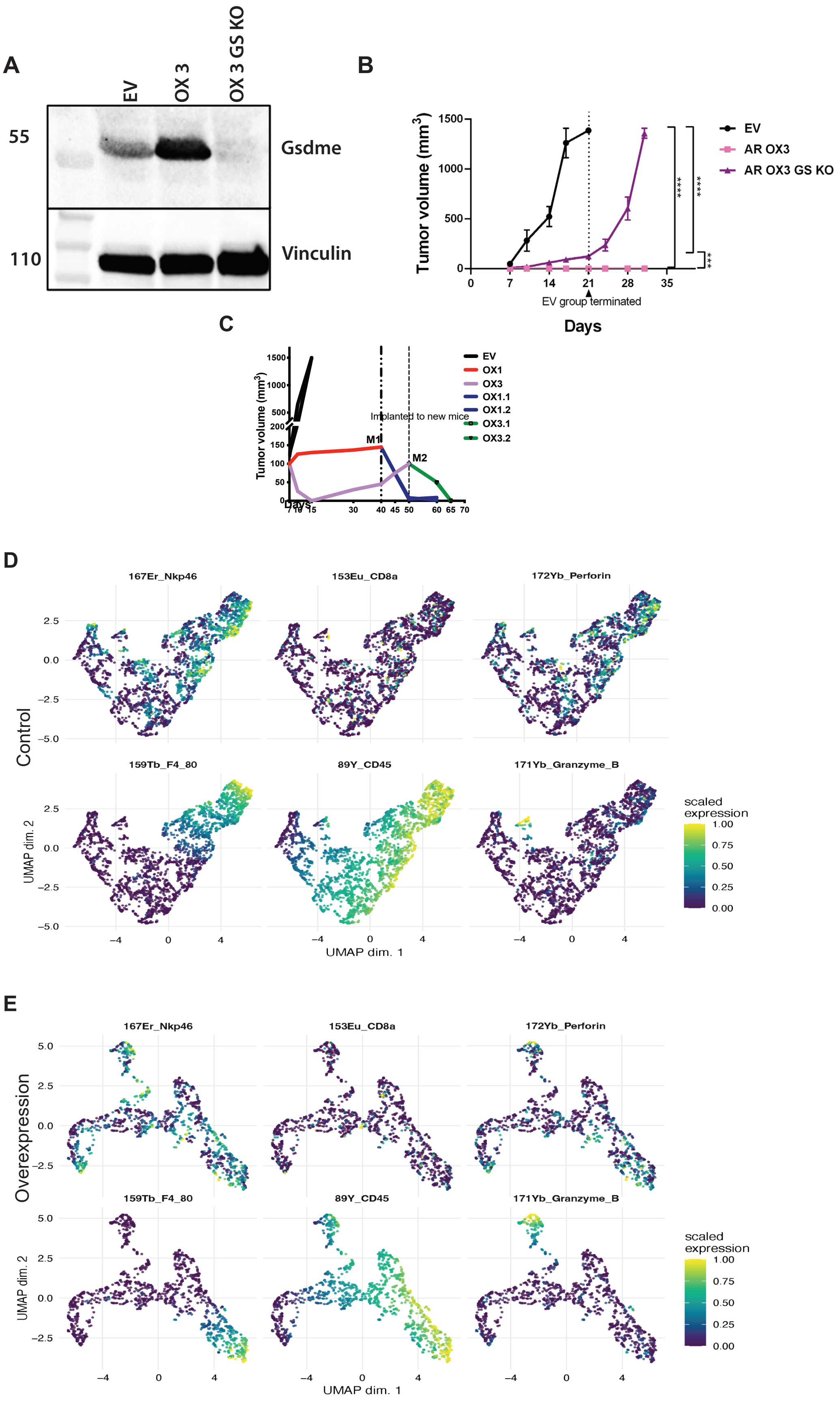
A
EV
OX 3
OX 3 GS KO
55
110
Gsdme
Vinculin
B
Tumor volume (mm3)
1500
1000
500
0
0
7
14
21
28
35
EV group terminated
Days
EV
AR OX3
AR OX3 GS KO
****
****
***
C
Tumor volume (mm3)
1500
1000
500
200
150
100
50
0
Days
7
10
15
30
40
45
50
60
65
70
EV
OX1
OX3
OX1.1
OX1.2
OX3.1
OX3.2
Implanted to new mice
M1
M2
D
Control
167Er_Nkp46
153Eu_CD8a
172Yb_Perforin
159Tb_F4_80
89Y_CD45
171Yb_Granzyme_B
UMAP dim. 2
UMAP dim. 1
2.5
0.0
−2.5
−5.0
−4
0
4
scaled expression
1.00
0.75
0.50
0.25
0.00
E
Overexpression
167Er_Nkp46
153Eu_CD8a
172Yb_Perforin
159Tb_F4_80
89Y_CD45
171Yb_Granzyme_B
UMAP dim. 2
UMAP dim. 1
5.0
2.5
0.0
−2.5
−4
0
4
scaled expression
1.00
0.75
0.50
0.25
0.00

**Figure S6: CyTOF analyses of transplanted tumors and in vivo tests of Gasdermin E knockout cells (related to Fig. 6).** (**A**) Extracts were prepared from 4T1 cells stably overexpressing either Arrdc4 (clone OX3), the respective control (EV) or the OX3 cells that are devoid of Gsdme (OX-3 GSKO). Following electrophoresis and transfer, the blot was probed for Gsdme and vinculin. (**B**) The following derivatives of 4T1 cells ($1.5x10^6$ cells per mouse) were implanted in the fat pads of Balb/c mice (6 animals per group): control cells (EV), Arrdc4-overexpressing cells (AR OX3) and the same cell line but lacking expression of Gsdme (AR OX3-GS KO). Tumor volumes were monitored for 30 days. (**C**) Line graphs showing the volumes of two Arrdc4-overexpressing tumors, M1 and M2, which did not completely regress when implanted in Balb/c mice (see Fig. 6D). Note that the corresponding tumors completely regressed later. (**D** and **E**) Control 4T1 (EV) and Arrdc4 OX tumors were transferred from NOG to Balb/c mice, but they were resected 10 days later. Single cell suspensions were prepared and the cells were subjected to CyTOF analysis after barcoding and incubating cells with heavy metal conjugated primary antibodies targeting anti-CD45, CD8a, F4/80 and Nkp46. In addition, we probed for two markers of pyroptosis, granzyme B and perforin. The analysis used a dimension reduction visualization method, along with the indicated pyroptotic markers, to resolve distinct cell clusters.

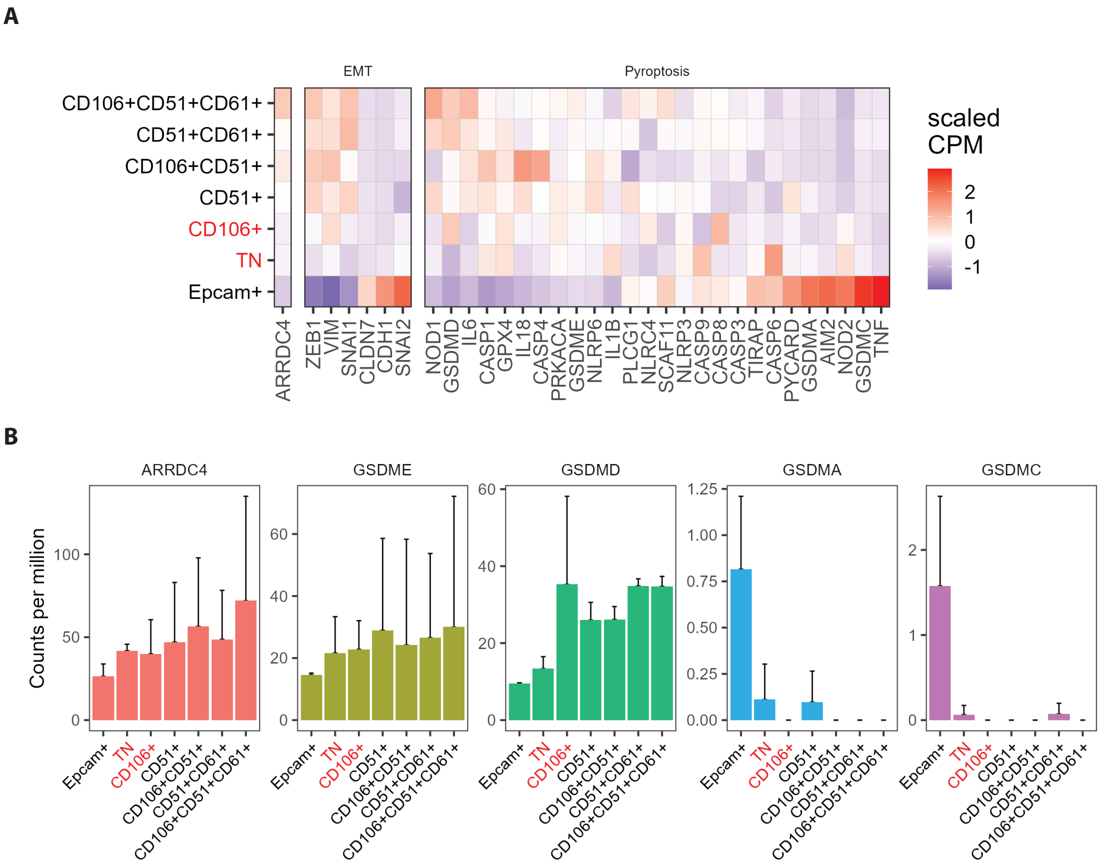


**Figure S7: Bioinformatic analyses support functional links between ARRDC4, GSDME and partial EMT (related to Fig. 7).** (**A**) Heatmaps tabluating mRNA expression levels of the indicated EMT (left part) and pyroptosis (right part) markers. The vertical axis lists the stable EMT transition states, alongside the fully epithelial and the fully mesenchymal states. In addition to ARRDC4's transcript levels, the scheme displays several main markers of EMT (Snail, SNAI1; Zinc Finger E-Box Binding Homeobox 1, ZEB1; Vimentin, VIM; SLUG/Snail2, SNAI2; Claudin 7, CLDN7 and E-cadherin, CDH1). The Pyroptosis box refers to previously identified markers of pyroptosis (see main text). The GSE110587 dataset was used to derive the indicated transcript levels. (**B**) Shown are the expression levels of ARRDC4 (in counts per million, CPM), as well as four additional gasdermins (GSDMA, GSDMC, GSDMD and GSDME/DFNA5), in the reported stable EMT states. TN refers to a triple negative state lacking expression of three cell adhesion molecules: CD51, CD61 and CD106, along with EpCAM. The data were derived from the previously reported GSE110587 dataset. Error bars refer to standad error values (n=3). Note that for comparison of all 4 gasdermins, the data for ARRDC4 are identical to data shown in main Figure 7.
