## Supplementary material for "Avoidance of pyroptosis accounts for the relatively high metastatic potential observed in early hybrid EMT states": Proteomic Table

| | | | | | | | |
|---|---|---|---|---|---|---|---|
| Protein IDs | Majority protein IDs | Protein names | Gene names | Fasta headers | Peptides | Unique peptides | Sequence coverage [%] |
| Q91W50 | Q91W50 | Cold shock | Csde1 | sp\|Q91W5 | 42 | 42 | 55,8 |
| Q62443 | Q62443 | Neuronal p | Nptx1 | sp\|Q62443 | 8 | 8 | 22,7 |
| Q8VCN5 | Q8VCN5 | Cystathion | Cth | sp\|Q8VCN | 13 | 13 | 50 |
| Q62469 | Q62469 | Integrin alp | Itga2 | sp\|Q62469 | 32 | 32 | 35,3 |
| P14824 | P14824 | Annexin A6 | Anxa6 | sp\|P14824 | 50 | 49 | 75,2 |
| O08709 | O08709 | Peroxiredo | Prdx6 | sp\|O08709 | 22 | 22 | 75,9 |
| Q9Z0U1;Q | Q9Z0U1 | Tight junct | Tjp2 | sp\|Q9Z0U1 | 36 | 36 | 42,8 |
| Q8K2B3 | Q8K2B3 | Succinate d | Sdha | sp\|Q8K2B3 | 29 | 29 | 56,8 |
| Q80TM9 | Q80TM9 | Nischarin | Nisch | sp\|Q80TM | 35 | 35 | 31 |
| Q8CI15 | Q8CI15 | Sphingosin | Sphk1 | sp\|Q8CI15 | 7 | 7 | 20,9 |
| P52196 | P52196 | Thiosulfate | Tst | sp\|P52196 | 10 | 10 | 40,4 |
| Q9Z0J0 | Q9Z0J0 | Epididyma | Npc2 | sp\|Q9Z0J0 | 8 | 8 | 50,3 |
| Q8C761 | Q8C761 | WD repeat | Wdr60 | sp\|Q8C761 | 16 | 16 | 23,2 |
| P10649 | P10649 | Glutathion | Gstm1 | sp\|P10649 | 14 | 8 | 62,8 |
| Q8BIK4 | Q8BIK4 | Dedicator o | Dock9 | sp\|Q8BIK4 | 48 | 45 | 30,9 |
| Q5PSV9 | Q5PSV9 | Mediator o | Mdc1 | sp\|Q5PSV9 | 45 | 45 | 46,5 |
| O08912;Q8 | O08912 | Polypeptid | Galnt1 | sp\|O08912 | 17 | 17 | 37 |
| Q9D8S9 | Q9D8S9 | BolA-like p | Bola1 | sp\|Q9D8S9 | 6 | 6 | 56,9 |
| Q8BH64 | Q8BH64 | EH domain | Ehd2 | sp\|Q8BH64 | 30 | 28 | 69,8 |
| Q61233 | Q61233 | Plastin-2 | Lcp1 | sp\|Q61233 | 39 | 34 | 71,6 |
| Q61001 | Q61001 | Laminin su | Lama5 | sp\|Q61001 | 68 | 68 | 31,8 |
| P33609 | P33609 | DNA polym | Pola1 | sp\|P33609 | 45 | 45 | 40,7 |
| P11440;Q3 | P11440 | Cyclin-dep | Cdk1 | sp\|P11440 | 20 | 18 | 67,7 |
| Q8C522 | Q8C522 | Endonucle | Endod1 | sp\|Q8C522 | 13 | 13 | 31,7 |
| Q3THK3 | Q3THK3 | General tra | Gtf2f1 | sp\|Q3THK3 | 15 | 15 | 30,7 |
| Q9QZE7 | Q9QZE7 | Translin-as | Tsnax | sp\|Q9QZE7 | 11 | 11 | 47,9 |
| Q9QZM4 | Q9QZM4 | Tumor nec | Tnfrsf10b | sp\|Q9QZM | 6 | 6 | 24,7 |
| P48722 | P48722 | Heat shock | Hspa4l | sp\|P48722 | 37 | 35 | 59,4 |
| P61028 | P61028 | Ras-related | Rab8b | sp\|P61028 | 7 | 1 | 33,3 |
| Q8VD75 | Q8VD75 | Huntingtin | Hip1 | sp\|Q8VD75 | 34 | 32 | 43,2 |
| P53568 | P53568 | CCAAT/enh | Cebpg | sp\|P53568 | 1 | 1 | 20,7 |
| P26040 | P26040 | Ezrin | Ezr | sp\|P26040 | 35 | 24 | 57,8 |
| Q9WTI7 | Q9WTI7 | Unconvent | Myo1c | sp\|Q9WTI7 | 51 | 50 | 54,5 |
| P70677 | P70677 | Caspase-3; | Casp3 | sp\|P70677 | 19 | 19 | 70 |
| Q9QXM0 | Q9QXM0 | Abhydrolas | Abhd2 | sp\|Q9QXM | 10 | 10 | 33,6 |
| Q9CQ45 | Q9CQ45 | Neudesin | Nenf | sp\|Q9CQ4 | 8 | 8 | 63,2 |
| CON__ENS | CON__ENSEMBL:ENSBTAP00000024146 | | | | 61 | 58 | 55,6 |
| Q9D379 | Q9D379 | Epoxide hy | Ephx1 | sp\|Q9D379 | 27 | 27 | 61,8 |
| O08663 | O08663 | Methionin | Metap2 | sp\|O08663 | 15 | 15 | 44,8 |
| P09103 | P09103 | Protein dis | P4hb | sp\|P09103 | 41 | 41 | 80,2 |
| P02468;Q9 | P02468 | Laminin su | Lamc1 | sp\|P02468 | 59 | 59 | 49,8 |
| Q58NB6 | Q58NB6 | Dehydroge | Dhrs9 | sp\|Q58NB | 10 | 10 | 34,8 |
| Q64449 | Q64449 | C-type ma | Mrc2 | sp\|Q64449 | 20 | 20 | 22,2 |
| P97346 | P97346 | Nucleored | Nxn | sp\|P97346 | 17 | 17 | 51 |
| Q6ZWR6 | Q6ZWR6 | Nesprin-1 | Syne1 | sp\|Q6ZWR | 84 | 82 | 13,3 |
| P07214 | P07214 | SPARC | Sparc | sp\|P07214 | 14 | 14 | 48,7 |

| | | | | | | | |
|---|---|---|---|---|---|---|---|
| P37913 | P37913 | DNA ligase | Lig1 | sp\|P37913 | 46 | 46 | 53,7 |
| Q8CIH5 | Q8CIH5 | 1-phospha | Plcg2 | sp\|Q8CIH5 | 35 | 34 | 41,6 |
| Q9Z129 | Q9Z129 | ATP-depen | Recql | sp\|Q9Z129 | 13 | 13 | 27,5 |
| Q11011 | Q11011 | Puromycin | Npepps | sp\|Q11011 | 41 | 41 | 56,5 |
| Q61210 | Q61210 | Rho guanir | Arhgef1 | sp\|Q61210 | 36 | 36 | 46,8 |
| P28474 | P28474 | Alcohol de | Adh5 | sp\|P28474 | 16 | 14 | 58,6 |
| P70697 | P70697 | Uroporphy | Urod | sp\|P70697 | 14 | 14 | 46,9 |
| Q9QWY8 | Q9QWY8 | Arf-GAP wi | Asap1 | sp\|Q9QWY | 24 | 21 | 26,7 |
| Q6DIC0 | Q6DIC0 | Probable g | Smarca2 | sp\|Q6DIC0 | 33 | 20 | 25,2 |
| Q8BWY9 | Q8BWY9 | Protein CIF | Kiaa1524 | sp\|Q8BWY | 26 | 26 | 33,3 |
| Q148V8 | Q148V8 | Protein FA | Fam83h | sp\|Q148V8 | 18 | 18 | 22,3 |
| Q5SSL4 | Q5SSL4 | Active brea | Abr | sp\|Q5SSL4 | 18 | 17 | 32,5 |
| Q9JMG7 | Q9JMG7 | Hepatoma | Hdgfrp3 | sp\|Q9JMG | 5 | 3 | 28,2 |
| Q9JHU9 | Q9JHU9 | Inositol-3-p | Isyna1 | sp\|Q9JHU9 | 18 | 18 | 49,4 |
| Q60790 | Q60790 | Ras GTPase | Rasa3 | sp\|Q60790 | 34 | 34 | 54 |
| Q6PGC1 | Q6PGC1 | ATP-depen | Dhx29 | sp\|Q6PGC | 38 | 37 | 35,2 |
| Q8BH86 | Q8BH86 | UPF0317 protein C14c | | sp\|Q8BH8 | 17 | 17 | 38,6 |
| Q9QZF2 | Q9QZF2 | Glypican-1 | Gpc1 | sp\|Q9QZF2 | 17 | 17 | 43,8 |
| P47739 | P47739 | Aldehyde c | Aldh3a1 | sp\|P47739 | 15 | 13 | 35,5 |
| P54923 | P54923 | [Protein AD | Adprh | sp\|P54923 | 12 | 12 | 49,2 |
| Q921H8;Q | Q921H8;Q8VC | 3-ketoacyl- | Acaa1a;Ac | sp\|Q921H8 | 17 | 17 | 56,4 |
| Q9R112 | Q9R112 | Sulfide:qui | Sqrdl | sp\|Q9R112 | 21 | 21 | 50 |
| Q3UTJ2 | Q3UTJ2 | Sorbin and | Sorbs2 | sp\|Q3UTJ2 | 18 | 17 | 20,3 |
| A2ALU4 | A2ALU4 | Protein Shr | Shroom2 | sp\|A2ALU4 | 8 | 8 | 7,8 |
| Q5JCS9 | Q5JCS9 | N-acetyllac | B3gnt3 | sp\|Q5JCS9 | 3 | 3 | 8,3 |
| P58242 | P58242 | Acid sphing | Smpdl3b | sp\|P58242 | 11 | 11 | 38,8 |
| Q60716 | Q60716 | Prolyl 4-hy | P4ha2 | sp\|Q60716 | 11 | 11 | 28,7 |
| Q6ZQ73 | Q6ZQ73 | Cullin-asso | Cand2 | sp\|Q6ZQ73 | 24 | 23 | 27,4 |
| Q91V61 | Q91V61 | Sideroflexi | Sfxn3 | sp\|Q91V61 | 13 | 12 | 52,6 |
| Q05BC3 | Q05BC3 | Echinoderr | Eml1 | sp\|Q05BC3 | 20 | 19 | 28,3 |
| Q9EPN1 | Q9EPN1 | Neurobeac | Nbea | sp\|Q9EPN1 | 27 | 22 | 12,9 |
| Q8BLU0 | Q8BLU0 | | Flrt2 | sp\|Q8BLU0 | 7 | 7 | 16,1 |
| Q8K368 | Q8K368 | Fanconi an | Fanci | sp\|Q8K368 | 32 | 31 | 29,3 |
| O35280 | O35280 | Serine/thre | Chek1 | sp\|O35280 | 13 | 13 | 28,6 |
| Q9CU65 | Q9CU65 | Zinc finger | Zmym2 | sp\|Q9CU65 | 29 | 27 | 28,3 |
| P97494 | P97494 | Glutamate | Gclc | sp\|P97494 | 25 | 25 | 49 |
| P00375 | P00375 | Dihydrofol | Dhfr | sp\|P00375 | 10 | 10 | 67,4 |
| Q9JLJ2 | Q9JLJ2 | 4-trimethy | Aldh9a1 | sp\|Q9JLJ2 | 21 | 21 | 57,7 |
| Q8QZT1 | Q8QZT1 | Acetyl-CoA | Acat1 | sp\|Q8QZT1 | 18 | 18 | 53,8 |
| Q08093 | Q08093 | Calponin-2 | Cnn2 | sp\|Q08093 | 12 | 11 | 62,6 |
| Q9DCN2 | Q9DCN2 | NADH-cytc | Cyb5r3 | sp\|Q9DCN | 24 | 24 | 66,4 |
| Q8K3H0 | Q8K3H0 | DCC-intera | Appl1 | sp\|Q8K3H0 | 25 | 25 | 49,6 |
| Q9D0B0 | Q9D0B0 | Serine/arg | Srsf9 | sp\|Q9D0B0 | 15 | 15 | 53,6 |
| Q571I9 | Q571I9 | Aldehyde c | Aldh16a1 | sp\|Q571I9 | 9 | 9 | 16,6 |
| E9PZM4 | E9PZM4 | Chromodo | Chd2 | sp\|E9PZM4 | 32 | 26 | 23 |
| Q8R0F8 | Q8R0F8 | Acylpyruva | Fahd1 | sp\|Q8R0F8 | 7 | 7 | 41 |
| Q8C398 | Q8C398 | Phosphatic | Pigw | sp\|Q8C398 | 5 | 5 | 10,5 |
| Q9JLZ8 | Q9JLZ8 | Single Ig IL | Sigirr | sp\|Q9JLZ8 | 16 | 16 | 50,6 |
| Q3TJD7 | Q3TJD7 | PDZ and LI | Pdlim7 | sp\|Q3TJD7 | 15 | 15 | 46 |
| Q02013 | Q02013 | Aquaporin- | Aqp1 | sp\|Q02013 | 4 | 4 | 17,1 |

| | | | | | | | |
|---|---|---|---|---|---|---|---|
| Q80XI3 | Q80XI3 | Eukaryotic | Eif4g3 | sp\|Q80XI3 | 26 | 22 | 23,4 |
| P37889 | P37889 | Fibulin-2 | Fbln2 | sp\|P37889 | 21 | 21 | 31 |
| O35640 | O35640 | Annexin A8 | Anxa8 | sp\|O35640 | 14 | 14 | 59 |
| Q07417 | Q07417 | Short-chai | Acads | sp\|Q07417 | 18 | 18 | 58,3 |
| P11352 | P11352 | Glutathion | Gpx1 | sp\|P11352 | 14 | 14 | 79,6 |
| Q8VEN2 | Q8VEN2 | Placenta-e | Plet1 | sp\|Q8VEN | 2 | 2 | 7,6 |
| Q91ZE0 | Q91ZE0 | Trimethylly | Tmlhe | sp\|Q91ZE0 | 15 | 15 | 41,6 |
| Q91W89 | Q91W89 | Alpha-man | Man2c1 | sp\|Q91W8 | 17 | 17 | 23,7 |
| Q9D2Y4 | Q9D2Y4 | Mixed line | Mlkl | sp\|Q9D2Y4 | 21 | 21 | 38,8 |
| Q62261 | Q62261 | Spectrin be | Sptbn1 | sp\|Q62261 | 123 | 120 | 64,6 |
| Q99JR1 | Q99JR1 | Sideroflexi | Sfxn1 | sp\|Q99JR1 | 8 | 7 | 34,8 |
| Q61333 | Q61333 | Tumor nec | Tnfaip2 | sp\|Q61333 | 18 | 18 | 33,6 |
| Q9ESC8 | Q9ESC8 | AF4/FMR2 | Aff4 | sp\|Q9ESC8 | 17 | 16 | 20,7 |
| P06745;CC | P06745 | Glucose-6- | Gpi | sp\|P06745 | 24 | 24 | 47,5 |
| Q9EPL8 | Q9EPL8 | Importin-7 | Ipo7 | sp\|Q9EPL8 | 33 | 32 | 40,7 |
| Q99J08 | Q99J08 | SEC14-like | Sec14l2 | sp\|Q99J08 | 6 | 6 | 17,9 |
| Q9CYH2 | Q9CYH2 | Redox-regu | Fam213a | sp\|Q9CYH | 11 | 11 | 43,1 |
| P52795 | P52795 | Ephrin-B1 | Efnb1 | sp\|P52795 | 9 | 8 | 44,6 |
| Q8R4N0 | Q8R4N0 | Citrate lyas | Clybl | sp\|Q8R4N0 | 10 | 10 | 39,3 |
| B2RUP2 | B2RUP2 | Protein un | Unc13d | sp\|B2RUP2 | 17 | 17 | 23,6 |
| Q80SU7 | Q80SU7 | Interferon- | Gvin1 | sp\|Q80SU7 | 13 | 13 | 5,9 |
| Q9DBR0 | Q9DBR0 | A-kinase ar | Akap8 | sp\|Q9DBR0 | 19 | 19 | 46,4 |
| P29391;P4 | P29391;P4994 | Ferritin ligh | Ftl1;Ftl2 | sp\|P29391 | 8 | 8 | 59,6 |
| Q8BYY4 | Q8BYY4 | Tetratricop | Ttc39b | sp\|Q8BYY4 | 14 | 14 | 31,6 |
| Q78IQ7 | Q78IQ7 | Zinc transp | Slc39a4 | sp\|Q78IQ7 | 6 | 6 | 15,8 |
| O08579 | O08579 | Emerin | Emd | sp\|O08579 | 15 | 15 | 78,8 |
| Q8K363 | Q8K363 | ATP-depen | Ddx18 | sp\|Q8K363 | 27 | 27 | 51,4 |
| Q6NSR8 | Q6NSR8 | Probable a | Npepl1 | sp\|Q6NSR8 | 11 | 11 | 34 |
| Q91VI7 | Q91VI7 | Ribonuclea | Rnh1 | sp\|Q91VI7 | 28 | 28 | 85,3 |
| Q93092 | Q93092 | Transaldola | Taldo1 | sp\|Q93092 | 15 | 15 | 39,8 |
| Q9ERA6 | Q9ERA6 | Tuftelin-int | Tfip11 | sp\|Q9ERA6 | 26 | 26 | 37,5 |
| Q02257 | Q02257 | Junction pl | Jup | sp\|Q02257 | 22 | 18 | 41,6 |
| Q9QXS1 | Q9QXS1 | Plectin | Plec | sp\|Q9QXS1 | 314 | 310 | 67,3 |
| Q8CIM8 | Q8CIM8 | Integrator | Ints4 | sp\|Q8CIM8 | 25 | 25 | 35,1 |
| P18242 | P18242 | Cathepsin | Ctsd | sp\|P18242 | 15 | 15 | 42,9 |
| Q61739 | Q61739 | Integrin alp | Itga6 | sp\|Q61739 | 39 | 39 | 43,4 |
| P53690 | P53690 | Matrix met | Mmp14 | sp\|P53690 | 16 | 16 | 31,1 |
| Q69ZR9 | Q69ZR9 | Protein FA | Fam208a | sp\|Q69ZR9 | 15 | 15 | 13,9 |
| Q6A065 | Q6A065 | Centrosom | Cep170 | sp\|Q6A065 | 39 | 38 | 35,1 |
| P42225 | P42225 | Signal tran | Stat1 | sp\|P42225 | 20 | 20 | 35,1 |
| Q9D620;G | Q9D620 | Rab11 fam | Rab11fip1 | sp\|Q9D620 | 14 | 14 | 29,3 |
| Q9D7H3 | Q9D7H3 | RNA 3-term | RtcA | sp\|Q9D7H | 14 | 14 | 46,4 |
| Q9CRY7 | Q9CRY7 | Glyceropho | Gdpd1 | sp\|Q9CRY7 | 10 | 10 | 47,8 |
| P52651 | P52651 | Homeobox | Rhox5 | sp\|P52651 | 11 | 11 | 61 |
| B2RXS4;Q9 | B2RXS4 | Plexin-B2 | Plxnb2 | sp\|B2RXS4 | 47 | 47 | 35,8 |
| P35456 | P35456 | Urokinase | Plaur | sp\|P35456 | 3 | 3 | 14,1 |
| Q9CRD0 | Q9CRD0 | OCIA doma | Ociad1 | sp\|Q9CRD0 | 16 | 16 | 72,5 |
| Q9JMK0 | Q9JMK0 | Beta-1,4-ga | B4galt5 | sp\|Q9JMK0 | 4 | 3 | 10,6 |
| Q8VCR7 | Q8VCR7 | Alpha/beta | Abhd14b | sp\|Q8VCR7 | 8 | 8 | 57,1 |
| Q71LX4 | Q71LX4 | Talin-2 | Tln2 | sp\|Q71LX4 | 62 | 51 | 36,3 |

| | | | | | | | |
|---|---|---|---|---|---|---|---|
| Q00519 | Q00519 | Xanthine d | Xdh | sp\|Q00519 | 36 | 35 | 39,2 |
| Q9DA03 | Q9DA03 | Complex II | Lyrm7 | sp\|Q9DA0 | 2 | 2 | 34,6 |
| O55229 | O55229 | Choline/et | Chkb | sp\|O55229 | 9 | 9 | 26,6 |
| Q9R0G7 | Q9R0G7 | Zinc finger | Zeb2 | sp\|Q9R0G | 16 | 16 | 19,3 |
| Q921C5 | Q921C5 | Protein bic | Bicd2 | sp\|Q921C5 | 23 | 20 | 36,3 |
| P16110 | P16110 | Galectin-3 | Lgals3 | sp\|P16110 | 10 | 10 | 35,6 |
| Q923D2 | Q923D2 | Flavin redu | Blvrb | sp\|Q923D2 | 10 | 10 | 71,8 |
| Q8C0D5 | Q8C0D5 | Elongation | Eftud1 | sp\|Q8C0D5 | 29 | 29 | 35,8 |
| P24527 | P24527 | Leukotrien | Lta4h | sp\|P24527 | 29 | 29 | 57,1 |
| Q9JII6 | Q9JII6 | Alcohol de | Akr1a1 | sp\|Q9JII6\| | 18 | 18 | 66,8 |
| Q8BK35 | Q8BK35 | | Gltscr2 | sp\|Q8BK35 | 16 | 16 | 40,3 |
| Q67FY2 | Q67FY2 | B-cell CLL/l | Bcl9l | sp\|Q67FY2 | 10 | 10 | 9,8 |
| P46735 | P46735 | Unconvent | Myo1b | sp\|P46735 | 35 | 31 | 36,5 |
| Q8BZ33 | Q8BZ33 | Tensin-4 | Tns4 | sp\|Q8BZ33 | 18 | 18 | 40,5 |
| P70227 | P70227 | Inositol 1,4 | Itpr3 | sp\|P70227 | 78 | 74 | 39,3 |
| Q05793 | Q05793 | Basement | Hspg2 | sp\|Q05793 | 74 | 74 | 31 |
| P97329 | P97329 | Kinesin-like | Kif20a | sp\|P97329 | 28 | 28 | 42,8 |
| Q9QUK4;Q | Q9QUK4 | Baculovira | Naip2 | sp\|Q9QUK | 11 | 10 | 12 |
| P11688 | P11688 | Integrin alp | Itga5 | sp\|P11688 | 25 | 25 | 28,4 |
| Q07113 | Q07113 | Cation-ind | Igf2r | sp\|Q07113 | 72 | 72 | 38,7 |
| P47811 | P47811 | Mitogen-a | Mapk14 | sp\|P47811 | 13 | 13 | 53,3 |
| Q9QXS6 | Q9QXS6 | Drebrin | Dbn1 | sp\|Q9QXS6 | 18 | 18 | 41,5 |
| Q8K072 | Q8K072 | Receptor e | Reep4 | sp\|Q8K072 | 4 | 4 | 19,5 |
| P43883 | P43883 | Perilipin-2 | Plin2 | sp\|P43883 | 14 | 14 | 47,8 |
| Q9EQP2 | Q9EQP2 | EH domain | Ehd4 | sp\|Q9EQP2 | 30 | 24 | 68,9 |
| Q9D0F9 | Q9D0F9 | Phosphogl | Pgm1 | sp\|Q9D0F9 | 32 | 32 | 66,5 |
| P59242 | P59242 | Cingulin | Cgn | sp\|P59242 | 17 | 16 | 21,9 |
| Q8R3H7 | Q8R3H7 | Heparan su | Hs2st1 | sp\|Q8R3H7 | 11 | 11 | 37,1 |
| Q9JHK4 | Q9JHK4 | Geranylger | Rabggta | sp\|Q9JHK4 | 19 | 19 | 39 |
| P97371 | P97371 | Proteasom | Psme1 | sp\|P97371 | 16 | 16 | 54,2 |
| Q8K411 | Q8K411 | Presequen | Pitrm1 | sp\|Q8K411 | 39 | 39 | 49,9 |
| P97770 | P97770 | THUMP do | Thumpd3 | sp\|P97770 | 17 | 17 | 43 |
| Q9D8C4 | Q9D8C4 | Interferon- | Ifi35 | sp\|Q9D8C4 | 9 | 9 | 51,7 |
| Q8BHG2 | Q8BHG2 | UPF0587 protein C1or | | sp\|Q8BHG | 4 | 4 | 45,6 |
| Q09200 | Q09200 | Beta-1,4 N | B4galnt1 | sp\|Q09200 | 11 | 11 | 26,8 |
| Q8K354 | Q8K354 | Carbonyl r | Cbr3 | sp\|Q8K354 | 11 | 10 | 54,2 |
| Q8BKC5 | Q8BKC5 | Importin-5 | Ipo5 | sp\|Q8BKC5 | 51 | 50 | 65,1 |
| Q02053;P3 | Q02053 | Ubiquitin-l | Uba1 | sp\|Q02053 | 47 | 47 | 60,9 |
| Q80TR8 | Q80TR8 | Protein VP | Vprbp | sp\|Q80TR8 | 35 | 35 | 30,1 |
| Q9DCX2 | Q9DCX2 | ATP syntha | Atp5h | sp\|Q9DCX2 | 15 | 15 | 77 |
| Q8BYR1 | Q8BYR1 | tRNA wybu | Lcmt2 | sp\|Q8BYR1 | 7 | 7 | 14,3 |
| Q3UZ09 | Q3UZ09 | Compleme | C1rl | sp\|Q3UZ09 | 4 | 4 | 13,5 |
| Q91Y74 | Q91Y74 | CMP-N-ace | St3gal4 | sp\|Q91Y74 | 7 | 7 | 27,6 |
| O35495 | O35495 | Cyclin-dep | Cdk14 | sp\|O35495 | 5 | 4 | 12,2 |
| Q8BY89 | Q8BY89 | Choline tra | Slc44a2 | sp\|Q8BY89 | 15 | 15 | 25,6 |
| Q8K2Q9 | Q8K2Q9 | Shootin-1 | Kiaa1598 | sp\|Q8K2Q | 25 | 25 | 42,6 |
| Q8BGD5 | Q8BGD5 | Carnitine O | Cpt1c | sp\|Q8BGD | 5 | 5 | 8,4 |
| Q8R326 | Q8R326 | Paraspeckl | Pspc1 | sp\|Q8R326 | 13 | 13 | 33,7 |
| CON__Q3S | CON__Q3SZV7 | | | | 14 | 14 | 42 |
| B1AY13 | B1AY13 | Ubiquitin c | Usp24 | sp\|B1AY13 | 44 | 44 | 22,6 |

| | | | | | | | |
|---|---|---|---|---|---|---|---|
| Q3TIU4 | Q3TIU4 | 2,5-phosph | Pde12 | sp\|Q3TIU4 | 16 | 16 | 39 |
| O88844 | O88844 | Isocitrate d | Idh1 | sp\|O88844 | 24 | 23 | 63,8 |
| Q9JLV5 | Q9JLV5 | Cullin-3 | Cul3 | sp\|Q9JLV5 | 37 | 37 | 49,2 |
| O88738;RE | O88738 | Baculovira | Birc6 | sp\|O88738 | 74 | 73 | 24,7 |
| B1ARD6;B1 | B1ARD6 | | Slfn9 | sp\|B1ARD6 | 33 | 33 | 43,1 |
| Q69ZR2 | Q69ZR2 | E3 ubiquiti | Hectd1 | sp\|Q69ZR2 | 81 | 81 | 41,4 |
| Q61411 | Q61411 | GTPase HR | Hras | sp\|Q61411 | 10 | 3 | 60,8 |
| P27612 | P27612 | Phospholip | Plaa | sp\|P27612 | 34 | 34 | 58,6 |
| Q3U7R1 | Q3U7R1 | Extended s | Esyt1 | sp\|Q3U7R | 36 | 36 | 43,9 |
| P70441 | P70441 | Na(+)/H(+) | Slc9a3r1 | sp\|P70441 | 14 | 14 | 54,9 |
| P28063 | P28063 | Proteasom | Psmb8 | sp\|P28063 | 11 | 11 | 48,9 |
| Q9ERK4 | Q9ERK4 | Exportin-2 | Cse1l | sp\|Q9ERK4 | 39 | 39 | 46,8 |
| Q8C878 | Q8C878 | NEDD8-act | Uba3 | sp\|Q8C878 | 18 | 18 | 51,1 |
| Q6PAJ1 | Q6PAJ1 | Breakpoint | Bcr | sp\|Q6PAJ1 | 20 | 19 | 19,7 |
| Q8BX90 | Q8BX90 | Fibronectir | Fndc3a | sp\|Q8BX90 | 30 | 30 | 40,7 |
| P60330 | P60330 | Separin | Espl1 | sp\|P60330 | 29 | 29 | 21,1 |
| P56389 | P56389 | Cytidine de | Cda | sp\|P56389 | 7 | 7 | 79,5 |
| CON__Q0I | CON__Q0IIK2;CON__Q29443 | | | ; | 38 | 38 | 65,6 |
| P09411 | P09411 | Phosphogly | Pgk1 | sp\|P09411 | 37 | 28 | 83 |
| Q7TNG5 | Q7TNG5 | Echinoderr | Eml2 | sp\|Q7TNG | 11 | 11 | 25 |
| Q8BHG1 | Q8BHG1 | Nardilysin | Nrd1 | sp\|Q8BHG | 38 | 38 | 40,2 |
| Q61089 | Q61089 | Frizzled-6 | Fzd6 | sp\|Q61089 | 7 | 7 | 13,8 |
| Q99LB6 | Q99LB6 | Methionin | Mat2b | sp\|Q99LB6 | 13 | 13 | 45,2 |
| Q3UM45 | Q3UM45 | Protein ph | Ppp1r7 | sp\|Q3UM4 | 17 | 17 | 61,8 |
| Q6ZQ38 | Q6ZQ38 | Cullin-asso | Cand1 | sp\|Q6ZQ38 | 56 | 55 | 48,4 |
| Q80XP8 | Q80XP8 | Protein FA | Fam76b | sp\|Q80XP8 | 8 | 8 | 33 |
| Q99MU3 | Q99MU3 | Double-str | Adar | sp\|Q99MU | 28 | 28 | 36,1 |
| Q9JKF1 | Q9JKF1 | Ras GTPase | Iqgap1 | sp\|Q9JKF1 | 91 | 88 | 65,4 |
| Q8BKX1 | Q8BKX1 | Brain-spec | Baiap2 | sp\|Q8BKX1 | 27 | 27 | 61,5 |
| Q6ZWQ0 | Q6ZWQ0 | Nesprin-2 | Syne2 | sp\|Q6ZWQ | 172 | 170 | 34,7 |
| Q99KG5 | Q99KG5 | Lipolysis-st | Lsr | sp\|Q99KG5 | 7 | 7 | 17,3 |
| Q78PY7 | Q78PY7 | Staphyloco | Snd1 | sp\|Q78PY7 | 49 | 49 | 62,5 |
| Q8BSY0 | Q8BSY0 | Aspartyl/as | Asph | sp\|Q8BSY0 | 24 | 24 | 35,2 |
| Q64133 | Q64133 | Amine oxid | Maoa | sp\|Q64133 | 18 | 18 | 45,4 |
| Q9CQ21 | Q9CQ21 | Malignant | Mcts2 | sp\|Q9CQ2 | 6 | 3 | 53 |
| Q80XQ2 | Q80XQ2 | TBC1 doma | Tbc1d5 | sp\|Q80XQ | 15 | 15 | 27,7 |
| Q7TPW1 | Q7TPW1 | Nexilin | Nexn | sp\|Q7TPW | 6 | 6 | 8,4 |
| Q501J6 | Q501J6 | Probable A | Ddx17 | sp\|Q501J6 | 31 | 24 | 48,2 |
| P43275 | P43275 | Histone H1 | Hist1h1a | sp\|P43275 | 12 | 10 | 35,7 |
| Q80UK8 | Q80UK8 | Integrator | Ints2 | sp\|Q80UK8 | 17 | 17 | 18,4 |
| P70202 | P70202 | Latexin | Lxn | sp\|P70202 | 5 | 5 | 34,2 |
| Q921G6 | Q921G6 | Leucine-ric | Lrch4 | sp\|Q921G6 | 16 | 16 | 39,9 |
| Q9WV92 | Q9WV92 | Band 4.1-li | Epb41l3 | sp\|Q9WV9 | 34 | 29 | 48,8 |
| Q923E4 | Q923E4 | NAD-deper | Sirt1 | sp\|Q923E4 | 14 | 14 | 33,5 |
| Q91YM2 | Q91YM2 | Rho GTPas | Arhgap35 | sp\|Q91YM | 27 | 27 | 23,9 |
| Q8VCL2 | Q8VCL2 | Protein SC | Sco2 | sp\|Q8VCL2 | 8 | 8 | 35,3 |
| P47968 | P47968 | Ribose-5-p | Rpia | sp\|P47968 | 8 | 8 | 38 |
| Q69ZK6 | Q69ZK6 | Probable J | Jmjd1c | sp\|Q69ZK6 | 35 | 35 | 20,5 |
| Q99K01 | Q99K01 | Pyridoxal-d | Pdxdc1 | sp\|Q99K01 | 24 | 24 | 42,3 |
| P09528 | P09528 | Ferritin he | Fth1 | sp\|P09528 | 15 | 15 | 72 |

| | | | | | | | |
|---|---|---|---|---|---|---|---|
| Q9D8V7 | Q9D8V7 | Signal pept | Sec11c | sp\|Q9D8V | 7 | 7 | 30,7 |
| Q91VJ2 | Q91VJ2 | Protein kin | Prkcdbp | sp\|Q91VJ2 | 11 | 11 | 56,9 |
| P69566 | P69566 | Ran-bindin | Ranbp9 | sp\|P69566 | 17 | 15 | 36,6 |
| Q8C854 | Q8C854 | Myelin exp | Myef2 | sp\|Q8C854 | 23 | 23 | 44,2 |
| Q9WU78 | Q9WU78 | Programm | Pdcd6ip | sp\|Q9WU7 | 50 | 50 | 66,6 |
| Q9CPV5 | Q9CPV5 | Polyamine | Pmf1 | sp\|Q9CPV5 | 10 | 10 | 59,4 |
| Q8BKY8 | Q8BKY8 | Transcripti | Mterf2 | sp\|Q8BKY8 | 5 | 5 | 17,7 |
| Q8BL66 | Q8BL66 | Early endo | Eea1 | sp\|Q8BL66 | 60 | 60 | 47,3 |
| Q8K2H1 | Q8K2H1 | Periphilin-1 | Pphln1 | sp\|Q8K2H1 | 4 | 4 | 11,5 |
| P70398 | P70398 | Probable u | Usp9x | sp\|P70398 | 57 | 57 | 27,2 |
| P28658 | P28658 | Ataxin-10 | Atxn10 | sp\|P28658 | 27 | 27 | 62,5 |
| Q5DW34 | Q5DW34 | Histone-lys | Ehmt1 | sp\|Q5DW3 | 22 | 22 | 28,6 |
| Q69ZS7 | Q69ZS7 | HBS1-like p | Hbs1l | sp\|Q69ZS7 | 21 | 21 | 45,9 |
| Q9JLV1 | Q9JLV1 | BAG family | Bag3 | sp\|Q9JLV1 | 33 | 33 | 73,1 |
| Q8C1D8 | Q8C1D8 | Protein IW | Iws1 | sp\|Q8C1D8 | 14 | 14 | 19,5 |
| Q8CAF4 | Q8CAF4 | NHS-like pr | Nhsl1 | sp\|Q8CAF4 | 8 | 8 | 9,1 |
| Q8C0P0 | Q8C0P0 | Serine/thre | Mastl | sp\|Q8C0P0 | 19 | 18 | 35 |
| Q8BPG6 | Q8BPG6 | Sulfatase-m | Sumf2 | sp\|Q8BPG | 8 | 8 | 31,5 |
| Q91VM9 | Q91VM9 | Inorganic p | Ppa2 | sp\|Q91VM | 17 | 17 | 60,3 |
| A6X919;P0 | A6X919 | Probable C | Dpy19l1 | sp\|A6X919 | 13 | 13 | 25,6 |
| Q99K51 | Q99K51 | Plastin-3 | Pls3 | sp\|Q99K51 | 37 | 30 | 61,9 |
| Q9EQQ9 | Q9EQQ9 | Protein O-( | Mgea5 | sp\|Q9EQQ | 31 | 31 | 50,5 |
| Q8BHZ4 | Q8BHZ4 | Zinc finger | Znf592 | sp\|Q8BHZ4 | 14 | 14 | 17,4 |
| Q99K30 | Q99K30 | Epidermal | Eps8l2 | sp\|Q99K30 | 29 | 29 | 49,8 |
| Q9JHL1 | Q9JHL1 | Na(+)/H(+) | Slc9a3r2 | sp\|Q9JHL1 | 12 | 12 | 40,9 |
| Q8BIJ7 | Q8BIJ7 | RUN and F | Rufy1 | sp\|Q8BIJ7 | 15 | 14 | 22,6 |
| Q9Z1Z0 | Q9Z1Z0 | General ve | Uso1 | sp\|Q9Z1Z0 | 36 | 36 | 48,2 |
| Q6PCM2 | Q6PCM2 | Integrator | Ints6 | sp\|Q6PCM | 22 | 20 | 37,6 |
| P59113 | P59113 | Fermitin fa | Fermt1 | sp\|P59113 | 14 | 14 | 27,9 |
| P56480 | P56480 | ATP syntha | Atp5b | sp\|P56480 | 33 | 33 | 85,3 |
| Q9CXT8 | Q9CXT8 | Mitochond | Pmpcb | sp\|Q9CXT8 | 19 | 18 | 54,2 |
| P11679;CC | P11679 | Keratin, ty | Krt8 | sp\|P11679 | 45 | 17 | 72,4 |
| E9Q7G0 | E9Q7G0 | | Numa1 | sp\|E9Q7G0 | 109 | 109 | 60,7 |
| Q9CZA6 | Q9CZA6 | Nuclear dis | Nde1 | sp\|Q9CZA6 | 9 | 7 | 41,6 |
| Q9QXX4 | Q9QXX4 | Calcium-bi | Slc25a13 | sp\|Q9QXX | 26 | 23 | 48,5 |
| Q9QXZ0 | Q9QXZ0 | Microtubu | Macf1 | sp\|Q9QXZ0 | 147 | 141 | 26,9 |
| Q8BGS1 | Q8BGS1 | Band 4.1-li | Epb41l5 | sp\|Q8BGS1 | 9 | 9 | 21,3 |
| Q8BZF2 | Q8BZF2 | Magnesiur | Nipal4 | sp\|Q8BZF2 | 3 | 3 | 10,8 |
| Q9WV34 | Q9WV34 | MAGUK p5 | Mpp2 | sp\|Q9WV3 | 9 | 8 | 25 |
| P35762 | P35762 | CD81 antig | Cd81 | sp\|P35762 | 3 | 3 | 21,2 |
| P28862;O5 | P28862 | Stromelysi | Mmp3 | sp\|P28862 | 15 | 15 | 44 |
| P32921 | P32921 | Tryptophar | Wars | sp\|P32921 | 26 | 26 | 64,4 |
| Q99MN9 | Q99MN9 | Propionyl-( | Pccb | sp\|Q99MN | 17 | 17 | 42,5 |
| Q9CQN1 | Q9CQN1 | Heat shock | Trap1 | sp\|Q9CQN | 33 | 33 | 55,4 |
| Q80TC5 | Q80TC5 | Pogo trans | Pogk | sp\|Q80TC5 | 3 | 3 | 6,8 |
| P28352 | P28352 | DNA-(apur | Apex1 | sp\|P28352 | 16 | 16 | 62,8 |
| Q9R1Z8 | Q9R1Z8 | Vinexin | Sorbs3 | sp\|Q9R1Z8 | 10 | 10 | 19,4 |
| O70293;O7 | O70293 | G protein-( | Grk6 | sp\|O70293 | 12 | 9 | 30,4 |
| Q6EDY6 | Q6EDY6 | Leucine-ric | Lrrc16a | sp\|Q6EDY6 | 23 | 23 | 22,6 |
| P08074 | P08074 | Carbonyl re | Cbr2 | sp\|P08074 | 6 | 6 | 33,2 |

| | | | | | | | |
|---|---|---|---|---|---|---|---|
| Q9DCM0 | Q9DCM0 | Persulfide | Ethe1 | sp\|Q9DCM | 12 | 12 | 70,1 |
| Q91XE4 | Q91XE4 | N-acyl-aro | Acy3 | sp\|Q91XE4 | 10 | 10 | 45,9 |
| Q9ER41 | Q9ER41 | Torsin-1B | Tor1b | sp\|Q9ER41 | 7 | 7 | 25 |
| E9Q555 | E9Q555 | E3 ubiquiti | Rnf213 | sp\|E9Q555 | 136 | 136 | 34 |
| Q9WU03 | Q9WU03 | Kunitz-type | Spint2 | sp\|Q9WU0 | 8 | 8 | 34,9 |
| Q61666 | Q61666 | Protein HIF | Hira | sp\|Q61666 | 17 | 17 | 24,1 |
| Q9JL26 | Q9JL26 | Formin-like | Fmnl1 | sp\|Q9JL26 | 29 | 26 | 33,6 |
| Q61048 | Q61048 | WW doma | Wbp4 | sp\|Q61048 | 5 | 5 | 22,9 |
| Q8C181 | Q8C181 | Muscleblin | Mbnl2 | sp\|Q8C181 | 7 | 2 | 26 |
| P97393 | P97393 | Rho GTPas | Arhgap5 | sp\|P97393 | 28 | 28 | 24,9 |
| Q6PDI5 | Q6PDI5 | Proteasom | Ecm29 | sp\|Q6PDI5 | 64 | 64 | 46,4 |
| Q8K4Q8 | Q8K4Q8 | Collectin-1 | Colec12 | sp\|Q8K4Q | 19 | 19 | 34 |
| P54763;Q8 | P54763 | Ephrin type | Ephb2 | sp\|P54763 | 11 | 8 | 14 |
| Q9D5T0 | Q9D5T0 | ATPase far | Atad1 | sp\|Q9D5T0 | 14 | 14 | 52,4 |
| Q8R502 | Q8R502 | Volume-re | Lrrc8c | sp\|Q8R502 | 30 | 27 | 44,1 |
| Q91YK2 | Q91YK2 | Ribosomal | Rrp1b | sp\|Q91YK2 | 23 | 23 | 43,1 |
| A2A863 | A2A863 | Integrin be | Itgb4 | sp\|A2A863 | 52 | 52 | 39,2 |
| P40142 | P40142 | Transketol | Tkt | sp\|P40142 | 37 | 37 | 65,8 |
| P59672 | P59672 | Ankyrin re | Anks1a | sp\|P59672 | 13 | 13 | 16,3 |
| Q7TQK1 | Q7TQK1 | Integrator | Ints7 | sp\|Q7TQK | 25 | 25 | 34,5 |
| Q9JI38 | Q9JI38 | tRNA pseu | Pus3 | sp\|Q9JI38 | 4 | 4 | 13,3 |
| Q922D8 | Q922D8 | C-1-tetrahy | Mthfd1 | sp\|Q922D8 | 47 | 47 | 61,7 |
| Q8CGK3 | Q8CGK3 | Lon protea | Lonp1 | sp\|Q8CGK | 41 | 41 | 49,7 |
| Q8VCT3 | Q8VCT3 | Aminopept | Rnpep | sp\|Q8VCT3 | 28 | 28 | 52,9 |
| Q8BU04 | Q8BU04 | Putative E3 | Ubr7 | sp\|Q8BU0 | 10 | 10 | 22,4 |
| CON__P19 | CON__P19001 | Keratin, ty | Krt19 | ;sp\|P1900 | 30 | 19 | 76,9 |
| Q99KQ4 | Q99KQ4 | Nicotinami | Nampt | sp\|Q99KQ | 23 | 23 | 66,4 |
| P25322;P3 | P25322 | G1/S-speci | Ccnd1 | sp\|P25322 | 15 | 15 | 63,1 |
| Q8K013 | Q8K013 | GTP-bindir | Gtpbp10 | sp\|Q8K013 | 7 | 7 | 30,6 |
| Q9CQE9 | Q9CQE9 | FLYWCH fa | Flywch2 | sp\|Q9CQE | 3 | 3 | 28,1 |
| Q8R310;Q8 | Q8R310 | Transmem | Tmcc3 | sp\|Q8R310 | 10 | 10 | 30,6 |
| Q8BJS4 | Q8BJS4 | SUN doma | Sun2 | sp\|Q8BJS4 | 25 | 25 | 51,4 |
| Q9JKA5 | Q9JKA5 | Cell surface | Gpa33 | sp\|Q9JKA5 | 6 | 6 | 17,2 |
| Q9CX34 | Q9CX34 | Suppressor | Sugt1 | sp\|Q9CX34 | 19 | 19 | 62,5 |
| Q6P5H2 | Q6P5H2 | Nestin | Nes | sp\|Q6P5H2 | 71 | 71 | 50,4 |
| P08122 | P08122 | Collagen al | Col4a2 | sp\|P08122 | 16 | 16 | 14,2 |
| P22437 | P22437 | Prostaglan | Ptgs1 | sp\|P22437 | 17 | 16 | 34,7 |
| P00493 | P00493 | Hypoxanth | Hprt1 | sp\|P00493 | 12 | 12 | 64,2 |
| Q3TVI8 | Q3TVI8 | Pre-B-cell l | Pbxip1 | sp\|Q3TVI8 | 11 | 11 | 20,4 |
| O70572 | O70572 | Sphingomy | Smpd2 | sp\|O70572 | 15 | 15 | 44,4 |
| Q9DAU7 | Q9DAU7 | WAP four-d | Wfdc2 | sp\|Q9DAU | 3 | 3 | 32,8 |
| Q8BRM2 | Q8BRM2 | RAB6-inter | Gorab | sp\|Q8BRM | 9 | 9 | 38 |
| P51881 | P51881 | ADP/ATP t | Slc25a5 | sp\|P51881 | 22 | 11 | 63,1 |
| F7BWT7 | F7BWT7 | Tetraspani | Tspan15 | sp\|F7BWT | 5 | 5 | 21,1 |
| P17879;Q6 | P17879;Q6169 | Heat shock | Hspa1b;Hs | sp\|P17879 | 28 | 16 | 54,8 |
| Q01965 | Q01965 | T-lymphoc | Ly9 | sp\|Q01965 | 6 | 6 | 15 |
| P58281;P5 | P58281 | Dynamin-li | Opa1 | sp\|P58281 | 31 | 31 | 39,7 |
| Q99KG3 | Q99KG3 | RNA-bindir | Rbm10 | sp\|Q99KG | 23 | 19 | 33 |
| O35350 | O35350 | Calpain-1 c | Capn1 | sp\|O35350 | 32 | 32 | 50,1 |
| Q8BMS1 | Q8BMS1 | Trifunction | Hadha | sp\|Q8BMS | 36 | 36 | 63 |

| | | | | | | | |
|---|---|---|---|---|---|---|---|
| Q5DTT8 | Q5DTT8 | Paraneopla | Pnma5 | sp\|Q5DTT8 | 8 | 8 | 20,9 |
| Q99JY0 | Q99JY0 | Trifunction | Hadhb | sp\|Q99JY0 | 23 | 23 | 52,4 |
| Q9R097 | Q9R097 | Kunitz-type | Spint1 | sp\|Q9R097 | 7 | 7 | 19,3 |
| Q91WG2 | Q91WG2 | Rab GTPas | Rabep2 | sp\|Q91WG | 15 | 15 | 36,3 |
| Q6P4S8 | Q6P4S8 | Integrator | Ints1 | sp\|Q6P4S8 | 48 | 48 | 29,1 |
| Q5S248 | Q5S248 | Transmem | Tmprss11e | sp\|Q5S248 | 16 | 16 | 47,8 |
| Q8R5H1 | Q8R5H1 | Ubiquitin c | Usp15 | sp\|Q8R5H | 39 | 37 | 51,8 |
| Q925B0 | Q925B0 | PRKC apop | Pawr | sp\|Q925B0 | 13 | 13 | 64 |
| Q99M71 | Q99M71 | Mammalia | Epdr1 | sp\|Q99M7 | 4 | 4 | 19,2 |
| Q99JR8 | Q99JR8 | SWI/SNF-r | Smarcd2 | sp\|Q99JR8 | 19 | 17 | 44,3 |
| Q9EQK5 | Q9EQK5 | Major vaul | Mvp | sp\|Q9EQK | 39 | 39 | 60,2 |
| Q812F8 | Q812F8 | Alpha-1,3- | Mgat4b | sp\|Q812F8 | 4 | 4 | 10,6 |
| P29533 | P29533 | Vascular ce | Vcam1 | sp\|P29533 | 23 | 23 | 39,6 |
| Q8VI75 | Q8VI75 | Importin-4 | Ipo4 | sp\|Q8VI75 | 29 | 29 | 33,3 |
| Q80U49 | Q80U49 | Centrosom | Cep170b | sp\|Q80U4 | 27 | 26 | 27,1 |
| Q8VE70 | Q8VE70 | Programm | Pdcd10 | sp\|Q8VE70 | 10 | 10 | 61,3 |
| Q64318 | Q64318 | Zinc finger | Zeb1 | sp\|Q64318 | 10 | 10 | 16 |
| P54822 | P54822 | Adenylosu | Adsl | sp\|P54822 | 23 | 23 | 59,9 |
| Q9R0Y5 | Q9R0Y5 | Adenylate | Ak1 | sp\|Q9R0Y5 | 10 | 10 | 59,3 |
| Q04736;P0 | Q04736 | Tyrosine-p | Yes1 | sp\|Q04736 | 18 | 11 | 46,2 |
| Q9WUU7 | Q9WUU7 | Cathepsin | Ctsz | sp\|Q9WU | 12 | 12 | 58,5 |
| P24547 | P24547 | Inosine-5-r | Impdh2 | sp\|P24547 | 29 | 28 | 55,6 |
| CON__REF | CON__REFSEQ:XP_001252647 | | | | 9 | 7 | 20,6 |
| E9Q5F9 | E9Q5F9 | Histone-lys | Setd2 | sp\|E9Q5F9 | 29 | 29 | 16,2 |
| Q05769 | Q05769 | Prostaglan | Ptgs2 | sp\|Q05769 | 24 | 23 | 54 |
| Q9WVL3 | Q9WVL3 | Solute carr | Slc12a7 | sp\|Q9WVL | 15 | 15 | 19,8 |
| O35379 | O35379 | Multidrug | Abcc1 | sp\|O35379 | 36 | 35 | 32,1 |
| Q9Z0L0 | Q9Z0L0 | Trophoblas | Tpbg | sp\|Q9Z0L0 | 6 | 6 | 20,7 |
| Q91ZA3 | Q91ZA3 | Propionyl- | Pcca | sp\|Q91ZA3 | 23 | 23 | 42,8 |
| Q9R0E2 | Q9R0E2 | Procollage | Plod1 | sp\|Q9R0E2 | 23 | 23 | 42 |
| Q61503 | Q61503 | 5-nucleoti | Nt5e | sp\|Q61503 | 21 | 21 | 47,9 |
| Q99LF4 | Q99LF4 | tRNA-splic | Rtcb | sp\|Q99LF4 | 22 | 22 | 49,7 |
| P51432 | P51432 | 1-phospha | Plcb3 | sp\|P51432 | 35 | 35 | 33,5 |
| Q80V03 | Q80V03 | Uncharact | Adck5 | sp\|Q80V03 | 13 | 13 | 26,5 |
| Q62383 | Q62383 | Transcripti | Supt6h | sp\|Q62383 | 54 | 53 | 38,5 |
| O89050 | O89050 | Muskelin | Mkln1 | sp\|O89050 | 25 | 25 | 45,3 |
| P06869 | P06869 | Urokinase- | Plau | sp\|P06869 | 9 | 9 | 32,6 |
| P06151;P0 | P06151 | L-lactate d | Ldha | sp\|P06151 | 26 | 25 | 71,1 |
| P27641 | P27641 | X-ray repai | Xrcc5 | sp\|P27641 | 32 | 32 | 54,8 |
| Q8VC28 | Q8VC28 | Aldo-keto | Akr1c13 | sp\|Q8VC28 | 13 | 13 | 62,8 |
| Q6NZR2 | Q6NZR2 | Myb/SANT | Msantd2 | sp\|Q6NZR | 12 | 12 | 33,1 |
| P70698 | P70698 | CTP syntha | Ctps1 | sp\|P70698 | 25 | 23 | 54,1 |
| P39447 | P39447 | Tight junct | Tjp1 | sp\|P39447 | 54 | 54 | 44,5 |
| Q8VBT9 | Q8VBT9 | Tether con | Aspscr1 | sp\|Q8VBT | 19 | 19 | 43,5 |
| Q9WV91 | Q9WV91 | Prostaglan | Ptgfrn | sp\|Q9WV9 | 28 | 28 | 38,8 |
| Q9WVC3 | Q9WVC3 | Caveolin-2 | Cav2 | sp\|Q9WV0 | 5 | 5 | 54,9 |
| Q9D820 | Q9D820 | Prolyl-tRN | Prorsd1 | sp\|Q9D82 | 4 | 4 | 32,5 |
| Q99J79 | Q99J79 | DNA dama | Ddb2 | sp\|Q99J79 | 7 | 7 | 21,1 |
| Q689Z5 | Q689Z5 | Protein str | Sbno1 | sp\|Q689Z5 | 29 | 26 | 32,3 |
| Q9CW46 | Q9CW46 | Ribonucle | Raver1 | sp\|Q9CW4 | 26 | 26 | 53,2 |

| | | | | | | | |
|---|---|---|---|---|---|---|---|
| Q9CZ91 | Q9CZ91 | Serum resp | Srfbp1 | sp\|Q9CZ91 | 4 | 4 | 10,7 |
| Q9ESN6 | Q9ESN6 | Tripartite r | Trim2 | sp\|Q9ESN6 | 14 | 13 | 28,8 |
| Q9CYA6 | Q9CYA6 | Zinc finger | Zcchc8 | sp\|Q9CYA6 | 19 | 19 | 49,6 |
| B9EJ86 | B9EJ86 | Oxysterol-l | Osbpl8 | sp\|B9EJ86 | 19 | 17 | 33,2 |
| Q6NV83 | Q6NV83 | U2 snRNP- | U2surp | sp\|Q6NV8 | 35 | 35 | 40,9 |
| Q9CR26 | Q9CR26 | Vacuolar p | Vta1 | sp\|Q9CR26 | 9 | 9 | 33,7 |
| O08528;Q3 | O08528 | Hexokinase | Hk2 | sp\|O08528 | 39 | 34 | 50,9 |
| Q9R045 | Q9R045 | Angiopoiet | Angptl2 | sp\|Q9R045 | 16 | 15 | 41,2 |
| CON__P15 | CON__P15497 | | | | 14 | 14 | 47,5 |
| Q3URE1 | Q3URE1 | Acyl-CoA s | Acsf3 | sp\|Q3URE | 7 | 7 | 21,6 |
| Q04750;Q8 | Q04750 | DNA topois | Top1 | sp\|Q04750 | 32 | 32 | 42,4 |
| Q91VE6 | Q91VE6 | MKI67 FHA | Nifk | sp\|Q91VE6 | 19 | 19 | 64,4 |
| Q6ZQ08 | Q6ZQ08 | CCR4-NOT | Cnot1 | sp\|Q6ZQ08 | 81 | 81 | 44,1 |
| P52479 | P52479 | Ubiquitin c | Usp10 | sp\|P52479 | 20 | 20 | 43,7 |
| P46935 | P46935 | E3 ubiquiti | Nedd4 | sp\|P46935 | 35 | 32 | 52,9 |
| O35639 | O35639 | Annexin A3 | Anxa3 | sp\|O35639 | 31 | 31 | 76,8 |
| P35951 | P35951 | Low-densit | Ldlr | sp\|P35951 | 22 | 21 | 37,4 |
| Q8BXV2 | Q8BXV2 | BRI3-bindi | Bri3bp | sp\|Q8BXV2 | 3 | 3 | 16,6 |
| Q8VIK5 | Q8VIK5 | Platelet en | Pear1 | sp\|Q8VIK5 | 6 | 6 | 10,6 |
| Q9JL35 | Q9JL35 | High mobil | Hmgn5 | sp\|Q9JL35 | 9 | 9 | 33 |
| Q9DBJ1;O7 | Q9DBJ1 | Phosphogly | Pgam1 | sp\|Q9DBJ1 | 20 | 20 | 80,7 |
| O88986 | O88986 | 2-amino-3- | Gcat | sp\|O88986 | 14 | 14 | 52,4 |
| G3X9J0 | G3X9J0 | | Sipa1l3 | sp\|G3X9J0 | 42 | 42 | 36,9 |
| P47856;Q9 | P47856 | Glutamine- | Gfpt1 | sp\|P47856 | 29 | 29 | 53,5 |
| Q60714 | Q60714 | Long-chain | Slc27a1 | sp\|Q60714 | 6 | 5 | 13,6 |
| Q3ULD5 | Q3ULD5 | Methylcrot | Mccc2 | sp\|Q3ULD | 18 | 18 | 44,4 |
| CON__P06 | CON__P06868 | | | | 25 | 25 | 43,2 |
| Q3TZZ7 | Q3TZZ7 | Extended s | Esyt2 | sp\|Q3TZZ7 | 25 | 25 | 45,4 |
| Q9CQT2 | Q9CQT2 | RNA-bindir | Rbm7 | sp\|Q9CQT | 9 | 9 | 47,9 |
| Q6NVF4 | Q6NVF4 | DNA helica | Helb | sp\|Q6NVF4 | 20 | 20 | 24 |
| Q9ET54;Q5 | Q9ET54 | Palladin | Palld | sp\|Q9ET54 | 21 | 21 | 22,9 |
| Q99KK9 | Q99KK9 | Probable h | Hars2 | sp\|Q99KK9 | 19 | 16 | 44,6 |
| Q80U35 | Q80U35 | Rho guanir | Arhgef17 | sp\|Q80U35 | 23 | 23 | 18,5 |
| P10107 | P10107 | Annexin A1 | Anxa1 | sp\|P10107 | 32 | 32 | 76,6 |
| Q62074 | Q62074 | Protein kin | Prkci | sp\|Q62074 | 10 | 10 | 29,4 |
| Q9CWS0 | Q9CWS0 | N(G),N(G)- | Ddah1 | sp\|Q9CWS | 16 | 15 | 69,1 |
| Q9JL16 | Q9JL16 | Interferon- | Isg20 | sp\|Q9JL16 | 4 | 4 | 14,3 |
| Q8R2Y2 | Q8R2Y2 | Cell surface | Mcam | sp\|Q8R2Y2 | 7 | 7 | 16 |
| Q00993;Q6 | Q00993 | Tyrosine-p | Axl | sp\|Q00993 | 31 | 31 | 46,8 |
| Q8BTT6 | Q8BTT6 | Digestive o | Diexf | sp\|Q8BTT6 | 21 | 21 | 33 |
| Q9D0B6 | Q9D0B6 | Protein PB | Pbdc1 | sp\|Q9D0B6 | 9 | 9 | 55,1 |
| Q9EQJ9 | Q9EQJ9 | Membrane | Magi3 | sp\|Q9EQJ9 | 17 | 16 | 14,6 |
| Q9Z2I8 | Q9Z2I8 | Succinyl-Co | Suclg2 | sp\|Q9Z2I8 | 21 | 21 | 56,8 |
| Q63870 | Q63870 | Collagen al | Col7a1 | sp\|Q63870 | 21 | 21 | 12,3 |
| P50171 | P50171 | Estradiol 1 | Hsd17b8 | sp\|P50171 | 6 | 6 | 35,9 |
| P51125 | P51125 | Calpastatin | Cast | sp\|P51125 | 27 | 27 | 42,1 |
| Q8K4L3 | Q8K4L3 | Supervillin | Svil | sp\|Q8K4L3 | 40 | 40 | 25,7 |
| P50396 | P50396 | Rab GDP d | Gdi1 | sp\|P50396 | 27 | 21 | 74,9 |
| P05627 | P05627 | Transcripti | Jun | sp\|P05627 | 10 | 9 | 51,5 |
| Q5SF07 | Q5SF07 | Insulin-like | Igf2bp2 | sp\|Q5SF07 | 23 | 21 | 44,1 |

| | | | | | | | |
|---|---|---|---|---|---|---|---|
| Q02248 | Q02248 | Catenin be | Ctnnb1 | sp\|Q02248 | 28 | 24 | 51,6 |
| Q8R0S2 | Q8R0S2 | IQ motif ar | Iqsec1 | sp\|Q8R0S2 | 10 | 8 | 12,7 |
| Q921T2 | Q921T2 | Torsin-1A-i | Tor1aip1 | sp\|Q921T2 | 20 | 20 | 52,3 |
| P97386 | P97386 | DNA ligase | Lig3 | sp\|P97386 | 30 | 30 | 38,5 |
| Q8JZK9;P5 | Q8JZK9 | Hydroxyme | Hmgcs1 | sp\|Q8JZK9 | 17 | 17 | 36,7 |
| Q99K43 | Q99K43 | Protein reg | Prc1 | sp\|Q99K43 | 25 | 25 | 54,9 |
| Q9EPL4 | Q9EPL4 | Methyltrar | Mettl9 | sp\|Q9EPL4 | 5 | 5 | 19,8 |
| Q8VHK9 | Q8VHK9 | ATP-deper | Dhx36 | sp\|Q8VHK | 31 | 30 | 39,8 |
| Q99LS3 | Q99LS3 | Phosphose | Psph | sp\|Q99LS3 | 11 | 11 | 69,3 |
| Q3THS6;Q | Q3THS6 | S-adenosyl | Mat2a | sp\|Q3THS6 | 15 | 15 | 51,1 |
| Q8BI22 | Q8BI22 | Centrosom | Cep128 | sp\|Q8BI22 | 4 | 4 | 4,5 |
| Q8BRV5 | Q8BRV5 | Uncharacte | Kiaa1671 | sp\|Q8BRV5 | 7 | 7 | 41,2 |
| O88736 | O88736 | 3-keto-ste | Hsd17b7 | sp\|O88736 | 7 | 7 | 33,2 |
| Q8K3W0 | Q8K3W0 | BRCA1-A c | Bre | sp\|Q8K3W | 10 | 10 | 29,2 |
| Q9R1T2 | Q9R1T2 | SUMO-acti | Sae1 | sp\|Q9R1T2 | 22 | 22 | 72 |
| P63013;Q0 | P63013 | Paired mes | Prrx1 | sp\|P63013 | 7 | 6 | 48,2 |
| Q9R0P3 | Q9R0P3 | S-formylglu | Esd | sp\|Q9R0P3 | 17 | 17 | 87,9 |
| Q8BML9 | Q8BML9 | | Qars | sp\|Q8BML | 39 | 39 | 64,8 |
| Q62351 | Q62351 | Transferrin | Tfrc | sp\|Q62351 | 30 | 30 | 47,8 |
| Q60953 | Q60953 | Protein PM | Pml | sp\|Q60953 | 24 | 24 | 40,3 |
| Q8BJ05 | Q8BJ05 | Zinc finger | Zc3h14 | sp\|Q8BJ05 | 21 | 21 | 38,9 |
| Q33DR3 | Q33DR3 | Decapreny | Pdss2 | sp\|Q33DR3 | 7 | 7 | 24,2 |
| Q569Z5 | Q569Z5 | Probable A | Ddx46 | sp\|Q569Z5 | 44 | 44 | 41,7 |
| Q8C0E3 | Q8C0E3 | Tripartite r | Trim47 | sp\|Q8C0E3 | 16 | 16 | 39,8 |
| Q91VD9 | Q91VD9 | NADH-ubic | Ndufs1 | sp\|Q91VD | 28 | 28 | 50,1 |
| Q9WV03;Q | Q9WV03 | Protein FA | Fam50a | sp\|Q9WV0 | 11 | 11 | 38,9 |
| Q8BKX6 | Q8BKX6 | Serine/thre | Smg1 | sp\|Q8BKX6 | 50 | 50 | 20,6 |
| O08739;Q | O08739 | AMP deam | Ampd3 | sp\|O08739 | 25 | 25 | 38,4 |
| P48760 | P48760 | Folylpolygl | Fpgs | sp\|P48760 | 11 | 11 | 27,3 |
| O08580 | O08580 | Steroid ho | Esrra | sp\|O08580 | 5 | 5 | 20,4 |
| Q8BHL4 | Q8BHL4 | Retinoic ac | Gprc5a | sp\|Q8BHL4 | 5 | 5 | 26,7 |
| P59997 | P59997 | Lysine-spe | Kdm2a | sp\|P59997 | 21 | 21 | 23,7 |
| Q9JLB0;RE | Q9JLB0 | MAGUK p5 | Mpp6 | sp\|Q9JLB0 | 26 | 25 | 58,2 |
| Q8VI94 | Q8VI94 | 2-5-oligoac | Oasl1 | sp\|Q8VI94 | 5 | 5 | 14,9 |
| Q8K2V6 | Q8K2V6 | Importin-1 | Ipo11 | sp\|Q8K2V6 | 16 | 16 | 20,3 |
| Q8R080 | Q8R080 | G2 and S p | Gtse1 | sp\|Q8R080 | 17 | 17 | 33,1 |
| Q91V64 | Q91V64 | Isochorism | Isoc1 | sp\|Q91V64 | 7 | 7 | 33,3 |
| Q91X52 | Q91X52 | L-xylulose | Dcxr | sp\|Q91X52 | 2 | 2 | 9,4 |
| Q9D8S3 | Q9D8S3 | ADP-ribosy | Arfgap3 | sp\|Q9D8S3 | 14 | 14 | 35,4 |
| Q9CYY7 | Q9CYY7 | Protein slo | Slmo2 | sp\|Q9CYY7 | 5 | 5 | 32,8 |
| O88792 | O88792 | Junctional | F11r | sp\|O88792 | 8 | 8 | 46,3 |
| Q1W617 | Q1W617 | Protein Sh | Shroom4 | sp\|Q1W61 | 8 | 8 | 7,5 |
| Q8BY02 | Q8BY02 | NF-kappa-I | Nkrf | sp\|Q8BY02 | 19 | 19 | 36,1 |
| Q8BIW1 | Q8BIW1 | Protein pru | Prune | sp\|Q8BIW | 10 | 10 | 30,6 |
| Q91VN6 | Q91VN6 | Probable A | Ddx41 | sp\|Q91VN | 25 | 25 | 45,2 |
| Q9R0M4 | Q9R0M4 | Podocalyxi | Podxl | sp\|Q9R0M | 8 | 8 | 19,7 |
| Q68FH0;O | Q68FH0 | Plakophilin | Pkp4 | sp\|Q68FH0 | 25 | 25 | 30,9 |
| Q6P9Z1 | Q6P9Z1 | SWI/SNF-r | Smarcd3 | sp\|Q6P9Z1 | 18 | 15 | 49,9 |
| P21956 | P21956 | Lactadheri | Mfge8 | sp\|P21956 | 15 | 15 | 38 |
| Q9WUA2 | Q9WUA2 | Phenylalan | Farsb | sp\|Q9WUA | 26 | 26 | 52,8 |

| | | | | | | | |
|---|---|---|---|---|---|---|---|
| Q7TSH6 | Q7TSH6 | | Scaf4 | sp\|Q7TSH6 | 16 | 13 | 20,8 |
| Q3TB82 | Q3TB82 | Pleckstrin | Plekhf1 | sp\|Q3TB82 | 7 | 7 | 36,9 |
| P16546 | P16546 | Spectrin al | Sptan1 | sp\|P16546 | 153 | 152 | 66,6 |
| Q9CZX0 | Q9CZX0 | Elongator | Elp3 | sp\|Q9CZX0 | 15 | 15 | 36,2 |
| Q62422 | Q62422 | Osteoclast | Ostf1 | sp\|Q62422 | 9 | 9 | 53,5 |
| Q9EQG7 | Q9EQG7 | Ectonucle | Enpp5 | sp\|Q9EQG | 7 | 7 | 25,8 |
| Q9Z0X1 | Q9Z0X1 | Apoptosis- | Aifm1 | sp\|Q9Z0X1 | 25 | 25 | 52 |
| Q9QZQ8;Q | Q9QZQ8 | Core histo | H2afy | sp\|Q9QZQ | 18 | 18 | 59,7 |
| Q6NVE8 | Q6NVE8 | WD repeat | Wdr44 | sp\|Q6NVE | 23 | 23 | 40,1 |
| Q8CAY6 | Q8CAY6 | Acetyl-CoA | Acat2 | sp\|Q8CAY6 | 11 | 11 | 45,8 |
| Q9EP71 | Q9EP71 | Ankycorbir | Rai14 | sp\|Q9EP71 | 36 | 36 | 47 |
| O88983 | O88983 | Syntaxin-8 | Stx8 | sp\|O88983 | 7 | 7 | 38,1 |
| Q8K2Z2 | Q8K2Z2 | Pre-mRNA- | Prpf39 | sp\|Q8K2Z2 | 16 | 16 | 29 |
| Q6PIP5 | Q6PIP5 | NudC dom | Nudcd1 | sp\|Q6PIP5 | 27 | 27 | 60,5 |
| Q6PG16 | Q6PG16 | Holliday ju | Hjurp | sp\|Q6PG16 | 14 | 14 | 29,1 |
| Q9DC50 | Q9DC50 | Peroxisom | Crot | sp\|Q9DC50 | 15 | 15 | 32,8 |
| P23475 | P23475 | X-ray repai | Xrcc6 | sp\|P23475 | 26 | 26 | 49,2 |
| P05063 | P05063 | Fructose-b | Aldoc | sp\|P05063 | 13 | 7 | 47,7 |
| P56677 | P56677 | Suppresso | St14 | sp\|P56677 | 12 | 12 | 22 |
| P70399 | P70399 | Tumor sup | Tp53bp1 | sp\|P70399 | 43 | 43 | 41,7 |
| P39428 | P39428 | TNF recept | Traf1 | sp\|P39428 | 7 | 7 | 24,9 |
| Q80T69 | Q80T69 | Round spe | Rsbn1 | sp\|Q80T69 | 14 | 14 | 24,5 |
| Q3UH06 | Q3UH06 | Ras-respor | Rreb1 | sp\|Q3UH0 | 12 | 12 | 12,5 |
| D2EAC2 | D2EAC2 | Zinc finger | Zbed6 | sp\|D2EAC2 | 12 | 12 | 19,8 |
| Q3UV55 | Q3UV55 | Nuclear re | Nr1d1 | sp\|Q3UV5 | 7 | 6 | 17,2 |
| Q03265 | Q03265 | ATP syntha | Atp5a1 | sp\|Q03265 | 32 | 32 | 59,1 |
| F8VPU2 | F8VPU2 | FERM, Rho | Farp1 | sp\|F8VPU2 | 20 | 20 | 29,3 |
| Q9WU42 | Q9WU42 | Nuclear re | Ncor2 | sp\|Q9WU4 | 60 | 59 | 35,6 |
| Q8BVU5 | Q8BVU5 | ADP-ribose | Nudt9 | sp\|Q8BVU | 9 | 9 | 34,3 |
| Q62470 | Q62470 | Integrin al | Itga3 | sp\|Q62470 | 32 | 32 | 39,7 |
| Q91YP2 | Q91YP2 | Neurolysin | Nln | sp\|Q91YP2 | 26 | 26 | 45 |
| Q9D4H1 | Q9D4H1 | Exocyst co | Exoc2 | sp\|Q9D4H | 15 | 15 | 19,4 |
| Q8JZR0 | Q8JZR0 | Long-chain | Acsl5 | sp\|Q8JZR0 | 25 | 22 | 47,7 |
| A2BE28 | A2BE28 | Ribosomal | Las1l | sp\|A2BE28 | 24 | 24 | 45,4 |
| Q99JY4 | Q99JY4 | TraB doma | Trabd | sp\|Q99JY4 | 9 | 9 | 34,6 |
| Q9WVG6 | Q9WVG6 | Histone-ar | Carm1 | sp\|Q9WV | 18 | 18 | 39,1 |
| Q9D0L4 | Q9D0L4 | Uncharact | Adck1 | sp\|Q9D0L4 | 11 | 11 | 32 |
| Q6NVG1 | Q6NVG1 | Lysophosp | Lpcat4 | sp\|Q6NVG | 11 | 11 | 40,6 |
| B2RY04 | B2RY04 | Dedicator | Dock5 | sp\|B2RY04 | 39 | 38 | 28,9 |
| Q91ZN5 | Q91ZN5 | Adenosine | Slc35b2 | sp\|Q91ZN5 | 6 | 6 | 17,2 |
| P51655 | P51655 | Glypican-4 | Gpc4 | sp\|P51655 | 16 | 14 | 41,3 |
| Q99JW5 | Q99JW5 | Epithelial | Epcam | sp\|Q99JW | 4 | 4 | 28,3 |
| O35955 | O35955 | Proteasom | Psmb10 | sp\|O35955 | 6 | 6 | 29,7 |
| Q9DBJ3 | Q9DBJ3 | Brain-spec | Baiap2l1 | sp\|Q9DBJ3 | 15 | 15 | 42 |
| Q5SSW2 | Q5SSW2 | Proteasom | Psme4 | sp\|Q5SSW | 39 | 39 | 25,7 |
| Q8BIJ6 | Q8BIJ6 | Isoleucine- | Iars2 | sp\|Q8BIJ6 | 34 | 34 | 44,4 |
| P17427 | P17427 | AP-2 comp | Ap2a2 | sp\|P17427 | 36 | 28 | 47,5 |
| Q80YS6;Q8 | Q80YS6 | Actin filam | Afap1 | sp\|Q80YS6 | 16 | 16 | 29 |
| Q8CIN4 | Q8CIN4 | Serine/thr | Pak2 | sp\|Q8CIN4 | 28 | 21 | 60,5 |
| Q5H8C4 | Q5H8C4 | Vacuolar p | Vps13a | sp\|Q5H8C | 44 | 44 | 19,8 |

| | | | | | | | |
|---|---|---|---|---|---|---|---|
| Q62448 | Q62448 | Eukaryotic | Eif4g2 | sp\|Q62448 | 41 | 41 | 52,4 |
| Q6KAR6 | Q6KAR6 | Exocyst co | Exoc3 | sp\|Q6KAR6 | 21 | 21 | 34,7 |
| Q99LJ7 | Q99LJ7 | RCC1 and E | Rcbtb2 | sp\|Q99LJ7 | 10 | 10 | 20,1 |
| P97310 | P97310 | DNA replic | Mcm2 | sp\|P97310 | 41 | 41 | 55,2 |
| P24270 | P24270 | Catalase | Cat | sp\|P24270 | 25 | 25 | 61,9 |
| Q60604 | Q60604 | Adseverin | Scin | sp\|Q60604 | 27 | 27 | 52,7 |
| P03975 | P03975 | IgE-binding | Iap | sp\|P03975 | 26 | 26 | 46,9 |
| Q91WT8;P | Q91WT8 | RNA-bindir | Rbm47 | sp\|Q91WT | 13 | 12 | 31,4 |
| P52332 | P52332 | Tyrosine-p | Jak1 | sp\|P52332 | 27 | 26 | 32,4 |
| O09172 | O09172 | Glutamate | Gclm | sp\|O09172 | 7 | 7 | 32,8 |
| Q8C2E4 | Q8C2E4 | Pentatrico | Ptcd1 | sp\|Q8C2E4 | 13 | 13 | 25,2 |
| A2AKG8 | A2AKG8 | Focadhesin | Focad | sp\|A2AKG8 | 25 | 25 | 17,7 |
| Q640N1;Q | Q640N1 | Adipocyte | Aebp1 | sp\|Q640N1 | 18 | 18 | 21,5 |
| P70175 | P70175 | Disks large | Dlg3 | sp\|P70175 | 10 | 8 | 16,1 |
| Q9ERU9 | Q9ERU9 | E3 SUMO-p | Ranbp2 | sp\|Q9ERU9 | 106 | 106 | 48,3 |
| Q925I1 | Q925I1 | ATPase fan | Atad3 | sp\|Q925I1 | 24 | 24 | 46 |
| Q8BFR5 | Q8BFR5 | Elongation | Tufm | sp\|Q8BFR5 | 26 | 26 | 68,8 |
| Q9CWX4 | Q9CWX4 | RNA pseud | Rpusd4 | sp\|Q9CWX | 9 | 9 | 29,7 |
| Q99L27 | Q99L27 | GMP reduc | Gmpr2 | sp\|Q99L27 | 7 | 7 | 27,9 |
| Q8CIG8 | Q8CIG8 | Protein arg | Prmt5 | sp\|Q8CIG8 | 22 | 22 | 44,7 |
| Q505D7 | Q505D7 | Optic atrop | Opa3 | sp\|Q505D7 | 3 | 3 | 15,6 |
| Q8VE95 | Q8VE95 | UPF0598 protein C8or | | sp\|Q8VE95 | 6 | 6 | 43,6 |
| Q9JLC8 | Q9JLC8 | Sacsin | Sacs | sp\|Q9JLC8 | 48 | 48 | 13,6 |
| O35114 | O35114 | Lysosome | Scarb2 | sp\|O35114 | 12 | 12 | 33,5 |
| Q99LB7 | Q99LB7 | Sarcosine d | Sardh | sp\|Q99LB7 | 8 | 8 | 12,1 |
| Q8VDG3 | Q8VDG3 | Poly(A)-spe | Parn | sp\|Q8VDG | 15 | 15 | 29,5 |
| P31324 | P31324 | cAMP-depe | Prkar2b | sp\|P31324 | 15 | 13 | 51,9 |
| Q8K019 | Q8K019 | Bcl-2-asso | Bclaf1 | sp\|Q8K019 | 24 | 23 | 27,4 |
| Q91WM1 | Q91WM1 | Spermatid | Strbp | sp\|Q91WM | 21 | 17 | 36,2 |
| P14094 | P14094 | Sodium/po | Atp1b1 | sp\|P14094 | 8 | 8 | 27,3 |
| Q6Q899 | Q6Q899 | Probable A | Ddx58 | sp\|Q6Q89 | 22 | 22 | 27,9 |
| CON__Q3S | CON__Q3SZ57 | | | | 23 | 23 | 54,8 |
| Q9Z1F9 | Q9Z1F9 | SUMO-acti | Uba2 | sp\|Q9Z1F9 | 33 | 33 | 68 |
| Q8K5C0 | Q8K5C0 | Grainyhead | Grhl2 | sp\|Q8K5C0 | 10 | 10 | 25,1 |
| Q9WVL0 | Q9WVL0 | Maleylacet | Gstz1 | sp\|Q9WVL | 9 | 9 | 60,6 |
| P13864 | P13864 | DNA (cytos | Dnmt1 | sp\|P13864 | 71 | 71 | 54,6 |
| Q07813 | Q07813 | Apoptosis | Bax | sp\|Q07813 | 10 | 10 | 60,4 |
| Q9DBY8 | Q9DBY8 | Nuclear va | Nvl | sp\|Q9DBY8 | 37 | 37 | 56 |
| Q3TDN2 | Q3TDN2 | FAS-associ | Faf2 | sp\|Q3TDN | 17 | 17 | 57,1 |
| G5E8K5;Q0 | G5E8K5 | Ankyrin-3 | Ank3 | sp\|G5E8K5 | 31 | 26 | 22,8 |
| Q3TJZ6 | Q3TJZ6 | Protein FA | Fam98a | sp\|Q3TJZ6 | 11 | 8 | 27,2 |
| Q9CWS4 | Q9CWS4 | Integrator | Cpsf3l | sp\|Q9CWS | 9 | 9 | 21,3 |
| Q91Z31 | Q91Z31 | Polypyrimi | Ptbp2 | sp\|Q91Z31 | 18 | 18 | 60,6 |
| Q920E5 | Q920E5 | Farnesyl py | Fdps | sp\|Q920E5 | 11 | 11 | 36,8 |
| Q9WU63 | Q9WU63 | Heme-bind | Hebp2 | sp\|Q9WU6 | 5 | 5 | 38 |
| Q3TIR3 | Q3TIR3 | Synembryn | Ric8a | sp\|Q3TIR3 | 16 | 15 | 34,3 |
| P24288 | P24288 | Branched-c | Bcat1 | sp\|P24288 | 8 | 8 | 26,9 |
| Q8R016 | Q8R016 | Bleomycin | Blmh | sp\|Q8R016 | 18 | 18 | 46,4 |
| Q8R2U2 | Q8R2U2 | | BC027231 | sp\|Q8R2U | 8 | 8 | 23 |
| O09061 | O09061 | Proteasom | Psmb1 | sp\|O09061 | 13 | 13 | 76,7 |

| | | | | | | | |
|---|---|---|---|---|---|---|---|
| Q8CH77 | Q8CH77 | Neuron na | Nav1 | sp\|Q8CH7 | 34 | 33 | 29,5 |
| F6ZDS4 | F6ZDS4 | Nucleopro | Tpr | sp\|F6ZDS4 | 97 | 97 | 42,3 |
| CON__Q9T | CON__Q9TTE1 | | | | 11 | 4 | 31,4 |
| Q80VI1 | Q80VI1 | E3 ubiquiti | Trim56 | sp\|Q80VI1 | 21 | 21 | 45,4 |
| Q9D8X5 | Q9D8X5 | CCR4-NOT | Cnot8 | sp\|Q9D8X | 5 | 5 | 26 |
| Q6P9L6 | Q6P9L6 | Kinesin-like | Kif15 | sp\|Q6P9L6 | 39 | 39 | 36,6 |
| P15626;O3 | P15626 | Glutathion | Gstm2 | sp\|P15626 | 16 | 11 | 68,3 |
| Q14AI0 | Q14AI0 | Sister chro | DSCC1 | sp\|Q14AI0 | 10 | 10 | 32,8 |
| Q8CG19 | Q8CG19 | Latent-tran | Ltbp1 | sp\|Q8CG1 | 17 | 17 | 15,2 |
| Q8CJ26 | Q8CJ26 | Death dom | Nradd | sp\|Q8CJ26 | 5 | 5 | 28,5 |
| Q91YW3 | Q91YW3 | DnaJ homo | Dnajc3 | sp\|Q91YW | 19 | 19 | 43,5 |
| Q60676 | Q60676 | Serine/thre | Ppp5c | sp\|Q60676 | 15 | 15 | 38,9 |
| CON__P34 | CON__P34955 | | | | 16 | 16 | 33,9 |
| P11835 | P11835 | Integrin be | Itgb2 | sp\|P11835 | 18 | 18 | 31,9 |
| Q9Z184 | Q9Z184 | Protein-arg | Padi3 | sp\|Q9Z184 | 11 | 8 | 18,7 |
| Q9CQI6 | Q9CQI6 | Coactosin- | Cotl1 | sp\|Q9CQI6 | 5 | 5 | 42,3 |
| Q80ZQ9 | Q80ZQ9 | Protein Sin | Fam206a | sp\|Q80ZQ | 8 | 8 | 65,8 |
| Q6P6M7 | Q6P6M7 | O-phospho | Sepsecs | sp\|Q6P6M | 6 | 6 | 16,3 |
| P11103 | P11103 | Poly [ADP- | Parp1 | sp\|P11103 | 34 | 34 | 34,9 |
| Q61033 | Q61033 | Lamina-ass | Tmpo | sp\|Q61033 | 33 | 23 | 63,3 |
| Q8K4G5 | Q8K4G5 | Actin-bindi | Ablim1 | sp\|Q8K4G | 18 | 18 | 26,9 |
| Q3UMF0 | Q3UMF0 | Cordon-ble | Cobll1 | sp\|Q3UMF | 43 | 43 | 50,4 |
| Q99KN1 | Q99KN1 | Arrestin do | Arrdc1 | sp\|Q99KN | 5 | 5 | 18,7 |
| Q8BM72 | Q8BM72 | Heat shock | Hspa13 | sp\|Q8BM7 | 13 | 13 | 29,5 |
| O09131;Q8 | O09131 | Glutathion | Gsto1 | sp\|O09131 | 17 | 17 | 70 |
| Q3TVI4 | Q3TVI4 | Protein HE | Hexim2 | sp\|Q3TVI4 | 5 | 5 | 24,3 |
| Q8CJF7 | Q8CJF7 | Protein ELY | Ahctf1 | sp\|Q8CJF7 | 76 | 76 | 47,8 |
| Q9CYR6 | Q9CYR6 | Phosphoac | Pgm3 | sp\|Q9CYR6 | 14 | 14 | 34,1 |
| Q8CGF7;Q | Q8CGF7 | Transcripti | Tcerg1 | sp\|Q8CGF | 35 | 35 | 29,3 |
| E9QAM5 | E9QAM5 | Helicase w | Helz2 | sp\|E9QAM | 14 | 14 | 7,4 |
| Q6ZPR6 | Q6ZPR6 | Inhibitor o | Ibtk | sp\|Q6ZPR6 | 16 | 16 | 16,1 |
| Q8BKT7 | Q8BKT7 | THO comp | Thoc5 | sp\|Q8BKT7 | 21 | 21 | 38,1 |
| Q9JIK5 | Q9JIK5 | Nucleolar R | Ddx21 | sp\|Q9JIK5 | 55 | 55 | 65,9 |
| Q61735 | Q61735 | Leukocyte | Cd47 | sp\|Q61735 | 6 | 6 | 20,1 |
| Q9CQE1 | Q9CQE1 | Protein Nip | Nipsnap3b | sp\|Q9CQE | 8 | 8 | 40,1 |
| A2AB59 | A2AB59 | Rho GTPas | Arhgap27 | sp\|A2AB59 | 12 | 12 | 24,4 |
| Q9WUA3 | Q9WUA3 | ATP-depen | Pfkp | sp\|Q9WUA | 26 | 23 | 44,5 |
| Q9Z0V7 | Q9Z0V7 | Mitochond | Timm17b | sp\|Q9Z0V7 | 3 | 3 | 30,8 |
| P09055 | P09055 | Integrin be | Itgb1 | sp\|P09055 | 25 | 25 | 39,3 |
| P29351 | P29351 | Tyrosine-p | Ptpn6 | sp\|P29351 | 18 | 18 | 42,5 |
| Q9WV85 | Q9WV85 | Nucleoside | Nme3 | sp\|Q9WV8 | 4 | 4 | 27,2 |
| Q8C033 | Q8C033 | Rho guanin | Arhgef10 | sp\|Q8C033 | 6 | 5 | 6,2 |
| Q3TMW1 | Q3TMW1 | Coiled-coil | Ccdc102a | sp\|Q3TMV | 9 | 9 | 23,1 |
| Q6ZPE2 | Q6ZPE2 | Myotubula | Sbf1 | sp\|Q6ZPE2 | 35 | 34 | 26,3 |
| Q60767 | Q60767 | Lymphocyt | Ly75 | sp\|Q60767 | 40 | 40 | 31,5 |
| B9EJR8 | B9EJR8 | Dynein ass | Dnaaf5 | sp\|B9EJR8 | 18 | 18 | 34,5 |
| Q99MR8 | Q99MR8 | Methylcrot | Mccc1 | sp\|Q99MR | 20 | 20 | 44,1 |
| Q8K268 | Q8K268 | ATP-bindin | Abcf3 | sp\|Q8K268 | 18 | 18 | 34,8 |
| Q99NH0 | Q99NH0 | Ankyrin re | Ankrd17 | sp\|Q99NH | 45 | 45 | 25,9 |
| Q8VBT6 | Q8VBT6 | Apolipopro | Apobr | sp\|Q8VBT6 | 26 | 26 | 40,9 |

| | | | | | | | |
|---|---|---|---|---|---|---|---|
| Q9JHJ3 | Q9JHJ3 | Glycosylate | Glmp | sp\|Q9JHJ3 | 2 | 2 | 11,6 |
| Q61879 | Q61879 | Myosin-10 | Myh10 | sp\|Q61879 | 105 | 84 | 52,1 |
| Q9JMA1 | Q9JMA1 | Ubiquitin c | Usp14 | sp\|Q9JMA | 28 | 28 | 63,1 |
| Q5UAK0 | Q5UAK0 | Mesoderm | Mier1 | sp\|Q5UAK | 11 | 11 | 31,1 |
| O35643 | O35643 | AP-1 comp | Ap1b1 | sp\|O35643 | 39 | 22 | 48 |
| O54724 | O54724 | Polymeras | Ptrf | sp\|O54724 | 18 | 18 | 44,4 |
| Q8CJ40 | Q8CJ40 | Rootletin | Crocc | sp\|Q8CJ40 | 2 | 2 | 0,9 |
| Q8CH18 | Q8CH18 | Cell divisio | Ccar1 | sp\|Q8CH18 | 28 | 28 | 25,7 |
| CON__Q9U | CON__Q9U6Y5 | | | | 11 | 11 | 60,6 |
| Q8BMF4 | Q8BMF4 | Dihydrolip | Dlat | sp\|Q8BMF | 23 | 23 | 48,4 |
| Q8R0J7 | Q8R0J7 | Vacuolar p | Vps37b | sp\|Q8R0J7 | 8 | 8 | 43,9 |
| Q3V0K9 | Q3V0K9 | Plastin-1 | Pls1 | sp\|Q3V0K9 | 13 | 10 | 28,4 |
| Q9QZ08 | Q9QZ08 | N-acetyl-D | Nagk | sp\|Q9QZ08 | 3 | 3 | 13,1 |
| Q9ESZ8 | Q9ESZ8 | General tra | Gtf2i | sp\|Q9ESZ8 | 41 | 40 | 49 |
| Q62312 | Q62312 | TGF-beta r | Tgfbr2 | sp\|Q62312 | 11 | 11 | 23,5 |
| Q9R0M6;C | Q9R0M6 | Ras-relatec | Rab9a | sp\|Q9R0M | 9 | 9 | 55,2 |
| P03911 | P03911 | NADH-ubic | Mtnd4 | sp\|P03911 | 3 | 3 | 9,4 |
| Q3UIA2 | Q3UIA2 | Rho GTPas | Arhgap17 | sp\|Q3UIA2 | 21 | 21 | 37,1 |
| Q9D7E3 | Q9D7E3 | Ovarian ca | Ovca2 | sp\|Q9D7E3 | 7 | 7 | 50,2 |
| Q8C7V8 | Q8C7V8 | Coiled-coil | Ccdc134 | sp\|Q8C7V8 | 5 | 5 | 31,9 |
| Q8CGB3 | Q8CGB3 | Uveal auto | Uaca | sp\|Q8CGB | 19 | 19 | 18,9 |
| Q07076 | Q07076 | Annexin A7 | Anxa7 | sp\|Q07076 | 22 | 22 | 39,1 |
| Q925H1 | Q925H1 | Zinc finger | Trps1 | sp\|Q925H1 | 31 | 31 | 37,2 |
| Q9CWW6 | Q9CWW6 | Peptidyl-pr | Pin4 | sp\|Q9CWV | 5 | 5 | 49,6 |
| Q8BND5 | Q8BND5 | Sulfhydryl | Qsox1 | sp\|Q8BND | 3 | 3 | 4,5 |
| Q6ZPU9 | Q6ZPU9 | KIF1-bindir | Kbp | sp\|Q6ZPU9 | 19 | 19 | 38,7 |
| Q6P5G6 | Q6P5G6 | UBX doma | Ubxn7 | sp\|Q6P5G6 | 11 | 11 | 37,3 |
| Q6PB66 | Q6PB66 | Leucine-ric | Lrpprc | sp\|Q6PB66 | 78 | 78 | 60 |
| D0QMC3 | D0QMC3 | Myeloid ce | Mndal | sp\|D0QMC | 18 | 8 | 37 |
| A2ASQ1 | A2ASQ1 | Agrin;Agrir | Agrn | sp\|A2ASQ1 | 27 | 27 | 21,6 |
| A6H5Z3 | A6H5Z3 | Exocyst co | Exoc6b | sp\|A6H5Z3 | 15 | 14 | 23,2 |
| Q9DBS1 | Q9DBS1 | Transmem | Tmem43 | sp\|Q9DBS1 | 16 | 16 | 52 |
| Q99PG2 | Q99PG2 | Opioid gro | Ogfr | sp\|Q99PG2 | 16 | 16 | 52,1 |
| Q8VCS3 | Q8VCS3 | Glycosami | Fam20b | sp\|Q8VCS3 | 10 | 10 | 34 |
| Q64727 | Q64727 | Vinculin | Vcl | sp\|Q64727 | 61 | 61 | 65,6 |
| Q9CRC9 | Q9CRC9 | Glucosami | Gnpda2 | sp\|Q9CRC9 | 7 | 6 | 37,3 |
| P29758 | P29758 | Ornithine a | Oat | sp\|P29758 | 22 | 22 | 68,3 |
| O70475 | O70475 | UDP-gluco | Ugdh | sp\|O70475 | 31 | 31 | 78,5 |
| Q9CYL5 | Q9CYL5 | Golgi-asso | Glipr2 | sp\|Q9CYL5 | 8 | 8 | 64,9 |
| Q8R5A3 | Q8R5A3 | Amyloid be | Apbb1ip | sp\|Q8R5A3 | 16 | 16 | 31,9 |
| Q80TP3 | Q80TP3 | E3 ubiquiti | Ubr5 | sp\|Q80TP3 | 66 | 66 | 33,6 |
| Q2KN98 | Q2KN98 | Cytospin-A | Specc1l | sp\|Q2KN98 | 24 | 24 | 28,9 |
| Q8CHG3 | Q8CHG3 | GRIP and c | Gcc2 | sp\|Q8CHG | 34 | 34 | 26,1 |
| Q9CXR1 | Q9CXR1 | Dehydroge | Dhrs7 | sp\|Q9CXR1 | 7 | 7 | 21,6 |
| Q9ES46 | Q9ES46 | Beta-parvi | Parvb | sp\|Q9ES46 | 9 | 8 | 31,5 |
| Q8CGB6 | Q8CGB6 | Tensin-2 | Tns2 | sp\|Q8CGB | 23 | 21 | 26,6 |
| Q8R1G2 | Q8R1G2 | Carboxyme | Cmbl | sp\|Q8R1G | 2 | 2 | 9 |
| P46978 | P46978 | Dolichyl-di | Stt3a | sp\|P46978 | 17 | 16 | 26,7 |
| Q9DAR7 | Q9DAR7 | m7GpppX | Dcps | sp\|Q9DAR | 16 | 16 | 58 |
| Q91YP3 | Q91YP3 | Deoxyribos | Dera | sp\|Q91YP3 | 10 | 10 | 45,9 |

| | | | | | | | |
|---|---|---|---|---|---|---|---|
| Q9CZU3 | Q9CZU3 | Superkiller | Skiv2l2 | sp\|Q9CZU3 | 46 | 46 | 49,4 |
| Q8BTG3 | Q8BTG3 | T-complex | Tcp11l1 | sp\|Q8BTG3 | 7 | 7 | 17,1 |
| E9Q394 | E9Q394 | | Akap13 | sp\|E9Q394 | 27 | 27 | 17,4 |
| Q9Z2I9 | Q9Z2I9 | Succinyl-Co | Sucla2 | sp\|Q9Z2I9 | 20 | 20 | 48,8 |
| Q9WU40 | Q9WU40 | Inner nucle | Lemd3 | sp\|Q9WU4 | 26 | 26 | 42,8 |
| Q9Z2D1 | Q9Z2D1 | Myotubula | Mtmr2 | sp\|Q9Z2D1 | 21 | 21 | 44,3 |
| P13020 | P13020 | Gelsolin | Gsn | sp\|P13020 | 32 | 19 | 48,1 |
| Q9DBG7 | Q9DBG7 | Signal reco | Srpr | sp\|Q9DBG | 26 | 26 | 51,3 |
| Q80X41 | Q80X41 | Serine/thre | Vrk1 | sp\|Q80X41 | 16 | 16 | 53,2 |
| P46062 | P46062 | Signal-indu | Sipa1 | sp\|P46062 | 13 | 13 | 22 |
| P52480 | P52480 | Pyruvate k | Pkm | sp\|P52480 | 46 | 45 | 79,8 |
| Q8CJ67 | Q8CJ67 | Double-str | Stau2 | sp\|Q8CJ67 | 10 | 10 | 26 |
| Q91UZ5 | Q91UZ5 | Inositol mo | Impa2 | sp\|Q91UZ5 | 7 | 7 | 31 |
| Q9JJ89 | Q9JJ89 | Coiled-coil | Ccdc86 | sp\|Q9JJ89 | 13 | 13 | 33,1 |
| Q5SVQ0 | Q5SVQ0 | Histone ac | Kat7 | sp\|Q5SVQ | 20 | 19 | 41,1 |
| Q8CHP8 | Q8CHP8 | Phosphogly | Pgp | sp\|Q8CHP | 11 | 11 | 44,5 |
| O70439 | O70439 | Syntaxin-7 | Stx7 | sp\|O70439 | 7 | 7 | 41,4 |
| P07091 | P07091 | Protein S10 | S100a4 | sp\|P07091 | 8 | 8 | 84,2 |
| Q8BHT6 | Q8BHT6 | Beta-1,3-gl | B3galtl | sp\|Q8BHT | 10 | 10 | 27,2 |
| Q3TJ91 | Q3TJ91 | Lethal(2) g | Llgl2 | sp\|Q3TJ91 | 20 | 20 | 29,6 |
| P97798 | P97798 | Neogenin | Neo1 | sp\|P97798 | 17 | 17 | 18,2 |
| Q61097 | Q61097 | Kinase sup | Ksr1 | sp\|Q61097 | 3 | 3 | 4 |
| Q8VCH6 | Q8VCH6 | Delta(24)-s | Dhcr24 | sp\|Q8VCH | 8 | 8 | 18,6 |
| O55091 | O55091 | Protein IM | Impact | sp\|O55091 | 7 | 7 | 30,8 |
| P60521 | P60521 | Gamma-an | Gabarapl2 | sp\|P60521 | 6 | 5 | 41,9 |
| Q9CY27 | Q9CY27 | Very-long-c | Tecr | sp\|Q9CY27 | 11 | 11 | 26,6 |
| Q6ZPJ3 | Q6ZPJ3 | E2/E3 hybr | Ube2o | sp\|Q6ZPJ3 | 34 | 34 | 38,2 |
| P15116;P3 | P15116 | Cadherin-2 | Cdh2 | sp\|P15116 | 17 | 17 | 29,8 |
| P47740 | P47740 | Fatty aldeh | Aldh3a2 | sp\|P47740 | 18 | 16 | 38,6 |
| Q9WUR9 | Q9WUR9 | Adenylate | Ak4 | sp\|Q9WUR | 7 | 7 | 49,3 |
| P25911;P1 | P25911 | Tyrosine-p | Lyn | sp\|P25911 | 20 | 18 | 49,6 |
| Q61136 | Q61136 | Serine/thre | Prpf4b | sp\|Q61136 | 23 | 23 | 23,8 |
| Q8CBH5 | Q8CBH5 | Major facil | Mfsd6 | sp\|Q8CBH | 3 | 3 | 6,6 |
| Q9CQ92 | Q9CQ92 | Mitochond | Fis1 | sp\|Q9CQ9 | 6 | 6 | 42,1 |
| Q5EG47;Q | Q5EG47 | 5-AMP-act | Prkaa1 | sp\|Q5EG47 | 18 | 18 | 44,4 |
| Q8BX70 | Q8BX70 | Vacuolar p | Vps13c | sp\|Q8BX70 | 67 | 67 | 24,6 |
| Q6DFV5 | Q6DFV5 | Probable h | Helz | sp\|Q6DFV5 | 20 | 20 | 13,5 |
| Q9Z1B7 | Q9Z1B7 | Mitogen-ac | Mapk13 | sp\|Q9Z1B7 | 13 | 12 | 43,7 |
| P19182 | P19182 | Interferon- | Ifrd1 | sp\|P19182 | 11 | 11 | 28,3 |
| Q9JI78 | Q9JI78 | Peptide-N( | Ngly1 | sp\|Q9JI78 | 6 | 6 | 15,8 |
| Q9D937 | Q9D937 | Uncharacterized prote | | sp\|Q9D937 | 5 | 5 | 33,3 |
| Q8BTI8 | Q8BTI8 | Serine/arg | Srrm2 | sp\|Q8BTI8 | 53 | 53 | 27,7 |
| Q9D771 | Q9D771 | Transmem | Tmem206 | sp\|Q9D771 | 7 | 7 | 25,4 |
| P54227;P6 | P54227 | Stathmin | Stmn1 | sp\|P54227 | 10 | 7 | 49 |
| Q0KL02 | Q0KL02 | Triple func | Trio | sp\|Q0KL02 | 41 | 38 | 17,2 |
| Q80XC3 | Q80XC3 | USP6 N-ter | Usp6nl | sp\|Q80XC3 | 8 | 8 | 13,2 |
| P20664 | P20664 | DNA prima | Prim1 | sp\|P20664 | 19 | 19 | 49,4 |
| P23242 | P23242 | Gap juncti | Gja1 | sp\|P23242 | 14 | 14 | 41,1 |
| A2CG63 | A2CG63 | AT-rich inte | Arid4b | sp\|A2CG63 | 14 | 13 | 14,1 |
| Q9Z0S1 | Q9Z0S1 | 3(2),5-bisp | Bpnt1 | sp\|Q9Z0S1 | 12 | 12 | 50,3 |

| | | | | | | | |
|---|---|---|---|---|---|---|---|
| P23249 | P23249 | Putative he | Mov10 | sp\|P23249 | 33 | 33 | 39,5 |
| O35231 | O35231 | Kinesin-like | Kifc3 | sp\|O35231 | 9 | 8 | 16,5 |
| Q9DBT5 | Q9DBT5 | AMP deam | Ampd2 | sp\|Q9DBT5 | 20 | 20 | 27,7 |
| Q9D1E6 | Q9D1E6 | Tubulin-fol | Tbcb | sp\|Q9D1E6 | 11 | 11 | 45,5 |
| Q8K3C3 | Q8K3C3 | Protein LZI | Lzic | sp\|Q8K3C3 | 7 | 7 | 48,9 |
| P21550 | P21550 | Beta-enola | Eno3 | sp\|P21550 | 21 | 15 | 59,4 |
| P35700 | P35700 | Peroxiredo | Prdx1 | sp\|P35700 | 17 | 16 | 69,8 |
| Q8BWU5 | Q8BWU5 | Probable t | Osgep | sp\|Q8BWU | 9 | 9 | 37 |
| Q9JIM1 | Q9JIM1 | Equilibrativ | Slc29a1 | sp\|Q9JIM1 | 7 | 7 | 17,4 |
| Q9CWP6 | Q9CWP6 | Motile spe | Mospd2 | sp\|Q9CWP | 8 | 8 | 16,8 |
| Q80V53 | Q80V53 | Carbohydr | Chst14 | sp\|Q80V53 | 7 | 7 | 22,9 |
| O70228 | O70228 | Probable p | Atp9a | sp\|O70228 | 13 | 11 | 19 |
| Q80UW2 | Q80UW2 | F-box only | Fbxo2 | sp\|Q80UW | 6 | 6 | 27,3 |
| P10922 | P10922 | Histone H1 | H1f0 | sp\|P10922 | 8 | 8 | 28,4 |
| Q9DCD0 | Q9DCD0 | 6-phospho | Pgd | sp\|Q9DCD | 21 | 21 | 53 |
| Q922S4 | Q922S4 | cGMP-dep | Pde2a | sp\|Q922S4 | 22 | 22 | 33 |
| Q8VHX6 | Q8VHX6 | Filamin-C | Flnc | sp\|Q8VHX | 119 | 106 | 59,6 |
| Q91YN9 | Q91YN9 | BAG family | Bag2 | sp\|Q91YN9 | 11 | 11 | 55,7 |
| O08539;D3 | O08539 | Myc box-d | Bin1 | sp\|O08539 | 18 | 18 | 39,6 |
| Q8R3Q6 | Q8R3Q6 | Coiled-coil | Ccdc58 | sp\|Q8R3Q | 9 | 9 | 66,7 |
| Q6KCD5 | Q6KCD5 | Nipped-B-l | Nipbl | sp\|Q6KCD | 51 | 51 | 24,2 |
| Q91VR5 | Q91VR5 | ATP-depen | Ddx1 | sp\|Q91VR5 | 41 | 41 | 64,5 |
| Q8BZS9 | Q8BZS9 | Putative pr | Dhx32 | sp\|Q8BZS9 | 9 | 9 | 16,7 |
| P70459 | P70459 | ETS domai | Erf | sp\|P70459 | 8 | 8 | 27,9 |
| P70451 | P70451 | Tyrosine-p | Fer | sp\|P70451 | 8 | 7 | 13,9 |
| CON__P01 | CON__P01030 | | | | 46 | 3 | 35,6 |
| Q63ZW7 | Q63ZW7 | InaD-like p | Inadl | sp\|Q63ZW | 9 | 8 | 8,2 |
| Q9CZB0 | Q9CZB0 | Succinate d | Sdhc | sp\|Q9CZB0 | 5 | 5 | 29 |
| P02469 | P02469 | Laminin su | Lamb1 | sp\|P02469 | 55 | 55 | 38,6 |
| P33174 | P33174 | Chromoso | Kif4 | sp\|P33174 | 45 | 44 | 47,1 |
| Q99J85 | Q99J85 | Neuronal p | Nptxr | sp\|Q99J85 | 8 | 8 | 29,2 |
| CON__ENS | CON__ENSEMBL:ENSBTAP00000007350 | | | | 57 | 14 | 43,2 |
| Q9CQA3 | Q9CQA3 | Succinate d | Sdhb | sp\|Q9CQA | 12 | 12 | 44 |
| Q8BGX0 | Q8BGX0 | E3 ubiquiti | Trim23 | sp\|Q8BGX0 | 1 | 1 | 1,7 |
| Q04735 | Q04735 | Cyclin-dep | Cdk16 | sp\|Q04735 | 11 | 6 | 25,6 |
| Q8CJ53 | Q8CJ53 | Cdc42-inte | Trip10 | sp\|Q8CJ53 | 17 | 17 | 33 |
| Q8K301 | Q8K301 | Probable A | Ddx52 | sp\|Q8K301 | 18 | 18 | 36,5 |
| Q09XV5 | Q09XV5 | Chromodo | Chd8 | sp\|Q09XV5 | 40 | 30 | 23,8 |
| P29268 | P29268 | Connective | Ctgf | sp\|P29268 | 6 | 6 | 20,7 |
| Q61599 | Q61599 | Rho GDP-d | Arhgdib | sp\|Q61599 | 6 | 6 | 55,5 |
| Q9Z185 | Q9Z185 | Protein-arg | Padi1 | sp\|Q9Z185 | 15 | 15 | 31,6 |
| Q9ESW4 | Q9ESW4 | Acylglycero | Agk | sp\|Q9ESW | 13 | 13 | 47 |
| Q9ESJ0 | Q9ESJ0 | Exportin-4 | Xpo4 | sp\|Q9ESJ0 | 26 | 26 | 31,3 |
| Q99KH8 | Q99KH8 | Serine/thre | Stk24 | sp\|Q99KH8 | 16 | 8 | 42,5 |
| Q9EPT5 | Q9EPT5 | Solute carr | Slco2a1 | sp\|Q9EPT5 | 9 | 9 | 13,4 |
| P70206;P7 | P70206 | Plexin-A1 | Plxna1 | sp\|P70206 | 35 | 32 | 24,8 |
| Q9DCW4 | Q9DCW4 | Electron tr | Etfb | sp\|Q9DCW | 16 | 16 | 67,8 |
| P97384 | P97384 | Annexin A1 | Anxa11 | sp\|P97384 | 20 | 20 | 37,6 |
| Q6NZC7 | Q6NZC7 | SEC23-inte | Sec23ip | sp\|Q6NZC7 | 23 | 23 | 28,7 |
| Q9QX47 | Q9QX47 | Protein SO | Son | sp\|Q9QX4 | 44 | 44 | 27,7 |

| | | | | | | | |
|---|---|---|---|---|---|---|---|
| Q6ZQA0 | Q6ZQA0 | Neurobeac | Nbeal2 | sp\|Q6ZQA | 26 | 26 | 14,6 |
| P18653 | P18653 | Ribosomal | Rps6ka1 | sp\|P18653 | 21 | 14 | 40,3 |
| CON__P02 | CON__P02769 | | | | 54 | 48 | 77,3 |
| P84086;P6 | P84086;P6304 | Complexin | Cplx2;Cplx | sp\|P84086 | 2 | 2 | 19,4 |
| Q8BVY0 | Q8BVY0 | Ribosomal | Rsl1d1 | sp\|Q8BVY( | 26 | 26 | 56,9 |
| Q9CQ06 | Q9CQ06 | 39S riboso | Mrpl24 | sp\|Q9CQ0 | 7 | 7 | 35,2 |
| P47738;Q6 | P47738 | Aldehyde d | Aldh2 | sp\|P47738 | 32 | 30 | 61,1 |
| Q61292 | Q61292 | Laminin su | Lamb2 | sp\|Q6129 | 9 | 9 | 7,3 |
| Q9D2V7 | Q9D2V7 | Coronin-7 | Coro7 | sp\|Q9D2V | 13 | 13 | 23,8 |
| Q8BG51 | Q8BG51 | Mitochonc | Rhot1 | sp\|Q8BG5 | 22 | 21 | 46,6 |
| P39429 | P39429 | TNF recept | Traf2 | sp\|P39429 | 9 | 9 | 25 |
| CON__REF | CON__REFSEQ:XP_585019 | | | | 5 | 5 | 10,4 |
| Q6ZPR5 | Q6ZPR5 | Sphingomy | Smpd4 | sp\|Q6ZPR5 | 25 | 25 | 40,8 |
| Q8K298 | Q8K298 | Actin-bindi | Anln | sp\|Q8K298 | 48 | 48 | 58,6 |
| P23506 | P23506 | Protein-L-i | Pcmt1 | sp\|P23506 | 12 | 12 | 62,1 |
| O54836 | O54836 | Zinc finger | Zmat3 | sp\|O5483( | 6 | 6 | 41,4 |
| Q69ZN7;RI | Q69ZN7 | Myoferlin | Myof | sp\|Q69ZN | 125 | 123 | 66,8 |
| Q8CG72 | Q8CG72 | Poly(ADP-r | Adprhl2 | sp\|Q8CG7 | 9 | 9 | 31,1 |
| Q9JMC3 | Q9JMC3 | DnaJ homo | Dnaja4 | sp\|Q9JMC | 6 | 6 | 20,9 |
| Q8K3Z9 | Q8K3Z9 | Nuclear en | Pom121 | sp\|Q8K3Z | 22 | 22 | 27,6 |
| P35821 | P35821 | Tyrosine-p | Ptpn1 | sp\|P35821 | 16 | 16 | 51,6 |
| Q9QYS9 | Q9QYS9 | Protein qu | Qki | sp\|Q9QYS | 15 | 15 | 48,1 |
| O35368 | O35368 | Interferon- | Ifi203 | sp\|O35368 | 18 | 12 | 46,6 |
| Q9CPP0 | Q9CPP0 | Nucleoplas | Npm3 | sp\|Q9CPP( | 6 | 6 | 47,4 |
| Q62523 | Q62523 | Zyxin | Zyx | sp\|Q6252 | 16 | 16 | 43,6 |
| P09450 | P09450 | Transcripti | Junb | sp\|P09450 | 11 | 11 | 52,3 |
| P14901 | P14901 | Heme oxyg | Hmox1 | sp\|P14901 | 9 | 9 | 49,5 |
| Q9CRT8 | Q9CRT8 | Exportin-T | Xpot | sp\|Q9CRT8 | 29 | 29 | 39,4 |
| Q7TSH2 | Q7TSH2 | Phosphory | Phkb | sp\|Q7TSH | 9 | 9 | 13,3 |
| P49443 | P49443 | Protein ph | Ppm1a | sp\|P49443 | 13 | 10 | 49,7 |
| Q922K9 | Q922K9 | Tyrosine-p | Frk | sp\|Q922K | 14 | 13 | 33,6 |
| Q61466 | Q61466 | SWI/SNF-r | Smarcd1 | sp\|Q6146 | 20 | 18 | 49,1 |
| P56376 | P56376 | Acylphospl | Acyp1 | sp\|P56376 | 6 | 6 | 71,7 |
| O88939 | O88939 | Zinc finger | Zbtb7a | sp\|O8893 | 9 | 9 | 27,6 |
| Q8CC27;P5 | Q8CC27 | Voltage-de | Cacnb2 | sp\|Q8CC2 | 9 | 7 | 18,9 |
| Q8BJ56 | Q8BJ56 | Patatin-like | Pnpla2 | sp\|Q8BJ56 | 7 | 7 | 21,2 |
| Q9Z0H8 | Q9Z0H8 | CAP-Gly do | Clip2 | sp\|Q9Z0H | 31 | 27 | 37,8 |
| Q69Z38 | Q69Z38 | Pseudopoc | Peak1 | sp\|Q69Z38 | 47 | 47 | 41,3 |
| Q9DBE9 | Q9DBE9 | pre-rRNA p | Ftsj3 | sp\|Q9DBE | 27 | 27 | 39,6 |
| CON__Q9D | CON__Q9DCV | Keratin, ty | Krt7 | ;sp\|Q9DCV | 36 | 27 | 74,8 |
| O89032 | O89032 | SH3 and P) | Sh3pxd2a | sp\|O89032 | 25 | 23 | 30,8 |
| Q8C570 | Q8C570 | mRNA exp | Rae1 | sp\|Q8C57( | 15 | 15 | 69,3 |
| Q9CRC8 | Q9CRC8 | Leucine-ric | Lrrc40 | sp\|Q9CRC | 24 | 24 | 51,2 |
| Q8CC35 | Q8CC35 | Synaptopo | Synpo | sp\|Q8CC3 | 13 | 13 | 22 |
| Q6PR54 | Q6PR54 | Telomere-a | Rif1 | sp\|Q6PR54 | 56 | 56 | 31,9 |
| Q6P9J9 | Q6P9J9 | Anoctamin | Ano6 | sp\|Q6P9J9 | 19 | 19 | 25,9 |
| Q80XL6 | Q80XL6 | Acyl-CoA d | Acad11 | sp\|Q80XL6 | 12 | 12 | 24,5 |
| CON__ENS | CON__ENSEMBL:ENSBTAP00000032840 | | | | 23 | 22 | 34,8 |
| Q6P549;Q | Q6P549 | Phosphatic | Inppl1 | sp\|Q6P54 | 32 | 31 | 35,5 |
| Q61559 | Q61559 | IgG recept | Fcgrt | sp\|Q6155 | 4 | 4 | 6,6 |

| | | | | | | | |
|---|---|---|---|---|---|---|---|
| Q8C6B2 | Q8C6B2 | Rhotekin | Rtkn | sp\|Q8C6B2 | 12 | 12 | 34,4 |
| P04925 | P04925 | Major prio | Prnp | sp\|P04925 | 4 | 4 | 16,5 |
| Q9JJT0 | Q9JJT0 | RNA 3-term | Rcl1 | sp\|Q9JJT0 | 16 | 16 | 53,6 |
| Q8BGF3 | Q8BGF3 | WD repeat | Wdr92 | sp\|Q8BGF3 | 8 | 8 | 32,2 |
| P12265 | P12265 | Beta-glucu | Gusb | sp\|P12265 | 13 | 13 | 25,9 |
| Q8VCH5 | Q8VCH5 | Rab9 effec | Rabepk | sp\|Q8VCH | 8 | 8 | 32,9 |
| P39689 | P39689 | Cyclin-dep | Cdkn1a | sp\|P39689 | 6 | 6 | 59,1 |
| Q8BQZ5 | Q8BQZ5 | Cleavage a | Cpsf4 | sp\|Q8BQZ | 11 | 11 | 59,7 |
| Q9QYR9;O | Q9QYR9;O551 | Acyl-coenz | Acot2;Acot | sp\|Q9QYR | 9 | 8 | 26,7 |
| Q91WQ3 | Q91WQ3 | Tyrosine--t | Yars | sp\|Q91WQ | 34 | 34 | 66,1 |
| Q8C050 | Q8C050 | Ribosomal | Rps6ka5 | sp\|Q8C050 | 8 | 8 | 13,1 |
| Q8BUR3 | Q8BUR3 | Forkhead b | Foxj3 | sp\|Q8BUR | 3 | 3 | 7,9 |
| Q9QXY9 | Q9QXY9 | Peroxisom | Pex3 | sp\|Q9QXY9 | 9 | 9 | 34,9 |
| Q9QYC0 | Q9QYC0 | Alpha-addu | Add1 | sp\|Q9QYC | 11 | 11 | 25,3 |
| Q68FL6 | Q68FL6 | Methionin | Mars | sp\|Q68FL6 | 39 | 39 | 53 |
| Q8K4L0 | Q8K4L0 | ATP-depen | Ddx54 | sp\|Q8K4L0 | 20 | 20 | 31,7 |
| Q9CXJ4 | Q9CXJ4 | ATP-bindin | Abcb8 | sp\|Q9CXJ4 | 11 | 11 | 18,4 |
| Q99KC8 | Q99KC8 | von Willeb | Vwa5a | sp\|Q99KC8 | 30 | 30 | 46,8 |
| P60335 | P60335 | Poly(rC)-bi | Pcbp1 | sp\|P60335 | 18 | 13 | 78,9 |
| P49769 | P49769 | Presenilin- | Psen1 | sp\|P49769 | 5 | 5 | 15,4 |
| Q9JIX8 | Q9JIX8 | Apoptotic | Acin1 | sp\|Q9JIX8 | 29 | 29 | 30,8 |
| Q8BPM0;C | Q8BPM0 | Disheveled | Daam1 | sp\|Q8BPM | 17 | 17 | 19,1 |
| Q8VI36 | Q8VI36 | Paxillin | Pxn | sp\|Q8VI36 | 17 | 17 | 39,9 |
| Q6WKZ8 | Q6WKZ8 | E3 ubiquiti | Ubr2 | sp\|Q6WKZ | 41 | 41 | 31,6 |
| Q8K212 | Q8K212 | Phosphofu | Pacs1 | sp\|Q8K212 | 13 | 11 | 20,6 |
| P47809 | P47809 | Dual specif | Map2k4 | sp\|P47809 | 12 | 12 | 44,3 |
| Q8VDF2 | Q8VDF2 | E3 ubiquiti | Uhrf1 | sp\|Q8VDF2 | 35 | 34 | 52,2 |
| Q8CJG0 | Q8CJG0 | Protein arg | Ago2 | sp\|Q8CJG0 | 29 | 22 | 44,4 |
| Q80V86 | Q80V86 | Integrator | Ints8 | sp\|Q80V86 | 13 | 13 | 19,3 |
| Q9Z0P5 | Q9Z0P5 | Twinfilin-2 | Twf2 | sp\|Q9Z0P5 | 18 | 16 | 67,6 |
| P70335 | P70335 | Rho-associ | Rock1 | sp\|P70335 | 24 | 19 | 22,7 |
| Q8CD19 | Q8CD19 | LanC-like p | Lancl3 | sp\|Q8CD19 | 5 | 5 | 21 |
| Q8BU30 | Q8BU30 | Isoleucine- | Iars | sp\|Q8BU30 | 48 | 48 | 48,4 |
| P07141 | P07141 | Macrophag | Csf1 | sp\|P07141 | 17 | 17 | 45,7 |
| Q80VA0 | Q80VA0 | N-acetylga | Galnt7 | sp\|Q80VA0 | 21 | 21 | 39,6 |
| Q8R5M8 | Q8R5M8 | Cell adhesi | Cadm1 | sp\|Q8R5M | 12 | 12 | 32,5 |
| O89001 | O89001 | Carboxype | Cpd | sp\|O89001 | 30 | 30 | 29,3 |
| Q99MD6 | Q99MD6 | Thioredoxi | Txnrd3 | sp\|Q99MD | 11 | 11 | 26,4 |
| Q9QZW0 | Q9QZW0 | Phospholip | Atp11c | sp\|Q9QZW | 19 | 16 | 21,3 |
| O70422 | O70422 | General tra | Gtf2h4 | sp\|O70422 | 8 | 8 | 24,8 |
| Q8BTY2 | Q8BTY2 | Sodium bic | Slc4a7 | sp\|Q8BTY2 | 17 | 16 | 17,9 |
| Q9ER72 | Q9ER72 | Cysteine--t | Cars | sp\|Q9ER72 | 44 | 44 | 57,9 |
| Q8BTY8 | Q8BTY8 | Sec1 family | Scfd2 | sp\|Q8BTY8 | 13 | 13 | 25 |
| Q91W92 | Q91W92 | Cdc42 effe | Cdc42ep1 | sp\|Q91W9 | 12 | 12 | 54,3 |
| Q99LD8 | Q99LD8 | N(G),N(G)- | Ddah2 | sp\|Q99LD8 | 7 | 6 | 37,9 |
| Q62417 | Q62417 | Sorbin and | Sorbs1 | sp\|Q62417 | 19 | 18 | 20,2 |
| Q9CQ33 | Q9CQ33 | Leucine-zip | Lztr1 | sp\|Q9CQ3 | 7 | 7 | 11,5 |
| Q99PU8 | Q99PU8 | Putative A | Dhx30 | sp\|Q99PU | 49 | 49 | 50,9 |
| Q9Z0N1;Q | Q9Z0N1;Q9Z0 | Eukaryotic | Eif2s3x;Eif | sp\|Q9Z0N | 23 | 23 | 57,4 |
| Q7TND5 | Q7TND5 | Ribosome | Rpf1 | sp\|Q7TND | 14 | 14 | 49,6 |

| | | | | | | | |
|---|---|---|---|---|---|---|---|
| Q9CR89 | Q9CR89 | Endoplasm | Ergic2 | sp\|Q9CR8 | 5 | 5 | 18 |
| Q921N6 | Q921N6 | Probable A | Ddx27 | sp\|Q921N | 35 | 35 | 45 |
| Q8C147 | Q8C147 | Dedicator | Dock8 | sp\|Q8C147 | 33 | 29 | 21,2 |
| Q8BHY3 | Q8BHY3 | Anoctamin | Ano1 | sp\|Q8BHY | 8 | 8 | 11,6 |
| P48758 | P48758 | Carbonyl r | Cbr1 | sp\|P48758 | 15 | 14 | 64,3 |
| Q6P1F6;Q | Q6P1F6 | Serine/thre | Ppp2r2a | sp\|Q6P1F6 | 19 | 10 | 51 |
| Q8C436 | Q8C436 | Protein-lys | Vcpkmt | sp\|Q8C436 | 4 | 4 | 26,3 |
| P09021;P0 | P09021 | Homeobox | Hoxa5 | sp\|P09021 | 8 | 8 | 44,4 |
| Q61398 | Q61398 | Procollage | Pcolce | sp\|Q61398 | 14 | 14 | 38,5 |
| Q9CR00 | Q9CR00 | 26S protea | Psmd9 | sp\|Q9CR00 | 9 | 9 | 44,1 |
| Q8R4G6 | Q8R4G6 | Alpha-1,6- | Mgat5 | sp\|Q8R4G | 10 | 9 | 18,6 |
| Q8VCM3 | Q8VCM3 | Zinc finger | Zfyve21 | sp\|Q8VCM | 4 | 4 | 27,8 |
| Q7TNB8 | Q7TNB8 | Protein str | Sbno2 | sp\|Q7TNB | 9 | 6 | 8,2 |
| Q9Z247 | Q9Z247 | Peptidyl-pr | Fkbp9 | sp\|Q9Z247 | 10 | 10 | 26 |
| P37040 | P37040 | NADPH--cy | Por | sp\|P37040 | 33 | 33 | 55,3 |
| Q9ERB0 | Q9ERB0 | Synaptoso | Snap29 | sp\|Q9ERB0 | 11 | 11 | 48,1 |
| G5E829;Q | G5E829 | Plasma me | Atp2b1 | sp\|G5E829 | 39 | 29 | 38,4 |
| Q922H1 | Q922H1 | Protein arg | Prmt3 | sp\|Q922H1 | 17 | 17 | 36,7 |
| Q69ZL1 | Q69ZL1 | FYVE, Rho | Fgd6 | sp\|Q69ZL1 | 8 | 8 | 5,9 |
| Q80WE4 | Q80WE4 | Kinesin-like | Kif20b | sp\|Q80WE | 36 | 36 | 25,5 |
| Q9D281 | Q9D281 | Protein No | Fam114a1 | sp\|Q9D28 | 13 | 13 | 35,9 |
| O35857 | O35857 | Mitochond | Timm44 | sp\|O35857 | 18 | 18 | 42,9 |
| Q8C6L5 | Q8C6L5 | Cyclic GMP | Mb21d1 | sp\|Q8C6L5 | 18 | 18 | 42,8 |
| Q9ERI5 | Q9ERI5 | Bifunction | Jmjd6 | sp\|Q9ERI5 | 12 | 12 | 42,2 |
| Q9BCZ4 | Q9BCZ4 | Selenoprot | Vimp | sp\|Q9BCZ4 | 6 | 6 | 46,8 |
| Q8K4B2 | Q8K4B2 | Interleukin | Irak3 | sp\|Q8K4B2 | 4 | 4 | 9,5 |
| Q9EPS3 | Q9EPS3 | D-glucuron | Glce | sp\|Q9EPS3 | 9 | 9 | 21,4 |
| P70290 | P70290 | 55 kDa ery | Mpp1 | sp\|P70290 | 21 | 21 | 65 |
| Q8BIP0 | Q8BIP0 | Aspartate- | Dars2 | sp\|Q8BIP0 | 15 | 15 | 29,6 |
| Q8C0C0 | Q8C0C0 | Zinc finger | Zhx2 | sp\|Q8C0C0 | 17 | 17 | 31,1 |
| Q8VDP4 | Q8VDP4 | Cell cycle a | Ccar2 | sp\|Q8VDP | 28 | 28 | 51,8 |
| Q9DBL2 | Q9DBL2 | Ganglioside | Gdap2 | sp\|Q9DBL | 7 | 7 | 18,9 |
| Q9DCS2 | Q9DCS2 | UPF0585 protein C16 | | sp\|Q9DCS | 5 | 5 | 39,7 |
| B2RUR8;Q | B2RUR8 | OTU doma | Otud7b | sp\|B2RUR | 17 | 17 | 30,5 |
| Q920A5 | Q920A5 | Retinoid-in | Scpep1 | sp\|Q920A5 | 7 | 7 | 17,9 |
| Q9CQ62 | Q9CQ62 | 2,4-dienoy | Decr1 | sp\|Q9CQ6 | 6 | 6 | 27,5 |
| Q6VN19 | Q6VN19 | Ran-bindin | Ranbp10 | sp\|Q6VN1 | 14 | 12 | 36,6 |
| O88643 | O88643 | Serine/thre | Pak1 | sp\|O88643 | 19 | 11 | 46,4 |
| Q9Z1X4 | Q9Z1X4 | Interleukin | Ilf3 | sp\|Q9Z1X4 | 31 | 27 | 39,6 |
| Q61140 | Q61140 | Breast can | Bcar1 | sp\|Q61140 | 19 | 19 | 36,5 |
| Q05920 | Q05920 | Pyruvate c | Pc | sp\|Q05920 | 24 | 24 | 27,5 |
| Q9JJK2 | Q9JJK2 | LanC-like p | Lancl2 | sp\|Q9JJK2 | 15 | 15 | 40,9 |
| P62331 | P62331 | ADP-ribosy | Arf6 | sp\|P62331 | 8 | 8 | 61,7 |
| P62627 | P62627 | Dynein ligh | Dynlrb1 | sp\|P62627 | 5 | 5 | 60,4 |
| Q8VC88 | Q8VC88 | Grancalcin | Gca | sp\|Q8VC8 | 3 | 3 | 13,6 |
| Q9CWR7 | Q9CWR7 | Metallored | Steap1 | sp\|Q9CWR | 6 | 6 | 17,7 |
| Q5M8N4 | Q5M8N4 | Epimerase | Sdr39u1 | sp\|Q5M8N | 5 | 5 | 19,5 |
| P34056;Q6 | P34056;Q6VU | Transcripti | Tfap2a;Tfa | sp\|P34056 | 6 | 5 | 19,5 |
| Q9CYH6 | Q9CYH6 | Ribosome | Rrs1 | sp\|Q9CYH | 15 | 15 | 39,7 |
| Q99JI1 | Q99JI1 | Musculosk | Mustn1 | sp\|Q99JI1 | 3 | 3 | 46,3 |

| | | | | | | | |
|---|---|---|---|---|---|---|---|
| P23591 | P23591 | GDP-L-fucc | Tsta3 | sp\|P23591 | 12 | 12 | 53,6 |
| Q91ZS8 | Q91ZS8 | Double-str | Adarb1 | sp\|Q91ZS8 | 12 | 12 | 26,6 |
| Q9JJ80 | Q9JJ80 | Ribosome | Rpf2 | sp\|Q9JJ80 | 17 | 17 | 51,3 |
| P59808;P5 | P59808 | SAM and S | Sash1 | sp\|P59808 | 9 | 9 | 10,7 |
| Q61102 | Q61102 | ATP-bindin | Abcb7 | sp\|Q61102 | 13 | 13 | 25,9 |
| Q9EQN3 | Q9EQN3 | TSC22 dom | Tsc22d4 | sp\|Q9EQN | 8 | 7 | 26,6 |
| P30285 | P30285 | Cyclin-dep | Cdk4 | sp\|P30285 | 19 | 18 | 70,3 |
| Q91YD3 | Q91YD3 | mRNA-dec | Dcp1a | sp\|Q91YD3 | 15 | 15 | 37,2 |
| Q99MY8 | Q99MY8 | Histone-lys | Ash1l | sp\|Q99MY | 14 | 14 | 6,9 |
| Q8R550 | Q8R550 | SH3 domai | Sh3kbp1 | sp\|Q8R550 | 20 | 20 | 32,7 |
| Q8K4F5 | Q8K4F5 | Alpha/beta | Abhd11 | sp\|Q8K4F5 | 5 | 5 | 24,8 |
| Q64437 | Q64437 | Alcohol de | Adh7 | sp\|Q64437 | 16 | 14 | 65,8 |
| Q8C129 | Q8C129 | Leucyl-cyst | Lnpep | sp\|Q8C129 | 31 | 31 | 39,4 |
| P35822 | P35822 | Receptor-t | Ptprk | sp\|P35822 | 28 | 27 | 27,6 |
| Q9WVQ5 | Q9WVQ5 | Methylthic | Apip | sp\|Q9WVC | 7 | 7 | 45,6 |
| Q64701 | Q64701 | Retinoblas | Rbl1 | sp\|Q64701 | 28 | 28 | 30,9 |
| Q14B71 | Q14B71 | Cell divisio | Cdca2 | sp\|Q14B71 | 18 | 18 | 32,8 |
| O55242 | O55242 | Sigma non- | Sigmar1 | sp\|O55242 | 4 | 4 | 32,3 |
| Q80UM7 | Q80UM7 | Mannosyl- | Mogs | sp\|Q80UM | 20 | 20 | 36,5 |
| Q99N93 | Q99N93 | 39S riboso | Mrpl16 | sp\|Q99N93 | 6 | 6 | 30,7 |
| Q8C052 | Q8C052 | Microtubu | Map1s | sp\|Q8C052 | 18 | 18 | 27,3 |
| Q9WV80 | Q9WV80 | Sorting ne | Snx1 | sp\|Q9WV8 | 22 | 20 | 46,2 |
| Q91XA2 | Q91XA2 | Golgi mem | Golm1 | sp\|Q91XA2 | 6 | 6 | 19,6 |
| Q8BYA0 | Q8BYA0 | Tubulin-sp | Tbcd | sp\|Q8BYA0 | 29 | 29 | 30,4 |
| Q9DCD5 | Q9DCD5 | Tight junct | Tjap1 | sp\|Q9DCD | 17 | 17 | 47,9 |
| Q6AXC6 | Q6AXC6 | Probable A | Ddx11 | sp\|Q6AXC6 | 14 | 14 | 22,8 |
| P04627 | P04627 | Serine/thre | Araf | sp\|P04627 | 12 | 9 | 26,2 |
| Q9JKK8 | Q9JKK8 | Serine/thre | Atr | sp\|Q9JKK8 | 41 | 41 | 21,3 |
| Q6PAV2 | Q6PAV2 | Probable E | Herc4 | sp\|Q6PAV2 | 32 | 32 | 40,5 |
| Q8C3S2 | Q8C3S2 | Transport a | Tango6 | sp\|Q8C3S2 | 14 | 14 | 17,4 |
| Q9Z0H1 | Q9Z0H1 | WD repeat | Wdr46 | sp\|Q9Z0H1 | 14 | 14 | 30,9 |
| Q9JKX6 | Q9JKX6 | ADP-sugar | Nudt5 | sp\|Q9JKX6 | 10 | 10 | 55 |
| Q91VS7 | Q91VS7 | Microsoma | Mgst1 | sp\|Q91VS7 | 2 | 2 | 18,7 |
| Q91ZU6 | Q91ZU6 | Dystonin | Dst | sp\|Q91ZU6 | 156 | 151 | 28,1 |
| Q69ZA1 | Q69ZA1 | Cyclin-dep | Cdk13 | sp\|Q69ZA1 | 21 | 17 | 22,5 |
| Q6P1G2 | Q6P1G2 | Lysine-spe | Kdm2b | sp\|Q6P1G2 | 7 | 7 | 6,4 |
| Q9CWG8 | Q9CWG8 | NADH deh | Ndufaf7 | sp\|Q9CWG | 17 | 17 | 48,4 |
| Q3TYD4 | Q3TYD4 | Arylsulfata | Arsg | sp\|Q3TYD4 | 5 | 5 | 18,1 |
| Q5SRX1 | Q5SRX1 | TOM1-like | Tom1l2 | sp\|Q5SRX1 | 14 | 14 | 40,6 |
| Q60738 | Q60738 | Zinc transp | Slc30a1 | sp\|Q60738 | 7 | 7 | 26 |
| Q91Z58 | Q91Z58 | Uncharacterized prote | | sp\|Q91Z58 | 13 | 13 | 19,2 |
| Q91YR9 | Q91YR9 | Prostaglan | Ptgr1 | sp\|Q91YR9 | 7 | 7 | 29,5 |
| Q91VE0 | Q91VE0 | Long-chain | Slc27a4 | sp\|Q91VE0 | 16 | 15 | 31,1 |
| Q9WUU8 | Q9WUU8 | TNFAIP3-ir | Tnip1 | sp\|Q9WUU | 7 | 7 | 21 |
| A2A9C3 | A2A9C3 | Protein SZ | Szt2 | sp\|A2A9C3 | 6 | 6 | 2,7 |
| Q8CGC6 | Q8CGC6 | RNA-bindir | Rbm28 | sp\|Q8CGC | 20 | 19 | 29,6 |
| Q0GNC1 | Q0GNC1 | Inverted fo | Inf2 | sp\|Q0GNC | 32 | 32 | 37,2 |
| O55125 | O55125 | Protein Nip | Nipsnap1 | sp\|O55125 | 8 | 8 | 33,1 |
| Q07797 | Q07797 | Galectin-3- | Lgals3bp | sp\|Q07797 | 5 | 5 | 15,3 |
| P19783 | P19783 | Cytochrom | Cox4i1 | sp\|P19783 | 11 | 11 | 53,8 |

| | | | | | | | |
|---|---|---|---|---|---|---|---|
| Q5S006 | Q5S006 | Leucine-ric | Lrrk2 | sp\|Q5S006 | 14 | 14 | 8 |
| P70365 | P70365 | Nuclear re | Ncoa1 | sp\|P70365 | 7 | 7 | 7,4 |
| Q8BZX4 | Q8BZX4 | Splicing reg | Srek1 | sp\|Q8BZX4 | 6 | 6 | 17,8 |
| Q8BVE8 | Q8BVE8 | Histone-lys | Whsc1 | sp\|Q8BVE8 | 38 | 38 | 37,4 |
| CON__Q9T | CON__Q9TRI1 | | | | 14 | 10 | 22,1 |
| Q61029 | Q61029 | Lamina-ass | Tmpo | sp\|Q61029 | 21 | 11 | 57,3 |
| Q8K2T1 | Q8K2T1 | NmrA-like | Nmral1 | sp\|Q8K2T1 | 10 | 10 | 39,5 |
| P45481 | P45481 | CREB-bindi | Crebbp | sp\|P45481 | 23 | 14 | 13,1 |
| P20152;P1 | P20152 | Vimentin | Vim | sp\|P20152 | 62 | 55 | 89,9 |
| Q7TPV4 | Q7TPV4 | Myb-bindir | Mybbp1a | sp\|Q7TPV4 | 69 | 69 | 57,5 |
| Q8BGT9 | Q8BGT9 | Polypeptid | Galnt12 | sp\|Q8BGT9 | 4 | 3 | 9,2 |
| P46664 | P46664 | Adenylosu | Adss | sp\|P46664 | 22 | 20 | 59,2 |
| Q9Z2E2 | Q9Z2E2 | Methyl-Cp | Mbd1 | sp\|Q9Z2E2 | 7 | 7 | 19,2 |
| Q9QZM0 | Q9QZM0 | Ubiquilin-2 | Ubqln2 | sp\|Q9QZM | 10 | 7 | 28,4 |
| Q921J2 | Q921J2 | GTP-bindir | Rheb | sp\|Q921J2 | 9 | 9 | 38 |
| Q9JIW9 | Q9JIW9 | Ras-related | Ralb | sp\|Q9JIW9 | 9 | 5 | 50 |
| Q7TMM9 | Q7TMM9 | Tubulin be | Tubb2a | sp\|Q7TMN | 26 | 1 | 69 |
| P10630 | P10630 | Eukaryotic | Eif4a2 | sp\|P10630 | 24 | 9 | 51,1 |
| Q8BYB9 | Q8BYB9 | Protein O-g | Poglut1 | sp\|Q8BYB9 | 11 | 11 | 29,8 |
| P17047 | P17047 | Lysosome- | Lamp2 | sp\|P17047 | 7 | 7 | 15,7 |
| P16125 | P16125 | L-lactate d | Ldhb | sp\|P16125 | 9 | 8 | 30,5 |
| Q8K0C4 | Q8K0C4 | Lanosterol | Cyp51a1 | sp\|Q8K0C4 | 16 | 16 | 39,2 |
| P26638 | P26638 | Serine--tRN | Sars | sp\|P26638 | 30 | 30 | 58,8 |
| Q8VCW8 | Q8VCW8 | Acyl-CoA s | Acsf2 | sp\|Q8VCW | 16 | 16 | 34,1 |
| Q9D4B1 | Q9D4B1 | Phosphatid | Sgms2 | sp\|Q9D4B | 7 | 7 | 24,1 |
| Q8VCN9 | Q8VCN9 | Tubulin-sp | Tbcc | sp\|Q8VCN | 16 | 16 | 64,5 |
| Q9D279 | Q9D279 | Mitotic int | Misp | sp\|Q9D279 | 10 | 10 | 23,5 |
| P03921 | P03921 | NADH-ubic | Mtnd5 | sp\|P03921 | 5 | 5 | 11,9 |
| Q8BYW9 | Q8BYW9 | EGF domai | Eogt | sp\|Q8BYW | 8 | 8 | 17,5 |
| Q3UHH8 | Q3UHH8 | Glucoside x | Gxylt1 | sp\|Q3UHH | 7 | 7 | 19,1 |
| Q8CCH7 | Q8CCH7 | Zinc finger | Zfpm2 | sp\|Q8CCH | 7 | 7 | 12,3 |
| P48036 | P48036 | Annexin A5 | Anxa5 | sp\|P48036 | 33 | 33 | 84,3 |
| Q91VB4 | Q91VB4 | Hermansky | Hps3 | sp\|Q91VB4 | 12 | 12 | 14,5 |
| Q8VE98 | Q8VE98 | CD276 ant | Cd276 | sp\|Q8VE98 | 2 | 2 | 6,3 |
| Q8BKG3 | Q8BKG3 | Inactive ty | Ptk7 | sp\|Q8BKG | 28 | 28 | 39,7 |
| Q8BY71 | Q8BY71 | Histone ac | Hat1 | sp\|Q8BY71 | 14 | 14 | 49 |
| Q9WU79 | Q9WU79 | Proline deh | Prodh | sp\|Q9WU7 | 11 | 11 | 29,9 |
| Q99J93 | Q99J93 | Interferon- | Ifitm2 | sp\|Q99J93 | 3 | 2 | 29,2 |
| Q9EQ06 | Q9EQ06 | Estradiol 1 | Hsd17b11 | sp\|Q9EQ0 | 3 | 3 | 14,8 |
| O89112 | O89112 | LanC-like p | Lancl1 | sp\|O89112 | 7 | 7 | 24,8 |
| Q99ME2 | Q99ME2 | WD repeat | Wdr6 | sp\|Q99ME | 21 | 21 | 29,8 |
| Q8BQM4 | Q8BQM4 | HEAT repe | Heatr3 | sp\|Q8BQM | 19 | 19 | 37,6 |
| B1AWL2 | B1AWL2 | | Zfp462 | sp\|B1AWL | 23 | 23 | 14,1 |
| P53995 | P53995 | Anaphase- | Anapc1 | sp\|P53995 | 43 | 43 | 29,9 |
| Q8BZH4 | Q8BZH4 | Pogo trans | Pogz | sp\|Q8BZH4 | 33 | 33 | 37,7 |
| Q8BX10 | Q8BX10 | Serine/thre | Pgam5 | sp\|Q8BX10 | 13 | 13 | 49,3 |
| P61957;Q9 | P61957;Q9Z17 | Small ubiq | Sumo2;Sur | sp\|P61957 | 2 | 2 | 23,2 |
| O88842 | O88842 | FYVE, RhoG | Fgd3 | sp\|O88842 | 7 | 7 | 17,3 |
| P59108 | P59108 | Copine-2 | Cpne2 | sp\|P59108 | 17 | 16 | 37,4 |
| Q8BIL5 | Q8BIL5 | Protein Ho | Hook1 | sp\|Q8BIL5 | 9 | 9 | 18,1 |

| | | | | | | | |
|---|---|---|---|---|---|---|---|
| P49117 | P49117 | Nuclear re | Nr2c2 | sp\|P49117 | 13 | 13 | 31,2 |
| Q9CXA2 | Q9CXA2 | Trans-L-3-h | L3hypdh | sp\|Q9CXA2 | 7 | 7 | 29,7 |
| Q9D7J9 | Q9D7J9 | Enoyl-CoA | Echdc3 | sp\|Q9D7J9 | 10 | 10 | 51,7 |
| Q64521 | Q64521 | Glycerol-3- | Gpd2 | sp\|Q64521 | 46 | 46 | 68,6 |
| Q91VH6 | Q91VH6 | Protein ME | Memo1 | sp\|Q91VH | 14 | 14 | 70,7 |
| Q9D1A2 | Q9D1A2 | Cytosolic n | Cndp2 | sp\|Q9D1A | 20 | 20 | 50,1 |
| Q9D071 | Q9D071 | MMS19 nu | Mms19 | sp\|Q9D07 | 27 | 27 | 40,5 |
| O35400 | O35400 | Sulfotransf | Sult2b1 | sp\|O35400 | 5 | 5 | 22,5 |
| Q60769 | Q60769 | Tumor nec | Tnfaip3 | sp\|Q60769 | 20 | 20 | 35,2 |
| Q8C561 | Q8C561 | LMBR1 dor | Lmbrd2 | sp\|Q8C561 | 6 | 6 | 13,4 |
| Q6PB93 | Q6PB93 | Polypeptid | Galnt2 | sp\|Q6PB93 | 19 | 18 | 41,9 |
| P18155 | P18155 | Bifunctiona | Mthfd2 | sp\|P18155 | 12 | 12 | 42 |
| CON__Q3S | CON__Q3SZR3 | | | | 7 | 7 | 37,6 |
| O09111 | O09111 | NADH deh | Ndufb11 | sp\|O09111 | 5 | 5 | 46,4 |
| Q80Y44 | Q80Y44 | Probable A | Ddx10 | sp\|Q80Y44 | 24 | 24 | 34,1 |
| Q9Z1Z2 | Q9Z1Z2 | Serine-thre | Strap | sp\|Q9Z1Z2 | 19 | 19 | 70 |
| Q8K2J0 | Q8K2J0 | 1-phospha | Plcd3 | sp\|Q8K2J0 | 22 | 22 | 41,4 |
| Q8BTI7 | Q8BTI7 | Serine/thre | Ankrd52 | sp\|Q8BTI7 | 15 | 15 | 20 |
| P52431 | P52431 | DNA polym | Pold1 | sp\|P52431 | 45 | 45 | 51,3 |
| Q8R059 | Q8R059 | UDP-gluco | Gale | sp\|Q8R059 | 10 | 10 | 42,4 |
| Q9Z0Z7 | Q9Z0Z7 | Krueppel-li | Klf5 | sp\|Q9Z0Z7 | 6 | 6 | 20,4 |
| Q91VK4 | Q91VK4 | Integral me | Itm2c | sp\|Q91VK4 | 7 | 7 | 43,1 |
| Q9R087 | Q9R087 | Glypican-6 | Gpc6 | sp\|Q9R087 | 7 | 5 | 18,7 |
| Q8BG17 | Q8BG17 | Nucleolar p | Nol12 | sp\|Q8BG17 | 5 | 5 | 37,3 |
| Q9CWT6 | Q9CWT6 | Probable A | Ddx28 | sp\|Q9CWT | 12 | 12 | 29,6 |
| Q60775 | Q60775 | ETS-related | Elf1 | sp\|Q60775 | 6 | 6 | 20,4 |
| Q61595 | Q61595 | Kinectin | Ktn1 | sp\|Q61595 | 71 | 71 | 52,1 |
| Q8BRH0 | Q8BRH0 | Transmem | Tmtc3 | sp\|Q8BRH | 27 | 25 | 33,7 |
| P29352 | P29352 | Tyrosine-p | Ptpn22 | sp\|P29352 | 8 | 8 | 16,2 |
| Q9WTL7 | Q9WTL7 | Acyl-protei | Lypla2 | sp\|Q9WTL | 6 | 6 | 44,6 |
| P35564 | P35564 | Calnexin | Canx | sp\|P35564 | 41 | 41 | 49,2 |
| Q9R078 | Q9R078 | 5-AMP-act | Prkab1 | sp\|Q9R078 | 6 | 5 | 35,2 |
| Q8C7Q4 | Q8C7Q4 | RNA-bindir | Rbm4 | sp\|Q8C7Q | 12 | 2 | 38,8 |
| P56183 | P56183 | Ribosomal | Rrp1 | sp\|P56183 | 16 | 16 | 47,8 |
| Q8BTS4 | Q8BTS4 | Nuclear po | Nup54 | sp\|Q8BTS4 | 21 | 21 | 50,4 |
| Q8BKZ9 | Q8BKZ9 | Pyruvate d | Pdhx | sp\|Q8BKZ9 | 9 | 9 | 28,7 |
| Q3UFY8 | Q3UFY8 | Mitochond | Trmt10c | sp\|Q3UFY8 | 14 | 14 | 43,2 |
| Q9EPK8 | Q9EPK8 | Transient r | Trpv4 | sp\|Q9EPK8 | 17 | 17 | 24,9 |
| P70419;Q8 | P70419 | Polypeptid | Galnt3 | sp\|P70419 | 17 | 17 | 34,9 |
| Q3V3R1 | Q3V3R1 | Monofunc | Mthfd1l | sp\|Q3V3R1 | 43 | 43 | 56 |
| Q810Q5 | Q810Q5 | Normal mu | Nmes1 | sp\|Q810Q | 3 | 3 | 47 |
| Q80XC2 | Q80XC2 | tRNA (aden | Trmt61a | sp\|Q80XC2 | 8 | 8 | 47,2 |
| Q9QUR6 | Q9QUR6 | Prolyl endo | Prep | sp\|Q9QUR | 38 | 38 | 67,5 |
| Q3ULF4 | Q3ULF4 | Paraplegin | Spg7 | sp\|Q3ULF4 | 12 | 12 | 21,8 |
| P57716 | P57716 | Nicastrin | Ncstn | sp\|P57716 | 12 | 12 | 21,6 |
| Q9DCZ4 | Q9DCZ4 | Apolipopro | Apoo | sp\|Q9DCZ4 | 5 | 5 | 34,3 |
| CON__P02 | CON__P02777 | | | | 3 | 3 | 5,7 |
| Q8BIQ5 | Q8BIQ5 | Cleavage s | Cstf2 | sp\|Q8BIQ5 | 13 | 9 | 38,1 |
| Q80X85 | Q80X85 | 28S riboso | Mrps7 | sp\|Q80X85 | 8 | 8 | 37,6 |
| Q80Y19 | Q80Y19 | Rho GTPas | Arhgap11a | sp\|Q80Y19 | 14 | 14 | 21 |

| | | | | | | | |
|---|---|---|---|---|---|---|---|
| Q9D0N7 | Q9D0N7 | Chromatin | Chaf1b | sp\|Q9D0N | 19 | 19 | 54,4 |
| Q64314 | Q64314 | Hematopo | Cd34 | sp\|Q64314 | 7 | 7 | 23,6 |
| Q8BJY1 | Q8BJY1 | 26S protea | Psmd5 | sp\|Q8BJY1 | 20 | 20 | 56,3 |
| Q8BFZ9 | Q8BFZ9 | Erlin-2 | Erlin2 | sp\|Q8BFZ9 | 17 | 15 | 53,5 |
| Q7TN12 | Q7TN12 | Uncharacterized prote | | sp\|Q7TN12 | 3 | 3 | 5,3 |
| Q9CQ26 | Q9CQ26 | STAM-bind | Stambp | sp\|Q9CQ2 | 13 | 13 | 37 |
| Q8BH59 | Q8BH59 | Calcium-bi | Slc25a12 | sp\|Q8BH5 | 33 | 30 | 65,6 |
| O70456 | O70456 | 14-3-3 pro | Sfn | sp\|O70456 | 12 | 9 | 55,6 |
| Q3TX08 | Q3TX08 | tRNA (guar | Trmt1 | sp\|Q3TX08 | 19 | 19 | 38,6 |
| Q9DBB1 | Q9DBB1 | Dual specif | Dusp6 | sp\|Q9DBB | 2 | 2 | 6,3 |
| Q571E4 | Q571E4 | N-acetylga | Galns | sp\|Q571E4 | 6 | 6 | 16,3 |
| Q0VEE6 | Q0VEE6 | Zinc finger | Znf800 | sp\|Q0VEE6 | 8 | 8 | 19,6 |
| Q8BG73 | Q8BG73 | SH3 domai | Sh3bgrl2 | sp\|Q8BG7 | 5 | 5 | 72 |
| Q5SW19 | Q5SW19 | Clustered r | Cluh | sp\|Q5SW1 | 34 | 34 | 35,1 |
| Q8BVG4 | Q8BVG4 | Dipeptidyl | Dpp9 | sp\|Q8BVG | 26 | 26 | 42,1 |
| Q6AW69 | Q6AW69 | Cingulin-lik | Cgnl1 | sp\|Q6AW6 | 32 | 31 | 30,9 |
| Q9D7X8 | Q9D7X8 | Gamma-gl | Ggct | sp\|Q9D7X8 | 6 | 6 | 47,3 |
| Q8R3Y8 | Q8R3Y8 | Interferon | Irf2bp1 | sp\|Q8R3Y8 | 19 | 18 | 44,9 |
| Q8R3C6 | Q8R3C6 | Probable R | Rbm19 | sp\|Q8R3C6 | 29 | 29 | 43,4 |
| Q811I0 | Q811I0 | ATP syntha | Atpaf1 | sp\|Q811I0 | 9 | 9 | 36,7 |
| P00405 | P00405 | Cytochrom | Mtco2 | sp\|P00405 | 5 | 5 | 27,8 |
| Q8R2M2 | Q8R2M2 | Deoxynucl | Dnttip2 | sp\|Q8R2M | 28 | 28 | 44,3 |
| Q9Z179 | Q9Z179 | SHC SH2 d | Shcbp1 | sp\|Q9Z179 | 18 | 18 | 36,8 |
| Q9JIF7 | Q9JIF7 | Coatomer | Copb1 | sp\|Q9JIF7 | 41 | 41 | 54,1 |
| Q5SUR0 | Q5SUR0 | Phosphorib | Pfas | sp\|Q5SUR0 | 40 | 40 | 54,9 |
| Q9DBF1 | Q9DBF1 | Alpha-amir | Aldh7a1 | sp\|Q9DBF1 | 13 | 13 | 36,4 |
| Q9CQE8 | Q9CQE8 | UPF0568 protein C14 | | sp\|Q9CQE | 16 | 16 | 66 |
| Q3U7U3 | Q3U7U3 | F-box only | Fbxo7 | sp\|Q3U7U | 5 | 5 | 14,7 |
| Q62425 | Q62425 | Cytochrom | Ndufa4 | sp\|Q62425 | 5 | 5 | 47,6 |
| Q9CZ42 | Q9CZ42 | ATP-depen | Carkd | sp\|Q9CZ42 | 14 | 14 | 65 |
| Q9Z2E1 | Q9Z2E1 | Methyl-Cp | Mbd2 | sp\|Q9Z2E1 | 11 | 11 | 35,3 |
| O08914 | O08914 | Fatty-acid | Faah | sp\|O08914 | 11 | 11 | 33,3 |
| O88958 | O88958 | Glucosami | Gnpda1 | sp\|O88958 | 10 | 9 | 44,6 |
| Q9JI99 | Q9JI99 | Sphingosin | Sgpp1 | sp\|Q9JI99 | 5 | 5 | 14,7 |
| Q99JW2 | Q99JW2 | Aminoacyl | Acy1 | sp\|Q99JW | 16 | 16 | 50,5 |
| Q8BX02 | Q8BX02 | KN motif a | Kank2 | sp\|Q8BX02 | 23 | 23 | 42,5 |
| P35922 | P35922 | Fragile X m | Fmr1 | sp\|P35922 | 22 | 20 | 38,9 |
| A2AJL3 | A2AJL3 | FGGY carb | Fggy | sp\|A2AJL3 | 12 | 12 | 31,2 |
| Q03160 | Q03160 | Growth fac | Grb7 | sp\|Q03160 | 4 | 4 | 9,7 |
| Q6P5D8 | Q6P5D8 | Structural | Smchd1 | sp\|Q6P5D8 | 84 | 84 | 51,4 |
| Q6IRU2 | Q6IRU2 | Tropomyo | Tpm4 | sp\|Q6IRU2 | 29 | 25 | 77,8 |
| P0DPE0;P0 | P0DPE0;P0DPD9 | | | sp\|P0DPE0 | 3 | 3 | 13,7 |
| P06801 | P06801 | NADP-dep | Me1 | sp\|P06801 | 22 | 22 | 50,2 |
| Q8CGA0 | Q8CGA0 | Protein ph | Ppm1f | sp\|Q8CGA | 8 | 8 | 23,7 |
| Q80UU1 | Q80UU1 | Ankyrin re | Ankzf1 | sp\|Q80UU | 13 | 13 | 23,8 |
| Q923D5 | Q923D5 | WW doma | Wbp11 | sp\|Q923D5 | 17 | 17 | 26,1 |
| P42128 | P42128 | Forkhead b | Foxk1 | sp\|P42128 | 15 | 14 | 28,8 |
| Q5XJE5 | Q5XJE5 | RNA polym | Leo1 | sp\|Q5XJE5 | 10 | 10 | 22,6 |
| Q8R001 | Q8R001 | Microtubu | Mapre2 | sp\|Q8R001 | 9 | 8 | 45,4 |
| Q9CZN8 | Q9CZN8 | Glutamyl-t | Qrsl1 | sp\|Q9CZN8 | 13 | 13 | 35,6 |

| | | | | | | | |
|---|---|---|---|---|---|---|---|
| Q8BG15 | Q8BG15 | CTD small | Ctdspl2 | sp\|Q8BG1 | 15 | 15 | 35,1 |
| Q8VD12 | Q8VD12 | Zinc finger | Znf385a | sp\|Q8VD1 | 5 | 5 | 19,9 |
| O08573 | O08573 | Galectin-9 | Lgals9 | sp\|O0857 | 9 | 9 | 32,9 |
| Q3UFY7 | Q3UFY7 | 7-methylgu | Nt5c3b | sp\|Q3UFY | 12 | 12 | 62,7 |
| Q61468 | Q61468 | Mesotheli | Msln | sp\|Q6146 | 12 | 12 | 24,6 |
| Q8C4X2;Q | Q8C4X2;Q8BV | Casein kina | Csnk1g3;C | sp\|Q8C4X | 12 | 6 | 31,4 |
| Q8VBX6 | Q8VBX6 | Multiple P | Mpdz | sp\|Q8VBX | 26 | 25 | 21,5 |
| Q9DCT1 | Q9DCT1 | 1,5-anhydr | Akr1e2 | sp\|Q9DCT | 7 | 7 | 35,9 |
| P40201 | P40201 | Chromodo | Chd1 | sp\|P40201 | 34 | 28 | 25,5 |
| Q9CQB2 | Q9CQB2 | Protein FA | Fam195a | sp\|Q9CQB | 2 | 2 | 22,5 |
| Q91VT1 | Q91VT1 | E3 SUMO- | Nsmce2 | sp\|Q91VT | 4 | 4 | 18,6 |
| CON__P01 | CON__P01966 | | | | 7 | 5 | 58,5 |
| Q8BW10 | Q8BW10 | RNA-bindi | Nob1 | sp\|Q8BW1 | 14 | 14 | 40,4 |
| Q9JLT4 | Q9JLT4 | Thioredoxi | Txnrd2 | sp\|Q9JLT4 | 8 | 7 | 22,3 |
| O88952;Q | O88952 | Protein lin | Lin7c | sp\|O88952 | 9 | 9 | 53,8 |
| Q8R1A4 | Q8R1A4 | Dedicator | Dock7 | sp\|Q8R1A | 56 | 51 | 33,8 |
| Q9JJR8 | Q9JJR8 | Transmem | Tmem9b | sp\|Q9JJR8 | 4 | 3 | 30,2 |
| P24369 | P24369 | Peptidyl-p | Ppib | sp\|P24369 | 17 | 17 | 60,6 |
| Q9DCS9 | Q9DCS9 | NADH deh | Ndufb10 | sp\|Q9DCS | 9 | 9 | 55,7 |
| Q9JLQ0 | Q9JLQ0 | CD2-associ | Cd2ap | sp\|Q9JLQ | 23 | 23 | 44,7 |
| Q6ZQ58 | Q6ZQ58 | La-related | Larp1 | sp\|Q6ZQ5 | 41 | 41 | 57,3 |
| Q9WTU0 | Q9WTU0 | Lysine-spe | Phf2 | sp\|Q9WTU | 14 | 13 | 18,1 |
| Q9ERU3 | Q9ERU3 | Zinc finger | Znf22 | sp\|Q9ERU | 1 | 1 | 5,1 |
| Q9D3U0 | Q9D3U0 | Putative tR | Pus10 | sp\|Q9D3U | 15 | 15 | 32,4 |
| Q8VCN6 | Q8VCN6 | CD99 antig | Cd99 | sp\|Q8VCN | 1 | 1 | 13,1 |
| Q9R233 | Q9R233 | Tapasin | Tapbp | sp\|Q9R23 | 10 | 10 | 27,3 |
| Q8VDM6 | Q8VDM6 | Heterogen | Hnrnpul1 | sp\|Q8VDM | 29 | 29 | 40,6 |
| Q69ZB0 | Q69ZB0 | Leucine-ric | Lrrcc1 | sp\|Q69ZB | 5 | 5 | 5,4 |
| Q68ED3 | Q68ED3 | Non-canon | Papd5 | sp\|Q68ED | 14 | 14 | 33,2 |
| Q05117 | Q05117 | Tartrate-re | Acp5 | sp\|Q0511 | 6 | 6 | 28,7 |
| Q6P5E6 | Q6P5E6 | ADP-ribosy | Gga2 | sp\|Q6P5E | 6 | 5 | 17,6 |
| CON__Q2K | CON__Q2KJF1 | | | | 12 | 12 | 40,8 |
| Q64737 | Q64737 | Trifunction | Gart | sp\|Q6473 | 42 | 42 | 56,2 |
| O55201 | O55201 | Transcripti | Supt5h | sp\|O55201 | 41 | 41 | 47,3 |
| Q9DBX2 | Q9DBX2 | Phosducin- | Pdcl | sp\|Q9DBX | 6 | 6 | 26,6 |
| Q9WUP7 | Q9WUP7 | Ubiquitin c | Uchl5 | sp\|Q9WU | 19 | 19 | 75,7 |
| Q9D832 | Q9D832 | DnaJ homo | Dnajb4 | sp\|Q9D83 | 15 | 14 | 42,4 |
| Q810S1 | Q810S1 | Calcium un | Ccdc109b | sp\|Q810S1 | 11 | 11 | 36,2 |
| A6H5Y3 | A6H5Y3 | Methionin | Mtr | sp\|A6H5Y | 11 | 11 | 11,9 |
| Q9D4H8 | Q9D4H8 | Cullin-2 | Cul2 | sp\|Q9D4H | 28 | 28 | 46 |
| Q9D4V4 | Q9D4V4 | TATA box-l | Taf1d | sp\|Q9D4V | 2 | 2 | 7,1 |
| Q8BNU0 | Q8BNU0 | Armadillo | Armc6 | sp\|Q8BNU | 9 | 9 | 29,5 |
| Q8BGA7 | Q8BGA7 | Uncharacterized prot | | sp\|Q8BGA | 6 | 6 | 20,5 |
| Q99KW3 | Q99KW3 | TRIO and F | Triobp | sp\|Q99KW | 24 | 24 | 15 |
| P28271 | P28271 | Cytoplasm | Aco1 | sp\|P28271 | 29 | 28 | 42,2 |
| Q01853 | Q01853 | Transition | Vcp | sp\|Q0185 | 61 | 61 | 81,8 |
| Q9CWK8 | Q9CWK8 | Sorting ne | Snx2 | sp\|Q9CWK | 21 | 19 | 47,2 |
| Q9DCJ5 | Q9DCJ5 | NADH deh | Ndufa8 | sp\|Q9DCJ5 | 9 | 9 | 61 |
| Q6RHR9 | Q6RHR9 | Membrane | Magi1 | sp\|Q6RHR | 20 | 19 | 24,9 |
| Q9JLV6 | Q9JLV6 | Bifunction | Pnkp | sp\|Q9JLV6 | 20 | 20 | 52,1 |

| | | | | | | | |
|---|---|---|---|---|---|---|---|
| Q8QZR4 | Q8QZR4 | Out at first | Oaf | sp\|Q8QZR4 | 7 | 7 | 27,7 |
| E9PXF8 | E9PXF8 | | Sbf2 | sp\|E9PXF8 | 15 | 14 | 11,6 |
| Q9JLQ2 | Q9JLQ2 | ARF GTPas | Git2 | sp\|Q9JLQ2 | 17 | 13 | 37,6 |
| Q00780 | Q00780 | Collagen al | Col8a1 | sp\|Q00780 | 10 | 10 | 17,3 |
| P11370 | P11370 | Retrovirus- | Fv4 | sp\|P11370 | 7 | 3 | 9,6 |
| Q64373 | Q64373 | Bcl-2-like p | Bcl2l1 | sp\|Q64373 | 6 | 6 | 36,9 |
| Q8C5L7 | Q8C5L7 | RNA-bindir | Rbm34 | sp\|Q8C5L7 | 18 | 18 | 49,8 |
| Q6A058 | Q6A058 | Armadillo r | Armcx2 | sp\|Q6A058 | 5 | 5 | 13,5 |
| Q9JHW2 | Q9JHW2 | Omega-am | Nit2 | sp\|Q9JHW | 14 | 14 | 60,1 |
| Q7TSE6 | Q7TSE6 | Serine/thre | Stk38l | sp\|Q7TSE6 | 5 | 4 | 13,8 |
| Q9Z2L6 | Q9Z2L6 | Multiple in | Minpp1 | sp\|Q9Z2L6 | 10 | 10 | 28,7 |
| Q9WTQ5 | Q9WTQ5 | A-kinase ar | Akap12 | sp\|Q9WTQ | 31 | 31 | 33,1 |
| Q9CYZ2 | Q9CYZ2 | Tumor pro | Tpd52l2 | sp\|Q9CYZ2 | 11 | 11 | 55 |
| Q8CI94;Q9 | Q8CI94 | Glycogen p | Pygb | sp\|Q8CI94 | 32 | 26 | 47,9 |
| Q61941 | Q61941 | NAD(P) tra | Nnt | sp\|Q61941 | 33 | 33 | 37 |
| B1AQJ2 | B1AQJ2 | Ubiquitin c | Usp36 | sp\|B1AQJ2 | 17 | 17 | 24,2 |
| Q9CSV6 | Q9CSV6 | Vesicle tra | Sft2d3 | sp\|Q9CSV6 | 2 | 2 | 14,2 |
| Q9WUR2;( | Q9WUR2 | Enoyl-CoA | Eci2 | sp\|Q9WUR | 11 | 11 | 36,6 |
| P22315 | P22315 | Ferrochela | Fech | sp\|P22315 | 11 | 11 | 36 |
| Q9Z2U0;Q | Q9Z2U0 | Proteasom | Psma7 | sp\|Q9Z2U0 | 15 | 15 | 56,5 |
| Q8BWB6 | Q8BWB6 | Metallored | Steap2 | sp\|Q8BWB | 13 | 13 | 34,4 |
| Q3UDE2 | Q3UDE2 | Tubulin--ty | Ttll12 | sp\|Q3UDE | 31 | 31 | 65,6 |
| Q6A037 | Q6A037 | NEDD4-bin | N4bp1 | sp\|Q6A037 | 8 | 8 | 12,3 |
| Q4G5Y1 | Q4G5Y1 | Kelch dom | Klhdc2 | sp\|Q4G5Y1 | 7 | 7 | 26,4 |
| Q61321 | Q61321 | Homeobox | Six4 | sp\|Q61321 | 5 | 4 | 13 |
| Q8BHX3 | Q8BHX3 | Borealin | Cdca8 | sp\|Q8BHX3 | 12 | 12 | 55 |
| P45878 | P45878 | Peptidyl-pr | Fkbp2 | sp\|P45878 | 6 | 6 | 50,7 |
| Q8R2Y0 | Q8R2Y0 | Monoacylg | Abhd6 | sp\|Q8R2Y0 | 11 | 11 | 43,2 |
| Q8K1X1 | Q8K1X1 | WD repeat | Wdr11 | sp\|Q8K1X1 | 30 | 30 | 33,2 |
| Q9DB20 | Q9DB20 | ATP syntha | Atp5o | sp\|Q9DB20 | 11 | 11 | 56,3 |
| Q8CE96 | Q8CE96 | tRNA (ader | Trmt6 | sp\|Q8CE96 | 13 | 13 | 32 |
| Q8R5F7 | Q8R5F7 | Interferon- | Ifih1 | sp\|Q8R5F7 | 7 | 7 | 9 |
| Q9QYL7 | Q9QYL7 | Activator c | Abt1 | sp\|Q9QYL7 | 10 | 10 | 54,6 |
| Q8K1E6 | Q8K1E6 | Alpha-keto | Alkbh3 | sp\|Q8K1E6 | 7 | 7 | 35,3 |
| Q6PA06 | Q6PA06 | Atlastin-2 | Atl2 | sp\|Q6PA06 | 14 | 13 | 38,9 |
| Q9CRC0 | Q9CRC0 | Vitamin K e | Vkorc1 | sp\|Q9CRC0 | 1 | 1 | 8,1 |
| Q9JK38 | Q9JK38 | Glucosami | Gnpnat1 | sp\|Q9JK38 | 8 | 8 | 64,7 |
| P54823 | P54823 | Probable A | Ddx6 | sp\|P54823 | 27 | 27 | 72,3 |
| Q8BWT5 | Q8BWT5 | Disco-inter | Dip2a | sp\|Q8BWT | 19 | 17 | 18,6 |
| Q3U2A8 | Q3U2A8 | Valine--tRN | Vars2 | sp\|Q3U2A | 16 | 16 | 18,5 |
| P05531 | P05531 | X-linked ly | Xlr | sp\|P05531 | 4 | 4 | 20,2 |
| Q922Q4 | Q922Q4 | Pyrroline-5 | Pycr2 | sp\|Q922Q | 15 | 12 | 55,6 |
| P17710 | P17710 | Hexokinase | Hk1 | sp\|P17710 | 42 | 36 | 47,9 |
| Q80Y84 | Q80Y84 | Lysine-spe | Kdm5b | sp\|Q80Y84 | 7 | 5 | 5,9 |
| P56818 | P56818 | Beta-secre | Bace1 | sp\|P56818 | 4 | 4 | 11,2 |
| Q9Z2R6 | Q9Z2R6 | Protein un | Unc119 | sp\|Q9Z2R6 | 6 | 5 | 36,2 |
| Q91VY5 | Q91VY5 | Lysine-spe | Kdm4b | sp\|Q91VY5 | 12 | 12 | 17 |
| Q60610 | Q60610 | T-lymphon | Tiam1 | sp\|Q60610 | 7 | 7 | 7,5 |
| P53395 | P53395 | Lipoamide | Dbt | sp\|P53395 | 6 | 6 | 15,6 |
| P10810 | P10810 | Monocyte | Cd14 | sp\|P10810 | 8 | 8 | 28,4 |

| | | | | | | | |
|---|---|---|---|---|---|---|---|
| Q61810 | Q61810 | Latent-trar | Ltbp3 | sp\|Q61810 | 16 | 16 | 21,9 |
| O08992 | O08992 | Syntenin-1 | Sdcbp | sp\|O08992 | 8 | 8 | 39,1 |
| Q64151 | Q64151 | Semaphori | Sema4c | sp\|Q64151 | 11 | 11 | 15,6 |
| Q9Z0T9 | Q9Z0T9 | Integrin be | Itgb6 | sp\|Q9Z0T9 | 1 | 1 | 1,1 |
| Q8R5K2 | Q8R5K2 | Ubiquitin c | Usp33 | sp\|Q8R5K2 | 5 | 5 | 11,2 |
| Q9CQ65 | Q9CQ65 | S-methyl-5 | Mtap | sp\|Q9CQ6 | 22 | 22 | 84,8 |
| Q80T85 | Q80T85 | DDB1- and | Dcaf5 | sp\|Q80T85 | 9 | 9 | 14 |
| Q9R062 | Q9R062 | Glycogenir | Gyg1 | sp\|Q9R062 | 8 | 8 | 34,5 |
| Q8K2F8 | Q8K2F8 | Protein LSI | Lsm14a | sp\|Q8K2F8 | 12 | 12 | 22,1 |
| Q8BYU6 | Q8BYU6 | Torsin-1A-i | Tor1aip2 | sp\|Q8BYU | 11 | 11 | 31,3 |
| P46656 | P46656 | Adrenodox | Fdx1 | sp\|P46656 | 4 | 4 | 20,2 |
| Q9DD02 | Q9DD02 | Protein Hik | L7rn6 | sp\|Q9DD0 | 5 | 5 | 27,4 |
| Q9D6Y9 | Q9D6Y9 | 1,4-alpha-g | Gbe1 | sp\|Q9D6Y | 13 | 13 | 25,6 |
| Q61194 | Q61194 | Phosphatic | Pik3c2a | sp\|Q61194 | 28 | 28 | 20,3 |
| Q8K135 | Q8K135 | Dyslexia-as | Kiaa0319l | sp\|Q8K135 | 6 | 6 | 7 |
| Q91YU8 | Q91YU8 | Suppressor | Ppan | sp\|Q91YU8 | 14 | 14 | 38,5 |
| Q03145;Q | Q03145 | Ephrin type | Epha2 | sp\|Q03145 | 34 | 32 | 45 |
| CON__Q58 | CON__Q58D62 | | | | 9 | 9 | 27,9 |
| Q9ER71 | Q9ER71 | Rho-relate | Rhoj | sp\|Q9ER71 | 2 | 1 | 11,7 |
| O35855 | O35855 | Branched-c | Bcat2 | sp\|O35855 | 11 | 11 | 36,9 |
| P70665 | P70665 | Sialate O-a | Siae | sp\|P70665 | 5 | 5 | 12 |
| Q8R3J4 | Q8R3J4 | Transcripti | Mterf3 | sp\|Q8R3J4 | 11 | 11 | 29,4 |
| Q8C0L6 | Q8C0L6 | Peroxisom | Paox | sp\|Q8C0L6 | 8 | 8 | 20,6 |
| Q8C569 | Q8C569 | Protein FA | Fam118b | sp\|Q8C569 | 7 | 7 | 21,1 |
| Q9D4V0 | Q9D4V0 | Ethanolam | Etnk1 | sp\|Q9D4V | 7 | 7 | 21,8 |
| P97358 | P97358 | TATA box-l | Taf1b | sp\|P97358 | 8 | 8 | 18,3 |
| Q8K2Q7 | Q8K2Q7 | BRO1 dom | Brox | sp\|Q8K2Q | 10 | 10 | 33,1 |
| Q6PFD9 | Q6PFD9 | Nuclear po | Nup98 | sp\|Q6PFD | 56 | 56 | 42,7 |
| Q9D8M4 | Q9D8M4 | 60S riboso | Rpl7l1 | sp\|Q9D8M | 10 | 10 | 34,6 |
| Q8BFY9 | Q8BFY9 | Transporti | Tnpo1 | sp\|Q8BFY9 | 31 | 24 | 44,1 |
| P23359 | P23359 | Bone morp | Bmp7 | sp\|P23359 | 2 | 2 | 5,3 |
| Q8C1E7 | Q8C1E7 | Transmem | Tmem120a | sp\|Q8C1E7 | 7 | 7 | 25,1 |
| O35730 | O35730 | E3 ubiquiti | Ring1 | sp\|O35730 | 8 | 6 | 28,8 |
| Q3TKT4 | Q3TKT4 | Transcripti | Smarca4 | sp\|Q3TKT4 | 46 | 33 | 34,9 |
| Q80TL7 | Q80TL7 | Protein MC | Mon2 | sp\|Q80TL7 | 32 | 32 | 22 |
| Q5DU41 | Q5DU41 | Volume-re | Lrrc8b | sp\|Q5DU4 | 7 | 6 | 12 |
| Q80U95 | Q80U95 | Ubiquitin-p | Ube3c | sp\|Q80U9 | 35 | 35 | 45 |
| Q8VEE4 | Q8VEE4 | Replicatior | Rpa1 | sp\|Q8VEE4 | 23 | 23 | 39,6 |
| Q78JW9 | Q78JW9 | Ubiquitin c | Ubfd1 | sp\|Q78JW | 7 | 7 | 29,1 |
| Q499X9 | Q499X9 | Methionin | Mars2 | sp\|Q499X9 | 12 | 12 | 31,1 |
| P30355 | P30355 | Arachidona | Alox5ap | sp\|P30355 | 3 | 3 | 20,5 |
| Q62141 | Q62141 | Paired amp | Sin3b | sp\|Q62141 | 25 | 24 | 33,5 |
| A2AN08 | A2AN08 | E3 ubiquiti | Ubr4 | sp\|A2AN0 | 104 | 104 | 27,9 |
| Q8CCP0 | Q8CCP0 | Nuclear ex | Nemf | sp\|Q8CCP | 23 | 23 | 29,2 |
| Q8BKI2 | Q8BKI2 | Trinucleoti | Tnrc6b | sp\|Q8BKI2 | 24 | 24 | 23,3 |
| CON__P05 | CON__P05784 | Keratin, ty | Krt18 | ;sp\|P0578 | 43 | 40 | 84,4 |
| Q811Q9 | Q811Q9 | Choline-ph | Pcyt1b | sp\|Q811Q | 7 | 3 | 30,4 |
| Q3V1H1 | Q3V1H1 | Cytoskelet | Ckap2 | sp\|Q3V1H | 14 | 14 | 31,9 |
| Q9Z103 | Q9Z103 | Activity-de | Adnp | sp\|Q9Z103 | 37 | 37 | 41,6 |
| P15535 | P15535 | Beta-1,4-ga | B4galt1 | sp\|P15535 | 4 | 4 | 16,5 |

| | | | | | | | |
|---|---|---|---|---|---|---|---|
| Q8BJT9 | Q8BJT9 | alpha-1,2-l | Edem2 | sp\|Q8BJT9 | 5 | 5 | 12,1 |
| Q148V7 | Q148V7 | LisH domai | Kiaa1468 | sp\|Q148V7 | 24 | 24 | 27,1 |
| O35604 | O35604 | Niemann-P | Npc1 | sp\|O35604 | 7 | 7 | 6,1 |
| Q8R3C0 | Q8R3C0 | Mini-chron | Mcmbp | sp\|Q8R3C0 | 22 | 22 | 45,8 |
| Q6PER3 | Q6PER3 | Microtubu | Mapre3 | sp\|Q6PER3 | 11 | 8 | 40,6 |
| Q8BHL3 | Q8BHL3 | TBC1 doma | Tbc1d10b | sp\|Q8BHL3 | 12 | 11 | 24,7 |
| Q8BH74 | Q8BH74 | Nuclear po | Nup107 | sp\|Q8BH74 | 39 | 39 | 52,7 |
| Q8VEM8 | Q8VEM8 | Phosphate | Slc25a3 | sp\|Q8VEM | 18 | 18 | 57,1 |
| Q8R409 | Q8R409 | Protein HE | Hexim1 | sp\|Q8R409 | 14 | 14 | 50 |
| Q3TW96 | Q3TW96 | UDP-N-ace | Uap1l1 | sp\|Q3TW9 | 25 | 23 | 58,6 |
| P47857 | P47857 | ATP-depen | Pfkm | sp\|P47857 | 24 | 21 | 40,1 |
| P35486 | P35486 | Pyruvate d | Pdha1 | sp\|P35486 | 23 | 23 | 63,8 |
| O09174 | O09174 | Alpha-met | Amacr | sp\|O09174 | 10 | 10 | 36,5 |
| Q9WUD1 | Q9WUD1 | STIP1 hom | Stub1 | sp\|Q9WUD | 12 | 12 | 52,6 |
| Q8BGX1;Q | Q8BGX1 | PC-esteras | Pced1b | sp\|Q8BGX | 5 | 5 | 12,9 |
| Q8CBY1 | Q8CBY1 | Protein Sm | Samd4a | sp\|Q8CBY1 | 17 | 16 | 35 |
| Q6PHZ5 | Q6PHZ5 | Putative RI | Rbm15b | sp\|Q6PHZ5 | 14 | 13 | 26,2 |
| Q9CXF7 | Q9CXF7 | Chromodo | Chd1l | sp\|Q9CXF7 | 18 | 18 | 24,6 |
| Q71FD5 | Q71FD5 | E3 ubiquiti | Znrf2 | sp\|Q71FD5 | 3 | 2 | 12,6 |
| Q52KI8 | Q52KI8 | Serine/arg | Srrm1 | sp\|Q52KI8 | 12 | 12 | 16 |
| CON__Q3K | CON__Q3KUS7 | | | | 6 | 6 | 13,7 |
| Q6P9R1 | Q6P9R1 | ATP-depen | Ddx51 | sp\|Q6P9R1 | 21 | 21 | 41 |
| O70493 | O70493 | Sorting nex | Snx12 | sp\|O70493 | 10 | 8 | 61,8 |
| Q8CGU1 | Q8CGU1 | Calcium-bi | Calcoco1 | sp\|Q8CGU | 6 | 6 | 10 |
| Q8C0V9 | Q8C0V9 | FERM dom | Frmd6 | sp\|Q8C0V9 | 10 | 10 | 17,5 |
| O08760 | O08760 | N-glycosyla | Ogg1 | sp\|O08760 | 4 | 4 | 15,4 |
| Q8CEE7 | Q8CEE7 | Retinol del | Rdh13 | sp\|Q8CEE7 | 9 | 9 | 31,7 |
| P98083;Q6 | P98083 | SHC-transf | Shc1 | sp\|P98083 | 13 | 13 | 32 |
| O08599 | O08599 | Syntaxin-b | Stxbp1 | sp\|O08599 | 18 | 18 | 43,3 |
| O08784 | O08784 | Treacle pro | Tcof1 | sp\|O08784 | 26 | 26 | 23,4 |
| Q9QXB9 | Q9QXB9 | Developme | Drg2 | sp\|Q9QXB | 17 | 17 | 51,4 |
| Q8R105 | Q8R105 | Vacuolar p | Vps37c | sp\|Q8R105 | 6 | 6 | 26,7 |
| Q9CVI2 | Q9CVI2 | Protein FA | Fam133b | sp\|Q9CVI2 | 3 | 3 | 11,8 |
| Q9ERA0 | Q9ERA0 | Alpha-glob | Tfcp2 | sp\|Q9ERA0 | 4 | 3 | 12,4 |
| O08738 | O08738 | Caspase-6; | Casp6 | sp\|O08738 | 13 | 13 | 45,7 |
| P43346 | P43346 | Deoxycytid | Dck | sp\|P43346 | 10 | 10 | 48,5 |
| P55302 | P55302 | Alpha-2-m | Lrpap1 | sp\|P55302 | 11 | 11 | 29,7 |
| Q5SNZ0 | Q5SNZ0 | Girdin | Ccdc88a | sp\|Q5SNZ0 | 31 | 31 | 21,5 |
| O35632 | O35632 | Hyaluronid | Hyal2 | sp\|O35632 | 5 | 5 | 18,4 |
| Q06831;Q7 | Q06831 | Transcripti | Sox4 | sp\|Q06831 | 4 | 4 | 16,6 |
| Q9WV84 | Q9WV84 | Nucleoside | Nme4 | sp\|Q9WV8 | 4 | 4 | 27,4 |
| P12382 | P12382 | ATP-depen | Pfkl | sp\|P12382 | 29 | 25 | 47,3 |
| Q9CQH3 | Q9CQH3 | NADH dehy | Ndufb5 | sp\|Q9CQH | 8 | 8 | 31,2 |
| P97364 | P97364 | Selenide, w | Sephs2 | sp\|P97364 | 9 | 9 | 26,1 |
| Q8CHY6 | Q8CHY6 | Transcripti | Gatad2a | sp\|Q8CHY6 | 18 | 17 | 39,9 |
| Q61147;C0 | Q61147 | Ceruloplas | Cp | sp\|Q61147 | 16 | 16 | 21,4 |
| Q9Z2B9 | Q9Z2B9 | Ribosomal | Rps6ka4 | sp\|Q9Z2B9 | 18 | 18 | 32,9 |
| P57724 | P57724 | Poly(rC)-bi | Pcbp4 | sp\|P57724 | 5 | 4 | 18,9 |
| Q9DCC4 | Q9DCC4 | Pyrroline-5 | Pycrl | sp\|Q9DCC4 | 6 | 6 | 31 |
| Q8K4X7 | Q8K4X7 | 1-acyl-sn-g | Agpat4 | sp\|Q8K4X7 | 10 | 10 | 26,2 |

| | | | | | | | |
|---|---|---|---|---|---|---|---|
| Q8R332 | Q8R332 | Nucleopor | Nupl1 | sp\|Q8R332 | 8 | 8 | 21,8 |
| Q99K85 | Q99K85 | Phosphose | Psat1 | sp\|Q99K85 | 18 | 18 | 52,2 |
| Q63961 | Q63961 | Endoglin | Eng | sp\|Q63961 | 6 | 6 | 13,5 |
| P14152 | P14152 | Malate del | Mdh1 | sp\|P14152 | 12 | 12 | 38,6 |
| Q4FK66 | Q4FK66 | Pre-mRNA- | Prpf38a | sp\|Q4FK66 | 14 | 14 | 31,4 |
| P19096 | P19096 | Fatty acid s | Fasn | sp\|P19096 | 106 | 106 | 60 |
| P97792 | P97792 | Coxsackiev | Cxadr | sp\|P97792 | 10 | 10 | 37,5 |
| Q8C0P5 | Q8C0P5 | Coronin-2A | Coro2a | sp\|Q8C0P5 | 6 | 5 | 13 |
| Q9QXP4 | Q9QXP4 | Protein do | Donson | sp\|Q9QXP | 4 | 4 | 11,2 |
| P18406 | P18406 | Protein CY | Cyr61 | sp\|P18406 | 11 | 11 | 39,8 |
| Q923B1 | Q923B1 | Lariat debr | Dbr1 | sp\|Q923B1 | 11 | 11 | 24,4 |
| P28704;P2 | P28704 | Retinoic ac | Rxrb | sp\|P28704 | 11 | 7 | 31,5 |
| P68433 | P68433 | Histone H3 | Hist1h3a | sp\|P68433 | 8 | 1 | 55,9 |
| Q9CYD3 | Q9CYD3 | Cartilage-a | Crtap | sp\|Q9CYD3 | 9 | 9 | 27 |
| Q3TMX7 | Q3TMX7 | Sulfhydryl | Qsox2 | sp\|Q3TMX | 9 | 9 | 19,1 |
| Q3TL44 | Q3TL44 | NLR family | Nlrx1 | sp\|Q3TL44 | 10 | 10 | 15,2 |
| Q9CZL5 | Q9CZL5 | Pterin-4-al | Pcbd2 | sp\|Q9CZL5 | 5 | 5 | 37,5 |
| Q1HFZ0 | Q1HFZ0 | tRNA (cyto | Nsun2 | sp\|Q1HFZ0 | 44 | 44 | 70 |
| P70336 | P70336 | Rho-associ | Rock2 | sp\|P70336 | 42 | 37 | 39,5 |
| Q8K3A9 | Q8K3A9 | 7SK snRNA | Mepce | sp\|Q8K3A9 | 19 | 19 | 48,6 |
| P54103 | P54103 | DnaJ homo | Dnajc2 | sp\|P54103 | 22 | 22 | 44,8 |
| P17182 | P17182 | Alpha-enol | Eno1 | sp\|P17182 | 33 | 28 | 78,6 |
| Q9WVA4 | Q9WVA4 | Transgelin- | Tagln2 | sp\|Q9WVA | 20 | 17 | 93,5 |
| Q921W4 | Q921W4 | Quinone o | Cryzl1 | sp\|Q921W | 10 | 10 | 37,6 |
| P98197 | P98197 | Probable p | Atp11a | sp\|P98197 | 11 | 8 | 10,3 |
| Q8BZ20 | Q8BZ20 | Poly [ADP- | Parp12 | sp\|Q8BZ20 | 12 | 12 | 19,8 |
| Q8CAK1 | Q8CAK1 | Putative tr | Iba57 | sp\|Q8CAK1 | 5 | 5 | 26,3 |
| Q64669 | Q64669 | NAD(P)H d | Nqo1 | sp\|Q64669 | 7 | 7 | 34,7 |
| Q8K1A6 | Q8K1A6 | Coiled-coil | Cc2d1a | sp\|Q8K1A6 | 17 | 17 | 23,3 |
| Q921U8 | Q921U8 | Smootheli | Smtn | sp\|Q921U8 | 23 | 23 | 34,3 |
| Q80XS6 | Q80XS6 | Protein Sm | Samd4b | sp\|Q80XS6 | 14 | 13 | 35,7 |
| Q7TSF4 | Q7TSF4 | Leucine-ric | Lrrc75a | sp\|Q7TSF4 | 5 | 5 | 21,8 |
| Q8BIW9 | Q8BIW9 | Chromosoi | Chtf18 | sp\|Q8BIW9 | 19 | 19 | 29,1 |
| Q9WUU9 | Q9WUU9 | Germinal-c | Mcm3ap | sp\|Q9WUU | 30 | 30 | 26 |
| P16330 | P16330 | 2,3-cyclic-r | Cnp | sp\|P16330 | 11 | 11 | 30,7 |
| Q8K330 | Q8K330 | Protein ph | Ssh3 | sp\|Q8K330 | 7 | 7 | 18 |
| Q64105 | Q64105 | Sepiapteri | Spr | sp\|Q64105 | 14 | 14 | 63,2 |
| Q9Z1K5 | Q9Z1K5 | E3 ubiquiti | Arih1 | sp\|Q9Z1K5 | 19 | 19 | 53,5 |
| Q9D906 | Q9D906 | Ubiquitin-l | Atg7 | sp\|Q9D906 | 21 | 21 | 40,8 |
| Q6P3A8 | Q6P3A8 | 2-oxoisova | Bckdhb | sp\|Q6P3A8 | 10 | 10 | 36,4 |
| A3KGF9 | A3KGF9 | Uncharacterized prote | | sp\|A3KGF9 | 5 | 5 | 18,2 |
| Q2HXL6 | Q2HXL6 | ER degrada | Edem3 | sp\|Q2HXL6 | 15 | 15 | 23,5 |
| Q60698 | Q60698 | Ski oncoge | Ski | sp\|Q60698 | 4 | 4 | 9,1 |
| A8C756 | A8C756 | Thyroid ad | Thada | sp\|A8C756 | 36 | 36 | 22,8 |
| P97363 | P97363 | Serine palr | Sptlc2 | sp\|P97363 | 16 | 16 | 40,7 |
| Q6ZQB6 | Q6ZQB6 | Inositol he | Ppip5k2 | sp\|Q6ZQB6 | 21 | 16 | 22,8 |
| Q8CFC7 | Q8CFC7 | CLK4-assoc | Clasrp | sp\|Q8CFC7 | 6 | 6 | 12,3 |
| Q68FD5 | Q68FD5 | Clathrin he | Cltc | sp\|Q68FD5 | 95 | 95 | 63,2 |
| Q9JLI8 | Q9JLI8 | Squamous | Sart3 | sp\|Q9JLI8\| | 41 | 41 | 50,5 |
| Q9D0C4 | Q9D0C4 | tRNA (guar | Trmt5 | sp\|Q9D0C4 | 16 | 16 | 35,7 |

| | | | | | | | |
|---|---|---|---|---|---|---|---|
| Q62036 | Q62036 | Centrosom | Cep131 | sp\|Q62036 | 16 | 16 | 20,4 |
| P54818 | P54818 | Galactocer | Galc | sp\|P54818 | 9 | 9 | 20,9 |
| Q9DB05;P1 | Q9DB05 | Alpha-solu | Napa | sp\|Q9DB0 | 19 | 19 | 75,3 |
| Q8BWY4 | Q8BWY4 | Metallo-be | Mblac1 | sp\|Q8BWY | 2 | 2 | 11,5 |
| Q62187 | Q62187 | Transcripti | Ttf1 | sp\|Q62187 | 9 | 9 | 16,2 |
| Q8BJF9 | Q8BJF9 | Charged m | Chmp2b | sp\|Q8BJF9 | 3 | 2 | 12,7 |
| P12023 | P12023 | Amyloid be | App | sp\|P12023 | 15 | 15 | 29,1 |
| Q9JI75 | Q9JI75 | Ribosyldih | Nqo2 | sp\|Q9JI75 | 11 | 11 | 61 |
| Q922S8 | Q922S8 | Kinesin-like | Kif2c | sp\|Q922S8 | 17 | 17 | 32,7 |
| Q61337 | Q61337 | Bcl2-associ | Bad | sp\|Q61337 | 5 | 5 | 39,2 |
| Q9DBL9 | Q9DBL9 | 1-acylglyce | Abhd5 | sp\|Q9DBL9 | 4 | 4 | 19,1 |
| Q9JMA2 | Q9JMA2 | Queuine tR | Qtrt1 | sp\|Q9JMA | 8 | 8 | 30,3 |
| Q71RI9 | Q71RI9 | Kynurenine | Ccbl2 | sp\|Q71RI9 | 10 | 10 | 27,9 |
| Q8VDN2;Q | Q8VDN2 | Sodium/po | Atp1a1 | sp\|Q8VDN | 41 | 41 | 44,8 |
| Q9CQX8 | Q9CQX8 | 28S riboso | Mrps36 | sp\|Q9CQX | 5 | 5 | 70,6 |
| P97499 | P97499 | Telomeras | Tep1 | sp\|P97499 | 16 | 16 | 9,4 |
| Q9JLR9 | Q9JLR9 | HIG1 doma | Higd1a | sp\|Q9JLR9 | 1 | 1 | 18,9 |
| Q60751 | Q60751 | Insulin-like | Igf1r | sp\|Q60751 | 20 | 18 | 19,4 |
| O54916;Q8 | O54916 | RalBP1-ass | Reps1 | sp\|O54916 | 16 | 16 | 31,2 |
| A2AWL7 | A2AWL7 | MAX gene- | Mga | sp\|A2AWL | 13 | 13 | 7 |
| Q8R322 | Q8R322 | Nucleopor | Gle1 | sp\|Q8R322 | 16 | 16 | 30,3 |
| A2APV2 | A2APV2 | Formin-like | Fmnl2 | sp\|A2APV2 | 21 | 14 | 20,8 |
| O70152 | O70152 | Dolichol-ph | Dpm1 | sp\|O70152 | 11 | 11 | 47,7 |
| Q3UI43 | Q3UI43 | BRISC and | Babam1 | sp\|Q3UI43 | 6 | 6 | 32,1 |
| Q7TT37 | Q7TT37 | Elongator c | Ikbkap | sp\|Q7TT37 | 34 | 34 | 35,7 |
| Q9ERS5 | Q9ERS5 | Pleckstrin h | Plekha2 | sp\|Q9ERS5 | 11 | 11 | 40,5 |
| Q7TSV9 | Q7TSV9 | Enkurin do | Enkd1 | sp\|Q7TSV9 | 3 | 3 | 12,7 |
| Q8JZM0 | Q8JZM0 | Dimethylad | Tfb1m | sp\|Q8JZM0 | 10 | 10 | 37,4 |
| Q8K297 | Q8K297 | Procollage | Colgalt1 | sp\|Q8K297 | 21 | 21 | 42,6 |
| Q62193 | Q62193 | Replication | Rpa2 | sp\|Q62193 | 7 | 7 | 39,3 |
| Q99MJ9 | Q99MJ9 | ATP-depen | Ddx50 | sp\|Q99MJ9 | 26 | 26 | 44,4 |
| Q3MI99 | Q3MI99 | Collagen ar | Ccbe1 | sp\|Q3MI99 | 7 | 7 | 29,2 |
| Q5FWI3 | Q5FWI3 | Transmem | Tmem2 | sp\|Q5FWI3 | 23 | 23 | 24,5 |
| Q9CXI0 | Q9CXI0 | 2-methoxy | Coq5 | sp\|Q9CXI0 | 11 | 11 | 48,9 |
| Q99LI5 | Q99LI5 | Zinc finger | Znf281 | sp\|Q99LI5 | 13 | 13 | 28,7 |
| Q9WTZ0 | Q9WTZ0 | Protein UX | Uxt | sp\|Q9WTZ | 2 | 2 | 15,3 |
| P70213 | P70213 | Friend viru | Fv1 | sp\|P70213 | 9 | 9 | 29 |
| Q921G8 | Q921G8 | Gamma-tu | Tubgcp2 | sp\|Q921G8 | 30 | 30 | 43,3 |
| Q9DC16 | Q9DC16 | Endoplasm | Ergic1 | sp\|Q9DC16 | 8 | 8 | 32,4 |
| Q99L47 | Q99L47 | Hsc70-inte | St13 | sp\|Q99L47 | 10 | 10 | 27,5 |
| P33610 | P33610 | DNA prima | Prim2 | sp\|P33610 | 20 | 20 | 47,3 |
| Q99NG0 | Q99NG0 | Helicase AI | Rad54l2 | sp\|Q99NG | 14 | 14 | 13,4 |
| P61979 | P61979 | Heterogen | Hnrnpk | sp\|P61979 | 27 | 27 | 65,7 |
| Q04863 | Q04863 | Transcripti | Relb | sp\|Q04863 | 8 | 8 | 21,1 |
| CON__P41 | CON__P41361 | | | | 13 | 13 | 32,7 |
| Q61329 | Q61329 | Zinc finger | Zfhx3 | sp\|Q61329 | 11 | 11 | 4,9 |
| Q99JY1 | Q99JY1 | Toll/interle | Tirap | sp\|Q99JY1 | 2 | 2 | 12 |
| P18826 | P18826 | Phosphory | Phka1 | sp\|P18826 | 11 | 11 | 14,2 |
| Q6NS46 | Q6NS46 | Protein RR | Pdcd11 | sp\|Q6NS46 | 70 | 70 | 42,5 |
| Q922P8 | Q922P8 | Transmem | Tmem132a | sp\|Q922P8 | 12 | 12 | 19 |

| | | | | | | | |
|---|---|---|---|---|---|---|---|
| Q3UUQ7 | Q3UUQ7 | GPI inosito | Pgap1 | sp\|Q3UUC | 7 | 7 | 8,7 |
| Q8VHE0 | Q8VHE0 | Translocati | Sec63 | sp\|Q8VHE( | 35 | 35 | 45,3 |
| Q920A7 | Q920A7 | AFG3-like | Afg3l1 | sp\|Q920A7 | 21 | 20 | 35,1 |
| Q99L48 | Q99L48 | 60S riboso | Nmd3 | sp\|Q99L48 | 16 | 16 | 36,2 |
| Q8K157 | Q8K157 | Aldose 1-e | Galm | sp\|Q8K157 | 4 | 4 | 17,5 |
| Q61881 | Q61881 | DNA replic | Mcm7 | sp\|Q61881 | 36 | 36 | 59 |
| Q91W82;P | Q91W82;P524 | Ubiquitin-c | Ube2e2;Ub | sp\|Q91W8 | 4 | 1 | 27,4 |
| Q8R164 | Q8R164 | Valacyclov | Bphl | sp\|Q8R164 | 12 | 12 | 54 |
| O88544 | O88544 | COP9 signa | Cops4 | sp\|O88544 | 24 | 24 | 78,6 |
| Q80VY9 | Q80VY9 | Putative A | Dhx33 | sp\|Q80VY9 | 18 | 18 | 26,2 |
| Q8R4R6 | Q8R4R6 | Nucleopor | Nup35 | sp\|Q8R4R( | 12 | 12 | 62,8 |
| Q60766 | Q60766 | Immunity- | Irgm1 | sp\|Q6076( | 6 | 6 | 18,1 |
| Q8JZU2 | Q8JZU2 | Tricarboxy | Slc25a1 | sp\|Q8JZU2 | 7 | 7 | 28,3 |
| P09671 | P09671 | Superoxide | Sod2 | sp\|P09671 | 8 | 8 | 38,3 |
| Q99P65 | Q99P65 | Equilibrati | Slc29a3 | sp\|Q99P65 | 3 | 3 | 10,3 |
| Q6PAL7 | Q6PAL7 | AT-hook D | Ahdc1 | sp\|Q6PAL7 | 28 | 28 | 27,3 |
| Q8R2E9 | Q8R2E9 | ERO1-like | Ero1lb | sp\|Q8R2E9 | 6 | 5 | 17,3 |
| Q62087 | Q62087 | Serum para | Pon3 | sp\|Q62087 | 12 | 12 | 51,1 |
| Q8BH47 | Q8BH47 | Vesicle-tra | Sec22a | sp\|Q8BH47 | 4 | 4 | 21,8 |
| P97372 | P97372 | Proteasom | Psme2 | sp\|P97372 | 12 | 12 | 53,6 |
| A2AI08 | A2AI08 | Taperin | Tprn | sp\|A2AI08 | 8 | 8 | 18,6 |
| O70279 | O70279 | Protein DG | Dgcr14 | sp\|O70279 | 11 | 11 | 35,5 |
| Q8K339 | Q8K339 | DNA/RNA- | Kin | sp\|Q8K339 | 7 | 7 | 22 |
| Q64695 | Q64695 | Endothelia | Procr | sp\|Q64695 | 2 | 2 | 12,4 |
| P49070 | P49070 | Calcium sig | Camlg | sp\|P49070 | 7 | 7 | 38,4 |
| O35382 | O35382 | Exocyst co | Exoc4 | sp\|O35382 | 28 | 28 | 33,7 |
| Q9CXJ1 | Q9CXJ1 | Probable g | Ears2 | sp\|Q9CXJ1 | 12 | 12 | 28,5 |
| P34152 | P34152 | Focal adhe | Ptk2 | sp\|P34152 | 26 | 25 | 37 |
| O08808 | O08808 | Protein dia | Diaph1 | sp\|O08808 | 26 | 26 | 26,7 |
| O54931 | O54931 | A-kinase a | Akap2 | sp\|O54931 | 23 | 23 | 40,1 |
| Q80ZK9 | Q80ZK9 | WD and te | Wdtc1 | sp\|Q80ZK9 | 9 | 9 | 17 |
| Q8R146 | Q8R146 | Acylamino- | Apeh | sp\|Q8R14( | 17 | 17 | 32,5 |
| B2RRE7 | B2RRE7 | OTU doma | Otud4 | sp\|B2RRE7 | 27 | 27 | 36,1 |
| P47879 | P47879 | Insulin-like | Igfbp4 | sp\|P47879 | 6 | 6 | 35,4 |
| P56391 | P56391 | Cytochrom | Cox6b1 | sp\|P56391 | 7 | 7 | 70,9 |
| Q8BFW7 | Q8BFW7 | Lipoma-pre | Lpp | sp\|Q8BFW | 18 | 18 | 43,2 |
| Q9JKY0 | Q9JKY0 | Cell differe | Rqcd1 | sp\|Q9JKY0 | 11 | 11 | 42,5 |
| Q9R269 | Q9R269 | Periplakin | Ppl | sp\|Q9R269 | 59 | 59 | 40,8 |
| Q05816 | Q05816 | Fatty acid- | Fabp5 | sp\|Q05816 | 12 | 12 | 72,6 |
| Q8BMI0 | Q8BMI0 | F-box only | Fbxo38 | sp\|Q8BMI( | 15 | 15 | 17,3 |
| P55194 | P55194 | SH3 domai | Sh3bp1 | sp\|P55194 | 10 | 10 | 19,4 |
| Q8C163 | Q8C163 | Nuclease E | Exog | sp\|Q8C163 | 9 | 9 | 43,8 |
| P98063;Q9 | P98063 | Bone morp | Bmp1 | sp\|P98063 | 13 | 13 | 16,8 |
| Q8BMK4 | Q8BMK4 | Cytoskelet | Ckap4 | sp\|Q8BMK | 31 | 31 | 59,5 |
| Q9CPQ1 | Q9CPQ1 | Cytochrom | Cox6c | sp\|Q9CPQ | 7 | 7 | 61,8 |
| Q61584 | Q61584 | Fragile X m | Fxr1 | sp\|Q61584 | 39 | 37 | 64,4 |
| Q7TNP2 | Q7TNP2 | Serine/thre | Ppp2r1b | sp\|Q7TNP2 | 26 | 17 | 55,2 |
| Q9WUQ2 | Q9WUQ2 | Prolactin r | Preb | sp\|Q9WU( | 14 | 14 | 47,7 |
| Q7TMA2 | Q7TMA2 | Zinc finger | Znf503 | sp\|Q7TMA | 3 | 2 | 7,1 |
| Q80ZW2 | Q80ZW2 | Protein TH | Them6 | sp\|Q80ZW | 10 | 10 | 43,5 |

| | | | | | | | |
|---|---|---|---|---|---|---|---|
| O35621 | O35621 | Phosphom | Pmm1 | sp\|O35621 | 10 | 8 | 46,9 |
| P21460 | P21460 | Cystatin-C | Cst3 | sp\|P21460 | 5 | 5 | 42,1 |
| Q9WV95 | Q9WV95 | Pleckstrin | Phlda3 | sp\|Q9WV9 | 3 | 3 | 20 |
| P28650 | P28650 | Adenylosu | Adssl1 | sp\|P28650 | 12 | 10 | 36,3 |
| Q9D0P8 | Q9D0P8 | Intraflagell | Ift27 | sp\|Q9D0P | 5 | 5 | 29 |
| Q8R5L3 | Q8R5L3 | Vam6/Vps | Vps39 | sp\|Q8R5L3 | 16 | 16 | 22,2 |
| P22907 | P22907 | Porphobili | Hmbs | sp\|P22907 | 12 | 12 | 41,8 |
| Q9EST8 | Q9EST8 | NF-kappa-I | Nfkbiz | sp\|Q9EST8 | 3 | 3 | 6,7 |
| Q6P5F7 | Q6P5F7 | Protein tw | Ttyh3 | sp\|Q6P5F7 | 3 | 3 | 9 |
| Q9WV55 | Q9WV55 | Vesicle-ass | Vapa | sp\|Q9WV5 | 13 | 12 | 48,6 |
| P55264 | P55264 | Adenosine | Adk | sp\|P55264 | 16 | 16 | 54,3 |
| Q9D720 | Q9D720 | Non-struct | Nsmce1 | sp\|Q9D72 | 4 | 4 | 25,6 |
| Q3UHC7 | Q3UHC7 | Disabled h | Dab2ip | sp\|Q3UHC | 10 | 10 | 12,6 |
| Q9Z2Z6 | Q9Z2Z6 | Mitochond | Slc25a20 | sp\|Q9Z2Z6 | 11 | 11 | 48,8 |
| Q9CQL0 | Q9CQL0 | Protein N-I | Mettl21A | sp\|Q9CQL | 2 | 2 | 11,5 |
| Q9DD18 | Q9DD18 | D-tyrosyl-t | Dtd1 | sp\|Q9DD1 | 2 | 2 | 20,1 |
| Q9DAW9 | Q9DAW9 | Calponin-3 | Cnn3 | sp\|Q9DAW | 14 | 13 | 51,2 |
| Q99LG2 | Q99LG2 | Transporti | Tnpo2 | sp\|Q99LG2 | 21 | 14 | 35,7 |
| D3YXK1 | D3YXK1 | Atherin | Samd1 | sp\|D3YXK1 | 9 | 9 | 21,8 |
| Q8BUM9 | Q8BUM9 | Ubiquitin c | Usp43 | sp\|Q8BUM | 10 | 10 | 13,6 |
| Q3SXD3 | Q3SXD3 | HD domain | Hddc2 | sp\|Q3SXD3 | 11 | 11 | 71,9 |
| A2A7S8 | A2A7S8 | Uncharacte | Kiaa1522 | sp\|A2A7S8 | 21 | 21 | 32,5 |
| Q3URD3 | Q3URD3 | Sarcolemm | Slmap | sp\|Q3URD | 23 | 23 | 33,1 |
| Q62371 | Q62371 | Discoidin d | Ddr2 | sp\|Q62371 | 17 | 17 | 30,3 |
| P28660;Q8 | P28660 | Nck-associ | Nckap1 | sp\|P28660 | 33 | 33 | 36,3 |
| Q8VCP8 | Q8VCP8 | Adenylate | Ak6 | sp\|Q8VCP8 | 4 | 4 | 29,7 |
| Q9Z183 | Q9Z183 | Protein-arg | Padi4 | sp\|Q9Z183 | 29 | 26 | 53,2 |
| Q5U462 | Q5U462 | CUB doma | Cdcp1 | sp\|Q5U462 | 18 | 18 | 26,9 |
| Q922J9 | Q922J9 | Fatty acyl-( | Far1 | sp\|Q922J9 | 26 | 26 | 51,5 |
| Q8BUH8 | Q8BUH8 | Sentrin-spe | Senp7 | sp\|Q8BUH | 13 | 13 | 20,3 |
| Q9R1R2 | Q9R1R2 | Tripartite r | Trim3 | sp\|Q9R1R2 | 19 | 18 | 34 |
| P70700 | P70700 | DNA-direct | Polr1b | sp\|P70700 | 25 | 25 | 25,5 |
| P58021 | P58021 | Transmem | Tm9sf2 | sp\|P58021 | 15 | 15 | 26,7 |
| Q9Z0W3 | Q9Z0W3 | Nuclear po | Nup160 | sp\|Q9Z0W | 39 | 39 | 37,1 |
| Q8C2Q3 | Q8C2Q3 | RNA-bindir | Rbm14 | sp\|Q8C2Q | 26 | 26 | 48 |
| P97742;Q9 | P97742 | Carnitine C | Cpt1a | sp\|P97742 | 17 | 17 | 27,3 |
| Q8R4U7 | Q8R4U7 | Leucine zip | Luzp1 | sp\|Q8R4U | 17 | 17 | 21,5 |
| Q6ZPJ0 | Q6ZPJ0 | Testis-expr | Tex2 | sp\|Q6ZPJ0 | 11 | 11 | 16,1 |
| Q8CBE3 | Q8CBE3 | WD repeat | Wdr37 | sp\|Q8CBE3 | 5 | 5 | 14,1 |
| Q7TSS2 | Q7TSS2 | Ubiquitin-c | Ube2q1 | sp\|Q7TSS2 | 10 | 8 | 31 |
| Q920Q8 | Q920Q8 | Influenza v | Ivns1abp | sp\|Q920Q | 14 | 14 | 34,6 |
| Q922B9 | Q922B9 | Sperm-spe | Ssfa2 | sp\|Q922B9 | 9 | 9 | 9,9 |
| P56546 | P56546 | C-terminal | Ctbp2 | sp\|P56546 | 15 | 11 | 41,6 |
| Q8K0Q5 | Q8K0Q5 | Rho GTPas | Arhgap18 | sp\|Q8K0Q | 16 | 16 | 31,2 |
| Q99PM9 | Q99PM9 | Uridine-cyt | Uck2 | sp\|Q99PM | 9 | 7 | 51 |
| Q8C5H8 | Q8C5H8 | NAD kinase | Nadk2 | sp\|Q8C5H8 | 15 | 15 | 47,1 |
| Q9Z1P7 | Q9Z1P7 | KN motif a | Kank3 | sp\|Q9Z1P7 | 3 | 3 | 6,3 |
| P62192 | P62192 | 26S protea | Psmc1 | sp\|P62192 | 24 | 24 | 59,8 |
| Q4VAE3 | Q4VAE3 | Transmem | Tmem65 | sp\|Q4VAE3 | 5 | 5 | 24,8 |
| Q9QYJ3 | Q9QYJ3 | DnaJ homo | Dnajb1 | sp\|Q9QYJ3 | 13 | 12 | 35,6 |

| | | | | | | | |
|---|---|---|---|---|---|---|---|
| P11031 | P11031 | Activated F | Sub1 | sp\|P11031 | 6 | 6 | 38,6 |
| Q8JZX4 | Q8JZX4 | Splicing fac | Rbm17 | sp\|Q8JZX4 | 16 | 16 | 44 |
| CON__ENS | CON__ENSEMBL:ENSBTAP00000034412 | | | | 6 | 6 | 30,7 |
| O08553;Q( | O08553 | Dihydropy | Dpysl2 | sp\|O08553 | 28 | 28 | 72 |
| Q9EP97 | Q9EP97 | Sentrin-spe | Senp3 | sp\|Q9EP97 | 17 | 17 | 41,2 |
| Q91VJ4 | Q91VJ4 | Serine/thre | Stk38 | sp\|Q91VJ4 | 10 | 9 | 31,4 |
| Q8BH02 | Q8BH02 | Torsin-4A | Tor4a | sp\|Q8BH0 | 7 | 7 | 24,4 |
| Q9D8N0 | Q9D8N0 | Elongation | Eef1g | sp\|Q9D8N | 27 | 27 | 63,2 |
| Q8R0W0 | Q8R0W0 | Epiplakin | Eppk1 | sp\|Q8R0W | 62 | 59 | 40,7 |
| Q91W40 | Q91W40 | Kinesin ligh | Klc3 | sp\|Q91W4 | 13 | 12 | 39,8 |
| Q7SIG6 | Q7SIG6 | Arf-GAP wi | Asap2 | sp\|Q7SIG6 | 24 | 21 | 39 |
| O70255 | O70255 | Myelin pro | Mpzl2 | sp\|O70255 | 2 | 2 | 10,2 |
| E9PVB5 | E9PVB5 | Tetratricop | Ttc17 | sp\|E9PVB5 | 10 | 10 | 14,1 |
| Q9WVM1 | Q9WVM1 | Rac GTPase | Racgap1 | sp\|Q9WVN | 25 | 25 | 50,5 |
| O08911 | O08911 | Mitogen-ac | Mapk12 | sp\|O08911 | 11 | 10 | 34,6 |
| Q9JI90 | Q9JI90 | E3 ubiquiti | Rnf14 | sp\|Q9JI90 | 6 | 6 | 15,3 |
| Q3UHU5 | Q3UHU5 | Microtubu | Mtcl1 | sp\|Q3UHU | 23 | 21 | 15,8 |
| Q99LX0 | Q99LX0 | Protein de | Park7 | sp\|Q99LX0 | 11 | 11 | 67,2 |
| Q9DBH5 | Q9DBH5 | Vesicular ir | Lman2 | sp\|Q9DBH | 14 | 14 | 46,9 |
| Q6PDX6 | Q6PDX6 | E3 ubiquiti | Rnf220 | sp\|Q6PDX( | 2 | 2 | 4,2 |
| P11672 | P11672 | Neutrophil | Lcn2 | sp\|P11672 | 3 | 3 | 22,5 |
| Q9JKZ2 | Q9JKZ2 | Sodium/m | Slc5a3 | sp\|Q9JKZ2 | 8 | 8 | 18,5 |
| Q8C5W3 | Q8C5W3 | Tubulin-sp | Tbcel | sp\|Q8C5W | 5 | 5 | 19,3 |
| Q3UJB9 | Q3UJB9 | Enhancer c | Edc4 | sp\|Q3UJB9 | 40 | 40 | 46,9 |
| O35969 | O35969 | Guanidino | Gamt | sp\|O35969 | 10 | 10 | 67,4 |
| Q60759 | Q60759 | Glutaryl-C | Gcdh | sp\|Q60759 | 8 | 8 | 23,1 |
| CON__P28 | CON__P28800 | | | | 5 | 5 | 13,8 |
| P59759 | P59759 | MKL/myoc | Mkl2 | sp\|P59759 | 13 | 12 | 20 |
| O08529 | O08529 | Calpain-2 c | Capn2 | sp\|O08529 | 36 | 36 | 60,1 |
| Q8VIM9 | Q8VIM9 | Immunity- | Irgq | sp\|Q8VIM9 | 8 | 8 | 19,9 |
| Q62426 | Q62426 | Cystatin-B | Cstb | sp\|Q62426 | 7 | 7 | 75,5 |
| Q8CFE2 | Q8CFE2 | UPF0609 protein C4or | | sp\|Q8CFE2 | 12 | 12 | 45,7 |
| Q8K009;Q8 | Q8K009 | Mitochond | Aldh1l2 | sp\|Q8K009 | 33 | 33 | 49,7 |
| P08030 | P08030 | Adenine ph | Aprt | sp\|P08030 | 10 | 10 | 71,1 |
| Q810A7 | Q810A7 | ATP-depen | Ddx42 | sp\|Q810A7 | 39 | 39 | 56,6 |
| P62849 | P62849 | 40S riboso | Rps24 | sp\|P62849 | 6 | 6 | 35,3 |
| Q80U04 | Q80U04 | E3 ubiquiti | Pja2 | sp\|Q80U04 | 6 | 6 | 18,5 |
| P43135 | P43135 | COUP tran | Nr2f2 | sp\|P43135 | 7 | 2 | 21 |
| O54956 | O54956 | DNA polym | Pole2 | sp\|O54956 | 11 | 11 | 31,5 |
| Q3UMC0;( | Q3UMC0 | Spermatog | Spata5 | sp\|Q3UMC | 27 | 27 | 41,3 |
| P34022 | P34022 | Ran-specifi | Ranbp1 | sp\|P34022 | 8 | 8 | 59,1 |
| Q7TMY8 | Q7TMY8 | E3 ubiquiti | Huwe1 | sp\|Q7TMY | 82 | 82 | 27,6 |
| Q9WVK4 | Q9WVK4 | EH domain | Ehd1 | sp\|Q9WVK | 38 | 25 | 70,2 |
| Q61624 | Q61624 | Zinc finger | Znf148 | sp\|Q61624 | 13 | 13 | 26,1 |
| P98078 | P98078 | Disabled h | Dab2 | sp\|P98078 | 19 | 19 | 45,3 |
| Q64435;Q( | Q64435 | UDP-glucu | Ugt1a6 | sp\|Q64435 | 11 | 10 | 27,1 |
| Q6P5E4 | Q6P5E4 | UDP-gluco | Uggt1 | sp\|Q6P5E4 | 71 | 71 | 58,8 |
| Q6PDG5 | Q6PDG5 | SWI/SNF c | Smarcc2 | sp\|Q6PDG | 33 | 24 | 31,5 |
| Q64674 | Q64674 | Spermidine | Srm | sp\|Q64674 | 20 | 20 | 78,8 |
| Q3TGF2 | Q3TGF2 | Protein FA | Fam107b | sp\|Q3TGF2 | 8 | 8 | 48,9 |

| | | | | | | | |
|---|---|---|---|---|---|---|---|
| P26443 | P26443 | Glutamate | Glud1 | sp\|P26443 | 27 | 27 | 51,4 |
| Q60770 | Q60770 | Syntaxin-b | Stxbp3 | sp\|Q6077C | 22 | 22 | 42,4 |
| Q68FH4 | Q68FH4 | N-acetylga | Galk2 | sp\|Q68FH4 | 9 | 9 | 32,1 |
| Q8K3K8 | Q8K3K8 | Optineurin | Optn | sp\|Q8K3K8 | 10 | 10 | 20,7 |
| Q3UFM5 | Q3UFM5 | Nucleolar I | Nom1 | sp\|Q3UFM | 17 | 17 | 26,9 |
| O35134 | O35134 | DNA-direct | Polr1a | sp\|O35134 | 51 | 51 | 45 |
| Q8C0D4 | Q8C0D4 | Rho GTPas | Arhgap12 | sp\|Q8C0D4 | 10 | 10 | 16,5 |
| Q61490 | Q61490 | CD166 ant | Alcam | sp\|Q6149C | 14 | 14 | 34,5 |
| A6H611 | A6H611 | Mitochond | Mipep | sp\|A6H611 | 16 | 16 | 30,4 |
| Q8K1R7 | Q8K1R7 | Serine/thr | Nek9 | sp\|Q8K1R7 | 21 | 21 | 27,1 |
| P17809 | P17809 | Solute carr | Slc2a1 | sp\|P17809 | 7 | 7 | 12,6 |
| Q6P5F9 | Q6P5F9 | Exportin-1 | Xpo1 | sp\|Q6P5F9 | 41 | 41 | 48,8 |
| Q91YJ5 | Q91YJ5 | Translation | Mtif2 | sp\|Q91YJ5 | 9 | 9 | 19,9 |
| Q8BVW3 | Q8BVW3 | Tripartite r | Trim14 | sp\|Q8BVW | 6 | 6 | 18,6 |
| Q924W5 | Q924W5 | Structural | Smc6 | sp\|Q924W | 28 | 28 | 32,7 |
| P60487 | P60487 | Pyridoxal p | Pdxp | sp\|P60487 | 5 | 5 | 27,4 |
| Q8BMC4 | Q8BMC4 | Nucleolar p | Nop9 | sp\|Q8BMC | 14 | 14 | 30,3 |
| Q3V384 | Q3V384 | Lactation e | Lace1 | sp\|Q3V384 | 9 | 9 | 28,5 |
| P70699 | P70699 | Lysosomal | Gaa | sp\|P70699 | 20 | 20 | 26,7 |
| Q8CAQ8 | Q8CAQ8 | MICOS con | Immt | sp\|Q8CAQ | 38 | 38 | 59,8 |
| Q8VDV8 | Q8VDV8 | MIT domai | Mitd1 | sp\|Q8VDV | 3 | 3 | 22,9 |
| Q9CQE7 | Q9CQE7 | Endoplasm | Ergic3 | sp\|Q9CQE | 3 | 3 | 13,8 |
| Q8VHI3 | Q8VHI3 | GDP-fucos | Pofut2 | sp\|Q8VHI3 | 13 | 13 | 35,4 |
| P0DMN7;D | P0DMN7;D3Z6 | S-adenosyl | Amd1;Amd | sp\|P0DMN | 4 | 4 | 14,1 |
| Q8JZM7 | Q8JZM7 | Parafibrom | Cdc73 | sp\|Q8JZM7 | 21 | 21 | 42,6 |
| Q9EQC5 | Q9EQC5 | N-terminal | Scyl1 | sp\|Q9EQC | 23 | 23 | 42,3 |
| Q8BSS9;P6 | Q8BSS9 | Liprin-alph | Ppfia2 | sp\|Q8BSS9 | 7 | 7 | 6,4 |
| Q80U38 | Q80U38 | Protein KH | Khnyn | sp\|Q80U38 | 7 | 7 | 15,1 |
| P97868 | P97868 | E3 ubiquiti | Rbbp6 | sp\|P97868 | 24 | 24 | 18,7 |
| Q8BY87 | Q8BY87 | Ubiquitin c | Usp47 | sp\|Q8BY87 | 45 | 45 | 45,9 |
| Q9R1C7 | Q9R1C7 | Pre-mRNA- | Prpf40a | sp\|Q9R1C7 | 23 | 23 | 25,7 |
| P28662 | P28662 | Brain prote | Bri3 | sp\|P28662 | 1 | 1 | 12,8 |
| Q8BT60;Q8 | Q8BT60 | Copine-3 | Cpne3 | sp\|Q8BT6C | 18 | 16 | 43 |
| O55111 | O55111 | Desmoglei | Dsg2 | sp\|O55111 | 11 | 11 | 15,3 |
| Q8CI11 | Q8CI11 | Guanine nu | Gnl3 | sp\|Q8CI11 | 27 | 27 | 56,1 |
| Q3UQA7 | Q3UQA7 | Selenoprot | Selh | sp\|Q3UQA | 4 | 4 | 30,2 |
| Q61112 | Q61112 | 45 kDa calc | Sdf4 | sp\|Q61112 | 10 | 10 | 41,8 |
| Q99LM2 | Q99LM2 | CDK5 regu | Cdk5rap3 | sp\|Q99LM | 17 | 17 | 38,8 |
| Q8CII2 | Q8CII2 | Cell divisio | Cdc123 | sp\|Q8CII2 | 9 | 9 | 31,5 |
| Q8BT14 | Q8BT14 | CCR4-NOT | Cnot4 | sp\|Q8BT14 | 8 | 8 | 20,5 |
| Q8CI32 | Q8CI32 | BAG family | Bag5 | sp\|Q8CI32 | 20 | 20 | 52,8 |
| Q9CYF5 | Q9CYF5 | Williams-B | Wbscr16 | sp\|Q9CYF5 | 7 | 7 | 26,2 |
| O88477 | O88477 | Insulin-like | Igf2bp1 | sp\|O88477 | 14 | 13 | 27,7 |
| Q8K1L2 | Q8K1L2 | Spindlin-4 | Spin4 | sp\|Q8K1L2 | 4 | 4 | 19,3 |
| Q3TFD2 | Q3TFD2 | Lysophosp | Lpcat1 | sp\|Q3TFD2 | 16 | 16 | 35 |
| P57722 | P57722 | Poly(rC)-bi | Pcbp3 | sp\|P57722 | 8 | 2 | 29,4 |
| Q6ZWX6 | Q6ZWX6 | Eukaryotic | Eif2s1 | sp\|Q6ZWX | 24 | 24 | 69,2 |
| E9Q5C9 | E9Q5C9 | | Nolc1 | sp\|E9Q5C9 | 15 | 15 | 20,5 |
| P06795;P2 | P06795 | Multidrug | Abcb1b | sp\|P06795 | 32 | 15 | 33,9 |
| Q9QZE5 | Q9QZE5 | Coatomer | Copg1 | sp\|Q9QZE5 | 41 | 37 | 69,5 |

| | | | | | | | |
|---|---|---|---|---|---|---|---|
| Q9Z1S0 | Q9Z1S0 | Mitotic che | Bub1b | sp\|Q9Z1S0 | 17 | 17 | 22,2 |
| Q8R317 | Q8R317 | Ubiquilin-1 | Ubqln1 | sp\|Q8R317 | 14 | 11 | 36,9 |
| Q9CQZ6 | Q9CQZ6 | NADH dehy | Ndufb3 | sp\|Q9CQZ | 2 | 2 | 17,3 |
| Q8VC16 | Q8VC16 | Leucine-ric | Lrrc14 | sp\|Q8VC16 | 9 | 9 | 21,9 |
| Q9JJU8 | Q9JJU8 | SH3 domai | Sh3bgrl | sp\|Q9JJU8 | 9 | 9 | 84,2 |
| Q8K0G5 | Q8K0G5 | Protein TSS | Tssc1 | sp\|Q8K0G | 10 | 10 | 38,9 |
| P55937 | P55937 | Golgin sub | Golga3 | sp\|P55937 | 43 | 43 | 39,8 |
| Q8CGN4 | Q8CGN4 | BCL-6 core | Bcor | sp\|Q8CGN | 5 | 5 | 5,3 |
| Q61655;Q9 | Q61655 | ATP-depen | Ddx19a | sp\|Q61655 | 20 | 20 | 53,8 |
| Q8BR63 | Q8BR63 | Protein FA | Fam177a1 | sp\|Q8BR63 | 6 | 6 | 37,2 |
| CON__Q0V | CON__Q0V8M9 | | | | 13 | 13 | 19,4 |
| Q3TN34 | Q3TN34 | MICAL-like | Micall2 | sp\|Q3TN34 | 33 | 33 | 47,8 |
| Q8BIG4 | Q8BIG4 | F-box only | Fbxo28 | sp\|Q8BIG4 | 6 | 6 | 22 |
| Q8BGT5 | Q8BGT5 | Alanine am | Gpt2 | sp\|Q8BGT | 9 | 9 | 28,2 |
| Q8K4F6 | Q8K4F6 | Probable 2 | Nsun5 | sp\|Q8K4F6 | 18 | 18 | 57,8 |
| Q8CHQ0 | Q8CHQ0 | F-box only | Fbxo4 | sp\|Q8CHQ | 11 | 11 | 35,3 |
| Q8K4P0 | Q8K4P0 | pre-mRNA | Wdr33 | sp\|Q8K4P0 | 22 | 22 | 22 |
| Q8BFP9 | Q8BFP9 | [Pyruvate d | Pdk1 | sp\|Q8BFP9 | 11 | 10 | 34,3 |
| Q6P9Q4 | Q6P9Q4 | FH1/FH2 d | Fhod1 | sp\|Q6P9Q | 37 | 37 | 48,9 |
| Q91X78 | Q91X78 | Erlin-1 | Erlin1 | sp\|Q91X78 | 11 | 9 | 40,8 |
| Q8BP40 | Q8BP40 | Lysophosp | Acp6 | sp\|Q8BP40 | 15 | 15 | 38,8 |
| O55126 | O55126 | Protein Nip | Gbas | sp\|O55126 | 6 | 6 | 27,4 |
| P10639 | P10639 | Thioredoxi | Txn | sp\|P10639 | 8 | 8 | 65,7 |
| O08715 | O08715 | A-kinase an | Akap1 | sp\|O08715 | 17 | 17 | 36,6 |
| Q9CRB3 | Q9CRB3 | 5-hydroxyi | Urah | sp\|Q9CRB | 3 | 3 | 45,8 |
| P63321 | P63321 | Ras-related | Rala | sp\|P63321 | 7 | 3 | 44,7 |
| CON__P00 | CON__P00978 | | | | 6 | 6 | 23 |
| Q9JHS9 | Q9JHS9 | Spliceosom | Cwc15 | sp\|Q9JHS9 | 7 | 7 | 28,4 |
| P50431 | P50431 | Serine hyd | Shmt1 | sp\|P50431 | 16 | 15 | 45,6 |
| Q9D7S7 | Q9D7S7 | 60S riboso | Rpl22l1 | sp\|Q9D7S7 | 2 | 2 | 19,7 |
| Q9WTX8 | Q9WTX8 | Mitotic spi | Mad1l1 | sp\|Q9WTX | 27 | 27 | 47,6 |
| Q8VEH5 | Q8VEH5 | EPM2A-int | Epm2aip1 | sp\|Q8VEH | 17 | 17 | 41,1 |
| Q9JJF3 | Q9JJF3 | Bifunctiona | No66 | sp\|Q9JJF3 | 13 | 13 | 31,8 |
| Q80UG5;R | Q80UG5 | Septin-9 | 09-set | sp\|Q80UG | 34 | 34 | 65,4 |
| A2A5Z6 | A2A5Z6 | E3 ubiquiti | Smurf2 | sp\|A2A5Z6 | 9 | 7 | 14 |
| Q3UB74 | Q3UB74 | Transformi | Tbrg1 | sp\|Q3UB74 | 4 | 4 | 16,7 |
| Q6QD59 | Q6QD59 | Vesicle tra | Bnip1 | sp\|Q6QD5 | 9 | 9 | 46,5 |
| O88379 | O88379 | Bromodom | Baz1a | sp\|O88379 | 52 | 52 | 36,9 |
| Q9CSH3 | Q9CSH3 | Exosome c | Dis3 | sp\|Q9CSH | 44 | 44 | 59,7 |
| P59438 | P59438 | Hermansky | Hps5 | sp\|P59438 | 16 | 16 | 22,1 |
| Q91WS7 | Q91WS7 | DEP domai | Depdc7 | sp\|Q91WS | 11 | 11 | 25,4 |
| P56395 | P56395 | Cytochrom | Cyb5a | sp\|P56395 | 4 | 4 | 41 |
| G5E8F4 | G5E8F4 | | Fpgt | sp\|G5E8F4 | 8 | 8 | 21,9 |
| P45376 | P45376 | Aldose red | Akr1b1 | sp\|P45376 | 19 | 19 | 61,1 |
| Q4FZC9 | Q4FZC9 | Nesprin-3 | Syne3 | sp\|Q4FZC9 | 42 | 42 | 51,4 |
| CON__ENS | CON__ENSEMBL:ENSBTAP0000024 | | ; | | 3 | 3 | 14 |
| Q9CXK9 | Q9CXK9 | RNA-bindin | Rbm33 | sp\|Q9CXK9 | 13 | 13 | 17,1 |
| Q8BI84 | Q8BI84 | Melanoma | Mia3 | sp\|Q8BI84 | 47 | 47 | 30,5 |
| O54890 | O54890 | Integrin be | Itgb3 | sp\|O54890 | 11 | 11 | 18,9 |
| Q9CZG9 | Q9CZG9 | PDZ domai | Pdzd11 | sp\|Q9CZG9 | 5 | 5 | 60 |

| | | | | | | | |
|---|---|---|---|---|---|---|---|
| Q8BUV3 | Q8BUV3 | Gephyrin;M | Gphn | sp\|Q8BUV | 18 | 18 | 33,9 |
| Q68FL4 | Q68FL4 | Putative ad | Ahcyl2 | sp\|Q68FL4 | 12 | 3 | 22,7 |
| P11928 | P11928 | 2-5-oligoad | Oas1a | sp\|P11928 | 3 | 3 | 13,1 |
| Q3TBT3 | Q3TBT3 | Stimulator | Tmem173 | sp\|Q3TBT3 | 11 | 11 | 42,6 |
| Q91WE1 | Q91WE1 | Sorting nex | Snx15 | sp\|Q91WE | 6 | 6 | 26,4 |
| P62484 | P62484 | Abl interac | Abi2 | sp\|P62484 | 9 | 5 | 24,4 |
| Q99NB9 | Q99NB9 | Splicing fac | Sf3b1 | sp\|Q99NB | 58 | 58 | 57,1 |
| E9Q7D5 | E9Q7D5 | | Arhgef5 | sp\|E9Q7D5 | 21 | 21 | 19,2 |
| Q9CQJ8 | Q9CQJ8 | NADH dehy | Ndufb9 | sp\|Q9CQJ8 | 9 | 9 | 65,4 |
| Q9D0K2 | Q9D0K2 | Succinyl-Co | Oxct1 | sp\|Q9D0K2 | 18 | 18 | 54,4 |
| Q01320 | Q01320 | DNA topois | Top2a | sp\|Q01320 | 84 | 68 | 52,9 |
| Q6PCP5 | Q6PCP5 | Mitochond | Mff | sp\|Q6PCP5 | 12 | 12 | 51,5 |
| Q8BZM1 | Q8BZM1 | Glomulin | Glmn | sp\|Q8BZM | 19 | 19 | 42,3 |
| CON__Q3M | CON__Q3MHN5 | | | | 14 | 2 | 39,7 |
| Q8BZ36 | Q8BZ36 | RAD50-inte | Rint1 | sp\|Q8BZ36 | 8 | 8 | 11,1 |
| Q62433 | Q62433 | Protein ND | Ndrg1 | sp\|Q62433 | 10 | 10 | 41,9 |
| Q8K0C9 | Q8K0C9 | GDP-mann | Gmds | sp\|Q8K0C9 | 12 | 12 | 44,9 |
| Q60872 | Q60872 | Eukaryotic | Eif1a | sp\|Q60872 | 7 | 1 | 59 |
| P48774 | P48774 | Glutathion | Gstm5 | sp\|P48774 | 12 | 11 | 52,7 |
| P98192 | P98192 | Dihydroxya | Gnpat | sp\|P98192 | 16 | 15 | 31,3 |
| Q5SYD0;Q5 | Q5SYD0 | Unconvent | Myo1d | sp\|Q5SYD0 | 21 | 21 | 26,5 |
| Q91ZP6 | Q91ZP6 | NEDD4 fam | Ndfip2 | sp\|Q91ZP6 | 2 | 2 | 7,7 |
| Q8R3B1 | Q8R3B1 | 1-phospha | Plcd1 | sp\|Q8R3B1 | 12 | 12 | 24,5 |
| Q91YL2 | Q91YL2 | E3 ubiquiti | Rnf126 | sp\|Q91YL2 | 4 | 4 | 15,3 |
| Q9D1P2 | Q9D1P2 | Histone ac | Kat8 | sp\|Q9D1P2 | 6 | 5 | 21,8 |
| P47226 | P47226 | Testin | Tes | sp\|P47226 | 16 | 16 | 42,6 |
| P97496 | P97496 | SWI/SNF co | Smarcc1 | sp\|P97496 | 40 | 31 | 43,8 |
| Q8BHC1 | Q8BHC1 | Ras-related | Rab39b | sp\|Q8BHC | 8 | 7 | 47,4 |
| Q99LW0 | Q99LW0 | Ankyrin re | Ankrd10 | sp\|Q99LW | 3 | 3 | 10,8 |
| Q6NZN0 | Q6NZN0 | RNA-bindin | Rbm26 | sp\|Q6NZN | 23 | 22 | 23,7 |
| Q91VW5 | Q91VW5 | Golgin sub | Golga4 | sp\|Q91VW | 54 | 54 | 31,9 |
| P47941 | P47941 | Crk-like pr | Crkl | sp\|P47941 | 15 | 15 | 62,7 |
| Q8VEJ9 | Q8VEJ9 | Vacuolar p | Vps4a | sp\|Q8VEJ9 | 11 | 8 | 33,2 |
| Q9D287 | Q9D287 | Pre-mRNA- | Bcas2 | sp\|Q9D287 | 14 | 14 | 76 |
| P48381 | P48381 | Transcripti | Rfx3 | sp\|P48381 | 9 | 5 | 20 |
| Q9CR62 | Q9CR62 | Mitochond | Slc25a11 | sp\|Q9CR62 | 14 | 14 | 53,5 |
| Q3KNM2 | Q3KNM2 | E3 ubiquiti | 05-mar | sp\|Q3KNM | 8 | 8 | 38,1 |
| Q9CQU0 | Q9CQU0 | Thioredoxi | Txndc12 | sp\|Q9CQU | 7 | 7 | 48,2 |
| Q571K4 | Q571K4 | TGF-beta-a | Tab3 | sp\|Q571K4 | 7 | 7 | 16,3 |
| Q9D938 | Q9D938 | Transmem | Tmem160 | sp\|Q9D938 | 3 | 3 | 16,5 |
| Q8BGB5 | Q8BGB5 | LIM domai | Limd2 | sp\|Q8BGB | 3 | 3 | 29,7 |
| Q8K2Z8 | Q8K2Z8 | Ubiquitin-c | Ube2q2 | sp\|Q8K2Z8 | 6 | 4 | 18,5 |
| Q9R0A0 | Q9R0A0 | Peroxisom | Pex14 | sp\|Q9R0A0 | 9 | 9 | 41,8 |
| Q9D903 | Q9D903 | Probable r | Ebna1bp2 | sp\|Q9D903 | 16 | 16 | 46,1 |
| Q8BGV0 | Q8BGV0 | Probable a | Nars2 | sp\|Q8BGV | 13 | 13 | 37,5 |
| Q7TSJ2 | Q7TSJ2 | Microtubu | Map6 | sp\|Q7TSJ2 | 14 | 14 | 21,3 |
| Q8BGQ7 | Q8BGQ7 | Alanine--tR | Aars | sp\|Q8BGQ | 45 | 45 | 62,1 |
| Q8BIH0 | Q8BIH0 | Histone de | Sap130 | sp\|Q8BIH0 | 9 | 9 | 13,4 |
| Q9CQC7 | Q9CQC7 | NADH dehy | Ndufb4 | sp\|Q9CQC | 6 | 6 | 46,5 |
| Q8BU03 | Q8BU03 | Periodic tr | Pwp2 | sp\|Q8BU0 | 21 | 21 | 35,4 |

| | | | | | | | |
|---|---|---|---|---|---|---|---|
| Q8R344 | Q8R344 | Coiled-coil | Ccdc12 | sp\|Q8R344 | 5 | 5 | 25,9 |
| L0N7N1 | L0N7N1 | Kinesin-like | Kif14 | sp\|L0N7N1 | 15 | 15 | 10,8 |
| Q9ESY9 | Q9ESY9 | Gamma-in | Ifi30 | sp\|Q9ESY9 | 5 | 5 | 28,2 |
| Q8BFQ8 | Q8BFQ8 | Parkinson | Pddc1 | sp\|Q8BFQ | 5 | 5 | 41,4 |
| P50136 | P50136 | 2-oxoisova | Bckdha | sp\|P50136 | 13 | 13 | 46,6 |
| Q7TMI3 | Q7TMI3 | E3 ubiquiti | Uhrf2 | sp\|Q7TMI | 15 | 14 | 23,4 |
| P61226 | P61226 | Ras-relatec | Rap2b | sp\|P61226 | 8 | 4 | 54,6 |
| Q6P9S7 | Q6P9S7 | Polypeptid | Galnt10 | sp\|Q6P9S7 | 3 | 3 | 8,6 |
| P15066 | P15066 | Transcripti | Jund | sp\|P15066 | 6 | 5 | 27 |
| P51829 | P51829 | Adenylate | Adcy7 | sp\|P51829 | 7 | 6 | 10,7 |
| Q9D0I4 | Q9D0I4 | Syntaxin-1 | Stx17 | sp\|Q9D0I4 | 7 | 7 | 41,5 |
| Q5SFM8 | Q5SFM8 | RNA-bindir | Rbm27 | sp\|Q5SFM | 20 | 19 | 24,8 |
| A7L9Z8 | A7L9Z8 | Calcium-tr | Atp2c2 | sp\|A7L9Z8 | 8 | 7 | 13 |
| Q3UPF5 | Q3UPF5 | Zinc finger | Zc3hav1 | sp\|Q3UPF | 26 | 26 | 39,1 |
| Q60932 | Q60932 | Voltage-de | Vdac1 | sp\|Q6093 | 21 | 21 | 79,1 |
| Q9Z0G9 | Q9Z0G9 | Claudin-3 | Cldn3 | sp\|Q9Z0G | 2 | 2 | 12,8 |
| Q6PDY0 | Q6PDY0 | Coiled-coil | Ccdc85b | sp\|Q6PDY | 3 | 3 | 23,3 |
| Q9WV30 | Q9WV30 | Nuclear fa | Nfat5 | sp\|Q9WV3 | 13 | 13 | 14,9 |
| P63044 | P63044 | Vesicle-ass | Vamp2 | sp\|P63044 | 3 | 1 | 34,5 |
| O08581 | O08581 | Potassium | Kcnk1 | sp\|O08581 | 2 | 2 | 11,6 |
| Q61263 | Q61263 | Sterol O-a | Soat1 | sp\|Q61263 | 11 | 11 | 23,3 |
| Q9JKF6 | Q9JKF6 | Nectin-1 | Pvrl1 | sp\|Q9JKF6 | 5 | 5 | 13,4 |
| Q9D0F3 | Q9D0F3 | Protein ER | Lman1 | sp\|Q9D0F | 18 | 18 | 43,1 |
| Q8C1B1 | Q8C1B1 | Calmodulir | Camsap2 | sp\|Q8C1B | 14 | 14 | 15,9 |
| P49312 | P49312 | Heterogen | Hnrnpa1 | sp\|P49312 | 24 | 22 | 73,8 |
| Q923Z3 | Q923Z3 | Protein M1 | Mto1 | sp\|Q923Z | 8 | 8 | 17,6 |
| Q9QXL2 | Q9QXL2 | Kinesin-like | Kif21a | sp\|Q9QXL | 5 | 4 | 3,9 |
| Q9ESD1 | Q9ESD1 | Prostasin;P | Prss8 | sp\|Q9ESD | 2 | 2 | 6,1 |
| Q8VCR3 | Q8VCR3 | Transmem | Tmem242 | sp\|Q8VCR | 3 | 3 | 29,3 |
| P47754 | P47754 | F-actin-cap | Capza2 | sp\|P47754 | 16 | 14 | 75,2 |
| O09164 | O09164 | Extracellul | Sod3 | sp\|O09164 | 4 | 4 | 25,9 |
| Q9D7V2 | Q9D7V2 | LysM and | Lysmd2 | sp\|Q9D7V | 4 | 4 | 20,5 |
| Q7TT18 | Q7TT18 | Activating | Atf7ip | sp\|Q7TT18 | 14 | 14 | 18,1 |
| Q9CZE3;Q9 | Q9CZE3 | Ras-relatec | Rab32 | sp\|Q9CZE | 8 | 8 | 48,4 |
| O70161 | O70161 | Phosphatid | Pip5k1c | sp\|O70161 | 13 | 11 | 26,3 |
| Q9ES00 | Q9ES00 | Ubiquitin c | Ube4b | sp\|Q9ES0 | 37 | 37 | 43,6 |
| Q80UW5 | Q80UW5 | Serine/thre | Cdc42bpg | sp\|Q80UW | 16 | 15 | 16,3 |
| Q9D2L9 | Q9D2L9 | Protein FA | Fam111a | sp\|Q9D2L | 8 | 8 | 15,7 |
| Q62311 | Q62311 | Transcripti | Taf6 | sp\|Q6231 | 17 | 17 | 32,6 |
| P62748;P8 | P62748 | Hippocalci | Hpcal1 | sp\|P62748 | 11 | 11 | 62,2 |
| Q9Z148 | Q9Z148 | Histone-lys | Ehmt2 | sp\|Q9Z148 | 25 | 25 | 28,6 |
| Q8VCY6 | Q8VCY6 | U3 small n | Utp6 | sp\|Q8VCY | 23 | 23 | 37,2 |
| P50637 | P50637 | Translocat | Tspo | sp\|P50637 | 2 | 2 | 8,3 |
| A6H8H2 | A6H8H2 | DENN dom | Dennd4c | sp\|A6H8H | 36 | 36 | 27,9 |
| P48972 | P48972 | Myb-relate | Mybl2 | sp\|P48972 | 7 | 7 | 15,6 |
| P59048 | P59048 | p53 and DI | Pdrg1 | sp\|P59048 | 3 | 3 | 30,8 |
| Q3UQ84 | Q3UQ84 | Threonine- | Tars2 | sp\|Q3UQ8 | 20 | 20 | 38,9 |
| Q9ER69 | Q9ER69 | Pre-mRNA- | Wtap | sp\|Q9ER6 | 15 | 15 | 48,5 |
| Q62077 | Q62077 | 1-phospha | Plcg1 | sp\|Q6207 | 25 | 25 | 27,4 |
| P35441 | P35441 | Thrombosp | Thbs1 | sp\|P35441 | 15 | 10 | 17,7 |

| | | | | | | | |
|---|---|---|---|---|---|---|---|
| Q8BTM8 | Q8BTM8 | Filamin-A | Flna | sp\|Q8BTM | 137 | 127 | 67,7 |
| Q6P6L0 | Q6P6L0 | Filamin A-i | Filip1l | sp\|Q6P6L0 | 15 | 14 | 16,9 |
| Q8R0X7 | Q8R0X7 | Sphingosin | Sgpl1 | sp\|Q8R0X7 | 16 | 16 | 34,2 |
| Q5Y5T1 | Q5Y5T1 | Probable p | Zdhhc20 | sp\|Q5Y5T1 | 5 | 5 | 20,3 |
| Q8C827 | Q8C827 | Zinc finger | Zfp62 | sp\|Q8C827 | 6 | 6 | 8,8 |
| Q9Z2G9 | Q9Z2G9 | Oxidoredu | Htatip2 | sp\|Q9Z2G9 | 5 | 5 | 27,3 |
| P49025 | P49025 | Citron Rho | Cit | sp\|P49025 | 40 | 40 | 25,1 |
| Q3UKC1 | Q3UKC1 | Tax1-bindi | Tax1bp1 | sp\|Q3UKC | 20 | 20 | 29,9 |
| Q8C547 | Q8C547 | HEAT repe | Heatr5b | sp\|Q8C547 | 29 | 29 | 23 |
| Q8BWG8 | Q8BWG8 | Beta-arres | Arrb1 | sp\|Q8BWG | 5 | 5 | 20,6 |
| Q61462 | Q61462 | Cytochrom | Cyba | sp\|Q61462 | 2 | 2 | 18,2 |
| Q56A08 | Q56A08 | G patch do | Gpkow | sp\|Q56A08 | 8 | 8 | 28,3 |
| Q99LZ3 | Q99LZ3 | DNA replic | Gins4 | sp\|Q99LZ3 | 8 | 8 | 44,8 |
| Q9DB25 | Q9DB25 | Dolichyl-pl | Alg5 | sp\|Q9DB2 | 10 | 10 | 35,2 |
| P07901 | P07901 | Heat shock | Hsp90aa1 | sp\|P07901 | 52 | 35 | 66,2 |
| Q9JLM8;O | Q9JLM8 | Serine/thre | Dclk1 | sp\|Q9JLM8 | 17 | 17 | 27,4 |
| Q7TPR4 | Q7TPR4 | Alpha-actin | Actn1 | sp\|Q7TPR4 | 59 | 35 | 71 |
| Q8BJU9 | Q8BJU9 | Peptide ch | Mtrf1l | sp\|Q8BJU9 | 3 | 3 | 12,6 |
| Q9D517 | Q9D517 | 1-acyl-sn-g | Agpat3 | sp\|Q9D517 | 10 | 10 | 22,3 |
| Q9JHQ5 | Q9JHQ5 | Leucine zip | Lztfl1 | sp\|Q9JHQ5 | 10 | 10 | 43,5 |
| Q9DBR7 | Q9DBR7 | Protein ph | Ppp1r12a | sp\|Q9DBR | 30 | 28 | 37 |
| Q922M3 | Q922M3 | BTB/POZ d | Kctd10 | sp\|Q922M | 7 | 6 | 25,4 |
| Q9Z2D3 | Q9Z2D3 | Non-syndr | Dfna5 | sp\|Q9Z2D3 | 15 | 15 | 37,9 |
| P70349 | P70349 | Histidine tr | Hint1 | sp\|P70349 | 8 | 8 | 78,6 |
| Q8BWW4 | Q8BWW4 | La-related | Larp4 | sp\|Q8BWW | 18 | 18 | 32,3 |
| Q9DC48 | Q9DC48 | Pre-mRNA- | Cdc40 | sp\|Q9DC48 | 25 | 25 | 53,4 |
| Q8BGF7 | Q8BGF7 | PAB-deper | Pan2 | sp\|Q8BGF7 | 8 | 8 | 8,5 |
| Q91YE5 | Q91YE5 | Bromodom | Baz2a | sp\|Q91YE5 | 21 | 21 | 16,1 |
| Q62101 | Q62101 | Serine/thre | Prkd1 | sp\|Q62101 | 10 | 6 | 18,8 |
| Q9DCS3 | Q9DCS3 | Trans-2-en | Mecr | sp\|Q9DCS3 | 10 | 10 | 41 |
| Q91W90 | Q91W90 | Thioredoxi | Txndc5 | sp\|Q91W9 | 18 | 18 | 52 |
| Q8CDA1 | Q8CDA1 | Phosphatid | Inpp5f | sp\|Q8CDA | 8 | 8 | 10,4 |
| Q9CR58;Q | Q9CR58 | Kidney mit | Slc25a30 | sp\|Q9CR58 | 6 | 6 | 29,9 |
| Q9D1I2 | Q9D1I2 | Bcl10-interacting CAR | | sp\|Q9D1I2 | 4 | 4 | 23 |
| Q8R1F9 | Q8R1F9 | Ribonuclea | Rpp40 | sp\|Q8R1F9 | 4 | 4 | 12,9 |
| Q3TLD5 | Q3TLD5 | Unconvent | Uri1 | sp\|Q3TLD5 | 12 | 12 | 39,5 |
| O08997 | O08997 | Copper tra | Atox1 | sp\|O08997 | 7 | 7 | 88,2 |
| Q3UBG2 | Q3UBG2 | PTB-contai | Pid1 | sp\|Q3UBG | 4 | 4 | 30 |
| A2AM29 | A2AM29 | Protein AF | Mllt3 | sp\|A2AM2 | 7 | 7 | 14,1 |
| Q9DCT8 | Q9DCT8 | Cysteine-ri | Crip2 | sp\|Q9DCT8 | 8 | 8 | 45,7 |
| P23772;O | P23772 | Trans-actin | Gata3 | sp\|P23772 | 6 | 6 | 17,2 |
| Q9DBE0 | Q9DBE0 | Cysteine su | Csad | sp\|Q9DBE0 | 3 | 3 | 7,3 |
| P07607 | P07607 | Thymidylat | Tyms | sp\|P07607 | 12 | 12 | 42,3 |
| Q61335 | Q61335 | B-cell rece | Bcap31 | sp\|Q61335 | 11 | 11 | 39,2 |
| Q9Z0H4 | Q9Z0H4 | CUGBP Ela | Celf2 | sp\|Q9Z0H4 | 9 | 8 | 20,7 |
| Q8R035 | Q8R035 | Peptidyl-tR | Ict1 | sp\|Q8R035 | 7 | 7 | 32,5 |
| Q9CWH5 | Q9CWH5 | tRNA (guar | Trmt11 | sp\|Q9CWH | 8 | 8 | 18,9 |
| Q8VEC4 | Q8VEC4 | Calcium ho | Calhm2 | sp\|Q8VEC4 | 4 | 4 | 15,5 |
| P97315 | P97315 | Cysteine ar | Csrp1 | sp\|P97315 | 9 | 9 | 59,6 |
| Q8BLN5 | Q8BLN5 | Lanosterol | Lss | sp\|Q8BLN5 | 19 | 19 | 32,6 |

| | | | | | | | |
|---|---|---|---|---|---|---|---|
| Q9JJV2 | Q9JJV2 | Profilin-2 | Pfn2 | sp\|Q9JJV2 | 3 | 3 | 30,7 |
| Q8CH02 | Q8CH02 | SURP and ( | Sugp1 | sp\|Q8CH0 | 20 | 20 | 47,7 |
| Q3UGS4 | Q3UGS4 | Protein FA | Fam195b | sp\|Q3UGS | 4 | 4 | 50,5 |
| Q62318 | Q62318 | Transcripti | Trim28 | sp\|Q62318 | 34 | 34 | 65,7 |
| Q8VD04 | Q8VD04 | GRIP1-asso | Gripap1 | sp\|Q8VD0 | 7 | 7 | 13,8 |
| Q03958 | Q03958 | Prefoldin s | Pfdn6 | sp\|Q03958 | 10 | 10 | 46,5 |
| Q8CIG3 | Q8CIG3 | Lysine-spe | Kdm1b | sp\|Q8CIG3 | 11 | 11 | 22,5 |
| P26041 | P26041 | Moesin | Msn | sp\|P26041 | 51 | 39 | 74,5 |
| Q9Z2Y8 | Q9Z2Y8 | Proline syn | Prosc | sp\|Q9Z2Y8 | 11 | 11 | 54,7 |
| E9Q7F2 | E9Q7F2 | E3 ubiquiti | Rnf169 | sp\|E9Q7F2 | 7 | 7 | 16,1 |
| Q9DAT2 | Q9DAT2 | MRG/MOR | Mrgbp | sp\|Q9DAT | 1 | 1 | 6,4 |
| Q5SWU9 | Q5SWU9 | Acetyl-CoA | Acaca | sp\|Q5SWU | 70 | 60 | 40 |
| Q69ZF7 | Q69ZF7 | Metal tran | Cnnm4 | sp\|Q69ZF7 | 12 | 10 | 20,4 |
| Q3UIR3 | Q3UIR3 | E3 ubiquiti | Dtx3l | sp\|Q3UIR3 | 6 | 6 | 10,6 |
| Q8VD26 | Q8VD26 | Transmem | Tmem143 | sp\|Q8VD2 | 7 | 7 | 22,5 |
| Q9JMH9 | Q9JMH9 | Unconvent | Myo18a | sp\|Q9JMH | 49 | 49 | 32,8 |
| Q9CXF4 | Q9CXF4 | TBC1 doma | Tbc1d15 | sp\|Q9CXF4 | 25 | 25 | 47,5 |
| P49935 | P49935 | Pro-cathep | Ctsh | sp\|P49935 | 7 | 7 | 33 |
| Q9EQ20 | Q9EQ20 | Methylmal | Aldh6a1 | sp\|Q9EQ2 | 15 | 15 | 46 |
| Q91W98 | Q91W98 | Solute carr | Slc15a4 | sp\|Q91W9 | 5 | 5 | 9,8 |
| Q8K2I4 | Q8K2I4 | Beta-mann | Manba | sp\|Q8K2I4 | 11 | 11 | 16 |
| Q8K224 | Q8K224 | N-acetyltra | Nat10 | sp\|Q8K224 | 41 | 41 | 47,9 |
| Q9EQG9 | Q9EQG9 | Collagen ty | Col4a3bp | sp\|Q9EQG | 20 | 20 | 44,9 |
| Q8R5A0 | Q8R5A0 | N-lysine m | Smyd2 | sp\|Q8R5A | 7 | 7 | 21,7 |
| O54984 | O54984 | ATPase Asr | Asna1 | sp\|O54984 | 12 | 12 | 44,3 |
| O08585 | O08585 | Clathrin lig | Clta | sp\|O08585 | 7 | 7 | 21,3 |
| P63028 | P63028 | Translation | Tpt1 | sp\|P63028 | 9 | 9 | 41,3 |
| Q3UNA4 | Q3UNA4 | NTF2-relat | Nxt2 | sp\|Q3UNA | 2 | 2 | 15,5 |
| Q99JY9;Q6 | Q99JY9 | Actin-relat | Actr3 | sp\|Q99JY9 | 20 | 20 | 62 |
| Q69ZP3 | Q69ZP3 | Probable h | Pnkd | sp\|Q69ZP3 | 1 | 1 | 3,9 |
| P57080 | P57080 | Ubiquitin c | Usp25 | sp\|P57080 | 15 | 15 | 19,1 |
| Q924D0 | Q924D0 | Reticulon-4 | Rtn4ip1 | sp\|Q924D | 6 | 6 | 21 |
| Q8K124 | Q8K124 | Pleckstrin I | Plekho2 | sp\|Q8K124 | 7 | 7 | 27,5 |
| Q6P8M1 | Q6P8M1 | Putative de | Tatdn1 | sp\|Q6P8M | 10 | 10 | 47,8 |
| Q9D1I5 | Q9D1I5 | Methylmal | Mcee | sp\|Q9D1I5 | 2 | 2 | 18 |
| P63254 | P63254 | Cysteine-ri | Crip1 | sp\|P63254 | 4 | 4 | 44,2 |
| Q8CJH3 | Q8CJH3 | Plexin-B1 | Plxnb1 | sp\|Q8CJH3 | 12 | 12 | 8,8 |
| P56135 | P56135 | ATP syntha | Atp5j2 | sp\|P56135 | 4 | 4 | 36,4 |
| Q3UL36 | Q3UL36 | Arginine ar | Arglu1 | sp\|Q3UL3 | 3 | 3 | 8,9 |
| Q8VDC1 | Q8VDC1 | FYVE and c | Fyco1 | sp\|Q8VDC | 26 | 26 | 25,7 |
| Q9D7G0 | Q9D7G0 | Ribose-pho | Prps1 | sp\|Q9D7G | 14 | 8 | 50,3 |
| O88845 | O88845 | A-kinase ar | Akap10 | sp\|O88845 | 10 | 10 | 20,4 |
| Q9EQQ0 | Q9EQQ0 | Histone-lys | Suv39h2 | sp\|Q9EQQ | 6 | 6 | 17,6 |
| Q8BGB7 | Q8BGB7 | Enolase-ph | Enoph1 | sp\|Q8BGB | 5 | 5 | 28 |
| Q8K1N2 | Q8K1N2 | Pleckstrin I | Phldb2 | sp\|Q8K1N | 14 | 14 | 16,2 |
| Q5DTW7 | Q5DTW7 | Uncharacte | Kiaa1551 | sp\|Q5DTW | 20 | 20 | 18,8 |
| Q8CIW5 | Q8CIW5 | Twinkle pr | Peo1 | sp\|Q8CIW | 13 | 13 | 29,5 |
| Q9QXV8 | Q9QXV8 | Protein spr | Spry2 | sp\|Q9QXV | 4 | 4 | 21,3 |
| Q8CFE4 | Q8CFE4 | SCY1-like p | Scyl2 | sp\|Q8CFE4 | 19 | 19 | 27,7 |
| Q9D0K1 | Q9D0K1 | Peroxisom | Pex13 | sp\|Q9D0K | 9 | 9 | 36,8 |

| | | | | | | | |
|---|---|---|---|---|---|---|---|
| Q8BGA9 | Q8BGA9 | Mitochond | Oxa1l | sp\|Q8BGA | 5 | 5 | 12,9 |
| Q62465 | Q62465 | Synaptic ve | Vat1 | sp\|Q62465 | 19 | 19 | 50,5 |
| Q64337 | Q64337 | Sequestoso | Sqstm1 | sp\|Q64337 | 15 | 15 | 60,6 |
| A2A6Q5 | A2A6Q5 | Cell divisio | Cdc27 | sp\|A2A6Q | 19 | 19 | 37,5 |
| Q99KF1 | Q99KF1 | Transmem | Tmed9 | sp\|Q99KF1 | 11 | 10 | 32,8 |
| Q99J83 | Q99J83 | Autophagy | Atg5 | sp\|Q99J83 | 8 | 8 | 33,1 |
| Q9CRA9 | Q9CRA9 | FGFR1 onc | Fgfr1op2 | sp\|Q9CRA | 8 | 8 | 46,2 |
| P97864 | P97864 | Caspase-7; | Casp7 | sp\|P97864 | 4 | 4 | 16,8 |
| Q61024 | Q61024 | Asparagine | Asns | sp\|Q61024 | 26 | 26 | 59 |
| Q8R570 | Q8R570 | Synaptoso | Snap47 | sp\|Q8R570 | 16 | 16 | 50,8 |
| Q9QUJ7 | Q9QUJ7 | Long-chain | Acsl4 | sp\|Q9QUJ7 | 21 | 21 | 41,9 |
| P70347 | P70347 | TRAF famil | Tank | sp\|P70347 | 5 | 5 | 19,9 |
| Q9D0M1 | Q9D0M1 | Phosphoril | Prpsap1 | sp\|Q9D0M | 13 | 11 | 48,3 |
| CON__ENS | CON__ENSEM | Hepatocyte | Hgfac | ;sp\|Q9R09 | 3 | 3 | 5,4 |
| B1AZP2 | B1AZP2 | Disks large | Dlgap4 | sp\|B1AZP2 | 8 | 8 | 11,4 |
| Q6R0H7;P6 | Q6R0H7;P6309 | Guanine nu | Gnas | sp\|Q6R0H7 | 18 | 17 | 20,2 |
| Q91X58 | Q91X58 | AN1-type z | Zfand2b | sp\|Q91X58 | 3 | 3 | 15,6 |
| Q9Z0L8 | Q9Z0L8 | Gamma-glu | Ggh | sp\|Q9Z0L8 | 7 | 7 | 28,4 |
| Q6L8S8 | Q6L8S8 | Beta-1,4-N | B4galnt3 | sp\|Q6L8S8 | 5 | 5 | 6 |
| Q9CZC8 | Q9CZC8 | Secernin-1 | Scrn1 | sp\|Q9CZC8 | 7 | 7 | 23,9 |
| Q8R0A0 | Q8R0A0 | General tra | Gtf2f2 | sp\|Q8R0A0 | 14 | 14 | 63,9 |
| Q9D0D5 | Q9D0D5 | General tra | Gtf2e1 | sp\|Q9D0D | 8 | 8 | 22,3 |
| Q9JHW4 | Q9JHW4 | Selenocyst | Eefsec | sp\|Q9JHW | 23 | 23 | 55,1 |
| Q8CHH9 | Q8CHH9 | Septin-8 | 08-set | sp\|Q8CHH | 22 | 19 | 57,3 |
| Q8K1M6 | Q8K1M6 | Dynamin-1 | Dnm1l | sp\|Q8K1M | 35 | 35 | 67,3 |
| Q7TMK6 | Q7TMK6 | Protein Ho | Hook2 | sp\|Q7TMK | 9 | 9 | 12,8 |
| Q61937 | Q61937 | Nucleopho | Npm1 | sp\|Q61937 | 22 | 22 | 77,7 |
| Q00560 | Q00560 | Interleukin | Il6st | sp\|Q00560 | 11 | 11 | 20,8 |
| P83887;Q8 | P83887;Q8VCK | Tubulin ga | Tubg1;Tub | sp\|P83887 | 18 | 18 | 62,3 |
| P45952 | P45952 | Medium-ch | Acadm | sp\|P45952 | 14 | 14 | 48,5 |
| Q3TEW6 | Q3TEW6 | Myelin pro | Mpzl1 | sp\|Q3TEW | 2 | 2 | 8,5 |
| Q9QYB1 | Q9QYB1 | Chloride in | Clic4 | sp\|Q9QYB | 17 | 17 | 77,5 |
| P54761 | P54761 | Ephrin type | Ephb4 | sp\|P54761 | 18 | 13 | 29,3 |
| Q8BZ03 | Q8BZ03 | Serine/thre | Prkd2 | sp\|Q8BZ03 | 9 | 7 | 17,3 |
| Q3U1V6 | Q3U1V6 | Ubiquitin-c | Uevld | sp\|Q3U1V | 16 | 16 | 41,4 |
| P84091 | P84091 | AP-2 comp | Ap2m1 | sp\|P84091 | 18 | 18 | 48,3 |
| Q8VE18 | Q8VE18 | Protein SM | Smg8 | sp\|Q8VE18 | 14 | 14 | 22,2 |
| O88207 | O88207 | Collagen al | Col5a1 | sp\|O88207 | 11 | 9 | 11 |
| Q9WTX2 | Q9WTX2 | Interferon- | Prkra | sp\|Q9WTX | 6 | 6 | 26,8 |
| Q3TWI9 | Q3TWI9 | CSC1-like p | Tmem63b | sp\|Q3TWI9 | 4 | 4 | 4,8 |
| Q6PAR5 | Q6PAR5 | GTPase-act | Gapvd1 | sp\|Q6PAR5 | 37 | 37 | 31,3 |
| Q7TQH0 | Q7TQH0 | Ataxin-2-lil | Atxn2l | sp\|Q7TQH | 35 | 35 | 41,4 |
| Q9QUN7 | Q9QUN7 | Toll-like re | Tlr2 | sp\|Q9QUN | 6 | 6 | 9,3 |
| Q8BUR4 | Q8BUR4 | Dedicator c | Dock1 | sp\|Q8BUR | 29 | 28 | 20,8 |
| Q571I4 | Q571I4 | Tyrosine-p | Sgk223 | sp\|Q571I4 | 4 | 4 | 3,3 |
| Q8VEK3 | Q8VEK3 | Heterogen | Hnrnpu | sp\|Q8VEK3 | 43 | 43 | 53,6 |
| A2ASA8 | A2ASA8 | Inositol 1,4 | Itpripl1 | sp\|A2ASA8 | 5 | 5 | 12,4 |
| P97377;Q8 | P97377 | Cyclin-dep | Cdk2 | sp\|P97377 | 17 | 15 | 54,6 |
| Q6A026 | Q6A026 | Sister chro | Pds5a | sp\|Q6A026 | 56 | 51 | 47,1 |
| P17439 | P17439 | Glucosylce | Gba | sp\|P17439 | 15 | 15 | 34,8 |

| | | | | | | | |
|---|---|---|---|---|---|---|---|
| Q8CCJ3 | Q8CCJ3 | E3 UFM1-p | Ufl1 | sp\|Q8CCJ3 | 31 | 31 | 46,7 |
| Q9R0Q7 | Q9R0Q7 | Prostaglan | Ptges3 | sp\|Q9R0Q | 9 | 9 | 60 |
| Q8VEL0 | Q8VEL0 | Motile spe | Mospd1 | sp\|Q8VEL( | 4 | 4 | 30 |
| Q5SVR0;Q | Q5SVR0 | TBC1 doma | Tbc1d9b | sp\|Q5SVR( | 11 | 11 | 12,1 |
| P70460 | P70460 | Vasodilato | Vasp | sp\|P70460 | 16 | 16 | 51,2 |
| O35682 | O35682 | Myeloid-as | Myadm | sp\|O35682 | 6 | 6 | 20,9 |
| P51660 | P51660 | Peroxisom | Hsd17b4 | sp\|P51660 | 26 | 26 | 48,7 |
| Q8JZR6;Q5 | Q8JZR6 | Electroneu | Slc4a8 | sp\|Q8JZR6 | 8 | 7 | 10,5 |
| E9Q5G3 | E9Q5G3 | Kinesin-like | Kif23 | sp\|E9Q5G3 | 30 | 30 | 42,5 |
| O35704 | O35704 | Serine palr | Sptlc1 | sp\|O35704 | 9 | 9 | 28,3 |
| Q9Z0G0;Q | Q9Z0G0 | PDZ domai | Gipc1 | sp\|Q9Z0G( | 10 | 10 | 44,4 |
| Q8R313 | Q8R313 | Exocyst co | Exoc6 | sp\|Q8R313 | 9 | 8 | 12,8 |
| P58501 | P58501 | PAX3- and | Paxbp1 | sp\|P58501 | 18 | 18 | 27,9 |
| Q61656 | Q61656 | Probable A | Ddx5 | sp\|Q61656 | 40 | 33 | 57,3 |
| P20065 | P20065 | Thymosin | Tmsb4x | sp\|P20065 | 4 | 3 | 38 |
| Q9CXC3 | Q9CXC3 | Mitochond | Mgme1 | sp\|Q9CXC3 | 3 | 3 | 13,9 |
| Q8C9B9 | Q8C9B9 | Death-indu | Dido1 | sp\|Q8C9B9 | 47 | 47 | 29,7 |
| Q9D7P6 | Q9D7P6 | Iron-sulfur | Iscu | sp\|Q9D7P( | 5 | 5 | 38,7 |
| O08901 | O08901 | Mitotic che | Bub1 | sp\|O08901 | 5 | 5 | 8,3 |
| O08688 | O08688 | Calpain-5 | Capn5 | sp\|O08688 | 3 | 3 | 7,3 |
| Q9DCF9 | Q9DCF9 | Translocon | Ssr3 | sp\|Q9DCF9 | 1 | 1 | 7,6 |
| Q6DFV3 | Q6DFV3 | Rho GTPas | Arhgap21 | sp\|Q6DFV3 | 17 | 17 | 16,1 |
| Q91WM3 | Q91WM3 | U3 small n | Rrp9 | sp\|Q91WM | 12 | 12 | 33,7 |
| Q5NBX1 | Q5NBX1 | Protein co | Cobl | sp\|Q5NBX | 12 | 12 | 14,4 |
| P12388 | P12388 | Plasminoge | Serpinb2 | sp\|P12388 | 12 | 12 | 43,4 |
| Q62419;Q( | Q62419 | Endophilin | Sh3gl1 | sp\|Q62419 | 16 | 16 | 54,9 |
| Q924K8 | Q924K8 | Metastasis | Mta3 | sp\|Q924K8 | 10 | 8 | 23,2 |
| Q9CQC8 | Q9CQC8 | Maspardin | Spg21 | sp\|Q9CQC | 11 | 11 | 41,6 |
| Q99LE6 | Q99LE6 | ATP-bindin | Abcf2 | sp\|Q99LE6 | 26 | 26 | 43,3 |
| B1AR13 | B1AR13 | CDGSH iro | Cisd3 | sp\|B1AR13 | 1 | 1 | 9,5 |
| Q640M6 | Q640M6 | Glyceroph | Gdpd5 | sp\|Q640M | 3 | 3 | 5,6 |
| Q9Z2W0 | Q9Z2W0 | Aspartyl ar | Dnpep | sp\|Q9Z2W | 15 | 15 | 45 |
| Q9DCT6 | Q9DCT6 | Chromatin | Bap18 | sp\|Q9DCT( | 4 | 4 | 36,3 |
| O54734 | O54734 | Dolichyl-di | Ddost | sp\|O54734 | 15 | 15 | 50,3 |
| Q8BKR5 | Q8BKR5 | Protein ph | Ppp1r37 | sp\|Q8BKR5 | 6 | 6 | 11,8 |
| Q9D8L5 | Q9D8L5 | Coiled-coil | Ccdc91 | sp\|Q9D8L5 | 6 | 6 | 15,4 |
| P16092 | P16092 | Fibroblast | Fgfr1 | sp\|P16092 | 3 | 3 | 7,3 |
| Q8K1H1 | Q8K1H1 | Tudor dom | Tdrd7 | sp\|Q8K1H1 | 26 | 26 | 36,3 |
| Q6P3D0 | Q6P3D0 | U8 snoRNA | Nudt16 | sp\|Q6P3D( | 3 | 2 | 15,9 |
| Q3TCH7 | Q3TCH7 | Cullin-4A | Cul4a | sp\|Q3TCH7 | 35 | 26 | 47,6 |
| Q8BLH7 | Q8BLH7 | HIRA-inter | Hirip3 | sp\|Q8BLH7 | 10 | 10 | 16,8 |
| P50295;P5 | P50295 | Arylamine | Nat2 | sp\|P50295 | 11 | 11 | 51,7 |
| E9PZJ8 | E9PZJ8 | Activating | Ascc3 | sp\|E9PZJ8 | 40 | 40 | 26,5 |
| Q05186 | Q05186 | Reticuloca | Rcn1 | sp\|Q05186 | 14 | 14 | 55,4 |
| P43136 | P43136 | Nuclear re | Nr2f6 | sp\|P43136 | 7 | 6 | 25,9 |
| Q6PGH2 | Q6PGH2 | Hematolog | Hn1l | sp\|Q6PGH | 6 | 6 | 48,4 |
| Q8C0T5 | Q8C0T5 | Signal-indu | Sipa1l1 | sp\|Q8C0T5 | 41 | 41 | 36,8 |
| Q8CI43 | Q8CI43 | Myosin ligh | Myl6b | sp\|Q8CI43 | 6 | 5 | 36,2 |
| P11276 | P11276 | Fibronectin | Fn1 | sp\|P11276 | 61 | 61 | 37,9 |
| Q99LQ7 | Q99LQ7 | Transmem | Tmem189 | sp\|Q99LQ7 | 2 | 2 | 9,2 |

| | | | | | | | |
|---|---|---|---|---|---|---|---|
| P61600 | P61600 | N-alpha-ac | Naa20 | sp\|P61600 | 6 | 6 | 47,2 |
| Q8BHB4 | Q8BHB4 | WD repeat | Wdr3 | sp\|Q8BHB | 40 | 40 | 57,1 |
| Q6ZQH8 | Q6ZQH8 | Nucleopor | Nup188 | sp\|Q6ZQH | 39 | 39 | 28 |
| Q8BGX2 | Q8BGX2 | Uncharacterized prot | | sp\|Q8BGX | 6 | 6 | 33,1 |
| Q8VDK1 | Q8VDK1 | Nitrilase h | Nit1 | sp\|Q8VDK | 10 | 10 | 44 |
| Q2NL51 | Q2NL51 | Glycogen s | Gsk3a | sp\|Q2NL51 | 13 | 11 | 35,3 |
| Q8R2Y8 | Q8R2Y8 | Peptidyl-tR | Ptrh2 | sp\|Q8R2Y8 | 8 | 8 | 59,1 |
| E9PYL2 | E9PYL2 | | Prr12 | sp\|E9PYL2 | 15 | 15 | 14,6 |
| CON__ENS | CON__ENSEMBL:ENSBTAP00000037665 | | | | 8 | 8 | 17,8 |
| Q9CQ85 | Q9CQ85 | Mitochond | Timm22 | sp\|Q9CQ8 | 3 | 3 | 31,4 |
| O88466 | O88466 | Zinc finger | Znf106 | sp\|O88466 | 12 | 12 | 10,8 |
| P14873 | P14873 | Microtubu | Map1b | sp\|P14873 | 77 | 74 | 46,5 |
| O35613 | O35613 | Death dom | Daxx | sp\|O35613 | 6 | 6 | 15,2 |
| Q6P1H6 | Q6P1H6 | Ankyrin re | Ankle2 | sp\|Q6P1H | 35 | 35 | 47,9 |
| Q0P5W1 | Q0P5W1 | Vacuolar p | Vps8 | sp\|Q0P5W | 8 | 8 | 6,9 |
| O08582 | O08582 | GTP-bindir | Gtpbp1 | sp\|O0858 | 18 | 18 | 43,4 |
| E9Q3L2 | E9Q3L2 | | Pi4ka | sp\|E9Q3L2 | 25 | 25 | 15,8 |
| P09922;Q9 | P09922;Q9WV | Interferon- | Mx1;Mx2 | sp\|P09922 | 2 | 2 | 4,6 |
| Q8R060 | Q8R060 | Protein zw | Zwilch | sp\|Q8R06 | 13 | 13 | 30,1 |
| Q8BWQ6 | Q8BWQ6 | UPF0505 protein C16 | | sp\|Q8BW | 19 | 19 | 24,2 |
| Q9WVE8 | Q9WVE8 | Protein kin | Pacsin2 | sp\|Q9WVE | 18 | 18 | 42,2 |
| Q9JHP7 | Q9JHP7 | KDEL moti | Kdelc1 | sp\|Q9JHP7 | 14 | 14 | 39,8 |
| P12032 | P12032 | Metallopro | Timp1 | sp\|P12032 | 4 | 4 | 33,2 |
| Q8VEH8 | Q8VEH8 | Endoplasm | Erlec1 | sp\|Q8VEH | 10 | 10 | 26,5 |
| Q5SUF2 | Q5SUF2 | Luc7-like p | Luc7l3 | sp\|Q5SUF2 | 12 | 12 | 31 |
| P07356 | P07356 | Annexin A | Anxa2 | sp\|P07356 | 36 | 36 | 81,4 |
| Q9JJX6 | Q9JJX6 | P2X purino | P2rx4 | sp\|Q9JJX6 | 4 | 4 | 11,6 |
| C0HKD9;C | C0HKD9;C0HKD8 | | | sp\|C0HKD | 8 | 8 | 18,7 |
| Q9ERI6 | Q9ERI6 | Retinol del | Rdh14 | sp\|Q9ERI6 | 8 | 8 | 38,6 |
| B7ZNG0 | B7ZNG0 | Kinesin-like | Kif7 | sp\|B7ZNG | 5 | 4 | 4,1 |
| P70236 | P70236 | Dual specif | Map2k6 | sp\|P70236 | 6 | 5 | 24,3 |
| Q8C3Y4 | Q8C3Y4 | Kinetochor | Kntc1 | sp\|Q8C3Y | 32 | 32 | 17,7 |
| Q9Z2A0 | Q9Z2A0 | 3-phospho | Pdpk1 | sp\|Q9Z2A | 14 | 14 | 35,6 |
| Q8R1Q8 | Q8R1Q8 | Cytoplasm | Dync1li1 | sp\|Q8R1Q | 22 | 21 | 55,6 |
| Q9JK92 | Q9JK92 | Heat shock | Hspb8 | sp\|Q9JK92 | 6 | 6 | 43,4 |
| O35874 | O35874 | Neutral am | Slc1a4 | sp\|O35874 | 7 | 7 | 20,1 |
| Q8BG81 | Q8BG81 | Polymeras | Poldip3 | sp\|Q8BG8 | 18 | 18 | 52,9 |
| Q8BTJ4 | Q8BTJ4 | Bis(5-aden | Enpp4 | sp\|Q8BTJ4 | 4 | 4 | 9,6 |
| Q8CG76 | Q8CG76 | Aflatoxin B | Akr7a2 | sp\|Q8CG7 | 8 | 8 | 30,5 |
| Q8CB77 | Q8CB77 | Transcripti | Tceb3 | sp\|Q8CB7 | 12 | 12 | 20,7 |
| Q6PD10 | Q6PD10 | Inositol he | Ip6k1 | sp\|Q6PD1 | 3 | 3 | 7,4 |
| Q3V1T4 | Q3V1T4 | Prolyl 3-hy | Lepre1 | sp\|Q3V1T | 8 | 8 | 12,2 |
| P53810 | P53810 | Phosphatic | Pitpna | sp\|P53810 | 16 | 14 | 59 |
| Q9WTJ4 | Q9WTJ4 | Flt3-intera | Fiz1 | sp\|Q9WTJ | 2 | 2 | 4 |
| Q7TPN9 | Q7TPN9 | Proline-ric | Prr14 | sp\|Q7TPN | 13 | 13 | 28,9 |
| O55022 | O55022 | Membrane | Pgrmc1 | sp\|O55022 | 10 | 9 | 61,5 |
| Q8VDM4 | Q8VDM4 | 26S protea | Psmd2 | sp\|Q8VDM | 45 | 45 | 60,9 |
| Q9D1C1 | Q9D1C1 | Ubiquitin-c | Ube2c | sp\|Q9D1C | 10 | 10 | 58,7 |
| P30412 | P30412 | Peptidyl-pr | Ppic | sp\|P30412 | 6 | 6 | 42,5 |
| Q60949 | Q60949 | TBC1 doma | Tbc1d1 | sp\|Q60949 | 30 | 30 | 30,8 |

| | | | | | | | |
|---|---|---|---|---|---|---|---|
| Q9R1T4 | Q9R1T4 | Septin-6 | 06-set | sp\|Q9R1T4 | 15 | 7 | 39,4 |
| Q9D0L7 | Q9D0L7 | Armadillo | Armc10 | sp\|Q9D0L7 | 8 | 8 | 42,8 |
| Q8BFY6 | Q8BFY6 | Peflin | Pef1 | sp\|Q8BFY6 | 5 | 5 | 21,1 |
| Q64455 | Q64455 | Receptor-t | Ptprj | sp\|Q64455 | 8 | 8 | 7,1 |
| Q8VDJ3 | Q8VDJ3 | Vigilin | Hdlbp | sp\|Q8VDJ3 | 56 | 56 | 53,9 |
| Q9JMH6 | Q9JMH6 | Thioredoxi | Txnrd1 | sp\|Q9JMH | 25 | 24 | 51,4 |
| P45377 | P45377 | Aldose red | Akr1b8 | sp\|P45377 | 16 | 13 | 58,9 |
| O35448 | O35448 | Lysosomal | Ppt2 | sp\|O35448 | 2 | 2 | 8,6 |
| Q3TLH4 | Q3TLH4 | Protein PR | Prrc2c | sp\|Q3TLH4 | 58 | 56 | 28,1 |
| Q3UHJ0 | Q3UHJ0 | AP2-associ | Aak1 | sp\|Q3UHJ0 | 21 | 20 | 33,5 |
| P97822 | P97822 | Acidic leuc | Anp32e | sp\|P97822 | 9 | 9 | 42,3 |
| O55143;Q8 | O55143 | Sarcoplasm | Atp2a2 | sp\|O55143 | 49 | 43 | 50,1 |
| Q8R1M0 | Q8R1M0 | Embryonic | Hmces | sp\|Q8R1M | 8 | 8 | 30,6 |
| Q61074 | Q61074 | Protein ph | Ppm1g | sp\|Q61074 | 20 | 20 | 43,7 |
| Q8VE96 | Q8VE96 | Solute carr | Slc35f6 | sp\|Q8VE96 | 2 | 2 | 9,1 |
| Q9D8U8 | Q9D8U8 | Sorting nex | Snx5 | sp\|Q9D8U | 20 | 20 | 51 |
| Q9D172 | Q9D172 | ES1 protei | D10Jhu81e | sp\|Q9D172 | 7 | 7 | 36,8 |
| Q8BH79 | Q8BH79 | Anoctamin | Ano10 | sp\|Q8BH79 | 17 | 17 | 32,3 |
| Q8R0H9 | Q8R0H9 | ADP-ribosy | Gga1 | sp\|Q8R0H9 | 12 | 11 | 28,3 |
| P63085 | P63085 | Mitogen-a | Mapk1 | sp\|P63085 | 19 | 14 | 61,7 |
| Q8K1R3 | Q8K1R3 | Polyribonu | Pnpt1 | sp\|Q8K1R3 | 27 | 27 | 40,9 |
| Q8VBW5 | Q8VBW5 | HMG box t | Bbx | sp\|Q8VBW | 13 | 13 | 22,6 |
| P15306 | P15306 | Thrombom | Thbd | sp\|P15306 | 3 | 3 | 9,4 |
| Q91WG4 | Q91WG4 | Elongator | Elp2 | sp\|Q91WG | 27 | 27 | 45,7 |
| Q8BKT3 | Q8BKT3 | GC-rich se | Gcfc2 | sp\|Q8BKT3 | 14 | 14 | 22,9 |
| Q3UHH1 | Q3UHH1 | Zinc finger | Zswim8 | sp\|Q3UHH | 15 | 15 | 12,4 |
| Q4FZF3 | Q4FZF3 | Probable A | Ddx49 | sp\|Q4FZF3 | 14 | 14 | 42,5 |
| Q3UBZ5 | Q3UBZ5 | MIF4G dor | Mif4gd | sp\|Q3UBZ5 | 6 | 6 | 34,7 |
| Q8VBZ3 | Q8VBZ3 | Cleft lip an | Clptm1 | sp\|Q8VBZ3 | 17 | 17 | 36,9 |
| Q8K0V4 | Q8K0V4 | CCR4-NOT | Cnot3 | sp\|Q8K0V4 | 10 | 10 | 19 |
| Q8BVU0 | Q8BVU0 | Leucine-ric | Lrch3 | sp\|Q8BVU | 14 | 14 | 26,5 |
| P97762 | P97762 | Retinitis pi | rp9 | sp\|P97762 | 3 | 3 | 14,6 |
| Q9CQE5 | Q9CQE5 | Regulator | Rgs10 | sp\|Q9CQE | 4 | 4 | 27,1 |
| Q69ZH9 | Q69ZH9 | Rho GTPas | Arhgap23 | sp\|Q69ZH9 | 8 | 8 | 9,2 |
| Q8K296;Q9 | Q8K296 | Myotubula | Mtmr3 | sp\|Q8K296 | 16 | 16 | 20,8 |
| Q61092 | Q61092 | Laminin su | Lamc2 | sp\|Q61092 | 27 | 27 | 31,5 |
| Q9DBP5 | Q9DBP5 | UMP-CMP | Cmpk1 | sp\|Q9DBP5 | 11 | 11 | 53,6 |
| Q9WU62 | Q9WU62 | Inner centr | Incenp | sp\|Q9WU6 | 18 | 18 | 34 |
| O35309 | O35309 | N-myc-inte | Nmi | sp\|O35309 | 8 | 8 | 35 |
| P28798 | P28798 | Granulins; | Grn | sp\|P28798 | 16 | 16 | 42,1 |
| Q9Z2C4 | Q9Z2C4 | Myotubula | Mtmr1 | sp\|Q9Z2C4 | 14 | 14 | 24,8 |
| Q99P88 | Q99P88 | Nuclear po | Nup155 | sp\|Q99P88 | 48 | 48 | 49 |
| P0DN34 | P0DN34 | | | sp\|P0DN34 | 3 | 3 | 33,3 |
| Q8CCS6;Q5 | Q8CCS6 | Polyadeny | Pabpn1 | sp\|Q8CCS6 | 11 | 11 | 44,4 |
| Q8BRF7 | Q8BRF7 | Sec1 family | Scfd1 | sp\|Q8BRF7 | 22 | 22 | 48,2 |
| Q99LM9 | Q99LM9 | Transcripti | Tada1 | sp\|Q99LM | 7 | 7 | 23,6 |
| Q99MS8 | Q99MS8 | Tubulin po | Tpgs1 | sp\|Q99MS | 5 | 5 | 19,5 |
| P04184 | P04184 | Thymidine | Tk1 | sp\|P04184 | 14 | 14 | 58,4 |
| P56695 | P56695 | Wolframin | Wfs1 | sp\|P56695 | 16 | 16 | 21,5 |
| Q9Z1J3 | Q9Z1J3 | Cysteine de | Nfs1 | sp\|Q9Z1J3 | 18 | 18 | 50,1 |

| | | | | | | | |
|---|---|---|---|---|---|---|---|
| P97311 | P97311 | DNA replic | Mcm6 | sp\|P97311 | 45 | 45 | 63,6 |
| Q4VAC9 | Q4VAC9 | Pleckstrin | Plekhg3 | sp\|Q4VAC | 20 | 20 | 22,6 |
| A2ALW5 | A2ALW5 | DnaJ homo | Dnajc25 | sp\|A2ALW | 5 | 5 | 21,6 |
| Q91VS8 | Q91VS8 | FERM, Rho | Farp2 | sp\|Q91VS | 12 | 12 | 18,7 |
| Q8C145 | Q8C145 | Zinc transp | Slc39a6 | sp\|Q8C14 | 3 | 3 | 4,4 |
| Q9DBW3 | Q9DBW3 | Protein NA | Natd1 | sp\|Q9DBW | 3 | 3 | 36,4 |
| Q8K400 | Q8K400 | Syntaxin-b | Stxbp5 | sp\|Q8K40 | 7 | 7 | 10,4 |
| P36371 | P36371 | Antigen pe | Tap2 | sp\|P36371 | 6 | 6 | 10,8 |
| Q91W36 | Q91W36 | Ubiquitin c | Usp3 | sp\|Q91W3 | 16 | 16 | 36,5 |
| O70281 | O70281 | Protein-tyr | Tpst1 | sp\|O7028 | 6 | 6 | 23 |
| Q8C4B4 | Q8C4B4 | Protein un | Unc119b | sp\|Q8C4B | 8 | 7 | 47,8 |
| Q9Z1Q5 | Q9Z1Q5 | Chloride in | Clic1 | sp\|Q9Z1Q | 15 | 15 | 81,7 |
| Q8CFE3 | Q8CFE3 | REST corep | Rcor1 | sp\|Q8CFE3 | 17 | 13 | 49 |
| Q9CWE0 | Q9CWE0 | Mitochond | Mtfr1l | sp\|Q9CWE | 9 | 9 | 39,1 |
| Q8K4R4 | Q8K4R4 | Cytoplasm | Pitpnc1 | sp\|Q8K4R | 5 | 5 | 15,1 |
| Q9DBD5 | Q9DBD5 | Proline-, gl | Pelp1 | sp\|Q9DBD | 19 | 19 | 28,4 |
| Q5SSI6 | Q5SSI6 | U3 small n | Utp18 | sp\|Q5SSI6 | 19 | 19 | 48,4 |
| P36916 | P36916 | Guanine nu | Gnl1 | sp\|P36916 | 21 | 21 | 41,5 |
| Q9JK30 | Q9JK30 | Origin reco | Orc3 | sp\|Q9JK30 | 18 | 18 | 36,9 |
| Q3UFS4 | Q3UFS4 | G patch do | Gpatch11 | sp\|Q3UFS | 3 | 3 | 11,5 |
| P08775 | P08775 | DNA-direct | Polr2a | sp\|P08775 | 65 | 65 | 42,1 |
| Q8K114 | Q8K114 | Integrator | Ints9 | sp\|Q8K11 | 18 | 18 | 37,4 |
| P51163 | P51163 | Uroporphy | Uros | sp\|P51163 | 7 | 7 | 43,4 |
| Q91XI1 | Q91XI1 | tRNA-dihy | Dus3l | sp\|Q91XI1 | 18 | 18 | 49,1 |
| Q6P4T0 | Q6P4T0 | Autophagy | Atg2a | sp\|Q6P4T | 11 | 11 | 8 |
| P26039 | P26039 | Talin-1 | Tln1 | sp\|P26039 | 118 | 107 | 64,6 |
| Q8K1N4 | Q8K1N4 | Spermatog | Spats2 | sp\|Q8K1N | 17 | 17 | 41,3 |
| Q8VHN8 | Q8VHN8 | Protein syr | Nudt16l1 | sp\|Q8VHN | 9 | 8 | 55 |
| Q99LC5 | Q99LC5 | Electron tr | Etfa | sp\|Q99LC5 | 15 | 15 | 60,4 |
| Q8BP00 | Q8BP00 | IQ calmodu | Iqcb1 | sp\|Q8BP0 | 5 | 5 | 9 |
| Q6P8H8 | Q6P8H8 | Probable d | Alg8 | sp\|Q6P8H | 6 | 6 | 14,8 |
| Q99ME9 | Q99ME9 | Nucleolar | Gtpbp4 | sp\|Q99ME | 30 | 30 | 51,1 |
| O35435 | O35435 | Dihydroor | Dhodh | sp\|O3543 | 17 | 17 | 53,7 |
| Q8C7R4 | Q8C7R4 | Ubiquitin-l | Uba6 | sp\|Q8C7R | 41 | 41 | 53 |
| Q80Y83 | Q80Y83 | Dixin | Dixdc1 | sp\|Q80Y8 | 3 | 3 | 6,9 |
| Q3UFK8 | Q3UFK8 | FERM dom | Frmd8 | sp\|Q3UFK | 10 | 10 | 30,5 |
| O55060 | O55060 | Thiopurine | Tpmt | sp\|O5506 | 7 | 7 | 40 |
| P31938 | P31938 | Dual speci | Map2k1 | sp\|P31938 | 17 | 12 | 53,4 |
| P28656 | P28656 | Nucleoson | Nap1l1 | sp\|P28656 | 16 | 15 | 47,8 |
| Q9D8T7 | Q9D8T7 | SRA stem-l | Slirp | sp\|Q9D8T | 8 | 8 | 76,8 |
| Q60I26 | Q60I26 | ALS2 C-ter | Als2cl | sp\|Q60I26 | 8 | 8 | 12,2 |
| Q61239 | Q61239 | Protein far | Fnta | sp\|Q6123 | 13 | 13 | 38,7 |
| Q9D709 | Q9D709 | Transmem | Tmem79 | sp\|Q9D70 | 6 | 6 | 19,7 |
| Q9CZG3 | Q9CZG3 | COMM dor | Commd8 | sp\|Q9CZG | 5 | 5 | 36,1 |
| Q3TC72 | Q3TC72 | Fumarylac | Fahd2 | sp\|Q3TC7 | 5 | 5 | 31,9 |
| Q8K4Z3 | Q8K4Z3 | NAD(P)H-h | Apoa1bp | sp\|Q8K4Z | 8 | 8 | 35,8 |
| P16056 | P16056 | Hepatocyt | Met | sp\|P16056 | 24 | 23 | 22,1 |
| Q8C4J7 | Q8C4J7 | Transducin | Tbl3 | sp\|Q8C4J7 | 33 | 33 | 55,7 |
| Q9QWV9 | Q9QWV9 | Cyclin-T1 | Ccnt1 | sp\|Q9QW | 14 | 13 | 26,9 |
| O70458 | O70458 | Oncostatin | Osmr | sp\|O7045 | 4 | 4 | 6,5 |

| | | | | | | | |
|---|---|---|---|---|---|---|---|
| Q6A0A9 | Q6A0A9 | Constitutiv | FAM120A | sp\|Q6A0A9 | 36 | 34 | 47,4 |
| Q60737 | Q60737 | Casein kina | Csnk2a1 | sp\|Q60737 | 18 | 18 | 55,2 |
| Q3UV70 | Q3UV70 | [Pyruvate d | Pdp1 | sp\|Q3UV7 | 14 | 14 | 40,5 |
| Q61249;Q9 | Q61249 | Immunoglo | Igbp1 | sp\|Q61249 | 14 | 14 | 50,6 |
| O08848 | O08848 | 60 kDa SS-A | Trove2 | sp\|O08848 | 15 | 15 | 38,1 |
| Q9CQR6 | Q9CQR6 | Serine/thre | Ppp6c | sp\|Q9CQR | 10 | 10 | 40,3 |
| Q8C263 | Q8C263 | Spindle and | Ska3 | sp\|Q8C263 | 9 | 9 | 33,3 |
| Q80VJ3 | Q80VJ3 | 2-deoxynu | Dnph1 | sp\|Q80VJ3 | 6 | 6 | 49,1 |
| Q8R3N6 | Q8R3N6 | THO comp | Thoc1 | sp\|Q8R3N | 19 | 19 | 38,1 |
| Q8BHK9 | Q8BHK9 | DNA excisi | Ercc6l | sp\|Q8BHK | 22 | 22 | 25,3 |
| Q9D2R6 | Q9D2R6 | Cytochrom | Coa3 | sp\|Q9D2R | 4 | 4 | 38 |
| Q9DCB8 | Q9DCB8 | Iron-sulfur | Isca2 | sp\|Q9DCB | 7 | 7 | 63,6 |
| Q7TSH3 | Q7TSH3 | Zinc finger | Znf516 | sp\|Q7TSH3 | 16 | 16 | 19,7 |
| Q8K1E0 | Q8K1E0 | Syntaxin-5 | Stx5 | sp\|Q8K1E0 | 9 | 9 | 35,5 |
| Q6PCN7 | Q6PCN7 | Helicase-lik | Hltf | sp\|Q6PCN | 18 | 18 | 23,9 |
| Q8BYI8 | Q8BYI8 | Uncharacte | Kiaa1467 | sp\|Q8BYI8 | 5 | 5 | 14,7 |
| B2RXV4 | B2RXV4 | Feline leuk | Flvcr1 | sp\|B2RXV4 | 2 | 2 | 6,6 |
| Q9QZD8 | Q9QZD8 | Mitochond | Slc25a10 | sp\|Q9QZD | 12 | 12 | 51,9 |
| P35288 | P35288 | Ras-related | Rab23 | sp\|P35288 | 12 | 12 | 50,2 |
| P59325 | P59325 | Eukaryotic | Eif5 | sp\|P59325 | 16 | 16 | 41,5 |
| Q3TTY0 | Q3TTY0 | Phospholip | Plb1 | sp\|Q3TTY0 | 12 | 12 | 12,7 |
| Q80ZJ1 | Q80ZJ1 | Ras-related | Rap2a | sp\|Q80ZJ1 | 8 | 3 | 54,6 |
| P47758 | P47758 | Signal reco | Srprb | sp\|P47758 | 15 | 15 | 56,1 |
| Q8R5K4 | Q8R5K4 | Nucleolar p | Nol6 | sp\|Q8R5K4 | 23 | 23 | 23,7 |
| Q2YDW2 | Q2YDW2 | Protein mi | Msto1 | sp\|Q2YDW | 23 | 23 | 55,4 |
| Q61572;Q6 | Q61572 | Forkhead b | Foxc1 | sp\|Q61572 | 4 | 3 | 10,7 |
| B0V2N1 | B0V2N1 | Receptor-t | Ptprs | sp\|B0V2N1 | 25 | 20 | 21,5 |
| Q99JP0 | Q99JP0 | Mitogen-ac | Map4k3 | sp\|Q99JP0 | 7 | 7 | 11,4 |
| Q9Z1G3;Q9 | Q9Z1G3 | V-type pro | Atp6v1c1 | sp\|Q9Z1G3 | 20 | 20 | 54,7 |
| P70297 | P70297 | Signal tran | Stam | sp\|P70297 | 14 | 12 | 35,9 |
| Q7TPS5 | Q7TPS5 | C2 domain | C2cd5 | sp\|Q7TPS5 | 6 | 6 | 9,4 |
| Q8VCX6 | Q8VCX6 | Kaptin | Kptn | sp\|Q8VCX6 | 2 | 2 | 6,7 |
| Q6PAM0 | Q6PAM0 | 5-AMP-act | Prkab2 | sp\|Q6PAM | 5 | 4 | 23,6 |
| Q61165 | Q61165 | Sodium/hy | Slc9a1 | sp\|Q61165 | 11 | 11 | 20,9 |
| Q6PDZ2 | Q6PDZ2 | TATA box-b | Taf1c | sp\|Q6PDZ2 | 6 | 6 | 12,7 |
| Q99MN1 | Q99MN1 | Lysine--tRN | Kars | sp\|Q99MN | 34 | 34 | 61 |
| Q8C7E9 | Q8C7E9 | Cleavage st | Cstf2t | sp\|Q8C7E9 | 15 | 11 | 34,2 |
| P56399 | P56399 | Ubiquitin c | Usp5 | sp\|P56399 | 33 | 33 | 50,3 |
| O55028 | O55028 | [3-methyl- | Bckdk | sp\|O55028 | 6 | 6 | 23,5 |
| P23949 | P23949 | Zinc finger | Zfp36l2 | sp\|P23949 | 13 | 12 | 31 |
| Q8VDQ9 | Q8VDQ9 | Protein KR | Kri1 | sp\|Q8VDQ | 11 | 11 | 23,7 |
| Q8R1X6 | Q8R1X6 | Spartin | Spg20 | sp\|Q8R1X6 | 15 | 15 | 28,5 |
| P49817;P5 | P49817 | Caveolin-1 | Cav1 | sp\|P49817 | 8 | 8 | 52,2 |
| Q9EPB5 | Q9EPB5 | Serine hyd | Serhl | sp\|Q9EPB5 | 10 | 10 | 41,8 |
| Q3UBX0 | Q3UBX0 | Transmem | Tmem109 | sp\|Q3UBX | 3 | 3 | 9,1 |
| Q60855 | Q60855 | Receptor-i | Ripk1 | sp\|Q60855 | 14 | 14 | 28,5 |
| CON__Q95 | CON__Q95121 | Pigment ep | Serpinf1 | ;sp\|P9729 | 4 | 4 | 10,8 |
| P97807 | P97807 | Fumarate h | Fh | sp\|P97807 | 20 | 20 | 61,1 |
| Q59J78 | Q59J78 | Mimitin, m | Ndufaf2 | sp\|Q59J78 | 10 | 10 | 56,5 |
| P53569 | P53569 | CCAAT/enh | Cebpz | sp\|P53569 | 37 | 37 | 40,7 |

| | | | | | | | |
|---|---|---|---|---|---|---|---|
| Q8BGT1;Q | Q8BGT1 | | Flrt3 | sp\|Q8BGT | 6 | 6 | 13,1 |
| Q8R3N1 | Q8R3N1 | Nucleolar p | Nop14 | sp\|Q8R3N | 27 | 27 | 32,6 |
| Q91XL9 | Q91XL9 | Oxysterol-l | Osbpl1a | sp\|Q91XL9 | 6 | 6 | 9,1 |
| Q9QYY8 | Q9QYY8 | Spastin | Spast | sp\|Q9QYY8 | 15 | 15 | 32,9 |
| P35283 | P35283 | Ras-related | Rab12 | sp\|P35283 | 12 | 12 | 50,2 |
| P56812 | P56812 | Programm | Pdcd5 | sp\|P56812 | 7 | 7 | 50,8 |
| P05064 | P05064 | Fructose-b | Aldoa | sp\|P05064 | 31 | 25 | 84,6 |
| Q8BJ71 | Q8BJ71 | Nuclear po | Nup93 | sp\|Q8BJ71 | 42 | 42 | 53 |
| Q9D5V6 | Q9D5V6 | Synapse-as | Syap1 | sp\|Q9D5V | 12 | 12 | 49,9 |
| Q02395 | Q02395 | Metal-resp | Mtf2 | sp\|Q02395 | 6 | 6 | 16,2 |
| Q8BWR4 | Q8BWR4 | Ubiquitin c | Usp40 | sp\|Q8BWR | 17 | 17 | 19,7 |
| O08602;O0 | O08602;O0860 | Retinoic ac | Raet1a;Rae | sp\|O08602 | 2 | 2 | 8,3 |
| P51949 | P51949 | CDK-activa | Mnat1 | sp\|P51949 | 11 | 11 | 51,5 |
| Q8VEK0;Q8 | Q8VEK0 | Cell cycle c | Tmem30a | sp\|Q8VEK0 | 8 | 8 | 23,9 |
| Q8VD31 | Q8VD31 | Tapasin-re | Tapbpl | sp\|Q8VD3 | 4 | 4 | 15,1 |
| P61025 | P61025 | Cyclin-dep | Cks1b | sp\|P61025 | 2 | 2 | 32,9 |
| Q6PGB6 | Q6PGB6 | N-alpha-ac | Naa50 | sp\|Q6PGB | 15 | 15 | 66,3 |
| Q61142 | Q61142 | Spindlin-1 | Spin1 | sp\|Q61142 | 7 | 7 | 36,3 |
| Q8BHG9 | Q8BHG9 | CGG triplet | Cggbp1 | sp\|Q8BHG | 7 | 7 | 55,7 |
| Q9D358 | Q9D358 | Low molec | Acp1 | sp\|Q9D358 | 6 | 6 | 44,3 |
| P08556 | P08556 | GTPase NR | Nras | sp\|P08556 | 9 | 2 | 52,9 |
| Q9CQ71 | Q9CQ71 | Replication | Rpa3 | sp\|Q9CQ7 | 8 | 8 | 68,6 |
| Q9D1J1 | Q9D1J1 | Adaptin ea | Necap2 | sp\|Q9D1J1 | 10 | 10 | 68,8 |
| Q5XJY5 | Q5XJY5 | Coatomer | Arcn1 | sp\|Q5XJY5 | 26 | 26 | 48,7 |
| Q9D9K3 | Q9D9K3 | Cell death | Aven | sp\|Q9D9K3 | 4 | 4 | 19 |
| Q9R0B9 | Q9R0B9 | Procollage | Plod2 | sp\|Q9R0B9 | 23 | 23 | 37,4 |
| Q61581 | Q61581 | Insulin-like | Igfbp7 | sp\|Q61581 | 6 | 6 | 28,8 |
| P22682 | P22682 | E3 ubiquiti | Cbl | sp\|P22682 | 17 | 16 | 31,2 |
| Q9DBL7 | Q9DBL7 | Bifunction | Coasy | sp\|Q9DBL7 | 9 | 9 | 27,9 |
| P16332 | P16332 | Methylmal | Mut | sp\|P16332 | 22 | 22 | 42,5 |
| Q99KR3 | Q99KR3 | Beta-lactar | Lactb2 | sp\|Q99KR3 | 11 | 11 | 41,7 |
| O70311 | O70311 | Glycylpept | Nmt2 | sp\|O70311 | 15 | 11 | 41,6 |
| Q9CWU2 | Q9CWU2 | Palmitoyltr | Zdhhc13 | sp\|Q9CWU | 13 | 12 | 26 |
| Q06335 | Q06335 | Amyloid-lik | Aplp2 | sp\|Q06335 | 9 | 9 | 17,7 |
| P02463 | P02463 | Collagen al | Col4a1 | sp\|P02463 | 10 | 10 | 9,1 |
| Q9JJ94 | Q9JJ94 | Sjoegren sy | Ssna1 | sp\|Q9JJ94 | 4 | 4 | 46,2 |
| Q5F285 | Q5F285 | Transmem | Tmem256 | sp\|Q5F285 | 2 | 2 | 31,9 |
| Q99JR5 | Q99JR5 | Tubulointe | Tinagl1 | sp\|Q99JR5 | 10 | 10 | 33,5 |
| O70318 | O70318 | Band 4.1-li | Epb41l2 | sp\|O70318 | 44 | 39 | 53,7 |
| Q8C5N5 | Q8C5N5 | Programm | Pdcd2l | sp\|Q8C5N | 2 | 2 | 11,5 |
| Q8CIL4 | Q8CIL4 | Uncharacterized prote | | sp\|Q8CIL4 | 10 | 10 | 38,8 |
| Q3TNL8 | Q3TNL8 | Inositol 1,4 | Itprip | sp\|Q3TNL8 | 9 | 9 | 21,4 |
| Q8R0G9 | Q8R0G9 | Nuclear po | Nup133 | sp\|Q8R0G | 38 | 38 | 46,6 |
| P63328 | P63328 | Serine/thre | Ppp3ca | sp\|P63328 | 11 | 6 | 25,9 |
| Q9D114 | Q9D114 | Guanosine | Hddc3 | sp\|Q9D114 | 5 | 5 | 40,8 |
| Q8K221 | Q8K221 | Arfaptin-2 | Arfip2 | sp\|Q8K221 | 6 | 6 | 19,9 |
| Q91V09 | Q91V09 | WD repeat | Wdr13 | sp\|Q91V09 | 6 | 6 | 14,2 |
| Q3U3T8 | Q3U3T8 | WD repeat | Wdr62 | sp\|Q3U3T8 | 14 | 14 | 13,9 |
| Q64331 | Q64331 | Unconvent | Myo6 | sp\|Q64331 | 24 | 24 | 23,5 |
| Q571H0 | Q571H0 | Nucleolar p | Urb1 | sp\|Q571H0 | 50 | 50 | 29,4 |

| | | | | | | | |
|---|---|---|---|---|---|---|---|
| Q9DBG6 | Q9DBG6 | Dolichyl-di | Rpn2 | sp\|Q9DBG | 26 | 26 | 53,9 |
| P42230 | P42230 | Signal tran | Stat5a | sp\|P42230 | 16 | 4 | 21,6 |
| P39053 | P39053 | Dynamin-1 | Dnm1 | sp\|P39053 | 13 | 5 | 19 |
| Q61171 | Q61171 | Peroxiredo | Prdx2 | sp\|Q61171 | 15 | 15 | 77,3 |
| P97808 | P97808 | FXYD doma | Fxyd5 | sp\|P97808 | 5 | 5 | 52,8 |
| P61208 | P61208 | ADP-ribosy | Arl4c | sp\|P61208 | 4 | 4 | 29,7 |
| Q9CZL2 | Q9CZL2 | Uncharacterized prote | | sp\|Q9CZL2 | 2 | 2 | 14,5 |
| Q9CR29 | Q9CR29 | Coiled-coil | Ccdc43 | sp\|Q9CR29 | 7 | 7 | 31,1 |
| Q80TQ2 | Q80TQ2 | Ubiquitin c | Cyld | sp\|Q80TQ | 15 | 15 | 19,3 |
| Q8BTX9 | Q8BTX9 | Inactive hy | Hsdl1 | sp\|Q8BTX | 9 | 9 | 33,3 |
| Q9QYB5 | Q9QYB5 | Gamma-ad | Add3 | sp\|Q9QYB | 11 | 11 | 22,2 |
| Q8JZS9 | Q8JZS9 | 39S riboso | Mrpl48 | sp\|Q8JZS9 | 2 | 2 | 10,9 |
| Q9Z0Z3 | Q9Z0Z3 | S-phase kir | Skp2 | sp\|Q9Z0Z3 | 8 | 8 | 26,2 |
| Q3UKJ7 | Q3UKJ7 | WD40 repe | Smu1 | sp\|Q3UKJ7 | 22 | 22 | 55 |
| Q8VC31 | Q8VC31 | Coiled-coil | Ccdc9 | sp\|Q8VC31 | 7 | 7 | 12,7 |
| Q6P2L7 | Q6P2L7 | Protein CA | Casc4 | sp\|Q6P2L7 | 9 | 9 | 27,4 |
| P11260 | P11260 | LINE-1 retrotransposa | | sp\|P11260 | 5 | 5 | 16 |
| P97450 | P97450 | ATP syntha | Atp5j | sp\|P97450 | 8 | 8 | 50,9 |
| Q91XU0 | Q91XU0 | ATPase WF | Wrnip1 | sp\|Q91XU | 16 | 16 | 33,8 |
| Q60710 | Q60710 | Deoxynucl | Samhd1 | sp\|Q6071 | 30 | 30 | 52,7 |
| Q8VDR9 | Q8VDR9 | Dedicator | Dock6 | sp\|Q8VDR | 12 | 8 | 7,5 |
| Q2VPA6 | Q2VPA6 | Helicase P | Helq | sp\|Q2VPA | 4 | 4 | 5,8 |
| Q91XU3 | Q91XU3 | Phosphatid | Pip4k2c | sp\|Q91XU | 7 | 7 | 21,9 |
| O88630 | O88630 | Golgi SNAF | Gosr1 | sp\|O8863 | 8 | 8 | 44,4 |
| Q8BMP6 | Q8BMP6 | Golgi resid | Acbd3 | sp\|Q8BMP | 17 | 17 | 42,9 |
| Q91XC8 | Q91XC8 | Death-asso | Dap | sp\|Q91XC | 2 | 2 | 47,1 |
| Q924H5 | Q924H5 | DNA repair | Rad51c | sp\|Q924H | 5 | 5 | 18 |
| Q8BTW8 | Q8BTW8 | CDK5 regu | Cdk5rap1 | sp\|Q8BTW | 9 | 9 | 22,4 |
| Q3TXS7 | Q3TXS7 | 26S protea | Psmd1 | sp\|Q3TXS7 | 45 | 45 | 56,6 |
| Q8CI61 | Q8CI61 | BAG family | Bag4 | sp\|Q8CI61 | 4 | 4 | 12,7 |
| Q8K288 | Q8K288 | Tumor nec | Tnfaip8l1 | sp\|Q8K288 | 3 | 3 | 20,4 |
| P45700 | P45700 | Mannosyl- | Man1a1 | sp\|P45700 | 10 | 10 | 21,7 |
| A2RTL5 | A2RTL5 | Arginine/s | Rsrc2 | sp\|A2RTL5 | 5 | 5 | 18,4 |
| Q8BI72 | Q8BI72 | CDKN2A-in | Cdkn2aip | sp\|Q8BI72 | 13 | 13 | 35,5 |
| Q99J09 | Q99J09 | Methyloso | Wdr77 | sp\|Q99J09 | 10 | 10 | 46,8 |
| Q8BMA5 | Q8BMA5 | Protein NP | Npat | sp\|Q8BMA | 13 | 13 | 14,7 |
| Q9CXW2 | Q9CXW2 | 28S riboso | Mrps22 | sp\|Q9CXW | 19 | 19 | 55,4 |
| O88307 | O88307 | Sortilin-rel | Sorl1 | sp\|O88307 | 8 | 8 | 5,2 |
| Q61578 | Q61578 | NADPH:ad | Fdxr | sp\|Q61578 | 18 | 18 | 47,4 |
| Q91ZX7 | Q91ZX7 | Prolow-de | Lrp1 | sp\|Q91ZX7 | 113 | 112 | 34,9 |
| Q9EPU0;Q | Q9EPU0 | Regulator | Upf1 | sp\|Q9EPU | 53 | 53 | 57,7 |
| Q8VCH7 | Q8VCH7 | Retinol del | Rdh10 | sp\|Q8VCH | 7 | 7 | 25,8 |
| Q3TWL2 | Q3TWL2 | Type 1 pho | Tmem55b | sp\|Q3TWL | 4 | 3 | 19 |
| Q61083 | Q61083 | Mitogen-a | Map3k2 | sp\|Q61083 | 14 | 10 | 33,1 |
| Q8CI75 | Q8CI75 | DIS3-like e | Dis3l2 | sp\|Q8CI75 | 5 | 5 | 8,2 |
| Q8R349 | Q8R349 | Cell divisio | Cdc16 | sp\|Q8R349 | 19 | 19 | 43,1 |
| O54804 | O54804 | Choline kir | Chka | sp\|O54804 | 9 | 9 | 22,5 |
| O35345 | O35345 | Importin s | Kpna6 | sp\|O35345 | 16 | 10 | 34 |
| Q8CAS9 | Q8CAS9 | Poly [ADP- | Parp9 | sp\|Q8CAS | 11 | 11 | 16,1 |
| P20029 | P20029 | 78 kDa glu | Hspa5 | sp\|P20029 | 47 | 45 | 64,4 |

| | | | | | | | |
|---|---|---|---|---|---|---|---|
| Q91W39 | Q91W39 | Nuclear re | Ncoa5 | sp\|Q91W3 | 21 | 21 | 48,4 |
| Q8C0C7 | Q8C0C7 | Phenylalan | Farsa | sp\|Q8C0C7 | 20 | 20 | 42,5 |
| P59016 | P59016 | Vacuolar p | Vps33b | sp\|P59016 | 15 | 15 | 29,2 |
| Q9QZN4 | Q9QZN4 | F-box only | Fbxo6 | sp\|Q9QZN | 9 | 9 | 49,2 |
| P84096 | P84096 | Rho-relate | Rhog | sp\|P84096 | 11 | 10 | 62,3 |
| Q9D142 | Q9D142 | Uridine dip | Nudt14 | sp\|Q9D142 | 4 | 4 | 21,2 |
| Q5SSH7 | Q5SSH7 | Zinc finger | Zzef1 | sp\|Q5SSH7 | 24 | 24 | 12,8 |
| Q6ZWZ2 | Q6ZWZ2 | Ubiquitin-c | Ube2r2 | sp\|Q6ZWZ | 3 | 3 | 12,2 |
| Q8VC03 | Q8VC03 | Echinoderr | Eml3 | sp\|Q8VC03 | 10 | 10 | 15,9 |
| P28667 | P28667 | MARCKS-re | Marcksl1 | sp\|P28667 | 7 | 7 | 70 |
| Q9Z1N5 | Q9Z1N5 | Spliceosom | Ddx39b | sp\|Q9Z1N5 | 24 | 14 | 64,3 |
| Q9CQX4 | Q9CQX4 | PCNA-asso | Paf | sp\|Q9CQX | 5 | 5 | 55,5 |
| Q9QYF1 | Q9QYF1 | Retinol del | Rdh11 | sp\|Q9QYF1 | 10 | 10 | 40,8 |
| Q6PAL0 | Q6PAL0 | BEN doma | Bend3 | sp\|Q6PAL0 | 17 | 17 | 26,4 |
| Q9CY52 | Q9CY52 | Probable t | Thg1l | sp\|Q9CY52 | 7 | 7 | 28,2 |
| O70492 | O70492 | Sorting ne | Snx3 | sp\|O70492 | 11 | 9 | 74,1 |
| O54785;P5 | O54785 | LIM domai | Limk2 | sp\|O54785 | 11 | 11 | 23,4 |
| Q80YD1 | Q80YD1 | ATP-depen | Supv3l1 | sp\|Q80YD1 | 15 | 15 | 25,9 |
| Q11136 | Q11136 | Xaa-Pro di | Pepd | sp\|Q11136 | 17 | 17 | 41,6 |
| E1U8D0 | E1U8D0 | Protein SO | Soga1 | sp\|E1U8D0 | 28 | 27 | 23,6 |
| Q6PAC3 | Q6PAC3 | DDB1- and | Dcaf13 | sp\|Q6PAC3 | 16 | 16 | 42,5 |
| Q99JT1 | Q99JT1 | Glutamyl-t | Gatb | sp\|Q99JT1 | 8 | 8 | 26,4 |
| P25976 | P25976 | Nucleolar t | Ubtf | sp\|P25976 | 24 | 24 | 31,1 |
| Q61151 | Q61151 | Serine/thre | Ppp2r5e | sp\|Q61151 | 19 | 14 | 39,6 |
| Q9CR14 | Q9CR14 | E3 ubiquiti | Fancl | sp\|Q9CR14 | 4 | 4 | 12,8 |
| O88597 | O88597 | Beclin-1 | Becn1 | sp\|O88597 | 6 | 6 | 18,8 |
| Q9CRA5 | Q9CRA5 | Golgi phos | Golph3 | sp\|Q9CRA5 | 10 | 8 | 46 |
| P62918 | P62918 | 60S riboso | Rpl8 | sp\|P62918 | 14 | 14 | 54,5 |
| Q9D2C2 | Q9D2C2 | Protein SA | Saal1 | sp\|Q9D2C2 | 13 | 13 | 43,2 |
| Q80ZS3 | Q80ZS3 | 28S riboso | Mrps26 | sp\|Q80ZS3 | 6 | 6 | 34 |
| Q8VI84 | Q8VI84 | Nucleolar c | Noc3l | sp\|Q8VI84 | 18 | 18 | 27,8 |
| Q8BHI7 | Q8BHI7 | Elongation | Elovl5 | sp\|Q8BHI7 | 3 | 3 | 16,4 |
| Q7TPD0 | Q7TPD0 | Integrator | Ints3 | sp\|Q7TPD0 | 30 | 30 | 42,3 |
| Q68FE6 | Q68FE6 | Protein FA | Fam65a | sp\|Q68FE6 | 14 | 14 | 15,9 |
| Q9QXL8 | Q9QXL8 | Nucleoside | Nme7 | sp\|Q9QXL8 | 11 | 11 | 35,9 |
| E9PVX6 | E9PVX6 | | Mki67 | sp\|E9PVX6 | 155 | 155 | 59,3 |
| P62827;Q6 | P62827 | GTP-bindir | Ran | sp\|P62827 | 13 | 13 | 58,8 |
| Q3UHN9 | Q3UHN9 | Bifunctiona | Ndst1 | sp\|Q3UHN | 5 | 4 | 7,6 |
| Q3TYA6 | Q3TYA6 | M-phase p | Mphosph8 | sp\|Q3TYA6 | 12 | 12 | 15,5 |
| Q91XC9 | Q91XC9 | Peroxisom | Pex16 | sp\|Q91XC9 | 5 | 5 | 20,8 |
| Q3ZK22 | Q3ZK22 | Vezatin | Vezt | sp\|Q3ZK22 | 15 | 15 | 31,4 |
| Q7TMM8 | Q7TMM8 | Mono [ADI | Parp16 | sp\|Q7TMN | 4 | 4 | 14 |
| Q99KE1;Q8 | Q99KE1 | NAD-deper | Me2 | sp\|Q99KE1 | 26 | 26 | 61,3 |
| Q3U9G9 | Q3U9G9 | Lamin-B re | Lbr | sp\|Q3U9G | 17 | 17 | 29,6 |
| Q8CCI5 | Q8CCI5 | RING1 and | Rybp | sp\|Q8CCI5 | 3 | 3 | 26,3 |
| P70403 | P70403 | Protein CA | Cux1 | sp\|P70403 | 19 | 4 | 36,4 |
| O35242 | O35242 | Protein FA | Nsmaf | sp\|O35242 | 6 | 6 | 8,7 |
| Q99JY8 | Q99JY8 | Lipid phos | Ppap2b | sp\|Q99JY8 | 6 | 6 | 20,8 |
| Q8R1K1 | Q8R1K1 | Ubiquitin-a | Ubac2 | sp\|Q8R1K1 | 6 | 6 | 21,4 |
| Q8BGP6 | Q8BGP6 | Solute carr | Slc25a40 | sp\|Q8BGP | 10 | 10 | 33,8 |

| | | | | | | | |
|---|---|---|---|---|---|---|---|
| Q99PL5 | Q99PL5 | Ribosome- | Rrbp1 | sp\|Q99PL5 | 50 | 50 | 45 |
| Q99J47 | Q99J47 | Dehydroge | Dhrs7b | sp\|Q99J47 | 15 | 15 | 45,8 |
| Q80UY1 | Q80UY1 | UPF0586 protein C9or | | sp\|Q80UY1 | 6 | 6 | 17,8 |
| P27048;P6 | P27048;P6316 | Small nucle | Snrpb;Snrp | sp\|P27048 | 10 | 10 | 34,6 |
| Q91V92 | Q91V92 | ATP-citrate | Acly | sp\|Q91V92 | 45 | 45 | 51,6 |
| O55236 | O55236 | mRNA-cap | Rngtt | sp\|O55236 | 14 | 14 | 32,5 |
| O35719 | O35719 | DNA repair | Rad51b | sp\|O35719 | 5 | 5 | 15,7 |
| Q6KAU4 | Q6KAU4 | Multivesic | Mvb12b | sp\|Q6KAU | 6 | 6 | 37,2 |
| Q921L5 | Q921L5 | Conserved | Cog2 | sp\|Q921L5 | 19 | 19 | 45,8 |
| CON__Q3S | CON__Q3SZH5 | | | | 7 | 7 | 25,5 |
| Q9CXL3 | Q9CXL3 | Uncharacterized prote | | sp\|Q9CXL3 | 4 | 4 | 39,5 |
| O70433 | O70433 | Four and a | Fhl2 | sp\|O70433 | 14 | 14 | 57,3 |
| Q8R395 | Q8R395 | COMM do | Commd5 | sp\|Q8R395 | 6 | 6 | 36,6 |
| Q80VD1 | Q80VD1 | Protein FA | Fam98b | sp\|Q80VD | 14 | 11 | 39,9 |
| Q3TDD9 | Q3TDD9 | Protein ph | Ppp1r21 | sp\|Q3TDD | 14 | 14 | 27,9 |
| Q9JHJ0;Q9 | Q9JHJ0 | Tropomod | Tmod3 | sp\|Q9JHJ0 | 15 | 15 | 45,2 |
| Q9JIG8 | Q9JIG8 | PRA1 famil | Praf2 | sp\|Q9JIG8 | 5 | 5 | 28,7 |
| Q8R422 | Q8R422 | CD109 ant | Cd109 | sp\|Q8R422 | 15 | 15 | 14 |
| Q3U3Q1 | Q3U3Q1 | Serine/thre | Ulk3 | sp\|Q3U3Q | 4 | 4 | 11 |
| Q9Z1M8 | Q9Z1M8 | Protein Re | Ik | sp\|Q9Z1M | 19 | 19 | 43,1 |
| Q566J8 | Q566J8 | AarF doma | Adck4 | sp\|Q566J8 | 7 | 7 | 20,1 |
| Q8K2Q5 | Q8K2Q5 | Coiled-coil | Chchd7 | sp\|Q8K2Q | 1 | 1 | 21,2 |
| Q9CZP5 | Q9CZP5 | Mitochond | Bcs1l | sp\|Q9CZP5 | 16 | 16 | 49,3 |
| Q9WUH1 | Q9WUH1 | Transmem | Tmem115 | sp\|Q9WUH | 4 | 4 | 16,3 |
| Q3V038 | Q3V038 | Tetratricop | Ttc9 | sp\|Q3V038 | 3 | 3 | 16 |
| O88327 | O88327 | Alpha-catu | Ctnnal1 | sp\|O88327 | 12 | 12 | 20,8 |
| Q04447 | Q04447 | Creatine ki | Ckb | sp\|Q04447 | 10 | 10 | 43,8 |
| Q3UIL6 | Q3UIL6 | Pleckstrin | Plekha7 | sp\|Q3UIL6 | 13 | 13 | 13,1 |
| Q80X50 | Q80X50 | Ubiquitin-a | Ubap2l | sp\|Q80X50 | 23 | 23 | 37,9 |
| O55234 | O55234 | Proteasom | Psmb5 | sp\|O55234 | 16 | 16 | 66,3 |
| P39061 | P39061 | Collagen al | Col18a1 | sp\|P39061 | 21 | 21 | 16 |
| O35988 | O35988 | Syndecan-4 | Sdc4 | sp\|O35988 | 4 | 4 | 22,7 |
| O70477 | O70477 | Homeobox | Pknox1 | sp\|O70477 | 4 | 4 | 13,8 |
| Q9CPR5 | Q9CPR5 | 39S riboso | Mrpl15 | sp\|Q9CPR5 | 16 | 16 | 64,1 |
| Q80WG5 | Q80WG5 | Volume-re | Lrrc8a | sp\|Q80WG | 20 | 18 | 30,5 |
| Q8BRG8 | Q8BRG8 | Transmem | Tmem209 | sp\|Q8BRG | 14 | 14 | 39 |
| Q8BTI9;O3 | Q8BTI9 | Phosphatid | Pik3cb | sp\|Q8BTI9 | 10 | 10 | 12 |
| B9EJ80 | B9EJ80 | | Pdzd8 | sp\|B9EJ80 | 21 | 21 | 25,8 |
| P70388 | P70388 | DNA repair | Rad50 | sp\|P70388 | 54 | 54 | 40,2 |
| P58465 | P58465 | CTD small | Ctdspl | sp\|P58465 | 3 | 2 | 12,3 |
| Q8K202 | Q8K202 | DNA-direct | Polr1e | sp\|Q8K202 | 17 | 17 | 47,7 |
| Q8K389 | Q8K389 | CDK5 regu | Cdk5rap2 | sp\|Q8K389 | 22 | 22 | 15,4 |
| Q8BK08 | Q8BK08 | Transmem | Tmem11 | sp\|Q8BK08 | 4 | 4 | 22,6 |
| Q9DAX2 | Q9DAX2 | Lipid phos | Ppap2c | sp\|Q9DAX | 4 | 4 | 15,9 |
| CON__A2I | CON__A2I7N0 | | | | 9 | 1 | 22,9 |
| Q9JJ06 | Q9JJ06 | Glycoprote | C1galt1 | sp\|Q9JJ06 | 4 | 4 | 14,6 |
| Q9D1G5 | Q9D1G5 | Leucine-ric | Lrrc57 | sp\|Q9D1G | 4 | 4 | 18,8 |
| Q8CEE6 | Q8CEE6 | PAS domai | Pask | sp\|Q8CEE6 | 14 | 14 | 14,6 |
| Q6ZQ93 | Q6ZQ93 | Ubiquitin c | Usp34 | sp\|Q6ZQ93 | 43 | 43 | 17,8 |
| Q8BYL4 | Q8BYL4 | Tyrosine--t | Yars2 | sp\|Q8BYL4 | 10 | 10 | 35 |

| | | | | | | | |
|---|---|---|---|---|---|---|---|
| Q7TMY7 | Q7TMY7 | Importin-8 | Ipo8 | sp\|Q7TMY | 21 | 20 | 23,5 |
| Q6ZWY3 | Q6ZWY3 | 40S riboso | Rps27l | sp\|Q6ZWY | 6 | 3 | 41,7 |
| Q91WJ8 | Q91WJ8 | Far upstrea | Fubp1 | sp\|Q91WJ | 27 | 24 | 51 |
| P35235 | P35235 | Tyrosine-p | Ptpn11 | sp\|P35235 | 22 | 22 | 51,9 |
| Q9WV70 | Q9WV70 | Nucleolar ( | Noc2l | sp\|Q9WV7 | 15 | 15 | 18,3 |
| Q80U78 | Q80U78 | Pumilio ho | Pum1 | sp\|Q80U78 | 21 | 18 | 25,7 |
| Q9D8E6 | Q9D8E6 | 60S riboso | Rpl4 | sp\|Q9D8E6 | 22 | 22 | 48,7 |
| Q9JK42 | Q9JK42 | [Pyruvate ( | Pdk2 | sp\|Q9JK42 | 1 | 1 | 3,9 |
| P11930 | P11930 | Nucleoside | Nudt19 | sp\|P11930 | 9 | 9 | 36,4 |
| O35654 | O35654 | DNA polym | Pold2 | sp\|O35654 | 15 | 15 | 47,5 |
| Q8VIK2 | Q8VIK2 | Mpv17-like | Mpv17l2 | sp\|Q8VIK2 | 3 | 3 | 21,5 |
| O35680 | O35680 | 28S riboso | Mrps12 | sp\|O35680 | 2 | 2 | 25,9 |
| Q08775;Q( | Q08775 | Runt-relate | Runx2 | sp\|Q08775 | 8 | 8 | 21,3 |
| Q9JJA7 | Q9JJA7 | Cyclin-L2 | Ccnl2 | sp\|Q9JJA7 | 4 | 3 | 12 |
| Q9QY66 | Q9QY66 | Zinc finger | Znhit2 | sp\|Q9QY6( | 6 | 6 | 19,8 |
| P46938 | P46938 | Transcripti | Yap1 | sp\|P46938 | 12 | 12 | 47,5 |
| Q00612 | Q00612 | Glucose-6- | G6pdx | sp\|Q00612 | 29 | 20 | 68,9 |
| Q61087 | Q61087 | Laminin su | Lamb3 | sp\|Q61087 | 34 | 34 | 41,4 |
| Q8R1S4 | Q8R1S4 | Metastasis | Mtss1 | sp\|Q8R1S4 | 7 | 7 | 16,5 |
| P46737 | P46737 | Lys-63-spe | Brcc3 | sp\|P46737 | 10 | 10 | 33,7 |
| Q9QY23 | Q9QY23 | Plakophilin | Pkp3 | sp\|Q9QY2 | 18 | 18 | 31 |
| Q61084 | Q61084 | Mitogen-ac | Map3k3 | sp\|Q61084 | 9 | 5 | 19,6 |
| Q3TPX4 | Q3TPX4 | Exocyst co | Exoc5 | sp\|Q3TPX4 | 17 | 17 | 33,2 |
| P30276 | P30276 | G2/mitotic | Ccnb2 | sp\|P30276 | 9 | 9 | 31,7 |
| Q8K2C6 | Q8K2C6 | NAD-deper | Sirt5 | sp\|Q8K2C6 | 5 | 5 | 23,9 |
| Q60989 | Q60989 | E3 ubiquiti | Xiap | sp\|Q60989 | 12 | 12 | 34,7 |
| Q91WC0 | Q91WC0 | Histone-lys | Setd3 | sp\|Q91WC | 21 | 21 | 41,1 |
| P01900;P0 | P01900;P0189 | H-2 class I | H2-D1 | sp\|P01900 | 13 | 7 | 39,7 |
| Q9D6R2 | Q9D6R2 | Isocitrate ( | Idh3a | sp\|Q9D6R | 18 | 18 | 49,5 |
| O55187 | O55187 | E3 SUMO-p | Cbx4 | sp\|O55187 | 11 | 11 | 23,2 |
| Q9D0E1;Q | Q9D0E1 | Heterogen | Hnrnpm | sp\|Q9D0E1 | 48 | 48 | 67,5 |
| A2RSY6 | A2RSY6 | TRMT1-like | Trmt1l | sp\|A2RSY6 | 21 | 21 | 47,3 |
| Q9CQ80 | Q9CQ80 | Vacuolar p | Vps25 | sp\|Q9CQ8 | 12 | 12 | 78,4 |
| P17751 | P17751 | Triosephos | Tpi1 | sp\|P17751 | 18 | 18 | 72,2 |
| Q8K409 | Q8K409 | DNA polym | Polb | sp\|Q8K409 | 9 | 9 | 32,2 |
| O35691 | O35691 | Pinin | Pnn | sp\|O35691 | 20 | 20 | 29,9 |
| Q921I6 | Q921I6 | SH3 domai | Sh3bp4 | sp\|Q921I6 | 19 | 19 | 28,1 |
| Q9DBV4 | Q9DBV4 | Matrix-rem | Mxra8 | sp\|Q9DBV | 7 | 7 | 17,9 |
| Q9R0X4;Q | Q9R0X4;Q32M | Acyl-coenz | Acot9;Acot | sp\|Q9R0X4 | 23 | 23 | 59,7 |
| Q6KAQ7 | Q6KAQ7 | ZZ-type zin | Zzz3 | sp\|Q6KAQ | 7 | 7 | 12,9 |
| Q811U4 | Q811U4 | Mitofusin- | Mfn1 | sp\|Q811U4 | 22 | 20 | 35,9 |
| A2AJI0 | A2AJI0 | MAP7 dom | Map7d1 | sp\|A2AJI0 | 13 | 13 | 21,6 |
| Q499E6 | Q499E6 | Uncharacterized prote | | sp\|Q499E6 | 2 | 2 | 13,8 |
| Q64324 | Q64324 | Syntaxin-b | Stxbp2 | sp\|Q64324 | 23 | 23 | 44,7 |
| O09159 | O09159 | Lysosomal | Man2b1 | sp\|O09159 | 15 | 15 | 21,2 |
| Q6P5B0 | Q6P5B0 | RRP12-like | Rrp12 | sp\|Q6P5B0 | 39 | 39 | 37,2 |
| P52912 | P52912 | Nucleolysin | Tia1 | sp\|P52912 | 8 | 5 | 28,8 |
| Q9CYX7 | Q9CYX7 | RRP15-like | Rrp15 | sp\|Q9CYX7 | 7 | 7 | 20,6 |
| Q8CFY5 | Q8CFY5 | Protoheme | Cox10 | sp\|Q8CFY5 | 3 | 3 | 12 |
| Q8K3G9 | Q8K3G9 | DCC-intera | Appl2 | sp\|Q8K3G9 | 6 | 6 | 14,2 |

| | | | | | | | |
|---|---|---|---|---|---|---|---|
| P62743 | P62743 | AP-2 comp | Ap2s1 | sp\|P62743 | 7 | 7 | 44,4 |
| Q8BP56 | Q8BP56 | Acid trehal | Athl1 | sp\|Q8BP56 | 8 | 8 | 15,1 |
| Q8CFI7 | Q8CFI7 | DNA-direct | Polr2b | sp\|Q8CFI7 | 49 | 49 | 52,2 |
| P59470 | P59470 | DNA-direct | Polr3b | sp\|P59470 | 11 | 11 | 12,1 |
| P56375 | P56375 | Acylphospl | Acyp2 | sp\|P56375 | 4 | 4 | 36,8 |
| Q8VE08 | Q8VE08 | F-box only | Fbxo33 | sp\|Q8VE08 | 2 | 2 | 3,7 |
| Q9Z2Q6 | Q9Z2Q6 | Septin-5 | 05-set | sp\|Q9Z2Q6 | 9 | 9 | 33,3 |
| Q9Z204;Q8 | Q9Z204 | Heterogen | Hnrnpc | sp\|Q9Z204 | 19 | 19 | 54,3 |
| P60824 | P60824 | Cold-induc | Cirbp | sp\|P60824 | 5 | 5 | 38,4 |
| Q80W47 | Q80W47 | WD repeat | Wipi2 | sp\|Q80W4 | 11 | 11 | 45,4 |
| P97930 | P97930 | Thymidylat | Dtymk | sp\|P97930 | 13 | 13 | 53,3 |
| Q9Z0S9 | Q9Z0S9 | Prenylated | Rabac1 | sp\|Q9Z0S9 | 2 | 2 | 15,7 |
| P85094 | P85094 | Isochorism | Isoc2a | sp\|P85094 | 5 | 5 | 46,1 |
| P42227 | P42227 | Signal tran | Stat3 | sp\|P42227 | 24 | 24 | 49,1 |
| Q9D1C8 | Q9D1C8 | Vacuolar p | Vps28 | sp\|Q9D1C8 | 10 | 10 | 62,4 |
| O35551 | O35551 | Rab GTPas | Rabep1 | sp\|O35551 | 16 | 16 | 23,3 |
| Q9D8V0 | Q9D8V0 | Minor hist | Hm13 | sp\|Q9D8V0 | 8 | 8 | 20,6 |
| Q64362 | Q64362 | AKT-intera | Aktip | sp\|Q64362 | 3 | 3 | 20,5 |
| Q3TPE9 | Q3TPE9 | Ankyrin re | Ankmy2 | sp\|Q3TPE9 | 10 | 10 | 22,5 |
| Q5RJY2 | Q5RJY2 | G2/M phas | G2e3 | sp\|Q5RJY2 | 1 | 1 | 2,5 |
| Q8BJA3 | Q8BJA3 | Homeobox | Hmbox1 | sp\|Q8BJA3 | 3 | 3 | 10,5 |
| Q9QYF9 | Q9QYF9 | Protein ND | Ndrg3 | sp\|Q9QYF9 | 11 | 11 | 44,3 |
| Q99LH1 | Q99LH1 | Nucleolar | Gnl2 | sp\|Q99LH1 | 19 | 19 | 27,6 |
| P0CB42 | P0CB42 | Alkylated D | Alkbh1 | sp\|P0CB42 | 4 | 4 | 15,4 |
| P62862 | P62862 | 40S riboso | Fau | sp\|P62862 | 3 | 3 | 20,3 |
| Q61543 | Q61543 | Golgi appa | Glg1 | sp\|Q61543 | 37 | 37 | 42,9 |
| Q8CCN5 | Q8CCN5 | Breast card | Bcas3 | sp\|Q8CCN | 9 | 9 | 14,1 |
| P63005 | P63005 | Platelet-ac | Pafah1b1 | sp\|P63005 | 23 | 23 | 64,1 |
| Q91YS8 | Q91YS8 | Calcium/ca | Camk1 | sp\|Q91YS8 | 10 | 8 | 42,2 |
| Q6PCM1 | Q6PCM1 | Lysine-spe | Kdm3a | sp\|Q6PCM | 36 | 35 | 38,4 |
| Q9CZH3 | Q9CZH3 | Proteasom | Psmg3 | sp\|Q9CZH3 | 4 | 4 | 50 |
| Q922L6 | Q922L6 | Negative e | Nelfcd | sp\|Q922L6 | 14 | 14 | 35,5 |
| Q9ER64 | Q9ER64 | Oxysterol- | Osbpl5 | sp\|Q9ER64 | 16 | 14 | 27,8 |
| P31750 | P31750 | RAC-alpha | Akt1 | sp\|P31750 | 21 | 15 | 45 |
| Q7TPM9 | Q7TPM9 | Myotubula | Mtmr10 | sp\|Q7TPM | 19 | 19 | 35,1 |
| Q9Z2M7 | Q9Z2M7 | Phosphom | Pmm2 | sp\|Q9Z2M | 12 | 10 | 49,6 |
| Q8CIE4 | Q8CIE4 | | Parp10 | sp\|Q8CIE4 | 10 | 10 | 19,1 |
| Q8BM75 | Q8BM75 | AT-rich int | Arid5b | sp\|Q8BM7 | 12 | 12 | 14,5 |
| Q91X88 | Q91X88 | Protein O- | Pomgnt1 | sp\|Q91X88 | 3 | 3 | 5,5 |
| Q9D3D9 | Q9D3D9 | ATP syntha | Atp5d | sp\|Q9D3D | 4 | 4 | 43,5 |
| P59481 | P59481 | VIP36-like | Lman2l | sp\|P59481 | 6 | 6 | 24,2 |
| Q9JJG9 | Q9JJG9 | Nitric oxid | Noa1 | sp\|Q9JJG9 | 14 | 14 | 27,6 |
| Q91YR7 | Q91YR7 | Pre-mRNA | Prpf6 | sp\|Q91YR7 | 43 | 43 | 48,2 |
| Q9JHS4 | Q9JHS4 | ATP-depen | Clpx | sp\|Q9JHS4 | 22 | 22 | 42 |
| G5E870 | G5E870 | E3 ubiquiti | Trip12 | sp\|G5E870 | 47 | 47 | 32,2 |
| Q9D8S4 | Q9D8S4 | Oligoribon | Rexo2 | sp\|Q9D8S4 | 8 | 8 | 34,6 |
| P97300 | P97300 | Neuroplast | Nptn | sp\|P97300 | 5 | 5 | 16,4 |
| Q6P8X1;Q8 | Q6P8X1 | Sorting ne | Snx6 | sp\|Q6P8X1 | 22 | 22 | 52,2 |
| Q9D061 | Q9D061 | Acyl-CoA-b | Acbd6 | sp\|Q9D061 | 11 | 11 | 44 |
| Q8VDC0 | Q8VDC0 | Probable le | Lars2 | sp\|Q8VDC | 18 | 18 | 24,9 |

| | | | | | | | |
|---|---|---|---|---|---|---|---|
| Q99MK9 | Q99MK9 | Ras associa | Rassf1 | sp\|Q99MK | 11 | 11 | 40,3 |
| Q80U93 | Q80U93 | Nuclear po | Nup214 | sp\|Q80U93 | 47 | 47 | 33,2 |
| P58801 | P58801 | Receptor-i | Ripk2 | sp\|P58801 | 9 | 9 | 19,9 |
| P97355 | P97355 | Spermine s | Sms | sp\|P97355 | 9 | 9 | 39,3 |
| Q9D2E2 | Q9D2E2 | Target of E | Toe1 | sp\|Q9D2E2 | 11 | 11 | 28,4 |
| CON__Q3Y | CON__Q3Y5Z3 | | | | 4 | 4 | 21,2 |
| Q7TT50 | Q7TT50 | Serine/thre | Cdc42bpb | sp\|Q7TT50 | 56 | 52 | 38,8 |
| Q64339 | Q64339 | Ubiquitin-l | Isg15 | sp\|Q64339 | 4 | 4 | 36 |
| P70671 | P70671 | Interferon | Irf3 | sp\|P70671 | 7 | 7 | 28,9 |
| Q8C5P5 | Q8C5P5 | 5-nucleotid | Nt5dc1 | sp\|Q8C5P5 | 13 | 13 | 39,6 |
| Q9CPU0 | Q9CPU0 | Lactoylglut | Glo1 | sp\|Q9CPU | 16 | 16 | 85,9 |
| Q8R0G7 | Q8R0G7 | Protein spi | Spns1 | sp\|Q8R0G | 7 | 7 | 24,4 |
| Q9CS00 | Q9CS00 | Cactin | Cactin | sp\|Q9CS00 | 14 | 14 | 24,7 |
| Q9DB96 | Q9DB96 | Neuroguid | Ngdn | sp\|Q9DB96 | 9 | 9 | 45,1 |
| A2APB8 | A2APB8 | Targeting p | Tpx2 | sp\|A2APB8 | 31 | 31 | 45,9 |
| Q9QY36 | Q9QY36 | N-alpha-ac | Naa10 | sp\|Q9QY36 | 11 | 8 | 65,1 |
| CON__Q2U | CON__Q2UVX4 | | | | 20 | 18 | 15,8 |
| Q9Z2D8 | Q9Z2D8 | Methyl-Cp | Mbd3 | sp\|Q9Z2D8 | 12 | 12 | 48,4 |
| O88572 | O88572 | Low-densit | Lrp6 | sp\|O88572 | 2 | 1 | 1,1 |
| Q78HU3 | Q78HU3 | Multivesicu | Mvb12a | sp\|Q78HU | 6 | 6 | 28,4 |
| Q6A098 | Q6A098 | Selenocyst | Secisbp2l | sp\|Q6A098 | 6 | 6 | 8,7 |
| A2A791 | A2A791 | Zinc finger | Zmym4 | sp\|A2A791 | 20 | 20 | 20,5 |
| Q9EP82 | Q9EP82 | tRNA (guar | Wdr4 | sp\|Q9EP82 | 16 | 16 | 58,6 |
| Q80TE0 | Q80TE0 | RNA polym | Rpap1 | sp\|Q80TE0 | 15 | 15 | 16,5 |
| Q7TQK5 | Q7TQK5 | Coiled-coil | Ccdc93 | sp\|Q7TQK | 16 | 16 | 30,4 |
| Q8BH07 | Q8BH07 | ADP-ribosy | Arl6ip6 | sp\|Q8BH0 | 3 | 3 | 24,8 |
| Q9D864 | Q9D864 | Actin-relat | Actr6 | sp\|Q9D864 | 9 | 9 | 34,3 |
| Q6PCQ0 | Q6PCQ0 | IQ domain- | Iqce | sp\|Q6PCQ | 7 | 7 | 15,3 |
| Q8K327 | Q8K327 | Chromoso | Champ1 | sp\|Q8K327 | 25 | 25 | 38,4 |
| P53702 | P53702 | Cytochrom | Hccs | sp\|P53702 | 13 | 13 | 51,1 |
| Q8BYC6 | Q8BYC6 | Serine/thre | Taok3 | sp\|Q8BYC6 | 10 | 8 | 12,4 |
| Q61211 | Q61211 | Eukaryotic | Eif2d | sp\|Q61211 | 18 | 18 | 48,8 |
| Q61792;Q9 | Q61792 | LIM and SH | Lasp1 | sp\|Q61792 | 14 | 14 | 49 |
| Q6ZQF0 | Q6ZQF0 | DNA topois | Topbp1 | sp\|Q6ZQF0 | 33 | 33 | 31 |
| Q8K3D3 | Q8K3D3 | DNA repair | Swi5 | sp\|Q8K3D3 | 5 | 5 | 44,9 |
| Q8VC65 | Q8VC65 | Nurim | Nrm | sp\|Q8VC65 | 7 | 7 | 28,6 |
| P97287 | P97287 | Induced m | Mcl1 | sp\|P97287 | 8 | 8 | 46,2 |
| A2A432 | A2A432 | Cullin-4B | Cul4b | sp\|A2A432 | 39 | 30 | 44,1 |
| Q5NC05 | Q5NC05 | Transcripti | Ttf2 | sp\|Q5NC05 | 13 | 13 | 17,3 |
| Q9D081 | Q9D081 | UDP-N-ace | Alg14 | sp\|Q9D081 | 3 | 3 | 18,9 |
| Q9CY62 | Q9CY62 | E3 ubiquiti | Rnf181 | sp\|Q9CY62 | 4 | 4 | 32,7 |
| Q9CQL5 | Q9CQL5 | 39S riboso | Mrpl18 | sp\|Q9CQL5 | 5 | 5 | 29,4 |
| Q3U2I3 | Q3U2I3 | FTS and Ho | Fam160a2 | sp\|Q3U2I3 | 6 | 6 | 8,1 |
| Q6P1Y8 | Q6P1Y8 | Type II ino | Inpp4b | sp\|Q6P1Y8 | 5 | 5 | 8 |
| Q9Z1Y4 | Q9Z1Y4 | Thyroid re | Trip6 | sp\|Q9Z1Y4 | 16 | 16 | 56,9 |
| Q9D219 | Q9D219 | B-cell CLL/ | Bcl9 | sp\|Q9D219 | 6 | 6 | 6,7 |
| P16951 | P16951 | Cyclic AMP | Atf2 | sp\|P16951 | 11 | 11 | 40 |
| Q91WA1 | Q91WA1 | TIMELESS-i | Tipin | sp\|Q91WA | 4 | 4 | 10,4 |
| A2AQ25 | A2AQ25 | Sickle tail p | Skt | sp\|A2AQ25 | 24 | 24 | 17,4 |
| O88856 | O88856 | Protein-tyr | Tpst2 | sp\|O88856 | 7 | 7 | 25 |

| | | | | | | | |
|---|---|---|---|---|---|---|---|
| Q8BSE0 | Q8BSE0 | Regulator | Rmdn2 | sp\|Q8BSE0 | 6 | 6 | 21 |
| P48410 | P48410 | ATP-bindin | Abcd1 | sp\|P48410 | 3 | 3 | 6 |
| P20067 | P20067 | DNA-bindi | Id1 | sp\|P20067 | 2 | 2 | 13,1 |
| Q8JZN5 | Q8JZN5 | Acyl-CoA d | Acad9 | sp\|Q8JZN5 | 31 | 31 | 53,4 |
| Q9DCE5 | Q9DCE5 | p21-activa | Pak1ip1 | sp\|Q9DCE | 18 | 18 | 63,4 |
| Q9DBZ1 | Q9DBZ1 | Inhibitor o | Ikbip | sp\|Q9DBZ | 10 | 10 | 35,1 |
| Q68FE8 | Q68FE8 | Zinc finger | Znf280d | sp\|Q68FE8 | 8 | 6 | 10,5 |
| Q8K3K7 | Q8K3K7 | 1-acyl-sn-g | Agpat2 | sp\|Q8K3K | 9 | 9 | 39,2 |
| Q6P5F6 | Q6P5F6 | Zinc transp | Slc39a10 | sp\|Q6P5F | 9 | 9 | 15,6 |
| Q99J23 | Q99J23 | GH3 doma | Ghdc | sp\|Q99J23 | 5 | 5 | 14,5 |
| Q80WQ6 | Q80WQ6 | Inactive rh | Rhbdf2 | sp\|Q80W | 6 | 6 | 11,5 |
| Q3TSG4 | Q3TSG4 | RNA deme | Alkbh5 | sp\|Q3TSG | 9 | 9 | 37 |
| Q6P069 | Q6P069 | Sorcin | Sri | sp\|Q6P06 | 10 | 10 | 55,1 |
| P29387 | P29387 | Guanine n | Gnb4 | sp\|P29387 | 8 | 3 | 36,2 |
| A2A8L5 | A2A8L5 | Receptor-t | Ptprf | sp\|A2A8L5 | 29 | 25 | 23,9 |
| Q6P3E7 | Q6P3E7 | Histone de | Hdac10 | sp\|Q6P3E7 | 3 | 3 | 5,9 |
| Q91XV3 | Q91XV3 | Brain acid | Basp1 | sp\|Q91XV | 15 | 15 | 81,4 |
| Q9D6K7 | Q9D6K7 | Tetratricop | Ttc33 | sp\|Q9D6K | 5 | 5 | 30,5 |
| Q6PGF3 | Q6PGF3 | Mediator d | Med16 | sp\|Q6PGF | 13 | 13 | 17,8 |
| A2A4P0 | A2A4P0 | ATP-depen | Dhx8 | sp\|A2A4P | 32 | 31 | 36,1 |
| Q91VU0 | Q91VU0 | Protein FA | Fam3c | sp\|Q91VU | 9 | 9 | 49,8 |
| Q99JP4 | Q99JP4 | Anaphase- | Cdc26 | sp\|Q99JP4 | 3 | 3 | 18,8 |
| CON__P12 | CON__P12763 | | | | 13 | 13 | 51 |
| O35609 | O35609 | Secretory | Scamp3 | sp\|O3560 | 7 | 7 | 31,2 |
| Q80TH2 | Q80TH2 | Protein LA | Erbb2ip | sp\|Q80TH | 31 | 30 | 31,7 |
| P61294 | P61294 | Ras-relate | Rab6b | sp\|P61294 | 11 | 4 | 57,2 |
| Q6ZPL9 | Q6ZPL9 | ATP-depen | Ddx55 | sp\|Q6ZPL9 | 8 | 8 | 18,5 |
| P97813 | P97813 | Phospholip | Pld2 | sp\|P97813 | 10 | 10 | 13,5 |
| Q8JZQ2 | Q8JZQ2 | AFG3-like | Afg3l2 | sp\|Q8JZQ2 | 26 | 24 | 38,7 |
| Q80UM3 | Q80UM3 | N-alpha-ac | Naa15 | sp\|Q80UM | 36 | 32 | 49,7 |
| Q99LP6 | Q99LP6 | GrpE prote | Grpel1 | sp\|Q99LP6 | 12 | 12 | 59,9 |
| P42859 | P42859 | Huntingtin | Htt | sp\|P42859 | 32 | 32 | 13,5 |
| Q8R4X3 | Q8R4X3 | RNA-bindi | Rbm12 | sp\|Q8R4X | 21 | 21 | 26 |
| Q8CIV8 | Q8CIV8 | Tubulin-sp | Tbce | sp\|Q8CIV8 | 11 | 11 | 28,2 |
| Q91V01 | Q91V01 | Lysophosp | Lpcat3 | sp\|Q91V01 | 4 | 4 | 10,5 |
| Q7TN37 | Q7TN37 | Transient | Trpm4 | sp\|Q7TN3 | 9 | 9 | 11,4 |
| Q8K3A0 | Q8K3A0 | Iron-sulfur | Hscb | sp\|Q8K3A | 6 | 6 | 29,9 |
| Q99M07 | Q99M07 | Cytochrom | Coa5 | sp\|Q99M0 | 2 | 2 | 37,8 |
| Q9CZJ1 | Q9CZJ1 | Probable U | Utp11l | sp\|Q9CZJ1 | 8 | 8 | 30 |
| O70310 | O70310 | Glycylpept | Nmt1 | sp\|O7031 | 16 | 12 | 37,3 |
| Q9D162 | Q9D162 | Coiled-coil | Ccdc167 | sp\|Q9D16 | 1 | 1 | 15,5 |
| Q8CG46 | Q8CG46 | Structural | Smc5 | sp\|Q8CG4 | 36 | 36 | 39,2 |
| Q3U487 | Q3U487 | E3 ubiquiti | Hectd3 | sp\|Q3U48 | 11 | 11 | 18,9 |
| Q04690 | Q04690 | Neurofibro | Nf1 | sp\|Q0469 | 34 | 34 | 13,6 |
| P61290 | P61290 | Proteasom | Psme3 | sp\|P61290 | 13 | 13 | 53,9 |
| O55003 | O55003 | BCL2/aden | Bnip3 | sp\|O5500 | 3 | 3 | 11,8 |
| Q8R0L1 | Q8R0L1 | Signal-tran | Stap2 | sp\|Q8R0L1 | 6 | 6 | 18,7 |
| Q9D787 | Q9D787 | Peptidyl-pr | Ppil2 | sp\|Q9D78 | 13 | 13 | 31,7 |
| Q922Y2 | Q922Y2 | Tripartite | Trim59 | sp\|Q922Y2 | 12 | 12 | 38,7 |
| D3Z7P3 | D3Z7P3 | Glutamina | Gls | sp\|D3Z7P3 | 28 | 28 | 50,7 |

| | | | | | | | |
|---|---|---|---|---|---|---|---|
| Q923K4 | Q923K4 | tRNA modi | Gtpbp3 | sp\|Q923K4 | 6 | 6 | 20,5 |
| Q9DB34 | Q9DB34 | Charged m | Chmp2a | sp\|Q9DB3 | 6 | 6 | 22,1 |
| Q811F1 | Q811F1 | Zinc finger | Zbtb41 | sp\|Q811F1 | 3 | 3 | 3,9 |
| Q8CG70 | Q8CG70 | Prolyl 3-hy | Leprel2 | sp\|Q8CG7 | 3 | 3 | 6,4 |
| Q5FWK3 | Q5FWK3 | Rho GTPas | Arhgap1 | sp\|Q5FWK | 16 | 16 | 49,4 |
| Q60862 | Q60862 | Origin reco | Orc2 | sp\|Q60862 | 12 | 12 | 22,6 |
| Q8BMA6 | Q8BMA6 | Signal reco | Srp68 | sp\|Q8BMA | 25 | 25 | 43,2 |
| Q91YP0 | Q91YP0 | L-2-hydrox | L2hgdh | sp\|Q91YP( | 16 | 16 | 39,2 |
| Q9CZP0 | Q9CZP0 | Ufm1-spec | Ufsp1 | sp\|Q9CZP( | 3 | 3 | 23 |
| Q60575 | Q60575 | Kinesin-like | Kif1b | sp\|Q60575 | 20 | 13 | 15,7 |
| Q3USZ8 | Q3USZ8 | Deleted in autism pro | | sp\|Q3USZ8 | 7 | 7 | 20,2 |
| P52293 | P52293 | Importin su | Kpna2 | sp\|P52293 | 24 | 24 | 56 |
| Q9CZX7 | Q9CZX7 | Type 2 pho | Tmem55a | sp\|Q9CZX7 | 8 | 7 | 47,1 |
| Q3TAE8 | Q3TAE8 | Dolichyl py | Alg6 | sp\|Q3TAE8 | 1 | 1 | 2,6 |
| Q9CWU9 | Q9CWU9 | Nucleopor | Nup37 | sp\|Q9CWU | 12 | 12 | 41,1 |
| Q9CQV4 | Q9CQV4 | Protein FA | Fam134c | sp\|Q9CQV | 6 | 6 | 18,5 |
| P15655 | P15655 | Fibroblast | Fgf2 | sp\|P15655 | 3 | 3 | 18,2 |
| O54781 | O54781 | SRSF prote | Srpk2 | sp\|O54781 | 14 | 11 | 28,3 |
| Q8K0Z7 | Q8K0Z7 | Translation | Taco1 | sp\|Q8K0Z7 | 8 | 8 | 38,1 |
| Q99L04 | Q99L04 | Dehydroge | Dhrs1 | sp\|Q99L04 | 14 | 14 | 46,6 |
| Q8C1Q6 | Q8C1Q6 | Small integ | Smim4 | sp\|Q8C1Q | 2 | 2 | 28,8 |
| Q8BNV1 | Q8BNV1 | tRNA (urac | Trmt2a | sp\|Q8BNV | 23 | 23 | 60,6 |
| Q8BHA3 | Q8BHA3 | Probable D | Dtd2 | sp\|Q8BHA | 8 | 8 | 57,7 |
| Q8K2I1 | Q8K2I1 | Protein far | Fntb | sp\|Q8K2I1 | 8 | 8 | 17,2 |
| Q9CZM2 | Q9CZM2 | 60S riboso | Rpl15 | sp\|Q9CZM | 14 | 14 | 50 |
| Q9QUG2 | Q9QUG2 | DNA polym | Polk | sp\|Q9QUG | 10 | 10 | 15,1 |
| Q505B7 | Q505B7 | Protein arc | Zbtb8os | sp\|Q505B7 | 2 | 2 | 16,1 |
| Q9D051 | Q9D051 | Pyruvate d | Pdhb | sp\|Q9D051 | 15 | 15 | 50,4 |
| Q62348 | Q62348 | Translin | Tsn | sp\|Q62348 | 10 | 10 | 42,5 |
| Q9D1Q4 | Q9D1Q4 | Dolichol-ph | Dpm3 | sp\|Q9D1Q | 2 | 2 | 23,9 |
| Q9D7W5 | Q9D7W5 | Mediator c | Med8 | sp\|Q9D7W | 5 | 5 | 29,1 |
| Q3URQ7 | Q3URQ7 | Methenylt | Mthfsd | sp\|Q3URQ | 3 | 3 | 12,6 |
| Q8K4Z5 | Q8K4Z5 | Splicing fac | Sf3a1 | sp\|Q8K4Z5 | 39 | 39 | 48,4 |
| P18760 | P18760 | Cofilin-1 | Cfl1 | sp\|P18760 | 23 | 21 | 80,1 |
| Q8K2D6 | Q8K2D6 | Deoxycytid | Dctd | sp\|Q8K2D | 5 | 5 | 32,6 |
| Q923X4 | Q923X4 | Glutaredox | Glrx2 | sp\|Q923X4 | 3 | 3 | 34 |
| Q9ES28;Q8 | Q9ES28 | Rho guanin | Arhgef7 | sp\|Q9ES28 | 26 | 26 | 37,2 |
| Q9JJZ4 | Q9JJZ4 | Ubiquitin-c | Ube2j1 | sp\|Q9JJZ4 | 8 | 8 | 34,3 |
| Q8CFI2 | Q8CFI2 | Ubiquitin-c | Cdc34 | sp\|Q8CFI2 | 7 | 7 | 22,6 |
| Q9CT10 | Q9CT10 | Ran-bindin | Ranbp3 | sp\|Q9CT1( | 18 | 18 | 53,4 |
| Q6DFX2 | Q6DFX2 | Anthrax to | Antxr2 | sp\|Q6DFX2 | 12 | 11 | 38,2 |
| Q9JJQ0 | Q9JJQ0 | GPI manno | Pigb | sp\|Q9JJQ0 | 7 | 7 | 16,6 |
| Q9CZD3 | Q9CZD3 | Glycine--tR | Gars | sp\|Q9CZD3 | 30 | 30 | 45,8 |
| Q8R1T1 | Q8R1T1 | Charged m | Chmp7 | sp\|Q8R1T1 | 3 | 3 | 10,2 |
| Q91YU6 | Q91YU6 | Leucine zip | Lzts2 | sp\|Q91YU( | 8 | 8 | 18,2 |
| Q91VX9 | Q91VX9 | Transmem | Tmem168 | sp\|Q91VX9 | 6 | 6 | 11 |
| O54750 | O54750 | Cytochrom | Cyp2j6 | sp\|O54750 | 3 | 3 | 8 |
| Q9D710 | Q9D710 | Thioredoxi | Tmx2 | sp\|Q9D71( | 8 | 8 | 30,2 |
| Q9CU62 | Q9CU62 | Structural | Smc1a | sp\|Q9CU6 | 54 | 54 | 48,7 |
| Q9JI39 | Q9JI39 | ATP-bindin | Abcb10 | sp\|Q9JI39 | 13 | 13 | 22,1 |

| | | | | | | | |
|---|---|---|---|---|---|---|---|
| Q3UK78 | Q3UK78 | Polycomb | Pcgf5 | sp\|Q3UK7 | 3 | 3 | 15,2 |
| Q9D1F4 | Q9D1F4 | Proline-ric | Akt1s1 | sp\|Q9D1F | 5 | 5 | 24,9 |
| P62962;C | P62962 | Profilin-1 | Pfn1 | sp\|P62962 | 13 | 13 | 91,4 |
| Q8C6G1 | Q8C6G1 | Protein C21orf2 hom | | sp\|Q8C6G | 2 | 2 | 11,2 |
| O35144 | O35144 | Telomeric | Terf2 | sp\|O3514 | 11 | 11 | 25,3 |
| Q8BVF2 | Q8BVF2 | Phosducin- | Pdcl3 | sp\|Q8BVF | 6 | 6 | 29,6 |
| Q9D8Y1 | Q9D8Y1 | Transmem | Tmem126 | sp\|Q9D8Y | 5 | 5 | 25 |
| Q6PD19 | Q6PD19 | UPF0668 protein C10 | | sp\|Q6PD1 | 7 | 7 | 13,4 |
| Q9DBQ7 | Q9DBQ7 | Protein-as | Scyl3 | sp\|Q9DBQ | 12 | 12 | 23,8 |
| Q9Z1Q9 | Q9Z1Q9 | Valine--tR | Vars | sp\|Q9Z1Q | 62 | 62 | 52,5 |
| Q9ES70 | Q9ES70 | Serine/thr | Nek6 | sp\|Q9ES7 | 2 | 1 | 7,7 |
| Q9JIK9 | Q9JIK9 | 28S riboso | Mrps34 | sp\|Q9JIK9 | 12 | 12 | 59,2 |
| Q99JX4 | Q99JX4 | Eukaryotic | Eif3m | sp\|Q99JX4 | 17 | 17 | 57 |
| Q60902 | Q60902 | Epidermal | Eps15l1 | sp\|Q6090 | 33 | 33 | 50,1 |
| A6H6E9 | A6H6E9 | Tetratrico | Ttc23l | sp\|A6H6E | 1 | 1 | 1,7 |
| Q9D7A8 | Q9D7A8 | Armadillo | Armc1 | sp\|Q9D7A | 12 | 12 | 60,3 |
| Q4VGL6 | Q4VGL6 | Roquin-1 | Rc3h1 | sp\|Q4VGL | 17 | 14 | 26,9 |
| Q8CFE6 | Q8CFE6 | Sodium-co | Slc38a2 | sp\|Q8CFE | 5 | 5 | 16,1 |
| O55023 | O55023 | Inositol m | Impa1 | sp\|O5502 | 14 | 14 | 46,6 |
| Q80UJ7 | Q80UJ7 | Rab3 GTPa | Rab3gap1 | sp\|Q80UJ | 22 | 22 | 28,7 |
| Q5SUQ9 | Q5SUQ9 | CST compl | Ctc1 | sp\|Q5SUQ | 23 | 23 | 28,7 |
| Q8CD26 | Q8CD26 | Solute car | Slc35e1 | sp\|Q8CD2 | 4 | 4 | 13,4 |
| Q8BZQ7 | Q8BZQ7 | Anaphase- | Anapc2 | sp\|Q8BZQ | 21 | 21 | 31,5 |
| P67871 | P67871 | Casein kin | Csnk2b | sp\|P67871 | 9 | 9 | 52,6 |
| P57780 | P57780 | Alpha-acti | Actn4 | sp\|P57780 | 62 | 41 | 71,2 |
| P27046 | P27046 | Alpha-man | Man2a1 | sp\|P27046 | 39 | 39 | 44,3 |
| Q91VY9 | Q91VY9 | Zinc finger | Znf622 | sp\|Q91VY | 14 | 14 | 40,1 |
| Q9DC29 | Q9DC29 | ATP-bindi | Abcb6 | sp\|Q9DC2 | 10 | 10 | 16,3 |
| Q9JLB2 | Q9JLB2 | MAGUK p5 | Mpp5 | sp\|Q9JLB2 | 15 | 15 | 34,1 |
| Q9CX48 | Q9CX48 | Zinc finger | Zcchc10 | sp\|Q9CX4 | 1 | 1 | 7,3 |
| Q8BM55 | Q8BM55 | Transmem | Tmem214 | sp\|Q8BM5 | 18 | 18 | 29,7 |
| Q8VEL9 | Q8VEL9 | GTP-bindi | Rem2 | sp\|Q8VEL | 7 | 7 | 27,9 |
| Q9WUP4 | Q9WUP4 | Polypreno | Srd5a3 | sp\|Q9WU | 3 | 3 | 10,6 |
| Q8BWR8 | Q8BWR8 | Rhophilin- | Rhpn2 | sp\|Q8BW | 6 | 6 | 9,8 |
| A2AR02 | A2AR02 | Peptidyl-p | Ppig | sp\|A2AR0 | 8 | 8 | 13,8 |
| Q8BJL1 | Q8BJL1 | F-box only | Fbxo30 | sp\|Q8BJL1 | 9 | 9 | 15 |
| Q3ULJ0;P1 | Q3ULJ0 | Glycerol-3- | Gpd1l | sp\|Q3ULJ | 14 | 14 | 57 |
| Q6PDL0 | Q6PDL0 | Cytoplasm | Dync1li2 | sp\|Q6PDL | 20 | 19 | 51,4 |
| Q8BWF0 | Q8BWF0 | Succinate- | Aldh5a1 | sp\|Q8BW | 9 | 9 | 27,5 |
| P61804 | P61804 | Dolichyl-d | Dad1 | sp\|P61804 | 3 | 3 | 28,3 |
| Q9R1S3 | Q9R1S3 | GPI ethano | Pign | sp\|Q9R1S | 9 | 9 | 11,8 |
| E9Q7E2 | E9Q7E2 | | Arid2 | sp\|E9Q7E | 30 | 30 | 24,5 |
| Q9JIF0;Q6 | Q9JIF0 | Protein arg | Prmt1 | sp\|Q9JIF0 | 23 | 23 | 58,5 |
| O35598 | O35598 | Disintegri | Adam10 | sp\|O3559 | 27 | 27 | 52,5 |
| Q9CQJ6 | Q9CQJ6 | Density-re | Denr | sp\|Q9CQJ | 8 | 8 | 47,5 |
| Q9QY81 | Q9QY81 | Nuclear po | Nup210 | sp\|Q9QY8 | 6 | 6 | 5,1 |
| Q3U2P1 | Q3U2P1 | Protein tra | Sec24a | sp\|Q3U2P | 17 | 17 | 21,7 |
| Q91X20 | Q91X20 | Set1/Ash2 | Ash2l | sp\|Q91X2 | 21 | 21 | 49,4 |
| Q8R0L9 | Q8R0L9 | Transcripti | Tada3 | sp\|Q8R0L | 8 | 8 | 21,3 |
| P99026 | P99026 | Proteasom | Psmb4 | sp\|P99026 | 10 | 10 | 59,8 |

| | | | | | | | |
|---|---|---|---|---|---|---|---|
| Q8VD65 | Q8VD65 | Phosphoin | Pik3r4 | sp\|Q8VD6 | 25 | 25 | 25,3 |
| Q8CFX1 | Q8CFX1 | GDH/6PGL | H6pd | sp\|Q8CFX1 | 16 | 16 | 27,1 |
| Q99PI5 | Q99PI5 | Phosphatic | Lpin2 | sp\|Q99PI5 | 6 | 6 | 9 |
| O88569 | O88569 | Heterogen | Hnrnpa2b1 | sp\|O8856 | 31 | 30 | 74,2 |
| Q58A65 | Q58A65 | C-Jun-amir | Spag9 | sp\|Q58A6 | 32 | 29 | 34,6 |
| Q9D1B9 | Q9D1B9 | 39S riboso | Mrpl28 | sp\|Q9D1B | 12 | 12 | 50,6 |
| O54879 | O54879 | High mobil | Hmgb3 | sp\|O5487 | 9 | 9 | 38,5 |
| Q8BGC0 | Q8BGC0 | HIV Tat-sp | Htatsf1 | sp\|Q8BGC | 20 | 20 | 39,1 |
| Q8BYH7 | Q8BYH7 | TBC1 doma | Tbc1d17 | sp\|Q8BYH | 16 | 16 | 35,3 |
| Q6A068 | Q6A068 | Cell divisio | Cdc5l | sp\|Q6A06 | 34 | 34 | 54,9 |
| P29416 | P29416 | Beta-hexos | Hexa | sp\|P29416 | 11 | 11 | 23,7 |
| Q9CX00 | Q9CX00 | IST1 homo | Ist1 | sp\|Q9CX0 | 9 | 9 | 26 |
| Q922P9 | Q922P9 | Putative o | Glyr1 | sp\|Q922P | 19 | 19 | 49,1 |
| Q8BVA5 | Q8BVA5 | UPF0554 protein C2or | | sp\|Q8BVA | 13 | 13 | 35,9 |
| Q9QXJ1 | Q9QXJ1 | Amyloid be | Apbb1 | sp\|Q9QXJ1 | 7 | 6 | 17,2 |
| O88895 | O88895 | Histone de | Hdac3 | sp\|O8889 | 6 | 5 | 18,2 |
| Q9D168 | Q9D168 | Integrator | Ints12 | sp\|Q9D16 | 13 | 13 | 39,7 |
| Q9JJ78 | Q9JJ78 | Lymphokin | Pbk | sp\|Q9JJ78 | 13 | 13 | 52,4 |
| Q9CR11 | Q9CR11 | YEATS dom | Yeats4 | sp\|Q9CR1 | 6 | 6 | 30,8 |
| Q5ND52 | Q5ND52 | rRNA meth | Rnmtl1 | sp\|Q5ND5 | 6 | 6 | 16,5 |
| Q6PE54 | Q6PE54 | Probable A | Dhx40 | sp\|Q6PE54 | 9 | 9 | 15 |
| Q9CR59 | Q9CR59 | Growth arr | Gadd45gip | sp\|Q9CR5 | 5 | 5 | 36 |
| P62996 | P62996 | Transform | Tra2b | sp\|P62996 | 9 | 8 | 37,2 |
| P09803 | P09803 | Cadherin-1 | Cdh1 | sp\|P09803 | 20 | 20 | 37,4 |
| P01902;P1 | P01902 | H-2 class I | H2-K1 | sp\|P01902 | 16 | 11 | 46,2 |
| P10417 | P10417 | Apoptosis | Bcl2 | sp\|P10417 | 2 | 2 | 11,4 |
| Q9DB00 | Q9DB00 | GON-4-like | Gon4l | sp\|Q9DB0 | 11 | 11 | 8,5 |
| P61961 | P61961 | Ubiquitin-f | Ufm1 | sp\|P61961 | 5 | 5 | 68,2 |
| Q9D6J6 | Q9D6J6 | NADH deh | Ndufv2 | sp\|Q9D6J6 | 12 | 12 | 53,2 |
| Q9D0R4 | Q9D0R4 | Probable A | Ddx56 | sp\|Q9D0R | 16 | 16 | 36,8 |
| Q9CQ54 | Q9CQ54 | NADH deh | Ndufc2 | sp\|Q9CQ5 | 5 | 5 | 31,7 |
| Q9CPY6 | Q9CPY6 | Glucose-in | Gid4 | sp\|Q9CPY | 4 | 4 | 30,4 |
| P23780 | P23780 | Beta-galac | Glb1 | sp\|P23780 | 15 | 14 | 27,7 |
| B8ZXI1 | B8ZXI1 | Queuine t | Qtrtd1 | sp\|B8ZXI1 | 10 | 10 | 32,3 |
| Q9ER10 | Q9ER10 | Brain-spec | Prss22 | sp\|Q9ER1 | 5 | 5 | 21,2 |
| Q8K2Y7 | Q8K2Y7 | 39S riboso | Mrpl47 | sp\|Q8K2Y | 10 | 10 | 34,5 |
| Q8R480 | Q8R480 | Nuclear po | Nup85 | sp\|Q8R48 | 22 | 22 | 37,8 |
| Q9CUX1 | Q9CUX1 | 52 kDa rep | Prkrir | sp\|Q9CUX | 11 | 11 | 22,7 |
| Q61823 | Q61823 | Programm | Pdcd4 | sp\|Q6182 | 9 | 9 | 31,6 |
| Q9D0A3 | Q9D0A3 | Arpin | Arpin | sp\|Q9D0A | 10 | 10 | 46,5 |
| P50580 | P50580 | Proliferati | Pa2g4 | sp\|P50580 | 26 | 26 | 69,8 |
| Q91Z96 | Q91Z96 | BMP-2-ind | Bmp2k | sp\|Q91Z9 | 18 | 17 | 26,4 |
| P25425;P3 | P25425 | POU doma | Pou2f1 | sp\|P25425 | 7 | 7 | 16,9 |
| P97434 | P97434 | Myosin ph | Mprip | sp\|P97434 | 42 | 42 | 58,2 |
| Q14C51 | Q14C51 | Pentatrico | Ptcd3 | sp\|Q14C5 | 22 | 22 | 43,6 |
| Q9D0M2 | Q9D0M2 | Cell divisio | Cdca7 | sp\|Q9D0M | 6 | 5 | 27 |
| Q99KW9 | Q99KW9 | T-cell imm | Itfg1 | sp\|Q99KW | 7 | 7 | 13,3 |
| P36552 | P36552 | Oxygen-de | Cpox | sp\|P36552 | 15 | 15 | 36,3 |
| P46460 | P46460 | Vesicle-fus | Nsf | sp\|P46460 | 36 | 36 | 55,1 |
| Q9D1P0 | Q9D1P0 | 39S riboso | Mrpl13 | sp\|Q9D1P | 8 | 8 | 50 |

| | | | | | | | |
|---|---|---|---|---|---|---|---|
| Q8BTV2 | Q8BTV2 | Cleavage a | Cpsf7 | sp\|Q8BTV2 | 9 | 9 | 21 |
| Q9CPR7 | Q9CPR7 | Suppressor | Sike1 | sp\|Q9CPR7 | 7 | 7 | 42 |
| Q6NWW9 | Q6NWW9 | Fibronectin | Fndc3b | sp\|Q6NWW | 26 | 26 | 34,5 |
| P27577 | P27577 | Protein C-e | Ets1 | sp\|P27577 | 2 | 2 | 5,9 |
| Q8BG67 | Q8BG67 | Protein EFI | Efr3a | sp\|Q8BG67 | 16 | 15 | 27,1 |
| Q8R4H2 | Q8R4H2 | Rho guanin | Arhgef12 | sp\|Q8R4H2 | 24 | 24 | 20,9 |
| B1AVZ0 | B1AVZ0 | Uracil phos | Uprt | sp\|B1AVZ0 | 6 | 6 | 33,9 |
| Q9D361 | Q9D361 | U11/U12 s | Snrnp48 | sp\|Q9D361 | 2 | 2 | 3,9 |
| Q9D7E4 | Q9D7E4 | UPF0449 protein C19o | | sp\|Q9D7E4 | 4 | 4 | 66,1 |
| Q9CYR0 | Q9CYR0 | Single-stra | Ssbp1 | sp\|Q9CYR0 | 6 | 6 | 38,8 |
| O35465 | O35465 | Peptidyl-pr | Fkbp8 | sp\|O35465 | 10 | 10 | 36,1 |
| Q9CWL8 | Q9CWL8 | Beta-caten | Ctnnbl1 | sp\|Q9CWL | 20 | 20 | 34,8 |
| Q7TSC1 | Q7TSC1 | Protein PR | Prrc2a | sp\|Q7TSC1 | 48 | 46 | 35,4 |
| Q9CX86 | Q9CX86 | Heterogen | Hnrnpa0 | sp\|Q9CX86 | 13 | 13 | 51,8 |
| Q9ERE7 | Q9ERE7 | LDLR chape | Mesdc2 | sp\|Q9ERE7 | 11 | 11 | 52,7 |
| Q9EQS3 | Q9EQS3 | C-Myc-bind | Mycbp | sp\|Q9EQS3 | 5 | 5 | 62,1 |
| O70551;Q9 | O70551 | SRSF prote | Srpk1 | sp\|O70551 | 17 | 14 | 39,5 |
| Q9CQM9 | Q9CQM9 | Glutaredox | Glrx3 | sp\|Q9CQM | 17 | 17 | 60,5 |
| P54116 | P54116 | Erythrocyt | Stom | sp\|P54116 | 8 | 8 | 36,6 |
| Q8CFQ3 | Q8CFQ3 | Intron-bind | Aqr | sp\|Q8CFQ3 | 44 | 44 | 42,4 |
| Q64704 | Q64704 | Syntaxin-3 | Stx3 | sp\|Q64704 | 2 | 2 | 9,3 |
| Q99JB2 | Q99JB2 | Stomatin-li | Stoml2 | sp\|Q99JB2 | 14 | 14 | 49,3 |
| Q9EPK2 | Q9EPK2 | Protein XR | Rp2 | sp\|Q9EPK2 | 4 | 4 | 15,3 |
| O35963;P9 | O35963 | Ras-related | Rab33b | sp\|O35963 | 5 | 5 | 28,8 |
| Q924C6 | Q924C6 | Lysyl oxida | Loxl4 | sp\|Q924C6 | 11 | 11 | 23,9 |
| Q9DB16 | Q9DB16 | Calcium-bi | Cab39l | sp\|Q9DB16 | 8 | 6 | 25,5 |
| Q8QZR5 | Q8QZR5 | Alanine am | Gpt | sp\|Q8QZR5 | 7 | 7 | 16,1 |
| Q9JMD3 | Q9JMD3 | PCTP-like p | Stard10 | sp\|Q9JMD | 4 | 4 | 20,3 |
| Q8K012 | Q8K012 | Formin-bin | Fnbp1l | sp\|Q8K012 | 15 | 15 | 29,6 |
| Q7TMK9 | Q7TMK9 | Heterogen | Syncrip | sp\|Q7TMK | 33 | 33 | 59,6 |
| P26883 | P26883 | Peptidyl-pr | Fkbp1a | sp\|P26883 | 6 | 6 | 41,7 |
| Q8CHP6 | Q8CHP6 | Polyhomeo | Phc3 | sp\|Q8CHP6 | 5 | 4 | 10,3 |
| P70268 | P70268 | Serine/thre | Pkn1 | sp\|P70268 | 16 | 15 | 22,5 |
| O35516 | O35516 | Neurogeni | Notch2 | sp\|O35516 | 26 | 25 | 16,4 |
| Q9CPQ8 | Q9CPQ8 | ATP syntha | Atp5l | sp\|Q9CPQ | 4 | 4 | 41,7 |
| Q06649 | Q06649 | SH3 domai | Sh3bp2 | sp\|Q06649 | 4 | 4 | 11,1 |
| Q80X73 | Q80X73 | Protein pe | Pelo | sp\|Q80X73 | 8 | 8 | 23,9 |
| Q8BJU0 | Q8BJU0 | Small gluta | Sgta | sp\|Q8BJU0 | 8 | 8 | 27,6 |
| P59326 | P59326 | YTH domai | Ythdf1 | sp\|P59326 | 11 | 7 | 27,7 |
| Q9CWJ9 | Q9CWJ9 | Bifunctiona | Atic | sp\|Q9CWJ | 38 | 38 | 77,9 |
| Q9DBY0 | Q9DBY0 | Forkhead b | Foxp4 | sp\|Q9DBY0 | 9 | 9 | 26,2 |
| Q6A078 | Q6A078 | Centrosom | Cep290 | sp\|Q6A078 | 6 | 6 | 2,3 |
| Q9QZQ1 | Q9QZQ1 | Afadin | Mllt4 | sp\|Q9QZQ | 55 | 55 | 38,4 |
| O88396 | O88396 | GrpE prote | Grpel2 | sp\|O88396 | 3 | 3 | 15,2 |
| Q8BK64 | Q8BK64 | Activator o | Ahsa1 | sp\|Q8BK64 | 17 | 17 | 61,8 |
| Q5XG71 | Q5XG71 | Small subu | Utp20 | sp\|Q5XG71 | 86 | 86 | 37,3 |
| Q61733 | Q61733 | 28S riboso | Mrps31 | sp\|Q61733 | 12 | 12 | 48,2 |
| Q9JHE7 | Q9JHE7 | Protein TSS | Tssc4 | sp\|Q9JHE7 | 6 | 6 | 29 |
| Q63943 | Q63943 | Myocyte-s | Mef2d | sp\|Q63943 | 5 | 4 | 14 |
| Q9JKX4 | Q9JKX4 | Protein AA | Aatf | sp\|Q9JKX4 | 23 | 23 | 51,7 |

| | | | | | | | |
|---|---|---|---|---|---|---|---|
| Q3V009 | Q3V009 | Transmem | Tmed1 | sp\|Q3V009 | 4 | 4 | 22,5 |
| O08586 | O08586 | Phosphatid | Pten | sp\|O08586 | 4 | 4 | 11,4 |
| Q9Z0P4 | Q9Z0P4 | Paralemmi | Palm | sp\|Q9Z0P4 | 9 | 9 | 39,2 |
| Q9DBE8 | Q9DBE8 | Alpha-1,3/ | Alg2 | sp\|Q9DBE8 | 14 | 14 | 47,7 |
| Q9D6U8 | Q9D6U8 | Protein FA | Fam162a | sp\|Q9D6U | 7 | 7 | 33,5 |
| Q9CRB9 | Q9CRB9 | MICOS con | Chchd3 | sp\|Q9CRB9 | 11 | 11 | 33 |
| Q63810 | Q63810 | Calcineurin | Ppp3r1 | sp\|Q63810 | 7 | 7 | 61,2 |
| Q91Z53 | Q91Z53 | Glyoxylate | Grhpr | sp\|Q91Z53 | 13 | 13 | 54,9 |
| Q8C3X2 | Q8C3X2 | Coiled-coil | Ccdc90b | sp\|Q8C3X2 | 9 | 9 | 34 |
| Q8BIZ6 | Q8BIZ6 | Smad nucle | Snip1 | sp\|Q8BIZ6 | 5 | 5 | 21,7 |
| Q6PAQ4 | Q6PAQ4 | RNA exonu | Rexo4 | sp\|Q6PAQ | 15 | 15 | 41,9 |
| Q9WTP7 | Q9WTP7 | GTP:AMP p | Ak3 | sp\|Q9WTP | 11 | 11 | 52,9 |
| Q99K23 | Q99K23 | Ufm1-spec | Ufsp2 | sp\|Q99K23 | 17 | 17 | 52,5 |
| P18052 | P18052 | Receptor-t | Ptpra | sp\|P18052 | 16 | 16 | 26,7 |
| P10518 | P10518 | Delta-amin | Alad | sp\|P10518 | 15 | 15 | 58,8 |
| P35761 | P35761 | Dual specif | Ttk | sp\|P35761 | 19 | 19 | 32,6 |
| Q61316 | Q61316 | Heat shock | Hspa4 | sp\|Q61316 | 56 | 53 | 75,9 |
| P52825 | P52825 | Carnitine O | Cpt2 | sp\|P52825 | 19 | 19 | 40,9 |
| Q99PV0 | Q99PV0 | Pre-mRNA- | Prpf8 | sp\|Q99PV0 | 114 | 114 | 57,4 |
| Q9D853 | Q9D853 | Protein-lys | Mettl10 | sp\|Q9D853 | 3 | 3 | 14,3 |
| Q99N84 | Q99N84 | 28S riboso | Mrps18b | sp\|Q99N84 | 6 | 6 | 26,8 |
| Q00422 | Q00422 | GA-binding | Gabpa | sp\|Q00422 | 18 | 18 | 50,9 |
| Q9JJW6 | Q9JJW6 | Aly/REF ex | Alyref2 | sp\|Q9JJW6 | 7 | 3 | 51,4 |
| Q64729 | Q64729 | TGF-beta r | Tgfbr1 | sp\|Q64729 | 6 | 6 | 17,5 |
| P43406 | P43406 | Integrin alp | Itgav | sp\|P43406 | 31 | 31 | 35,8 |
| Q99LV7 | Q99LV7 | Phosphatid | Pigx | sp\|Q99LV7 | 2 | 2 | 11,8 |
| O08750 | O08750 | Nuclear fac | Nfil3 | sp\|O08750 | 5 | 5 | 13,9 |
| Q8CA72 | Q8CA72 | Gigaxonin | Gan | sp\|Q8CA72 | 10 | 10 | 23,6 |
| Q91X43 | Q91X43 | SH3 domai | Sh3d19 | sp\|Q91X43 | 11 | 11 | 19,3 |
| Q8BLB7 | Q8BLB7 | Lethal(3)m | L3mbtl3 | sp\|Q8BLB7 | 15 | 15 | 30,9 |
| CON__Q9T | CON__Q9TT36 | | | | 2 | 2 | 6,8 |
| Q80WJ7 | Q80WJ7 | Protein LYR | Mtdh | sp\|Q80WJ | 26 | 26 | 52,8 |
| P58137 | P58137 | Acyl-coenz | Acot8 | sp\|P58137 | 7 | 7 | 30,9 |
| Q8C863 | Q8C863 | E3 ubiquiti | Itch | sp\|Q8C863 | 20 | 18 | 31,1 |
| Q6PE15 | Q6PE15 | Mycophen | Abhd10 | sp\|Q6PE15 | 8 | 8 | 42,4 |
| Q6NZL6 | Q6NZL6 | Tonsoku-lil | Tonsl | sp\|Q6NZL6 | 14 | 14 | 14,6 |
| Q6XUX1 | Q6XUX1 | Dual serine | Dstyk | sp\|Q6XUX | 2 | 2 | 3,2 |
| P49717 | P49717 | DNA replic | Mcm4 | sp\|P49717 | 43 | 42 | 55,6 |
| Q9D0I9 | Q9D0I9 | Arginine--t | Rars | sp\|Q9D0I9 | 46 | 46 | 61,8 |
| Q8R1F5 | Q8R1F5 | Putative hy | Hyi | sp\|Q8R1F5 | 6 | 6 | 33,6 |
| P49446 | P49446 | Receptor-t | Ptpre | sp\|P49446 | 19 | 19 | 32,6 |
| Q6ZPS6 | Q6ZPS6 | Ankyrin re | Ankib1 | sp\|Q6ZPS6 | 4 | 4 | 6 |
| O88712 | O88712 | C-terminal | Ctbp1 | sp\|O88712 | 15 | 11 | 44,7 |
| Q9CR50 | Q9CR50 | RING finge | Rchy1 | sp\|Q9CR50 | 2 | 2 | 9,6 |
| Q8VCM4 | Q8VCM4 | Lipoyltrans | Lipt1 | sp\|Q8VCM | 3 | 3 | 10,5 |
| Q8R3F9 | Q8R3F9 | Speckle tar | Tut1 | sp\|Q8R3F9 | 13 | 13 | 27,6 |
| CON__P00 | CON__P00761 | | | | 10 | 10 | 54,5 |
| O54962 | O54962 | Barrier-to- | Banf1 | sp\|O54962 | 7 | 7 | 66,3 |
| Q3UVL4 | Q3UVL4 | Vacuolar p | Vps51 | sp\|Q3UVL4 | 20 | 20 | 34,3 |
| Q9D479 | Q9D479 | UV-stimula | Uvssa | sp\|Q9D479 | 4 | 4 | 7,3 |

| | | | | | | | |
|---|---|---|---|---|---|---|---|
| P45591 | P45591 | Cofilin-2 | Cfl2 | sp\|P45591 | 13 | 11 | 65,7 |
| Q8C6C7 | Q8C6C7 | Protein FA | Fam204a | sp\|Q8C6C7 | 2 | 2 | 22,5 |
| Q9DBL1 | Q9DBL1 | Short/bran | Acadsb | sp\|Q9DBL1 | 15 | 15 | 46,5 |
| Q8R2U0 | Q8R2U0 | Nucleopor | Seh1l | sp\|Q8R2U( | 16 | 16 | 62,2 |
| Q3UZ01 | Q3UZ01 | RNA-bindir | Rnpc3 | sp\|Q3UZ01 | 9 | 9 | 25,5 |
| Q99LJ1 | Q99LJ1 | Tissue alph | Fuca1 | sp\|Q99LJ1 | 1 | 1 | 2,4 |
| Q9CQK1 | Q9CQK1 | Zinc finger | Znhit3 | sp\|Q9CQK | 4 | 4 | 25,8 |
| D3YZP9 | D3YZP9 | Coiled-coil | Ccdc6 | sp\|D3YZP9 | 15 | 15 | 40,3 |
| O88286 | O88286 | Protein Wi | Wiz | sp\|O88286 | 28 | 28 | 23,6 |
| Q8VED9 | Q8VED9 | Galectin-re | Lgalsl | sp\|Q8VED9 | 7 | 7 | 56,4 |
| Q5U431 | Q5U431 | G-protein c | Gpr39 | sp\|Q5U431 | 2 | 2 | 5 |
| Q3TXX3 | Q3TXX3 | Protrudin | Zfyve27 | sp\|Q3TXX3 | 5 | 5 | 19,3 |
| P26011 | P26011 | Integrin be | Itgb7 | sp\|P26011 | 4 | 4 | 6,3 |
| Q9D1L0 | Q9D1L0 | Coiled-coil | Chchd2 | sp\|Q9D1L( | 4 | 4 | 30,7 |
| Q9WTK3 | Q9WTK3 | Glycosylph | Gpaa1 | sp\|Q9WTK | 8 | 8 | 13,8 |
| P27546 | P27546 | Microtubu | Map4 | sp\|P27546 | 40 | 40 | 54 |
| Q04887;Q( | Q04887;Q048 | Transcripti | Sox9;Sox8; | sp\|Q04887 | 2 | 2 | 5,3 |
| Q8C3P7 | Q8C3P7 | N6-adenos | Mettl3 | sp\|Q8C3P7 | 20 | 20 | 44,7 |
| Q8BGD9 | Q8BGD9 | Eukaryotic | Eif4b | sp\|Q8BGD | 24 | 24 | 46 |
| Q8BQ33 | Q8BQ33 | Treslin | Ticrr | sp\|Q8BQ3 | 6 | 6 | 4,8 |
| Q9CXS4 | Q9CXS4 | Centromer | Cenpv | sp\|Q9CXS4 | 11 | 11 | 54 |
| P51174 | P51174 | Long-chain | Acadl | sp\|P51174 | 20 | 20 | 47 |
| O08788 | O08788 | Dynactin s | Dctn1 | sp\|O08788 | 42 | 42 | 44,3 |
| Q8VEB4 | Q8VEB4 | Group XV p | Pla2g15 | sp\|Q8VEB4 | 7 | 7 | 21,8 |
| Q64311 | Q64311 | Protein N-t | Ntan1 | sp\|Q64311 | 10 | 10 | 41,9 |
| P49135 | P49135 | TFIIH basal | Ercc3 | sp\|P49135 | 19 | 19 | 33,8 |
| Q922E4 | Q922E4 | Ethanolam | Pcyt2 | sp\|Q922E4 | 5 | 5 | 14,6 |
| B9EKI3 | B9EKI3 | TATA elem | Tmf1 | sp\|B9EKI3 | 19 | 19 | 24,5 |
| Q61146 | Q61146 | Occludin | Ocln | sp\|Q61146 | 4 | 4 | 11,1 |
| Q6PFD6 | Q6PFD6 | Kinesin-like | Kif18b | sp\|Q6PFD6 | 19 | 18 | 35,1 |
| Q8VEB3 | Q8VEB3 | UNC119-bi | D1Ertd622 | sp\|Q8VEB3 | 3 | 3 | 20,8 |
| P58252 | P58252 | Elongation | Eef2 | sp\|P58252 | 61 | 60 | 76,3 |
| Q4PZA2 | Q4PZA2 | Endothelin | Ece1 | sp\|Q4PZA2 | 14 | 14 | 24,3 |
| Q01514 | Q01514 | Interferon- | Gbp1 | sp\|Q01514 | 7 | 6 | 14,8 |
| Q9WUI1 | Q9WUI1 | Mitogen-a | Mapk11 | sp\|Q9WUI | 6 | 6 | 24,7 |
| O08600 | O08600 | Endonucle | Endog | sp\|O08600 | 6 | 6 | 25,2 |
| Q6ZQ88 | Q6ZQ88 | Lysine-spe | Kdm1a | sp\|Q6ZQ88 | 25 | 25 | 40,6 |
| Q64127 | Q64127 | Transcripti | Trim24 | sp\|Q64127 | 15 | 15 | 22,4 |
| Q61035 | Q61035 | Histidine--t | Hars | sp\|Q61035 | 23 | 20 | 53,4 |
| Q8VBZ0 | Q8VBZ0 | Dehydroge | Dhrsx | sp\|Q8VBZ( | 6 | 6 | 32,5 |
| Q9WVA3 | Q9WVA3 | Mitotic che | Bub3 | sp\|Q9WVA | 11 | 11 | 42 |
| Q9DAK9 | Q9DAK9 | 14 kDa pho | Phpt1 | sp\|Q9DAK | 6 | 6 | 55,6 |
| CON__Q9N | CON__Q9N2I2 | | | | 5 | 5 | 18,6 |
| P83741;Q8 | P83741 | Serine/thre | Wnk1 | sp\|P83741 | 30 | 27 | 17,6 |
| Q9JLI6 | Q9JLI6 | Selenocyst | Scly | sp\|Q9JLI6 \| | 11 | 11 | 32,9 |
| Q9CQC6 | Q9CQC6 | Basic leuci | Bzw1 | sp\|Q9CQC | 16 | 15 | 39,9 |
| Q8CBY0 | Q8CBY0 | Glutamyl-t | Gatc | sp\|Q8CBY( | 2 | 2 | 23,9 |
| Q91VR2 | Q91VR2 | ATP syntha | Atp5c1 | sp\|Q91VR2 | 11 | 11 | 34,6 |
| P68368 | P68368 | Tubulin alp | Tuba4a | sp\|P68368 | 26 | 5 | 62,7 |
| Q99JZ7 | Q99JZ7 | ERBB recep | Errfi1 | sp\|Q99JZ7 | 6 | 6 | 26 |

| | | | | | | | |
|---|---|---|---|---|---|---|---|
| Q9EPU4 | Q9EPU4 | Cleavage a | Cpsf1 | sp\|Q9EPU4 | 40 | 40 | 35,2 |
| P41216 | P41216 | Long-chain | Acsl1 | sp\|P41216 | 24 | 21 | 43,3 |
| Q8VDQ8 | Q8VDQ8 | NAD-deper | Sirt2 | sp\|Q8VDQ | 9 | 9 | 30,8 |
| Q5U3K5 | Q5U3K5 | Rab-like pr | Rabl6 | sp\|Q5U3K | 12 | 12 | 25,2 |
| Q6PE01 | Q6PE01 | U5 small n | Snrnp40 | sp\|Q6PE01 | 16 | 16 | 67,3 |
| Q8BMG1 | Q8BMG1 | ATR-intera | Atrip | sp\|Q8BMG | 6 | 6 | 14 |
| O89023 | O89023 | Tripeptidyl | Tpp1 | sp\|O89023 | 5 | 5 | 14,2 |
| P58044 | P58044 | Isopenteny | Idi1 | sp\|P58044 | 9 | 9 | 58,1 |
| Q03347 | Q03347 | Runt-relate | Runx1 | sp\|Q03347 | 12 | 12 | 39,5 |
| Q6PDJ6 | Q6PDJ6 | F-box only | Fbxo42 | sp\|Q6PDJ6 | 4 | 4 | 9,1 |
| Q8BYH8 | Q8BYH8 | Chromodo | Chd9 | sp\|Q8BYH | 14 | 6 | 6,2 |
| P17918 | P17918 | Proliferatir | Pcna | sp\|P17918 | 14 | 14 | 72,8 |
| Q80U63 | Q80U63 | Mitofusin-: | Mfn2 | sp\|Q80U6 | 21 | 19 | 32,1 |
| Q5SXA9 | Q5SXA9 | Protein KIE | Wwc1 | sp\|Q5SXA | 4 | 4 | 6,5 |
| Q8K304 | Q8K304 | E3 ubiquiti | Tmem129 | sp\|Q8K304 | 4 | 4 | 15,5 |
| Q9D023 | Q9D023 | Mitochond | Mpc2 | sp\|Q9D02 | 6 | 5 | 42,5 |
| P61089 | P61089 | Ubiquitin-c | Ube2n | sp\|P61089 | 10 | 9 | 77,6 |
| Q7M759 | Q7M759 | Alpha/beta | Abhd17b | sp\|Q7M75 | 8 | 8 | 49 |
| Q99PT1 | Q99PT1 | Rho GDP-d | Arhgdia | sp\|Q99PT1 | 13 | 13 | 74,5 |
| Q8BM39 | Q8BM39 | Pre-mRNA- | Prpf18 | sp\|Q8BM3 | 9 | 9 | 37,4 |
| Q9QZ73 | Q9QZ73 | DCN1-like | Dcun1d1 | sp\|Q9QZ7 | 5 | 5 | 24,7 |
| Q6P3B9 | Q6P3B9 | Putative ri | Rbfa | sp\|Q6P3B | 5 | 5 | 24,6 |
| Q9D7M1 | Q9D7M1 | Glucose-in | Gid8 | sp\|Q9D7M | 12 | 12 | 64,9 |
| Q8BP48 | Q8BP48 | Methionin | Metap1 | sp\|Q8BP4 | 18 | 18 | 69,2 |
| Q3TXU5 | Q3TXU5 | Deoxyhypu | Dhps | sp\|Q3TXU | 8 | 8 | 26,3 |
| Q91XC0 | Q91XC0 | LIM domai | Ajuba | sp\|Q91XC | 3 | 3 | 10,8 |
| Q8K310 | Q8K310 | Matrin-3 | Matr3 | sp\|Q8K31 | 38 | 38 | 43,9 |
| Q6RT24 | Q6RT24 | Centromer | Cenpe | sp\|Q6RT24 | 39 | 39 | 22 |
| P41158 | P41158 | ETS domai | Elk4 | sp\|P41158 | 1 | 1 | 4 |
| Q810B6 | Q810B6 | Rabankyrir | Ankfy1 | sp\|Q810B | 34 | 34 | 40,5 |
| P62301 | P62301 | 40S riboso | Rps13 | sp\|P62301 | 15 | 15 | 62,3 |
| P51855 | P51855 | Glutathion | Gss | sp\|P51855 | 13 | 13 | 36,3 |
| Q99JN2 | Q99JN2 | Kelch-like | Klhl22 | sp\|Q99JN2 | 15 | 15 | 32 |
| E9Q557 | E9Q557 | Desmoplak | Dsp | sp\|E9Q557 | 43 | 43 | 20 |
| Q9WTM5 | Q9WTM5 | RuvB-like 2 | Ruvbl2 | sp\|Q9WTN | 29 | 29 | 71,9 |
| Q99LJ0 | Q99LJ0 | CTTNBP2 N | Cttnbp2nl | sp\|Q99LJ0 | 22 | 22 | 45,1 |
| Q7TPE5 | Q7TPE5 | Probable R | Slc7a6os | sp\|Q7TPE5 | 4 | 4 | 16,3 |
| P80316 | P80316 | T-complex | Cct5 | sp\|P80316 | 40 | 40 | 80,4 |
| Q9CYA0 | Q9CYA0 | Cysteine-ri | Creld2 | sp\|Q9CYA | 11 | 11 | 43,1 |
| Q99PL6 | Q99PL6 | UBX doma | Ubxn6 | sp\|Q99PL6 | 16 | 16 | 51,6 |
| Q8CIE6 | Q8CIE6 | Coatomer | Copa | sp\|Q8CIE6 | 62 | 62 | 57,8 |
| Q91YL3 | Q91YL3 | Uridine-cyt | Uckl1 | sp\|Q91YL3 | 6 | 6 | 14,4 |
| Q91WG8 | Q91WG8 | Bifunction | Gne | sp\|Q91WG | 19 | 19 | 34,5 |
| P05201 | P05201 | Aspartate | Got1 | sp\|P05201 | 18 | 18 | 54,2 |
| Q9EQX4 | Q9EQX4 | Allograft ir | Aif1l | sp\|Q9EQX | 4 | 4 | 26,7 |
| Q9CR47 | Q9CR47 | Ribosome | Nsa2 | sp\|Q9CR4 | 11 | 11 | 43,1 |
| Q9WVK0 | Q9WVK0 | Type-1 ang | Agtrap | sp\|Q9WVK | 2 | 2 | 30,4 |
| Q91YN1 | Q91YN1 | Protein FA | Fam118a | sp\|Q91YN | 4 | 4 | 12,9 |
| Q3U3R4 | Q3U3R4 | Lipase mat | Lmf1 | sp\|Q3U3R | 5 | 5 | 12 |
| Q9WVM3 | Q9WVM3 | Anaphase- | Anapc7 | sp\|Q9WVN | 19 | 19 | 43,4 |

| | | | | | | | |
|---|---|---|---|---|---|---|---|
| Q8BR90 | Q8BR90 | UPF0600 protein C5or | | sp\|Q8BR9( | 6 | 6 | 32,3 |
| P46471 | P46471 | 26S protea | Psmc2 | sp\|P46471 | 35 | 35 | 73,7 |
| Q61457;Q9 | Q61457;Q9Z2: | G1/S-speci | Ccne1;Ccn | sp\|Q61457 | 2 | 2 | 5,9 |
| Q9JM05 | Q9JM05 | E3 SUMO-p | Pias4 | sp\|Q9JM0 | 4 | 4 | 11,8 |
| Q9CPX6 | Q9CPX6 | Ubiquitin-l | Atg3 | sp\|Q9CPX6 | 7 | 7 | 28 |
| Q3UHX0 | Q3UHX0 | Nucleolar p | Nol8 | sp\|Q3UHX | 19 | 19 | 26,3 |
| Q8BVL3 | Q8BVL3 | Sorting ne: | Snx17 | sp\|Q8BVL3 | 11 | 11 | 29,6 |
| Q9EST5 | Q9EST5 | Acidic leuc | Anp32b | sp\|Q9EST5 | 8 | 6 | 38,6 |
| D3YZG8 | D3YZG8 | Probable b | Mthfd2l | sp\|D3YZG8 | 7 | 7 | 28,7 |
| Q80SY5 | Q80SY5 | Pre-mRNA- | Prpf38b | sp\|Q80SY5 | 8 | 8 | 14,2 |
| P06537 | P06537 | Glucocortic | Nr3c1 | sp\|P06537 | 18 | 18 | 37,9 |
| B7ZMP1 | B7ZMP1 | Probable X | Xpnpep3 | sp\|B7ZMP | 12 | 12 | 37,4 |
| Q7TMB8 | Q7TMB8 | Cytoplasm | Cyfip1 | sp\|Q7TMB | 41 | 27 | 39,5 |
| Q9CZX5 | Q9CZX5 | PIN2/TERF | Pinx1 | sp\|Q9CZX5 | 3 | 3 | 13,6 |
| Q80UW8 | Q80UW8 | DNA-direct | Polr2e | sp\|Q80UW | 8 | 8 | 58,6 |
| P53564 | P53564 | Homeobox | Cux1 | sp\|P53564 | 41 | 24 | 42,7 |
| Q9R1C6 | Q9R1C6 | Diacylglyce | Dgke | sp\|Q9R1C6 | 5 | 5 | 12,4 |
| Q91VT4 | Q91VT4 | Carbonyl r | Cbr4 | sp\|Q91VT4 | 6 | 6 | 35,6 |
| Q9D968 | Q9D968 | Host cell fa | Hcfc2 | sp\|Q9D968 | 15 | 14 | 27,7 |
| Q9Z130 | Q9Z130 | Heterogen | Hnrnpdl | sp\|Q9Z130 | 13 | 12 | 32,9 |
| Q9CPT3 | Q9CPT3 | N-acylneur | Nanp | sp\|Q9CPT3 | 7 | 6 | 34,3 |
| Q8VCA8 | Q8VCA8 | Secernin-2 | Scrn2 | sp\|Q8VCA8 | 10 | 10 | 42,8 |
| Q80U87 | Q80U87 | Ubiquitin c | Usp8 | sp\|Q80U87 | 12 | 12 | 15,5 |
| Q91WD1 | Q91WD1 | DNA-direct | Polr3d | sp\|Q91WD | 4 | 4 | 12,1 |
| Q9CQ48 | Q9CQ48 | NudC dom | Nudcd2 | sp\|Q9CQ4 | 8 | 8 | 63,1 |
| Q8VDD9 | Q8VDD9 | PH-interac | Phip | sp\|Q8VDD | 42 | 40 | 31,5 |
| P35505 | P35505 | Fumarylac | Fah | sp\|P35505 | 13 | 13 | 45,6 |
| P97760 | P97760 | DNA-direct | Polr2c | sp\|P97760 | 10 | 10 | 41,5 |
| P58404 | P58404 | Striatin-4 | Strn4 | sp\|P58404 | 16 | 14 | 34,1 |
| Q8VI33;Q6 | Q8VI33;Q6NZA | Transcripti | Taf9;Taf9b | sp\|Q8VI33 | 10 | 10 | 45,5 |
| Q9QZR0 | Q9QZR0 | E3 ubiquiti | Rnf25 | sp\|Q9QZR( | 8 | 8 | 26,5 |
| Q99MB2 | Q99MB2 | Mitochond | Mtfr1 | sp\|Q99MB | 5 | 5 | 25,6 |
| P56212 | P56212 | cAMP-regu | Arpp19 | sp\|P56212 | 8 | 7 | 68,8 |
| Q3TIV5 | Q3TIV5 | Zinc finger | Zc3h15 | sp\|Q3TIV5 | 14 | 14 | 42,5 |
| P51859 | P51859 | Hepatoma | Hdgf | sp\|P51859 | 22 | 21 | 84,4 |
| Q80TA9 | Q80TA9 | Ectopic P g | Epg5 | sp\|Q80TA9 | 10 | 10 | 5,2 |
| Q3TCN2 | Q3TCN2 | Putative ph | Plbd2 | sp\|Q3TCN2 | 10 | 10 | 24,6 |
| B1AUR6 | B1AUR6 | Protein MN | Mms22l | sp\|B1AUR6 | 11 | 11 | 12,1 |
| Q91VH2 | Q91VH2 | Sorting ne: | Snx9 | sp\|Q91VH | 21 | 21 | 45,7 |
| Q9DB98 | Q9DB98 | Leukocyte | Leng1 | sp\|Q9DB9 | 1 | 1 | 8,8 |
| Q9D967 | Q9D967 | Magnesiun | Mdp1 | sp\|Q9D967 | 7 | 7 | 53,7 |
| P97360 | P97360 | Transcripti | Etv6 | sp\|P97360 | 7 | 7 | 21 |
| Q8BGU5 | Q8BGU5 | Cyclin-Y | Ccny | sp\|Q8BGU | 6 | 6 | 22,9 |
| Q60520 | Q60520 | Paired amp | Sin3a | sp\|Q60520 | 38 | 37 | 37,6 |
| Q8VE88 | Q8VE88 | Protein FA | Fam114a2 | sp\|Q8VE88 | 14 | 14 | 37,8 |
| O88554 | O88554 | Poly [ADP- | Parp2 | sp\|O88554 | 11 | 11 | 28,6 |
| O08919 | O08919 | Numb-like | Numbl | sp\|O08919 | 3 | 3 | 6,3 |
| Q924Z5 | Q924Z5 | Translocati | Tram2 | sp\|Q924Z5 | 2 | 2 | 5,7 |
| Q99LN9 | Q99LN9 | Deoxyhypu | Dohh | sp\|Q99LN9 | 14 | 14 | 66,9 |
| Q8C4Y3 | Q8C4Y3 | Negative e | Nelfb | sp\|Q8C4Y3 | 17 | 17 | 43,3 |

| | | | | | | | |
|---|---|---|---|---|---|---|---|
| O54692 | O54692 | Centromer | Zw10 | sp\|O54692 | 25 | 25 | 45,4 |
| Q99LB2 | Q99LB2 | Dehydroge | Dhrs4 | sp\|Q99LB2 | 11 | 11 | 50,9 |
| Q99N87 | Q99N87 | 28S riboso | Mrps5 | sp\|Q99N87 | 15 | 15 | 42,6 |
| P58069 | P58069 | Ras GTPase | Rasa2 | sp\|P58069 | 11 | 11 | 16,3 |
| Q3U6N9 | Q3U6N9 | UPF0488 protein C8or | | sp\|Q3U6N | 6 | 6 | 44,6 |
| P23950 | P23950 | Zinc finger | Zfp36l1 | sp\|P23950 | 3 | 2 | 12,7 |
| A9JSM3 | A9JSM3 | Transmem | Tmem238 | sp\|A9JSM3 | 5 | 5 | 40,9 |
| Q9CZD5 | Q9CZD5 | Translation | Mtif3 | sp\|Q9CZD5 | 4 | 4 | 19,2 |
| P54265 | P54265 | Myotonin- | Dmpk | sp\|P54265 | 5 | 4 | 11,1 |
| Q9DB90 | Q9DB90 | Protein SM | Smg9 | sp\|Q9DB90 | 13 | 13 | 42,9 |
| O35071 | O35071 | Kinesin-like | Kif1c | sp\|O35071 | 16 | 13 | 19,2 |
| Q9ESL4 | Q9ESL4 | Mitogen-ac | Zak | sp\|Q9ESL4 | 23 | 23 | 39 |
| Q8CHG7 | Q8CHG7 | Rap guanin | Rapgef2 | sp\|Q8CHG | 14 | 14 | 14,4 |
| Q9CPW7 | Q9CPW7 | Zinc finger | Zmat2 | sp\|Q9CPW | 3 | 3 | 16,6 |
| Q9WV86 | Q9WV86 | Katanin p6 | Katna1 | sp\|Q9WV8 | 12 | 11 | 34,2 |
| Q8CEC0 | Q8CEC0 | Nuclear po | Nup88 | sp\|Q8CEC0 | 18 | 18 | 34,7 |
| Q9CRA4 | Q9CRA4 | Methylster | Msmo1 | sp\|Q9CRA4 | 4 | 4 | 15 |
| Q6P5C5 | Q6P5C5 | Single-stra | Smug1 | sp\|Q6P5C5 | 7 | 7 | 49,1 |
| Q9Z2D0 | Q9Z2D0 | Myotubula | Mtmr9 | sp\|Q9Z2D0 | 9 | 9 | 20,2 |
| P38585 | P38585 | Tubulin--ty | Ttl | sp\|P38585 | 5 | 5 | 21,2 |
| O54782 | O54782 | Epididymis | Man2b2 | sp\|O54782 | 13 | 13 | 18,5 |
| Q8BHW2 | Q8BHW2 | Protein OS | Oscp1 | sp\|Q8BHW | 4 | 4 | 13,2 |
| Q9WTP2 | Q9WTP2 | Protein spr | Spry4 | sp\|Q9WTP | 4 | 4 | 21,3 |
| P01582 | P01582 | Interleukin | Il1a | sp\|P01582 | 5 | 5 | 28,1 |
| Q8CI59 | Q8CI59 | Metallored | Steap3 | sp\|Q8CI59 | 12 | 12 | 26,8 |
| Q61070 | Q61070 | Etoposide- | Ei24 | sp\|Q61070 | 4 | 4 | 12,6 |
| Q9D1R1 | Q9D1R1 | Complex I | Tmem126b | sp\|Q9D1R | 2 | 2 | 10,4 |
| Q9WV68 | Q9WV68 | Peroxisom | Decr2 | sp\|Q9WV6 | 5 | 5 | 26 |
| Q80YV4 | Q80YV4 | Pantothen | Pank4 | sp\|Q80YV4 | 11 | 11 | 16,3 |
| Q8QZY1 | Q8QZY1 | Eukaryotic | Eif3l | sp\|Q8QZY1 | 27 | 27 | 52 |
| Q9DC63 | Q9DC63 | F-box only | Fbxo3 | sp\|Q9DC63 | 11 | 11 | 25,8 |
| Q61152 | Q61152 | Tyrosine-p | Ptpn18 | sp\|Q61152 | 1 | 1 | 3,5 |
| P11627 | P11627 | Neural cell | L1cam | sp\|P11627 | 3 | 3 | 4,4 |
| Q3UY34 | Q3UY34 | Uncharacterized prote | | sp\|Q3UY34 | 7 | 7 | 42,2 |
| Q56A10 | Q56A10 | Zinc finger | Znf608 | sp\|Q56A10 | 12 | 12 | 12,5 |
| Q8VDY9 | Q8VDY9 | Caspase ac | Caap1 | sp\|Q8VDY9 | 5 | 5 | 24,2 |
| Q8BHS3 | Q8BHS3 | Pre-mRNA- | Rbm22 | sp\|Q8BHS3 | 15 | 15 | 47,6 |
| Q8BIV3 | Q8BIV3 | Ran-bindin | Ranbp6 | sp\|Q8BIV3 | 7 | 6 | 7,6 |
| Q61521 | Q61521 | Proepiregu | Ereg | sp\|Q61521 | 1 | 1 | 6,8 |
| CON__P08 | CON__P08727 | | | | 11 | 2 | 21 |
| Q9CZB3 | Q9CZB3 | THUMP do | Thumpd2 | sp\|Q9CZB3 | 4 | 4 | 10 |
| Q04207 | Q04207 | Transcripti | Rela | sp\|Q04207 | 13 | 13 | 36,4 |
| Q9Z0F8 | Q9Z0F8 | Disintegrin | Adam17 | sp\|Q9Z0F8 | 17 | 17 | 32 |
| P36536 | P36536 | GTP-bindin | Sar1a | sp\|P36536 | 7 | 6 | 57,6 |
| Q3TUF7 | Q3TUF7 | YEATS dom | Yeats2 | sp\|Q3TUF7 | 17 | 17 | 21 |
| Q9CQT5 | Q9CQT5 | Proteasom | Pomp | sp\|Q9CQT | 4 | 4 | 42,6 |
| Q3U155 | Q3U155 | Coiled-coil | Ccdc174 | sp\|Q3U15 | 5 | 5 | 13,3 |
| Q8BLX7 | Q8BLX7 | Collagen al | Col16a1 | sp\|Q8BLX7 | 2 | 2 | 2,3 |
| Q922B1 | Q922B1 | O-acetyl-A | Macrod1 | sp\|Q922B1 | 3 | 3 | 14,9 |
| Q9D2Y9 | Q9D2Y9 | Elongation | Elovl7 | sp\|Q9D2Y9 | 2 | 2 | 5,3 |

| | | | | | | | |
|---|---|---|---|---|---|---|---|
| B2RPV6 | B2RPV6 | Multimerir | Mmrn1 | sp\|B2RPV6 | 2 | 2 | 1,7 |
| Q9D6L8 | Q9D6L8 | Peptidyl-pr | Ppil3 | sp\|Q9D6L8 | 8 | 8 | 52,8 |
| Q9CQX5 | Q9CQX5 | Claudin do | Cldnd1 | sp\|Q9CQX | 2 | 2 | 8,7 |
| Q8CHT3 | Q8CHT3 | Integrator | Ints5 | sp\|Q8CHT3 | 15 | 15 | 18,8 |
| Q61749 | Q61749 | Translation | Eif2b4 | sp\|Q61749 | 19 | 19 | 48,9 |
| Q8BH95 | Q8BH95 | Enoyl-CoA | Echs1 | sp\|Q8BH95 | 12 | 12 | 46,9 |
| Q5RJH6 | Q5RJH6 | Protein SM | Smg7 | sp\|Q5RJH6 | 9 | 9 | 11,7 |
| Q8R0W6 | Q8R0W6 | NEDD4 fan | Ndfip1 | sp\|Q8R0W | 5 | 5 | 29 |
| Q8BFQ4 | Q8BFQ4 | WD repeat | Wdr82 | sp\|Q8BFQ4 | 14 | 14 | 57,2 |
| CON__Q2K | CON__Q2KIS7 | | | | 1 | 1 | 6,9 |
| P42932 | P42932 | T-complex | Cct8 | sp\|P42932 | 40 | 40 | 76,1 |
| P46662 | P46662 | Merlin | Nf2 | sp\|P46662 | 3 | 3 | 5,9 |
| P18828 | P18828 | Syndecan-1 | Sdc1 | sp\|P18828 | 3 | 3 | 10 |
| P70452 | P70452 | Syntaxin-4 | Stx4 | sp\|P70452 | 5 | 5 | 24,5 |
| Q9R1X4 | Q9R1X4 | Protein tim | Timeless | sp\|Q9R1X4 | 27 | 27 | 33,3 |
| Q9JMB7 | Q9JMB7 | Piwi-like pr | Piwil1 | sp\|Q9JMB | 2 | 2 | 3 |
| Q7TN98 | Q7TN98 | Cytoplasm | Cpeb4 | sp\|Q7TN98 | 11 | 6 | 21 |
| Q91WG3 | Q91WG3 | Probable t | Trub2 | sp\|Q91WG | 5 | 5 | 17,5 |
| Q8R149 | Q8R149 | BUD13 hor | Bud13 | sp\|Q8R149 | 5 | 5 | 10,4 |
| P62046 | P62046 | Leucine-ric | Lrch1 | sp\|P62046 | 7 | 7 | 13 |
| P23336 | P23336 | N-acetylla | Ggta1 | sp\|P23336 | 3 | 3 | 10,2 |
| P49718 | P49718 | DNA replic | Mcm5 | sp\|P49718 | 44 | 44 | 64,2 |
| Q8C0J2 | Q8C0J2 | Autophagy | Atg16l1 | sp\|Q8C0J2 | 7 | 7 | 21,9 |
| Q91WD0 | Q91WD0 | Protein GP | Gpr108 | sp\|Q91WD | 2 | 2 | 6 |
| Q8BFW4 | Q8BFW4 | Tripartite r | Trim65 | sp\|Q8BFW | 10 | 10 | 30,7 |
| Q7TSQ8 | Q7TSQ8 | Pyruvate d | Pdpr | sp\|Q7TSQ8 | 17 | 17 | 23,9 |
| Q8CD15 | Q8CD15 | Bifunctiona | Mina | sp\|Q8CD15 | 11 | 11 | 28 |
| Q8BJ37 | Q8BJ37 | Tyrosyl-DN | Tdp1 | sp\|Q8BJ37 | 12 | 12 | 22,2 |
| Q8VD57 | Q8VD57 | Vesicle tra | Sft2d2 | sp\|Q8VD5 | 1 | 1 | 10,7 |
| Q9DC61 | Q9DC61 | Mitochond | Pmpca | sp\|Q9DC6 | 24 | 24 | 51,1 |
| Q9JJC6 | Q9JJC6 | RILP-like pr | Rilpl1 | sp\|Q9JJC6 | 4 | 4 | 10,6 |
| P62305 | P62305 | Small nucle | Snrpe | sp\|P62305 | 4 | 4 | 65,2 |
| O35047 | O35047 | Homologo | Psmc3ip | sp\|O35047 | 4 | 4 | 28,1 |
| Q5SV85 | Q5SV85 | Synergin ga | Synrg | sp\|Q5SV85 | 18 | 18 | 21,1 |
| Q9CQ56 | Q9CQ56 | Vesicle tra | Use1 | sp\|Q9CQ5 | 9 | 9 | 39,6 |
| Q99L43 | Q99L43 | Phosphatid | Cds2 | sp\|Q99L43 | 4 | 4 | 12,2 |
| Q8BHD7 | Q8BHD7 | Polypyrimi | Ptbp3 | sp\|Q8BHD | 16 | 13 | 45,7 |
| Q8BP78 | Q8BP78 | Protein FR/ | Fra10ac1 | sp\|Q8BP78 | 6 | 6 | 23,8 |
| Q00493 | Q00493 | Carboxype | Cpe | sp\|Q00493 | 10 | 10 | 28,8 |
| Q80TV8 | Q80TV8 | CLIP-assoc | Clasp1 | sp\|Q80TV8 | 37 | 32 | 30,7 |
| Q8BTN6 | Q8BTN6 | Leukocyte | Leng9 | sp\|Q8BTN | 3 | 3 | 9,9 |
| Q03173 | Q03173 | Protein en | Enah | sp\|Q03173 | 11 | 11 | 19,5 |
| Q62388 | Q62388 | Serine-pro | Atm | sp\|Q62388 | 34 | 34 | 15,4 |
| Q3U1J1 | Q3U1J1 | TCF3 fusio | Tfpt | sp\|Q3U1J1 | 4 | 4 | 23,9 |
| Q91YU3 | Q91YU3 | Myb/SANT | Msantd4 | sp\|Q91YU3 | 4 | 4 | 19,4 |
| Q9D4F8 | Q9D4F8 | Gamma-tu | Tubgcp4 | sp\|Q9D4F8 | 9 | 9 | 20,4 |
| P97432 | P97432 | Next to BR | Nbr1 | sp\|P97432 | 8 | 8 | 13,9 |
| Q8BK67 | Q8BK67 | Protein RC | Rcc2 | sp\|Q8BK67 | 26 | 26 | 63,7 |
| Q7TMR0 | Q7TMR0 | Lysosomal | Prcp | sp\|Q7TMR | 8 | 8 | 25,9 |
| Q9D706 | Q9D706 | RNA polym | Rpap3 | sp\|Q9D70 | 27 | 27 | 50,2 |

| | | | | | | | |
|---|---|---|---|---|---|---|---|
| P47713 | P47713 | Cytosolic p | Pla2g4a | sp\|P47713 | 19 | 19 | 31,4 |
| Q8R1V4 | Q8R1V4 | Transmem | Tmed4 | sp\|Q8R1V4 | 8 | 7 | 41,4 |
| Q5I043 | Q5I043 | Ubiquitin c | Usp28 | sp\|Q5I043 | 12 | 12 | 14,6 |
| P14211 | P14211 | Calreticulin | Calr | sp\|P14211 | 26 | 26 | 77,2 |
| Q9D1C9 | Q9D1C9 | Ribosomal | Rrp7a | sp\|Q9D1C9 | 11 | 11 | 49,6 |
| Q6P3Y5 | Q6P3Y5 | Zinc finger | Znf280c | sp\|Q6P3Y5 | 13 | 11 | 24,9 |
| Q64511 | Q64511 | DNA topois | Top2b | sp\|Q64511 | 70 | 54 | 47,8 |
| Q6P4S6 | Q6P4S6 | Serine/thre | Sik3 | sp\|Q6P4S6 | 20 | 20 | 22,8 |
| Q3TAP4 | Q3TAP4 | AP-5 comp | Ap5b1 | sp\|Q3TAP4 | 4 | 4 | 6,7 |
| Q8BGS0 | Q8BGS0 | Protein MA | Mak16 | sp\|Q8BGS0 | 11 | 11 | 39,2 |
| Q9CZJ9 | Q9CZJ9 | DnaJ homo | Dnajc18 | sp\|Q9CZJ9 | 6 | 6 | 21 |
| Q9EPR4 | Q9EPR4 | Solute carr | Slc23a2 | sp\|Q9EPR4 | 3 | 3 | 4,8 |
| Q9DBT3 | Q9DBT3 | Coiled-coil | Ccdc97 | sp\|Q9DBT3 | 9 | 9 | 45,6 |
| Q8C176 | Q8C176 | Transcripti | Taf2 | sp\|Q8C176 | 12 | 12 | 15 |
| Q80Z37 | Q80Z37 | E3 ubiquiti | Topors | sp\|Q80Z37 | 4 | 4 | 4,7 |
| Q8BX17 | Q8BX17 | Gem-assoc | Gemin5 | sp\|Q8BX17 | 37 | 37 | 31,1 |
| CON__Q3T | CON__Q3T052 | | | | 13 | 13 | 16,5 |
| Q9CXW3 | Q9CXW3 | Calcyclin-b | Cacybp | sp\|Q9CXW | 16 | 16 | 71,2 |
| P52624;Q8 | P52624 | Uridine ph | Upp1 | sp\|P52624 | 15 | 15 | 63 |
| Q8BUY9 | Q8BUY9 | Geranylger | Pggt1b | sp\|Q8BUY9 | 6 | 6 | 18 |
| Q8R1U1 | Q8R1U1 | Conserved | Cog4 | sp\|Q8R1U | 26 | 26 | 41,7 |
| Q8BG40 | Q8BG40 | Katanin p8 | Katnb1 | sp\|Q8BG40 | 10 | 10 | 23,7 |
| Q924H7 | Q924H7 | WW doma | Wac | sp\|Q924H7 | 10 | 10 | 23,4 |
| P13405 | P13405 | Retinoblas | Rb1 | sp\|P13405 | 6 | 6 | 7,9 |
| P60229 | P60229 | Eukaryotic | Eif3e | sp\|P60229 | 26 | 26 | 67,2 |
| P58389 | P58389 | Serine/thre | Ppp2r4 | sp\|P58389 | 10 | 10 | 50,8 |
| Q4LDD4 | Q4LDD4 | Arf-GAP wi | Arap1 | sp\|Q4LDD4 | 16 | 16 | 15,3 |
| Q8BIF9 | Q8BIF9 | Zinc finger | Znf787 | sp\|Q8BIF9 | 4 | 4 | 15 |
| Q8K4J6 | Q8K4J6 | MKL/myoc | Mkl1 | sp\|Q8K4J6 | 11 | 10 | 16,8 |
| A6H630 | A6H630 | Protein-glu | Armt1 | sp\|A6H630 | 8 | 8 | 17,3 |
| Q91ZW3 | Q91ZW3 | SWI/SNF-r | Smarca5 | sp\|Q91ZW | 56 | 43 | 54,5 |
| A2A6A1;A2 | A2A6A1 | G patch do | Gpatch8 | sp\|A2A6A1 | 12 | 12 | 12 |
| Q6PD28 | Q6PD28 | | Ppp2r5b | sp\|Q6PD28 | 5 | 3 | 12,1 |
| Q9DBU6 | Q9DBU6 | Serine/Arg | Rsrc1 | sp\|Q9DBU | 7 | 7 | 25,4 |
| Q8BRT1 | Q8BRT1 | CLIP-assoc | Clasp2 | sp\|Q8BRT1 | 32 | 27 | 35,1 |
| P97390 | P97390 | Vacuolar p | Vps45 | sp\|P97390 | 17 | 17 | 34,2 |
| P13745;P1 | P13745;P1064 | Glutathion | Gsta1;Gsta | sp\|P13745 | 5 | 4 | 23,3 |
| O70546 | O70546 | Lysine-spe | Kdm6a | sp\|O70546 | 13 | 11 | 13,4 |
| P11438 | P11438 | Lysosome- | Lamp1 | sp\|P11438 | 6 | 6 | 19,2 |
| Q64012 | Q64012 | RNA-bindir | Raly | sp\|Q64012 | 16 | 16 | 55,4 |
| Q5DTX6 | Q5DTX6 | Junctional | Jcad | sp\|Q5DTX6 | 11 | 11 | 13,9 |
| Q61187 | Q61187 | Tumor sus | Tsg101 | sp\|Q61187 | 14 | 14 | 38,9 |
| Q91VC3 | Q91VC3 | Eukaryotic | Eif4a3 | sp\|Q91VC3 | 28 | 22 | 63,3 |
| Q3U0V2 | Q3U0V2 | Tumor nec | Tradd | sp\|Q3U0V | 8 | 8 | 41,9 |
| Q8CBC4 | Q8CBC4 | Consortin | Cnst | sp\|Q8CBC4 | 5 | 5 | 10,8 |
| O09012 | O09012 | Peroxisom | Pex5 | sp\|O09012 | 15 | 15 | 35,5 |
| Q9D0F4;Q | Q9D0F4 | NF-kappa-I | Nkap | sp\|Q9D0F4 | 5 | 5 | 16,1 |
| Q80TA6 | Q80TA6 | Myotubula | Mtmr12 | sp\|Q80TA6 | 11 | 11 | 18,6 |
| P31266;O0 | P31266 | Recombini | Rbpj | sp\|P31266 | 16 | 16 | 37,6 |
| Q4VAA7 | Q4VAA7 | Sorting ne | Snx33 | sp\|Q4VAA | 6 | 6 | 13,4 |

| | | | | | | | |
|---|---|---|---|---|---|---|---|
| Q8VC34 | Q8VC34 | Putative RI | Rpap2 | sp\|Q8VC34 | 11 | 11 | 25,9 |
| Q60974 | Q60974 | Nuclear re | Ncor1 | sp\|Q60974 | 23 | 22 | 15,2 |
| Q923J1;Q2 | Q923J1 | Transient r | Trpm7 | sp\|Q923J1 | 10 | 9 | 8,7 |
| P53798 | P53798 | Squalene s | Fdft1 | sp\|P53798 | 9 | 9 | 31,2 |
| Q9Z160 | Q9Z160 | Conserved | Cog1 | sp\|Q9Z160 | 25 | 25 | 34,1 |
| P61965 | P61965 | WD repeat | Wdr5 | sp\|P61965 | 13 | 13 | 57,8 |
| P62880 | P62880 | Guanine nu | Gnb2 | sp\|P62880 | 14 | 6 | 57,4 |
| Q9DBH0 | Q9DBH0 | NEDD4-like | Wwp2 | sp\|Q9DBH | 11 | 10 | 18,3 |
| Q99K74 | Q99K74 | Mediator c | Med24 | sp\|Q99K74 | 14 | 14 | 19,8 |
| Q8R1B4 | Q8R1B4 | Eukaryotic | Eif3c | sp\|Q8R1B4 | 30 | 30 | 33,7 |
| Q9CXP4 | Q9CXP4 | Rho GTPas | Arhgap8 | sp\|Q9CXP4 | 3 | 3 | 11,1 |
| Q9CR24 | Q9CR24 | Nucleoside | Nudt8 | sp\|Q9CR24 | 5 | 5 | 32,9 |
| Q8BH04 | Q8BH04 | Phosphoer | Pck2 | sp\|Q8BH04 | 26 | 26 | 56,7 |
| Q63850 | Q63850 | Nuclear po | Nup62 | sp\|Q63850 | 13 | 13 | 28,9 |
| P47810 | P47810 | Wee1-like | Wee1 | sp\|P47810 | 3 | 3 | 6,3 |
| Q8CHS2 | Q8CHS2 | DnaJ homo | Dnajc22 | sp\|Q8CHS2 | 4 | 4 | 17,7 |
| Q8VD62 | Q8VD62 | UPF0696 p | Bles03 | sp\|Q8VD6 | 9 | 9 | 33,6 |
| O88697 | O88697 | Serine/thr | Stk16 | sp\|O88697 | 3 | 3 | 17 |
| Q9CPR8 | Q9CPR8 | Melanoma | Ndnl2 | sp\|Q9CPR8 | 11 | 11 | 45,9 |
| P62702 | P62702 | 40S riboso | Rps4x | sp\|P62702 | 21 | 21 | 64,6 |
| Q62415 | Q62415 | Apoptosis- | Ppp1r13b | sp\|Q62415 | 6 | 5 | 7,1 |
| Q80TF3 | Q80TF3 | Protocadh | Pcdh19 | sp\|Q80TF3 | 2 | 2 | 2,6 |
| Q8BG32 | Q8BG32 | 26S protea | Psmd11 | sp\|Q8BG3 | 32 | 32 | 70,9 |
| Q9CQC5 | Q9CQC5 | Cdc42 effe | Cdc42ep3 | sp\|Q9CQC | 3 | 3 | 24,4 |
| Q69Z66 | Q69Z66 | Histone H2 | Mysm1 | sp\|Q69Z66 | 6 | 6 | 10,3 |
| CON__Q3S | CON__Q3SX09 | | | | 11 | 9 | 60,7 |
| Q8VH51 | Q8VH51 | RNA-bindir | Rbm39 | sp\|Q8VH5 | 21 | 21 | 43 |
| Q91Z92 | Q91Z92 | Beta-1,3-g | B3galt6 | sp\|Q91Z92 | 4 | 4 | 15,1 |
| Q9CYI4 | Q9CYI4 | Putative RI | Luc7l | sp\|Q9CYI4 | 11 | 7 | 32,1 |
| Q921Q7 | Q921Q7 | Ras and Ra | Rin1 | sp\|Q921Q | 17 | 17 | 32,8 |
| Q9D0R2 | Q9D0R2 | Threonine- | Tars | sp\|Q9D0R | 36 | 34 | 55,7 |
| O54918 | O54918 | Bcl-2-like p | Bcl2l11 | sp\|O54918 | 3 | 3 | 17,3 |
| Q8C6G8 | Q8C6G8 | WD repeat | Wdr26 | sp\|Q8C6G | 21 | 21 | 47,3 |
| Q9D6Z0 | Q9D6Z0 | Alpha-keto | Alkbh7 | sp\|Q9D6Z0 | 3 | 3 | 25,8 |
| Q6NXW6 | Q6NXW6 | Cell cycle c | Rad17 | sp\|Q6NXW | 5 | 5 | 7,8 |
| Q9JME7 | Q9JME7 | Trafficking | Trappc2l | sp\|Q9JME7 | 2 | 2 | 19,4 |
| Q91YR5 | Q91YR5 | Methyltrar | Mettl13 | sp\|Q91YR5 | 17 | 17 | 34,7 |
| Q9CQ60 | Q9CQ60 | 6-phospho | Pgls | sp\|Q9CQ6 | 9 | 9 | 61,1 |
| P58929 | P58929 | Glucocorti | Gmeb2 | sp\|P58929 | 8 | 6 | 15,3 |
| Q6ZPI0 | Q6ZPI0 | Protein Jac | Jade1 | sp\|Q6ZPI0 | 8 | 8 | 13,3 |
| O35638 | O35638 | Cohesin su | Stag2 | sp\|O35638 | 35 | 31 | 35,2 |
| Q99M04 | Q99M04 | Lipoyl synt | Lias | sp\|Q99M0 | 10 | 10 | 42,6 |
| Q9EPK7 | Q9EPK7 | Exportin-7 | Xpo7 | sp\|Q9EPK7 | 28 | 27 | 34,4 |
| Q9D7B7 | Q9D7B7 | Probable g | Gpx8 | sp\|Q9D7B | 6 | 6 | 37,3 |
| P28867;RE | P28867 | Protein kin | Prkcd | sp\|P28867 | 17 | 16 | 35,8 |
| O54833 | O54833 | Casein kina | Csnk2a2 | sp\|O54833 | 18 | 18 | 65,1 |
| Q33DR2 | Q33DR2 | Decapreny | Pdss1 | sp\|Q33DR | 4 | 4 | 14,9 |
| Q3UGP8 | Q3UGP8 | Putative D | Alg10b | sp\|Q3UGP | 4 | 4 | 8,4 |
| Q8BGB2 | Q8BGB2 | Tetratricop | Ttc7a | sp\|Q8BGB | 6 | 6 | 8,4 |
| Q9ERF3 | Q9ERF3 | WD repeat | Wdr61 | sp\|Q9ERF3 | 15 | 15 | 58,7 |

| | | | | | | | |
|---|---|---|---|---|---|---|---|
| Q8BYJ6 | Q8BYJ6 | TBC1 doma | Tbc1d4 | sp\|Q8BYJ6 | 14 | 14 | 16 |
| O08807 | O08807 | Peroxiredo | Prdx4 | sp\|O08807 | 14 | 13 | 60,9 |
| Q9JL15 | Q9JL15 | Galectin-8 | Lgals8 | sp\|Q9JL15 | 7 | 7 | 28,2 |
| P18872 | P18872 | Guanine nu | Gnao1 | sp\|P18872 | 6 | 4 | 22 |
| Q9D0V7 | Q9D0V7 | Receptor-b | Ebag9 | sp\|Q9D0V | 3 | 3 | 28,2 |
| P81117 | P81117 | Nucleobind | Nucb2 | sp\|P81117 | 18 | 18 | 47,9 |
| A2AF47 | A2AF47 | Dedicator d | Dock11 | sp\|A2AF47 | 24 | 21 | 15,7 |
| Q91VX2 | Q91VX2 | Ubiquitin-a | Ubap2 | sp\|Q91VX2 | 19 | 19 | 29,7 |
| Q8VDW0 | Q8VDW0 | ATP-depen | Ddx39a | sp\|Q8VDW | 22 | 12 | 63,9 |
| Q9DC28 | Q9DC28 | Casein kina | Csnk1d | sp\|Q9DC28 | 12 | 6 | 34,9 |
| Q6ZQ11 | Q6ZQ11 | Chondroiti | Chsy1 | sp\|Q6ZQ11 | 6 | 6 | 10,5 |
| Q9DA08 | Q9DA08 | SAGA-asso | Ccdc101 | sp\|Q9DA08 | 6 | 6 | 28,3 |
| Q99J36 | Q99J36 | THUMP do | Thumpd1 | sp\|Q99J36 | 16 | 16 | 53,7 |
| Q9D2D7 | Q9D2D7 | Zinc finger | Znf687 | sp\|Q9D2D | 11 | 11 | 13,6 |
| O08756 | O08756 | 3-hydroxya | Hsd17b10 | sp\|O08756 | 10 | 10 | 51,7 |
| Q61712 | Q61712 | DnaJ homo | Dnajc1 | sp\|Q61712 | 9 | 9 | 17,6 |
| Q64519 | Q64519 | Syndecan-3 | Sdc3 | sp\|Q64519 | 1 | 1 | 5 |
| O88271 | O88271 | Craniofacia | Cfdp1 | sp\|O88271 | 6 | 6 | 21,7 |
| Q99M01 | Q99M01 | Phenylalan | Fars2 | sp\|Q99M0 | 7 | 7 | 23,1 |
| Q9DB32 | Q9DB32 | Hydroxyac | Haghl | sp\|Q9DB32 | 4 | 4 | 21,9 |
| O35099 | O35099 | Mitogen-ac | Map3k5 | sp\|O35099 | 6 | 6 | 4,9 |
| Q8K5B2 | Q8K5B2 | Multiple co | Mcfd2 | sp\|Q8K5B2 | 2 | 2 | 28,3 |
| P70302 | P70302 | Stromal int | Stim1 | sp\|P70302 | 16 | 15 | 34,3 |
| Q91YH5 | Q91YH5 | Atlastin-3 | Atl3 | sp\|Q91YH5 | 25 | 24 | 57,7 |
| Q9JLY7 | Q9JLY7 | Dual specif | Dusp14 | sp\|Q9JLY7 | 2 | 2 | 13,6 |
| Q8BJS8 | Q8BJS8 | Mdm2-bin | Mtbp | sp\|Q8BJS8 | 11 | 11 | 20,4 |
| Q9D8W5 | Q9D8W5 | 26S protea | Psmd12 | sp\|Q9D8W | 27 | 27 | 57 |
| Q9Z1N2 | Q9Z1N2 | Origin reco | Orc1 | sp\|Q9Z1N2 | 23 | 23 | 33,2 |
| Q9CQA0 | Q9CQA0 | Centromer | Cenpm | sp\|Q9CQA | 2 | 2 | 16,7 |
| Q99JH1 | Q99JH1 | Ribonuclea | Rpp25l | sp\|Q99JH1 | 3 | 3 | 25,8 |
| G3X9K3 | G3X9K3 | Brefeldin A | Arfgef1 | sp\|G3X9K3 | 33 | 26 | 23,6 |
| Q3UX10 | Q3UX10 | Tubulin alp | Tubal3 | sp\|Q3UX10 | 5 | 2 | 13 |
| Q9CQJ7 | Q9CQJ7 | Securin | Pttg1 | sp\|Q9CQJ7 | 5 | 5 | 26,6 |
| Q9WUZ9 | Q9WUZ9 | Ectonucleo | Entpd5 | sp\|Q9WUZ | 6 | 6 | 17,3 |
| Q9D7J6 | Q9D7J6 | Deoxyribon | Dnase1l1 | sp\|Q9D7J6 | 3 | 3 | 13,7 |
| Q7TMC8 | Q7TMC8 | | Fuk | sp\|Q7TMC | 11 | 11 | 16,5 |
| Q01405 | Q01405 | Protein tra | Sec23a | sp\|Q01405 | 24 | 21 | 45,8 |
| Q6P542 | Q6P542 | ATP-bindin | Abcf1 | sp\|Q6P542 | 29 | 29 | 48 |
| Q9DCA5 | Q9DCA5 | Ribosome | Brix1 | sp\|Q9DCA | 14 | 14 | 52,4 |
| A2A8U2 | A2A8U2 | Transmem | Tmem201 | sp\|A2A8U2 | 8 | 8 | 14,2 |
| Q9CR63 | Q9CR63 | Cytochrom | Cox16 | sp\|Q9CR63 | 2 | 2 | 16 |
| Q9WUJ8 | Q9WUJ8 | Origin reco | Orc6 | sp\|Q9WUJ | 3 | 3 | 13,4 |
| Q3UM18 | Q3UM18 | Large subu | Lsg1 | sp\|Q3UM1 | 18 | 18 | 32 |
| P23116 | P23116 | Eukaryotic | Eif3a | sp\|P23116 | 63 | 63 | 46,4 |
| O89079 | O89079 | Coatomer | Cope | sp\|O89079 | 14 | 14 | 59,1 |
| P53986 | P53986 | Monocarbo | Slc16a1 | sp\|P53986 | 7 | 7 | 17 |
| Q64685 | Q64685 | Beta-galac | St6gal1 | sp\|Q64685 | 2 | 2 | 6,5 |
| Q9CQR7 | Q9CQR7 | Gamma-se | Psenen | sp\|Q9CQR | 1 | 1 | 14,9 |
| P21958 | P21958 | Antigen pe | Tap1 | sp\|P21958 | 7 | 7 | 13,3 |
| Q9Z277 | Q9Z277 | Tyrosine-p | Baz1b | sp\|Q9Z277 | 55 | 55 | 41,4 |

| | | | | | | | |
|---|---|---|---|---|---|---|---|
| Q8R3Q2 | Q8R3Q2 | Serine/thre | Ppp6r2 | sp\|Q8R3Q | 3 | 3 | 5 |
| P15806 | P15806 | Transcripti | Tcf3 | sp\|P15806 | 5 | 5 | 14 |
| O54950 | O54950 | 5-AMP-act | Prkag1 | sp\|O5495( | 12 | 11 | 47,9 |
| Q8BKS9 | Q8BKS9 | Pumilio do | Kiaa0020 | sp\|Q8BKS9 | 28 | 28 | 47,8 |
| Q6PDH0 | Q6PDH0 | Pleckstrin | Phldb1 | sp\|Q6PDH | 34 | 34 | 35,2 |
| O88508 | O88508 | DNA (cytos | Dnmt3a | sp\|O8850{ | 12 | 9 | 14,9 |
| P04768;P0 | P04768;P0409 | Prolactin-2 | Prl2c3;Prl2 | sp\|P04768 | 1 | 1 | 6,7 |
| Q9D6J5 | Q9D6J5 | NADH deh | Ndufb8 | sp\|Q9D6J5 | 6 | 6 | 46,2 |
| Q60739 | Q60739 | BAG family | Bag1 | sp\|Q6073! | 8 | 8 | 25,1 |
| Q3TWF6 | Q3TWF6 | WD repeat | Wdr70 | sp\|Q3TWF | 23 | 23 | 46,4 |
| Q811C2 | Q811C2 | Cysteine p | Atg4c | sp\|Q811C2 | 3 | 3 | 7,6 |
| Q8BWW9 | Q8BWW9 | Serine/thre | Pkn2 | sp\|Q8BW\ | 39 | 38 | 49,8 |
| Q6A4J8 | Q6A4J8 | Ubiquitin c | Usp7 | sp\|Q6A4J8 | 48 | 48 | 50,1 |
| Q0P5V9 | Q0P5V9 | Solute carr | Slc45a4 | sp\|Q0P5V! | 1 | 1 | 2,9 |
| P47199 | P47199 | Quinone o | Cryz | sp\|P47199 | 13 | 13 | 56,8 |
| Q8CGI1 | Q8CGI1 | Protein FA | Fam193a | sp\|Q8CGI1 | 7 | 7 | 8,4 |
| O08736;P7 | O08736 | Caspase-12 | Casp12 | sp\|O0873€ | 7 | 7 | 18,6 |
| Q8K2W3 | Q8K2W3 | Thioredoxi | Txndc11 | sp\|Q8K2W | 6 | 6 | 8,2 |
| Q91YN0 | Q91YN0 | Protein C1 | D6Wsu163 | sp\|Q91YN( | 4 | 4 | 11,6 |
| P01027 | P01027 | Compleme | C3 | sp\|P01027 | 8 | 6 | 6,3 |
| P41230;Q6 | P41230 | Lysine-spe | Kdm5c | sp\|P41230 | 16 | 13 | 15,1 |
| P30415 | P30415 | NK-tumor | Nktr | sp\|P30415 | 7 | 7 | 6,7 |
| Q61553 | Q61553 | Fascin | Fscn1 | sp\|Q6155: | 30 | 30 | 66,1 |
| Q8BLY7 | Q8BLY7 | Hermansky | Hps6 | sp\|Q8BLY7 | 14 | 14 | 26,3 |
| P70265 | P70265 | 6-phospho | Pfkfb2 | sp\|P70265 | 12 | 11 | 23,9 |
| Q8C5L3 | Q8C5L3 | CCR4-NOT | Cnot2 | sp\|Q8C5L3 | 13 | 13 | 35 |
| Q8VCQ3 | Q8VCQ3 | Nuclear re | Nrbf2 | sp\|Q8VCQ | 2 | 2 | 15,3 |
| Q9WV98 | Q9WV98 | Mitochond | Timm9 | sp\|Q9WV9 | 6 | 6 | 77,5 |
| Q80UX8 | Q80UX8 | Alpha/beta | Abhd13 | sp\|Q80UX8 | 4 | 4 | 14,5 |
| Q8BY46 | Q8BY46 | Zinc finger | Znf574 | sp\|Q8BY4€ | 4 | 4 | 8,3 |
| Q9JJE7 | Q9JJE7 | Fatty acid | Fads3 | sp\|Q9JJE7 | 2 | 2 | 7,8 |
| Q60749;Q9 | Q60749 | KH domain | Khdrbs1 | sp\|Q6074! | 12 | 12 | 30,9 |
| Q99PP9 | Q99PP9 | Tripartite r | Trim16 | sp\|Q99PP! | 18 | 18 | 40,5 |
| P63166 | P63166 | Small ubiq | Sumo1 | sp\|P63166 | 4 | 4 | 34,7 |
| Q9JHI5 | Q9JHI5 | Isovaleryl- | Ivd | sp\|Q9JHI5 | 15 | 15 | 36,8 |
| O55033 | O55033 | Cytoplasm | Nck2 | sp\|O5503: | 9 | 9 | 38,7 |
| Q8CB62 | Q8CB62 | Centrobin | Cntrob | sp\|Q8CB62 | 3 | 3 | 5,2 |
| Q640M1 | Q640M1 | U3 small n | Utp14a | sp\|Q640M | 26 | 23 | 39,6 |
| Q9ET30 | Q9ET30 | Transmem | Tm9sf3 | sp\|Q9ET3( | 13 | 13 | 23,5 |
| Q6ZQA6 | Q6ZQA6 | Immunogl | Igsf3 | sp\|Q6ZQA | 21 | 21 | 24,9 |
| P17225;B2 | P17225 | Polypyrimi | Ptbp1 | sp\|P17225 | 20 | 17 | 56,4 |
| Q80YQ2 | Q80YQ2 | Mediator c | Med23 | sp\|Q80YQ | 24 | 24 | 21,8 |
| E9Q4N7 | E9Q4N7 | | Arid1b | sp\|E9Q4N | 35 | 33 | 20,7 |
| Q3UZ39 | Q3UZ39 | Leucine-ric | Lrrfip1 | sp\|Q3UZ3! | 28 | 27 | 53,8 |
| Q8R3S6 | Q8R3S6 | Exocyst co | Exoc1 | sp\|Q8R3S€ | 18 | 18 | 26,1 |
| Q8BSN5 | Q8BSN5 | PHD finger | Phf23 | sp\|Q8BSN! | 5 | 5 | 20,4 |
| Q8C0E2 | Q8C0E2 | Vacuolar p | Vps26b | sp\|Q8C0E2 | 12 | 11 | 46,1 |
| Q8CFI0 | Q8CFI0 | E3 ubiquiti | Nedd4l | sp\|Q8CFI0 | 22 | 19 | 29 |
| Q99J45 | Q99J45 | Nuclear re | Nrbp1 | sp\|Q99J45 | 14 | 14 | 35 |
| P15208 | P15208 | Insulin rec | Insr | sp\|P15208 | 10 | 7 | 9 |

| | | | | | | | |
|---|---|---|---|---|---|---|---|
| P56581 | P56581 | G/T misma | Tdg | sp\|P56581 | 5 | 5 | 13,5 |
| Q3URS9 | Q3URS9 | Coiled-coil | Ccdc51 | sp\|Q3URS | 9 | 9 | 23,2 |
| Q9CQN3 | Q9CQN3 | Mitochond | Tomm6 | sp\|Q9CQN | 1 | 1 | 18,9 |
| P62317 | P62317 | Small nucle | Snrpd2 | sp\|P62317 | 9 | 9 | 61,9 |
| P31650 | P31650 | Sodium- ar | Slc6a11 | sp\|P31650 | 2 | 1 | 2,6 |
| P40336 | P40336 | Vacuolar p | Vps26a | sp\|P40336 | 17 | 16 | 64,5 |
| Q9D8H7 | Q9D8H7 | Metalloen | Oma1 | sp\|Q9D8H | 5 | 5 | 12,1 |
| Q62167;P1 | Q62167;P1638 | ATP-depen | Ddx3x;D1P | sp\|Q62167 | 40 | 12 | 71,3 |
| Q924A2 | Q924A2 | Protein cap | Cic | sp\|Q924A2 | 17 | 17 | 12,4 |
| Q9DBC3 | Q9DBC3 | Cap-specifi | Cmtr1 | sp\|Q9DBC | 25 | 25 | 38,4 |
| Q9D0D3 | Q9D0D3 | Poly(A) RN | Mtpap | sp\|Q9D0D | 15 | 15 | 37,8 |
| Q7TN02 | Q7TN02 | Mediator c | Med26 | sp\|Q7TN0 | 3 | 3 | 8,5 |
| Q9WUB4 | Q9WUB4 | Dynactin s | Dctn6 | sp\|Q9WUE | 2 | 2 | 16,8 |
| Q9D1H6 | Q9D1H6 | NADH deh | Ndufaf4 | sp\|Q9D1H | 6 | 6 | 33,5 |
| Q9D0R8 | Q9D0R8 | Protein LSI | Lsm12 | sp\|Q9D0R | 4 | 4 | 32,8 |
| Q921X6 | Q921X6 | DNA-direct | Polr3f | sp\|Q921X6 | 5 | 5 | 19 |
| Q8C7V3 | Q8C7V3 | U3 small n | Utp15 | sp\|Q8C7V | 22 | 21 | 52,1 |
| Q61216 | Q61216 | Double-str | Mre11a | sp\|Q61216 | 24 | 24 | 40,8 |
| Q61081 | Q61081 | Hsp90 co-c | Cdc37 | sp\|Q61081 | 16 | 16 | 39,1 |
| Q9CWI3 | Q9CWI3 | BRCA2 and | Bccip | sp\|Q9CWI | 10 | 10 | 43,4 |
| O35316;P3 | O35316 | Sodium- ar | Slc6a6 | sp\|O35316 | 5 | 4 | 9,7 |
| E9PY46 | E9PY46 | | Ift140 | sp\|E9PY46 | 6 | 6 | 5,7 |
| Q91YI0 | Q91YI0 | Argininosu | Asl | sp\|Q91YI0 | 11 | 11 | 32,3 |
| P41971 | P41971 | ETS domai | Elk3 | sp\|P41971 | 3 | 3 | 9,5 |
| P09041 | P09041 | Phosphogl | Pgk2 | sp\|P09041 | 10 | 1 | 19,9 |
| P80313 | P80313 | T-complex | Cct7 | sp\|P80313 | 38 | 38 | 74,4 |
| Q8BN78 | Q8BN78 | Transcripti | Zbtb33 | sp\|Q8BN7 | 6 | 6 | 14,6 |
| Q8VEE0 | Q8VEE0 | Ribulose-p | Rpe | sp\|Q8VEE | 2 | 2 | 17,5 |
| Q91ZM2 | Q91ZM2 | SH2B adap | Sh2b1 | sp\|Q91ZM | 10 | 9 | 24,3 |
| Q9CRW3 | Q9CRW3 | UPF0538 protein C2or | | sp\|Q9CRW | 3 | 3 | 29,4 |
| Q8CIF6 | Q8CIF6 | SID1 trans | Sidt2 | sp\|Q8CIF6 | 5 | 5 | 8,3 |
| Q7TQH7 | Q7TQH7 | Low-densit | Lrp10 | sp\|Q7TQH | 4 | 4 | 11,9 |
| E9Q4P1 | E9Q4P1 | | Wdfy1 | sp\|E9Q4P1 | 8 | 8 | 28,8 |
| Q6SKR2 | Q6SKR2 | | N6amt1 | sp\|Q6SKR2 | 6 | 6 | 41,6 |
| Q91WT4 | Q91WT4 | DnaJ homo | Dnajc17 | sp\|Q91WT | 5 | 5 | 20,1 |
| O08967 | O08967 | Cytohesin- | Cyth3 | sp\|O08967 | 10 | 6 | 31,1 |
| Q8R2R1 | Q8R2R1 | Protein O- | Pomt1 | sp\|Q8R2R1 | 2 | 2 | 3,6 |
| E9Q634 | E9Q634 | Unconvent | Myo1e | sp\|E9Q634 | 40 | 36 | 45,1 |
| P62488 | P62488 | DNA-direct | Polr2g | sp\|P62488 | 7 | 7 | 54,7 |
| P41234 | P41234 | ATP-bindin | Abca2 | sp\|P41234 | 4 | 3 | 2,5 |
| CON__Q03 | CON__Q03247 | | | | 6 | 6 | 26,9 |
| Q8K2K6 | Q8K2K6 | Arf-GAP do | Agfg1 | sp\|Q8K2K6 | 8 | 8 | 23,4 |
| Q63829 | Q63829 | COMM dor | Commd3 | sp\|Q63829 | 12 | 12 | 66,2 |
| Q8BH43 | Q8BH43 | Wiskott-Al | Wasf2 | sp\|Q8BH4 | 11 | 10 | 27,4 |
| Q52KR3 | Q52KR3 | Protein pru | Prune2 | sp\|Q52KR3 | 6 | 6 | 3,5 |
| Q3TUH1 | Q3TUH1 | Phosphatid | Tamm41 | sp\|Q3TUH | 10 | 10 | 31,2 |
| Q80Y81 | Q80Y81 | Zinc phosp | Elac2 | sp\|Q80Y81 | 23 | 23 | 37,8 |
| Q91YE7 | Q91YE7 | RNA-bindir | Rbm5 | sp\|Q91YE7 | 16 | 12 | 21,3 |
| Q9CR08 | Q9CR08 | Ribonuclea | Pop4 | sp\|Q9CR08 | 6 | 6 | 29,4 |
| Q9D6Y7 | Q9D6Y7 | Mitochond | Msra | sp\|Q9D6Y | 7 | 7 | 44,6 |

| | | | | | | | |
|---|---|---|---|---|---|---|---|
| Q91WL8 | Q91WL8 | WW doma | Wwox | sp\|Q91WL | 6 | 6 | 18,8 |
| Q80VP1 | Q80VP1 | Epsin-1 | Epn1 | sp\|Q80VP1 | 7 | 6 | 16,2 |
| Q8K0L9 | Q8K0L9 | Zinc finger | Zbtb20 | sp\|Q8K0L9 | 8 | 8 | 22 |
| Q9DBR4 | Q9DBR4 | Amyloid be | Apbb2 | sp\|Q9DBR | 15 | 14 | 23,7 |
| Q3TZX8 | Q3TZX8 | Polynucleo | Nol9 | sp\|Q3TZX8 | 13 | 13 | 22,7 |
| Q924M7 | Q924M7 | Mannose-6 | Mpi | sp\|Q924M | 11 | 11 | 39,7 |
| P25444 | P25444 | 40S riboso | Rps2 | sp\|P25444 | 18 | 18 | 61,8 |
| Q62189 | Q62189 | U1 small n | Snrpa | sp\|Q62189 | 11 | 9 | 40,4 |
| Q91WG5 | Q91WG5 | 5-AMP-act | Prkag2 | sp\|Q91WG | 6 | 5 | 16,3 |
| Q9JKP8 | Q9JKP8 | Chromatin | Chrac1 | sp\|Q9JKP8 | 5 | 5 | 52,7 |
| Q80U56 | Q80U56 | Late secret | Avl9 | sp\|Q80U56 | 3 | 3 | 4,9 |
| Q9D7B1 | Q9D7B1 | tRNA-dihyd | Dus2 | sp\|Q9D7B | 6 | 6 | 17,8 |
| P11881 | P11881 | Inositol 1,4 | Itpr1 | sp\|P11881 | 27 | 20 | 13,5 |
| P43274 | P43274 | Histone H1 | Hist1h1e | sp\|P43274 | 14 | 4 | 47 |
| Q80XJ3 | Q80XJ3 | Tetratricop | Ttc28 | sp\|Q80XJ3 | 2 | 2 | 1,4 |
| Q5U4D9 | Q5U4D9 | THO comp | Thoc6 | sp\|Q5U4D | 11 | 11 | 48,1 |
| P11983 | P11983 | T-complex | Tcp1 | sp\|P11983 | 36 | 36 | 80,6 |
| Q8VC57;Q | Q8VC57 | BTB/POZ d | Kctd5 | sp\|Q8VC57 | 9 | 9 | 62,8 |
| Q9D1K7 | Q9D1K7 | UPF0687 protein C20 | | sp\|Q9D1K7 | 4 | 4 | 39,1 |
| P51150 | P51150 | Ras-related | Rab7a | sp\|P51150 | 18 | 18 | 79,7 |
| Q8VEA8 | Q8VEA8 | Ras-related | Rab7b | sp\|Q8VEA8 | 5 | 5 | 36,2 |
| Q8BH60 | Q8BH60 | Golgi-asso | Gopc | sp\|Q8BH60 | 8 | 8 | 24 |
| Q3UVG3 | Q3UVG3 | Protein FA | Fam91a1 | sp\|Q3UVG | 17 | 17 | 29,7 |
| Q91X21 | Q91X21 | Uncharacte | Kiaa2013 | sp\|Q91X21 | 10 | 10 | 25,1 |
| Q9QXK2 | Q9QXK2 | E3 ubiquiti | Rad18 | sp\|Q9QXK | 13 | 13 | 37,3 |
| P70122 | P70122 | Ribosome | Sbds | sp\|P70122 | 17 | 17 | 59,6 |
| P13808;P1 | P13808 | Anion exch | Slc4a2 | sp\|P13808 | 3 | 3 | 4,5 |
| O70325 | O70325 | Phospholip | Gpx4 | sp\|O70325 | 8 | 8 | 40,1 |
| Q8BSL7 | Q8BSL7 | ADP-ribosy | Arf2 | sp\|Q8BSL7 | 11 | 2 | 58,6 |
| Q5RJG1 | Q5RJG1 | Nucleolar p | Nol10 | sp\|Q5RJG1 | 25 | 25 | 48,6 |
| Q9D273 | Q9D273 | Cob(I)yrini | Mmab | sp\|Q9D273 | 4 | 4 | 23,2 |
| Q69ZQ2 | Q69ZQ2 | Pre-mRNA- | Isy1 | sp\|Q69ZQ2 | 7 | 7 | 28,8 |
| Q6NZQ2 | Q6NZQ2 | | Ddx31 | sp\|Q6NZQ | 20 | 20 | 41,2 |
| Q8C172 | Q8C172 | Ceramide s | Cers6 | sp\|Q8C172 | 5 | 5 | 18,2 |
| Q8R5F3 | Q8R5F3 | O-acetyl-A | Oard1 | sp\|Q8R5F3 | 4 | 4 | 28,3 |
| Q9JHR7 | Q9JHR7 | Insulin-deg | Ide | sp\|Q9JHR7 | 35 | 35 | 33,7 |
| P37172 | P37172 | Activin rec | Acvr1 | sp\|P37172 | 3 | 3 | 7,3 |
| Q8VC70 | Q8VC70 | RNA-bindir | Rbms2 | sp\|Q8VC70 | 6 | 6 | 19,3 |
| Q64442 | Q64442 | Sorbitol de | Sord | sp\|Q64442 | 6 | 6 | 24,9 |
| Q0VBD2 | Q0VBD2 | Protein MC | Mcm10 | sp\|Q0VBD | 5 | 5 | 10,1 |
| Q9D6N1 | Q9D6N1 | Carbonic a | Ca13 | sp\|Q9D6N | 7 | 7 | 42,7 |
| Q9QZA0 | Q9QZA0 | Carbonic a | Ca5b | sp\|Q9QZA | 4 | 4 | 15,5 |
| P58158 | P58158 | Galactosylg | B3gat3 | sp\|P58158 | 2 | 2 | 11,6 |
| Q91WE4 | Q91WE4 | UPF0729 protein C18 | | sp\|Q91WE | 1 | 1 | 16,7 |
| Q3UEB3 | Q3UEB3 | Poly(U)-bir | Puf60 | sp\|Q3UEB | 26 | 26 | 64,4 |
| Q3TQQ9 | Q3TQQ9 | Uncharacterized prote | | sp\|Q3TQQ | 6 | 6 | 9 |
| Q61072 | Q61072 | Disintegrin | Adam9 | sp\|Q61072 | 17 | 17 | 27 |
| Q9D009 | Q9D009 | Putative lip | Lipt2 | sp\|Q9D009 | 1 | 1 | 8,2 |
| Q5RL79 | Q5RL79 | Keratinocy | Krtcap2 | sp\|Q5RL79 | 1 | 1 | 12,5 |
| Q9D0I6 | Q9D0I6 | WD repeat | Wdsub1 | sp\|Q9D0I6 | 4 | 4 | 11 |

| | | | | | | | |
|---|---|---|---|---|---|---|---|
| Q8BMJ2 | Q8BMJ2 | Leucine--tR | Lars | sp\|Q8BMJ | 53 | 53 | 52,8 |
| Q8VCI5 | Q8VCI5 | Peroxisom | Pex19 | sp\|Q8VCI5 | 8 | 8 | 48,2 |
| O88456;Q9 | O88456 | Calpain sm | Capns1 | sp\|O88456 | 12 | 12 | 58,7 |
| Q9DB28 | Q9DB28 | Ribonuclea | Pop5 | sp\|Q9DB28 | 3 | 3 | 18,3 |
| Q8BSQ9 | Q8BSQ9 | Protein po | Pbrm1 | sp\|Q8BSQ | 49 | 49 | 40,1 |
| Q91YX5 | Q91YX5 | Acyl-CoA:ly | Lpgat1 | sp\|Q91YX5 | 11 | 11 | 32,7 |
| Q8R2U6;P0 | Q8R2U6 | Diphospho | Nudt4 | sp\|Q8R2U | 6 | 5 | 39,7 |
| Q08639 | Q08639 | Transcripti | Tfdp1 | sp\|Q08639 | 7 | 7 | 30 |
| Q9CW79 | Q9CW79 | Golgin sub | Golga1 | sp\|Q9CW7 | 19 | 19 | 35,1 |
| Q80WQ2 | Q80WQ2 | Protein VA | Vac14 | sp\|Q80WQ | 17 | 17 | 30,1 |
| Q9ES34 | Q9ES34 | Ubiquitin-p | Ube3b | sp\|Q9ES34 | 2 | 2 | 1,8 |
| Q80UK0 | Q80UK0 | SEC14 dom | Sestd1 | sp\|Q80UK0 | 14 | 14 | 28,3 |
| Q9D8L0 | Q9D8L0 | Uncharacterized prote | | sp\|Q9D8L0 | 2 | 2 | 21,3 |
| P28076 | P28076 | Proteasom | Psmb9 | sp\|P28076 | 4 | 4 | 29,7 |
| Q8BR65 | Q8BR65 | Sin3 histor | Suds3 | sp\|Q8BR65 | 6 | 6 | 26,8 |
| Q61026 | Q61026 | Nuclear re | Ncoa2 | sp\|Q61026 | 7 | 7 | 6,8 |
| P47801 | P47801 | Epithelial r | Emp1 | sp\|P47801 | 1 | 1 | 12,5 |
| P80318 | P80318 | T-complex | Cct3 | sp\|P80318 | 37 | 37 | 63,7 |
| A2RSX7 | A2RSX7 | tRNA wybu | Tyw5 | sp\|A2RSX7 | 6 | 6 | 27,9 |
| Q9CZI9 | Q9CZI9 | Apoptosis- | Aen | sp\|Q9CZI9 | 8 | 8 | 33,3 |
| Q9R049 | Q9R049 | E3 ubiquiti | Amfr | sp\|Q9R049 | 11 | 11 | 24 |
| Q9CQF3 | Q9CQF3 | Cleavage a | Nudt21 | sp\|Q9CQF3 | 12 | 12 | 44,9 |
| Q8CBW3 | Q8CBW3 | Abl interac | Abi1 | sp\|Q8CBW | 17 | 13 | 34,5 |
| Q5XG73 | Q5XG73 | Acyl-CoA-b | Acbd5 | sp\|Q5XG73 | 10 | 10 | 24 |
| Q8R3L2 | Q8R3L2 | Transcripti | Tcf25 | sp\|Q8R3L2 | 12 | 12 | 25,4 |
| Q9D7N9 | Q9D7N9 | Adipocyte | Apmap | sp\|Q9D7N | 16 | 16 | 54,7 |
| P70333 | P70333 | Heterogen | Hnrnph2 | sp\|P70333 | 15 | 7 | 41,9 |
| Q9D1Z3 | Q9D1Z3 | Protein FA | Fam173b | sp\|Q9D1Z3 | 4 | 4 | 27,1 |
| Q9D4H2 | Q9D4H2 | GRIP and c | Gcc1 | sp\|Q9D4H | 8 | 8 | 14,8 |
| A6X8Z5 | A6X8Z5 | Rho GTPas | Arhgap31 | sp\|A6X8Z5 | 8 | 8 | 8,4 |
| P84104 | P84104 | Serine/arg | Srsf3 | sp\|P84104 | 8 | 7 | 43,3 |
| Q80SW1 | Q80SW1 | Putative ad | Ahcyl1 | sp\|Q80SW | 17 | 8 | 35,7 |
| Q9DAS9 | Q9DAS9 | Guanine nu | Gng12 | sp\|Q9DAS9 | 5 | 5 | 69,4 |
| Q91WR3 | Q91WR3 | Activating | Ascc2 | sp\|Q91WR | 12 | 12 | 21,8 |
| P97477 | P97477 | Aurora kin | Aurka | sp\|P97477 | 9 | 8 | 32,9 |
| Q8BG79 | Q8BG79 | CWF19-like | Cwf19l2 | sp\|Q8BG79 | 14 | 14 | 18,6 |
| Q8BFV2 | Q8BFV2 | PCI domain | Pcid2 | sp\|Q8BFV2 | 17 | 17 | 57,1 |
| Q9JKB1;P5 | Q9JKB1;P5832 | Ubiquitin c | Uchl3;Uchl | sp\|Q9JKB1 | 12 | 12 | 72,6 |
| Q922R8 | Q922R8 | Protein dis | Pdia6 | sp\|Q922R8 | 21 | 21 | 61,1 |
| Q8CGZ0 | Q8CGZ0 | Calcium ho | Cherp | sp\|Q8CGZ0 | 23 | 23 | 26 |
| Q925J9 | Q925J9 | Mediator o | Med1 | sp\|Q925J9 | 29 | 29 | 27,6 |
| P70288 | P70288 | Histone de | Hdac2 | sp\|P70288 | 12 | 8 | 36,5 |
| Q8BH65 | Q8BH65 | Protein DE | Dennd6a | sp\|Q8BH65 | 4 | 4 | 10,7 |
| P83940 | P83940 | Transcripti | Tceb1 | sp\|P83940 | 5 | 5 | 57,1 |
| P32883 | P32883 | GTPase KR | Kras | sp\|P32883 | 7 | 2 | 41,8 |
| Q91VN0 | Q91VN0 | Low-densit | Lrp5 | sp\|Q91VN | 5 | 4 | 4,2 |
| Q3TFK5 | Q3TFK5 | G patch do | Gpatch4 | sp\|Q3TFK5 | 11 | 11 | 34,9 |
| Q920I9 | Q920I9 | WD repeat | Wdr7 | sp\|Q920I9 | 10 | 10 | 11,6 |
| O88522 | O88522 | NF-kappa-I | Ikbkg | sp\|O88522 | 7 | 7 | 22,8 |
| P15105 | P15105 | Glutamine | Glul | sp\|P15105 | 6 | 6 | 22,8 |

| | | | | | | | |
|---|---|---|---|---|---|---|---|
| Q8BP92 | Q8BP92 | Reticulocal | Rcn2 | sp\|Q8BP92 | 18 | 18 | 55,9 |
| Q9CQF6 | Q9CQF6 | L-aminoad | Aasdhppt | sp\|Q9CQF( | 9 | 9 | 37,2 |
| Q8C1M2 | Q8C1M2 | Zinc finger | Znf428 | sp\|Q8C1M | 3 | 3 | 27,3 |
| Q8C166 | Q8C166 | Copine-1 | Cpne1 | sp\|Q8C16( | 14 | 13 | 39 |
| Q61818 | Q61818 | Retinoic ac | Rai1 | sp\|Q61818 | 30 | 30 | 28,5 |
| Q9EPE9 | Q9EPE9 | Manganes( | Atp13a1 | sp\|Q9EPES | 38 | 38 | 41,2 |
| Q6P2L6 | Q6P2L6 | Histone-lys | Whsc1l1 | sp\|Q6P2L6 | 20 | 20 | 19 |
| Q91YM4 | Q91YM4 | Protein TB | Tbrg4 | sp\|Q91YM | 19 | 19 | 39,2 |
| Q3UPL0 | Q3UPL0 | Protein tra | Sec31a | sp\|Q3UPL( | 39 | 39 | 36,7 |
| Q7TPD2 | Q7TPD2 | Protein FA | Fam185a | sp\|Q7TPD2 | 3 | 3 | 10,3 |
| P59222 | P59222 | Scavenger | Scarf2 | sp\|P59222 | 5 | 5 | 8,5 |
| Q8R5J9 | Q8R5J9 | PRA1 famil | Arl6ip5 | sp\|Q8R5J9 | 6 | 6 | 28,7 |
| Q6NZB1 | Q6NZB1 | Protein arg | Prmt6 | sp\|Q6NZB1 | 6 | 6 | 24,6 |
| Q7TPD1 | Q7TPD1 | F-box only | Fbxo11 | sp\|Q7TPD1 | 9 | 9 | 13,9 |
| Q61672 | Q61672 | Equilibrativ | Slc29a2 | sp\|Q61672 | 1 | 1 | 2,6 |
| P52019 | P52019 | Squalene r | Sqle | sp\|P52019 | 6 | 6 | 15,9 |
| Q99KJ8 | Q99KJ8 | Dynactin s | Dctn2 | sp\|Q99KJ8 | 19 | 19 | 57,7 |
| Q9JLV2 | Q9JLV2 | Short trans | Trpc4ap | sp\|Q9JLV2 | 6 | 6 | 12,7 |
| O89016 | O89016 | ATP-bindin | Abcd4 | sp\|O89016 | 6 | 6 | 10,7 |
| Q8K1J5 | Q8K1J5 | Protein SD | Sde2 | sp\|Q8K1J5 | 7 | 7 | 17,6 |
| O09005 | O09005 | Sphingolipi | Degs1 | sp\|O09005 | 6 | 6 | 20,1 |
| Q91ZD4 | Q91ZD4 | Vang-like p | Vangl2 | sp\|Q91ZD4 | 2 | 2 | 4,8 |
| Q3URQ0 | Q3URQ0 | Testis-expr | Tex10 | sp\|Q3URQ | 21 | 21 | 29,2 |
| Q922R1 | Q922R1 | UPF0183 protein C16c | | sp\|Q922R1 | 5 | 5 | 17,3 |
| Q8VCG3 | Q8VCG3 | WD repeat | Wdr74 | sp\|Q8VCG | 15 | 15 | 58,6 |
| Q8BHJ5 | Q8BHJ5 | F-box-like/ | Tbl1xr1 | sp\|Q8BHJ5 | 17 | 11 | 48,2 |
| Q80TJ7 | Q80TJ7 | Histone lys | Phf8 | sp\|Q80TJ7 | 8 | 7 | 9,5 |
| Q60823 | Q60823 | RAC-beta s | Akt2 | sp\|Q60823 | 11 | 8 | 31,8 |
| P08003 | P08003 | Protein dis | Pdia4 | sp\|P08003 | 34 | 33 | 53,6 |
| Q3UPH7 | Q3UPH7 | Rho guanin | Arhgef40 | sp\|Q3UPH | 39 | 39 | 41,5 |
| D3YX43 | D3YX43 | V-set and i | Vsig10 | sp\|D3YX43 | 5 | 5 | 9,5 |
| P11499 | P11499 | Heat shock | Hsp90ab1 | sp\|P11499 | 50 | 31 | 67 |
| Q3TEI4 | Q3TEI4 | Uncharacterized prote | | sp\|Q3TEI4 | 6 | 6 | 9,4 |
| Q8VBW6 | Q8VBW6 | NEDD8-act | Nae1 | sp\|Q8VBW | 17 | 17 | 43,4 |
| P60122 | P60122 | RuvB-like 1 | Ruvbl1 | sp\|P60122 | 27 | 27 | 69,7 |
| Q8K2Z4 | Q8K2Z4 | Condensin | Ncapd2 | sp\|Q8K2Z4 | 48 | 48 | 46,8 |
| Q8K305 | Q8K305 | Kinetochor | Nsl1 | sp\|Q8K305 | 6 | 6 | 35,9 |
| Q80TN5 | Q80TN5 | Palmitoyltr | Zdhhc17 | sp\|Q80TN5 | 7 | 6 | 13 |
| Q9CQ91 | Q9CQ91 | NADH dehy | Ndufa3 | sp\|Q9CQ9 | 3 | 3 | 41,7 |
| Q8VCE9 | Q8VCE9 | Pleckstrin I | Plekhh3 | sp\|Q8VCES | 7 | 7 | 12,4 |
| Q91WK2 | Q91WK2 | Eukaryotic | Eif3h | sp\|Q91WK | 24 | 24 | 76,4 |
| Q61207 | Q61207 | Prosaposin | Psap | sp\|Q61207 | 16 | 16 | 35,9 |
| Q9DBB4 | Q9DBB4 | N-alpha-ac | Naa16 | sp\|Q9DBB | 11 | 7 | 14,6 |
| O88796 | O88796 | Ribonuclea | Rpp30 | sp\|O88796 | 12 | 12 | 57,1 |
| Q9CQA9 | Q9CQA9 | Cancer-rela | Ntpcr | sp\|Q9CQA | 7 | 7 | 55,3 |
| Q5NCE8 | Q5NCE8 | Magnesiur | Mrs2 | sp\|Q5NCE | 5 | 5 | 17,3 |
| Q99KY4 | Q99KY4 | Cyclin-G-as | Gak | sp\|Q99KY4 | 21 | 21 | 21,7 |
| Q8BYR2;Q | Q8BYR2 | Serine/thre | Lats1 | sp\|Q8BYR2 | 8 | 8 | 8,6 |
| Q99J10 | Q99J10 | Cytoplasm | Ctu1 | sp\|Q99J10 | 4 | 4 | 14,5 |
| Q61598 | Q61598 | Rab GDP d | Gdi2 | sp\|Q61598 | 38 | 32 | 77,8 |

| | | | | | | | |
|---|---|---|---|---|---|---|---|
| Q9DBR1 | Q9DBR1 | 5-3 exoribo | Xrn2 | sp\|Q9DBR | 41 | 41 | 46,5 |
| Q2TBE6 | Q2TBE6 | Phosphatid | Pi4k2a | sp\|Q2TBE6 | 16 | 16 | 40,3 |
| Q921M4 | Q921M4 | Golgin sub | Golga2 | sp\|Q921M | 22 | 22 | 35,8 |
| Q8C5K5 | Q8C5K5 | Uncharacterized prote | | sp\|Q8C5K5 | 7 | 7 | 26,6 |
| P59266 | P59266 | Fat storage | Fitm2 | sp\|P59266 | 2 | 2 | 8,4 |
| Q9CR98 | Q9CR98 | Protein FA | Fam136a | sp\|Q9CR98 | 5 | 5 | 46,4 |
| Q80YR7 | Q80YR7 | Claspin | Clspn | sp\|Q80YR7 | 10 | 10 | 10,5 |
| Q3TKY6 | Q3TKY6 | Peptidyl-pr | Cwc27 | sp\|Q3TKY6 | 11 | 11 | 35,6 |
| O35685 | O35685 | Nuclear mi | Nudc | sp\|O35685 | 21 | 21 | 53,6 |
| P60904 | P60904 | DnaJ homo | Dnajc5 | sp\|P60904 | 6 | 6 | 54,5 |
| O35215 | O35215 | D-dopachr | Ddt | sp\|O35215 | 6 | 6 | 70,3 |
| Q3UVK0 | Q3UVK0 | Endoplasm | Ermp1 | sp\|Q3UVK | 15 | 15 | 24,4 |
| P48678 | P48678 | Prelamin-A | Lmna | sp\|P48678 | 60 | 59 | 78,6 |
| Q91ZW2 | Q91ZW2 | GDP-fucos | Pofut1 | sp\|Q91ZW | 14 | 14 | 45,5 |
| Q924Z6 | Q924Z6 | Exportin-6 | Xpo6 | sp\|Q924Z6 | 14 | 14 | 16,7 |
| Q9D1M7 | Q9D1M7 | Peptidyl-pr | Fkbp11 | sp\|Q9D1M | 7 | 7 | 35,3 |
| Q78ZA7 | Q78ZA7 | Nucleosom | Nap1l4 | sp\|Q78ZA7 | 14 | 13 | 57,6 |
| Q8BFR6 | Q8BFR6 | AN1-type z | Zfand1 | sp\|Q8BFR6 | 5 | 5 | 27,2 |
| P97470 | P97470 | Serine/thre | Ppp4c | sp\|P97470 | 13 | 11 | 59,3 |
| Q8BWQ4 | Q8BWQ4 | Cap-specifi | Cmtr2 | sp\|Q8BWC | 7 | 7 | 13,6 |
| Q62172 | Q62172 | RalA-bindi | Ralbp1 | sp\|Q62172 | 3 | 3 | 7,3 |
| Q61090 | Q61090 | Frizzled-7 | Fzd7 | sp\|Q61090 | 3 | 2 | 4,4 |
| Q8BQ47 | Q8BQ47 | Protein car | Cnpy4 | sp\|Q8BQ4 | 8 | 8 | 33,5 |
| Q61464 | Q61464 | Zinc finger | Znf638 | sp\|Q61464 | 41 | 41 | 27,5 |
| Q8BJW5 | Q8BJW5 | Nucleolar p | Nol11 | sp\|Q8BJW | 20 | 20 | 32,2 |
| P52432 | P52432 | DNA-direct | Polr1c | sp\|P52432 | 9 | 9 | 35,3 |
| Q0VGB7 | Q0VGB7 | Serine/thre | Ppp4r2 | sp\|Q0VGB | 17 | 17 | 56,4 |
| Q8VEG4 | Q8VEG4 | Exonucleas | Exd2 | sp\|Q8VEG4 | 11 | 11 | 24 |
| O88532 | O88532 | Zinc finger | Zfr | sp\|O88532 | 35 | 35 | 48,8 |
| Q9DBU0 | Q9DBU0 | Transmem | Tm9sf1 | sp\|Q9DBU | 7 | 7 | 15,8 |
| Q80TE4 | Q80TE4 | Signal-indu | Sipa1l2 | sp\|Q80TE4 | 9 | 9 | 6,3 |
| Q99M87 | Q99M87 | DnaJ homo | Dnaja3 | sp\|Q99M8 | 10 | 10 | 23,3 |
| Q9Z0R4 | Q9Z0R4 | Intersectin | Itsn1 | sp\|Q9Z0R4 | 21 | 14 | 16,7 |
| Q9CX84 | Q9CX84 | Regulator o | Rgs19 | sp\|Q9CX84 | 2 | 2 | 9,7 |
| Q9CQY6 | Q9CQY6 | Ubiquinol-c | Uqcc2 | sp\|Q9CQY | 5 | 5 | 41,9 |
| Q60660 | Q60660 | Killer cell le | Klra2 | sp\|Q60660 | 2 | 2 | 9,7 |
| Q8C170 | Q8C170 | Unconvent | Myo9a | sp\|Q8C170 | 8 | 6 | 3,8 |
| Q9CR75 | Q9CR75 | Tumor nec | Tnfrsf12a | sp\|Q9CR75 | 2 | 2 | 29,5 |
| Q922U1;R | Q922U1 | U4/U6 sma | Prpf3 | sp\|Q922U | 27 | 27 | 45,7 |
| Q8C6I2 | Q8C6I2 | Succinate d | Sdhaf2 | sp\|Q8C6I2 | 5 | 5 | 37,2 |
| Q8CH72 | Q8CH72 | E3 ubiquiti | Trim32 | sp\|Q8CH72 | 19 | 19 | 43,5 |
| P25446 | P25446 | Tumor nec | Fas | sp\|P25446 | 2 | 2 | 7,6 |
| Q8BL80 | Q8BL80 | Rho GTPas | Arhgap22 | sp\|Q8BL80 | 15 | 15 | 28,5 |
| O70503 | O70503 | Very-long-c | Hsd17b12 | sp\|O70503 | 14 | 14 | 53,2 |
| Q9CSN1 | Q9CSN1 | SNW doma | Snw1 | sp\|Q9CSN1 | 23 | 23 | 47 |
| Q8CHE4 | Q8CHE4 | PH domain | Phlpp1 | sp\|Q8CHE4 | 5 | 5 | 5,8 |
| Q99KI0 | Q99KI0 | Aconitate h | Aco2 | sp\|Q99KI0 | 37 | 37 | 55,1 |
| P21981 | P21981 | Protein-glu | Tgm2 | sp\|P21981 | 18 | 18 | 39,9 |
| Q3UMT1 | Q3UMT1 | Protein ph | Ppp1r12c | sp\|Q3UMT | 13 | 13 | 21,4 |
| Q5SYL3 | Q5SYL3 | Protein KIA | Kiaa0100 | sp\|Q5SYL3 | 6 | 6 | 3,2 |

| | | | | | | | |
|---|---|---|---|---|---|---|---|
| Q9QZ05 | Q9QZ05 | Eukaryotic | Eif2ak4 | sp\|Q9QZ0 | 9 | 9 | 6,7 |
| Q32P12 | Q32P12 | Uncharacterized prot | | sp\|Q32P12 | 7 | 7 | 19,8 |
| O70401 | O70401 | Tetraspani | Tspan6 | sp\|O70401 | 2 | 2 | 12,2 |
| P59114 | P59114 | Phosphory | Pcif1 | sp\|P59114 | 14 | 14 | 27,3 |
| Q9Z0E6 | Q9Z0E6 | Interferon- | Gbp2 | sp\|Q9Z0E6 | 4 | 3 | 10,7 |
| Q6IQX8 | Q6IQX8 | | Zfp219 | sp\|Q6IQX8 | 8 | 8 | 20,5 |
| Q925E7 | Q925E7 | Serine/thre | Ppp2r2d | sp\|Q925E7 | 14 | 5 | 33,6 |
| Q3TTL0 | Q3TTL0 | Uncharacterized prot | | sp\|Q3TTL0 | 4 | 4 | 16,7 |
| Q9Z2U4 | Q9Z2U4 | ETS-relate | Elf4 | sp\|Q9Z2U | 3 | 3 | 7,6 |
| P09925 | P09925 | Surfeit loc | Surf1 | sp\|P09925 | 5 | 5 | 23,5 |
| Q8R2T8 | Q8R2T8 | General tr | Gtf3c5 | sp\|Q8R2T | 11 | 11 | 26,9 |
| Q8BJU2 | Q8BJU2 | Tetraspani | Tspan9 | sp\|Q8BJU2 | 2 | 2 | 10,9 |
| Q7TSZ8 | Q7TSZ8 | Nucleus ac | Nacc1 | sp\|Q7TSZ8 | 15 | 14 | 37,5 |
| Q6F3F9 | Q6F3F9 | G-protein | Gpr126 | sp\|Q6F3F9 | 3 | 3 | 3,5 |
| Q75NR7 | Q75NR7 | ATP-depen | Recql4 | sp\|Q75NR | 6 | 6 | 6,8 |
| P70168 | P70168 | Importin s | Kpnb1 | sp\|P70168 | 39 | 39 | 54,5 |
| Q61188 | Q61188 | Histone-ly | Ezh2 | sp\|Q6118 | 20 | 18 | 37,7 |
| Q6PDM2 | Q6PDM2 | Serine/arg | Srsf1 | sp\|Q6PDM | 18 | 18 | 54,4 |
| Q06138 | Q06138 | Calcium-bi | Cab39 | sp\|Q0613 | 13 | 11 | 37,2 |
| P55200 | P55200 | Histone-ly | Kmt2a | sp\|P55200 | 39 | 39 | 18 |
| CON__P00 | CON__P00735 | | | | 6 | 6 | 10,6 |
| Q6NZQ4 | Q6NZQ4 | PAX-intera | Paxip1 | sp\|Q6NZQ | 12 | 12 | 17 |
| Q8K353 | Q8K353 | Cysteine-ri | Cystm1 | sp\|Q8K353 | 1 | 1 | 9,6 |
| B2RWS6 | B2RWS6 | Histone ac | Ep300 | sp\|B2RWS | 25 | 16 | 13,7 |
| Q91VA6 | Q91VA6 | Polymeras | Poldip2 | sp\|Q91VA | 15 | 15 | 55,2 |
| Q69Z37 | Q69Z37 | Sterile alp | Samd9l | sp\|Q69Z37 | 10 | 10 | 7,9 |
| Q8BMI3 | Q8BMI3 | ADP-ribos | Gga3 | sp\|Q8BMI | 11 | 10 | 25,1 |
| Q9QY06 | Q9QY06 | Unconvent | Myo9b | sp\|Q9QY0 | 41 | 39 | 28,4 |
| Q3U108 | Q3U108 | AT-rich int | Arid5a | sp\|Q3U10 | 3 | 3 | 6,4 |
| Q9Z2A5 | Q9Z2A5 | Arginyl-tR | Ate1 | sp\|Q9Z2A5 | 10 | 10 | 30,6 |
| P40237 | P40237 | CD82 antig | Cd82 | sp\|P40237 | 3 | 3 | 12,8 |
| P97411 | P97411 | Islet cell a | Ica1 | sp\|P97411 | 5 | 5 | 12,1 |
| Q9QY30 | Q9QY30 | Bile salt ex | Abcb11 | sp\|Q9QY3 | 1 | 1 | 1,3 |
| Q7TMF2 | Q7TMF2 | 3-5 exorib | Eri1 | sp\|Q7TMF | 9 | 9 | 35,4 |
| Q91ZU1 | Q91ZU1 | Ankyrin re | Asb6 | sp\|Q91ZU | 7 | 7 | 21,1 |
| Q9DAT5 | Q9DAT5 | Mitochond | Trmu | sp\|Q9DAT | 11 | 11 | 33,8 |
| Q05512 | Q05512 | Serine/thr | Mark2 | sp\|Q05512 | 19 | 14 | 32,9 |
| Q505F1 | Q505F1 | Nuclear re | Nr2c1 | sp\|Q505F1 | 7 | 7 | 17,5 |
| O70472 | O70472 | Transmem | Tmem131 | sp\|O70472 | 18 | 18 | 12,7 |
| Q9DC53;Q | Q9DC53 | Copine-8 | Cpne8 | sp\|Q9DC5 | 19 | 18 | 46,6 |
| Q9WUM4; | Q9WUM4 | Coronin-1 | Coro1c | sp\|Q9WU | 15 | 15 | 43,5 |
| Q8BH49 | Q8BH49 | Sesquiped | Fam109a | sp\|Q8BH4 | 3 | 3 | 22,6 |
| Q64735 | Q64735 | Compleme | Cr1l | sp\|Q64735 | 4 | 4 | 15,5 |
| CON__Q28 | CON__Q28085 | | | | 6 | 6 | 8,3 |
| Q9DBM1 | Q9DBM1 | G patch do | Gpatch1 | sp\|Q9DBM | 19 | 19 | 30,9 |
| Q8VE11 | Q8VE11 | Myotubula | Mtmr6 | sp\|Q8VE1 | 10 | 10 | 20,7 |
| Q8VI63 | Q8VI63 | MOB kinas | Mob2 | sp\|Q8VI63 | 6 | 6 | 37 |
| Q8VEA4 | Q8VEA4 | Mitochond | Chchd4 | sp\|Q8VEA | 3 | 3 | 36,7 |
| Q8VCF0 | Q8VCF0 | Mitochond | Mavs | sp\|Q8VCF | 13 | 13 | 59,4 |
| Q7TPQ3 | Q7TPQ3 | E3 ubiquit | Shprh | sp\|Q7TPQ | 22 | 22 | 15,7 |

| | | | | | | | |
|---|---|---|---|---|---|---|---|
| Q8R2U4 | Q8R2U4 | N-terminal | Ntmt1 | sp\|Q8R2U | 8 | 8 | 45,3 |
| Q8CEI1 | Q8CEI1 | BolA-like p | Bola3 | sp\|Q8CEI1 | 3 | 3 | 40 |
| Q8VCH8 | Q8VCH8 | UBX doma | Ubxn4 | sp\|Q8VCH | 7 | 7 | 26,5 |
| P97452 | P97452 | Ribosome | Bop1 | sp\|P97452 | 28 | 28 | 57,7 |
| Q8K0D0 | Q8K0D0 | Cyclin-dep | Cdk17 | sp\|Q8K0D | 13 | 8 | 32,9 |
| Q91WS0 | Q91WS0 | CDGSH iro | Cisd1 | sp\|Q91WS | 5 | 5 | 44,4 |
| Q3U821 | Q3U821 | | Wdr75 | sp\|Q3U82 | 27 | 27 | 46,1 |
| Q6P2B1 | Q6P2B1 | Transporti | Tnpo3 | sp\|Q6P2B | 23 | 22 | 35 |
| Q9D7S9 | Q9D7S9 | Charged m | Chmp5 | sp\|Q9D7S | 5 | 5 | 28,3 |
| P27773 | P27773 | Protein dis | Pdia3 | sp\|P27773 | 44 | 43 | 72,5 |
| A2RSY1 | A2RSY1 | KAT8 regul | Kansl3 | sp\|A2RSY1 | 12 | 12 | 21,9 |
| Q6PAT0 | Q6PAT0 | Probable i | Adat3 | sp\|Q6PAT | 6 | 6 | 28,9 |
| Q3UMU9 | Q3UMU9 | Hepatoma | Hdgfrp2 | sp\|Q3UM | 13 | 11 | 21,8 |
| P97825 | P97825 | Hematolog | Hn1 | sp\|P97825 | 4 | 4 | 37,7 |
| Q8C7E7 | Q8C7E7 | Starch-bin | Stbd1 | sp\|Q8C7E | 7 | 7 | 30,8 |
| Q9JJ66 | Q9JJ66 | Cell divisio | Cdc20 | sp\|Q9JJ66 | 10 | 10 | 38,5 |
| P32067 | P32067 | Lupus La p | Ssb | sp\|P32067 | 30 | 30 | 53 |
| Q9CR88 | Q9CR88 | 28S riboso | Mrps14 | sp\|Q9CR8 | 3 | 3 | 28,9 |
| P97819 | P97819 | 85/88 kDa | Pla2g6 | sp\|P97819 | 4 | 4 | 9,7 |
| Q9CQV6 | Q9CQV6 | Microtubu | Map1lc3b | sp\|Q9CQV | 3 | 1 | 22,4 |
| Q3UHK8 | Q3UHK8 | Trinucleoti | Tnrc6a | sp\|Q3UHK | 12 | 11 | 10,8 |
| Q8C3Q9 | Q8C3Q9 | Caspase-9; | Casp9 | sp\|Q8C3Q | 5 | 5 | 14,3 |
| Q8JZY2 | Q8JZY2 | COMM do | Commd10 | sp\|Q8JZY2 | 7 | 7 | 50,5 |
| Q9R0P4 | Q9R0P4 | Small acidi | Smap | sp\|Q9R0P | 7 | 7 | 30,4 |
| Q9Z223 | Q9Z223 | Molybdopt | Mocs2 | sp\|Q9Z223 | 2 | 2 | 16,4 |
| Q9D1L9 | Q9D1L9 | Ragulator | Lamtor5 | sp\|Q9D1L | 3 | 3 | 59,3 |
| Q9CR41 | Q9CR41 | Huntingtin | Hypk | sp\|Q9CR4 | 7 | 7 | 51,9 |
| P62325 | P62325 | Protein BT | Btg1 | sp\|P62325 | 2 | 2 | 16,4 |
| Q8BL97 | Q8BL97 | Serine/arg | Srsf7 | sp\|Q8BL97 | 9 | 8 | 27,7 |
| Q70FJ1 | Q70FJ1 | A-kinase a | Akap9 | sp\|Q70FJ1 | 59 | 59 | 21 |
| Q923W1 | Q923W1 | Trimethylg | Tgs1 | sp\|Q923W | 10 | 10 | 17,1 |
| G5E8B9 | G5E8B9 | | Zbtb11 | sp\|G5E8B | 4 | 4 | 5 |
| Q78IS1 | Q78IS1 | Transmem | Tmed3 | sp\|Q78IS1 | 4 | 4 | 29,4 |
| Q8C9S8 | Q8C9S8 | Cysteine p | Atg4a | sp\|Q8C9S | 4 | 4 | 12,1 |
| Q91V12 | Q91V12 | Cytosolic a | Acot7 | sp\|Q91V1 | 14 | 14 | 37,5 |
| O89090 | O89090 | Transcripti | Sp1 | sp\|O8909 | 5 | 5 | 13,6 |
| B2RY56 | B2RY56 | RNA-bindi | Rbm25 | sp\|B2RY56 | 23 | 23 | 29,1 |
| Q9WVJ2 | Q9WVJ2 | 26S protea | Psmd13 | sp\|Q9WVJ | 25 | 25 | 69,9 |
| Q8BXL7 | Q8BXL7 | ADP-ribosy | Arfrp1 | sp\|Q8BXL | 7 | 7 | 49,3 |
| P56671 | P56671 | Myc-assoc | Maz | sp\|P56671 | 2 | 2 | 6,1 |
| Q9QWF0 | Q9QWF0 | Chromatin | Chaf1a | sp\|Q9QW | 21 | 21 | 39,5 |
| Q8BGA5 | Q8BGA5 | KRR1 smal | Krr1 | sp\|Q8BGA | 15 | 15 | 36,3 |
| P27808 | P27808 | Alpha-1,3- | Mgat1 | sp\|P27808 | 13 | 13 | 36,7 |
| O35954 | O35954 | Membrane | Pitpnm1 | sp\|O35954 | 15 | 15 | 16,6 |
| Q9R0D8 | Q9R0D8 | WD repeat | Wdr54 | sp\|Q9R0D | 7 | 7 | 38,6 |
| Q922B6 | Q922B6 | E3 ubiquiti | Traf7 | sp\|Q922B | 5 | 5 | 13,1 |
| O35166 | O35166 | Golgi SNAF | Gosr2 | sp\|O3516 | 3 | 3 | 15,6 |
| Q9WUV0 | Q9WUV0 | Origin reco | Orc5 | sp\|Q9WU | 13 | 13 | 36,8 |
| Q9QYH6 | Q9QYH6 | Melanoma | Maged1 | sp\|Q9QYH | 8 | 8 | 16,9 |
| Q9D819 | Q9D819 | Inorganic p | Ppa1 | sp\|Q9D81 | 23 | 23 | 81 |

| | | | | | | | |
|---|---|---|---|---|---|---|---|
| Q9DBN5 | Q9DBN5 | Lon protea | Lonp2 | sp\|Q9DBN | 19 | 18 | 32,3 |
| Q69ZX6 | Q69ZX6 | MORC fam | Morc2a | sp\|Q69ZX6 | 30 | 26 | 36 |
| Q8BTE5 | Q8BTE5 | Protein CE | Cebpzos | sp\|Q8BTE5 | 2 | 2 | 36,2 |
| Q8BWM0 | Q8BWM0 | Prostaglan | Ptges2 | sp\|Q8BWN | 10 | 10 | 36,2 |
| Q5SWW4 | Q5SWW4 | Mediator c | Med13 | sp\|Q5SWV | 21 | 21 | 13,7 |
| Q9CZT4 | Q9CZT4 | DNA-direct | Polr3e | sp\|Q9CZT4 | 6 | 6 | 14,6 |
| Q9DCH6 | Q9DCH6 | AN1-type z | Zfand6 | sp\|Q9DCH | 3 | 3 | 22 |
| P19157;P4 | P19157;P4642 | Glutathion | Gstp1;Gstp | sp\|P19157 | 16 | 16 | 66,2 |
| P20357 | P20357 | Microtubu | Map2 | sp\|P20357 | 5 | 5 | 3,8 |
| Q8BGT7 | Q8BGT7 | Survival of | Smndc1 | sp\|Q8BGT | 5 | 5 | 38,2 |
| Q922H4 | Q922H4 | Mannose-1 | Gmppa | sp\|Q922H | 10 | 10 | 30,2 |
| P19324 | P19324 | Serpin H1 | Serpinh1 | sp\|P19324 | 22 | 22 | 56,1 |
| P97352 | P97352 | Protein S1 | S100a13 | sp\|P97352 | 3 | 3 | 33,7 |
| Q8C080 | Q8C080 | Sorting ne | Snx16 | sp\|Q8C08 | 10 | 10 | 33,7 |
| Q0P678 | Q0P678 | Zinc finger | Zc3h18 | sp\|Q0P678 | 9 | 9 | 14,9 |
| Q6PGH1 | Q6PGH1 | Protein BU | Bud31 | sp\|Q6PGH | 10 | 10 | 54,2 |
| Q920Q6;Q | Q920Q6 | RNA-bindir | Msi2 | sp\|Q920Q | 7 | 7 | 27,7 |
| Q9R0Q6 | Q9R0Q6 | Actin-relat | Arpc1a | sp\|Q9R0Q | 7 | 5 | 25,1 |
| P49615 | P49615 | Cyclin-dep | Cdk5 | sp\|P49615 | 14 | 13 | 52,4 |
| P22777 | P22777 | Plasminog | Serpine1 | sp\|P22777 | 7 | 7 | 19,7 |
| O08601 | O08601 | Microsoma | Mttp | sp\|O08601 | 17 | 17 | 23,4 |
| Q62011 | Q62011 | Podoplanir | Pdpn | sp\|Q62011 | 3 | 3 | 25,6 |
| A3KGB4 | A3KGB4 | TBC1 doma | Tbc1d8b | sp\|A3KGB | 11 | 11 | 13,8 |
| Q60876 | Q60876 | Eukaryotic | Eif4ebp1 | sp\|Q60876 | 3 | 3 | 37,6 |
| A2ADA5 | A2ADA5 | tRNA pseu | Pusl1 | sp\|A2ADA | 3 | 3 | 15,5 |
| Q9CR56 | Q9CR56 | NF-kappa-I | Nkiras2 | sp\|Q9CR56 | 3 | 3 | 17,8 |
| Q3TJM4 | Q3TJM4 | Centromer | Cenpt | sp\|Q3TJM | 7 | 7 | 22,7 |
| P10493 | P10493 | Nidogen-1 | Nid1 | sp\|P10493 | 2 | 2 | 2,3 |
| Q60596 | Q60596 | DNA repair | Xrcc1 | sp\|Q60596 | 9 | 9 | 24,4 |
| Q6NZJ6 | Q6NZJ6 | Eukaryotic | Eif4g1 | sp\|Q6NZJ6 | 60 | 56 | 42,6 |
| Q62191 | Q62191 | E3 ubiquiti | Trim21 | sp\|Q62191 | 3 | 3 | 10,4 |
| Q9ESU6 | Q9ESU6 | Bromodom | Brd4 | sp\|Q9ESU | 23 | 17 | 22,5 |
| P35276;Q9 | P35276 | Ras-relate | Rab3d | sp\|P35276 | 6 | 3 | 32,9 |
| Q8CI95 | Q8CI95 | Oxysterol-l | Osbpl11 | sp\|Q8CI95 | 11 | 9 | 26,5 |
| Q9D8B4 | Q9D8B4 | NADH deh | Ndufa11 | sp\|Q9D8B | 3 | 3 | 35,5 |
| Q68FD7 | Q68FD7 | Folliculin-i | Fnip1 | sp\|Q68FD | 5 | 5 | 5,7 |
| O08811 | O08811 | TFIIH basal | Ercc2 | sp\|O08811 | 11 | 11 | 23,8 |
| Q80Y14 | Q80Y14 | Glutaredox | Glrx5 | sp\|Q80Y14 | 2 | 2 | 21,7 |
| Q9QY76 | Q9QY76 | Vesicle-ass | Vapb | sp\|Q9QY7 | 11 | 10 | 50,2 |
| Q8K370 | Q8K370 | Acyl-CoA d | Acad10 | sp\|Q8K37 | 13 | 13 | 16 |
| Q8R086 | Q8R086 | Sulfite oxid | Suox | sp\|Q8R08 | 6 | 6 | 14,7 |
| Q91VM5 | Q91VM5 | RNA bindir | Rbmxl1 | sp\|Q91VM | 19 | 4 | 46,9 |
| Q99LI8 | Q99LI8 | Hepatocyt | Hgs | sp\|Q99LI8 | 17 | 17 | 26,3 |
| Q8VE97 | Q8VE97 | Serine/arg | Srsf4 | sp\|Q8VE9 | 10 | 7 | 19 |
| Q14AX6 | Q14AX6 | Cyclin-dep | Cdk12 | sp\|Q14AX | 18 | 14 | 17,9 |
| Q9JI19 | Q9JI19 | Acidic fibr | Fibp | sp\|Q9JI19 | 9 | 9 | 33,1 |
| Q8VDH1 | Q8VDH1 | F-box only | Fbxo21 | sp\|Q8VDH | 2 | 2 | 6,5 |
| Q6IRU5 | Q6IRU5 | Clathrin lig | Cltb | sp\|Q6IRU5 | 6 | 6 | 20,5 |
| P61327;Q9 | P61327;Q9CQ | Protein ma | Magoh;Ma | sp\|P61327 | 9 | 9 | 52,7 |
| Q9Z2H5 | Q9Z2H5 | Band 4.1-li | Epb41l1 | sp\|Q9Z2H | 16 | 14 | 26,3 |

| | | | | | | | |
|---|---|---|---|---|---|---|---|
| Q8R574 | Q8R574 | Phosphoril | Prpsap2 | sp\|Q8R574 | 16 | 14 | 55,3 |
| Q9Z2V5 | Q9Z2V5 | Histone de | Hdac6 | sp\|Q9Z2V5 | 8 | 8 | 9,4 |
| P57759 | P57759 | Endoplasm | Erp29 | sp\|P57759 | 12 | 12 | 48,9 |
| Q8BGE6 | Q8BGE6 | Cysteine p | Atg4b | sp\|Q8BGE | 11 | 11 | 39,7 |
| Q8BYI6 | Q8BYI6 | Lysophosp | Lpcat2 | sp\|Q8BYI6 | 9 | 9 | 20,6 |
| Q9JIY0 | Q9JIY0 | Pleckstrin | Plekho1 | sp\|Q9JIY0 | 4 | 4 | 16,2 |
| P70424;Q6 | P70424 | Receptor t | Erbb2 | sp\|P70424 | 11 | 11 | 11,9 |
| Q61990 | Q61990 | Poly(rC)-bi | Pcbp2 | sp\|Q61990 | 20 | 13 | 65,2 |
| O35343 | O35343 | Importin s | Kpna4 | sp\|O35343 | 14 | 9 | 40,7 |
| O35459 | O35459 | Delta(3,5)- | Ech1 | sp\|O35459 | 12 | 12 | 44,6 |
| Q8BN82 | Q8BN82 | Sialin | Slc17a5 | sp\|Q8BN8 | 1 | 1 | 4,2 |
| Q8BL74 | Q8BL74 | General tr | Gtf3c2 | sp\|Q8BL74 | 15 | 15 | 27,3 |
| P17742 | P17742 | Peptidyl-p | Ppia | sp\|P17742 | 13 | 13 | 71,3 |
| Q8R242 | Q8R242 | Di-N-acety | Ctbs | sp\|Q8R242 | 6 | 6 | 22,4 |
| P61924 | P61924 | Coatomer | Copz1 | sp\|P61924 | 6 | 6 | 41,8 |
| O35130 | O35130 | Ribosomal | Emg1 | sp\|O35130 | 16 | 16 | 74,6 |
| Q3TLP5 | Q3TLP5 | Enoyl-CoA | Echdc2 | sp\|Q3TLP5 | 6 | 6 | 28,4 |
| Q7TS72 | Q7TS72 | Inositol-tri | Itpkc | sp\|Q7TS72 | 3 | 3 | 7,7 |
| Q6P6J4 | Q6P6J4 | Alpha-keto | Alkbh2 | sp\|Q6P6J4 | 2 | 2 | 14,6 |
| Q8CBG9 | Q8CBG9 | E3 ubiquiti | Rnf170 | sp\|Q8CBG | 3 | 3 | 13,3 |
| Q9CPW4 | Q9CPW4 | Actin-relat | Arpc5 | sp\|Q9CPW | 9 | 9 | 66,9 |
| Q8C525 | Q8C525 | Protein MB | Mb21d2 | sp\|Q8C525 | 5 | 5 | 16,4 |
| Q80YQ8 | Q80YQ8 | Protein RM | Rmnd5a | sp\|Q80YQ | 5 | 5 | 15,3 |
| Q6PEV3 | Q6PEV3 | WAS/WAS | Wipf2 | sp\|Q6PEV3 | 7 | 7 | 21,1 |
| Q99JT9 | Q99JT9 | 1,2-dihydr | Adi1 | sp\|Q99JT9 | 9 | 9 | 59,2 |
| Q9CQQ7 | Q9CQQ7 | ATP syntha | Atp5f1 | sp\|Q9CQQ | 16 | 16 | 39,8 |
| P33611 | P33611 | DNA polym | Pola2 | sp\|P33611 | 17 | 17 | 41,5 |
| P62342 | P62342 | Selenoprot | Selt | sp\|P62342 | 4 | 4 | 28,7 |
| Q9CXV1 | Q9CXV1 | Succinate | Sdhd | sp\|Q9CXV1 | 2 | 2 | 10,7 |
| Q91W53 | Q91W53 | Golgin sub | Golga7 | sp\|Q91W5 | 4 | 4 | 32,1 |
| Q8BH83 | Q8BH83 | Ankyrin re | Ankrd9 | sp\|Q8BH8 | 4 | 4 | 18,1 |
| D3Z4I3;Q6 | D3Z4I3;Q6217 | RNA-bindi | Rbm24;Rb | sp\|D3Z4I3 | 1 | 1 | 6,8 |
| Q8VID5 | Q8VID5 | ATP-depen | Recql5 | sp\|Q8VID5 | 3 | 3 | 3,8 |
| Q8K248 | Q8K248 | 4-hydroxy | Hpdl | sp\|Q8K248 | 2 | 2 | 5,9 |
| Q9WTR2 | Q9WTR2 | Mitogen-a | Map3k6 | sp\|Q9WTR | 4 | 4 | 4 |
| Q9R0U0 | Q9R0U0 | Serine/arg | Srsf10 | sp\|Q9R0U | 9 | 9 | 26,7 |
| Q9D394 | Q9D394 | Protein RU | Rufy3 | sp\|Q9D394 | 6 | 5 | 14,9 |
| Q9CZ57 | Q9CZ57 | 5-methylcy | Nsun4 | sp\|Q9CZ57 | 9 | 9 | 28,3 |
| Q80YR4 | Q80YR4 | Zinc finger | Znf598 | sp\|Q80YR4 | 17 | 17 | 27,9 |
| Q9WUN2 | Q9WUN2 | Serine/thre | Tbk1 | sp\|Q9WUN | 16 | 16 | 29,9 |
| Q8VIJ8 | Q8VIJ8 | Nitrogen p | Nprl3 | sp\|Q8VIJ8 | 6 | 6 | 15,3 |
| Q8R3L0 | Q8R3L0 | Membrane | Mmgt2 | sp\|Q8R3L0 | 2 | 2 | 26 |
| O09130 | O09130 | NFATC2-in | Nfatc2ip | sp\|O09130 | 5 | 5 | 22,1 |
| Q8K2Q0 | Q8K2Q0 | COMM dor | Commd9 | sp\|Q8K2Q | 7 | 7 | 50,5 |
| Q8BZ47 | Q8BZ47 | Zinc finger | Znf609 | sp\|Q8BZ47 | 15 | 15 | 16,8 |
| Q8BGS3 | Q8BGS3 | Zinc finger | Zkscan1 | sp\|Q8BGS3 | 3 | 3 | 9,3 |
| O35286 | O35286 | Pre-mRNA- | Dhx15 | sp\|O35286 | 37 | 36 | 50,2 |
| P46414 | P46414 | Cyclin-dep | Cdkn1b | sp\|P46414 | 5 | 5 | 35 |
| Q9D600 | Q9D600 | DNA replic | Gins2 | sp\|Q9D600 | 5 | 5 | 37,3 |
| Q91V83 | Q91V83 | TELO2-inte | Tti1 | sp\|Q91V83 | 4 | 4 | 5,1 |

| | | | | | | | |
|---|---|---|---|---|---|---|---|
| Q68EF0 | Q68EF0 | Rab-3A-int | Rab3ip | sp\|Q68EF0 | 6 | 6 | 21 |
| Q9EQU5 | Q9EQU5 | Protein SE | Set | sp\|Q9EQU | 13 | 13 | 44,6 |
| Q3UQN2 | Q3UQN2 | F-BAR dom | Fcho2 | sp\|Q3UQN | 19 | 19 | 38,2 |
| Q6PD03 | Q6PD03 | Serine/thre | Ppp2r5a | sp\|Q6PD0 | 14 | 12 | 32,7 |
| P26516 | P26516 | 26S protea | Psmd7 | sp\|P26516 | 12 | 12 | 46,7 |
| Q9CZ44 | Q9CZ44 | NSFL1 cofa | Nsfl1c | sp\|Q9CZ44 | 18 | 18 | 56,2 |
| Q9EPJ9 | Q9EPJ9 | ADP-ribosy | Arfgap1 | sp\|Q9EPJ9 | 12 | 12 | 43,5 |
| P97789 | P97789 | 5-3 exoribo | Xrn1 | sp\|P97789 | 29 | 29 | 22,6 |
| Q8BJL0 | Q8BJL0 | SWI/SNF-re | Smarcal1 | sp\|Q8BJL0 | 17 | 17 | 27,4 |
| Q80UP5;Q | Q80UP5 | Ankyrin re | Ankrd13a | sp\|Q80UP | 4 | 4 | 10,5 |
| Q76MZ3 | Q76MZ3 | Serine/thre | Ppp2r1a | sp\|Q76MZ | 33 | 24 | 64 |
| Q8BR07 | Q8BR07 | Protein bic | Bicd1 | sp\|Q8BR0 | 4 | 1 | 4,8 |
| Q9CQN7 | Q9CQN7 | 39S riboso | Mrpl41 | sp\|Q9CQN | 6 | 6 | 50,4 |
| Q9CYN2 | Q9CYN2 | Signal pept | Spcs2 | sp\|Q9CYN | 8 | 8 | 42 |
| P31230 | P31230 | Aminoacyl | Aimp1 | sp\|P31230 | 19 | 19 | 72,3 |
| Q9D3D0 | Q9D3D0 | Alpha-toco | Ttpal | sp\|Q9D3D | 5 | 5 | 17,5 |
| Q9D880 | Q9D880 | Mitochond | Timm50 | sp\|Q9D88 | 12 | 12 | 43,9 |
| Q3TYS2 | Q3TYS2 | Uncharacterized prote | | sp\|Q3TYS2 | 5 | 5 | 29,9 |
| O35188 | O35188 | Fractalkine | Cx3cl1 | sp\|O35188 | 2 | 2 | 6,6 |
| Q6RUT7 | Q6RUT7 | Protein CC | Ccsmst1 | sp\|Q6RUT | 1 | 1 | 18,4 |
| Q80V62 | Q80V62 | Fanconi an | Fancd2 | sp\|Q80V62 | 11 | 11 | 10,6 |
| Q91VW9 | Q91VW9 | Zinc finger | Zkscan3 | sp\|Q91VW | 6 | 6 | 19,2 |
| P62141 | P62141 | Serine/thre | Ppp1cb | sp\|P62141 | 20 | 7 | 70,6 |
| Q8VIJ6 | Q8VIJ6 | Splicing fac | Sfpq | sp\|Q8VIJ6 | 33 | 32 | 50,1 |
| CON__ENS | CON__ENSEM | Fibulin-1 | Fbln1 | ;sp\|Q0887 | 2 | 2 | 2,7 |
| Q9EP72 | Q9EP72 | ER membra | Emc7 | sp\|Q9EP72 | 6 | 6 | 39,8 |
| Q8BGR9 | Q8BGR9 | Ubiquitin-l | Ublcp1 | sp\|Q8BGR | 12 | 12 | 38,7 |
| P49442 | P49442 | Inositol po | Inpp1 | sp\|P49442 | 5 | 5 | 16,4 |
| Q78YY6 | Q78YY6 | DnaJ homo | Dnajc15 | sp\|Q78YY6 | 2 | 2 | 16,8 |
| Q9D7X3 | Q9D7X3 | Dual specif | Dusp3 | sp\|Q9D7X | 4 | 4 | 33,5 |
| Q3UQ28 | Q3UQ28 | Peroxidasi | Pxdn | sp\|Q3UQ2 | 32 | 32 | 32 |
| P14733 | P14733 | Lamin-B1 | Lmnb1 | sp\|P14733 | 39 | 37 | 62,6 |
| Q9R1P3 | Q9R1P3 | Proteasom | Psmb2 | sp\|Q9R1P3 | 9 | 9 | 49,3 |
| Q8BK63 | Q8BK63 | Casein kina | Csnk1a1 | sp\|Q8BK63 | 11 | 11 | 38,6 |
| Q8QZR8 | Q8QZR8 | Cyclin-rela | Fam58b | sp\|Q8QZR | 3 | 3 | 18,8 |
| Q8BVQ5 | Q8BVQ5 | Protein ph | Ppme1 | sp\|Q8BVQ | 14 | 14 | 43,8 |
| Q8R0F6 | Q8R0F6 | Integrin-lin | Ilkap | sp\|Q8R0F6 | 9 | 9 | 42,3 |
| Q02780 | Q02780 | Nuclear fac | Nfia | sp\|Q02780 | 10 | 7 | 27,6 |
| O89017 | O89017 | Legumain | Lgmn | sp\|O89017 | 8 | 8 | 30,1 |
| Q9Z1T1 | Q9Z1T1 | AP-3 comp | Ap3b1 | sp\|Q9Z1T1 | 31 | 26 | 34,6 |
| Q9D0B5 | Q9D0B5 | Thiosulfate | Tstd3 | sp\|Q9D0B | 5 | 5 | 35,7 |
| Q6Y7W8 | Q6Y7W8 | PERQ amir | Gigyf2 | sp\|Q6Y7W | 26 | 26 | 30 |
| P47955 | P47955 | 60S acidic | Rplp1 | sp\|P47955 | 5 | 5 | 94,7 |
| Q9QZN1 | Q9QZN1 | F-box/LRR- | Fbxl17 | sp\|Q9QZN | 5 | 5 | 10,8 |
| Q6P5D3 | Q6P5D3 | Putative A | Dhx57 | sp\|Q6P5D | 15 | 13 | 14,3 |
| Q3TC93 | Q3TC93 | HCLS1-bind | Hs1bp3 | sp\|Q3TC93 | 14 | 14 | 46,6 |
| Q6ZWY8;C | Q6ZWY8 | Thymosin | Tmsb10 | sp\|Q6ZWY | 5 | 4 | 63,6 |
| Q9Z321 | Q9Z321 | DNA topoi | Top3b | sp\|Q9Z321 | 23 | 22 | 39,9 |
| Q9QX60 | Q9QX60 | Deoxyguar | Dguok | sp\|Q9QX6 | 3 | 3 | 15,9 |
| Q9ER88 | Q9ER88 | 28S riboso | Dap3 | sp\|Q9ER88 | 13 | 13 | 41,4 |

| | | | | | | | |
|---|---|---|---|---|---|---|---|
| P15864;Q( | P15864 | Histone H1 | Hist1h1c | sp\|P15864 | 15 | 6 | 46,2 |
| P28033 | P28033 | CCAAT/enh | Cebpb | sp\|P28033 | 4 | 4 | 18,6 |
| Q9CQF9 | Q9CQF9 | Prenylcyste | Pcyox1 | sp\|Q9CQF9 | 17 | 17 | 55 |
| Q8VCF1 | Q8VCF1 | Soluble cal | Cant1 | sp\|Q8VCF1 | 10 | 10 | 33 |
| Q9D5K4 | Q9D5K4 | S100P-binc | S100pbp | sp\|Q9D5K4 | 4 | 4 | 14,6 |
| Q91VN4 | Q91VN4 | MICOS con | Chchd6 | sp\|Q91VN | 6 | 6 | 25,6 |
| B2RS91 | B2RS91 | RNA polym | Rrn3 | sp\|B2RS91 | 6 | 6 | 9,5 |
| P20934 | P20934 | Protein EV | Evi2a | sp\|P20934 | 1 | 1 | 8,5 |
| Q9JKK1 | Q9JKK1 | Syntaxin-6 | Stx6 | sp\|Q9JKK1 | 9 | 9 | 45,1 |
| Q8BMS4 | Q8BMS4 | Ubiquinon | Coq3 | sp\|Q8BMS | 3 | 3 | 11,9 |
| Q8BS95 | Q8BS95 | Golgi pH re | Gpr89a | sp\|Q8BS95 | 3 | 3 | 7,9 |
| Q8C5Q4 | Q8C5Q4 | G-rich sequ | Grsf1 | sp\|Q8C5Q | 15 | 15 | 45,1 |
| Q8CG79 | Q8CG79 | Apoptosis- | Tp53bp2 | sp\|Q8CG7 | 14 | 13 | 21,4 |
| A2AIL4 | A2AIL4 | NADH deh | Ndufaf6 | sp\|A2AIL4 | 3 | 3 | 13,2 |
| P70371 | P70371 | Telomeric | Terf1 | sp\|P70371 | 6 | 6 | 18,1 |
| Q9JJY4 | Q9JJY4 | Probable A | Ddx20 | sp\|Q9JJY4 | 25 | 25 | 41 |
| Q61550;A2 | Q61550 | Double-str | Rad21 | sp\|Q61550 | 24 | 24 | 48,5 |
| Q8VDQ1 | Q8VDQ1 | Prostaglan | Ptgr2 | sp\|Q8VDQ | 10 | 10 | 38,7 |
| Q8K2F0 | Q8K2F0 | Bromodom | Brd3 | sp\|Q8K2F0 | 16 | 12 | 28,1 |
| P21107 | P21107 | Tropomyo | Tpm3 | sp\|P21107 | 25 | 12 | 55,4 |
| O35344 | O35344 | Importin su | Kpna3 | sp\|O35344 | 16 | 11 | 43,4 |
| Q8BHS8 | Q8BHS8 | Syntabulin | Sybu | sp\|Q8BHS8 | 1 | 1 | 2,6 |
| Q8R1J9;P0 | Q8R1J9 | Torsin-2A | Tor2a | sp\|Q8R1J9 | 5 | 5 | 22,1 |
| Q02614 | Q02614 | SAP30-binc | Sap30bp | sp\|Q02614 | 8 | 8 | 33,4 |
| P10923 | P10923 | Osteopont | Spp1 | sp\|P10923 | 5 | 5 | 28,2 |
| B1AVH7 | B1AVH7 | TBC1 doma | Tbc1d2 | sp\|B1AVH7 | 4 | 4 | 5,4 |
| Q9R1Q6 | Q9R1Q6 | Transmem | Tmem176b | sp\|Q9R1Q | 3 | 3 | 10,3 |
| Q91YI4 | Q91YI4 | Beta-arres | Arrb2 | sp\|Q91YI4 | 10 | 10 | 27,3 |
| Q6NZQ6 | Q6NZQ6 | Zinc finger | Znf740 | sp\|Q6NZQ | 3 | 3 | 14,4 |
| Q9CQW1 | Q9CQW1 | Synaptobre | Ykt6 | sp\|Q9CQW | 10 | 10 | 62,6 |
| Q924T3 | Q924T3 | DNA repair | Xrcc4 | sp\|Q924T3 | 8 | 8 | 28,5 |
| Q8R366 | Q8R366 | Immunogl | Igsf8 | sp\|Q8R366 | 7 | 7 | 17 |
| Q62018 | Q62018 | RNA polym | Ctr9 | sp\|Q62018 | 20 | 20 | 22,1 |
| Q8R0K4 | Q8R0K4 | Coiled-coil | Ccdc137 | sp\|Q8R0K4 | 11 | 11 | 37,6 |
| Q922G0 | Q922G0 | Solute carr | Slc25a36 | sp\|Q922G0 | 3 | 2 | 12,2 |
| Q6ZWU9 | Q6ZWU9 | 40S riboso | Rps27 | sp\|Q6ZWU | 6 | 3 | 41,7 |
| Q9CS42 | Q9CS42 | Ribose-pho | Prps2 | sp\|Q9CS42 | 13 | 7 | 49,1 |
| Q9DAP7 | Q9DAP7 | Histone ch | Asf1b | sp\|Q9DAP | 5 | 5 | 48 |
| Q8VD00 | Q8VD00 | Transmem | Tmem97 | sp\|Q8VD0 | 2 | 2 | 8,5 |
| Q9D8T2 | Q9D8T2 | Gasdermin | Gsdmdc1 | sp\|Q9D8T2 | 11 | 11 | 28,3 |
| Q91W34 | Q91W34 | RUS1 family protein C | | sp\|Q91W3 | 3 | 3 | 8,4 |
| Q8CDJ8 | Q8CDJ8 | Stonin-1 | Ston1 | sp\|Q8CDJ8 | 14 | 14 | 24,1 |
| P13439 | P13439 | Uridine 5-r | Umps | sp\|P13439 | 26 | 26 | 76,1 |
| Q9JK48 | Q9JK48 | Endophilin | Sh3glb1 | sp\|Q9JK48 | 12 | 11 | 38,6 |
| Q8BUH7 | Q8BUH7 | Ring finger | Rnf26 | sp\|Q8BUH | 3 | 3 | 6,8 |
| Q64008 | Q64008 | Ras-relate | Rab34 | sp\|Q64008 | 10 | 10 | 46,3 |
| Q91XD6 | Q91XD6 | Vacuolar p | Vps36 | sp\|Q91XD | 18 | 18 | 52,6 |
| P00158 | P00158 | Cytochrom | Mt-Cyb | sp\|P00158 | 2 | 2 | 5 |
| Q8BX07 | Q8BX07 | Carboxy-te | Ctdsp2 | sp\|Q8BX07 | 2 | 2 | 7,4 |
| Q8VE47 | Q8VE47 | Ubiquitin-l | Uba5 | sp\|Q8VE47 | 9 | 9 | 31,3 |

| | | | | | | | |
|---|---|---|---|---|---|---|---|
| Q8CHU3 | Q8CHU3 | Epsin-2 | Epn2 | sp\|Q8CHU | 18 | 17 | 45,4 |
| Q8C460 | Q8C460 | ERI1 exorib | Eri3 | sp\|Q8C46( | 5 | 5 | 23,1 |
| Q9D554 | Q9D554 | Splicing fac | Sf3a3 | sp\|Q9D554 | 22 | 22 | 49,5 |
| Q3UV16 | Q3UV16 | Inositol 1,4 | Itpripl2 | sp\|Q3UV1 | 7 | 7 | 18,5 |
| A3KFM7 | A3KFM7 | Chromodo | Chd6 | sp\|A3KFM | 14 | 9 | 7 |
| P12367 | P12367 | cAMP-dep | Prkar2a | sp\|P12367 | 10 | 8 | 32,7 |
| Q6P4T2 | Q6P4T2 | U5 small n | Snrnp200 | sp\|Q6P4T2 | 110 | 110 | 58 |
| Q8BGZ4 | Q8BGZ4 | Cell divisio | Cdc23 | sp\|Q8BGZ4 | 20 | 20 | 45,1 |
| P57016 | P57016 | Ladinin-1 | Lad1 | sp\|P57016 | 21 | 21 | 44,7 |
| Q3UU96 | Q3UU96 | Serine/thre | Cdc42bpa | sp\|Q3UU9 | 26 | 22 | 20,2 |
| P70255 | P70255 | Nuclear fac | Nfic | sp\|P70255 | 10 | 8 | 25,1 |
| Q99LG0 | Q99LG0 | Ubiquitin c | Usp16 | sp\|Q99LG( | 12 | 12 | 24,6 |
| Q9ERG2 | Q9ERG2 | Striatin-3 | Strn3 | sp\|Q9ERG2 | 17 | 15 | 26,5 |
| Q9CYU6 | Q9CYU6 | Diphthine | Dph7 | sp\|Q9CYU( | 5 | 5 | 14,3 |
| Q02819 | Q02819 | Nucleobind | Nucb1 | sp\|Q02819 | 27 | 27 | 59,9 |
| Q8BZA9 | Q8BZA9 | Fructose-2 | Tigar | sp\|Q8BZA9 | 2 | 2 | 14,1 |
| Q8R0S1 | Q8R0S1 | Cyclic AMP | Atf7 | sp\|Q8R0S1 | 5 | 5 | 17,4 |
| Q6P9R2 | Q6P9R2 | Serine/thre | Oxsr1 | sp\|Q6P9R2 | 15 | 13 | 39,5 |
| Q8BPS4 | Q8BPS4 | Integral me | Gpr180 | sp\|Q8BPS4 | 2 | 2 | 5,4 |
| Q9CZ69 | Q9CZ69 | CKLF-like M | Cmtm6 | sp\|Q9CZ69 | 2 | 2 | 15,8 |
| Q5U4F6 | Q5U4F6 | WD repeat | Wdr34 | sp\|Q5U4F( | 7 | 7 | 22,1 |
| Q9D1P4 | Q9D1P4 | Cysteine a | Chordc1 | sp\|Q9D1P4 | 15 | 15 | 53,5 |
| O88879 | O88879 | Apoptotic | Apaf1 | sp\|O88879 | 18 | 18 | 21,9 |
| Q791T5 | Q791T5 | Mitochond | Mtch1 | sp\|Q791T5 | 9 | 9 | 26,5 |
| Q8CAA7 | Q8CAA7 | Glucose 1,( | Pgm2l1 | sp\|Q8CAA | 4 | 4 | 8,1 |
| O08609 | O08609 | Max-like p | Mlx | sp\|O08609 | 2 | 2 | 10,4 |
| Q9D291 | Q9D291 | Desumoyla | Desi2 | sp\|Q9D291 | 2 | 2 | 17 |
| Q91WD4 | Q91WD4 | UPF0415 protein C7or | | sp\|Q91WD | 6 | 6 | 18,5 |
| Q8C142 | Q8C142 | Low densit | Ldlrap1 | sp\|Q8C142 | 5 | 5 | 20,5 |
| Q8BPE4 | Q8BPE4 | Transmem | Tmem177 | sp\|Q8BPE4 | 3 | 3 | 9,6 |
| Q9JHS3 | Q9JHS3 | Ragulator | Lamtor2 | sp\|Q9JHS3 | 6 | 6 | 60 |
| Q9DCX1 | Q9DCX1 | MAD2L1-b | Mad2l1bp | sp\|Q9DCX | 7 | 7 | 39,5 |
| Q9Z2B5 | Q9Z2B5 | Eukaryotic | Eif2ak3 | sp\|Q9Z2B5 | 4 | 4 | 5 |
| Q9JKP5 | Q9JKP5 | Muscleblin | Mbnl1 | sp\|Q9JKP5 | 8 | 2 | 23,5 |
| Q8CE33 | Q8CE33 | Kelch-like | Klhl11 | sp\|Q8CE33 | 6 | 6 | 12,6 |
| O55029 | O55029 | Coatomer | Copb2 | sp\|O55029 | 38 | 38 | 60,1 |
| O88907 | O88907 | E3 SUMO- | Pias1 | sp\|O88907 | 9 | 7 | 17,7 |
| Q9CYK1 | Q9CYK1 | Tryptophar | Wars2 | sp\|Q9CYK1 | 6 | 6 | 19,4 |
| Q60952 | Q60952 | Centrosom | Cep250 | sp\|Q60952 | 11 | 11 | 6 |
| Q9QZH6 | Q9QZH6 | Evolutiona | Ecsit | sp\|Q9QZH | 11 | 11 | 37,9 |
| Q69ZB8 | Q69ZB8 | Zinc finger | Zcchc2 | sp\|Q69ZB8 | 5 | 5 | 5,7 |
| Q9WVA2;( | Q9WVA2 | Mitochond | Timm8a1 | sp\|Q9WVA | 5 | 5 | 79,4 |
| Q8BUL5 | Q8BUL5 | Kelch-like | Klhl7 | sp\|Q8BUL5 | 5 | 5 | 11,1 |
| O70252 | O70252 | Heme oxyg | Hmox2 | sp\|O70252 | 17 | 17 | 65,1 |
| O35943 | O35943 | Frataxin, m | Fxn | sp\|O35943 | 4 | 4 | 25,1 |
| Q6IE82 | Q6IE82 | Protein Jad | Jade3 | sp\|Q6IE82 | 5 | 5 | 8,3 |
| P51863 | P51863 | V-type pro | Atp6v0d1 | sp\|P51863 | 11 | 11 | 43,3 |
| Q9DCU9 | Q9DCU9 | 4-hydroxy- | Hoga1 | sp\|Q9DCU | 4 | 4 | 13,7 |
| Q8VDB2 | Q8VDB2 | Dol-P-Man | Alg12 | sp\|Q8VDB | 1 | 1 | 3,3 |
| Q8BTZ5 | Q8BTZ5 | Ankyrin re | Ankrd46 | sp\|Q8BTZ5 | 4 | 4 | 31,6 |

| | | | | | | | |
|---|---|---|---|---|---|---|---|
| Q3UFF7 | Q3UFF7 | Lysophosp | Lyplal1 | sp\|Q3UFF7 | 3 | 3 | 18,8 |
| Q8BVF7 | Q8BVF7 | Gamma-se | Aph1a | sp\|Q8BVF7 | 1 | 1 | 4,9 |
| Q571G4 | Q571G4 | Protein lin- | Lin54 | sp\|Q571G4 | 11 | 11 | 22,2 |
| Q99LD4 | Q99LD4 | COP9 signa | Gps1 | sp\|Q99LD4 | 17 | 17 | 39,5 |
| Q8R1F1 | Q8R1F1 | Niban-like | Fam129b | sp\|Q8R1F1 | 26 | 26 | 50,5 |
| Q9JMB0 | Q9JMB0 | G kinase-a | Gkap1 | sp\|Q9JMB | 7 | 7 | 24,3 |
| Q3TC33 | Q3TC33 | Coiled-coil | Ccdc127 | sp\|Q3TC33 | 5 | 5 | 26,5 |
| Q9ERN0 | Q9ERN0 | Secretory c | Scamp2 | sp\|Q9ERN0 | 2 | 2 | 9,7 |
| Q80VC6 | Q80VC6 | tRNA selen | Trnau1ap | sp\|Q80VC6 | 2 | 2 | 13,2 |
| Q9D8N2 | Q9D8N2 | Protein FA | Fam45a | sp\|Q9D8N | 9 | 9 | 31,9 |
| Q00558 | Q00558 | Factor VIII | F8a1 | sp\|Q00558 | 3 | 3 | 13,4 |
| Q7TQK0 | Q7TQK0 | | Ccnt2 | sp\|Q7TQK0 | 3 | 2 | 5,9 |
| Q8CD92 | Q8CD92 | Tetratricop | Ttc27 | sp\|Q8CD92 | 22 | 22 | 32 |
| Q9EQ61 | Q9EQ61 | Pescadillo | Pes1 | sp\|Q9EQ6 | 20 | 20 | 34,2 |
| P59110 | P59110 | Sentrin-spe | Senp1 | sp\|P59110 | 9 | 9 | 19,4 |
| Q60793 | Q60793 | Krueppel-li | Klf4 | sp\|Q60793 | 2 | 2 | 8,3 |
| Q9CXI5 | Q9CXI5 | Mesencepl | Manf | sp\|Q9CXI5 | 10 | 10 | 41,9 |
| Q8JZV7 | Q8JZV7 | Putative N- | Amdhd2 | sp\|Q8JZV7 | 3 | 3 | 11,5 |
| Q60854 | Q60854 | Serpin B6 | Serpinb6 | sp\|Q60854 | 20 | 20 | 64,8 |
| Q3TDX8 | Q3TDX8 | Cytochrom | Cyb5r4 | sp\|Q3TDX8 | 5 | 5 | 16,3 |
| Q9CXU0 | Q9CXU0 | Mediator c | Med10 | sp\|Q9CXU0 | 4 | 4 | 45,9 |
| Q8CDM1 | Q8CDM1 | ATPase far | Atad2 | sp\|Q8CDM | 33 | 33 | 38,7 |
| Q8C0J6 | Q8C0J6 | Ankyrin re | Sowahc | sp\|Q8C0J6 | 2 | 2 | 4,3 |
| Q62136 | Q62136 | Tyrosine-p | Ptpn21 | sp\|Q62136 | 13 | 13 | 13,3 |
| Q9Z120 | Q9Z120 | tRNA (guar | Mettl1 | sp\|Q9Z120 | 8 | 8 | 36,9 |
| Q9CX99 | Q9CX99 | GRB2-relat | Grap | sp\|Q9CX99 | 2 | 2 | 10,1 |
| Q9JIG7 | Q9JIG7 | Coiled-coil | Ccdc22 | sp\|Q9JIG7 | 23 | 23 | 51,5 |
| Q9WTZ2 | Q9WTZ2 | Membrane | Mbtps1 | sp\|Q9WTZ | 4 | 4 | 4,2 |
| Q9R0B7 | Q9R0B7 | Zinc finger | Znf346 | sp\|Q9R0B7 | 4 | 4 | 20,1 |
| Q8JZS6 | Q8JZS6 | NEDD4-bin | N4bp2l2 | sp\|Q8JZS6 | 7 | 7 | 16 |
| Q00420 | Q00420 | GA-binding | Gabpb1 | sp\|Q00420 | 2 | 1 | 7 |
| Q8BVE3 | Q8BVE3 | V-type pro | Atp6v1h | sp\|Q8BVE3 | 11 | 11 | 31,3 |
| Q8R3G1 | Q8R3G1 | Nuclear inh | Ppp1r8 | sp\|Q8R3G | 12 | 12 | 53 |
| Q9JKS5 | Q9JKS5 | Intracellula | Habp4 | sp\|Q9JKS5 | 3 | 3 | 15,3 |
| Q9DB73 | Q9DB73 | NADH-cyto | Cyb5r1 | sp\|Q9DB73 | 12 | 12 | 50,2 |
| Q80Y56 | Q80Y56 | Rabenosyn | Rbsn | sp\|Q80Y56 | 9 | 9 | 15,3 |
| Q6PD21 | Q6PD21 | SH2 domai | Shb | sp\|Q6PD21 | 1 | 1 | 2,8 |
| Q9QUM9 | Q9QUM9 | Proteasom | Psma6 | sp\|Q9QUM | 17 | 17 | 70,7 |
| Q9Z2M6 | Q9Z2M6 | Ubiquitin-l | Ubl3 | sp\|Q9Z2M | 3 | 3 | 33,3 |
| Q8BKE9 | Q8BKE9 | Intraflagell | Ift74 | sp\|Q8BKE9 | 4 | 4 | 8,2 |
| P97314 | P97314 | Cysteine ar | Csrp2 | sp\|P97314 | 6 | 6 | 40,9 |
| Q9CR61 | Q9CR61 | NADH deh | Ndufb7 | sp\|Q9CR61 | 7 | 7 | 58,4 |
| P62259 | P62259 | 14-3-3 pro | Ywhae | sp\|P62259 | 27 | 24 | 77,6 |
| Q3U0V1 | Q3U0V1 | Far upstrea | Khsrp | sp\|Q3U0V | 31 | 28 | 51,3 |
| Q9EQY0 | Q9EQY0 | Serine/thre | Ern1 | sp\|Q9EQY0 | 8 | 8 | 18 |
| Q6NZF1 | Q6NZF1 | Zinc finger | Zc3h11a | sp\|Q6NZF1 | 35 | 35 | 52,5 |
| Q9R0N0 | Q9R0N0 | Galactokin | Galk1 | sp\|Q9R0N0 | 14 | 14 | 32,7 |
| Q6NXN1 | Q6NXN1 | SUZ domai | Szrd1 | sp\|Q6NXN | 4 | 4 | 48 |
| A2AIV2 | A2AIV2 | Protein vir | Kiaa1429 | sp\|A2AIV2 | 33 | 33 | 26,5 |
| Q6P9P6 | Q6P9P6 | Kinesin-like | Kif11 | sp\|Q6P9P6 | 32 | 32 | 38,8 |

| | | | | | | | |
|---|---|---|---|---|---|---|---|
| P35689 | P35689 | DNA repair | Ercc5 | sp\|P35689 | 4 | 4 | 6,1 |
| Q9D7Z3 | Q9D7Z3 | Nucleolar p | Nol7 | sp\|Q9D7Z3 | 10 | 10 | 31,1 |
| P34884 | P34884 | Macrophag | Mif | sp\|P34884 | 7 | 7 | 90,4 |
| Q4PJX1 | Q4PJX1 | Protein od | Odr4 | sp\|Q4PJX1 | 11 | 11 | 28 |
| P62073 | P62073 | Mitochond | Timm10 | sp\|P62073 | 6 | 6 | 32,2 |
| Q99LG4 | Q99LG4 | Tetratricop | Ttc5 | sp\|Q99LG4 | 6 | 6 | 20,2 |
| Q8BUE4 | Q8BUE4 | Apoptosis- | Aifm2 | sp\|Q8BUE4 | 6 | 6 | 21,2 |
| Q3V1L4 | Q3V1L4 | Cytosolic p | Nt5c2 | sp\|Q3V1L4 | 19 | 19 | 46,2 |
| Q63918 | Q63918 | Serum dep | Sdpr | sp\|Q63918 | 13 | 13 | 54,3 |
| P70182;P7 | P70182 | Phosphatid | Pip5k1a | sp\|P70182 | 11 | 9 | 23,8 |
| Q9WUM3 | Q9WUM3 | Coronin-1B | Coro1b | sp\|Q9WUM | 13 | 13 | 30,6 |
| P70444 | P70444 | BH3-intera | Bid | sp\|P70444 | 4 | 4 | 31,3 |
| Q6ZPF4 | Q6ZPF4 | Formin-like | Fmnl3 | sp\|Q6ZPF4 | 21 | 16 | 23,2 |
| O88425 | O88425 | Nucleoside | Nme6 | sp\|O88425 | 3 | 3 | 18 |
| Q9D2P4 | Q9D2P4 | Ubiquitin-r | Urm1 | sp\|Q9D2P4 | 3 | 3 | 36,6 |
| Q3U186 | Q3U186 | Probable a | Rars2 | sp\|Q3U186 | 15 | 15 | 31 |
| Q922R5 | Q922R5 | Serine/thre | Smek2 | sp\|Q922R5 | 24 | 17 | 36,7 |
| Q9D5V5 | Q9D5V5 | Cullin-5 | Cul5 | sp\|Q9D5V | 34 | 34 | 47,9 |
| O35623 | O35623 | BET1 homo | Bet1 | sp\|O35623 | 2 | 2 | 24,6 |
| Q9D0T1 | Q9D0T1 | NHP2-like | Nhp2l1 | sp\|Q9D0T1 | 9 | 9 | 53,1 |
| P62717 | P62717 | 60S riboso | Rpl18a | sp\|P62717 | 8 | 8 | 39,2 |
| Q80XJ2 | Q80XJ2 | E3 ubiquiti | Rnf168 | sp\|Q80XJ2 | 3 | 3 | 5 |
| Q924C1 | Q924C1 | Exportin-5 | Xpo5 | sp\|Q924C1 | 38 | 38 | 40 |
| Q501J2 | Q501J2 | Protein FA | Fam173a | sp\|Q501J2 | 7 | 7 | 45,4 |
| Q3UFB2 | Q3UFB2 | Box C/D sn | Znhit6 | sp\|Q3UFB2 | 7 | 7 | 23,9 |
| Q80TN7 | Q80TN7 | Neuron na | Nav3 | sp\|Q80TN7 | 7 | 6 | 4,2 |
| Q91Z67;Q9 | Q91Z67 | SLIT-ROBO | Srgap2 | sp\|Q91Z67 | 28 | 28 | 36,5 |
| P30999 | P30999 | Catenin de | Ctnnd1 | sp\|P30999 | 37 | 37 | 54,5 |
| Q9DCV4 | Q9DCV4 | Regulator | Rmdn1 | sp\|Q9DCV4 | 7 | 7 | 31,1 |
| Q99KL7 | Q99KL7 | Ras-related | Rab28 | sp\|Q99KL7 | 6 | 6 | 32,6 |
| Q8VDI7 | Q8VDI7 | Ubiquitin-a | Ubac1 | sp\|Q8VDI7 | 2 | 2 | 5,9 |
| E9Q4F7 | E9Q4F7 | | Ankrd11 | sp\|E9Q4F7 | 8 | 8 | 4,5 |
| Q921J4 | Q921J4 | Ubiquitin-c | Ube2s | sp\|Q921J4 | 11 | 11 | 67,3 |
| P83870 | P83870 | PHD finger | Phf5a | sp\|P83870 | 7 | 7 | 61,8 |
| O88746 | O88746 | Target of M | Tom1 | sp\|O88746 | 10 | 10 | 33,3 |
| Q9ESN9 | Q9ESN9 | C-Jun-amin | Mapk8ip3 | sp\|Q9ESN9 | 6 | 3 | 5,8 |
| Q8C156 | Q8C156 | Condensin | Ncaph | sp\|Q8C156 | 22 | 22 | 43,9 |
| Q8BJ34 | Q8BJ34 | Meiosis arr | Marf1 | sp\|Q8BJ34 | 11 | 11 | 7,9 |
| Q99MR1 | Q99MR1 | PERQ amin | Gigyf1 | sp\|Q99MR | 7 | 7 | 8,6 |
| P14685 | P14685 | 26S protea | Psmd3 | sp\|P14685 | 33 | 33 | 64 |
| Q9ESP1 | Q9ESP1 | Stromal ce | Sdf2l1 | sp\|Q9ESP1 | 5 | 5 | 35,7 |
| Q8CJ27 | Q8CJ27 | Abnormal s | Aspm | sp\|Q8CJ27 | 15 | 15 | 6,5 |
| O88393 | O88393 | Transformi | Tgfbr3 | sp\|O88393 | 3 | 3 | 4,2 |
| Q60967;O8 | Q60967 | Bifunctiona | Papss1 | sp\|Q60967 | 23 | 23 | 49,5 |
| Q8BL95 | Q8BL95 | UPF0769 protein C21o | | sp\|Q8BL95 | 2 | 2 | 16,6 |
| Q3THG9 | Q3THG9 | Alanyl-tRN | Aarsd1 | sp\|Q3THG | 16 | 16 | 58,3 |
| Q8VCB2 | Q8VCB2 | Mediator o | Med25 | sp\|Q8VCB2 | 3 | 3 | 7,2 |
| Q99J25 | Q99J25 | rRNA meth | Mrm1 | sp\|Q99J25 | 2 | 2 | 10,6 |
| Q62241 | Q62241 | U1 small n | Snrpc | sp\|Q62241 | 4 | 4 | 30,2 |
| Q9Z2Y3 | Q9Z2Y3 | Homer pro | Homer1 | sp\|Q9Z2Y3 | 4 | 4 | 13,1 |

| | | | | | | | |
|---|---|---|---|---|---|---|---|
| Q9QXN3 | Q9QXN3 | Activating | Trip4 | sp\|Q9QXN | 12 | 12 | 31,2 |
| P58468 | P58468 | Protein FA | Fam207a | sp\|P58468 | 8 | 8 | 50,2 |
| Q91V41 | Q91V41 | Ras-related | Rab14 | sp\|Q91V41 | 16 | 16 | 82,3 |
| Q9CQ74 | Q9CQ74 | Leptin rece | Leprotl1 | sp\|Q9CQ7 | 1 | 1 | 9,9 |
| P14576 | P14576 | Signal reco | Srp54 | sp\|P14576 | 20 | 20 | 47,2 |
| Q6DID7 | Q6DID7 | Protein wn | Wls | sp\|Q6DID7 | 9 | 9 | 16,6 |
| Q8K385 | Q8K385 | Ferric-chel | FRRS1 | sp\|Q8K385 | 6 | 6 | 12,7 |
| Q60634 | Q60634 | Flotillin-2 | Flot2 | sp\|Q60634 | 21 | 21 | 56,8 |
| O89084;Q( | O89084 | cAMP-spec | Pde4a | sp\|O89084 | 3 | 3 | 5,2 |
| Q8BXZ1 | Q8BXZ1 | Protein dis | Tmx3 | sp\|Q8BXZ1 | 18 | 18 | 45,8 |
| E9Q8I9 | E9Q8I9 | Protein fur | Fry | sp\|E9Q8I9 | 4 | 4 | 1,6 |
| Q8CFV9 | Q8CFV9 | Riboflavin | Rfk | sp\|Q8CFV9 | 4 | 4 | 32,3 |
| P62320 | P62320 | Small nucle | Snrpd3 | sp\|P62320 | 5 | 5 | 53,2 |
| Q9WV60 | Q9WV60 | Glycogen s | Gsk3b | sp\|Q9WV6 | 12 | 10 | 45,2 |
| Q8BH24 | Q8BH24 | Transmem | Tm9sf4 | sp\|Q8BH24 | 10 | 10 | 19,4 |
| Q64261 | Q64261 | Cyclin-dep | Cdk6 | sp\|Q64261 | 16 | 15 | 56,1 |
| Q91WE2 | Q91WE2 | Protein FA | Fam192a | sp\|Q91WE | 6 | 6 | 18,1 |
| Q9Z0R9 | Q9Z0R9 | Fatty acid ( | Fads2 | sp\|Q9Z0R9 | 11 | 11 | 27,5 |
| Q99JW4;Q | Q99JW4 | LIM and se | Lims1 | sp\|Q99JW | 10 | 10 | 34,8 |
| Q9ES97 | Q9ES97 | Reticulon-3 | Rtn3 | sp\|Q9ES97 | 4 | 4 | 7,3 |
| Q9CR21 | Q9CR21 | Acyl carrie | Ndufab1 | sp\|Q9CR21 | 2 | 2 | 9,6 |
| Q9CXU9 | Q9CXU9 | Eukaryotic | Eif1b | sp\|Q9CXU | 8 | 1 | 66,4 |
| P62082 | P62082 | 40S riboso | Rps7 | sp\|P62082 | 13 | 13 | 64,9 |
| Q8R5A6 | Q8R5A6 | TBC1 doma | Tbc1d22a | sp\|Q8R5A( | 2 | 2 | 5,6 |
| Q8C9S4 | Q8C9S4 | Coiled-coil | Ccdc186 | sp\|Q8C9S4 | 13 | 13 | 18,1 |
| Q9D1H7 | Q9D1H7 | Golgi to ER | Get4 | sp\|Q9D1H | 9 | 9 | 31,2 |
| Q9D1H8 | Q9D1H8 | 39S riboso | Mrpl53 | sp\|Q9D1H | 6 | 6 | 66,1 |
| Q9D136 | Q9D136 | 2-oxogluta | Ogfod3 | sp\|Q9D13( | 3 | 3 | 14,9 |
| Q66JV4 | Q66JV4 | RNA-bindir | Rbm12b2 | sp\|Q66JV4 | 18 | 6 | 24,2 |
| P53811 | P53811 | Phosphatid | Pitpnb | sp\|P53811 | 14 | 12 | 61,3 |
| Q5XJY4 | Q5XJY4 | Presenilins | Parl | sp\|Q5XJY4 | 4 | 4 | 13,3 |
| Q923U0 | Q923U0 | TOM1-like | Tom1l1 | sp\|Q923U( | 2 | 2 | 5,3 |
| P0DOV2;Q | P0DOV2 | | | sp\|P0DOV | 26 | 14 | 54 |
| P47757 | P47757 | F-actin-cap | Capzb | sp\|P47757 | 16 | 16 | 61 |
| B1AX39 | B1AX39 | Zinc finger | Zcchc7 | sp\|B1AX39 | 1 | 1 | 2,6 |
| Q8CJF9 | Q8CJF9 | Protein arg | Ago3 | sp\|Q8CJF9 | 14 | 3 | 24,1 |
| Q9EPC1 | Q9EPC1 | Alpha-parv | Parva | sp\|Q9EPC1 | 10 | 9 | 29,8 |
| P97478 | P97478 | 5-demetho | Coq7 | sp\|P97478 | 5 | 5 | 29,5 |
| Q8CEE0 | Q8CEE0 | Centrosom | Cep57 | sp\|Q8CEE( | 7 | 7 | 21,4 |
| Q9JI57 | Q9JI57 | General tra | Gtf2ird1 | sp\|Q9JI57 | 2 | 2 | 3 |
| Q8K190 | Q8K190 | SAYSvFN d | Saysd1 | sp\|Q8K19( | 2 | 2 | 28,2 |
| Q924H2 | Q924H2 | Mediator o | Med15 | sp\|Q924H2 | 13 | 13 | 23,8 |
| Q9CXV9;Q | Q9CXV9 | DCN1-like | Dcun1d5 | sp\|Q9CXV9 | 8 | 8 | 31,2 |
| Q9R190 | Q9R190 | Metastasis | Mta2 | sp\|Q9R19( | 27 | 26 | 53 |
| Q9ESW8 | Q9ESW8 | Pyroglutan | Pgpep1 | sp\|Q9ESW | 6 | 6 | 44,5 |
| P49282 | P49282 | Natural res | Slc11a2 | sp\|P49282 | 4 | 4 | 11,1 |
| Q8BWD8 | Q8BWD8 | Cyclin-dep | Cdk19 | sp\|Q8BWD | 5 | 2 | 13,2 |
| Q8BVH9 | Q8BVH9 | Methyltrar | Mettl6 | sp\|Q8BVH | 4 | 4 | 12,8 |
| P30051;P4 | P30051 | Transcripti | Tead1 | sp\|P30051 | 10 | 6 | 27,7 |
| Q5NCI0 | Q5NCI0 | Up-regulat | Urgcp | sp\|Q5NCI( | 18 | 18 | 26,2 |

| | | | | | | | |
|---|---|---|---|---|---|---|---|
| Q5SXY1 | Q5SXY1 | Cytospin-B | Specc1 | sp\|Q5SXY1 | 24 | 24 | 31 |
| Q9Z0H7 | Q9Z0H7 | B-cell lymp | Bcl10 | sp\|Q9Z0H7 | 5 | 5 | 21,9 |
| Q9Z2V6 | Q9Z2V6 | Histone de | Hdac5 | sp\|Q9Z2V6 | 4 | 4 | 3,9 |
| P60603 | P60603 | Reactive o | Romo1 | sp\|P60603 | 1 | 1 | 21,5 |
| P42567 | P42567 | Epidermal | Eps15 | sp\|P42567 | 27 | 27 | 44,7 |
| Q3UHC2 | Q3UHC2 | Leucine-ric | Lrrk1 | sp\|Q3UHC | 20 | 20 | 12,2 |
| Q8R2N0 | Q8R2N0 | Thyroid tra | Ccdc59 | sp\|Q8R2N | 4 | 4 | 18,8 |
| Q9QZM2 | Q9QZM2 | DNA polym | Polg2 | sp\|Q9QZM | 2 | 2 | 6,8 |
| Q9Z1E3 | Q9Z1E3 | NF-kappa-I | Nfkbia | sp\|Q9Z1E3 | 3 | 3 | 15 |
| Q8R2R3 | Q8R2R3 | Alpha- and | Aagab | sp\|Q8R2R3 | 6 | 6 | 32,6 |
| Q80WW9 | Q80WW9 | DDRGK do | Ddrgk1 | sp\|Q80WV | 5 | 5 | 30,8 |
| A2A8Z1 | A2A8Z1 | Oxysterol-l | Osbpl9 | sp\|A2A8Z1 | 16 | 15 | 28,5 |
| Q99K70;Q | Q99K70 | Ras-related | Rragc | sp\|Q99K70 | 12 | 12 | 39,2 |
| Q91WN1 | Q91WN1 | DnaJ homo | Dnajc9 | sp\|Q91WN | 14 | 14 | 49 |
| Q4VAA2 | Q4VAA2 | Protein CD | Cdv3 | sp\|Q4VAA | 15 | 15 | 67,3 |
| Q8K2M0 | Q8K2M0 | 39S riboso | Mrpl38 | sp\|Q8K2M | 10 | 10 | 37,6 |
| P67778 | P67778 | Prohibitin | Phb | sp\|P67778 | 18 | 18 | 72,8 |
| P50429 | P50429 | Arylsulfata | Arsb | sp\|P50429 | 8 | 8 | 15,4 |
| Q9CWZ7 | Q9CWZ7 | Gamma-so | Napg | sp\|Q9CWZ | 12 | 12 | 44,2 |
| Q8C6B9 | Q8C6B9 | Active regu | Rps19bp1 | sp\|Q8C6B9 | 6 | 6 | 41,3 |
| A2AJK6 | A2AJK6 | Chromodo | Chd7 | sp\|A2AJK6 | 21 | 12 | 8,6 |
| P61953 | P61953 | Guanine nu | Gng11 | sp\|P61953 | 2 | 2 | 17,8 |
| Q8BW70 | Q8BW70 | Ubiquitin c | Usp38 | sp\|Q8BW7 | 7 | 7 | 9 |
| Q8K2C8 | Q8K2C8 | Glycerol-3- | Agpat6 | sp\|Q8K2C8 | 8 | 7 | 14 |
| Q8VHH5 | Q8VHH5 | Arf-GAP wi | Agap3 | sp\|Q8VHH | 11 | 9 | 16,8 |
| P28571 | P28571 | Sodium- ar | Slc6a9 | sp\|P28571 | 3 | 3 | 4,2 |
| P61021 | P61021 | Ras-related | Rab5b | sp\|P61021 | 9 | 6 | 66,5 |
| Q8CIB5 | Q8CIB5 | Fermitin fa | Fermt2 | sp\|Q8CIB5 | 24 | 24 | 45,9 |
| P39749 | P39749 | Flap endon | Fen1 | sp\|P39749 | 12 | 11 | 37,6 |
| O70274 | O70274 | Protein tyr | Ptp4a2 | sp\|O70274 | 8 | 5 | 52,1 |
| Q91W18 | Q91W18 | Tudor dom | Tdrd3 | sp\|Q91W1 | 14 | 14 | 26,8 |
| Q8R0K9 | Q8R0K9 | Transcripti | E2f4 | sp\|Q8R0K9 | 7 | 7 | 25,6 |
| Q8VE09 | Q8VE09 | Tetratricop | Ttc39c | sp\|Q8VE09 | 8 | 8 | 18,6 |
| Q9CZ30 | Q9CZ30 | Obg-like AT | Ola1 | sp\|Q9CZ30 | 24 | 24 | 68,2 |
| Q3TQB2 | Q3TQB2 | FAD-deper | Foxred1 | sp\|Q3TQB | 5 | 5 | 16,6 |
| Q5SXJ3 | Q5SXJ3 | Fanconi an | Brip1 | sp\|Q5SXJ3 | 15 | 15 | 17,1 |
| Q9ER62 | Q9ER62 | Tumor nec | Tnfrsf22 | sp\|Q9ER62 | 3 | 3 | 20,2 |
| Q8BP27 | Q8BP27 | Swi5-depe | Sfr1 | sp\|Q8BP27 | 10 | 10 | 38,6 |
| Q3U6U5 | Q3U6U5 | Putative G | Gtpbp6 | sp\|Q3U6U | 4 | 4 | 10,1 |
| Q99NB8 | Q99NB8 | Ubiquilin-4 | Ubqln4 | sp\|Q99NB | 11 | 9 | 29,9 |
| Q08288 | Q08288 | Cell growth | Lyar | sp\|Q08288 | 19 | 19 | 44,8 |
| P62915 | P62915 | Transcripti | Gtf2b | sp\|P62915 | 15 | 15 | 56,3 |
| Q60929;Q8 | Q60929;Q8CFI | Myocyte-s | Mef2a;Me | sp\|Q60929 | 4 | 3 | 11,8 |
| Q6P9Q6 | Q6P9Q6 | FK506-binc | Fkbp15 | sp\|Q6P9Q | 16 | 16 | 21,1 |
| Q6X893 | Q6X893 | Choline tra | Slc44a1 | sp\|Q6X893 | 10 | 10 | 21 |
| Q9DBG5 | Q9DBG5 | Perilipin-3 | Plin3 | sp\|Q9DBG | 17 | 17 | 70 |
| Q8BGF6 | Q8BGF6 | ELMO dom | Elmod2 | sp\|Q8BGF | 4 | 4 | 14,7 |
| O88693 | O88693 | Ceramide g | Ugcg | sp\|O88693 | 3 | 3 | 11,9 |
| Q8K1J6 | Q8K1J6 | CCA tRNA r | Trnt1 | sp\|Q8K1J6 | 16 | 16 | 46,1 |
| Q9EPK5 | Q9EPK5 | WW doma | Wwtr1 | sp\|Q9EPK5 | 6 | 6 | 18,7 |

| | | | | | | | |
|---|---|---|---|---|---|---|---|
| Q60838 | Q60838 | Segment p | Dvl2 | sp\|Q60838 | 12 | 10 | 26,8 |
| Q8CIC2 | Q8CIC2 | Nucleopor | Nupl2 | sp\|Q8CIC2 | 8 | 8 | 33,6 |
| Q6P1G0 | Q6P1G0 | HEAT repe | Heatr6 | sp\|Q6P1G( | 18 | 18 | 23,2 |
| Q91VZ6 | Q91VZ6 | Stromal m | Smap1 | sp\|Q91VZ( | 4 | 4 | 13,2 |
| Q6NXH3 | Q6NXH3 | Vasculin | Gpbp1 | sp\|Q6NXH | 2 | 2 | 8 |
| Q9JKY5 | Q9JKY5 | Huntingtin | Hip1r | sp\|Q9JKY5 | 7 | 5 | 8,6 |
| P01897;P0 | P01897 | H-2 class I | H2-L | sp\|P01897 | 15 | 2 | 44,8 |
| Q8C341 | Q8C341 | SUN doma | Suco | sp\|Q8C341 | 4 | 4 | 4,2 |
| Q6IR34 | Q6IR34 | G-protein-s | Gpsm1 | sp\|Q6IR34 | 6 | 5 | 15,2 |
| Q8CGF6 | Q8CGF6 | WD repeat | Wdr47 | sp\|Q8CGF( | 7 | 7 | 9,1 |
| Q9CQQ8 | Q9CQQ8 | U6 snRNA- | Lsm7 | sp\|Q9CQQ | 4 | 4 | 57,3 |
| Q8BX57 | Q8BX57 | PX domain | Pxk | sp\|Q8BX57 | 9 | 9 | 22,5 |
| Q8R050;Q | Q8R050 | Eukaryotic | Gspt1 | sp\|Q8R05( | 25 | 25 | 46,1 |
| Q6P6J9 | Q6P6J9 | Thioredoxi | Txndc15 | sp\|Q6P6J9 | 10 | 10 | 44,5 |
| Q8C886 | Q8C886 | Probable p | Plekhn1 | sp\|Q8C88( | 2 | 2 | 4,8 |
| Q8C0L0 | Q8C0L0 | Thioredoxi | Tmx4 | sp\|Q8C0L( | 6 | 6 | 20,6 |
| Q61687 | Q61687 | Transcripti | Atrx | sp\|Q61687 | 41 | 41 | 24,6 |
| Q9DCL9 | Q9DCL9 | Multifunct | Paics | sp\|Q9DCL9 | 26 | 26 | 65,9 |
| Q3UHB1 | Q3UHB1 | 5-nucleoti | Nt5dc3 | sp\|Q3UHB | 21 | 21 | 45,4 |
| Q922K7 | Q922K7 | Probable 2 | Nop2 | sp\|Q922K7 | 26 | 26 | 40,6 |
| Q149F1 | Q149F1 | RNA pseud | Rpusd2 | sp\|Q149F1 | 10 | 10 | 28,6 |
| Q8BGE4 | Q8BGE4 | Sushi doma | Susd6 | sp\|Q8BGE4 | 1 | 1 | 4,6 |
| P62196 | P62196 | 26S protea | Psmc5 | sp\|P62196 | 31 | 31 | 68,2 |
| Q9D338 | Q9D338 | 39S riboso | Mrpl19 | sp\|Q9D338 | 11 | 11 | 49,7 |
| O08789 | O08789 | Max-bindir | Mnt | sp\|O08789 | 6 | 6 | 13,4 |
| CON__Q3Z | CON__Q3ZBS7 | | | | 8 | 8 | 20,2 |
| Q99J27 | Q99J27 | Acetyl-coe | Slc33a1 | sp\|Q99J27 | 10 | 10 | 19,3 |
| O70481 | O70481 | E3 ubiquiti | Ubr1 | sp\|O70481 | 22 | 22 | 16,8 |
| Q8C3X4 | Q8C3X4 | Translation | Guf1 | sp\|Q8C3X4 | 13 | 13 | 30,1 |
| Q925N2 | Q925N2 | Sideroflexi | Sfxn2 | sp\|Q925N2 | 5 | 5 | 18,9 |
| P62334 | P62334 | 26S protea | Psmc6 | sp\|P62334 | 20 | 20 | 59,6 |
| Q99L00 | Q99L00 | HAUS augn | Haus8 | sp\|Q99L0( | 2 | 2 | 6,7 |
| Q8BGC4 | Q8BGC4 | Zinc-bindir | Zadh2 | sp\|Q8BGC | 10 | 10 | 31,8 |
| A2ALK8 | A2ALK8 | Tyrosine-p | Ptpn3 | sp\|A2ALK8 | 3 | 3 | 3,7 |
| Q8VDL4 | Q8VDL4 | ADP-deper | Adpgk | sp\|Q8VDL4 | 14 | 14 | 39,3 |
| Q9D786 | Q9D786 | HAUS augn | Haus5 | sp\|Q9D78( | 14 | 14 | 30,2 |
| Q3V4B5 | Q3V4B5 | COMM dor | Commd6 | sp\|Q3V4B5 | 3 | 3 | 47,1 |
| Q9D2C6 | Q9D2C6 | DNA-direct | Polr3h | sp\|Q9D2C( | 7 | 7 | 35,8 |
| Q7TN79 | Q7TN79 | A-kinase a | Akap7 | sp\|Q7TN79 | 4 | 4 | 14 |
| Q922V4 | Q922V4 | Pleiotropic | Plrg1 | sp\|Q922V4 | 20 | 20 | 55,8 |
| P61082 | P61082 | NEDD8-cor | Ube2m | sp\|P61082 | 8 | 8 | 43,2 |
| Q9ERG0 | Q9ERG0 | LIM domai | Lima1 | sp\|Q9ERG( | 22 | 22 | 39,7 |
| Q9QWH1 | Q9QWH1 | Polyhomeo | Phc2 | sp\|Q9QWI | 11 | 10 | 17,9 |
| Q61103 | Q61103 | Zinc finger | Dpf2 | sp\|Q61103 | 17 | 17 | 54 |
| P63034 | P63034 | Cytohesin- | Cyth2 | sp\|P63034 | 11 | 6 | 32,2 |
| Q6NSQ9 | Q6NSQ9 | Glucose-6- | G6pc3 | sp\|Q6NSQ | 2 | 2 | 9,5 |
| Q8QZS1 | Q8QZS1 | 3-hydroxyi | Hibch | sp\|Q8QZS1 | 12 | 12 | 39 |
| Q9CPY1 | Q9CPY1 | 39S riboso | Mrpl51 | sp\|Q9CPY1 | 3 | 3 | 22,7 |
| A2A9A2 | A2A9A2 | Doublesex | Dmrta2 | sp\|A2A9A2 | 2 | 2 | 5,5 |
| O35083 | O35083 | 1-acyl-sn-g | Agpat1 | sp\|O35083 | 4 | 4 | 23,5 |

| | | | | | | | |
|---|---|---|---|---|---|---|---|
| O88573 | O88573 | AF4/FMR2 | Aff1 | sp\|O88573 | 7 | 7 | 9 |
| Q6PDK2 | Q6PDK2 | Histone-lys | Kmt2d | sp\|Q6PDK2 | 29 | 29 | 10,1 |
| O88351 | O88351 | Inhibitor o | Ikbkb | sp\|O88351 | 16 | 15 | 29,5 |
| Q9CTH6 | Q9CTH6 | rRNA-proc | Fcf1 | sp\|Q9CTH6 | 5 | 5 | 31,3 |
| Q9CWW7 | Q9CWW7 | CXXC-type | Cxxc1 | sp\|Q9CWW | 12 | 12 | 25,8 |
| Q3THE2 | Q3THE2 | Myosin reg | Myl12b | sp\|Q3THE2 | 10 | 5 | 62,8 |
| O35381;Q( | O35381 | Acidic leuc | Anp32a | sp\|O35381 | 12 | 10 | 43,3 |
| Q5DU37 | Q5DU37 | Zinc finger | Zfyve26 | sp\|Q5DU3 | 5 | 5 | 3,4 |
| Q8BVI5 | Q8BVI5 | Syntaxin-1 | Stx16 | sp\|Q8BVI5 | 4 | 4 | 16 |
| P80315 | P80315 | T-complex | Cct4 | sp\|P80315 | 41 | 41 | 80,1 |
| Q62313;Q( | Q62313;Q623 | Trans-Golg | Tgoln1;Tgo | sp\|Q62313 | 4 | 4 | 15,9 |
| Q8N9S3 | Q8N9S3 | Activator o | Ahsa2 | sp\|Q8N9S3 | 11 | 11 | 42,6 |
| Q8K2A7 | Q8K2A7 | Integrator | Ints10 | sp\|Q8K2A7 | 21 | 21 | 34,5 |
| Q9DBC7;P | Q9DBC7 | cAMP-depe | Prkar1a | sp\|Q9DBC | 18 | 18 | 53,3 |
| Q8C167 | Q8C167 | Prolyl endo | Prepl | sp\|Q8C167 | 10 | 10 | 21,5 |
| Q9JL56 | Q9JL56 | Glyceropho | Gde1 | sp\|Q9JL56 | 6 | 6 | 20,8 |
| P58043 | P58043 | Sestrin-2 | Sesn2 | sp\|P58043 | 4 | 4 | 10,4 |
| Q8C6E0 | Q8C6E0 | Cilia- and f | Cfap36 | sp\|Q8C6E0 | 9 | 9 | 39,4 |
| Q9JLR1 | Q9JLR1 | Protein tra | Sec61a2 | sp\|Q9JLR1 | 6 | 3 | 14,3 |
| Q8R2W9 | Q8R2W9 | Pantothen | Pank3 | sp\|Q8R2W | 9 | 7 | 31,4 |
| Q3TMV7 | Q3TMV7 | Pyridine nu | Pyroxd1 | sp\|Q3TMV | 4 | 4 | 13,1 |
| Q9CY97 | Q9CY97 | RNA polym | Ssu72 | sp\|Q9CY97 | 7 | 7 | 41,8 |
| Q9CXY6 | Q9CXY6 | Interleukin | Ilf2 | sp\|Q9CXY6 | 18 | 18 | 63,8 |
| Q0VBL3 | Q0VBL3 | | Rbm15 | sp\|Q0VBL3 | 25 | 24 | 31,3 |
| Q9CQT1 | Q9CQT1 | Methylthio | Mri1 | sp\|Q9CQT | 12 | 12 | 57,5 |
| Q8R1J3 | Q8R1J3 | Zinc finger | Zcchc9 | sp\|Q8R1J3 | 4 | 4 | 23,2 |
| Q9CR16 | Q9CR16 | Peptidyl-pr | Ppid | sp\|Q9CR16 | 25 | 25 | 72,4 |
| Q8BLY2 | Q8BLY2 | Probable t | Tarsl2 | sp\|Q8BLY2 | 8 | 6 | 12 |
| Q9QUI1 | Q9QUI1 | Protein FA | Fam89b | sp\|Q9QUI1 | 2 | 2 | 14,3 |
| Q8CHW4 | Q8CHW4 | Translation | Eif2b5 | sp\|Q8CHW | 19 | 19 | 37 |
| Q8C0D0 | Q8C0D0 | Probable t | Trub1 | sp\|Q8C0D0 | 4 | 4 | 13 |
| Q8BS39 | Q8BS39 | Uncharacte | Ccdc32 | sp\|Q8BS39 | 1 | 1 | 17,9 |
| Q9JJK5 | Q9JJK5 | Homocyste | Herpud1 | sp\|Q9JJK5 | 2 | 2 | 7,7 |
| P62983;P6 | P62983;P6298 | Ubiquitin-4 | Rps27a;Ub | sp\|P62983 | 10 | 10 | 60,3 |
| Q9R002 | Q9R002 | Interferon- | Ifi202 | sp\|Q9R002 | 10 | 10 | 23,8 |
| Q8BH15 | Q8BH15 | CCR4-NOT | Cnot10 | sp\|Q8BH1 | 21 | 21 | 40,9 |
| Q80Z96 | Q80Z96 | Vang-like p | Vangl1 | sp\|Q80Z96 | 8 | 8 | 24,9 |
| Q8CHK3 | Q8CHK3 | Lysophosp | Mboat7 | sp\|Q8CHK | 6 | 6 | 19,2 |
| Q9JK83 | Q9JK83 | Partitionin | Pard6b | sp\|Q9JK83 | 6 | 6 | 17,5 |
| Q8C6U2 | Q8C6U2 | PQ-loop re | Pqlc3 | sp\|Q8C6U | 3 | 3 | 20,8 |
| Q62384 | Q62384 | Zinc finger | Zpr1 | sp\|Q62384 | 13 | 13 | 45,5 |
| Q6P6J0 | Q6P6J0 | tRNA-speci | Adat2 | sp\|Q6P6J0 | 4 | 4 | 33,5 |
| Q922Y1 | Q922Y1 | UBX doma | Ubxn1 | sp\|Q922Y1 | 12 | 12 | 50,5 |
| Q8CC88 | Q8CC88 | von Willeb | Vwa8 | sp\|Q8CC88 | 46 | 46 | 32,4 |
| Q64516;Q9 | Q64516 | Glycerol ki | Gk | sp\|Q64516 | 19 | 19 | 44,2 |
| Q5BLK4 | Q5BLK4 | Terminal u | Zcchc6 | sp\|Q5BLK4 | 18 | 16 | 16,9 |
| P62774 | P62774 | Myotrophi | Mtpn | sp\|P62774 | 6 | 6 | 75,4 |
| P12787 | P12787 | Cytochrom | Cox5a | sp\|P12787 | 9 | 9 | 54,1 |
| Q8C0M9 | Q8C0M9 | Isoaspartyl | Asrgl1 | sp\|Q8C0M | 7 | 7 | 25,8 |
| P55012;P5 | P55012 | Solute carr | Slc12a2 | sp\|P55012 | 30 | 30 | 28,7 |

| | | | | | | | |
|---|---|---|---|---|---|---|---|
| P62482 | P62482 | Voltage-ga | Kcnab2 | sp\|P62482 | 9 | 9 | 38,7 |
| Q8R464 | Q8R464 | Cell adhesi | Cadm4 | sp\|Q8R464 | 5 | 5 | 24 |
| F7BJB9 | F7BJB9 | | Morc3 | sp\|F7BJB9 | 31 | 31 | 46,1 |
| Q8BGG7 | Q8BGG7 | Ubiquitin-a | Ubash3b | sp\|Q8BGG | 2 | 2 | 5 |
| Q923G2 | Q923G2 | DNA-direct | Polr2h | sp\|Q923G2 | 6 | 6 | 57,3 |
| Q5PRF0 | Q5PRF0 | HEAT repe | Heatr5a | sp\|Q5PRF0 | 30 | 30 | 21 |
| Q3U319 | Q3U319 | E3 ubiquiti | Rnf40 | sp\|Q3U319 | 32 | 29 | 34,6 |
| P55258 | P55258 | Ras-relatec | Rab8a | sp\|P55258 | 10 | 5 | 52,2 |
| P39688 | P39688 | Tyrosine-p | Fyn | sp\|P39688 | 6 | 2 | 11,9 |
| Q8BWY7 | Q8BWY7 | Zinc transp | Slc39a11 | sp\|Q8BWY | 2 | 2 | 12,9 |
| Q80YA3 | Q80YA3 | Phospholip | Ddhd1 | sp\|Q80YA3 | 4 | 4 | 9,5 |
| Q8R3V5 | Q8R3V5 | Endophilin | Sh3glb2 | sp\|Q8R3V5 | 12 | 11 | 41 |
| Q9CQ22 | Q9CQ22 | Ragulator c | Lamtor1 | sp\|Q9CQ2 | 9 | 9 | 75,2 |
| Q9DBA6 | Q9DBA6 | Peroxisom | Tysnd1 | sp\|Q9DBA | 3 | 3 | 8,3 |
| P70188 | P70188 | Kinesin-ass | Kifap3 | sp\|P70188 | 8 | 8 | 12,6 |
| O35491 | O35491 | Dual specif | Clk2 | sp\|O35491 | 3 | 3 | 6 |
| Q80ZM7;R | Q80ZM7 | Transcripti | Gtf2a2 | sp\|Q80ZM | 3 | 3 | 29,4 |
| Q9D211 | Q9D211 | CDKN2AIP | Cdkn2aipn | sp\|Q9D211 | 3 | 3 | 35,3 |
| Q91WC9 | Q91WC9 | Sn1-specifi | Daglb | sp\|Q91WC | 3 | 3 | 5,7 |
| Q9CVD2 | Q9CVD2 | Ataxin-3 | Atxn3 | sp\|Q9CVD | 5 | 5 | 21,1 |
| P10833 | P10833 | Ras-relatec | Rras | sp\|P10833 | 8 | 6 | 48,2 |
| Q9R0A5 | Q9R0A5 | Serine/thre | Nek3 | sp\|Q9R0A5 | 3 | 3 | 7,4 |
| Q9Z1W9 | Q9Z1W9 | STE20/SPS | Stk39 | sp\|Q9Z1W | 9 | 7 | 30,6 |
| Q9QXK7 | Q9QXK7 | Cleavage a | Cpsf3 | sp\|Q9QXK | 25 | 25 | 44,3 |
| Q61221 | Q61221 | Hypoxia-in | Hif1a | sp\|Q61221 | 4 | 4 | 7,3 |
| Q9JK81 | Q9JK81 | UPF0160 p | Myg1 | sp\|Q9JK81 | 15 | 15 | 46,6 |
| Q61166 | Q61166 | Microtubu | Mapre1 | sp\|Q61166 | 18 | 16 | 63,8 |
| Q99LJ5 | Q99LJ5 | CKLF-like M | Cmtm3 | sp\|Q99LJ5 | 1 | 1 | 10,3 |
| P58871 | P58871 | 182 kDa ta | Tnks1bp1 | sp\|P58871 | 54 | 54 | 49,8 |
| Q923Z0 | Q923Z0 | G-protein c | Gprc5b | sp\|Q923Z0 | 2 | 2 | 6,1 |
| O08832 | O08832 | Polypeptid | Galnt4 | sp\|O08832 | 7 | 5 | 14 |
| Q8CHS8 | Q8CHS8 | Vacuolar p | Vps37a | sp\|Q8CHS8 | 7 | 7 | 24,9 |
| Q99KI3 | Q99KI3 | ER membra | Emc3 | sp\|Q99KI3 | 13 | 13 | 49,8 |
| Q5SV77 | Q5SV77 | Gametoge | Ggnbp2 | sp\|Q5SV77 | 4 | 4 | 7,9 |
| Q8BXA5 | Q8BXA5 | Cleft lip an | Clptm1l | sp\|Q8BXA5 | 5 | 5 | 17,4 |
| P24668 | P24668 | Cation-dep | M6pr | sp\|P24668 | 6 | 6 | 25,2 |
| O70479 | O70479 | BTB/POZ d | Tnfaip1 | sp\|O70479 | 3 | 2 | 9,8 |
| Q9D8K8 | Q9D8K8 | Solute carr | Slc25a39 | sp\|Q9D8K8 | 5 | 5 | 18,4 |
| Q923A2 | Q923A2 | Protein Spi | Spdl1 | sp\|Q923A2 | 13 | 13 | 30,6 |
| Q03141;Q8 | Q03141 | MAP/micro | Mark3 | sp\|Q03141 | 22 | 17 | 40 |
| Q3TFQ1 | Q3TFQ1 | SPRY doma | Spryd7 | sp\|Q3TFQ1 | 5 | 5 | 35,7 |
| Q8CBQ5 | Q8CBQ5 | Phosphatid | Pi4k2b | sp\|Q8CBQ | 7 | 7 | 16 |
| Q8BG60 | Q8BG60 | Thioredoxi | Txnip | sp\|Q8BG60 | 6 | 6 | 15,6 |
| Q8VBW1 | Q8VBW1 | Sodium- ar | Slc6a8 | sp\|Q8VBW | 4 | 3 | 7 |
| Q9DBA9 | Q9DBA9 | General tra | Gtf2h1 | sp\|Q9DBA | 5 | 5 | 10,6 |
| Q60605 | Q60605 | Myosin ligh | Myl6 | sp\|Q60605 | 9 | 8 | 60,3 |
| Q6PEE3 | Q6PEE3 | Ribonuclec | Rrm2b | sp\|Q6PEE3 | 4 | 2 | 13,4 |
| Q6DFV7 | Q6DFV7 | Nuclear re | Ncoa7 | sp\|Q6DFV7 | 6 | 6 | 8,5 |
| Q8C2A2 | Q8C2A2 | tRNA-splici | Tsen54 | sp\|Q8C2A2 | 7 | 7 | 19,8 |
| Q8BQR4 | Q8BQR4 | KAT8 regul | Kansl2 | sp\|Q8BQR | 6 | 6 | 18,5 |

| | | | | | | | |
|---|---|---|---|---|---|---|---|
| Q812A2 | Q812A2 | SLIT-ROBO | Srgap3 | sp\|Q812A2 | 8 | 8 | 9,6 |
| Q99KV1 | Q99KV1 | DnaJ homo | Dnajb11 | sp\|Q99KV1 | 13 | 13 | 33,5 |
| P19426 | P19426 | Negative e | Nelfe | sp\|P19426 | 13 | 13 | 43,2 |
| Q9CYW4 | Q9CYW4 | Haloacid d | Hdhd3 | sp\|Q9CYW | 8 | 8 | 47,8 |
| Q91Z49 | Q91Z49 | UAP56-inte | Fyttd1 | sp\|Q91Z49 | 8 | 8 | 31,9 |
| E9QAT4 | E9QAT4 | | Sec16a | sp\|E9QAT4 | 27 | 27 | 20,7 |
| Q8BJ03 | Q8BJ03 | Cytochrom | Cox15 | sp\|Q8BJ03 | 8 | 8 | 25,7 |
| Q8R3I3 | Q8R3I3 | Conserved | Cog6 | sp\|Q8R3I3 | 14 | 14 | 27,5 |
| B1AZI6 | B1AZI6 | THO comp | Thoc2 | sp\|B1AZI6 | 33 | 33 | 26,4 |
| Q8BWY3 | Q8BWY3 | Eukaryotic | Etf1 | sp\|Q8BWY | 23 | 23 | 60,6 |
| Q8BHF7 | Q8BHF7 | CDP-diacyl | Pgs1 | sp\|Q8BHF7 | 19 | 19 | 46,3 |
| Q9JM51 | Q9JM51 | Prostaglan | Ptges | sp\|Q9JM5 | 4 | 4 | 14,4 |
| Q921X9 | Q921X9 | Protein dis | Pdia5 | sp\|Q921X9 | 14 | 14 | 32,7 |
| Q8BH55 | Q8BH55 | Threonine | Thnsl1 | sp\|Q8BH5 | 7 | 7 | 13 |
| Q8BMB3 | Q8BMB3 | Eukaryotic | Eif4e2 | sp\|Q8BMB | 6 | 6 | 34,7 |
| Q9DBZ5 | Q9DBZ5 | Eukaryotic | Eif3k | sp\|Q9DBZ5 | 10 | 10 | 61,9 |
| P60840 | P60840 | Alpha-endo | Ensa | sp\|P60840 | 3 | 2 | 36,4 |
| Q9JIS8 | Q9JIS8 | Solute carr | Slc12a4 | sp\|Q9JIS8 | 22 | 18 | 26 |
| P81269 | P81269 | Cyclic AMP | Atf1 | sp\|P81269 | 3 | 1 | 15,6 |
| Q91V81 | Q91V81 | RNA-bindin | Rbm42 | sp\|Q91V81 | 9 | 9 | 31,6 |
| Q80U72 | Q80U72 | Protein scr | Scrib | sp\|Q80U72 | 44 | 41 | 41,8 |
| Q3U3W5 | Q3U3W5 | Putative pr | Prmt9 | sp\|Q3U3W | 10 | 10 | 16 |
| Q5XF89 | Q5XF89 | Probable c | Atp13a3 | sp\|Q5XF89 | 8 | 8 | 10,4 |
| Q9D773 | Q9D773 | 39S riboso | Mrpl2 | sp\|Q9D773 | 7 | 7 | 33 |
| Q8BLF1 | Q8BLF1 | Neutral ch | Nceh1 | sp\|Q8BLF1 | 13 | 13 | 38,5 |
| Q9DBV3 | Q9DBV3 | Probable A | Dhx34 | sp\|Q9DBV | 6 | 6 | 8,5 |
| Q9QZ06 | Q9QZ06 | Toll-intera | Tollip | sp\|Q9QZ0 | 8 | 8 | 44,5 |
| P10605 | P10605 | Cathepsin | Ctsb | sp\|P10605 | 13 | 13 | 45,4 |
| Q61193 | Q61193 | Ral guanin | Rgl2 | sp\|Q61193 | 7 | 7 | 14,7 |
| Q7TNC4 | Q7TNC4 | Putative RI | Luc7l2 | sp\|Q7TNC | 11 | 7 | 28,6 |
| Q91YN5 | Q91YN5 | UDP-N-ace | Uap1 | sp\|Q91YN | 20 | 19 | 44,1 |
| Q80UZ2 | Q80UZ2 | Protein SD | Sdad1 | sp\|Q80UZ | 17 | 17 | 30,7 |
| Q8K274 | Q8K274 | Ketosamin | Fn3krp | sp\|Q8K274 | 4 | 4 | 11 |
| O35126 | O35126 | Atrophin-1 | Atn1 | sp\|O35126 | 3 | 3 | 3,8 |
| Q99LD9 | Q99LD9 | Translation | Eif2b2 | sp\|Q99LD9 | 12 | 12 | 41 |
| O35451 | O35451 | Cyclic AMP | Atf6b | sp\|O35451 | 5 | 5 | 8,6 |
| Q91W67 | Q91W67 | Ubiquitin-l | Ubl7 | sp\|Q91W6 | 6 | 6 | 23,4 |
| P54775 | P54775 | 26S protea | Psmc4 | sp\|P54775 | 30 | 30 | 63,9 |
| Q91WK5 | Q91WK5 | Glycine cle | Gcsh | sp\|Q91WK | 3 | 3 | 34,7 |
| Q8CG47 | Q8CG47 | Structural | Smc4 | sp\|Q8CG47 | 56 | 56 | 46,7 |
| CON__Q05 | CON__Q05B55 | | | | 3 | 3 | 23,3 |
| Q9QYI4 | Q9QYI4 | DnaJ homo | Dnajb12 | sp\|Q9QYI4 | 8 | 8 | 33,5 |
| Q8R5H6 | Q8R5H6 | Wiskott-Al | Wasf1 | sp\|Q8R5H | 7 | 6 | 16,1 |
| Q80U70 | Q80U70 | Polycomb | Suz12 | sp\|Q80U7 | 24 | 24 | 51,1 |
| Q8R3P2 | Q8R3P2 | Probable E | Dtx2 | sp\|Q8R3P2 | 3 | 3 | 6,8 |
| Q8R1K4 | Q8R1K4 | 5-phospho | Phykpl | sp\|Q8R1K4 | 6 | 6 | 15,6 |
| Q9D8C3 | Q9D8C3 | Putative bi | Alg13 | sp\|Q9D8C | 2 | 2 | 2,2 |
| P54729 | P54729 | NEDD8 ulti | Nub1 | sp\|P54729 | 20 | 20 | 41 |
| P20444;P6 | P20444 | Protein kin | Prkca | sp\|P20444 | 12 | 8 | 25,3 |
| Q8BVN4 | Q8BVN4 | Transcripti | Mterf4 | sp\|Q8BVN | 4 | 4 | 11,6 |

| | | | | | | | |
|---|---|---|---|---|---|---|---|
| Q99LC8 | Q99LC8 | Translation | Eif2b1 | sp\|Q99LC8 | 9 | 9 | 35,1 |
| Q9D6T0 | Q9D6T0 | Nitric oxide | Nosip | sp\|Q9D6T0 | 11 | 11 | 51,5 |
| P53762 | P53762 | Aryl hydro | Arnt | sp\|P53762 | 12 | 12 | 21,9 |
| CON__P50 | CON__P50448 | | | | 4 | 4 | 12 |
| Q3UD01 | Q3UD01 | Putative at | Atxn7l3b | sp\|Q3UD0 | 3 | 3 | 44,3 |
| Q9JJA4 | Q9JJA4 | Ribosome | Wdr12 | sp\|Q9JJA4 | 14 | 14 | 45,9 |
| P70303 | P70303 | CTP syntha | Ctps2 | sp\|P70303 | 13 | 11 | 30 |
| Q9QXK3 | Q9QXK3 | Coatomer | Copg2 | sp\|Q9QXK | 25 | 21 | 39,5 |
| Q9CXG3 | Q9CXG3 | Peptidyl-pr | Ppil4 | sp\|Q9CXG | 18 | 18 | 45,1 |
| O88839 | O88839 | Disintegrin | Adam15 | sp\|O88839 | 4 | 4 | 6,5 |
| Q8C3W1 | Q8C3W1 | Uncharacterized prote | | sp\|Q8C3W | 7 | 7 | 32,3 |
| Q8K284 | Q8K284 | General tra | Gtf3c1 | sp\|Q8K284 | 48 | 48 | 31,9 |
| Q7TSI3 | Q7TSI3 | Serine/thre | Ppp6r1 | sp\|Q7TSI3 | 16 | 16 | 25,7 |
| Q99KK2 | Q99KK2 | N-acylneur | Cmas | sp\|Q99KK2 | 12 | 12 | 33,3 |
| Q9CPV1 | Q9CPV1 | Spindle an | Ska1 | sp\|Q9CPV1 | 1 | 1 | 4,3 |
| Q2VPU4 | Q2VPU4 | MLX-intera | Mlxip | sp\|Q2VPU | 2 | 2 | 3,6 |
| P19536 | P19536 | Cytochrom | Cox5b | sp\|P19536 | 6 | 6 | 35,9 |
| Q9D8T0;P9 | Q9D8T0 | Protein FA | Fam3a | sp\|Q9D8T0 | 7 | 7 | 43 |
| O55100 | O55100 | Synaptogy | Syngr1 | sp\|O55100 | 1 | 1 | 5,1 |
| Q9DCI9 | Q9DCI9 | 39S riboso | Mrpl32 | sp\|Q9DCI9 | 3 | 3 | 23 |
| Q3UFY0 | Q3UFY0 | Ribosomal | Rrp36 | sp\|Q3UFY0 | 3 | 3 | 13,1 |
| Q9D8X2 | Q9D8X2 | Coiled-coil | Ccdc124 | sp\|Q9D8X2 | 7 | 7 | 30,4 |
| Q9JM52;P8 | Q9JM52;P8351 | Misshapen | Mink1;Tnik | sp\|Q9JM5 | 12 | 5 | 10,9 |
| O08810 | O08810 | 116 kDa U5 | Eftud2 | sp\|O08810 | 52 | 51 | 70 |
| Q8C0R0 | Q8C0R0 | Ubiquitin c | Usp37 | sp\|Q8C0R0 | 4 | 4 | 5,7 |
| Q9JKC8;Q8 | Q9JKC8 | AP-3 comp | Ap3m1 | sp\|Q9JKC8 | 14 | 14 | 52,2 |
| Q9CZS3 | Q9CZS3 | LisH doma | Fopnl | sp\|Q9CZS3 | 1 | 1 | 12,1 |
| Q60668 | Q60668 | Heterogen | Hnrnpd | sp\|Q60668 | 24 | 23 | 44,5 |
| Q61771 | Q61771 | Kinesin-like | Kif3b | sp\|Q61771 | 10 | 7 | 17,1 |
| Q7TQC7 | Q7TQC7 | G patch do | Gpatch2 | sp\|Q7TQC | 4 | 4 | 12,7 |
| P59328 | P59328 | WD repeat | Wdhd1 | sp\|P59328 | 37 | 37 | 47,4 |
| Q99L28 | Q99L28 | Probable ri | Rsl24d1 | sp\|Q99L28 | 4 | 4 | 28,2 |
| O35658 | O35658 | Compleme | C1qbp | sp\|O35658 | 7 | 7 | 34,5 |
| Q924T7 | Q924T7 | E3 ubiquiti | Rnf31 | sp\|Q924T7 | 8 | 8 | 12,5 |
| Q8CI51 | Q8CI51 | PDZ and LI | Pdlim5 | sp\|Q8CI51 | 26 | 26 | 59,1 |
| Q791V5 | Q791V5 | Mitochond | Mtch2 | sp\|Q791V5 | 12 | 12 | 55,8 |
| O89051 | O89051 | Integral me | Itm2b | sp\|O89051 | 9 | 9 | 47,7 |
| Q3UJU9 | Q3UJU9 | Regulator c | Rmdn3 | sp\|Q3UJU9 | 8 | 8 | 24,3 |
| Q99M28 | Q99M28 | RNA-bindir | Rnps1 | sp\|Q99M2 | 7 | 7 | 27,9 |
| Q9WVJ9 | Q9WVJ9 | EGF-contai | Efemp2 | sp\|Q9WVJ | 2 | 2 | 5,4 |
| Q9Z222 | Q9Z222 | N-acetyllac | B3GNT2 | sp\|Q9Z222 | 5 | 5 | 15,4 |
| Q810V0 | Q810V0 | U3 small n | Mphosph1 | sp\|Q810V0 | 15 | 15 | 30,4 |
| Q9WV72 | Q9WV72 | Ankyrin re | Asb3 | sp\|Q9WV7 | 5 | 5 | 16,6 |
| Q9JI44 | Q9JI44 | DNA meth | Dmap1 | sp\|Q9JI44 | 13 | 13 | 39,3 |
| Q62511 | Q62511 | E3 ubiquiti | Zfp91 | sp\|Q62511 | 4 | 4 | 10 |
| Q9Z1E4;Q8 | Q9Z1E4 | Glycogen [ | Gys1 | sp\|Q9Z1E4 | 20 | 20 | 32,9 |
| O88874 | O88874 | Cyclin-K | Ccnk | sp\|O88874 | 9 | 9 | 26,9 |
| P61620 | P61620 | Protein tra | Sec61a1 | sp\|P61620 | 8 | 5 | 17,6 |
| P70353 | P70353 | Nuclear tra | Nfyc | sp\|P70353 | 5 | 5 | 20,6 |
| Q9CY16 | Q9CY16 | 28S riboso | Mrps28 | sp\|Q9CY16 | 7 | 7 | 45,7 |

| | | | | | | | |
|---|---|---|---|---|---|---|---|
| Q922Q2 | Q922Q2 | Serine/thre | Riok1 | sp\|Q922Q | 7 | 7 | 15 |
| Q9DB42 | Q9DB42 | Zinc finger | Znf593 | sp\|Q9DB4 | 5 | 5 | 46,3 |
| Q7TMW6 | Q7TMW6 | Cytosolic F | Narfl | sp\|Q7TMV | 6 | 6 | 20,8 |
| P53996 | P53996 | Cellular nu | Cnbp | sp\|P53996 | 5 | 5 | 35,4 |
| O35372 | O35372 | Protein cor | Cnih1 | sp\|O3537 | 1 | 1 | 17,4 |
| Q3TZM9 | Q3TZM9 | GDP-Man:I | Alg11 | sp\|Q3TZM | 7 | 7 | 19,9 |
| Q922Q1 | Q922Q1 | Mitochond | 02-mar | sp\|Q922Q | 12 | 12 | 33,4 |
| Q8K2V1 | Q8K2V1 | Serine/thre | Ppp4r1 | sp\|Q8K2V | 27 | 27 | 43,5 |
| Q8C4Q6 | Q8C4Q6 | Axin intera | Aida | sp\|Q8C4Q | 10 | 10 | 44,3 |
| Q8CI78 | Q8CI78 | Required fo | Rmnd1 | sp\|Q8CI78 | 4 | 4 | 10,9 |
| Q8R1N4 | Q8R1N4 | NudC dom | Nudcd3 | sp\|Q8R1N | 11 | 11 | 38,3 |
| Q8BRH4 | Q8BRH4 | Histone-lys | Kmt2c | sp\|Q8BRH | 3 | 3 | 1,2 |
| Q3UA37 | Q3UA37 | Glutamine- | Qrich1 | sp\|Q3UA3 | 11 | 11 | 17,5 |
| Q5EBH1 | Q5EBH1 | Ras associa | Rassf5 | sp\|Q5EBH | 5 | 5 | 16,7 |
| Q7TNE3 | Q7TNE3 | Sperm-ass | Spag7 | sp\|Q7TNE | 6 | 6 | 35,7 |
| Q9EQM6 | Q9EQM6 | Microproc | Dgcr8 | sp\|Q9EQM | 14 | 14 | 20,6 |
| Q8BH58 | Q8BH58 | TIP41-like | Tiprl | sp\|Q8BH5 | 11 | 11 | 57,6 |
| Q9Z280 | Q9Z280 | Phospholip | Pld1 | sp\|Q9Z280 | 11 | 11 | 15,5 |
| Q91YR1 | Q91YR1 | Twinfilin-1 | Twf1 | sp\|Q91YR | 18 | 16 | 57,4 |
| Q9EPL2 | Q9EPL2 | Calsynteni | Clstn1 | sp\|Q9EPL2 | 8 | 8 | 11,3 |
| Q80UU9 | Q80UU9 | Membrane | Pgrmc2 | sp\|Q80UU | 14 | 13 | 62,7 |
| Q9JJG0 | Q9JJG0 | Transformi | Tacc2 | sp\|Q9JJG0 | 13 | 12 | 17,8 |
| Q8BLK9 | Q8BLK9 | Ribosomal | Rps6kc1 | sp\|Q8BLK | 5 | 5 | 7,1 |
| Q4VA36 | Q4VA36 | Coiled-coil | Ccdc84 | sp\|Q4VA3 | 2 | 2 | 9,3 |
| Q99JF5 | Q99JF5 | Diphospho | Mvd | sp\|Q99JF5 | 4 | 4 | 16,2 |
| Q9QYC7 | Q9QYC7 | Vitamin K- | Ggcx | sp\|Q9QYC | 5 | 5 | 8,9 |
| Q8R2R6 | Q8R2R6 | Mitochond | Mtg1 | sp\|Q8R2R | 8 | 8 | 27,9 |
| Q61458 | Q61458 | Cyclin-H | Ccnh | sp\|Q6145 | 10 | 10 | 33,4 |
| Q8BSP2 | Q8BSP2 | Condensin | Ncaph2 | sp\|Q8BSP | 12 | 12 | 29,8 |
| Q62084 | Q62084 | Protein ph | Ppp1r14b | sp\|Q6208 | 7 | 7 | 58,5 |
| P25785 | P25785 | Metallopro | Timp2 | sp\|P25785 | 4 | 4 | 23,2 |
| E9Q414 | E9Q414 | Apolipopro | Apob | sp\|E9Q41 | 7 | 6 | 2,1 |
| Q9D2G5 | Q9D2G5 | Synaptojan | Synj2 | sp\|Q9D2G | 14 | 14 | 14,3 |
| Q9QZD9 | Q9QZD9 | Eukaryotic | Eif3i | sp\|Q9QZD | 20 | 20 | 80,3 |
| P08207 | P08207 | Protein S10 | S100a10 | sp\|P08207 | 5 | 5 | 72,2 |
| Q8VE19 | Q8VE19 | WD repeat | Mios | sp\|Q8VE1 | 10 | 10 | 16,5 |
| Q3UUI3 | Q3UUI3 | Acyl-coenz | Them4 | sp\|Q3UUI | 3 | 3 | 14,3 |
| O35615 | O35615 | Zinc finger | Zfpm1 | sp\|O3561 | 3 | 3 | 5,4 |
| O88520 | O88520 | Leucine-ric | Shoc2 | sp\|O8852 | 11 | 11 | 30,6 |
| Q9EST3 | Q9EST3 | Eukaryotic | Eif4enif1 | sp\|Q9EST3 | 15 | 15 | 27,7 |
| Q3UCQ1 | Q3UCQ1 | Forkhead b | Foxk2 | sp\|Q3UCQ | 11 | 10 | 25 |
| Q9CPX9 | Q9CPX9 | Anaphase- | Anapc11 | sp\|Q9CPX | 1 | 1 | 17,9 |
| Q791N7 | Q791N7 | DNA-direct | Znrd1 | sp\|Q791N | 3 | 3 | 31,7 |
| Q3U829 | Q3U829 | AP-5 comp | Ap5z1 | sp\|Q3U82 | 6 | 6 | 10 |
| Q9CQ88 | Q9CQ88 | Tetraspani | Tspan31 | sp\|Q9CQ8 | 2 | 2 | 12,4 |
| Q62432 | Q62432 | Mothers ag | Smad2 | sp\|Q6243 | 7 | 3 | 15,2 |
| Q8BJW6 | Q8BJW6 | Eukaryotic | Eif2a | sp\|Q8BJW | 28 | 28 | 64,2 |
| P84078;P6 | P84078;P6120 | ADP-ribosy | Arf1;Arf3 | sp\|P84078 | 12 | 2 | 63,5 |
| Q5SWK7 | Q5SWK7 | RING finge | Rnf145 | sp\|Q5SWK | 5 | 5 | 11,2 |
| P70279 | P70279 | Surfeit locu | Surf6 | sp\|P70279 | 11 | 11 | 29,3 |

| | | | | | | | |
|---|---|---|---|---|---|---|---|
| Q9CQS9 | Q9CQS9 | HAUS augr | Haus2 | sp\|Q9CQS | 2 | 2 | 12,8 |
| P97765 | P97765 | WW doma | Wbp2 | sp\|P97765 | 5 | 5 | 16,9 |
| P48771 | P48771 | Cytochrom | Cox7a2 | sp\|P48771 | 1 | 1 | 15,7 |
| Q8N7N5 | Q8N7N5 | DDB1- and | Dcaf8 | sp\|Q8N7N | 13 | 13 | 29,6 |
| Q3UHX9 | Q3UHX9 | Putative m | D2Wsu81e | sp\|Q3UHX | 3 | 3 | 8,8 |
| Q9DB52 | Q9DB52 | Protein FA | Fam122a | sp\|Q9DB5 | 4 | 3 | 20,4 |
| Q9QYA2 | Q9QYA2 | Mitochond | Tomm40 | sp\|Q9QYA | 10 | 10 | 42,4 |
| Q9ESG9 | Q9ESG9 | Membrane | Pkmyt1 | sp\|Q9ESG | 5 | 5 | 16,9 |
| P29452 | P29452 | Caspase-1; | Casp1 | sp\|P29452 | 6 | 6 | 15,2 |
| Q5SSK3 | Q5SSK3 | Transcripti | Tefm | sp\|Q5SSK3 | 8 | 8 | 28,6 |
| Q8JZZ7;Q8 | Q8JZZ7 | Latrophilin | Lphn2 | sp\|Q8JZZ7 | 26 | 26 | 26,4 |
| Q8BR76 | Q8BR76 | Meckelin | Tmem67 | sp\|Q8BR7 | 2 | 2 | 3,5 |
| Q9DCN1 | Q9DCN1 | Peroxisom | Nudt12 | sp\|Q9DCN | 4 | 4 | 12,3 |
| Q9Z255 | Q9Z255 | Ubiquitin-c | Ube2a | sp\|Q9Z255 | 2 | 2 | 17,8 |
| Q9D9Z1 | Q9D9Z1 | Small kinet | Knstrn | sp\|Q9D9Z | 6 | 6 | 31,7 |
| P51569 | P51569 | Alpha-gala | Gla | sp\|P51569 | 5 | 5 | 15 |
| Q922H2 | Q922H2 | [Pyruvate c | Pdk3 | sp\|Q922H | 16 | 15 | 46 |
| Q91W96 | Q91W96 | Anaphase- | Anapc4 | sp\|Q91W9 | 22 | 22 | 36,2 |
| B2RX14 | B2RX14 | Terminal u | Zcchc11 | sp\|B2RX14 | 12 | 10 | 9,4 |
| Q9QZ23 | Q9QZ23 | NFU1 iron- | Nfu1 | sp\|Q9QZ2 | 6 | 6 | 29,4 |
| Q99JX1 | Q99JX1 | Transcripti | Taf11 | sp\|Q99JX1 | 2 | 2 | 17,5 |
| A2AGH6 | A2AGH6 | Mediator c | Med12 | sp\|A2AGH | 43 | 40 | 27,3 |
| P16460 | P16460 | Argininosu | Ass1 | sp\|P16460 | 4 | 4 | 12,4 |
| Q3UPH1 | Q3UPH1 | Protein PR | Prrc1 | sp\|Q3UPH | 10 | 10 | 39,7 |
| Q9CR02 | Q9CR02 | Translation | Tma16 | sp\|Q9CR0 | 9 | 9 | 44,8 |
| Q4JIM5 | Q4JIM5 | Abelson ty | Abl2 | sp\|Q4JIM5 | 8 | 8 | 10,4 |
| P97823 | P97823 | Acyl-prote | Lypla1 | sp\|P97823 | 5 | 5 | 33 |
| Q00PI9 | Q00PI9 | Heterogen | Hnrnpul2 | sp\|Q00PI9 | 27 | 27 | 38,9 |
| Q9DCH4 | Q9DCH4 | Eukaryotic | Eif3f | sp\|Q9DCH | 10 | 10 | 33,5 |
| Q3UGP9 | Q3UGP9 | Leucine-ric | Lrrc58 | sp\|Q3UGP | 12 | 12 | 51,9 |
| Q9CQP0 | Q9CQP0 | 39S riboso | Mrpl33 | sp\|Q9CQP | 4 | 4 | 52,3 |
| Q8C2B3 | Q8C2B3 | Histone de | Hdac7 | sp\|Q8C2B | 10 | 10 | 15,2 |
| Q32M08 | Q32M08 | tRNA-dihyc | Dus4l | sp\|Q32M0 | 3 | 3 | 16,4 |
| Q78PG9 | Q78PG9 | Coiled-coil | Ccdc25 | sp\|Q78PG | 5 | 5 | 28,8 |
| Q9ESX5 | Q9ESX5 | H/ACA ribo | Dkc1 | sp\|Q9ESX5 | 22 | 22 | 50,9 |
| Q9D2M8 | Q9D2M8 | Ubiquitin-c | Ube2v2 | sp\|Q9D2M | 9 | 4 | 68,3 |
| Q9JJC9 | Q9JJC9 | Homocyste | Herpud2 | sp\|Q9JJC9 | 2 | 2 | 6,2 |
| P27600 | P27600 | Guanine nu | Gna12 | sp\|P27600 | 2 | 1 | 4,7 |
| Q3U3C9 | Q3U3C9 | Genetic su | Gse1 | sp\|Q3U3C | 5 | 5 | 9,1 |
| Q8VCM8 | Q8VCM8 | Nicalin | Ncln | sp\|Q8VCM | 17 | 17 | 32 |
| Q91ZJ0 | Q91ZJ0 | Crossover j | Mus81 | sp\|Q91ZJ0 | 1 | 1 | 2,9 |
| Q80W00 | Q80W00 | Serine/thre | Ppp1r10 | sp\|Q80W0 | 18 | 18 | 29,7 |
| Q9CR67 | Q9CR67 | Transmem | Tmem33 | sp\|Q9CR6 | 10 | 10 | 38,9 |
| Q91ZR2 | Q91ZR2 | Sorting ne | Snx18 | sp\|Q91ZR | 9 | 9 | 16,3 |
| Q9CXE7 | Q9CXE7 | Transmem | Tmed5 | sp\|Q9CXE | 5 | 5 | 18,8 |
| Q3U182 | Q3U182 | CREB-regu | Crtc2 | sp\|Q3U18 | 9 | 9 | 20,8 |
| Q922H9 | Q922H9 | Zinc finger | Znf330 | sp\|Q922H | 3 | 3 | 15,5 |
| G5E8P0 | G5E8P0 | | Tubgcp6 | sp\|G5E8P0 | 8 | 8 | 8,3 |
| Q99KX1 | Q99KX1 | Myeloid le | Mlf2 | sp\|Q99KX | 4 | 4 | 26,7 |
| Q8BHY2 | Q8BHY2 | Nucleolar c | Noc4l | sp\|Q8BHY | 15 | 15 | 38,2 |

| | | | | | | | |
|---|---|---|---|---|---|---|---|
| Q9CPU9 | Q9CPU9 | Probable lo | Slc31a2 | sp\|Q9CPU | 1 | 1 | 17,5 |
| Q9JIB4 | Q9JIB4 | General tra | Gtf2h2 | sp\|Q9JIB4 | 8 | 8 | 30,1 |
| Q9ES74 | Q9ES74 | Serine/thre | Nek7 | sp\|Q9ES74 | 15 | 14 | 53,6 |
| Q64310 | Q64310 | Surfeit locu | Surf4 | sp\|Q64310 | 5 | 5 | 24,5 |
| Q6P9N1;Q | Q6P9N1 | Hyccin | Fam126a | sp\|Q6P9N | 7 | 7 | 21,1 |
| Q8CIA9 | Q8CIA9 | Hippocamp | Hiatl1 | sp\|Q8CIA9 | 1 | 1 | 3,6 |
| Q8VE94 | Q8VE94 | Protein FA | Fam110c | sp\|Q8VE94 | 4 | 4 | 10,7 |
| O35375 | O35375 | Neuropilin | Nrp2 | sp\|O35375 | 25 | 25 | 34,9 |
| P52800 | P52800 | Ephrin-B2 | Efnb2 | sp\|P52800 | 8 | 7 | 36,3 |
| Q68FE2 | Q68FE2 | Autophagy | Atg9a | sp\|Q68FE2 | 5 | 5 | 12,7 |
| Q8VHR5 | Q8VHR5 | Transcripti | Gatad2b | sp\|Q8VHR | 20 | 19 | 47,6 |
| Q9DBG9 | Q9DBG9 | Tax1-bindi | Tax1bp3 | sp\|Q9DBG | 3 | 3 | 39,5 |
| Q60648 | Q60648 | Gangliosid | Gm2a | sp\|Q60648 | 5 | 5 | 29,5 |
| Q9D662 | Q9D662 | Protein tra | Sec23b | sp\|Q9D662 | 29 | 26 | 57,8 |
| Q9CQK7 | Q9CQK7 | RWD doma | Rwdd1 | sp\|Q9CQK | 6 | 6 | 19,3 |
| Q9ESD6 | Q9ESD6 | CKLF-like N | Cmtm7 | sp\|Q9ESD6 | 2 | 2 | 19,2 |
| Q62203 | Q62203 | Splicing fac | Sf3a2 | sp\|Q62203 | 8 | 8 | 21,7 |
| P61222 | P61222 | ATP-bindin | Abce1 | sp\|P61222 | 28 | 28 | 51,4 |
| Q6ZWV7 | Q6ZWV7 | 60S riboso | Rpl35 | sp\|Q6ZWV | 3 | 3 | 18,7 |
| Q8K2T8 | Q8K2T8 | RNA polym | Paf1 | sp\|Q8K2T8 | 19 | 19 | 44,1 |
| Q9QXE7 | Q9QXE7 | F-box-like/ | Tbl1x | sp\|Q9QXE | 17 | 11 | 51,8 |
| Q9Z110 | Q9Z110 | Delta-1-py | Aldh18a1 | sp\|Q9Z110 | 41 | 41 | 62,1 |
| P35601 | P35601 | Replication | Rfc1 | sp\|P35601 | 32 | 32 | 37 |
| Q8R1Z9 | Q8R1Z9 | RING finge | Rnf121 | sp\|Q8R1Z9 | 3 | 3 | 14,1 |
| Q9CZR3 | Q9CZR3 | Mitochond | Tomm40l | sp\|Q9CZR3 | 4 | 4 | 16,9 |
| P18654 | P18654 | Ribosomal | Rps6ka3 | sp\|P18654 | 23 | 15 | 38,2 |
| Q9D958 | Q9D958 | Signal pept | Spcs1 | sp\|Q9D958 | 2 | 2 | 8,1 |
| Q8K039 | Q8K039 | Uncharacterized prote | | sp\|Q8K039 | 2 | 2 | 10,3 |
| Q99K48 | Q99K48 | Non-POU d | Nono | sp\|Q99K48 | 25 | 24 | 51,6 |
| Q8VE80 | Q8VE80 | THO comp | Thoc3 | sp\|Q8VE80 | 6 | 6 | 31,9 |
| Q8CG64 | Q8CG64 | Fukutin-rel | Fkrp | sp\|Q8CG64 | 2 | 2 | 5,1 |
| Q91XL3 | Q91XL3 | UDP-glucu | Uxs1 | sp\|Q91XL3 | 10 | 10 | 36 |
| Q9D666 | Q9D666 | SUN doma | Sun1 | sp\|Q9D666 | 29 | 29 | 42,5 |
| Q8C2E7 | Q8C2E7 | WASH com | Kiaa0196 | sp\|Q8C2E7 | 33 | 33 | 32,4 |
| Q60649 | Q60649 | Caseinolyti | Clpb | sp\|Q60649 | 14 | 14 | 31,3 |
| Q8BHC7 | Q8BHC7 | Rhomboid- | Rhbdd1 | sp\|Q8BHC | 1 | 1 | 5,4 |
| O88700 | O88700 | Bloom syn | Blm | sp\|O88700 | 13 | 13 | 15,1 |
| P11416;P1 | P11416 | Retinoic ac | Rara | sp\|P11416 | 4 | 3 | 15,8 |
| Q78IK4 | Q78IK4 | MICOS con | Apool | sp\|Q78IK4 | 11 | 11 | 46 |
| P56380 | P56380 | Bis(5-nucle | Nudt2 | sp\|P56380 | 8 | 8 | 74,8 |
| Q00262 | Q00262 | Syntaxin-2 | Stx2 | sp\|Q00262 | 6 | 6 | 26,6 |
| Q80TN4 | Q80TN4 | DnaJ homo | Dnajc16 | sp\|Q80TN4 | 9 | 9 | 17,5 |
| Q6P6I6 | Q6P6I6 | DNA-direct | Polr2m | sp\|Q6P6I6 | 5 | 5 | 28,4 |
| Q9D404 | Q9D404 | 3-oxoacyl-[ | Oxsm | sp\|Q9D404 | 7 | 7 | 28,1 |
| Q8BSK8 | Q8BSK8 | Ribosomal | Rps6kb1 | sp\|Q8BSK8 | 9 | 8 | 26,3 |
| Q8R4C2 | Q8R4C2 | RUN and F | Rufy2 | sp\|Q8R4C2 | 9 | 8 | 15,7 |
| P41241;P4 | P41241 | Tyrosine-p | Csk | sp\|P41241 | 18 | 18 | 49,8 |
| Q8CDN6 | Q8CDN6 | Thioredoxi | Txnl1 | sp\|Q8CDN | 19 | 19 | 67,1 |
| Q8CGF5 | Q8CGF5 | Transmem | Tmem56 | sp\|Q8CGF5 | 2 | 2 | 7,2 |
| Q3UCV8 | Q3UCV8 | Ubiquitin t | Otulin | sp\|Q3UCV | 3 | 3 | 9,4 |

| | | | | | | | |
|---|---|---|---|---|---|---|---|
| Q811S7 | Q811S7 | Upstream- | Ubp1 | sp\|Q811S7 | 11 | 10 | 27,6 |
| Q9CWJ3 | Q9CWJ3 | KATNB1-lik | Katnbl1 | sp\|Q9CWJ | 3 | 3 | 17,4 |
| Q9CZT6 | Q9CZT6 | Protein CM | Cmss1 | sp\|Q9CZT6 | 6 | 6 | 26,4 |
| Q8QZS3 | Q8QZS3 | Folliculin | Flcn | sp\|Q8QZS3 | 7 | 7 | 16,9 |
| P05480;Q6 | P05480 | Neuronal p | Src | sp\|P05480 | 18 | 12 | 40,5 |
| Q6PIX5 | Q6PIX5 | Inactive rh | Rhbdf1 | sp\|Q6PIX5 | 7 | 7 | 13,2 |
| Q3UG98 | Q3UG98 | N-acetyltra | Nat9 | sp\|Q3UG9 | 3 | 3 | 11,6 |
| Q9CTN4 | Q9CTN4 | Rho-relate | Rhobtb3 | sp\|Q9CTN4 | 8 | 8 | 20,9 |
| Q9JJI8 | Q9JJI8 | 60S riboso | Rpl38 | sp\|Q9JJI8\| | 6 | 6 | 50 |
| Q8CBR6 | Q8CBR6 | Tsukushin | Tsku | sp\|Q8CBR6 | 4 | 4 | 20,9 |
| Q9Z175 | Q9Z175 | Lysyl oxida | Loxl3 | sp\|Q9Z175 | 7 | 7 | 13,7 |
| Q8BV57 | Q8BV57 | Soluble sca | Ssc5d | sp\|Q8BV57 | 3 | 3 | 4,2 |
| B1AZA5 | B1AZA5 | Transmem | Tmem245 | sp\|B1AZA5 | 5 | 5 | 5,4 |
| Q6DI86 | Q6DI86 | FAST kinas | Fastkd1 | sp\|Q6DI86 | 2 | 2 | 3 |
| Q8BHC9 | Q8BHC9 | Alpha-(1,3) | Fut11 | sp\|Q8BHC | 4 | 4 | 14,3 |
| Q9CZ96 | Q9CZ96 | Zinc finger | Zcrb1 | sp\|Q9CZ96 | 3 | 3 | 23 |
| Q9CVB6 | Q9CVB6 | Actin-relat | Arpc2 | sp\|Q9CVB6 | 24 | 24 | 77 |
| Q8BYM8 | Q8BYM8 | Probable c | Cars2 | sp\|Q8BYM | 12 | 12 | 25 |
| Q8R3Q0 | Q8R3Q0 | Store-oper | Saraf | sp\|Q8R3Q | 5 | 5 | 20,4 |
| O70194 | O70194 | Eukaryotic | Eif3d | sp\|O70194 | 24 | 24 | 55,1 |
| Q8K3G5 | Q8K3G5 | Inactive se | Vrk3 | sp\|Q8K3G | 6 | 6 | 20,5 |
| P41731 | P41731 | CD63 antig | Cd63 | sp\|P41731 | 4 | 4 | 23,5 |
| Q9CWU4 | Q9CWU4 | UPF0690 protein C1or | | sp\|Q9CWU | 6 | 6 | 35 |
| Q99MP8 | Q99MP8 | BRCA1-ass | Brap | sp\|Q99MP | 6 | 6 | 13,5 |
| Q80VQ1 | Q80VQ1 | Leucine-ric | Lrrc1 | sp\|Q80VQ | 15 | 12 | 42,6 |
| Q9QXV3 | Q9QXV3 | Inhibitor o | Ing1 | sp\|Q9QXV | 3 | 3 | 16,5 |
| P07742 | P07742 | Ribonucleo | Rrm1 | sp\|P07742 | 36 | 36 | 65,2 |
| Q99LR1 | Q99LR1 | Monoacylg | Abhd12 | sp\|Q99LR1 | 10 | 10 | 31,2 |
| Q9D8Z1;RE | Q9D8Z1 | Activating | Ascc1 | sp\|Q9D8Z1 | 7 | 7 | 19,7 |
| Q7TMQ7 | Q7TMQ7 | WD repeat | Wdr91 | sp\|Q7TMQ | 9 | 9 | 13,9 |
| Q62245 | Q62245 | Son of seve | Sos1 | sp\|Q62245 | 18 | 14 | 17,2 |
| Q9D0M5 | Q9D0M5 | Dynein ligh | Dynll2 | sp\|Q9D0M | 5 | 3 | 62,9 |
| Q99M54 | Q99M54 | Cell divisio | Cdca3 | sp\|Q99M5 | 1 | 1 | 6,8 |
| Q8BWZ3 | Q8BWZ3 | N-alpha-ac | Naa25 | sp\|Q8BWZ | 30 | 30 | 39,3 |
| Q8C006 | Q8C006 | Tripartite r | Trim35 | sp\|Q8C006 | 3 | 3 | 7,6 |
| P24452 | P24452 | Macrophag | Capg | sp\|P24452 | 11 | 11 | 42 |
| G3X9G7 | G3X9G7 | Zinc finger | Zfp809 | sp\|G3X9G7 | 4 | 4 | 15,4 |
| P63101 | P63101 | 14-3-3 pro | Ywhaz | sp\|P63101 | 20 | 15 | 73,1 |
| E9Q9R9 | E9Q9R9 | | Dlg5 | sp\|E9Q9R9 | 8 | 8 | 5,7 |
| Q9CX53 | Q9CX53 | Gem-assoc | Gemin6 | sp\|Q9CX53 | 3 | 3 | 21,1 |
| A2AUM9 | A2AUM9 | Centrosom | Cep152 | sp\|A2AUM | 8 | 8 | 6,5 |
| Q3THW5;P | Q3THW5;P0C0 | Histone H2 | H2afv;H2a | sp\|Q3THW | 6 | 4 | 53,9 |
| Q3TCJ1 | Q3TCJ1 | BRISC com | Fam175b | sp\|Q3TCJ1 | 5 | 5 | 21,9 |
| Q9CY94 | Q9CY94 | DNA replic | Gins3 | sp\|Q9CY94 | 9 | 9 | 56 |
| Q62083 | Q62083 | PRKCA-bin | Pick1 | sp\|Q62083 | 3 | 3 | 9,4 |
| Q9JJ48 | Q9JJ48 | Zinc finger | Zc3h8 | sp\|Q9JJ48 | 3 | 3 | 15,1 |
| Q62280 | Q62280 | Protein SS | Ss18 | sp\|Q62280 | 3 | 3 | 9,6 |
| Q78J03 | Q78J03 | Methionin | Msrb2 | sp\|Q78J03 | 3 | 3 | 18,9 |
| Q99J62 | Q99J62 | Replicatior | Rfc4 | sp\|Q99J62 | 17 | 17 | 48,6 |
| P32233 | P32233 | Developme | Drg1 | sp\|P32233 | 19 | 19 | 54,5 |

| | | | | | | | |
|---|---|---|---|---|---|---|---|
| Q9DBU3 | Q9DBU3 | Serine/thre | Riok3 | sp\|Q9DBU | 8 | 8 | 22,7 |
| Q9DCR2 | Q9DCR2 | AP-3 comp | Ap3s1 | sp\|Q9DCR | 6 | 6 | 38,3 |
| E2JF22 | E2JF22 | Piezo-type | Piezo1 | sp\|E2JF22 | 27 | 27 | 17,7 |
| Q9CWX9 | Q9CWX9 | Probable A | Ddx47 | sp\|Q9CWX | 13 | 13 | 35,2 |
| Q9R0I7 | Q9R0I7 | YLP motif- | Ylpm1 | sp\|Q9R0I7 | 33 | 33 | 32,9 |
| Q6PHN9 | Q6PHN9 | Ras-related | Rab35 | sp\|Q6PHN | 6 | 6 | 41,8 |
| Q8VIG1 | Q8VIG1 | RE1-silenci | Rest | sp\|Q8VIG1 | 6 | 6 | 8,2 |
| P70124 | P70124 | Serpin B5 | Serpinb5 | sp\|P70124 | 4 | 4 | 15,7 |
| Q9D2C7 | Q9D2C7 | Bax inhibit | Tmbim6 | sp\|Q9D2C7 | 2 | 2 | 8,9 |
| Q9WTX5 | Q9WTX5 | S-phase kir | Skp1 | sp\|Q9WTX | 13 | 13 | 76,1 |
| Q8BGD6 | Q8BGD6 | Sodium-co | Slc38a9 | sp\|Q8BGD | 1 | 1 | 4,6 |
| Q99LH2 | Q99LH2 | Phosphatid | Ptdss1 | sp\|Q99LH2 | 4 | 4 | 13,5 |
| Q91WM2 | Q91WM2 | Cat eye syr | Cecr5 | sp\|Q91WN | 6 | 6 | 19,8 |
| P54731 | P54731 | FAS-associ | Faf1 | sp\|P54731 | 18 | 18 | 33,4 |
| Q8BWR2 | Q8BWR2 | PITH doma | Pithd1 | sp\|Q8BWF | 5 | 5 | 39,3 |
| Q9CY45 | Q9CY45 | Protein-lys | N6amt2 | sp\|Q9CY45 | 5 | 5 | 26,6 |
| Q8C8M1 | Q8C8M1 | Protein FA | Fam60a | sp\|Q8C8M | 2 | 2 | 14,9 |
| Q5I012 | Q5I012 | Putative so | Slc38a10 | sp\|Q5I012 | 22 | 22 | 31,1 |
| P70296;Q8 | P70296 | Phosphatid | Pebp1 | sp\|P70296 | 13 | 13 | 88,8 |
| Q9WUA6 | Q9WUA6 | RAC-gamm | Akt3 | sp\|Q9WUA | 9 | 4 | 21,3 |
| Q921Q3 | Q921Q3 | Chitobiosy | Alg1 | sp\|Q921Q | 6 | 6 | 18,7 |
| Q9DB85 | Q9DB85 | Ribosomal | Rrp8 | sp\|Q9DB8 | 13 | 13 | 40,9 |
| P70257 | P70257 | Nuclear fac | Nfix | sp\|P70257 | 14 | 11 | 42,8 |
| O35129 | O35129 | Prohibitin- | Phb2 | sp\|O35129 | 22 | 22 | 72,9 |
| E9Q8D0 | E9Q8D0 | | Dnajc21 | sp\|E9Q8D0 | 7 | 7 | 17,5 |
| Q99NH2 | Q99NH2 | Partitionin | Pard3 | sp\|Q99NH | 21 | 21 | 22,7 |
| Q8BFQ6 | Q8BFQ6 | Disrupted i | Dirc2 | sp\|Q8BFQ | 1 | 1 | 3,1 |
| Q80T14 | Q80T14 | Extracellula | Fras1 | sp\|Q80T14 | 1 | 1 | 0,2 |
| Q8BMB0 | Q8BMB0 | Protein EM | Emsy | sp\|Q8BME | 13 | 13 | 15,1 |
| Q8BUK6 | Q8BUK6 | Protein Ho | Hook3 | sp\|Q8BUK | 17 | 17 | 29 |
| P59235 | P59235 | Nucleopor | Nup43 | sp\|P59235 | 11 | 11 | 39,7 |
| O70309 | O70309 | Integrin be | Itgb5 | sp\|O70309 | 12 | 12 | 17,7 |
| Q9DCL8 | Q9DCL8 | Protein ph | Ppp1r2 | sp\|Q9DCL8 | 6 | 6 | 28,2 |
| Q9CQ36 | Q9CQ36 | DNA polym | Pole4 | sp\|Q9CQ3 | 4 | 4 | 55,9 |
| Q5NCF2 | Q5NCF2 | Trafficking | Trappc1 | sp\|Q5NCF2 | 6 | 6 | 46,9 |
| Q5SWD9 | Q5SWD9 | Pre-rRNA-p | Tsr1 | sp\|Q5SWD | 24 | 24 | 44,6 |
| Q6PD26 | Q6PD26 | GPI transa | Pigs | sp\|Q6PD26 | 20 | 20 | 56,6 |
| P52927 | P52927 | High mobil | Hmga2 | sp\|P52927 | 9 | 9 | 51,9 |
| P47753 | P47753 | F-actin-cap | Capza1 | sp\|P47753 | 12 | 10 | 65 |
| A2ABV5 | A2ABV5 | Mediator c | Med14 | sp\|A2ABV5 | 20 | 20 | 21 |
| Q9D187 | Q9D187 | Mitotic spi | Fam96b | sp\|Q9D187 | 6 | 6 | 67,5 |
| Q8BKE6 | Q8BKE6 | Cytochrom | Cyp20a1 | sp\|Q8BKE6 | 18 | 18 | 51,9 |
| Q99LI7 | Q99LI7 | Cleavage s | Cstf3 | sp\|Q99LI7 | 26 | 26 | 43,7 |
| Q80XU3 | Q80XU3 | Nuclear ub | Nucks1 | sp\|Q80XU | 5 | 5 | 21,8 |
| Q8K4D3 | Q8K4D3 | Proton-cou | Slc36a1 | sp\|Q8K4D | 2 | 2 | 6,1 |
| Q8R3Y5 | Q8R3Y5 | Uncharacterized prote | | sp\|Q8R3Y5 | 5 | 5 | 22,5 |
| Q80Y17 | Q80Y17 | Lethal(2) g | Llgl1 | sp\|Q80Y17 | 26 | 26 | 39,9 |
| Q8C092 | Q8C092 | Transcripti | Taf5 | sp\|Q8C092 | 13 | 13 | 28 |
| Q60996 | Q60996 | Serine/thre | Ppp2r5c | sp\|Q60996 | 15 | 13 | 39,5 |
| Q8VEH6 | Q8VEH6 | COBW don | Cbwd1 | sp\|Q8VEH | 11 | 11 | 38,9 |

| | | | | | | | |
|---|---|---|---|---|---|---|---|
| P62311 | P62311 | U6 snRNA- | Lsm3 | sp\|P62311 | 3 | 3 | 40,2 |
| P50096 | P50096 | Inosine-5-r | Impdh1 | sp\|P50096 | 11 | 10 | 27,8 |
| Q3TLS3 | Q3TLS3 | GDP-D-glu | Gdpgp1 | sp\|Q3TLS3 | 6 | 6 | 19,7 |
| Q9CXT7 | Q9CXT7 | Transmem | Tmem192 | sp\|Q9CXT7 | 5 | 5 | 25,2 |
| Q921F4 | Q921F4 | Heterogen | Hnrnpll | sp\|Q921F4 | 24 | 24 | 56,7 |
| Q91WK0 | Q91WK0 | Leucine-ric | Lrrfip2 | sp\|Q91WK | 6 | 5 | 17,3 |
| Q5SSZ5 | Q5SSZ5 | Tensin-3 | Tns3 | sp\|Q5SSZ5 | 34 | 32 | 39,4 |
| Q9R207 | Q9R207 | Nibrin | Nbn | sp\|Q9R207 | 18 | 18 | 30,1 |
| Q99PI4 | Q99PI4 | Phosphatic | Lpin3 | sp\|Q99PI4 | 5 | 5 | 9,7 |
| Q80XC6 | Q80XC6 | Protein NR | Nrde2 | sp\|Q80XC6 | 15 | 15 | 19 |
| F8VPZ9 | F8VPZ9 | | Gltscr1 | sp\|F8VPZ9 | 5 | 5 | 4,6 |
| Q99J21 | Q99J21 | Mucolipin- | Mcoln1 | sp\|Q99J21 | 3 | 3 | 10 |
| P03958 | P03958 | Adenosine | Ada | sp\|P03958 | 12 | 12 | 42,6 |
| Q0VBL1 | Q0VBL1 | Tigger tran | Tigd2 | sp\|Q0VBL1 | 4 | 4 | 12,4 |
| Q8BJ48 | Q8BJ48 | N-acetylglu | Nagpa | sp\|Q8BJ48 | 2 | 2 | 6,2 |
| Q0VEJ0 | Q0VEJ0 | Centrosom | Cep76 | sp\|Q0VEJ0 | 2 | 2 | 3,8 |
| Q9CPY3 | Q9CPY3 | Sororin | Cdca5 | sp\|Q9CPY3 | 7 | 7 | 43,2 |
| Q8VDG5 | Q8VDG5 | Phosphopa | Ppcs | sp\|Q8VDG | 6 | 6 | 18,3 |
| Q9CWX2 | Q9CWX2 | Complex I | Ndufaf1 | sp\|Q9CWX | 10 | 10 | 37,8 |
| Q60692 | Q60692 | Proteasom | Psmb6 | sp\|Q60692 | 11 | 11 | 60,9 |
| Q9CWP8 | Q9CWP8 | DNA polym | Pold4 | sp\|Q9CWP | 4 | 4 | 63,6 |
| Q99LL5 | Q99LL5 | Periodic tr | Pwp1 | sp\|Q99LL5 | 10 | 10 | 34,9 |
| Q99J56 | Q99J56 | Derlin-1 | Derl1 | sp\|Q99J56 | 3 | 3 | 13,1 |
| B2RX12 | B2RX12 | Canalicular | Abcc3 | sp\|B2RX12 | 2 | 1 | 1,4 |
| Q3U3N0 | Q3U3N0 | Mesoderm | Mier2 | sp\|Q3U3N | 2 | 2 | 5,7 |
| P48193 | P48193 | Protein 4.1 | Epb41 | sp\|P48193 | 16 | 13 | 25,9 |
| Q9ER73 | Q9ER73 | Elongator c | Elp4 | sp\|Q9ER73 | 14 | 14 | 41,5 |
| P50247 | P50247 | Adenosylh | Ahcy | sp\|P50247 | 27 | 27 | 69,4 |
| Q99JT6 | Q99JT6 | Calfacilitin | Tlcd1 | sp\|Q99JT6 | 3 | 3 | 12,6 |
| P46718 | P46718 | Programm | Pdcd2 | sp\|P46718 | 8 | 8 | 29,7 |
| O35607 | O35607 | Bone morp | Bmpr2 | sp\|O35607 | 4 | 4 | 5,5 |
| P70428 | P70428 | Exostosin-2 | Ext2 | sp\|P70428 | 10 | 10 | 19,6 |
| Q9CPS7 | Q9CPS7 | RNA-bindir | Pno1 | sp\|Q9CPS7 | 8 | 8 | 39,5 |
| Q6NS45 | Q6NS45 | Coiled-coil | Ccdc66 | sp\|Q6NS45 | 2 | 2 | 4 |
| Q8K2D3 | Q8K2D3 | Enhancer c | Edc3 | sp\|Q8K2D3 | 13 | 13 | 33,9 |
| Q9D2N9 | Q9D2N9 | Vacuolar p | Vps33a | sp\|Q9D2N | 15 | 15 | 27,6 |
| Q921M3 | Q921M3 | Splicing fac | Sf3b3 | sp\|Q921M | 45 | 45 | 48,5 |
| Q8K021 | Q8K021 | Secretory c | Scamp1 | sp\|Q8K021 | 6 | 6 | 34,3 |
| Q7TQK4 | Q7TQK4 | Exosome c | Exosc3 | sp\|Q7TQK | 15 | 15 | 76,6 |
| P10711;Q9 | P10711 | Transcripti | Tcea1 | sp\|P10711 | 17 | 17 | 51,5 |
| Q6DFV1 | Q6DFV1 | Condensin | Ncapg2 | sp\|Q6DFV1 | 34 | 34 | 39,3 |
| Q9D083 | Q9D083 | Kinetochor | Spc24 | sp\|Q9D083 | 8 | 8 | 61,2 |
| P83093 | P83093 | Stromal int | Stim2 | sp\|P83093 | 17 | 16 | 34,3 |
| Q9D1G1 | Q9D1G1 | Ras-related | Rab1b | sp\|Q9D1G | 14 | 8 | 74,1 |
| P50228 | P50228 | C-X-C moti | Cxcl5 | sp\|P50228 | 2 | 2 | 25,8 |
| Q8BHN0 | Q8BHN0 | Protein ph | Ppm1l | sp\|Q8BHN | 6 | 6 | 29,7 |
| Q9CWV1 | Q9CWV1 | DNA helica | Mcm8 | sp\|Q9CWV | 5 | 4 | 8,8 |
| A2ADY9;Q9 | A2ADY9 | Protein DD | Ddi2 | sp\|A2ADY9 | 13 | 13 | 46,1 |
| O35218 | O35218 | Cleavage a | Cpsf2 | sp\|O35218 | 35 | 35 | 56,9 |
| Q6A009 | Q6A009 | E3 ubiquiti | Ltn1 | sp\|Q6A009 | 28 | 28 | 21,3 |

| | | | | | | | |
|---|---|---|---|---|---|---|---|
| O35657 | O35657 | Sialidase-1 | Neu1 | sp\|O35657 | 3 | 3 | 13,7 |
| Q99KP6 | Q99KP6 | Pre-mRNA- | Prpf19 | sp\|Q99KP6 | 19 | 19 | 58,7 |
| Q9D4H9 | Q9D4H9 | PHD finger | Phf14 | sp\|Q9D4H | 14 | 14 | 25,1 |
| Q924S7 | Q924S7 | Sprouty-re | Spred2 | sp\|Q924S7 | 6 | 5 | 22,4 |
| Q9CYG7 | Q9CYG7 | Mitochond | Tomm34 | sp\|Q9CYG7 | 12 | 12 | 45,3 |
| Q5SU73 | Q5SU73 | Coilin | Coil | sp\|Q5SU73 | 9 | 9 | 23,2 |
| Q8C407 | Q8C407 | Protein YIP | Yipf4 | sp\|Q8C407 | 3 | 3 | 13,4 |
| P16054 | P16054 | Protein kin | Prkce | sp\|P16054 | 3 | 3 | 5,6 |
| P97376 | P97376 | Protein FR | Frg1 | sp\|P97376 | 10 | 10 | 38,4 |
| O88708;C | O88708 | Origin reco | Orc4 | sp\|O88708 | 10 | 10 | 32,3 |
| E9Q286 | E9Q286 | Little elong | Ice1 | sp\|E9Q286 | 12 | 11 | 7,8 |
| P50543 | P50543 | Protein S1 | S100a11 | sp\|P50543 | 7 | 7 | 51 |
| Q9CPW2 | Q9CPW2 | Adrenodox | Fdx1l | sp\|Q9CPW | 4 | 4 | 31 |
| Q9WTS2 | Q9WTS2 | Alpha-(1,6) | Fut8 | sp\|Q9WTS | 14 | 14 | 34,8 |
| CON__P35 | CON__P35908v2;CON__P35908 | | | ; | 21 | 12 | 43,3 |
| Q9R0H0 | Q9R0H0 | Peroxisom | Acox1 | sp\|Q9R0H | 22 | 22 | 51,1 |
| Q3TQI7 | Q3TQI7 | Uncharacterized prote | | sp\|Q3TQI7 | 6 | 6 | 21,1 |
| Q6VNB8 | Q6VNB8 | WD repeat | Wdfy3 | sp\|Q6VNB | 17 | 17 | 7,8 |
| Q60972 | Q60972 | Histone-bi | Rbbp4 | sp\|Q60972 | 20 | 13 | 69,9 |
| Q9D3P8 | Q9D3P8 | Plasminoge | Plgrkt | sp\|Q9D3P8 | 3 | 3 | 21,8 |
| Q3TPJ7 | Q3TPJ7 | Midnolin | Midn | sp\|Q3TPJ7 | 2 | 2 | 6,2 |
| Q8BS90 | Q8BS90 | Protein au | Bora | sp\|Q8BS90 | 4 | 4 | 15,8 |
| Q9EQH2 | Q9EQH2 | Endoplasm | Erap1 | sp\|Q9EQH | 27 | 27 | 34,2 |
| P97493 | P97493 | Thioredoxi | Txn2 | sp\|P97493 | 6 | 6 | 41,6 |
| Q8C1Y8 | Q8C1Y8 | Vacuolar fu | Ccz1 | sp\|Q8C1Y8 | 13 | 13 | 32,9 |
| Q66JQ7 | Q66JQ7 | Protein CA | Casc5 | sp\|Q66JQ7 | 14 | 14 | 12,5 |
| Q9D3E6 | Q9D3E6 | Cohesin su | Stag1 | sp\|Q9D3E6 | 30 | 26 | 28,9 |
| Q9D6J3 | Q9D6J3 | Coiled-coil | Ccdc94 | sp\|Q9D6J3 | 10 | 10 | 40,8 |
| Q3UMY5;Q | Q3UMY5 | Echinoderm | Eml4 | sp\|Q3UMY | 19 | 18 | 23,2 |
| Q3TP92 | Q3TP92 | CTD nuclea | Ctdnep1 | sp\|Q3TP92 | 4 | 4 | 16,4 |
| O08915 | O08915 | AH recepto | Aip | sp\|O08915 | 13 | 13 | 50,9 |
| Q64FW2 | Q64FW2 | All-trans-re | Retsat | sp\|Q64FW | 7 | 7 | 14,9 |
| P81069 | P81069 | GA-binding | Gabpb2 | sp\|P81069 | 8 | 7 | 22 |
| Q8K4Q6 | Q8K4Q6 | Endonucle | Neil1 | sp\|Q8K4Q | 1 | 1 | 5,7 |
| Q8C8T8 | Q8C8T8 | Pre-rRNA-p | Tsr2 | sp\|Q8C8T8 | 3 | 3 | 17,8 |
| Q8K1L5 | Q8K1L5 | Protein ph | Ppp1r11 | sp\|Q8K1L5 | 2 | 2 | 12,2 |
| Q9EQ28 | Q9EQ28 | DNA polym | Pold3 | sp\|Q9EQ28 | 9 | 9 | 27,5 |
| Q01147 | Q01147 | Cyclic AMP | Creb1 | sp\|Q01147 | 6 | 4 | 25,8 |
| Q61846 | Q61846 | Maternal e | Melk | sp\|Q61846 | 14 | 14 | 29,9 |
| Q91VJ5 | Q91VJ5 | Polyglutam | Pqbp1 | sp\|Q91VJ5 | 9 | 9 | 39,9 |
| Q8C0S1 | Q8C0S1 | DIS3-like e | Dis3l | sp\|Q8C0S1 | 13 | 13 | 16,7 |
| Q9QYI5 | Q9QYI5 | DnaJ homo | Dnajb2 | sp\|Q9QYI5 | 5 | 4 | 18,5 |
| Q80UF4 | Q80UF4 | Serological | Sdccag8 | sp\|Q80UF4 | 2 | 2 | 3,9 |
| Q8K1C9 | Q8K1C9 | Leucine-ric | Lrrc41 | sp\|Q8K1C9 | 16 | 16 | 26,8 |
| P33215 | P33215 | Protein NE | Nedd1 | sp\|P33215 | 19 | 19 | 46,5 |
| Q80YR5 | Q80YR5 | Scaffold at | Safb2 | sp\|Q80YR5 | 25 | 14 | 27,5 |
| Q922W5 | Q922W5 | Pyrroline-5 | Pycr1 | sp\|Q922W | 10 | 7 | 49,2 |
| P35285 | P35285 | Ras-related | Rab22a | sp\|P35285 | 7 | 5 | 49 |
| Q99N85 | Q99N85 | 28S riboso | Mrps18a | sp\|Q99N8 | 3 | 3 | 20,9 |
| Q9R1K9;P4 | Q9R1K9 | Centrin-2 | Cetn2 | sp\|Q9R1K9 | 6 | 6 | 53,5 |

| | | | | | | | |
|---|---|---|---|---|---|---|---|
| Q7TS74 | Q7TS74 | Cytoskelet | Ckap2l | sp\|Q7TS74 | 5 | 5 | 10,3 |
| Q8VEH3 | Q8VEH3 | ADP-ribosy | Arl8a | sp\|Q8VEH | 11 | 6 | 67,2 |
| P68373 | P68373 | Tubulin alp | Tuba1c | sp\|P68373 | 27 | 3 | 65,7 |
| Q9WU28 | Q9WU28 | Prefoldin s | Pfdn5 | sp\|Q9WU2 | 8 | 8 | 73,4 |
| Q9CWV6 | Q9CWV6 | PRKR-inter | Prkrip1 | sp\|Q9CWV | 4 | 4 | 28 |
| Q03147;Q8 | Q03147 | Cyclin-dep | Cdk7 | sp\|Q03147 | 10 | 10 | 42,2 |
| O08696 | O08696 | Forkhead b | Foxm1 | sp\|O08696 | 4 | 4 | 9,5 |
| Q0VGY8 | Q0VGY8 | Protein TA | Tanc1 | sp\|Q0VGY | 27 | 27 | 20,7 |
| Q7TME2 | Q7TME2 | Sperm-ass | Spag5 | sp\|Q7TME | 16 | 16 | 17,9 |
| Q5IRJ6 | Q5IRJ6 | Zinc transp | Slc30a9 | sp\|Q5IRJ6 | 10 | 10 | 21,3 |
| O08740 | O08740 | DNA-direct | Polr2j | sp\|O08740 | 6 | 6 | 45,3 |
| P25799 | P25799 | Nuclear fac | Nfkb1 | sp\|P25799 | 27 | 27 | 38,4 |
| Q8R331 | Q8R331 | Zinc finger | Znhit1 | sp\|Q8R331 | 4 | 4 | 24 |
| Q6ZQI3 | Q6ZQI3 | Malectin | Mlec | sp\|Q6ZQI3 | 12 | 12 | 41,6 |
| Q9CRA7 | Q9CRA7 | ATP syntha | Atp5s | sp\|Q9CRA | 4 | 4 | 31 |
| O54784;Q8 | O54784 | Death-ass | Dapk3 | sp\|O54784 | 8 | 8 | 23,2 |
| P58462 | P58462 | Forkhead b | Foxp1 | sp\|P58462 | 5 | 5 | 14 |
| Q9DBU5 | Q9DBU5 | E3 ubiquiti | Rnf6 | sp\|Q9DBU | 5 | 5 | 12,1 |
| Q8BTH8 | Q8BTH8 | Casein kina | Csnk1g1 | sp\|Q8BTH | 9 | 3 | 20,7 |
| Q9CPT4 | Q9CPT4 | Myeloid-de | Mydgf | sp\|Q9CPT4 | 6 | 6 | 38,6 |
| Q9R060 | Q9R060 | Cytosolic F | Nubp1 | sp\|Q9R060 | 4 | 4 | 17,5 |
| Q3UJP5 | Q3UJP5 | Protein C8orf37 homo | | sp\|Q3UJP5 | 4 | 4 | 22 |
| Q99LI2 | Q99LI2 | Chloride ch | Clcc1 | sp\|Q99LI2 | 12 | 12 | 34,7 |
| P97865 | P97865 | Peroxisom | Pex7 | sp\|P97865 | 2 | 2 | 9,7 |
| Q8BL48 | Q8BL48 | RING finge | Unk | sp\|Q8BL48 | 4 | 3 | 7,8 |
| P49586 | P49586 | Choline-ph | Pcyt1a | sp\|P49586 | 9 | 5 | 37,3 |
| A2AAY5 | A2AAY5 | SH3 and PX | Sh3pxd2b | sp\|A2AAY5 | 23 | 21 | 32,4 |
| P48453 | P48453 | Serine/thre | Ppp3cb | sp\|P48453 | 11 | 6 | 30,5 |
| Q922M7 | Q922M7 | Ashwin | | sp\|Q922M | 9 | 9 | 56,5 |
| Q8BGQ6 | Q8BGQ6 | EF-hand ca | Efcab14 | sp\|Q8BGQ | 6 | 6 | 17,1 |
| Q9Z0U0 | Q9Z0U0 | Xenotropic | Xpr1 | sp\|Q9Z0U0 | 6 | 6 | 13,1 |
| Q91WD5 | Q91WD5 | NADH deh | Ndufs2 | sp\|Q91WD | 19 | 19 | 59,6 |
| Q9R008 | Q9R008 | Mevalonat | Mvk | sp\|Q9R008 | 6 | 6 | 26,1 |
| Q9D4V7 | Q9D4V7 | Rab-like pr | Rabl3 | sp\|Q9D4V | 5 | 5 | 30,1 |
| Q9CYC6 | Q9CYC6 | m7GpppN- | Dcp2 | sp\|Q9CYC6 | 5 | 5 | 15,4 |
| Q8C754 | Q8C754 | Vacuolar p | Vps52 | sp\|Q8C754 | 15 | 15 | 28,1 |
| P54276 | P54276 | DNA mism | Msh6 | sp\|P54276 | 51 | 51 | 49,6 |
| A2AGT5 | A2AGT5 | Cytoskelet | Ckap5 | sp\|A2AGT5 | 79 | 79 | 48,2 |
| P19258 | P19258 | Protein Mp | Mpv17 | sp\|P19258 | 2 | 2 | 19,3 |
| Q8C7D2 | Q8C7D2 | Protein cer | Crbn | sp\|Q8C7D2 | 6 | 6 | 21,6 |
| E9Q6P5 | E9Q6P5 | | Ttc7b | sp\|E9Q6P5 | 3 | 3 | 4,9 |
| Q6NVF9 | Q6NVF9 | Cleavage a | Cpsf6 | sp\|Q6NVF | 13 | 13 | 28,5 |
| Q9QWR8 | Q9QWR8 | Alpha-N-ac | Naga | sp\|Q9QW | 4 | 4 | 13 |
| Q5SUE8 | Q5SUE8 | Ankyrin re | Ankrd40 | sp\|Q5SUE | 4 | 4 | 19 |
| Q8VC48 | Q8VC48 | Peroxisom | Pex12 | sp\|Q8VC48 | 3 | 3 | 14,5 |
| Q7TSI1 | Q7TSI1 | Pleckstrin | Plekhm1 | sp\|Q7TSI1 | 7 | 7 | 8,9 |
| Q9DAV9 | Q9DAV9 | Trimeric in | Tmem38b | sp\|Q9DAV | 2 | 2 | 9,6 |
| Q8BYH3 | Q8BYH3 | tRNA:m(4) | Trmt13 | sp\|Q8BYH | 7 | 7 | 20,4 |
| Q8JZW5 | Q8JZW5 | SH2 domai | Sh2d5 | sp\|Q8JZW | 6 | 6 | 18,4 |
| Q66JX5;P0 | Q66JX5 | FGFR1 onc | Fgfr1op | sp\|Q66JX5 | 10 | 10 | 44,4 |

| | | | | | | | |
|---|---|---|---|---|---|---|---|
| P26350 | P26350 | Prothymos | Ptma | sp\|P26350 | 9 | 9 | 40,5 |
| Q8BHY8 | Q8BHY8 | Sorting ne: | Snx14 | sp\|Q8BHY | 8 | 8 | 10 |
| Q8VDX6 | Q8VDX6 | Transmem | Tmem5 | sp\|Q8VDX | 3 | 3 | 7,4 |
| Q8BGS2 | Q8BGS2 | BolA-like p | Bola2 | sp\|Q8BGS | 4 | 4 | 65,1 |
| Q9CY34 | Q9CY34 | NEDD8-cor | Ube2f | sp\|Q9CY34 | 3 | 3 | 24,9 |
| Q99L20;Q6 | Q99L20 | | Gstt3 | sp\|Q99L20 | 7 | 7 | 35,7 |
| Q9D868 | Q9D868 | Peptidyl-pr | Ppih | sp\|Q9D86 | 8 | 8 | 60,1 |
| Q07235 | Q07235 | Glia-derive | Serpine2 | sp\|Q0723 | 3 | 3 | 9,6 |
| Q66JZ4 | Q66JZ4 | T-cell activ | TCAIM | sp\|Q66JZ4 | 2 | 2 | 5,2 |
| Q60631 | Q60631 | Growth fac | Grb2 | sp\|Q6063 | 10 | 10 | 44,2 |
| Q6NSU3 | Q6NSU3 | Glycosyltra | Glt8d1 | sp\|Q6NSU | 7 | 7 | 27,8 |
| P14069 | P14069 | Protein S1 | S100a6 | sp\|P14069 | 13 | 13 | 97,8 |
| Q8VDS8 | Q8VDS8 | Syntaxin-1 | Stx18 | sp\|Q8VDS | 11 | 11 | 35,3 |
| P48455 | P48455 | Serine/thre | Ppp3cc | sp\|P48455 | 6 | 3 | 13,6 |
| Q9JJ28 | Q9JJ28 | Protein flig | Flii | sp\|Q9JJ28 | 41 | 41 | 41,1 |
| Q8K245 | Q8K245 | | Uvrag | sp\|Q8K245 | 5 | 5 | 7,7 |
| Q99KR7 | Q99KR7 | Peptidyl-pr | Ppif | sp\|Q99KR | 4 | 4 | 39,3 |
| Q66JY6 | Q66JY6 | Rho guanir | Arhgef39 | sp\|Q66JY6 | 1 | 1 | 4,1 |
| Q9CXX9 | Q9CXX9 | CUE domai | Cuedc2 | sp\|Q9CXX | 6 | 6 | 25,7 |
| P81122 | P81122 | Insulin rec | Irs2 | sp\|P81122 | 3 | 3 | 4,1 |
| P70324 | P70324 | T-box tran: | Tbx3 | sp\|P70324 | 3 | 3 | 6,9 |
| Q9DCS1 | Q9DCS1 | Transmem | Tmem176a | sp\|Q9DCS | 3 | 3 | 18 |
| Q6ZQ29 | Q6ZQ29 | Serine/thre | Taok2 | sp\|Q6ZQ2 | 9 | 7 | 9,8 |
| Q5SQX6 | Q5SQX6 | Cytoplasm | Cyfip2 | sp\|Q5SQX | 16 | 2 | 14 |
| Q8R3E3 | Q8R3E3 | WD repeat | Wipi1 | sp\|Q8R3E | 8 | 8 | 35 |
| Q9WV32 | Q9WV32 | Actin-relat | Arpc1b | sp\|Q9WV3 | 17 | 15 | 55,6 |
| Q52KE7 | Q52KE7 | Cyclin-L1 | Ccnl1 | sp\|Q52KE | 11 | 10 | 32,9 |
| P18572 | P18572 | Basigin | Bsg | sp\|P18572 | 10 | 10 | 25,7 |
| Q921E6 | Q921E6 | Polycomb | Eed | sp\|Q921E | 17 | 17 | 50,3 |
| Q922Q9 | Q922Q9 | Chitinase d | Chid1 | sp\|Q922Q | 8 | 8 | 27,7 |
| Q3TEA8 | Q3TEA8 | Heterochr | Hp1bp3 | sp\|Q3TEA | 21 | 21 | 41,3 |
| P04370 | P04370 | Myelin bas | Mbp | sp\|P04370 | 2 | 2 | 25,6 |
| Q9DBG3 | Q9DBG3 | AP-2 comp | Ap2b1 | sp\|Q9DBG | 42 | 25 | 49,2 |
| Q570Y9 | Q570Y9 | DEP domai | Deptor | sp\|Q570Y | 8 | 8 | 26,9 |
| Q61609 | Q61609 | Sodium-de | Slc20a1 | sp\|Q6160 | 2 | 2 | 3,7 |
| Q497H0 | Q497H0 | AN1-type z | Zfand3 | sp\|Q497H | 1 | 1 | 5,3 |
| Q8K0C1 | Q8K0C1 | Importin-1 | Ipo13 | sp\|Q8K0C | 8 | 8 | 10,9 |
| Q8C079;Q8 | Q8C079 | Striatin-int | Strip1 | sp\|Q8C07 | 17 | 17 | 27,7 |
| Q6DVA0 | Q6DVA0 | LEM doma | Lemd2 | sp\|Q6DVA | 14 | 14 | 37,4 |
| Q9DCT5 | Q9DCT5 | Stromal ce | Sdf2 | sp\|Q9DCT | 5 | 5 | 36 |
| Q924T2 | Q924T2 | 28S riboso | Mrps2 | sp\|Q924T2 | 7 | 7 | 35,4 |
| Q9QYM8 | Q9QYM8 | Centromer | Cenph | sp\|Q9QYM | 5 | 5 | 24,1 |
| Q9CUN6 | Q9CUN6 | E3 ubiquiti | Smurf1 | sp\|Q9CUN | 4 | 2 | 4,8 |
| Q64514 | Q64514 | Tripeptidyl | Tpp2 | sp\|Q64514 | 52 | 52 | 53,1 |
| Q6A051 | Q6A051 | Attractin-li | Atrnl1 | sp\|Q6A05 | 3 | 3 | 2,6 |
| Q8VEJ4 | Q8VEJ4 | Notchless | Nle1 | sp\|Q8VEJ4 | 15 | 15 | 45,8 |
| P97370 | P97370 | Sodium/pc | Atp1b3 | sp\|P97370 | 15 | 15 | 61,9 |
| Q05AH6 | Q05AH6 | Uncharacterized prote | | sp\|Q05AH | 3 | 3 | 13,1 |
| P52760 | P52760 | Ribonuclea | Hrsp12 | sp\|P52760 | 8 | 8 | 84,4 |
| Q3UHF3 | Q3UHF3 | Mesoderm | Mier3 | sp\|Q3UHF | 3 | 3 | 7,1 |

| | | | | | | | |
|---|---|---|---|---|---|---|---|
| Q9R0E1 | Q9R0E1 | Procollage | Plod3 | sp\|Q9R0E1 | 20 | 20 | 35,4 |
| Q80XR2 | Q80XR2 | Calcium-tr | Atp2c1 | sp\|Q80XR2 | 12 | 11 | 20,5 |
| Q62095;Q6 | Q62095 | ATP-depen | Ddx3y | sp\|Q62095 | 29 | 1 | 42,7 |
| Q3UJ81 | Q3UJ81 | Nuclear en | Cnep1r1 | sp\|Q3UJ81 | 4 | 4 | 40 |
| O09044;P6 | O09044 | Synaptoso | Snap23 | sp\|O09044 | 10 | 10 | 68,1 |
| Q8BQS4 | Q8BQS4 | Protein FA | Fam102b | sp\|Q8BQS4 | 8 | 8 | 39,8 |
| Q8BW96 | Q8BW96 | Calcium/ca | Camk1d | sp\|Q8BW9 | 3 | 1 | 11,7 |
| Q9R1X5 | Q9R1X5 | Multidrug | Abcc5 | sp\|Q9R1X5 | 3 | 3 | 3,1 |
| Q06186 | Q06186 | Proheparin | Hbegf | sp\|Q06186 | 3 | 3 | 19,7 |
| Q99L13 | Q99L13 | 3-hydroxyi | Hibadh | sp\|Q99L13 | 11 | 11 | 43,3 |
| Q9D8B3 | Q9D8B3 | Charged m | Chmp4b | sp\|Q9D8B3 | 5 | 5 | 34,8 |
| Q3UIW5 | Q3UIW5 | RING finge | Rnf10 | sp\|Q3UIW | 3 | 3 | 5,5 |
| Q9ESV0 | Q9ESV0 | ATP-depen | Ddx24 | sp\|Q9ESV0 | 21 | 21 | 30,2 |
| Q9D5D8 | Q9D5D8 | Chromodo | Cdyl2 | sp\|Q9D5D | 2 | 2 | 4,4 |
| E9Q8T2 | E9Q8T2 | | Prdm15 | sp\|E9Q8T2 | 2 | 2 | 2,1 |
| Q8BHN5 | Q8BHN5 | RNA-bindir | Rbm45 | sp\|Q8BHN | 8 | 8 | 22,5 |
| Q80UU2 | Q80UU2 | Ribonuclea | Rpp38 | sp\|Q80UU | 12 | 12 | 61,8 |
| P41778;O3 | P41778 | Pre-B-cell l | Pbx1 | sp\|P41778 | 3 | 3 | 13,3 |
| Q3ULW8 | Q3ULW8 | | Parp3 | sp\|Q3ULW | 11 | 11 | 31,7 |
| Q8BKF1 | Q8BKF1 | DNA-direct | Polrmt | sp\|Q8BKF1 | 22 | 22 | 26,4 |
| Q6PFY1 | Q6PFY1 | F-BAR and | Fchsd1 | sp\|Q6PFY1 | 1 | 1 | 1,6 |
| Q9CX56 | Q9CX56 | 26S protea | Psmd8 | sp\|Q9CX56 | 11 | 11 | 33,7 |
| Q9CY57 | Q9CY57 | Chromatin | Chtop | sp\|Q9CY57 | 6 | 6 | 28,9 |
| Q8VC19 | Q8VC19 | 5-aminolev | Alas1 | sp\|Q8VC19 | 3 | 3 | 5,5 |
| Q3TMH2 | Q3TMH2 | Secernin-3 | Scrn3 | sp\|Q3TMH | 5 | 5 | 18,9 |
| E9Q1P8 | E9Q1P8 | Interferon | Irf2bp2 | sp\|E9Q1P8 | 19 | 15 | 56,5 |
| Q9CPR1 | Q9CPR1 | RWD doma | Rwdd4 | sp\|Q9CPR1 | 5 | 5 | 39,9 |
| Q9WUK2 | Q9WUK2 | Eukaryotic | Eif4h | sp\|Q9WUK | 15 | 15 | 57,3 |
| P51410 | P51410 | 60S riboso | Rpl9 | sp\|P51410 | 10 | 10 | 60,9 |
| Q8VBU8 | Q8VBU8 | Protein BA | Banp | sp\|Q8VBU | 5 | 5 | 11,1 |
| P13705 | P13705 | DNA mism | Msh3 | sp\|P13705 | 3 | 3 | 3 |
| P54728 | P54728 | UV excisio | Rad23b | sp\|P54728 | 16 | 13 | 56,2 |
| Q8C5D8 | Q8C5D8 | E3 SUMO-p | Pias2 | sp\|Q8C5D8 | 8 | 6 | 15,8 |
| Q99L45 | Q99L45 | Eukaryotic | Eif2s2 | sp\|Q99L45 | 27 | 27 | 69,2 |
| Q61753 | Q61753 | D-3-phospl | Phgdh | sp\|Q61753 | 27 | 27 | 53,1 |
| Q8BUI3 | Q8BUI3 | Leucine-ric | LRWD1 | sp\|Q8BUI3 | 21 | 21 | 45,1 |
| Q8K4R9 | Q8K4R9 | Disks large | Dlgap5 | sp\|Q8K4R9 | 19 | 19 | 31,2 |
| Q8BXK4 | Q8BXK4 | Probable methyltrans | | sp\|Q8BXK4 | 1 | 1 | 3,2 |
| Q8BJ90 | Q8BJ90 | Zinc finger | Znf771 | sp\|Q8BJ90 | 2 | 2 | 6,9 |
| Q8C5L6 | Q8C5L6 | Inositol po | Inpp5k | sp\|Q8C5L6 | 6 | 6 | 15 |
| G5E897 | G5E897 | | Kdelc2 | sp\|G5E897 | 11 | 11 | 26,6 |
| Q6JPI3 | Q6JPI3 | Mediator o | Med13l | sp\|Q6JPI3 | 6 | 6 | 4,1 |
| Q99KS2 | Q99KS2 | Neugrin | Ngrn | sp\|Q99KS2 | 7 | 7 | 32,1 |
| Q9D024 | Q9D024 | Coiled-coil | Ccdc47 | sp\|Q9D024 | 18 | 18 | 45,8 |
| Q9EP53 | Q9EP53 | Hamartin | Tsc1 | sp\|Q9EP53 | 7 | 7 | 8,7 |
| Q4VBE8 | Q4VBE8 | WD repeat | Wdr18 | sp\|Q4VBE8 | 12 | 12 | 38,5 |
| Q3UMR0 | Q3UMR0 | Ankyrin re | Ankrd27 | sp\|Q3UMR | 2 | 2 | 2,2 |
| P53026 | P53026 | 60S riboso | Rpl10a | sp\|P53026 | 13 | 13 | 53 |
| Q8CFI5 | Q8CFI5 | Probable p | Pars2 | sp\|Q8CFI5 | 9 | 9 | 28,6 |
| Q6ZPY2 | Q6ZPY2 | Protein SM | Smg5 | sp\|Q6ZPY2 | 12 | 12 | 13,5 |

| | | | | | | | |
|---|---|---|---|---|---|---|---|
| Q9QYR6 | Q9QYR6 | Microtubu | Map1a | sp\|Q9QYR | 8 | 5 | 3,2 |
| Q9JM93 | Q9JM93 | ADP-ribosy | Arl6ip4 | sp\|Q9JM9 | 3 | 3 | 21 |
| O35207;Q9 | O35207 | Cyclin-dep | Cdk2ap1 | sp\|O35207 | 3 | 3 | 43 |
| Q8BW72 | Q8BW72 | Lysine-spe | Kdm4a | sp\|Q8BW7 | 13 | 13 | 14,3 |
| P54310 | P54310 | Hormone-s | Lipe | sp\|P54310 | 6 | 6 | 9,4 |
| Q9CZH8 | Q9CZH8 | Coiled-coil | Ccdc77 | sp\|Q9CZH8 | 3 | 3 | 10,2 |
| Q9D4H4 | Q9D4H4 | Angiomoti | Amotl1 | sp\|Q9D4H | 5 | 5 | 9,1 |
| Q9JMD0 | Q9JMD0 | BUB3-inter | Znf207 | sp\|Q9JMD | 7 | 7 | 17 |
| Q8VE91 | Q8VE91 | Protein FA | Fam134b | sp\|Q8VE91 | 5 | 5 | 19,4 |
| Q6PF93 | Q6PF93 | Phosphatid | Pik3c3 | sp\|Q6PF93 | 14 | 14 | 17,9 |
| P61514 | P61514 | 60S riboso | Rpl37a | sp\|P61514 | 4 | 4 | 51,1 |
| Q3UIU2 | Q3UIU2 | NADH deh | Ndufb6 | sp\|Q3UIU2 | 3 | 3 | 27,3 |
| Q3UHA3 | Q3UHA3 | Spatacsin | Spg11 | sp\|Q3UHA | 4 | 4 | 2,6 |
| Q9D486 | Q9D486 | C-Maf-indu | Cmip | sp\|Q9D48 | 6 | 6 | 10,2 |
| Q8BTZ4 | Q8BTZ4 | Anaphase- | Anapc5 | sp\|Q8BTZ4 | 19 | 19 | 35,8 |
| Q8BN58 | Q8BN58 | Rho GTPas | Arhgap28 | sp\|Q8BN5 | 2 | 2 | 4,3 |
| Q6R891 | Q6R891 | Neurabin-2 | Ppp1r9b | sp\|Q6R891 | 15 | 15 | 30,1 |
| O35841 | O35841 | Apoptosis | Api5 | sp\|O35841 | 22 | 22 | 46,2 |
| Q7TQE6 | Q7TQE6 | Macoilin | Tmem57 | sp\|Q7TQE | 5 | 5 | 11,7 |
| O35245 | O35245 | Polycystin- | Pkd2 | sp\|O35245 | 17 | 17 | 23,1 |
| E9PYK3 | E9PYK3 | | Parp4 | sp\|E9PYK3 | 12 | 12 | 10,3 |
| Q9Z199;P6 | Q9Z199;P6327 | Transcripti | Supt4h1b;S | sp\|Q9Z199 | 6 | 6 | 50,4 |
| Q9D6K8 | Q9D6K8 | FUN14 dor | Fundc2 | sp\|Q9D6K8 | 5 | 5 | 30,5 |
| Q9D818 | Q9D818 | Suppressor | Sapcd2 | sp\|Q9D818 | 6 | 6 | 22,8 |
| Q8R216 | Q8R216 | NAD-deper | Sirt4 | sp\|Q8R216 | 3 | 3 | 9,6 |
| Q9Z2C5 | Q9Z2C5 | Myotubula | Mtm1 | sp\|Q9Z2C5 | 4 | 4 | 8,1 |
| Q91W59 | Q91W59 | RNA-bindir | Rbms1 | sp\|Q91W5 | 7 | 6 | 33 |
| Q62130 | Q62130 | Tyrosine-p | Ptpn14 | sp\|Q62130 | 10 | 10 | 15,6 |
| Q62073 | Q62073 | Mitogen-a | Map3k7 | sp\|Q62073 | 9 | 9 | 21,2 |
| Q69ZS0 | Q69ZS0 | E3 ubiquiti | Pdzrn3 | sp\|Q69ZS0 | 5 | 5 | 6,4 |
| Q99JI4 | Q99JI4 | 26S protea | Psmd6 | sp\|Q99JI4 | 29 | 29 | 66,1 |
| Q80WT5 | Q80WT5 | Aftiphilin | Aftph | sp\|Q80WT | 14 | 14 | 25 |
| Q9CQF4 | Q9CQF4 | Uncharacterized prote | | sp\|Q9CQF4 | 3 | 3 | 15,4 |
| Q8BJM3 | Q8BJM3 | Coiled-coil | R3hcc1l | sp\|Q8BJM | 7 | 7 | 11,9 |
| Q9JL61 | Q9JL61 | DNA-bindi | Rfx5 | sp\|Q9JL61 | 4 | 4 | 7,4 |
| P41133 | P41133 | DNA-bindi | Id3 | sp\|P41133 | 1 | 1 | 10,1 |
| Q921V5 | Q921V5 | Alpha-1,6- | Mgat2 | sp\|Q921V5 | 13 | 13 | 31,7 |
| Q61508 | Q61508 | Extracellul | Ecm1 | sp\|Q61508 | 18 | 18 | 45,1 |
| O70445 | O70445 | BRCA1-ass | Bard1 | sp\|O70445 | 7 | 7 | 17,3 |
| Q80SZ7 | Q80SZ7 | Guanine nu | Gng5 | sp\|Q80SZ7 | 3 | 3 | 63,2 |
| Q3TXT3 | Q3TXT3 | SOSS comp | Inip | sp\|Q3TXT3 | 2 | 2 | 29,8 |
| Q80X71 | Q80X71 | Transmem | Tmem106l | sp\|Q80X71 | 5 | 5 | 28,7 |
| Q8VC42 | Q8VC42 | Uncharacte | Mic1 | sp\|Q8VC42 | 10 | 10 | 23,3 |
| Q9QUR7 | Q9QUR7 | Peptidyl-pr | Pin1 | sp\|Q9QUR | 10 | 10 | 71,5 |
| O08664 | O08664 | B-cell CLL/ | Bcl7c | sp\|O08664 | 5 | 4 | 30,9 |
| Q8R1S0 | Q8R1S0 | Ubiquinon | Coq6 | sp\|Q8R1S0 | 7 | 7 | 18,9 |
| Q8VD76 | Q8VD76 | General tra | Gtf2h3 | sp\|Q8VD7 | 3 | 3 | 12 |
| Q80VQ0 | Q80VQ0 | Aldehyde d | Aldh3b1 | sp\|Q80VQ | 4 | 4 | 10,3 |
| Q9Z2X1 | Q9Z2X1 | Heterogen | Hnrnpf | sp\|Q9Z2X1 | 20 | 18 | 61,9 |
| Q8BFS6 | Q8BFS6 | Serine/thre | Cpped1 | sp\|Q8BFS6 | 7 | 7 | 36,9 |

| | | | | | | | |
|---|---|---|---|---|---|---|---|
| A2AJ88 | A2AJ88 | Patatin-like | Pnpla7 | sp\|A2AJ88 | 8 | 7 | 8,9 |
| Q3TA59 | Q3TA59 | JmjC doma | Jmjd8 | sp\|Q3TA59 | 1 | 1 | 4,8 |
| Q9CWD8 | Q9CWD8 | Iron-sulfur | Nubpl | sp\|Q9CWD | 5 | 5 | 23,5 |
| Q61025 | Q61025 | Intraflagell | Ift20 | sp\|Q61025 | 6 | 6 | 62,1 |
| Q8C0I1 | Q8C0I1 | Alkyldihyd | Agps | sp\|Q8C0I1 | 28 | 28 | 62,6 |
| Q8C7K6 | Q8C7K6 | Prenylcyst | Pcyox1l | sp\|Q8C7K6 | 8 | 8 | 21 |
| Q00547 | Q00547 | Hyalurona | Hmmr | sp\|Q00547 | 22 | 22 | 39,5 |
| Q60960 | Q60960 | Importin su | Kpna1 | sp\|Q60960 | 11 | 5 | 22,9 |
| Q64433 | Q64433 | 10 kDa hea | Hspe1 | sp\|Q64433 | 9 | 9 | 75,5 |
| Q8CFK2 | Q8CFK2 | Transcripti | Brf1 | sp\|Q8CFK2 | 7 | 7 | 13,6 |
| Q62093 | Q62093 | Serine/arg | Srsf2 | sp\|Q62093 | 8 | 8 | 33,9 |
| Q8VE73 | Q8VE73 | Cullin-7 | Cul7 | sp\|Q8VE73 | 4 | 4 | 3,8 |
| Q9D753 | Q9D753 | Exosome c | Exosc8 | sp\|Q9D753 | 12 | 12 | 52,5 |
| Q3UN02 | Q3UN02 | Lysocardio | Lclat1 | sp\|Q3UN0 | 7 | 7 | 22,9 |
| P62897;CC | P62897 | Cytochrom | Cycs | sp\|P62897 | 7 | 7 | 56,2 |
| Q9CUU3 | Q9CUU3 | Synaptone | Sycp2 | sp\|Q9CUU | 2 | 2 | 1,8 |
| Q8BSA9 | Q8BSA9 | tRNA wybu | Tyw3 | sp\|Q8BSA9 | 5 | 5 | 28 |
| Q8BX13 | Q8BX13 | E3 ubiquiti | Ube3d | sp\|Q8BX13 | 2 | 2 | 8,2 |
| P57776 | P57776 | Elongation | Eef1d | sp\|P57776 | 15 | 13 | 58,4 |
| Q9CWQ0 | Q9CWQ0 | Diphthine s | Dph5 | sp\|Q9CWQ | 11 | 11 | 41,6 |
| Q9CZN7 | Q9CZN7 | Serine hyd | Shmt2 | sp\|Q9CZN7 | 29 | 28 | 65,1 |
| Q7TT79 | Q7TT79 | Microceph | Mcph1 | sp\|Q7TT79 | 11 | 11 | 19,6 |
| Q91ZJ5 | Q91ZJ5 | UTP--gluco | Ugp2 | sp\|Q91ZJ5 | 16 | 16 | 41,3 |
| Q9R0L6 | Q9R0L6 | Pericentric | Pcm1 | sp\|Q9R0L6 | 46 | 46 | 31,1 |
| Q924L1 | Q924L1 | LETM1 dor | Letmd1 | sp\|Q924L1 | 2 | 2 | 6,9 |
| Q9QUR8 | Q9QUR8 | Semaphori | Sema7a | sp\|Q9QUR | 5 | 5 | 11,6 |
| Q99LC2 | Q99LC2 | Cleavage s | Cstf1 | sp\|Q99LC2 | 16 | 16 | 52,7 |
| Q69ZW3 | Q69ZW3 | EH domain | Ehbp1 | sp\|Q69ZW | 2 | 2 | 2,8 |
| P51791;Q6 | P51791 | H(+)/Cl(-) e | Clcn3 | sp\|P51791 | 8 | 7 | 12,6 |
| Q3UDW8 | Q3UDW8 | Heparan-a | Hgsnat | sp\|Q3UDW | 4 | 4 | 7,6 |
| Q00899;Q3 | Q00899 | Transcripti | Yy1 | sp\|Q00899 | 15 | 15 | 44,9 |
| Q6DTY7;P7 | Q6DTY7 | 6-phospho | Pfkfb4 | sp\|Q6DTY7 | 7 | 6 | 22 |
| Q62048 | Q62048 | Astrocytic | Pea15 | sp\|Q62048 | 6 | 6 | 44,6 |
| Q8K396 | Q8K396 | Meiotic nu | Mnd1 | sp\|Q8K396 | 4 | 4 | 25,9 |
| P28574 | P28574 | Protein ma | Max | sp\|P28574 | 4 | 4 | 31,2 |
| Q8CGC7 | Q8CGC7 | Bifunctiona | Eprs | sp\|Q8CGC | 76 | 76 | 58,6 |
| Q9CWV0 | Q9CWV0 | Mitochond | Malsu1 | sp\|Q9CWV | 3 | 3 | 16,2 |
| Q9QXD8 | Q9QXD8 | LIM domai | Limd1 | sp\|Q9QXD | 13 | 13 | 41,5 |
| Q9QXV1 | Q9QXV1 | Chromobo | Cbx8 | sp\|Q9QXV | 10 | 9 | 39 |
| Q9D0J8 | Q9D0J8 | Parathymo | Ptms | sp\|Q9D0J8 | 2 | 2 | 22,8 |
| Q9QY93 | Q9QY93 | dCTP pyrop | Dctpp1 | sp\|Q9QY93 | 3 | 3 | 23,5 |
| CON__ENS | CON__ENSEMBL:ENSBTAP00000018574 | | | | 5 | 5 | 12,1 |
| P62274 | P62274 | 40S riboso | Rps29 | sp\|P62274 | 4 | 4 | 58,9 |
| P22892 | P22892 | AP-1 comp | Ap1g1 | sp\|P22892 | 24 | 24 | 36,1 |
| D3YXK2 | D3YXK2 | Scaffold at | Safb | sp\|D3YXK2 | 27 | 16 | 29,2 |
| P28659 | P28659 | CUGBP Ela | Celf1 | sp\|P28659 | 11 | 10 | 25,3 |
| Q8CF66 | Q8CF66 | Regulator c | Lamtor4 | sp\|Q8CF66 | 1 | 1 | 10,1 |
| Q9Z2H7 | Q9Z2H7 | PDZ domai | Gipc2 | sp\|Q9Z2H7 | 1 | 1 | 5,4 |
| Q3UDR8 | Q3UDR8 | Protein YIP | Yipf3 | sp\|Q3UDR | 5 | 5 | 17,9 |
| P40240 | P40240 | CD9 antige | Cd9 | sp\|P40240 | 9 | 9 | 26,5 |

| | | | | | | | |
|---|---|---|---|---|---|---|---|
| P63082 | P63082 | V-type pro | Atp6v0c | sp\|P63082 | 1 | 1 | 11,6 |
| Q9D074;Q | Q9D074 | E3 ubiquiti | Mgrn1 | sp\|Q9D074 | 8 | 8 | 25,4 |
| P62814 | P62814 | V-type pro | Atp6v1b2 | sp\|P62814 | 22 | 22 | 66,7 |
| Q8VD72 | Q8VD72 | Tetratricop | Ttc8 | sp\|Q8VD7 | 2 | 2 | 7,4 |
| Q09143 | Q09143 | High affinit | Slc7a1 | sp\|Q09143 | 7 | 7 | 15,4 |
| Q91WD7 | Q91WD7 | Kinesin-like | Kif18a | sp\|Q91WD | 3 | 2 | 4,5 |
| Q80ZU0;Q | Q80ZU0 | ADP-ribosy | Arl5a | sp\|Q80ZU0 | 3 | 2 | 18,4 |
| O35326 | O35326 | Serine/arg | Srsf5 | sp\|O35326 | 12 | 11 | 30,5 |
| Q8BWP8 | Q8BWP8 | Beta-1,4-gl | B4gat1 | sp\|Q8BWP | 9 | 9 | 29,9 |
| Q9CQY5 | Q9CQY5 | Magnesium | Magt1 | sp\|Q9CQY | 6 | 6 | 18,8 |
| Q9CR13 | Q9CR13 | UPF0562 protein C7or | | sp\|Q9CR13 | 2 | 2 | 20,4 |
| Q9JK91 | Q9JK91 | DNA mism | Mlh1 | sp\|Q9JK91 | 14 | 14 | 26,6 |
| Q9JI11 | Q9JI11 | Serine/thre | Stk4 | sp\|Q9JI11 | 13 | 7 | 33,3 |
| O35737 | O35737 | Heterogen | Hnrnph1 | sp\|O35737 | 19 | 10 | 56,1 |
| O55057 | O55057 | Retinal roc | Pde6d | sp\|O55057 | 4 | 4 | 33,3 |
| Q6DFW4 | Q6DFW4 | Nucleolar p | Nop58 | sp\|Q6DFW | 24 | 24 | 53,4 |
| Q8CI04 | Q8CI04 | Conserved | Cog3 | sp\|Q8CI04 | 23 | 23 | 38 |
| Q91X96 | Q91X96 | Guanine nu | Rabif | sp\|Q91X96 | 4 | 4 | 45,5 |
| Q9CZW5 | Q9CZW5 | Mitochond | Tomm70a | sp\|Q9CZW | 25 | 25 | 55,3 |
| Q9D7N6 | Q9D7N6 | 39S riboso | Mrpl30 | sp\|Q9D7N | 2 | 2 | 22,5 |
| P62746 | P62746 | Rho-relate | Rhob | sp\|P62746 | 4 | 2 | 26,5 |
| Q9WTP6 | Q9WTP6 | Adenylate | Ak2 | sp\|Q9WTP | 13 | 13 | 57,3 |
| Q6NS82 | Q6NS82 | Protein FA | Fam134a | sp\|Q6NS82 | 5 | 5 | 15,7 |
| Q6DID5 | Q6DID5 | PWWP dor | Mum1 | sp\|Q6DID5 | 13 | 13 | 31,2 |
| Q99K90 | Q99K90 | TGF-beta-a | Tab2 | sp\|Q99K90 | 3 | 3 | 7,4 |
| Q8K3J5 | Q8K3J5 | Zinc finger | Znf131 | sp\|Q8K3J5 | 3 | 3 | 9,4 |
| Q9R0P6 | Q9R0P6 | Signal pept | Sec11a | sp\|Q9R0P6 | 6 | 6 | 35,8 |
| Q9QXG2 | Q9QXG2 | Rab protei | Chm | sp\|Q9QXG | 9 | 8 | 23,8 |
| Q8BG48 | Q8BG48 | Serine/thre | Stk17b | sp\|Q8BG48 | 2 | 2 | 7,3 |
| Q9D032;Q | Q9D032;Q9CY | Single-stra | Ssbp3;Ssbp | sp\|Q9D032 | 2 | 2 | 7 |
| Q5RKZ7 | Q5RKZ7 | Molybdenu | Mocs1 | sp\|Q5RKZ7 | 3 | 3 | 7,7 |
| P58466 | P58466 | Carboxy-te | Ctdsp1 | sp\|P58466 | 4 | 3 | 16,1 |
| O09043 | O09043 | Napsin-A | Napsa | sp\|O09043 | 2 | 2 | 6,4 |
| Q8C5N3 | Q8C5N3 | Pre-mRNA- | Cwc22 | sp\|Q8C5N3 | 13 | 13 | 13,4 |
| CON__P02 | CON__P02754 | | | | 2 | 2 | 21 |
| O88967 | O88967 | ATP-depen | Yme1l1 | sp\|O88967 | 22 | 22 | 39,9 |
| Q9CQY2 | Q9CQY2 | RNMT-acti | Fam103a1 | sp\|Q9CQY | 5 | 5 | 44,5 |
| O70496 | O70496 | H(+)/Cl(-) e | Clcn7 | sp\|O70496 | 5 | 5 | 12,8 |
| P70193 | P70193 | Leucine-ric | Lrig1 | sp\|P70193 | 24 | 24 | 34,1 |
| P29594 | P29594 | Caspase-2; | Casp2 | sp\|P29594 | 7 | 7 | 22,3 |
| Q9WU84 | Q9WU84 | Copper cha | Ccs | sp\|Q9WU8 | 4 | 4 | 15,3 |
| P26618 | P26618 | Platelet-de | Pdgfra | sp\|P26618 | 4 | 4 | 6,7 |
| P56960 | P56960 | Exosome c | Exosc10 | sp\|P56960 | 39 | 39 | 60,4 |
| P21126 | P21126 | Ubiquitin-l | Ubl4a | sp\|P21126 | 8 | 8 | 58 |
| Q91W86 | Q91W86 | Vacuolar p | Vps11 | sp\|Q91W8 | 15 | 15 | 21,5 |
| Q61387 | Q61387 | Cytochrom | Cox7a2l | sp\|Q61387 | 2 | 2 | 40,5 |
| Q9ERH4 | Q9ERH4 | Nucleolar a | Nusap1 | sp\|Q9ERH4 | 9 | 9 | 21,8 |
| Q8VEK6 | Q8VEK6 | Inhibitor o | Ing3 | sp\|Q8VEK6 | 2 | 2 | 9,5 |
| Q920B9 | Q920B9 | FACT comp | Supt16h | sp\|Q920B9 | 44 | 44 | 45,4 |
| P27601 | P27601 | Guanine nu | Gna13 | sp\|P27601 | 16 | 15 | 50,1 |

| | | | | | | | |
|---|---|---|---|---|---|---|---|
| Q9JL62 | Q9JL62 | Glycolipid | Gltp | sp\|Q9JL62 | 4 | 4 | 22 |
| P63017 | P63017 | Heat shock | Hspa8 | sp\|P63017 | 46 | 31 | 78,8 |
| P42232 | P42232 | Signal tran | Stat5b | sp\|P42232 | 17 | 5 | 24,6 |
| Q8R1B0;P9 | Q8R1B0 | SH3 and cy | Stac2 | sp\|Q8R1B( | 3 | 3 | 11,8 |
| A2AUU0 | A2AUU0 | Methyltrar | Mettl8 | sp\|A2AUU | 2 | 2 | 12,5 |
| Q9CQ13 | Q9CQ13 | Coordinato | Coprs | sp\|Q9CQ1 | 1 | 1 | 17,3 |
| Q8VHL1 | Q8VHL1 | Histone-lys | Setd7 | sp\|Q8VHL | 9 | 9 | 40,7 |
| Q99KN9 | Q99KN9 | Clathrin int | Clint1 | sp\|Q99KN | 16 | 16 | 28,5 |
| Q9CV28 | Q9CV28 | Protein FA | Fam188a | sp\|Q9CV2 | 6 | 6 | 21,8 |
| Q8CFD4 | Q8CFD4 | Sorting ne | Snx8 | sp\|Q8CFD | 3 | 3 | 8,9 |
| Q6NXY1 | Q6NXY1 | TBC1 doma | Tbc1d31 | sp\|Q6NXY | 4 | 4 | 6,5 |
| Q9DC70 | Q9DC70 | NADH deh | Ndufs7 | sp\|Q9DC7 | 5 | 5 | 30,4 |
| Q99J99 | Q99J99 | 3-mercapt | Mpst | sp\|Q99J99 | 13 | 13 | 57,2 |
| Q61164 | Q61164 | Transcripti | Ctcf | sp\|Q61164 | 14 | 14 | 25,8 |
| Q11204 | Q11204 | CMP-N-ace | St3gal2 | sp\|Q11204 | 3 | 3 | 9,1 |
| Q8R3P6 | Q8R3P6 | von Willeb | Vwa9 | sp\|Q8R3P | 12 | 12 | 36,1 |
| P47934 | P47934 | Carnitine C | Crat | sp\|P47934 | 9 | 9 | 17,3 |
| Q9CYN9 | Q9CYN9 | Renin rece | Atp6ap2 | sp\|Q9CYN | 8 | 8 | 24,6 |
| P99027 | P99027 | 60S acidic | Rplp2 | sp\|P99027 | 12 | 12 | 95,7 |
| Q14B70 | Q14B70 | Highly dive | Hdx | sp\|Q14B7 | 2 | 2 | 3,2 |
| P00397 | P00397 | Cytochrom | Mtco1 | sp\|P00397 | 1 | 1 | 6,2 |
| Q99KD5 | Q99KD5 | Protein un | Unc45a | sp\|Q99KD | 24 | 24 | 32,5 |
| Q8BQY8 | Q8BQY8 | Checkpoin | Hus1 | sp\|Q8BQY | 4 | 4 | 19,6 |
| Q9CQ37 | Q9CQ37 | Ubiquitin-c | Ube2t | sp\|Q9CQ3 | 5 | 4 | 34,3 |
| Q9WUM5 | Q9WUM5 | Succinyl-Co | Suclg1 | sp\|Q9WU | 9 | 9 | 35,5 |
| Q9D110 | Q9D110 | 5-formylte | Mthfs | sp\|Q9D11 | 6 | 6 | 38,4 |
| Q3UR70 | Q3UR70 | Transformi | Tgfbrap1 | sp\|Q3UR7 | 2 | 2 | 3,6 |
| Q99LC9 | Q99LC9 | Peroxisom | Pex6 | sp\|Q99LC | 11 | 11 | 17,7 |
| P07724 | P07724 | Serum albu | Alb | sp\|P07724 | 22 | 20 | 45,9 |
| Q61730 | Q61730 | Interleukin | Il1rap | sp\|Q6173 | 6 | 6 | 11,4 |
| Q9EQH3 | Q9EQH3 | Vacuolar p | Vps35 | sp\|Q9EQH | 27 | 27 | 42,1 |
| Q61037 | Q61037 | Tuberin | Tsc2 | sp\|Q61037 | 8 | 8 | 6,6 |
| Q99KK1 | Q99KK1 | Receptor e | Reep3 | sp\|Q99KK | 5 | 5 | 24,8 |
| Q9CY64 | Q9CY64 | Biliverdin r | Blvra | sp\|Q9CY64 | 14 | 14 | 48,8 |
| Q9D385 | Q9D385 | ADP-ribosy | Arl2bp | sp\|Q9D38 | 2 | 2 | 14,7 |
| P97464 | P97464 | Exostosin-1 | Ext1 | sp\|P97464 | 3 | 2 | 4,4 |
| Q9CZ15 | Q9CZ15 | DNA replic | Gins1 | sp\|Q9CZ1 | 7 | 7 | 46,9 |
| Q9R0C8;P2 | Q9R0C8 | Guanine nu | Vav3 | sp\|Q9R0C | 13 | 13 | 21,1 |
| Q8K1C0 | Q8K1C0 | Protein an | Angel2 | sp\|Q8K1C | 7 | 7 | 15,4 |
| P99029 | P99029 | Peroxiredo | Prdx5 | sp\|P99029 | 13 | 13 | 57,6 |
| Q9QXA5 | Q9QXA5 | U6 snRNA- | Lsm4 | sp\|Q9QXA | 6 | 6 | 46 |
| Q6S5J6 | Q6S5J6 | Krev intera | Krit1 | sp\|Q6S5J6 | 8 | 8 | 14,9 |
| Q8VCE4 | Q8VCE4 | Uncharacterized prote | | sp\|Q8VCE | 1 | 1 | 12,9 |
| Q3UE17 | Q3UE17 | RNA-bindir | Mex3d | sp\|Q3UE1 | 6 | 4 | 15,4 |
| Q8BKT8 | Q8BKT8 | HAUS augr | Haus7 | sp\|Q8BKT | 8 | 8 | 37,4 |
| Q6GYP7 | Q6GYP7 | Ral GTPase | Ralgapa1 | sp\|Q6GYP | 14 | 14 | 9,6 |
| P35290 | P35290 | Ras-relatec | Rab24 | sp\|P35290 | 5 | 5 | 38,9 |
| E9Q309 | E9Q309 | | Cep350 | sp\|E9Q30 | 7 | 7 | 2,9 |
| Q3U3D7 | Q3U3D7 | Transmem | Kiaa0922 | sp\|Q3U3D | 6 | 6 | 5,6 |
| Q99KP3 | Q99KP3 | Lambda-cr | Cryl1 | sp\|Q99KP | 5 | 5 | 22,3 |

| | | | | | | | |
|---|---|---|---|---|---|---|---|
| Q5ND34 | Q5ND34 | WD repeat | Wdr81 | sp\|Q5ND3 | 10 | 10 | 8,5 |
| Q6ZWM4 | Q6ZWM4 | U6 snRNA- | Lsm8 | sp\|Q6ZWN | 6 | 6 | 79,2 |
| Q63844 | Q63844 | Mitogen-a | Mapk3 | sp\|Q63844 | 17 | 12 | 50 |
| Q8C0L8 | Q8C0L8 | Conserved | Cog5 | sp\|Q8C0L8 | 17 | 17 | 32,6 |
| Q8BK03 | Q8BK03 | Protein FA | Fam73b | sp\|Q8BK03 | 3 | 3 | 5,7 |
| Q9D8P7 | Q9D8P7 | General tra | Gtf3c6 | sp\|Q9D8P7 | 2 | 2 | 7 |
| Q8BHN7 | Q8BHN7 | Uncharacterized prote | | sp\|Q8BHN | 7 | 7 | 31,5 |
| O88685 | O88685 | 26S protea | Psmc3 | sp\|O88685 | 29 | 29 | 66,1 |
| P60898 | P60898 | DNA-direct | Polr2i | sp\|P60898 | 7 | 7 | 78,4 |
| P58774;CC | P58774;CON__ | Tropomyos | Tpm2 | sp\|P58774 | 15 | 4 | 32,4 |
| Q78WZ7 | Q78WZ7 | DNA-direct | Twistnb | sp\|Q78WZ | 8 | 8 | 40 |
| Q8BU33 | Q8BU33 | Acetolacta | Ilvbl | sp\|Q8BU3 | 15 | 15 | 33,2 |
| Q9JJ00 | Q9JJ00 | Phospholip | Plscr1 | sp\|Q9JJ00 | 6 | 6 | 20,4 |
| Q9CXE2 | Q9CXE2 | B-cell CLL/ | Bcl7a | sp\|Q9CXE2 | 4 | 2 | 21,9 |
| Q8CBY8 | Q8CBY8 | Dynactin s | Dctn4 | sp\|Q8CBY8 | 15 | 15 | 40,9 |
| Q921I9 | Q921I9 | Exosome c | Exosc4 | sp\|Q921I9 | 9 | 9 | 50,6 |
| Q8BJ83 | Q8BJ83 | TM2 doma | Tm2d3 | sp\|Q8BJ83 | 2 | 2 | 11,5 |
| Q8K2A8 | Q8K2A8 | Dol-P-Man | Alg3 | sp\|Q8K2A8 | 2 | 2 | 4,6 |
| Q8R1N0 | Q8R1N0 | Zinc finger | Znf830 | sp\|Q8R1N | 11 | 11 | 49 |
| Q91ZX6 | Q91ZX6 | Sentrin-spe | Senp2 | sp\|Q91ZX6 | 2 | 2 | 6,1 |
| Q60715 | Q60715 | Prolyl 4-hy | P4ha1 | sp\|Q60715 | 26 | 26 | 52,8 |
| P57784 | P57784 | U2 small n | Snrpa1 | sp\|P57784 | 16 | 16 | 55,7 |
| P43277 | P43277 | Histone H1 | Hist1h1d | sp\|P43277 | 14 | 1 | 46,6 |
| Q8CG50 | Q8CG50 | Ras-relatec | Rab43 | sp\|Q8CG5 | 7 | 7 | 44,3 |
| P62855 | P62855 | 40S riboso | Rps26 | sp\|P62855 | 4 | 4 | 37,4 |
| Q8BU85 | Q8BU85 | Methionin | Msrb3 | sp\|Q8BU8 | 6 | 6 | 39,5 |
| Q9DCG9 | Q9DCG9 | Multifunct | Trmt112 | sp\|Q9DCG | 6 | 6 | 64 |
| Q8C1Z8 | Q8C1Z8 | tRNA meth | Trmt10a | sp\|Q8C1Z8 | 7 | 7 | 28 |
| Q8BVR6 | Q8BVR6 | RING finge | Rspry1 | sp\|Q8BVR | 3 | 3 | 9,2 |
| Q9EQQ2 | Q9EQQ2 | Protein YIP | Yipf5 | sp\|Q9EQQ | 2 | 2 | 17,5 |
| Q8VCD7 | Q8VCD7 | Lysine-spe | Kdm4c | sp\|Q8VCD | 6 | 6 | 9,4 |
| Q9JMG2 | Q9JMG2 | C1GALT1-s | C1galt1c1 | sp\|Q9JMG | 3 | 3 | 13,3 |
| Q9CQR2 | Q9CQR2 | 40S riboso | Rps21 | sp\|Q9CQR | 6 | 6 | 59 |
| Q5HZJ0 | Q5HZJ0 | Ribonuclea | Drosha | sp\|Q5HZJ0 | 18 | 18 | 18,6 |
| P01942 | P01942 | Hemoglobi | Hba | sp\|P01942 | 5 | 3 | 34,5 |
| O88622 | O88622 | Poly(ADP-r | Parg | sp\|O88622 | 17 | 17 | 23,1 |
| Q78IK2 | Q78IK2 | Up-regulat | Usmg5 | sp\|Q78IK2 | 3 | 3 | 44,8 |
| Q9Z2X2 | Q9Z2X2 | 26S protea | Psmd10 | sp\|Q9Z2X2 | 4 | 4 | 28,1 |
| Q8BW00 | Q8BW00 | Probable p | Ptrh1 | sp\|Q8BW0 | 4 | 4 | 34,3 |
| CON__Q28 | CON__Q28065 | | | | 8 | 8 | 17 |
| Q91WP6 | Q91WP6 | Serine prot | Serpina3n | sp\|Q91WP | 2 | 1 | 7,9 |
| Q9D8X1 | Q9D8X1 | Copper ho | Cutc | sp\|Q9D8X1 | 2 | 2 | 12,1 |
| Q8R3B7 | Q8R3B7 | Bromodom | Brd8 | sp\|Q8R3B7 | 12 | 12 | 18,7 |
| Q9JIZ9 | Q9JIZ9 | Phospholip | Plscr3 | sp\|Q9JIZ9 | 4 | 4 | 20,3 |
| P12970 | P12970 | 60S riboso | Rpl7a | sp\|P12970 | 20 | 20 | 47,7 |
| Q64010 | Q64010 | Adapter m | Crk | sp\|Q64010 | 15 | 15 | 54,9 |
| Q8CH36 | Q8CH36 | Proton-cou | Slc36a4 | sp\|Q8CH3 | 2 | 2 | 6,6 |
| Q9DBY1 | Q9DBY1 | E3 ubiquiti | Syvn1 | sp\|Q9DBY1 | 4 | 4 | 6 |
| Q9CQH7 | Q9CQH7 | Transcripti | Btf3l4 | sp\|Q9CQH | 7 | 7 | 55,1 |
| Q9D2Z4 | Q9D2Z4 | Sentrin-spe | Senp8 | sp\|Q9D2Z4 | 2 | 2 | 14 |

| | | | | | | | |
|---|---|---|---|---|---|---|---|
| Q9CQV1 | Q9CQV1 | Mitochond | Pam16 | sp\|Q9CQV | 2 | 2 | 32 |
| Q9D8C6 | Q9D8C6 | Mediator c | Med11 | sp\|Q9D8C | 3 | 3 | 39,3 |
| P03930 | P03930 | ATP syntha | Mtatp8 | sp\|P03930 | 2 | 2 | 34,3 |
| Q91ZR1 | Q91ZR1 | Ras-relatec | Rab4b | sp\|Q91ZR1 | 8 | 6 | 50,2 |
| Q8K371 | Q8K371 | Angiomoti | Amotl2 | sp\|Q8K371 | 5 | 5 | 6,9 |
| Q99JW1 | Q99JW1 | Alpha/beta | Abhd17a | sp\|Q99JW | 2 | 2 | 11 |
| Q6P8I4 | Q6P8I4 | PEST prote | Pcnp | sp\|Q6P8I4 | 6 | 6 | 38,2 |
| P70180 | P70180 | Atrial natri | Npr3 | sp\|P70180 | 9 | 9 | 21,1 |
| Q08024 | Q08024 | Core-bindi | Cbfb | sp\|Q08024 | 7 | 7 | 41,2 |
| O35261;P5 | O35261 | Transcripti | E2f3 | sp\|O35261 | 4 | 4 | 11,8 |
| Q6A028 | Q6A028 | Switch-ass | Swap70 | sp\|Q6A028 | 26 | 26 | 45,6 |
| Q9JKF7 | Q9JKF7 | 39S riboso | Mrpl39 | sp\|Q9JKF7 | 17 | 17 | 52,4 |
| Q9DCD6 | Q9DCD6 | Gamma-ar | Gabarap | sp\|Q9DCD | 2 | 1 | 15,4 |
| Q7TMF3 | Q7TMF3 | NADH deh | Ndufa12 | sp\|Q7TMF | 9 | 9 | 60,7 |
| Q9D7I8 | Q9D7I8 | Protein FA | Fam83d | sp\|Q9D7I8 | 6 | 6 | 11,5 |
| Q8K1A0 | Q8K1A0 | Methyltrar | Mettl5 | sp\|Q8K1A | 4 | 4 | 28,2 |
| Q9CWM4 | Q9CWM4 | Prefoldin s | Pfdn1 | sp\|Q9CWN | 7 | 7 | 43,4 |
| Q8BGH2 | Q8BGH2 | Sorting and | Samm50 | sp\|Q8BGH | 18 | 18 | 50,7 |
| P25119 | P25119 | Tumor nec | Tnfrsf1b | sp\|P25119 | 2 | 2 | 7 |
| Q9CY50 | Q9CY50 | Translocon | Ssr1 | sp\|Q9CY5C | 5 | 5 | 26,9 |
| Q9CR76 | Q9CR76 | Transmem | Tmem186 | sp\|Q9CR76 | 2 | 2 | 8,8 |
| Q99KU0 | Q99KU0 | Vacuole m | Vmp1 | sp\|Q99KU | 3 | 3 | 11,6 |
| Q80WC3 | Q80WC3 | Trinucleoti | Tnrc18 | sp\|Q80WC | 2 | 2 | 0,9 |
| Q8R527 | Q8R527 | Rho-relate | Rhoq | sp\|Q8R527 | 5 | 4 | 29,3 |
| Q8BMQ8 | Q8BMQ8 | Vacuolar fu | Mon1b | sp\|Q8BMC | 2 | 2 | 6,1 |
| Q9CXH7 | Q9CXH7 | Shugoshin- | Sgol1 | sp\|Q9CXH | 7 | 7 | 15,9 |
| Q91VU7 | Q91VU7 | | Pus7 | sp\|Q91VU | 30 | 30 | 60,2 |
| Q9JK23 | Q9JK23 | Proteasom | Psmg1 | sp\|Q9JK23 | 13 | 13 | 54 |
| Q7TS68 | Q7TS68 | Putative m | Nsun6 | sp\|Q7TS68 | 2 | 2 | 6,1 |
| Q9D842 | Q9D842 | Aprataxin a | Aplf | sp\|Q9D842 | 9 | 9 | 29,3 |
| Q5SQY2 | Q5SQY2 | Biorientati | Bod1 | sp\|Q5SQY2 | 4 | 2 | 29,5 |
| Q9Z2X8 | Q9Z2X8 | Kelch-like | Keap1 | sp\|Q9Z2X8 | 18 | 18 | 32,9 |
| Q9CR68 | Q9CR68 | Cytochrom | Uqcrfs1 | sp\|Q9CR68 | 16 | 16 | 56,2 |
| P17012;P2 | P17012;P2360 | Zinc finger | Zfx;Zfa | sp\|P17012 | 8 | 8 | 14,3 |
| O70566 | O70566 | Protein dia | Diaph2 | sp\|O70566 | 7 | 7 | 7,1 |
| Q80XI4 | Q80XI4 | Phosphatic | Pip4k2b | sp\|Q80XI4 | 12 | 8 | 38,2 |
| Q80YV2 | Q80YV2 | Nuclear-int | Zc3hc1 | sp\|Q80YV2 | 11 | 11 | 36,7 |
| P59017 | P59017 | Bcl-2-like p | Bcl2l13 | sp\|P59017 | 9 | 9 | 39,6 |
| Q78XR0 | Q78XR0 | Trafficking | Trappc6a | sp\|Q78XR0 | 2 | 2 | 17 |
| Q9QZB1 | Q9QZB1 | Regulator c | Rgs20 | sp\|Q9QZB | 2 | 2 | 9,2 |
| P63037 | P63037 | DnaJ homo | Dnaja1 | sp\|P63037 | 16 | 16 | 57,7 |
| Q8VCB1 | Q8VCB1 | Nucleopor | Ndc1 | sp\|Q8VCB1 | 21 | 21 | 40,7 |
| Q80Y98 | Q80Y98 | Phospholip | Ddhd2 | sp\|Q80Y98 | 12 | 12 | 27,8 |
| Q9EQK7 | Q9EQK7 | Protein-S-i | Icmt | sp\|Q9EQK | 3 | 3 | 9,9 |
| Q9CXZ1 | Q9CXZ1 | NADH deh | Ndufs4 | sp\|Q9CXZ1 | 6 | 6 | 31,4 |
| D3Z4S3 | D3Z4S3 | Putative pe | Ptrhd1 | sp\|D3Z4S3 | 4 | 4 | 21,4 |
| Q5HZI9 | Q5HZI9 | Solute carr | Slc25a51 | sp\|Q5HZI9 | 4 | 4 | 14,4 |
| Q9CWY3 | Q9CWY3 | N-lysine m | Setd6 | sp\|Q9CWY | 9 | 9 | 27,7 |
| Q9QY33 | Q9QY33 | Tetraspani | Tspan3 | sp\|Q9QY33 | 2 | 2 | 9,1 |
| Q7TT28 | Q7TT28 | RNA exonu | Rexo1 | sp\|Q7TT28 | 3 | 3 | 4,5 |

| | | | | | | | |
|---|---|---|---|---|---|---|---|
| Q8VHZ7 | Q8VHZ7 | U3 small n | Imp4 | sp\|Q8VHZ | 9 | 9 | 37,8 |
| Q8CI70 | Q8CI70 | Leucine-ric | Lrrc20 | sp\|Q8CI70 | 3 | 3 | 19 |
| Q80TM6 | Q80TM6 | R3H doma | R3hdm2 | sp\|Q80TM | 3 | 3 | 4,3 |
| Q99MZ7 | Q99MZ7 | Peroxisom | Pecr | sp\|Q99MZ | 6 | 6 | 27,7 |
| Q8BGS7 | Q8BGS7 | Choline/et | Cept1 | sp\|Q8BGS | 4 | 4 | 12,5 |
| Q4U2R1 | Q4U2R1 | E3 ubiquiti | Herc2 | sp\|Q4U2R | 35 | 35 | 11,2 |
| Q8BV13 | Q8BV13 | COP9 signa | Cops7b | sp\|Q8BV1 | 7 | 7 | 31,1 |
| P51162 | P51162 | Gastrotrop | Fabp6 | sp\|P51162 | 7 | 7 | 65,6 |
| Q9CWY8 | Q9CWY8 | Ribonuclea | Rnaseh2a | sp\|Q9CWY | 11 | 11 | 41,2 |
| P51943;Q6 | P51943 | Cyclin-A2 | Ccna2 | sp\|P51943 | 11 | 11 | 43,6 |
| E9PVA8 | E9PVA8 | | Gcn1l1 | sp\|E9PVA8 | 116 | 116 | 49 |
| P61982 | P61982 | 14-3-3 pro | Ywhag | sp\|P61982 | 17 | 12 | 70,4 |
| Q3UMB5 | Q3UMB5 | Smith-Mag | Smcr8 | sp\|Q3UME | 9 | 9 | 12,4 |
| Q3UUJ4 | Q3UUJ4 | STE20-rela | Strada | sp\|Q3UUJ | 4 | 4 | 10,9 |
| Q9EPA7 | Q9EPA7 | Nicotinami | Nmnat1 | sp\|Q9EPA | 4 | 4 | 19,3 |
| O88653 | O88653 | Ragulator c | Lamtor3 | sp\|O88653 | 7 | 7 | 62,9 |
| Q8BUV6 | Q8BUV6 | U7 snRNA- | Lsm11 | sp\|Q8BUV | 4 | 4 | 17,7 |
| Q8K3P5 | Q8K3P5 | CCR4-NOT | Cnot6 | sp\|Q8K3P5 | 6 | 6 | 11,8 |
| Q3U1Y4 | Q3U1Y4 | DENN dom | Dennd4b | sp\|Q3U1Y | 7 | 7 | 6,3 |
| Q8K0H5 | Q8K0H5 | Transcripti | Taf10 | sp\|Q8K0H | 4 | 4 | 30,7 |
| O09053 | O09053 | Werner syı | Wrn | sp\|O09053 | 9 | 9 | 11,8 |
| Q9R0X0 | Q9R0X0 | Mediator c | Med20 | sp\|Q9R0X | 5 | 5 | 38,2 |
| O55102 | O55102 | Biogenesis | Bloc1s1 | sp\|O5510 | 3 | 3 | 25,6 |
| Q5F2E7 | Q5F2E7 | Nuclear fra | Nufip2 | sp\|Q5F2E7 | 17 | 17 | 39,6 |
| Q5HZH2 | Q5HZH2 | Ribosome | Tsr3 | sp\|Q5HZH | 7 | 7 | 31,6 |
| Q3UW53 | Q3UW53 | Protein Nib | Fam129a | sp\|Q3UW5 | 3 | 3 | 6,7 |
| Q9Z0E0 | Q9Z0E0 | Neurochor | Ncdn | sp\|Q9Z0E0 | 12 | 12 | 22,9 |
| Q6PIJ4 | Q6PIJ4 | Nuclear fac | Nfrkb | sp\|Q6PIJ4 | 19 | 19 | 20,8 |
| Q9JHU4 | Q9JHU4 | Cytoplasm | Dync1h1 | sp\|Q9JHU | 216 | 216 | 51,7 |
| P51612 | P51612 | DNA repair | Xpc | sp\|P51612 | 9 | 9 | 15,6 |
| Q8BGR6 | Q8BGR6 | ADP-ribosy | Arl15 | sp\|Q8BGR | 5 | 5 | 42,2 |
| O35892;Q9 | O35892 | Nuclear au | Sp100 | sp\|O35892 | 3 | 3 | 8,5 |
| Q61009 | Q61009 | Scavenger | Scarb1 | sp\|Q61009 | 3 | 3 | 5,7 |
| Q9Z0P7 | Q9Z0P7 | Suppressor | Sufu | sp\|Q9Z0P7 | 8 | 8 | 26 |
| Q8CGY8 | Q8CGY8 | UDP-N-ace | Ogt | sp\|Q8CGY | 37 | 37 | 43,5 |
| Q9JKV5 | Q9JKV5 | Secretory c | Scamp4 | sp\|Q9JKV5 | 1 | 1 | 4,8 |
| Q7TSG2 | Q7TSG2 | RNA polym | Ctdp1 | sp\|Q7TSG | 26 | 26 | 40,6 |
| Q8C804 | Q8C804 | Spindle an | Spice1 | sp\|Q8C80 | 8 | 8 | 14,2 |
| Q9CWR2 | Q9CWR2 | Histone-lys | Smyd3 | sp\|Q9CWR | 7 | 7 | 20,6 |
| O88843 | O88843 | Death dom | Cradd | sp\|O88843 | 3 | 3 | 36,2 |
| P56382 | P56382 | ATP syntha | Atp5e | sp\|P56382 | 2 | 2 | 28,8 |
| Q8BPY9 | Q8BPY9 | Fidgetin-lik | Fignl1 | sp\|Q8BPY | 10 | 9 | 23,9 |
| P06797 | P06797 | Cathepsin | Ctsl | sp\|P06797 | 11 | 11 | 51,2 |
| Q8VDM1 | Q8VDM1 | Zinc finger | Zgpat | sp\|Q8VDM | 11 | 11 | 28,6 |
| Q3THJ3 | Q3THJ3 | Probable R | Eif1ad | sp\|Q3THJ3 | 6 | 6 | 32,9 |
| A2BDX3 | A2BDX3 | Adenylyltra | Mocs3 | sp\|A2BDX | 9 | 9 | 26,7 |
| O55222 | O55222 | Integrin-lin | Ilk | sp\|O5522 | 16 | 16 | 43,1 |
| Q8CHP5 | Q8CHP5 | Partner of | Wibg | sp\|Q8CHP | 7 | 7 | 59,6 |
| Q9EQ80 | Q9EQ80 | NIF3-like p | Nif3l1 | sp\|Q9EQ8 | 10 | 10 | 35,4 |
| Q99ML4 | Q99ML4 | Protein FA | Fam69b | sp\|Q99ML | 5 | 5 | 15,1 |

| | | | | | | | |
|---|---|---|---|---|---|---|---|
| Q8BTV1 | Q8BTV1 | Tumor sup | Tusc3 | sp\|Q8BTV1 | 3 | 3 | 8,4 |
| Q99KU6 | Q99KU6 | Bombesin receptor-ac | | sp\|Q99KU | 2 | 2 | 6,3 |
| Q8K0T2 | Q8K0T2 | Cytoplasm | Dync2li1 | sp\|Q8K0T2 | 3 | 3 | 11,1 |
| CON__P17 | CON__P17690 | | | | 12 | 12 | 47,5 |
| Q9QZD4 | Q9QZD4 | DNA repair | Ercc4 | sp\|Q9QZD | 14 | 14 | 21,8 |
| P07903 | P07903 | DNA excisi | Ercc1 | sp\|P07903 | 7 | 7 | 29,5 |
| Q3U0M1 | Q3U0M1 | Trafficking | Trappc9 | sp\|Q3U0M | 12 | 12 | 12 |
| Q8K1Z0 | Q8K1Z0 | Ubiquinone | Coq9 | sp\|Q8K1Z0 | 6 | 6 | 30 |
| Q6PNC0 | Q6PNC0 | DmX-like p | Dmxl1 | sp\|Q6PNC | 13 | 13 | 6,2 |
| Q14CH7 | Q14CH7 | Alanine--tR | Aars2 | sp\|Q14CH7 | 17 | 17 | 27 |
| Q3TAS6 | Q3TAS6 | ER membra | Emc10 | sp\|Q3TAS6 | 3 | 3 | 20,5 |
| Q61107 | Q61107 | Guanylate- | Gbp4 | sp\|Q61107 | 3 | 3 | 7,1 |
| Q9R061 | Q9R061 | Cytosolic F | Nubp2 | sp\|Q9R061 | 6 | 6 | 26,5 |
| Q60817;P7 | Q60817;P7067 | Nascent po | Naca | sp\|Q60817 | 7 | 7 | 52,1 |
| Q61686 | Q61686 | Chromobo | Cbx5 | sp\|Q61686 | 10 | 9 | 47,1 |
| Q6PHQ8 | Q6PHQ8 | N-alpha-ac | Naa35 | sp\|Q6PHQ | 18 | 18 | 29,7 |
| Q8BXN9 | Q8BXN9 | Transmem | Tmem87a | sp\|Q8BXN | 9 | 9 | 19,6 |
| Q9D945 | Q9D945 | Protein LLF | Llph | sp\|Q9D945 | 5 | 5 | 29,2 |
| Q8CJG1 | Q8CJG1 | Protein arg | Ago1 | sp\|Q8CJG1 | 13 | 3 | 21,1 |
| Q8BP47 | Q8BP47 | Asparagine | Nars | sp\|Q8BP47 | 24 | 24 | 47,9 |
| Q921E2 | Q921E2 | Ras-related | Rab31 | sp\|Q921E2 | 11 | 9 | 72,7 |
| Q9ERR7 | Q9ERR7 | 15 kDa sele | 15-set | sp\|Q9ERR7 | 5 | 5 | 32,7 |
| Q6PB44 | Q6PB44 | Tyrosine-p | Ptpn23 | sp\|Q6PB44 | 45 | 45 | 33 |
| Q9Z1J2 | Q9Z1J2 | Serine/thre | Nek4 | sp\|Q9Z1J2 | 5 | 5 | 9,3 |
| Q9WTK5 | Q9WTK5 | Nuclear fac | Nfkb2 | sp\|Q9WTK | 26 | 26 | 42,4 |
| Q8CI08 | Q8CI08 | SLAIN mot | Slain2 | sp\|Q8CI08 | 12 | 12 | 31,3 |
| Q8CGP2;Q | Q8CGP2;Q8CG | Histone H2 | Hist1h2bp; | sp\|Q8CGP | 10 | 2 | 57,9 |
| Q9D8Z6 | Q9D8Z6 | Autophagy | Atg101 | sp\|Q9D8Z6 | 4 | 4 | 22,5 |
| P38060 | P38060 | Hydroxyme | Hmgcl | sp\|P38060 | 8 | 8 | 36 |
| Q9R099 | Q9R099 | Transducin | Tbl2 | sp\|Q9R099 | 15 | 15 | 36,4 |
| Q9Z108 | Q9Z108 | Double-str | Stau1 | sp\|Q9Z108 | 23 | 23 | 47,6 |
| Q9Z1Q2 | Q9Z1Q2 | Abhydrolas | Abhd16a | sp\|Q9Z1Q | 14 | 14 | 33 |
| Q61738 | Q61738 | Integrin alp | Itga7 | sp\|Q61738 | 10 | 10 | 13,1 |
| Q8BGH7 | Q8BGH7 | CDC42 sma | Cdc42se2 | sp\|Q8BGH | 1 | 1 | 22,6 |
| P84244;P0 | P84244;P0230 | Histone H3 | H3f3a;H3f3 | sp\|P84244 | 7 | 2 | 55,1 |
| Q9D7K2 | Q9D7K2 | CST compl | Ten1 | sp\|Q9D7K | 3 | 3 | 38,5 |
| Q9D8Y8 | Q9D8Y8 | Inhibitor o | Ing5 | sp\|Q9D8Y8 | 3 | 3 | 14,6 |
| Q9JLL0 | Q9JLL0 | Cysteine-ri | Crim1 | sp\|Q9JLL0 | 1 | 1 | 1,2 |
| P08752;B2 | P08752 | Guanine nu | Gnai2 | sp\|P08752 | 17 | 12 | 59,4 |
| Q9WU56 | Q9WU56 | tRNA pseu | Pus1 | sp\|Q9WU5 | 16 | 16 | 49,2 |
| Q9CYC5 | Q9CYC5 | Kinetochor | Dsn1 | sp\|Q9CYC5 | 4 | 4 | 15,5 |
| Q9D898 | Q9D898 | Actin-relat | Arpc5l | sp\|Q9D898 | 6 | 6 | 58,2 |
| Q9EPX5 | Q9EPX5 | F-box/LRR- | Fbxl12 | sp\|Q9EPX5 | 3 | 3 | 15,6 |
| Q8CJ96 | Q8CJ96 | Ras associa | Rassf8 | sp\|Q8CJ96 | 5 | 5 | 15,5 |
| P63011 | P63011 | Ras-related | Rab3a | sp\|P63011 | 6 | 4 | 39,1 |
| Q3UDP0 | Q3UDP0 | WD repeat | Wdr41 | sp\|Q3UDP | 4 | 4 | 16,3 |
| P97801 | P97801 | Survival m | Smn1 | sp\|P97801 | 5 | 5 | 31,2 |
| Q8K2X3 | Q8K2X3 | CST compl | Obfc1 | sp\|Q8K2X3 | 12 | 12 | 49,2 |
| Q8CHC8 | Q8CHC8 | ADNP hom | Adnp2 | sp\|Q8CHC | 9 | 9 | 10,4 |
| Q3UHD2 | Q3UHD2 | Glucose-fr | Gfod1 | sp\|Q3UHD | 2 | 2 | 4,9 |

| | | | | | | | |
|---|---|---|---|---|---|---|---|
| P56565 | P56565 | Protein S1( | S100a1 | sp\|P56565 | 4 | 4 | 40,4 |
| Q810A3 | Q810A3 | Tetratricop | Ttc9c | sp\|Q810A3 | 3 | 3 | 24 |
| Q6WQJ1 | Q6WQJ1 | Sn1-specifi | Dagla | sp\|Q6WQJ | 2 | 2 | 2,1 |
| Q8C4M7 | Q8C4M7 | Centromer | Cenpu | sp\|Q8C4M | 3 | 3 | 8,3 |
| P25206 | P25206 | DNA replic | Mcm3 | sp\|P25206 | 50 | 50 | 74,6 |
| Q8BKN5 | Q8BKN5 | Gamma-tu | Tubgcp5 | sp\|Q8BKN | 5 | 5 | 6,1 |
| Q9QXG4 | Q9QXG4 | Acetyl-coe | Acss2 | sp\|Q9QXG | 8 | 8 | 17,3 |
| Q9DBS5 | Q9DBS5 | Kinesin ligh | Klc4 | sp\|Q9DBS! | 10 | 4 | 21,3 |
| Q99K95 | Q99K95 | Protein RT | Rtfdc1 | sp\|Q99K95 | 4 | 4 | 14,7 |
| P41105 | P41105 | 60S riboso | Rpl28 | sp\|P41105 | 7 | 7 | 40,9 |
| Q9D125 | Q9D125 | 28S riboso | Mrps25 | sp\|Q9D12! | 7 | 7 | 43,3 |
| Q8CIV2 | Q8CIV2 | Membralin | Tmem259 | sp\|Q8CIV2 | 8 | 8 | 22,1 |
| Q9Z265 | Q9Z265 | Serine/thre | Chek2 | sp\|Q9Z265 | 7 | 7 | 16,3 |
| Q8C735 | Q8C735 | Protein lin- | Lin9 | sp\|Q8C735 | 8 | 8 | 27,5 |
| O89110 | O89110 | Caspase-8; | Casp8 | sp\|O89110 | 16 | 16 | 41,2 |
| A2AQ19 | A2AQ19 | RNA polym | Rtf1 | sp\|A2AQ19 | 13 | 13 | 21,3 |
| P61255 | P61255 | 60S riboso | Rpl26 | sp\|P61255 | 9 | 9 | 44,8 |
| Q9CRA8 | Q9CRA8 | Exosome c | Exosc5 | sp\|Q9CRA8 | 9 | 9 | 62,1 |
| Q80US4 | Q80US4 | Actin-relat | Actr5 | sp\|Q80US4 | 8 | 8 | 18,3 |
| Q61301 | Q61301 | Catenin alp | Ctnna2 | sp\|Q61301 | 8 | 2 | 11,5 |
| O70378 | O70378 | ER membra | Emc8 | sp\|O70378 | 9 | 9 | 57 |
| Q9CQJ2 | Q9CQJ2 | PIH1 doma | Pih1d1 | sp\|Q9CQJ2 | 10 | 10 | 46,2 |
| Q91V04 | Q91V04 | Translocati | Tram1 | sp\|Q91V04 | 7 | 7 | 19,3 |
| Q8BFT2 | Q8BFT2 | HAUS augr | Haus4 | sp\|Q8BFT2 | 8 | 8 | 30,3 |
| Q99L02 | Q99L02 | PAXIP1-ass | Pagr1a | sp\|Q99L02 | 2 | 2 | 28,1 |
| A2AWA9 | A2AWA9 | Rab GTPas | Rabgap1 | sp\|A2AWA | 21 | 21 | 27,4 |
| Q8BPA8 | Q8BPA8 | Protein DP | Dpcd | sp\|Q8BPA8 | 5 | 5 | 24,1 |
| C0HKE9;C0 | C0HKE9;C0HKE | Histone H2 | Hist1h2ah; | sp\|C0HKE9 | 9 | 1 | 68,5 |
| P62254 | P62254 | Ubiquitin-c | Ube2g1 | sp\|P62254 | 7 | 7 | 54,1 |
| Q69ZZ6 | Q69ZZ6 | Transmem | Tmcc1 | sp\|Q69ZZ6 | 3 | 3 | 6,8 |
| Q08890 | Q08890 | Iduronate : | Ids | sp\|Q08890 | 1 | 1 | 1,8 |
| Q8BT51 | Q8BT51 | Cytochrom | COA4 | sp\|Q8BT51 | 2 | 2 | 17,2 |
| P35980 | P35980 | 60S riboso | Rpl18 | sp\|P35980 | 9 | 9 | 41,5 |
| Q3UZ18 | Q3UZ18 | Little elong | Ice2 | sp\|Q3UZ18 | 7 | 7 | 9,6 |
| Q99MR6 | Q99MR6 | Serrate RN | Srrt | sp\|Q99MR | 32 | 32 | 36 |
| Q9DBF7 | Q9DBF7 | Pre-mRNA- | Cwc25 | sp\|Q9DBF7 | 6 | 6 | 17,5 |
| Q9CR39 | Q9CR39 | WD repeat | Wdr45b | sp\|Q9CR39 | 8 | 8 | 25,3 |
| Q8BGZ3 | Q8BGZ3 | DDB1- and | Dcaf12 | sp\|Q8BGZ3 | 2 | 2 | 6 |
| Q9CQ69 | Q9CQ69 | Cytochrom | Uqcrq | sp\|Q9CQ6 | 5 | 5 | 70,7 |
| Q6NSQ7 | Q6NSQ7 | Protein LT\ | Ltv1 | sp\|Q6NSQ | 6 | 6 | 18,5 |
| O08999 | O08999 | Latent-trar | Ltbp2 | sp\|O08999 | 13 | 13 | 10,1 |
| Q9CZR8 | Q9CZR8 | Elongation | Tsfm | sp\|Q9CZR8 | 11 | 11 | 41 |
| P47962 | P47962 | 60S riboso | Rpl5 | sp\|P47962 | 15 | 15 | 55,2 |
| Q9D8N6 | Q9D8N6 | Protein lin- | Lin37 | sp\|Q9D8N | 3 | 3 | 18,7 |
| P51141 | P51141 | Segment p | Dvl1 | sp\|P51141 | 5 | 4 | 11,1 |
| Q80UV9 | Q80UV9 | Transcripti | Taf1 | sp\|Q80UV | 15 | 15 | 10,7 |
| Q8CDG3 | Q8CDG3 | Deubiquiti | Vcpip1 | sp\|Q8CDG | 28 | 28 | 32,4 |
| Q60973 | Q60973 | Histone-bi | Rbbp7 | sp\|Q60973 | 20 | 13 | 64,9 |
| Q6PDY2 | Q6PDY2 | 2-aminoet | Ado | sp\|Q6PDY2 | 9 | 9 | 52,3 |
| Q9JHD2 | Q9JHD2 | Histone ac | Kat2a | sp\|Q9JHD2 | 8 | 6 | 11,6 |

| | | | | | | | |
|---|---|---|---|---|---|---|---|
| Q9DBY5 | Q9DBY5 | Chromobo | Cbx6 | sp\|Q9DBY! | 7 | 6 | 25,8 |
| O08795 | O08795 | Glucosidas | Prkcsh | sp\|O08795 | 21 | 21 | 51,6 |
| Q5SYH2 | Q5SYH2 | Transmem | Tmem199 | sp\|Q5SYH2 | 6 | 6 | 37,5 |
| Q3U2C5 | Q3U2C5 | E3 ubiquiti | Rnf149 | sp\|Q3U2C | 3 | 3 | 11,4 |
| Q9D2H6 | Q9D2H6 | Transcripti | Sp2 | sp\|Q9D2H | 4 | 4 | 10,9 |
| Q08376 | Q08376 | Zinc finger | Zbtb14 | sp\|Q08376 | 1 | 1 | 3,3 |
| P60843 | P60843 | Eukaryotic | Eif4a1 | sp\|P60843 | 33 | 18 | 68,7 |
| Q9DCD2 | Q9DCD2 | Pre-mRNA- | Xab2 | sp\|Q9DCD | 29 | 29 | 44,7 |
| O35593 | O35593 | 26S protea | Psmd14 | sp\|O35593 | 11 | 11 | 56,1 |
| P97480 | P97480 | Eyes abser | Eya3 | sp\|P97480 | 9 | 9 | 21,4 |
| Q8K4B0 | Q8K4B0 | Metastasis | Mta1 | sp\|Q8K4B( | 24 | 22 | 49 |
| A2RSJ4 | A2RSJ4 | UHRF1-bin | Uhrf1bp1l | sp\|A2RSJ4 | 13 | 13 | 15,4 |
| Q8CHZ9;B9 | Q8CHZ9;B9EJ5 | Transcripti | Mterf1a;M | sp\|Q8CHZ! | 3 | 3 | 9 |
| Q9DCK3 | Q9DCK3 | Tetraspani | Tspan4 | sp\|Q9DCK | 2 | 2 | 13,9 |
| P10126;P6 | P10126 | Elongation | Eef1a1 | sp\|P10126 | 30 | 30 | 73,6 |
| P56371 | P56371 | Ras-relate | Rab4a | sp\|P56371 | 7 | 5 | 37,2 |
| Q8K057 | Q8K057 | Intraflagell | Ift80 | sp\|Q8K057 | 4 | 4 | 5,5 |
| Q8R173 | Q8R173 | Palmitoyltr | Zdhhc3 | sp\|Q8R173 | 3 | 3 | 16,4 |
| Q920D3 | Q920D3 | Mediator c | Med28 | sp\|Q920D | 4 | 4 | 30,3 |
| P50544 | P50544 | Very long-c | Acadvl | sp\|P50544 | 33 | 33 | 59,5 |
| Q9CR57 | Q9CR57 | 60S riboso | Rpl14 | sp\|Q9CR57 | 5 | 5 | 25,8 |
| O35405 | O35405 | Phospholip | Pld3 | sp\|O35405 | 5 | 5 | 16 |
| P56873 | P56873 | Sjoegren sy | Sssca1 | sp\|P56873 | 6 | 6 | 64,3 |
| Q61542 | Q61542 | StAR-relate | Stard3 | sp\|Q61542 | 5 | 5 | 20 |
| Q80WS3 | Q80WS3 | rRNA/tRNA | Fbll1 | sp\|Q80WS | 5 | 1 | 20,1 |
| O08917 | O08917 | Flotillin-1 | Flot1 | sp\|O08917 | 17 | 17 | 55,1 |
| Q80ZJ6 | Q80ZJ6 | Protein zer | Zer1 | sp\|Q80ZJ6 | 4 | 4 | 6,7 |
| P47964 | P47964 | 60S riboso | Rpl36 | sp\|P47964 | 2 | 2 | 18,1 |
| Q9ESJ4 | Q9ESJ4 | NCK-intera | Nckipsd | sp\|Q9ESJ4 | 6 | 6 | 10,6 |
| Q08297 | Q08297 | DNA repair | Rad51 | sp\|Q08297 | 11 | 11 | 42,5 |
| Q9CW03 | Q9CW03 | Structural | Smc3 | sp\|Q9CW0 | 54 | 54 | 45,4 |
| B2RQC6 | B2RQC6 | CAD protei | Cad | sp\|B2RQC | 81 | 79 | 52,9 |
| Q6S5C2 | Q6S5C2 | N-acetylglu | Gnptg | sp\|Q6S5C2 | 3 | 3 | 14,3 |
| CON__P13 | CON__P13645 | | | | 21 | 16 | 37,3 |
| Q640Q5 | Q640Q5 | PAB-deper | Pan3 | sp\|Q640Q | 11 | 11 | 16,5 |
| Q922G2 | Q922G2 | Protein FA | Fam76a | sp\|Q922G | 2 | 2 | 8,5 |
| Q8BZ98 | Q8BZ98 | Dynamin-3 | Dnm3 | sp\|Q8BZ98 | 13 | 2 | 13,3 |
| Q8BHA0 | Q8BHA0 | INO80 com | Ino80c | sp\|Q8BHA | 6 | 6 | 52,9 |
| P14869 | P14869 | 60S acidic | Rplp0 | sp\|P14869 | 14 | 14 | 50,5 |
| Q9QYI6 | Q9QYI6 | DnaJ homo | Dnajb9 | sp\|Q9QYI6 | 1 | 1 | 6,8 |
| A2CG49 | A2CG49 | Kalirin | Kalrn | sp\|A2CG49 | 5 | 1 | 1,8 |
| Q9CRB2 | Q9CRB2 | H/ACA ribo | Nhp2 | sp\|Q9CRB | 5 | 5 | 49 |
| Q9CR95 | Q9CR95 | Adaptin ea | Necap1 | sp\|Q9CR95 | 7 | 7 | 44,4 |
| Q9Z306 | Q9Z306 | Solute carr | Slc22a4 | sp\|Q9Z306 | 3 | 3 | 7,4 |
| Q80YT7 | Q80YT7 | Myomegal | Pde4dip | sp\|Q80YT7 | 8 | 8 | 4,4 |
| Q91WV0 | Q91WV0 | Protein Dr | Dr1 | sp\|Q91WV | 4 | 4 | 33 |
| Q91YE6 | Q91YE6 | Importin-9 | Ipo9 | sp\|Q91YE6 | 28 | 28 | 37,7 |
| P97863 | P97863 | Nuclear fac | Nfib | sp\|P97863 | 14 | 12 | 36,7 |
| Q8BPI1 | Q8BPI1 | Protein kin | Dnaaf2 | sp\|Q8BPI1 | 4 | 4 | 9,6 |
| P08228 | P08228 | Superoxide | Sod1 | sp\|P08228 | 9 | 9 | 54,5 |

| | | | | | | | |
|---|---|---|---|---|---|---|---|
| Q58FA4 | Q58FA4 | Transcripti | E2f8 | sp\|Q58FA4 | 7 | 7 | 12,9 |
| Q9EQJ0 | Q9EQJ0 | Two pore c | Tpcn1 | sp\|Q9EQJ0 | 3 | 3 | 3,8 |
| Q4VBF2 | Q4VBF2 | R3H doma | R3hdm4 | sp\|Q4VBF2 | 2 | 2 | 8,4 |
| Q9Z315 | Q9Z315 | U4/U6.U5 | Sart1 | sp\|Q9Z315 | 36 | 36 | 50,9 |
| Q9ESE1 | Q9ESE1 | Lipopolysa | Lrba | sp\|Q9ESE1 | 36 | 31 | 18 |
| O88342 | O88342 | WD repeat | Wdr1 | sp\|O88342 | 32 | 32 | 62,2 |
| Q9WTN0 | Q9WTN0 | Geranylger | Ggps1 | sp\|Q9WTN | 8 | 8 | 29,7 |
| Q8BHK1 | Q8BHK1 | Magnesiur | Nipa1 | sp\|Q8BHK | 3 | 3 | 13,9 |
| Q91YJ2 | Q91YJ2 | Sorting ne: | Snx4 | sp\|Q91YJ2 | 13 | 13 | 39,6 |
| A2AWP8 | A2AWP8 | Rho guanir | Arhgef10l | sp\|A2AWP | 21 | 20 | 27 |
| Q9QXM1 | Q9QXM1 | Junction-m | Jmy | sp\|Q9QXM | 3 | 3 | 4,7 |
| O88413 | O88413 | Tubby-rela | Tulp3 | sp\|O88413 | 7 | 7 | 28,5 |
| Q920Q4 | Q920Q4 | Vacuolar p | Vps16 | sp\|Q920Q | 15 | 15 | 24,4 |
| Q3TYX3 | Q3TYX3 | SET and M | Smyd5 | sp\|Q3TYX3 | 10 | 10 | 31,5 |
| Q9D4G9 | Q9D4G9 | RecQ-med | Rmi1 | sp\|Q9D4G | 5 | 5 | 14,3 |
| Q8BKU8 | Q8BKU8 | Transmem | Tmem87b | sp\|Q8BKU | 2 | 2 | 4,7 |
| Q7TPH6 | Q7TPH6 | E3 ubiquiti | Mycbp2 | sp\|Q7TPH | 22 | 22 | 6,8 |
| Q9CXW4 | Q9CXW4 | 60S riboso | Rpl11 | sp\|Q9CXW | 9 | 9 | 52,8 |
| P15533 | P15533 | Tripartite r | Trim30a | sp\|P15533 | 4 | 4 | 11,9 |
| Q8VCD5 | Q8VCD5 | Mediator c | Med17 | sp\|Q8VCD | 16 | 16 | 36,2 |
| Q8R1G6 | Q8R1G6 | PDZ and LI | Pdlim2 | sp\|Q8R1G | 14 | 14 | 54,2 |
| Q922E6 | Q922E6 | FAST kinas | Fastkd2 | sp\|Q922E6 | 11 | 11 | 21,6 |
| O35075 | O35075 | Down synd | Dscr3 | sp\|O35075 | 8 | 8 | 39,1 |
| A2ALS5 | A2ALS5 | Rap1 GTPa | Rap1gap | sp\|A2ALS5 | 6 | 6 | 11,3 |
| Q99KK7 | Q99KK7 | Dipeptidyl | Dpp3 | sp\|Q99KK7 | 25 | 25 | 49,2 |
| Q99M08 | Q99M08 | Uncharacterized prote | | sp\|Q99M0 | 2 | 2 | 44,6 |
| Q9CQ40 | Q9CQ40 | 39S riboso | Mrpl49 | sp\|Q9CQ4 | 7 | 7 | 48,8 |
| Q91WZ8 | Q91WZ8 | Dysbindin | Dtnbp1 | sp\|Q91WZ | 3 | 3 | 9,4 |
| P35278 | P35278 | Ras-relatec | Rab5c | sp\|P35278 | 13 | 10 | 80,1 |
| Q8VCC8 | Q8VCC8 | Rap guanir | Rapgef3 | sp\|Q8VCC | 3 | 3 | 5,9 |
| Q8BP67 | Q8BP67 | 60S riboso | Rpl24 | sp\|Q8BP67 | 10 | 10 | 43,9 |
| Q8BGQ1 | Q8BGQ1 | Spermatog | Vipas39 | sp\|Q8BGQ | 7 | 7 | 22,4 |
| Q9CQV8 | Q9CQV8 | 14-3-3 pro | Ywhab | sp\|Q9CQV | 15 | 9 | 68,7 |
| P62315 | P62315 | Small nucle | Snrpd1 | sp\|P62315 | 5 | 5 | 54,6 |
| Q8VDP2 | Q8VDP2 | UPF0428 protein CXor | | sp\|Q8VDP | 6 | 6 | 35,1 |
| Q921N7 | Q921N7 | Transmem | Tmem70 | sp\|Q921N | 1 | 1 | 3,6 |
| O70589 | O70589 | Peripheral | Cask | sp\|O70589 | 15 | 15 | 24,7 |
| Q3UHG7 | Q3UHG7 | Protein LCI | Lchn | sp\|Q3UHG | 4 | 4 | 11,4 |
| Q60864 | Q60864 | Stress-indu | Stip1 | sp\|Q60864 | 44 | 44 | 64,8 |
| Q5RJ54 | Q5RJ54 | Zinc finger | Zscan26 | sp\|Q5RJ54 | 2 | 2 | 4,9 |
| Q8C1B7 | Q8C1B7 | Septin-11 | 11-set | sp\|Q8C1B | 20 | 11 | 49,7 |
| Q80TA1 | Q80TA1 | Ethanolam | Ept1 | sp\|Q80TA1 | 2 | 2 | 4,5 |
| Q9Z2D6 | Q9Z2D6 | Methyl-Cp | Mecp2 | sp\|Q9Z2D | 8 | 8 | 24,8 |
| Q9CQS8 | Q9CQS8 | Protein tra | Sec61b | sp\|Q9CQS | 3 | 3 | 37,5 |
| P63139 | P63139 | Nuclear tra | Nfyb | sp\|P63139 | 3 | 3 | 19,8 |
| Q8C1A5 | Q8C1A5 | Thimet olig | Thop1 | sp\|Q8C1A | 30 | 30 | 51,7 |
| O88545 | O88545 | COP9 signa | Cops6 | sp\|O88545 | 15 | 15 | 58,3 |
| Q8CHB8 | Q8CHB8 | Tubulin po | Ttll5 | sp\|Q8CHB | 2 | 2 | 2,5 |
| Q8BJQ9 | Q8BJQ9 | Chondroiti | Csgalnact1 | sp\|Q8BJQ | 3 | 2 | 7,2 |
| P59941 | P59941 | NAD-deper | Sirt6 | sp\|P59941 | 4 | 4 | 15 |

| | | | | | | | |
|---|---|---|---|---|---|---|---|
| Q6NZB0 | Q6NZB0 | DnaJ homo | Dnajc8 | sp\|Q6NZB0 | 11 | 11 | 45,8 |
| Q8BPZ8 | Q8BPZ8 | BRCA1-A c | Fam175a | sp\|Q8BPZ8 | 4 | 4 | 15 |
| P48024 | P48024 | Eukaryotic | Eif1 | sp\|P48024 | 11 | 4 | 78,8 |
| Q99JH8;Q9 | Q99JH8;Q9CQ | ER lumen p | Kdelr1;Kde | sp\|Q99JH8 | 2 | 2 | 14,2 |
| Q9JJE4 | Q9JJE4 | Progestin a | Paqr4 | sp\|Q9JJE4 | 2 | 2 | 8,1 |
| Q8CEC6 | Q8CEC6 | Peptidylpr | Ppwd1 | sp\|Q8CEC6 | 24 | 24 | 46,7 |
| P08113 | P08113 | Endoplasm | Hsp90b1 | sp\|P08113 | 56 | 54 | 64,2 |
| Q3U3I9 | Q3U3I9 | Zinc finger | Znf865 | sp\|Q3U3I9 | 3 | 3 | 3 |
| Q5RI75 | Q5RI75 | Ras and EF | Rasef | sp\|Q5RI75 | 2 | 2 | 4,1 |
| O54791 | O54791 | Transcripti | Maff | sp\|O54791 | 6 | 5 | 48,1 |
| Q6GV12 | Q6GV12 | 3-ketodihy | Kdsr | sp\|Q6GV1 | 8 | 8 | 33,7 |
| Q9QWT9 | Q9QWT9 | Kinesin-like | Kifc1 | sp\|Q9QWT | 20 | 20 | 36,5 |
| P0C090 | P0C090 | Roquin-2 | Rc3h2 | sp\|P0C090 | 5 | 2 | 5,3 |
| Q8BYN3 | Q8BYN3 | Inositol-te | Itpk1 | sp\|Q8BYN | 4 | 4 | 17,4 |
| Q6ZWV3;P | Q6ZWV3;P860 | 60S riboso | Rpl10;Rpl1 | sp\|Q6ZWV | 11 | 11 | 46,7 |
| Q8BXQ2 | Q8BXQ2 | GPI transa | Pigt | sp\|Q8BXQ | 10 | 10 | 19,8 |
| Q64191 | Q64191 | N(4)-(beta- | Aga | sp\|Q64191 | 6 | 6 | 24,3 |
| Q9CRD2 | Q9CRD2 | ER membra | Emc2 | sp\|Q9CRD | 9 | 9 | 36 |
| Q8VCE2 | Q8VCE2 | GPN-loop G | Gpn1 | sp\|Q8VCE2 | 8 | 8 | 33,6 |
| Q8BHX1 | Q8BHX1 | HAUS augm | Haus1 | sp\|Q8BHX | 8 | 8 | 36 |
| O35143 | O35143 | ATPase inh | Atpif1 | sp\|O35143 | 2 | 2 | 8,5 |
| Q8BMD6 | Q8BMD6 | Transmem | Tmem260 | sp\|Q8BMD | 2 | 2 | 4 |
| P60766 | P60766 | Cell divisio | Cdc42 | sp\|P60766 | 10 | 9 | 49,2 |
| Q6PGG2 | Q6PGG2 | GEM-intera | Gmip | sp\|Q6PGG | 6 | 6 | 8 |
| P0CG14 | P0CG14 | Chromoso | Chtf8 | sp\|P0CG14 | 5 | 5 | 15,6 |
| P50518;Q9 | P50518 | V-type pro | Atp6v1e1 | sp\|P50518 | 7 | 7 | 29,2 |
| Q9Z2U2 | Q9Z2U2 | Zinc finger | Zfp292 | sp\|Q9Z2U2 | 18 | 18 | 10,5 |
| Q91Y44 | Q91Y44 | Bromodom | Brdt | sp\|Q91Y44 | 4 | 2 | 3,6 |
| Q9JM96 | Q9JM96 | Cdc42 effe | Cdc42ep4 | sp\|Q9JM96 | 3 | 3 | 14,6 |
| E9Q6J5 | E9Q6J5 | | Bod1l | sp\|E9Q6J5 | 50 | 48 | 27,4 |
| Q9Z1G4 | Q9Z1G4 | V-type pro | Atp6v0a1 | sp\|Q9Z1G4 | 15 | 14 | 22,3 |
| Q91W78 | Q91W78 | U6 snRNA | Usb1 | sp\|Q91W7 | 2 | 2 | 4,9 |
| Q8K337 | Q8K337 | Type II ino | Inpp5b | sp\|Q8K337 | 15 | 15 | 20,5 |
| P62830 | P62830 | 60S riboso | Rpl23 | sp\|P62830 | 9 | 9 | 70,7 |
| P63280 | P63280 | SUMO-con | Ube2i | sp\|P63280 | 10 | 10 | 58,9 |
| Q6QI06 | Q6QI06 | Rapamycin | Rictor | sp\|Q6QI06 | 17 | 17 | 13,9 |
| Q6TEK5 | Q6TEK5 | Vitamin K e | Vkorc1l1 | sp\|Q6TEK5 | 4 | 4 | 15,3 |
| Q3U1C6 | Q3U1C6 | Putative de | Tatdn3 | sp\|Q3U1C | 5 | 5 | 25,2 |
| P09405 | P09405 | Nucleolin | Ncl | sp\|P09405 | 41 | 41 | 48,8 |
| Q8BTZ7 | Q8BTZ7 | Mannose-1 | Gmppb | sp\|Q8BTZ7 | 11 | 11 | 43,1 |
| Q9R210 | Q9R210 | Transcripti | Tfeb | sp\|Q9R210 | 2 | 1 | 5,7 |
| Q61127 | Q61127 | NGFI-A-bin | Nab2 | sp\|Q61127 | 8 | 7 | 22,9 |
| P02340 | P02340 | Cellular tu | Tp53 | sp\|P02340 | 5 | 5 | 17,9 |
| Q05D44 | Q05D44 | Eukaryotic | Eif5b | sp\|Q05D44 | 35 | 35 | 30,8 |
| Q99N57 | Q99N57 | RAF proto- | Raf1 | sp\|Q99N57 | 12 | 8 | 27 |
| Q9R1J0 | Q9R1J0 | Sterol-4-al | Nsdhl | sp\|Q9R1J0 | 13 | 13 | 48,6 |
| O88848 | O88848 | ADP-ribosy | Arl6 | sp\|O88848 | 11 | 11 | 68,3 |
| O88696 | O88696 | ATP-depen | Clpp | sp\|O88696 | 8 | 8 | 38,2 |
| Q14AI6 | Q14AI6 | RNA pseud | Rpusd3 | sp\|Q14AI6 | 5 | 5 | 19,8 |
| Q80SY4 | Q80SY4 | E3 ubiquiti | Mib1 | sp\|Q80SY4 | 6 | 6 | 10,1 |

| | | | | | | | |
|---|---|---|---|---|---|---|---|
| O35149 | O35149 | Zinc transp | Slc30a4 | sp\|O35149 | 2 | 2 | 10 |
| Q9Z2G1;Q | Q9Z2G1;Q8C0 | Protein fer | Fem1aa;Fe | sp\|Q9Z2G1 | 2 | 1 | 4,7 |
| Q9QZH3 | Q9QZH3 | Peptidyl-pr | Ppie | sp\|Q9QZH | 14 | 14 | 60,5 |
| Q8R1R3 | Q8R1R3 | StAR-relate | Stard7 | sp\|Q8R1R3 | 3 | 3 | 11 |
| Q8QZY6 | Q8QZY6 | Tetraspani | Tspan14 | sp\|Q8QZY6 | 3 | 3 | 14,4 |
| P35282 | P35282 | Ras-related | Rab21 | sp\|P35282 | 12 | 12 | 59,5 |
| P97348 | P97348 | Rho-relate | Rhod | sp\|P97348 | 4 | 4 | 34,3 |
| Q8VE92 | Q8VE92 | RNA-bindir | Rbm4b | sp\|Q8VE92 | 12 | 2 | 39,2 |
| Q69ZJ7 | Q69ZJ7 | RAB6A-GEI | Ric1 | sp\|Q69ZJ7 | 8 | 8 | 7,7 |
| P60060 | P60060 | Protein tra | Sec61g | sp\|P60060 | 2 | 2 | 36,8 |
| Q8BK30 | Q8BK30 | NADH deh | Ndufv3 | sp\|Q8BK30 | 1 | 1 | 16,3 |
| Q68FF6 | Q68FF6 | ARF GTPas | Git1 | sp\|Q68FF6 | 23 | 19 | 42,1 |
| Q6P9R4 | Q6P9R4 | Rho guanir | Arhgef18 | sp\|Q6P9R4 | 9 | 9 | 8,8 |
| Q4KMM3 | Q4KMM3 | Oxidation | Oxr1 | sp\|Q4KMN | 14 | 14 | 27,5 |
| Q99MS7 | Q99MS7 | EH domain | Ehbp1l1 | sp\|Q99MS | 27 | 27 | 24,7 |
| Q9JHN8 | Q9JHN8 | Serine/thre | Stk19 | sp\|Q9JHN8 | 5 | 5 | 25,2 |
| P21278;P3 | P21278 | Guanine nu | Gna11 | sp\|P21278 | 14 | 7 | 43,5 |
| Q9DB40 | Q9DB40 | Mediator c | Med27 | sp\|Q9DB40 | 8 | 8 | 45 |
| Q9D2X5 | Q9D2X5 | MAU2 chro | Mau2 | sp\|Q9D2X5 | 13 | 13 | 27,1 |
| Q9JK24 | Q9JK24 | Serine/thre | Ppp2r3c | sp\|Q9JK24 | 2 | 2 | 5,7 |
| Q8K0F1 | Q8K0F1 | TBC1 doma | Tbc1d23 | sp\|Q8K0F1 | 12 | 12 | 24,3 |
| Q9CWZ3 | Q9CWZ3 | RNA-bindir | Rbm8a | sp\|Q9CWZ | 9 | 9 | 71,3 |
| P97357 | P97357 | TATA box-b | Taf1a | sp\|P97357 | 6 | 6 | 14,8 |
| Q04899 | Q04899 | Cyclin-depe | Cdk18 | sp\|Q04899 | 6 | 4 | 18,6 |
| Q8BML1 | Q8BML1 | Protein-me | Mical2 | sp\|Q8BML | 7 | 6 | 10,2 |
| Q3TL26 | Q3TL26 | Dimethylad | Tfb2m | sp\|Q3TL26 | 6 | 6 | 20,7 |
| Q9D7J4 | Q9D7J4 | Cytochrom | Cox20 | sp\|Q9D7J4 | 3 | 3 | 23,1 |
| Q923Q2 | Q923Q2 | StAR-relate | Stard13 | sp\|Q923Q | 7 | 7 | 9,3 |
| Q8R2P1 | Q8R2P1 | Kelch-like | Klhl25 | sp\|Q8R2P1 | 3 | 3 | 5,6 |
| P26645 | P26645 | Myristoyla | Marcks | sp\|P26645 | 10 | 10 | 59,5 |
| P61460 | P61460 | DEP domai | Depdc5 | sp\|P61460 | 5 | 5 | 4,7 |
| Q6RI63 | Q6RI63 | Constitutiv | Fam120b | sp\|Q6RI63 | 6 | 6 | 10,4 |
| B2RQC2 | B2RQC2 | Ubiquitin c | Usp42 | sp\|B2RQC2 | 6 | 6 | 7,4 |
| Q9QZB7 | Q9QZB7 | Actin-relat | Actr10 | sp\|Q9QZB | 11 | 11 | 39,1 |
| Q99MK8 | Q99MK8 | Beta-adren | Adrbk1 | sp\|Q99MK | 13 | 8 | 22,6 |
| Q8R4E9 | Q8R4E9 | DNA replic | Cdt1 | sp\|Q8R4E9 | 7 | 7 | 20,1 |
| Q91XY4 | Q91XY4 | Protocadhe | Pcdhga4 | sp\|Q91XY4 | 2 | 2 | 3,4 |
| Q9ER00 | Q9ER00 | Syntaxin-1 | Stx12 | sp\|Q9ER00 | 6 | 6 | 31,8 |
| Q9CPT5 | Q9CPT5 | Nucleolar p | Nop16 | sp\|Q9CPT5 | 7 | 7 | 41 |
| A2BH40 | A2BH40 | AT-rich inte | Arid1a | sp\|A2BH40 | 35 | 33 | 25,2 |
| Q9ET22 | Q9ET22 | Dipeptidyl | Dpp7 | sp\|Q9ET22 | 4 | 4 | 10,5 |
| Q80ZK0 | Q80ZK0 | 28S riboso | Mrps10 | sp\|Q80ZK0 | 8 | 8 | 46,9 |
| Q8BL86 | Q8BL86 | Metallo-be | Mblac2 | sp\|Q8BL86 | 3 | 3 | 18,3 |
| Q6NVF0 | Q6NVF0 | Inositol po | Ocrl | sp\|Q6NVF0 | 9 | 9 | 15,4 |
| O55101 | O55101 | Synaptogy | Syngr2 | sp\|O55101 | 3 | 3 | 12,9 |
| Q8VC51 | Q8VC51 | Telomeras | Wrap53 | sp\|Q8VC51 | 6 | 6 | 16,2 |
| Q80TY5 | Q80TY5 | Vacuolar p | Vps13b | sp\|Q80TY5 | 5 | 5 | 1,5 |
| O70404 | O70404 | Vesicle-ass | Vamp8 | sp\|O70404 | 5 | 5 | 43,6 |
| Q64705 | Q64705 | Upstream s | Usf2 | sp\|Q64705 | 6 | 4 | 15 |
| P62313 | P62313 | U6 snRNA- | Lsm6 | sp\|P62313 | 5 | 5 | 55 |

| | | | | | | | |
|---|---|---|---|---|---|---|---|
| Q9D5R2 | Q9D5R2 | WD repeat | Wdr20 | sp\|Q9D5R2 | 6 | 6 | 11,8 |
| P51175 | P51175 | Protoporpl | Ppox | sp\|P51175 | 13 | 13 | 43,8 |
| Q9R1P0 | Q9R1P0 | Proteasom | Psma4 | sp\|Q9R1P0 | 12 | 12 | 68,2 |
| P21619 | P21619 | Lamin-B2 | Lmnb2 | sp\|P21619 | 20 | 18 | 43,6 |
| Q3UXZ9 | Q3UXZ9 | Lysine-spe | Kdm5a | sp\|Q3UXZ9 | 24 | 21 | 19,7 |
| O54825 | O54825 | Bystin | Bysl | sp\|O54825 | 17 | 17 | 47,2 |
| Q8VE99 | Q8VE99 | Coiled-coil | Ccdc115 | sp\|Q8VE99 | 8 | 8 | 48,9 |
| Q99JP7 | Q99JP7 | Gamma-glu | Ggt7 | sp\|Q99JP7 | 2 | 2 | 3,6 |
| Q69Z98 | Q69Z98 | Serine/thre | Brsk2 | sp\|Q69Z98 | 1 | 1 | 1,6 |
| P29341 | P29341 | Polyadenyl | Pabpc1 | sp\|P29341 | 38 | 38 | 67,8 |
| Q80WC1 | Q80WC1 | Ubinuclein | Ubn2 | sp\|Q80WC | 5 | 5 | 7,4 |
| Q5EE38 | Q5EE38 | Adrenocor | Acd | sp\|Q5EE38 | 4 | 4 | 20,2 |
| Q8BZR9 | Q8BZR9 | Uncharacterized prote | | sp\|Q8BZR9 | 12 | 12 | 24,7 |
| O35972 | O35972 | 39S riboso | Mrpl23 | sp\|O35972 | 7 | 7 | 51,4 |
| Q9CQV7 | Q9CQV7 | Mitochond | Dnajc19 | sp\|Q9CQV | 4 | 4 | 43,1 |
| Q9D7M8 | Q9D7M8 | DNA-direct | Polr2d | sp\|Q9D7M | 8 | 8 | 69 |
| Q91YJ3 | Q91YJ3 | Thymocyte | Thyn1 | sp\|Q91YJ3 | 10 | 10 | 36,7 |
| Q9D1J3 | Q9D1J3 | SAP domai | Sarnp | sp\|Q9D1J3 | 9 | 9 | 39,5 |
| Q8CFT2 | Q8CFT2 | Histone-lys | Setd1b | sp\|Q8CFT2 | 8 | 7 | 6,3 |
| Q3UPE3 | Q3UPE3 | RecQ-med | Rmi2 | sp\|Q3UPE3 | 1 | 1 | 11,4 |
| P22366 | P22366 | Myeloid di | Myd88 | sp\|P22366 | 7 | 7 | 27,4 |
| Q8CC21 | Q8CC21 | Tetratricop | Ttc19 | sp\|Q8CC21 | 2 | 2 | 7,1 |
| P35293 | P35293 | Ras-related | Rab18 | sp\|P35293 | 13 | 13 | 71,4 |
| Q922B2 | Q922B2 | Aspartate- | Dars | sp\|Q922B2 | 35 | 35 | 69,1 |
| Q6PIU9 | Q6PIU9 | Uncharacterized prote | | sp\|Q6PIU9 | 9 | 9 | 40,4 |
| Q921W0 | Q921W0 | Charged m | Chmp1a | sp\|Q921W | 3 | 3 | 13,3 |
| P23492 | P23492 | Purine nuc | Pnp | sp\|P23492 | 15 | 15 | 69,2 |
| Q7TQI3 | Q7TQI3 | Ubiquitin t | Otub1 | sp\|Q7TQI3 | 12 | 12 | 53,1 |
| Q8BH66 | Q8BH66 | Atlastin-1 | Atl1 | sp\|Q8BH66 | 5 | 3 | 9,9 |
| Q8BGR2 | Q8BGR2 | Volume-re | Lrrc8d | sp\|Q8BGR | 22 | 22 | 32,9 |
| Q9Z2Q5 | Q9Z2Q5 | 39S riboso | Mrpl40 | sp\|Q9Z2Q | 7 | 7 | 37,9 |
| Q9QZB9 | Q9QZB9 | Dynactin s | Dctn5 | sp\|Q9QZB | 4 | 4 | 23,1 |
| Q7TSY8 | Q7TSY8 | Shugoshin- | Sgol2 | sp\|Q7TSY8 | 14 | 14 | 21,2 |
| Q8VC30 | Q8VC30 | Bifunction | Dak | sp\|Q8VC30 | 15 | 15 | 39,4 |
| Q4VBD2 | Q4VBD2 | Transmem | Tapt1 | sp\|Q4VBD | 12 | 12 | 35,1 |
| Q8CID0 | Q8CID0 | Cysteine-ri | Csrp2bp | sp\|Q8CID0 | 4 | 4 | 5,8 |
| Q4KML4 | Q4KML4 | Costars far | Abracl | sp\|Q4KML | 3 | 3 | 46,9 |
| Q76KJ5 | Q76KJ5 | DNA-direct | Cd3eap | sp\|Q76KJ5 | 10 | 10 | 37,8 |
| P25916 | P25916 | Polycomb | Bmi1 | sp\|P25916 | 5 | 3 | 16,7 |
| Q8BJQ2 | Q8BJQ2 | Ubiquitin c | Usp1 | sp\|Q8BJQ2 | 9 | 9 | 15,1 |
| Q811J3 | Q811J3 | Iron-respo | Ireb2 | sp\|Q811J3 | 9 | 8 | 15,3 |
| Q9D0R9 | Q9D0R9 | WD repeat | Wdr89 | sp\|Q9D0R9 | 10 | 10 | 37 |
| Q80WQ9 | Q80WQ9 | Zinc finger | Zbed4 | sp\|Q80WQ | 1 | 1 | 1,2 |
| Q9CPQ3 | Q9CPQ3 | Mitochond | Tomm22 | sp\|Q9CPQ | 7 | 7 | 76,8 |
| Q80XK6 | Q80XK6 | Autophagy | Atg2b | sp\|Q80XK6 | 19 | 19 | 11,8 |
| Q9R0L7 | Q9R0L7 | A-kinase ar | Akap8l | sp\|Q9R0L7 | 9 | 9 | 19 |
| P28231 | P28231 | Gap juncti | Gjb3 | sp\|P28231 | 4 | 4 | 23 |
| Q8BH73 | Q8BH73 | Glutaminyl | Qpctl | sp\|Q8BH73 | 2 | 2 | 5,5 |
| Q99J64 | Q99J64 | RNA-bindir | Rbm43 | sp\|Q99J64 | 3 | 3 | 11,1 |
| Q9JKN1 | Q9JKN1 | Zinc transp | Slc30a7 | sp\|Q9JKN1 | 5 | 5 | 15,9 |

| | | | | | | | |
|---|---|---|---|---|---|---|---|
| Q9D902 | Q9D902 | General tra | Gtf2e2 | sp\|Q9D902 | 10 | 10 | 44,5 |
| P63276 | P63276 | 40S riboso | Rps17 | sp\|P63276 | 12 | 12 | 60 |
| Q65Z40 | Q65Z40 | Wings apar | Wapal | sp\|Q65Z40 | 30 | 30 | 34,9 |
| O70126 | O70126 | Aurora kin | Aurkb | sp\|O70126 | 14 | 13 | 47,2 |
| Q8CH25 | Q8CH25 | SAFB-like t | Sltm | sp\|Q8CH25 | 19 | 19 | 21 |
| Q9D2G2 | Q9D2G2 | Dihydrolip | Dlst | sp\|Q9D2G | 16 | 16 | 45,8 |
| Q99P69 | Q99P69 | Kinetochor | Nuf2 | sp\|Q99P69 | 12 | 12 | 23,8 |
| Q3UWW6 | Q3UWW6 | GAS2-like p | Gas2l3 | sp\|Q3UWV | 6 | 6 | 11,1 |
| Q9CQD1 | Q9CQD1 | Ras-relatec | Rab5a | sp\|Q9CQD | 10 | 7 | 72,1 |
| Q6P1E1;Q8 | Q6P1E1 | Zinc finger | Zmiz1 | sp\|Q6P1E1 | 9 | 9 | 11,2 |
| Q8VE37 | Q8VE37 | Regulator c | Rcc1 | sp\|Q8VE37 | 17 | 17 | 64,1 |
| A2AH22 | A2AH22 | Activating | Ambra1 | sp\|A2AH22 | 7 | 7 | 7,5 |
| Q9ERV1 | Q9ERV1 | Probable E | Mkrn2 | sp\|Q9ERV1 | 4 | 4 | 16,1 |
| Q99PP7 | Q99PP7 | E3 ubiquiti | Trim33 | sp\|Q99PP7 | 27 | 27 | 33,6 |
| Q8R107 | Q8R107 | PRELI dom | Prelid1 | sp\|Q8R107 | 3 | 3 | 14,7 |
| Q8R015 | Q8R015 | Biogenesis | Bloc1s5 | sp\|Q8R015 | 5 | 5 | 36,2 |
| Q8R081 | Q8R081 | Heterogen | Hnrnpl | sp\|Q8R081 | 27 | 27 | 61,3 |
| Q9Z266 | Q9Z266 | SNARE-ass | Snapin | sp\|Q9Z266 | 4 | 4 | 41,9 |
| P97784;Q9 | P97784;Q9R19 | Cryptochro | Cry1;Cry2 | sp\|P97784 | 3 | 3 | 5,4 |
| P97929 | P97929 | Breast can | Brca2 | sp\|P97929 | 3 | 3 | 1,3 |
| Q3UR97 | Q3UR97 | | Snx21 | sp\|Q3UR9 | 1 | 1 | 4,4 |
| Q9EP52 | Q9EP52 | Twisted ga | Twsg1 | sp\|Q9EP52 | 3 | 3 | 24,8 |
| Q6PEB6 | Q6PEB6 | MOB-like p | Mob4 | sp\|Q6PEB6 | 7 | 7 | 49,3 |
| Q9JMH7 | Q9JMH7 | Sialidase-3 | Neu3 | sp\|Q9JMH | 2 | 2 | 5,5 |
| Q8K2C9 | Q8K2C9 | Very-long-c | Hacd3 | sp\|Q8K2C9 | 8 | 8 | 30,7 |
| P61164 | P61164 | Alpha-cent | Actr1a | sp\|P61164 | 13 | 7 | 52,7 |
| O88942 | O88942 | Nuclear fac | Nfatc1 | sp\|O88942 | 8 | 8 | 18,5 |
| Q5U5M8 | Q5U5M8 | Biogenesis | Bloc1s3 | sp\|Q5U5M | 2 | 2 | 17,4 |
| Q5RL51 | Q5RL51 | Glutathion | Gstcd | sp\|Q5RL51 | 7 | 7 | 19,1 |
| O88455 | O88455 | 7-dehydro | Dhcr7 | sp\|O88455 | 5 | 5 | 14 |
| Q8BMG8 | Q8BMG8 | Mitochond | Slc25a32 | sp\|Q8BMG | 6 | 6 | 23,1 |
| Q03963 | Q03963 | Interferon- | Eif2ak2 | sp\|Q03963 | 15 | 15 | 28,7 |
| Q9DB30 | Q9DB30 | Phosphory | Phkg2 | sp\|Q9DB30 | 3 | 3 | 12,1 |
| Q8BK75 | Q8BK75 | Elongator c | Elp6 | sp\|Q8BK75 | 7 | 7 | 26,3 |
| O88384 | O88384 | Vesicle tra | Vti1b | sp\|O88384 | 6 | 6 | 31,5 |
| Q6ZQ06 | Q6ZQ06 | Centrosom | Cep162 | sp\|Q6ZQ06 | 7 | 7 | 5,7 |
| Q5KU39 | Q5KU39 | Vacuolar p | Vps41 | sp\|Q5KU39 | 8 | 8 | 11,7 |
| Q80WC7 | Q80WC7 | Arf-GAP do | Agfg2 | sp\|Q80WC | 4 | 4 | 14 |
| Q6P7W0 | Q6P7W0 | Sentrin-spe | Senp6 | sp\|Q6P7W | 4 | 4 | 4,5 |
| O70172 | O70172 | Phosphatid | Pip4k2a | sp\|O70172 | 8 | 4 | 23,7 |
| P56213 | P56213 | FAD-linked | Gfer | sp\|P56213 | 7 | 7 | 31,3 |
| P63001;P6 | P63001;P6076 | Ras-relatec | Rac1;Rac3; | sp\|P63001 | 9 | 8 | 50 |
| Q99KR8 | Q99KR8 | Plasma alp | Fuca2 | sp\|Q99KR8 | 3 | 3 | 8,2 |
| Q78T54 | Q78T54 | Vacuolar A | Vma21 | sp\|Q78T54 | 3 | 3 | 34,7 |
| Q3UTQ7 | Q3UTQ7 | PR domain | Prdm10 | sp\|Q3UTQ | 4 | 4 | 4,5 |
| Q9D9V3 | Q9D9V3 | Ethylmalor | Echdc1 | sp\|Q9D9V | 7 | 7 | 26,7 |
| P97429 | P97429 | Annexin A4 | Anxa4 | sp\|P97429 | 23 | 22 | 72,4 |
| Q9DC60 | Q9DC60 | UbiA preny | Ubiad1 | sp\|Q9DC60 | 2 | 2 | 5,7 |
| Q8R151 | Q8R151 | NFX1-type | Znfx1 | sp\|Q8R151 | 20 | 20 | 13,9 |
| P38647 | P38647 | Stress-70 p | Hspa9 | sp\|P38647 | 48 | 47 | 68,2 |

| | | | | | | | |
|---|---|---|---|---|---|---|---|
| Q6PGG6 | Q6PGG6 | Guanine nu | Gnl3l | sp\|Q6PGG | 11 | 11 | 26,5 |
| Q3UHX2 | Q3UHX2 | 28 kDa hea | Pdap1 | sp\|Q3UHX | 7 | 7 | 28,2 |
| Q6P8I6 | Q6P8I6 | Cytochrom | Cox11 | sp\|Q6P8I6 | 3 | 3 | 21,8 |
| Q03137 | Q03137 | Ephrin type | Epha4 | sp\|Q03137 | 6 | 4 | 7,8 |
| Q9CSU0 | Q9CSU0 | Regulation | Rprd1b | sp\|Q9CSU0 | 12 | 10 | 48,5 |
| Q64213 | Q64213 | Splicing fac | Sf1 | sp\|Q64213 | 17 | 17 | 36,4 |
| Q91WK7 | Q91WK7 | Ankyrin re | Ankrd54 | sp\|Q91WK | 3 | 3 | 22,4 |
| O08734 | O08734 | Bcl-2 homo | Bak1 | sp\|O08734 | 6 | 6 | 51,7 |
| P80317;Q6 | P80317 | T-complex | Cct6a | sp\|P80317 | 25 | 25 | 53,1 |
| Q61586 | Q61586 | Glycerol-3- | Gpam | sp\|Q61586 | 10 | 10 | 16,8 |
| Q9QXN0 | Q9QXN0 | Protein Shi | Shroom3 | sp\|Q9QXN | 2 | 2 | 1,5 |
| Q9JKR6 | Q9JKR6 | Hypoxia up | Hyou1 | sp\|Q9JKR6 | 37 | 37 | 45,5 |
| Q91W52 | Q91W52 | Transmem | Tmem19 | sp\|Q91W5 | 2 | 2 | 7,4 |
| Q6NXK5 | Q6NXK5 | RNA/RNP c | Dusp11 | sp\|Q6NXK | 4 | 4 | 19,9 |
| Q8BGC1 | Q8BGC1 | UPF0489 protein C5or | | sp\|Q8BGC | 3 | 3 | 10,2 |
| Q91WQ5 | Q91WQ5 | TAF5-like F | Taf5l | sp\|Q91WQ | 12 | 12 | 26,8 |
| P99024 | P99024 | Tubulin be | Tubb5 | sp\|P99024 | 27 | 4 | 77,9 |
| Q8BTU1 | Q8BTU1 | Cilia- and f | Cfap20 | sp\|Q8BTU | 8 | 8 | 55,4 |
| Q9CPV4 | Q9CPV4 | Glyoxalase | Glod4 | sp\|Q9CPV4 | 17 | 17 | 60,7 |
| Q8BX09 | Q8BX09 | Retinoblas | Rbbp5 | sp\|Q8BX09 | 13 | 13 | 33,5 |
| Q921H9 | Q921H9 | Cytochrom | Coa7 | sp\|Q921H9 | 6 | 6 | 39 |
| Q8CCF0 | Q8CCF0 | U4/U6 sma | Prpf31 | sp\|Q8CCF0 | 17 | 17 | 46,5 |
| Q61191 | Q61191 | Host cell fa | Hcfc1 | sp\|Q61191 | 53 | 52 | 36,8 |
| Q9Z0Y1 | Q9Z0Y1 | Dynactin s | Dctn3 | sp\|Q9Z0Y1 | 6 | 6 | 39,2 |
| Q8R404 | Q8R404 | Protein QIl | Qil1 | sp\|Q8R404 | 3 | 3 | 41,2 |
| Q8C8N2 | Q8C8N2 | Protein SC | Scai | sp\|Q8C8N | 3 | 3 | 7,1 |
| P59999 | P59999 | Actin-relat | Arpc4 | sp\|P59999 | 9 | 9 | 63,1 |
| Q8CDD9 | Q8CDD9 | Ligand-dep | Lrif1 | sp\|Q8CDD | 3 | 3 | 6,9 |
| Q80X32 | Q80X32 | UPF0461 protein C5or | | sp\|Q80X32 | 4 | 4 | 29,3 |
| Q9D9M5 | Q9D9M5 | Pyridoxal p | Phospho2 | sp\|Q9D9M | 2 | 2 | 11,6 |
| Q8BKJ9 | Q8BKJ9 | NAD-depe | Sirt7 | sp\|Q8BKJ9 | 4 | 4 | 13,9 |
| Q9D0S9 | Q9D0S9 | Histidine t | Hint2 | sp\|Q9D0S9 | 4 | 4 | 39,9 |
| Q9WTQ8 | Q9WTQ8 | Mitochond | Timm23 | sp\|Q9WTQ | 4 | 4 | 26,3 |
| Q7TSV4 | Q7TSV4 | Phosphogl | Pgm2 | sp\|Q7TSV4 | 24 | 24 | 45,6 |
| Q91W69 | Q91W69 | Epsin-3 | Epn3 | sp\|Q91W6 | 4 | 2 | 6,1 |
| P98199 | P98199 | Phospholip | Atp8b2 | sp\|P98199 | 3 | 3 | 2,5 |
| Q9JJF9 | Q9JJF9 | Signal pept | Sppl2a | sp\|Q9JJF9 | 4 | 4 | 10,7 |
| P68040 | P68040 | Guanine nu | Gnb2l1 | sp\|P68040 | 21 | 21 | 79,2 |
| Q9D0F6 | Q9D0F6 | Replication | Rfc5 | sp\|Q9D0F6 | 15 | 15 | 54 |
| Q8CC12 | Q8CC12 | Codanin-1 | Cdan1 | sp\|Q8CC12 | 7 | 7 | 8,9 |
| Q61122 | Q61122 | NGFI-A-bin | Nab1 | sp\|Q61122 | 5 | 4 | 15,2 |
| Q9D8T4 | Q9D8T4 | Golgi appa | Tvp23b | sp\|Q9D8T4 | 2 | 2 | 19 |
| P97872 | P97872 | Dimethylar | Fmo5 | sp\|P97872 | 3 | 3 | 7,7 |
| O88531 | O88531 | Palmitoyl-p | Ppt1 | sp\|O88531 | 7 | 7 | 29,1 |
| P53994 | P53994 | Ras-related | Rab2a | sp\|P53994 | 15 | 7 | 70,8 |
| Q60961 | Q60961 | Lysosomal- | Laptm4a | sp\|Q60961 | 2 | 2 | 17,2 |
| Q64430;Q6 | Q64430 | Copper-tra | Atp7a | sp\|Q64430 | 12 | 12 | 12,3 |
| Q8R123 | Q8R123 | FAD syntha | Flad1 | sp\|Q8R123 | 13 | 13 | 38,2 |
| Q7TSG3 | Q7TSG3 | F-box only | Fbxo5 | sp\|Q7TSG3 | 2 | 2 | 5 |
| P47963 | P47963 | 60S riboso | Rpl13 | sp\|P47963 | 12 | 12 | 49,8 |

| | | | | | | | |
|---|---|---|---|---|---|---|---|
| Q9D0M0 | Q9D0M0 | Exosome c | Exosc7 | sp\|Q9D0M | 11 | 11 | 52,9 |
| Q63932 | Q63932 | Dual specif | Map2k2 | sp\|Q63932 | 15 | 10 | 46,6 |
| Q8CES0 | Q8CES0 | N-alpha-ac | Naa30 | sp\|Q8CES0 | 5 | 5 | 15,4 |
| Q7TQI7 | Q7TQI7 | Ankyrin re | Abtb2 | sp\|Q7TQI7 | 2 | 2 | 3,2 |
| Q8BZ21 | Q8BZ21 | Histone ac | Kat6a | sp\|Q8BZ21 | 10 | 8 | 6,9 |
| Q8BZW8 | Q8BZW8 | NHL repea | Nhlrc2 | sp\|Q8BZW | 15 | 15 | 40,4 |
| Q8BRK9 | Q8BRK9 | Alpha-man | Man2a2 | sp\|Q8BRK9 | 3 | 3 | 4,2 |
| P70362 | P70362 | Ubiquitin f | Ufd1l | sp\|P70362 | 13 | 13 | 48,2 |
| Q3TB48 | Q3TB48 | Transmem | Tmem104 | sp\|Q3TB48 | 3 | 3 | 5,8 |
| O54774 | O54774 | AP-3 comp | Ap3d1 | sp\|O54774 | 34 | 34 | 37,9 |
| Q9CQ02 | Q9CQ02 | COMM do | Commd4 | sp\|Q9CQ0 | 5 | 5 | 27,1 |
| Q9JHI2 | Q9JHI2 | tRNA-speci | Adat1 | sp\|Q9JHI2 | 7 | 7 | 19,8 |
| Q3UGC7;Q | Q3UGC7;Q66J | Eukaryotic | Eif3j1;Eif3j | sp\|Q3UGC | 13 | 13 | 48,3 |
| Q80WQ8 | Q80WQ8 | Mis18-bind | Mis18bp1 | sp\|Q80WC | 12 | 12 | 16,8 |
| O35387 | O35387 | HCLS1-ass | Hax1 | sp\|O35387 | 9 | 9 | 52,5 |
| Q99104 | Q99104 | Unconvent | Myo5a | sp\|Q99104 | 53 | 50 | 33 |
| Q62276 | Q62276 | Mediator c | Med22 | sp\|Q62276 | 3 | 3 | 19 |
| Q80X90 | Q80X90 | Filamin-B | Flnb | sp\|Q80X90 | 140 | 130 | 72,9 |
| Q9D3W4 | Q9D3W4 | GPN-loop G | Gpn3 | sp\|Q9D3W | 6 | 6 | 34,5 |
| Q9ERS2 | Q9ERS2 | NADH deh | Ndufa13 | sp\|Q9ERS2 | 7 | 7 | 43,1 |
| Q99LT0 | Q99LT0 | Protein dp | Dpy30 | sp\|Q99LT0 | 2 | 2 | 38,4 |
| Q8R3A2 | Q8R3A2 | Cysteine-ri | Cdpf1 | sp\|Q8R3A2 | 1 | 1 | 11,8 |
| Q9CQW0 | Q9CQW0 | ER membr | Emc6 | sp\|Q9CQW | 3 | 3 | 20,9 |
| Q7TN29 | Q7TN29 | Stromal m | Smap2 | sp\|Q7TN29 | 4 | 4 | 11,2 |
| Q99LQ4 | Q99LQ4 | Coiled-coil | Ccdc23 | sp\|Q99LQ4 | 1 | 1 | 22,7 |
| P70295 | P70295 | Ancient ub | Aup1 | sp\|P70295 | 8 | 8 | 26,1 |
| Q61161 | Q61161 | Mitogen-a | Map4k2 | sp\|Q61161 | 5 | 5 | 9,4 |
| P40124;Q9 | P40124 | Adenylyl cy | Cap1 | sp\|P40124 | 26 | 26 | 64,3 |
| Q9DB77 | Q9DB77 | Cytochrom | Uqcrc2 | sp\|Q9DB77 | 19 | 19 | 49,2 |
| Q3TIX9 | Q3TIX9 | U4/U6.U5 | Usp39 | sp\|Q3TIX9 | 19 | 19 | 43,3 |
| Q9D6M3;Q | Q9D6M3 | Mitochond | Slc25a22 | sp\|Q9D6M | 6 | 6 | 19,2 |
| Q8VBT0 | Q8VBT0 | Thioredoxi | Tmx1 | sp\|Q8VBT0 | 10 | 10 | 25,2 |
| Q8BGB8 | Q8BGB8 | Ubiquinone | Coq4 | sp\|Q8BGB | 6 | 6 | 23,3 |
| Q8K2H2 | Q8K2H2 | OTU doma | Otud6b | sp\|Q8K2H2 | 10 | 10 | 53,1 |
| Q3TBW2 | Q3TBW2 | 39S riboso | Mrpl10 | sp\|Q3TBW | 8 | 8 | 46,2 |
| Q5SUC9 | Q5SUC9 | Protein SC | Sco1 | sp\|Q5SUC9 | 5 | 5 | 20,1 |
| Q8BKW4 | Q8BKW4 | Zinc finger | Zcchc4 | sp\|Q8BKW | 7 | 7 | 15,6 |
| Q8VEG6 | Q8VEG6 | CCR4-NOT | Cnot6l | sp\|Q8VEG | 4 | 4 | 8,3 |
| Q91VR8 | Q91VR8 | Protein BR | Brk1 | sp\|Q91VR8 | 2 | 2 | 32 |
| Q921F2 | Q921F2 | TAR DNA-b | Tardbp | sp\|Q921F2 | 17 | 17 | 56 |
| Q9JIX0 | Q9JIX0 | Transcripti | Eny2 | sp\|Q9JIX0 | 6 | 6 | 46,5 |
| Q99KB8 | Q99KB8 | Hydroxyac | Hagh | sp\|Q99KB8 | 8 | 8 | 38,2 |
| Q9CR46 | Q9CR46 | Spindle an | Ska2 | sp\|Q9CR46 | 1 | 1 | 9,2 |
| Q8BTW3 | Q8BTW3 | Exosome c | Exosc6 | sp\|Q8BTW | 8 | 8 | 32,6 |
| Q6NZM9 | Q6NZM9 | Histone de | Hdac4 | sp\|Q6NZM | 7 | 7 | 10,1 |
| Q9WVD4 | Q9WVD4 | H(+)/Cl(-) e | Clcn5 | sp\|Q9WVD | 4 | 3 | 6,3 |
| Q69ZN6 | Q69ZN6 | N-acetylglu | Gnptab | sp\|Q69ZN6 | 4 | 4 | 3,9 |
| Q6IQX7 | Q6IQX7 | Chondroiti | Chpf | sp\|Q6IQX7 | 7 | 7 | 11,4 |
| P61079;P6 | P61079;P6283 | Ubiquitin-c | Ube2d3;Ub | sp\|P61079 | 4 | 4 | 40,8 |
| Q9JM13 | Q9JM13 | Rab5 GDP/ | Rabgef1 | sp\|Q9JM1 | 5 | 5 | 13,6 |

| | | | | | | | |
|---|---|---|---|---|---|---|---|
| Q80UY2 | Q80UY2 | E3 ubiquiti | Kcmf1 | sp\|Q80UY | 3 | 3 | 11,5 |
| P22518 | P22518 | Dual specif | Clk1 | sp\|P22518 | 8 | 4 | 22,6 |
| Q9CQ90 | Q9CQ90 | Uncharacterized prot | | sp\|Q9CQ9 | 3 | 3 | 23,9 |
| Q8CGC4 | Q8CGC4 | Protein LSI | Lsm14b | sp\|Q8CGC | 15 | 15 | 49,4 |
| Q6P2K6 | Q6P2K6 | Serine/thre | Smek1 | sp\|Q6P2K | 30 | 23 | 45 |
| Q9Z2L7 | Q9Z2L7 | Cytokine re | Crlf3 | sp\|Q9Z2L7 | 8 | 8 | 30,8 |
| O54967 | O54967 | Activated | Tnk2 | sp\|O5496 | 9 | 9 | 14,5 |
| Q61510 | Q61510 | E3 ubiquiti | Trim25 | sp\|Q6151 | 26 | 26 | 49,8 |
| Q8C196 | Q8C196 | Carbamoyl | Cps1 | sp\|Q8C19 | 10 | 8 | 9 |
| Q61789 | Q61789 | Laminin su | Lama3 | sp\|Q6178 | 4 | 4 | 1,6 |
| Q9WTU6 | Q9WTU6 | Mitogen-a | Mapk9 | sp\|Q9WTU | 5 | 3 | 17,3 |
| Q80W68 | Q80W68 | Kin of IRRE | Kirrel | sp\|Q80W6 | 17 | 17 | 36,1 |
| Q0VGT4 | Q0VGT4 | Protein ZG | Zgrf1 | sp\|Q0VGT | 2 | 2 | 1,4 |
| Q8BKH7 | Q8BKH7 | Target of r | Mapkap1 | sp\|Q8BKH | 6 | 6 | 17,6 |
| Q8VEF1 | Q8VEF1 | GRAM don | Gramd1a | sp\|Q8VEF1 | 5 | 5 | 13,9 |
| P63038 | P63038 | 60 kDa hea | Hspd1 | sp\|P63038 | 48 | 48 | 77,5 |
| Q8VH37 | Q8VH37 | Histone de | Hdac8 | sp\|Q8VH3 | 2 | 2 | 6,9 |
| A8Y5H7 | A8Y5H7 | | Sec14l1 | sp\|A8Y5H7 | 12 | 12 | 21 |
| O35386 | O35386 | Phytanoyl- | Phyh | sp\|O3538 | 4 | 4 | 20,1 |
| Q9CY73 | Q9CY73 | 39S riboso | Mrpl44 | sp\|Q9CY7 | 11 | 11 | 35,7 |
| Q8BH57 | Q8BH57 | WD repeat | Wdr48 | sp\|Q8BH5 | 18 | 18 | 34,8 |
| Q9JHZ2 | Q9JHZ2 | Progressive | Ankh | sp\|Q9JHZ2 | 7 | 7 | 16,7 |
| Q91VL8 | Q91VL8 | Telomeric | Terf2ip | sp\|Q91VL8 | 9 | 9 | 35,6 |
| Q8R2N2 | Q8R2N2 | Cirhin | Cirh1a | sp\|Q8R2N | 23 | 23 | 42,6 |
| Q91V57 | Q91V57 | N-chimaer | Chn1 | sp\|Q91V5 | 2 | 2 | 5,7 |
| Q8BTW9;C | Q8BTW9 | Serine/thre | Pak4 | sp\|Q8BTW | 10 | 10 | 26,3 |
| Q8BXA1 | Q8BXA1 | Golgi integ | Golim4 | sp\|Q8BXA | 9 | 9 | 18,8 |
| Q9CPW9 | Q9CPW9 | Methionin | Metap1d | sp\|Q9CPW | 3 | 3 | 14 |
| P15327 | P15327 | Bisphosph | Bpgm | sp\|P15327 | 1 | 1 | 7,3 |
| P32507 | P32507 | Nectin-2 | Pvrl2 | sp\|P32507 | 9 | 9 | 32,6 |
| Q8WTY4 | Q8WTY4 | Anamorsin | Ciapin1 | sp\|Q8WTY | 15 | 15 | 69,9 |
| Q61699 | Q61699 | Heat shock | Hsph1 | sp\|Q6169 | 40 | 38 | 62,9 |
| Q8BRN9 | Q8BRN9 | Coiled-coil | Cc2d1b | sp\|Q8BRN | 22 | 22 | 40,8 |
| Q78YZ6 | Q78YZ6 | Short coile | Scoc | sp\|Q78YZ6 | 1 | 1 | 14,4 |
| Q9R1Q7 | Q9R1Q7 | Proteolipid | Plp2 | sp\|Q9R1Q | 2 | 2 | 17,8 |
| Q8VC66 | Q8VC66 | Afadin- an | Ssx2ip | sp\|Q8VC6 | 3 | 3 | 7,3 |
| Q9JKV1 | Q9JKV1 | Proteasom | Adrm1 | sp\|Q9JKV1 | 7 | 7 | 15 |
| O35594 | O35594 | Intraflagell | Ift81 | sp\|O3559 | 5 | 5 | 9,5 |
| P62257 | P62257 | Ubiquitin-c | Ube2h | sp\|P62257 | 4 | 4 | 26,8 |
| O54998 | O54998 | Peptidyl-p | Fkbp7 | sp\|O5499 | 4 | 4 | 21,1 |
| O88851 | O88851 | Putative hy | Rbbp9 | sp\|O8885 | 8 | 8 | 74,2 |
| Q9DBN9 | Q9DBN9 | Probable A | Ddx59 | sp\|Q9DBN | 4 | 4 | 8,4 |
| Q61768;P2 | Q61768 | Kinesin-1 h | Kif5b | sp\|Q6176 | 51 | 43 | 58,7 |
| Q9JIH2 | Q9JIH2 | Nuclear po | Nup50 | sp\|Q9JIH2 | 19 | 19 | 54,1 |
| Q99N94 | Q99N94 | 39S riboso | Mrpl9 | sp\|Q99N9 | 9 | 9 | 38,1 |
| Q9Z2A7 | Q9Z2A7 | Diacylglyce | Dgat1 | sp\|Q9Z2A | 3 | 3 | 10,4 |
| Q64520 | Q64520 | Guanylate | Guk1 | sp\|Q6452 | 8 | 8 | 56,6 |
| P61166 | P61166 | Transmem | Tmem258 | sp\|P61166 | 1 | 1 | 10,1 |
| Q6KAU8 | Q6KAU8 | Autophagy | Atg16l2 | sp\|Q6KAU | 1 | 1 | 2,9 |
| Q8BGY4 | Q8BGY4 | Kelch-like | Klhl26 | sp\|Q8BGY | 3 | 3 | 7,3 |

| | | | | | | | |
|---|---|---|---|---|---|---|---|
| O08908 | O08908 | Phosphatid | Pik3r2 | sp\|O08908 | 9 | 8 | 21,3 |
| P03893 | P03893 | NADH-ubiq | Mtnd2 | sp\|P03893 | 2 | 2 | 7,5 |
| Q80U30 | Q80U30 | Protein CLI | Clec16a | sp\|Q80U30 | 3 | 3 | 6,1 |
| Q06180 | Q06180 | Tyrosine-p | Ptpn2 | sp\|Q06180 | 11 | 11 | 38,2 |
| Q05CL8 | Q05CL8 | La-related | Larp7 | sp\|Q05CL8 | 18 | 18 | 36,8 |
| Q8BN21 | Q8BN21 | Serine/thre | Vrk2 | sp\|Q8BN2 | 11 | 11 | 28,8 |
| O54826 | O54826 | Protein AF | Mllt10 | sp\|O54826 | 5 | 5 | 11,2 |
| P97333 | P97333 | Neuropilin | Nrp1 | sp\|P97333 | 5 | 5 | 8,8 |
| Q922F4;CC | Q922F4 | Tubulin be | Tubb6 | sp\|Q922F4 | 26 | 12 | 77,6 |
| Q8BHC4 | Q8BHC4 | Dephosph | Dcakd | sp\|Q8BHC | 12 | 12 | 50,2 |
| Q7TT00 | Q7TT00 | Transcripti | Supt20h | sp\|Q7TT0 | 7 | 7 | 18,7 |
| Q9CQC9 | Q9CQC9 | GTP-bindin | Sar1b | sp\|Q9CQC | 10 | 9 | 55,1 |
| Q9WV54 | Q9WV54 | Acid ceram | Asah1 | sp\|Q9WV5 | 14 | 14 | 35,3 |
| P59729 | P59729 | Ras and Ra | Rin3 | sp\|P59729 | 8 | 8 | 10,8 |
| Q9D0G0 | Q9D0G0 | 28S riboso | Mrps30 | sp\|Q9D0G | 16 | 16 | 48,9 |
| Q8C3B8 | Q8C3B8 | Protein RF | Rft1 | sp\|Q8C3B | 2 | 2 | 3,3 |
| Q8BGD8 | Q8BGD8 | Cytochrom | Coa6 | sp\|Q8BGD | 3 | 3 | 38 |
| Q99K82 | Q99K82 | Spermine | Smox | sp\|Q99K8 | 4 | 4 | 12,3 |
| Q9QZ67 | Q9QZ67 | Protein ph | Ppm1d | sp\|Q9QZ6 | 2 | 2 | 5,4 |
| P39098 | P39098 | Mannosyl- | Man1a2 | sp\|P39098 | 13 | 13 | 27,5 |
| O88196 | O88196 | E3 ubiquiti | Ttc3 | sp\|O88196 | 11 | 11 | 7 |
| P58771 | P58771 | Tropomyo | Tpm1 | sp\|P58771 | 28 | 12 | 60,6 |
| Q64378 | Q64378 | Peptidyl-p | Fkbp5 | sp\|Q64378 | 15 | 15 | 37,5 |
| Q05860 | Q05860 | Formin-1 | Fmn1 | sp\|Q05860 | 5 | 5 | 4,4 |
| Q8R0F3 | Q8R0F3 | Sulfatase- | Sumf1 | sp\|Q8R0F3 | 7 | 7 | 24,7 |
| P62911 | P62911 | 60S riboso | Rpl32 | sp\|P62911 | 8 | 8 | 45,9 |
| Q9D799 | Q9D799 | Methionyl- | Mtfmt | sp\|Q9D79 | 3 | 3 | 9,3 |
| P05132 | P05132 | cAMP-dep | Prkaca | sp\|P05132 | 14 | 8 | 47,6 |
| Q9CQS4 | Q9CQS4 | Solute carr | Slc25a46 | sp\|Q9CQS | 5 | 5 | 17 |
| P14206 | P14206 | 40S riboso | Rpsa | sp\|P14206 | 16 | 16 | 58,6 |
| Q8R307 | Q8R307 | Vacuolar p | Vps18 | sp\|Q8R30 | 13 | 13 | 18,6 |
| Q08122;Q | Q08122 | Transducin | Tle3 | sp\|Q08122 | 15 | 11 | 29,1 |
| Q91Y86 | Q91Y86 | Mitogen-a | Mapk8 | sp\|Q91Y86 | 8 | 6 | 28,4 |
| O35710 | O35710 | Nocturnin | Ccrn4l | sp\|O35710 | 6 | 6 | 23,8 |
| Q8BWH0 | Q8BWH0 | Putative so | Slc38a7 | sp\|Q8BWH | 3 | 3 | 7,3 |
| Q80X95;Q | Q80X95;Q6NT | Ras-relate | Rraga;Rrag | sp\|Q80X95 | 11 | 11 | 38 |
| Q91ZV3 | Q91ZV3 | Discoidin, | Dcbld2 | sp\|Q91ZV3 | 8 | 8 | 16,3 |
| Q9DB29 | Q9DB29 | Isoamyl ac | Iah1 | sp\|Q9DB2 | 6 | 6 | 35,7 |
| Q921S7 | Q921S7 | 39S riboso | Mrpl37 | sp\|Q921S7 | 22 | 22 | 54,8 |
| Q62393 | Q62393 | Tumor pro | Tpd52 | sp\|Q62393 | 12 | 12 | 63,4 |
| Q8VCQ6 | Q8VCQ6 | Phosphatid | Sgms1 | sp\|Q8VCQ | 2 | 2 | 6,2 |
| Q9DD03 | Q9DD03 | Ras-relate | Rab13 | sp\|Q9DD0 | 6 | 3 | 30,2 |
| Q01341 | Q01341 | Adenylate | Adcy6 | sp\|Q01341 | 4 | 4 | 5,1 |
| Q8K0S9 | Q8K0S9 | snRNA-acti | Snapc1 | sp\|Q8K0S9 | 3 | 3 | 7,2 |
| Q3UK10 | Q3UK10 | B9 domain | B9d2 | sp\|Q3UK1 | 1 | 1 | 12 |
| P99028 | P99028 | Cytochrom | Uqcrh | sp\|P99028 | 5 | 5 | 57,3 |
| P62806 | P62806 | Histone H4 | Hist1h4a | sp\|P62806 | 14 | 14 | 57,3 |
| Q8BYK6 | Q8BYK6 | YTH domai | Ythdf3 | sp\|Q8BYK | 12 | 9 | 29,6 |
| Q810L3 | Q810L3 | E3 ubiquiti | Chfr | sp\|Q810L3 | 1 | 1 | 2,1 |
| Q9JL60 | Q9JL60 | Glucocorti | Gmeb1 | sp\|Q9JL60 | 9 | 7 | 17,6 |

| | | | | | | | |
|---|---|---|---|---|---|---|---|
| Q8BZT9 | Q8BZT9 | Laccase do | Lacc1 | sp\|Q8BZT9 | 1 | 1 | 4 |
| Q61827 | Q61827 | Transcripti | Mafk | sp\|Q61827 | 4 | 3 | 30,1 |
| Q9D8U2 | Q9D8U2 | Transmem | Tmem41a | sp\|Q9D8U | 1 | 1 | 4,5 |
| Q8BG05 | Q8BG05 | Heterogen | Hnrnpa3 | sp\|Q8BG0 | 26 | 25 | 54,9 |
| Q9CWY4 | Q9CWY4 | Gem-assoc | Gemin7 | sp\|Q9CWY | 2 | 2 | 28,7 |
| O54988 | O54988 | STE20-like | Slk | sp\|O54988 | 34 | 32 | 43,2 |
| Q9D9M2 | Q9D9M2 | Ubiquitin c | Usp12 | sp\|Q9D9M | 4 | 1 | 15,1 |
| P61202 | P61202 | COP9 signa | Cops2 | sp\|P61202 | 25 | 25 | 59,4 |
| Q9CQG3 | Q9CQG3 | Zinc finger | Zmynd19 | sp\|Q9CQG | 2 | 2 | 12,8 |
| P63024 | P63024 | Vesicle-ass | Vamp3 | sp\|P63024 | 4 | 2 | 54,4 |
| Q61334 | Q61334 | B-cell rece | Bcap29 | sp\|Q61334 | 5 | 5 | 20,8 |
| Q8BGW1 | Q8BGW1 | Alpha-keto | Fto | sp\|Q8BGW | 15 | 15 | 37,5 |
| P0CL69 | P0CL69 | Zinc finger | Znf703 | sp\|P0CL69 | 4 | 3 | 10,6 |
| P70261 | P70261 | Paladin | Pald1 | sp\|P70261 | 13 | 13 | 18,2 |
| CON__P02 | CON__P02662 | | | | 3 | 3 | 18,6 |
| Q8BUM6 | Q8BUM6 | Protein FA | Fam163b | sp\|Q8BUM | 1 | 1 | 18 |
| Q6A0A2 | Q6A0A2 | La-related | Larp4b | sp\|Q6A0A | 15 | 15 | 26,6 |
| Q8VI59 | Q8VI59 | Pecanex-lik | Pcnxl3 | sp\|Q8VI59 | 5 | 4 | 2,9 |
| Q921Y4 | Q921Y4 | Molybdate | Mfsd5 | sp\|Q921Y4 | 4 | 4 | 12 |
| O35445 | O35445 | E3 ubiquiti | Rnf5 | sp\|O35445 | 1 | 1 | 8,3 |
| P62077 | P62077 | Mitochond | Timm8b | sp\|P62077 | 3 | 3 | 37,3 |
| P97797 | P97797 | Tyrosine-p | Sirpa | sp\|P97797 | 4 | 4 | 11,9 |
| Q9ER39 | Q9ER39 | Torsin-1A | Tor1a | sp\|Q9ER39 | 7 | 7 | 25,2 |
| P62075 | P62075 | Mitochond | Timm13 | sp\|P62075 | 5 | 5 | 62,1 |
| Q4QY64 | Q4QY64 | ATPase far | Atad5 | sp\|Q4QY6 | 33 | 33 | 25,8 |
| P32020 | P32020 | Non-specif | Scp2 | sp\|P32020 | 19 | 19 | 37,7 |
| Q9WVG9 | Q9WVG9 | Male-speci | Msl3 | sp\|Q9WVG | 1 | 1 | 2,3 |
| Q8C1F4 | Q8C1F4 | Chondroiti | Csgalnact2 | sp\|Q8C1F4 | 6 | 5 | 14,4 |
| Q9R0P5 | Q9R0P5 | Destrin | Dstn | sp\|Q9R0P5 | 13 | 12 | 78,2 |
| Q9CZV8;Q8 | Q9CZV8 | F-box/LRR- | Fbxl20 | sp\|Q9CZV8 | 8 | 8 | 21,3 |
| Q3UFS0 | Q3UFS0 | Protein zyg | Zyg11b | sp\|Q3UFS0 | 12 | 12 | 23 |
| Q8BHN3 | Q8BHN3 | Neutral alp | Ganab | sp\|Q8BHN | 45 | 45 | 59,5 |
| E9PYH6 | E9PYH6 | Histone-lys | Setd1a | sp\|E9PYH6 | 22 | 21 | 17,8 |
| Q91WB7;Q | Q91WB7;Q6P0 | Ubiquitin d | Ubtd1;Ubt | sp\|Q91WB | 4 | 4 | 20,7 |
| Q9JMG1 | Q9JMG1 | Endothelia | Edf1 | sp\|Q9JMG | 4 | 4 | 30,4 |
| Q9CZU6 | Q9CZU6 | Citrate syn | Cs | sp\|Q9CZU | 19 | 19 | 47,4 |
| Q8R4U1 | Q8R4U1 | Myb-relate | Mypop | sp\|Q8R4U | 1 | 1 | 5,9 |
| Q9QZI9 | Q9QZI9 | Serine inco | Serinc3 | sp\|Q9QZI9 | 1 | 1 | 2,5 |
| Q2TPA8 | Q2TPA8 | Hydroxyste | Hsdl2 | sp\|Q2TPA8 | 6 | 6 | 18 |
| Q8BH75 | Q8BH75 | E3 ubiquiti | Rnf41 | sp\|Q8BH7 | 1 | 1 | 6,9 |
| Q9D892 | Q9D892 | Inosine trip | Itpa | sp\|Q9D89 | 8 | 8 | 62,1 |
| Q9D975 | Q9D975 | Sulfiredoxi | Srxn1 | sp\|Q9D97 | 4 | 4 | 60,3 |
| Q91VL9 | Q91VL9 | Zinc finger | Zbtb1 | sp\|Q91VL9 | 5 | 5 | 10 |
| Q8K1S3;O0 | Q8K1S3 | Netrin rece | Unc5b | sp\|Q8K1S3 | 18 | 18 | 33,8 |
| Q9CZX8 | Q9CZX8 | 40S riboso | Rps19 | sp\|Q9CZX8 | 12 | 12 | 57,2 |
| P62242 | P62242 | 40S riboso | Rps8 | sp\|P62242 | 12 | 12 | 58,2 |
| Q8BND3 | Q8BND3 | WD repeat | Wdr35 | sp\|Q8BND | 6 | 6 | 8,6 |
| Q62418 | Q62418 | Drebrin-lik | Dbnl | sp\|Q62418 | 15 | 15 | 44 |
| P70372 | P70372 | ELAV-like p | Elavl1 | sp\|P70372 | 18 | 18 | 62,9 |
| P84084 | P84084 | ADP-ribosy | Arf5 | sp\|P84084 | 10 | 4 | 60,6 |

| | | | | | | | |
|---|---|---|---|---|---|---|---|
| Q80ZV0 | Q80ZV0 | Ribonuclea | Rnaseh2b | sp\|Q80ZV( | 6 | 6 | 24 |
| P26450;Q9 | P26450 | Phosphatic | Pik3r1 | sp\|P26450 | 9 | 6 | 18,9 |
| Q64152 | Q64152 | Transcripti | Btf3 | sp\|Q64152 | 7 | 7 | 57,4 |
| Q8VED2 | Q8VED2 | Biogenesis | Bloc1s4 | sp\|Q8VED | 3 | 3 | 16,3 |
| Q9CPQ5 | Q9CPQ5 | Centromer | Cenpq | sp\|Q9CPQ | 4 | 4 | 22,5 |
| Q8CHC4 | Q8CHC4 | Synaptojar | Synj1 | sp\|Q8CHC | 10 | 10 | 10,2 |
| P83882 | P83882 | 60S riboso | Rpl36a | sp\|P83882 | 6 | 6 | 44,3 |
| P40630 | P40630 | Transcripti | Tfam | sp\|P40630 | 8 | 8 | 31,3 |
| Q91VK1 | Q91VK1 | Basic leuci | Bzw2 | sp\|Q91VK1 | 19 | 18 | 44,4 |
| Q6P5D4 | Q6P5D4 | Centrosom | Cep135 | sp\|Q6P5D | 5 | 5 | 5,4 |
| Q9JM76 | Q9JM76 | Actin-relat | Arpc3 | sp\|Q9JM7 | 7 | 7 | 36,5 |
| Q9Z1R2 | Q9Z1R2 | Large proli | Bag6 | sp\|Q9Z1R2 | 27 | 27 | 31,8 |
| Q9CZ09 | Q9CZ09 | Histidine p | Mettl18 | sp\|Q9CZ09 | 4 | 4 | 14,9 |
| P62071 | P62071 | Ras-relatec | Rras2 | sp\|P62071 | 8 | 6 | 43,6 |
| Q9CR27 | Q9CR27 | WASH com | Ccdc53 | sp\|Q9CR27 | 5 | 5 | 38,7 |
| Q3U5F4 | Q3U5F4 | YrdC doma | Yrdc | sp\|Q3U5F4 | 7 | 7 | 32,5 |
| Q9QYJ0 | Q9QYJ0 | DnaJ homo | Dnaja2 | sp\|Q9QYJ( | 15 | 15 | 48,5 |
| Q99J95 | Q99J95 | Cyclin-dep | Cdk9 | sp\|Q99J95 | 10 | 9 | 33,6 |
| Q9CYH5 | Q9CYH5 | Glucose-fr | Gfod2 | sp\|Q9CYH | 3 | 3 | 11,7 |
| Q8BSU7 | Q8BSU7 | MOB kinas | Mob3a | sp\|Q8BSU | 1 | 1 | 11,5 |
| O35492 | O35492 | Dual speci | Clk3 | sp\|O35492 | 12 | 12 | 24,9 |
| Q3U1T3 | Q3U1T3 | Breast can | Brms1l | sp\|Q3U1T | 7 | 7 | 30,3 |
| P61087 | P61087 | Ubiquitin-c | Ube2k | sp\|P61087 | 11 | 11 | 63,5 |
| O88811 | O88811 | Signal tran | Stam2 | sp\|O88811 | 17 | 15 | 43,8 |
| P62960;Q9 | P62960 | Nuclease-s | Ybx1 | sp\|P62960 | 20 | 15 | 68 |
| Q9CZX9 | Q9CZX9 | ER membra | Emc4 | sp\|Q9CZX9 | 7 | 7 | 51,4 |
| Q08509 | Q08509 | Epidermal | Eps8 | sp\|Q08509 | 29 | 29 | 45,3 |
| A2A5R2 | A2A5R2 | Brefeldin A | Arfgef2 | sp\|A2A5R2 | 23 | 16 | 16,7 |
| P97473 | P97473 | RISC-loadir | Tarbp2 | sp\|P97473 | 5 | 5 | 17,5 |
| P61161 | P61161 | Actin-relat | Actr2 | sp\|P61161 | 16 | 16 | 44,9 |
| Q8QZV7 | Q8QZV7 | Protein asu | Asun | sp\|Q8QZV | 18 | 18 | 29,6 |
| Q99JP6 | Q99JP6 | Homer pro | Homer3 | sp\|Q99JP6 | 5 | 5 | 19,4 |
| Q5SSG4 | Q5SSG4 | GAS2-like | Gas2l2 | sp\|Q5SSG4 | 1 | 1 | 1,5 |
| A6H603 | A6H603 | NACHT dor | Nwd1 | sp\|A6H603 | 2 | 2 | 1,4 |
| Q8K3X4 | Q8K3X4 | Interferon | Irf2bpl | sp\|Q8K3X4 | 20 | 16 | 35,2 |
| Q8R2Y9 | Q8R2Y9 | SOSS comp | Nabp2 | sp\|Q8R2Y9 | 4 | 3 | 31,6 |
| Q9EP69 | Q9EP69 | Phosphatic | Sacm1l | sp\|Q9EP69 | 22 | 22 | 49,6 |
| Q9JLZ3 | Q9JLZ3 | Methylglut | Auh | sp\|Q9JLZ3 | 6 | 6 | 23,2 |
| A2AGA4 | A2AGA4 | Rhomboid- | Rhbdl2 | sp\|A2AGA | 2 | 2 | 8,3 |
| P62900 | P62900 | 60S riboso | Rpl31 | sp\|P62900 | 8 | 8 | 47,2 |
| Q921Y2 | Q921Y2 | U3 small n | Imp3 | sp\|Q921Y2 | 11 | 11 | 66,8 |
| Q8BMK0 | Q8BMK0 | Centrosom | Cep85 | sp\|Q8BMK | 7 | 7 | 12,9 |
| O70622 | O70622 | Reticulon-2 | Rtn2 | sp\|O70622 | 2 | 2 | 7,2 |
| O89116 | O89116 | Vesicle tra | Vti1a | sp\|O89116 | 3 | 3 | 18,9 |
| Q9D735 | Q9D735 | Uncharacterized prote | | sp\|Q9D735 | 5 | 5 | 28,3 |
| Q3UUG6 | Q3UUG6 | TBC1 doma | Tbc1d24 | sp\|Q3UUG | 4 | 4 | 8,6 |
| Q9WVS7 | Q9WVS7 | Dual speci | Map2k5 | sp\|Q9WVS | 5 | 5 | 18,5 |
| Q8JZZ6 | Q8JZZ6 | AMMECR1 | Ammecr1l | sp\|Q8JZZ6 | 5 | 4 | 20,3 |
| Q9DD19 | Q9DD19 | Ras associa | Rassf7 | sp\|Q9DD1 | 2 | 2 | 8,9 |
| Q91YK8 | Q91YK8 | Ly6/PLAUR | Lypd3 | sp\|Q91YK8 | 1 | 1 | 5,2 |

| | | | | | | | |
|---|---|---|---|---|---|---|---|
| Q8VDG7 | Q8VDG7 | Platelet-ac | Pafah2 | sp\|Q8VDG | 6 | 6 | 20,5 |
| Q91XE8 | Q91XE8 | Transmem | Tmem205 | sp\|Q91XE8 | 1 | 1 | 9 |
| O88559 | O88559 | Menin | Men1 | sp\|O88559 | 10 | 10 | 29 |
| Q9D8Y0;Q | Q9D8Y0 | EF-hand dc | Efhd2 | sp\|Q9D8Y( | 11 | 11 | 60 |
| Q9CR25 | Q9CR25 | Diphthami | Dph2 | sp\|Q9CR25 | 5 | 5 | 13,3 |
| Q5XF90 | Q5XF90 | Probable c | Atp13a4 | sp\|Q5XF90 | 3 | 3 | 3,9 |
| Q9D6K5 | Q9D6K5 | Synaptojar | Synj2bp | sp\|Q9D6K5 | 4 | 4 | 32,4 |
| Q5XKN4 | Q5XKN4 | Protein jag | Jagn1 | sp\|Q5XKN4 | 3 | 3 | 15,8 |
| Q9EPQ8 | Q9EPQ8 | Transcripti | Tcf20 | sp\|Q9EPQ8 | 23 | 23 | 20,2 |
| Q9CXG9 | Q9CXG9 | PHD finger | Phf19 | sp\|Q9CXG9 | 5 | 5 | 10,6 |
| Q80YR2 | Q80YR2 | Protein FA | Fam160b2 | sp\|Q80YR2 | 5 | 4 | 7,3 |
| Q9D084 | Q9D084 | Centromer | Apitd1 | sp\|Q9D084 | 2 | 2 | 15,5 |
| O54940 | O54940 | BCL2/aden | Bnip2 | sp\|O54940 | 5 | 4 | 16,3 |
| Q9JI13 | Q9JI13 | Something | Utp3 | sp\|Q9JI13 | 13 | 13 | 35,6 |
| Q8BZB2 | Q8BZB2 | Phosphopa | Ppcdc | sp\|Q8BZB2 | 1 | 1 | 6,9 |
| Q9CWG9 | Q9CWG9 | Biogenesis | Bloc1s2 | sp\|Q9CWG | 2 | 2 | 19,6 |
| Q3UMQ8 | Q3UMQ8 | H/ACA ribc | Naf1 | sp\|Q3UMQ | 5 | 5 | 16,4 |
| Q9Z2I2 | Q9Z2I2 | Peptidyl-pr | Fkbp1b | sp\|Q9Z2I2 | 2 | 2 | 25 |
| Q3U8K7 | Q3U8K7 | Histone-lys | Suv420h1 | sp\|Q3U8K7 | 4 | 4 | 7,7 |
| Q6TYB5 | Q6TYB5 | Fasciculati | Fez2 | sp\|Q6TYB5 | 5 | 5 | 18,4 |
| Q8R184 | Q8R184 | Protein FA | Fam110a | sp\|Q8R184 | 2 | 2 | 11,1 |
| O54786 | O54786 | DNA fragm | Dffa | sp\|O54786 | 4 | 4 | 15,1 |
| Q3TYD6 | Q3TYD6 | Serine/thre | Lmtk2 | sp\|Q3TYD6 | 10 | 10 | 10,1 |
| Q8BMD8 | Q8BMD8 | Calcium-bi | Slc25a24 | sp\|Q8BMD | 27 | 27 | 63,2 |
| Q9WTK2 | Q9WTK2 | Chromodo | Cdyl | sp\|Q9WTK | 6 | 6 | 13,2 |
| Q80Y55 | Q80Y55 | BSD domai | Bsdc1 | sp\|Q80Y55 | 6 | 6 | 25,5 |
| P52623 | P52623 | Uridine-cyt | Uck1 | sp\|P52623 | 8 | 6 | 41,5 |
| Q8C0N2 | Q8C0N2 | Glycerol-3- | Agpat9 | sp\|Q8C0N | 11 | 10 | 27,2 |
| Q8BJM5 | Q8BJM5 | Zinc transp | Slc30a6 | sp\|Q8BJM | 4 | 4 | 14,3 |
| Q8K2C7 | Q8K2C7 | Protein OS | Os9 | sp\|Q8K2C7 | 12 | 12 | 28,7 |
| Q8R088 | Q8R088 | Golgi phos | Golph3l | sp\|Q8R088 | 6 | 4 | 29,8 |
| A2RT67 | A2RT67 | DENN dom | Dennd3 | sp\|A2RT67 | 3 | 3 | 3,3 |
| P56394 | P56394 | Cytochrom | Cox17 | sp\|P56394 | 2 | 2 | 57,1 |
| Q9D1I6 | Q9D1I6 | 39S riboso | Mrpl14 | sp\|Q9D1I6 | 4 | 4 | 37,2 |
| Q8C1S0 | Q8C1S0 | Mediator c | Med19 | sp\|Q8C1S0 | 3 | 3 | 25,4 |
| P47791 | P47791 | Glutathion | Gsr | sp\|P47791 | 16 | 16 | 46 |
| Q9EQC1 | Q9EQC1 | 3 beta-hyd | Hsd3b7 | sp\|Q9EQC | 7 | 7 | 22,8 |
| Q920G9 | Q920G9 | Germ cell-l | Gmcl1 | sp\|Q920G9 | 3 | 3 | 9,4 |
| Q99N91 | Q99N91 | 39S riboso | Mrpl34 | sp\|Q99N9 | 1 | 1 | 13 |
| Q9EST4 | Q9EST4 | Proteasom | Psmg2 | sp\|Q9EST4 | 8 | 8 | 31,4 |
| Q8K1K4 | Q8K1K4 | Centromer | Cenpi | sp\|Q8K1K4 | 4 | 4 | 7,9 |
| Q7TNV0 | Q7TNV0 | Protein DE | Dek | sp\|Q7TNV( | 13 | 13 | 29,5 |
| Q9CZ13 | Q9CZ13 | Cytochrom | Uqcrc1 | sp\|Q9CZ13 | 20 | 19 | 55,4 |
| Q8R4K2 | Q8R4K2 | Interleukin | Irak4 | sp\|Q8R4K2 | 7 | 7 | 22,4 |
| Q99LI9 | Q99LI9 | Polyribonu | Clp1 | sp\|Q99LI9 | 8 | 8 | 26,8 |
| P16045 | P16045 | Galectin-1 | Lgals1 | sp\|P16045 | 14 | 14 | 94,1 |
| P62245 | P62245 | 40S riboso | Rps15a | sp\|P62245 | 9 | 9 | 65,4 |
| O35317 | O35317 | Pre-B-cell l | Pbx3 | sp\|O35317 | 2 | 2 | 9 |
| O09106 | O09106 | Histone de | Hdac1 | sp\|O09106 | 12 | 8 | 40,2 |
| B1AS29 | B1AS29 | Glutamate | Grik3 | sp\|B1AS29 | 3 | 3 | 5,4 |

| | | | | | | | |
|---|---|---|---|---|---|---|---|
| Q9CQW9 | Q9CQW9 | Interferon- | Ifitm3 | sp\|Q9CQW | 4 | 3 | 48,2 |
| Q505G8 | Q505G8 | Zinc finger | Znf827 | sp\|Q505G8 | 7 | 7 | 8,2 |
| P59279 | P59279 | Ras-related | Rab2b | sp\|P59279 | 11 | 3 | 56 |
| Q99NF3 | Q99NF3 | Centrosom | Cep41 | sp\|Q99NF3 | 5 | 5 | 22 |
| Q9D7F7 | Q9D7F7 | Charged m | Chmp4c | sp\|Q9D7F7 | 3 | 3 | 21,6 |
| P15532 | P15532 | Nucleoside | Nme1 | sp\|P15532 | 17 | 11 | 86,2 |
| Q9JJL8 | Q9JJL8 | Serine--tRN | Sars2 | sp\|Q9JJL8 | 14 | 14 | 43,4 |
| Q9DB70 | Q9DB70 | FUN14 dor | Fundc1 | sp\|Q9DB70 | 1 | 1 | 11 |
| Q8R4Y8 | Q8R4Y8 | Rotatin | Rttn | sp\|Q8R4Y8 | 7 | 7 | 5,4 |
| Q8VCZ6 | Q8VCZ6 | Small G pro | Sgsm3 | sp\|Q8VCZ6 | 2 | 2 | 3,2 |
| Q99LY9 | Q99LY9 | NADH deh | Ndufs5 | sp\|Q99LY9 | 5 | 5 | 48,1 |
| O35566 | O35566 | CD151 ant | Cd151 | sp\|O35566 | 4 | 4 | 22,1 |
| Q8BHT7 | Q8BHT7 | RAB6A-GEI | Rgp1 | sp\|Q8BHT7 | 1 | 1 | 8,4 |
| P62492 | P62492 | Ras-related | Rab11a | sp\|P62492 | 13 | 2 | 73,1 |
| Q9D0T2 | Q9D0T2 | Dual specif | Dusp12 | sp\|Q9D0T2 | 4 | 4 | 16,2 |
| P0C7N9 | P0C7N9 | Proteasom | Psmg4 | sp\|P0C7N9 | 3 | 3 | 45,5 |
| Q8BVA2 | Q8BVA2 | Transmem | Tmem222 | sp\|Q8BVA2 | 3 | 3 | 27,4 |
| O54864 | O54864 | Histone-lys | Suv39h1 | sp\|O54864 | 2 | 2 | 6,1 |
| Q8C8U0;O | Q8C8U0 | Liprin-beta | Ppfibp1 | sp\|Q8C8U0 | 35 | 35 | 51,2 |
| Q9CY21 | Q9CY21 | Probable 1 | Wbscr22 | sp\|Q9CY21 | 3 | 3 | 21,4 |
| Q921D4 | Q921D4 | Mediator c | Med6 | sp\|Q921D4 | 3 | 3 | 12,6 |
| P56390 | P56390 | Cyclin-dep | Cks2 | sp\|P56390 | 2 | 2 | 24,1 |
| Q8VDD5 | Q8VDD5 | Myosin-9 | Myh9 | sp\|Q8VDD | 141 | 118 | 61,2 |
| Q61206 | Q61206 | Platelet-ac | Pafah1b2 | sp\|Q61206 | 9 | 9 | 49,8 |
| Q99PW4 | Q99PW4 | TP53-regul | Tp53rk | sp\|Q99PW | 5 | 5 | 32,8 |
| Q5SV80 | Q5SV80 | Unconvent | Myo19 | sp\|Q5SV80 | 7 | 7 | 10,2 |
| O70480 | O70480 | Vesicle-ass | Vamp4 | sp\|O70480 | 3 | 3 | 28,4 |
| Q8R5C5 | Q8R5C5 | Beta-centr | Actr1b | sp\|Q8R5C5 | 13 | 7 | 56,6 |
| A2ASS6 | A2ASS6 | Titin | Ttn | sp\|A2ASS6 | 20 | 20 | 0,7 |
| Q3V0C5 | Q3V0C5 | Ubiquitin c | Usp48 | sp\|Q3V0C5 | 18 | 18 | 23,7 |
| Q8C551 | Q8C551 | RAD51-ass | Rad51ap1 | sp\|Q8C551 | 3 | 3 | 11,3 |
| P83917 | P83917 | Chromobo | Cbx1 | sp\|P83917 | 8 | 7 | 54,1 |
| A3KGS3 | A3KGS3 | Ral GTPase | Ralgapa2 | sp\|A3KGS3 | 6 | 6 | 3,8 |
| P55821 | P55821 | Stathmin-2 | Stmn2 | sp\|P55821 | 4 | 1 | 15,1 |
| Q9CY24 | Q9CY24 | Transmem | Tmem179b | sp\|Q9CY24 | 1 | 1 | 9,1 |
| Q9QXA1 | Q9QXA1 | Cysteine ar | Cyhr1 | sp\|Q9QXA | 4 | 4 | 15,1 |
| Q3TT38 | Q3TT38 | Uncharacte | D17Wsu92 | sp\|Q3TT38 | 3 | 3 | 18,2 |
| P29037;Q6 | P29037;Q6SJ9 | TATA-box-l | Tbp;Tbpl2 | sp\|P29037 | 4 | 4 | 19,3 |
| Q8VDS4 | Q8VDS4 | Regulation | Rprd1a | sp\|Q8VDS4 | 13 | 11 | 51,3 |
| Q9WUF3 | Q9WUF3 | CASP8-ass | Casp8ap2 | sp\|Q9WUF | 6 | 6 | 4,3 |
| Q9Z127 | Q9Z127 | Large neut | Slc7a5 | sp\|Q9Z127 | 5 | 4 | 8,8 |
| O55135 | O55135 | Eukaryotic | Eif6 | sp\|O55135 | 10 | 10 | 60 |
| P70351 | P70351 | Histone-lys | Ezh1 | sp\|P70351 | 5 | 3 | 9,8 |
| P35428 | P35428 | Transcripti | Hes1 | sp\|P35428 | 2 | 2 | 12,1 |
| Q9JM62 | Q9JM62 | Receptor e | Reep6 | sp\|Q9JM6 | 1 | 1 | 6,5 |
| P38532 | P38532 | Heat shock | Hsf1 | sp\|P38532 | 6 | 6 | 11,2 |
| P70275 | P70275 | Semaphori | Sema3e | sp\|P70275 | 4 | 4 | 5,7 |
| Q9CX97 | Q9CX97 | WD repeat | Wdr55 | sp\|Q9CX97 | 9 | 9 | 34 |
| Q9R1C0 | Q9R1C0 | Transcripti | Taf7 | sp\|Q9R1C0 | 6 | 6 | 24,6 |
| Q8R179 | Q8R179 | Kelch repe | Kbtbd4 | sp\|Q8R179 | 6 | 6 | 12,9 |

| | | | | | | | |
|---|---|---|---|---|---|---|---|
| P57791 | P57791 | CAAX pren | Rce1 | sp\|P57791 | 3 | 3 | 10,9 |
| Q9Z2G6 | Q9Z2G6 | Protein sel | Sel1l | sp\|Q9Z2G6 | 13 | 13 | 26,7 |
| Q5SPL2 | Q5SPL2 | PHD finger | Phf12 | sp\|Q5SPL2 | 9 | 9 | 15,4 |
| Q6P9P0 | Q6P9P0 | Protein FA | Fam178a | sp\|Q6P9P0 | 1 | 1 | 0,8 |
| O54965 | O54965 | E3 ubiquiti | Rnf13 | sp\|O54965 | 5 | 5 | 15 |
| Q9QX11 | Q9QX11 | Cytohesin- | Cyth1 | sp\|Q9QX1 | 7 | 2 | 19,3 |
| Q8CAB8 | Q8CAB8 | GATS-like | Gatsl2 | sp\|Q8CAB | 4 | 3 | 17 |
| Q9WU60 | Q9WU60 | Attractin | Atrn | sp\|Q9WU6 | 8 | 8 | 7,5 |
| Q569Z6 | Q569Z6 | Thyroid ho | Thrap3 | sp\|Q569Z6 | 28 | 27 | 26,6 |
| Q8CF89 | Q8CF89 | TGF-beta-a | Tab1 | sp\|Q8CF89 | 9 | 9 | 32,9 |
| Q9DBZ9 | Q9DBZ9 | Rho GTPas | Syde1 | sp\|Q9DBZ | 2 | 2 | 3,7 |
| O35127 | O35127 | Protein C1 | Grcc10 | sp\|O35127 | 3 | 3 | 37,3 |
| Q9CQL7 | Q9CQL7 | MORF4 fan | Mrfap1 | sp\|Q9CQL7 | 4 | 4 | 42,4 |
| Q6ZPK0 | Q6ZPK0 | PHD finger | Phf21a | sp\|Q6ZPK0 | 6 | 6 | 13,2 |
| Q3U308 | Q3U308 | Cytoplasm | Ctu2 | sp\|Q3U30 | 13 | 13 | 31,3 |
| Q3UN04 | Q3UN04 | Ubiquitin c | Usp30 | sp\|Q3UN0 | 6 | 6 | 19,3 |
| Q9R020 | Q9R020 | Zinc finger | Zranb2 | sp\|Q9R020 | 10 | 10 | 28,8 |
| E9Q784 | E9Q784 | | Zc3h13 | sp\|E9Q784 | 14 | 14 | 9,3 |
| Q9JKL4 | Q9JKL4 | NADH deh | Ndufaf3 | sp\|Q9JKL4 | 4 | 4 | 28,1 |
| Q91VM3 | Q91VM3 | WD repeat | Wdr45 | sp\|Q91VM | 4 | 4 | 12,8 |
| P42337 | P42337 | Phosphatid | Pik3ca | sp\|P42337 | 9 | 9 | 11,3 |
| P68510 | P68510 | 14-3-3 pro | Ywhah | sp\|P68510 | 18 | 14 | 69,5 |
| Q8CHI8 | Q8CHI8 | E1A-bindin | Ep400 | sp\|Q8CHI8 | 45 | 45 | 20,9 |
| Q9R1Q9 | Q9R1Q9 | V-type pro | Atp6ap1 | sp\|Q9R1Q | 12 | 12 | 27,4 |
| Q8R3W5 | Q8R3W5 | tRNA-splici | Tsen15 | sp\|Q8R3W | 2 | 2 | 21,4 |
| P54071 | P54071 | Isocitrate d | Idh2 | sp\|P54071 | 26 | 25 | 54,2 |
| P23798 | P23798 | Polycomb | Pcgf2 | sp\|P23798 | 5 | 3 | 16,7 |
| Q924N4 | Q924N4 | Solute carr | Slc12a6 | sp\|Q924N | 6 | 2 | 5,3 |
| Q6PB70 | Q6PB70 | Anoctamin | Ano8 | sp\|Q6PB70 | 6 | 6 | 6,8 |
| Q9CQS5 | Q9CQS5 | Serine/thre | Riok2 | sp\|Q9CQS | 6 | 6 | 18,5 |
| Q9CXY1 | Q9CXY1 | Transmem | Tmem175 | sp\|Q9CXY1 | 2 | 2 | 9 |
| P61969 | P61969 | LIM domai | Lmo4 | sp\|P61969 | 2 | 2 | 26,1 |
| Q9D0D4 | Q9D0D4 | Probable d | Dimt1 | sp\|Q9D0D | 13 | 13 | 46,3 |
| Q8C0L9 | Q8C0L9 | Glyceroph | Gpcpd1 | sp\|Q8C0L9 | 5 | 5 | 8,9 |
| Q9WV89 | Q9WV89 | Syntaxin-b | Stxbp4 | sp\|Q9WV8 | 5 | 5 | 9,7 |
| Q8VEL2 | Q8VEL2 | Myotubula | Mtmr14 | sp\|Q8VEL2 | 6 | 6 | 10,3 |
| Q69ZF8 | Q69ZF8 | E3 ubiquiti | Msl2 | sp\|Q69ZF8 | 2 | 2 | 4,2 |
| P62270 | P62270 | 40S riboso | Rps18 | sp\|P62270 | 9 | 9 | 48,7 |
| Q9CQS2 | Q9CQS2 | H/ACA ribo | Nop10 | sp\|Q9CQS | 4 | 4 | 57,8 |
| B2RR83 | B2RR83 | Probable A | Ythdc2 | sp\|B2RR83 | 24 | 24 | 21,7 |
| Q9CZB9 | Q9CZB9 | Transmem | Tmem128 | sp\|Q9CZB9 | 1 | 1 | 15,3 |
| Q8BVI4 | Q8BVI4 | Dihydropte | Qdpr | sp\|Q8BVI4 | 9 | 9 | 57,3 |
| P03899 | P03899 | NADH-ubiq | Mtnd3 | sp\|P03899 | 1 | 1 | 13 |
| Q9JII5 | Q9JII5 | DAZ-associ | Dazap1 | sp\|Q9JII5\| | 13 | 13 | 39,7 |
| Q8BXC6 | Q8BXC6 | COMM dor | Commd2 | sp\|Q8BXC6 | 5 | 5 | 29,1 |
| Q8BSF4 | Q8BSF4 | Phosphatid | Pisd | sp\|Q8BSF4 | 9 | 9 | 25,9 |
| Q9CPP6 | Q9CPP6 | NADH deh | Ndufa5 | sp\|Q9CPP6 | 5 | 5 | 63,8 |
| Q5BL07 | Q5BL07 | Peroxisom | Pex1 | sp\|Q5BL07 | 8 | 8 | 8,3 |
| CON__ENS | CON__ENSEMBL:ENSBTAP00000014147 | | | | 1 | 1 | 13,3 |
| Q3TNH5 | Q3TNH5 | Protein FA | Fam172a | sp\|Q3TNH | 11 | 11 | 30,9 |

| | | | | | | | |
|---|---|---|---|---|---|---|---|
| O55131;Q9 | O55131 | Septin-7 | 07-set | sp\|O55131 | 25 | 24 | 53,2 |
| Q9CWR1 | Q9CWR1 | WD repeat | Wdr73 | sp\|Q9CWR | 6 | 6 | 22,1 |
| Q80YX1 | Q80YX1 | Tenascin | Tnc | sp\|Q80YX1 | 9 | 9 | 6,1 |
| O88665 | O88665 | Bromodom | Brd7 | sp\|O88665 | 12 | 12 | 35,8 |
| P11157 | P11157 | Ribonucleo | Rrm2 | sp\|P11157 | 20 | 18 | 58,2 |
| Q8C0K5 | Q8C0K5 | Graves dis | Slc25a16 | sp\|Q8C0K5 | 2 | 2 | 7,8 |
| Q8BW41 | Q8BW41 | Protein O-l | Pomgnt2 | sp\|Q8BW4 | 5 | 5 | 12,9 |
| Q8VDZ4 | Q8VDZ4 | Palmitoyltr | Zdhhc5 | sp\|Q8VDZ4 | 9 | 9 | 21,3 |
| Q9QUG3 | Q9QUG3 | Prion-like p | Prnd | sp\|Q9QUG | 3 | 3 | 20,1 |
| O08856 | O08856 | RNA polym | Ell | sp\|O08856 | 8 | 8 | 16,6 |
| Q8C2K1 | Q8C2K1 | Differentia | Def6 | sp\|Q8C2K1 | 3 | 3 | 5,4 |
| Q8R180 | Q8R180 | ERO1-like p | Ero1l | sp\|Q8R180 | 26 | 25 | 58,6 |
| Q62481 | Q62481 | Vacuolar p | Vps72 | sp\|Q62481 | 9 | 9 | 23,6 |
| Q3UMR5 | Q3UMR5 | Calcium un | Mcu | sp\|Q3UMR | 11 | 11 | 30,6 |
| Q9D7B6 | Q9D7B6 | Isobutyryl- | Acad8 | sp\|Q9D7B6 | 13 | 13 | 40,4 |
| Q62446 | Q62446 | Peptidyl-pr | Fkbp3 | sp\|Q62446 | 13 | 13 | 51,8 |
| Q9DBN4 | Q9DBN4 | Putative m | P33monox | sp\|Q9DBN | 5 | 5 | 24,4 |
| Q8BNI4;Q9 | Q8BNI4 | Derlin-2 | Derl2 | sp\|Q8BNI4 | 5 | 5 | 20,5 |
| Q9DAZ9 | Q9DAZ9 | Abscission/ | Zfyve19 | sp\|Q9DAZ9 | 2 | 2 | 10,8 |
| A6H619 | A6H619 | PHD and R | Phrf1 | sp\|A6H619 | 12 | 12 | 13,7 |
| Q60925 | Q60925 | D site-bind | Dbp | sp\|Q60925 | 2 | 2 | 12,6 |
| P14115 | P14115 | 60S riboso | Rpl27a | sp\|P14115 | 8 | 8 | 44,6 |
| P48755 | P48755 | Fos-related | Fosl1 | sp\|P48755 | 8 | 8 | 35,2 |
| Q9JIQ3 | Q9JIQ3 | Diablo hon | Diablo | sp\|Q9JIQ3 | 3 | 3 | 19,8 |
| Q9ERI2 | Q9ERI2 | Ras-related | Rab27a | sp\|Q9ERI2 | 5 | 4 | 24 |
| A2AJ15 | A2AJ15 | Endoplasm | Man1b1 | sp\|A2AJ15 | 10 | 10 | 17,9 |
| Q99020 | Q99020 | Heterogen | Hnrnpab | sp\|Q99020 | 20 | 20 | 48,1 |
| Q9ET26 | Q9ET26 | E3 ubiquiti | Rnf114 | sp\|Q9ET26 | 5 | 5 | 26,6 |
| P97858 | P97858 | Solute carr | Slc35b1 | sp\|P97858 | 1 | 1 | 3,1 |
| Q8BQX5 | Q8BQX5 | Transmem | Tmco6 | sp\|Q8BQX | 5 | 5 | 14,8 |
| Q3U0K8 | Q3U0K8 | Prolyl 3-hy | Ogfod1 | sp\|Q3U0K8 | 12 | 12 | 35,2 |
| Q91YI6 | Q91YI6 | Poly(A) RN | Papd4 | sp\|Q91YI6 | 3 | 3 | 11 |
| Q9CR60 | Q9CR60 | Vesicle tra | Golt1b | sp\|Q9CR60 | 2 | 2 | 16,7 |
| Q6DID3 | Q6DID3 | Protein SCA | Scaf8 | sp\|Q6DID3 | 20 | 17 | 26,7 |
| Q8C9X6 | Q8C9X6 | Enhancer o | Epc1 | sp\|Q8C9X6 | 4 | 4 | 5 |
| P62892 | P62892 | 60S riboso | Rpl39 | sp\|P62892 | 2 | 2 | 19,6 |
| Q3U0J8 | Q3U0J8 | TBC1 doma | Tbc1d2b | sp\|Q3U0J8 | 4 | 4 | 7,5 |
| P67984 | P67984 | 60S riboso | Rpl22 | sp\|P67984 | 4 | 4 | 50,8 |
| Q8BM85 | Q8BM85 | TBC domai | Tbck | sp\|Q8BM8 | 9 | 9 | 12,6 |
| Q8C3X8 | Q8C3X8 | Lipase mat | Lmf2 | sp\|Q8C3X8 | 9 | 9 | 16,2 |
| P48377 | P48377 | MHC class | Rfx1 | sp\|P48377 | 19 | 16 | 28,2 |
| P27699 | P27699 | cAMP-resp | Crem | sp\|P27699 | 1 | 1 | 8,7 |
| P29595 | P29595 | NEDD8 | Nedd8 | sp\|P29595 | 3 | 3 | 28,4 |
| Q8BVW0 | Q8BVW0 | Neutral alp | Ganc | sp\|Q8BVW | 2 | 2 | 2,2 |
| Q8VDT9 | Q8VDT9 | 39S riboso | Mrpl50 | sp\|Q8VDT9 | 7 | 7 | 47,8 |
| Q6GQT6 | Q6GQT6 | Sterol regu | Scap | sp\|Q6GQT | 6 | 6 | 6,7 |
| Q9Z210 | Q9Z210 | Peroxisom | Pex11b | sp\|Q9Z210 | 6 | 6 | 29 |
| Q497V5 | Q497V5 | S1 RNA-bin | Srbd1 | sp\|Q497V5 | 24 | 24 | 33 |
| Q9D0W5 | Q9D0W5 | Peptidyl-pr | Ppil1 | sp\|Q9D0W | 6 | 6 | 37,3 |
| Q80YA7 | Q80YA7 | Dipeptidyl | Dpp8 | sp\|Q80YA7 | 8 | 8 | 13,5 |

| | | | | | | | |
|---|---|---|---|---|---|---|---|
| Q9CQA5 | Q9CQA5 | Mediator c | Med4 | sp\|Q9CQA | 10 | 10 | 53,3 |
| Q9CPV7 | Q9CPV7 | Palmitoyltr | Zdhhc6 | sp\|Q9CPV7 | 5 | 5 | 13,8 |
| Q9CPY7 | Q9CPY7 | Cytosol am | Lap3 | sp\|Q9CPY7 | 26 | 26 | 62,8 |
| Q9CPV2 | Q9CPV2 | Anaphase- | Anapc16 | sp\|Q9CPV2 | 3 | 3 | 40,9 |
| Q8K1N1 | Q8K1N1 | Calcium-in | Pnpla8 | sp\|Q8K1N | 8 | 8 | 15,7 |
| P63323 | P63323 | 40S riboso | Rps12 | sp\|P63323 | 8 | 8 | 59,8 |
| Q3UHC0 | Q3UHC0 | Trinucleoti | Tnrc6c | sp\|Q3UHC | 5 | 4 | 4 |
| P62821;Q8 | P62821 | Ras-related | Rab1A | sp\|P62821 | 15 | 9 | 76,1 |
| P58058 | P58058 | NAD kinase | Nadk | sp\|P58058 | 7 | 7 | 21,9 |
| Q78XF5 | Q78XF5 | Oligosacch | Ostc | sp\|Q78XF5 | 2 | 2 | 13,4 |
| O35427 | O35427 | DNA-direct | Crcp | sp\|O35427 | 4 | 4 | 53,4 |
| Q91ZV0 | Q91ZV0 | Melanoma | Mia2 | sp\|Q91ZV0 | 23 | 23 | 21,6 |
| Q9CQU3 | Q9CQU3 | Protein RE | Rer1 | sp\|Q9CQU | 5 | 5 | 29,6 |
| Q9D8U7 | Q9D8U7 | DTW doma | Dtwd1 | sp\|Q9D8U | 4 | 4 | 19,7 |
| Q9QZ88 | Q9QZ88 | Vacuolar p | Vps29 | sp\|Q9QZ88 | 8 | 8 | 47,8 |
| Q8VDY4 | Q8VDY4 | EF-hand ca | Efcab7 | sp\|Q8VDY4 | 6 | 6 | 13,2 |
| Q6NZP1 | Q6NZP1 | DNA annea | Zranb3 | sp\|Q6NZP1 | 3 | 3 | 4,4 |
| Q8K0D5 | Q8K0D5 | Elongation | Gfm1 | sp\|Q8K0D5 | 30 | 30 | 46,3 |
| P97855 | P97855 | Ras GTPase | G3bp1 | sp\|P97855 | 22 | 19 | 58,1 |
| Q9JLN9 | Q9JLN9 | Serine/thre | Mtor | sp\|Q9JLN9 | 39 | 39 | 23,1 |
| P62137 | P62137 | Serine/thre | Ppp1ca | sp\|P62137 | 23 | 9 | 72,7 |
| Q9EPL9 | Q9EPL9 | Peroxisom | Acox3 | sp\|Q9EPL9 | 6 | 6 | 12,6 |
| Q9CYZ6 | Q9CYZ6 | Uncharacterized prote | | sp\|Q9CYZ6 | 8 | 8 | 69,2 |
| Q64327 | Q64327 | Male-enha | Mea1 | sp\|Q64327 | 1 | 1 | 7,5 |
| Q9CR80 | Q9CR80 | Protein FA | Fam32a | sp\|Q9CR80 | 4 | 4 | 35,7 |
| O88512 | O88512 | AP-1 comp | Ap1g2 | sp\|O88512 | 5 | 5 | 8,6 |
| Q61286 | Q61286 | Transcripti | Tcf12 | sp\|Q61286 | 5 | 5 | 15,3 |
| Q3U2K0 | Q3U2K0 | Protein FA | Fam193b | sp\|Q3U2K0 | 2 | 2 | 5,3 |
| Q6TXD4 | Q6TXD4 | Dynamin-b | Dnmbp | sp\|Q6TXD4 | 10 | 10 | 7,8 |
| Q6NXH2 | Q6NXH2 | Glycoprote | Manea | sp\|Q6NXH | 3 | 3 | 11,9 |
| Q60870 | Q60870 | Receptor e | Reep5 | sp\|Q60870 | 2 | 2 | 10,3 |
| Q8CB44 | Q8CB44 | GRAM dom | Gramd4 | sp\|Q8CB44 | 14 | 14 | 27,8 |
| P23708 | P23708 | Nuclear tra | Nfya | sp\|P23708 | 2 | 2 | 12,1 |
| Q9CWU6 | Q9CWU6 | Ubiquinol- | Uqcc1 | sp\|Q9CWU | 6 | 6 | 25,8 |
| P42669 | P42669 | Transcripti | Pura | sp\|P42669 | 8 | 8 | 40,2 |
| Q52KR2 | Q52KR2 | Leucine-ric | Lrig2 | sp\|Q52KR2 | 2 | 2 | 2,3 |
| Q9CPV3 | Q9CPV3 | 39S riboso | Mrpl42 | sp\|Q9CPV3 | 2 | 2 | 25,4 |
| Q9CY25 | Q9CY25 | Protein MI | Mis12 | sp\|Q9CY25 | 6 | 6 | 35,9 |
| Q9CQ49 | Q9CQ49 | Nuclear ca | Ncbp2 | sp\|Q9CQ4 | 4 | 4 | 25 |
| Q99LH9 | Q99LH9 | SH3 domai | Sh3bp5l | sp\|Q99LH9 | 5 | 5 | 14,5 |
| Q99KN2 | Q99KN2 | Probable c | Ciao1 | sp\|Q99KN | 9 | 9 | 43,4 |
| E9Q8T7 | E9Q8T7 | | Dnah1 | sp\|E9Q8T7 | 4 | 4 | 1,4 |
| Q6ZQ89 | Q6ZQ89 | E3 ubiquiti | 06-mar | sp\|Q6ZQ8 | 7 | 7 | 12,7 |
| Q9WVS8 | Q9WVS8 | Mitogen-a | Mapk7 | sp\|Q9WVS | 5 | 5 | 10,3 |
| Q8R1I1 | Q8R1I1 | Cytochrom | Uqcr10 | sp\|Q8R1I1 | 3 | 3 | 50 |
| Q6PGF7 | Q6PGF7 | Exocyst co | Exoc8 | sp\|Q6PGF7 | 11 | 11 | 20,8 |
| Q9JKJ9 | Q9JKJ9 | 24-hydroxy | Cyp39a1 | sp\|Q9JKJ9 | 1 | 1 | 3,2 |
| Q9CX30 | Q9CX30 | Protein YIF | Yif1b | sp\|Q9CX30 | 5 | 5 | 20,6 |
| Q9CXY9 | Q9CXY9 | GPI-anchor | Pigk | sp\|Q9CXY9 | 9 | 9 | 30,6 |
| Q6A039 | Q6A039 | TBC1 doma | Tbc1d12 | sp\|Q6A039 | 4 | 3 | 8,2 |

| | | | | | | | |
|---|---|---|---|---|---|---|---|
| Q9JL19 | Q9JL19 | Nuclear re | Ncoa6 | sp\|Q9JL19 | 5 | 5 | 4,1 |
| Q9JLJ5 | Q9JLJ5 | Elongation | Elovl1 | sp\|Q9JLJ5 | 3 | 3 | 14 |
| O55128 | O55128 | Histone de | Sap18 | sp\|O55128 | 7 | 7 | 45,8 |
| Q9CQL4 | Q9CQL4 | 39S riboso | Mrpl20 | sp\|Q9CQL4 | 4 | 4 | 35,6 |
| Q9CPZ6 | Q9CPZ6 | ORM1-like | Ormdl3 | sp\|Q9CPZ6 | 2 | 1 | 17 |
| Q61425 | Q61425 | Hydroxyac | Hadh | sp\|Q61425 | 9 | 9 | 48,4 |
| P42125 | P42125 | Enoyl-CoA | Eci1 | sp\|P42125 | 12 | 12 | 54,3 |
| Q6GQT9 | Q6GQT9 | Nodal mod | Nomo1 | sp\|Q6GQT | 43 | 43 | 52,5 |
| Q61214 | Q61214 | Dual specif | Dyrk1a | sp\|Q61214 | 6 | 5 | 10 |
| Q62186 | Q62186 | Translocon | Ssr4 | sp\|Q62186 | 4 | 4 | 30,8 |
| P62715 | P62715 | Serine/thre | Ppp2cb | sp\|P62715 | 14 | 1 | 61,2 |
| Q8VCM5 | Q8VCM5 | Mitochond | Mul1 | sp\|Q8VCM | 3 | 3 | 17,3 |
| Q9EPL0 | Q9EPL0 | Xylosyltran | Xylt2 | sp\|Q9EPL0 | 4 | 4 | 6,5 |
| Q9Z0M5 | Q9Z0M5 | Lysosomal | Lipa | sp\|Q9Z0M | 3 | 3 | 6,8 |
| P63168 | P63168 | Dynein ligh | Dynll1 | sp\|P63168 | 5 | 3 | 61,8 |
| P97465 | P97465 | Docking pr | Dok1 | sp\|P97465 | 13 | 13 | 44,2 |
| Q6URW6 | Q6URW6 | Myosin-14 | Myh14 | sp\|Q6URW | 17 | 9 | 9 |
| P42867 | P42867 | UDP-N-ace | Dpagt1 | sp\|P42867 | 1 | 1 | 6,8 |
| Q9R1Q8 | Q9R1Q8 | Transgelin- | Tagln3 | sp\|Q9R1Q | 5 | 2 | 22,1 |
| Q9CR51 | Q9CR51 | V-type pro | Atp6v1g1 | sp\|Q9CR51 | 4 | 4 | 46,6 |
| P68181 | P68181 | cAMP-dep | Prkacb | sp\|P68181 | 12 | 6 | 45,6 |
| O70133 | O70133 | ATP-depen | Dhx9 | sp\|O70133 | 51 | 51 | 41,2 |
| O35066 | O35066 | Kinesin-like | Kif3c | sp\|O35066 | 10 | 7 | 15,8 |
| A2A8L1 | A2A8L1 | Chromodo | Chd5 | sp\|A2A8L1 | 17 | 2 | 9,7 |
| P63087 | P63087 | Serine/thre | Ppp1cc | sp\|P63087 | 18 | 3 | 56,7 |
| Q9JJD0 | Q9JJD0 | THAP dom | Thap11 | sp\|Q9JJD0 | 3 | 3 | 8,2 |
| Q8K4M5 | Q8K4M5 | COMM dor | Commd1 | sp\|Q8K4M | 4 | 4 | 24,5 |
| P52482 | P52482 | Ubiquitin-c | Ube2e1 | sp\|P52482 | 4 | 1 | 28,5 |
| Q08943 | Q08943 | FACT comp | Ssrp1 | sp\|Q08943 | 24 | 24 | 41,7 |
| P49452 | P49452 | Centromer | Cenpc | sp\|P49452 | 17 | 17 | 25,1 |
| O08710 | O08710 | Thyroglobu | Tg | sp\|O08710 | 1 | 1 | 0,4 |
| O70201 | O70201 | Baculovira | Birc5 | sp\|O70201 | 7 | 7 | 67,1 |
| A2AKX3 | A2AKX3 | Probable h | Setx | sp\|A2AKX3 | 19 | 19 | 10,1 |
| O88447 | O88447 | Kinesin ligh | Klc1 | sp\|O88447 | 27 | 20 | 52,1 |
| Q9Z0V8 | Q9Z0V8 | Mitochond | Timm17a | sp\|Q9Z0V8 | 3 | 3 | 45,6 |
| P60605 | P60605 | Ubiquitin-c | Ube2g2 | sp\|P60605 | 4 | 4 | 42,4 |
| Q9QXT0 | Q9QXT0 | Protein car | Cnpy2 | sp\|Q9QXT0 | 10 | 10 | 56 |
| CON__Q3S | CON__Q3SX14 | | | | 18 | 5 | 31,6 |
| Q91W61 | Q91W61 | F-box/LRR- | Fbxl15 | sp\|Q91W6 | 2 | 2 | 13,7 |
| P62889 | P62889 | 60S riboso | Rpl30 | sp\|P62889 | 8 | 8 | 66,1 |
| Q6ZQ82 | Q6ZQ82 | Rho GTPas | Arhgap26 | sp\|Q6ZQ82 | 1 | 1 | 1,8 |
| Q99M96 | Q99M96 | Suppresso | St7 | sp\|Q99M9 | 4 | 2 | 8,5 |
| Q7M6Y3;Q | Q7M6Y3 | Phosphatid | Picalm | sp\|Q7M6Y | 14 | 14 | 36,8 |
| O70585;Q9 | O70585 | Dystrobrev | Dtnb | sp\|O70585 | 13 | 13 | 31,3 |
| Q8QZX5 | Q8QZX5 | U7 snRNA- | Lsm10 | sp\|Q8QZX5 | 3 | 3 | 28,7 |
| Q9DCW2 | Q9DCW2 | Phospholip | Plscr2 | sp\|Q9DCW | 1 | 1 | 6,5 |
| Q9DB50 | Q9DB50 | AP-1 comp | Ap1s2 | sp\|Q9DB50 | 4 | 2 | 21,2 |
| Q8BG95 | Q8BG95 | Protein ph | Ppp1r12b | sp\|Q8BG95 | 4 | 2 | 4,6 |
| Q8R0I4 | Q8R0I4 | TM2 doma | Tm2d2 | sp\|Q8R0I4 | 1 | 1 | 8 |
| Q8CFH6 | Q8CFH6 | Serine/thre | Sik2 | sp\|Q8CFH6 | 4 | 4 | 6,4 |

| | | | | | | | |
|---|---|---|---|---|---|---|---|
| Q9QYK7 | Q9QYK7 | RING finge | Rnf11 | sp\|Q9QYK | 2 | 2 | 18,8 |
| Q9CZB6 | Q9CZB6 | Mediator c | Med7 | sp\|Q9CZB6 | 4 | 4 | 30,9 |
| Q9JIA7 | Q9JIA7 | Sphingosin | Sphk2 | sp\|Q9JIA7 | 4 | 4 | 11 |
| O70338 | O70338 | Ribonuclea | Rnaseh1 | sp\|O70338 | 2 | 2 | 9,5 |
| Q9CPX7 | Q9CPX7 | 28S riboso | Mrps16 | sp\|Q9CPX7 | 3 | 3 | 28,9 |
| Q9D2U5 | Q9D2U5 | N-alpha-ac | Naa38 | sp\|Q9D2U | 4 | 4 | 36 |
| P62700 | P62700 | Protein yip | Ypel5 | sp\|P62700 | 5 | 5 | 52,9 |
| D0G895 | D0G895 | | Ace3 | sp\|D0G895 | 2 | 2 | 3,4 |
| P52503 | P52503 | NADH deh | Ndufs6 | sp\|P52503 | 7 | 7 | 62,1 |
| P99025 | P99025 | GTP cycloh | Gchfr | sp\|P99025 | 2 | 2 | 50 |
| Q920L5 | Q920L5 | Elongation | Elovl6 | sp\|Q920L5 | 2 | 2 | 10,5 |
| Q9Z2Q2 | Q9Z2Q2 | Lysine-rich | Knop1 | sp\|Q9Z2Q2 | 10 | 10 | 27 |
| Q8VHK1 | Q8VHK1 | Caskin-2 | Caskin2 | sp\|Q8VHK | 8 | 8 | 12 |
| Q99L85 | Q99L85 | Elongator | Elp5 | sp\|Q99L85 | 3 | 3 | 13,7 |
| Q8R2K4 | Q8R2K4 | TAF6-like R | Taf6l | sp\|Q8R2K4 | 9 | 9 | 21,4 |
| Q920S3 | Q920S3 | GATA zinc | Gatad1 | sp\|Q920S3 | 4 | 4 | 22,9 |
| Q6PHS6 | Q6PHS6 | Sorting nex | Snx13 | sp\|Q6PHS6 | 7 | 7 | 9,5 |
| Q8C078 | Q8C078 | Calcium/ca | Camkk2 | sp\|Q8C078 | 8 | 8 | 25,2 |
| P98203 | P98203 | Armadillo | Arvcf | sp\|P98203 | 8 | 8 | 11,6 |
| Q60866 | Q60866 | Phosphotri | Pter | sp\|Q60866 | 14 | 14 | 51,9 |
| Q9JJZ6 | Q9JJZ6 | Krueppel-li | Klf13 | sp\|Q9JJZ6 | 3 | 3 | 19,4 |
| Q8BUY5 | Q8BUY5 | Complex I | Timmdc1 | sp\|Q8BUY | 5 | 5 | 22,1 |
| Q9D7N3 | Q9D7N3 | 28S riboso | Mrps9 | sp\|Q9D7N | 18 | 18 | 53,3 |
| Q5SS80 | Q5SS80 | Dehydroge | Dhrs13 | sp\|Q5SS80 | 12 | 12 | 50,3 |
| Q922X9 | Q922X9 | Protein arg | Prmt7 | sp\|Q922X9 | 9 | 9 | 21,2 |
| P49138;Q3 | P49138 | MAP kinas | Mapkapk2 | sp\|P49138 | 8 | 7 | 33,4 |
| Q811D0;Q | Q811D0 | Disks large | Dlg1 | sp\|Q811D0 | 22 | 20 | 38,7 |
| Q80UK7 | Q80UK7 | Spindle ass | Sass6 | sp\|Q80UK7 | 5 | 5 | 9 |
| Q8C180;Q | Q8C180 | Fibroblast | Frs2 | sp\|Q8C180 | 4 | 4 | 12,2 |
| P70662;O5 | P70662;O5520 | LIM domai | Ldb1;Ldb2 | sp\|P70662 | 2 | 2 | 7,1 |
| P06336 | P06336 | Ig epsilon chain C regi | | sp\|P06336 | 1 | 1 | 3,3 |
| Q9WVR4 | Q9WVR4 | Fragile X m | Fxr2 | sp\|Q9WVR | 22 | 19 | 42,2 |
| P37804 | P37804 | Transgelin | Tagln | sp\|P37804 | 7 | 7 | 35,3 |
| Q01730 | Q01730 | Ras suppre | Rsu1 | sp\|Q01730 | 11 | 11 | 50,9 |
| F8VPQ2 | F8VPQ2 | | Arid4a | sp\|F8VPQ2 | 12 | 11 | 14,7 |
| P54751 | P54751 | CMP-N-ace | St3gal1 | sp\|P54751 | 5 | 5 | 19,3 |
| Q9D198 | Q9D198 | Pre-mRNA- | Syf2 | sp\|Q9D198 | 12 | 12 | 55,4 |
| Q8CIG9 | Q8CIG9 | F-box/LRR- | Fbxl8 | sp\|Q8CIG9 | 6 | 6 | 22,2 |
| P61406 | P61406 | Telomeras | Smg6 | sp\|P61406 | 12 | 12 | 12,9 |
| P51912 | P51912 | Neutral am | Slc1a5 | sp\|P51912 | 9 | 9 | 20,1 |
| Q9CQ75 | Q9CQ75 | NADH deh | Ndufa2 | sp\|Q9CQ7 | 3 | 3 | 32,3 |
| Q9EP89 | Q9EP89 | Serine beta | Lactb | sp\|Q9EP89 | 2 | 2 | 4,9 |
| Q8C351 | Q8C351 | Layilin | Layn | sp\|Q8C351 | 3 | 3 | 8,9 |
| Q62179 | Q62179 | Semaphori | Sema4b | sp\|Q62179 | 6 | 6 | 12,4 |
| Q9CXU1 | Q9CXU1 | Mediator c | Med31 | sp\|Q9CXU | 3 | 3 | 27,5 |
| B1AY10 | B1AY10 | Transcripti | Nfx1 | sp\|B1AY10 | 5 | 5 | 7,4 |
| P63002 | P63002 | Amino-ter | Aes | sp\|P63002 | 2 | 2 | 14,2 |
| Q61189 | Q61189 | Methyloso | Clns1a | sp\|Q61189 | 6 | 6 | 41,9 |
| Q8BMI4 | Q8BMI4 | Flap endor | Gen1 | sp\|Q8BMI4 | 6 | 6 | 9,7 |
| Q9DAI2 | Q9DAI2 | Intraflagell | Ift22 | sp\|Q9DAI2 | 7 | 7 | 56,2 |

| | | | | | | | |
|---|---|---|---|---|---|---|---|
| Q8BXJ2 | Q8BXJ2 | Transcripti | Trerf1 | sp\|Q8BXJ2 | 4 | 4 | 6,3 |
| Q9CXD6 | Q9CXD6 | Mitochond | Mcur1 | sp\|Q9CXD | 2 | 2 | 5,9 |
| Q8VDD8 | Q8VDD8 | WAS prote | Wash1 | sp\|Q8VDD | 10 | 10 | 31,8 |
| Q9CQ79 | Q9CQ79 | Thioredoxi | Txndc9 | sp\|Q9CQ7 | 7 | 7 | 38,9 |
| P62340 | P62340 | TATA box-l | Tbpl1 | sp\|P62340 | 3 | 3 | 22,6 |
| Q99JB8 | Q99JB8 | Protein kin | Pacsin3 | sp\|Q99JB8 | 10 | 10 | 35,4 |
| Q8QZZ7 | Q8QZZ7 | EKC/KEOPS | Tprkb | sp\|Q8QZZ | 5 | 5 | 49,1 |
| Q6ZPY7 | Q6ZPY7 | Lysine-spe | Kdm3b | sp\|Q6ZPY7 | 36 | 35 | 38 |
| Q8C0I4 | Q8C0I4 | Enhancer c | Epc2 | sp\|Q8C0I4 | 7 | 7 | 10,9 |
| Q8VC74 | Q8VC74 | Mitochond | Cox18 | sp\|Q8VC7 | 4 | 4 | 15,4 |
| P51954 | P51954 | Serine/thre | Nek1 | sp\|P51954 | 4 | 4 | 4,9 |
| Q9WTX6 | Q9WTX6 | Cullin-1 | Cul1 | sp\|Q9WTX | 31 | 31 | 51,2 |
| CON__P08 | CON__P08779 | | | | 17 | 7 | 40,8 |
| Q8K4Q0 | Q8K4Q0 | Regulatory | Rptor | sp\|Q8K4Q | 16 | 16 | 15,3 |
| Q3UGR5 | Q3UGR5 | Haloacid d | Hdhd2 | sp\|Q3UGR | 7 | 7 | 41,7 |
| P47915 | P47915 | 60S riboso | Rpl29 | sp\|P47915 | 3 | 3 | 18,8 |
| Q80TD3 | Q80TD3 | Folliculin-ir | Fnip2 | sp\|Q80TD | 4 | 4 | 3,7 |
| A2AHC3 | A2AHC3 | Calmodulir | Camsap1 | sp\|A2AHC | 16 | 16 | 17 |
| Q8BWT1 | Q8BWT1 | 3-ketoacyl- | Acaa2 | sp\|Q8BWT | 23 | 23 | 76,3 |
| P60003 | P60003 | Transcripti | Elof1 | sp\|P60003 | 1 | 1 | 21,7 |
| Q9JKB3 | Q9JKB3 | Y-box-bind | Ybx3 | sp\|Q9JKB3 | 18 | 13 | 67 |
| Q9Z2I0 | Q9Z2I0 | LETM1 and | Letm1 | sp\|Q9Z2I0 | 29 | 29 | 42,1 |
| P41245 | P41245 | Matrix met | Mmp9 | sp\|P41245 | 4 | 4 | 7,7 |
| P0C0A3 | P0C0A3 | Charged m | Chmp6 | sp\|P0C0A3 | 3 | 3 | 19,5 |
| Q5F2E8 | Q5F2E8 | Serine/thre | Taok1 | sp\|Q5F2E8 | 20 | 17 | 25,7 |
| Q91YE3;Q9 | Q91YE3 | Egl nine ho | Egln1 | sp\|Q91YE3 | 6 | 6 | 20,2 |
| Q5BU09 | Q5BU09 | E2F-associ | Eapp | sp\|Q5BU0 | 3 | 3 | 15,3 |
| Q99MD9 | Q99MD9 | Nuclear au | Nasp | sp\|Q99MD | 33 | 33 | 54,1 |
| Q99JF8 | Q99JF8 | PC4 and SF | Psip1 | sp\|Q99JF8 | 19 | 18 | 39 |
| Q9DC69 | Q9DC69 | NADH deh | Ndufa9 | sp\|Q9DC6 | 15 | 15 | 49,1 |
| E9Q4Z2 | E9Q4Z2 | Acetyl-CoA | Acacb | sp\|E9Q4Z2 | 13 | 3 | 7,1 |
| P58802 | P58802 | TBC1 doma | Tbc1d10a | sp\|P58802 | 12 | 11 | 29,2 |
| Q71FD7 | Q71FD7 | Filamin-bir | Fblim1 | sp\|Q71FD | 10 | 10 | 33,9 |
| Q4VC33 | Q4VC33 | Macrophag | Maea | sp\|Q4VC3 | 9 | 9 | 28 |
| Q8K211 | Q8K211 | High affinit | Slc31a1 | sp\|Q8K21 | 2 | 2 | 15,3 |
| Q6ZPZ3 | Q6ZPZ3 | Zinc finger | Zc3h4 | sp\|Q6ZPZ3 | 23 | 23 | 30,9 |
| P24638 | P24638 | Lysosomal | Acp2 | sp\|P24638 | 7 | 7 | 17,3 |
| Q8CCH2 | Q8CCH2 | NHL repea | Nhlrc3 | sp\|Q8CCH | 6 | 6 | 17,9 |
| O54714 | O54714 | E3 SUMO-p | Pias3 | sp\|O54714 | 2 | 2 | 4,3 |
| Q9DAJ4 | Q9DAJ4 | WD repeat | Wdr83 | sp\|Q9DAJ4 | 6 | 6 | 31,7 |
| Q8BU31 | Q8BU31 | Ras-relatec | Rap2c | sp\|Q8BU3 | 11 | 6 | 68,3 |
| P70404 | P70404 | Isocitrate d | Idh3g | sp\|P70404 | 13 | 13 | 50,6 |
| P30416 | P30416 | Peptidyl-pr | Fkbp4 | sp\|P30416 | 33 | 33 | 70,5 |
| Q99L90 | Q99L90 | Microsphe | Mcrs1 | sp\|Q99L90 | 6 | 6 | 21,4 |
| P57746 | P57746 | V-type pro | Atp6v1d | sp\|P57746 | 9 | 9 | 44,1 |
| Q3B7Z2;Q5 | Q3B7Z2 | Oxysterol-l | Osbp | sp\|Q3B7Z2 | 18 | 18 | 26,6 |
| P97379 | P97379 | Ras GTPase | G3bp2 | sp\|P97379 | 18 | 15 | 47,1 |
| P68254 | P68254 | 14-3-3 pro | Ywhaq | sp\|P68254 | 19 | 14 | 66,1 |
| Q8VDU0 | Q8VDU0 | G-protein-s | Gpsm2 | sp\|Q8VDU | 15 | 14 | 32,8 |
| Q9D9Z5 | Q9D9Z5 | DET1- and | Dda1 | sp\|Q9D9Z5 | 3 | 3 | 36,3 |

| | | | | | | | |
|---|---|---|---|---|---|---|---|
| Q811L6;Q6 | Q811L6 | Microtubu | Mast4 | sp\|Q811L6 | 15 | 13 | 9,4 |
| Q0HA38 | Q0HA38 | Tetratricop | Ttc21b | sp\|Q0HA3 | 9 | 9 | 10 |
| Q9ES56 | Q9ES56 | Trafficking | Trappc4 | sp\|Q9ES56 | 8 | 8 | 38,4 |
| Q3TDQ1 | Q3TDQ1 | Dolichyl-di | Stt3b | sp\|Q3TDQ | 12 | 11 | 18,6 |
| Q9CY18 | Q9CY18 | Sorting ne | Snx7 | sp\|Q9CY18 | 9 | 9 | 26,9 |
| Q9D174 | Q9D174 | SS18-like p | Ss18l2 | sp\|Q9D174 | 1 | 1 | 13 |
| Q921K9 | Q921K9 | B-cell CLL/ | Bcl7b | sp\|Q921K9 | 7 | 5 | 45 |
| Q3TLI0 | Q3TLI0 | Trafficking | Trappc10 | sp\|Q3TLI0 | 6 | 6 | 6,6 |
| Q9D270 | Q9D270 | Probable p | Zdhhc21 | sp\|Q9D27( | 3 | 3 | 11,7 |
| P97931 | P97931 | Uracil-DNA | Ung | sp\|P97931 | 2 | 2 | 9,8 |
| Q8R0X2 | Q8R0X2 | CDK2-asso | Cacul1 | sp\|Q8R0X2 | 5 | 5 | 21,8 |
| Q99N96 | Q99N96 | 39S riboso | Mrpl1 | sp\|Q99N9( | 12 | 12 | 41,7 |
| O54941 | O54941 | SWI/SNF-r | Smarce1 | sp\|O54941 | 14 | 14 | 38,9 |
| Q9QZV9 | Q9QZV9 | NTF2-relat | Nxt1 | sp\|Q9QZV | 3 | 3 | 25,7 |
| O88487;O8 | O88487 | Cytoplasm | Dync1i2 | sp\|O88487 | 19 | 19 | 46,7 |
| Q99N92 | Q99N92 | 39S riboso | Mrpl27 | sp\|Q99N9 | 6 | 6 | 50 |
| Q9CZ83 | Q9CZ83 | 39S riboso | Mrpl55 | sp\|Q9CZ83 | 4 | 4 | 30,7 |
| Q9WTL8 | Q9WTL8 | Aryl hydro | Arntl | sp\|Q9WTL | 4 | 4 | 7,9 |
| Q99JX7 | Q99JX7 | Nuclear RN | Nxf1 | sp\|Q99JX7 | 29 | 29 | 58,3 |
| Q99LW6 | Q99LW6 | YY1-associ | Yaf2 | sp\|Q99LW | 2 | 2 | 20,7 |
| Q9WV96 | Q9WV96 | Mitochond | Timm10b | sp\|Q9WV9 | 1 | 1 | 19 |
| Q9D7H4 | Q9D7H4 | Arginine va | Avpi1 | sp\|Q9D7H | 2 | 2 | 35,9 |
| P39054 | P39054 | Dynamin-2 | Dnm2 | sp\|P39054 | 42 | 28 | 57,8 |
| P0DP28;P0 | P0DP28;P0DP27;P0DP26 | | | sp\|P0DP28 | 14 | 14 | 67,8 |
| Q9JM73 | Q9JM73 | Serum resp | Srf | sp\|Q9JM7 | 1 | 1 | 5 |
| Q8CHH5 | Q8CHH5 | GLTSCR1-li | Gltscr1l | sp\|Q8CHH | 4 | 4 | 5,8 |
| Q9DC51;P2 | Q9DC51 | Guanine nu | Gnai3 | sp\|Q9DC5 | 13 | 8 | 44,1 |
| Q9ERY9 | Q9ERY9 | Probable e | ORF11 | sp\|Q9ERY9 | 2 | 2 | 17,1 |
| Q3USH5 | Q3USH5 | Splicing fac | Sfswap | sp\|Q3USH | 11 | 11 | 16,1 |
| P46467 | P46467 | Vacuolar p | Vps4b | sp\|P46467 | 16 | 13 | 44,4 |
| P0DOV1 | P0DOV1 | | | sp\|P0DOV | 8 | 1 | 28 |
| P62876 | P62876 | DNA-direct | Polr2l | sp\|P62876 | 2 | 2 | 29,9 |
| Q9Z1D1 | Q9Z1D1 | Eukaryotic | Eif3g | sp\|Q9Z1D1 | 19 | 19 | 59,7 |
| P48962 | P48962 | ADP/ATP t | Slc25a4 | sp\|P48962 | 22 | 11 | 67,4 |
| Q8K194 | Q8K194 | U4/U6.U5 | Snrnp27 | sp\|Q8K194 | 4 | 4 | 36,1 |
| Q8BXK9;Q8 | Q8BXK9;Q8BH | Chloride in | Clic5;Clic6 | sp\|Q8BXK9 | 1 | 1 | 6,8 |
| Q9R118 | Q9R118 | Serine prot | Htra1 | sp\|Q9R118 | 13 | 13 | 37,3 |
| Q8K273 | Q8K273 | Membrane | Mmgt1 | sp\|Q8K273 | 3 | 3 | 26,7 |
| Q9JJX7 | Q9JJX7 | Tyrosyl-DN | Tdp2 | sp\|Q9JJX7 | 6 | 6 | 24,3 |
| Q9CZP7 | Q9CZP7 | Hsp90 co-c | Cdc37l1 | sp\|Q9CZP7 | 2 | 2 | 7,5 |
| Q3UM29 | Q3UM29 | Conserved | Cog7 | sp\|Q3UM2 | 20 | 20 | 31,6 |
| Q9D0I8 | Q9D0I8 | mRNA turn | Mrto4 | sp\|Q9D0I8 | 18 | 18 | 57,7 |
| Q80U44 | Q80U44 | Zinc finger | Zfyve16 | sp\|Q80U4 | 4 | 4 | 4,1 |
| Q922D4 | Q922D4 | Serine/thr | Ppp6r3 | sp\|Q922D4 | 28 | 28 | 38,5 |
| Q99MV1 | Q99MV1 | Tudor dom | Tdrd1 | sp\|Q99MV | 2 | 2 | 1,9 |
| Q9D6Z1 | Q9D6Z1 | Nucleolar p | Nop56 | sp\|Q9D6Z1 | 40 | 40 | 74,7 |
| Q9JKC7 | Q9JKC7 | AP-4 comp | Ap4m1 | sp\|Q9JKC7 | 5 | 5 | 14,3 |
| P63242 | P63242 | Eukaryotic | Eif5a | sp\|P63242 | 18 | 9 | 88,3 |
| Q8BLR9 | Q8BLR9 | Hypoxia-in | Hif1an | sp\|Q8BLR9 | 6 | 6 | 24,9 |
| Q8CCB4 | Q8CCB4 | Vacuolar p | Vps53 | sp\|Q8CCB4 | 21 | 21 | 33,9 |

| | | | | | | | |
|---|---|---|---|---|---|---|---|
| P24472 | P24472 | Glutathion | Gsta4 | sp\|P24472 | 8 | 7 | 36 |
| Q9Z1B5 | Q9Z1B5 | Mitotic spi | Mad2l1 | sp\|Q9Z1B5 | 8 | 8 | 43,9 |
| F8VQB6 | F8VQB6 | Unconvent | Myo10 | sp\|F8VQB6 | 9 | 9 | 5,2 |
| Q01768 | Q01768 | Nucleoside | Nme2 | sp\|Q01768 | 14 | 8 | 84,2 |
| Q8BQU3 | Q8BQU3 | Succinate d | Sdhaf3 | sp\|Q8BQU | 2 | 2 | 28 |
| P16254 | P16254 | Signal reco | Srp14 | sp\|P16254 | 5 | 5 | 42,7 |
| Q8BN57 | Q8BN57 | Protein C3orf33 homo | | sp\|Q8BN5 | 4 | 4 | 18 |
| Q6ZQL4 | Q6ZQL4 | WD repeat | Wdr43 | sp\|Q6ZQL4 | 22 | 22 | 51,3 |
| P58854 | P58854 | Gamma-tu | Tubgcp3 | sp\|P58854 | 31 | 31 | 45,5 |
| Q68FG3 | Q68FG3 | Protein SP | Spty2d1 | sp\|Q68FG3 | 10 | 10 | 16,4 |
| Q5NCQ5 | Q5NCQ5 | Diphthami | Dph1 | sp\|Q5NCQ | 7 | 7 | 17,6 |
| Q921I2 | Q921I2 | Kelch dom | Klhdc4 | sp\|Q921I2 | 16 | 16 | 29,5 |
| Q91YQ5 | Q91YQ5 | Dolichyl-di | Rpn1 | sp\|Q91YQ | 39 | 39 | 67,6 |
| Q8BHL5;Q | Q8BHL5 | Engulfmen | Elmo2 | sp\|Q8BHL5 | 13 | 13 | 27,2 |
| Q9JHI7 | Q9JHI7 | Exosome c | Exosc9 | sp\|Q9JHI7 | 10 | 10 | 33,8 |
| Q9QXT1 | Q9QXT1 | Dual adapt | Dapp1 | sp\|Q9QXT | 3 | 3 | 16,8 |
| Q8C3R1 | Q8C3R1 | BRCA1-ass | Brat1 | sp\|Q8C3R1 | 10 | 10 | 16,7 |
| O88448 | O88448 | Kinesin ligh | Klc2 | sp\|O88448 | 19 | 12 | 42,4 |
| Q9JHT5 | Q9JHT5 | AMME syn | Ammecr1 | sp\|Q9JHT5 | 3 | 2 | 11 |
| Q9CQJ4 | Q9CQJ4 | E3 ubiquiti | Rnf2 | sp\|Q9CQJ4 | 11 | 9 | 43,8 |
| Q9Z1K6 | Q9Z1K6 | E3 ubiquiti | Arih2 | sp\|Q9Z1K6 | 9 | 9 | 23,6 |
| Q8BFZ3 | Q8BFZ3 | Beta-actin- | Actbl2 | sp\|Q8BFZ3 | 8 | 3 | 29,8 |
| Q920R0 | Q920R0 | Alsin | Als2 | sp\|Q920R0 | 5 | 5 | 5,1 |
| Q9DCM2 | Q9DCM2 | Glutathion | Gstk1 | sp\|Q9DCM | 2 | 2 | 11,9 |
| Q5U419 | Q5U419 | Major facil | Mfsd3 | sp\|Q5U419 | 2 | 2 | 5,6 |
| Q80W49 | Q80W49 | Beta/gamr | Crybg3 | sp\|Q80W4 | 3 | 3 | 4,2 |
| Q9D020 | Q9D020 | Cytosolic 5 | Nt5c3a | sp\|Q9D020 | 14 | 14 | 52 |
| Q8BMG7 | Q8BMG7 | Rab3 GTPa | Rab3gap2 | sp\|Q8BMG | 31 | 31 | 28,3 |
| Q3UH60 | Q3UH60 | Disco-inter | Dip2b | sp\|Q3UH6 | 22 | 20 | 19,3 |
| Q9D1F5 | Q9D1F5 | UPF0544 protein C5or | | sp\|Q9D1F5 | 2 | 2 | 8,4 |
| Q60809 | Q60809 | CCR4-NOT | Cnot7 | sp\|Q60809 | 8 | 8 | 45,6 |
| B2RXC1 | B2RXC1 | Trafficking | Trappc11 | sp\|B2RXC1 | 21 | 21 | 27,4 |
| P61027 | P61027 | Ras-relate | Rab10 | sp\|P61027 | 15 | 13 | 65,5 |
| Q5U458 | Q5U458 | DnaJ homo | Dnajc11 | sp\|Q5U458 | 15 | 15 | 34,3 |
| Q9DAW6 | Q9DAW6 | U4/U6 sma | Prpf4 | sp\|Q9DAW | 23 | 23 | 59,7 |
| Q9D708 | Q9D708 | | S100a16 | sp\|Q9D708 | 2 | 2 | 16,1 |
| Q99JZ0 | Q99JZ0 | Syntenin-2 | Sdcbp2 | sp\|Q99JZ0 | 1 | 1 | 3,8 |
| Q8BX65 | Q8BX65 | MyoD fam | Mdfic | sp\|Q8BX65 | 1 | 1 | 7,3 |
| Q91Z22 | Q91Z22 | Porimin | Tmem123 | sp\|Q91Z22 | 1 | 1 | 4,6 |
| O09000 | O09000 | Nuclear re | Ncoa3 | sp\|O09000 | 9 | 9 | 8,9 |
| Q8R3H9 | Q8R3H9 | Tetratricop | Ttc4 | sp\|Q8R3H9 | 4 | 4 | 13,2 |
| P15379 | P15379 | CD44 antig | Cd44 | sp\|P15379 | 13 | 13 | 22,5 |
| O55013 | O55013 | Trafficking | Trappc3 | sp\|O55013 | 5 | 5 | 30,6 |
| Q80Y20 | Q80Y20 | Alkylated D | Alkbh8 | sp\|Q80Y20 | 4 | 4 | 8,3 |
| Q9QVP9 | Q9QVP9 | Protein-tyr | Ptk2b | sp\|Q9QVP | 4 | 3 | 4,8 |
| O54790 | O54790 | Transcripti | Mafg | sp\|O54790 | 6 | 5 | 33,3 |
| Q9Z1R4 | Q9Z1R4 | Uncharacte | D17h6s53e | sp\|Q9Z1R4 | 4 | 4 | 14,7 |
| P56959 | P56959 | RNA-bindir | Fus | sp\|P56959 | 12 | 12 | 23,4 |
| Q9DBC0 | Q9DBC0 | Selenoprot | Selo | sp\|Q9DBC | 3 | 3 | 6,7 |
| Q8K2H6 | Q8K2H6 | Anaphase- | Anapc10 | sp\|Q8K2H | 5 | 5 | 41,1 |

| | | | | | | | |
|---|---|---|---|---|---|---|---|
| Q80TZ9 | Q80TZ9 | Arginine-gl | Rere | sp\|Q80TZ9 | 7 | 7 | 7,3 |
| P97454 | P97454 | Mothers ag | Smad5 | sp\|P97454 | 6 | 2 | 17 |
| Q921Z5 | Q921Z5 | Tumor neo | Tnfaip8 | sp\|Q921Z5 | 3 | 3 | 16,7 |
| Q505D1 | Q505D1 | Serine/thre | Ankrd28 | sp\|Q505D1 | 18 | 18 | 26,3 |
| P97471 | P97471 | Mothers ag | Smad4 | sp\|P97471 | 14 | 14 | 36,7 |
| Q9R0Q3 | Q9R0Q3 | Transmem | Tmed2 | sp\|Q9R0Q | 7 | 7 | 38,8 |
| Q8BK58 | Q8BK58 | HSPB1-ass | Hspbap1 | sp\|Q8BK58 | 10 | 10 | 30 |
| Q8BJZ4 | Q8BJZ4 | 28S ribosc | Mrps35 | sp\|Q8BJZ4 | 11 | 11 | 51,9 |
| Q9DAU1 | Q9DAU1 | Protein ca | Cnpy3 | sp\|Q9DAU | 6 | 6 | 23,6 |
| Q8VC85 | Q8VC85 | U6 snRNA- | Lsm1 | sp\|Q8VC85 | 4 | 4 | 44,4 |
| Q9Z0R6 | Q9Z0R6 | Intersectin | Itsn2 | sp\|Q9Z0R6 | 34 | 31 | 27 |
| Q9CQA6 | Q9CQA6 | Coiled-coil | Chchd1 | sp\|Q9CQA | 2 | 2 | 22,9 |
| Q9CPW3 | Q9CPW3 | 39S ribosc | Mrpl54 | sp\|Q9CPW | 3 | 3 | 43,7 |
| A2A6T1 | A2A6T1 | Cerebellar | Cdr2l | sp\|A2A6T1 | 2 | 2 | 4,1 |
| Q8K3J1 | Q8K3J1 | NADH deh | Ndufs8 | sp\|Q8K3J1 | 4 | 4 | 21,2 |
| Q91YD9 | Q91YD9 | Neural Wis | Wasl | sp\|Q91YD9 | 11 | 11 | 35,5 |
| Q06185 | Q06185 | ATP syntha | Atp5i | sp\|Q06185 | 4 | 4 | 52,1 |
| Q6DG52 | Q6DG52 | Protein Ch | Churc1 | sp\|Q6DG5 | 2 | 2 | 27,7 |
| Q9ER81 | Q9ER81 | Torsin-1A-i | Tor1aip2 | sp\|Q9ER81 | 3 | 3 | 32,1 |
| Q8BMQ2 | Q8BMQ2 | General tra | Gtf3c4 | sp\|Q8BMC | 28 | 28 | 43,2 |
| P47802 | P47802 | Metaxin-1 | Mtx1 | sp\|P47802 | 11 | 11 | 32,5 |
| Q8R3P9 | Q8R3P9 | Ankyrin re | Ankrd32 | sp\|Q8R3P9 | 4 | 4 | 4,7 |
| Q6Y5D8 | Q6Y5D8 | Rho GTPas | Arhgap10 | sp\|Q6Y5D8 | 14 | 14 | 24,2 |
| Q3TV65 | Q3TV65 | MPN doma | Mpnd | sp\|Q3TV65 | 2 | 2 | 5,1 |
| Q149L6 | Q149L6 | DnaJ homo | Dnajb14 | sp\|Q149L6 | 3 | 3 | 7,9 |
| P31996 | P31996 | Macrosiali | Cd68 | sp\|P31996 | 2 | 2 | 5,5 |
| O89086 | O89086 | RNA-bindir | Rbm3 | sp\|O89086 | 6 | 6 | 55,6 |
| P14404 | P14404 | MDS1 and | Mecom | sp\|P14404 | 2 | 2 | 3,9 |
| Q9QZS3 | Q9QZS3 | Protein nu | Numb | sp\|Q9QZS3 | 3 | 3 | 6,6 |
| P62869 | P62869 | Transcripti | Tceb2 | sp\|P62869 | 14 | 14 | 91,5 |
| Q5U5Q9 | Q5U5Q9 | BRCA1-A c | Uimc1 | sp\|Q5U5Q | 8 | 8 | 16,9 |
| P70695 | P70695 | Fructose-1 | Fbp2 | sp\|P70695 | 2 | 2 | 6,5 |
| Q8BGK6 | Q8BGK6 | Y+L amino | Slc7a6 | sp\|Q8BGK | 5 | 4 | 12,6 |
| Q8BGX7;R | Q8BGX7 | DNA repair | Spidr | sp\|Q8BGX | 5 | 5 | 5,1 |
| O35887 | O35887 | Calumenin | Calu | sp\|O35887 | 20 | 20 | 61,9 |
| Q04692 | Q04692 | SWI/SNF-r | Smarcad1 | sp\|Q04692 | 23 | 23 | 31,4 |
| Q91YY4 | Q91YY4 | ATP syntha | Atpaf2 | sp\|Q91YY4 | 9 | 9 | 46,7 |
| Q6PFQ7 | Q6PFQ7 | Ras GTPase | Rasa4 | sp\|Q6PFQ7 | 23 | 23 | 39,7 |
| P20060 | P20060 | Beta-hexos | Hexb | sp\|P20060 | 7 | 7 | 16,4 |
| P60762 | P60762 | Mortality f | Morf4l1 | sp\|P60762 | 14 | 12 | 47 |
| Q61205 | Q61205 | Platelet-ac | Pafah1b3 | sp\|Q61205 | 11 | 11 | 60,3 |
| Q6ZPV2 | Q6ZPV2 | DNA helica | Ino80 | sp\|Q6ZPV2 | 16 | 16 | 12,4 |
| Q9QXD1 | Q9QXD1 | Peroxisom | Acox2 | sp\|Q9QXD | 2 | 2 | 3,8 |
| Q9CQZ0 | Q9CQZ0 | ORM1-like | Ormdl2 | sp\|Q9CQZ0 | 5 | 3 | 38,6 |
| Q9ES89 | Q9ES89 | Exostosin-l | Extl2 | sp\|Q9ES89 | 5 | 5 | 20,6 |
| Q9QY73 | Q9QY73 | Transmem | Tmem59 | sp\|Q9QY73 | 7 | 7 | 36,5 |
| Q810D6 | Q810D6 | Glutamate | Grwd1 | sp\|Q810D6 | 16 | 16 | 57,8 |
| Q8K2T4 | Q8K2T4 | Ubiquinol- | Uqcc3 | sp\|Q8K2T4 | 1 | 1 | 11,2 |
| Q8JZL3 | Q8JZL3 | Thiamine-t | Thtpa | sp\|Q8JZL3 | 3 | 3 | 17,4 |
| Q8BXK8 | Q8BXK8 | Arf-GAP wi | Agap1 | sp\|Q8BXK8 | 9 | 7 | 14,6 |

| | | | | | | | |
|---|---|---|---|---|---|---|---|
| Q8BU14 | Q8BU14 | Translocati | Sec62 | sp\|Q8BU14 | 6 | 6 | 17,6 |
| Q6T264 | Q6T264 | Mastermin | Maml1 | sp\|Q6T264 | 2 | 2 | 3,9 |
| Q60848 | Q60848 | Lymphocyt | Hells | sp\|Q60848 | 30 | 30 | 40,3 |
| CON__P07 | CON__P07477 | | | | 2 | 2 | 12,1 |
| Q3UJV1 | Q3UJV1 | Coiled-coil | Ccdc61 | sp\|Q3UJV1 | 6 | 6 | 14,7 |
| Q9ESK4 | Q9ESK4 | Inhibitor o | Ing2 | sp\|Q9ESK4 | 4 | 4 | 21,7 |
| Q99KR6 | Q99KR6 | E3 ubiquiti | Rnf34 | sp\|Q99KR6 | 2 | 2 | 8 |
| E9Q9D5 | E9Q9D5 | Rab-like pr | Rabl2 | sp\|E9Q9D5 | 3 | 3 | 18,8 |
| Q61235;Q6 | Q61235 | Beta-2-syn | Sntb2 | sp\|Q61235 | 18 | 16 | 41,2 |
| P62069 | P62069 | Ubiquitin c | Usp46 | sp\|P62069 | 5 | 2 | 18,9 |
| Q62086 | Q62086 | Serum para | Pon2 | sp\|Q62086 | 10 | 10 | 37,6 |
| Q9R0Q9 | Q9R0Q9 | Mannose-P | Mpdu1 | sp\|Q9R0Q | 3 | 3 | 13,8 |
| P48428 | P48428 | Tubulin-sp | Tbca | sp\|P48428 | 10 | 10 | 49,1 |
| Q9CZ82 | Q9CZ82 | Mediator c | Med18 | sp\|Q9CZ82 | 3 | 3 | 16,3 |
| Q9DCT2 | Q9DCT2 | NADH deh | Ndufs3 | sp\|Q9DCT2 | 9 | 9 | 30,8 |
| Q9CR23 | Q9CR23 | Transmem | Tmem9 | sp\|Q9CR23 | 3 | 2 | 26,2 |
| Q9CYI0 | Q9CYI0 | Protein Njmu-R1 | | sp\|Q9CYI0 | 8 | 8 | 33,8 |
| P27661 | P27661 | Histone H2 | H2afx | sp\|P27661 | 10 | 3 | 65 |
| P27659 | P27659 | 60S riboso | Rpl3 | sp\|P27659 | 24 | 24 | 52,9 |
| Q4VA53 | Q4VA53 | Sister chro | Pds5b | sp\|Q4VA53 | 50 | 45 | 37,2 |
| Q9R1P4 | Q9R1P4 | Proteasom | Psma1 | sp\|Q9R1P4 | 16 | 16 | 62,4 |
| Q9EQH4 | Q9EQH4 | Transcripti | Taf8 | sp\|Q9EQH | 5 | 5 | 23,7 |
| Q8C726 | Q8C726 | BTB/POZ d | Btbd9 | sp\|Q8C726 | 3 | 3 | 8,2 |
| Q9JHC9 | Q9JHC9 | ETS-relatec | Elf2 | sp\|Q9JHC9 | 12 | 12 | 34,4 |
| Q78S06 | Q78S06 | Protein SYS | Sys1 | sp\|Q78S06 | 2 | 2 | 19,9 |
| Q923T9 | Q923T9 | Calcium/ca | Camk2g | sp\|Q923T9 | 14 | 9 | 39,5 |
| Q8K0B2 | Q8K0B2 | Probable ly | Lmbrd1 | sp\|Q8K0B2 | 1 | 1 | 3,4 |
| Q922Q8 | Q922Q8 | Leucine-ric | Lrrc59 | sp\|Q922Q | 14 | 14 | 52,4 |
| Q9WV76 | Q9WV76 | AP-4 comp | Ap4b1 | sp\|Q9WV7 | 8 | 8 | 14,9 |
| Q9D2V5 | Q9D2V5 | Protein AA | Aar2 | sp\|Q9D2V | 9 | 9 | 34,4 |
| Q91YI1 | Q91YI1 | Autophagy | Atg13 | sp\|Q91YI1 | 3 | 3 | 9,1 |
| Q9D6K9 | Q9D6K9 | Ceramide s | Cers5 | sp\|Q9D6K9 | 6 | 6 | 21,3 |
| Q14AM7 | Q14AM7 | UPF0472 protein C16o | | sp\|Q14AM | 1 | 1 | 6,5 |
| Q9DCC8 | Q9DCC8 | Mitochond | Tomm20 | sp\|Q9DCC | 4 | 4 | 35,9 |
| O08638 | O08638 | Myosin-11 | Myh11 | sp\|O08638 | 56 | 40 | 30,2 |
| Q99P58 | Q99P58 | Ras-relatec | Rab27b | sp\|Q99P58 | 5 | 4 | 28,9 |
| O35648 | O35648 | Centrin-3 | Cetn3 | sp\|O35648 | 2 | 2 | 19,2 |
| Q04519 | Q04519 | Sphingomy | Smpd1 | sp\|Q04519 | 2 | 2 | 3,3 |
| Q8CIB9 | Q8CIB9 | N-acetyltra | Esco2 | sp\|Q8CIB9 | 6 | 6 | 13,5 |
| Q8CBY3 | Q8CBY3 | Leukocyte | Leng8 | sp\|Q8CBY3 | 11 | 11 | 22,3 |
| Q3KNJ2 | Q3KNJ2 | Non-homo | Nhej1 | sp\|Q3KNJ2 | 1 | 1 | 10,8 |
| Q99PM3 | Q99PM3 | Transcripti | Gtf2a1 | sp\|Q99PM | 1 | 1 | 2,9 |
| Q99K41 | Q99K41 | EMILIN-1 | Emilin1 | sp\|Q99K41 | 3 | 3 | 3,9 |
| Q9D8M7 | Q9D8M7 | PHD finger | Phf10 | sp\|Q9D8M | 6 | 6 | 17,1 |
| Q91V76 | Q91V76 | Ester hydrolase C11or | | sp\|Q91V76 | 12 | 12 | 46,7 |
| Q6IRU7 | Q6IRU7 | Centrosom | Cep78 | sp\|Q6IRU7 | 3 | 3 | 6,7 |
| P46061 | P46061 | Ran GTPas | Rangap1 | sp\|P46061 | 30 | 30 | 53,5 |
| P14426 | P14426 | H-2 class I | H2-D1 | sp\|P14426 | 4 | 1 | 14,6 |
| Q921I0 | Q921I0 | ORM1-like | Ormdl1 | sp\|Q921I0 | 3 | 2 | 28,8 |
| P28700 | P28700 | Retinoic ac | Rxra | sp\|P28700 | 7 | 3 | 17,8 |

| | | | | | | | |
|---|---|---|---|---|---|---|---|
| Q9Z2U1 | Q9Z2U1 | Proteasom | Psma5 | sp\|Q9Z2U1 | 14 | 14 | 69,3 |
| Q8VCA6 | Q8VCA6 | Transmem | Tmem161a | sp\|Q8VCA | 3 | 3 | 7,1 |
| P60867 | P60867 | 40S riboso | Rps20 | sp\|P60867 | 6 | 6 | 37 |
| P14131 | P14131 | 40S riboso | Rps16 | sp\|P14131 | 13 | 13 | 70,5 |
| P40338 | P40338 | Von Hippe | Vhl | sp\|P40338 | 2 | 2 | 12,7 |
| Q80V94 | Q80V94 | AP-4 comp | Ap4e1 | sp\|Q80V9 | 13 | 13 | 17,6 |
| Q8R3F5 | Q8R3F5 | Malonyl-C | Mcat | sp\|Q8R3F | 14 | 14 | 52,5 |
| Q6ZQ03 | Q6ZQ03 | Formin-bin | Fnbp4 | sp\|Q6ZQ0 | 14 | 14 | 14,2 |
| Q8CIF4 | Q8CIF4 | Biotinidase | Btd | sp\|Q8CIF4 | 2 | 2 | 5,4 |
| Q62217;Q6 | Q62217 | Semaphori | Sema5a | sp\|Q62217 | 3 | 3 | 3,6 |
| Q8C2P3 | Q8C2P3 | tRNA-dihy | Dus1l | sp\|Q8C2P | 3 | 3 | 6,3 |
| Q8CIM3 | Q8CIM3 | D-2-hydro | D2hgdh | sp\|Q8CIM | 8 | 8 | 21,3 |
| Q9CWK3 | Q9CWK3 | CD2 antige | Cd2bp2 | sp\|Q9CWK | 11 | 11 | 53,2 |
| Q9DBS2 | Q9DBS2 | Tumor pro | Tprg1l | sp\|Q9DBS | 6 | 6 | 35,7 |
| Q9CZ28 | Q9CZ28 | Vacuolar-s | Snf8 | sp\|Q9CZ2 | 6 | 6 | 31 |
| O70494 | O70494 | Transcripti | Sp3 | sp\|O7049 | 1 | 1 | 3,6 |
| Q9WVL6 | Q9WVL6 | Exostosin-l | Extl3 | sp\|Q9WVL | 6 | 6 | 10 |
| Q01721 | Q01721 | Growth arr | Gas1 | sp\|Q0172 | 2 | 2 | 12 |
| Q6ZQK5 | Q6ZQK5 | Arf-GAP wi | Acap2 | sp\|Q6ZQK | 11 | 11 | 21,8 |
| Q9D0U6 | Q9D0U6 | Repressor | Maf1 | sp\|Q9D0U | 5 | 5 | 41,1 |
| Q8C7H1 | Q8C7H1 | Methylmal | Mmaa | sp\|Q8C7H | 7 | 7 | 22,2 |
| O35450 | O35450 | FK506-bind | Fkbpl | sp\|O3545 | 4 | 4 | 14,4 |
| Q8VE62 | Q8VE62 | Polyadenyl | Paip1 | sp\|Q8VE6 | 11 | 11 | 34,2 |
| Q501J7 | Q501J7 | Phosphata | Phactr4 | sp\|Q501J7 | 15 | 15 | 35,3 |
| Q9QXE0 | Q9QXE0 | 2-hydroxya | Hacl1 | sp\|Q9QXE | 2 | 2 | 5 |
| B2RVL6 | B2RVL6 | Zinc finger | Zcchc24 | sp\|B2RVL6 | 1 | 1 | 7,1 |
| P98195 | P98195 | Probable p | Atp9b | sp\|P98195 | 12 | 10 | 12,4 |
| O88878 | O88878 | AN1-type z | Zfand5 | sp\|O8887 | 3 | 3 | 25,4 |
| Q9CQI3;Q9 | Q9CQI3 | Glia matur | Gmfb | sp\|Q9CQI3 | 6 | 6 | 65,5 |
| Q2VPQ9 | Q2VPQ9 | Chromatin | Meaf6 | sp\|Q2VPQ | 7 | 7 | 38,7 |
| Q8R4Z4 | Q8R4Z4 | ETS translo | Etv3 | sp\|Q8R4Z | 3 | 3 | 10,1 |
| Q9DAX9 | Q9DAX9 | Amyloid pr | Appbp2 | sp\|Q9DAX | 7 | 7 | 17,9 |
| Q3UYV9 | Q3UYV9 | Nuclear ca | Ncbp1 | sp\|Q3UYV | 23 | 23 | 33,8 |
| P10852 | P10852 | 4F2 cell-su | Slc3a2 | sp\|P10852 | 22 | 22 | 49,8 |
| E9Q5K9 | E9Q5K9 | YTH domai | Ythdc1 | sp\|E9Q5K | 12 | 12 | 18,3 |
| Q0VBK2;C | Q0VBK2;CON_ | Keratin, ty | Krt80 | sp\|Q0VBK | 2 | 2 | 6 |
| Q6ZQK0 | Q6ZQK0 | Condensin | Ncapd3 | sp\|Q6ZQK | 31 | 31 | 26,8 |
| Q6PAM1 | Q6PAM1 | Alpha-taxil | Txlna | sp\|Q6PAM | 19 | 19 | 40,1 |
| Q8BZ09 | Q8BZ09 | Mitochond | Slc25a21 | sp\|Q8BZ0 | 1 | 1 | 4,4 |
| Q9DAM7 | Q9DAM7 | Transmem | Tmem263 | sp\|Q9DAM | 4 | 4 | 54,8 |
| Q8BFU3 | Q8BFU3 | RING finge | Rnf214 | sp\|Q8BFU | 14 | 14 | 31,3 |
| Q9DB91 | Q9DB91 | Mediator c | Med29 | sp\|Q9DB9 | 4 | 4 | 31,2 |
| Q9DB26 | Q9DB26 | Phytanoyl- | Phyhd1 | sp\|Q9DB2 | 1 | 1 | 10 |
| Q3TZX3 | Q3TZX3 | Solute carr | Slc25a33 | sp\|Q3TZX3 | 3 | 2 | 10,6 |
| Q5DTT3 | Q5DTT3 | Protein FA | Fam208b | sp\|Q5DTT | 17 | 17 | 12,4 |
| P35831 | P35831 | Tyrosine-p | Ptpn12 | sp\|P35831 | 19 | 19 | 35,5 |
| Q6P1D7 | Q6P1D7 | Structure-s | Slx4 | sp\|Q6P1D | 4 | 4 | 4 |
| P81066 | P81066 | Iroquois-cl | Irx2 | sp\|P81066 | 1 | 1 | 2,7 |
| Q3THK7 | Q3THK7 | GMP synth | Gmps | sp\|Q3THK | 40 | 40 | 72,3 |
| P36993 | P36993 | Protein ph | Ppm1b | sp\|P36993 | 13 | 10 | 51,5 |

| | | | | | | | |
|---|---|---|---|---|---|---|---|
| Q5U430 | Q5U430 | E3 ubiquiti | Ubr3 | sp\|Q5U430 | 9 | 9 | 6,2 |
| O70251 | O70251 | Elongation | Eef1b | sp\|O70251 | 11 | 9 | 66,7 |
| Q9QZK2 | Q9QZK2 | Breast can | Bcar3 | sp\|Q9QZK2 | 6 | 6 | 9,6 |
| O09167 | O09167 | 60S riboso | Rpl21 | sp\|O09167 | 6 | 6 | 32,5 |
| Q8QZX2 | Q8QZX2 | HAUS augr | Haus3 | sp\|Q8QZX2 | 18 | 18 | 42,6 |
| Q9D2R8 | Q9D2R8 | 28S riboso | Mrps33 | sp\|Q9D2R8 | 3 | 3 | 26,4 |
| Q8BGT6 | Q8BGT6 | MICAL-like | Micall1 | sp\|Q8BGT6 | 9 | 9 | 14,8 |
| Q8BPM2 | Q8BPM2 | Mitogen-a | Map4k5 | sp\|Q8BPM | 23 | 23 | 35,2 |
| Q8K2L8 | Q8K2L8 | Trafficking | Trappc12 | sp\|Q8K2L8 | 14 | 14 | 29,9 |
| Q60598 | Q60598 | Src substra | Cttn | sp\|Q60598 | 21 | 21 | 42,1 |
| Q3UA06 | Q3UA06 | Pachytene | Trip13 | sp\|Q3UA0 | 14 | 14 | 38 |
| Q9DC33 | Q9DC33 | High mobil | Hmg20a | sp\|Q9DC33 | 4 | 4 | 16,8 |
| Q80VJ2 | Q80VJ2 | Steroid rec | Sra1 | sp\|Q80VJ2 | 3 | 3 | 22,8 |
| Q9CQM5 | Q9CQM5 | Thioredoxi | Txndc17 | sp\|Q9CQM | 5 | 5 | 55,3 |
| Q99LX5 | Q99LX5 | Multiple m | Mmtag2 | sp\|Q99LX5 | 8 | 8 | 34,2 |
| Q9D1Q6 | Q9D1Q6 | Endoplasm | Erp44 | sp\|Q9D1Q | 18 | 18 | 62,3 |
| Q60680 | Q60680 | Inhibitor o | Chuk | sp\|Q60680 | 12 | 11 | 20,4 |
| Q8BFR4 | Q8BFR4 | N-acetylglu | Gns | sp\|Q8BFR4 | 17 | 17 | 43,8 |
| P05202 | P05202 | Aspartate a | Got2 | sp\|P05202 | 29 | 29 | 65,6 |
| Q9DCH2 | Q9DCH2 | Ribonuclea | Pop7 | sp\|Q9DCH | 1 | 1 | 10,7 |
| P70340 | P70340 | Mothers ag | Smad1 | sp\|P70340 | 9 | 4 | 26,2 |
| Q62447 | Q62447 | Cyclin-C | Ccnc | sp\|Q62447 | 3 | 3 | 13,4 |
| P35123;Q9 | P35123 | Ubiquitin c | Usp4 | sp\|P35123 | 39 | 37 | 50,1 |
| Q8K201 | Q8K201 | Keratinocy | Kct2 | sp\|Q8K201 | 3 | 3 | 13,1 |
| P97351 | P97351 | 40S riboso | Rps3a | sp\|P97351 | 21 | 21 | 68,6 |
| CON__P35 | CON__P35527 | | | | 21 | 20 | 44,1 |
| Q9D1R2 | Q9D1R2 | Protein KT | Kti12 | sp\|Q9D1R2 | 12 | 12 | 46,7 |
| Q9D4J7 | Q9D4J7 | PHD finger | Phf6 | sp\|Q9D4J7 | 8 | 8 | 31 |
| Q8C0Z1 | Q8C0Z1 | Protein ITF | Itfg3 | sp\|Q8C0Z1 | 8 | 8 | 19,6 |
| Q9CQV5 | Q9CQV5 | 28S riboso | Mrps24 | sp\|Q9CQV | 4 | 4 | 24 |
| Q9JL70 | Q9JL70 | Fanconi an | Fanca | sp\|Q9JL70 | 10 | 10 | 8,1 |
| P63073 | P63073 | Eukaryotic | Eif4e | sp\|P63073 | 10 | 10 | 43,8 |
| Q9CY28 | Q9CY28 | GTP-bindir | Gtpbp8 | sp\|Q9CY28 | 5 | 5 | 23,9 |
| Q9DAA6 | Q9DAA6 | Exosome c | Exosc1 | sp\|Q9DAA | 6 | 6 | 40 |
| Q8VEB2 | Q8VEB2 | Protein sal | Sav1 | sp\|Q8VEB2 | 6 | 6 | 27,5 |
| B1AXD8 | B1AXD8 | Akirin-2 | Akirin2 | sp\|B1AXD8 | 4 | 4 | 24,4 |
| Q9R1A8 | Q9R1A8 | E3 ubiquiti | Rfwd2 | sp\|Q9R1A8 | 3 | 3 | 5,6 |
| P62080 | P62080 | Tetraspani | Tspan5 | sp\|P62080 | 1 | 1 | 6 |
| Q99J72 | Q99J72 | DNA dC->d | Apobec3 | sp\|Q99J72 | 9 | 9 | 19,3 |
| B1AVD1 | B1AVD1 | | Xpnpep2 | sp\|B1AVD1 | 1 | 1 | 1,8 |
| Q9JHE3 | Q9JHE3 | Neutral ce | Asah2 | sp\|Q9JHE3 | 4 | 4 | 7,4 |
| Q3UMM4 | Q3UMM4 | Cyclin-dep | Cdk10 | sp\|Q3UMI | 2 | 2 | 6,7 |
| Q3U2V3 | Q3U2V3 | 8-oxo-dGD | Nudt18 | sp\|Q3U2V | 1 | 1 | 5 |
| Q9DB43 | Q9DB43 | Zinc finger | Zfpl1 | sp\|Q9DB43 | 6 | 6 | 25,8 |
| Q8CGE9 | Q8CGE9 | Regulator o | Rgs12 | sp\|Q8CGE9 | 4 | 4 | 3,3 |
| O88852 | O88852 | Testis-spec | Tspyl1 | sp\|O88852 | 3 | 3 | 7,4 |
| Q80UG8 | Q80UG8 | Tubulin po | Ttll4 | sp\|Q80UG | 2 | 2 | 2,3 |
| P14430;P1 | P14430;P1442 | H-2 class I | H2-Q8;H2- | sp\|P14430 | 6 | 2 | 20,2 |
| Q8BK72 | Q8BK72 | 28S riboso | Mrps27 | sp\|Q8BK72 | 18 | 18 | 43,6 |
| Q9D0L8 | Q9D0L8 | mRNA cap | Rnmt | sp\|Q9D0L8 | 19 | 19 | 45,6 |

| | | | | | | | |
|---|---|---|---|---|---|---|---|
| Q5SSN7 | Q5SSN7 | Vesicle tra | Sft2d1 | sp\|Q5SSN7 | 1 | 1 | 17 |
| Q80TQ5 | Q80TQ5 | Pleckstrin I | Plekhm2 | sp\|Q80TQ | 6 | 6 | 7,4 |
| Q62159 | Q62159 | Rho-relate | Rhoc | sp\|Q62159 | 10 | 2 | 65,8 |
| Q9WTV7 | Q9WTV7 | E3 ubiquiti | Rlim | sp\|Q9WTV | 8 | 8 | 23,2 |
| P43247 | P43247 | DNA mism | Msh2 | sp\|P43247 | 38 | 38 | 52,2 |
| P46638;Q9 | P46638 | Ras-related | Rab11b | sp\|P46638 | 14 | 3 | 73,9 |
| Q3UIK4 | Q3UIK4 | N6-adenos | Mettl14 | sp\|Q3UIK4 | 3 | 3 | 11 |
| CON__P13 | CON__P13647 | | | | 10 | 1 | 15,8 |
| Q8CHK4 | Q8CHK4 | Histone ac | Kat5 | sp\|Q8CHK | 8 | 8 | 26,9 |
| Q80V26 | Q80V26 | Inositol mo | Impad1 | sp\|Q80V26 | 9 | 9 | 38,8 |
| Q76N33 | Q76N33 | AMSH-like | Stambpl1 | sp\|Q76N33 | 9 | 9 | 26,6 |
| Q62108 | Q62108 | Disks large | Dlg4 | sp\|Q62108 | 1 | 1 | 1,4 |
| Q99J77 | Q99J77 | | Nans | sp\|Q99J77 | 16 | 16 | 53,8 |
| Q7TQC5 | Q7TQC5 | Aprataxin | Aptx | sp\|Q7TQC | 4 | 4 | 14,6 |
| P04202 | P04202 | Transformi | Tgfb1 | sp\|P04202 | 3 | 3 | 10,8 |
| Q9CRB0 | Q9CRB0 | Sorting ne | Snx24 | sp\|Q9CRB0 | 3 | 3 | 21,9 |
| Q3U3N6 | Q3U3N6 | AP-4 comp | Enthd2 | sp\|Q3U3N | 3 | 3 | 7,9 |
| P97461 | P97461 | 40S riboso | Rps5 | sp\|P97461 | 12 | 12 | 47,5 |
| P40694 | P40694 | DNA-bindi | Ighmbp2 | sp\|P40694 | 8 | 8 | 12,6 |
| Q9DB15 | Q9DB15 | 39S riboso | Mrpl12 | sp\|Q9DB15 | 8 | 8 | 46,8 |
| Q2EMV9 | Q2EMV9 | Poly [ADP- | Parp14 | sp\|Q2EMV | 3 | 3 | 2 |
| Q91XB0 | Q91XB0 | Three-prim | Trex1 | sp\|Q91XB0 | 9 | 9 | 35 |
| P31786 | P31786 | Acyl-CoA-b | Dbi | sp\|P31786 | 8 | 8 | 70,1 |
| Q9D1M0 | Q9D1M0 | Protein SE | Sec13 | sp\|Q9D1M | 12 | 12 | 58,4 |
| P28322;P4 | P28322 | ETS translo | Etv4 | sp\|P28322 | 5 | 5 | 14,2 |
| Q9CX66 | Q9CX66 | Uncharacte | D10Wsu10 | sp\|Q9CX66 | 2 | 2 | 24,9 |
| Q9CQE6 | Q9CQE6 | Histone ch | Asf1a | sp\|Q9CQE | 6 | 6 | 44,1 |
| P53368 | P53368 | 7,8-dihydr | Nudt1 | sp\|P53368 | 2 | 2 | 17,9 |
| Q8BL06 | Q8BL06 | Inactive ub | Usp54 | sp\|Q8BL06 | 1 | 1 | 1 |
| Q91XL2 | Q91XL2 | Ligand of N | Lnx2 | sp\|Q91XL2 | 2 | 2 | 4,8 |
| Q8BP71;Q9 | Q8BP71 | RNA bindir | Rbfox2 | sp\|Q8BP71 | 9 | 9 | 36,5 |
| O88668 | O88668 | Protein CR | Creg1 | sp\|O88668 | 2 | 2 | 13,2 |
| P54279 | P54279 | Mismatch | Pms2 | sp\|P54279 | 7 | 7 | 10,5 |
| O35900 | O35900 | U6 snRNA- | Lsm2 | sp\|O35900 | 6 | 6 | 75,8 |
| Q5HZG4 | Q5HZG4 | Transcripti | Taf3 | sp\|Q5HZG | 7 | 7 | 10,1 |
| Q0VGM9 | Q0VGM9 | Regulator | Rtel1 | sp\|Q0VGM | 7 | 7 | 8,1 |
| Q4VBD9 | Q4VBD9 | GDNF-indu | Gzf1 | sp\|Q4VBD | 6 | 6 | 10,6 |
| Q9D9G3 | Q9D9G3 | Cysteine-ri | Chic2 | sp\|Q9D9G | 3 | 3 | 21,8 |
| Q3U4G3 | Q3U4G3 | Xyloside xy | Xxylt1 | sp\|Q3U4G | 13 | 13 | 45,7 |
| Q62468 | Q62468 | Villin-1 | Vil1 | sp\|Q62468 | 12 | 12 | 18,1 |
| B2RRD7 | B2RRD7 | | Brpf1 | sp\|B2RRD7 | 8 | 8 | 9,2 |
| P60670 | P60670 | Nuclear pr | Nploc4 | sp\|P60670 | 30 | 30 | 60 |
| A6PWY4 | A6PWY4 | WD repeat | Wdr76 | sp\|A6PWY | 9 | 9 | 18,6 |
| P97440 | P97440 | Histone RN | Slbp | sp\|P97440 | 2 | 2 | 9,8 |
| Q9CQG2 | Q9CQG2 | Methyltran | Mettl16 | sp\|Q9CQG | 19 | 19 | 49,7 |
| P24788 | P24788 | Cyclin-dep | Cdk11b | sp\|P24788 | 17 | 16 | 22,2 |
| Q8CD10 | Q8CD10 | Calcium up | Micu2 | sp\|Q8CD10 | 12 | 12 | 33,3 |
| Q9CR70 | Q9CR70 | EKC/KEOPS | Lage3 | sp\|Q9CR70 | 2 | 2 | 22,3 |
| P62874 | P62874 | Guanine nu | Gnb1 | sp\|P62874 | 15 | 10 | 58,5 |
| Q9CQQ4 | Q9CQQ4 | Gem-assoc | Gemin2 | sp\|Q9CQQ | 8 | 8 | 42,8 |

| | | | | | | | |
|---|---|---|---|---|---|---|---|
| Q925F4 | Q925F4 | RING finge | Ubox5 | sp\|Q925F4 | 1 | 1 | 2,8 |
| P54726 | P54726 | UV excisio | Rad23a | sp\|P54726 | 11 | 8 | 40,8 |
| Q62376 | Q62376 | U1 small n | Snrnp70 | sp\|Q62376 | 19 | 19 | 51,8 |
| Q8BU11;Q | Q8BU11 | TOX high n | Tox4 | sp\|Q8BU1 | 8 | 8 | 15,8 |
| P70284 | P70284 | Homeobox | Tgif1 | sp\|P70284 | 1 | 1 | 5,1 |
| Q9CWN7 | Q9CWN7 | CCR4-NOT | Cnot11 | sp\|Q9CWN | 6 | 6 | 20,4 |
| Q91WE6 | Q91WE6 | Threonylca | Cdkal1 | sp\|Q91WE | 8 | 8 | 19,4 |
| Q8BLX4 | Q8BLX4 | GDP-fucos | Slc35c1 | sp\|Q8BLX4 | 3 | 3 | 13,2 |
| Q8BT07 | Q8BT07 | Centrosom | Cep55 | sp\|Q8BT07 | 20 | 20 | 39,8 |
| P62878 | P62878 | E3 ubiquiti | Rbx1 | sp\|P62878 | 4 | 4 | 30,6 |
| Q8R420;E9 | Q8R420 | ATP-bindin | Abca3 | sp\|Q8R420 | 12 | 12 | 9,6 |
| O88574 | O88574 | Histone de | Sap30 | sp\|O88574 | 3 | 3 | 13,6 |
| Q3TUA9 | Q3TUA9 | Protein O-I | Pomk | sp\|Q3TUA | 2 | 2 | 8,6 |
| Q6NXJ0 | Q6NXJ0 | Protein W\ | Wwc2 | sp\|Q6NXJ0 | 12 | 12 | 15,2 |
| P62309 | P62309 | Small nucle | Snrpg | sp\|P62309 | 3 | 3 | 26,3 |
| Q8BMJ3 | Q8BMJ3 | Eukaryotic | Eif1ax | sp\|Q8BMJ | 8 | 2 | 68,8 |
| Q9D824 | Q9D824 | Pre-mRNA | Fip1l1 | sp\|Q9D824 | 15 | 15 | 43,4 |
| O88543 | O88543 | COP9 signa | Cops3 | sp\|O88543 | 15 | 15 | 53,2 |
| P68037 | P68037 | Ubiquitin-c | Ube2l3 | sp\|P68037 | 10 | 10 | 69,5 |
| Q9DBR3 | Q9DBR3 | Armadillo r | Armc8 | sp\|Q9DBR | 18 | 18 | 31,6 |
| Q9D6S7 | Q9D6S7 | Ribosome- | Mrrf | sp\|Q9D6S7 | 7 | 7 | 43,9 |
| Q8R0A2 | Q8R0A2 | Zinc finger | Zbtb44 | sp\|Q8R0A2 | 2 | 2 | 8,2 |
| Q5U4C3 | Q5U4C3 | Splicing fac | Scaf1 | sp\|Q5U4C3 | 10 | 10 | 11,5 |
| Q8BX94 | Q8BX94 | Oxysterol-l | Osbpl2 | sp\|Q8BX94 | 8 | 8 | 26,4 |
| Q60980 | Q60980 | Krueppel-li | Klf3 | sp\|Q60980 | 3 | 3 | 10,5 |
| CON__Q1R | CON__Q1RMK2 | | | | 5 | 5 | 11,1 |
| P56485 | P56485 | Atypical ch | Ackr3 | sp\|P56485 | 1 | 1 | 3,9 |
| Q9CXK8 | Q9CXK8 | 60S riboso | Nip7 | sp\|Q9CXK8 | 11 | 11 | 77,8 |
| Q9JLJ1 | Q9JLJ1 | Selenoprot | Selk | sp\|Q9JLJ1 | 1 | 1 | 12,8 |
| Q9CQI9 | Q9CQI9 | Mediator c | Med30 | sp\|Q9CQI9 | 5 | 5 | 39,9 |
| P23906 | P23906 | Interferon | Irf2 | sp\|P23906 | 2 | 2 | 9,5 |
| P21279 | P21279 | Guanine nu | Gnaq | sp\|P21279 | 14 | 7 | 45,1 |
| A2AJQ3 | A2AJQ3 | Probable C | Dpy19l4 | sp\|A2AJQ3 | 5 | 5 | 9,8 |
| Q8BHJ9 | Q8BHJ9 | Pre-mRNA- | Slu7 | sp\|Q8BHJ9 | 13 | 13 | 34,5 |
| Q7JJ13 | Q7JJ13 | Bromodom | Brd2 | sp\|Q7JJ13 | 26 | 23 | 38,7 |
| Q9QWZ1 | Q9QWZ1 | Cell cycle c | Rad1 | sp\|Q9QWZ | 3 | 3 | 17,9 |
| Q8VE65 | Q8VE65 | Transcripti | Taf12 | sp\|Q8VE65 | 2 | 2 | 18 |
| Q9Z206 | Q9Z206 | Neuroepith | Net1 | sp\|Q9Z206 | 2 | 2 | 5,2 |
| Q3TUU5 | Q3TUU5 | Testis-expr | Tex30 | sp\|Q3TUU | 6 | 6 | 34,2 |
| O08648 | O08648 | Mitogen-ac | Map3k4 | sp\|O08648 | 13 | 13 | 10,1 |
| Q99LC3 | Q99LC3 | NADH deh | Ndufa10 | sp\|Q99LC3 | 13 | 13 | 49,9 |
| Q9D4C5 | Q9D4C5 | ELL-associa | Eaf1 | sp\|Q9D4C5 | 5 | 5 | 36,2 |
| Q9D1N9 | Q9D1N9 | 39S riboso | Mrpl21 | sp\|Q9D1N | 5 | 5 | 31,6 |
| Q9D2R0 | Q9D2R0 | Acetoacety | Aacs | sp\|Q9D2R0 | 21 | 21 | 37,9 |
| O70348 | O70348 | Decapping | Dxo | sp\|O70348 | 4 | 4 | 12,6 |
| P17426 | P17426 | AP-2 comp | Ap2a1 | sp\|P17426 | 42 | 34 | 56,5 |
| Q9CQN6 | Q9CQN6 | Transmem | Tmem14c | sp\|Q9CQN | 2 | 2 | 24,6 |
| P12815 | P12815 | Programm | Pdcd6 | sp\|P12815 | 11 | 11 | 73,8 |
| P62858 | P62858 | 40S riboso | Rps28 | sp\|P62858 | 4 | 4 | 52,2 |
| Q9WUE3 | Q9WUE3 | Cytochrom | Cyb561d2 | sp\|Q9WUE | 1 | 1 | 5,4 |

| | | | | | | | |
|---|---|---|---|---|---|---|---|
| Q9CPS6 | Q9CPS6 | Histidine tr | Hint3 | sp\|Q9CPS6 | 1 | 1 | 9,1 |
| Q3V0J1 | Q3V0J1 | Transmem | Tmem237 | sp\|Q3V0J1 | 3 | 3 | 14,8 |
| P84089 | P84089 | Enhancer c | Erh | sp\|P84089 | 4 | 4 | 41,3 |
| Q07139 | Q07139 | Protein EC | Ect2 | sp\|Q07139 | 23 | 23 | 36,8 |
| Q61069 | Q61069 | Upstream s | Usf1 | sp\|Q61069 | 4 | 2 | 12,9 |
| Q9CQU5 | Q9CQU5 | ZW10 inter | Zwint | sp\|Q9CQU | 6 | 6 | 31,7 |
| Q80WR5 | Q80WR5 | UPF0688 protein C1or | | sp\|Q80WR | 5 | 5 | 43 |
| P59913 | P59913 | Protein-L-i | Pcmtd1 | sp\|P59913 | 2 | 2 | 6,4 |
| CON__P04 | CON__P04264 | | | | 30 | 19 | 41,3 |
| Q8R2X8 | Q8R2X8 | Golgin-45 | Blzf1 | sp\|Q8R2X8 | 2 | 2 | 6,2 |
| Q921Y0 | Q921Y0 | MOB kinas | Mob1a | sp\|Q921Y0 | 7 | 1 | 26,4 |
| Q99LE3 | Q99LE3 | LysM and p | Lysmd3 | sp\|Q99LE3 | 1 | 1 | 3,9 |
| Q6P0X2 | Q6P0X2 | Zinc finger | Znf511 | sp\|Q6P0X2 | 2 | 2 | 12,3 |
| Q8BUL6 | Q8BUL6 | Pleckstrin l | Plekha1 | sp\|Q8BUL6 | 2 | 2 | 9,9 |
| Q3U1J4 | Q3U1J4 | DNA dama | Ddb1 | sp\|Q3U1J4 | 57 | 57 | 49,6 |
| Q3TD49 | Q3TD49 | Signal pept | Sppl2b | sp\|Q3TD49 | 2 | 2 | 5,7 |
| P09602 | P09602 | Non-histor | Hmgn2 | sp\|P09602 | 3 | 3 | 30 |
| O55098 | O55098 | Serine/thre | Stk10 | sp\|O55098 | 14 | 12 | 16,3 |
| Q8BP86 | Q8BP86 | snRNA-acti | Snapc4 | sp\|Q8BP86 | 2 | 2 | 2,7 |
| G5E8Z2 | G5E8Z2 | Transcripti | Taf4b | sp\|G5E8Z2 | 2 | 2 | 2,7 |
| Q9D2V8 | Q9D2V8 | Major facil | Mfsd10 | sp\|Q9D2V | 6 | 6 | 16,7 |
| P10404 | P10404 | MLV-related proviral E | | sp\|P10404 | 6 | 2 | 10,3 |
| P61967 | P61967 | AP-1 comp | Ap1s1 | sp\|P61967 | 7 | 5 | 37,3 |
| Q3TGW2 | Q3TGW2 | Endonucle | Eepd1 | sp\|Q3TGW | 5 | 5 | 12,7 |
| Q8BG09 | Q8BG09 | Transmem | Tmem184l | sp\|Q8BG0 | 1 | 1 | 2,7 |
| P41233;Q9 | P41233 | ATP-bindin | Abca1 | sp\|P41233 | 5 | 4 | 3,5 |
| Q3TWW8 | Q3TWW8 | Serine/arg | Srsf6 | sp\|Q3TWV | 12 | 9 | 31 |
| Q9CQB5 | Q9CQB5 | CDGSH iro | Cisd2 | sp\|Q9CQB | 7 | 7 | 53,3 |
| Q8R2L5 | Q8R2L5 | 28S riboso | Mrps18c | sp\|Q8R2L5 | 2 | 2 | 10,5 |
| Q9WU00 | Q9WU00 | Nuclear re | Nrf1 | sp\|Q9WU0 | 5 | 5 | 11,5 |
| Q9CQF0 | Q9CQF0 | 39S riboso | Mrpl11 | sp\|Q9CQF | 10 | 10 | 54,7 |
| Q5DTM8 | Q5DTM8 | E3 ubiquiti | Rnf20 | sp\|Q5DTM | 35 | 32 | 41,3 |
| Q8VEM9 | Q8VEM9 | Kelch dom | Klhdc3 | sp\|Q8VEM | 1 | 1 | 5 |
| Q99KU1 | Q99KU1 | Dehydrodo | Dhdds | sp\|Q99KU | 4 | 4 | 14,7 |
| P49722 | P49722 | Proteasom | Psma2 | sp\|P49722 | 13 | 13 | 72,6 |
| P47930 | P47930 | Fos-related | Fosl2 | sp\|P47930 | 3 | 3 | 13,8 |
| Q6PEE2 | Q6PEE2 | CBP80/20- | Ctif | sp\|Q6PEE2 | 6 | 6 | 17,7 |
| Q924Z4 | Q924Z4 | Ceramide s | Cers2 | sp\|Q924Z4 | 7 | 7 | 26,1 |
| Q9Z2G0 | Q9Z2G0 | Protein fer | Fem1b | sp\|Q9Z2G | 10 | 10 | 17,1 |
| Q9DC23 | Q9DC23 | DnaJ homo | Dnajc10 | sp\|Q9DC2 | 24 | 24 | 41,5 |
| Q6EJB6 | Q6EJB6 | U3 small n | Utp14b | sp\|Q6EJB6 | 8 | 5 | 12,2 |
| P84099 | P84099 | 60S riboso | Rpl19 | sp\|P84099 | 8 | 8 | 31,6 |
| Q99MI1 | Q99MI1 | ELKS/Rab6 | Erc1 | sp\|Q99MI | 22 | 16 | 23,8 |
| Q920F6 | Q920F6 | Structural | Smc1b | sp\|Q920F6 | 4 | 4 | 3,8 |
| Q9CQP3 | Q9CQP3 | Coiled-coil | Chchd5 | sp\|Q9CQP | 4 | 4 | 49,1 |
| P58742 | P58742 | Aladin | Aaas | sp\|P58742 | 17 | 17 | 49,6 |
| Q8BH48 | Q8BH48 | Ubiquitin-a | Ubap1 | sp\|Q8BH4 | 8 | 8 | 27,5 |
| P97305 | P97305 | Nuclear fa | Nfatc3 | sp\|P97305 | 4 | 4 | 5,8 |
| P63147 | P63147 | Ubiquitin-c | Ube2b | sp\|P63147 | 2 | 2 | 19,7 |
| Q61644 | Q61644 | Protein kin | Pacsin1 | sp\|Q61644 | 1 | 1 | 2 |

| | | | | | | | |
|---|---|---|---|---|---|---|---|
| P26231 | P26231 | Catenin alp | Ctnna1 | sp\|P26231 | 44 | 37 | 61,3 |
| O08550 | O08550 | Histone-lys | Kmt2b | sp\|O08550 | 16 | 16 | 8,4 |
| Q3TLR7 | Q3TLR7 | Denticleles | Dtl | sp\|Q3TLR7 | 8 | 8 | 15,1 |
| Q8BH70 | Q8BH70 | F-box/LRR- | Fbxl4 | sp\|Q8BH70 | 4 | 4 | 7,4 |
| Q8CDM8 | Q8CDM8 | Protein FA | Fam160b1 | sp\|Q8CDM | 13 | 12 | 24,1 |
| P30658 | P30658 | Chromobo | Cbx2 | sp\|P30658 | 6 | 6 | 17,7 |
| G5E861 | G5E861 | Sodium ch | Sclt1 | sp\|G5E861 | 2 | 2 | 3,5 |
| P42208 | P42208 | Septin-2 | 02-set | sp\|P42208 | 17 | 17 | 66,5 |
| Q3UE37 | Q3UE37 | Ubiquitin-c | Ube2z | sp\|Q3UE37 | 13 | 13 | 36,5 |
| Q9D0M3 | Q9D0M3 | Cytochrom | Cyc1 | sp\|Q9D0M | 9 | 9 | 46,8 |
| Q8CIB6 | Q8CIB6 | Transmem | Tmem230 | sp\|Q8CIB6 | 1 | 1 | 11,7 |
| P01887 | P01887 | Beta-2-mic | B2m | sp\|P01887 | 2 | 2 | 16 |
| Q91VU8 | Q91VU8 | Prostate tu | Ptov1 | sp\|Q91VU | 2 | 2 | 8,9 |
| Q9CQ18 | Q9CQ18 | Ribonuclea | Rnaseh2c | sp\|Q9CQ1 | 3 | 3 | 31,3 |
| Q6ZQE4 | Q6ZQE4 | Transmem | Tmem194a | sp\|Q6ZQE4 | 3 | 3 | 6,9 |
| Q8BV49 | Q8BV49 | Pyrin and H | Pyhin1 | sp\|Q8BV49 | 3 | 2 | 14 |
| Q8K3W3 | Q8K3W3 | Protein CA | Casc3 | sp\|Q8K3W | 2 | 2 | 5,4 |
| P47911 | P47911 | 60S riboso | Rpl6 | sp\|P47911 | 18 | 18 | 46,3 |
| Q921R4 | Q921R4 | DnaJ homo | Dnajc14 | sp\|Q921R4 | 3 | 3 | 7,1 |
| F8VPZ5 | F8VPZ5 | | Ercc6 | sp\|F8VPZ5 | 9 | 9 | 8,9 |
| Q5PRE5 | Q5PRE5 | Proline and | Proser1 | sp\|Q5PRE5 | 3 | 3 | 2,3 |
| Q9CZ62 | Q9CZ62 | Centrosom | Cep97 | sp\|Q9CZ62 | 3 | 3 | 6 |
| Q6PAR0 | Q6PAR0 | Kelch dom | Klhdc10 | sp\|Q6PAR0 | 2 | 2 | 6,4 |
| Q62406 | Q62406 | Interleukin | Irak1 | sp\|Q62406 | 2 | 2 | 3,9 |
| Q8K1I7 | Q8K1I7 | WAS/WAS | Wipf1 | sp\|Q8K1I7 | 4 | 4 | 11,8 |
| Q8R092 | Q8R092 | Uncharacterized prote | | sp\|Q8R092 | 2 | 2 | 10,7 |
| Q9D0C1 | Q9D0C1 | E3 ubiquiti | Rnf115 | sp\|Q9D0C1 | 1 | 1 | 5,9 |
| Q80TY0 | Q80TY0 | Formin-bin | Fnbp1 | sp\|Q80TY0 | 10 | 10 | 20,9 |
| Q9Z0F6 | Q9Z0F6 | Cell cycle c | Rad9a | sp\|Q9Z0F6 | 3 | 3 | 6,2 |
| Q9Z1X9 | Q9Z1X9 | Cell divisio | Cdc45 | sp\|Q9Z1X9 | 7 | 7 | 15,4 |
| E9Q735 | E9Q735 | Ubiquitin c | Ube4a | sp\|E9Q735 | 14 | 14 | 19,5 |
| Q8BNY6 | Q8BNY6 | Neuronal c | Ncs1 | sp\|Q8BNY6 | 9 | 9 | 55,3 |
| P80314 | P80314 | T-complex | Cct2 | sp\|P80314 | 36 | 36 | 75,1 |
| P35550 | P35550 | rRNA 2-O-r | Fbl | sp\|P35550 | 18 | 14 | 59,3 |
| Q9CZY3 | Q9CZY3 | Ubiquitin-c | Ube2v1 | sp\|Q9CZY3 | 7 | 2 | 57,8 |
| Q6ZWS8 | Q6ZWS8 | Speckle-typ | Spop | sp\|Q6ZWS | 4 | 4 | 13,6 |
| Q8K2W9 | Q8K2W9 | Uncharacterized prote | | sp\|Q8K2W | 2 | 2 | 8,3 |
| Q61602 | Q61602 | Transcripti | Gli3 | sp\|Q61602 | 11 | 11 | 12,8 |
| Q8BKC8 | Q8BKC8 | Phosphatid | Pi4kb | sp\|Q8BKC8 | 15 | 15 | 22,7 |
| P97467 | P97467 | Peptidyl-gl | Pam | sp\|P97467 | 8 | 8 | 13,1 |
| O35239 | O35239 | Tyrosine-p | Ptpn9 | sp\|O35239 | 14 | 14 | 31,7 |
| Q8VCK5 | Q8VCK5 | Kelch-like p | Klhl20 | sp\|Q8VCK5 | 2 | 2 | 3,6 |
| O70157 | O70157 | DNA topois | Top3a | sp\|O70157 | 5 | 4 | 6,9 |
| P63158 | P63158 | High mobil | Hmgb1 | sp\|P63158 | 18 | 17 | 55,8 |
| Q6P3Z3 | Q6P3Z3 | THAP dom | Thap4 | sp\|Q6P3Z3 | 4 | 4 | 7,7 |
| Q8BGW5 | Q8BGW5 | SOSS comp | Nabp1 | sp\|Q8BGW | 3 | 2 | 20,7 |
| Q9QZI8 | Q9QZI8 | Serine inco | Serinc1 | sp\|Q9QZI8 | 3 | 3 | 9,3 |
| Q9D1D4 | Q9D1D4 | Transmem | Tmed10 | sp\|Q9D1D | 7 | 7 | 33,3 |
| P58059 | P58059 | 28S riboso | Mrps21 | sp\|P58059 | 2 | 2 | 31 |
| Q8JZX3 | Q8JZX3 | POC1 cent | Poc1a | sp\|Q8JZX3 | 3 | 3 | 10,4 |

| | | | | | | | |
|---|---|---|---|---|---|---|---|
| O08759 | O08759 | Ubiquitin-p | Ube3a | sp\|O08759 | 27 | 27 | 37,1 |
| Q9DC40 | Q9DC40 | Telomere l | Telo2 | sp\|Q9DC40 | 13 | 13 | 20,6 |
| P61022 | P61022 | Calcineurin | Chp1 | sp\|P61022 | 10 | 10 | 66,7 |
| Q8BH50 | Q8BH50 | Uncharacterized prote | | sp\|Q8BH50 | 3 | 3 | 20,8 |
| P61750 | P61750 | ADP-ribosy | Arf4 | sp\|P61750 | 9 | 4 | 64,4 |
| Q60930 | Q60930 | Voltage-de | Vdac2 | sp\|Q60930 | 18 | 18 | 73,2 |
| Q62165 | Q62165 | Dystroglyc | Dag1 | sp\|Q62165 | 12 | 12 | 19 |
| Q9JIL5 | Q9JIL5 | Tubby-rela | Tulp4 | sp\|Q9JIL5 | 3 | 3 | 4 |
| Q8BGG6 | Q8BGG6 | Motile spe | Mospd3 | sp\|Q8BGG | 3 | 3 | 21,3 |
| Q91WK1 | Q91WK1 | SPRY doma | Spryd4 | sp\|Q91WK | 9 | 9 | 50,7 |
| Q99N95 | Q99N95 | 39S riboso | Mrpl3 | sp\|Q99N95 | 8 | 8 | 27,9 |
| P68369;P0 | P68369;P0521 | Tubulin alp | Tuba1a;Tu | sp\|P68369 | 27 | 1 | 65,9 |
| Q8R2M0 | Q8R2M0 | Transcripti | Tceanc2 | sp\|Q8R2M | 4 | 4 | 18,8 |
| O70591 | O70591 | Prefoldin s | Pfdn2 | sp\|O70591 | 6 | 6 | 44,8 |
| Q8K2I9 | Q8K2I9 | F-box only | Fbxo18 | sp\|Q8K2I9 | 7 | 7 | 8,2 |
| Q923S9 | Q923S9 | Ras-relatec | Rab30 | sp\|Q923S9 | 5 | 5 | 31 |
| Q99P31 | Q99P31 | Hsp70-bin | Hspbp1 | sp\|Q99P31 | 13 | 13 | 58 |
| Q8CI33 | Q8CI33 | CWF19-like | Cwf19l1 | sp\|Q8CI33 | 15 | 15 | 40,4 |
| Q8R361 | Q8R361 | Rab11 fam | Rab11fip5 | sp\|Q8R361 | 20 | 20 | 49 |
| Q9CZW4 | Q9CZW4 | Long-chain | Acsl3 | sp\|Q9CZW | 20 | 20 | 36,9 |
| Q9ESK9 | Q9ESK9 | RB1-induci | Rb1cc1 | sp\|Q9ESK9 | 18 | 18 | 15 |
| P61211 | P61211 | ADP-ribosy | Arl1 | sp\|P61211 | 11 | 11 | 72,9 |
| P58064 | P58064 | 28S riboso | Mrps6 | sp\|P58064 | 6 | 6 | 44,8 |
| P70195 | P70195 | Proteasom | Psmb7 | sp\|P70195 | 14 | 14 | 71,8 |
| Q9D7G4 | Q9D7G4 | Protein EU | Eurl | sp\|Q9D7G | 1 | 1 | 7,2 |
| Q9CQG1 | Q9CQG1 | Putative gl | Chac2 | sp\|Q9CQG | 2 | 2 | 18,5 |
| Q3UQU0 | Q3UQU0 | Bromodom | Brd9 | sp\|Q3UQU | 7 | 7 | 19,8 |
| Q99N89 | Q99N89 | 39S riboso | Mrpl43 | sp\|Q99N89 | 3 | 3 | 18,6 |
| Q8VBY2 | Q8VBY2 | Calcium/ca | Camkk1 | sp\|Q8VBY2 | 1 | 1 | 4 |
| Q9D1E8 | Q9D1E8 | 1-acyl-sn-g | Agpat5 | sp\|Q9D1E8 | 5 | 5 | 20,3 |
| Q8R010 | Q8R010 | Aminoacyl | Aimp2 | sp\|Q8R010 | 8 | 8 | 44,7 |
| Q91YT7 | Q91YT7 | YTH domai | Ythdf2 | sp\|Q91YT7 | 12 | 8 | 28 |
| Q8C3I8 | Q8C3I8 | Protein HG | Hgh1 | sp\|Q8C3I8 | 11 | 11 | 41,2 |
| P52875 | P52875 | Transmem | Tmem165 | sp\|P52875 | 6 | 6 | 22,9 |
| P50516 | P50516 | V-type pro | Atp6v1a | sp\|P50516 | 32 | 32 | 66,6 |
| Q3U1G5 | Q3U1G5 | Interferon- | Isg20l2 | sp\|Q3U1G | 13 | 13 | 39,9 |
| Q9Z248 | Q9Z248 | Zinc finger | Aebp2 | sp\|Q9Z248 | 6 | 6 | 19,8 |
| Q8K3M5 | Q8K3M5 | CDK5 and A | Cables2 | sp\|Q8K3M | 2 | 2 | 6,1 |
| Q9Z2N8;Q | Q9Z2N8 | Actin-like p | Actl6a | sp\|Q9Z2N8 | 16 | 16 | 52,9 |
| Q8VE22 | Q8VE22 | 28S riboso | Mrps23 | sp\|Q8VE22 | 10 | 10 | 65 |
| Q9DA19 | Q9DA19 | Corepresso | Cir1 | sp\|Q9DA1 | 2 | 2 | 5,1 |
| Q8BUV8 | Q8BUV8 | Protein GP | Gpr107 | sp\|Q8BUV | 4 | 4 | 10,9 |
| P55096 | P55096 | ATP-bindin | Abcd3 | sp\|P55096 | 18 | 18 | 35,1 |
| Q99LB0 | Q99LB0 | Deoxynucl | Dnttip1 | sp\|Q99LB0 | 8 | 8 | 36 |
| Q9D8X0 | Q9D8X0 | Protein MA | Manbal | sp\|Q9D8X0 | 2 | 2 | 40 |
| P26043 | P26043 | Radixin | Rdx | sp\|P26043 | 31 | 18 | 55,6 |
| Q62377 | Q62377 | U2 small n | Zrsr2 | sp\|Q62377 | 3 | 2 | 9,1 |
| Q8BQZ4 | Q8BQZ4 | Ral GTPase | Ralgapb | sp\|Q8BQZ | 12 | 12 | 15 |
| Q9D1R9 | Q9D1R9 | 60S riboso | Rpl34 | sp\|Q9D1R | 8 | 8 | 43,6 |
| CON__Q0V | CON__Q0VCM5 | | | | 2 | 2 | 4,1 |

| | | | | | | | |
|---|---|---|---|---|---|---|---|
| Q9CQA1 | Q9CQA1 | Trafficking | Trappc5 | sp\|Q9CQA | 3 | 3 | 15,4 |
| A6H687 | A6H687 | SAC3 doma | Sac3d1 | sp\|A6H687 | 2 | 2 | 9,8 |
| Q6PDM1 | Q6PDM1 | Male-speci | Msl1 | sp\|Q6PDM | 7 | 7 | 17,7 |
| Q6GQV7 | Q6GQV7 | Erythroid d | Edrf1 | sp\|Q6GQV | 3 | 3 | 4,2 |
| P61971 | P61971 | Nuclear tra | Nutf2 | sp\|P61971 | 5 | 5 | 69,3 |
| Q6P1B1 | Q6P1B1 | Xaa-Pro an | Xpnpep1 | sp\|Q6P1B1 | 28 | 27 | 63,4 |
| Q80YV3 | Q80YV3 | Transforma | Trrap | sp\|Q80YV3 | 64 | 64 | 36,1 |
| O70579 | O70579 | Peroxisom | Slc25a17 | sp\|O70579 | 10 | 10 | 33,9 |
| Q810J8 | Q810J8 | Zinc finger | Zfyve1 | sp\|Q810J8 | 3 | 3 | 4,8 |
| Q8C0P7 | Q8C0P7 | Zinc finger | Znf451 | sp\|Q8C0P7 | 4 | 4 | 2,9 |
| Q6PFR5 | Q6PFR5 | Transforme | Tra2a | sp\|Q6PFR5 | 8 | 7 | 29,5 |
| Q9D289 | Q9D289 | Trafficking | Trappc6b | sp\|Q9D289 | 6 | 6 | 41,8 |
| Q8BHE8 | Q8BHE8 | Uncharacterized prote | | sp\|Q8BHE8 | 3 | 3 | 12 |
| Q8BG19 | Q8BG19 | Transmem | Tmtc4 | sp\|Q8BG19 | 6 | 4 | 9,3 |
| Q9CQH8 | Q9CQH8 | Ribonuclea | Rpp14 | sp\|Q9CQH | 2 | 2 | 22,1 |
| Q9CQ28 | Q9CQ28 | Diphthine- | Dph6 | sp\|Q9CQ2 | 4 | 4 | 27,3 |
| O70230 | O70230 | Zinc finger | Znf143 | sp\|O70230 | 5 | 5 | 16,6 |
| Q91Z38 | Q91Z38 | Tetratricop | Ttc1 | sp\|Q91Z38 | 7 | 7 | 39,4 |
| Q8R205 | Q8R205 | Zinc finger | Zc3h10 | sp\|Q8R205 | 3 | 3 | 9 |
| Q6PDN3 | Q6PDN3 | Myosin ligh | Mylk | sp\|Q6PDN | 13 | 13 | 7,8 |
| Q9D8Z2 | Q9D8Z2 | TP53-regul | Triap1 | sp\|Q9D8Z2 | 1 | 1 | 18,4 |
| Q8R003 | Q8R003 | Muscleblin | Mbnl3 | sp\|Q8R003 | 7 | 4 | 15,8 |
| O35153 | O35153 | BET1-like p | Bet1l | sp\|O35153 | 3 | 3 | 44,1 |
| Q8K2R5 | Q8K2R5 | Zinc finger | Znf668 | sp\|Q8K2R5 | 6 | 6 | 11,1 |
| Q8CHY3 | Q8CHY3 | Dymeclin | Dym | sp\|Q8CHY3 | 3 | 3 | 3,9 |
| Q9Z0H3 | Q9Z0H3 | SWI/SNF-re | Smarcb1 | sp\|Q9Z0H3 | 13 | 13 | 51,2 |
| Q924S8 | Q924S8 | Sprouty-re | Spred1 | sp\|Q924S8 | 5 | 4 | 14,6 |
| O55106 | O55106 | Striatin | Strn | sp\|O55106 | 25 | 24 | 52,1 |
| Q9WTZ1 | Q9WTZ1 | RING-box p | Rnf7 | sp\|Q9WTZ | 3 | 3 | 38,1 |
| Q61908 | Q61908 | Cx9C motif | Cmc4 | sp\|Q61908 | 1 | 1 | 20,6 |
| Q8BHL8 | Q8BHL8 | Proteasom | Psmf1 | sp\|Q8BHL8 | 4 | 4 | 20,7 |
| Q9CQZ7 | Q9CQZ7 | DNA-direct | Polr3k | sp\|Q9CQZ7 | 4 | 4 | 55,6 |
| Q52KG4 | Q52KG4 | | Zbtb45 | sp\|Q52KG4 | 2 | 2 | 6,3 |
| Q9CZW2 | Q9CZW2 | Centromer | Cenpn | sp\|Q9CZW | 4 | 4 | 16 |
| Q9WVF7 | Q9WVF7 | DNA polym | Pole | sp\|Q9WVF | 45 | 45 | 25,6 |
| Q9CZ92 | Q9CZ92 | Centromer | Cenpp | sp\|Q9CZ92 | 1 | 1 | 4,5 |
| Q9CR20 | Q9CR20 | Immediate | Ier3ip1 | sp\|Q9CR20 | 2 | 2 | 34,1 |
| Q80ZI6 | Q80ZI6 | E3 ubiquiti | Lrsam1 | sp\|Q80ZI6 | 5 | 5 | 10 |
| Q6KAU7 | Q6KAU7 | Pleckstrin h | Plekhg2 | sp\|Q6KAU | 1 | 1 | 1,2 |
| Q8BH69 | Q8BH69 | Selenide, w | Sephs1 | sp\|Q8BH69 | 10 | 10 | 39 |
| Q9JLB9 | Q9JLB9 | Nectin-3 | Pvrl3 | sp\|Q9JLB9 | 3 | 3 | 9,1 |
| Q5NCC3 | Q5NCC3 | E3 ubiquiti | Trim41 | sp\|Q5NCC | 3 | 3 | 7,1 |
| P30681 | P30681 | High mobil | Hmgb2 | sp\|P30681 | 13 | 12 | 50,5 |
| P08249 | P08249 | Malate deh | Mdh2 | sp\|P08249 | 20 | 20 | 67,8 |
| P15508 | P15508 | Spectrin be | Sptb | sp\|P15508 | 7 | 4 | 4,3 |
| O08989 | O08989 | Ras-related | Mras | sp\|O08989 | 2 | 2 | 14,4 |
| O88974 | O88974 | Histone-lys | Setdb1 | sp\|O88974 | 12 | 12 | 12,9 |
| Q99LQ1 | Q99LQ1 | MAP3K12- | Mbip | sp\|Q99LQ1 | 5 | 5 | 20,8 |
| Q9JMK2 | Q9JMK2 | Casein kina | Csnk1e | sp\|Q9JMK | 7 | 1 | 18,5 |
| P28740 | P28740 | Kinesin-like | Kif2a | sp\|P28740 | 20 | 19 | 32,8 |

| | | | | | | | |
|---|---|---|---|---|---|---|---|
| P51807 | P51807 | Dynein ligh | Dynlt1 | sp\|P51807 | 4 | 4 | 70,8 |
| Q5XPI3 | Q5XPI3 | E3 ubiquiti | Rnf123 | sp\|Q5XPI3 | 6 | 6 | 7,5 |
| P60882 | P60882 | Multiple e | Megf8 | sp\|P60882 | 8 | 8 | 5 |
| P52633 | P52633 | Signal tran | Stat6 | sp\|P52633 | 19 | 19 | 29,3 |
| Q8JZY4 | Q8JZY4 | Mitochond | Kiaa0391 | sp\|Q8JZY4 | 11 | 11 | 25,7 |
| Q921M7;Q | Q921M7 | Protein FA | Fam49b | sp\|Q921M | 16 | 16 | 65,4 |
| Q9DB07 | Q9DB07 | Intraflagell | Ift46 | sp\|Q9DB0 | 1 | 1 | 4,7 |
| Q9WUK4 | Q9WUK4 | Replication | Rfc2 | sp\|Q9WU | 19 | 19 | 65,9 |
| Q8BUN5 | Q8BUN5 | Mothers a | Smad3 | sp\|Q8BUN | 8 | 4 | 22,6 |
| Q8BSZ2 | Q8BSZ2 | AP-3 comp | Ap3s2 | sp\|Q8BSZ2 | 3 | 3 | 15,5 |
| P12850 | P12850 | Growth-re | Cxcl1 | sp\|P12850 | 2 | 2 | 32,3 |
| Q8R323 | Q8R323 | Replication | Rfc3 | sp\|Q8R323 | 21 | 21 | 67,4 |
| Q9CQR4 | Q9CQR4 | Acyl-coenz | Acot13 | sp\|Q9CQR | 5 | 5 | 43,6 |
| Q9R1Z7 | Q9R1Z7 | 6-pyruvoyl | Pts | sp\|Q9R1Z7 | 4 | 4 | 25 |
| Q3URV1 | Q3URV1 | Protein bro | Tbc1d32 | sp\|Q3URV | 4 | 4 | 4,3 |
| Q921I1 | Q921I1 | Serotransf | Tf | sp\|Q921I1 | 12 | 12 | 23,1 |
| Q99P72 | Q99P72 | Reticulon-4 | Rtn4 | sp\|Q99P72 | 43 | 43 | 62,9 |
| Q07832 | Q07832 | Serine/thre | Plk1 | sp\|Q07832 | 18 | 18 | 38,6 |
| Q9D2Q3 | Q9D2Q3 | Uncharacterized prote | | sp\|Q9D2Q | 4 | 4 | 10,6 |
| Q9D8P4 | Q9D8P4 | 39S riboso | Mrpl17 | sp\|Q9D8P4 | 6 | 6 | 28,4 |
| CON__P02 | CON__P02533 | Keratin, ty | Krt42 | ;;sp\|Q6IFX | 14 | 3 | 28,2 |
| Q7TQ95 | Q7TQ95 | Protein lun | Lnp | sp\|Q7TQ9 | 17 | 17 | 52,9 |
| CON__P17 | CON__P17697 | | | | 2 | 2 | 5,5 |
| Q8BP97 | Q8BP97 | Rhomboid | Rhbdd3 | sp\|Q8BP97 | 2 | 2 | 11,2 |
| Q6P073 | Q6P073 | Ubiquitin-c | Ube2j2 | sp\|Q6P073 | 7 | 7 | 38,2 |
| Q6NZK5 | Q6NZK5 | Uncharacte | Kiaa1328 | sp\|Q6NZK5 | 1 | 1 | 2,6 |
| Q91WC1 | Q91WC1 | Protection | Pot1 | sp\|Q91WC | 6 | 6 | 14,2 |
| O35493 | O35493 | Dual specif | Clk4 | sp\|O35493 | 7 | 3 | 19,5 |
| O35625 | O35625 | Axin-1 | Axin1 | sp\|O35625 | 2 | 2 | 3,2 |
| P48725 | P48725 | Pericentrin | Pcnt | sp\|P48725 | 46 | 46 | 23,7 |
| Q8BH31 | Q8BH31 | Major facil | Mfsd8 | sp\|Q8BH3 | 2 | 2 | 5,6 |
| P62908 | P62908 | 40S riboso | Rps3 | sp\|P62908 | 22 | 22 | 78,6 |
| P09926 | P09926 | Surfeit locu | Surf2 | sp\|P09926 | 6 | 6 | 35,4 |
| Q6ZPG2 | Q6ZPG2 | WD repeat | Wdr90 | sp\|Q6ZPG2 | 4 | 4 | 3,3 |
| Q9CPR4 | Q9CPR4 | 60S riboso | Rpl17 | sp\|Q9CPR4 | 11 | 11 | 49,5 |
| Q9D115 | Q9D115 | Zinc finger | Znf706 | sp\|Q9D115 | 2 | 2 | 30,3 |
| Q8VDI9 | Q8VDI9 | Alpha-1,2-r | Alg9 | sp\|Q8VDI9 | 8 | 8 | 15,5 |
| Q8BM86 | Q8BM86 | Lipoma HN | Lhfp | sp\|Q8BM8 | 2 | 2 | 10,5 |
| Q9JM14 | Q9JM14 | 5(3)-deoxy | Nt5c | sp\|Q9JM14 | 10 | 10 | 59,5 |
| Q9JIY5 | Q9JIY5 | Serine prot | Htra2 | sp\|Q9JIY5 | 11 | 10 | 34,1 |
| Q61532 | Q61532 | Mitogen-ac | Mapk6 | sp\|Q61532 | 4 | 3 | 7,1 |
| Q8C650 | Q8C650 | Septin-10 | 10-set | sp\|Q8C650 | 12 | 11 | 44 |
| O09117 | O09117 | Synaptoph | Sypl1 | sp\|O09117 | 5 | 5 | 24,5 |
| Q3V1V3 | Q3V1V3 | ESF1 homo | Esf1 | sp\|Q3V1V3 | 23 | 23 | 29,7 |
| Q8BTF7 | Q8BTF7 | DNA ligase | Lig4 | sp\|Q8BTF7 | 4 | 4 | 5,7 |
| Q8VE38 | Q8VE38 | Oxidoredu | Oxnad1 | sp\|Q8VE38 | 8 | 8 | 29,6 |
| Q9JI10 | Q9JI10 | Serine/thre | Stk3 | sp\|Q9JI10 | 21 | 15 | 47,7 |
| Q9QYT7 | Q9QYT7 | Phosphatid | Pigq | sp\|Q9QYT | 3 | 3 | 8,1 |
| Q9R1K5 | Q9R1K5 | Fizzy-relate | Fzr1 | sp\|Q9R1K5 | 3 | 3 | 8,3 |
| Q9R059 | Q9R059 | Four and a | Fhl3 | sp\|Q9R059 | 8 | 8 | 37,4 |

| | | | | | | | |
|---|---|---|---|---|---|---|---|
| Q99PU7 | Q99PU7 | Ubiquitin c | Bap1 | sp\|Q99PU7 | 7 | 7 | 12,8 |
| Q8VDP6 | Q8VDP6 | CDP-diacyl | Cdipt | sp\|Q8VDP | 4 | 4 | 21,1 |
| Q9Z1P6 | Q9Z1P6 | NADH deh | Ndufa7 | sp\|Q9Z1P6 | 2 | 2 | 21,2 |
| Q8BFY7 | Q8BFY7 | Protein FA | Fam64a | sp\|Q8BFY7 | 4 | 4 | 26,8 |
| P61358 | P61358 | 60S riboso | Rpl27 | sp\|P61358 | 9 | 9 | 51,5 |
| Q9CR55 | Q9CR55 | UPF0547 protein C16o | | sp\|Q9CR55 | 2 | 2 | 16,9 |
| Q9D2I5 | Q9D2I5 | LisH domai | Armc9 | sp\|Q9D2I5 | 4 | 4 | 6,1 |
| P70280 | P70280 | Vesicle-ass | Vamp7 | sp\|P70280 | 11 | 11 | 57,7 |
| P62852 | P62852 | 40S riboso | Rps25 | sp\|P62852 | 6 | 6 | 31,2 |
| Q505F5 | Q505F5 | Leucine-ric | Lrrc47 | sp\|Q505F5 | 22 | 22 | 55,6 |
| P83877 | P83877 | Thioredoxi | Txnl4a | sp\|P83877 | 7 | 7 | 67,6 |
| Q6NXY9 | Q6NXY9 | DNA-direct | Polr3g | sp\|Q6NXY9 | 2 | 2 | 15,7 |
| Q8BMJ7 | Q8BMJ7 | Cell growth | Cgrrf1 | sp\|Q8BMJ | 2 | 2 | 10,2 |
| Q9D8F1 | Q9D8F1 | Alpha-keto | Alkbh4 | sp\|Q9D8F1 | 2 | 2 | 9 |
| Q571F8 | Q571F8 | Glutamina | Gls2 | sp\|Q571F8 | 3 | 3 | 9,1 |
| Q923B0 | Q923B0 | Gamma-gl | Ggact | sp\|Q923B0 | 3 | 3 | 27,5 |
| Q8C0D9 | Q8C0D9 | Centrosom | Cep68 | sp\|Q8C0D9 | 1 | 1 | 2,3 |
| Q9CZU4 | Q9CZU4 | GTPase Era | Eral1 | sp\|Q9CZU4 | 14 | 14 | 50,6 |
| Q9R0Q4 | Q9R0Q4 | Mortality f | Morf4l2 | sp\|Q9R0Q | 12 | 10 | 51,4 |
| O08749 | O08749 | Dihydrolipo | Dld | sp\|O08749 | 19 | 19 | 55 |
| Q3USB7 | Q3USB7 | Inactive ph | Plcl1 | sp\|Q3USB7 | 4 | 4 | 6,3 |
| Q8CJ61 | Q8CJ61 | CKLF-like M | Cmtm4 | sp\|Q8CJ61 | 1 | 1 | 16,8 |
| Q61160 | Q61160 | FAS-associ | Fadd | sp\|Q61160 | 2 | 2 | 17,6 |
| Q9EQN9 | Q9EQN9 | | Slc19a2 | sp\|Q9EQN | 2 | 2 | 9 |
| Q9CQE3 | Q9CQE3 | 28S riboso | Mrps17 | sp\|Q9CQE | 6 | 6 | 61,7 |
| Q6ZWQ7 | Q6ZWQ7 | | Spcs3 | sp\|Q6ZWQ | 4 | 4 | 21,7 |
| Q8K2Y9 | Q8K2Y9 | Cerebral ca | Ccm2 | sp\|Q8K2Y9 | 6 | 6 | 19,6 |
| Q80TG1 | Q80TG1 | KAT8 regul | Kansl1 | sp\|Q80TG1 | 9 | 9 | 12,4 |
| P14148 | P14148 | 60S riboso | Rpl7 | sp\|P14148 | 25 | 25 | 55,6 |
| P97366 | P97366 | Ecotropic v | Evi5 | sp\|P97366 | 6 | 6 | 8,4 |
| Q9WUD8 | Q9WUD8 | Fas apopto | Faim | sp\|Q9WUD | 4 | 4 | 22,9 |
| Q62504 | Q62504 | Msx2-inter | Spen | sp\|Q62504 | 32 | 32 | 12,7 |
| A2APY7 | A2APY7 | NADH deh | Ndufaf5 | sp\|A2APY7 | 7 | 7 | 25,9 |
| Q99M31 | Q99M31 | Heat shock | Hspa14 | sp\|Q99M3 | 23 | 23 | 54,2 |
| Q91YT0 | Q91YT0 | NADH deh | Ndufv1 | sp\|Q91YT0 | 16 | 16 | 50,9 |
| Q6PHZ2;P1 | Q6PHZ2 | Calcium/ca | Camk2d | sp\|Q6PHZ2 | 14 | 11 | 38,5 |
| Q9CZJ6 | Q9CZJ6 | Protein Mi | Mis18a | sp\|Q9CZJ6 | 2 | 2 | 14,2 |
| Q99MD8 | Q99MD8 | Myoneurin | Mynn | sp\|Q99MD | 3 | 3 | 6,9 |
| Q6P7W5 | Q6P7W5 | tRNA-splici | Tsen2 | sp\|Q6P7W | 3 | 3 | 8 |
| Q9JI46 | Q9JI46 | Diphospho | Nudt3 | sp\|Q9JI46 | 5 | 4 | 34,5 |
| Q60632 | Q60632 | COUP tran | Nr2f1 | sp\|Q60632 | 8 | 3 | 25,4 |
| Q8K214 | Q8K214 | Polycomb | Scmh1 | sp\|Q8K214 | 5 | 5 | 8,5 |
| Q9CWG1 | Q9CWG1 | Glioma pat | Glipr1 | sp\|Q9CWG | 2 | 2 | 9,4 |
| Q9DCA2 | Q9DCA2 | 28S riboso | Mrps11 | sp\|Q9DCA | 5 | 5 | 38,2 |
| Q8C0V0 | Q8C0V0 | Serine/thre | Tlk1 | sp\|Q8C0V0 | 7 | 5 | 14,9 |
| Q99LJ8 | Q99LJ8 | Dehydrodo | Nus1 | sp\|Q99LJ8 | 2 | 2 | 12,8 |
| Q8VBV7 | Q8VBV7 | COP9 signa | Cops8 | sp\|Q8VBV7 | 8 | 8 | 72,2 |
| Q91WB4 | Q91WB4 | Pleckstrin | Plekhf2 | sp\|Q91WB | 3 | 3 | 21,7 |
| P56387 | P56387 | Dynein ligh | Dynlt3 | sp\|P56387 | 4 | 4 | 49,1 |
| Q9CX11 | Q9CX11 | rRNA-proc | Utp23 | sp\|Q9CX11 | 4 | 4 | 21,3 |

| | | | | | | | |
|---|---|---|---|---|---|---|---|
| A2AKB9 | A2AKB9 | DDB1- and | Dcaf10 | sp\|A2AKB9 | 7 | 7 | 17,8 |
| Q91VW3 | Q91VW3 | SH3 domai | Sh3bgrl3 | sp\|Q91VW | 5 | 5 | 45,2 |
| P48754 | P48754 | Breast can | Brca1 | sp\|P48754 | 20 | 20 | 16,3 |
| Q99JX3 | Q99JX3 | Golgi reass | Gorasp2 | sp\|Q99JX3 | 9 | 9 | 29,7 |
| Q8QZY9 | Q8QZY9 | Splicing fac | Sf3b4 | sp\|Q8QZY9 | 6 | 6 | 24,1 |
| Q3UGF1 | Q3UGF1 | WD repeat | Wdr19 | sp\|Q3UGF | 3 | 3 | 3 |
| P70318 | P70318 | Nucleolysi | Tial1 | sp\|P70318 | 10 | 7 | 34,2 |
| P97431 | P97431 | Interferon | Irf6 | sp\|P97431 | 4 | 4 | 14,1 |
| Q6PJN8 | Q6PJN8 | DALR antic | Dalrd3 | sp\|Q6PJN8 | 1 | 1 | 2,8 |
| Q8BZH0 | Q8BZH0 | Zinc transp | Slc39a13 | sp\|Q8BZH0 | 3 | 3 | 12,5 |
| Q8BG30 | Q8BG30 | Negative e | Nelfa | sp\|Q8BG30 | 14 | 14 | 40,4 |
| Q62440;Q6 | Q62440;Q624 | Transducin | Tle1;Tle4 | sp\|Q62440 | 8 | 4 | 14,8 |
| P46694 | P46694 | Radiation-i | Ier3 | sp\|P46694 | 2 | 2 | 18,3 |
| Q9D855 | Q9D855 | Cytochrom | Uqcrb | sp\|Q9D855 | 8 | 8 | 49,5 |
| P63325 | P63325 | 40S riboso | Rps10 | sp\|P63325 | 11 | 11 | 58,8 |
| Q9D0F1 | Q9D0F1 | Kinetochor | Ndc80 | sp\|Q9D0F1 | 19 | 19 | 35,4 |
| Q9CQX2 | Q9CQX2 | Cytochrom | Cyb5b | sp\|Q9CQX | 7 | 7 | 58,9 |
| Q8VE10 | Q8VE10 | N-alpha-ac | Naa40 | sp\|Q8VE10 | 3 | 3 | 15,2 |
| Q922J3 | Q922J3 | CAP-Gly do | Clip1 | sp\|Q922J3 | 42 | 38 | 35,6 |
| Q9CS74 | Q9CS74 | Protein SG | Ecd | sp\|Q9CS74 | 7 | 7 | 16,2 |
| Q9CR86 | Q9CR86 | Calcium-re | Carhsp1 | sp\|Q9CR86 | 5 | 5 | 66,9 |
| Q80UP8 | Q80UP8 | Sodium-de | Slc20a2 | sp\|Q80UP8 | 5 | 5 | 13 |
| Q922M5 | Q922M5 | Cell divisio | Cdca7l | sp\|Q922M | 9 | 8 | 20,3 |
| Q78JE5 | Q78JE5 | F-box only | Fbxo22 | sp\|Q78JE5 | 8 | 8 | 31,8 |
| Q9CQI7 | Q9CQI7 | U2 small n | Snrpb2 | sp\|Q9CQI7 | 9 | 7 | 39,6 |
| Q99M51 | Q99M51 | Cytoplasm | Nck1 | sp\|Q99M5 | 13 | 13 | 47,7 |
| Q9D1G2 | Q9D1G2 | Phosphom | Pmvk | sp\|Q9D1G | 6 | 6 | 28,1 |
| Q64267 | Q64267 | DNA repair | Xpa | sp\|Q64267 | 3 | 3 | 12,5 |
| Q60597 | Q60597 | 2-oxogluta | Ogdh | sp\|Q60597 | 48 | 48 | 55 |
| A2AL36 | A2AL36 | Centriolin | Cntrl | sp\|A2AL36 | 4 | 4 | 2 |
| P15307 | P15307 | Proto-onco | Rel | sp\|P15307 | 9 | 9 | 18,6 |
| Q6ZPF3 | Q6ZPF3 | T-lymphon | Tiam2 | sp\|Q6ZPF3 | 1 | 1 | 1,3 |
| P35279 | P35279 | Ras-relatec | Rab6a | sp\|P35279 | 11 | 4 | 52,9 |
| P50428 | P50428 | Arylsulfata | Arsa | sp\|P50428 | 5 | 5 | 25,1 |
| O35864 | O35864 | COP9 signa | Cops5 | sp\|O35864 | 17 | 17 | 65,3 |
| Q9R1S8 | Q9R1S8 | Calpain-7 | Capn7 | sp\|Q9R1S8 | 6 | 6 | 9,2 |
| O89109 | O89109 | Intermedia | Kcnn4 | sp\|O89109 | 2 | 2 | 5,2 |
| Q80X66 | Q80X66 | BTB/POZ d | Btbd10 | sp\|Q80X66 | 2 | 2 | 5,1 |
| Q923D4 | Q923D4 | Splicing fac | Sf3b5 | sp\|Q923D4 | 5 | 5 | 83,7 |
| CON__P02 | CON__P02663 | | | | 4 | 4 | 18,8 |
| Q9D0J4 | Q9D0J4 | ADP-ribosy | Arl2 | sp\|Q9D0J4 | 8 | 8 | 55,4 |
| Q63739 | Q63739 | Protein tyr | Ptp4a1 | sp\|Q63739 | 6 | 3 | 40,5 |
| Q9DD16 | Q9DD16 | Nucleoside | Nudt22 | sp\|Q9DD1 | 2 | 2 | 12,7 |
| P97304 | P97304 | DNA-direct | Polr1d | sp\|P97304 | 6 | 6 | 50,4 |
| Q9JKP7 | Q9JKP7 | DNA polym | Pole3 | sp\|Q9JKP7 | 4 | 4 | 31,7 |
| P70245 | P70245 | 3-beta-hyd | Ebp | sp\|P70245 | 3 | 3 | 15,2 |
| P03888 | P03888 | NADH-ubic | Mtnd1 | sp\|P03888 | 4 | 4 | 11,9 |
| Q9CSP9 | Q9CSP9 | Tetratricop | Ttc14 | sp\|Q9CSP9 | 9 | 9 | 15,6 |
| Q5NCR9 | Q5NCR9 | Nuclear sp | Nsrp1 | sp\|Q5NCR | 6 | 6 | 17,2 |
| P43276 | P43276 | Histone H1 | Hist1h1b | sp\|P43276 | 12 | 9 | 41,3 |

| | | | | | | | |
|---|---|---|---|---|---|---|---|
| Q8R4H9 | Q8R4H9 | Zinc transp | Slc30a5 | sp\|Q8R4H | 5 | 5 | 10,6 |
| Q8R3K3 | Q8R3K3 | Pentatrico | Ptcd2 | sp\|Q8R3K | 7 | 7 | 17,6 |
| Q99JI6 | Q99JI6 | Ras-relate | Rap1b | sp\|Q99JI6 | 15 | 6 | 78,3 |
| Q6PDQ2 | Q6PDQ2 | Chromodo | Chd4 | sp\|Q6PDQ | 81 | 66 | 47,1 |
| Q8BIE6;Q9 | Q8BIE6 | FERM dom | Frmd4a | sp\|Q8BIE6 | 17 | 17 | 23,2 |
| Q9DBS9 | Q9DBS9 | Oxysterol-l | Osbpl3 | sp\|Q9DBS | 22 | 21 | 37,8 |
| O08604 | O08604 | Retinoic ac | Raet1c | sp\|O0860 | 1 | 1 | 5,1 |
| P54099 | P54099 | DNA polym | Polg | sp\|P54099 | 5 | 5 | 4,6 |
| Q91XF0 | Q91XF0 | Pyridoxine | Pnpo | sp\|Q91XF | 3 | 3 | 18 |
| Q8BIX3 | Q8BIX3 | ARL14 effe | Arl14ep | sp\|Q8BIX3 | 5 | 5 | 25,7 |
| P20108 | P20108 | Thioredoxi | Prdx3 | sp\|P20108 | 9 | 9 | 43,2 |
| Q3UA16 | Q3UA16 | Kinetochor | Spc25 | sp\|Q3UA1 | 6 | 6 | 31,9 |
| Q8CHT0 | Q8CHT0 | Delta-1-py | Aldh4a1 | sp\|Q8CHT | 2 | 2 | 6 |
| O54992 | O54992 | MAP kinas | Mapkapk5 | sp\|O5499 | 3 | 3 | 8,2 |
| P53612 | P53612 | Geranylger | Rabggtb | sp\|P53612 | 5 | 5 | 17,1 |
| Q69ZC8 | Q69ZC8 | GPALPP m | Gpalpp1 | sp\|Q69ZC | 2 | 2 | 6,4 |
| Q8R127 | Q8R127 | Saccharopi | Sccpdh | sp\|Q8R12 | 11 | 11 | 39,9 |
| Q91VC9 | Q91VC9 | Growth ho | Ghitm | sp\|Q91VC | 3 | 3 | 9,2 |
| A2AG58 | A2AG58 | Uncharacterized prote | | sp\|A2AG5 | 4 | 4 | 8,4 |
| Q80U28 | Q80U28 | MAP kinas | Madd | sp\|Q80U2 | 1 | 1 | 1,5 |
| Q8BU88 | Q8BU88 | 39S riboso | Mrpl22 | sp\|Q8BU8 | 8 | 8 | 42,7 |
| Q80X82 | Q80X82 | Symplekin | Sympk | sp\|Q80X8 | 41 | 41 | 41 |
| Q9QZC8 | Q9QZC8 | Abhydrola | Abhd1 | sp\|Q9QZC | 6 | 6 | 24,5 |
| Q8K183 | Q8K183 | Pyridoxal k | Pdxk | sp\|Q8K18 | 9 | 9 | 31,7 |
| Q9CQ25 | Q9CQ25 | Mitotic-spi | Mzt2 | sp\|Q9CQ2 | 3 | 3 | 37,7 |
| Q75N73 | Q75N73 | Zinc transp | Slc39a14 | sp\|Q75N7 | 3 | 3 | 12,9 |
| Q9WUL7 | Q9WUL7 | ADP-ribosy | Arl3 | sp\|Q9WUL | 7 | 7 | 63,2 |
| Q9CQ10 | Q9CQ10 | Charged m | Chmp3 | sp\|Q9CQ1 | 4 | 4 | 16,1 |
| P17095 | P17095 | High mobil | Hmga1 | sp\|P17095 | 7 | 7 | 38,3 |
| P23198 | P23198 | Chromobo | Cbx3 | sp\|P23198 | 9 | 7 | 48,6 |
| Q3TCX3 | Q3TCX3 | UPF0469 p | Kiaa0907 | sp\|Q3TCX | 10 | 10 | 25,5 |
| Q9DCJ1 | Q9DCJ1 | Target of r | Mlst8 | sp\|Q9DCJ1 | 6 | 6 | 29,1 |
| O55047 | O55047 | Serine/thre | Tlk2 | sp\|O5504 | 12 | 10 | 21,2 |
| Q80W37 | Q80W37 | Snurportin | Snupn | sp\|Q80W3 | 6 | 6 | 22,9 |
| Q9D7A6 | Q9D7A6 | Signal reco | Srp19 | sp\|Q9D7A | 3 | 3 | 35,4 |
| Q8BJH1 | Q8BJH1 | Zinc finger | Zc2hc1a | sp\|Q8BJH1 | 6 | 6 | 28,4 |
| Q8CE46 | Q8CE46 | Pseudouri | Pus7l | sp\|Q8CE4 | 7 | 7 | 15,7 |
| A2ACP1 | A2ACP1 | Tetratrico | Ttc39a | sp\|A2ACP1 | 1 | 1 | 1,9 |
| Q8K203 | Q8K203 | Endonucle | Neil3 | sp\|Q8K20 | 3 | 3 | 5,4 |
| O55142 | O55142 | 60S riboso | Rpl35a | sp\|O5514 | 10 | 10 | 54,5 |
| O88632 | O88632 | Semaphori | Sema3f | sp\|O8863 | 3 | 3 | 5,2 |
| A2AI05 | A2AI05 | NADPH-de | Ndor1 | sp\|A2AI05 | 2 | 2 | 4,3 |
| Q8VCX5 | Q8VCX5 | Calcium up | Micu1 | sp\|Q8VCX | 7 | 7 | 20,8 |
| Q6PCL9 | Q6PCL9 | Poly(A) pol | Papolg | sp\|Q6PCL | 15 | 12 | 27,2 |
| Q9JJT9 | Q9JJT9 | Phosphory | Phax | sp\|Q9JJT9 | 13 | 13 | 39,2 |
| Q8K0T4 | Q8K0T4 | Katanin p6 | Katnal1 | sp\|Q8K0T | 4 | 3 | 8,6 |
| Q9DB94 | Q9DB94 | WD repeat | Wdr53 | sp\|Q9DB9 | 2 | 2 | 14,5 |
| Q99K28 | Q99K28 | ADP-ribosy | Arfgap2 | sp\|Q99K2 | 13 | 13 | 31 |
| Q8BZ32 | Q8BZ32 | Putative P | Asxl2 | sp\|Q8BZ3 | 3 | 3 | 2,6 |
| Q9D6N5 | Q9D6N5 | Dr1-associ | Drap1 | sp\|Q9D6N | 5 | 5 | 29,3 |

| | | | | | | | |
|---|---|---|---|---|---|---|---|
| Q6PGL7 | Q6PGL7 | WASH com | Fam21 | sp\|Q6PGL7 | 37 | 37 | 48,6 |
| P97820 | P97820 | Mitogen-a | Map4k4 | sp\|P97820 | 21 | 14 | 20,8 |
| Q9D173 | Q9D173 | Mitochond | Tomm7 | sp\|Q9D173 | 1 | 1 | 30,9 |
| P23188 | P23188 | Furin | Furin | sp\|P23188 | 7 | 7 | 14,8 |
| Q8CG73 | Q8CG73 | Protein far | Rpgrip1l | sp\|Q8CG73 | 3 | 3 | 2,8 |
| Q9CQW2 | Q9CQW2 | ADP-ribosy | Arl8b | sp\|Q9CQW | 9 | 4 | 57 |
| Q8K1Y2 | Q8K1Y2 | Serine/thre | Prkd3 | sp\|Q8K1Y2 | 7 | 4 | 12,3 |
| Q9D6H2 | Q9D6H2 | Intraflagell | Hspb11 | sp\|Q9D6H | 5 | 5 | 44,1 |
| P16858;Q6 | P16858 | Glyceralde | Gapdh | sp\|P16858 | 26 | 26 | 85,3 |
| A2AQH4 | A2AQH4 | BCL-6 core | Bcorl1 | sp\|A2AQH | 6 | 6 | 5 |
| Q8CG48 | Q8CG48 | Structural | Smc2 | sp\|Q8CG48 | 49 | 49 | 49,2 |
| Q9R0T8 | Q9R0T8 | Inhibitor of | Ikbke | sp\|Q9R0T8 | 4 | 4 | 7,4 |
| Q8R3D1 | Q8R3D1 | TBC1 doma | Tbc1d13 | sp\|Q8R3D | 5 | 5 | 20 |
| Q9Z2W1 | Q9Z2W1 | Serine/thre | Stk25 | sp\|Q9Z2W | 8 | 2 | 20,4 |
| P27790 | P27790 | Major cent | Cenpb | sp\|P27790 | 4 | 4 | 10,2 |
| O70305 | O70305 | Ataxin-2 | Atxn2 | sp\|O70305 | 22 | 22 | 25,5 |
| Q8K215 | Q8K215 | LYR motif- | Lyrm4 | sp\|Q8K215 | 5 | 5 | 40,7 |
| Q8R0F5 | Q8R0F5 | RNA-bindir | Rbmx2 | sp\|Q8R0F5 | 4 | 4 | 13,8 |
| Q9D1Q1 | Q9D1Q1 | M-phase p | Mphosph6 | sp\|Q9D1Q | 4 | 4 | 21,1 |
| Q6NXI6 | Q6NXI6 | Regulation | Rprd2 | sp\|Q6NXI6 | 33 | 33 | 39,7 |
| Q8R5C8 | Q8R5C8 | Zinc finger | Zmynd11 | sp\|Q8R5C8 | 7 | 7 | 15,9 |
| Q3UMB9 | Q3UMB9 | WASH com | Kiaa1033 | sp\|Q3UME | 20 | 20 | 22,1 |
| Q61545 | Q61545 | RNA-bindir | Ewsr1 | sp\|Q61545 | 8 | 8 | 16 |
| Q920L1 | Q920L1 | Fatty acid | Fads1 | sp\|Q920L1 | 11 | 11 | 28 |
| Q61420 | Q61420 | CMP-sialic | Slc35a1 | sp\|Q61420 | 2 | 2 | 7,7 |
| C0HKG6;C0 | C0HKG6;C0HKG5 | | | sp\|C0HKG6 | 5 | 5 | 19,3 |
| Q3UDK1 | Q3UDK1 | TRAF-type | Trafd1 | sp\|Q3UDK | 2 | 2 | 5,2 |
| Q60931 | Q60931 | Voltage-de | Vdac3 | sp\|Q60931 | 14 | 14 | 58,7 |
| Q9CPZ8 | Q9CPZ8 | COX assem | Cmc1 | sp\|Q9CPZ8 | 4 | 4 | 43,4 |
| P49962 | P49962 | Signal reco | Srp9 | sp\|P49962 | 3 | 3 | 34,9 |
| Q69Z99 | Q69Z99 | Zinc finger | Znf512 | sp\|Q69Z99 | 17 | 17 | 44,5 |
| Q5SRY7;Q3 | Q5SRY7;Q3UL | F-box/WD | Fbxw11;Bt | sp\|Q5SRY7 | 10 | 10 | 22,1 |
| Q8K358 | Q8K358 | Phosphatid | Pigu | sp\|Q8K358 | 4 | 4 | 11,1 |
| P62835 | P62835 | Ras-related | Rap1a | sp\|P62835 | 14 | 5 | 78,3 |
| Q149C2 | Q149C2 | TRAF3-inte | Traf3ip1 | sp\|Q149C2 | 3 | 3 | 4,8 |
| Q9DB27 | Q9DB27 | Malignant | Mcts1 | sp\|Q9DB27 | 7 | 4 | 44,8 |
| Q9WVD5 | Q9WVD5 | Mitochond | Slc25a15 | sp\|Q9WVD | 4 | 4 | 18,3 |
| P61759 | P61759 | Prefoldin s | Vbp1 | sp\|P61759 | 9 | 9 | 52 |
| P16675 | P16675 | Lysosomal | Ctsa | sp\|P16675 | 11 | 11 | 27,2 |
| Q9CY66 | Q9CY66 | H/ACA ribo | Gar1 | sp\|Q9CY66 | 8 | 8 | 32,5 |
| Q9CPU4 | Q9CPU4 | Microsoma | Mgst3 | sp\|Q9CPU4 | 4 | 4 | 25,5 |
| P21300 | P21300 | Aldose red | Akr1b7 | sp\|P21300 | 6 | 3 | 14,6 |
| Q8R2Y3 | Q8R2Y3 | Dolichol ki | Dolk | sp\|Q8R2Y3 | 2 | 2 | 4,1 |
| Q8VBV3 | Q8VBV3 | Exosome c | Exosc2 | sp\|Q8VBV3 | 10 | 10 | 45,7 |
| Q3U3E2 | Q3U3E2 | Protein FA | Fam117b | sp\|Q3U3E2 | 2 | 2 | 5,1 |
| Q8BFQ9 | Q8BFQ9 | Kelch-like | Klhl42 | sp\|Q8BFQ9 | 7 | 7 | 16,8 |
| Q8BJM7 | Q8BJM7 | S-adenosyl | Tyw1 | sp\|Q8BJM | 12 | 12 | 22,1 |
| P62307 | P62307 | Small nucle | Snrpf | sp\|P62307 | 7 | 7 | 48,8 |
| Q921L3 | Q921L3 | Transmem | Tmco1 | sp\|Q921L3 | 2 | 2 | 11,7 |
| Q8C0M0 | Q8C0M0 | WD repeat | Wdr59 | sp\|Q8C0M | 4 | 4 | 5,5 |

| | | | | | | | |
|---|---|---|---|---|---|---|---|
| Q9WVJ3 | Q9WVJ3 | Carboxype | Cpq | sp\|Q9WVJ | 7 | 7 | 21,3 |
| Q9WTK7 | Q9WTK7 | Serine/thre | Stk11 | sp\|Q9WTK | 4 | 4 | 13,5 |
| Q8R418 | Q8R418 | Endoribonu | Dicer1 | sp\|Q8R418 | 33 | 33 | 23,1 |
| Q64321 | Q64321 | Zinc finger | Zbtb7b | sp\|Q64321 | 4 | 4 | 13,2 |
| Q8VCG9 | Q8VCG9 | Regulatory | Rfxap | sp\|Q8VCG | 2 | 2 | 13,9 |
| P24860 | P24860 | G2/mitotic | Ccnb1 | sp\|P24860 | 9 | 9 | 27,7 |
| O88441 | O88441 | Metaxin-2 | Mtx2 | sp\|O88441 | 8 | 8 | 46 |
| Q9D011 | Q9D011 | M-phase-s | Mplkip | sp\|Q9D011 | 1 | 1 | 14 |
| O35683 | O35683 | NADH dehy | Ndufa1 | sp\|O35683 | 1 | 1 | 14,3 |
| Q9CQ86 | Q9CQ86 | Migration a | Mien1 | sp\|Q9CQ8 | 3 | 3 | 24,3 |
| Q9WVH4 | Q9WVH4 | Forkhead b | Foxo3 | sp\|Q9WVH | 2 | 2 | 4,9 |
| Q76LS9 | Q76LS9 | Protein FA | Fam63a | sp\|Q76LS9 | 6 | 6 | 17,7 |
| Q80W54 | Q80W54 | CAAX pren | Zmpste24 | sp\|Q80W5 | 11 | 11 | 24,6 |
| Q9D924 | Q9D924 | Iron-sulfur | Isca1 | sp\|Q9D924 | 5 | 5 | 51,9 |
| Q3UZP0 | Q3UZP0 | MARVEL do | Marveld2 | sp\|Q3UZP0 | 1 | 1 | 2,9 |
| Q6PFX9 | Q6PFX9 | Tankyrase- | Tnks | sp\|Q6PFX9 | 2 | 1 | 1,7 |
| Q9JJI6 | Q9JJI6 | GPI ethano | Pigo | sp\|Q9JJI6\| | 5 | 5 | 6,3 |
| Q9D883 | Q9D883 | Splicing fac | U2af1 | sp\|Q9D883 | 11 | 7 | 46,4 |
| Q9CZ04 | Q9CZ04 | COP9 signa | Cops7a | sp\|Q9CZ04 | 12 | 12 | 53,1 |
| Q9CQG6 | Q9CQG6 | Transmem | Tmem147 | sp\|Q9CQG | 4 | 4 | 20,1 |
| Q9D1K2 | Q9D1K2 | V-type pro | Atp6v1f | sp\|Q9D1K2 | 6 | 6 | 68,1 |
| Q61183;Q9 | Q61183 | Poly(A) pol | Papola | sp\|Q61183 | 27 | 24 | 47 |
| CON__P02 | CON__P02668 | | | | 2 | 2 | 16,6 |
| Q9DC71 | Q9DC71 | 28S riboso | Mrps15 | sp\|Q9DC71 | 10 | 10 | 36 |
| O35295 | O35295 | Transcripti | Purb | sp\|O35295 | 10 | 10 | 49,1 |
| Q5XPT3 | Q5XPT3 | Glycosyltra | Gyltl1b | sp\|Q5XPT3 | 1 | 1 | 2 |
| P56379 | P56379 | 6.8 kDa mi | Mp68 | sp\|P56379 | 2 | 2 | 29,3 |
| O70443 | O70443 | Guanine nu | Gnaz | sp\|O70443 | 3 | 2 | 10,7 |
| Q921G7 | Q921G7 | Electron tr | Etfdh | sp\|Q921G7 | 16 | 16 | 34,4 |
| B2RRL2 | B2RRL2 | Jerky prote | Jrkl | sp\|B2RRL2 | 4 | 4 | 11,7 |
| Q9D0Y8 | Q9D0Y8 | 39S riboso | Mrpl52 | sp\|Q9D0Y8 | 3 | 3 | 40,5 |
| Q66GT5 | Q66GT5 | Phosphatid | Ptpmt1 | sp\|Q66GT5 | 6 | 6 | 35,8 |
| Q9DCU6 | Q9DCU6 | 39S riboso | Mrpl4 | sp\|Q9DCU | 8 | 8 | 32 |
| P62281 | P62281 | 40S riboso | Rps11 | sp\|P62281 | 12 | 12 | 55,7 |
| P35979 | P35979 | 60S riboso | Rpl12 | sp\|P35979 | 9 | 9 | 63,6 |
| Q9Z104 | Q9Z104 | SWI/SNF-re | Hmg20b | sp\|Q9Z104 | 4 | 4 | 19,6 |
| Q05928 | Q05928 | Probetacel | Btc | sp\|Q05928 | 1 | 1 | 7,9 |
| Q9CQL6 | Q9CQL6 | 39S riboso | Mrpl35 | sp\|Q9CQL6 | 1 | 1 | 7,4 |
| Q60865 | Q60865 | Caprin-1 | Caprin1 | sp\|Q60865 | 18 | 18 | 30,8 |
| Q8BG94 | Q8BG94 | COMM dom | Commd7 | sp\|Q8BG94 | 5 | 5 | 28,5 |
| A6H6A9 | A6H6A9 | Rab GTPas | Rabgap1l | sp\|A6H6A9 | 5 | 5 | 7,6 |
| Q4G0F8 | Q4G0F8 | Ubinuclein | Ubn1 | sp\|Q4G0F8 | 3 | 3 | 3,2 |
| Q3TRM4 | Q3TRM4 | Neuropath | Pnpla6 | sp\|Q3TRM | 16 | 15 | 17,4 |
| O09110 | O09110 | Dual specif | Map2k3 | sp\|O09110 | 13 | 12 | 53 |
| Q9DCI3 | Q9DCI3 | MLN64 N-t | Stard3nl | sp\|Q9DCI3 | 2 | 2 | 9,8 |
| Q8BMZ5 | Q8BMZ5 | tRNA-splici | Tsen34 | sp\|Q8BMZ | 12 | 12 | 40,8 |
| Q9JLG8 | Q9JLG8 | Calpain-15 | Capn15 | sp\|Q9JLG8 | 1 | 1 | 2,3 |
| Q3UJD6 | Q3UJD6 | Ubiquitin c | Usp19 | sp\|Q3UJD6 | 34 | 34 | 37,1 |
| P28741 | P28741 | Kinesin-like | Kif3a | sp\|P28741 | 12 | 10 | 23,3 |
| Q9D0Q7 | Q9D0Q7 | 39S riboso | Mrpl45 | sp\|Q9D0Q | 13 | 13 | 51,3 |

| | | | | | | | |
|---|---|---|---|---|---|---|---|
| Q8BQ30 | Q8BQ30 | Phostensin | Ppp1r18 | sp\|Q8BQ3 | 19 | 19 | 44,4 |
| P36895;P3 | P36895;P3689 | Bone morp | Bmpr1a;Br | sp\|P36895 | 4 | 4 | 10 |
| Q7TPM1 | Q7TPM1 | Protein PR | Prrc2b | sp\|Q7TPM | 23 | 21 | 25,2 |
| P61963 | P61963 | DDB1- and | Dcaf7 | sp\|P61963 | 10 | 10 | 35,1 |
| Q9EPK6 | Q9EPK6 | Nucleotide | Sil1 | sp\|Q9EPK6 | 9 | 9 | 24,9 |
| Q7TMV3 | Q7TMV3 | FAST kinas | Fastkd5 | sp\|Q7TMV | 11 | 11 | 22,7 |
| Q9D1M4 | Q9D1M4 | Eukaryotic | Eef1e1 | sp\|Q9D1M | 8 | 8 | 52,9 |
| Q8VCS6 | Q8VCS6 | Mediator c | Med9 | sp\|Q8VCS6 | 2 | 2 | 26,8 |
| O88513 | O88513 | Geminin | Gmnn | sp\|O88513 | 4 | 4 | 27,7 |
| Q3UZW7 | Q3UZW7 | Protein-lys | Eef2kmt | sp\|Q3UZW | 6 | 6 | 27,8 |
| Q8BIA4 | Q8BIA4 | F-box/WD | Fbxw8 | sp\|Q8BIA4 | 5 | 5 | 12,5 |
| Q8R3U1 | Q8R3U1 | HRAS-like s | Pla2g16 | sp\|Q8R3U | 2 | 2 | 14,2 |
| Q8JZQ9 | Q8JZQ9 | Eukaryotic | Eif3b | sp\|Q8JZQ9 | 46 | 46 | 61,4 |
| Q9JJ11 | Q9JJ11 | Transformi | Tacc3 | sp\|Q9JJ11 | 14 | 14 | 29,6 |
| Q8BFV3 | Q8BFV3 | Dual specif | Dusp4 | sp\|Q8BFV3 | 2 | 2 | 7,3 |
| Q80U58 | Q80U58 | Pumilio ho | Pum2 | sp\|Q80U58 | 20 | 17 | 25 |
| Q9DCM7 | Q9DCM7 | Nucleus ac | Nacc2 | sp\|Q9DCM | 3 | 2 | 6,7 |
| Q8R2Q4 | Q8R2Q4 | Ribosome- | Gfm2 | sp\|Q8R2Q | 14 | 14 | 24,1 |
| Q64028 | Q64028 | Polyhomeo | Phc1 | sp\|Q64028 | 4 | 4 | 10,8 |
| Q9JIY2 | Q9JIY2 | E3 ubiquiti | Cbll1 | sp\|Q9JIY2 | 5 | 5 | 18,5 |
| Q60772 | Q60772 | Cyclin-dep | Cdkn2c | sp\|Q60772 | 5 | 5 | 42,9 |
| Q6P5E8 | Q6P5E8 | Diacylglyce | Dgkq | sp\|Q6P5E8 | 1 | 1 | 1,5 |
| P35585;Q9 | P35585 | AP-1 comp | Ap1m1 | sp\|P35585 | 20 | 20 | 58,4 |
| Q9Z207 | Q9Z207 | Protein dia | Diaph3 | sp\|Q9Z207 | 29 | 29 | 31,6 |
| Q91YT8 | Q91YT8 | CSC1-like p | Tmem63a | sp\|Q91YT8 | 11 | 11 | 17,2 |
| Q99PN3 | Q99PN3 | Tripartite r | Trim26 | sp\|Q99PN3 | 2 | 2 | 5,3 |
| Q9CR09 | Q9CR09 | Ubiquitin-f | Ufc1 | sp\|Q9CR09 | 5 | 5 | 29,3 |
| Q9CQW7 | Q9CQW7 | Apoptogen | Apopt1 | sp\|Q9CQW | 2 | 2 | 16,1 |
| P26369 | P26369 | Splicing fac | U2af2 | sp\|P26369 | 15 | 15 | 48,6 |
| O54946;Q9 | O54946 | DnaJ homo | Dnajb6 | sp\|O54946 | 8 | 7 | 31,5 |
| O88587 | O88587 | Catechol O | Comt | sp\|O88587 | 13 | 13 | 61,5 |
| Q9EQW7 | Q9EQW7 | Kinesin-like | Kif13a | sp\|Q9EQW | 7 | 6 | 5,4 |
| P62267 | P62267 | 40S riboso | Rps23 | sp\|P62267 | 9 | 9 | 49,7 |
| Q9JLM4 | Q9JLM4 | Zinc finger | Zmym3 | sp\|Q9JLM4 | 4 | 4 | 4,2 |
| Q9QYI3 | Q9QYI3 | DnaJ homo | Dnajc7 | sp\|Q9QYI3 | 29 | 29 | 55,1 |
| Q6ZWN5 | Q6ZWN5 | 40S riboso | Rps9 | sp\|Q6ZWN | 17 | 17 | 55,2 |
| O55176 | O55176 | E3 ubiquiti | Pja1 | sp\|O55176 | 5 | 5 | 14,2 |
| Q810U5 | Q810U5 | Coiled-coil | Ccdc50 | sp\|Q810U5 | 5 | 5 | 17,7 |
| Q9CQY1 | Q9CQY1 | Ubiquitin-l | Atg12 | sp\|Q9CQY | 1 | 1 | 7,1 |
| Q6PB51 | Q6PB51 | Coiled-coil | Ccdc117 | sp\|Q6PB51 | 4 | 4 | 27,1 |
| Q99MR3 | Q99MR3 | Solute carr | Slc12a9 | sp\|Q99MR | 12 | 12 | 18,5 |
| O70435 | O70435 | Proteasom | Psma3 | sp\|O70435 | 11 | 11 | 42,4 |
| Q9EQI8 | Q9EQI8 | 39S riboso | Mrpl46 | sp\|Q9EQI8 | 12 | 12 | 47,7 |
| Q8BIG7 | Q8BIG7 | Catechol O | Comtd1 | sp\|Q8BIG7 | 5 | 5 | 27,5 |
| Q7TMX5 | Q7TMX5 | Protein SH | Shq1 | sp\|Q7TMX | 8 | 8 | 25,7 |
| Q3UHD6 | Q3UHD6 | Sorting nex | Snx27 | sp\|Q3UHD | 19 | 18 | 48,4 |
| Q9CWQ3 | Q9CWQ3 | Mitochond | Xrcc6bp1 | sp\|Q9CWQ | 1 | 1 | 8 |
| P62264 | P62264 | 40S riboso | Rps14 | sp\|P62264 | 9 | 9 | 65,6 |
| Q99PP2 | Q99PP2 | Zinc finger | Znf318 | sp\|Q99PP2 | 4 | 4 | 2,2 |
| P62751 | P62751 | 60S riboso | Rpl23a | sp\|P62751 | 11 | 11 | 49,4 |

| | | | | | | | |
|---|---|---|---|---|---|---|---|
| Q9Z0R0 | Q9Z0R0 | Serine/thre | Gsg2 | sp\|Q9Z0R0 | 10 | 10 | 20,4 |
| O08583 | O08583 | THO comp | Alyref | sp\|O08583 | 10 | 6 | 46,7 |
| Q9EQR6 | Q9EQR6 | Fanconi an | Fancg | sp\|Q9EQR | 5 | 5 | 11,9 |
| Q8R143 | Q8R143 | Pituitary tu | Pttg1ip | sp\|Q8R143 | 4 | 4 | 18,4 |
| Q9QUI0 | Q9QUI0 | Transformi | Rhoa | sp\|Q9QUI0 | 11 | 3 | 72 |
| Q5I1X5 | Q5I1X5 | RelA-assoc | Ppp1r13l | sp\|Q5I1X5 | 14 | 14 | 26,1 |
| Q91X84 | Q91X84 | CREB-regu | Crtc3 | sp\|Q91X84 | 5 | 5 | 12,6 |
| Q9D483 | Q9D483 | DNA-direct | Polr3c | sp\|Q9D483 | 9 | 9 | 23,3 |
| O35226 | O35226 | 26S protea | Psmd4 | sp\|O35226 | 14 | 14 | 45,5 |
| P05977;P0 | P05977;P0954 | Myosin ligh | Myl1;Myl3 | sp\|P05977 | 2 | 2 | 8,5 |
| Q62158 | Q62158 | Zinc finger | Trim27 | sp\|Q62158 | 4 | 4 | 10,5 |
| P15920 | P15920 | V-type pro | Atp6v0a2 | sp\|P15920 | 11 | 10 | 17,2 |
| Q9JJA2 | Q9JJA2 | Conserved | Cog8 | sp\|Q9JJA2 | 12 | 12 | 29,5 |
| Q8VDN4 | Q8VDN4 | Coiled-coil | Ccdc92 | sp\|Q8VDN | 3 | 3 | 17,8 |
| Q3V300 | Q3V300 | Kinesin-like | Kif22 | sp\|Q3V300 | 22 | 22 | 53,6 |
| Q6NXM2 | Q6NXM2 | RCC1 and B | Rcbtb1 | sp\|Q6NXM | 3 | 3 | 8,7 |
| Q9Z1T6 | Q9Z1T6 | 1-phospha | Pikfyve | sp\|Q9Z1T6 | 5 | 5 | 3,4 |
| Q8C7X2 | Q8C7X2 | ER membra | Emc1 | sp\|Q8C7X2 | 29 | 29 | 38,8 |
| Q32NY4 | Q32NY4 | Metal tran | Cnnm3 | sp\|Q32NY4 | 9 | 9 | 16,8 |
| Q66T02 | Q66T02 | Pleckstrin | Plekhg5 | sp\|Q66T02 | 1 | 1 | 1,1 |
| Q9CQZ5 | Q9CQZ5 | NADH dehy | Ndufa6 | sp\|Q9CQZ | 6 | 6 | 51,9 |
| Q8K2Y0 | Q8K2Y0 | RING finge | Rnf219 | sp\|Q8K2Y0 | 6 | 6 | 10,2 |
| B1AUE5 | B1AUE5 | Peroxisom | Pex10 | sp\|B1AUE5 | 2 | 2 | 9,6 |
| Q8BSI6 | Q8BSI6 | R3H and co | R3hcc1 | sp\|Q8BSI6 | 8 | 8 | 28,3 |
| Q8K382 | Q8K382 | DENN dom | Dennd1a | sp\|Q8K382 | 5 | 5 | 6,7 |
| O08547 | O08547 | Vesicle-tra | Sec22b | sp\|O08547 | 14 | 14 | 60,9 |
| O88888 | O88888 | Amyloid be | Apba3 | sp\|O88888 | 2 | 2 | 3,5 |
| CON__Q28 | CON__Q28107 | | | | 8 | 8 | 5,8 |
| D3Z2R5 | D3Z2R5 | | Sepn1 | sp\|D3Z2R5 | 4 | 4 | 12,7 |
| Q9CY58 | Q9CY58 | Plasminoge | Serbp1 | sp\|Q9CY58 | 28 | 28 | 60,4 |
| Q9D9I4 | Q9D9I4 | TBC1 doma | Tbc1d20 | sp\|Q9D9I4 | 6 | 6 | 20,1 |
| Q7TMY4 | Q7TMY4 | THO comp | Thoc7 | sp\|Q7TMY | 9 | 9 | 36,8 |
| Q60875 | Q60875 | Rho guanir | Arhgef2 | sp\|Q60875 | 41 | 41 | 48,7 |
| Q8R2S9 | Q8R2S9 | Actin-relat | Actr8 | sp\|Q8R2S9 | 10 | 10 | 21,3 |
| Q8BGY7 | Q8BGY7 | Protein FA | Fam210a | sp\|Q8BGY7 | 3 | 3 | 11,7 |
| Q8JZN7 | Q8JZN7 | Mitochond | Rhot2 | sp\|Q8JZN7 | 16 | 15 | 33,7 |
| P00860 | P00860 | Ornithine c | Odc1 | sp\|P00860 | 8 | 8 | 23,6 |
| P62754 | P62754 | 40S riboso | Rps6 | sp\|P62754 | 11 | 11 | 39,4 |
| Q9ER38 | Q9ER38 | Torsin-3A | Tor3a | sp\|Q9ER38 | 8 | 8 | 30,9 |
| Q6NVE9 | Q6NVE9 | Protein ph | Pptc7 | sp\|Q6NVE | 6 | 6 | 29,4 |
| Q9QUH0 | Q9QUH0 | Glutaredox | Glrx | sp\|Q9QUH | 5 | 5 | 50,5 |
| P63154 | P63154 | Crooked ne | Crnkl1 | sp\|P63154 | 27 | 27 | 43,5 |
| Q9WUB0 | Q9WUB0 | RanBP-type | Rbck1 | sp\|Q9WUE | 8 | 8 | 24 |
| Q9CQ82 | Q9CQ82 | Centromer | Itgb3bp | sp\|Q9CQ8 | 2 | 2 | 13,1 |
| Q8VED8 | Q8VED8 | Mitochond | Mtfr2 | sp\|Q8VED | 3 | 3 | 11,9 |
| O88291 | O88291 | DBIRD com | Znf326 | sp\|O88291 | 15 | 15 | 31,4 |
| Q9R1P1 | Q9R1P1 | Proteasom | Psmb3 | sp\|Q9R1P1 | 10 | 10 | 50,2 |
| P19253 | P19253 | 60S riboso | Rpl13a | sp\|P19253 | 10 | 10 | 35 |
| Q8CB27 | Q8CB27 | Ubiquitin t | Yod1 | sp\|Q8CB27 | 2 | 2 | 7,9 |
| Q8CI71 | Q8CI71 | Coiled-coil | Ccdc132 | sp\|Q8CI71 | 20 | 20 | 30,1 |

| | | | | | | | |
|---|---|---|---|---|---|---|---|
| Q8BGV4 | Q8BGV4 | TELO2-inte | Tti2 | sp\|Q8BGV | 4 | 4 | 10,7 |
| P59708 | P59708 | Splicing fac | Sf3b6 | sp\|P59708 | 6 | 6 | 54,4 |
| Q3TC46 | Q3TC46 | Protein PA | Patl1 | sp\|Q3TC46 | 16 | 16 | 26,4 |
| Q9DCL4 | Q9DCL4 | Probable n | Mettl15 | sp\|Q9DCL4 | 3 | 3 | 12,6 |
| Q9QYE6 | Q9QYE6 | Golgin sub | Golga5 | sp\|Q9QYE6 | 19 | 19 | 35,8 |
| P31001 | P31001 | Desmin | Des | sp\|P31001 | 9 | 3 | 12,8 |
| O35250 | O35250 | Exocyst co | Exoc7 | sp\|O35250 | 20 | 20 | 33,9 |
| Q9D0V8 | Q9D0V8 | Cyclin-dep | Cinp | sp\|Q9D0V | 2 | 2 | 11,8 |
| Q9CZV5 | Q9CZV5 | STAGA con | Supt7l | sp\|Q9CZV5 | 6 | 6 | 28,6 |
| Q9DCL2 | Q9DCL2 | MIP18 fam | Fam96a | sp\|Q9DCL2 | 4 | 4 | 30 |
| P58334 | P58334 | Krueppel-li | Klf16 | sp\|P58334 | 2 | 2 | 19,1 |
| Q80XH1 | Q80XH1 | KxDL motif | Kxd1 | sp\|Q80XH1 | 3 | 3 | 22,6 |
| P62843 | P62843 | 40S riboso | Rps15 | sp\|P62843 | 6 | 6 | 49 |
| Q6ZPT1 | Q6ZPT1 | Kelch-like | Klhl9 | sp\|Q6ZPT1 | 12 | 9 | 26,7 |

| | | | | | C | |
|---|---|---|---|---|---|---|
| Unique sequence coverage [%] | Potential contaminant | p-value 'OE'/'C' | q-value 'OE'/'C' | Ratio 'OE'/'C' | LFQ intensity C1 | LFQ intensity C2 |
| 55,8 | | 0,0000001 | 0,0000000 | 1,32 | 1,31E+09 | 1,31E+09 |
| 22,7 | | 0,0000001 | 0,0000000 | 0,04 | 1,39E+08 | 1,26E+08 |
| 50 | | 0,0000002 | 0,0000000 | 0,12 | 1,18E+08 | 1,27E+08 |
| 35,3 | | 0,0000002 | 0,0000000 | 0,01 | 1,33E+09 | 1,50E+09 |
| 75,2 | | 0,0000002 | 0,0000000 | 0,38 | 4,70E+09 | 4,81E+09 |
| 75,9 | | 0,0000003 | 0,0000000 | 0,25 | 7,87E+09 | 8,06E+09 |
| 42,8 | | 0,0000003 | 0,0000000 | 0,24 | 8,16E+08 | 8,15E+08 |
| 56,8 | | 0,0000005 | 0,0000000 | 0,57 | 1,84E+09 | 1,84E+09 |
| 31 | | 0,0000005 | 0,0000000 | 0,47 | 4,16E+08 | 4,34E+08 |
| 20,9 | | 0,0000006 | 0,0000000 | 2,83 | 3,44E+07 | 3,57E+07 |
| 40,4 | | 0,0000006 | 0,0000000 | 2,79 | 8,39E+07 | 8,59E+07 |
| 50,3 | | 0,0000007 | 0,0000000 | 0,26 | 1,40E+09 | 1,41E+09 |
| 23,2 | | 0,0000007 | 0,0000000 | 0,36 | 7,88E+07 | 8,32E+07 |
| 32,1 | | 0,0000007 | 0,0000000 | 21,44 | 6,58E+07 | 6,22E+07 |
| 29,6 | | 0,0000007 | 0,0000000 | 0,22 | 4,94E+08 | 4,97E+08 |
| 46,5 | | 0,0000008 | 0,0000000 | 2,87 | 3,16E+08 | 3,30E+08 |
| 37 | | 0,0000008 | 0,0000000 | 0,34 | 2,82E+08 | 2,65E+08 |
| 56,9 | | 0,0000009 | 0,0000000 | 0,34 | 1,31E+08 | 1,31E+08 |
| 66,3 | | 0,0000010 | 0,0000000 | 4,61 | 4,45E+08 | 4,52E+08 |
| 64,9 | | 0,0000010 | 0,0000000 | 0,00 | 1,91E+09 | 1,93E+09 |
| 31,8 | | 0,0000010 | 0,0000000 | 0,00 | 9,63E+08 | 1,42E+09 |
| 40,7 | | 0,0000011 | 0,0000000 | 1,45 | 5,42E+08 | 5,48E+08 |
| 61,3 | | 0,0000011 | 0,0000000 | 0,60 | 2,67E+09 | 2,64E+09 |
| 31,7 | | 0,0000014 | 0,0000000 | 0,21 | 8,07E+08 | 8,59E+08 |
| 30,7 | | 0,0000017 | 0,0000000 | 0,71 | 3,33E+08 | 3,28E+08 |
| 47,9 | | 0,0000017 | 0,0000000 | 0,02 | 2,25E+08 | 2,08E+08 |
| 24,7 | | 0,0000019 | 0,0000000 | 0,21 | 7,20E+07 | 6,88E+07 |
| 55,8 | | 0,0000019 | 0,0000000 | 2,39 | 6,25E+08 | 5,93E+08 |
| 6,8 | | 0,0000019 | 0,0000000 | 0,44 | 3,82E+07 | 3,66E+07 |
| 40,9 | | 0,0000021 | 0,0000000 | 0,18 | 4,07E+08 | 4,23E+08 |
| 20,7 | | 0,0000021 | 0,0000000 | 1,82 | 1,35E+07 | 1,29E+07 |
| 42,3 | | 0,0000021 | 0,0000000 | 0,24 | 8,88E+08 | 8,45E+08 |
| 53,8 | | 0,0000021 | 0,0000000 | 2,13 | 1,59E+09 | 1,60E+09 |
| 70 | | 0,0000022 | 0,0000000 | 0,19 | 1,50E+09 | 1,52E+09 |
| 33,6 | | 0,0000023 | 0,0000000 | 13,44 | 4,23E+06 | 3,97E+06 |
| 63,2 | | 0,0000023 | 0,0000000 | 0,21 | 4,62E+08 | 4,19E+08 |
| 53,3 | + | 0,0000023 | 0,0000000 | 3,56 | 7,70E+08 | 8,35E+08 |
| 61,8 | | 0,0000025 | 0,0000000 | 78,82 | 4,23E+07 | 5,48E+07 |
| 44,8 | | 0,0000025 | 0,0000000 | 2,05 | 1,60E+08 | 1,63E+08 |
| 80,2 | | 0,0000025 | 0,0000000 | 1,83 | 5,67E+09 | 5,64E+09 |
| 49,8 | | 0,0000026 | 0,0000000 | 2,94 | 6,80E+08 | 7,17E+08 |
| 34,8 | | 0,0000032 | 0,0000000 | 1,92 | 7,76E+07 | 8,16E+07 |
| 22,2 | | 0,0000033 | 0,0000000 | 40,24 | 3,25E+06 | 4,49E+06 |
| 51 | | 0,0000034 | 0,0000000 | 0,04 | 2,17E+08 | 2,48E+08 |
| 13,1 | | 0,0000036 | 0,0000000 | 0,48 | 5,41E+08 | 5,63E+08 |
| 48,7 | | 0,0000038 | 0,0000000 | 3,43 | 2,07E+08 | 1,90E+08 |

| | | | | | | |
|---|---|---|---|---|---|---|
| 53,7 | | 0,0000038 | 0,0000000 | 1,77 | 9,17E+08 | 9,05E+08 |
| 41 | | 0,0000039 | 0,0000000 | 0,55 | 2,28E+08 | 2,21E+08 |
| 27,5 | | 0,0000039 | 0,0000000 | 6,28 | 1,32E+07 | 1,26E+07 |
| 56,5 | | 0,0000040 | 0,0000000 | 1,67 | 1,46E+09 | 1,45E+09 |
| 46,8 | | 0,0000042 | 0,0000000 | 0,40 | 7,86E+08 | 7,78E+08 |
| 55,9 | | 0,0000044 | 0,0000000 | 1,60 | 6,94E+08 | 7,03E+08 |
| 46,9 | | 0,0000045 | 0,0000000 | 1,34 | 1,89E+08 | 1,93E+08 |
| 24,9 | | 0,0000045 | 0,0000000 | 2,77 | 4,82E+07 | 5,08E+07 |
| 15,9 | | 0,0000045 | 0,0000000 | 4,39 | 6,67E+07 | 6,26E+07 |
| 33,3 | | 0,0000046 | 0,0000000 | 0,40 | 2,81E+08 | 2,63E+08 |
| 22,3 | | 0,0000047 | 0,0000000 | 0,51 | 1,36E+08 | 1,38E+08 |
| 31,3 | | 0,0000050 | 0,0000000 | 2,45 | 6,52E+07 | 6,65E+07 |
| 15,8 | | 0,0000050 | 0,0000000 | 63,65 | 1,73E+06 | 2,63E+06 |
| 49,4 | | 0,0000051 | 0,0000000 | 0,01 | 5,91E+08 | 6,43E+08 |
| 54 | | 0,0000051 | 0,0000000 | 2,42 | 2,41E+08 | 2,53E+08 |
| 34,4 | | 0,0000053 | 0,0000000 | 1,85 | 2,47E+08 | 2,60E+08 |
| 38,6 | | 0,0000053 | 0,0000000 | 3,28 | 2,09E+08 | 1,88E+08 |
| 43,8 | | 0,0000054 | 0,0000000 | 3,25 | 2,32E+08 | 2,15E+08 |
| 32,9 | | 0,0000054 | 0,0000000 | 4,99 | 8,75E+07 | 8,35E+07 |
| 49,2 | | 0,0000056 | 0,0000000 | 2,12 | 1,20E+08 | 1,19E+08 |
| 56,4 | | 0,0000058 | 0,0000000 | 0,30 | 1,50E+09 | 1,66E+09 |
| 50 | | 0,0000063 | 0,0000000 | 2,83 | 1,11E+08 | 1,08E+08 |
| 19,3 | | 0,0000064 | 0,0000000 | 0,01 | 4,14E+08 | 3,75E+08 |
| 7,8 | | 0,0000067 | 0,0000000 | 0,01 | 1,55E+08 | 1,42E+08 |
| 8,3 | | 0,0000067 | 0,0000000 | 0,10 | 3,66E+07 | 3,58E+07 |
| 38,8 | | 0,0000068 | 0,0000000 | 0,13 | 2,16E+08 | 2,08E+08 |
| 28,7 | | 0,0000068 | 0,0000000 | 0,02 | 1,02E+08 | 9,54E+07 |
| 26,8 | | 0,0000068 | 0,0000000 | 0,25 | 3,18E+08 | 2,99E+08 |
| 46,7 | | 0,0000070 | 0,0000000 | 2,38 | 4,09E+08 | 4,43E+08 |
| 27,3 | | 0,0000070 | 0,0000000 | 3,33 | 6,90E+07 | 7,45E+07 |
| 10,7 | | 0,0000070 | 0,0000000 | 7,77 | 8,66E+06 | 8,15E+06 |
| 16,1 | | 0,0000073 | 0,0000000 | 11,62 | 2,33E+06 | 2,27E+06 |
| 28,8 | | 0,0000074 | 0,0000000 | 2,68 | 1,03E+08 | 1,09E+08 |
| 28,6 | | 0,0000074 | 0,0000000 | 0,41 | 1,24E+08 | 1,26E+08 |
| 26,2 | | 0,0000075 | 0,0000000 | 0,07 | 1,28E+08 | 1,24E+08 |
| 49 | | 0,0000077 | 0,0000000 | 3,06 | 1,54E+08 | 1,51E+08 |
| 67,4 | | 0,0000080 | 0,0000000 | 1,86 | 2,49E+08 | 2,61E+08 |
| 57,7 | | 0,0000082 | 0,0000000 | 0,22 | 7,64E+08 | 6,99E+08 |
| 53,8 | | 0,0000083 | 0,0000000 | 0,48 | 2,57E+09 | 2,62E+09 |
| 59 | | 0,0000084 | 0,0000000 | 0,39 | 8,87E+08 | 8,39E+08 |
| 66,4 | | 0,0000084 | 0,0000000 | 2,88 | 3,06E+09 | 2,89E+09 |
| 49,6 | | 0,0000085 | 0,0000000 | 3,27 | 1,67E+08 | 1,58E+08 |
| 53,6 | | 0,0000085 | 0,0000000 | 0,61 | 4,29E+08 | 4,43E+08 |
| 16,6 | | 0,0000086 | 0,0000000 | 2,22 | 5,89E+07 | 5,61E+07 |
| 20 | | 0,0000087 | 0,0000000 | 0,11 | 5,20E+08 | 4,65E+08 |
| 41 | | 0,0000088 | 0,0000000 | 4,45 | 3,20E+07 | 3,52E+07 |
| 10,5 | | 0,0000088 | 0,0000000 | 7,49 | 4,80E+06 | 5,56E+06 |
| 50,6 | | 0,0000090 | 0,0000000 | 0,00 | 2,48E+08 | 2,70E+08 |
| 46 | | 0,0000091 | 0,0000000 | 0,38 | 3,69E+08 | 3,70E+08 |
| 17,1 | | 0,0000092 | 0,0000000 | 198,42 | 1,87E+06 | 2,24E+06 |

| | | | | | | |
|---|---|---|---|---|---|---|
| 20,7 | | 0,0000093 | 0,0000000 | 0,04 | 8,82E+07 | 7,38E+07 |
| 31 | | 0,0000093 | 0,0000000 | 0,21 | 2,31E+08 | 2,06E+08 |
| 59 | | 0,0000094 | 0,0000000 | 0,15 | 2,56E+08 | 2,59E+08 |
| 58,3 | | 0,0000095 | 0,0000000 | 0,36 | 6,15E+08 | 6,42E+08 |
| 79,6 | | 0,0000097 | 0,0000000 | 2,54 | 1,97E+08 | 1,98E+08 |
| 7,6 | | 0,0000102 | 0,0000000 | 0,02 | 1,49E+08 | 1,44E+08 |
| 41,6 | | 0,0000109 | 0,0000000 | 83,57 | 2,73E+06 | 4,72E+06 |
| 23,7 | | 0,0000110 | 0,0000000 | 1,61 | 3,19E+07 | 3,14E+07 |
| 38,8 | | 0,0000110 | 0,0000000 | 1,50 | 3,54E+08 | 3,67E+08 |
| 63 | | 0,0000114 | 0,0000000 | 0,62 | 5,34E+09 | 5,32E+09 |
| 28,9 | | 0,0000114 | 0,0000000 | 0,41 | 9,35E+08 | 8,65E+08 |
| 33,6 | | 0,0000119 | 0,0000000 | 2,64 | 7,67E+07 | 8,18E+07 |
| 20,1 | | 0,0000119 | 0,0000000 | 0,68 | 7,61E+07 | 7,56E+07 |
| 47,5 | | 0,0000120 | 0,0000000 | 2,35 | 1,76E+09 | 1,89E+09 |
| 39,7 | | 0,0000122 | 0,0000000 | 1,40 | 1,57E+09 | 1,62E+09 |
| 17,9 | | 0,0000123 | 0,0000000 | 0,22 | 3,07E+07 | 2,64E+07 |
| 43,1 | | 0,0000124 | 0,0000000 | 0,00 | 3,08E+08 | 4,25E+08 |
| 37,1 | | 0,0000124 | 0,0000000 | 0,02 | 1,10E+08 | 1,11E+08 |
| 39,3 | | 0,0000124 | 0,0000000 | 0,04 | 1,06E+08 | 9,45E+07 |
| 23,6 | | 0,0000128 | 0,0000000 | 0,04 | 8,21E+07 | 7,88E+07 |
| 5,9 | | 0,0000135 | 0,0000000 | 0,11 | 1,49E+07 | 1,79E+07 |
| 46,4 | | 0,0000138 | 0,0000000 | 0,68 | 3,77E+08 | 3,67E+08 |
| 59,6 | | 0,0000140 | 0,0000000 | 2,63 | 1,75E+08 | 1,85E+08 |
| 31,6 | | 0,0000142 | 0,0000000 | 1,91 | 8,61E+07 | 8,72E+07 |
| 15,8 | | 0,0000145 | 0,0000000 | 0,02 | 1,06E+08 | 1,44E+08 |
| 78,8 | | 0,0000145 | 0,0000000 | 1,37 | 6,39E+08 | 6,51E+08 |
| 51,4 | | 0,0000151 | 0,0000000 | 0,52 | 1,17E+09 | 1,18E+09 |
| 34 | | 0,0000152 | 0,0000000 | 0,02 | 1,67E+08 | 1,53E+08 |
| 85,3 | | 0,0000157 | 0,0000000 | 2,20 | 1,64E+09 | 1,70E+09 |
| 39,8 | | 0,0000158 | 0,0000000 | 2,60 | 1,10E+09 | 1,01E+09 |
| 37,5 | | 0,0000160 | 0,0000000 | 0,68 | 3,15E+08 | 3,10E+08 |
| 35,2 | | 0,0000161 | 0,0000000 | 0,02 | 1,51E+08 | 1,42E+08 |
| 65,7 | | 0,0000161 | 0,0000000 | 1,71 | 2,42E+10 | 2,30E+10 |
| 35,1 | | 0,0000163 | 0,0000000 | 0,63 | 1,86E+08 | 1,89E+08 |
| 42,9 | | 0,0000168 | 0,0000000 | 3,09 | 6,93E+08 | 6,69E+08 |
| 43,4 | | 0,0000168 | 0,0000000 | 0,26 | 1,57E+09 | 1,70E+09 |
| 31,1 | | 0,0000172 | 0,0000000 | 0,17 | 2,20E+08 | 2,49E+08 |
| 13,9 | | 0,0000174 | 0,0000000 | 0,06 | 3,38E+07 | 3,62E+07 |
| 34,3 | | 0,0000174 | 0,0000000 | 0,52 | 6,01E+08 | 5,53E+08 |
| 35,1 | | 0,0000175 | 0,0000000 | 1,84 | 8,18E+07 | 8,58E+07 |
| 29,3 | | 0,0000175 | 0,0000000 | 0,06 | 1,40E+08 | 1,06E+08 |
| 46,4 | | 0,0000175 | 0,0000000 | 1,87 | 2,32E+08 | 2,31E+08 |
| 47,8 | | 0,0000176 | 0,0000000 | 0,02 | 1,11E+08 | 1,00E+08 |
| 61 | | 0,0000176 | 0,0000000 | 67,65 | 4,63E+06 | 3,60E+06 |
| 35,8 | | 0,0000177 | 0,0000000 | 0,50 | 1,02E+09 | 9,26E+08 |
| 14,1 | | 0,0000179 | 0,0000000 | 0,37 | 3,19E+08 | 3,27E+08 |
| 72,5 | | 0,0000182 | 0,0000000 | 2,02 | 3,46E+08 | 3,23E+08 |
| 8,5 | | 0,0000184 | 0,0000000 | 0,19 | 6,11E+07 | 7,07E+07 |
| 57,1 | | 0,0000184 | 0,0000000 | 3,21 | 9,77E+07 | 1,01E+08 |
| 32 | | 0,0000191 | 0,0000000 | 5,54 | 6,25E+07 | 5,69E+07 |

| | | | | | | |
|---|---|---|---|---|---|---|
| 38,4 | | 0,0000196 | 0,0000000 | 3,20 | 2,63E+08 | 2,90E+08 |
| 34,6 | | 0,0000196 | 0,0000000 | 1,80 | 1,96E+07 | 1,96E+07 |
| 26,6 | | 0,0000202 | 0,0000000 | 2,88 | 4,36E+07 | 4,37E+07 |
| 19,3 | | 0,0000202 | 0,0000000 | 2,34 | 4,33E+07 | 4,28E+07 |
| 34,8 | | 0,0000209 | 0,0000000 | 0,70 | 1,65E+08 | 1,62E+08 |
| 35,6 | | 0,0000211 | 0,0000000 | 3,31 | 2,01E+09 | 1,80E+09 |
| 71,8 | | 0,0000212 | 0,0000000 | 8,11 | 9,43E+07 | 9,99E+07 |
| 35,8 | | 0,0000213 | 0,0000000 | 1,61 | 2,15E+08 | 2,07E+08 |
| 57,1 | | 0,0000216 | 0,0000000 | 1,48 | 1,07E+09 | 1,04E+09 |
| 66,8 | | 0,0000216 | 0,0000000 | 1,73 | 1,21E+09 | 1,19E+09 |
| 40,3 | | 0,0000219 | 0,0000000 | 0,56 | 1,55E+08 | 1,54E+08 |
| 9,8 | | 0,0000220 | 0,0000000 | 1,28 | 3,96E+07 | 3,82E+07 |
| 32,2 | | 0,0000221 | 0,0000000 | 0,01 | 4,88E+08 | 5,67E+08 |
| 40,5 | | 0,0000226 | 0,0000000 | 0,01 | 3,38E+08 | 2,81E+08 |
| 37,9 | | 0,0000230 | 0,0000000 | 0,54 | 8,13E+08 | 8,66E+08 |
| 31 | | 0,0000233 | 0,0000000 | 0,62 | 6,74E+08 | 6,58E+08 |
| 42,8 | | 0,0000236 | 0,0000000 | 0,52 | 3,66E+08 | 3,69E+08 |
| 10,8 | | 0,0000240 | 0,0000000 | 8,43 | 2,84E+06 | 3,20E+06 |
| 28,4 | | 0,0000244 | 0,0000000 | 3,45 | 1,72E+08 | 2,05E+08 |
| 38,7 | | 0,0000247 | 0,0000000 | 0,51 | 1,14E+09 | 1,23E+09 |
| 53,3 | | 0,0000248 | 0,0000000 | 0,70 | 2,19E+08 | 2,17E+08 |
| 41,5 | | 0,0000249 | 0,0000000 | 0,00 | 6,28E+08 | 7,25E+08 |
| 19,5 | | 0,0000249 | 0,0000000 | 0,33 | 6,61E+07 | 5,82E+07 |
| 47,8 | | 0,0000249 | 0,0000000 | 2,17 | 1,06E+08 | 1,12E+08 |
| 58,6 | | 0,0000258 | 0,0000000 | 0,30 | 8,29E+08 | 7,67E+08 |
| 66,5 | | 0,0000259 | 0,0000000 | 2,97 | 9,02E+08 | 7,87E+08 |
| 20,9 | | 0,0000261 | 0,0000000 | 0,03 | 8,60E+07 | 8,38E+07 |
| 37,1 | | 0,0000262 | 0,0000000 | 0,36 | 4,30E+08 | 4,60E+08 |
| 39 | | 0,0000262 | 0,0000000 | 0,47 | 2,73E+08 | 2,57E+08 |
| 54,2 | | 0,0000268 | 0,0000000 | 0,43 | 1,19E+09 | 1,22E+09 |
| 49,9 | | 0,0000270 | 0,0000000 | 0,56 | 1,16E+09 | 1,09E+09 |
| 43 | | 0,0000272 | 0,0000000 | 1,71 | 1,43E+08 | 1,34E+08 |
| 51,7 | | 0,0000274 | 0,0000000 | 0,54 | 1,61E+08 | 1,63E+08 |
| 45,6 | | 0,0000280 | 0,0000000 | 2,22 | 8,07E+07 | 8,00E+07 |
| 26,8 | | 0,0000284 | 0,0000000 | 0,14 | 1,19E+08 | 9,02E+07 |
| 51,3 | | 0,0000285 | 0,0000000 | 85,17 | 2,70E+06 | 1,40E+06 |
| 65,1 | | 0,0000289 | 0,0000000 | 0,34 | 3,79E+09 | 4,26E+09 |
| 60,9 | | 0,0000295 | 0,0000000 | 2,52 | 3,42E+09 | 3,41E+09 |
| 30,1 | | 0,0000296 | 0,0000000 | 0,74 | 3,19E+08 | 3,17E+08 |
| 77 | | 0,0000296 | 0,0000000 | 0,67 | 3,25E+09 | 3,36E+09 |
| 14,3 | | 0,0000299 | 0,0000000 | 10,12 | 2,18E+06 | 2,50E+06 |
| 13,5 | | 0,0000300 | 0,0000000 | 4,38 | 3,12E+06 | 3,33E+06 |
| 27,6 | | 0,0000302 | 0,0000000 | 0,21 | 1,03E+08 | 8,71E+07 |
| 10,4 | | 0,0000304 | 0,0000000 | 2,00 | 8,55E+06 | 8,30E+06 |
| 25,6 | | 0,0000318 | 0,0000000 | 0,11 | 2,19E+08 | 2,68E+08 |
| 42,6 | | 0,0000320 | 0,0000000 | 0,01 | 3,13E+08 | 2,91E+08 |
| 8,4 | | 0,0000324 | 0,0000000 | 16,50 | 2,50E+06 | 3,70E+06 |
| 33,7 | | 0,0000324 | 0,0000000 | 0,37 | 9,99E+08 | 1,00E+09 |
| 42 | + | 0,0000329 | 0,0000000 | 2,35 | 1,10E+08 | 1,15E+08 |
| 22,6 | | 0,0000330 | 0,0000000 | 1,75 | 1,91E+08 | 2,06E+08 |

| | | | | | | |
|---|---|---|---|---|---|---|
| 39 | | 0,0000332 | 0,0000000 | 1,71 | 2,21E+08 | 2,19E+08 |
| 61,6 | | 0,0000337 | 0,0000000 | 0,46 | 3,97E+08 | 3,85E+08 |
| 49,2 | | 0,0000348 | 0,0000000 | 1,51 | 4,77E+08 | 4,58E+08 |
| 24,6 | | 0,0000351 | 0,0000000 | 1,51 | 6,03E+08 | 5,81E+08 |
| 43,1 | | 0,0000352 | 0,0000000 | 1,94 | 5,29E+08 | 5,12E+08 |
| 41,4 | | 0,0000353 | 0,0000000 | 1,46 | 8,91E+08 | 9,21E+08 |
| 19 | | 0,0000354 | 0,0000000 | 13,47 | 2,03E+07 | 1,69E+07 |
| 58,6 | | 0,0000354 | 0,0000000 | 1,87 | 7,96E+08 | 7,42E+08 |
| 43,9 | | 0,0000356 | 0,0000000 | 0,37 | 1,32E+09 | 1,48E+09 |
| 54,9 | | 0,0000356 | 0,0000000 | 0,40 | 3,64E+08 | 3,54E+08 |
| 48,9 | | 0,0000357 | 0,0000000 | 0,01 | 3,34E+08 | 3,67E+08 |
| 46,8 | | 0,0000360 | 0,0000000 | 1,51 | 3,54E+09 | 3,44E+09 |
| 51,1 | | 0,0000360 | 0,0000000 | 0,72 | 3,43E+08 | 3,52E+08 |
| 18,9 | | 0,0000360 | 0,0000000 | 0,33 | 1,30E+08 | 1,26E+08 |
| 40,7 | | 0,0000371 | 0,0000000 | 0,24 | 3,20E+08 | 3,94E+08 |
| 21,1 | | 0,0000374 | 0,0000000 | 1,51 | 8,92E+07 | 9,06E+07 |
| 79,5 | | 0,0000374 | 0,0000000 | 2,92 | 1,08E+08 | 1,11E+08 |
| 65,6 | + | 0,0000375 | 0,0000000 | 2,10 | 1,05E+09 | 1,03E+09 |
| 72,9 | | 0,0000378 | 0,0000000 | 2,55 | 9,78E+09 | 9,37E+09 |
| 25 | | 0,0000378 | 0,0000000 | 0,02 | 7,91E+07 | 8,21E+07 |
| 40,2 | | 0,0000381 | 0,0000000 | 1,89 | 7,78E+08 | 8,11E+08 |
| 13,8 | | 0,0000382 | 0,0000000 | 0,18 | 1,75E+07 | 1,86E+07 |
| 45,2 | | 0,0000401 | 0,0000000 | 0,39 | 4,31E+08 | 4,18E+08 |
| 61,8 | | 0,0000402 | 0,0000000 | 0,48 | 4,09E+08 | 4,61E+08 |
| 47,7 | | 0,0000403 | 0,0000000 | 1,60 | 3,37E+09 | 3,15E+09 |
| 33 | | 0,0000407 | 0,0000000 | 0,04 | 1,72E+08 | 1,66E+08 |
| 36,1 | | 0,0000409 | 0,0000000 | 0,50 | 1,93E+08 | 1,81E+08 |
| 63,9 | | 0,0000411 | 0,0000000 | 1,64 | 4,94E+09 | 5,10E+09 |
| 61,5 | | 0,0000414 | 0,0000000 | 0,74 | 1,09E+09 | 1,14E+09 |
| 34,5 | | 0,0000414 | 0,0000000 | 1,39 | 1,06E+09 | 1,04E+09 |
| 17,3 | | 0,0000415 | 0,0000000 | 0,04 | 8,26E+07 | 6,31E+07 |
| 62,5 | | 0,0000418 | 0,0000000 | 1,26 | 2,80E+09 | 2,88E+09 |
| 35,2 | | 0,0000421 | 0,0000000 | 1,55 | 5,21E+08 | 4,95E+08 |
| 45,4 | | 0,0000426 | 0,0000000 | 2,38 | 2,78E+08 | 2,99E+08 |
| 26 | | 0,0000429 | 0,0000000 | 16,42 | 2,54E+06 | 4,01E+06 |
| 27,7 | | 0,0000430 | 0,0000000 | 1,91 | 1,03E+08 | 1,02E+08 |
| 8,4 | | 0,0000431 | 0,0000000 | 0,01 | 4,80E+08 | 6,85E+08 |
| 36,5 | | 0,0000432 | 0,0000000 | 0,55 | 1,08E+09 | 1,02E+09 |
| 31,5 | | 0,0000434 | 0,0000000 | 1516,21 | 5,62E+06 | 6,93E+06 |
| 18,4 | | 0,0000434 | 0,0000000 | 0,49 | 1,11E+08 | 1,02E+08 |
| 34,2 | | 0,0000438 | 0,0000000 | 0,01 | 3,39E+08 | 3,08E+08 |
| 39,9 | | 0,0000438 | 0,0000000 | 0,64 | 1,10E+08 | 1,03E+08 |
| 43,8 | | 0,0000439 | 0,0000000 | 193,96 | 3,64E+06 | 8,38E+06 |
| 33,5 | | 0,0000439 | 0,0000000 | 0,10 | 9,23E+07 | 9,59E+07 |
| 23,9 | | 0,0000442 | 0,0005145 | 1,84 | 1,00E+08 | 1,08E+08 |
| 35,3 | | 0,0000446 | 0,0005124 | 2,12 | 1,53E+08 | 1,63E+08 |
| 38 | | 0,0000448 | 0,0005103 | 0,28 | 1,33E+08 | 1,28E+08 |
| 20,5 | | 0,0000453 | 0,0005082 | 0,51 | 2,07E+08 | 1,96E+08 |
| 42,3 | | 0,0000454 | 0,0005061 | 0,59 | 3,64E+08 | 3,82E+08 |
| 72 | | 0,0000457 | 0,0005041 | 2,11 | 6,46E+08 | 6,14E+08 |

| | | | | | | |
|---|---|---|---|---|---|---|
| 30,7 | | 0,0000469 | 0,0005020 | 0,15 | 1,78E+08 | 2,01E+08 |
| 56,9 | | 0,0000476 | 0,0005000 | 5,71 | 1,05E+08 | 7,83E+07 |
| 31,4 | | 0,0000476 | 0,0004980 | 0,69 | 2,00E+08 | 2,08E+08 |
| 44,2 | | 0,0000481 | 0,0008800 | 0,39 | 9,85E+08 | 1,17E+09 |
| 66,6 | | 0,0000486 | 0,0008765 | 0,72 | 3,26E+09 | 3,23E+09 |
| 59,4 | | 0,0000487 | 0,0008730 | 1,99 | 1,15E+08 | 1,06E+08 |
| 17,7 | | 0,0000493 | 0,0008696 | 0,62 | 1,77E+07 | 1,82E+07 |
| 47,3 | | 0,0000495 | 0,0008661 | 1,92 | 8,03E+08 | 7,65E+08 |
| 11,5 | | 0,0000503 | 0,0008627 | 0,59 | 2,53E+07 | 2,59E+07 |
| 27,2 | | 0,0000503 | 0,0008594 | 2,04 | 4,13E+08 | 4,18E+08 |
| 62,5 | | 0,0000504 | 0,0008560 | 1,68 | 6,75E+08 | 6,76E+08 |
| 28,6 | | 0,0000504 | 0,0008527 | 0,63 | 2,36E+08 | 2,48E+08 |
| 45,9 | | 0,0000505 | 0,0008494 | 0,63 | 2,62E+08 | 2,75E+08 |
| 73,1 | | 0,0000506 | 0,0008462 | 2,19 | 4,87E+08 | 4,30E+08 |
| 19,5 | | 0,0000507 | 0,0008429 | 0,61 | 2,51E+08 | 2,45E+08 |
| 9,1 | | 0,0000513 | 0,0008397 | 0,07 | 4,24E+07 | 3,17E+07 |
| 34,1 | | 0,0000518 | 0,0008365 | 0,63 | 7,87E+07 | 7,75E+07 |
| 31,5 | | 0,0000523 | 0,0008333 | 0,16 | 8,55E+07 | 8,17E+07 |
| 60,3 | | 0,0000530 | 0,0008302 | 0,26 | 3,67E+08 | 4,42E+08 |
| 25,6 | | 0,0000531 | 0,0008271 | 1,47 | 1,62E+08 | 1,54E+08 |
| 51,1 | | 0,0000533 | 0,0008240 | 1,76 | 1,46E+09 | 1,58E+09 |
| 50,5 | | 0,0000535 | 0,0008209 | 1,98 | 3,10E+08 | 3,36E+08 |
| 17,4 | | 0,0000537 | 0,0008178 | 1,96 | 2,84E+07 | 2,75E+07 |
| 49,8 | | 0,0000538 | 0,0008148 | 0,47 | 7,14E+08 | 7,05E+08 |
| 40,9 | | 0,0000540 | 0,0008118 | 0,02 | 9,94E+07 | 7,84E+07 |
| 21,5 | | 0,0000544 | 0,0008088 | 1,63 | 4,59E+07 | 4,56E+07 |
| 48,2 | | 0,0000548 | 0,0008059 | 1,82 | 8,43E+08 | 7,85E+08 |
| 35 | | 0,0000551 | 0,0008029 | 0,36 | 1,30E+08 | 1,37E+08 |
| 27,9 | | 0,0000564 | 0,0008000 | 0,06 | 5,64E+07 | 6,14E+07 |
| 85,3 | | 0,0000586 | 0,0007971 | 0,66 | 1,68E+10 | 1,79E+10 |
| 52,8 | | 0,0000591 | 0,0007942 | 0,77 | 6,45E+08 | 6,76E+08 |
| 32,9 | | 0,0000591 | 0,0007914 | 0,00 | 6,49E+09 | 4,32E+09 |
| 60,7 | | 0,0000597 | 0,0007885 | 1,40 | 3,29E+09 | 3,14E+09 |
| 34,9 | | 0,0000598 | 0,0007857 | 0,42 | 4,90E+07 | 4,79E+07 |
| 44,2 | | 0,0000606 | 0,0007829 | 0,28 | 8,93E+08 | 9,86E+08 |
| 25,9 | | 0,0000615 | 0,0007801 | 2,06 | 3,73E+08 | 4,01E+08 |
| 21,3 | | 0,0000615 | 0,0007774 | 0,05 | 3,94E+07 | 4,35E+07 |
| 10,8 | | 0,0000618 | 0,0007746 | 0,05 | 2,89E+07 | 2,92E+07 |
| 22,8 | | 0,0000622 | 0,0007719 | 8,29 | 7,28E+06 | 5,72E+06 |
| 21,2 | | 0,0000626 | 0,0007692 | 0,04 | 6,15E+07 | 5,39E+07 |
| 44 | | 0,0000627 | 0,0007666 | 0,03 | 1,74E+08 | 1,77E+08 |
| 64,4 | | 0,0000631 | 0,0007639 | 1,57 | 8,14E+08 | 8,35E+08 |
| 42,5 | | 0,0000632 | 0,0007612 | 0,36 | 1,99E+08 | 1,95E+08 |
| 55,4 | | 0,0000632 | 0,0007586 | 0,46 | 1,56E+09 | 1,74E+09 |
| 6,8 | | 0,0000638 | 0,0007560 | 0,26 | 9,54E+06 | 1,05E+07 |
| 62,8 | | 0,0000645 | 0,0007534 | 1,79 | 8,38E+08 | 9,04E+08 |
| 19,4 | | 0,0000652 | 0,0010785 | 15,96 | 2,83E+06 | 2,18E+06 |
| 23,8 | | 0,0000659 | 0,0010748 | 0,44 | 7,42E+07 | 7,23E+07 |
| 22,6 | | 0,0000662 | 0,0010712 | 0,07 | 1,53E+08 | 1,72E+08 |
| 33,2 | | 0,0000665 | 0,0010676 | 0,18 | 1,40E+08 | 1,32E+08 |

| | | | | | | |
|---|---|---|---|---|---|---|
| 70,1 | | 0,0000667 | 0,0010640 | 2,69 | 4,00E+08 | 4,17E+08 |
| 45,9 | | 0,0000669 | 0,0010604 | 57,45 | 3,87E+06 | 2,13E+06 |
| 25 | | 0,0000675 | 0,0010569 | 1,54 | 1,19E+08 | 1,27E+08 |
| 34 | | 0,0000676 | 0,0010533 | 1,39 | 1,40E+09 | 1,43E+09 |
| 34,9 | | 0,0000684 | 0,0010498 | 0,07 | 9,24E+07 | 1,05E+08 |
| 24,1 | | 0,0000689 | 0,0010464 | 0,59 | 5,59E+07 | 5,53E+07 |
| 31,2 | | 0,0000694 | 0,0010429 | 0,01 | 3,64E+08 | 3,29E+08 |
| 22,9 | | 0,0000703 | 0,0010395 | 0,61 | 9,33E+07 | 8,66E+07 |
| 9,9 | | 0,0000708 | 0,0010361 | 0,27 | 4,81E+07 | 4,75E+07 |
| 24,9 | | 0,0000709 | 0,0010327 | 1,43 | 8,35E+07 | 8,62E+07 |
| 46,4 | | 0,0000709 | 0,0010293 | 1,36 | 9,51E+08 | 9,78E+08 |
| 34 | | 0,0000710 | 0,0010260 | 53,13 | 2,02E+06 | 3,63E+06 |
| 11,1 | | 0,0000716 | 0,0010227 | 0,34 | 2,99E+07 | 3,20E+07 |
| 52,4 | | 0,0000736 | 0,0010194 | 1,17 | 3,53E+08 | 3,48E+08 |
| 40,2 | | 0,0000741 | 0,0010161 | 0,32 | 5,43E+08 | 6,16E+08 |
| 43,1 | | 0,0000745 | 0,0010128 | 0,38 | 4,41E+08 | 3,78E+08 |
| 39,2 | | 0,0000763 | 0,0010096 | 0,00 | 1,42E+09 | 1,33E+09 |
| 65,8 | | 0,0000774 | 0,0010064 | 0,58 | 5,29E+09 | 5,33E+09 |
| 16,3 | | 0,0000779 | 0,0010032 | 0,42 | 3,93E+07 | 4,07E+07 |
| 34,5 | | 0,0000782 | 0,0010000 | 0,58 | 3,34E+08 | 3,25E+08 |
| 13,3 | | 0,0000782 | 0,0009968 | 0,31 | 3,08E+07 | 2,83E+07 |
| 61,7 | | 0,0000783 | 0,0009937 | 1,75 | 1,21E+09 | 1,17E+09 |
| 49,7 | | 0,0000795 | 0,0009906 | 1,44 | 1,64E+09 | 1,69E+09 |
| 52,9 | | 0,0000800 | 0,0009875 | 0,55 | 6,58E+08 | 6,34E+08 |
| 22,4 | | 0,0000809 | 0,0009844 | 1,77 | 7,09E+07 | 7,33E+07 |
| 58,8 | + | 0,0000814 | 0,0009814 | 0,00 | 5,71E+08 | 4,74E+08 |
| 66,4 | | 0,0000831 | 0,0009783 | 1,52 | 6,54E+08 | 6,87E+08 |
| 63,1 | | 0,0000835 | 0,0009753 | 2,48 | 1,52E+08 | 1,36E+08 |
| 30,6 | | 0,0000838 | 0,0009723 | 0,03 | 6,36E+07 | 7,00E+07 |
| 28,1 | | 0,0000838 | 0,0009693 | 0,19 | 1,90E+07 | 1,46E+07 |
| 30,6 | | 0,0000856 | 0,0009664 | 0,63 | 1,05E+08 | 1,11E+08 |
| 51,4 | | 0,0000866 | 0,0009634 | 2,64 | 3,95E+08 | 4,30E+08 |
| 17,2 | | 0,0000869 | 0,0009605 | 0,01 | 4,40E+08 | 5,29E+08 |
| 62,5 | | 0,0000871 | 0,0009576 | 0,59 | 8,43E+08 | 7,84E+08 |
| 50,4 | | 0,0000872 | 0,0009547 | 3,34 | 1,50E+09 | 1,31E+09 |
| 14,2 | | 0,0000873 | 0,0009518 | 0,59 | 1,21E+08 | 1,21E+08 |
| 33,2 | | 0,0000875 | 0,0009489 | 2,14 | 3,29E+08 | 3,72E+08 |
| 64,2 | | 0,0000875 | 0,0009461 | 238,58 | 1,76E+07 | 8,58E+06 |
| 20,4 | | 0,0000880 | 0,0009433 | 2,81 | 2,03E+07 | 2,09E+07 |
| 44,4 | | 0,0000889 | 0,0009405 | 0,46 | 1,58E+08 | 1,43E+08 |
| 32,8 | | 0,0000889 | 0,0009377 | 0,03 | 1,37E+08 | 1,38E+08 |
| 38 | | 0,0000891 | 0,0009349 | 0,62 | 7,15E+07 | 7,31E+07 |
| 25,8 | | 0,0000898 | 0,0009322 | 2,06 | 6,20E+09 | 6,38E+09 |
| 21,1 | | 0,0000901 | 0,0009294 | 0,55 | 6,17E+07 | 5,78E+07 |
| 32,6 | | 0,0000909 | 0,0009267 | 2,38 | 3,66E+08 | 3,46E+08 |
| 15 | | 0,0000915 | 0,0009240 | 0,11 | 4,16E+07 | 4,15E+07 |
| 39,7 | | 0,0000921 | 0,0009213 | 0,63 | 5,12E+08 | 5,52E+08 |
| 29,9 | | 0,0000924 | 0,0009186 | 1,52 | 2,32E+08 | 2,49E+08 |
| 50,1 | | 0,0000924 | 0,0009159 | 0,45 | 5,44E+08 | 5,65E+08 |
| 63 | | 0,0000928 | 0,0011792 | 1,52 | 1,84E+09 | 1,75E+09 |

| | | | | | | |
|---|---|---|---|---|---|---|
| 20,9 | | 0,0000933 | 0,0011758 | 12,82 | 4,16E+06 | 5,41E+06 |
| 52,4 | | 0,0000934 | 0,0011724 | 1,46 | 1,46E+09 | 1,53E+09 |
| 19,3 | | 0,0000935 | 0,0011691 | 0,04 | 4,77E+07 | 4,31E+07 |
| 36,3 | | 0,0000936 | 0,0011657 | 2,03 | 5,39E+07 | 4,67E+07 |
| 29,1 | | 0,0000936 | 0,0011624 | 0,58 | 4,23E+08 | 4,39E+08 |
| 47,8 | | 0,0000940 | 0,0011591 | 0,00 | 4,58E+08 | 1,48E+08 |
| 49,8 | | 0,0000942 | 0,0011558 | 1,29 | 3,84E+08 | 4,01E+08 |
| 64 | | 0,0000948 | 0,0011525 | 0,01 | 1,42E+08 | 1,14E+08 |
| 19,2 | | 0,0000950 | 0,0011493 | 31,47 | 6,19E+06 | 3,55E+06 |
| 38,8 | | 0,0000954 | 0,0011461 | 0,41 | 2,68E+08 | 3,10E+08 |
| 60,2 | | 0,0000955 | 0,0011429 | 2,01 | 8,73E+08 | 8,13E+08 |
| 10,6 | | 0,0000956 | 0,0011397 | 0,46 | 3,42E+07 | 3,97E+07 |
| 39,6 | | 0,0000958 | 0,0011365 | 0,01 | 2,93E+08 | 3,34E+08 |
| 33,3 | | 0,0000960 | 0,0011333 | 0,49 | 5,34E+08 | 5,76E+08 |
| 26,2 | | 0,0000964 | 0,0011302 | 0,02 | 1,12E+08 | 9,99E+07 |
| 61,3 | | 0,0000965 | 0,0011271 | 0,46 | 1,63E+08 | 1,76E+08 |
| 16 | | 0,0000967 | 0,0011240 | 22,72 | 2,96E+06 | 3,12E+06 |
| 59,9 | | 0,0000978 | 0,0011209 | 1,56 | 8,64E+08 | 8,94E+08 |
| 59,3 | | 0,0000979 | 0,0011178 | 322,49 | 1,71E+06 | 7,50E+05 |
| 35,7 | | 0,0000980 | 0,0011148 | 0,50 | 1,39E+08 | 1,35E+08 |
| 58,5 | | 0,0000985 | 0,0011117 | 0,28 | 3,59E+08 | 2,84E+08 |
| 54,1 | | 0,0000986 | 0,0011087 | 0,53 | 4,25E+09 | 4,17E+09 |
| 16,3 | + | 0,0000990 | 0,0011057 | 11,89 | 1,11E+07 | 8,25E+06 |
| 16,2 | | 0,0000991 | 0,0011027 | 0,66 | 1,34E+08 | 1,28E+08 |
| 52,5 | | 0,0001001 | 0,0010997 | 0,33 | 1,48E+09 | 1,56E+09 |
| 19,8 | | 0,0001016 | 0,0010968 | 0,08 | 2,45E+08 | 1,99E+08 |
| 31,2 | | 0,0001017 | 0,0010938 | 0,40 | 3,47E+08 | 3,92E+08 |
| 20,7 | | 0,0001019 | 0,0010909 | 0,44 | 8,05E+07 | 8,11E+07 |
| 42,8 | | 0,0001033 | 0,0010880 | 0,35 | 2,11E+08 | 2,04E+08 |
| 42 | | 0,0001039 | 0,0010851 | 1,95 | 2,43E+08 | 2,74E+08 |
| 47,9 | | 0,0001052 | 0,0010822 | 0,01 | 6,02E+08 | 5,73E+08 |
| 49,7 | | 0,0001061 | 0,0010794 | 1,35 | 1,58E+09 | 1,54E+09 |
| 33,5 | | 0,0001067 | 0,0010765 | 0,52 | 4,15E+08 | 3,89E+08 |
| 26,5 | | 0,0001080 | 0,0010737 | 3,25 | 2,55E+07 | 2,30E+07 |
| 38,1 | | 0,0001083 | 0,0010709 | 0,75 | 7,43E+08 | 7,31E+08 |
| 45,3 | | 0,0001097 | 0,0013927 | 0,66 | 1,40E+08 | 1,49E+08 |
| 32,6 | | 0,0001108 | 0,0013890 | 0,02 | 9,14E+07 | 1,17E+08 |
| 67,5 | | 0,0001113 | 0,0013854 | 2,02 | 1,32E+10 | 1,38E+10 |
| 54,8 | | 0,0001115 | 0,0013818 | 1,43 | 4,98E+08 | 4,89E+08 |
| 62,8 | | 0,0001116 | 0,0013782 | 11,26 | 4,72E+07 | 3,93E+07 |
| 33,1 | | 0,0001116 | 0,0013747 | 0,56 | 4,97E+07 | 5,38E+07 |
| 49,7 | | 0,0001119 | 0,0013711 | 1,60 | 9,58E+08 | 9,78E+08 |
| 44,5 | | 0,0001124 | 0,0013676 | 1,59 | 7,45E+08 | 7,10E+08 |
| 43,5 | | 0,0001129 | 0,0013641 | 1,50 | 1,98E+08 | 1,98E+08 |
| 38,8 | | 0,0001157 | 0,0013606 | 0,64 | 4,08E+08 | 3,80E+08 |
| 54,9 | | 0,0001165 | 0,0013571 | 7,43 | 3,93E+07 | 2,54E+07 |
| 32,5 | | 0,0001167 | 0,0013537 | 0,20 | 1,66E+07 | 1,83E+07 |
| 21,1 | | 0,0001169 | 0,0013503 | 0,46 | 2,46E+07 | 2,52E+07 |
| 29,6 | | 0,0001172 | 0,0013468 | 3,23 | 6,05E+07 | 5,36E+07 |
| 53,2 | | 0,0001179 | 0,0013434 | 0,72 | 6,25E+08 | 6,14E+08 |

| | | | | | | |
|---|---|---|---|---|---|---|
| 10,7 | | 0,0001190 | 0,0013401 | 0,64 | 4,47E+07 | 4,25E+07 |
| 27,3 | | 0,0001192 | 0,0013367 | 2,49 | 2,52E+07 | 2,81E+07 |
| 49,6 | | 0,0001195 | 0,0013333 | 0,31 | 3,04E+08 | 3,13E+08 |
| 30,9 | | 0,0001198 | 0,0013300 | 1,60 | 1,57E+08 | 1,71E+08 |
| 40,9 | | 0,0001198 | 0,0013267 | 0,71 | 9,32E+08 | 8,80E+08 |
| 33,7 | | 0,0001198 | 0,0013234 | 0,58 | 2,90E+08 | 3,13E+08 |
| 45,6 | | 0,0001211 | 0,0013201 | 0,68 | 1,39E+09 | 1,49E+09 |
| 38,9 | | 0,0001213 | 0,0013168 | 1,94 | 8,29E+07 | 9,56E+07 |
| 47,5 | + | 0,0001220 | 0,0013136 | 3,32 | 1,34E+08 | 1,63E+08 |
| 21,6 | | 0,0001225 | 0,0013103 | 0,32 | 2,62E+07 | 2,51E+07 |
| 42,4 | | 0,0001234 | 0,0013071 | 0,70 | 1,42E+09 | 1,38E+09 |
| 64,4 | | 0,0001235 | 0,0013039 | 0,61 | 8,55E+08 | 9,23E+08 |
| 44,1 | | 0,0001235 | 0,0013007 | 1,36 | 9,37E+08 | 9,38E+08 |
| 43,7 | | 0,0001237 | 0,0012976 | 0,61 | 6,06E+08 | 6,05E+08 |
| 49,8 | | 0,0001239 | 0,0012944 | 1,25 | 1,88E+09 | 1,96E+09 |
| 76,8 | | 0,0001247 | 0,0012913 | 0,55 | 1,15E+10 | 1,11E+10 |
| 36,5 | | 0,0001256 | 0,0012881 | 0,30 | 5,13E+08 | 5,35E+08 |
| 16,6 | | 0,0001263 | 0,0012850 | 0,44 | 2,00E+08 | 1,67E+08 |
| 10,6 | | 0,0001266 | 0,0012819 | 0,07 | 2,68E+07 | 2,34E+07 |
| 33 | | 0,0001269 | 0,0012788 | 1,83 | 9,43E+07 | 9,74E+07 |
| 80,7 | | 0,0001273 | 0,0012758 | 1,65 | 4,39E+09 | 4,08E+09 |
| 52,4 | | 0,0001274 | 0,0012727 | 0,51 | 2,85E+08 | 2,55E+08 |
| 36,9 | | 0,0001275 | 0,0012697 | 0,85 | 2,59E+08 | 2,51E+08 |
| 53,5 | | 0,0001279 | 0,0012667 | 0,54 | 6,08E+08 | 5,43E+08 |
| 11,5 | | 0,0001282 | 0,0012637 | 0,47 | 1,05E+07 | 1,07E+07 |
| 44,4 | | 0,0001283 | 0,0012607 | 0,40 | 2,70E+08 | 2,67E+08 |
| 43,2 | + | 0,0001285 | 0,0012577 | 0,44 | 2,93E+08 | 2,55E+08 |
| 45,4 | | 0,0001285 | 0,0012547 | 0,47 | 1,10E+09 | 1,21E+09 |
| 47,9 | | 0,0001306 | 0,0012518 | 0,43 | 2,23E+08 | 1,99E+08 |
| 24 | | 0,0001308 | 0,0012488 | 2,99 | 8,39E+07 | 9,55E+07 |
| 22,9 | | 0,0001310 | 0,0012459 | 0,45 | 3,30E+08 | 3,37E+08 |
| 39 | | 0,0001312 | 0,0012430 | 0,72 | 1,54E+08 | 1,51E+08 |
| 18,5 | | 0,0001334 | 0,0012401 | 1,96 | 5,36E+07 | 5,88E+07 |
| 76,6 | | 0,0001336 | 0,0012372 | 0,59 | 1,45E+10 | 1,40E+10 |
| 29,4 | | 0,0001338 | 0,0012343 | 0,67 | 1,32E+08 | 1,31E+08 |
| 66,3 | | 0,0001346 | 0,0012315 | 0,54 | 5,96E+08 | 6,17E+08 |
| 14,3 | | 0,0001372 | 0,0014781 | 0,23 | 8,11E+07 | 5,91E+07 |
| 16 | | 0,0001397 | 0,0014747 | 0,08 | 5,16E+07 | 5,90E+07 |
| 46,8 | | 0,0001409 | 0,0014713 | 1,71 | 2,72E+08 | 2,81E+08 |
| 33 | | 0,0001409 | 0,0014679 | 0,74 | 2,78E+08 | 2,76E+08 |
| 55,1 | | 0,0001413 | 0,0014645 | 1,49 | 2,87E+08 | 3,13E+08 |
| 14,2 | | 0,0001422 | 0,0014612 | 5,17 | 1,13E+07 | 9,07E+06 |
| 56,8 | | 0,0001426 | 0,0014579 | 1,29 | 8,02E+08 | 7,88E+08 |
| 12,3 | | 0,0001427 | 0,0014545 | 0,06 | 1,43E+08 | 8,49E+07 |
| 35,9 | | 0,0001430 | 0,0014512 | 0,03 | 7,70E+07 | 8,54E+07 |
| 42,1 | | 0,0001441 | 0,0014480 | 1,36 | 7,29E+08 | 6,91E+08 |
| 25,7 | | 0,0001442 | 0,0014447 | 0,43 | 3,03E+08 | 2,67E+08 |
| 60 | | 0,0001444 | 0,0014414 | 2,08 | 2,57E+08 | 2,19E+08 |
| 49,1 | | 0,0001454 | 0,0014382 | 2,07 | 7,95E+07 | 7,55E+07 |
| 40,2 | | 0,0001475 | 0,0014350 | 0,50 | 8,38E+08 | 8,04E+08 |

| | | | | | | |
|---|---|---|---|---|---|---|
| 47,1 | | 0,0001483 | 0,0014318 | 1,27 | 4,82E+08 | 4,62E+08 |
| 11 | | 0,0001486 | 0,0014286 | 2,11 | 1,26E+07 | 1,30E+07 |
| 52,3 | | 0,0001490 | 0,0014254 | 0,60 | 5,13E+08 | 5,54E+08 |
| 38,5 | | 0,0001494 | 0,0014222 | 0,43 | 3,27E+08 | 2,84E+08 |
| 36,7 | | 0,0001496 | 0,0014191 | 0,36 | 2,40E+08 | 2,77E+08 |
| 54,9 | | 0,0001497 | 0,0014159 | 1,34 | 3,63E+08 | 3,69E+08 |
| 19,8 | | 0,0001500 | 0,0014128 | 1,62 | 2,24E+07 | 2,22E+07 |
| 38,9 | | 0,0001501 | 0,0014097 | 1,43 | 3,67E+08 | 3,95E+08 |
| 69,3 | | 0,0001513 | 0,0014066 | 0,12 | 4,21E+08 | 4,92E+08 |
| 51,1 | | 0,0001515 | 0,0014035 | 0,63 | 1,86E+09 | 1,70E+09 |
| 4,5 | | 0,0001520 | 0,0014004 | 2,78 | 8,39E+06 | 7,52E+06 |
| 41,2 | | 0,0001523 | 0,0013974 | 0,07 | 4,60E+07 | 4,16E+07 |
| 33,2 | | 0,0001528 | 0,0013943 | 0,52 | 6,19E+07 | 6,92E+07 |
| 29,2 | | 0,0001529 | 0,0013913 | 1,72 | 8,09E+07 | 7,69E+07 |
| 72 | | 0,0001531 | 0,0013883 | 1,61 | 8,33E+08 | 9,12E+08 |
| 44,9 | | 0,0001541 | 0,0013853 | 33,29 | 5,86E+06 | 3,54E+06 |
| 87,9 | | 0,0001544 | 0,0013823 | 1,47 | 2,85E+09 | 2,85E+09 |
| 64,8 | | 0,0001547 | 0,0013793 | 0,60 | 1,76E+09 | 1,91E+09 |
| 47,8 | | 0,0001553 | 0,0013763 | 0,37 | 1,87E+09 | 1,65E+09 |
| 40,3 | | 0,0001554 | 0,0013734 | 1,99 | 2,68E+08 | 3,10E+08 |
| 38,9 | | 0,0001576 | 0,0013704 | 0,66 | 2,92E+08 | 2,69E+08 |
| 24,2 | | 0,0001584 | 0,0013675 | 0,42 | 3,83E+07 | 3,38E+07 |
| 41,7 | | 0,0001586 | 0,0013646 | 0,71 | 1,28E+09 | 1,23E+09 |
| 39,8 | | 0,0001592 | 0,0013617 | 2,24 | 8,24E+07 | 6,81E+07 |
| 50,1 | | 0,0001594 | 0,0013588 | 0,79 | 7,57E+08 | 7,57E+08 |
| 38,9 | | 0,0001604 | 0,0016186 | 1,84 | 1,45E+08 | 1,61E+08 |
| 20,6 | | 0,0001632 | 0,0016152 | 1,42 | 1,47E+08 | 1,38E+08 |
| 38,4 | | 0,0001636 | 0,0016118 | 2,03 | 1,58E+08 | 1,45E+08 |
| 27,3 | | 0,0001637 | 0,0016084 | 0,45 | 4,01E+07 | 4,22E+07 |
| 20,4 | | 0,0001640 | 0,0016050 | 0,10 | 4,59E+07 | 4,09E+07 |
| 26,7 | | 0,0001642 | 0,0016017 | 0,15 | 2,36E+08 | 2,31E+08 |
| 23,7 | | 0,0001651 | 0,0015983 | 0,59 | 2,53E+08 | 2,72E+08 |
| 56,1 | | 0,0001652 | 0,0015950 | 1,85 | 3,05E+08 | 2,98E+08 |
| 14,9 | | 0,0001671 | 0,0015917 | 0,04 | 3,94E+07 | 3,97E+07 |
| 20,3 | | 0,0001681 | 0,0015884 | 1,67 | 1,44E+08 | 1,40E+08 |
| 33,1 | | 0,0001684 | 0,0015851 | 2,95 | 5,03E+07 | 4,04E+07 |
| 33,3 | | 0,0001689 | 0,0015818 | 0,57 | 8,97E+07 | 8,46E+07 |
| 9,4 | | 0,0001693 | 0,0015785 | 3,98 | 3,15E+06 | 2,89E+06 |
| 35,4 | | 0,0001694 | 0,0015753 | 2,24 | 5,36E+07 | 5,26E+07 |
| 32,8 | | 0,0001695 | 0,0015720 | 1,76 | 1,07E+07 | 9,66E+06 |
| 46,3 | | 0,0001697 | 0,0015688 | 0,01 | 2,22E+08 | 2,52E+08 |
| 7,5 | | 0,0001699 | 0,0015656 | 0,02 | 1,24E+08 | 7,72E+07 |
| 36,1 | | 0,0001708 | 0,0015624 | 2,34 | 1,11E+08 | 1,22E+08 |
| 30,6 | | 0,0001715 | 0,0015592 | 0,71 | 8,61E+07 | 9,34E+07 |
| 45,2 | | 0,0001737 | 0,0015560 | 0,42 | 1,52E+08 | 1,55E+08 |
| 19,7 | | 0,0001738 | 0,0015528 | 0,02 | 2,08E+08 | 2,00E+08 |
| 30,9 | | 0,0001744 | 0,0015497 | 2,01 | 6,51E+07 | 6,19E+07 |
| 42,4 | | 0,0001746 | 0,0015466 | 1,48 | 1,12E+08 | 1,09E+08 |
| 38 | | 0,0001751 | 0,0015434 | 2,34 | 2,52E+08 | 2,57E+08 |
| 52,8 | | 0,0001758 | 0,0015403 | 1,43 | 8,52E+08 | 8,82E+08 |

| | | | | | | |
|---|---|---|---|---|---|---|
| 18,6 | | 0,0001761 | 0,0015372 | 0,62 | 7,06E+07 | 7,93E+07 |
| 36,9 | | 0,0001764 | 0,0015341 | 1,63 | 4,53E+07 | 4,20E+07 |
| 66,6 | | 0,0001771 | 0,0015311 | 0,62 | 6,80E+09 | 6,56E+09 |
| 36,2 | | 0,0001772 | 0,0015280 | 0,71 | 2,08E+08 | 2,14E+08 |
| 53,5 | | 0,0001778 | 0,0015250 | 1,82 | 2,20E+08 | 2,42E+08 |
| 25,8 | | 0,0001790 | 0,0015219 | 0,06 | 2,63E+07 | 3,51E+07 |
| 52 | | 0,0001805 | 0,0015189 | 1,72 | 1,14E+09 | 1,18E+09 |
| 59,7 | | 0,0001808 | 0,0015159 | 0,36 | 1,82E+09 | 1,77E+09 |
| 40,1 | | 0,0001810 | 0,0015129 | 2,30 | 5,03E+07 | 4,49E+07 |
| 45,8 | | 0,0001813 | 0,0015099 | 0,47 | 3,52E+08 | 3,86E+08 |
| 47 | | 0,0001814 | 0,0015069 | 0,64 | 5,11E+08 | 4,59E+08 |
| 38,1 | | 0,0001826 | 0,0015039 | 0,65 | 2,60E+08 | 2,39E+08 |
| 29 | | 0,0001833 | 0,0015010 | 1,79 | 1,10E+08 | 1,03E+08 |
| 60,5 | | 0,0001838 | 0,0014980 | 1,26 | 4,47E+08 | 4,71E+08 |
| 29,1 | | 0,0001845 | 0,0014951 | 0,51 | 1,30E+08 | 1,19E+08 |
| 32,8 | | 0,0001851 | 0,0014922 | 1,86 | 2,09E+08 | 2,11E+08 |
| 49,2 | | 0,0001864 | 0,0014893 | 1,62 | 4,03E+08 | 4,45E+08 |
| 29,8 | | 0,0001864 | 0,0014864 | 0,01 | 1,54E+08 | 1,53E+08 |
| 22 | | 0,0001869 | 0,0014835 | 0,03 | 9,22E+07 | 8,62E+07 |
| 41,7 | | 0,0001872 | 0,0014806 | 0,53 | 4,69E+08 | 4,59E+08 |
| 24,9 | | 0,0001882 | 0,0014778 | 2,55 | 3,10E+07 | 3,06E+07 |
| 24,5 | | 0,0001884 | 0,0014749 | 0,55 | 1,59E+08 | 1,47E+08 |
| 12,5 | | 0,0001892 | 0,0014721 | 0,28 | 2,66E+07 | 3,42E+07 |
| 19,8 | | 0,0001919 | 0,0014692 | 0,01 | 5,29E+07 | 5,87E+07 |
| 15,6 | | 0,0001939 | 0,0014664 | 6,96 | 2,71E+06 | 2,70E+06 |
| 59,1 | | 0,0001940 | 0,0014636 | 0,63 | 1,59E+10 | 1,59E+10 |
| 29,3 | | 0,0001942 | 0,0014608 | 0,38 | 2,99E+08 | 3,30E+08 |
| 35,2 | | 0,0001946 | 0,0014580 | 0,62 | 5,64E+08 | 5,40E+08 |
| 34,3 | | 0,0001947 | 0,0014552 | 0,64 | 1,12E+08 | 1,10E+08 |
| 39,7 | | 0,0001954 | 0,0014525 | 0,42 | 3,05E+09 | 3,79E+09 |
| 45 | | 0,0001960 | 0,0014497 | 0,63 | 7,20E+08 | 7,76E+08 |
| 19,4 | | 0,0001976 | 0,0014470 | 1,33 | 8,52E+07 | 8,26E+07 |
| 43,6 | | 0,0001986 | 0,0014442 | 0,51 | 7,09E+08 | 7,28E+08 |
| 45,4 | | 0,0002002 | 0,0014415 | 0,61 | 4,18E+08 | 4,49E+08 |
| 34,6 | | 0,0002014 | 0,0014388 | 1,56 | 5,11E+07 | 5,43E+07 |
| 39,1 | | 0,0002015 | 0,0014361 | 0,66 | 2,86E+08 | 2,82E+08 |
| 32 | | 0,0002015 | 0,0014334 | 2,24 | 5,41E+07 | 4,52E+07 |
| 40,6 | | 0,0002021 | 0,0014307 | 0,56 | 7,25E+07 | 8,21E+07 |
| 28,3 | | 0,0002022 | 0,0014280 | 0,58 | 2,16E+08 | 2,10E+08 |
| 17,2 | | 0,0002024 | 0,0016269 | 1,57 | 8,14E+07 | 7,63E+07 |
| 38,2 | | 0,0002030 | 0,0016238 | 0,29 | 2,14E+08 | 2,03E+08 |
| 28,3 | | 0,0002032 | 0,0016208 | 0,05 | 6,23E+07 | 7,60E+07 |
| 29,7 | | 0,0002060 | 0,0016178 | 0,34 | 5,84E+07 | 6,60E+07 |
| 42 | | 0,0002071 | 0,0016148 | 0,02 | 1,23E+08 | 1,40E+08 |
| 25,7 | | 0,0002078 | 0,0016118 | 1,52 | 2,72E+08 | 2,92E+08 |
| 44,4 | | 0,0002080 | 0,0016089 | 0,78 | 9,92E+08 | 1,03E+09 |
| 36,8 | | 0,0002082 | 0,0016059 | 2,03 | 6,09E+08 | 5,52E+08 |
| 29 | | 0,0002083 | 0,0016029 | 0,36 | 2,81E+08 | 2,22E+08 |
| 45,2 | | 0,0002085 | 0,0016000 | 0,44 | 1,09E+09 | 1,05E+09 |
| 19,8 | | 0,0002091 | 0,0015971 | 1,57 | 1,39E+08 | 1,48E+08 |

| | | | | | | |
|---|---|---|---|---|---|---|
| 52,4 | | 0,0002115 | 0,0015941 | 1,28 | 1,13E+09 | 1,09E+09 |
| 34,7 | | 0,0002117 | 0,0015912 | 1,46 | 1,32E+08 | 1,23E+08 |
| 20,1 | | 0,0002119 | 0,0015883 | 0,02 | 6,25E+07 | 6,51E+07 |
| 55,2 | | 0,0002124 | 0,0015855 | 1,24 | 2,50E+09 | 2,57E+09 |
| 61,9 | | 0,0002151 | 0,0015826 | 0,64 | 8,74E+08 | 8,69E+08 |
| 52,7 | | 0,0002154 | 0,0015797 | 0,00 | 6,65E+08 | 6,92E+08 |
| 46,9 | | 0,0002164 | 0,0015769 | 0,26 | 2,72E+09 | 2,87E+09 |
| 29,5 | | 0,0002179 | 0,0015740 | 0,08 | 1,27E+08 | 1,34E+08 |
| 31,7 | | 0,0002181 | 0,0015712 | 1,48 | 1,12E+08 | 1,06E+08 |
| 32,8 | | 0,0002190 | 0,0015683 | 2,14 | 1,07E+08 | 9,87E+07 |
| 25,2 | | 0,0002209 | 0,0015655 | 0,55 | 7,22E+07 | 7,28E+07 |
| 17,7 | | 0,0002217 | 0,0015627 | 2,20 | 5,13E+07 | 4,46E+07 |
| 21,5 | | 0,0002222 | 0,0015599 | 76,91 | 3,48E+06 | 1,50E+06 |
| 13,7 | | 0,0002229 | 0,0015571 | 0,07 | 4,19E+07 | 4,37E+07 |
| 48,3 | | 0,0002237 | 0,0015544 | 0,74 | 2,12E+09 | 2,08E+09 |
| 46 | | 0,0002242 | 0,0015516 | 1,18 | 6,25E+08 | 6,32E+08 |
| 68,8 | | 0,0002246 | 0,0015488 | 1,42 | 2,19E+09 | 2,22E+09 |
| 29,7 | | 0,0002251 | 0,0015461 | 0,37 | 8,99E+07 | 9,60E+07 |
| 27,9 | | 0,0002260 | 0,0015434 | 0,66 | 5,92E+07 | 5,61E+07 |
| 44,7 | | 0,0002270 | 0,0015406 | 0,65 | 7,42E+08 | 7,30E+08 |
| 15,6 | | 0,0002272 | 0,0015379 | 2,15 | 2,99E+07 | 2,48E+07 |
| 43,6 | | 0,0002283 | 0,0015352 | 7,59 | 4,13E+06 | 2,43E+06 |
| 13,6 | | 0,0002304 | 0,0015325 | 21,25 | 1,25E+07 | 6,08E+06 |
| 33,5 | | 0,0002308 | 0,0015298 | 1,50 | 1,33E+08 | 1,19E+08 |
| 12,1 | | 0,0002311 | 0,0015271 | 11,65 | 3,73E+06 | 2,28E+06 |
| 29,5 | | 0,0002350 | 0,0015245 | 0,50 | 1,99E+08 | 1,95E+08 |
| 45 | | 0,0002353 | 0,0015218 | 2,26 | 1,78E+08 | 1,99E+08 |
| 26,2 | | 0,0002365 | 0,0015192 | 0,67 | 7,41E+08 | 6,83E+08 |
| 31,1 | | 0,0002365 | 0,0015165 | 0,60 | 1,56E+08 | 1,39E+08 |
| 27,3 | | 0,0002370 | 0,0015139 | 0,02 | 1,54E+08 | 4,16E+08 |
| 27,9 | | 0,0002372 | 0,0015113 | 1,49 | 1,12E+08 | 1,02E+08 |
| 54,8 | + | 0,0002380 | 0,0015087 | 52,17 | 6,13E+06 | 6,38E+06 |
| 68 | | 0,0002401 | 0,0015060 | 1,52 | 1,04E+09 | 9,47E+08 |
| 25,1 | | 0,0002406 | 0,0015034 | 0,07 | 4,00E+07 | 7,29E+07 |
| 60,6 | | 0,0002423 | 0,0015009 | 2,62 | 6,00E+07 | 5,70E+07 |
| 54,6 | | 0,0002434 | 0,0014983 | 0,77 | 2,64E+09 | 2,67E+09 |
| 60,4 | | 0,0002439 | 0,0014957 | 3,29 | 2,74E+08 | 2,66E+08 |
| 56 | | 0,0002446 | 0,0014932 | 0,72 | 8,56E+08 | 8,91E+08 |
| 57,1 | | 0,0002476 | 0,0014906 | 0,73 | 6,24E+08 | 6,65E+08 |
| 20,1 | | 0,0002495 | 0,0014881 | 9,24 | 1,68E+07 | 1,80E+07 |
| 20,8 | | 0,0002499 | 0,0014855 | 1,77 | 1,42E+08 | 1,38E+08 |
| 21,3 | | 0,0002500 | 0,0014830 | 0,49 | 6,47E+07 | 7,05E+07 |
| 60,6 | | 0,0002521 | 0,0014805 | 2,11 | 1,51E+08 | 1,46E+08 |
| 36,8 | | 0,0002526 | 0,0014780 | 0,47 | 9,69E+08 | 1,04E+09 |
| 38 | | 0,0002528 | 0,0014755 | 0,34 | 3,09E+07 | 3,57E+07 |
| 31,7 | | 0,0002544 | 0,0014730 | 1,73 | 8,02E+07 | 7,33E+07 |
| 26,9 | | 0,0002547 | 0,0014705 | 31,18 | 4,32E+06 | 4,45E+06 |
| 46,4 | | 0,0002558 | 0,0014680 | 1,26 | 4,93E+08 | 5,22E+08 |
| 23 | | 0,0002567 | 0,0014655 | 0,61 | 6,53E+07 | 6,89E+07 |
| 76,7 | | 0,0002568 | 0,0014631 | 0,91 | 2,01E+09 | 2,01E+09 |

| | | | | | | |
|---|---|---|---|---|---|---|
| 28,9 | | 0,0002574 | 0,0014606 | 0,11 | 1,51E+08 | 1,44E+08 |
| 42,3 | | 0,0002580 | 0,0014582 | 0,65 | 3,22E+09 | 2,89E+09 |
| 14,1 | + | 0,0002582 | 0,0014558 | 16,37 | 5,27E+06 | 9,30E+06 |
| 45,4 | | 0,0002587 | 0,0014533 | 0,81 | 1,77E+08 | 1,68E+08 |
| 26 | | 0,0002606 | 0,0014509 | 1,97 | 3,20E+07 | 3,61E+07 |
| 36,6 | | 0,0002614 | 0,0014485 | 0,54 | 2,94E+08 | 2,63E+08 |
| 44 | | 0,0002615 | 0,0014461 | 3,36 | 1,73E+08 | 2,21E+08 |
| 32,8 | | 0,0002629 | 0,0014437 | 1,43 | 5,02E+07 | 4,82E+07 |
| 15,2 | | 0,0002634 | 0,0014413 | 1,75 | 5,78E+07 | 5,49E+07 |
| 28,5 | | 0,0002655 | 0,0014389 | 14,37 | 4,25E+06 | 3,35E+06 |
| 43,5 | | 0,0002662 | 0,0014366 | 0,25 | 2,53E+08 | 3,63E+08 |
| 38,9 | | 0,0002666 | 0,0014342 | 1,48 | 1,52E+08 | 1,49E+08 |
| 33,9 | + | 0,0002674 | 0,0014319 | 7,11 | 2,63E+08 | 2,84E+08 |
| 31,9 | | 0,0002690 | 0,0014295 | 0,02 | 2,29E+08 | 1,86E+08 |
| 14,5 | | 0,0002691 | 0,0014272 | 1,72 | 1,99E+07 | 1,86E+07 |
| 42,3 | | 0,0002700 | 0,0015817 | 0,10 | 5,32E+07 | 6,14E+07 |
| 65,8 | | 0,0002719 | 0,0015791 | 0,68 | 8,52E+07 | 8,62E+07 |
| 16,3 | | 0,0002724 | 0,0015765 | 2,54 | 1,26E+07 | 1,12E+07 |
| 34,9 | | 0,0002725 | 0,0015740 | 0,53 | 1,34E+09 | 1,46E+09 |
| 44,2 | | 0,0002730 | 0,0015714 | 1,98 | 8,12E+08 | 6,68E+08 |
| 26,9 | | 0,0002744 | 0,0015689 | 0,21 | 1,15E+08 | 9,08E+07 |
| 50,4 | | 0,0002767 | 0,0015663 | 2,63 | 2,58E+08 | 2,01E+08 |
| 18,7 | | 0,0002805 | 0,0017189 | 0,43 | 4,05E+07 | 3,60E+07 |
| 29,5 | | 0,0002830 | 0,0017161 | 0,64 | 2,20E+08 | 2,32E+08 |
| 70 | | 0,0002831 | 0,0017134 | 1,63 | 1,06E+09 | 9,74E+08 |
| 24,3 | | 0,0002862 | 0,0017106 | 2,96 | 1,28E+07 | 9,55E+06 |
| 47,8 | | 0,0002864 | 0,0017079 | 0,67 | 1,28E+09 | 1,31E+09 |
| 34,1 | | 0,0002865 | 0,0017051 | 0,49 | 1,40E+08 | 1,51E+08 |
| 29,3 | | 0,0002869 | 0,0017024 | 0,78 | 1,06E+09 | 1,07E+09 |
| 7,4 | | 0,0002878 | 0,0016997 | 0,16 | 2,60E+07 | 3,25E+07 |
| 16,1 | | 0,0002880 | 0,0016970 | 8,40 | 1,39E+07 | 1,03E+07 |
| 38,1 | | 0,0002920 | 0,0016943 | 0,78 | 1,95E+08 | 1,97E+08 |
| 65,9 | | 0,0002924 | 0,0016916 | 0,77 | 6,56E+09 | 6,82E+09 |
| 20,1 | | 0,0002949 | 0,0016889 | 0,39 | 2,82E+08 | 3,57E+08 |
| 40,1 | | 0,0002954 | 0,0016862 | 2,09 | 9,96E+07 | 1,13E+08 |
| 24,4 | | 0,0002967 | 0,0016835 | 0,16 | 4,16E+07 | 3,13E+07 |
| 39,8 | | 0,0002968 | 0,0016809 | 0,31 | 4,28E+08 | 3,81E+08 |
| 30,8 | | 0,0002983 | 0,0016782 | 1,60 | 3,58E+07 | 4,04E+07 |
| 39,3 | | 0,0002998 | 0,0016756 | 0,59 | 3,07E+09 | 2,96E+09 |
| 42,5 | | 0,0003013 | 0,0016730 | 0,60 | 1,82E+08 | 1,62E+08 |
| 27,2 | | 0,0003017 | 0,0016703 | 0,10 | 5,93E+07 | 3,70E+07 |
| 5,6 | | 0,0003031 | 0,0016677 | 6,53 | 3,20E+06 | 3,32E+06 |
| 23,1 | | 0,0003056 | 0,0016651 | 0,06 | 4,17E+07 | 3,68E+07 |
| 25,8 | | 0,0003065 | 0,0016625 | 1,34 | 1,27E+08 | 1,34E+08 |
| 31,5 | | 0,0003076 | 0,0016599 | 0,58 | 5,16E+08 | 5,51E+08 |
| 34,5 | | 0,0003094 | 0,0016573 | 0,53 | 1,51E+08 | 1,66E+08 |
| 44,1 | | 0,0003099 | 0,0016547 | 0,45 | 4,03E+08 | 4,12E+08 |
| 34,8 | | 0,0003122 | 0,0016522 | 0,69 | 1,59E+08 | 1,54E+08 |
| 25,9 | | 0,0003123 | 0,0016496 | 1,29 | 4,30E+08 | 4,46E+08 |
| 40,9 | | 0,0003124 | 0,0016471 | 0,38 | 2,93E+08 | 2,46E+08 |

| | | | | | | |
|---|---|---|---|---|---|---|
| 11,6 | | 0,0003133 | 0,0016445 | 0,51 | 4,57E+07 | 5,17E+07 |
| 44,3 | | 0,0003140 | 0,0016420 | 0,00 | 1,91E+09 | 1,94E+09 |
| 63,1 | | 0,0003142 | 0,0016394 | 0,53 | 1,47E+09 | 1,39E+09 |
| 31,1 | | 0,0003159 | 0,0016369 | 1,96 | 2,98E+07 | 2,83E+07 |
| 33,1 | | 0,0003160 | 0,0016344 | 1,30 | 3,45E+08 | 3,45E+08 |
| 44,4 | | 0,0003161 | 0,0016319 | 2,04 | 1,30E+09 | 1,13E+09 |
| 0,9 | | 0,0003163 | 0,0016294 | 0,26 | 2,41E+08 | 2,65E+08 |
| 25,7 | | 0,0003173 | 0,0016269 | 0,83 | 7,12E+08 | 7,22E+08 |
| 60,6 | + | 0,0003190 | 0,0016244 | 0,00 | 6,00E+09 | 4,97E+09 |
| 48,4 | | 0,0003194 | 0,0016220 | 0,66 | 1,19E+09 | 1,29E+09 |
| 43,9 | | 0,0003203 | 0,0016195 | 2,09 | 5,15E+07 | 4,53E+07 |
| 22,9 | | 0,0003218 | 0,0016170 | 0,03 | 1,00E+08 | 3,69E+07 |
| 13,1 | | 0,0003218 | 0,0016146 | 0,28 | 3,21E+07 | 3,58E+07 |
| 48,1 | | 0,0003271 | 0,0016121 | 0,68 | 1,18E+09 | 1,08E+09 |
| 23,5 | | 0,0003304 | 0,0016097 | 0,09 | 4,14E+07 | 6,54E+07 |
| 55,2 | | 0,0003305 | 0,0016073 | 1,38 | 1,18E+08 | 1,14E+08 |
| 9,4 | | 0,0003314 | 0,0016048 | 2,22 | 2,22E+07 | 2,58E+07 |
| 37,1 | | 0,0003322 | 0,0016024 | 1,75 | 1,74E+08 | 1,73E+08 |
| 50,2 | | 0,0003330 | 0,0016000 | 1,74 | 5,25E+07 | 5,49E+07 |
| 31,9 | | 0,0003332 | 0,0015976 | 1,81 | 2,64E+07 | 2,56E+07 |
| 18,9 | | 0,0003340 | 0,0015952 | 1,10 | 6,78E+07 | 6,76E+07 |
| 39,1 | | 0,0003344 | 0,0015928 | 0,79 | 1,89E+09 | 1,84E+09 |
| 37,2 | | 0,0003348 | 0,0015904 | 0,55 | 3,48E+08 | 2,97E+08 |
| 49,6 | | 0,0003353 | 0,0015881 | 2,03 | 1,33E+08 | 1,15E+08 |
| 4,5 | | 0,0003358 | 0,0015857 | 0,33 | 1,06E+07 | 1,46E+07 |
| 38,7 | | 0,0003369 | 0,0015833 | 0,67 | 2,10E+08 | 2,32E+08 |
| 37,3 | | 0,0003371 | 0,0015810 | 0,53 | 1,23E+08 | 1,22E+08 |
| 60 | | 0,0003371 | 0,0015786 | 1,25 | 3,30E+09 | 3,46E+09 |
| 19,5 | | 0,0003390 | 0,0015763 | 0,51 | 9,92E+07 | 1,12E+08 |
| 21,6 | | 0,0003393 | 0,0015740 | 0,06 | 7,34E+07 | 7,05E+07 |
| 22,2 | | 0,0003429 | 0,0015716 | 2,17 | 2,61E+07 | 3,07E+07 |
| 52 | | 0,0003440 | 0,0015693 | 1,46 | 4,87E+08 | 5,15E+08 |
| 52,1 | | 0,0003455 | 0,0015670 | 1,69 | 1,56E+08 | 1,38E+08 |
| 34 | | 0,0003465 | 0,0015647 | 0,46 | 5,14E+07 | 5,05E+07 |
| 65,6 | | 0,0003477 | 0,0015624 | 1,47 | 2,78E+09 | 2,50E+09 |
| 33 | | 0,0003494 | 0,0015601 | 0,05 | 3,30E+07 | 3,65E+07 |
| 68,3 | | 0,0003495 | 0,0015578 | 1,34 | 2,00E+09 | 2,16E+09 |
| 78,5 | | 0,0003497 | 0,0015556 | 2,55 | 1,74E+09 | 2,16E+09 |
| 64,9 | | 0,0003510 | 0,0015533 | 0,52 | 1,33E+08 | 1,11E+08 |
| 31,9 | | 0,0003520 | 0,0015510 | 0,57 | 2,62E+08 | 2,83E+08 |
| 33,6 | | 0,0003533 | 0,0015488 | 0,81 | 7,79E+08 | 7,54E+08 |
| 28,9 | | 0,0003563 | 0,0015465 | 0,44 | 2,18E+08 | 1,79E+08 |
| 26,1 | | 0,0003564 | 0,0015443 | 0,76 | 1,80E+08 | 1,74E+08 |
| 21,6 | | 0,0003570 | 0,0015420 | 2,09 | 7,32E+07 | 6,59E+07 |
| 31,5 | | 0,0003577 | 0,0015398 | 1,91 | 7,18E+07 | 6,19E+07 |
| 24,9 | | 0,0003585 | 0,0015376 | 1,32 | 6,92E+07 | 7,24E+07 |
| 9 | | 0,0003598 | 0,0015354 | 4,48 | 2,31E+06 | 2,11E+06 |
| 25,7 | | 0,0003605 | 0,0015331 | 0,48 | 9,88E+08 | 1,07E+09 |
| 58 | | 0,0003606 | 0,0015309 | 0,56 | 1,60E+09 | 1,42E+09 |
| 45,9 | | 0,0003619 | 0,0015287 | 2,05 | 7,46E+07 | 6,77E+07 |

| | | | | | | |
|---|---|---|---|---|---|---|
| 49,4 | | 0,0003629 | 0,0015265 | 0,70 | 1,26E+09 | 1,38E+09 |
| 17,1 | | 0,0003648 | 0,0015244 | 0,13 | 1,63E+07 | 2,91E+07 |
| 17,4 | | 0,0003659 | 0,0015222 | 5,88 | 1,63E+07 | 2,54E+07 |
| 48,8 | | 0,0003668 | 0,0015200 | 0,66 | 1,17E+09 | 1,14E+09 |
| 42,8 | | 0,0003673 | 0,0015178 | 0,79 | 4,32E+08 | 4,17E+08 |
| 44,3 | | 0,0003678 | 0,0015157 | 0,45 | 2,93E+08 | 2,45E+08 |
| 31,2 | | 0,0003689 | 0,0015135 | 1,60 | 1,64E+09 | 1,65E+09 |
| 51,3 | | 0,0003732 | 0,0015114 | 0,43 | 7,84E+08 | 8,67E+08 |
| 53,2 | | 0,0003736 | 0,0015092 | 0,73 | 5,05E+08 | 5,01E+08 |
| 22 | | 0,0003739 | 0,0015071 | 0,44 | 4,86E+07 | 5,89E+07 |
| 77,8 | | 0,0003748 | 0,0015050 | 1,56 | 3,05E+10 | 3,43E+10 |
| 26 | | 0,0003750 | 0,0015028 | 0,04 | 5,30E+07 | 4,53E+07 |
| 31 | | 0,0003772 | 0,0015007 | 0,58 | 7,38E+07 | 6,48E+07 |
| 33,1 | | 0,0003780 | 0,0014986 | 0,71 | 1,90E+08 | 1,80E+08 |
| 39,3 | | 0,0003791 | 0,0014965 | 1,46 | 1,42E+08 | 1,43E+08 |
| 44,5 | | 0,0003799 | 0,0014944 | 2,16 | 1,29E+08 | 1,24E+08 |
| 41,4 | | 0,0003805 | 0,0014923 | 0,56 | 2,15E+08 | 2,10E+08 |
| 84,2 | | 0,0003832 | 0,0014902 | 5,49 | 3,68E+09 | 3,02E+09 |
| 27,2 | | 0,0003835 | 0,0014881 | 0,05 | 4,71E+07 | 1,05E+08 |
| 29,6 | | 0,0003837 | 0,0014860 | 0,82 | 1,15E+08 | 1,14E+08 |
| 18,2 | | 0,0003842 | 0,0014840 | 1,32 | 7,85E+07 | 8,40E+07 |
| 4 | | 0,0003855 | 0,0014819 | 1,55 | 4,65E+06 | 4,73E+06 |
| 18,6 | | 0,0003856 | 0,0014798 | 2,06 | 7,16E+07 | 6,96E+07 |
| 30,8 | | 0,0003857 | 0,0014778 | 3,32 | 3,72E+07 | 2,96E+07 |
| 35,9 | | 0,0003857 | 0,0014757 | 0,36 | 5,82E+07 | 6,19E+07 |
| 26,6 | | 0,0003904 | 0,0014737 | 0,48 | 4,51E+08 | 4,24E+08 |
| 38,2 | | 0,0003926 | 0,0014716 | 1,87 | 2,92E+08 | 3,04E+08 |
| 29,8 | | 0,0003955 | 0,0014696 | 127,28 | 1,50E+06 | 6,73E+06 |
| 36,2 | | 0,0003975 | 0,0014676 | 1,32 | 5,23E+08 | 5,23E+08 |
| 49,3 | | 0,0004007 | 0,0014656 | 0,02 | 1,35E+08 | 1,43E+08 |
| 46,7 | | 0,0004011 | 0,0014635 | 0,02 | 1,49E+08 | 1,54E+08 |
| 23,8 | | 0,0004021 | 0,0014615 | 0,58 | 4,06E+08 | 4,72E+08 |
| 6,6 | | 0,0004040 | 0,0014595 | 0,21 | 7,82E+06 | 1,15E+07 |
| 42,1 | | 0,0004041 | 0,0014575 | 1,77 | 1,87E+08 | 2,20E+08 |
| 44,4 | | 0,0004078 | 0,0014555 | 0,62 | 2,36E+08 | 2,51E+08 |
| 24,6 | | 0,0004104 | 0,0014536 | 1,80 | 2,03E+08 | 1,80E+08 |
| 13,5 | | 0,0004114 | 0,0015989 | 0,54 | 5,07E+07 | 4,33E+07 |
| 39,3 | | 0,0004128 | 0,0015967 | 0,49 | 1,02E+08 | 9,14E+07 |
| 28,3 | | 0,0004136 | 0,0015946 | 1,49 | 1,26E+08 | 1,21E+08 |
| 15,8 | | 0,0004140 | 0,0015924 | 3,87 | 1,37E+07 | 1,04E+07 |
| 33,3 | | 0,0004141 | 0,0015902 | 0,62 | 1,55E+08 | 1,36E+08 |
| 27,7 | | 0,0004148 | 0,0015881 | 0,64 | 3,02E+09 | 2,64E+09 |
| 25,4 | | 0,0004150 | 0,0015859 | 0,30 | 6,79E+07 | 8,62E+07 |
| 37,6 | | 0,0004153 | 0,0015838 | 1,97 | 4,95E+08 | 4,76E+08 |
| 16,2 | | 0,0004167 | 0,0015816 | 1,90 | 7,30E+07 | 8,62E+07 |
| 13,2 | | 0,0004172 | 0,0015795 | 0,30 | 2,11E+07 | 2,58E+07 |
| 49,4 | | 0,0004173 | 0,0015774 | 1,42 | 2,21E+08 | 2,28E+08 |
| 41,1 | | 0,0004195 | 0,0015753 | 0,34 | 3,54E+08 | 2,70E+08 |
| 13,3 | | 0,0004224 | 0,0015732 | 0,51 | 3,28E+07 | 3,50E+07 |
| 50,3 | | 0,0004230 | 0,0015710 | 0,56 | 3,81E+08 | 3,24E+08 |

| | | | | | | |
|---|---|---|---|---|---|---|
| 39,5 | | 0,0004253 | 0,0015689 | 0,05 | 6,13E+08 | 7,93E+08 |
| 15,2 | | 0,0004280 | 0,0015668 | 0,29 | 1,44E+07 | 1,60E+07 |
| 27,7 | | 0,0004297 | 0,0015648 | 1,23 | 1,78E+08 | 1,86E+08 |
| 45,5 | | 0,0004306 | 0,0015627 | 1,65 | 1,44E+08 | 1,32E+08 |
| 48,9 | | 0,0004311 | 0,0015606 | 2,08 | 8,70E+07 | 8,27E+07 |
| 42,6 | | 0,0004342 | 0,0015585 | 0,42 | 1,57E+09 | 1,32E+09 |
| 69,8 | | 0,0004343 | 0,0015564 | 1,52 | 1,02E+10 | 9,01E+09 |
| 37 | | 0,0004343 | 0,0015544 | 0,65 | 1,14E+08 | 1,08E+08 |
| 17,4 | | 0,0004348 | 0,0015523 | 1,55 | 1,14E+08 | 1,18E+08 |
| 16,8 | | 0,0004350 | 0,0015503 | 0,60 | 4,19E+07 | 4,84E+07 |
| 22,9 | | 0,0004377 | 0,0015482 | 0,41 | 4,43E+07 | 4,54E+07 |
| 16,8 | | 0,0004388 | 0,0015462 | 0,07 | 7,39E+07 | 6,65E+07 |
| 27,3 | | 0,0004398 | 0,0015441 | 0,15 | 4,80E+07 | 5,86E+07 |
| 28,4 | | 0,0004405 | 0,0015421 | 1,76 | 1,27E+09 | 1,18E+09 |
| 53 | | 0,0004408 | 0,0015401 | 1,29 | 2,25E+09 | 2,24E+09 |
| 33 | | 0,0004441 | 0,0015381 | 13,54 | 1,97E+07 | 3,10E+07 |
| 55,1 | | 0,0004469 | 0,0015360 | 1,89 | 3,15E+09 | 2,89E+09 |
| 55,7 | | 0,0004492 | 0,0015340 | 0,54 | 2,84E+08 | 3,11E+08 |
| 39,6 | | 0,0004497 | 0,0015320 | 0,70 | 5,75E+08 | 5,54E+08 |
| 66,7 | | 0,0004511 | 0,0015300 | 0,59 | 3,86E+08 | 4,48E+08 |
| 24,2 | | 0,0004563 | 0,0015280 | 0,85 | 2,89E+08 | 2,95E+08 |
| 64,5 | | 0,0004575 | 0,0015260 | 1,48 | 1,74E+09 | 1,63E+09 |
| 16,7 | | 0,0004577 | 0,0015241 | 0,49 | 6,15E+07 | 6,58E+07 |
| 27,9 | | 0,0004580 | 0,0015221 | 1,61 | 5,88E+07 | 5,41E+07 |
| 12,5 | | 0,0004589 | 0,0015201 | 1,65 | 1,46E+07 | 1,64E+07 |
| 3 | + | 0,0004592 | 0,0015181 | 4,16 | 9,11E+06 | 6,97E+06 |
| 7,7 | | 0,0004596 | 0,0015162 | 0,14 | 2,52E+07 | 2,21E+07 |
| 29 | | 0,0004610 | 0,0015142 | 0,56 | 1,99E+08 | 2,23E+08 |
| 38,6 | | 0,0004610 | 0,0015123 | 1,48 | 1,82E+09 | 1,86E+09 |
| 46,2 | | 0,0004614 | 0,0015103 | 1,51 | 4,83E+08 | 5,26E+08 |
| 29,2 | | 0,0004620 | 0,0015084 | 0,18 | 1,12E+08 | 1,67E+08 |
| 11,1 | + | 0,0004668 | 0,0015064 | 4,92 | 1,30E+08 | 1,95E+08 |
| 44 | | 0,0004688 | 0,0015045 | 0,57 | 5,27E+08 | 6,17E+08 |
| 1,7 | | 0,0004690 | 0,0015026 | 1,72 | 5,16E+07 | 5,69E+07 |
| 15,5 | | 0,0004710 | 0,0015006 | 8,72 | 3,22E+06 | 3,52E+06 |
| 33 | | 0,0004721 | 0,0014987 | 0,66 | 4,74E+08 | 4,28E+08 |
| 36,5 | | 0,0004737 | 0,0014968 | 0,73 | 3,04E+08 | 3,32E+08 |
| 20 | | 0,0004743 | 0,0014949 | 0,72 | 1,86E+08 | 1,71E+08 |
| 20,7 | | 0,0004786 | 0,0014930 | 0,04 | 5,58E+07 | 7,16E+07 |
| 55,5 | | 0,0004792 | 0,0014911 | 0,02 | 1,95E+08 | 1,80E+08 |
| 31,6 | | 0,0004796 | 0,0014892 | 0,02 | 1,35E+08 | 1,21E+08 |
| 47 | | 0,0004802 | 0,0014873 | 1,48 | 5,44E+08 | 5,25E+08 |
| 31,3 | | 0,0004824 | 0,0014854 | 0,54 | 1,50E+08 | 1,54E+08 |
| 25,3 | | 0,0004850 | 0,0014835 | 0,40 | 1,48E+08 | 1,33E+08 |
| 13,4 | | 0,0004854 | 0,0014817 | 0,18 | 2,10E+08 | 2,01E+08 |
| 23,2 | | 0,0004865 | 0,0014798 | 1,58 | 1,08E+08 | 1,10E+08 |
| 67,8 | | 0,0004879 | 0,0014779 | 1,67 | 1,19E+09 | 1,40E+09 |
| 37,6 | | 0,0004883 | 0,0014761 | 0,82 | 2,20E+09 | 2,17E+09 |
| 28,7 | | 0,0004924 | 0,0014742 | 1,69 | 3,83E+08 | 4,03E+08 |
| 27,7 | | 0,0004936 | 0,0014724 | 0,69 | 1,05E+09 | 1,04E+09 |

| | | | | | | |
|---|---|---|---|---|---|---|
| 14,6 | | 0,0004938 | 0,0014705 | 0,03 | 1,26E+08 | 1,29E+08 |
| 29,7 | | 0,0004948 | 0,0014687 | 0,01 | 2,53E+08 | 2,29E+08 |
| 71 | + | 0,0004977 | 0,0014668 | 10,85 | 1,32E+09 | 1,38E+09 |
| 19,4 | | 0,0004994 | 0,0014650 | 4,65 | 4,36E+06 | 6,19E+06 |
| 56,9 | | 0,0005031 | 0,0014632 | 0,64 | 3,25E+09 | 3,13E+09 |
| 35,2 | | 0,0005043 | 0,0014613 | 0,78 | 1,13E+08 | 1,17E+08 |
| 59,5 | | 0,0005071 | 0,0014595 | 0,63 | 8,01E+09 | 8,35E+09 |
| 7,3 | | 0,0005073 | 0,0014577 | 5,40 | 2,15E+06 | 3,71E+06 |
| 23,8 | | 0,0005073 | 0,0014559 | 0,27 | 6,73E+07 | 7,55E+07 |
| 44,8 | | 0,0005075 | 0,0014541 | 1,17 | 2,76E+08 | 2,73E+08 |
| 25 | | 0,0005080 | 0,0014523 | 1,28 | 6,06E+07 | 6,38E+07 |
| 10,4 | + | 0,0005112 | 0,0014505 | 3,10 | 6,12E+06 | 5,02E+06 |
| 40,8 | | 0,0005152 | 0,0014487 | 0,55 | 3,26E+08 | 3,45E+08 |
| 58,6 | | 0,0005161 | 0,0014469 | 0,86 | 1,40E+09 | 1,37E+09 |
| 62,1 | | 0,0005169 | 0,0014451 | 0,82 | 2,93E+08 | 2,89E+08 |
| 41,4 | | 0,0005170 | 0,0014433 | 3,63 | 9,70E+06 | 6,79E+06 |
| 66,1 | | 0,0005189 | 0,0015990 | 1,17 | 1,08E+10 | 1,05E+10 |
| 31,1 | | 0,0005196 | 0,0015971 | 1,72 | 5,97E+07 | 5,72E+07 |
| 20,9 | | 0,0005207 | 0,0015951 | 0,11 | 3,63E+07 | 2,65E+07 |
| 27,6 | | 0,0005225 | 0,0015931 | 0,66 | 1,35E+08 | 1,36E+08 |
| 51,6 | | 0,0005241 | 0,0015912 | 0,71 | 3,50E+08 | 3,24E+08 |
| 48,1 | | 0,0005244 | 0,0015892 | 1,47 | 5,03E+08 | 4,71E+08 |
| 35,8 | | 0,0005261 | 0,0015873 | 0,59 | 1,98E+08 | 2,23E+08 |
| 47,4 | | 0,0005272 | 0,0015854 | 0,52 | 6,38E+08 | 6,13E+08 |
| 43,6 | | 0,0005285 | 0,0015834 | 0,63 | 4,40E+08 | 4,26E+08 |
| 52,3 | | 0,0005293 | 0,0015815 | 0,50 | 3,11E+08 | 3,14E+08 |
| 49,5 | | 0,0005300 | 0,0015796 | 1,59 | 1,44E+08 | 1,32E+08 |
| 39,4 | | 0,0005305 | 0,0015777 | 1,88 | 3,82E+08 | 4,30E+08 |
| 13,3 | | 0,0005314 | 0,0015758 | 3,56 | 1,12E+07 | 1,23E+07 |
| 42,1 | | 0,0005375 | 0,0015738 | 2,17 | 1,09E+08 | 1,32E+08 |
| 32 | | 0,0005376 | 0,0015719 | 0,07 | 1,40E+08 | 8,07E+07 |
| 44,5 | | 0,0005393 | 0,0015700 | 2,35 | 7,65E+07 | 6,14E+07 |
| 71,7 | | 0,0005401 | 0,0015682 | 1,56 | 9,11E+07 | 8,62E+07 |
| 27,6 | | 0,0005409 | 0,0015663 | 1,41 | 7,16E+07 | 6,44E+07 |
| 16,8 | | 0,0005419 | 0,0015644 | 4,96 | 3,46E+06 | 4,61E+06 |
| 21,2 | | 0,0005423 | 0,0015625 | 2,10 | 2,14E+07 | 2,24E+07 |
| 34,2 | | 0,0005440 | 0,0015606 | 0,48 | 1,92E+08 | 1,61E+08 |
| 41,3 | | 0,0005460 | 0,0015588 | 1,16 | 5,55E+08 | 5,65E+08 |
| 39,6 | | 0,0005475 | 0,0015569 | 0,53 | 1,25E+09 | 1,30E+09 |
| 65 | + | 0,0005495 | 0,0015550 | 0,00 | 1,19E+09 | 4,75E+09 |
| 29,2 | | 0,0005513 | 0,0015532 | 1,63 | 6,33E+07 | 5,38E+07 |
| 69,3 | | 0,0005524 | 0,0015513 | 0,81 | 4,85E+08 | 5,05E+08 |
| 51,2 | | 0,0005550 | 0,0015495 | 0,77 | 5,59E+08 | 5,18E+08 |
| 22 | | 0,0005553 | 0,0015476 | 0,66 | 5,79E+07 | 6,03E+07 |
| 31,9 | | 0,0005569 | 0,0016742 | 0,84 | 4,94E+08 | 4,86E+08 |
| 25,9 | | 0,0005581 | 0,0016722 | 0,63 | 3,70E+08 | 4,21E+08 |
| 24,5 | | 0,0005603 | 0,0016702 | 1,72 | 5,30E+07 | 4,79E+07 |
| 33,3 | + | 0,0005613 | 0,0016682 | 0,57 | 2,24E+08 | 2,44E+08 |
| 34,9 | | 0,0005620 | 0,0016663 | 1,61 | 1,50E+08 | 1,31E+08 |
| 6,6 | | 0,0005660 | 0,0016643 | 3,86 | 3,54E+06 | 5,23E+06 |

| | | | | | | | |
|---|---|---|---|---|---|---|---|
| 34,4 | | | 0,0005673 | 0,0016623 | 0,31 | 7,74E+07 | 7,43E+07 |
| 16,5 | | | 0,0005717 | 0,0016604 | 0,28 | 3,07E+07 | 4,03E+07 |
| 53,6 | | | 0,0005717 | 0,0016584 | 0,64 | 3,33E+08 | 3,31E+08 |
| 32,2 | | | 0,0005728 | 0,0016565 | 1,27 | 7,06E+07 | 6,76E+07 |
| 25,9 | | | 0,0005731 | 0,0016545 | 0,45 | 4,39E+08 | 4,86E+08 |
| 32,9 | | | 0,0005736 | 0,0016526 | 0,78 | 4,41E+07 | 4,29E+07 |
| 59,1 | | | 0,0005741 | 0,0016506 | 25,51 | 2,06E+06 | 5,91E+06 |
| 59,7 | | | 0,0005761 | 0,0016487 | 0,79 | 1,56E+08 | 1,55E+08 |
| 24,3 | | | 0,0005777 | 0,0016468 | 0,03 | 6,60E+07 | 8,54E+07 |
| 66,1 | | | 0,0005778 | 0,0016449 | 1,47 | 6,37E+08 | 6,47E+08 |
| 13,1 | | | 0,0005783 | 0,0016429 | 2,01 | 2,82E+07 | 2,47E+07 |
| 7,9 | | | 0,0005805 | 0,0016410 | 1,91 | 1,20E+07 | 1,23E+07 |
| 34,9 | | | 0,0005814 | 0,0016391 | 0,59 | 6,61E+07 | 6,23E+07 |
| 25,3 | | | 0,0005833 | 0,0016372 | 1,27 | 8,63E+07 | 8,32E+07 |
| 53 | | | 0,0005841 | 0,0016353 | 0,80 | 1,68E+09 | 1,61E+09 |
| 31,7 | | | 0,0005848 | 0,0016334 | 0,46 | 2,88E+08 | 3,13E+08 |
| 18,4 | | | 0,0005851 | 0,0016315 | 2,92 | 2,01E+07 | 2,21E+07 |
| 46,8 | | | 0,0005925 | 0,0016296 | 0,65 | 4,96E+08 | 5,18E+08 |
| 65,7 | | | 0,0005928 | 0,0016277 | 1,27 | 1,50E+09 | 1,49E+09 |
| 15,4 | | | 0,0005950 | 0,0016259 | 0,64 | 4,42E+07 | 4,46E+07 |
| 30,8 | | | 0,0005980 | 0,0017347 | 0,73 | 1,16E+09 | 1,27E+09 |
| 19,1 | | | 0,0005981 | 0,0017327 | 0,04 | 7,16E+07 | 7,67E+07 |
| 39,9 | | | 0,0005986 | 0,0017307 | 0,56 | 2,29E+08 | 2,02E+08 |
| 31,6 | | | 0,0006016 | 0,0017287 | 1,22 | 2,62E+08 | 2,50E+08 |
| 18,9 | | | 0,0006021 | 0,0017268 | 0,44 | 1,51E+08 | 1,27E+08 |
| 44,3 | | | 0,0006029 | 0,0017248 | 1,27 | 2,34E+08 | 2,40E+08 |
| 50,9 | | | 0,0006030 | 0,0017228 | 1,32 | 8,95E+08 | 9,80E+08 |
| 33,7 | | | 0,0006036 | 0,0017208 | 1,37 | 1,90E+08 | 1,91E+08 |
| 19,3 | | | 0,0006036 | 0,0017189 | 0,69 | 9,43E+07 | 1,04E+08 |
| 63,6 | | | 0,0006042 | 0,0017169 | 1,27 | 6,35E+08 | 6,31E+08 |
| 18,4 | | | 0,0006046 | 0,0017149 | 0,86 | 7,51E+07 | 7,84E+07 |
| 21 | | | 0,0006064 | 0,0017130 | 0,12 | 2,68E+07 | 3,35E+07 |
| 48,4 | | | 0,0006076 | 0,0017110 | 0,71 | 1,81E+09 | 1,99E+09 |
| 45,7 | | | 0,0006085 | 0,0017091 | 4,53 | 3,67E+07 | 6,04E+07 |
| 39,6 | | | 0,0006099 | 0,0017072 | 0,68 | 3,75E+08 | 4,01E+08 |
| 32,5 | | | 0,0006100 | 0,0017052 | 68,98 | 7,12E+06 | 4,86E+06 |
| 29,3 | | | 0,0006130 | 0,0017033 | 0,59 | 4,59E+08 | 4,90E+08 |
| 26,4 | | | 0,0006144 | 0,0017014 | 5,45 | 1,26E+07 | 8,26E+06 |
| 19,1 | | | 0,0006150 | 0,0016994 | 0,43 | 7,47E+07 | 6,73E+07 |
| 24,8 | | | 0,0006161 | 0,0016975 | 0,57 | 4,92E+07 | 4,63E+07 |
| 16,6 | | | 0,0006198 | 0,0016956 | 0,40 | 1,75E+08 | 2,21E+08 |
| 57,9 | | | 0,0006198 | 0,0016937 | 1,59 | 1,07E+09 | 9,27E+08 |
| 25 | | | 0,0006199 | 0,0016918 | 1,42 | 7,45E+07 | 6,98E+07 |
| 54,3 | | | 0,0006206 | 0,0016899 | 0,57 | 2,18E+08 | 1,88E+08 |
| 35,1 | | | 0,0006207 | 0,0016880 | 0,02 | 1,55E+08 | 1,62E+08 |
| 19,4 | | | 0,0006208 | 0,0016861 | 0,72 | 7,73E+07 | 7,58E+07 |
| 11,5 | | | 0,0006212 | 0,0016842 | 0,35 | 7,70E+06 | 9,41E+06 |
| 50,9 | | | 0,0006245 | 0,0016823 | 0,74 | 8,37E+08 | 8,91E+08 |
| 57,4 | | | 0,0006247 | 0,0016804 | 1,16 | 3,59E+09 | 3,49E+09 |
| 49,6 | | | 0,0006392 | 0,0019241 | 0,56 | 1,51E+08 | 1,60E+08 |

| | | | | | | |
|---|---|---|---|---|---|---|
| 18 | | 0,0006426 | 0,0020424 | 0,12 | 5,96E+07 | 7,69E+07 |
| 45 | | 0,0006428 | 0,0020401 | 0,73 | 7,10E+08 | 7,42E+08 |
| 19,6 | | 0,0006440 | 0,0020378 | 0,63 | 1,30E+08 | 1,30E+08 |
| 11,6 | | 0,0006487 | 0,0020356 | 0,07 | 4,51E+07 | 4,75E+07 |
| 61,4 | | 0,0006492 | 0,0020333 | 0,77 | 2,03E+08 | 1,98E+08 |
| 34,7 | | 0,0006502 | 0,0020310 | 0,74 | 2,88E+08 | 2,98E+08 |
| 26,3 | | 0,0006502 | 0,0020288 | 2,98 | 2,56E+07 | 2,13E+07 |
| 44,4 | | 0,0006514 | 0,0020265 | 0,06 | 7,31E+07 | 5,97E+07 |
| 38,5 | | 0,0006519 | 0,0020243 | 86,82 | 1,72E+06 | 5,43E+05 |
| 44,1 | | 0,0006524 | 0,0020221 | 0,67 | 2,49E+08 | 2,47E+08 |
| 17,6 | | 0,0006540 | 0,0020198 | 0,54 | 3,22E+07 | 2,84E+07 |
| 27,8 | | 0,0006564 | 0,0020176 | 2,93 | 3,31E+06 | 4,76E+06 |
| 5,4 | | 0,0006580 | 0,0020154 | 7,51 | 5,53E+06 | 3,59E+06 |
| 26 | | 0,0006596 | 0,0020132 | 0,60 | 2,29E+08 | 2,58E+08 |
| 55,3 | | 0,0006600 | 0,0020110 | 0,68 | 1,97E+09 | 2,09E+09 |
| 48,1 | | 0,0006609 | 0,0020088 | 0,49 | 1,06E+08 | 1,15E+08 |
| 29,4 | | 0,0006622 | 0,0020066 | 1,34 | 1,11E+09 | 1,11E+09 |
| 36,7 | | 0,0006647 | 0,0020044 | 2,01 | 1,52E+08 | 1,60E+08 |
| 5,9 | | 0,0006694 | 0,0020022 | 0,61 | 1,27E+07 | 1,28E+07 |
| 25,5 | | 0,0006701 | 0,0020000 | 0,81 | 1,93E+08 | 1,98E+08 |
| 35,9 | | 0,0006711 | 0,0019978 | 1,27 | 7,11E+07 | 7,22E+07 |
| 42,9 | | 0,0006770 | 0,0019956 | 0,78 | 5,51E+08 | 5,52E+08 |
| 42,8 | | 0,0006814 | 0,0019935 | 3,00 | 4,88E+07 | 3,48E+07 |
| 42,2 | | 0,0006822 | 0,0019913 | 1,64 | 1,96E+08 | 2,19E+08 |
| 46,8 | | 0,0006840 | 0,0019891 | 5,09 | 1,05E+07 | 5,80E+06 |
| 9,5 | | 0,0006863 | 0,0019870 | 3,19 | 3,20E+06 | 2,93E+06 |
| 21,4 | | 0,0006869 | 0,0020888 | 2,21 | 4,17E+07 | 5,20E+07 |
| 65 | | 0,0006886 | 0,0020866 | 1,76 | 3,43E+08 | 4,19E+08 |
| 29,6 | | 0,0006938 | 0,0020843 | 0,72 | 1,71E+08 | 1,61E+08 |
| 31,1 | | 0,0006960 | 0,0020821 | 3,34 | 1,46E+07 | 2,00E+07 |
| 51,8 | | 0,0006973 | 0,0020798 | 0,55 | 5,93E+08 | 6,17E+08 |
| 18,9 | | 0,0006975 | 0,0020776 | 1,56 | 1,62E+07 | 1,74E+07 |
| 39,7 | | 0,0006991 | 0,0020753 | 0,14 | 6,94E+07 | 1,02E+08 |
| 30,5 | | 0,0006993 | 0,0020731 | 1,37 | 1,02E+08 | 1,09E+08 |
| 17,9 | | 0,0007009 | 0,0022041 | 1,61 | 7,52E+07 | 8,48E+07 |
| 27,5 | | 0,0007033 | 0,0022017 | 0,07 | 5,73E+07 | 5,52E+07 |
| 31,1 | | 0,0007103 | 0,0023151 | 0,61 | 1,86E+08 | 1,95E+08 |
| 29,2 | | 0,0007122 | 0,0023126 | 1,65 | 1,78E+08 | 1,74E+08 |
| 35,9 | | 0,0007131 | 0,0023102 | 0,71 | 2,23E+09 | 2,04E+09 |
| 36,5 | | 0,0007139 | 0,0023077 | 0,63 | 1,78E+08 | 1,77E+08 |
| 27,5 | | 0,0007184 | 0,0023052 | 70,59 | 3,46E+06 | 3,69E+06 |
| 40,9 | | 0,0007188 | 0,0023028 | 1,75 | 8,73E+07 | 8,40E+07 |
| 61,7 | | 0,0007188 | 0,0023003 | 1,29 | 4,66E+08 | 4,92E+08 |
| 60,4 | | 0,0007244 | 0,0022979 | 1,50 | 1,31E+08 | 1,45E+08 |
| 13,6 | | 0,0007306 | 0,0022954 | 0,15 | 1,95E+07 | 2,15E+07 |
| 17,7 | | 0,0007320 | 0,0022930 | 1,34 | 5,73E+07 | 6,29E+07 |
| 19,5 | | 0,0007352 | 0,0022906 | 0,12 | 2,69E+07 | 3,78E+07 |
| 17,4 | | 0,0007356 | 0,0022881 | 0,25 | 1,05E+08 | 1,23E+08 |
| 39,7 | | 0,0007371 | 0,0022857 | 0,47 | 4,69E+08 | 5,13E+08 |
| 46,3 | | 0,0007374 | 0,0022833 | 9,00 | 2,89E+06 | 5,33E+06 |

| | | | | | | |
|---|---|---|---|---|---|---|
| 53,6 | | 0,0007401 | 0,0022809 | 2,13 | 2,15E+08 | 2,24E+08 |
| 26,6 | | 0,0007409 | 0,0022785 | 0,64 | 8,70E+07 | 1,01E+08 |
| 51,3 | | 0,0007412 | 0,0022761 | 0,45 | 4,33E+08 | 5,73E+08 |
| 10,7 | | 0,0007433 | 0,0022737 | 0,07 | 2,85E+07 | 3,47E+07 |
| 25,9 | | 0,0007451 | 0,0022713 | 1,78 | 9,31E+07 | 1,05E+08 |
| 24,8 | | 0,0007465 | 0,0022689 | 0,63 | 3,28E+07 | 2,96E+07 |
| 67,7 | | 0,0007486 | 0,0022665 | 1,66 | 5,58E+08 | 6,21E+08 |
| 37,2 | | 0,0007512 | 0,0022642 | 0,84 | 1,35E+08 | 1,33E+08 |
| 6,9 | | 0,0007517 | 0,0022618 | 1,28 | 2,47E+07 | 2,50E+07 |
| 32,7 | | 0,0007543 | 0,0022594 | 2,54 | 1,15E+08 | 1,09E+08 |
| 24,8 | | 0,0007578 | 0,0022571 | 0,68 | 7,87E+07 | 8,46E+07 |
| 63,1 | | 0,0007590 | 0,0022547 | 12,66 | 3,81E+07 | 5,37E+07 |
| 39,4 | | 0,0007608 | 0,0022523 | 0,78 | 7,73E+08 | 7,52E+08 |
| 26,8 | | 0,0007616 | 0,0022500 | 0,52 | 1,55E+08 | 1,61E+08 |
| 45,6 | | 0,0007643 | 0,0022477 | 1,28 | 8,05E+07 | 8,04E+07 |
| 30,9 | | 0,0007656 | 0,0022453 | 1,64 | 9,98E+07 | 9,52E+07 |
| 32,8 | | 0,0007660 | 0,0022430 | 0,40 | 9,02E+07 | 9,42E+07 |
| 32,3 | | 0,0007673 | 0,0022407 | 0,61 | 1,42E+08 | 1,42E+08 |
| 36,5 | | 0,0007697 | 0,0022383 | 1,81 | 4,42E+08 | 4,89E+08 |
| 30,7 | | 0,0007706 | 0,0022360 | 1,66 | 6,54E+07 | 7,07E+07 |
| 27,3 | | 0,0007711 | 0,0022337 | 1,53 | 2,83E+08 | 2,43E+08 |
| 42,1 | | 0,0007712 | 0,0022314 | 0,83 | 4,24E+08 | 4,13E+08 |
| 19,6 | | 0,0007719 | 0,0022291 | 0,03 | 6,93E+07 | 5,69E+07 |
| 30,4 | | 0,0007771 | 0,0022268 | 1,32 | 2,48E+08 | 2,32E+08 |
| 47,9 | | 0,0007783 | 0,0022245 | 1,44 | 1,44E+08 | 1,39E+08 |
| 22,8 | | 0,0007793 | 0,0022222 | 1,61 | 2,55E+07 | 2,86E+07 |
| 20,9 | | 0,0007794 | 0,0022199 | 2,15 | 2,57E+07 | 2,24E+07 |
| 21,3 | | 0,0007795 | 0,0022177 | 0,56 | 1,12E+08 | 1,06E+08 |
| 40,5 | | 0,0007832 | 0,0022154 | 0,76 | 2,52E+08 | 2,48E+08 |
| 17,4 | | 0,0007834 | 0,0022131 | 20,95 | 1,25E+06 | 3,33E+06 |
| 30,9 | | 0,0007860 | 0,0022108 | 0,66 | 4,72E+08 | 4,56E+08 |
| 55 | | 0,0007890 | 0,0022086 | 0,81 | 4,65E+08 | 4,64E+08 |
| 18,7 | | 0,0007898 | 0,0022063 | 0,38 | 2,02E+07 | 1,90E+07 |
| 27,2 | | 0,0007898 | 0,0022041 | 0,78 | 1,14E+09 | 1,08E+09 |
| 18,8 | | 0,0007919 | 0,0022018 | 0,70 | 1,09E+08 | 1,08E+08 |
| 6,4 | | 0,0007935 | 0,0021996 | 0,50 | 2,28E+07 | 2,09E+07 |
| 48,4 | | 0,0007961 | 0,0021974 | 0,64 | 9,97E+07 | 1,10E+08 |
| 18,1 | | 0,0007986 | 0,0021951 | 4,97 | 3,55E+06 | 3,35E+06 |
| 40,6 | | 0,0008024 | 0,0021929 | 1,46 | 7,57E+07 | 6,96E+07 |
| 26 | | 0,0008055 | 0,0021907 | 0,50 | 2,85E+07 | 3,63E+07 |
| 19,2 | | 0,0008065 | 0,0021884 | 0,04 | 8,53E+07 | 7,04E+07 |
| 29,5 | | 0,0008086 | 0,0022794 | 0,08 | 5,93E+07 | 7,21E+07 |
| 28,9 | | 0,0008106 | 0,0022770 | 0,62 | 4,53E+07 | 5,02E+07 |
| 21 | | 0,0008121 | 0,0022747 | 2,26 | 1,94E+07 | 1,62E+07 |
| 2,7 | | 0,0008133 | 0,0022725 | 4,55 | 2,87E+06 | 2,52E+06 |
| 28,7 | | 0,0008145 | 0,0023669 | 0,78 | 2,75E+08 | 2,71E+08 |
| 37,2 | | 0,0008171 | 0,0023646 | 1,91 | 5,90E+08 | 5,98E+08 |
| 33,1 | | 0,0008204 | 0,0023622 | 0,45 | 4,68E+08 | 6,15E+08 |
| 15,3 | | 0,0008208 | 0,0023598 | 0,39 | 3,32E+07 | 2,89E+07 |
| 53,8 | | 0,0008237 | 0,0023574 | 0,60 | 1,51E+09 | 1,57E+09 |

| | | | | | | |
|---|---|---|---|---|---|---|
| 8 | | 0,0008254 | 0,0023551 | 13,56 | 1,23E+06 | 1,51E+06 |
| 7,4 | | 0,0008283 | 0,0023527 | 4,13 | 9,32E+06 | 6,70E+06 |
| 17,8 | | 0,0008335 | 0,0023504 | 0,79 | 1,20E+08 | 1,26E+08 |
| 37,4 | | 0,0008348 | 0,0023480 | 1,31 | 2,87E+08 | 3,06E+08 |
| 17,4 | + | 0,0008366 | 0,0023457 | 9,10 | 1,55E+07 | 8,73E+06 |
| 27,9 | | 0,0008376 | 0,0023433 | 0,82 | 2,33E+09 | 2,28E+09 |
| 39,5 | | 0,0008429 | 0,0023410 | 0,64 | 3,30E+08 | 2,91E+08 |
| 9,1 | | 0,0008449 | 0,0023386 | 0,66 | 4,16E+07 | 4,15E+07 |
| 86,3 | | 0,0008463 | 0,0023363 | 1,56 | 4,24E+10 | 3,75E+10 |
| 57,5 | | 0,0008508 | 0,0023340 | 1,35 | 6,69E+09 | 7,26E+09 |
| 6,9 | | 0,0008511 | 0,0023317 | 0,24 | 1,28E+07 | 1,04E+07 |
| 54,6 | | 0,0008512 | 0,0023294 | 0,61 | 1,23E+09 | 1,37E+09 |
| 19,2 | | 0,0008517 | 0,0023271 | 0,08 | 2,59E+07 | 3,66E+07 |
| 21,3 | | 0,0008527 | 0,0023248 | 34,57 | 2,04E+06 | 8,00E+06 |
| 38 | | 0,0008535 | 0,0023225 | 1,35 | 2,17E+08 | 2,02E+08 |
| 19,4 | | 0,0008543 | 0,0023202 | 0,47 | 3,96E+08 | 3,77E+08 |
| 2,7 | | 0,0008545 | 0,0023179 | 0,02 | 1,28E+08 | 1,45E+08 |
| 24,3 | | 0,0008546 | 0,0023156 | 0,26 | 1,88E+08 | 1,34E+08 |
| 29,8 | | 0,0008562 | 0,0023133 | 0,42 | 1,58E+08 | 1,70E+08 |
| 15,7 | | 0,0008658 | 0,0023110 | 2,64 | 7,99E+08 | 7,57E+08 |
| 26,9 | | 0,0008670 | 0,0023088 | 0,04 | 8,84E+07 | 7,87E+07 |
| 39,2 | | 0,0008738 | 0,0023065 | 0,59 | 2,11E+08 | 2,36E+08 |
| 58,8 | | 0,0008756 | 0,0023042 | 1,85 | 1,57E+09 | 1,71E+09 |
| 34,1 | | 0,0008779 | 0,0023020 | 1,34 | 1,12E+08 | 1,01E+08 |
| 24,1 | | 0,0008782 | 0,0022997 | 0,40 | 2,65E+07 | 2,46E+07 |
| 64,5 | | 0,0008786 | 0,0022975 | 1,73 | 1,45E+08 | 1,54E+08 |
| 23,5 | | 0,0008825 | 0,0022952 | 0,05 | 5,90E+07 | 5,90E+07 |
| 11,9 | | 0,0008828 | 0,0022930 | 1,86 | 1,17E+07 | 1,19E+07 |
| 17,5 | | 0,0008848 | 0,0022907 | 0,41 | 8,94E+07 | 1,07E+08 |
| 19,1 | | 0,0008859 | 0,0022885 | 1,70 | 2,89E+07 | 2,88E+07 |
| 12,3 | | 0,0008863 | 0,0022863 | 8,58 | 5,57E+06 | 3,24E+06 |
| 84,3 | | 0,0008869 | 0,0023774 | 0,71 | 1,62E+10 | 1,59E+10 |
| 14,5 | | 0,0008877 | 0,0023751 | 4,89 | 1,13E+07 | 7,00E+06 |
| 6,3 | | 0,0008878 | 0,0023728 | 0,09 | 5,00E+07 | 5,70E+07 |
| 39,7 | | 0,0008951 | 0,0023705 | 0,01 | 1,02E+09 | 9,33E+08 |
| 49 | | 0,0008966 | 0,0023682 | 1,47 | 4,11E+08 | 4,28E+08 |
| 29,9 | | 0,0009045 | 0,0025479 | 0,02 | 1,23E+08 | 1,43E+08 |
| 23,6 | | 0,0009064 | 0,0025455 | 1,58 | 6,44E+07 | 6,86E+07 |
| 14,8 | | 0,0009072 | 0,0025430 | 0,27 | 9,54E+06 | 9,63E+06 |
| 24,8 | | 0,0009076 | 0,0025405 | 1,86 | 3,42E+07 | 4,22E+07 |
| 29,8 | | 0,0009087 | 0,0025381 | 0,10 | 2,64E+08 | 2,42E+08 |
| 37,6 | | 0,0009139 | 0,0025356 | 0,56 | 1,98E+08 | 2,21E+08 |
| 14,1 | | 0,0009278 | 0,0027449 | 0,63 | 7,88E+07 | 8,82E+07 |
| 29,9 | | 0,0009285 | 0,0027423 | 0,72 | 3,58E+08 | 3,46E+08 |
| 37,7 | | 0,0009296 | 0,0027397 | 0,87 | 3,01E+08 | 3,14E+08 |
| 49,3 | | 0,0009302 | 0,0027370 | 0,75 | 3,11E+08 | 2,81E+08 |
| 23,2 | | 0,0009305 | 0,0027344 | 1,52 | 3,11E+08 | 3,18E+08 |
| 17,3 | | 0,0009306 | 0,0027318 | 0,06 | 3,76E+07 | 2,18E+07 |
| 35,8 | | 0,0009331 | 0,0027292 | 1,78 | 2,95E+08 | 3,33E+08 |
| 18,1 | | 0,0009335 | 0,0027266 | 0,16 | 2,52E+07 | 2,11E+07 |

| | | | | | | |
|---|---|---|---|---|---|---|
| 31,2 | | 0,0009350 | 0,0027240 | 1,55 | 4,34E+07 | 3,94E+07 |
| 29,7 | | 0,0009350 | 0,0027214 | 0,03 | 8,00E+07 | 6,15E+07 |
| 51,7 | | 0,0009359 | 0,0027188 | 0,55 | 1,39E+08 | 1,67E+08 |
| 68,6 | | 0,0009365 | 0,0027162 | 0,72 | 1,57E+09 | 1,79E+09 |
| 70,7 | | 0,0009369 | 0,0027136 | 1,65 | 3,78E+08 | 4,50E+08 |
| 50,1 | | 0,0009372 | 0,0027110 | 1,34 | 6,31E+08 | 5,65E+08 |
| 40,5 | | 0,0009405 | 0,0027085 | 1,43 | 1,20E+08 | 1,21E+08 |
| 22,5 | | 0,0009420 | 0,0027059 | 0,18 | 2,35E+07 | 2,33E+07 |
| 35,2 | | 0,0009454 | 0,0027033 | 0,02 | 1,46E+08 | 1,15E+08 |
| 13,4 | | 0,0009488 | 0,0027008 | 0,17 | 8,30E+07 | 6,51E+07 |
| 40,2 | | 0,0009540 | 0,0026982 | 0,75 | 2,27E+08 | 2,26E+08 |
| 42 | | 0,0009543 | 0,0026957 | 0,59 | 3,52E+08 | 4,02E+08 |
| 37,6 | + | 0,0009569 | 0,0028102 | 64,71 | 3,18E+06 | 1,45E+06 |
| 46,4 | | 0,0009570 | 0,0028075 | 2,31 | 1,96E+07 | 1,81E+07 |
| 34,1 | | 0,0009647 | 0,0028049 | 0,72 | 2,62E+08 | 2,68E+08 |
| 70 | | 0,0009663 | 0,0028023 | 1,63 | 1,04E+09 | 9,03E+08 |
| 41,4 | | 0,0009744 | 0,0029012 | 0,63 | 1,85E+08 | 1,70E+08 |
| 20 | | 0,0009762 | 0,0028985 | 0,84 | 5,27E+07 | 4,95E+07 |
| 51,3 | | 0,0009770 | 0,0028958 | 1,45 | 4,95E+08 | 4,40E+08 |
| 42,4 | | 0,0009803 | 0,0028931 | 0,19 | 1,99E+08 | 1,43E+08 |
| 20,4 | | 0,0009819 | 0,0028903 | 0,46 | 3,08E+07 | 3,67E+07 |
| 43,1 | | 0,0009903 | 0,0029738 | 0,22 | 8,11E+07 | 6,46E+07 |
| 15,7 | | 0,0009907 | 0,0029710 | 6,68 | 5,50E+06 | 2,72E+06 |
| 37,3 | | 0,0009928 | 0,0029682 | 0,57 | 7,06E+07 | 6,67E+07 |
| 29,6 | | 0,0009947 | 0,0029655 | 0,60 | 1,10E+08 | 1,10E+08 |
| 20,4 | | 0,0009965 | 0,0029627 | 0,38 | 5,66E+07 | 4,35E+07 |
| 52,1 | | 0,0009983 | 0,0029599 | 1,30 | 1,47E+09 | 1,41E+09 |
| 32,5 | | 0,0009986 | 0,0029572 | 1,41 | 3,91E+08 | 4,01E+08 |
| 16,2 | | 0,0010001 | 0,0029544 | 2,22 | 2,18E+07 | 1,62E+07 |
| 44,6 | | 0,0010076 | 0,0029517 | 0,13 | 8,06E+07 | 5,97E+07 |
| 49,2 | | 0,0010079 | 0,0029489 | 1,38 | 5,88E+09 | 6,41E+09 |
| 31,5 | | 0,0010091 | 0,0029462 | 0,28 | 1,12E+07 | 1,58E+07 |
| 12,2 | | 0,0010108 | 0,0029435 | 0,40 | 4,22E+07 | 3,96E+07 |
| 47,8 | | 0,0010112 | 0,0029407 | 0,56 | 1,21E+08 | 1,31E+08 |
| 50,4 | | 0,0010170 | 0,0029380 | 0,50 | 4,88E+08 | 4,11E+08 |
| 28,7 | | 0,0010217 | 0,0029353 | 0,64 | 1,53E+08 | 1,51E+08 |
| 43,2 | | 0,0010240 | 0,0029326 | 0,61 | 1,69E+08 | 1,96E+08 |
| 24,9 | | 0,0010258 | 0,0029299 | 0,05 | 1,17E+08 | 1,13E+08 |
| 34,9 | | 0,0010284 | 0,0029272 | 0,02 | 2,58E+08 | 8,00E+07 |
| 56 | | 0,0010290 | 0,0029245 | 0,87 | 2,40E+09 | 2,46E+09 |
| 47 | | 0,0010304 | 0,0029218 | 8,59 | 8,07E+06 | 3,57E+06 |
| 47,2 | | 0,0010344 | 0,0029191 | 1,37 | 1,35E+08 | 1,28E+08 |
| 67,5 | | 0,0010355 | 0,0029164 | 0,76 | 2,28E+09 | 2,51E+09 |
| 21,8 | | 0,0010411 | 0,0029138 | 0,52 | 5,40E+07 | 6,00E+07 |
| 21,6 | | 0,0010436 | 0,0029111 | 0,59 | 3,35E+08 | 3,79E+08 |
| 34,3 | | 0,0010442 | 0,0029084 | 1,68 | 1,70E+08 | 1,48E+08 |
| 5,7 | + | 0,0010452 | 0,0029058 | 0,07 | 4,91E+07 | 3,49E+07 |
| 27,6 | | 0,0010464 | 0,0029031 | 0,78 | 3,41E+08 | 3,50E+08 |
| 37,6 | | 0,0010468 | 0,0029005 | 0,86 | 2,96E+08 | 2,98E+08 |
| 21 | | 0,0010504 | 0,0030109 | 0,63 | 5,34E+07 | 4,82E+07 |

| | | | | | | |
|---|---|---|---|---|---|---|
| 54,4 | | 0,0010510 | 0,0030082 | 0,72 | 1,95E+08 | 2,09E+08 |
| 23,6 | | 0,0010519 | 0,0030055 | 30,57 | 7,11E+06 | 3,17E+06 |
| 56,3 | | 0,0010524 | 0,0030027 | 1,29 | 1,05E+09 | 1,01E+09 |
| 50 | | 0,0010621 | 0,0030000 | 0,60 | 6,41E+08 | 6,54E+08 |
| 5,3 | | 0,0010728 | 0,0029973 | 0,23 | 6,80E+06 | 7,64E+06 |
| 37 | | 0,0010765 | 0,0030817 | 1,52 | 6,87E+07 | 6,47E+07 |
| 61,3 | | 0,0010781 | 0,0030789 | 1,13 | 6,37E+08 | 6,47E+08 |
| 45,6 | | 0,0010793 | 0,0030761 | 0,27 | 1,53E+08 | 2,02E+08 |
| 38,6 | | 0,0010835 | 0,0030733 | 0,70 | 2,26E+08 | 2,10E+08 |
| 6,3 | | 0,0010875 | 0,0031537 | 0,45 | 6,43E+06 | 6,11E+06 |
| 16,3 | | 0,0010912 | 0,0031509 | 1,26 | 2,03E+07 | 2,22E+07 |
| 19,6 | | 0,0010912 | 0,0031480 | 0,53 | 6,09E+07 | 4,77E+07 |
| 72 | | 0,0010923 | 0,0031452 | 0,03 | 7,70E+07 | 1,11E+08 |
| 35,1 | | 0,0010938 | 0,0031423 | 0,74 | 4,48E+08 | 4,36E+08 |
| 42,1 | | 0,0010980 | 0,0031395 | 1,45 | 1,11E+08 | 1,04E+08 |
| 30 | | 0,0010985 | 0,0031367 | 0,01 | 2,47E+08 | 2,37E+08 |
| 47,3 | | 0,0011013 | 0,0031339 | 0,29 | 7,89E+07 | 5,49E+07 |
| 43,7 | | 0,0011020 | 0,0031311 | 1,56 | 1,20E+08 | 1,12E+08 |
| 43,4 | | 0,0011095 | 0,0031283 | 0,76 | 3,16E+08 | 3,51E+08 |
| 36,7 | | 0,0011135 | 0,0031254 | 0,05 | 6,50E+07 | 6,47E+07 |
| 27,8 | | 0,0011149 | 0,0031226 | 0,49 | 4,74E+08 | 6,10E+08 |
| 44,3 | | 0,0011198 | 0,0031199 | 0,72 | 4,95E+08 | 5,14E+08 |
| 36,8 | | 0,0011218 | 0,0031171 | 0,01 | 2,12E+08 | 1,39E+08 |
| 54,1 | | 0,0011233 | 0,0031143 | 1,16 | 1,87E+09 | 1,84E+09 |
| 54,9 | | 0,0011237 | 0,0031115 | 1,31 | 1,04E+09 | 1,13E+09 |
| 36,4 | | 0,0011262 | 0,0031087 | 1,93 | 7,21E+07 | 9,13E+07 |
| 66 | | 0,0011265 | 0,0031060 | 1,44 | 5,39E+08 | 6,22E+08 |
| 14,7 | | 0,0011297 | 0,0031032 | 1,55 | 3,64E+07 | 4,01E+07 |
| 47,6 | | 0,0011344 | 0,0031004 | 0,61 | 6,72E+08 | 7,68E+08 |
| 65 | | 0,0011353 | 0,0030977 | 0,62 | 2,90E+08 | 3,15E+08 |
| 35,3 | | 0,0011359 | 0,0030949 | 1,40 | 1,88E+08 | 2,02E+08 |
| 33,3 | | 0,0011359 | 0,0030922 | 0,02 | 6,96E+07 | 8,23E+07 |
| 40,5 | | 0,0011368 | 0,0030895 | 1,35 | 7,17E+07 | 7,84E+07 |
| 14,7 | | 0,0011413 | 0,0030867 | 2,06 | 3,35E+07 | 2,67E+07 |
| 50,5 | | 0,0011441 | 0,0030840 | 0,63 | 3,65E+08 | 3,20E+08 |
| 42,5 | | 0,0011457 | 0,0030813 | 1,43 | 1,88E+08 | 1,66E+08 |
| 35,8 | | 0,0011459 | 0,0030786 | 1,81 | 3,03E+08 | 3,78E+08 |
| 31,2 | | 0,0011547 | 0,0030758 | 2,36 | 5,44E+07 | 4,62E+07 |
| 9,7 | | 0,0011583 | 0,0030731 | 0,27 | 1,16E+07 | 1,41E+07 |
| 51,4 | | 0,0011587 | 0,0030704 | 1,38 | 1,48E+09 | 1,42E+09 |
| 75 | | 0,0011606 | 0,0030677 | 0,67 | 2,20E+09 | 2,15E+09 |
| 13,7 | | 0,0011607 | 0,0030650 | 0,19 | 1,55E+07 | 1,56E+07 |
| 50,2 | | 0,0011626 | 0,0030623 | 1,86 | 1,96E+08 | 2,51E+08 |
| 23,7 | | 0,0011649 | 0,0031544 | 0,59 | 5,55E+07 | 4,98E+07 |
| 23,8 | | 0,0011673 | 0,0031516 | 1,72 | 6,21E+07 | 5,57E+07 |
| 26,1 | | 0,0011759 | 0,0031489 | 0,86 | 3,53E+08 | 3,62E+08 |
| 27,8 | | 0,0011768 | 0,0031461 | 0,79 | 1,04E+08 | 1,01E+08 |
| 22,6 | | 0,0011817 | 0,0031434 | 0,79 | 1,32E+08 | 1,38E+08 |
| 42,3 | | 0,0011824 | 0,0031406 | 1,47 | 1,30E+08 | 1,11E+08 |
| 35,6 | | 0,0011884 | 0,0031379 | 0,11 | 4,67E+07 | 8,11E+07 |

| | | | | | | |
|---|---|---|---|---|---|---|
| 35,1 | | 0,0011885 | 0,0031351 | 0,60 | 2,03E+08 | 1,88E+08 |
| 19,9 | | 0,0011890 | 0,0031324 | 2,08 | 8,82E+06 | 1,09E+07 |
| 32,9 | | 0,0011921 | 0,0031297 | 2,30 | 3,47E+07 | 4,47E+07 |
| 62,7 | | 0,0011961 | 0,0031270 | 0,48 | 1,38E+08 | 1,43E+08 |
| 24,6 | | 0,0011993 | 0,0031242 | 0,03 | 2,25E+08 | 2,44E+08 |
| 17,2 | | 0,0011997 | 0,0031215 | 0,30 | 4,64E+07 | 4,69E+07 |
| 21 | | 0,0012006 | 0,0031188 | 4,93 | 1,91E+07 | 3,11E+07 |
| 35,9 | | 0,0012142 | 0,0033380 | 1,46 | 3,51E+07 | 3,95E+07 |
| 22,3 | | 0,0012146 | 0,0033351 | 0,72 | 2,22E+08 | 2,41E+08 |
| 22,5 | | 0,0012154 | 0,0033322 | 0,56 | 9,56E+06 | 8,99E+06 |
| 18,6 | | 0,0012176 | 0,0033293 | 1,58 | 2,27E+07 | 2,45E+07 |
| 52,1 | + | 0,0012187 | 0,0033264 | 2,99 | 9,44E+07 | 1,03E+08 |
| 40,4 | | 0,0012196 | 0,0033236 | 1,50 | 2,81E+08 | 2,84E+08 |
| 19,5 | | 0,0012200 | 0,0033207 | 0,20 | 4,16E+07 | 6,81E+07 |
| 53,8 | | 0,0012228 | 0,0033178 | 1,66 | 1,13E+08 | 1,03E+08 |
| 31,8 | | 0,0012236 | 0,0033150 | 1,38 | 5,13E+08 | 4,96E+08 |
| 26,6 | | 0,0012257 | 0,0033121 | 2,00 | 2,54E+07 | 2,89E+07 |
| 60,6 | | 0,0012288 | 0,0033093 | 0,52 | 8,55E+09 | 7,02E+09 |
| 55,7 | | 0,0012295 | 0,0033064 | 2,24 | 2,17E+08 | 2,44E+08 |
| 44,7 | | 0,0012295 | 0,0033036 | 1,31 | 5,06E+08 | 5,19E+08 |
| 57,3 | | 0,0012296 | 0,0033008 | 1,60 | 9,66E+08 | 8,40E+08 |
| 16,8 | | 0,0012324 | 0,0033801 | 0,77 | 7,11E+07 | 7,29E+07 |
| 5,1 | | 0,0012346 | 0,0033772 | 0,17 | 3,81E+07 | 4,62E+07 |
| 32,4 | | 0,0012352 | 0,0033744 | 0,68 | 1,11E+08 | 1,24E+08 |
| 13,1 | | 0,0012358 | 0,0033715 | 17,86 | 1,28E+06 | 2,00E+06 |
| 27,3 | | 0,0012369 | 0,0033686 | 1,84 | 1,17E+08 | 1,37E+08 |
| 40,6 | | 0,0012375 | 0,0033657 | 1,35 | 9,11E+08 | 8,03E+08 |
| 5,4 | | 0,0012428 | 0,0033629 | 1,40 | 6,73E+06 | 6,36E+06 |
| 33,2 | | 0,0012454 | 0,0033600 | 1,71 | 3,72E+07 | 3,65E+07 |
| 28,7 | | 0,0012457 | 0,0033571 | 0,12 | 3,65E+07 | 4,63E+07 |
| 16,4 | | 0,0012515 | 0,0033543 | 0,16 | 1,45E+07 | 1,59E+07 |
| 40,8 | + | 0,0012578 | 0,0033514 | 121,02 | 6,15E+05 | 4,78E+05 |
| 56,2 | | 0,0012605 | 0,0033486 | 0,79 | 2,02E+09 | 2,13E+09 |
| 47,3 | | 0,0012626 | 0,0033458 | 1,09 | 1,01E+09 | 1,05E+09 |
| 26,6 | | 0,0012634 | 0,0033429 | 1,55 | 8,62E+07 | 7,51E+07 |
| 75,7 | | 0,0012663 | 0,0033401 | 0,51 | 7,88E+08 | 9,64E+08 |
| 38,9 | | 0,0012676 | 0,0033373 | 1,32 | 1,73E+08 | 1,74E+08 |
| 36,2 | | 0,0012691 | 0,0033345 | 1,90 | 1,15E+08 | 1,35E+08 |
| 11,9 | | 0,0012749 | 0,0033316 | 10,72 | 4,58E+06 | 9,19E+06 |
| 46 | | 0,0012812 | 0,0033288 | 0,87 | 4,04E+08 | 4,10E+08 |
| 7,1 | | 0,0012820 | 0,0033260 | 0,16 | 2,74E+07 | 1,25E+07 |
| 29,5 | | 0,0012823 | 0,0033232 | 1,72 | 5,35E+07 | 6,25E+07 |
| 20,5 | | 0,0012894 | 0,0033204 | 0,61 | 2,70E+07 | 2,42E+07 |
| 15 | | 0,0012950 | 0,0033176 | 1,51 | 9,53E+07 | 1,08E+08 |
| 41,3 | | 0,0012977 | 0,0033149 | 0,70 | 3,80E+09 | 3,41E+09 |
| 81,8 | | 0,0012985 | 0,0033121 | 1,27 | 9,97E+09 | 9,37E+09 |
| 43,2 | | 0,0013016 | 0,0033093 | 0,61 | 8,09E+08 | 7,19E+08 |
| 61 | | 0,0013022 | 0,0033065 | 1,17 | 4,41E+08 | 4,41E+08 |
| 24,5 | | 0,0013022 | 0,0033038 | 0,70 | 1,20E+08 | 1,13E+08 |
| 52,1 | | 0,0013026 | 0,0033010 | 1,30 | 3,14E+08 | 2,95E+08 |

| | | | | | | |
|---|---|---|---|---|---|---|
| 27,7 | | 0,0013139 | 0,0032982 | 0,29 | 1,59E+07 | 2,49E+07 |
| 11,1 | | 0,0013208 | 0,0032955 | 1,95 | 1,96E+07 | 2,07E+07 |
| 32,1 | | 0,0013272 | 0,0032927 | 0,55 | 8,95E+07 | 8,49E+07 |
| 17,3 | | 0,0013288 | 0,0033933 | 3,50 | 1,55E+07 | 2,60E+07 |
| 6,3 | | 0,0013335 | 0,0033905 | 1,52 | 1,82E+08 | 1,87E+08 |
| 36,9 | | 0,0013438 | 0,0033877 | 2,57 | 2,60E+07 | 3,46E+07 |
| 49,8 | | 0,0013456 | 0,0033849 | 0,70 | 2,36E+08 | 2,26E+08 |
| 13,5 | | 0,0013456 | 0,0033821 | 8,30 | 4,68E+06 | 1,98E+06 |
| 60,1 | | 0,0013468 | 0,0033793 | 0,85 | 2,02E+08 | 2,14E+08 |
| 11,4 | | 0,0013480 | 0,0033765 | 4,07 | 6,79E+06 | 8,72E+06 |
| 28,7 | | 0,0013487 | 0,0033737 | 1,79 | 1,08E+08 | 1,27E+08 |
| 33,1 | | 0,0013521 | 0,0033709 | 0,02 | 2,31E+08 | 2,34E+08 |
| 55 | | 0,0013526 | 0,0033681 | 0,76 | 7,05E+08 | 6,32E+08 |
| 39,6 | | 0,0013572 | 0,0034678 | 0,72 | 4,06E+08 | 4,42E+08 |
| 37 | | 0,0013625 | 0,0034649 | 1,50 | 1,12E+09 | 1,21E+09 |
| 24,2 | | 0,0013660 | 0,0034620 | 0,64 | 1,12E+08 | 9,61E+07 |
| 14,2 | | 0,0013666 | 0,0034592 | 4,40 | 4,56E+06 | 2,63E+06 |
| 36,6 | | 0,0013677 | 0,0034563 | 0,71 | 2,92E+08 | 3,14E+08 |
| 36 | | 0,0013680 | 0,0034535 | 0,70 | 1,30E+08 | 1,14E+08 |
| 56,5 | | 0,0013694 | 0,0034507 | 0,88 | 3,47E+09 | 3,51E+09 |
| 34,4 | | 0,0013721 | 0,0034478 | 0,52 | 2,35E+08 | 2,84E+08 |
| 65,6 | | 0,0013740 | 0,0034450 | 1,36 | 6,48E+08 | 7,30E+08 |
| 12,3 | | 0,0013748 | 0,0034422 | 1,87 | 1,10E+07 | 8,41E+06 |
| 26,4 | | 0,0013905 | 0,0035180 | 1,76 | 1,87E+07 | 2,22E+07 |
| 11,4 | | 0,0013914 | 0,0035152 | 0,48 | 1,39E+07 | 1,74E+07 |
| 55 | | 0,0013934 | 0,0035123 | 0,78 | 2,74E+08 | 2,67E+08 |
| 50,7 | | 0,0013934 | 0,0035094 | 1,53 | 3,97E+08 | 4,02E+08 |
| 43,2 | | 0,0013940 | 0,0035065 | 0,02 | 1,81E+08 | 1,89E+08 |
| 33,2 | | 0,0013949 | 0,0035037 | 1,38 | 1,38E+08 | 1,51E+08 |
| 56,3 | | 0,0013978 | 0,0035008 | 0,69 | 2,31E+09 | 2,33E+09 |
| 32 | | 0,0013993 | 0,0034980 | 1,43 | 1,36E+08 | 1,51E+08 |
| 9 | | 0,0014075 | 0,0034951 | 0,18 | 1,22E+07 | 1,13E+07 |
| 54,6 | | 0,0014111 | 0,0035932 | 0,56 | 1,21E+08 | 1,25E+08 |
| 35,3 | | 0,0014218 | 0,0035902 | 2,50 | 2,69E+07 | 2,11E+07 |
| 37 | | 0,0014234 | 0,0035873 | 1,43 | 2,74E+08 | 2,62E+08 |
| 8,1 | | 0,0014247 | 0,0035844 | 1,89 | 2,08E+07 | 2,00E+07 |
| 64,7 | | 0,0014290 | 0,0035815 | 1,56 | 5,45E+07 | 5,30E+07 |
| 72,3 | | 0,0014340 | 0,0035786 | 1,49 | 1,22E+09 | 1,28E+09 |
| 17,5 | | 0,0014344 | 0,0035757 | 0,36 | 5,54E+07 | 7,64E+07 |
| 18,5 | | 0,0014378 | 0,0035728 | 0,68 | 8,29E+07 | 8,83E+07 |
| 20,2 | | 0,0014431 | 0,0035699 | 8,51 | 3,33E+06 | 4,86E+06 |
| 45 | | 0,0014520 | 0,0035670 | 0,76 | 1,42E+09 | 1,48E+09 |
| 42,4 | | 0,0014554 | 0,0035642 | 1,15 | 1,40E+09 | 1,39E+09 |
| 3,8 | | 0,0014591 | 0,0035613 | 0,21 | 1,11E+07 | 1,04E+07 |
| 11,2 | | 0,0014614 | 0,0035584 | 0,19 | 1,59E+07 | 1,74E+07 |
| 32,5 | | 0,0014638 | 0,0035556 | 0,48 | 2,43E+07 | 2,42E+07 |
| 17 | | 0,0014650 | 0,0035527 | 2,86 | 2,73E+07 | 1,90E+07 |
| 7,5 | | 0,0014673 | 0,0035498 | 0,34 | 1,48E+07 | 1,15E+07 |
| 15,6 | | 0,0014699 | 0,0035470 | 0,05 | 7,90E+07 | 9,42E+07 |
| 28,4 | | 0,0014729 | 0,0036212 | 0,09 | 6,64E+07 | 7,51E+07 |

| | | | | | | |
|---|---|---|---|---|---|---|
| 21,9 | | 0,0014761 | 0,0036183 | 0,59 | 9,89E+07 | 9,25E+07 |
| 39,1 | | 0,0014761 | 0,0036154 | 0,62 | 3,47E+08 | 3,25E+08 |
| 15,6 | | 0,0014856 | 0,0036125 | 0,07 | 6,24E+07 | 5,85E+07 |
| 1,1 | | 0,0014891 | 0,0036096 | 0,45 | 1,01E+09 | 7,41E+08 |
| 11,2 | | 0,0014917 | 0,0036067 | 0,51 | 4,63E+07 | 5,15E+07 |
| 84,8 | | 0,0014937 | 0,0036038 | 0,75 | 4,48E+09 | 4,72E+09 |
| 14 | | 0,0014964 | 0,0036010 | 0,64 | 6,73E+07 | 6,38E+07 |
| 34,5 | | 0,0014998 | 0,0035981 | 0,01 | 1,32E+08 | 1,32E+08 |
| 22,1 | | 0,0015047 | 0,0035952 | 1,46 | 1,32E+08 | 1,53E+08 |
| 31,3 | | 0,0015242 | 0,0037803 | 0,60 | 1,48E+08 | 1,71E+08 |
| 20,2 | | 0,0015256 | 0,0037772 | 0,06 | 4,58E+07 | 3,32E+07 |
| 27,4 | | 0,0015310 | 0,0037742 | 1,96 | 6,63E+07 | 4,98E+07 |
| 25,6 | | 0,0015317 | 0,0037712 | 1,65 | 1,12E+08 | 9,17E+07 |
| 20,3 | | 0,0015342 | 0,0037683 | 0,82 | 1,76E+08 | 1,83E+08 |
| 7 | | 0,0015353 | 0,0037653 | 0,79 | 2,84E+07 | 2,92E+07 |
| 38,5 | | 0,0015434 | 0,0037623 | 0,65 | 2,50E+08 | 2,78E+08 |
| 43,1 | | 0,0015531 | 0,0037593 | 1,59 | 8,72E+08 | 9,60E+08 |
| 27,9 | + | 0,0015556 | 0,0037563 | 16,57 | 8,54E+06 | 1,06E+07 |
| 6,5 | | 0,0015641 | 0,0037534 | 2,03 | 3,80E+06 | 2,98E+06 |
| 36,9 | | 0,0015659 | 0,0037504 | 0,62 | 2,81E+08 | 2,68E+08 |
| 12 | | 0,0015696 | 0,0037474 | 0,62 | 7,08E+07 | 8,37E+07 |
| 29,4 | | 0,0015731 | 0,0037445 | 0,73 | 8,22E+07 | 8,75E+07 |
| 20,6 | | 0,0015732 | 0,0037415 | 0,04 | 5,47E+07 | 7,03E+07 |
| 21,1 | | 0,0015773 | 0,0037386 | 0,39 | 4,00E+07 | 3,60E+07 |
| 21,8 | | 0,0015809 | 0,0037356 | 0,81 | 4,12E+07 | 4,22E+07 |
| 18,3 | | 0,0015824 | 0,0037327 | 0,07 | 2,28E+07 | 3,88E+07 |
| 33,1 | | 0,0015846 | 0,0037298 | 0,72 | 1,65E+08 | 1,77E+08 |
| 42,7 | | 0,0015850 | 0,0037268 | 0,78 | 1,11E+09 | 1,13E+09 |
| 34,6 | | 0,0015855 | 0,0037239 | 0,65 | 4,54E+08 | 5,27E+08 |
| 35,9 | | 0,0015871 | 0,0037210 | 1,36 | 4,26E+08 | 4,54E+08 |
| 5,3 | | 0,0015873 | 0,0037181 | 0,25 | 9,37E+06 | 9,27E+06 |
| 25,1 | | 0,0015876 | 0,0037152 | 0,40 | 8,79E+07 | 7,32E+07 |
| 21,4 | | 0,0015896 | 0,0037123 | 1,86 | 3,17E+07 | 3,87E+07 |
| 26,3 | | 0,0015922 | 0,0037094 | 0,70 | 6,67E+08 | 6,95E+08 |
| 22 | | 0,0015927 | 0,0037065 | 1,50 | 1,36E+08 | 1,41E+08 |
| 10,8 | | 0,0015946 | 0,0037036 | 0,51 | 1,28E+07 | 1,09E+07 |
| 45 | | 0,0015961 | 0,0037007 | 1,46 | 2,68E+08 | 2,36E+08 |
| 39,6 | | 0,0016017 | 0,0036978 | 1,48 | 8,32E+08 | 8,48E+08 |
| 29,1 | | 0,0016102 | 0,0036949 | 1,65 | 7,13E+07 | 6,50E+07 |
| 31,1 | | 0,0016128 | 0,0036921 | 2,11 | 3,64E+07 | 2,96E+07 |
| 20,5 | | 0,0016159 | 0,0036892 | 13,51 | 3,71E+06 | 1,45E+06 |
| 31,4 | | 0,0016202 | 0,0036863 | 0,70 | 1,16E+08 | 1,17E+08 |
| 27,9 | | 0,0016261 | 0,0036835 | 1,21 | 1,03E+09 | 9,69E+08 |
| 29,2 | | 0,0016337 | 0,0036806 | 1,35 | 1,71E+08 | 1,66E+08 |
| 23,3 | | 0,0016378 | 0,0036778 | 2,56 | 6,43E+07 | 4,38E+07 |
| 79 | + | 0,0016389 | 0,0036749 | 0,00 | 8,77E+09 | 7,49E+09 |
| 14,4 | | 0,0016393 | 0,0036721 | 0,20 | 1,65E+07 | 9,09E+06 |
| 31,9 | | 0,0016396 | 0,0036692 | 2,22 | 1,08E+08 | 8,97E+07 |
| 41,6 | | 0,0016396 | 0,0036664 | 0,67 | 4,23E+08 | 3,77E+08 |
| 16,5 | | 0,0016441 | 0,0036636 | 0,15 | 2,25E+07 | 2,66E+07 |

| | | | | | | |
|---|---|---|---|---|---|---|
| 12,1 | | 0,0016498 | 0,0036608 | 0,41 | 1,42E+07 | 1,41E+07 |
| 27,1 | | 0,0016564 | 0,0036579 | 1,21 | 5,71E+07 | 5,39E+07 |
| 6,1 | | 0,0016575 | 0,0036551 | 1,75 | 4,68E+07 | 4,88E+07 |
| 45,8 | | 0,0016578 | 0,0036523 | 1,38 | 3,30E+08 | 3,80E+08 |
| 36,7 | | 0,0016582 | 0,0036495 | 2,59 | 1,93E+07 | 2,94E+07 |
| 23,3 | | 0,0016608 | 0,0036467 | 1,72 | 2,14E+07 | 2,59E+07 |
| 52,7 | | 0,0016654 | 0,0036439 | 0,77 | 6,50E+08 | 7,28E+08 |
| 57,1 | | 0,0016666 | 0,0036411 | 0,52 | 2,98E+09 | 3,07E+09 |
| 50 | | 0,0016720 | 0,0036383 | 1,54 | 3,28E+08 | 3,14E+08 |
| 54,6 | | 0,0016764 | 0,0036355 | 1,20 | 7,27E+08 | 6,95E+08 |
| 36,4 | | 0,0016832 | 0,0036327 | 1,39 | 2,25E+08 | 2,28E+08 |
| 63,8 | | 0,0016835 | 0,0036300 | 0,81 | 8,85E+08 | 9,37E+08 |
| 36,5 | | 0,0016850 | 0,0036272 | 3,80 | 3,05E+07 | 2,65E+07 |
| 52,6 | | 0,0016953 | 0,0036244 | 1,58 | 2,21E+08 | 2,23E+08 |
| 12,9 | | 0,0016958 | 0,0036217 | 0,33 | 7,85E+06 | 6,98E+06 |
| 32,8 | | 0,0016996 | 0,0036189 | 4,07 | 3,32E+07 | 4,87E+07 |
| 25,4 | | 0,0017066 | 0,0036161 | 0,65 | 9,30E+07 | 9,89E+07 |
| 24,6 | | 0,0017081 | 0,0036134 | 1,37 | 6,11E+07 | 5,96E+07 |
| 8,8 | | 0,0017084 | 0,0036106 | 1,93 | 4,03E+06 | 3,71E+06 |
| 16 | | 0,0017107 | 0,0036079 | 0,67 | 5,59E+08 | 5,61E+08 |
| 13,7 | + | 0,0017109 | 0,0036052 | 7,47 | 3,00E+06 | 1,72E+06 |
| 41 | | 0,0017113 | 0,0036024 | 0,66 | 3,17E+08 | 3,40E+08 |
| 57,6 | | 0,0017131 | 0,0035997 | 1,70 | 1,21E+08 | 1,34E+08 |
| 10 | | 0,0017151 | 0,0035970 | 1,56 | 9,76E+06 | 1,07E+07 |
| 17,5 | | 0,0017175 | 0,0035942 | 2,33 | 4,52E+07 | 5,04E+07 |
| 15,4 | | 0,0017205 | 0,0036732 | 1,16 | 1,69E+07 | 1,76E+07 |
| 31,7 | | 0,0017242 | 0,0036704 | 1,96 | 5,04E+07 | 4,55E+07 |
| 32 | | 0,0017266 | 0,0036677 | 0,68 | 1,29E+08 | 1,30E+08 |
| 43,3 | | 0,0017279 | 0,0036649 | 1,79 | 1,45E+08 | 1,71E+08 |
| 23,4 | | 0,0017295 | 0,0036621 | 0,72 | 4,44E+08 | 4,53E+08 |
| 51,4 | | 0,0017360 | 0,0036594 | 0,76 | 5,98E+08 | 5,72E+08 |
| 26,7 | | 0,0017364 | 0,0036566 | 1,32 | 5,90E+07 | 6,05E+07 |
| 11,8 | | 0,0017400 | 0,0036539 | 2,33 | 1,09E+07 | 1,14E+07 |
| 8,8 | | 0,0017456 | 0,0036511 | 2,68 | 4,46E+06 | 3,21E+06 |
| 45,7 | | 0,0017475 | 0,0036484 | 0,41 | 1,35E+08 | 1,56E+08 |
| 48,5 | | 0,0017508 | 0,0036456 | 1,35 | 1,10E+08 | 1,06E+08 |
| 29,7 | | 0,0017601 | 0,0036429 | 0,64 | 1,70E+08 | 1,94E+08 |
| 21,5 | | 0,0017637 | 0,0036402 | 1,51 | 1,05E+08 | 9,19E+07 |
| 18,4 | | 0,0017644 | 0,0036375 | 0,07 | 6,02E+07 | 6,36E+07 |
| 16,6 | | 0,0017658 | 0,0036347 | 0,17 | 1,67E+07 | 1,65E+07 |
| 27,4 | | 0,0017659 | 0,0036320 | 0,10 | 4,55E+07 | 3,89E+07 |
| 41,2 | | 0,0017716 | 0,0036293 | 0,72 | 2,07E+09 | 2,19E+09 |
| 31,2 | | 0,0017733 | 0,0036266 | 2,34 | 1,27E+08 | 1,47E+08 |
| 26,1 | | 0,0017761 | 0,0036239 | 4,48 | 7,40E+06 | 1,38E+07 |
| 38,3 | | 0,0017794 | 0,0036212 | 0,74 | 3,78E+08 | 3,86E+08 |
| 21,4 | | 0,0017850 | 0,0036185 | 0,04 | 8,08E+07 | 6,67E+07 |
| 32,9 | | 0,0017865 | 0,0036158 | 1,52 | 2,04E+08 | 1,99E+08 |
| 16,9 | | 0,0017882 | 0,0036131 | 0,67 | 3,19E+07 | 2,85E+07 |
| 31 | | 0,0017984 | 0,0036104 | 1,85 | 2,00E+08 | 1,91E+08 |
| 26,2 | | 0,0018000 | 0,0036077 | 2,24 | 3,06E+07 | 4,00E+07 |

| | | | | | | |
|---|---|---|---|---|---|---|
| 21,8 | | 0,0018048 | 0,0036050 | 0,53 | 1,72E+08 | 2,00E+08 |
| 52,2 | | 0,0018066 | 0,0036024 | 1,96 | 1,82E+09 | 2,31E+09 |
| 13,5 | | 0,0018111 | 0,0035997 | 0,04 | 6,03E+07 | 6,13E+07 |
| 38,6 | | 0,0018152 | 0,0035970 | 1,71 | 8,95E+08 | 8,18E+08 |
| 31,4 | | 0,0018157 | 0,0035944 | 0,70 | 3,55E+08 | 4,15E+08 |
| 60 | | 0,0018187 | 0,0035917 | 0,86 | 7,87E+09 | 8,05E+09 |
| 37,5 | | 0,0018193 | 0,0035891 | 0,02 | 1,05E+08 | 2,86E+08 |
| 11,6 | | 0,0018212 | 0,0035864 | 0,18 | 3,60E+07 | 3,43E+07 |
| 11,2 | | 0,0018324 | 0,0035838 | 0,19 | 2,17E+07 | 1,52E+07 |
| 39,8 | | 0,0018383 | 0,0035811 | 0,49 | 1,08E+08 | 1,01E+08 |
| 24,4 | | 0,0018406 | 0,0035785 | 1,53 | 2,85E+08 | 2,96E+08 |
| 22,7 | | 0,0018492 | 0,0035758 | 1,32 | 4,02E+07 | 3,63E+07 |
| 23,5 | | 0,0018604 | 0,0036439 | 0,07 | 2,78E+07 | 5,51E+07 |
| 27 | | 0,0018618 | 0,0036412 | 0,06 | 9,18E+07 | 1,03E+08 |
| 19,1 | | 0,0018653 | 0,0036385 | 0,57 | 2,68E+07 | 3,08E+07 |
| 15,2 | | 0,0018664 | 0,0037063 | 0,38 | 1,20E+07 | 1,34E+07 |
| 37,5 | | 0,0018682 | 0,0037036 | 2,27 | 1,33E+07 | 1,67E+07 |
| 70 | | 0,0018688 | 0,0037009 | 1,17 | 1,55E+09 | 1,65E+09 |
| 35,3 | | 0,0018735 | 0,0036982 | 1,98 | 2,59E+08 | 2,03E+08 |
| 48,6 | | 0,0018741 | 0,0036955 | 1,40 | 1,69E+08 | 1,75E+08 |
| 44,8 | | 0,0018745 | 0,0036928 | 1,08 | 3,72E+08 | 3,75E+08 |
| 63,8 | | 0,0018776 | 0,0036901 | 1,67 | 1,86E+10 | 1,65E+10 |
| 84,9 | | 0,0018838 | 0,0036874 | 0,61 | 2,47E+09 | 2,29E+09 |
| 37,6 | | 0,0018965 | 0,0036847 | 0,55 | 9,40E+07 | 8,53E+07 |
| 8,3 | | 0,0019020 | 0,0036820 | 0,32 | 3,03E+08 | 3,89E+08 |
| 19,8 | | 0,0019030 | 0,0036793 | 0,04 | 7,07E+07 | 5,70E+07 |
| 26,3 | | 0,0019046 | 0,0036766 | 1,95 | 9,36E+06 | 9,88E+06 |
| 34,7 | | 0,0019112 | 0,0036739 | 2,24 | 7,32E+07 | 5,70E+07 |
| 23,3 | | 0,0019118 | 0,0036713 | 1,27 | 9,52E+07 | 9,55E+07 |
| 34,3 | | 0,0019138 | 0,0036686 | 0,43 | 2,87E+08 | 2,24E+08 |
| 33,3 | | 0,0019141 | 0,0036659 | 1,68 | 5,03E+07 | 5,40E+07 |
| 21,8 | | 0,0019177 | 0,0036633 | 3,23 | 3,15E+06 | 2,92E+06 |
| 29,1 | | 0,0019179 | 0,0036606 | 1,34 | 8,24E+07 | 9,20E+07 |
| 26 | | 0,0019278 | 0,0036580 | 0,61 | 2,11E+08 | 1,88E+08 |
| 30,7 | | 0,0019314 | 0,0036553 | 0,05 | 5,12E+07 | 5,83E+07 |
| 18 | | 0,0019314 | 0,0036527 | 0,07 | 3,33E+07 | 2,68E+07 |
| 63,2 | | 0,0019324 | 0,0036500 | 1,74 | 3,24E+08 | 3,63E+08 |
| 53,5 | | 0,0019335 | 0,0036474 | 1,39 | 1,65E+08 | 1,61E+08 |
| 40,8 | | 0,0019392 | 0,0036448 | 1,77 | 7,48E+07 | 7,85E+07 |
| 36,4 | | 0,0019433 | 0,0036421 | 1,63 | 1,09E+08 | 1,09E+08 |
| 18,2 | | 0,0019479 | 0,0036395 | 8,69 | 3,90E+06 | 4,03E+06 |
| 23,5 | | 0,0019506 | 0,0036369 | 1,86 | 5,22E+07 | 6,13E+07 |
| 9,1 | | 0,0019543 | 0,0036343 | 4,83 | 3,23E+06 | 3,86E+06 |
| 22,8 | | 0,0019545 | 0,0036317 | 1,37 | 1,39E+08 | 1,43E+08 |
| 40,7 | | 0,0019570 | 0,0036290 | 0,80 | 2,90E+08 | 3,17E+08 |
| 17,4 | | 0,0019600 | 0,0036264 | 0,48 | 9,25E+07 | 8,34E+07 |
| 12,3 | | 0,0019606 | 0,0036238 | 0,65 | 2,59E+07 | 2,73E+07 |
| 63,2 | | 0,0019621 | 0,0036212 | 1,21 | 8,28E+09 | 8,98E+09 |
| 50,5 | | 0,0019625 | 0,0036186 | 0,71 | 9,40E+08 | 7,97E+08 |
| 35,7 | | 0,0019661 | 0,0036160 | 1,79 | 9,90E+07 | 9,12E+07 |

| | | | | | | |
|---|---|---|---|---|---|---|
| 20,4 | | 0,0019665 | 0,0036135 | 0,55 | 8,08E+07 | 6,48E+07 |
| 20,9 | | 0,0019719 | 0,0036881 | 0,64 | 8,93E+07 | 8,27E+07 |
| 75,3 | | 0,0019807 | 0,0037513 | 1,67 | 5,33E+08 | 5,54E+08 |
| 11,5 | | 0,0019813 | 0,0037486 | 0,61 | 3,63E+06 | 4,41E+06 |
| 16,2 | | 0,0019879 | 0,0037459 | 0,57 | 2,61E+07 | 2,26E+07 |
| 9,4 | | 0,0019937 | 0,0037432 | 0,23 | 1,18E+07 | 1,17E+07 |
| 29,1 | | 0,0019970 | 0,0037406 | 0,54 | 1,23E+08 | 1,24E+08 |
| 61 | | 0,0020016 | 0,0037379 | 1,62 | 9,05E+07 | 1,12E+08 |
| 32,7 | | 0,0020067 | 0,0037352 | 0,66 | 2,44E+08 | 2,40E+08 |
| 39,2 | | 0,0020130 | 0,0037326 | 2,08 | 3,77E+07 | 3,46E+07 |
| 19,1 | | 0,0020154 | 0,0037299 | 1,99 | 9,30E+06 | 1,10E+07 |
| 30,3 | | 0,0020190 | 0,0037273 | 0,63 | 1,40E+08 | 1,51E+08 |
| 27,9 | | 0,0020268 | 0,0037246 | 1,33 | 3,91E+07 | 3,83E+07 |
| 44,8 | | 0,0020271 | 0,0037220 | 0,67 | 2,65E+09 | 2,46E+09 |
| 70,6 | | 0,0020324 | 0,0037193 | 1,51 | 4,92E+08 | 4,41E+08 |
| 9,4 | | 0,0020352 | 0,0037167 | 1,56 | 3,49E+07 | 2,95E+07 |
| 18,9 | | 0,0020380 | 0,0037141 | 2,03 | 8,72E+07 | 6,96E+07 |
| 18 | | 0,0020397 | 0,0037115 | 0,79 | 8,43E+07 | 9,05E+07 |
| 31,2 | | 0,0020414 | 0,0037088 | 0,78 | 6,68E+07 | 6,92E+07 |
| 7 | | 0,0020418 | 0,0037062 | 0,37 | 3,53E+07 | 3,25E+07 |
| 30,3 | | 0,0020435 | 0,0037036 | 0,72 | 6,44E+07 | 6,53E+07 |
| 14 | | 0,0020517 | 0,0037010 | 0,76 | 6,27E+07 | 6,19E+07 |
| 47,7 | | 0,0020520 | 0,0036984 | 1,37 | 2,51E+08 | 2,59E+08 |
| 32,1 | | 0,0020525 | 0,0036958 | 1,44 | 7,62E+07 | 6,65E+07 |
| 35,7 | | 0,0020537 | 0,0036932 | 0,86 | 3,57E+08 | 3,56E+08 |
| 40,5 | | 0,0020564 | 0,0036906 | 0,13 | 7,90E+07 | 7,21E+07 |
| 12,7 | | 0,0020648 | 0,0036880 | 0,73 | 6,44E+06 | 5,84E+06 |
| 37,4 | | 0,0020652 | 0,0036854 | 1,76 | 7,08E+07 | 6,91E+07 |
| 42,6 | | 0,0020667 | 0,0036828 | 0,77 | 7,60E+08 | 8,00E+08 |
| 39,3 | | 0,0020695 | 0,0036802 | 1,44 | 2,44E+08 | 2,72E+08 |
| 44,4 | | 0,0020700 | 0,0036776 | 0,58 | 2,67E+08 | 2,77E+08 |
| 29,2 | | 0,0020706 | 0,0036751 | 0,11 | 4,59E+07 | 4,98E+07 |
| 24,5 | | 0,0020720 | 0,0036725 | 0,27 | 1,43E+08 | 1,76E+08 |
| 48,9 | | 0,0020733 | 0,0036699 | 0,66 | 9,68E+07 | 8,12E+07 |
| 28,7 | | 0,0020736 | 0,0036674 | 0,58 | 5,98E+07 | 5,82E+07 |
| 15,3 | | 0,0020777 | 0,0036648 | 1,47 | 1,37E+07 | 1,48E+07 |
| 29 | | 0,0020783 | 0,0036622 | 2,24 | 2,29E+07 | 2,06E+07 |
| 43,3 | | 0,0020787 | 0,0036597 | 1,25 | 1,90E+08 | 1,89E+08 |
| 32,4 | | 0,0020810 | 0,0036571 | 0,79 | 4,97E+08 | 5,24E+08 |
| 27,5 | | 0,0020851 | 0,0036546 | 1,51 | 1,89E+09 | 1,57E+09 |
| 47,3 | | 0,0020921 | 0,0036521 | 1,38 | 3,62E+08 | 3,77E+08 |
| 13,4 | | 0,0020945 | 0,0036495 | 0,60 | 3,73E+07 | 4,29E+07 |
| 65,7 | | 0,0021061 | 0,0036470 | 0,80 | 6,47E+09 | 6,60E+09 |
| 21,1 | | 0,0021074 | 0,0036444 | 3,65 | 9,79E+06 | 9,32E+06 |
| 32,7 | + | 0,0021112 | 0,0036419 | 2,01 | 4,17E+07 | 3,38E+07 |
| 4,9 | | 0,0021114 | 0,0036394 | 0,82 | 3,99E+07 | 4,27E+07 |
| 12 | | 0,0021139 | 0,0036369 | 0,45 | 8,26E+06 | 9,51E+06 |
| 14,2 | | 0,0021221 | 0,0037202 | 7,64 | 1,14E+07 | 4,38E+06 |
| 42,5 | | 0,0021249 | 0,0037176 | 0,68 | 1,63E+09 | 1,84E+09 |
| 19 | | 0,0021272 | 0,0037151 | 0,11 | 3,64E+07 | 4,30E+07 |

| | | | | | | |
|---|---|---|---|---|---|---|
| 8,7 | | 0,0021284 | 0,0037125 | 0,59 | 1,46E+07 | 1,77E+07 |
| 45,3 | | 0,0021299 | 0,0037099 | 0,61 | 7,08E+08 | 8,55E+08 |
| 33,7 | | 0,0021305 | 0,0037074 | 0,86 | 1,46E+08 | 1,49E+08 |
| 36,2 | | 0,0021330 | 0,0037048 | 1,23 | 1,87E+08 | 1,82E+08 |
| 17,5 | | 0,0021397 | 0,0037684 | 6,63 | 5,81E+06 | 3,68E+06 |
| 59 | | 0,0021424 | 0,0037658 | 1,14 | 2,68E+09 | 2,65E+09 |
| 9 | | 0,0021555 | 0,0037632 | 1,89 | 1,06E+07 | 1,36E+07 |
| 54 | | 0,0021558 | 0,0037607 | 0,72 | 1,32E+08 | 1,37E+08 |
| 78,6 | | 0,0021613 | 0,0037581 | 0,91 | 9,36E+08 | 9,61E+08 |
| 26,2 | | 0,0021622 | 0,0037555 | 0,60 | 1,18E+08 | 1,28E+08 |
| 62,8 | | 0,0021731 | 0,0037529 | 0,68 | 4,34E+08 | 4,67E+08 |
| 18,1 | | 0,0021737 | 0,0037503 | 1,51 | 5,45E+07 | 4,96E+07 |
| 28,3 | | 0,0021820 | 0,0037478 | 0,04 | 7,94E+07 | 1,26E+08 |
| 38,3 | | 0,0021856 | 0,0037452 | 1,63 | 4,34E+08 | 4,37E+08 |
| 10,3 | | 0,0021944 | 0,0037426 | 0,44 | 1,26E+07 | 1,72E+07 |
| 27,3 | | 0,0021970 | 0,0037401 | 0,71 | 1,49E+08 | 1,41E+08 |
| 15 | | 0,0022025 | 0,0037375 | 0,51 | 2,20E+07 | 2,18E+07 |
| 51,1 | | 0,0022034 | 0,0037350 | 0,02 | 1,56E+08 | 5,64E+08 |
| 21,8 | | 0,0022079 | 0,0037324 | 0,65 | 1,54E+07 | 1,78E+07 |
| 53,6 | | 0,0022105 | 0,0037299 | 0,42 | 4,68E+08 | 4,86E+08 |
| 18,6 | | 0,0022136 | 0,0037273 | 0,33 | 3,81E+07 | 3,23E+07 |
| 35,5 | | 0,0022156 | 0,0037248 | 0,57 | 3,64E+07 | 3,84E+07 |
| 22 | | 0,0022165 | 0,0037223 | 0,68 | 8,10E+07 | 7,62E+07 |
| 12,4 | | 0,0022203 | 0,0037197 | 0,08 | 6,26E+07 | 8,96E+07 |
| 38,4 | | 0,0022212 | 0,0037172 | 0,51 | 3,94E+07 | 4,43E+07 |
| 33,7 | | 0,0022254 | 0,0037147 | 1,74 | 1,12E+08 | 1,03E+08 |
| 28,5 | | 0,0022310 | 0,0037122 | 1,47 | 3,06E+07 | 3,68E+07 |
| 36,2 | | 0,0022375 | 0,0037096 | 1,13 | 1,96E+08 | 1,93E+08 |
| 26,7 | | 0,0022476 | 0,0037071 | 0,76 | 2,33E+08 | 2,43E+08 |
| 40,1 | | 0,0022507 | 0,0037046 | 1,62 | 2,76E+08 | 2,29E+08 |
| 17 | | 0,0022510 | 0,0037021 | 5,30 | 3,63E+06 | 8,22E+06 |
| 32,5 | | 0,0022556 | 0,0036996 | 0,63 | 4,60E+08 | 3,75E+08 |
| 36,1 | | 0,0022642 | 0,0036971 | 0,78 | 3,53E+08 | 3,39E+08 |
| 35,4 | | 0,0022647 | 0,0036946 | 0,44 | 6,99E+07 | 6,82E+07 |
| 70,9 | | 0,0022693 | 0,0036921 | 0,63 | 3,33E+08 | 4,00E+08 |
| 43,2 | | 0,0022697 | 0,0036896 | 1,33 | 2,71E+08 | 2,58E+08 |
| 42,5 | | 0,0022734 | 0,0036871 | 1,58 | 1,70E+08 | 1,87E+08 |
| 40,8 | | 0,0022739 | 0,0036846 | 0,01 | 6,62E+08 | 7,08E+08 |
| 72,6 | | 0,0022750 | 0,0036822 | 0,66 | 2,01E+09 | 2,27E+09 |
| 17,3 | | 0,0022758 | 0,0036797 | 1,28 | 9,05E+07 | 9,00E+07 |
| 19,4 | | 0,0022773 | 0,0036772 | 0,05 | 7,69E+07 | 6,31E+07 |
| 43,8 | | 0,0022891 | 0,0036747 | 0,55 | 1,13E+08 | 1,42E+08 |
| 16,8 | | 0,0022906 | 0,0036723 | 0,06 | 1,14E+08 | 1,32E+08 |
| 59,5 | | 0,0022922 | 0,0036698 | 0,81 | 1,61E+09 | 1,58E+09 |
| 61,8 | | 0,0022939 | 0,0036673 | 0,44 | 2,96E+08 | 2,89E+08 |
| 61,6 | | 0,0022953 | 0,0036649 | 1,29 | 1,21E+09 | 1,32E+09 |
| 39,9 | | 0,0022964 | 0,0036624 | 1,32 | 4,10E+08 | 3,86E+08 |
| 47,7 | | 0,0022996 | 0,0036600 | 0,70 | 3,31E+08 | 3,31E+08 |
| 5,2 | | 0,0023011 | 0,0036575 | 0,27 | 2,64E+07 | 1,98E+07 |
| 43,5 | | 0,0023032 | 0,0036551 | 157,47 | 5,76E+05 | 1,47E+06 |

| | | | | | | |
|---|---|---|---|---|---|---|
| 40,1 | | 0,0023036 | 0,0036526 | 10,25 | 4,59E+06 | 1,48E+06 |
| 42,1 | | 0,0023062 | 0,0036502 | 0,65 | 7,90E+07 | 8,52E+07 |
| 20 | | 0,0023098 | 0,0037118 | 6,44 | 3,78E+06 | 1,69E+06 |
| 31,7 | | 0,0023145 | 0,0037093 | 0,06 | 4,11E+07 | 5,72E+07 |
| 29 | | 0,0023181 | 0,0037069 | 7,51 | 5,68E+06 | 7,21E+06 |
| 22,2 | | 0,0023211 | 0,0037044 | 0,65 | 4,78E+07 | 5,17E+07 |
| 41,8 | | 0,0023391 | 0,0038563 | 0,80 | 1,80E+08 | 1,84E+08 |
| 6,7 | | 0,0023417 | 0,0038537 | 0,56 | 4,51E+06 | 4,23E+06 |
| 9 | | 0,0023430 | 0,0038512 | 0,12 | 2,02E+07 | 2,22E+07 |
| 48,6 | | 0,0023433 | 0,0038486 | 1,71 | 5,01E+08 | 4,21E+08 |
| 54,3 | | 0,0023448 | 0,0038461 | 1,43 | 3,50E+08 | 3,94E+08 |
| 25,6 | | 0,0023478 | 0,0038435 | 1,67 | 4,92E+07 | 5,71E+07 |
| 12,6 | | 0,0023487 | 0,0038410 | 0,42 | 2,36E+07 | 2,80E+07 |
| 48,8 | | 0,0023503 | 0,0038384 | 0,06 | 9,98E+07 | 7,80E+07 |
| 11,5 | | 0,0023662 | 0,0038359 | 2,16 | 3,97E+06 | 4,12E+06 |
| 20,1 | | 0,0023686 | 0,0038333 | 0,54 | 1,75E+07 | 1,97E+07 |
| 51,2 | | 0,0023752 | 0,0038308 | 1,30 | 6,51E+08 | 6,99E+08 |
| 27,4 | | 0,0023825 | 0,0038283 | 0,67 | 1,01E+08 | 1,22E+08 |
| 21,8 | | 0,0023918 | 0,0038257 | 2,84 | 1,69E+07 | 2,40E+07 |
| 13,6 | | 0,0023978 | 0,0038232 | 0,16 | 2,40E+07 | 2,33E+07 |
| 71,9 | | 0,0023986 | 0,0038207 | 0,01 | 2,57E+08 | 2,34E+08 |
| 32,5 | | 0,0024115 | 0,0038182 | 0,71 | 1,61E+08 | 1,45E+08 |
| 33,1 | | 0,0024164 | 0,0038157 | 1,29 | 9,34E+07 | 9,49E+07 |
| 30,3 | | 0,0024208 | 0,0038132 | 1,97 | 6,99E+07 | 8,05E+07 |
| 36,3 | | 0,0024222 | 0,0038107 | 1,19 | 4,42E+08 | 4,12E+08 |
| 29,7 | | 0,0024256 | 0,0038081 | 1,53 | 2,72E+07 | 2,98E+07 |
| 48,9 | | 0,0024259 | 0,0038056 | 0,00 | 1,72E+09 | 1,79E+09 |
| 26,9 | | 0,0024286 | 0,0038661 | 0,02 | 1,89E+08 | 1,94E+08 |
| 51,5 | | 0,0024294 | 0,0038636 | 1,12 | 8,82E+08 | 8,63E+08 |
| 20,3 | | 0,0024308 | 0,0038611 | 0,29 | 6,18E+07 | 4,55E+07 |
| 32,5 | | 0,0024328 | 0,0038585 | 15,83 | 4,38E+06 | 1,32E+07 |
| 25,5 | | 0,0024409 | 0,0038560 | 0,75 | 1,83E+08 | 1,86E+08 |
| 26,7 | | 0,0024500 | 0,0038535 | 0,01 | 2,00E+08 | 1,64E+09 |
| 37,1 | | 0,0024523 | 0,0038510 | 0,67 | 6,33E+08 | 7,21E+08 |
| 48 | | 0,0024557 | 0,0038485 | 0,79 | 1,11E+09 | 1,09E+09 |
| 27,3 | | 0,0024559 | 0,0038460 | 1,90 | 1,34E+08 | 1,64E+08 |
| 21,5 | | 0,0024570 | 0,0038434 | 0,73 | 7,48E+07 | 7,07E+07 |
| 16,1 | | 0,0024587 | 0,0039035 | 0,57 | 4,15E+07 | 4,34E+07 |
| 14,1 | | 0,0024606 | 0,0039010 | 0,51 | 3,19E+07 | 2,95E+07 |
| 28,4 | | 0,0024685 | 0,0038984 | 1,29 | 6,56E+07 | 6,82E+07 |
| 34,6 | | 0,0024692 | 0,0038959 | 0,47 | 1,09E+08 | 1,22E+08 |
| 9,9 | | 0,0024729 | 0,0038934 | 1,71 | 1,72E+07 | 1,83E+07 |
| 35,5 | | 0,0024735 | 0,0038908 | 1,40 | 5,39E+08 | 6,28E+08 |
| 31,2 | | 0,0024762 | 0,0038883 | 0,58 | 1,22E+08 | 1,06E+08 |
| 43,7 | | 0,0024769 | 0,0038858 | 0,62 | 2,25E+08 | 2,69E+08 |
| 47,1 | | 0,0024836 | 0,0038833 | 1,56 | 1,47E+08 | 1,80E+08 |
| 6,3 | | 0,0024851 | 0,0038808 | 0,14 | 2,18E+07 | 1,69E+07 |
| 59,8 | | 0,0024855 | 0,0038782 | 0,86 | 2,12E+09 | 2,22E+09 |
| 24,8 | | 0,0024946 | 0,0038757 | 1,82 | 4,91E+07 | 6,21E+07 |
| 32,1 | | 0,0024972 | 0,0038732 | 1,55 | 1,81E+08 | 1,56E+08 |

| | | | | | | |
|---|---|---|---|---|---|---|
| 38,6 | | 0,0025008 | 0,0038707 | 0,56 | 4,26E+08 | 4,77E+08 |
| 44 | | 0,0025009 | 0,0038682 | 0,67 | 4,80E+08 | 4,18E+08 |
| 30,7 | + | 0,0025094 | 0,0038657 | 0,46 | 4,37E+07 | 5,95E+07 |
| 72 | | 0,0025118 | 0,0038632 | 0,72 | 2,86E+09 | 3,16E+09 |
| 41,2 | | 0,0025169 | 0,0038607 | 0,51 | 1,57E+08 | 1,99E+08 |
| 29 | | 0,0025206 | 0,0038582 | 1,19 | 1,08E+08 | 9,98E+07 |
| 24,4 | | 0,0025268 | 0,0039176 | 0,17 | 5,02E+07 | 3,62E+07 |
| 63,2 | | 0,0025286 | 0,0039151 | 0,76 | 8,01E+09 | 7,84E+09 |
| 39,7 | | 0,0025326 | 0,0039125 | 0,64 | 8,75E+08 | 9,46E+08 |
| 37 | | 0,0025461 | 0,0039100 | 0,06 | 7,68E+07 | 7,65E+07 |
| 37 | | 0,0025522 | 0,0039075 | 0,26 | 1,80E+08 | 1,87E+08 |
| 10,2 | | 0,0025559 | 0,0039050 | 0,29 | 1,13E+07 | 1,47E+07 |
| 14,1 | | 0,0025566 | 0,0039025 | 2,80 | 1,40E+07 | 9,77E+06 |
| 50,5 | | 0,0025607 | 0,0039000 | 0,89 | 3,04E+08 | 3,12E+08 |
| 30,2 | | 0,0025637 | 0,0038975 | 0,05 | 5,02E+07 | 5,47E+07 |
| 15,3 | | 0,0025647 | 0,0038950 | 1,88 | 1,40E+07 | 1,29E+07 |
| 15,3 | | 0,0025669 | 0,0038925 | 0,79 | 8,08E+07 | 7,53E+07 |
| 67,2 | | 0,0025690 | 0,0038900 | 0,58 | 8,90E+08 | 8,33E+08 |
| 46,9 | | 0,0025709 | 0,0038875 | 0,64 | 1,08E+09 | 9,37E+08 |
| 4,2 | | 0,0025757 | 0,0038851 | 1,25 | 1,43E+07 | 1,52E+07 |
| 22,5 | | 0,0025777 | 0,0039617 | 0,06 | 3,29E+07 | 3,85E+07 |
| 18,5 | | 0,0025850 | 0,0039592 | 0,38 | 3,47E+07 | 4,37E+07 |
| 19,3 | | 0,0025906 | 0,0039567 | 4,86 | 7,42E+06 | 3,49E+06 |
| 46,9 | | 0,0025968 | 0,0039541 | 0,67 | 7,28E+08 | 8,51E+08 |
| 67,4 | | 0,0025974 | 0,0039516 | 1,65 | 9,59E+07 | 1,18E+08 |
| 23,1 | | 0,0025978 | 0,0039491 | 0,30 | 4,93E+07 | 3,44E+07 |
| 13,8 | + | 0,0026057 | 0,0039466 | 8,27 | 3,00E+06 | 7,01E+06 |
| 18,8 | | 0,0026079 | 0,0039441 | 0,61 | 6,91E+07 | 7,27E+07 |
| 60,1 | | 0,0026124 | 0,0039416 | 1,18 | 1,74E+09 | 1,70E+09 |
| 19,9 | | 0,0026131 | 0,0039391 | 1,80 | 3,57E+07 | 2,87E+07 |
| 75,5 | | 0,0026169 | 0,0039366 | 1,64 | 2,10E+08 | 2,06E+08 |
| 45,7 | | 0,0026178 | 0,0039341 | 0,66 | 1,34E+08 | 1,54E+08 |
| 49,7 | | 0,0026353 | 0,0039316 | 59,19 | 3,27E+07 | 9,04E+06 |
| 71,1 | | 0,0026353 | 0,0039291 | 0,65 | 1,74E+09 | 1,56E+09 |
| 56,6 | | 0,0026361 | 0,0039266 | 0,82 | 1,00E+09 | 9,99E+08 |
| 35,3 | | 0,0026365 | 0,0039241 | 0,89 | 1,52E+09 | 1,53E+09 |
| 18,5 | | 0,0026534 | 0,0039823 | 1,30 | 1,34E+07 | 1,37E+07 |
| 8,2 | | 0,0026553 | 0,0039798 | 3,05 | 3,53E+06 | 4,71E+06 |
| 31,5 | | 0,0026662 | 0,0039773 | 1,22 | 6,93E+07 | 6,56E+07 |
| 41,3 | | 0,0026691 | 0,0040530 | 1,53 | 9,27E+08 | 8,91E+08 |
| 59,1 | | 0,0026713 | 0,0041285 | 0,43 | 2,08E+08 | 1,39E+08 |
| 27,6 | | 0,0026746 | 0,0041259 | 1,23 | 4,75E+08 | 4,90E+08 |
| 49,3 | | 0,0026846 | 0,0041233 | 1,08 | 1,60E+09 | 1,56E+09 |
| 26,1 | | 0,0026976 | 0,0042365 | 0,63 | 1,49E+08 | 1,58E+08 |
| 45,3 | | 0,0027063 | 0,0042338 | 0,67 | 5,06E+08 | 4,44E+08 |
| 24,9 | | 0,0027076 | 0,0042312 | 18,02 | 1,85E+06 | 8,11E+06 |
| 58,8 | | 0,0027122 | 0,0042285 | 0,70 | 2,76E+09 | 3,29E+09 |
| 25,2 | | 0,0027252 | 0,0042258 | 1,26 | 4,34E+08 | 4,26E+08 |
| 78,8 | | 0,0027294 | 0,0042232 | 1,43 | 9,19E+08 | 1,10E+09 |
| 48,9 | | 0,0027321 | 0,0042206 | 0,60 | 1,24E+08 | 1,09E+08 |

| | | | | | | |
|---|---|---|---|---|---|---|
| 51,4 | | 0,0027377 | 0,0042179 | 1,29 | 2,88E+09 | 3,10E+09 |
| 42,4 | | 0,0027552 | 0,0042829 | 1,67 | 2,15E+08 | 2,07E+08 |
| 32,1 | | 0,0027628 | 0,0044353 | 0,46 | 1,37E+08 | 1,04E+08 |
| 20,7 | | 0,0027654 | 0,0044325 | 0,12 | 4,27E+07 | 3,31E+07 |
| 26,9 | | 0,0027665 | 0,0044297 | 0,49 | 2,32E+08 | 1,95E+08 |
| 45 | | 0,0027685 | 0,0044270 | 0,82 | 4,11E+08 | 4,52E+08 |
| 16,5 | | 0,0027758 | 0,0044841 | 0,11 | 5,37E+07 | 6,36E+07 |
| 34,5 | | 0,0027814 | 0,0044813 | 0,01 | 1,75E+08 | 2,02E+08 |
| 30,4 | | 0,0027815 | 0,0044785 | 0,43 | 9,57E+07 | 1,30E+08 |
| 27,1 | | 0,0027994 | 0,0044757 | 1,24 | 1,55E+08 | 1,52E+08 |
| 12,6 | | 0,0027995 | 0,0044729 | 0,66 | 5,12E+08 | 5,66E+08 |
| 48,8 | | 0,0028010 | 0,0044701 | 1,28 | 2,66E+09 | 2,91E+09 |
| 19,9 | | 0,0028167 | 0,0044674 | 1,18 | 7,89E+07 | 7,95E+07 |
| 18,6 | | 0,0028237 | 0,0045242 | 0,18 | 2,61E+07 | 1,30E+07 |
| 32,7 | | 0,0028258 | 0,0045214 | 1,37 | 1,35E+08 | 1,54E+08 |
| 27,4 | | 0,0028349 | 0,0045186 | 0,09 | 1,95E+07 | 3,00E+07 |
| 30,3 | | 0,0028429 | 0,0045158 | 0,67 | 3,24E+08 | 3,81E+08 |
| 28,5 | | 0,0028430 | 0,0045130 | 0,12 | 3,32E+07 | 3,17E+07 |
| 26,7 | | 0,0028466 | 0,0045102 | 0,57 | 1,50E+08 | 1,26E+08 |
| 59,8 | | 0,0028531 | 0,0045074 | 1,19 | 2,05E+09 | 1,91E+09 |
| 22,9 | | 0,0028542 | 0,0045046 | 0,66 | 3,59E+07 | 3,03E+07 |
| 13,8 | | 0,0028587 | 0,0045019 | 0,14 | 3,75E+07 | 4,75E+07 |
| 35,4 | | 0,0028656 | 0,0044991 | 0,03 | 8,64E+07 | 1,05E+08 |
| 14,1 | | 0,0028662 | 0,0044963 | 0,49 | 1,23E+07 | 1,31E+07 |
| 42,6 | | 0,0028696 | 0,0044935 | 0,91 | 2,98E+08 | 2,90E+08 |
| 42,3 | | 0,0028726 | 0,0044908 | 1,20 | 1,94E+08 | 1,92E+08 |
| 6,4 | | 0,0028739 | 0,0044880 | 1,46 | 2,10E+07 | 2,43E+07 |
| 15,1 | | 0,0028801 | 0,0044852 | 0,21 | 1,04E+07 | 8,72E+06 |
| 18,7 | | 0,0028837 | 0,0044825 | 0,90 | 2,35E+08 | 2,26E+08 |
| 45,9 | | 0,0028888 | 0,0044797 | 1,48 | 4,19E+08 | 3,67E+08 |
| 25,7 | | 0,0028963 | 0,0044770 | 0,67 | 6,45E+08 | 7,38E+08 |
| 12,8 | | 0,0028974 | 0,0044742 | 0,34 | 8,43E+06 | 7,34E+06 |
| 39,6 | | 0,0029050 | 0,0044715 | 0,53 | 4,18E+08 | 5,36E+08 |
| 15,3 | | 0,0029071 | 0,0044687 | 0,06 | 5,15E+07 | 4,47E+07 |
| 56,1 | | 0,0029076 | 0,0044660 | 0,62 | 1,26E+09 | 1,20E+09 |
| 30,2 | | 0,0029096 | 0,0044632 | 3,34 | 9,77E+06 | 7,65E+06 |
| 41,8 | | 0,0029114 | 0,0044605 | 1,26 | 8,32E+07 | 7,40E+07 |
| 38,8 | | 0,0029195 | 0,0044578 | 1,37 | 1,58E+08 | 1,75E+08 |
| 31,5 | | 0,0029197 | 0,0044550 | 0,69 | 1,74E+08 | 1,79E+08 |
| 20,5 | | 0,0029231 | 0,0044523 | 1,30 | 3,05E+07 | 2,91E+07 |
| 52,8 | | 0,0029485 | 0,0044496 | 1,53 | 1,97E+08 | 1,77E+08 |
| 26,2 | | 0,0029503 | 0,0044469 | 0,60 | 2,51E+07 | 3,18E+07 |
| 26 | | 0,0029583 | 0,0044442 | 0,07 | 1,23E+08 | 2,00E+08 |
| 19,3 | | 0,0029598 | 0,0044415 | 0,44 | 1,89E+07 | 2,84E+07 |
| 35 | | 0,0029629 | 0,0044388 | 0,59 | 2,23E+08 | 2,53E+08 |
| 9,7 | | 0,0029657 | 0,0044361 | 0,19 | 8,54E+06 | 1,94E+07 |
| 69,2 | | 0,0029671 | 0,0044334 | 1,38 | 1,73E+09 | 1,87E+09 |
| 20,5 | | 0,0029701 | 0,0044307 | 0,71 | 8,58E+08 | 8,46E+08 |
| 16,3 | | 0,0029704 | 0,0044280 | 2,27 | 1,60E+08 | 1,29E+08 |
| 64,4 | | 0,0029720 | 0,0044253 | 1,24 | 1,35E+09 | 1,44E+09 |

| | | | | | | |
|---|---|---|---|---|---|---|
| 22,2 | | 0,0029730 | 0,0044979 | 0,69 | 5,32E+07 | 5,54E+07 |
| 29,2 | | 0,0029800 | 0,0044951 | 0,61 | 3,05E+08 | 2,59E+08 |
| 17,3 | | 0,0029842 | 0,0044924 | 1,74 | 3,63E+07 | 3,11E+07 |
| 21,9 | | 0,0029989 | 0,0044897 | 2,27 | 1,20E+07 | 1,51E+07 |
| 84,2 | | 0,0030006 | 0,0044870 | 1,84 | 2,78E+08 | 3,51E+08 |
| 38,9 | | 0,0030036 | 0,0044843 | 0,65 | 7,10E+07 | 7,96E+07 |
| 39,8 | | 0,0030077 | 0,0044815 | 0,83 | 3,81E+08 | 3,45E+08 |
| 5,3 | | 0,0030082 | 0,0044788 | 0,28 | 1,15E+07 | 1,10E+07 |
| 53,8 | | 0,0030205 | 0,0045341 | 0,73 | 2,89E+08 | 3,29E+08 |
| 37,2 | | 0,0030212 | 0,0045314 | 0,23 | 7,84E+07 | 7,47E+07 |
| 19,4 | + | 0,0030274 | 0,0046687 | 4,59 | 9,95E+06 | 1,42E+07 |
| 47,8 | | 0,0030313 | 0,0046659 | 0,58 | 5,02E+08 | 4,35E+08 |
| 22 | | 0,0030355 | 0,0046631 | 0,63 | 5,20E+07 | 5,29E+07 |
| 28,2 | | 0,0030441 | 0,0046602 | 0,50 | 4,98E+07 | 5,36E+07 |
| 57,8 | | 0,0030445 | 0,0046574 | 0,79 | 1,77E+08 | 1,97E+08 |
| 35,3 | | 0,0030453 | 0,0046546 | 1,69 | 4,71E+07 | 4,66E+07 |
| 22 | | 0,0030617 | 0,0046518 | 0,77 | 2,68E+08 | 2,70E+08 |
| 31,8 | | 0,0030740 | 0,0047067 | 0,62 | 6,84E+07 | 7,96E+07 |
| 48,9 | | 0,0030758 | 0,0047039 | 0,70 | 4,35E+08 | 3,98E+08 |
| 33,9 | | 0,0030821 | 0,0047011 | 1,40 | 7,62E+07 | 7,41E+07 |
| 38,8 | | 0,0030879 | 0,0046983 | 1,40 | 2,11E+08 | 1,91E+08 |
| 27,4 | | 0,0030882 | 0,0046954 | 0,57 | 2,42E+08 | 1,97E+08 |
| 65,7 | | 0,0031113 | 0,0046926 | 1,76 | 3,58E+09 | 3,77E+09 |
| 36,6 | | 0,0031149 | 0,0046898 | 1,81 | 1,79E+08 | 1,62E+08 |
| 45,8 | | 0,0031181 | 0,0046870 | 0,10 | 2,37E+07 | 7,67E+07 |
| 14,1 | | 0,0031184 | 0,0046842 | 0,68 | 1,88E+08 | 2,20E+08 |
| 23 | + | 0,0031185 | 0,0046814 | 10,84 | 2,64E+06 | 6,40E+06 |
| 28,4 | | 0,0031268 | 0,0046786 | 0,73 | 1,74E+08 | 1,77E+08 |
| 42,7 | | 0,0031277 | 0,0046758 | 1,37 | 3,87E+08 | 3,65E+08 |
| 19,7 | | 0,0031308 | 0,0046730 | 0,32 | 3,11E+08 | 1,91E+08 |
| 47,6 | | 0,0031423 | 0,0046702 | 0,85 | 2,46E+08 | 2,35E+08 |
| 41,1 | | 0,0031446 | 0,0046675 | 0,68 | 2,12E+08 | 2,34E+08 |
| 31,8 | | 0,0031506 | 0,0046647 | 0,49 | 7,48E+07 | 6,82E+07 |
| 65,4 | | 0,0031566 | 0,0046619 | 0,86 | 1,26E+09 | 1,32E+09 |
| 12,2 | | 0,0031576 | 0,0046591 | 0,67 | 3,59E+07 | 3,98E+07 |
| 16,7 | | 0,0031606 | 0,0046564 | 0,23 | 5,91E+07 | 5,08E+07 |
| 46,5 | | 0,0031617 | 0,0046536 | 0,78 | 9,80E+07 | 1,03E+08 |
| 36,9 | | 0,0031660 | 0,0046508 | 0,90 | 6,72E+08 | 6,38E+08 |
| 59,7 | | 0,0031678 | 0,0046481 | 0,83 | 1,08E+09 | 1,05E+09 |
| 22,1 | | 0,0031787 | 0,0046453 | 2,21 | 3,34E+07 | 3,26E+07 |
| 25,4 | | 0,0031796 | 0,0046426 | 0,63 | 5,79E+07 | 5,45E+07 |
| 41 | | 0,0031809 | 0,0046398 | 0,60 | 4,26E+08 | 3,31E+08 |
| 21,9 | | 0,0031816 | 0,0046371 | 3,21 | 9,94E+06 | 1,79E+07 |
| 61,1 | | 0,0031885 | 0,0046343 | 1,88 | 5,44E+08 | 6,17E+08 |
| 51,4 | | 0,0032067 | 0,0047617 | 0,92 | 5,20E+08 | 5,27E+08 |
| 14 | + | 0,0032109 | 0,0048156 | 2,34 | 1,92E+07 | 2,23E+07 |
| 17,1 | | 0,0032115 | 0,0048128 | 1,24 | 9,12E+07 | 8,37E+07 |
| 30,5 | | 0,0032400 | 0,0048831 | 0,88 | 4,23E+08 | 4,38E+08 |
| 18,9 | | 0,0032407 | 0,0048802 | 0,05 | 6,26E+07 | 5,10E+07 |
| 60 | | 0,0032673 | 0,0048774 | 1,49 | 1,31E+07 | 1,35E+07 |

| | | | | | | |
|---|---|---|---|---|---|---|
| 33,9 | | 0,0032766 | 0,0048745 | 1,42 | 1,14E+08 | 1,17E+08 |
| 7,5 | | 0,0032868 | 0,0048716 | 0,50 | 9,08E+06 | 1,06E+07 |
| 13,1 | | 0,0032963 | 0,0049441 | 0,26 | 1,33E+07 | 1,47E+07 |
| 42,6 | | 0,0033220 | 0,0050047 | 0,03 | 2,07E+08 | 2,38E+08 |
| 26,4 | | 0,0033238 | 0,0050018 | 1,90 | 2,52E+07 | 1,92E+07 |
| 16,6 | | 0,0033285 | 0,0049988 | 1,56 | 3,00E+07 | 2,51E+07 |
| 57,1 | | 0,0033371 | 0,0049959 | 0,87 | 3,16E+09 | 3,13E+09 |
| 19,2 | | 0,0033379 | 0,0049930 | 0,61 | 7,17E+07 | 8,29E+07 |
| 65,4 | | 0,0033444 | 0,0049900 | 1,31 | 1,35E+08 | 1,27E+08 |
| 54,4 | | 0,0033455 | 0,0049871 | 1,55 | 6,59E+08 | 6,48E+08 |
| 45 | | 0,0033520 | 0,0049842 | 0,79 | 2,90E+09 | 3,17E+09 |
| 51,5 | | 0,0033552 | 0,0049813 | 1,57 | 1,05E+08 | 1,21E+08 |
| 42,3 | | 0,0033729 | 0,0049783 | 0,69 | 1,18E+08 | 1,31E+08 |
| 4 | + | 0,0033764 | 0,0049754 | 7,57 | 3,44E+06 | 1,02E+07 |
| 11,1 | | 0,0033785 | 0,0049725 | 0,58 | 3,64E+07 | 3,23E+07 |
| 41,9 | | 0,0033857 | 0,0049696 | 2,00 | 2,48E+08 | 2,16E+08 |
| 44,9 | | 0,0033901 | 0,0049667 | 1,67 | 1,30E+08 | 9,91E+07 |
| 6,9 | | 0,0033917 | 0,0049638 | 0,67 | 7,06E+07 | 6,60E+07 |
| 48,7 | | 0,0033941 | 0,0049609 | 1,82 | 6,42E+07 | 7,19E+07 |
| 30,2 | | 0,0033986 | 0,0049580 | 0,53 | 9,74E+07 | 1,25E+08 |
| 26,5 | | 0,0033999 | 0,0049552 | 0,04 | 1,77E+08 | 1,50E+08 |
| 7,7 | | 0,0034049 | 0,0049523 | 0,39 | 1,02E+07 | 6,21E+06 |
| 24,5 | | 0,0034183 | 0,0049494 | 0,57 | 2,67E+07 | 2,59E+07 |
| 15,3 | | 0,0034184 | 0,0049465 | 0,90 | 5,33E+07 | 5,20E+07 |
| 19,4 | | 0,0034229 | 0,0049436 | 1,10 | 2,79E+07 | 2,85E+07 |
| 42,6 | | 0,0034423 | 0,0049408 | 0,77 | 2,73E+08 | 2,88E+08 |
| 36,9 | | 0,0034458 | 0,0049379 | 0,88 | 1,38E+09 | 1,35E+09 |
| 42,3 | | 0,0034478 | 0,0049350 | 10,45 | 3,17E+06 | 3,44E+06 |
| 10,8 | | 0,0034506 | 0,0049322 | 0,12 | 1,25E+07 | 1,65E+07 |
| 22,5 | | 0,0034548 | 0,0049293 | 0,71 | 3,59E+08 | 3,45E+08 |
| 31,9 | | 0,0034646 | 0,0049890 | 0,68 | 3,72E+08 | 3,36E+08 |
| 62,7 | | 0,0034673 | 0,0049861 | 0,69 | 2,95E+08 | 2,98E+08 |
| 26,8 | | 0,0034681 | 0,0049832 | 1,51 | 7,29E+07 | 7,21E+07 |
| 76 | | 0,0034739 | 0,0049803 | 0,70 | 2,58E+08 | 2,56E+08 |
| 12,6 | | 0,0034880 | 0,0051046 | 0,25 | 1,53E+07 | 1,46E+07 |
| 53,5 | | 0,0035102 | 0,0051016 | 2,34 | 4,69E+08 | 6,91E+08 |
| 38,1 | | 0,0035103 | 0,0050987 | 1,24 | 1,17E+08 | 1,24E+08 |
| 48,2 | | 0,0035209 | 0,0050957 | 1,62 | 7,48E+08 | 8,00E+08 |
| 16,3 | | 0,0035218 | 0,0050928 | 0,35 | 2,76E+07 | 2,11E+07 |
| 16,5 | | 0,0035229 | 0,0050899 | 2,18 | 1,63E+07 | 2,27E+07 |
| 29,7 | | 0,0035304 | 0,0050869 | 0,10 | 2,05E+07 | 2,57E+07 |
| 13,5 | | 0,0035361 | 0,0050840 | 0,21 | 2,16E+07 | 1,51E+07 |
| 41,8 | | 0,0035371 | 0,0050811 | 0,66 | 1,73E+08 | 1,62E+08 |
| 46,1 | | 0,0035422 | 0,0050782 | 0,64 | 8,46E+08 | 8,39E+08 |
| 37,5 | | 0,0035483 | 0,0050752 | 0,73 | 7,15E+07 | 7,73E+07 |
| 21,3 | | 0,0035507 | 0,0050723 | 12,74 | 3,87E+06 | 1,45E+07 |
| 62,1 | | 0,0035514 | 0,0050694 | 1,65 | 1,54E+09 | 1,52E+09 |
| 13,4 | | 0,0035544 | 0,0050665 | 0,69 | 8,66E+07 | 7,53E+07 |
| 46,5 | | 0,0035560 | 0,0050636 | 1,33 | 1,80E+08 | 1,71E+08 |
| 35,4 | | 0,0035726 | 0,0050607 | 0,58 | 3,39E+08 | 4,33E+08 |

| | | | | | | |
|---|---|---|---|---|---|---|
| 25,9 | | 0,0035773 | 0,0050578 | 0,60 | 1,39E+08 | 1,66E+08 |
| 10,8 | | 0,0035912 | 0,0050549 | 0,61 | 4,40E+07 | 4,16E+07 |
| 28,2 | | 0,0035934 | 0,0050520 | 0,61 | 4,07E+07 | 3,86E+07 |
| 41,4 | | 0,0036153 | 0,0050491 | 1,24 | 6,53E+07 | 7,17E+07 |
| 46,6 | | 0,0036205 | 0,0051171 | 1,57 | 8,29E+07 | 8,24E+07 |
| 22,2 | | 0,0036296 | 0,0051142 | 0,02 | 1,28E+08 | 1,12E+08 |
| 28,4 | | 0,0036313 | 0,0051112 | 2,07 | 7,49E+07 | 1,02E+08 |
| 8,6 | | 0,0036320 | 0,0051083 | 1,48 | 2,69E+06 | 2,80E+06 |
| 24,6 | | 0,0036339 | 0,0051054 | 0,30 | 6,85E+07 | 6,47E+07 |
| 9,3 | | 0,0036357 | 0,0051025 | 0,13 | 3,27E+07 | 3,77E+07 |
| 41,5 | | 0,0036372 | 0,0050996 | 0,67 | 4,35E+07 | 3,60E+07 |
| 23,7 | | 0,0036392 | 0,0051581 | 0,81 | 2,41E+08 | 2,67E+08 |
| 11,1 | | 0,0036406 | 0,0051552 | 0,10 | 1,67E+07 | 2,81E+07 |
| 39,1 | | 0,0036514 | 0,0051523 | 1,32 | 3,47E+08 | 3,54E+08 |
| 79,1 | | 0,0036589 | 0,0051493 | 1,29 | 4,17E+09 | 4,05E+09 |
| 12,8 | | 0,0036607 | 0,0051464 | 0,15 | 2,53E+07 | 3,49E+07 |
| 23,3 | | 0,0036660 | 0,0052138 | 0,46 | 8,02E+06 | 6,90E+06 |
| 14,9 | | 0,0036818 | 0,0052109 | 1,39 | 7,37E+07 | 6,97E+07 |
| 13,8 | | 0,0036948 | 0,0052782 | 3,71 | 3,26E+06 | 5,89E+06 |
| 11,6 | | 0,0036960 | 0,0052752 | 0,12 | 2,16E+07 | 2,64E+07 |
| 23,3 | | 0,0036981 | 0,0052722 | 0,50 | 7,38E+08 | 9,63E+08 |
| 13,4 | | 0,0037088 | 0,0052692 | 1,62 | 2,93E+07 | 3,05E+07 |
| 43,1 | | 0,0037158 | 0,0052663 | 1,28 | 7,45E+08 | 6,91E+08 |
| 15,9 | | 0,0037173 | 0,0052633 | 0,60 | 3,46E+07 | 3,75E+07 |
| 73,8 | | 0,0037331 | 0,0053145 | 0,76 | 1,32E+10 | 1,50E+10 |
| 17,6 | | 0,0037423 | 0,0053115 | 0,11 | 1,69E+07 | 3,24E+07 |
| 3,2 | | 0,0037464 | 0,0053085 | 0,18 | 1,82E+07 | 2,61E+07 |
| 6,1 | | 0,0037620 | 0,0053596 | 0,31 | 6,48E+06 | 7,93E+06 |
| 29,3 | | 0,0037703 | 0,0054175 | 3,00 | 8,06E+06 | 5,75E+06 |
| 65,7 | | 0,0037731 | 0,0054144 | 1,49 | 4,20E+08 | 5,14E+08 |
| 25,9 | | 0,0037753 | 0,0054114 | 6,36 | 4,32E+06 | 1,68E+06 |
| 20,5 | | 0,0037811 | 0,0054083 | 0,36 | 7,86E+06 | 1,23E+07 |
| 18,1 | | 0,0037942 | 0,0054053 | 0,66 | 7,61E+07 | 7,43E+07 |
| 48,4 | | 0,0038033 | 0,0054022 | 1,96 | 2,49E+07 | 2,94E+07 |
| 23 | | 0,0038053 | 0,0053992 | 0,77 | 5,18E+07 | 5,67E+07 |
| 43,6 | | 0,0038198 | 0,0053962 | 1,12 | 3,82E+08 | 3,69E+08 |
| 15,9 | | 0,0038230 | 0,0053932 | 0,11 | 5,05E+07 | 3,46E+07 |
| 15,7 | | 0,0038273 | 0,0053901 | 1,97 | 6,08E+07 | 4,75E+07 |
| 32,6 | | 0,0038284 | 0,0053871 | 0,82 | 1,03E+08 | 9,67E+07 |
| 62,2 | | 0,0038298 | 0,0053841 | 1,52 | 3,79E+08 | 3,74E+08 |
| 28,6 | | 0,0038375 | 0,0053811 | 0,71 | 2,04E+08 | 1,96E+08 |
| 37,2 | | 0,0038394 | 0,0053781 | 0,77 | 4,16E+08 | 4,43E+08 |
| 8,3 | | 0,0038404 | 0,0053751 | 3,47 | 8,36E+07 | 1,45E+08 |
| 27,9 | | 0,0038440 | 0,0053721 | 1,34 | 1,78E+08 | 1,70E+08 |
| 15,6 | | 0,0038467 | 0,0053691 | 0,07 | 4,17E+07 | 3,32E+07 |
| 30,8 | | 0,0038582 | 0,0053661 | 2,33 | 2,24E+07 | 1,55E+07 |
| 38,9 | | 0,0038603 | 0,0053631 | 0,74 | 3,21E+08 | 3,63E+08 |
| 48,5 | | 0,0038620 | 0,0053601 | 0,89 | 2,84E+08 | 2,72E+08 |
| 27,4 | | 0,0038640 | 0,0054173 | 0,61 | 1,85E+08 | 1,54E+08 |
| 13,1 | | 0,0038786 | 0,0054143 | 0,14 | 5,47E+07 | 4,66E+07 |

| | | | | | | |
|---|---|---|---|---|---|---|
| 64,7 | | 0,0038786 | 0,0054112 | 1,49 | 1,63E+10 | 1,41E+10 |
| 16,3 | | 0,0038821 | 0,0054082 | 0,05 | 5,49E+08 | 7,81E+08 |
| 34,2 | | 0,0038874 | 0,0054052 | 0,62 | 1,04E+09 | 1,05E+09 |
| 20,3 | | 0,0039013 | 0,0054022 | 0,41 | 5,61E+07 | 3,51E+07 |
| 8,8 | | 0,0039379 | 0,0055125 | 1,50 | 1,00E+07 | 1,10E+07 |
| 27,3 | | 0,0039463 | 0,0055094 | 0,09 | 4,38E+07 | 4,13E+07 |
| 25,1 | | 0,0039488 | 0,0055064 | 0,83 | 1,83E+08 | 1,90E+08 |
| 29,9 | | 0,0039670 | 0,0055033 | 1,47 | 8,94E+07 | 9,19E+07 |
| 23 | | 0,0039725 | 0,0055003 | 1,29 | 1,08E+08 | 1,17E+08 |
| 20,6 | | 0,0039796 | 0,0054972 | 2,90 | 4,13E+06 | 2,50E+06 |
| 18,2 | | 0,0039992 | 0,0054942 | 0,17 | 1,37E+07 | 1,77E+07 |
| 28,3 | | 0,0040007 | 0,0054912 | 0,79 | 5,45E+07 | 5,54E+07 |
| 44,8 | | 0,0040034 | 0,0054881 | 1,36 | 8,87E+07 | 8,67E+07 |
| 35,2 | | 0,0040038 | 0,0054851 | 0,81 | 1,99E+08 | 2,14E+08 |
| 47,6 | | 0,0040039 | 0,0054821 | 1,48 | 4,80E+09 | 4,66E+09 |
| 27,4 | | 0,0040046 | 0,0054790 | 40,79 | 1,10E+07 | 2,51E+07 |
| 49 | | 0,0040090 | 0,0054760 | 1,38 | 2,39E+09 | 2,51E+09 |
| 12,6 | | 0,0040146 | 0,0054730 | 0,24 | 1,89E+07 | 1,67E+07 |
| 22,3 | | 0,0040199 | 0,0054700 | 0,59 | 1,31E+08 | 1,49E+08 |
| 43,5 | | 0,0040252 | 0,0054670 | 0,83 | 7,66E+07 | 7,74E+07 |
| 35,5 | | 0,0040480 | 0,0054640 | 1,24 | 5,73E+08 | 5,26E+08 |
| 23,2 | | 0,0040530 | 0,0054609 | 0,23 | 3,45E+07 | 3,70E+07 |
| 37,9 | | 0,0040585 | 0,0054579 | 61,07 | 3,50E+06 | 1,83E+06 |
| 78,6 | | 0,0040587 | 0,0054549 | 1,83 | 1,75E+09 | 2,37E+09 |
| 32,3 | | 0,0040641 | 0,0054519 | 1,24 | 2,98E+08 | 3,04E+08 |
| 53,4 | | 0,0040739 | 0,0054490 | 0,56 | 2,38E+08 | 3,24E+08 |
| 8,5 | | 0,0040875 | 0,0054460 | 1,42 | 1,98E+07 | 1,89E+07 |
| 16,1 | | 0,0040926 | 0,0054430 | 0,61 | 1,75E+08 | 1,68E+08 |
| 11,5 | | 0,0040958 | 0,0054400 | 7,06 | 2,40E+06 | 1,56E+06 |
| 41 | | 0,0041021 | 0,0054962 | 0,65 | 2,46E+08 | 2,67E+08 |
| 52 | | 0,0041062 | 0,0054932 | 0,77 | 2,19E+09 | 2,35E+09 |
| 10,4 | | 0,0041093 | 0,0054902 | 1,53 | 1,63E+07 | 1,87E+07 |
| 29,9 | | 0,0041103 | 0,0054872 | 0,69 | 4,22E+07 | 5,05E+07 |
| 23 | | 0,0041107 | 0,0054842 | 1,95 | 1,67E+07 | 1,61E+07 |
| 12,9 | | 0,0041283 | 0,0055991 | 0,64 | 2,77E+07 | 2,95E+07 |
| 39,5 | | 0,0041351 | 0,0055961 | 1,71 | 3,88E+07 | 3,23E+07 |
| 88,2 | | 0,0041451 | 0,0056519 | 1,88 | 3,95E+08 | 3,54E+08 |
| 30 | | 0,0041478 | 0,0056489 | 5,52 | 2,25E+06 | 4,99E+06 |
| 14,1 | | 0,0041563 | 0,0056458 | 3,53 | 1,32E+07 | 2,54E+07 |
| 45,7 | | 0,0041585 | 0,0056427 | 36,15 | 1,72E+06 | 5,81E+06 |
| 17,2 | | 0,0041636 | 0,0056396 | 0,42 | 1,90E+07 | 1,74E+07 |
| 7,3 | | 0,0041649 | 0,0056366 | 2,15 | 4,33E+06 | 4,78E+06 |
| 42,3 | | 0,0041718 | 0,0056335 | 1,20 | 2,79E+08 | 3,04E+08 |
| 39,2 | | 0,0041799 | 0,0056891 | 1,58 | 2,99E+08 | 3,26E+08 |
| 18,7 | | 0,0041926 | 0,0057382 | 0,71 | 1,03E+08 | 1,18E+08 |
| 32,5 | | 0,0042035 | 0,0057351 | 0,85 | 7,24E+07 | 7,86E+07 |
| 18,9 | | 0,0042116 | 0,0058014 | 0,60 | 2,41E+07 | 2,70E+07 |
| 15,5 | | 0,0042187 | 0,0057983 | 2,58 | 4,02E+06 | 6,71E+06 |
| 59,6 | | 0,0042208 | 0,0057951 | 0,62 | 6,64E+08 | 5,44E+08 |
| 32,6 | | 0,0042236 | 0,0057920 | 0,47 | 1,65E+08 | 2,09E+08 |

| | | | | | | |
|---|---|---|---|---|---|---|
| 30,7 | | 0,0042326 | 0,0057888 | 0,39 | 1,25E+07 | 1,03E+07 |
| 47,7 | | 0,0042353 | 0,0057857 | 0,82 | 1,91E+08 | 2,12E+08 |
| 50,5 | | 0,0042357 | 0,0057826 | 2,33 | 7,61E+07 | 5,02E+07 |
| 65,7 | | 0,0042376 | 0,0057795 | 1,45 | 3,70E+09 | 4,11E+09 |
| 13,8 | | 0,0042395 | 0,0057763 | 1,63 | 6,49E+06 | 6,95E+06 |
| 46,5 | | 0,0042576 | 0,0057732 | 1,14 | 3,45E+08 | 3,39E+08 |
| 22,5 | | 0,0042583 | 0,0057701 | 0,56 | 6,11E+07 | 5,81E+07 |
| 57,9 | | 0,0042586 | 0,0057670 | 0,72 | 5,36E+09 | 4,93E+09 |
| 54,7 | | 0,0042645 | 0,0057639 | 1,48 | 2,05E+08 | 2,14E+08 |
| 16,1 | | 0,0042754 | 0,0057608 | 14,70 | 4,18E+06 | 1,82E+06 |
| 6,4 | | 0,0042779 | 0,0057577 | 0,78 | 1,61E+07 | 1,70E+07 |
| 34,6 | | 0,0042839 | 0,0057546 | 0,91 | 7,42E+08 | 7,43E+08 |
| 17,3 | | 0,0042912 | 0,0057515 | 0,18 | 4,71E+07 | 7,11E+07 |
| 10,6 | | 0,0042943 | 0,0057484 | 0,12 | 1,52E+07 | 2,40E+07 |
| 22,5 | | 0,0042958 | 0,0057453 | 19,19 | 2,00E+06 | 8,69E+05 |
| 32,8 | | 0,0043011 | 0,0057422 | 1,31 | 2,28E+08 | 2,20E+08 |
| 47,5 | | 0,0043026 | 0,0057391 | 1,18 | 5,86E+08 | 6,25E+08 |
| 33 | | 0,0043118 | 0,0057361 | 0,03 | 1,51E+08 | 1,91E+08 |
| 46 | | 0,0043138 | 0,0057330 | 3,18 | 2,38E+07 | 2,51E+07 |
| 9,8 | | 0,0043161 | 0,0057299 | 0,65 | 3,43E+07 | 3,53E+07 |
| 16 | | 0,0043208 | 0,0057268 | 0,11 | 3,38E+07 | 6,17E+07 |
| 47,9 | | 0,0043218 | 0,0057238 | 0,74 | 1,11E+09 | 1,19E+09 |
| 44,9 | | 0,0043448 | 0,0057207 | 0,81 | 1,39E+08 | 1,52E+08 |
| 21,7 | | 0,0043477 | 0,0057176 | 0,41 | 3,34E+07 | 5,40E+07 |
| 44,3 | | 0,0043544 | 0,0057146 | 0,75 | 4,09E+08 | 4,48E+08 |
| 21,3 | | 0,0043597 | 0,0057115 | 1,30 | 4,03E+08 | 3,71E+08 |
| 41,3 | | 0,0043602 | 0,0057085 | 0,60 | 4,37E+09 | 3,32E+09 |
| 15,5 | | 0,0043788 | 0,0057054 | 1,57 | 5,95E+06 | 6,54E+06 |
| 62 | | 0,0043805 | 0,0057024 | 0,87 | 2,15E+09 | 2,06E+09 |
| 3,9 | | 0,0043836 | 0,0056994 | 0,51 | 1,00E+07 | 1,25E+07 |
| 19,1 | | 0,0043837 | 0,0056963 | 0,46 | 2,99E+07 | 3,42E+07 |
| 21 | | 0,0043859 | 0,0056933 | 0,29 | 1,44E+07 | 1,47E+07 |
| 27,5 | | 0,0043894 | 0,0056903 | 4,32 | 8,00E+06 | 1,42E+07 |
| 47,8 | | 0,0043896 | 0,0056872 | 2,14 | 7,08E+07 | 4,86E+07 |
| 18 | | 0,0043932 | 0,0056842 | 1,76 | 7,72E+06 | 1,01E+07 |
| 44,2 | | 0,0044089 | 0,0057322 | 0,03 | 1,21E+08 | 4,87E+07 |
| 8,8 | | 0,0044140 | 0,0057292 | 0,18 | 2,63E+07 | 3,46E+07 |
| 36,4 | | 0,0044141 | 0,0057261 | 0,64 | 1,50E+09 | 1,18E+09 |
| 8,9 | | 0,0044196 | 0,0057231 | 7,96 | 4,43E+06 | 2,67E+06 |
| 25,7 | | 0,0044221 | 0,0057200 | 4,75 | 3,85E+07 | 1,60E+07 |
| 27,4 | | 0,0044247 | 0,0057170 | 1,41 | 2,05E+08 | 2,32E+08 |
| 20,4 | | 0,0044293 | 0,0057140 | 1,41 | 3,82E+07 | 3,58E+07 |
| 17,6 | | 0,0044509 | 0,0057110 | 0,55 | 2,19E+08 | 1,65E+08 |
| 28 | | 0,0044549 | 0,0057079 | 0,68 | 1,09E+08 | 8,98E+07 |
| 16,2 | | 0,0044586 | 0,0057049 | 0,40 | 4,93E+07 | 4,81E+07 |
| 18,8 | | 0,0044635 | 0,0057019 | 28,06 | 8,87E+06 | 3,44E+06 |
| 29,5 | | 0,0044985 | 0,0056989 | 0,56 | 2,60E+07 | 2,32E+07 |
| 21,3 | | 0,0045121 | 0,0056959 | 0,21 | 1,83E+07 | 2,08E+07 |
| 27,7 | | 0,0045298 | 0,0056929 | 0,78 | 2,09E+08 | 2,05E+08 |
| 36,8 | | 0,0045364 | 0,0056899 | 0,71 | 5,63E+07 | 6,51E+07 |

| | | | | | | |
|---|---|---|---|---|---|---|
| 12,9 | | 0,0045758 | 0,0057438 | 0,64 | 6,21E+07 | 5,52E+07 |
| 50,5 | | 0,0045760 | 0,0057408 | 1,30 | 1,80E+09 | 1,73E+09 |
| 60,6 | | 0,0045773 | 0,0057378 | 1,34 | 4,54E+08 | 5,22E+08 |
| 37,5 | | 0,0045899 | 0,0058000 | 0,77 | 2,37E+08 | 2,55E+08 |
| 32,8 | | 0,0045988 | 0,0057969 | 0,83 | 5,50E+08 | 5,91E+08 |
| 33,1 | | 0,0046075 | 0,0057939 | 0,73 | 8,39E+07 | 7,29E+07 |
| 46,2 | | 0,0046201 | 0,0057909 | 1,23 | 3,53E+07 | 3,27E+07 |
| 16,8 | | 0,0046269 | 0,0057878 | 0,60 | 1,54E+07 | 1,43E+07 |
| 59 | | 0,0046293 | 0,0057848 | 1,36 | 1,39E+09 | 1,40E+09 |
| 50,8 | | 0,0046353 | 0,0057817 | 1,43 | 1,94E+08 | 2,00E+08 |
| 41,9 | | 0,0046458 | 0,0057787 | 0,65 | 2,87E+08 | 2,38E+08 |
| 19,9 | | 0,0046491 | 0,0057757 | 0,21 | 7,21E+06 | 5,28E+06 |
| 44,4 | | 0,0046499 | 0,0057727 | 1,33 | 2,48E+08 | 2,38E+08 |
| 5,4 | + | 0,0046541 | 0,0057696 | 0,13 | 3,37E+07 | 1,75E+07 |
| 11,4 | | 0,0046577 | 0,0057666 | 0,44 | 3,23E+07 | 3,52E+07 |
| 19,2 | | 0,0046674 | 0,0057636 | 0,75 | 1,03E+09 | 1,17E+09 |
| 15,6 | | 0,0046822 | 0,0058819 | 3,36 | 4,90E+06 | 4,82E+06 |
| 28,4 | | 0,0046830 | 0,0058788 | 6,04 | 3,51E+07 | 3,00E+07 |
| 6 | | 0,0046831 | 0,0058757 | 0,15 | 1,46E+07 | 1,38E+07 |
| 23,9 | | 0,0046887 | 0,0058727 | 0,46 | 4,16E+07 | 4,51E+07 |
| 63,9 | | 0,0046893 | 0,0058696 | 0,77 | 1,62E+08 | 1,62E+08 |
| 22,3 | | 0,0047074 | 0,0059312 | 0,81 | 1,19E+08 | 1,23E+08 |
| 55,1 | | 0,0047335 | 0,0061136 | 1,61 | 1,76E+08 | 2,00E+08 |
| 49,4 | | 0,0047378 | 0,0061104 | 1,35 | 3,37E+08 | 3,27E+08 |
| 67,3 | | 0,0047659 | 0,0062884 | 0,85 | 9,26E+08 | 9,51E+08 |
| 12,8 | | 0,0047668 | 0,0062851 | 0,50 | 3,19E+07 | 2,94E+07 |
| 77,7 | | 0,0047699 | 0,0062819 | 0,45 | 5,75E+09 | 8,00E+09 |
| 20,8 | | 0,0047821 | 0,0062786 | 1,69 | 3,12E+07 | 2,71E+07 |
| 62,3 | | 0,0047907 | 0,0062753 | 1,28 | 2,75E+08 | 2,81E+08 |
| 48,5 | | 0,0047982 | 0,0062721 | 0,72 | 6,65E+08 | 7,75E+08 |
| 8,5 | | 0,0047985 | 0,0062688 | 0,12 | 1,67E+07 | 1,36E+07 |
| 77,5 | | 0,0048060 | 0,0063154 | 1,86 | 2,18E+08 | 2,49E+08 |
| 22,6 | | 0,0048068 | 0,0063121 | 0,76 | 4,32E+07 | 4,15E+07 |
| 12,9 | | 0,0048163 | 0,0063088 | 0,09 | 2,63E+07 | 2,63E+07 |
| 41,4 | | 0,0048190 | 0,0063055 | 1,43 | 1,22E+08 | 1,05E+08 |
| 48,3 | | 0,0048547 | 0,0063520 | 1,30 | 5,55E+08 | 5,95E+08 |
| 22,2 | | 0,0048576 | 0,0063487 | 1,36 | 6,50E+07 | 6,71E+07 |
| 9,2 | | 0,0048649 | 0,0063454 | 14,91 | 2,20E+06 | 1,07E+07 |
| 26,8 | | 0,0048677 | 0,0063421 | 0,64 | 1,70E+08 | 1,33E+08 |
| 4,8 | | 0,0048687 | 0,0063388 | 0,53 | 1,88E+07 | 1,36E+07 |
| 31,3 | | 0,0048883 | 0,0063356 | 1,18 | 3,39E+08 | 3,18E+08 |
| 41,4 | | 0,0049034 | 0,0063323 | 1,40 | 6,64E+08 | 5,61E+08 |
| 9,3 | | 0,0049036 | 0,0063290 | 0,20 | 1,14E+07 | 1,14E+07 |
| 20,3 | | 0,0049116 | 0,0063258 | 2,01 | 4,04E+07 | 4,83E+07 |
| 3,3 | | 0,0049246 | 0,0063225 | 0,28 | 1,12E+07 | 8,25E+06 |
| 53,6 | | 0,0049308 | 0,0063193 | 0,87 | 1,09E+10 | 1,11E+10 |
| 12,4 | | 0,0049546 | 0,0063634 | 1,62 | 9,28E+06 | 1,10E+07 |
| 49,1 | | 0,0049609 | 0,0063601 | 0,79 | 3,62E+08 | 3,85E+08 |
| 43,6 | | 0,0049887 | 0,0064062 | 1,26 | 8,78E+08 | 9,11E+08 |
| 34,8 | | 0,0050000 | 0,0064666 | 1,85 | 1,26E+08 | 1,80E+08 |

| | | | | | | |
|---|---|---|---|---|---|---|
| 46,7 | | 0,0050023 | 0,0064633 | 1,36 | 5,05E+08 | 4,89E+08 |
| 60 | | 0,0050060 | 0,0064600 | 1,39 | 6,89E+08 | 5,83E+08 |
| 30 | | 0,0050060 | 0,0064566 | 0,51 | 1,23E+07 | 1,26E+07 |
| 12,1 | | 0,0050094 | 0,0064533 | 1,57 | 3,73E+07 | 2,96E+07 |
| 51,2 | | 0,0050164 | 0,0064500 | 0,67 | 2,68E+08 | 2,94E+08 |
| 20,9 | | 0,0050308 | 0,0064467 | 1,31 | 3,27E+08 | 3,25E+08 |
| 48,7 | | 0,0050326 | 0,0064434 | 0,81 | 8,85E+08 | 9,66E+08 |
| 9,3 | | 0,0050352 | 0,0064401 | 0,07 | 1,88E+07 | 2,47E+07 |
| 42,5 | | 0,0050399 | 0,0064368 | 0,73 | 4,49E+08 | 4,50E+08 |
| 28,3 | | 0,0050455 | 0,0064826 | 0,69 | 1,65E+08 | 1,72E+08 |
| 44,4 | | 0,0050500 | 0,0064793 | 0,43 | 1,10E+08 | 8,71E+07 |
| 11,8 | | 0,0050593 | 0,0065393 | 0,41 | 1,62E+07 | 1,59E+07 |
| 27,9 | | 0,0050672 | 0,0066013 | 0,67 | 1,23E+08 | 1,38E+08 |
| 45 | | 0,0050672 | 0,0065980 | 0,86 | 5,15E+09 | 5,17E+09 |
| 38 | | 0,0050763 | 0,0065946 | 0,03 | 3,42E+08 | 5,76E+07 |
| 13,9 | | 0,0050801 | 0,0065912 | 0,83 | 3,20E+07 | 3,47E+07 |
| 29,7 | | 0,0050806 | 0,0065879 | 0,80 | 4,18E+08 | 4,01E+08 |
| 38,7 | | 0,0051019 | 0,0065845 | 0,73 | 6,81E+07 | 6,32E+07 |
| 8,3 | | 0,0051028 | 0,0065812 | 0,65 | 2,61E+07 | 2,21E+07 |
| 7,3 | | 0,0051119 | 0,0065778 | 9,15 | 4,32E+06 | 1,58E+06 |
| 7,6 | | 0,0051161 | 0,0065745 | 0,64 | 2,75E+08 | 2,50E+08 |
| 16,1 | | 0,0051198 | 0,0065711 | 0,09 | 4,66E+07 | 5,48E+07 |
| 33,7 | | 0,0051204 | 0,0065678 | 0,66 | 9,87E+07 | 9,38E+07 |
| 14,4 | | 0,0051218 | 0,0065645 | 9,84 | 3,47E+06 | 1,04E+07 |
| 43,4 | | 0,0051283 | 0,0066261 | 0,17 | 4,48E+07 | 6,74E+07 |
| 54,9 | | 0,0051306 | 0,0066227 | 1,12 | 2,33E+08 | 2,47E+08 |
| 18,4 | | 0,0051339 | 0,0066194 | 0,69 | 9,83E+07 | 1,23E+08 |
| 41,6 | | 0,0051351 | 0,0066160 | 1,57 | 6,83E+07 | 8,13E+07 |
| 43,3 | | 0,0051379 | 0,0066127 | 1,35 | 9,11E+08 | 1,03E+09 |
| 9,5 | | 0,0051383 | 0,0066093 | 0,22 | 1,65E+07 | 1,90E+07 |
| 5,6 | | 0,0051395 | 0,0066545 | 0,45 | 1,37E+07 | 1,20E+07 |
| 45 | | 0,0051405 | 0,0066512 | 1,35 | 3,34E+08 | 3,30E+08 |
| 36,3 | | 0,0051501 | 0,0066478 | 0,53 | 7,47E+07 | 7,91E+07 |
| 50,3 | | 0,0051509 | 0,0066444 | 0,66 | 2,12E+09 | 2,60E+09 |
| 11,8 | | 0,0051599 | 0,0066875 | 2,46 | 3,94E+06 | 4,97E+06 |
| 15,4 | | 0,0051608 | 0,0066842 | 3,06 | 8,03E+06 | 1,48E+07 |
| 7,3 | | 0,0051897 | 0,0067352 | 3,42 | 2,99E+06 | 2,70E+06 |
| 36,3 | | 0,0052102 | 0,0067319 | 1,40 | 1,13E+08 | 9,33E+07 |
| 10,3 | | 0,0052122 | 0,0067285 | 2,73 | 3,76E+06 | 3,33E+06 |
| 37,4 | | 0,0052136 | 0,0067251 | 0,71 | 2,26E+08 | 2,51E+08 |
| 16,8 | | 0,0052203 | 0,0067217 | 1,67 | 1,27E+08 | 1,26E+08 |
| 51,7 | | 0,0052258 | 0,0067183 | 2,46 | 1,07E+07 | 1,37E+07 |
| 26,5 | | 0,0052328 | 0,0067149 | 1,32 | 3,35E+08 | 3,34E+08 |
| 55,4 | | 0,0052334 | 0,0067116 | 1,74 | 3,69E+08 | 3,24E+08 |
| 23,8 | | 0,0052337 | 0,0067082 | 0,39 | 1,86E+07 | 2,32E+07 |
| 48,4 | | 0,0052344 | 0,0067048 | 0,59 | 6,95E+08 | 5,63E+08 |
| 36,8 | | 0,0052380 | 0,0067015 | 1,56 | 2,43E+08 | 1,85E+08 |
| 30 | | 0,0052553 | 0,0066981 | 2,29 | 1,27E+07 | 1,49E+07 |
| 37,9 | | 0,0052619 | 0,0066947 | 1,29 | 8,75E+08 | 9,58E+08 |
| 9,2 | | 0,0052763 | 0,0066914 | 0,52 | 6,46E+06 | 6,57E+06 |

| | | | | | | |
|---|---|---|---|---|---|---|
| 47,2 | | 0,0052928 | 0,0067521 | 0,87 | 1,19E+08 | 1,13E+08 |
| 57,1 | | 0,0052956 | 0,0067487 | 0,63 | 6,77E+08 | 8,98E+08 |
| 28 | | 0,0053083 | 0,0068074 | 0,83 | 4,30E+08 | 4,24E+08 |
| 33,1 | | 0,0053104 | 0,0068040 | 1,79 | 4,82E+07 | 3,51E+07 |
| 44 | | 0,0053140 | 0,0068006 | 1,17 | 7,71E+07 | 7,43E+07 |
| 31,4 | | 0,0053144 | 0,0067972 | 1,45 | 1,10E+08 | 1,20E+08 |
| 59,1 | | 0,0053152 | 0,0067938 | 1,89 | 5,52E+08 | 8,00E+08 |
| 14,6 | | 0,0053356 | 0,0067904 | 1,52 | 7,75E+07 | 6,87E+07 |
| 17,8 | + | 0,0053525 | 0,0069566 | 5,40 | 3,16E+06 | 7,71E+06 |
| 31,4 | | 0,0053539 | 0,0069531 | 1,76 | 3,50E+07 | 4,69E+07 |
| 10,8 | | 0,0053544 | 0,0069497 | 1,23 | 2,27E+07 | 2,27E+07 |
| 45,5 | | 0,0053666 | 0,0069462 | 1,36 | 1,34E+09 | 1,34E+09 |
| 15,2 | | 0,0053737 | 0,0069428 | 0,66 | 3,98E+07 | 4,24E+07 |
| 47,9 | | 0,0053842 | 0,0069393 | 0,84 | 2,55E+08 | 2,77E+08 |
| 6,9 | | 0,0053943 | 0,0069359 | 2,74 | 2,70E+06 | 3,70E+06 |
| 43,4 | | 0,0053976 | 0,0069324 | 1,51 | 3,37E+08 | 2,73E+08 |
| 15,8 | | 0,0053977 | 0,0069290 | 1,38 | 7,96E+07 | 7,36E+07 |
| 4,6 | | 0,0054007 | 0,0069255 | 0,76 | 2,08E+07 | 1,96E+07 |
| 30,1 | | 0,0054010 | 0,0069221 | 1,48 | 7,20E+07 | 9,08E+07 |
| 24,2 | | 0,0054066 | 0,0069643 | 1,20 | 1,01E+08 | 9,98E+07 |
| 42,2 | | 0,0054067 | 0,0069608 | 1,23 | 3,90E+08 | 4,34E+08 |
| 39,8 | | 0,0054110 | 0,0069574 | 1,49 | 9,70E+07 | 7,88E+07 |
| 33,2 | | 0,0054180 | 0,0069539 | 4,91 | 7,77E+06 | 3,39E+06 |
| 26,5 | | 0,0054494 | 0,0069505 | 1,82 | 2,63E+07 | 1,89E+07 |
| 31 | | 0,0054502 | 0,0069471 | 1,18 | 2,41E+08 | 2,55E+08 |
| 81,4 | | 0,0054540 | 0,0069436 | 1,23 | 2,52E+10 | 2,55E+10 |
| 11,6 | | 0,0054603 | 0,0069402 | 0,30 | 1,27E+07 | 7,24E+06 |
| 18,7 | | 0,0054641 | 0,0069368 | 0,78 | 3,31E+08 | 3,22E+08 |
| 38,6 | | 0,0054775 | 0,0069333 | 1,59 | 5,55E+07 | 6,51E+07 |
| 3,3 | | 0,0054894 | 0,0069299 | 0,21 | 9,02E+06 | 1,26E+07 |
| 19,5 | | 0,0054990 | 0,0069265 | 2,49 | 4,02E+06 | 5,49E+06 |
| 17,7 | | 0,0055076 | 0,0069231 | 5,28 | 5,37E+07 | 2,18E+07 |
| 35,6 | | 0,0055260 | 0,0069197 | 0,58 | 1,22E+08 | 1,43E+08 |
| 53,2 | | 0,0055263 | 0,0069163 | 0,80 | 7,77E+08 | 7,00E+08 |
| 43,4 | | 0,0055360 | 0,0069129 | 1,65 | 1,07E+08 | 1,08E+08 |
| 20,1 | | 0,0055377 | 0,0069094 | 0,70 | 8,98E+07 | 8,90E+07 |
| 52,9 | | 0,0055445 | 0,0069061 | 1,32 | 3,86E+08 | 3,41E+08 |
| 9,6 | | 0,0055504 | 0,0069027 | 0,59 | 3,54E+07 | 4,02E+07 |
| 30,5 | | 0,0055521 | 0,0068993 | 1,16 | 1,43E+08 | 1,53E+08 |
| 20,7 | | 0,0055584 | 0,0068959 | 1,54 | 1,11E+08 | 1,29E+08 |
| 7,4 | | 0,0055627 | 0,0068925 | 0,53 | 7,56E+06 | 6,66E+06 |
| 12,2 | | 0,0055685 | 0,0068891 | 0,14 | 1,85E+07 | 1,10E+07 |
| 52,8 | | 0,0055695 | 0,0068857 | 1,22 | 8,09E+08 | 7,64E+08 |
| 4 | | 0,0055853 | 0,0069353 | 1,64 | 1,52E+07 | 1,25E+07 |
| 28,9 | | 0,0055856 | 0,0069319 | 2,89 | 5,07E+07 | 7,81E+07 |
| 56,9 | | 0,0055942 | 0,0069285 | 1,39 | 3,41E+08 | 3,14E+08 |
| 60,9 | | 0,0056026 | 0,0069251 | 0,77 | 2,69E+09 | 2,79E+09 |
| 58,7 | | 0,0056031 | 0,0069217 | 0,56 | 2,87E+08 | 2,24E+08 |
| 42,5 | | 0,0056126 | 0,0069790 | 35,47 | 8,33E+06 | 4,76E+06 |
| 30,8 | | 0,0056180 | 0,0069756 | 0,11 | 9,75E+07 | 1,20E+08 |

| | | | | | | |
|---|---|---|---|---|---|---|
| 25,3 | | 0,0056198 | 0,0069722 | 0,68 | 6,92E+07 | 8,64E+07 |
| 42,8 | | 0,0056288 | 0,0069688 | 0,68 | 1,61E+08 | 2,01E+08 |
| 21,1 | | 0,0056319 | 0,0069653 | 1,42 | 5,35E+07 | 6,47E+07 |
| 7,1 | | 0,0056457 | 0,0069620 | 2,71 | 1,36E+07 | 2,06E+07 |
| 53,9 | | 0,0056568 | 0,0070054 | 0,84 | 3,00E+09 | 2,80E+09 |
| 48,9 | | 0,0056752 | 0,0070019 | 1,11 | 1,23E+09 | 1,22E+09 |
| 53,5 | | 0,0056756 | 0,0069985 | 1,43 | 1,74E+08 | 2,00E+08 |
| 8,6 | | 0,0056849 | 0,0069951 | 0,71 | 1,03E+07 | 1,24E+07 |
| 27,7 | | 0,0056855 | 0,0069917 | 0,78 | 7,92E+08 | 7,00E+08 |
| 32,2 | | 0,0056911 | 0,0069883 | 1,19 | 1,96E+08 | 1,82E+08 |
| 42,3 | | 0,0056976 | 0,0070452 | 1,61 | 1,00E+09 | 1,15E+09 |
| 44,1 | | 0,0056977 | 0,0070418 | 0,89 | 2,62E+09 | 2,58E+09 |
| 30,6 | | 0,0056984 | 0,0070384 | 0,49 | 2,62E+07 | 4,02E+07 |
| 43,7 | | 0,0057098 | 0,0071476 | 1,30 | 4,79E+08 | 5,23E+08 |
| 9,1 | | 0,0057258 | 0,0071441 | 2,06 | 2,34E+07 | 2,58E+07 |
| 51 | | 0,0057403 | 0,0071406 | 0,71 | 3,18E+08 | 3,00E+08 |
| 36,8 | | 0,0057415 | 0,0071372 | 0,60 | 3,09E+08 | 3,64E+08 |
| 32,3 | | 0,0057568 | 0,0071337 | 1,56 | 1,46E+08 | 1,79E+08 |
| 27,2 | | 0,0057582 | 0,0071303 | 0,82 | 1,12E+08 | 1,06E+08 |
| 48,6 | | 0,0057582 | 0,0071268 | 0,85 | 1,02E+09 | 9,61E+08 |
| 40,9 | | 0,0057594 | 0,0071234 | 0,64 | 4,13E+08 | 3,73E+08 |
| 22,6 | | 0,0057813 | 0,0071199 | 1,58 | 4,39E+07 | 3,99E+07 |
| 9,4 | | 0,0057852 | 0,0071165 | 3,71 | 4,64E+06 | 2,04E+06 |
| 45,7 | | 0,0057855 | 0,0071130 | 0,77 | 2,49E+08 | 2,79E+08 |
| 22,9 | | 0,0057969 | 0,0071096 | 0,75 | 5,69E+07 | 6,30E+07 |
| 12,4 | | 0,0057998 | 0,0071062 | 1,37 | 4,96E+07 | 4,57E+07 |
| 42,5 | | 0,0058033 | 0,0071027 | 0,66 | 1,85E+08 | 1,84E+08 |
| 34,7 | | 0,0058108 | 0,0070993 | 1,34 | 2,96E+07 | 2,94E+07 |
| 36,9 | | 0,0058313 | 0,0071865 | 1,61 | 3,72E+08 | 3,83E+08 |
| 19 | | 0,0058432 | 0,0071830 | 1,53 | 1,02E+08 | 8,34E+07 |
| 26,5 | | 0,0058484 | 0,0071796 | 0,70 | 1,46E+08 | 1,38E+08 |
| 14,6 | | 0,0058493 | 0,0071761 | 0,47 | 2,12E+07 | 2,38E+07 |
| 27,1 | | 0,0058810 | 0,0072650 | 3,39 | 1,43E+07 | 6,97E+06 |
| 9,2 | | 0,0058893 | 0,0072615 | 1,76 | 8,59E+06 | 9,51E+06 |
| 20,8 | | 0,0058895 | 0,0072580 | 1,39 | 3,04E+07 | 2,74E+07 |
| 31,5 | | 0,0058989 | 0,0072546 | 0,02 | 2,01E+08 | 7,59E+08 |
| 53,6 | | 0,0059045 | 0,0072511 | 1,34 | 4,33E+08 | 3,73E+08 |
| 34 | | 0,0059052 | 0,0072476 | 0,74 | 1,49E+08 | 1,73E+08 |
| 35 | | 0,0059112 | 0,0072441 | 0,53 | 1,03E+08 | 1,04E+08 |
| 42,1 | | 0,0059178 | 0,0073001 | 1,46 | 3,91E+08 | 3,57E+08 |
| 24,8 | | 0,0059254 | 0,0072966 | 5,58 | 7,33E+06 | 1,62E+07 |
| 49 | | 0,0059257 | 0,0072931 | 0,63 | 1,32E+09 | 1,76E+09 |
| 33,3 | | 0,0059579 | 0,0073356 | 1,87 | 4,02E+07 | 5,05E+07 |
| 44,4 | | 0,0059887 | 0,0074679 | 0,62 | 9,52E+08 | 7,10E+08 |
| 48,2 | | 0,0059917 | 0,0074644 | 0,82 | 7,16E+08 | 7,71E+08 |
| 23,6 | | 0,0060112 | 0,0074608 | 0,82 | 5,66E+07 | 5,16E+07 |
| 19,5 | | 0,0060171 | 0,0074572 | 2,57 | 3,67E+06 | 2,69E+06 |
| 58,4 | | 0,0060237 | 0,0074537 | 0,74 | 2,88E+08 | 2,84E+08 |
| 21,5 | | 0,0060293 | 0,0074501 | 2,00 | 5,70E+07 | 7,06E+07 |
| 50,1 | | 0,0060382 | 0,0074466 | 0,75 | 1,25E+08 | 1,29E+08 |

| | | | | | | |
|---|---|---|---|---|---|---|
| 63,6 | | 0,0060443 | 0,0074430 | 1,18 | 2,22E+09 | 2,37E+09 |
| 22,6 | | 0,0060476 | 0,0074395 | 0,58 | 9,90E+07 | 8,99E+07 |
| 21,6 | | 0,0060615 | 0,0074359 | 1,97 | 2,53E+07 | 2,96E+07 |
| 18,7 | | 0,0060653 | 0,0074914 | 5,50 | 8,76E+06 | 2,64E+07 |
| 4,4 | | 0,0060779 | 0,0074879 | 0,48 | 3,81E+07 | 2,36E+07 |
| 36,4 | | 0,0060834 | 0,0074843 | 1,78 | 9,77E+06 | 1,19E+07 |
| 10,4 | | 0,0060872 | 0,0074807 | 0,17 | 2,95E+07 | 3,27E+07 |
| 10,8 | | 0,0060910 | 0,0074772 | 0,08 | 3,96E+07 | 3,75E+07 |
| 36,5 | | 0,0060948 | 0,0074736 | 0,70 | 1,19E+08 | 9,81E+07 |
| 23 | | 0,0061093 | 0,0074701 | 0,19 | 1,78E+07 | 1,91E+07 |
| 44,2 | | 0,0061101 | 0,0074665 | 0,54 | 7,74E+07 | 1,08E+08 |
| 81,7 | | 0,0061177 | 0,0074630 | 0,82 | 3,16E+09 | 3,44E+09 |
| 38,8 | | 0,0061334 | 0,0074595 | 0,70 | 1,67E+08 | 1,87E+08 |
| 39,1 | | 0,0061370 | 0,0074559 | 1,54 | 1,08E+08 | 1,00E+08 |
| 15,1 | | 0,0061427 | 0,0074524 | 0,13 | 1,87E+07 | 2,05E+07 |
| 28,4 | | 0,0061559 | 0,0074489 | 0,80 | 5,96E+08 | 5,71E+08 |
| 48,4 | | 0,0061568 | 0,0074453 | 0,74 | 3,19E+08 | 3,40E+08 |
| 41,5 | | 0,0061662 | 0,0074418 | 1,18 | 3,17E+08 | 3,06E+08 |
| 36,9 | | 0,0061700 | 0,0074383 | 1,24 | 1,39E+08 | 1,45E+08 |
| 11,5 | | 0,0061725 | 0,0074348 | 1,45 | 2,86E+07 | 3,29E+07 |
| 42,1 | | 0,0061827 | 0,0074313 | 0,86 | 1,14E+09 | 1,10E+09 |
| 37,4 | | 0,0061880 | 0,0074882 | 0,61 | 2,37E+08 | 2,20E+08 |
| 43,4 | | 0,0061978 | 0,0075432 | 10,56 | 1,02E+07 | 4,68E+06 |
| 49,1 | | 0,0062075 | 0,0075396 | 0,75 | 3,68E+08 | 3,83E+08 |
| 8 | | 0,0062101 | 0,0075945 | 7,11 | 3,68E+06 | 1,71E+06 |
| 60,6 | | 0,0062101 | 0,0075910 | 1,20 | 6,01E+09 | 5,55E+09 |
| 41,3 | | 0,0062308 | 0,0076326 | 0,70 | 1,20E+08 | 1,12E+08 |
| 49,8 | | 0,0062347 | 0,0076798 | 0,56 | 1,70E+08 | 1,87E+08 |
| 60,4 | | 0,0062363 | 0,0076762 | 1,30 | 1,70E+09 | 1,61E+09 |
| 9 | | 0,0062593 | 0,0076726 | 1,79 | 2,46E+07 | 1,75E+07 |
| 14,8 | | 0,0062600 | 0,0076690 | 2,09 | 4,60E+07 | 3,51E+07 |
| 51,1 | | 0,0062635 | 0,0076654 | 0,67 | 1,11E+09 | 1,28E+09 |
| 53,7 | | 0,0062721 | 0,0076618 | 0,75 | 2,66E+08 | 3,00E+08 |
| 53 | | 0,0062886 | 0,0077014 | 1,27 | 6,89E+08 | 8,02E+08 |
| 6,9 | | 0,0062942 | 0,0076978 | 1,72 | 4,48E+06 | 3,76E+06 |
| 30,5 | | 0,0062976 | 0,0076942 | 0,85 | 9,45E+07 | 8,89E+07 |
| 40 | | 0,0062978 | 0,0076906 | 0,11 | 1,80E+07 | 3,58E+07 |
| 40,2 | | 0,0063070 | 0,0076870 | 1,31 | 2,82E+08 | 3,01E+08 |
| 45,3 | | 0,0063153 | 0,0076834 | 1,33 | 2,93E+09 | 3,24E+09 |
| 76,8 | | 0,0063202 | 0,0076798 | 1,22 | 1,45E+08 | 1,34E+08 |
| 12,2 | | 0,0063274 | 0,0076762 | 0,17 | 1,96E+07 | 1,85E+07 |
| 38,7 | | 0,0063346 | 0,0076726 | 1,45 | 8,98E+07 | 1,12E+08 |
| 19,7 | | 0,0063451 | 0,0077288 | 0,12 | 2,24E+07 | 2,16E+07 |
| 36,1 | | 0,0063491 | 0,0077252 | 1,46 | 4,44E+07 | 4,34E+07 |
| 31,9 | | 0,0063508 | 0,0077216 | 0,10 | 3,24E+07 | 3,64E+07 |
| 35,8 | | 0,0063581 | 0,0077180 | 1,38 | 4,23E+08 | 4,72E+08 |
| 21,5 | | 0,0063796 | 0,0077144 | 0,78 | 1,13E+08 | 1,15E+08 |
| 55,7 | | 0,0063828 | 0,0077108 | 0,66 | 9,12E+08 | 1,01E+09 |
| 25,1 | | 0,0063843 | 0,0077072 | 1,22 | 1,97E+08 | 1,91E+08 |
| 6,5 | | 0,0063857 | 0,0077036 | 2,39 | 4,68E+06 | 8,02E+06 |

| | | | | | | |
|---|---|---|---|---|---|---|
| 45,4 | | 0,0063896 | 0,0077578 | 1,23 | 6,07E+08 | 5,40E+08 |
| 55,2 | | 0,0063911 | 0,0077542 | 0,75 | 8,73E+08 | 1,02E+09 |
| 40,5 | | 0,0063923 | 0,0077506 | 1,52 | 5,62E+07 | 4,64E+07 |
| 50,6 | | 0,0064045 | 0,0077470 | 1,15 | 1,19E+08 | 1,22E+08 |
| 38,1 | | 0,0064094 | 0,0077434 | 0,77 | 2,78E+08 | 3,01E+08 |
| 40,3 | | 0,0064436 | 0,0077900 | 1,29 | 1,51E+08 | 1,56E+08 |
| 33,3 | | 0,0064472 | 0,0077863 | 0,47 | 4,16E+07 | 3,39E+07 |
| 49,1 | | 0,0064529 | 0,0078849 | 1,41 | 2,00E+08 | 2,33E+08 |
| 38,1 | | 0,0064565 | 0,0078812 | 0,74 | 4,79E+08 | 5,35E+08 |
| 25,3 | | 0,0064569 | 0,0078776 | 1,70 | 1,04E+08 | 1,46E+08 |
| 38 | | 0,0064622 | 0,0078739 | 0,57 | 1,29E+08 | 1,19E+08 |
| 63,6 | | 0,0064764 | 0,0078703 | 0,28 | 5,35E+07 | 4,88E+07 |
| 19,7 | | 0,0065017 | 0,0078666 | 0,20 | 6,24E+07 | 5,45E+07 |
| 35,5 | | 0,0065057 | 0,0078630 | 0,88 | 1,27E+08 | 1,27E+08 |
| 23,9 | | 0,0065077 | 0,0078593 | 1,30 | 7,05E+07 | 6,40E+07 |
| 14,7 | | 0,0065147 | 0,0078557 | 0,23 | 2,51E+07 | 1,60E+07 |
| 6,6 | | 0,0065266 | 0,0078964 | 0,56 | 2,10E+07 | 2,52E+07 |
| 51,9 | | 0,0065279 | 0,0078928 | 1,57 | 2,13E+08 | 2,01E+08 |
| 50,2 | | 0,0065448 | 0,0079760 | 0,71 | 1,78E+08 | 1,54E+08 |
| 41,5 | | 0,0065472 | 0,0080148 | 1,27 | 1,33E+09 | 1,25E+09 |
| 12,7 | | 0,0065474 | 0,0080111 | 8,53 | 3,16E+06 | 1,15E+07 |
| 20,2 | | 0,0065548 | 0,0080074 | 0,55 | 1,86E+07 | 1,81E+07 |
| 56,1 | | 0,0065569 | 0,0080609 | 0,69 | 1,19E+09 | 1,38E+09 |
| 23,7 | | 0,0065587 | 0,0080571 | 0,79 | 3,88E+08 | 3,88E+08 |
| 55,4 | | 0,0065594 | 0,0080534 | 1,28 | 3,91E+08 | 3,56E+08 |
| 8,7 | | 0,0065716 | 0,0081436 | 0,31 | 1,23E+07 | 1,25E+07 |
| 18,2 | | 0,0065721 | 0,0081399 | 1,37 | 4,32E+07 | 4,60E+07 |
| 11,4 | | 0,0065807 | 0,0081362 | 0,69 | 1,27E+07 | 1,17E+07 |
| 54,7 | | 0,0065824 | 0,0081324 | 1,21 | 3,09E+08 | 3,35E+08 |
| 32,5 | | 0,0065832 | 0,0081287 | 0,78 | 1,81E+08 | 2,05E+08 |
| 9,4 | | 0,0065866 | 0,0081249 | 2,04 | 5,33E+06 | 7,48E+06 |
| 6,7 | | 0,0065913 | 0,0081212 | 1,38 | 3,24E+06 | 3,37E+06 |
| 19,9 | | 0,0065939 | 0,0081175 | 4,99 | 2,86E+06 | 4,62E+06 |
| 20,9 | | 0,0065946 | 0,0081138 | 1,44 | 3,76E+07 | 3,60E+07 |
| 12,7 | | 0,0065959 | 0,0081100 | 0,16 | 1,10E+07 | 1,40E+07 |
| 61 | | 0,0066039 | 0,0081063 | 0,75 | 1,88E+09 | 1,73E+09 |
| 24,5 | | 0,0066065 | 0,0081026 | 1,68 | 9,48E+07 | 9,04E+07 |
| 50,3 | | 0,0066076 | 0,0080989 | 1,28 | 1,14E+09 | 1,03E+09 |
| 23,5 | | 0,0066089 | 0,0080952 | 1,26 | 6,86E+07 | 7,60E+07 |
| 28,3 | | 0,0066248 | 0,0080915 | 0,73 | 1,02E+08 | 9,65E+07 |
| 23,7 | | 0,0066327 | 0,0080878 | 0,80 | 1,55E+08 | 1,48E+08 |
| 28,5 | | 0,0066356 | 0,0081408 | 1,38 | 1,99E+08 | 1,95E+08 |
| 52,2 | | 0,0066433 | 0,0081370 | 46,87 | 4,08E+06 | 3,06E+07 |
| 41,8 | | 0,0066476 | 0,0081333 | 1,76 | 5,42E+07 | 6,90E+07 |
| 9,1 | | 0,0066562 | 0,0081296 | 1,41 | 4,42E+07 | 4,13E+07 |
| 28,5 | | 0,0066659 | 0,0081259 | 1,52 | 7,92E+07 | 6,20E+07 |
| 10,8 | + | 0,0066695 | 0,0081222 | 4,76 | 3,29E+06 | 1,92E+06 |
| 61,1 | | 0,0066737 | 0,0081185 | 0,82 | 2,06E+09 | 2,28E+09 |
| 56,5 | | 0,0066972 | 0,0081148 | 0,84 | 1,61E+08 | 1,53E+08 |
| 40,7 | | 0,0067047 | 0,0081111 | 0,80 | 7,56E+08 | 7,88E+08 |

| | | | | | | |
|---|---|---|---|---|---|---|
| 13,1 | | 0,0067137 | 0,0081074 | 0,11 | 2,81E+07 | 3,48E+07 |
| 32,6 | | 0,0067368 | 0,0082020 | 0,75 | 5,35E+08 | 4,93E+08 |
| 9,1 | | 0,0067605 | 0,0081983 | 0,69 | 3,70E+07 | 3,24E+07 |
| 32,9 | | 0,0067621 | 0,0081945 | 1,32 | 6,53E+07 | 6,15E+07 |
| 50,2 | | 0,0067680 | 0,0081908 | 0,74 | 1,57E+08 | 1,62E+08 |
| 50,8 | | 0,0067684 | 0,0081871 | 1,35 | 6,31E+08 | 5,42E+08 |
| 70,9 | | 0,0067745 | 0,0081834 | 1,88 | 5,44E+09 | 6,06E+09 |
| 53 | | 0,0067914 | 0,0081797 | 0,80 | 1,19E+09 | 1,24E+09 |
| 49,9 | | 0,0067969 | 0,0081760 | 1,38 | 6,40E+07 | 6,67E+07 |
| 16,2 | | 0,0068198 | 0,0081723 | 0,66 | 1,93E+07 | 2,08E+07 |
| 19,7 | | 0,0068260 | 0,0081686 | 1,69 | 4,59E+07 | 3,54E+07 |
| 8,3 | | 0,0068336 | 0,0081649 | 0,41 | 9,50E+06 | 1,04E+07 |
| 51,5 | | 0,0068451 | 0,0081612 | 0,89 | 1,52E+08 | 1,49E+08 |
| 23,9 | | 0,0068598 | 0,0082136 | 0,47 | 4,78E+07 | 6,11E+07 |
| 15,1 | | 0,0068628 | 0,0082099 | 0,63 | 1,33E+07 | 1,52E+07 |
| 32,9 | | 0,0068824 | 0,0082061 | 0,61 | 1,18E+08 | 9,01E+07 |
| 66,3 | | 0,0069087 | 0,0082024 | 1,42 | 3,82E+08 | 3,27E+08 |
| 36,3 | | 0,0069088 | 0,0081987 | 0,54 | 9,39E+07 | 8,61E+07 |
| 55,7 | | 0,0069122 | 0,0081950 | 0,76 | 1,29E+08 | 1,23E+08 |
| 44,3 | | 0,0069190 | 0,0081913 | 1,46 | 6,08E+08 | 5,83E+08 |
| 12,7 | | 0,0069282 | 0,0082291 | 0,59 | 4,39E+07 | 5,49E+07 |
| 68,6 | | 0,0069308 | 0,0082254 | 1,52 | 2,41E+08 | 2,80E+08 |
| 68,8 | | 0,0069331 | 0,0082217 | 1,22 | 8,39E+07 | 9,53E+07 |
| 48,7 | | 0,0069340 | 0,0082180 | 1,10 | 6,53E+08 | 6,21E+08 |
| 19 | | 0,0069485 | 0,0082143 | 1,55 | 3,01E+07 | 2,44E+07 |
| 37,4 | | 0,0069641 | 0,0082106 | 1,47 | 3,69E+08 | 4,39E+08 |
| 28,8 | | 0,0069675 | 0,0082069 | 0,77 | 1,03E+08 | 1,06E+08 |
| 30,3 | | 0,0070027 | 0,0083417 | 0,76 | 9,20E+07 | 1,01E+08 |
| 27,9 | | 0,0070325 | 0,0083380 | 0,67 | 1,51E+08 | 1,62E+08 |
| 42,5 | | 0,0070440 | 0,0083342 | 1,29 | 4,01E+08 | 4,23E+08 |
| 41,7 | | 0,0070557 | 0,0083305 | 1,85 | 1,18E+08 | 8,59E+07 |
| 31,9 | | 0,0070833 | 0,0084129 | 1,32 | 1,87E+08 | 1,88E+08 |
| 24,4 | | 0,0070868 | 0,0084092 | 0,69 | 1,26E+08 | 1,35E+08 |
| 17,7 | | 0,0071112 | 0,0084915 | 0,58 | 8,52E+07 | 7,54E+07 |
| 9,1 | | 0,0071124 | 0,0084877 | 6,94 | 9,17E+06 | 6,45E+06 |
| 46,2 | | 0,0071345 | 0,0084839 | 1,46 | 3,12E+07 | 3,05E+07 |
| 31,9 | | 0,0071384 | 0,0084801 | 1,98 | 5,03E+07 | 6,40E+07 |
| 33,5 | | 0,0071393 | 0,0084763 | 0,27 | 6,76E+07 | 7,52E+07 |
| 50,1 | | 0,0071490 | 0,0084725 | 0,70 | 7,47E+08 | 8,61E+08 |
| 11,5 | | 0,0071530 | 0,0084687 | 1,95 | 4,08E+06 | 4,54E+06 |
| 38,8 | | 0,0071531 | 0,0084649 | 0,79 | 9,04E+07 | 8,53E+07 |
| 21,4 | | 0,0071706 | 0,0084611 | 0,68 | 9,46E+07 | 9,18E+07 |
| 46,6 | | 0,0071737 | 0,0084573 | 0,74 | 7,61E+08 | 9,22E+08 |
| 16,3 | | 0,0071737 | 0,0084536 | 3,79 | 1,51E+08 | 6,28E+07 |
| 40,8 | | 0,0071838 | 0,0084498 | 14,98 | 1,29E+06 | 1,41E+06 |
| 19,9 | | 0,0071905 | 0,0084460 | 1,49 | 3,00E+07 | 3,15E+07 |
| 14,2 | | 0,0071964 | 0,0084423 | 5,81 | 1,63E+06 | 5,15E+06 |
| 13,9 | | 0,0072363 | 0,0084813 | 18,69 | 7,94E+06 | 1,24E+06 |
| 23,5 | | 0,0072787 | 0,0085898 | 0,80 | 1,59E+08 | 1,65E+08 |
| 29,4 | | 0,0072947 | 0,0086340 | 0,82 | 2,73E+08 | 2,75E+08 |

| | | | | | | |
|---|---|---|---|---|---|---|
| 53,9 | | 0,0072950 | 0,0086302 | 0,70 | 3,39E+09 | 3,85E+09 |
| 7,4 | | 0,0072970 | 0,0086263 | 0,33 | 1,00E+07 | 9,16E+06 |
| 10,5 | | 0,0072982 | 0,0086225 | 0,24 | 1,08E+07 | 1,27E+07 |
| 77,3 | | 0,0072994 | 0,0086187 | 0,76 | 4,90E+09 | 5,80E+09 |
| 52,8 | | 0,0073445 | 0,0086628 | 2,26 | 1,26E+07 | 1,72E+07 |
| 29,7 | | 0,0073527 | 0,0086590 | 0,16 | 2,95E+07 | 2,09E+07 |
| 14,5 | | 0,0073591 | 0,0086551 | 0,32 | 1,07E+07 | 1,03E+07 |
| 31,1 | | 0,0073789 | 0,0086513 | 0,59 | 6,06E+07 | 6,05E+07 |
| 19,3 | | 0,0073907 | 0,0086475 | 1,45 | 2,77E+07 | 2,71E+07 |
| 33,3 | | 0,0073936 | 0,0086436 | 0,70 | 1,36E+08 | 1,53E+08 |
| 22,2 | | 0,0074073 | 0,0086398 | 1,84 | 4,97E+07 | 7,25E+07 |
| 10,9 | | 0,0074092 | 0,0086360 | 0,87 | 9,03E+07 | 8,41E+07 |
| 26,2 | | 0,0074229 | 0,0086321 | 0,06 | 7,00E+07 | 8,21E+07 |
| 55 | | 0,0074386 | 0,0086283 | 0,76 | 1,04E+09 | 1,12E+09 |
| 12,7 | | 0,0074481 | 0,0086245 | 1,46 | 3,43E+07 | 3,18E+07 |
| 27,4 | | 0,0074525 | 0,0086207 | 0,10 | 7,52E+07 | 8,55E+07 |
| 16 | | 0,0074660 | 0,0086169 | 7,50 | 2,04E+06 | 6,75E+06 |
| 50,9 | | 0,0074904 | 0,0086131 | 0,50 | 1,21E+09 | 9,77E+08 |
| 33,8 | | 0,0074983 | 0,0086093 | 0,77 | 1,36E+08 | 1,23E+08 |
| 52,7 | | 0,0075095 | 0,0086055 | 0,89 | 6,61E+08 | 7,09E+08 |
| 5,6 | | 0,0075097 | 0,0086017 | 4,23 | 3,57E+06 | 2,41E+06 |
| 5,8 | | 0,0075319 | 0,0085979 | 0,44 | 8,55E+06 | 1,03E+07 |
| 21,9 | | 0,0075791 | 0,0085941 | 0,71 | 8,30E+07 | 9,67E+07 |
| 44,4 | | 0,0076032 | 0,0085903 | 0,79 | 1,91E+08 | 2,20E+08 |
| 42,9 | | 0,0076101 | 0,0085865 | 0,75 | 3,14E+08 | 3,48E+08 |
| 47,1 | | 0,0076205 | 0,0085827 | 1,39 | 1,07E+08 | 1,33E+08 |
| 18 | | 0,0076325 | 0,0086335 | 1,96 | 1,16E+07 | 1,66E+07 |
| 22,4 | | 0,0076330 | 0,0086297 | 0,64 | 2,15E+07 | 2,61E+07 |
| 56,6 | | 0,0076358 | 0,0086259 | 1,09 | 2,52E+09 | 2,50E+09 |
| 12,7 | | 0,0076371 | 0,0086221 | 0,92 | 3,17E+07 | 3,28E+07 |
| 20,4 | | 0,0076477 | 0,0086658 | 0,24 | 8,44E+06 | 1,39E+07 |
| 21,7 | | 0,0076498 | 0,0086620 | 0,58 | 3,74E+07 | 4,09E+07 |
| 18,4 | | 0,0076544 | 0,0086582 | 0,72 | 5,52E+07 | 5,63E+07 |
| 35,5 | | 0,0076622 | 0,0086544 | 0,77 | 2,42E+08 | 2,25E+08 |
| 46,8 | | 0,0076634 | 0,0086506 | 0,74 | 2,11E+08 | 2,42E+08 |
| 14,7 | | 0,0076702 | 0,0086468 | 0,64 | 4,56E+07 | 4,57E+07 |
| 55,4 | | 0,0076713 | 0,0086430 | 0,83 | 4,71E+08 | 5,30E+08 |
| 5,2 | | 0,0076808 | 0,0086392 | 0,42 | 1,25E+07 | 1,18E+07 |
| 47,4 | | 0,0076844 | 0,0086354 | 1,30 | 1,89E+08 | 1,71E+08 |
| 34,7 | | 0,0076860 | 0,0086317 | 0,88 | 2,63E+09 | 2,60E+09 |
| 57,7 | | 0,0076889 | 0,0086279 | 0,83 | 1,69E+09 | 1,76E+09 |
| 25,8 | | 0,0077001 | 0,0086241 | 0,15 | 1,92E+07 | 5,57E+07 |
| 15,5 | | 0,0077349 | 0,0086623 | 1,77 | 1,97E+07 | 2,89E+07 |
| 26,5 | | 0,0077355 | 0,0086585 | 0,83 | 4,40E+07 | 4,35E+07 |
| 8,2 | | 0,0077429 | 0,0087089 | 0,48 | 1,98E+07 | 1,52E+07 |
| 43,1 | | 0,0077659 | 0,0087051 | 0,75 | 1,48E+08 | 1,65E+08 |
| 22,5 | | 0,0077889 | 0,0087013 | 1,29 | 4,31E+07 | 4,10E+07 |
| 24,4 | | 0,0077918 | 0,0086975 | 1,14 | 1,50E+08 | 1,40E+08 |
| 16,1 | | 0,0077932 | 0,0086937 | 0,10 | 2,21E+07 | 2,64E+07 |
| 64,4 | | 0,0077952 | 0,0086899 | 1,31 | 1,57E+10 | 1,39E+10 |

| | | | | | | |
|---|---|---|---|---|---|---|
| 48,4 | | 0,0078229 | 0,0086861 | 1,15 | 3,19E+08 | 3,19E+08 |
| 42,5 | | 0,0078469 | 0,0086823 | 1,34 | 5,80E+08 | 5,94E+08 |
| 29,2 | | 0,0078782 | 0,0087255 | 1,61 | 5,17E+07 | 3,78E+07 |
| 49,2 | | 0,0078820 | 0,0087217 | 1,77 | 5,86E+07 | 7,25E+07 |
| 56,5 | | 0,0078860 | 0,0087179 | 1,44 | 4,08E+08 | 4,06E+08 |
| 21,2 | | 0,0078996 | 0,0087142 | 0,22 | 2,38E+07 | 2,15E+07 |
| 12,8 | | 0,0079122 | 0,0087104 | 1,73 | 4,06E+07 | 5,97E+07 |
| 12,2 | | 0,0079163 | 0,0087066 | 1,79 | 3,02E+07 | 2,20E+07 |
| 15,9 | | 0,0079167 | 0,0087028 | 0,62 | 7,46E+07 | 5,53E+07 |
| 70 | | 0,0079201 | 0,0086990 | 0,04 | 5,64E+07 | 8,84E+07 |
| 41,8 | | 0,0079250 | 0,0086953 | 1,53 | 2,53E+08 | 3,02E+08 |
| 55,5 | | 0,0079297 | 0,0086915 | 0,57 | 4,41E+07 | 3,60E+07 |
| 40,8 | | 0,0079328 | 0,0086877 | 0,63 | 3,04E+08 | 2,72E+08 |
| 26,4 | | 0,0079406 | 0,0087238 | 0,05 | 5,82E+07 | 6,79E+07 |
| 28,2 | | 0,0079411 | 0,0087200 | 1,47 | 3,18E+07 | 3,82E+07 |
| 69,8 | | 0,0079516 | 0,0087163 | 0,68 | 3,60E+08 | 3,60E+08 |
| 23,4 | | 0,0079649 | 0,0087125 | 0,44 | 3,68E+07 | 4,37E+07 |
| 25,9 | | 0,0079677 | 0,0087087 | 0,80 | 1,27E+08 | 1,46E+08 |
| 41,6 | | 0,0079684 | 0,0087050 | 1,42 | 1,68E+08 | 1,37E+08 |
| 22,9 | | 0,0079840 | 0,0087012 | 0,05 | 5,42E+07 | 5,68E+07 |
| 42,5 | | 0,0079907 | 0,0086975 | 0,71 | 2,36E+08 | 2,60E+08 |
| 26,4 | | 0,0079930 | 0,0086937 | 0,36 | 1,69E+07 | 3,20E+07 |
| 31,1 | | 0,0080025 | 0,0086900 | 1,18 | 7,73E+08 | 7,79E+08 |
| 30 | | 0,0080066 | 0,0086862 | 1,24 | 1,56E+08 | 1,43E+08 |
| 12,8 | | 0,0080262 | 0,0086825 | 0,61 | 8,37E+06 | 6,98E+06 |
| 18,8 | | 0,0080296 | 0,0086787 | 1,25 | 2,60E+07 | 2,51E+07 |
| 40,6 | | 0,0080345 | 0,0086750 | 0,65 | 8,12E+08 | 7,73E+08 |
| 54,5 | | 0,0080430 | 0,0087263 | 1,27 | 1,21E+09 | 1,22E+09 |
| 43,2 | | 0,0080651 | 0,0087226 | 2,29 | 9,44E+07 | 6,52E+07 |
| 34 | | 0,0080879 | 0,0087584 | 0,67 | 3,02E+08 | 3,47E+08 |
| 27,8 | | 0,0081302 | 0,0087546 | 0,66 | 3,71E+08 | 3,84E+08 |
| 16,4 | | 0,0081306 | 0,0087509 | 0,65 | 4,82E+07 | 4,99E+07 |
| 42,3 | | 0,0081319 | 0,0087471 | 0,72 | 3,94E+08 | 4,58E+08 |
| 15,9 | | 0,0081322 | 0,0087433 | 6,13 | 1,22E+07 | 5,75E+06 |
| 35,9 | | 0,0081408 | 0,0087396 | 0,65 | 8,90E+07 | 9,74E+07 |
| 59,3 | | 0,0081437 | 0,0087358 | 0,87 | 3,99E+09 | 3,66E+09 |
| 58,8 | | 0,0081538 | 0,0087321 | 1,20 | 2,74E+09 | 2,98E+09 |
| 6,2 | | 0,0081553 | 0,0087284 | 0,21 | 6,89E+06 | 6,97E+06 |
| 15,5 | | 0,0081683 | 0,0087246 | 0,28 | 6,25E+07 | 5,97E+07 |
| 20,8 | | 0,0081754 | 0,0087209 | 0,47 | 2,36E+07 | 3,87E+07 |
| 31,4 | | 0,0081833 | 0,0087172 | 0,81 | 5,44E+07 | 4,91E+07 |
| 14 | | 0,0081887 | 0,0087134 | 0,18 | 2,16E+07 | 1,89E+07 |
| 61,3 | | 0,0081998 | 0,0087097 | 1,48 | 5,81E+08 | 7,23E+08 |
| 29,6 | | 0,0082126 | 0,0087060 | 0,55 | 1,01E+09 | 7,06E+08 |
| 26,3 | | 0,0082327 | 0,0087023 | 1,76 | 1,95E+07 | 2,81E+07 |
| 9,6 | | 0,0082386 | 0,0087447 | 0,67 | 4,98E+07 | 3,87E+07 |
| 8,7 | | 0,0082637 | 0,0087409 | 1,96 | 1,33E+07 | 1,37E+07 |
| 20,8 | | 0,0082651 | 0,0087372 | 13,38 | 1,12E+06 | 6,40E+06 |
| 21,4 | | 0,0082864 | 0,0087335 | 0,42 | 2,10E+07 | 2,85E+07 |
| 33,8 | | 0,0083103 | 0,0087298 | 1,87 | 3,01E+07 | 3,96E+07 |

| | | | | | | |
|---|---|---|---|---|---|---|
| 45 | | 0,0083114 | 0,0087260 | 1,27 | 1,56E+09 | 1,33E+09 |
| 45,8 | | 0,0083309 | 0,0088160 | 1,42 | 2,39E+08 | 2,70E+08 |
| 17,8 | | 0,0083666 | 0,0088514 | 1,37 | 7,24E+07 | 7,53E+07 |
| 34,6 | | 0,0083935 | 0,0088477 | 0,79 | 1,27E+09 | 1,35E+09 |
| 51,6 | | 0,0083979 | 0,0088439 | 1,20 | 1,56E+09 | 1,67E+09 |
| 32,5 | | 0,0084069 | 0,0088401 | 1,37 | 7,26E+07 | 7,49E+07 |
| 15,7 | | 0,0084181 | 0,0088364 | 3,70 | 3,53E+06 | 4,16E+06 |
| 37,2 | | 0,0084236 | 0,0088326 | 1,52 | 6,46E+06 | 8,02E+06 |
| 45,8 | | 0,0084291 | 0,0088289 | 0,87 | 1,18E+08 | 1,15E+08 |
| 25,5 | + | 0,0084350 | 0,0088251 | 19,45 | 1,59E+06 | 1,05E+06 |
| 39,5 | | 0,0084358 | 0,0088214 | 0,65 | 6,16E+07 | 6,16E+07 |
| 57,3 | | 0,0084374 | 0,0088176 | 1,36 | 1,34E+08 | 1,41E+08 |
| 36,6 | | 0,0084513 | 0,0088139 | 1,23 | 4,79E+07 | 5,00E+07 |
| 32,2 | | 0,0084633 | 0,0088627 | 1,31 | 3,12E+08 | 3,67E+08 |
| 27,9 | | 0,0084694 | 0,0088979 | 1,23 | 1,21E+08 | 1,17E+08 |
| 45,2 | | 0,0084979 | 0,0089467 | 0,69 | 8,53E+08 | 7,12E+08 |
| 28,7 | | 0,0085086 | 0,0089429 | 0,66 | 1,29E+08 | 1,34E+08 |
| 14 | | 0,0085173 | 0,0089391 | 11,90 | 1,57E+07 | 1,15E+07 |
| 11 | | 0,0085359 | 0,0089353 | 0,43 | 5,65E+06 | 6,82E+06 |
| 43,1 | | 0,0085748 | 0,0089315 | 0,75 | 4,18E+08 | 4,14E+08 |
| 20,1 | | 0,0085762 | 0,0089278 | 0,42 | 1,84E+07 | 2,32E+07 |
| 21,2 | | 0,0085862 | 0,0089628 | 0,45 | 2,51E+07 | 1,95E+07 |
| 49,3 | | 0,0085940 | 0,0089591 | 1,75 | 5,29E+07 | 7,87E+07 |
| 16,3 | | 0,0086038 | 0,0089553 | 0,54 | 7,29E+07 | 6,58E+07 |
| 16 | | 0,0086100 | 0,0089903 | 0,06 | 6,10E+07 | 4,86E+07 |
| 20,8 | | 0,0086176 | 0,0089865 | 0,52 | 5,08E+07 | 4,97E+07 |
| 43,8 | | 0,0086262 | 0,0090232 | 0,04 | 2,31E+08 | 1,49E+08 |
| 13,1 | | 0,0086486 | 0,0090598 | 0,05 | 4,75E+07 | 3,80E+07 |
| 37,9 | | 0,0086716 | 0,0090560 | 1,10 | 8,64E+08 | 8,70E+08 |
| 66,3 | | 0,0086741 | 0,0090522 | 1,16 | 1,09E+09 | 9,87E+08 |
| 16 | | 0,0086929 | 0,0091409 | 0,02 | 1,40E+08 | 5,88E+08 |
| 22,7 | | 0,0087255 | 0,0091371 | 0,28 | 4,45E+07 | 3,49E+07 |
| 13,8 | | 0,0087261 | 0,0091332 | 0,52 | 1,44E+07 | 1,73E+07 |
| 64,1 | | 0,0087559 | 0,0091294 | 0,88 | 3,02E+08 | 3,18E+08 |
| 27,9 | | 0,0087576 | 0,0091256 | 0,76 | 1,08E+08 | 1,22E+08 |
| 39 | | 0,0087612 | 0,0091217 | 0,78 | 1,47E+08 | 1,68E+08 |
| 12 | | 0,0087717 | 0,0091179 | 1,36 | 3,36E+07 | 3,29E+07 |
| 25,8 | | 0,0087745 | 0,0091141 | 0,71 | 1,71E+08 | 1,56E+08 |
| 40,2 | | 0,0087791 | 0,0091488 | 0,90 | 9,43E+08 | 9,29E+08 |
| 8,7 | | 0,0087797 | 0,0091450 | 0,52 | 5,38E+06 | 7,44E+06 |
| 47,7 | | 0,0087828 | 0,0091412 | 0,86 | 1,20E+08 | 1,11E+08 |
| 15,4 | | 0,0088406 | 0,0091893 | 1,24 | 4,27E+07 | 3,72E+07 |
| 22,6 | | 0,0088525 | 0,0091854 | 1,36 | 1,64E+08 | 1,86E+08 |
| 15,9 | | 0,0088647 | 0,0091816 | 0,16 | 3,96E+07 | 3,57E+07 |
| 4,6 | + | 0,0088970 | 0,0092564 | 3,94 | 3,43E+06 | 5,21E+06 |
| 14,6 | | 0,0089357 | 0,0092525 | 0,74 | 1,43E+07 | 1,55E+07 |
| 18,8 | | 0,0089427 | 0,0092486 | 0,78 | 3,17E+07 | 3,06E+07 |
| 14,6 | | 0,0089434 | 0,0092448 | 1,31 | 2,13E+07 | 2,48E+07 |
| 17,8 | | 0,0089474 | 0,0092409 | 1,20 | 1,34E+08 | 1,49E+08 |
| 35 | | 0,0089506 | 0,0092371 | 0,56 | 1,96E+08 | 2,30E+08 |

| | | | | | | |
|---|---|---|---|---|---|---|
| 22,5 | | 0,0089557 | 0,0092332 | 1,30 | 9,23E+07 | 1,10E+08 |
| 15,5 | | 0,0089748 | 0,0092294 | 2,01 | 4,83E+08 | 6,94E+08 |
| 48,5 | | 0,0089830 | 0,0092255 | 0,86 | 1,38E+09 | 1,40E+09 |
| 51,9 | | 0,0089902 | 0,0092217 | 0,90 | 3,30E+08 | 3,16E+08 |
| 18,3 | | 0,0089910 | 0,0092178 | 0,70 | 3,88E+08 | 3,97E+08 |
| 22,7 | | 0,0090127 | 0,0092523 | 1,26 | 1,42E+08 | 1,21E+08 |
| 48,7 | | 0,0090917 | 0,0092484 | 0,93 | 9,48E+09 | 9,05E+09 |
| 3,9 | | 0,0090964 | 0,0092446 | 0,70 | 4,21E+07 | 4,12E+07 |
| 36,4 | | 0,0091107 | 0,0092407 | 1,70 | 2,76E+07 | 3,91E+07 |
| 47,5 | | 0,0091124 | 0,0092369 | 1,14 | 2,13E+08 | 2,27E+08 |
| 21,5 | | 0,0091909 | 0,0092846 | 1,73 | 6,24E+06 | 8,16E+06 |
| 25,9 | | 0,0091939 | 0,0092807 | 0,70 | 2,61E+07 | 2,42E+07 |
| 21,3 | | 0,0092278 | 0,0093151 | 1,35 | 4,12E+07 | 3,45E+07 |
| 9,5 | | 0,0092402 | 0,0093112 | 0,29 | 1,11E+07 | 1,70E+07 |
| 19,8 | | 0,0092411 | 0,0093073 | 1,48 | 2,31E+07 | 2,35E+07 |
| 47,5 | | 0,0092456 | 0,0093035 | 0,66 | 2,14E+08 | 1,86E+08 |
| 52,2 | | 0,0092799 | 0,0092996 | 1,25 | 7,77E+08 | 7,49E+08 |
| 41,4 | | 0,0093007 | 0,0092958 | 0,02 | 9,27E+08 | 3,00E+08 |
| 16,5 | | 0,0093140 | 0,0093366 | 0,25 | 2,06E+07 | 2,10E+07 |
| 33,7 | | 0,0093166 | 0,0093328 | 1,50 | 7,95E+07 | 6,17E+07 |
| 31 | | 0,0093254 | 0,0093289 | 0,02 | 7,31E+07 | 9,32E+07 |
| 11,7 | | 0,0093334 | 0,0093251 | 10,13 | 1,44E+06 | 6,60E+06 |
| 33,2 | | 0,0093500 | 0,0093212 | 1,26 | 7,44E+07 | 7,71E+07 |
| 31,7 | | 0,0093554 | 0,0093174 | 0,70 | 4,40E+07 | 4,22E+07 |
| 23,9 | | 0,0093766 | 0,0093647 | 0,11 | 3,09E+07 | 3,45E+07 |
| 34,7 | | 0,0093787 | 0,0093609 | 1,95 | 3,26E+07 | 2,88E+07 |
| 41,1 | | 0,0093906 | 0,0093570 | 0,79 | 3,09E+08 | 3,47E+08 |
| 24,4 | | 0,0093958 | 0,0093531 | 0,78 | 6,18E+08 | 5,58E+08 |
| 49,5 | | 0,0094110 | 0,0094004 | 0,72 | 2,51E+09 | 2,72E+09 |
| 23,2 | | 0,0094277 | 0,0093965 | 0,76 | 6,87E+07 | 7,74E+07 |
| 67,5 | | 0,0094337 | 0,0093927 | 0,79 | 5,39E+09 | 5,47E+09 |
| 47,3 | | 0,0094633 | 0,0094267 | 0,91 | 2,51E+08 | 2,57E+08 |
| 78,4 | | 0,0094792 | 0,0094228 | 1,21 | 1,04E+08 | 1,13E+08 |
| 72,2 | | 0,0094797 | 0,0094189 | 1,53 | 7,01E+09 | 8,02E+09 |
| 32,2 | | 0,0094799 | 0,0094151 | 0,62 | 7,64E+07 | 5,76E+07 |
| 29,9 | | 0,0094996 | 0,0094112 | 0,86 | 4,49E+08 | 4,49E+08 |
| 28,1 | | 0,0095022 | 0,0094073 | 0,80 | 8,41E+07 | 8,51E+07 |
| 17,9 | | 0,0095201 | 0,0094429 | 3,05 | 9,83E+06 | 4,69E+06 |
| 59,7 | | 0,0095409 | 0,0094390 | 1,23 | 5,66E+08 | 6,37E+08 |
| 12,9 | | 0,0095521 | 0,0094351 | 0,69 | 3,46E+07 | 2,83E+07 |
| 35 | | 0,0095531 | 0,0094313 | 1,17 | 1,31E+08 | 1,36E+08 |
| 21,6 | | 0,0095595 | 0,0094274 | 0,57 | 2,14E+08 | 1,76E+08 |
| 13,8 | | 0,0095635 | 0,0094235 | 0,61 | 1,13E+07 | 1,05E+07 |
| 44,7 | | 0,0095637 | 0,0094197 | 0,24 | 4,03E+08 | 1,64E+08 |
| 21,2 | | 0,0095646 | 0,0094158 | 0,06 | 1,14E+08 | 1,21E+08 |
| 37,2 | | 0,0095670 | 0,0094120 | 0,80 | 8,07E+08 | 8,44E+08 |
| 20,2 | | 0,0095703 | 0,0094081 | 0,55 | 4,26E+07 | 5,74E+07 |
| 20,6 | | 0,0096051 | 0,0094877 | 0,70 | 6,19E+07 | 6,83E+07 |
| 12 | | 0,0096201 | 0,0094838 | 0,86 | 1,78E+07 | 1,87E+07 |
| 14,2 | | 0,0096301 | 0,0094800 | 0,72 | 1,01E+07 | 1,07E+07 |

| | | | | | | |
|---|---|---|---|---|---|---|
| 44,4 | | 0,0096703 | 0,0094761 | 1,44 | 1,84E+08 | 2,28E+08 |
| 15,1 | | 0,0096726 | 0,0094722 | 5,13 | 3,74E+06 | 3,79E+06 |
| 52,2 | | 0,0096752 | 0,0094684 | 0,82 | 1,03E+09 | 1,13E+09 |
| 12,1 | | 0,0096785 | 0,0094645 | 1,60 | 3,19E+07 | 3,79E+07 |
| 36,8 | | 0,0097017 | 0,0095047 | 1,98 | 1,46E+07 | 1,40E+07 |
| 3,7 | | 0,0097159 | 0,0095008 | 1,68 | 3,80E+06 | 3,35E+06 |
| 33,3 | | 0,0097164 | 0,0094969 | 1,53 | 6,88E+07 | 6,66E+07 |
| 54,3 | | 0,0097513 | 0,0094931 | 0,76 | 3,06E+09 | 2,68E+09 |
| 38,4 | | 0,0097515 | 0,0094892 | 0,73 | 2,79E+08 | 3,29E+08 |
| 45,4 | | 0,0097544 | 0,0094853 | 0,65 | 1,82E+08 | 1,61E+08 |
| 53,3 | | 0,0097577 | 0,0094815 | 1,22 | 2,31E+08 | 2,03E+08 |
| 15,7 | | 0,0097640 | 0,0094776 | 1,77 | 4,96E+07 | 5,29E+07 |
| 46,1 | | 0,0097735 | 0,0094738 | 14,69 | 3,76E+06 | 2,63E+07 |
| 49,1 | | 0,0098268 | 0,0094699 | 1,15 | 2,45E+08 | 2,55E+08 |
| 62,4 | | 0,0098316 | 0,0094661 | 1,48 | 1,56E+08 | 1,22E+08 |
| 23,3 | | 0,0098415 | 0,0094622 | 1,61 | 1,50E+08 | 1,18E+08 |
| 20,6 | | 0,0098543 | 0,0094584 | 0,82 | 2,26E+08 | 2,40E+08 |
| 20,5 | | 0,0098657 | 0,0094545 | 0,73 | 3,23E+07 | 3,33E+07 |
| 22,5 | | 0,0098677 | 0,0094507 | 1,55 | 6,17E+07 | 5,21E+07 |
| 2,5 | | 0,0098747 | 0,0094469 | 0,48 | 3,09E+06 | 2,07E+06 |
| 10,5 | | 0,0099099 | 0,0094430 | 0,25 | 1,39E+07 | 1,52E+07 |
| 44,3 | | 0,0099203 | 0,0094392 | 1,81 | 5,94E+07 | 5,00E+07 |
| 27,6 | | 0,0099301 | 0,0094354 | 0,69 | 2,77E+08 | 3,54E+08 |
| 15,4 | | 0,0099323 | 0,0094316 | 2,88 | 6,10E+06 | 1,00E+07 |
| 20,3 | | 0,0099582 | 0,0095233 | 1,27 | 1,42E+08 | 1,22E+08 |
| 42,9 | | 0,0099620 | 0,0095194 | 1,22 | 8,10E+08 | 8,90E+08 |
| 14,1 | | 0,0100230 | 0,0095528 | 1,81 | 2,05E+07 | 1,74E+07 |
| 64,1 | | 0,0100234 | 0,0095489 | 1,25 | 9,65E+08 | 8,55E+08 |
| 34,8 | | 0,0100245 | 0,0095451 | 0,74 | 1,13E+08 | 1,39E+08 |
| 37,5 | | 0,0100314 | 0,0095412 | 1,26 | 1,41E+08 | 1,55E+08 |
| 50 | | 0,0100373 | 0,0095809 | 0,83 | 9,97E+07 | 9,83E+07 |
| 35,5 | | 0,0100447 | 0,0095771 | 0,80 | 1,38E+08 | 1,61E+08 |
| 25,5 | | 0,0100501 | 0,0095732 | 1,58 | 4,60E+07 | 6,03E+07 |
| 33,5 | | 0,0100542 | 0,0096194 | 0,83 | 1,94E+08 | 1,98E+08 |
| 35,1 | | 0,0100551 | 0,0096155 | 1,28 | 1,07E+08 | 1,05E+08 |
| 42,1 | | 0,0100634 | 0,0096116 | 0,67 | 1,99E+08 | 1,57E+08 |
| 19,1 | | 0,0100692 | 0,0096077 | 1,43 | 1,72E+07 | 1,73E+07 |
| 14,5 | | 0,0100922 | 0,0096039 | 1,25 | 3,51E+07 | 3,58E+07 |
| 5,5 | | 0,0100926 | 0,0096000 | 0,28 | 8,26E+06 | 9,33E+06 |
| 43,5 | | 0,0100933 | 0,0095961 | 0,52 | 2,16E+08 | 2,38E+08 |
| 24,2 | | 0,0101083 | 0,0095923 | 0,42 | 3,92E+07 | 5,19E+07 |
| 27,6 | | 0,0101119 | 0,0095884 | 1,23 | 9,64E+07 | 1,07E+08 |
| 48,2 | | 0,0101174 | 0,0095846 | 0,79 | 1,02E+09 | 9,96E+08 |
| 42 | | 0,0101302 | 0,0095807 | 0,82 | 6,11E+08 | 5,36E+08 |
| 32,2 | | 0,0101363 | 0,0095769 | 1,23 | 5,75E+08 | 6,65E+08 |
| 34,6 | | 0,0101705 | 0,0096228 | 1,21 | 1,46E+08 | 1,48E+08 |
| 16,4 | | 0,0102304 | 0,0096623 | 1,55 | 3,25E+08 | 4,33E+08 |
| 52,2 | | 0,0102360 | 0,0096584 | 0,79 | 2,89E+08 | 3,07E+08 |
| 44 | | 0,0102536 | 0,0096930 | 0,85 | 9,32E+07 | 8,30E+07 |
| 24,9 | | 0,0102538 | 0,0096891 | 0,69 | 1,34E+08 | 1,28E+08 |

| | | | | | | |
|---|---|---|---|---|---|---|
| 40,3 | | 0,0102575 | 0,0096852 | 0,80 | 6,54E+07 | 5,96E+07 |
| 33,2 | | 0,0102747 | 0,0097310 | 0,85 | 6,47E+08 | 5,78E+08 |
| 19,9 | | 0,0102800 | 0,0097271 | 0,84 | 4,19E+07 | 3,97E+07 |
| 39,3 | | 0,0102962 | 0,0097232 | 0,59 | 6,46E+07 | 7,40E+07 |
| 28,4 | | 0,0103649 | 0,0097961 | 1,23 | 1,41E+08 | 1,55E+08 |
| 21,2 | + | 0,0103674 | 0,0097922 | 2,37 | 9,42E+06 | 7,31E+06 |
| 36,6 | | 0,0104237 | 0,0098698 | 0,94 | 5,81E+08 | 5,67E+08 |
| 36 | | 0,0104412 | 0,0098658 | 0,11 | 2,92E+07 | 3,45E+07 |
| 28,9 | | 0,0104591 | 0,0098986 | 1,48 | 3,73E+07 | 4,77E+07 |
| 39,6 | | 0,0104811 | 0,0099441 | 0,36 | 5,80E+07 | 6,26E+07 |
| 85,9 | | 0,0104998 | 0,0099402 | 0,70 | 1,07E+09 | 1,33E+09 |
| 24,4 | | 0,0105038 | 0,0099362 | 1,35 | 2,97E+07 | 3,64E+07 |
| 24,7 | | 0,0105127 | 0,0099322 | 0,75 | 1,03E+08 | 1,28E+08 |
| 45,1 | | 0,0105281 | 0,0099283 | 0,65 | 2,07E+08 | 1,86E+08 |
| 45,9 | | 0,0105296 | 0,0099243 | 0,87 | 5,74E+08 | 6,05E+08 |
| 48,5 | | 0,0105472 | 0,0099697 | 1,66 | 4,04E+07 | 4,50E+07 |
| 14,6 | + | 0,0105588 | 0,0099658 | 20,85 | 1,79E+06 | 1,18E+07 |
| 48,4 | | 0,0105840 | 0,0099618 | 0,82 | 3,07E+08 | 2,97E+08 |
| 0,7 | | 0,0105912 | 0,0100565 | 0,34 | 4,87E+06 | 6,27E+06 |
| 28,4 | | 0,0106011 | 0,0100525 | 1,46 | 4,80E+07 | 4,18E+07 |
| 8,7 | | 0,0106112 | 0,0100485 | 0,20 | 4,09E+07 | 2,80E+07 |
| 20,5 | | 0,0106358 | 0,0100445 | 1,39 | 8,66E+07 | 7,19E+07 |
| 58,6 | | 0,0106373 | 0,0100405 | 0,80 | 1,70E+08 | 1,78E+08 |
| 16,5 | | 0,0106451 | 0,0100857 | 0,91 | 5,65E+07 | 5,62E+07 |
| 30,4 | | 0,0106475 | 0,0100817 | 1,24 | 8,82E+07 | 7,65E+07 |
| 24,8 | | 0,0106809 | 0,0100777 | 1,28 | 1,39E+07 | 1,19E+07 |
| 34,3 | | 0,0107448 | 0,0101482 | 1,30 | 3,52E+07 | 3,55E+07 |
| 15,3 | | 0,0107592 | 0,0101442 | 0,65 | 1,50E+07 | 1,79E+07 |
| 38,4 | | 0,0107598 | 0,0101402 | 0,79 | 2,35E+08 | 2,18E+08 |
| 51,1 | | 0,0107746 | 0,0101362 | 1,16 | 1,08E+08 | 1,00E+08 |
| 10 | | 0,0107766 | 0,0101322 | 1,61 | 2,34E+07 | 2,26E+07 |
| 48,8 | | 0,0107928 | 0,0101661 | 1,24 | 1,65E+08 | 1,73E+08 |
| 49 | | 0,0107953 | 0,0101621 | 0,62 | 1,26E+09 | 9,00E+08 |
| 31 | | 0,0108110 | 0,0101581 | 1,04 | 1,52E+08 | 1,50E+08 |
| 44,9 | | 0,0108316 | 0,0101541 | 1,20 | 5,30E+07 | 5,35E+07 |
| 28,6 | | 0,0108362 | 0,0101501 | 1,46 | 9,12E+07 | 1,02E+08 |
| 46,2 | | 0,0108462 | 0,0101461 | 0,78 | 1,32E+08 | 1,21E+08 |
| 36,2 | | 0,0108473 | 0,0101421 | 1,35 | 3,79E+08 | 4,29E+08 |
| 17,3 | | 0,0108617 | 0,0101381 | 1,20 | 4,42E+07 | 4,01E+07 |
| 18,9 | | 0,0109003 | 0,0101830 | 2,91 | 1,41E+07 | 1,34E+07 |
| 32,7 | | 0,0109542 | 0,0102278 | 1,70 | 1,60E+07 | 1,17E+07 |
| 29,4 | | 0,0109997 | 0,0103215 | 0,66 | 7,18E+07 | 6,53E+07 |
| 8,1 | | 0,0110073 | 0,0103553 | 1,44 | 8,84E+06 | 1,06E+07 |
| 8 | | 0,0110266 | 0,0103512 | 0,39 | 7,86E+06 | 1,31E+07 |
| 56,9 | | 0,0110627 | 0,0103959 | 0,63 | 1,20E+08 | 1,07E+08 |
| 6,7 | | 0,0110651 | 0,0103918 | 0,11 | 3,25E+07 | 3,80E+07 |
| 40 | | 0,0111077 | 0,0103877 | 0,59 | 2,82E+07 | 3,42E+07 |
| 10,4 | | 0,0111239 | 0,0104324 | 1,36 | 1,37E+07 | 1,55E+07 |
| 17,4 | | 0,0111557 | 0,0104283 | 0,27 | 8,70E+07 | 8,19E+07 |
| 25 | | 0,0111668 | 0,0104729 | 0,67 | 7,32E+07 | 6,41E+07 |

| | | | | | | |
|---|---|---|---|---|---|---|
| 21 | | 0,0111962 | 0,0104688 | 2,30 | 1,43E+07 | 1,15E+07 |
| 6 | | 0,0112319 | 0,0105432 | 0,63 | 7,08E+06 | 6,14E+06 |
| 13,1 | | 0,0112356 | 0,0105390 | 0,47 | 9,95E+06 | 1,11E+07 |
| 53,4 | | 0,0112402 | 0,0105349 | 0,86 | 1,09E+09 | 1,12E+09 |
| 63,4 | | 0,0112454 | 0,0105308 | 0,72 | 3,37E+08 | 3,87E+08 |
| 35,1 | | 0,0112654 | 0,0105266 | 2,17 | 9,22E+07 | 7,21E+07 |
| 8,6 | | 0,0112849 | 0,0105711 | 0,16 | 1,92E+07 | 3,78E+07 |
| 39,2 | | 0,0112923 | 0,0105670 | 0,07 | 5,76E+07 | 6,57E+07 |
| 15,6 | | 0,0112924 | 0,0105628 | 0,72 | 6,06E+07 | 6,49E+07 |
| 14,5 | | 0,0112937 | 0,0105587 | 1,99 | 1,02E+07 | 1,31E+07 |
| 11,5 | | 0,0112987 | 0,0105546 | 0,42 | 1,86E+07 | 3,09E+07 |
| 37 | | 0,0113060 | 0,0105504 | 1,49 | 1,84E+08 | 1,82E+08 |
| 55,1 | | 0,0113076 | 0,0105463 | 1,45 | 2,96E+08 | 2,64E+08 |
| 22,1 | | 0,0113304 | 0,0105797 | 0,22 | 9,11E+06 | 2,13E+07 |
| 21,5 | | 0,0113532 | 0,0105756 | 0,06 | 1,50E+08 | 5,07E+07 |
| 5,9 | | 0,0113732 | 0,0106073 | 0,06 | 1,31E+08 | 1,37E+08 |
| 81,4 | | 0,0113780 | 0,0106032 | 1,44 | 8,22E+08 | 8,02E+08 |
| 30,5 | | 0,0114050 | 0,0106474 | 1,85 | 1,91E+07 | 2,17E+07 |
| 17,8 | | 0,0114147 | 0,0106433 | 0,76 | 1,07E+08 | 1,09E+08 |
| 35,4 | | 0,0114351 | 0,0106391 | 0,78 | 2,67E+08 | 2,35E+08 |
| 49,8 | | 0,0114361 | 0,0106350 | 0,52 | 9,48E+07 | 1,52E+08 |
| 18,8 | | 0,0114619 | 0,0106308 | 0,64 | 3,43E+07 | 3,24E+07 |
| 51 | + | 0,0114809 | 0,0106750 | 7,48 | 4,08E+08 | 3,85E+08 |
| 31,2 | | 0,0114880 | 0,0107066 | 0,74 | 2,02E+08 | 1,94E+08 |
| 30,5 | | 0,0114951 | 0,0107025 | 1,29 | 1,29E+08 | 1,53E+08 |
| 27,9 | | 0,0114960 | 0,0106983 | 0,13 | 3,36E+07 | 2,78E+07 |
| 18,5 | | 0,0114981 | 0,0106941 | 0,58 | 5,22E+07 | 5,61E+07 |
| 13,5 | | 0,0115063 | 0,0106900 | 0,13 | 1,70E+07 | 7,99E+06 |
| 37,7 | | 0,0115228 | 0,0106858 | 0,79 | 3,94E+08 | 3,55E+08 |
| 46,2 | | 0,0115259 | 0,0106817 | 1,21 | 9,65E+08 | 9,90E+08 |
| 59,9 | | 0,0115519 | 0,0106775 | 1,29 | 1,16E+09 | 1,22E+09 |
| 13,5 | | 0,0115613 | 0,0107091 | 1,41 | 5,21E+07 | 5,02E+07 |
| 26 | | 0,0115771 | 0,0107049 | 0,83 | 3,78E+08 | 4,00E+08 |
| 28,2 | | 0,0115910 | 0,0107426 | 0,65 | 6,06E+07 | 6,55E+07 |
| 10,5 | | 0,0116099 | 0,0107757 | 0,49 | 7,04E+07 | 7,58E+07 |
| 11,4 | | 0,0116355 | 0,0107715 | 0,11 | 9,24E+07 | 7,32E+07 |
| 29,9 | | 0,0116467 | 0,0107673 | 1,52 | 2,50E+07 | 2,94E+07 |
| 37,8 | | 0,0116703 | 0,0107632 | 0,14 | 2,15E+07 | 1,67E+07 |
| 30 | | 0,0116719 | 0,0107590 | 0,70 | 1,56E+08 | 1,79E+08 |
| 27 | | 0,0116742 | 0,0107548 | 1,40 | 4,20E+08 | 4,26E+08 |
| 15,5 | | 0,0116863 | 0,0107507 | 0,55 | 2,26E+07 | 2,23E+07 |
| 39,2 | | 0,0116987 | 0,0107465 | 1,33 | 2,12E+08 | 2,57E+08 |
| 18,9 | | 0,0117087 | 0,0107424 | 1,56 | 2,54E+07 | 2,86E+07 |
| 13,6 | | 0,0117298 | 0,0107382 | 6,94 | 1,26E+07 | 3,51E+07 |
| 53,9 | | 0,0117319 | 0,0107341 | 1,37 | 5,49E+08 | 5,45E+08 |
| 11,8 | | 0,0117390 | 0,0107299 | 3,25 | 2,74E+06 | 6,17E+06 |
| 18,7 | | 0,0117791 | 0,0107628 | 0,18 | 1,76E+07 | 1,95E+07 |
| 31,7 | | 0,0117888 | 0,0107587 | 0,75 | 7,57E+07 | 8,20E+07 |
| 38,7 | | 0,0118087 | 0,0108655 | 1,34 | 8,69E+07 | 8,15E+07 |
| 50,7 | | 0,0118115 | 0,0108613 | 1,12 | 9,14E+08 | 9,49E+08 |

| | | | | | | |
|---|---|---|---|---|---|---|
| 20,5 | | 0,0118229 | 0,0108571 | 0,48 | 2,21E+07 | 2,06E+07 |
| 22,1 | | 0,0118233 | 0,0108530 | 2,93 | 7,94E+07 | 3,79E+07 |
| 3,9 | | 0,0118474 | 0,0108857 | 0,36 | 2,42E+07 | 2,33E+07 |
| 6,4 | | 0,0118970 | 0,0109231 | 0,40 | 4,99E+06 | 6,17E+06 |
| 49,4 | | 0,0119067 | 0,0109666 | 0,93 | 3,04E+08 | 2,99E+08 |
| 22,6 | | 0,0120345 | 0,0111207 | 1,46 | 6,70E+07 | 8,45E+07 |
| 43,2 | | 0,0120558 | 0,0111656 | 0,75 | 6,13E+08 | 5,21E+08 |
| 39,2 | | 0,0120943 | 0,0112089 | 1,44 | 7,15E+07 | 9,34E+07 |
| 23 | | 0,0120943 | 0,0112046 | 2,84 | 6,31E+06 | 9,50E+06 |
| 10,7 | | 0,0121075 | 0,0112003 | 1,67 | 2,69E+07 | 2,11E+07 |
| 20,2 | | 0,0121284 | 0,0111960 | 5,51 | 1,09E+07 | 1,15E+07 |
| 56 | | 0,0121361 | 0,0111917 | 0,89 | 2,36E+09 | 2,56E+09 |
| 43,2 | | 0,0121503 | 0,0112288 | 0,77 | 1,89E+08 | 1,61E+08 |
| 2,6 | | 0,0121620 | 0,0112245 | 2,56 | 1,25E+07 | 1,26E+07 |
| 41,1 | | 0,0122004 | 0,0112984 | 0,74 | 3,53E+08 | 3,48E+08 |
| 18,5 | | 0,0122293 | 0,0113675 | 1,31 | 3,10E+07 | 3,00E+07 |
| 18,2 | | 0,0122393 | 0,0113999 | 5,17 | 3,46E+06 | 1,55E+06 |
| 24,1 | | 0,0123162 | 0,0115272 | 1,32 | 3,15E+07 | 3,68E+07 |
| 38,1 | | 0,0123248 | 0,0116054 | 1,18 | 8,73E+07 | 9,29E+07 |
| 46,6 | | 0,0123777 | 0,0116376 | 0,69 | 3,68E+08 | 3,42E+08 |
| 28,8 | | 0,0124144 | 0,0116332 | 2,30 | 1,61E+07 | 1,56E+07 |
| 60,6 | | 0,0124230 | 0,0116287 | 0,78 | 2,48E+08 | 2,47E+08 |
| 57,7 | | 0,0124234 | 0,0116243 | 0,53 | 5,18E+07 | 7,59E+07 |
| 17,2 | | 0,0124602 | 0,0116198 | 1,63 | 5,64E+07 | 4,10E+07 |
| 50 | | 0,0124840 | 0,0116994 | 0,87 | 2,66E+09 | 2,57E+09 |
| 15,1 | | 0,0124938 | 0,0117300 | 3,21 | 1,91E+07 | 1,28E+07 |
| 16,1 | | 0,0124969 | 0,0117255 | 1,68 | 2,58E+07 | 2,06E+07 |
| 50,4 | | 0,0125301 | 0,0117561 | 0,81 | 9,58E+08 | 1,12E+09 |
| 42,5 | | 0,0125540 | 0,0117989 | 0,81 | 4,86E+08 | 5,45E+08 |
| 23,9 | | 0,0125799 | 0,0117944 | 1,30 | 2,02E+07 | 1,71E+07 |
| 29,1 | | 0,0125799 | 0,0117899 | 0,81 | 1,18E+08 | 1,08E+08 |
| 12,6 | | 0,0125823 | 0,0118219 | 0,61 | 4,75E+06 | 5,94E+06 |
| 48,4 | | 0,0125849 | 0,0118174 | 0,87 | 2,20E+09 | 2,19E+09 |
| 69,3 | | 0,0126010 | 0,0118129 | 1,19 | 1,27E+10 | 1,30E+10 |
| 32,6 | | 0,0126042 | 0,0118084 | 0,15 | 3,08E+07 | 4,07E+07 |
| 34 | | 0,0126053 | 0,0118040 | 0,64 | 2,36E+07 | 2,74E+07 |
| 37,2 | | 0,0126322 | 0,0117995 | 0,65 | 2,17E+08 | 2,67E+08 |
| 34,3 | | 0,0126430 | 0,0118421 | 1,27 | 6,93E+07 | 5,88E+07 |
| 22,6 | | 0,0126486 | 0,0118376 | 0,81 | 6,58E+07 | 6,98E+07 |
| 53,4 | | 0,0126514 | 0,0118331 | 1,34 | 8,38E+08 | 7,10E+08 |
| 35,7 | | 0,0126534 | 0,0118286 | 0,68 | 1,26E+08 | 1,62E+08 |
| 16,6 | | 0,0126588 | 0,0118241 | 0,42 | 2,28E+07 | 2,82E+07 |
| 45,8 | | 0,0126614 | 0,0118560 | 1,24 | 1,23E+09 | 1,42E+09 |
| 10,2 | | 0,0126659 | 0,0118515 | 0,58 | 1,41E+07 | 1,18E+07 |
| 18,2 | | 0,0127128 | 0,0118834 | 2,28 | 2,31E+07 | 1,70E+07 |
| 11 | | 0,0127227 | 0,0118789 | 1,44 | 1,30E+07 | 1,43E+07 |
| 8 | | 0,0127295 | 0,0118744 | 2,26 | 4,10E+06 | 3,80E+06 |
| 30,2 | | 0,0127558 | 0,0118699 | 1,66 | 7,79E+07 | 1,04E+08 |
| 48,7 | | 0,0127784 | 0,0118654 | 1,12 | 1,07E+09 | 1,02E+09 |
| 22,1 | | 0,0127862 | 0,0118609 | 0,82 | 6,13E+07 | 5,47E+07 |

| | | | | | | |
|---|---|---|---|---|---|---|
| 15,2 | | 0,0128259 | 0,0120151 | 1,82 | 7,82E+06 | 9,80E+06 |
| 24,9 | | 0,0128374 | 0,0120514 | 1,83 | 3,78E+07 | 5,35E+07 |
| 91,4 | | 0,0128460 | 0,0120468 | 1,45 | 6,08E+09 | 5,27E+09 |
| 11,2 | | 0,0128541 | 0,0120423 | 1,42 | 3,64E+06 | 3,43E+06 |
| 25,3 | | 0,0128627 | 0,0120377 | 1,26 | 7,13E+07 | 8,49E+07 |
| 29,6 | | 0,0128663 | 0,0120332 | 0,66 | 1,35E+08 | 1,27E+08 |
| 25 | | 0,0129198 | 0,0120754 | 1,98 | 3,38E+07 | 5,55E+07 |
| 13,4 | | 0,0129640 | 0,0121176 | 0,73 | 3,22E+07 | 3,20E+07 |
| 23,8 | | 0,0129940 | 0,0121883 | 0,63 | 4,11E+07 | 3,97E+07 |
| 52,5 | | 0,0130041 | 0,0122319 | 0,87 | 5,75E+09 | 5,48E+09 |
| 5,1 | | 0,0131042 | 0,0123628 | 0,45 | 4,94E+06 | 7,38E+06 |
| 59,2 | | 0,0131146 | 0,0124048 | 0,87 | 3,73E+08 | 3,48E+08 |
| 57 | | 0,0131165 | 0,0124002 | 1,22 | 7,89E+08 | 8,81E+08 |
| 50,1 | | 0,0131494 | 0,0123955 | 0,82 | 2,99E+08 | 3,02E+08 |
| 1,7 | | 0,0131515 | 0,0123908 | 1,46 | 1,54E+07 | 1,71E+07 |
| 60,3 | | 0,0131660 | 0,0124793 | 0,85 | 1,62E+08 | 1,63E+08 |
| 23,3 | | 0,0131844 | 0,0125227 | 0,80 | 5,15E+07 | 4,39E+07 |
| 16,1 | | 0,0132139 | 0,0125180 | 0,69 | 3,78E+08 | 3,60E+08 |
| 46,6 | | 0,0132392 | 0,0125133 | 1,22 | 9,47E+08 | 9,31E+08 |
| 28,7 | | 0,0132421 | 0,0125086 | 0,87 | 2,12E+08 | 2,24E+08 |
| 28,7 | | 0,0132461 | 0,0125039 | 1,21 | 3,23E+08 | 3,07E+08 |
| 13,4 | | 0,0132505 | 0,0124993 | 0,52 | 1,96E+07 | 2,67E+07 |
| 31,5 | | 0,0132522 | 0,0124946 | 0,74 | 2,72E+08 | 2,92E+08 |
| 52,6 | | 0,0132661 | 0,0125303 | 0,74 | 5,18E+08 | 6,08E+08 |
| 50 | | 0,0132672 | 0,0125256 | 1,35 | 3,89E+09 | 3,36E+09 |
| 44,3 | | 0,0132828 | 0,0125210 | 1,42 | 6,68E+08 | 7,82E+08 |
| 40,1 | | 0,0132943 | 0,0125567 | 1,34 | 9,70E+07 | 1,08E+08 |
| 16,3 | | 0,0133086 | 0,0125520 | 6,83 | 4,46E+06 | 1,98E+07 |
| 34,1 | | 0,0133563 | 0,0125473 | 1,79 | 2,49E+07 | 3,33E+07 |
| 7,3 | | 0,0133669 | 0,0125770 | 0,59 | 4,91E+07 | 3,87E+07 |
| 29,7 | | 0,0133704 | 0,0125723 | 0,76 | 4,34E+08 | 4,45E+08 |
| 27,9 | | 0,0133834 | 0,0126378 | 0,08 | 2,72E+07 | 3,43E+07 |
| 10,6 | | 0,0134184 | 0,0126734 | 1,67 | 1,17E+07 | 1,73E+07 |
| 9,8 | | 0,0134289 | 0,0126687 | 1,64 | 1,73E+07 | 1,48E+07 |
| 13,8 | | 0,0134302 | 0,0126639 | 0,88 | 1,92E+08 | 1,90E+08 |
| 15 | | 0,0134428 | 0,0126592 | 0,69 | 9,90E+07 | 7,69E+07 |
| 57 | | 0,0134620 | 0,0126545 | 1,40 | 8,72E+07 | 9,52E+07 |
| 48,8 | | 0,0134966 | 0,0126841 | 1,28 | 2,37E+08 | 2,14E+08 |
| 27,5 | | 0,0135119 | 0,0126793 | 0,05 | 1,62E+08 | 2,45E+07 |
| 28,3 | | 0,0135316 | 0,0126746 | 0,64 | 4,71E+08 | 4,59E+08 |
| 11,8 | | 0,0135561 | 0,0127101 | 0,64 | 2,27E+08 | 2,03E+08 |
| 24,5 | | 0,0135585 | 0,0127054 | 1,15 | 1,41E+08 | 1,42E+08 |
| 58,5 | | 0,0135651 | 0,0127006 | 1,34 | 3,33E+09 | 3,56E+09 |
| 52,5 | | 0,0136185 | 0,0127822 | 1,39 | 3,11E+08 | 3,49E+08 |
| 47,5 | | 0,0136293 | 0,0127774 | 1,79 | 8,67E+07 | 1,15E+08 |
| 5,1 | | 0,0136429 | 0,0127727 | 0,18 | 1,82E+07 | 1,89E+07 |
| 21,7 | | 0,0136436 | 0,0127679 | 0,85 | 2,32E+08 | 2,34E+08 |
| 49,4 | | 0,0136439 | 0,0127632 | 0,79 | 2,96E+08 | 2,58E+08 |
| 21,3 | | 0,0136470 | 0,0127584 | 1,21 | 2,67E+07 | 2,69E+07 |
| 59,8 | | 0,0136475 | 0,0127537 | 0,89 | 8,36E+08 | 9,00E+08 |

| | | | | | | |
|---|---|---|---|---|---|---|
| 25,3 | | 0,0136514 | 0,0127490 | 1,33 | 1,31E+08 | 1,19E+08 |
| 27,1 | | 0,0136557 | 0,0127443 | 1,29 | 1,31E+08 | 1,31E+08 |
| 9 | | 0,0136638 | 0,0127395 | 1,65 | 1,70E+07 | 1,29E+07 |
| 74,2 | | 0,0136775 | 0,0127348 | 0,78 | 1,27E+10 | 1,39E+10 |
| 32,6 | | 0,0136797 | 0,0127301 | 1,28 | 2,72E+08 | 2,52E+08 |
| 50,6 | | 0,0137265 | 0,0127609 | 0,68 | 2,31E+08 | 1,84E+08 |
| 38,5 | | 0,0137655 | 0,0128021 | 1,39 | 4,34E+08 | 5,39E+08 |
| 39,1 | | 0,0138076 | 0,0129231 | 1,41 | 1,35E+08 | 1,72E+08 |
| 35,3 | | 0,0138220 | 0,0129183 | 1,49 | 5,90E+07 | 5,78E+07 |
| 54,9 | | 0,0138240 | 0,0129135 | 0,84 | 1,63E+09 | 1,60E+09 |
| 23,7 | | 0,0138317 | 0,0129088 | 24,56 | 9,53E+06 | 5,76E+06 |
| 26 | | 0,0138581 | 0,0129513 | 0,79 | 3,05E+08 | 2,70E+08 |
| 49,1 | | 0,0138656 | 0,0129465 | 0,75 | 2,74E+08 | 3,31E+08 |
| 35,9 | | 0,0138725 | 0,0129417 | 1,30 | 4,99E+07 | 5,99E+07 |
| 16,2 | | 0,0138891 | 0,0129369 | 2,16 | 1,67E+07 | 1,45E+07 |
| 16,3 | | 0,0139069 | 0,0129322 | 0,67 | 3,97E+07 | 5,05E+07 |
| 39,7 | | 0,0139241 | 0,0129274 | 0,59 | 1,30E+08 | 1,02E+08 |
| 52,4 | | 0,0139271 | 0,0129226 | 0,71 | 1,72E+08 | 2,21E+08 |
| 30,8 | | 0,0139489 | 0,0129532 | 1,50 | 1,83E+07 | 1,77E+07 |
| 16,5 | | 0,0139600 | 0,0129485 | 0,75 | 3,17E+07 | 2,85E+07 |
| 15 | | 0,0139887 | 0,0129437 | 0,09 | 4,44E+07 | 3,83E+07 |
| 36 | | 0,0140034 | 0,0129742 | 0,77 | 1,08E+08 | 1,11E+08 |
| 34 | | 0,0140158 | 0,0129695 | 0,74 | 7,27E+08 | 8,76E+08 |
| 37,4 | | 0,0140289 | 0,0130103 | 0,02 | 2,21E+08 | 6,02E+08 |
| 33,4 | | 0,0140929 | 0,0130408 | 0,81 | 9,56E+08 | 9,46E+08 |
| 11,4 | | 0,0140930 | 0,0130360 | 0,60 | 1,15E+07 | 1,34E+07 |
| 8,5 | | 0,0141063 | 0,0130312 | 0,31 | 2,22E+07 | 1,68E+07 |
| 68,2 | | 0,0141122 | 0,0130264 | 1,56 | 2,81E+08 | 2,29E+08 |
| 53,2 | | 0,0141256 | 0,0130613 | 0,91 | 2,69E+08 | 2,67E+08 |
| 36,8 | | 0,0141603 | 0,0130565 | 0,75 | 3,22E+08 | 2,86E+08 |
| 31,7 | | 0,0141746 | 0,0130517 | 1,27 | 1,03E+08 | 1,04E+08 |
| 30,4 | | 0,0141813 | 0,0130469 | 1,50 | 1,30E+07 | 1,49E+07 |
| 25,8 | | 0,0142603 | 0,0131609 | 1,41 | 1,39E+08 | 1,69E+08 |
| 32,3 | | 0,0142613 | 0,0131560 | 0,51 | 5,91E+07 | 9,64E+07 |
| 21,2 | | 0,0142681 | 0,0131512 | 0,19 | 1,49E+07 | 2,18E+07 |
| 34,5 | | 0,0142693 | 0,0131464 | 0,86 | 1,04E+08 | 1,09E+08 |
| 37,8 | | 0,0142765 | 0,0131416 | 0,80 | 4,14E+08 | 4,86E+08 |
| 22,7 | | 0,0142767 | 0,0131368 | 1,38 | 3,23E+07 | 3,54E+07 |
| 31,6 | | 0,0143041 | 0,0131320 | 1,67 | 3,62E+07 | 3,06E+07 |
| 46,5 | | 0,0143174 | 0,0131272 | 1,32 | 1,42E+08 | 1,70E+08 |
| 69,8 | | 0,0143646 | 0,0131911 | 1,29 | 2,96E+09 | 2,47E+09 |
| 25,4 | | 0,0143927 | 0,0132316 | 1,34 | 6,42E+07 | 6,12E+07 |
| 16,9 | | 0,0144558 | 0,0132267 | 0,70 | 9,44E+07 | 8,06E+07 |
| 58,2 | | 0,0144603 | 0,0132219 | 0,77 | 3,35E+08 | 3,53E+08 |
| 43,6 | | 0,0144636 | 0,0132171 | 0,87 | 6,33E+08 | 6,98E+08 |
| 24,6 | | 0,0144727 | 0,0132123 | 0,24 | 1,47E+07 | 1,87E+07 |
| 13,3 | | 0,0144759 | 0,0132074 | 0,37 | 6,04E+07 | 7,46E+07 |
| 36,3 | | 0,0144959 | 0,0132026 | 0,70 | 2,36E+08 | 2,30E+08 |
| 55,1 | | 0,0144994 | 0,0131978 | 1,40 | 9,93E+08 | 9,74E+08 |
| 50 | | 0,0145218 | 0,0131930 | 1,11 | 1,38E+08 | 1,32E+08 |

| | | | | | | |
|---|---|---|---|---|---|---|
| 21 | | 0,0145230 | 0,0131882 | 0,77 | 2,84E+08 | 2,75E+08 |
| 42 | | 0,0145550 | 0,0132227 | 1,31 | 2,58E+07 | 2,11E+07 |
| 34,5 | | 0,0145638 | 0,0132179 | 1,16 | 2,83E+08 | 2,73E+08 |
| 5,9 | | 0,0145741 | 0,0132480 | 0,65 | 2,96E+07 | 3,31E+07 |
| 25,5 | | 0,0145810 | 0,0132432 | 1,50 | 8,02E+07 | 8,34E+07 |
| 20,9 | | 0,0145961 | 0,0133227 | 1,33 | 8,14E+07 | 8,95E+07 |
| 33,9 | | 0,0146122 | 0,0134021 | 1,45 | 4,64E+07 | 3,85E+07 |
| 3,9 | | 0,0146189 | 0,0133972 | 0,71 | 4,08E+06 | 3,63E+06 |
| 66,1 | | 0,0146291 | 0,0133924 | 0,64 | 7,86E+07 | 6,18E+07 |
| 38,8 | | 0,0146366 | 0,0133875 | 1,31 | 1,30E+08 | 1,30E+08 |
| 36,1 | | 0,0146395 | 0,0133827 | 0,79 | 4,07E+08 | 3,63E+08 |
| 34,8 | | 0,0146459 | 0,0133778 | 0,73 | 3,70E+08 | 3,68E+08 |
| 34,9 | | 0,0146707 | 0,0133730 | 0,79 | 5,71E+08 | 5,10E+08 |
| 51,8 | | 0,0146971 | 0,0134812 | 0,73 | 2,66E+09 | 3,08E+09 |
| 52,7 | | 0,0147431 | 0,0135212 | 1,50 | 4,34E+08 | 6,02E+08 |
| 62,1 | | 0,0147689 | 0,0135163 | 0,71 | 1,62E+08 | 1,61E+08 |
| 35 | | 0,0147753 | 0,0135114 | 0,80 | 2,73E+08 | 2,89E+08 |
| 60,5 | | 0,0147967 | 0,0135991 | 1,52 | 6,49E+08 | 7,49E+08 |
| 36,6 | | 0,0148251 | 0,0135942 | 0,72 | 1,40E+08 | 1,40E+08 |
| 42,4 | | 0,0148404 | 0,0136341 | 0,81 | 6,18E+08 | 7,11E+08 |
| 9,3 | | 0,0148442 | 0,0136292 | 0,23 | 1,59E+07 | 1,89E+07 |
| 49,3 | | 0,0148533 | 0,0136243 | 0,87 | 6,47E+08 | 6,31E+08 |
| 15,3 | | 0,0148590 | 0,0136194 | 1,70 | 3,64E+07 | 4,20E+07 |
| 28,8 | | 0,0148636 | 0,0136144 | 0,85 | 4,95E+07 | 4,78E+07 |
| 23,9 | | 0,0148926 | 0,0136095 | 0,73 | 5,78E+07 | 6,36E+07 |
| 21,1 | | 0,0149200 | 0,0136494 | 0,57 | 3,12E+07 | 2,68E+07 |
| 16,1 | | 0,0149350 | 0,0137339 | 9,69 | 1,29E+06 | 5,36E+06 |
| 20,3 | | 0,0149388 | 0,0137289 | 0,19 | 1,78E+07 | 1,82E+07 |
| 29,6 | | 0,0149410 | 0,0137240 | 1,37 | 6,41E+07 | 5,82E+07 |
| 59,6 | | 0,0149597 | 0,0137190 | 0,86 | 5,38E+09 | 5,45E+09 |
| 41,7 | | 0,0149609 | 0,0137141 | 1,18 | 1,17E+09 | 1,04E+09 |
| 9 | | 0,0149796 | 0,0137480 | 0,69 | 3,84E+07 | 4,65E+07 |
| 21,7 | | 0,0150047 | 0,0137431 | 0,74 | 7,83E+07 | 6,90E+07 |
| 16 | | 0,0150191 | 0,0137770 | 1,50 | 8,27E+07 | 6,26E+07 |
| 41,7 | | 0,0150399 | 0,0138109 | 0,61 | 6,08E+08 | 4,08E+08 |
| 11,1 | | 0,0150534 | 0,0138059 | 0,51 | 2,00E+07 | 2,18E+07 |
| 23,9 | | 0,0150718 | 0,0138009 | 23,96 | 4,08E+05 | 5,31E+06 |
| 27,6 | | 0,0150732 | 0,0137960 | 0,75 | 3,18E+08 | 2,52E+08 |
| 19,5 | | 0,0151583 | 0,0138355 | 0,85 | 8,30E+07 | 8,03E+07 |
| 77,9 | | 0,0152416 | 0,0138636 | 1,34 | 2,98E+09 | 3,10E+09 |
| 26,2 | | 0,0152621 | 0,0138586 | 0,60 | 2,63E+07 | 3,66E+07 |
| 2,3 | | 0,0152898 | 0,0138537 | 0,35 | 1,90E+07 | 1,48E+07 |
| 38,4 | | 0,0152976 | 0,0138946 | 0,83 | 1,17E+09 | 1,04E+09 |
| 15,2 | | 0,0153015 | 0,0138896 | 0,59 | 1,88E+07 | 2,23E+07 |
| 61,8 | | 0,0153041 | 0,0138846 | 1,26 | 9,36E+08 | 1,11E+09 |
| 37,3 | | 0,0153176 | 0,0138797 | 0,85 | 9,39E+08 | 1,07E+09 |
| 48,2 | | 0,0153393 | 0,0138747 | 0,80 | 1,80E+08 | 1,92E+08 |
| 29 | | 0,0153428 | 0,0138697 | 1,69 | 1,22E+08 | 1,15E+08 |
| 11,9 | | 0,0153430 | 0,0138648 | 1,45 | 3,50E+07 | 2,61E+07 |
| 51,7 | | 0,0154055 | 0,0139828 | 0,77 | 3,96E+08 | 4,68E+08 |

| | | | | | | |
|---|---|---|---|---|---|---|
| 22,5 | | 0,0154481 | 0,0140107 | 1,62 | 3,65E+07 | 2,68E+07 |
| 11,4 | | 0,0154503 | 0,0140057 | 2,06 | 7,37E+06 | 9,61E+06 |
| 39,2 | | 0,0154592 | 0,0140007 | 1,26 | 1,08E+08 | 9,56E+07 |
| 47,7 | | 0,0154611 | 0,0139957 | 0,69 | 2,02E+08 | 2,44E+08 |
| 33,5 | | 0,0154911 | 0,0139907 | 1,16 | 5,83E+08 | 6,08E+08 |
| 33 | | 0,0154959 | 0,0139857 | 1,62 | 4,40E+08 | 3,88E+08 |
| 61,2 | | 0,0155233 | 0,0140578 | 4,09 | 1,76E+07 | 4,56E+07 |
| 54,9 | | 0,0155611 | 0,0140870 | 0,82 | 4,91E+08 | 4,24E+08 |
| 34 | | 0,0156347 | 0,0142446 | 11,59 | 2,25E+06 | 1,39E+07 |
| 21,7 | | 0,0156701 | 0,0142395 | 0,79 | 4,79E+07 | 5,77E+07 |
| 41,9 | | 0,0157485 | 0,0142344 | 0,84 | 4,06E+08 | 3,53E+08 |
| 52,9 | | 0,0157651 | 0,0142293 | 1,20 | 3,70E+08 | 3,63E+08 |
| 52,5 | | 0,0157677 | 0,0142243 | 0,70 | 2,95E+08 | 2,92E+08 |
| 26,7 | | 0,0158076 | 0,0142577 | 1,73 | 3,38E+07 | 4,74E+07 |
| 58,8 | | 0,0158191 | 0,0142526 | 1,55 | 2,99E+08 | 3,44E+08 |
| 32,6 | | 0,0158478 | 0,0142817 | 0,82 | 1,03E+08 | 9,34E+07 |
| 71,3 | | 0,0158551 | 0,0142766 | 1,17 | 5,82E+09 | 5,34E+09 |
| 40,9 | | 0,0159290 | 0,0143426 | 1,16 | 2,97E+08 | 3,06E+08 |
| 57,4 | | 0,0160026 | 0,0144256 | 0,83 | 5,66E+09 | 6,54E+09 |
| 14,3 | | 0,0160114 | 0,0144929 | 0,57 | 8,99E+06 | 8,83E+06 |
| 26,8 | | 0,0160154 | 0,0144878 | 0,85 | 6,12E+07 | 5,69E+07 |
| 50,9 | | 0,0160278 | 0,0144826 | 1,22 | 2,68E+08 | 2,64E+08 |
| 28,4 | | 0,0160493 | 0,0145215 | 3,84 | 5,61E+06 | 2,12E+06 |
| 17,5 | | 0,0160723 | 0,0145546 | 0,53 | 1,97E+07 | 2,49E+07 |
| 35,8 | | 0,0160870 | 0,0145495 | 0,87 | 3,40E+08 | 3,33E+08 |
| 11,8 | | 0,0161557 | 0,0145882 | 0,19 | 1,02E+07 | 1,34E+07 |
| 13,9 | | 0,0161876 | 0,0145831 | 0,42 | 1,54E+07 | 1,20E+07 |
| 23,6 | | 0,0161934 | 0,0146105 | 0,56 | 2,78E+07 | 2,54E+07 |
| 19,3 | | 0,0162637 | 0,0146053 | 0,79 | 3,18E+07 | 3,26E+07 |
| 30,9 | | 0,0162836 | 0,0146001 | 6,70 | 1,39E+08 | 4,33E+07 |
| 6,8 | + | 0,0162851 | 0,0145950 | 2,77 | 2,41E+06 | 3,11E+06 |
| 52,8 | | 0,0163185 | 0,0145898 | 0,71 | 9,31E+08 | 9,42E+08 |
| 30,9 | | 0,0163284 | 0,0145847 | 0,77 | 5,38E+07 | 5,13E+07 |
| 28,6 | | 0,0163447 | 0,0145795 | 0,68 | 1,11E+08 | 9,94E+07 |
| 42,4 | | 0,0163590 | 0,0145744 | 0,80 | 4,47E+07 | 4,82E+07 |
| 14,6 | | 0,0164375 | 0,0146031 | 1,41 | 4,02E+07 | 3,26E+07 |
| 3,2 | | 0,0164402 | 0,0145980 | 0,27 | 1,18E+07 | 3,96E+06 |
| 54,8 | | 0,0164769 | 0,0146267 | 1,10 | 2,40E+09 | 2,46E+09 |
| 61,8 | | 0,0165024 | 0,0146215 | 1,21 | 2,75E+09 | 3,17E+09 |
| 33,6 | | 0,0165541 | 0,0147264 | 0,04 | 1,10E+08 | 1,61E+08 |
| 32,6 | | 0,0165864 | 0,0148650 | 0,63 | 9,29E+07 | 1,08E+08 |
| 6 | | 0,0165932 | 0,0148598 | 1,88 | 9,59E+06 | 9,23E+06 |
| 38,5 | | 0,0165964 | 0,0148545 | 1,22 | 3,51E+08 | 3,55E+08 |
| 9,6 | | 0,0165976 | 0,0148493 | 1,59 | 4,95E+06 | 3,38E+06 |
| 10,5 | | 0,0166097 | 0,0148441 | 0,62 | 6,98E+06 | 9,22E+06 |
| 27,6 | | 0,0166252 | 0,0148388 | 0,76 | 1,23E+08 | 1,15E+08 |
| 54,5 | + | 0,0166627 | 0,0148336 | 1,93 | 7,41E+09 | 4,23E+09 |
| 66,3 | | 0,0166748 | 0,0148284 | 1,40 | 7,05E+08 | 8,14E+08 |
| 34,3 | | 0,0166841 | 0,0148668 | 0,88 | 1,61E+08 | 1,55E+08 |
| 7,3 | | 0,0166933 | 0,0148616 | 1,48 | 4,93E+06 | 5,02E+06 |

| | | | | | | |
|---|---|---|---|---|---|---|
| 54,8 | | 0,0167002 | 0,0148563 | 1,25 | 5,92E+08 | 5,15E+08 |
| 22,5 | | 0,0167329 | 0,0148511 | 0,46 | 6,85E+06 | 7,53E+06 |
| 46,5 | | 0,0167425 | 0,0148459 | 1,23 | 1,85E+08 | 1,65E+08 |
| 62,2 | | 0,0167651 | 0,0148744 | 0,76 | 2,52E+08 | 3,15E+08 |
| 25,5 | | 0,0167743 | 0,0149127 | 0,78 | 2,64E+07 | 2,40E+07 |
| 2,4 | | 0,0167988 | 0,0149453 | 0,22 | 8,59E+06 | 1,03E+07 |
| 25,8 | | 0,0168086 | 0,0149401 | 0,65 | 2,20E+07 | 1,87E+07 |
| 40,3 | | 0,0168160 | 0,0149348 | 0,64 | 1,76E+08 | 1,75E+08 |
| 23,6 | | 0,0169101 | 0,0150543 | 0,76 | 3,35E+08 | 2,73E+08 |
| 56,4 | | 0,0169246 | 0,0150490 | 0,07 | 8,36E+07 | 7,33E+07 |
| 5 | | 0,0169527 | 0,0150774 | 0,67 | 2,85E+07 | 2,97E+07 |
| 19,3 | | 0,0169644 | 0,0150721 | 0,62 | 1,02E+07 | 1,04E+07 |
| 6,3 | | 0,0169714 | 0,0151102 | 5,34 | 3,02E+06 | 1,43E+06 |
| 30,7 | | 0,0169980 | 0,0151049 | 0,78 | 4,62E+08 | 4,22E+08 |
| 13,8 | | 0,0170379 | 0,0151374 | 0,74 | 4,63E+07 | 5,80E+07 |
| 54 | | 0,0170484 | 0,0152145 | 1,49 | 1,02E+09 | 8,70E+08 |
| 5,3 | | 0,0170681 | 0,0153126 | 1,31 | 1,98E+07 | 2,20E+07 |
| 44,7 | | 0,0170944 | 0,0153073 | 1,12 | 1,04E+08 | 1,08E+08 |
| 46 | | 0,0171090 | 0,0153019 | 1,35 | 8,14E+08 | 6,54E+08 |
| 4,8 | | 0,0171149 | 0,0152966 | 1,34 | 7,73E+06 | 6,57E+06 |
| 54 | | 0,0171266 | 0,0152912 | 0,65 | 3,48E+08 | 3,74E+08 |
| 47 | | 0,0171718 | 0,0153180 | 1,27 | 1,50E+09 | 1,54E+09 |
| 44,3 | | 0,0172323 | 0,0153559 | 1,11 | 7,00E+08 | 6,46E+08 |
| 21,8 | | 0,0172439 | 0,0153505 | 0,75 | 6,23E+07 | 6,84E+07 |
| 41,9 | | 0,0172489 | 0,0153452 | 0,22 | 5,98E+08 | 6,07E+08 |
| 33,8 | | 0,0172553 | 0,0153398 | 0,69 | 7,48E+07 | 8,14E+07 |
| 14,6 | | 0,0172849 | 0,0153777 | 0,09 | 5,83E+07 | 4,29E+07 |
| 24,5 | | 0,0173308 | 0,0154043 | 0,75 | 7,40E+07 | 7,59E+07 |
| 11,1 | | 0,0173571 | 0,0154365 | 0,48 | 7,63E+06 | 1,24E+07 |
| 33,8 | | 0,0173785 | 0,0154312 | 1,22 | 7,54E+07 | 7,64E+07 |
| 20,8 | | 0,0173853 | 0,0154258 | 1,27 | 6,51E+06 | 7,45E+06 |
| 75,2 | | 0,0174184 | 0,0154204 | 0,81 | 2,17E+10 | 2,33E+10 |
| 24,3 | | 0,0174334 | 0,0154526 | 0,64 | 2,01E+08 | 2,47E+08 |
| 11,5 | | 0,0174379 | 0,0154472 | 5,62 | 5,48E+06 | 2,75E+06 |
| 24,7 | | 0,0174436 | 0,0154419 | 0,30 | 7,97E+06 | 1,42E+07 |
| 25,2 | | 0,0174468 | 0,0154684 | 0,09 | 4,08E+07 | 4,15E+07 |
| 40,6 | | 0,0174773 | 0,0154631 | 0,74 | 3,93E+08 | 4,34E+08 |
| 22,4 | | 0,0174836 | 0,0154577 | 1,39 | 6,91E+07 | 5,34E+07 |
| 47,9 | | 0,0175316 | 0,0154523 | 0,82 | 9,11E+08 | 9,51E+08 |
| 32,5 | | 0,0175494 | 0,0154470 | 1,52 | 2,31E+07 | 3,06E+07 |
| 42 | | 0,0175937 | 0,0155871 | 0,90 | 1,15E+09 | 1,06E+09 |
| 55,6 | | 0,0176079 | 0,0155817 | 0,69 | 9,14E+07 | 8,60E+07 |
| 18,6 | + | 0,0176090 | 0,0155763 | 1,73 | 1,20E+07 | 1,36E+07 |
| 16,3 | | 0,0176461 | 0,0156484 | 1,21 | 2,37E+08 | 2,37E+08 |
| 32,9 | | 0,0177216 | 0,0156430 | 0,73 | 5,03E+07 | 4,86E+07 |
| 38,2 | | 0,0177954 | 0,0156376 | 1,20 | 4,86E+08 | 4,87E+08 |
| 23,9 | | 0,0178121 | 0,0156322 | 0,26 | 8,66E+06 | 1,11E+07 |
| 34,6 | | 0,0178463 | 0,0156268 | 0,79 | 1,34E+09 | 1,37E+09 |
| 13,2 | | 0,0178520 | 0,0156214 | 0,55 | 3,46E+08 | 2,57E+08 |
| 26 | | 0,0178611 | 0,0156160 | 0,37 | 6,95E+06 | 7,35E+06 |

| | | | | | | |
|---|---|---|---|---|---|---|
| 35,2 | | 0,0179043 | 0,0156534 | 0,85 | 6,71E+08 | 7,32E+08 |
| 39,3 | | 0,0179241 | 0,0156853 | 1,27 | 1,17E+08 | 1,05E+08 |
| 30,8 | | 0,0179350 | 0,0156799 | 1,32 | 6,28E+07 | 5,21E+07 |
| 25,2 | | 0,0179618 | 0,0156745 | 1,34 | 4,93E+07 | 5,71E+07 |
| 67,3 | | 0,0179860 | 0,0156691 | 0,81 | 1,11E+09 | 1,20E+09 |
| 14 | | 0,0180050 | 0,0157064 | 0,27 | 1,50E+07 | 1,28E+07 |
| 14,2 | | 0,0180717 | 0,0157437 | 1,47 | 8,98E+07 | 1,21E+08 |
| 58,1 | | 0,0180813 | 0,0157383 | 0,68 | 1,64E+08 | 1,80E+08 |
| 39,5 | | 0,0181119 | 0,0157329 | 0,84 | 1,20E+08 | 1,36E+08 |
| 9,1 | | 0,0181168 | 0,0157275 | 1,83 | 2,30E+06 | 2,12E+06 |
| 3,5 | | 0,0181202 | 0,0157221 | 0,35 | 1,74E+07 | 1,05E+07 |
| 72,8 | | 0,0181548 | 0,0157923 | 1,52 | 1,38E+09 | 1,78E+09 |
| 30 | | 0,0181695 | 0,0157869 | 0,86 | 6,70E+07 | 7,39E+07 |
| 6,5 | | 0,0182054 | 0,0157814 | 0,34 | 6,88E+06 | 7,30E+06 |
| 15,5 | | 0,0182221 | 0,0158186 | 2,17 | 3,01E+06 | 5,88E+06 |
| 42,5 | | 0,0182313 | 0,0158132 | 1,51 | 8,55E+07 | 1,13E+08 |
| 73 | | 0,0182962 | 0,0159053 | 1,16 | 1,68E+09 | 1,68E+09 |
| 49 | | 0,0183568 | 0,0160178 | 0,63 | 4,21E+07 | 5,15E+07 |
| 74,5 | | 0,0183679 | 0,0160123 | 1,60 | 2,70E+09 | 3,52E+09 |
| 37,4 | | 0,0184162 | 0,0160069 | 0,81 | 4,49E+07 | 5,28E+07 |
| 24,7 | | 0,0184693 | 0,0160014 | 0,81 | 3,67E+08 | 3,65E+08 |
| 24,6 | | 0,0184944 | 0,0159959 | 0,40 | 4,27E+07 | 4,14E+07 |
| 64,9 | | 0,0185260 | 0,0160699 | 0,69 | 2,41E+08 | 2,45E+08 |
| 69,2 | | 0,0185272 | 0,0160644 | 0,82 | 8,46E+08 | 8,27E+08 |
| 26,3 | | 0,0185340 | 0,0160589 | 0,60 | 4,06E+07 | 3,95E+07 |
| 10,8 | | 0,0185629 | 0,0160849 | 1,79 | 6,75E+06 | 4,67E+06 |
| 43,9 | | 0,0186349 | 0,0160794 | 0,88 | 4,14E+09 | 3,75E+09 |
| 22 | | 0,0186440 | 0,0160739 | 0,82 | 2,04E+08 | 1,87E+08 |
| 4 | | 0,0187459 | 0,0161121 | 0,42 | 2,60E+06 | 2,37E+06 |
| 40,5 | | 0,0187866 | 0,0161763 | 1,19 | 4,53E+08 | 4,17E+08 |
| 62,3 | | 0,0188306 | 0,0162132 | 1,09 | 5,94E+09 | 5,81E+09 |
| 36,3 | | 0,0189028 | 0,0162814 | 1,20 | 9,77E+07 | 9,64E+07 |
| 32 | | 0,0189151 | 0,0162759 | 0,95 | 1,23E+08 | 1,22E+08 |
| 20 | | 0,0189972 | 0,0163072 | 0,04 | 1,40E+08 | 1,47E+08 |
| 71,9 | | 0,0190744 | 0,0164367 | 0,89 | 2,16E+09 | 2,06E+09 |
| 45,1 | | 0,0191191 | 0,0165375 | 0,82 | 2,19E+08 | 2,17E+08 |
| 16,3 | | 0,0191514 | 0,0165687 | 1,21 | 3,06E+07 | 3,19E+07 |
| 80,4 | | 0,0191584 | 0,0165631 | 0,92 | 6,69E+09 | 6,22E+09 |
| 43,1 | | 0,0191854 | 0,0165574 | 1,35 | 1,90E+08 | 2,31E+08 |
| 51,6 | | 0,0192303 | 0,0165518 | 1,37 | 7,74E+07 | 6,74E+07 |
| 57,8 | | 0,0192681 | 0,0165884 | 1,10 | 2,69E+09 | 2,65E+09 |
| 14,4 | | 0,0193131 | 0,0165827 | 0,81 | 2,48E+07 | 2,39E+07 |
| 34,5 | | 0,0193144 | 0,0165771 | 1,27 | 1,16E+08 | 1,25E+08 |
| 54,2 | | 0,0193313 | 0,0166041 | 1,17 | 8,28E+08 | 7,46E+08 |
| 26,7 | | 0,0193579 | 0,0166352 | 0,14 | 3,29E+07 | 3,45E+07 |
| 43,1 | | 0,0193684 | 0,0166295 | 0,69 | 2,29E+08 | 2,56E+08 |
| 30,4 | | 0,0193889 | 0,0166986 | 0,52 | 1,60E+07 | 1,79E+07 |
| 12,9 | | 0,0194147 | 0,0166929 | 3,44 | 5,81E+06 | 7,29E+06 |
| 12 | | 0,0194194 | 0,0166873 | 0,62 | 3,55E+07 | 3,79E+07 |
| 43,4 | | 0,0194289 | 0,0166816 | 0,80 | 1,42E+08 | 1,34E+08 |

| | | | | | | |
|---|---|---|---|---|---|---|
| 32,3 | | 0,0194363 | 0,0166759 | 1,32 | 2,70E+07 | 2,90E+07 |
| 73,7 | | 0,0194821 | 0,0167015 | 0,89 | 2,97E+09 | 2,70E+09 |
| 5,9 | | 0,0194942 | 0,0166958 | 1,70 | 2,68E+06 | 3,19E+06 |
| 11,8 | | 0,0195022 | 0,0166902 | 0,08 | 7,84E+07 | 1,06E+08 |
| 28 | | 0,0195370 | 0,0167279 | 1,30 | 5,47E+07 | 6,45E+07 |
| 26,3 | | 0,0195377 | 0,0167222 | 0,72 | 8,84E+07 | 1,08E+08 |
| 29,6 | | 0,0195942 | 0,0167166 | 1,28 | 8,94E+07 | 9,17E+07 |
| 33,1 | | 0,0196482 | 0,0167475 | 1,39 | 1,15E+09 | 1,39E+09 |
| 28,7 | | 0,0196808 | 0,0167418 | 0,12 | 1,32E+07 | 3,47E+07 |
| 14,2 | | 0,0197048 | 0,0168119 | 0,65 | 1,71E+08 | 2,19E+08 |
| 37,9 | | 0,0197148 | 0,0168062 | 1,91 | 1,65E+08 | 9,34E+07 |
| 37,4 | | 0,0197229 | 0,0168005 | 1,21 | 1,28E+08 | 1,14E+08 |
| 26,7 | | 0,0197496 | 0,0168273 | 1,32 | 4,68E+08 | 5,28E+08 |
| 13,6 | | 0,0197545 | 0,0168216 | 0,65 | 7,43E+07 | 8,20E+07 |
| 58,6 | | 0,0197952 | 0,0168484 | 0,74 | 2,28E+08 | 2,52E+08 |
| 29 | | 0,0198003 | 0,0168427 | 0,86 | 1,46E+08 | 1,37E+08 |
| 12,4 | | 0,0198044 | 0,0168370 | 0,13 | 2,80E+07 | 3,12E+07 |
| 35,6 | | 0,0198429 | 0,0168947 | 0,65 | 6,37E+07 | 6,41E+07 |
| 26,5 | | 0,0199096 | 0,0168890 | 1,27 | 5,47E+07 | 5,78E+07 |
| 30,2 | | 0,0199270 | 0,0168833 | 0,79 | 7,52E+08 | 9,06E+08 |
| 31,5 | | 0,0199885 | 0,0169141 | 2,52 | 8,25E+06 | 1,60E+07 |
| 42,8 | | 0,0200053 | 0,0169084 | 0,71 | 6,33E+07 | 5,93E+07 |
| 15,5 | | 0,0200179 | 0,0169444 | 0,82 | 8,44E+07 | 8,47E+07 |
| 12,1 | | 0,0200744 | 0,0170128 | 1,62 | 1,53E+07 | 1,68E+07 |
| 63,1 | | 0,0201708 | 0,0170744 | 1,46 | 1,42E+08 | 1,34E+08 |
| 30,1 | | 0,0202256 | 0,0171050 | 0,93 | 6,70E+08 | 6,57E+08 |
| 45,6 | | 0,0202553 | 0,0170992 | 0,76 | 4,48E+08 | 4,63E+08 |
| 41,5 | | 0,0202576 | 0,0170935 | 0,75 | 2,00E+08 | 2,12E+08 |
| 31,8 | | 0,0203100 | 0,0172087 | 1,29 | 1,29E+08 | 1,17E+08 |
| 45,5 | | 0,0203179 | 0,0172030 | 0,89 | 1,13E+08 | 1,22E+08 |
| 26,5 | | 0,0203561 | 0,0172388 | 1,58 | 8,03E+06 | 5,45E+06 |
| 25,6 | | 0,0203581 | 0,0172330 | 0,79 | 3,67E+07 | 3,80E+07 |
| 58,9 | | 0,0204252 | 0,0173360 | 0,83 | 2,35E+08 | 2,45E+08 |
| 42,5 | | 0,0205430 | 0,0173987 | 1,17 | 3,08E+08 | 3,19E+08 |
| 81 | | 0,0205869 | 0,0173928 | 1,42 | 1,68E+09 | 1,90E+09 |
| 5,2 | | 0,0206100 | 0,0173870 | 0,75 | 3,17E+07 | 3,83E+07 |
| 24,6 | | 0,0206550 | 0,0173812 | 1,33 | 6,60E+07 | 7,11E+07 |
| 12,1 | | 0,0207118 | 0,0173753 | 1,36 | 3,30E+07 | 3,40E+07 |
| 45,7 | | 0,0207229 | 0,0173695 | 1,10 | 5,44E+08 | 5,72E+08 |
| 8,8 | | 0,0207658 | 0,0174052 | 1,79 | 4,14E+06 | 2,52E+06 |
| 53,7 | | 0,0207763 | 0,0173994 | 1,46 | 8,17E+07 | 9,68E+07 |
| 21 | | 0,0208398 | 0,0174257 | 1,39 | 3,57E+07 | 2,78E+07 |
| 22,9 | | 0,0208471 | 0,0174199 | 0,77 | 3,67E+07 | 3,62E+07 |
| 35,8 | | 0,0208522 | 0,0174140 | 0,89 | 7,74E+08 | 8,18E+08 |
| 37,8 | | 0,0209378 | 0,0175219 | 1,28 | 1,07E+08 | 1,17E+08 |
| 28,6 | | 0,0209552 | 0,0175521 | 0,82 | 5,79E+07 | 5,65E+07 |
| 6,3 | | 0,0209648 | 0,0175463 | 1,49 | 5,66E+06 | 4,47E+06 |
| 5,7 | | 0,0209741 | 0,0175404 | 1,50 | 1,15E+07 | 8,41E+06 |
| 66,9 | | 0,0210454 | 0,0176681 | 0,77 | 2,11E+08 | 2,18E+08 |
| 43,3 | | 0,0210733 | 0,0177036 | 0,73 | 1,38E+08 | 1,43E+08 |

| | | | | | | |
|---|---|---|---|---|---|---|
| 45,4 | | 0,0210824 | 0,0176977 | 0,91 | 3,66E+08 | 3,49E+08 |
| 50,9 | | 0,0211303 | 0,0177238 | 1,53 | 1,27E+08 | 9,69E+07 |
| 42,6 | | 0,0211748 | 0,0177593 | 0,73 | 1,33E+08 | 1,59E+08 |
| 16,3 | | 0,0211771 | 0,0177533 | 0,80 | 3,95E+07 | 3,94E+07 |
| 44,6 | | 0,0211965 | 0,0177474 | 1,69 | 3,25E+07 | 2,87E+07 |
| 8,9 | | 0,0212553 | 0,0177415 | 8,19 | 8,68E+04 | 3,75E+05 |
| 40,9 | | 0,0213504 | 0,0177356 | 0,31 | 2,46E+07 | 4,58E+07 |
| 19,2 | | 0,0214167 | 0,0177603 | 0,68 | 5,92E+07 | 6,60E+07 |
| 10 | | 0,0214730 | 0,0177544 | 1,70 | 6,21E+06 | 5,93E+06 |
| 42,9 | | 0,0215207 | 0,0177844 | 1,52 | 6,32E+07 | 4,38E+07 |
| 16,3 | | 0,0215433 | 0,0178144 | 1,45 | 2,28E+07 | 2,24E+07 |
| 39 | | 0,0216245 | 0,0178870 | 0,77 | 3,38E+08 | 3,78E+08 |
| 14,4 | | 0,0216644 | 0,0178810 | 1,67 | 1,12E+07 | 1,28E+07 |
| 16,6 | | 0,0216661 | 0,0178751 | 0,70 | 2,30E+07 | 2,07E+07 |
| 32,8 | | 0,0216671 | 0,0178691 | 0,55 | 3,28E+07 | 3,59E+07 |
| 34,7 | | 0,0216908 | 0,0178632 | 0,87 | 3,49E+08 | 3,71E+08 |
| 15 | | 0,0217115 | 0,0178573 | 0,59 | 8,70E+07 | 8,99E+07 |
| 49,1 | | 0,0217233 | 0,0178938 | 2,98 | 5,57E+06 | 1,04E+07 |
| 20,2 | | 0,0217253 | 0,0178879 | 0,55 | 2,55E+07 | 2,71E+07 |
| 21,2 | | 0,0217344 | 0,0178820 | 0,31 | 9,76E+06 | 9,73E+06 |
| 18,5 | | 0,0217522 | 0,0178760 | 3,19 | 6,31E+07 | 4,45E+07 |
| 13,2 | | 0,0217702 | 0,0179112 | 0,76 | 2,24E+07 | 2,32E+07 |
| 21,3 | | 0,0218381 | 0,0179410 | 0,09 | 5,75E+07 | 4,54E+07 |
| 28,1 | | 0,0218818 | 0,0180079 | 0,21 | 2,47E+07 | 2,18E+07 |
| 26,8 | | 0,0218989 | 0,0180020 | 20,06 | 7,74E+06 | 1,26E+08 |
| 12,6 | | 0,0219109 | 0,0180688 | 0,15 | 4,93E+07 | 7,78E+07 |
| 10,4 | | 0,0219298 | 0,0181039 | 1,56 | 2,59E+07 | 2,31E+07 |
| 26 | | 0,0219726 | 0,0181389 | 0,22 | 5,68E+07 | 3,53E+07 |
| 16,3 | | 0,0219795 | 0,0181329 | 0,80 | 1,84E+07 | 1,59E+07 |
| 52 | | 0,0220049 | 0,0182049 | 1,19 | 2,16E+09 | 2,40E+09 |
| 25,8 | | 0,0220085 | 0,0181989 | 1,28 | 4,69E+07 | 4,52E+07 |
| 3,5 | | 0,0221142 | 0,0184333 | 0,52 | 5,57E+06 | 6,42E+06 |
| 4,4 | | 0,0222637 | 0,0187072 | 3,11 | 2,79E+06 | 2,44E+06 |
| 42,2 | | 0,0222827 | 0,0187314 | 0,67 | 3,82E+07 | 4,07E+07 |
| 12,5 | | 0,0223114 | 0,0187252 | 0,63 | 3,84E+07 | 3,08E+07 |
| 24,2 | | 0,0224278 | 0,0189472 | 1,25 | 2,15E+07 | 1,94E+07 |
| 47,6 | | 0,0224495 | 0,0189410 | 0,77 | 2,34E+08 | 2,41E+08 |
| 6,5 | | 0,0224616 | 0,0189347 | 4,61 | 4,26E+06 | 1,27E+06 |
| 6,8 | | 0,0224868 | 0,0189285 | 6,01 | 2,47E+06 | 1,68E+06 |
| 5,5 | + | 0,0224888 | 0,0189223 | 0,08 | 1,45E+07 | 9,01E+06 |
| 10 | | 0,0225583 | 0,0189924 | 2,15 | 2,27E+06 | 4,52E+06 |
| 36,4 | | 0,0225661 | 0,0190178 | 1,17 | 1,29E+08 | 1,28E+08 |
| 32 | | 0,0225764 | 0,0190115 | 0,74 | 1,57E+08 | 1,48E+08 |
| 52 | | 0,0225776 | 0,0190053 | 0,76 | 2,39E+08 | 2,45E+08 |
| 21 | | 0,0226479 | 0,0191108 | 0,71 | 7,25E+07 | 7,85E+07 |
| 42,6 | | 0,0226507 | 0,0191045 | 3,01 | 9,55E+06 | 1,46E+07 |
| 13,3 | | 0,0226708 | 0,0191692 | 2,28 | 5,29E+06 | 9,66E+06 |
| 2,3 | | 0,0227575 | 0,0192247 | 1,80 | 6,36E+06 | 4,36E+06 |
| 14,9 | | 0,0228029 | 0,0192184 | 2,39 | 1,46E+07 | 1,86E+07 |
| 5,3 | | 0,0228561 | 0,0192436 | 0,16 | 1,82E+07 | 2,07E+07 |

| | | | | | | |
|---|---|---|---|---|---|---|
| 1,7 | | 0,0228832 | 0,0192688 | 2,38 | 7,82E+06 | 6,41E+06 |
| 52,8 | | 0,0229348 | 0,0193255 | 1,11 | 7,63E+07 | 7,48E+07 |
| 8,7 | | 0,0229788 | 0,0193191 | 0,62 | 1,21E+07 | 8,53E+06 |
| 18,8 | | 0,0230579 | 0,0193797 | 0,72 | 1,08E+08 | 1,32E+08 |
| 48,9 | | 0,0231555 | 0,0194808 | 1,07 | 3,67E+08 | 3,50E+08 |
| 46,9 | | 0,0232214 | 0,0195098 | 0,75 | 3,84E+08 | 3,85E+08 |
| 11,7 | | 0,0232982 | 0,0195441 | 0,61 | 2,06E+07 | 2,28E+07 |
| 29 | | 0,0233195 | 0,0195783 | 0,69 | 1,99E+07 | 2,42E+07 |
| 57,2 | | 0,0233334 | 0,0195718 | 0,88 | 1,82E+08 | 1,76E+08 |
| 6,9 | + | 0,0233373 | 0,0195654 | 1,62 | 3,43E+06 | 2,17E+06 |
| 76,1 | | 0,0233381 | 0,0195590 | 0,88 | 4,89E+09 | 4,81E+09 |
| 5,9 | | 0,0233972 | 0,0195526 | 1,23 | 7,28E+06 | 8,18E+06 |
| 10 | | 0,0234369 | 0,0196613 | 0,18 | 1,91E+07 | 2,10E+07 |
| 24,5 | | 0,0235024 | 0,0197216 | 1,75 | 9,90E+07 | 9,90E+07 |
| 33,3 | | 0,0235031 | 0,0197151 | 1,35 | 1,61E+08 | 2,07E+08 |
| 3 | | 0,0235040 | 0,0197087 | 1,65 | 6,24E+06 | 6,37E+06 |
| 10,2 | | 0,0235297 | 0,0197023 | 2,62 | 4,11E+06 | 3,97E+06 |
| 17,5 | | 0,0235526 | 0,0196958 | 0,40 | 1,21E+07 | 1,24E+07 |
| 10,4 | | 0,0236229 | 0,0196894 | 0,61 | 2,09E+07 | 1,76E+07 |
| 13 | | 0,0236618 | 0,0197143 | 0,32 | 2,47E+07 | 1,25E+07 |
| 10,2 | | 0,0236757 | 0,0197079 | 0,40 | 2,24E+07 | 1,03E+07 |
| 64,2 | | 0,0237180 | 0,0197419 | 1,17 | 2,56E+09 | 2,88E+09 |
| 21,9 | | 0,0237947 | 0,0197758 | 0,64 | 2,85E+07 | 3,03E+07 |
| 6 | | 0,0238133 | 0,0197694 | 2,12 | 7,05E+06 | 9,75E+06 |
| 30,7 | | 0,0238698 | 0,0198750 | 1,31 | 5,06E+07 | 6,55E+07 |
| 23,9 | | 0,0238760 | 0,0198685 | 1,63 | 3,15E+07 | 4,22E+07 |
| 28 | | 0,0238896 | 0,0198620 | 0,09 | 3,94E+07 | 7,66E+07 |
| 22,2 | | 0,0239497 | 0,0199271 | 1,50 | 3,27E+07 | 3,94E+07 |
| 10,7 | | 0,0239763 | 0,0199610 | 0,45 | 1,39E+07 | 2,09E+07 |
| 51,1 | | 0,0239767 | 0,0199545 | 0,79 | 7,21E+08 | 7,58E+08 |
| 10,6 | | 0,0239817 | 0,0199480 | 5,61 | 4,52E+06 | 1,25E+07 |
| 65,2 | | 0,0240418 | 0,0199766 | 0,54 | 5,26E+08 | 5,37E+08 |
| 28,1 | | 0,0241332 | 0,0200805 | 1,29 | 2,25E+07 | 2,21E+07 |
| 21,1 | | 0,0241776 | 0,0201052 | 1,24 | 6,00E+07 | 6,50E+07 |
| 39,6 | | 0,0242091 | 0,0201389 | 0,70 | 8,78E+07 | 8,70E+07 |
| 12,2 | | 0,0242425 | 0,0201726 | 0,75 | 1,02E+08 | 9,96E+07 |
| 40,9 | | 0,0242777 | 0,0202011 | 0,72 | 2,38E+08 | 3,09E+08 |
| 23,8 | | 0,0243046 | 0,0202348 | 1,88 | 1,32E+07 | 2,01E+07 |
| 28,8 | | 0,0243350 | 0,0202684 | 0,83 | 1,48E+08 | 1,33E+08 |
| 27,4 | | 0,0243642 | 0,0203020 | 0,85 | 2,78E+08 | 2,51E+08 |
| 9,9 | | 0,0243831 | 0,0203265 | 2,54 | 2,94E+06 | 3,74E+06 |
| 19,5 | | 0,0244567 | 0,0204003 | 0,70 | 7,04E+07 | 5,04E+07 |
| 15,4 | | 0,0244878 | 0,0204286 | 1,25 | 6,84E+07 | 7,27E+07 |
| 23,9 | | 0,0244985 | 0,0204220 | 0,52 | 5,06E+06 | 3,28E+06 |
| 19,4 | | 0,0245399 | 0,0204905 | 0,69 | 1,72E+07 | 1,72E+07 |
| 20,4 | | 0,0245435 | 0,0204838 | 1,20 | 6,06E+07 | 5,93E+07 |
| 13,9 | | 0,0245465 | 0,0204772 | 0,72 | 2,99E+07 | 2,72E+07 |
| 63,7 | | 0,0245786 | 0,0204706 | 1,31 | 2,12E+09 | 1,64E+09 |
| 25,9 | | 0,0245925 | 0,0204640 | 1,33 | 6,21E+07 | 6,72E+07 |
| 50,2 | | 0,0246032 | 0,0204574 | 1,09 | 3,72E+08 | 3,99E+08 |

| | | | | | | |
|---|---|---|---|---|---|---|
| 31,4 | | 0,0246316 | 0,0205166 | 0,80 | 6,45E+08 | 7,53E+08 |
| 33,9 | | 0,0246329 | 0,0205100 | 0,88 | 2,98E+08 | 3,12E+08 |
| 14,6 | | 0,0246445 | 0,0205744 | 0,44 | 3,13E+07 | 3,00E+07 |
| 77,2 | | 0,0247168 | 0,0206490 | 1,49 | 4,56E+09 | 6,05E+09 |
| 49,6 | | 0,0247349 | 0,0206424 | 0,83 | 2,85E+08 | 2,40E+08 |
| 23,7 | | 0,0247824 | 0,0206357 | 1,38 | 9,94E+07 | 1,08E+08 |
| 39,8 | | 0,0248606 | 0,0207000 | 1,18 | 9,52E+08 | 9,93E+08 |
| 22,8 | | 0,0248854 | 0,0206933 | 1,42 | 7,28E+07 | 8,93E+07 |
| 6,7 | | 0,0249460 | 0,0207163 | 2,47 | 5,76E+06 | 6,95E+06 |
| 39,2 | | 0,0249616 | 0,0207096 | 0,70 | 2,45E+08 | 3,25E+08 |
| 21 | | 0,0250377 | 0,0207686 | 5,85 | 2,97E+06 | 3,51E+06 |
| 4,8 | | 0,0250426 | 0,0207619 | 1,72 | 1,35E+07 | 9,04E+06 |
| 45,6 | | 0,0250774 | 0,0208196 | 1,32 | 9,47E+07 | 8,20E+07 |
| 15 | | 0,0250832 | 0,0208129 | 0,76 | 5,10E+07 | 4,69E+07 |
| 4,7 | | 0,0251021 | 0,0208357 | 0,71 | 9,45E+06 | 9,78E+06 |
| 31,1 | | 0,0251107 | 0,0208290 | 1,13 | 3,83E+08 | 4,14E+08 |
| 16,5 | + | 0,0251362 | 0,0208224 | 9,07 | 5,04E+07 | 1,54E+07 |
| 71,2 | | 0,0251555 | 0,0208157 | 0,77 | 1,40E+09 | 1,13E+09 |
| 63 | | 0,0251667 | 0,0208090 | 1,29 | 4,34E+08 | 5,13E+08 |
| 18 | | 0,0252295 | 0,0208331 | 0,59 | 3,38E+07 | 3,19E+07 |
| 41,7 | | 0,0252442 | 0,0208264 | 0,84 | 1,21E+08 | 1,14E+08 |
| 23,7 | | 0,0252553 | 0,0208198 | 0,63 | 4,42E+07 | 3,50E+07 |
| 23,4 | | 0,0252888 | 0,0208528 | 0,80 | 1,54E+08 | 1,50E+08 |
| 7,9 | | 0,0252976 | 0,0208859 | 3,00 | 5,98E+06 | 3,42E+06 |
| 67,2 | | 0,0253193 | 0,0208792 | 1,10 | 9,49E+08 | 8,84E+08 |
| 50,8 | | 0,0253432 | 0,0208725 | 1,20 | 3,21E+08 | 3,70E+08 |
| 15,3 | | 0,0253990 | 0,0208966 | 1,28 | 3,47E+07 | 3,59E+07 |
| 15 | | 0,0254129 | 0,0208899 | 1,70 | 1,10E+07 | 1,35E+07 |
| 15,5 | | 0,0254447 | 0,0209139 | 1,12 | 4,61E+07 | 4,36E+07 |
| 17,3 | | 0,0254719 | 0,0209072 | 0,73 | 3,41E+07 | 3,17E+07 |
| 42,9 | | 0,0255297 | 0,0210451 | 0,88 | 1,53E+09 | 1,41E+09 |
| 12 | | 0,0255299 | 0,0210384 | 0,73 | 1,02E+08 | 8,64E+07 |
| 8,2 | | 0,0255473 | 0,0210623 | 3,46 | 4,79E+06 | 1,52E+06 |
| 25,4 | | 0,0255583 | 0,0210556 | 0,68 | 1,08E+08 | 1,02E+08 |
| 31,3 | | 0,0255605 | 0,0210489 | 0,79 | 2,67E+08 | 2,44E+08 |
| 34,2 | | 0,0255734 | 0,0210421 | 1,36 | 1,33E+08 | 1,45E+08 |
| 19,7 | | 0,0256756 | 0,0211874 | 0,12 | 6,70E+07 | 5,38E+07 |
| 11,3 | | 0,0257040 | 0,0212202 | 3,38 | 5,78E+06 | 5,23E+06 |
| 19,2 | | 0,0257043 | 0,0212134 | 1,56 | 2,34E+08 | 3,11E+08 |
| 55,4 | | 0,0257400 | 0,0212462 | 0,85 | 1,26E+09 | 1,19E+09 |
| 13,9 | | 0,0257637 | 0,0212394 | 1,44 | 2,75E+07 | 2,51E+07 |
| 38,9 | | 0,0257764 | 0,0212326 | 1,33 | 2,29E+08 | 2,35E+08 |
| 57,2 | | 0,0257884 | 0,0212998 | 0,86 | 2,27E+09 | 2,33E+09 |
| 41,9 | | 0,0258416 | 0,0213529 | 0,75 | 9,68E+07 | 8,76E+07 |
| 10,8 | | 0,0258991 | 0,0214097 | 0,66 | 1,00E+07 | 9,05E+06 |
| 35,5 | | 0,0259558 | 0,0214335 | 0,78 | 9,27E+07 | 1,12E+08 |
| 16,1 | | 0,0259777 | 0,0214954 | 0,55 | 2,48E+07 | 3,31E+07 |
| 18,6 | | 0,0260099 | 0,0214885 | 0,55 | 4,75E+07 | 3,34E+07 |
| 37,6 | | 0,0260160 | 0,0214817 | 1,18 | 9,67E+08 | 9,34E+08 |
| 13,4 | | 0,0260481 | 0,0214749 | 0,76 | 1,18E+07 | 1,30E+07 |

| | | | | | | |
|---|---|---|---|---|---|---|
| 25,9 | | 0,0261117 | 0,0215672 | 0,55 | 1,26E+07 | 1,52E+07 |
| 14,8 | | 0,0261681 | 0,0216302 | 0,81 | 9,64E+07 | 8,36E+07 |
| 7,9 | | 0,0262234 | 0,0216526 | 1,30 | 2,97E+07 | 2,52E+07 |
| 31,2 | | 0,0262705 | 0,0216762 | 0,51 | 2,93E+07 | 5,46E+07 |
| 34,1 | | 0,0262738 | 0,0216998 | 0,85 | 1,04E+08 | 1,04E+08 |
| 57,8 | | 0,0263129 | 0,0217665 | 0,88 | 4,71E+08 | 4,97E+08 |
| 31,2 | | 0,0263248 | 0,0217938 | 0,69 | 1,14E+09 | 1,50E+09 |
| 17,1 | | 0,0263252 | 0,0217869 | 0,42 | 3,42E+07 | 2,86E+07 |
| 19,8 | | 0,0263650 | 0,0218092 | 0,75 | 1,50E+08 | 1,43E+08 |
| 33,7 | | 0,0263662 | 0,0218023 | 1,15 | 1,31E+09 | 1,40E+09 |
| 11,1 | | 0,0263795 | 0,0218258 | 0,36 | 1,10E+07 | 1,10E+07 |
| 32,9 | | 0,0263864 | 0,0218189 | 0,29 | 1,10E+07 | 1,89E+07 |
| 56,7 | | 0,0264484 | 0,0218512 | 0,84 | 7,28E+08 | 8,03E+08 |
| 28,9 | | 0,0264562 | 0,0218443 | 0,85 | 5,33E+08 | 6,25E+08 |
| 6,3 | | 0,0264744 | 0,0218716 | 1,29 | 1,50E+07 | 1,65E+07 |
| 17,7 | | 0,0264986 | 0,0219039 | 0,24 | 1,64E+07 | 2,01E+07 |
| 33,6 | | 0,0265400 | 0,0218969 | 1,71 | 7,97E+07 | 1,23E+08 |
| 17 | | 0,0265669 | 0,0218900 | 1,54 | 7,01E+06 | 8,27E+06 |
| 45,9 | | 0,0265699 | 0,0218831 | 1,66 | 6,03E+07 | 6,52E+07 |
| 64,6 | | 0,0266152 | 0,0218762 | 1,15 | 6,22E+09 | 5,88E+09 |
| 6,3 | | 0,0266624 | 0,0218693 | 0,42 | 1,74E+07 | 1,10E+07 |
| 2,6 | | 0,0267612 | 0,0219015 | 2,04 | 2,85E+06 | 4,21E+06 |
| 70,9 | | 0,0268292 | 0,0219741 | 1,13 | 3,19E+09 | 2,90E+09 |
| 24,4 | | 0,0268484 | 0,0220076 | 3,11 | 1,89E+06 | 3,24E+06 |
| 10,3 | | 0,0269333 | 0,0220990 | 4,83 | 6,18E+06 | 1,82E+06 |
| 51,2 | + | 0,0269636 | 0,0221311 | 1,74 | 1,52E+08 | 1,19E+08 |
| 43 | | 0,0269961 | 0,0221242 | 0,80 | 1,03E+09 | 1,23E+09 |
| 15,1 | | 0,0270011 | 0,0221172 | 1,83 | 7,46E+06 | 1,15E+07 |
| 21,6 | | 0,0270460 | 0,0222035 | 0,86 | 2,06E+08 | 2,29E+08 |
| 32,8 | | 0,0271294 | 0,0222267 | 1,15 | 2,24E+08 | 1,98E+08 |
| 52,4 | | 0,0271361 | 0,0222197 | 1,15 | 1,63E+09 | 1,69E+09 |
| 17,3 | | 0,0271714 | 0,0222530 | 0,80 | 1,80E+07 | 2,10E+07 |
| 47,3 | | 0,0271954 | 0,0223580 | 0,78 | 3,01E+08 | 3,06E+08 |
| 25,8 | | 0,0272032 | 0,0223509 | 0,51 | 1,47E+07 | 1,22E+07 |
| 7,8 | | 0,0272293 | 0,0223439 | 1,29 | 6,12E+06 | 5,99E+06 |
| 19,4 | | 0,0273267 | 0,0223671 | 2,11 | 2,17E+06 | 3,56E+06 |
| 34,7 | | 0,0273590 | 0,0224669 | 1,23 | 1,07E+08 | 1,26E+08 |
| 61,1 | | 0,0273734 | 0,0224598 | 0,85 | 7,80E+08 | 7,56E+08 |
| 13,4 | | 0,0273766 | 0,0224527 | 0,81 | 2,27E+07 | 2,53E+07 |
| 13,3 | | 0,0273864 | 0,0224457 | 1,93 | 1,31E+07 | 1,02E+07 |
| 32,1 | | 0,0274790 | 0,0224387 | 1,39 | 1,78E+08 | 2,32E+08 |
| 42,6 | | 0,0275552 | 0,0225395 | 1,31 | 1,00E+08 | 1,19E+08 |
| 33,4 | | 0,0276050 | 0,0226002 | 0,83 | 2,80E+08 | 3,12E+08 |
| 37,3 | | 0,0276719 | 0,0226320 | 1,45 | 1,37E+08 | 1,06E+08 |
| 34,3 | | 0,0276977 | 0,0226637 | 1,32 | 1,08E+08 | 9,70E+07 |
| 65,1 | | 0,0277190 | 0,0226566 | 0,71 | 7,32E+08 | 8,03E+08 |
| 14,9 | | 0,0278019 | 0,0227134 | 0,44 | 2,10E+07 | 2,71E+07 |
| 8,4 | | 0,0278365 | 0,0227401 | 2,02 | 2,57E+07 | 3,42E+07 |
| 8,4 | | 0,0278618 | 0,0227330 | 2,95 | 8,16E+06 | 4,35E+06 |
| 58,7 | | 0,0278679 | 0,0227259 | 1,19 | 2,88E+08 | 3,29E+08 |

| | | | | | | |
|---|---|---|---|---|---|---|
| 16 | | 0,0279077 | 0,0227914 | 2,02 | 1,08E+07 | 1,71E+07 |
| 60,9 | | 0,0279860 | 0,0228530 | 0,84 | 2,30E+09 | 2,46E+09 |
| 28,2 | | 0,0279911 | 0,0228459 | 0,72 | 5,12E+07 | 4,51E+07 |
| 15,5 | | 0,0280171 | 0,0228388 | 7,64 | 3,52E+06 | 2,63E+06 |
| 28,2 | | 0,0280540 | 0,0228316 | 0,52 | 1,07E+07 | 1,31E+07 |
| 47,9 | | 0,0280583 | 0,0228245 | 0,84 | 4,21E+08 | 3,95E+08 |
| 14,5 | | 0,0280637 | 0,0228174 | 10,09 | 7,74E+06 | 2,44E+06 |
| 29,7 | | 0,0281107 | 0,0228777 | 0,79 | 2,60E+08 | 2,90E+08 |
| 38,6 | | 0,0281113 | 0,0228705 | 0,74 | 3,42E+08 | 3,83E+08 |
| 19,8 | | 0,0281799 | 0,0229657 | 1,83 | 7,54E+07 | 4,68E+07 |
| 10,5 | | 0,0282132 | 0,0229585 | 0,34 | 5,14E+06 | 8,26E+06 |
| 28,3 | | 0,0282297 | 0,0229514 | 1,48 | 9,89E+06 | 1,24E+07 |
| 53,7 | | 0,0282907 | 0,0229779 | 2,15 | 5,16E+07 | 7,10E+07 |
| 13,6 | | 0,0282990 | 0,0229707 | 1,38 | 5,13E+07 | 4,96E+07 |
| 51,7 | | 0,0283131 | 0,0229636 | 0,74 | 5,06E+08 | 5,69E+08 |
| 17,6 | | 0,0283580 | 0,0230884 | 1,31 | 6,32E+07 | 6,82E+07 |
| 5 | | 0,0283756 | 0,0231111 | 2,74 | 5,66E+06 | 2,02E+06 |
| 21,7 | | 0,0284952 | 0,0231425 | 0,80 | 4,38E+08 | 3,83E+08 |
| 23,1 | | 0,0285197 | 0,0231353 | 0,74 | 2,18E+07 | 2,68E+07 |
| 21,9 | | 0,0285450 | 0,0231667 | 0,72 | 5,02E+06 | 5,58E+06 |
| 4,9 | | 0,0285800 | 0,0232316 | 0,38 | 2,98E+07 | 1,14E+07 |
| 28,3 | | 0,0287296 | 0,0233648 | 1,97 | 1,80E+07 | 2,27E+07 |
| 32 | | 0,0287431 | 0,0233576 | 0,85 | 1,38E+08 | 1,40E+08 |
| 56,4 | | 0,0287851 | 0,0234174 | 1,38 | 7,07E+08 | 7,81E+08 |
| 13,6 | | 0,0288417 | 0,0235070 | 0,45 | 4,29E+07 | 2,45E+07 |
| 20,4 | | 0,0288442 | 0,0234997 | 1,82 | 1,83E+07 | 1,77E+07 |
| 57 | | 0,0289072 | 0,0235222 | 1,14 | 1,41E+09 | 1,57E+09 |
| 33,2 | | 0,0289445 | 0,0235149 | 0,88 | 1,35E+08 | 1,45E+08 |
| 16,7 | | 0,0289808 | 0,0235758 | 1,79 | 2,21E+06 | 3,25E+06 |
| 25,8 | | 0,0290121 | 0,0236069 | 1,38 | 4,10E+07 | 4,06E+07 |
| 20 | | 0,0290143 | 0,0235996 | 0,78 | 8,45E+07 | 8,24E+07 |
| 8,5 | | 0,0290295 | 0,0236208 | 1,91 | 3,00E+07 | 3,98E+07 |
| 26,6 | | 0,0290634 | 0,0236135 | 1,27 | 4,34E+07 | 4,51E+07 |
| 17,3 | | 0,0291425 | 0,0236446 | 4,78 | 1,18E+07 | 3,25E+06 |
| 13,7 | | 0,0291680 | 0,0236373 | 0,25 | 8,23E+06 | 8,78E+06 |
| 16,5 | | 0,0291759 | 0,0236300 | 0,46 | 3,22E+07 | 4,25E+07 |
| 40,3 | | 0,0292133 | 0,0236820 | 0,80 | 3,74E+08 | 3,60E+08 |
| 48 | | 0,0292383 | 0,0237081 | 1,09 | 5,48E+08 | 5,59E+08 |
| 52,4 | | 0,0292548 | 0,0237008 | 0,69 | 9,08E+08 | 1,19E+09 |
| 14,2 | | 0,0292788 | 0,0237614 | 1,44 | 4,26E+07 | 4,97E+07 |
| 16 | | 0,0295160 | 0,0238925 | 0,74 | 6,63E+06 | 6,55E+06 |
| 13,4 | | 0,0295426 | 0,0239246 | 0,54 | 3,07E+07 | 1,78E+07 |
| 32 | | 0,0295578 | 0,0239173 | 0,85 | 1,59E+08 | 1,45E+08 |
| 46,4 | | 0,0297391 | 0,0240198 | 1,05 | 3,95E+09 | 3,89E+09 |
| 59,1 | | 0,0297493 | 0,0240123 | 1,18 | 4,80E+08 | 5,56E+08 |
| 17 | | 0,0297773 | 0,0240049 | 0,55 | 6,07E+08 | 7,06E+08 |
| 6,5 | | 0,0298828 | 0,0240358 | 0,67 | 4,64E+06 | 4,30E+06 |
| 14,9 | | 0,0299001 | 0,0240284 | 0,59 | 8,38E+06 | 1,43E+07 |
| 13,3 | | 0,0299819 | 0,0240838 | 0,26 | 3,77E+07 | 1,86E+07 |
| 41,4 | | 0,0299973 | 0,0240764 | 0,90 | 1,27E+09 | 1,36E+09 |

| | | | | | | |
|---|---|---|---|---|---|---|
| 5 | | 0,0300414 | 0,0241565 | 2,16 | 2,58E+06 | 3,48E+06 |
| 14 | | 0,0300538 | 0,0241490 | 0,48 | 3,80E+07 | 3,32E+07 |
| 44,8 | | 0,0301017 | 0,0241416 | 0,48 | 1,00E+08 | 1,61E+08 |
| 47,8 | | 0,0302629 | 0,0242868 | 0,76 | 9,05E+08 | 9,34E+08 |
| 35,2 | | 0,0302772 | 0,0242793 | 0,91 | 2,82E+08 | 2,71E+08 |
| 12,4 | | 0,0303306 | 0,0243432 | 0,72 | 3,00E+07 | 2,54E+07 |
| 6,7 | | 0,0303415 | 0,0243357 | 0,40 | 5,67E+06 | 6,81E+06 |
| 46,2 | | 0,0303670 | 0,0243282 | 1,76 | 4,38E+07 | 5,70E+07 |
| 25,1 | | 0,0304302 | 0,0243920 | 1,62 | 2,64E+07 | 4,19E+07 |
| 46,4 | | 0,0304302 | 0,0243845 | 0,80 | 2,74E+08 | 2,31E+08 |
| 7,6 | | 0,0304961 | 0,0244912 | 2,17 | 3,52E+06 | 1,80E+06 |
| 49 | | 0,0305729 | 0,0245930 | 1,19 | 3,40E+08 | 3,40E+08 |
| 50,1 | | 0,0305899 | 0,0246530 | 0,84 | 9,30E+08 | 1,06E+09 |
| 2,9 | | 0,0306020 | 0,0246454 | 0,62 | 5,74E+06 | 4,50E+06 |
| 56,8 | | 0,0306118 | 0,0246378 | 0,84 | 3,79E+08 | 4,34E+08 |
| 8,4 | | 0,0306119 | 0,0246303 | 1,29 | 2,78E+07 | 2,20E+07 |
| 18,6 | | 0,0306417 | 0,0246902 | 2,51 | 8,81E+06 | 1,06E+07 |
| 8,2 | | 0,0307106 | 0,0247831 | 0,47 | 1,03E+07 | 1,87E+07 |
| 11,6 | | 0,0307557 | 0,0247755 | 1,25 | 1,42E+07 | 1,67E+07 |
| 5,1 | | 0,0307626 | 0,0247679 | 0,33 | 2,35E+07 | 1,25E+07 |
| 12,9 | | 0,0307979 | 0,0247983 | 1,28 | 1,63E+07 | 1,57E+07 |
| 6,7 | | 0,0308039 | 0,0247907 | 0,70 | 2,28E+07 | 2,18E+07 |
| 66,1 | | 0,0308170 | 0,0247831 | 0,72 | 2,11E+09 | 2,73E+09 |
| 26,3 | | 0,0308279 | 0,0248037 | 1,58 | 5,02E+07 | 3,85E+07 |
| 22 | | 0,0308641 | 0,0247961 | 0,32 | 2,68E+07 | 3,49E+07 |
| 35 | | 0,0309075 | 0,0248924 | 1,52 | 1,43E+08 | 1,01E+08 |
| 15,3 | | 0,0309438 | 0,0248848 | 0,39 | 1,10E+07 | 9,38E+06 |
| 77,5 | | 0,0309867 | 0,0249163 | 1,73 | 3,97E+08 | 2,29E+08 |
| 14,5 | | 0,0310267 | 0,0249380 | 0,69 | 1,25E+07 | 1,18E+07 |
| 8,3 | | 0,0310508 | 0,0249963 | 1,34 | 7,41E+06 | 7,27E+06 |
| 7,8 | | 0,0310924 | 0,0249887 | 0,31 | 1,59E+07 | 1,66E+07 |
| 30,9 | | 0,0310960 | 0,0249811 | 1,30 | 7,74E+08 | 6,12E+08 |
| 40,5 | | 0,0311591 | 0,0250064 | 0,92 | 1,88E+08 | 1,86E+08 |
| 34,7 | | 0,0311943 | 0,0250366 | 1,24 | 1,92E+08 | 2,14E+08 |
| 36,8 | | 0,0312275 | 0,0250667 | 1,08 | 3,32E+08 | 3,16E+08 |
| 38,7 | | 0,0312329 | 0,0250591 | 1,55 | 3,81E+07 | 2,49E+07 |
| 5,2 | | 0,0312866 | 0,0250905 | 0,50 | 8,36E+06 | 4,48E+06 |
| 35,5 | | 0,0313369 | 0,0251108 | 1,23 | 2,82E+08 | 3,51E+08 |
| 23,5 | | 0,0315794 | 0,0252651 | 0,82 | 6,68E+08 | 5,84E+08 |
| 24,9 | | 0,0315984 | 0,0252575 | 0,05 | 8,39E+07 | 7,64E+07 |
| 51,6 | | 0,0316514 | 0,0253264 | 0,78 | 1,16E+09 | 1,24E+09 |
| 21,8 | | 0,0317461 | 0,0254428 | 0,75 | 1,71E+08 | 1,97E+08 |
| 19 | | 0,0317554 | 0,0254351 | 1,17 | 1,28E+08 | 1,34E+08 |
| 52,1 | | 0,0318403 | 0,0255161 | 0,84 | 3,83E+08 | 4,02E+08 |
| 26,1 | | 0,0318739 | 0,0255084 | 1,24 | 1,13E+08 | 9,64E+07 |
| 20,4 | | 0,0319773 | 0,0255711 | 0,80 | 9,84E+07 | 8,61E+07 |
| 46,1 | | 0,0320351 | 0,0255633 | 1,31 | 1,22E+08 | 1,19E+08 |
| 26,3 | | 0,0320527 | 0,0255556 | 1,19 | 2,22E+08 | 2,14E+08 |
| 35 | | 0,0320886 | 0,0255478 | 0,79 | 2,34E+08 | 2,77E+08 |
| 6,9 | | 0,0321164 | 0,0256068 | 0,23 | 3,64E+07 | 3,94E+07 |

| | | | | | | |
|---|---|---|---|---|---|---|
| 13,5 | | 0,0321819 | 0,0256281 | 0,62 | 2,94E+07 | 2,84E+07 |
| 23,2 | | 0,0322089 | 0,0256204 | 0,78 | 8,44E+07 | 8,36E+07 |
| 18,9 | | 0,0322230 | 0,0256126 | 1,66 | 2,34E+07 | 1,97E+07 |
| 61,9 | | 0,0322576 | 0,0256339 | 0,77 | 1,55E+09 | 1,21E+09 |
| 1,4 | | 0,0322888 | 0,0256262 | 0,68 | 6,87E+07 | 6,73E+07 |
| 64,5 | | 0,0323469 | 0,0257505 | 0,83 | 5,31E+08 | 5,90E+08 |
| 12,1 | | 0,0324142 | 0,0258517 | 2,16 | 3,72E+06 | 8,11E+06 |
| 32,6 | | 0,0325904 | 0,0259794 | 1,44 | 4,99E+08 | 6,70E+08 |
| 12,4 | | 0,0326224 | 0,0260381 | 1,41 | 5,54E+07 | 4,59E+07 |
| 38,4 | | 0,0326447 | 0,0261295 | 0,89 | 2,91E+08 | 3,15E+08 |
| 37,8 | | 0,0327002 | 0,0261506 | 0,80 | 1,49E+08 | 1,86E+08 |
| 8,5 | | 0,0327243 | 0,0261427 | 0,55 | 1,89E+07 | 1,32E+07 |
| 16,8 | | 0,0327734 | 0,0261674 | 1,22 | 3,96E+07 | 3,80E+07 |
| 33,5 | | 0,0327860 | 0,0261595 | 1,10 | 3,64E+07 | 3,89E+07 |
| 32,8 | | 0,0327895 | 0,0261516 | 1,30 | 1,81E+08 | 1,79E+08 |
| 19 | | 0,0328710 | 0,0262017 | 1,45 | 2,74E+07 | 3,30E+07 |
| 50,6 | | 0,0328750 | 0,0261938 | 0,85 | 7,41E+08 | 7,15E+08 |
| 40,8 | | 0,0328915 | 0,0261859 | 0,89 | 5,15E+08 | 4,87E+08 |
| 39,1 | | 0,0329253 | 0,0262166 | 0,73 | 1,36E+09 | 1,16E+09 |
| 43,4 | | 0,0329301 | 0,0262087 | 1,42 | 4,29E+07 | 5,14E+07 |
| 8,5 | | 0,0329933 | 0,0262997 | 1,54 | 3,80E+06 | 3,34E+06 |
| 5,7 | | 0,0330400 | 0,0262917 | 2,98 | 3,06E+06 | 4,96E+06 |
| 32,3 | | 0,0330614 | 0,0263537 | 0,80 | 1,48E+08 | 1,75E+08 |
| 9,5 | | 0,0330662 | 0,0263458 | 0,66 | 2,03E+07 | 1,48E+07 |
| 3,1 | | 0,0331267 | 0,0264776 | 2,29 | 1,05E+07 | 4,42E+06 |
| 74,4 | | 0,0331498 | 0,0264696 | 0,89 | 4,61E+09 | 4,59E+09 |
| 14,6 | | 0,0332302 | 0,0264616 | 1,30 | 1,34E+07 | 1,49E+07 |
| 17,5 | | 0,0332304 | 0,0264537 | 1,55 | 8,86E+06 | 9,56E+06 |
| 23,3 | | 0,0332539 | 0,0264457 | 1,46 | 3,62E+07 | 2,57E+07 |
| 29,4 | | 0,0332714 | 0,0264378 | 2,01 | 3,33E+06 | 5,65E+06 |
| 8,3 | | 0,0333291 | 0,0265332 | 0,79 | 1,90E+07 | 2,17E+07 |
| 11,9 | | 0,0333743 | 0,0265901 | 0,34 | 1,11E+07 | 1,58E+07 |
| 28,8 | | 0,0333873 | 0,0265822 | 0,41 | 5,92E+07 | 7,77E+07 |
| 41,6 | | 0,0333896 | 0,0265742 | 0,76 | 1,87E+07 | 1,79E+07 |
| 20,1 | | 0,0334074 | 0,0265662 | 0,62 | 2,58E+07 | 1,83E+07 |
| 20,8 | | 0,0334106 | 0,0265582 | 0,54 | 1,46E+07 | 1,25E+07 |
| 3,6 | | 0,0334820 | 0,0265791 | 1,83 | 2,06E+06 | 3,90E+06 |
| 40,3 | | 0,0335161 | 0,0266371 | 1,09 | 4,89E+08 | 4,51E+08 |
| 54,7 | | 0,0335674 | 0,0267418 | 0,81 | 2,05E+08 | 2,22E+08 |
| 1,8 | | 0,0335827 | 0,0267710 | 2,60 | 2,36E+06 | 5,40E+06 |
| 26,9 | + | 0,0336141 | 0,0267953 | 1,24 | 2,58E+07 | 2,39E+07 |
| 23,4 | | 0,0336249 | 0,0267873 | 1,20 | 1,67E+08 | 1,83E+08 |
| 66,2 | | 0,0336428 | 0,0268164 | 1,20 | 2,07E+08 | 2,41E+08 |
| 24,7 | | 0,0337764 | 0,0269210 | 1,28 | 1,48E+08 | 1,19E+08 |
| 3,5 | | 0,0338133 | 0,0269129 | 3,13 | 4,12E+06 | 4,54E+06 |
| 31,2 | | 0,0338135 | 0,0269048 | 0,80 | 2,07E+08 | 1,94E+08 |
| 37,8 | | 0,0339210 | 0,0270284 | 1,19 | 2,98E+08 | 3,09E+08 |
| 17,8 | | 0,0339466 | 0,0270203 | 0,83 | 1,52E+08 | 1,44E+08 |
| 29,4 | | 0,0339485 | 0,0270123 | 0,74 | 8,76E+07 | 9,01E+07 |
| 44,6 | | 0,0340865 | 0,0270783 | 1,42 | 6,94E+07 | 6,95E+07 |

| | | | | | | |
|---|---|---|---|---|---|---|
| 18,8 | | 0,0341164 | 0,0271085 | 1,73 | 1,64E+07 | 1,52E+07 |
| 14,1 | | 0,0341462 | 0,0271004 | 0,41 | 1,79E+07 | 1,45E+07 |
| 22 | | 0,0341853 | 0,0271472 | 4,15 | 5,92E+06 | 3,24E+06 |
| 22,8 | | 0,0342089 | 0,0271678 | 0,79 | 8,17E+07 | 6,96E+07 |
| 22,7 | | 0,0342865 | 0,0272516 | 0,76 | 1,72E+08 | 2,08E+08 |
| 39,7 | | 0,0343117 | 0,0272434 | 1,21 | 5,29E+07 | 6,09E+07 |
| 61,8 | | 0,0343984 | 0,0272962 | 1,08 | 1,12E+10 | 1,08E+10 |
| 35,2 | | 0,0344785 | 0,0273250 | 0,75 | 5,50E+08 | 5,86E+08 |
| 14,5 | | 0,0345668 | 0,0273538 | 0,46 | 1,74E+07 | 3,24E+07 |
| 52,7 | | 0,0346559 | 0,0274100 | 1,24 | 1,04E+08 | 1,25E+08 |
| 4,9 | | 0,0347298 | 0,0274578 | 0,67 | 1,73E+07 | 1,89E+07 |
| 17,8 | | 0,0347950 | 0,0274866 | 1,16 | 1,03E+07 | 1,15E+07 |
| 11,2 | | 0,0348062 | 0,0274784 | 1,37 | 3,50E+07 | 4,87E+07 |
| 17,4 | | 0,0348476 | 0,0274702 | 0,81 | 1,11E+10 | 1,04E+10 |
| 1,4 | | 0,0348500 | 0,0274621 | 0,52 | 4,28E+06 | 5,75E+06 |
| 48,1 | | 0,0348780 | 0,0274908 | 0,79 | 2,69E+08 | 2,91E+08 |
| 80,6 | | 0,0348997 | 0,0274826 | 0,90 | 5,06E+09 | 5,59E+09 |
| 62,8 | | 0,0349341 | 0,0275125 | 0,41 | 3,55E+07 | 4,21E+07 |
| 39,1 | | 0,0349402 | 0,0275043 | 0,63 | 6,91E+07 | 5,45E+07 |
| 79,7 | | 0,0349697 | 0,0274961 | 1,25 | 2,35E+09 | 2,94E+09 |
| 36,2 | | 0,0349975 | 0,0274880 | 5,32 | 1,21E+07 | 1,93E+06 |
| 24 | | 0,0351238 | 0,0275487 | 0,77 | 9,91E+07 | 1,18E+08 |
| 29,7 | | 0,0351301 | 0,0275785 | 1,31 | 6,60E+07 | 6,19E+07 |
| 25,1 | | 0,0351690 | 0,0276024 | 0,75 | 5,58E+07 | 5,51E+07 |
| 37,3 | | 0,0351855 | 0,0276215 | 1,22 | 7,83E+07 | 8,24E+07 |
| 59,6 | | 0,0352325 | 0,0276453 | 1,19 | 3,71E+08 | 3,25E+08 |
| 4,5 | | 0,0352572 | 0,0277106 | 0,56 | 3,57E+06 | 5,14E+06 |
| 40,1 | | 0,0353147 | 0,0277024 | 1,73 | 2,53E+08 | 1,42E+08 |
| 9,9 | | 0,0353215 | 0,0276942 | 1,66 | 7,58E+06 | 1,03E+07 |
| 48,6 | | 0,0353436 | 0,0277227 | 0,76 | 5,15E+08 | 5,64E+08 |
| 23,2 | | 0,0353783 | 0,0277513 | 0,45 | 1,08E+07 | 1,33E+07 |
| 28,8 | | 0,0353980 | 0,0278165 | 0,78 | 1,37E+08 | 1,25E+08 |
| 41,2 | | 0,0354111 | 0,0278082 | 0,84 | 2,46E+08 | 2,80E+08 |
| 18,2 | | 0,0355082 | 0,0278686 | 1,84 | 2,02E+07 | 2,56E+07 |
| 28,3 | | 0,0355873 | 0,0278971 | 1,42 | 2,50E+07 | 2,49E+07 |
| 33,7 | | 0,0356307 | 0,0278888 | 1,15 | 1,01E+09 | 9,90E+08 |
| 7,3 | | 0,0357168 | 0,0279172 | 1,37 | 2,05E+07 | 2,32E+07 |
| 19,3 | | 0,0357230 | 0,0279374 | 0,76 | 1,93E+08 | 1,45E+08 |
| 24,9 | | 0,0357830 | 0,0279941 | 14,95 | 1,29E+07 | 8,02E+05 |
| 10,1 | | 0,0358003 | 0,0279858 | 0,65 | 1,14E+07 | 1,33E+07 |
| 42,7 | | 0,0358838 | 0,0280709 | 1,29 | 3,43E+08 | 3,20E+08 |
| 15,5 | | 0,0359523 | 0,0280626 | 7,40 | 1,68E+06 | 1,10E+07 |
| 11,6 | | 0,0359630 | 0,0280543 | 0,74 | 1,25E+07 | 1,34E+07 |
| 16,7 | | 0,0360025 | 0,0281109 | 2,15 | 1,10E+07 | 1,65E+07 |
| 64,4 | | 0,0360064 | 0,0281026 | 0,89 | 1,60E+09 | 1,62E+09 |
| 9 | | 0,0360162 | 0,0280943 | 0,62 | 1,51E+07 | 2,09E+07 |
| 27 | | 0,0361978 | 0,0281816 | 0,78 | 1,66E+08 | 1,42E+08 |
| 8,2 | | 0,0362116 | 0,0281732 | 1,49 | 2,44E+06 | 3,25E+06 |
| 12,5 | | 0,0362707 | 0,0282015 | 0,49 | 1,15E+08 | 1,12E+08 |
| 11 | | 0,0363386 | 0,0282980 | 1,36 | 3,89E+07 | 4,70E+07 |

| | | | | | | |
|---|---|---|---|---|---|---|
| 52,8 | | 0,0363598 | 0,0283627 | 0,83 | 2,08E+09 | 1,87E+09 |
| 48,2 | | 0,0363900 | 0,0283543 | 0,74 | 1,37E+08 | 1,58E+08 |
| 58,7 | | 0,0365415 | 0,0284519 | 1,16 | 4,65E+08 | 4,36E+08 |
| 18,3 | | 0,0366331 | 0,0285718 | 0,70 | 1,53E+07 | 2,01E+07 |
| 40,1 | | 0,0366852 | 0,0285634 | 1,14 | 3,50E+08 | 3,92E+08 |
| 32,7 | | 0,0366998 | 0,0285550 | 0,66 | 1,92E+08 | 1,83E+08 |
| 35,8 | | 0,0368216 | 0,0286112 | 1,22 | 5,17E+07 | 5,84E+07 |
| 30 | | 0,0368737 | 0,0286310 | 0,41 | 6,14E+07 | 7,11E+07 |
| 35,1 | | 0,0369167 | 0,0286590 | 0,79 | 5,69E+07 | 4,62E+07 |
| 30,1 | | 0,0369864 | 0,0287798 | 1,09 | 4,63E+07 | 4,88E+07 |
| 1,8 | | 0,0370069 | 0,0287995 | 0,81 | 4,39E+06 | 3,83E+06 |
| 28,3 | | 0,0370142 | 0,0287911 | 1,35 | 5,00E+07 | 6,49E+07 |
| 21,3 | | 0,0370279 | 0,0288108 | 0,25 | 1,30E+07 | 1,04E+07 |
| 29,7 | | 0,0370282 | 0,0288023 | 0,26 | 1,33E+07 | 1,26E+07 |
| 26,8 | | 0,0371787 | 0,0290026 | 0,75 | 4,20E+07 | 3,66E+07 |
| 6,8 | | 0,0372023 | 0,0289941 | 0,40 | 2,64E+07 | 1,65E+07 |
| 12,5 | | 0,0372138 | 0,0290220 | 1,68 | 7,09E+06 | 7,81E+06 |
| 63,7 | | 0,0372410 | 0,0290135 | 0,93 | 6,51E+09 | 6,87E+09 |
| 27,9 | | 0,0374245 | 0,0291877 | 1,41 | 6,34E+06 | 5,06E+06 |
| 33,3 | | 0,0374333 | 0,0291792 | 1,24 | 8,93E+07 | 1,02E+08 |
| 24 | | 0,0374822 | 0,0292245 | 0,75 | 1,02E+08 | 1,21E+08 |
| 44,9 | | 0,0375502 | 0,0293879 | 0,88 | 1,04E+09 | 1,09E+09 |
| 27,2 | | 0,0376150 | 0,0294355 | 1,16 | 3,78E+08 | 3,37E+08 |
| 24 | | 0,0376797 | 0,0294632 | 0,70 | 5,30E+07 | 7,88E+07 |
| 25,4 | | 0,0377110 | 0,0294545 | 1,23 | 7,49E+07 | 6,65E+07 |
| 54,7 | | 0,0377510 | 0,0294459 | 1,23 | 2,32E+08 | 2,55E+08 |
| 21,8 | | 0,0377563 | 0,0294373 | 1,96 | 1,09E+08 | 1,80E+08 |
| 27,1 | | 0,0379381 | 0,0294883 | 0,44 | 1,33E+07 | 8,71E+06 |
| 14,8 | | 0,0379745 | 0,0294797 | 0,78 | 1,45E+07 | 1,48E+07 |
| 8,4 | | 0,0379756 | 0,0294711 | 2,87 | 6,69E+06 | 9,63E+06 |
| 43,3 | | 0,0379773 | 0,0294625 | 0,68 | 1,98E+09 | 1,47E+09 |
| 18,1 | | 0,0380095 | 0,0294539 | 1,18 | 4,90E+08 | 4,34E+08 |
| 69,4 | | 0,0380135 | 0,0294453 | 1,71 | 7,90E+07 | 5,12E+07 |
| 21,8 | | 0,0381474 | 0,0296012 | 1,21 | 5,71E+07 | 6,38E+07 |
| 30,9 | | 0,0381556 | 0,0295925 | 0,72 | 7,45E+07 | 8,00E+07 |
| 18,6 | | 0,0381740 | 0,0295839 | 0,70 | 9,92E+07 | 1,32E+08 |
| 57,1 | | 0,0382377 | 0,0296033 | 0,76 | 1,17E+08 | 1,39E+08 |
| 72,6 | | 0,0382925 | 0,0295946 | 0,83 | 3,08E+08 | 3,64E+08 |
| 61,1 | | 0,0382995 | 0,0295860 | 1,16 | 3,83E+09 | 3,38E+09 |
| 26 | | 0,0383147 | 0,0295774 | 0,82 | 7,23E+08 | 6,82E+08 |
| 27,6 | | 0,0383367 | 0,0295688 | 0,96 | 2,31E+08 | 2,26E+08 |
| 27,3 | | 0,0383460 | 0,0296230 | 0,74 | 5,51E+08 | 5,61E+08 |
| 10,7 | | 0,0384421 | 0,0296889 | 1,22 | 9,57E+06 | 1,13E+07 |
| 57,1 | | 0,0384538 | 0,0296802 | 1,21 | 4,06E+08 | 3,94E+08 |
| 7,9 | | 0,0385946 | 0,0297902 | 0,59 | 8,96E+06 | 1,07E+07 |
| 3,8 | | 0,0386311 | 0,0297815 | 0,39 | 6,25E+06 | 1,38E+07 |
| 34,9 | | 0,0386832 | 0,0299239 | 1,14 | 9,71E+07 | 8,93E+07 |
| 11,6 | | 0,0387169 | 0,0299152 | 1,28 | 1,80E+07 | 2,37E+07 |
| 22,8 | | 0,0387287 | 0,0299065 | 1,33 | 6,84E+07 | 6,92E+07 |
| 22,8 | | 0,0387904 | 0,0299292 | 1,34 | 2,03E+07 | 2,12E+07 |

| | | | | | | |
|---|---|---|---|---|---|---|
| 55,9 | | 0,0387994 | 0,0299205 | 0,59 | 4,81E+08 | 6,80E+08 |
| 37,2 | | 0,0388415 | 0,0299118 | 0,74 | 5,04E+07 | 4,12E+07 |
| 27,3 | | 0,0388638 | 0,0299032 | 1,40 | 2,36E+07 | 2,18E+07 |
| 37,3 | | 0,0388775 | 0,0299223 | 1,46 | 3,56E+08 | 5,11E+08 |
| 28,5 | | 0,0389448 | 0,0300041 | 0,82 | 1,65E+08 | 1,47E+08 |
| 41,2 | | 0,0389934 | 0,0300545 | 0,86 | 6,17E+08 | 6,09E+08 |
| 19 | | 0,0390212 | 0,0301454 | 0,83 | 1,22E+08 | 1,16E+08 |
| 39,2 | | 0,0390266 | 0,0301367 | 1,31 | 3,61E+08 | 4,14E+08 |
| 36,7 | | 0,0390741 | 0,0301904 | 1,05 | 9,49E+08 | 9,24E+08 |
| 10,3 | | 0,0390904 | 0,0301817 | 0,30 | 1,67E+07 | 2,03E+07 |
| 8,5 | | 0,0391356 | 0,0302366 | 0,29 | 2,02E+07 | 1,61E+07 |
| 28,7 | | 0,0391413 | 0,0302279 | 1,42 | 4,53E+08 | 6,54E+08 |
| 24,6 | | 0,0391652 | 0,0302191 | 1,58 | 3,65E+07 | 3,67E+07 |
| 13,9 | | 0,0392538 | 0,0303283 | 0,79 | 2,91E+07 | 3,01E+07 |
| 2,6 | | 0,0394682 | 0,0305403 | 0,41 | 8,97E+06 | 4,17E+06 |
| 15,9 | | 0,0395693 | 0,0306528 | 1,16 | 4,67E+07 | 5,13E+07 |
| 57,7 | | 0,0395907 | 0,0306751 | 1,22 | 7,67E+08 | 6,45E+08 |
| 12,7 | | 0,0396296 | 0,0306663 | 0,67 | 9,85E+06 | 1,26E+07 |
| 10,7 | | 0,0396356 | 0,0306574 | 0,71 | 9,76E+06 | 1,17E+07 |
| 17,6 | | 0,0397048 | 0,0307109 | 0,76 | 5,96E+07 | 6,80E+07 |
| 20,1 | | 0,0398004 | 0,0308647 | 0,69 | 3,67E+08 | 3,47E+08 |
| 4,8 | | 0,0398361 | 0,0308558 | 0,44 | 9,30E+06 | 5,71E+06 |
| 29,2 | | 0,0398725 | 0,0308838 | 0,87 | 3,66E+08 | 3,64E+08 |
| 17,3 | | 0,0399537 | 0,0309683 | 1,48 | 1,00E+07 | 1,21E+07 |
| 58,6 | | 0,0399810 | 0,0309594 | 0,75 | 3,28E+08 | 2,61E+08 |
| 32,7 | | 0,0399988 | 0,0309505 | 0,90 | 1,29E+08 | 1,23E+08 |
| 8,1 | | 0,0400070 | 0,0309415 | 0,48 | 2,13E+07 | 1,63E+07 |
| 23,1 | | 0,0400432 | 0,0310697 | 1,34 | 4,47E+07 | 4,85E+07 |
| 53,6 | | 0,0400809 | 0,0310607 | 0,88 | 2,89E+09 | 2,95E+09 |
| 41,5 | | 0,0401238 | 0,0310794 | 0,88 | 3,22E+08 | 3,14E+08 |
| 9,5 | | 0,0401400 | 0,0310705 | 0,19 | 3,69E+07 | 3,30E+07 |
| 44,3 | | 0,0401564 | 0,0310615 | 0,80 | 2,30E+10 | 2,11E+10 |
| 9,4 | | 0,0401975 | 0,0311147 | 2,21 | 4,62E+06 | 4,76E+06 |
| 43,4 | | 0,0403462 | 0,0312552 | 0,68 | 2,50E+08 | 3,55E+08 |
| 69,7 | | 0,0403498 | 0,0312462 | 0,90 | 2,13E+09 | 2,40E+09 |
| 46,8 | | 0,0404710 | 0,0313038 | 1,12 | 1,10E+09 | 1,17E+09 |
| 35,9 | | 0,0404752 | 0,0312949 | 0,75 | 6,58E+07 | 6,90E+07 |
| 11,4 | | 0,0405076 | 0,0312859 | 0,55 | 1,84E+07 | 3,33E+07 |
| 41,7 | | 0,0405536 | 0,0313125 | 4,74 | 2,79E+06 | 4,60E+06 |
| 12,4 | | 0,0406608 | 0,0314664 | 0,24 | 6,29E+06 | 9,39E+06 |
| 76,4 | | 0,0406681 | 0,0314574 | 1,19 | 7,38E+08 | 8,69E+08 |
| 35,9 | | 0,0407477 | 0,0315814 | 1,23 | 2,86E+08 | 2,86E+08 |
| 10,3 | | 0,0407803 | 0,0316079 | 0,52 | 2,97E+07 | 3,06E+07 |
| 57,1 | | 0,0407822 | 0,0315989 | 0,79 | 3,24E+08 | 3,99E+08 |
| 55,3 | | 0,0408441 | 0,0316803 | 0,73 | 3,61E+07 | 3,49E+07 |
| 17,3 | | 0,0408781 | 0,0316712 | 0,58 | 4,80E+06 | 8,03E+06 |
| 21,7 | | 0,0410134 | 0,0316977 | 0,81 | 1,95E+08 | 1,56E+08 |
| 8,6 | | 0,0410718 | 0,0316886 | 0,64 | 3,21E+07 | 3,73E+07 |
| 14,5 | | 0,0411394 | 0,0317516 | 1,32 | 2,14E+07 | 2,53E+07 |
| 62,7 | | 0,0412449 | 0,0318272 | 0,91 | 3,97E+09 | 4,29E+09 |

| | | | | | | |
|---|---|---|---|---|---|---|
| 46,5 | | 0,0412586 | 0,0318181 | 0,92 | 1,16E+09 | 1,22E+09 |
| 40,3 | | 0,0413492 | 0,0319726 | 1,26 | 1,13E+08 | 9,76E+07 |
| 35,8 | | 0,0413862 | 0,0319634 | 0,87 | 3,34E+08 | 3,45E+08 |
| 26,6 | | 0,0413935 | 0,0319543 | 2,49 | 6,20E+06 | 9,51E+06 |
| 8,4 | | 0,0414629 | 0,0319806 | 1,75 | 1,12E+07 | 2,02E+07 |
| 46,4 | | 0,0414846 | 0,0321005 | 0,70 | 9,17E+07 | 7,98E+07 |
| 10,5 | | 0,0415300 | 0,0320914 | 1,27 | 1,62E+07 | 1,57E+07 |
| 35,6 | | 0,0415430 | 0,0320822 | 0,81 | 1,49E+08 | 1,47E+08 |
| 53,6 | | 0,0415726 | 0,0320730 | 0,82 | 7,97E+08 | 6,93E+08 |
| 54,5 | | 0,0415971 | 0,0320639 | 0,66 | 3,34E+08 | 2,98E+08 |
| 70,3 | | 0,0416757 | 0,0321084 | 0,87 | 3,45E+08 | 3,74E+08 |
| 24,4 | | 0,0417339 | 0,0321266 | 1,27 | 4,43E+08 | 3,85E+08 |
| 77,3 | | 0,0417718 | 0,0321174 | 1,20 | 3,25E+10 | 2,67E+10 |
| 45,5 | | 0,0417753 | 0,0321083 | 1,34 | 1,32E+08 | 1,79E+08 |
| 16,7 | | 0,0418716 | 0,0321299 | 1,30 | 5,70E+07 | 5,67E+07 |
| 35,3 | | 0,0419274 | 0,0321560 | 1,41 | 3,44E+08 | 4,95E+08 |
| 54,9 | | 0,0419681 | 0,0321776 | 1,40 | 6,08E+08 | 5,09E+08 |
| 27,2 | | 0,0420077 | 0,0321958 | 1,50 | 3,50E+07 | 3,76E+07 |
| 54,4 | | 0,0420609 | 0,0322526 | 1,17 | 3,47E+08 | 3,55E+08 |
| 13,6 | | 0,0421124 | 0,0323140 | 0,78 | 1,79E+07 | 1,73E+07 |
| 7,3 | | 0,0421312 | 0,0323048 | 0,77 | 1,18E+07 | 1,35E+07 |
| 3 | | 0,0421820 | 0,0323843 | 0,23 | 1,59E+07 | 1,76E+07 |
| 33,5 | | 0,0422323 | 0,0324910 | 0,81 | 2,15E+08 | 2,11E+08 |
| 27,5 | | 0,0422355 | 0,0324818 | 1,07 | 4,28E+08 | 4,26E+08 |
| 32,2 | | 0,0422545 | 0,0324987 | 0,81 | 4,97E+08 | 4,88E+08 |
| 35,3 | | 0,0423015 | 0,0325599 | 0,88 | 2,29E+08 | 2,50E+08 |
| 56,4 | | 0,0423334 | 0,0325813 | 1,12 | 3,19E+08 | 3,05E+08 |
| 24 | | 0,0423547 | 0,0325993 | 1,19 | 4,66E+07 | 5,43E+07 |
| 48,8 | | 0,0424127 | 0,0326173 | 1,12 | 6,51E+08 | 7,31E+08 |
| 15,8 | | 0,0424151 | 0,0326081 | 0,68 | 1,19E+08 | 9,58E+07 |
| 6,3 | | 0,0424921 | 0,0326340 | 5,61 | 1,86E+06 | 1,41E+07 |
| 23,3 | | 0,0424939 | 0,0326247 | 0,87 | 1,77E+08 | 1,74E+08 |
| 11,7 | | 0,0425178 | 0,0326506 | 0,17 | 6,53E+07 | 4,92E+07 |
| 9,7 | | 0,0425374 | 0,0326765 | 0,33 | 1,73E+07 | 1,75E+07 |
| 41,9 | | 0,0425974 | 0,0327567 | 0,69 | 3,09E+07 | 3,97E+07 |
| 9,7 | | 0,0426637 | 0,0328177 | 2,34 | 2,76E+06 | 3,09E+06 |
| 3,2 | | 0,0427555 | 0,0328084 | 1,48 | 1,53E+07 | 1,91E+07 |
| 29,5 | | 0,0429284 | 0,0329621 | 0,60 | 9,74E+06 | 1,48E+07 |
| 45,7 | | 0,0429326 | 0,0329833 | 0,81 | 7,39E+08 | 7,07E+08 |
| 37,2 | | 0,0429369 | 0,0329740 | 1,78 | 3,69E+07 | 6,73E+07 |
| 43,5 | | 0,0429766 | 0,0330269 | 1,22 | 1,14E+08 | 9,24E+07 |
| 7,6 | | 0,0430102 | 0,0330526 | 2,48 | 2,00E+06 | 5,29E+06 |
| 28,5 | | 0,0430528 | 0,0331133 | 1,21 | 8,43E+07 | 7,23E+07 |
| 53,2 | | 0,0430712 | 0,0331040 | 1,30 | 1,52E+09 | 1,56E+09 |
| 47 | | 0,0430792 | 0,0331206 | 0,94 | 8,33E+08 | 8,29E+08 |
| 5,8 | | 0,0431193 | 0,0331112 | 0,18 | 6,48E+06 | 1,28E+07 |
| 55,1 | | 0,0433606 | 0,0334056 | 1,13 | 4,38E+09 | 4,27E+09 |
| 39,9 | | 0,0434199 | 0,0334661 | 0,82 | 4,15E+08 | 3,67E+08 |
| 21,4 | | 0,0434574 | 0,0334567 | 2,06 | 4,06E+07 | 1,85E+07 |
| 3,2 | | 0,0434625 | 0,0334473 | 0,73 | 2,03E+07 | 1,70E+07 |

| | | | | | | |
|---|---|---|---|---|---|---|
| 6,7 | | 0,0435162 | 0,0334739 | 0,82 | 2,57E+07 | 3,15E+07 |
| 19,8 | | 0,0436306 | 0,0335693 | 1,76 | 1,08E+07 | 9,42E+06 |
| 12,2 | | 0,0436502 | 0,0335599 | 2,77 | 3,03E+06 | 1,47E+06 |
| 27,3 | | 0,0436674 | 0,0335763 | 1,21 | 4,67E+07 | 4,00E+07 |
| 7,5 | | 0,0437394 | 0,0337280 | 1,33 | 1,92E+07 | 1,81E+07 |
| 20,5 | | 0,0438233 | 0,0338063 | 2,95 | 1,95E+07 | 1,23E+07 |
| 17,4 | | 0,0438344 | 0,0337968 | 1,26 | 2,16E+07 | 2,20E+07 |
| 16,7 | | 0,0438480 | 0,0338132 | 0,59 | 4,85E+06 | 6,25E+06 |
| 7,6 | | 0,0438530 | 0,0338037 | 1,55 | 2,75E+06 | 2,69E+06 |
| 23,5 | | 0,0438729 | 0,0338211 | 0,76 | 5,46E+07 | 5,82E+07 |
| 26,9 | | 0,0440089 | 0,0338814 | 0,86 | 8,67E+07 | 1,03E+08 |
| 10,9 | | 0,0440644 | 0,0339258 | 1,26 | 1,59E+07 | 1,44E+07 |
| 35,2 | | 0,0440971 | 0,0339511 | 0,81 | 1,49E+08 | 1,28E+08 |
| 3,5 | | 0,0441198 | 0,0339685 | 0,23 | 2,24E+07 | 1,73E+07 |
| 6,8 | | 0,0441906 | 0,0340567 | 1,29 | 1,65E+07 | 1,34E+07 |
| 54,5 | | 0,0442616 | 0,0341078 | 1,14 | 4,27E+09 | 4,88E+09 |
| 34,2 | | 0,0442835 | 0,0340982 | 0,82 | 1,83E+08 | 2,11E+08 |
| 54,4 | | 0,0442895 | 0,0341190 | 0,87 | 2,76E+09 | 2,57E+09 |
| 32,8 | | 0,0442970 | 0,0341094 | 1,38 | 2,74E+08 | 2,94E+08 |
| 18 | | 0,0443094 | 0,0341346 | 0,86 | 2,06E+08 | 2,07E+08 |
| 10,6 | + | 0,0443432 | 0,0341598 | 4,29 | 5,94E+06 | 2,36E+06 |
| 17 | | 0,0443817 | 0,0341502 | 0,77 | 7,19E+07 | 6,66E+07 |
| 9,6 | | 0,0443842 | 0,0341407 | 0,11 | 3,16E+07 | 2,13E+07 |
| 9,7 | | 0,0443942 | 0,0341311 | 0,82 | 4,35E+07 | 4,42E+07 |
| 55,2 | | 0,0444043 | 0,0341518 | 0,86 | 3,33E+08 | 3,72E+08 |
| 7,9 | | 0,0444110 | 0,0341422 | 3,24 | 1,31E+07 | 3,22E+06 |
| 24,1 | | 0,0444187 | 0,0341327 | 0,75 | 4,01E+07 | 3,58E+07 |
| 28,1 | | 0,0445651 | 0,0342507 | 1,18 | 2,09E+08 | 2,13E+08 |
| 6,4 | | 0,0445913 | 0,0342713 | 2,11 | 1,87E+06 | 1,27E+06 |
| 30,6 | | 0,0447083 | 0,0345000 | 1,63 | 3,92E+07 | 5,80E+07 |
| 12,8 | | 0,0447094 | 0,0344904 | 0,66 | 5,02E+07 | 4,52E+07 |
| 12,1 | | 0,0447227 | 0,0345064 | 0,22 | 2,15E+07 | 1,56E+07 |
| 1,3 | | 0,0448621 | 0,0345873 | 3,18 | 3,47E+06 | 1,10E+07 |
| 35,4 | | 0,0448778 | 0,0346123 | 1,33 | 9,13E+07 | 7,18E+07 |
| 21,1 | | 0,0449518 | 0,0346976 | 1,23 | 3,01E+07 | 3,35E+07 |
| 33,8 | | 0,0450224 | 0,0347884 | 1,44 | 3,19E+07 | 2,28E+07 |
| 24,7 | | 0,0450911 | 0,0348401 | 0,81 | 1,27E+08 | 1,09E+08 |
| 17,5 | | 0,0450921 | 0,0348304 | 0,20 | 4,41E+07 | 4,20E+07 |
| 12,7 | | 0,0452251 | 0,0349166 | 1,42 | 6,08E+07 | 7,84E+07 |
| 45,1 | | 0,0452934 | 0,0349426 | 0,76 | 1,68E+08 | 1,96E+08 |
| 43,5 | | 0,0453369 | 0,0349941 | 0,85 | 8,94E+08 | 9,52E+08 |
| 22,6 | | 0,0454045 | 0,0350111 | 0,25 | 6,49E+06 | 1,13E+07 |
| 15,5 | | 0,0454090 | 0,0350014 | 0,59 | 1,39E+08 | 1,47E+08 |
| 8,3 | + | 0,0455025 | 0,0350184 | 2,41 | 8,36E+06 | 3,96E+06 |
| 30,9 | | 0,0456267 | 0,0351423 | 1,34 | 8,63E+07 | 7,67E+07 |
| 20,7 | | 0,0456946 | 0,0351626 | 0,63 | 2,23E+07 | 2,13E+07 |
| 37 | | 0,0456973 | 0,0351528 | 2,49 | 2,28E+07 | 2,09E+07 |
| 36,7 | | 0,0457184 | 0,0351430 | 1,36 | 7,44E+07 | 5,20E+07 |
| 59,4 | | 0,0458100 | 0,0351955 | 1,28 | 1,25E+08 | 1,36E+08 |
| 15,7 | | 0,0458281 | 0,0351858 | 0,37 | 6,06E+07 | 5,02E+07 |

| | | | | | | |
|---|---|---|---|---|---|---|
| 45,3 | | 0,0459052 | 0,0352449 | 0,68 | 1,14E+08 | 9,86E+07 |
| 40 | | 0,0459233 | 0,0352351 | 1,28 | 3,22E+07 | 2,78E+07 |
| 26,5 | | 0,0459499 | 0,0352253 | 0,75 | 2,46E+08 | 3,24E+08 |
| 57,7 | | 0,0459993 | 0,0352500 | 1,08 | 1,18E+09 | 1,17E+09 |
| 23,3 | | 0,0460025 | 0,0352702 | 0,68 | 3,34E+07 | 3,25E+07 |
| 44,4 | | 0,0460986 | 0,0352871 | 1,25 | 4,83E+08 | 4,12E+08 |
| 46,1 | | 0,0461835 | 0,0353417 | 0,86 | 5,50E+08 | 6,16E+08 |
| 34,2 | | 0,0462829 | 0,0353663 | 1,43 | 3,27E+08 | 4,06E+08 |
| 28,3 | | 0,0463150 | 0,0353908 | 0,92 | 1,40E+08 | 1,45E+08 |
| 69,5 | | 0,0464078 | 0,0354498 | 1,10 | 1,56E+10 | 1,45E+10 |
| 21,9 | | 0,0464102 | 0,0354400 | 0,79 | 1,04E+08 | 1,09E+08 |
| 28,9 | | 0,0465512 | 0,0356164 | 0,77 | 2,47E+07 | 2,28E+07 |
| 20,6 | | 0,0466005 | 0,0356409 | 1,22 | 9,99E+07 | 1,05E+08 |
| 37,7 | | 0,0466520 | 0,0356909 | 0,72 | 4,43E+08 | 3,79E+08 |
| 30,8 | | 0,0467149 | 0,0356810 | 2,42 | 6,14E+06 | 1,06E+07 |
| 38,5 | | 0,0467202 | 0,0356711 | 1,24 | 4,27E+07 | 4,28E+07 |
| 53 | | 0,0467360 | 0,0356612 | 1,15 | 2,26E+09 | 2,66E+09 |
| 28,9 | | 0,0467403 | 0,0356514 | 0,81 | 4,19E+07 | 4,58E+07 |
| 9,7 | | 0,0467893 | 0,0356669 | 0,50 | 1,34E+07 | 1,26E+07 |
| 11,2 | | 0,0468412 | 0,0357135 | 1,72 | 3,06E+07 | 5,63E+07 |
| 10,1 | | 0,0468776 | 0,0357335 | 1,18 | 2,30E+07 | 2,03E+07 |
| 14,3 | | 0,0469835 | 0,0357877 | 1,37 | 1,85E+07 | 1,51E+07 |
| 50,5 | | 0,0470103 | 0,0357778 | 1,22 | 3,00E+07 | 3,15E+07 |
| 30,4 | | 0,0470459 | 0,0357680 | 1,66 | 4,38E+07 | 2,52E+07 |
| 16,4 | | 0,0470466 | 0,0357581 | 1,62 | 5,66E+06 | 6,39E+06 |
| 59,3 | | 0,0472730 | 0,0359602 | 1,60 | 5,28E+07 | 6,83E+07 |
| 51,9 | | 0,0472880 | 0,0359503 | 1,13 | 2,59E+08 | 2,26E+08 |
| 16,4 | | 0,0472966 | 0,0359404 | 1,59 | 7,21E+06 | 8,70E+06 |
| 27,7 | | 0,0473809 | 0,0362273 | 0,84 | 1,46E+09 | 1,29E+09 |
| 21 | | 0,0473850 | 0,0362173 | 1,28 | 1,29E+08 | 1,42E+08 |
| 17,1 | | 0,0473990 | 0,0362073 | 0,92 | 2,86E+07 | 2,72E+07 |
| 5 | | 0,0474140 | 0,0361974 | 0,61 | 1,70E+07 | 1,33E+07 |
| 29,4 | | 0,0474791 | 0,0361874 | 1,87 | 1,85E+07 | 2,71E+07 |
| 12,1 | | 0,0475930 | 0,0362116 | 0,52 | 6,09E+06 | 3,19E+06 |
| 37,5 | | 0,0476415 | 0,0362313 | 1,13 | 1,09E+09 | 1,22E+09 |
| 13,6 | | 0,0476632 | 0,0362214 | 1,22 | 5,87E+07 | 5,41E+07 |
| 29,1 | | 0,0476982 | 0,0362114 | 0,77 | 1,37E+09 | 1,35E+09 |
| 69,9 | | 0,0477784 | 0,0362312 | 1,21 | 1,33E+09 | 1,27E+09 |
| 49,3 | | 0,0477862 | 0,0362509 | 0,73 | 3,87E+07 | 4,89E+07 |
| 6,1 | | 0,0478216 | 0,0362673 | 2,27 | 3,41E+06 | 4,37E+06 |
| 39,5 | | 0,0479541 | 0,0364102 | 0,86 | 2,68E+08 | 2,73E+08 |
| 36,3 | | 0,0481199 | 0,0366531 | 0,79 | 2,53E+08 | 3,16E+08 |
| 36,7 | | 0,0482044 | 0,0367552 | 1,26 | 8,57E+07 | 8,49E+07 |
| 16,6 | | 0,0483440 | 0,0368527 | 1,16 | 5,30E+07 | 5,92E+07 |
| 38,6 | | 0,0483996 | 0,0368767 | 1,46 | 2,22E+07 | 3,37E+07 |
| 13,1 | | 0,0484321 | 0,0369270 | 0,34 | 1,48E+07 | 5,27E+06 |
| 15,6 | | 0,0484870 | 0,0369168 | 0,87 | 6,52E+07 | 7,17E+07 |
| 36,8 | | 0,0484905 | 0,0369067 | 1,29 | 1,01E+08 | 9,62E+07 |
| 16,9 | | 0,0485365 | 0,0368966 | 0,57 | 5,78E+07 | 5,27E+07 |
| 81 | | 0,0485512 | 0,0369205 | 0,86 | 2,97E+09 | 3,27E+09 |

| | | | | | | |
|---|---|---|---|---|---|---|
| 31,5 | | 0,0486413 | 0,0369740 | 0,81 | 5,67E+07 | 4,84E+07 |
| 33 | | 0,0486692 | 0,0369638 | 0,89 | 1,66E+08 | 1,64E+08 |
| 36,2 | | 0,0487330 | 0,0370063 | 1,58 | 6,31E+06 | 5,54E+06 |
| 36,2 | | 0,0487622 | 0,0369962 | 1,19 | 1,25E+08 | 1,35E+08 |
| 13,7 | | 0,0488362 | 0,0370759 | 0,91 | 8,98E+07 | 8,78E+07 |
| 14,6 | | 0,0488500 | 0,0370920 | 3,55 | 1,74E+06 | 4,30E+06 |
| 22 | | 0,0489036 | 0,0371935 | 1,11 | 3,57E+07 | 3,49E+07 |
| 66,2 | | 0,0489574 | 0,0372173 | 1,25 | 1,56E+09 | 1,71E+09 |
| 3,8 | | 0,0489800 | 0,0372071 | 0,77 | 2,54E+07 | 2,15E+07 |
| 38,2 | | 0,0489846 | 0,0371969 | 0,80 | 2,61E+08 | 2,83E+08 |
| 30,2 | | 0,0490541 | 0,0373049 | 1,53 | 6,31E+07 | 6,77E+07 |
| 56,1 | | 0,0490831 | 0,0372947 | 0,85 | 6,36E+09 | 6,18E+09 |
| 33,7 | | 0,0490915 | 0,0372845 | 0,75 | 2,41E+08 | 1,82E+08 |
| 33,7 | | 0,0491636 | 0,0373038 | 1,18 | 3,53E+07 | 3,25E+07 |
| 14,9 | | 0,0492190 | 0,0373275 | 0,62 | 2,88E+08 | 1,89E+08 |
| 54,2 | | 0,0493808 | 0,0374451 | 0,78 | 1,82E+08 | 2,18E+08 |
| 27,7 | | 0,0494773 | 0,0375364 | 1,16 | 9,52E+07 | 8,31E+07 |
| 18,9 | | 0,0495156 | 0,0375600 | 1,27 | 2,16E+07 | 2,47E+07 |
| 49,7 | | 0,0496258 | 0,0375498 | 0,88 | 1,01E+08 | 1,13E+08 |
| 19,7 | | 0,0496362 | 0,0375396 | 0,34 | 2,87E+07 | 2,49E+07 |
| 23,4 | | 0,0496413 | 0,0375544 | 2,66 | 7,63E+07 | 3,89E+07 |
| 25,6 | | 0,0496886 | 0,0375442 | 3,04 | 3,43E+06 | 8,82E+06 |
| 13,8 | | 0,0497191 | 0,0375339 | 1,28 | 3,71E+07 | 2,89E+07 |
| 37,6 | | 0,0497224 | 0,0375237 | 1,67 | 6,48E+06 | 6,89E+06 |
| 15,5 | | 0,0497286 | 0,0375135 | 2,77 | 3,89E+06 | 1,14E+07 |
| 17,8 | | 0,0497523 | 0,0375294 | 0,72 | 1,91E+07 | 1,95E+07 |
| 22,7 | | 0,0498317 | 0,0376379 | 0,70 | 1,66E+07 | 2,27E+07 |
| 2,3 | | 0,0498553 | 0,0376277 | 0,48 | 1,13E+07 | 9,27E+06 |
| 24,4 | | 0,0498584 | 0,0376174 | 0,76 | 1,39E+08 | 1,45E+08 |
| 39,9 | | 0,0498603 | 0,0376072 | 0,91 | 2,34E+09 | 2,14E+09 |
| 10,4 | | 0,0498623 | 0,0375970 | 1,42 | 5,69E+06 | 5,53E+06 |
| 18,8 | | 0,0498743 | 0,0375867 | 0,75 | 2,94E+08 | 2,45E+08 |
| 17,4 | | 0,0499143 | 0,0376026 | 0,82 | 3,61E+07 | 3,15E+07 |
| 23,4 | | 0,0499246 | 0,0375924 | 0,59 | 3,81E+07 | 4,28E+07 |
| 35,5 | | 0,0499306 | 0,0375822 | 1,36 | 1,16E+08 | 1,07E+08 |
| 5,7 | | 0,0499628 | 0,0376067 | 1,61 | 4,46E+06 | 3,99E+06 |
| 23,8 | | 0,0500270 | 0,0375965 | 1,12 | 5,30E+07 | 5,94E+07 |
| 21,7 | | 0,0501326 | 0,0377003 | 0,72 | 2,42E+08 | 2,20E+08 |
| 50,2 | | 0,0502044 | 0,0377531 | 0,90 | 7,90E+08 | 7,21E+08 |
| 16 | | 0,0503143 | 0,0379034 | 2,08 | 2,42E+07 | 2,97E+07 |
| 14,7 | | 0,0505144 | 0,0380146 | 1,88 | 1,69E+07 | 9,42E+06 |
| 11,1 | | 0,0506677 | 0,0381020 | 0,76 | 1,80E+08 | 1,57E+08 |
| 26,3 | | 0,0506900 | 0,0380916 | 0,80 | 2,69E+08 | 2,78E+08 |
| 13,3 | | 0,0507040 | 0,0381409 | 1,24 | 7,32E+07 | 6,81E+07 |
| 14,2 | | 0,0507720 | 0,0382422 | 0,83 | 5,27E+07 | 4,32E+07 |
| 33,1 | | 0,0509065 | 0,0384171 | 1,32 | 4,38E+07 | 5,38E+07 |
| 6,5 | | 0,0510634 | 0,0385919 | 0,74 | 1,18E+07 | 1,63E+07 |
| 20,5 | | 0,0510731 | 0,0385815 | 1,27 | 7,57E+07 | 6,82E+07 |
| 52,7 | | 0,0510762 | 0,0385710 | 0,90 | 7,04E+08 | 7,43E+08 |
| 23,2 | | 0,0511395 | 0,0386147 | 0,91 | 8,84E+07 | 8,10E+07 |

| | | | | | | |
|---|---|---|---|---|---|---|
| 51,5 | | 0,0512696 | 0,0386562 | 1,26 | 1,99E+08 | 2,55E+08 |
| 9,4 | | 0,0513491 | 0,0388221 | 1,33 | 2,25E+07 | 3,00E+07 |
| 48,9 | | 0,0513738 | 0,0388116 | 0,91 | 8,33E+08 | 7,75E+08 |
| 39,7 | | 0,0514305 | 0,0388530 | 1,22 | 6,55E+07 | 7,68E+07 |
| 20,6 | | 0,0514344 | 0,0388425 | 0,09 | 4,36E+07 | 5,14E+07 |
| 16,2 | | 0,0515320 | 0,0388568 | 2,28 | 2,12E+06 | 4,30E+06 |
| 11,9 | | 0,0515834 | 0,0388723 | 0,71 | 5,38E+07 | 5,10E+07 |
| 45 | | 0,0515851 | 0,0388618 | 1,22 | 5,29E+08 | 6,03E+08 |
| 29,6 | | 0,0516057 | 0,0388772 | 1,36 | 2,36E+08 | 3,17E+08 |
| 44,6 | | 0,0516137 | 0,0388667 | 0,83 | 1,70E+08 | 1,77E+08 |
| 4,2 | | 0,0516477 | 0,0389112 | 1,58 | 3,01E+06 | 1,97E+06 |
| 27,3 | | 0,0517755 | 0,0390992 | 0,88 | 1,21E+08 | 1,22E+08 |
| 71,3 | | 0,0517878 | 0,0390887 | 1,14 | 1,75E+10 | 1,61E+10 |
| 22,4 | | 0,0518039 | 0,0390782 | 4,21 | 1,90E+06 | 7,98E+06 |
| 41,8 | | 0,0519462 | 0,0391226 | 1,26 | 3,63E+08 | 3,98E+08 |
| 74,6 | | 0,0519838 | 0,0391121 | 0,79 | 5,48E+08 | 6,80E+08 |
| 28,4 | | 0,0521204 | 0,0392491 | 0,07 | 2,14E+08 | 1,67E+08 |
| 7,7 | | 0,0522376 | 0,0393904 | 1,66 | 4,69E+06 | 2,98E+06 |
| 14,6 | | 0,0523946 | 0,0395133 | 1,73 | 2,10E+06 | 1,49E+06 |
| 13,3 | | 0,0524730 | 0,0395027 | 0,83 | 1,67E+07 | 1,39E+07 |
| 66,9 | | 0,0526254 | 0,0397557 | 0,64 | 5,80E+08 | 3,91E+08 |
| 16,4 | | 0,0526469 | 0,0397708 | 1,87 | 2,88E+06 | 3,02E+06 |
| 15,3 | | 0,0526874 | 0,0398204 | 0,71 | 1,62E+07 | 2,45E+07 |
| 21,1 | | 0,0527203 | 0,0398355 | 1,39 | 9,50E+06 | 8,41E+06 |
| 59,2 | | 0,0527967 | 0,0399205 | 0,76 | 6,54E+07 | 7,86E+07 |
| 39,8 | | 0,0528850 | 0,0399355 | 0,72 | 7,23E+08 | 8,61E+08 |
| 41,5 | | 0,0529242 | 0,0399538 | 1,23 | 3,31E+08 | 2,69E+08 |
| 28,7 | | 0,0530571 | 0,0400193 | 1,21 | 7,25E+07 | 7,06E+07 |
| 10,7 | | 0,0531300 | 0,0400344 | 0,60 | 5,26E+07 | 5,60E+07 |
| 32,1 | | 0,0531520 | 0,0400236 | 0,67 | 5,44E+07 | 5,48E+07 |
| 18,1 | | 0,0532437 | 0,0400676 | 1,42 | 7,03E+06 | 9,66E+06 |
| 6,8 | | 0,0534463 | 0,0401910 | 0,75 | 6,71E+06 | 7,92E+06 |
| 3,8 | | 0,0534994 | 0,0402467 | 0,44 | 3,81E+06 | 9,56E+06 |
| 5,9 | | 0,0535420 | 0,0402617 | 0,64 | 4,61E+06 | 5,15E+06 |
| 4 | | 0,0536107 | 0,0402798 | 3,42 | 1,20E+07 | 3,55E+06 |
| 26,7 | | 0,0536548 | 0,0402690 | 0,75 | 4,13E+08 | 5,46E+08 |
| 13,2 | | 0,0536888 | 0,0402925 | 1,55 | 1,00E+07 | 1,50E+07 |
| 28,3 | | 0,0537049 | 0,0403149 | 0,84 | 3,93E+07 | 3,24E+07 |
| 27,9 | | 0,0538264 | 0,0403545 | 0,89 | 1,01E+08 | 8,99E+07 |
| 29,9 | | 0,0539461 | 0,0403940 | 0,75 | 1,18E+08 | 8,83E+07 |
| 15,3 | | 0,0541091 | 0,0404999 | 1,95 | 8,36E+06 | 1,02E+07 |
| 26 | | 0,0541277 | 0,0404890 | 0,61 | 1,25E+07 | 1,43E+07 |
| 22,1 | | 0,0541564 | 0,0405028 | 1,42 | 2,17E+07 | 1,48E+07 |
| 50,5 | | 0,0542883 | 0,0406449 | 1,21 | 5,73E+07 | 6,86E+07 |
| 16,8 | | 0,0542938 | 0,0406341 | 0,68 | 7,46E+07 | 5,92E+07 |
| 9,3 | | 0,0543609 | 0,0406895 | 1,28 | 9,40E+06 | 1,02E+07 |
| 49,1 | | 0,0543961 | 0,0408026 | 1,13 | 2,81E+09 | 3,05E+09 |
| 35 | | 0,0544460 | 0,0408248 | 1,35 | 3,97E+07 | 5,21E+07 |
| 37,3 | | 0,0544551 | 0,0408395 | 1,22 | 6,06E+07 | 5,21E+07 |
| 5,1 | | 0,0544607 | 0,0408286 | 0,79 | 3,55E+07 | 3,55E+07 |

| | | | | | | |
|---|---|---|---|---|---|---|
| 21 | | 0,0545814 | 0,0409384 | 0,73 | 2,84E+07 | 2,54E+07 |
| 44,6 | | 0,0545967 | 0,0409605 | 0,80 | 3,08E+09 | 2,40E+09 |
| 38,2 | | 0,0546163 | 0,0409827 | 1,14 | 1,23E+08 | 1,22E+08 |
| 30,9 | | 0,0547590 | 0,0410048 | 1,15 | 1,23E+08 | 1,03E+08 |
| 46,7 | | 0,0549023 | 0,0411922 | 1,21 | 3,18E+08 | 3,50E+08 |
| 56,2 | | 0,0549562 | 0,0412324 | 1,14 | 8,73E+08 | 7,61E+08 |
| 43,5 | | 0,0549939 | 0,0412886 | 0,79 | 2,11E+08 | 1,67E+08 |
| 22,6 | | 0,0550694 | 0,0413031 | 1,12 | 1,97E+08 | 2,20E+08 |
| 27,4 | | 0,0551303 | 0,0414413 | 3,10 | 1,01E+07 | 3,88E+07 |
| 10,5 | | 0,0551588 | 0,0415218 | 0,55 | 7,08E+06 | 1,10E+07 |
| 48,4 | | 0,0552063 | 0,0415363 | 0,81 | 2,76E+09 | 2,50E+09 |
| 2,2 | | 0,0552717 | 0,0416338 | 2,12 | 4,63E+06 | 1,99E+06 |
| 50,4 | | 0,0553013 | 0,0416802 | 0,67 | 1,70E+08 | 1,41E+08 |
| 42 | | 0,0554659 | 0,0417617 | 0,80 | 3,93E+08 | 4,96E+08 |
| 72,3 | | 0,0556042 | 0,0418293 | 1,12 | 9,60E+08 | 9,72E+08 |
| 17,5 | | 0,0556905 | 0,0418756 | 1,26 | 1,72E+07 | 1,85E+07 |
| 43,9 | | 0,0557094 | 0,0418900 | 1,15 | 4,06E+08 | 4,58E+08 |
| 29,9 | | 0,0557773 | 0,0419118 | 0,70 | 1,53E+08 | 1,82E+08 |
| 6,6 | | 0,0558156 | 0,0419623 | 0,28 | 7,96E+06 | 3,18E+07 |
| 18,4 | | 0,0558241 | 0,0419511 | 0,64 | 1,53E+07 | 2,21E+07 |
| 10,6 | | 0,0558380 | 0,0419400 | 1,63 | 3,10E+07 | 1,76E+07 |
| 19,2 | | 0,0558853 | 0,0419947 | 3,78 | 3,27E+06 | 9,02E+06 |
| 33 | | 0,0559062 | 0,0419836 | 0,83 | 4,35E+08 | 4,71E+08 |
| 48,9 | | 0,0559196 | 0,0419724 | 0,88 | 6,30E+09 | 5,93E+09 |
| 2,7 | + | 0,0559643 | 0,0420154 | 2,22 | 1,11E+07 | 5,36E+06 |
| 39,8 | | 0,0559818 | 0,0420042 | 1,22 | 2,95E+08 | 3,44E+08 |
| 38,7 | | 0,0560669 | 0,0422104 | 1,14 | 1,20E+08 | 1,19E+08 |
| 16,4 | | 0,0561317 | 0,0422862 | 0,80 | 3,42E+07 | 3,39E+07 |
| 16,8 | | 0,0561844 | 0,0422750 | 0,24 | 1,98E+07 | 1,92E+07 |
| 33,5 | | 0,0562221 | 0,0422966 | 0,76 | 1,05E+08 | 1,04E+08 |
| 32 | | 0,0563258 | 0,0424008 | 0,89 | 2,11E+08 | 2,45E+08 |
| 59,2 | | 0,0563446 | 0,0423896 | 0,87 | 3,89E+09 | 3,28E+09 |
| 49,3 | | 0,0564639 | 0,0425509 | 0,88 | 9,91E+08 | 1,13E+09 |
| 38,6 | | 0,0565217 | 0,0425397 | 0,82 | 3,00E+08 | 3,24E+08 |
| 18,8 | | 0,0565709 | 0,0425284 | 1,30 | 2,33E+07 | 1,86E+07 |
| 43,8 | | 0,0566905 | 0,0426282 | 1,10 | 2,10E+08 | 2,24E+08 |
| 42,3 | | 0,0567449 | 0,0427322 | 1,26 | 7,69E+07 | 8,86E+07 |
| 21,2 | | 0,0567845 | 0,0427717 | 1,15 | 9,01E+07 | 9,29E+07 |
| 30,1 | | 0,0568316 | 0,0427889 | 1,29 | 1,29E+08 | 1,43E+08 |
| 30,1 | | 0,0568537 | 0,0427776 | 0,95 | 4,20E+08 | 3,98E+08 |
| 35,7 | | 0,0569630 | 0,0428445 | 1,94 | 8,40E+06 | 1,99E+07 |
| 30 | | 0,0570086 | 0,0429419 | 1,29 | 1,79E+08 | 1,79E+08 |
| 94,7 | | 0,0570985 | 0,0429844 | 1,56 | 1,63E+09 | 1,02E+09 |
| 10,8 | | 0,0571189 | 0,0430069 | 2,21 | 8,11E+06 | 3,25E+06 |
| 12,8 | | 0,0571320 | 0,0430282 | 0,70 | 3,48E+07 | 3,28E+07 |
| 46,6 | | 0,0571646 | 0,0430169 | 1,21 | 1,23E+08 | 1,45E+08 |
| 63,6 | | 0,0571743 | 0,0430055 | 0,25 | 1,63E+09 | 1,33E+09 |
| 38,6 | | 0,0571815 | 0,0429942 | 1,18 | 1,04E+08 | 1,26E+08 |
| 15,9 | | 0,0572763 | 0,0430440 | 0,71 | 3,39E+07 | 4,00E+07 |
| 41,4 | | 0,0572920 | 0,0430327 | 0,89 | 4,52E+08 | 5,07E+08 |

| | | | | | | |
|---|---|---|---|---|---|---|
| 19,8 | | 0,0573596 | 0,0430866 | 0,69 | 2,95E+09 | 2,01E+09 |
| 18,6 | | 0,0573761 | 0,0430753 | 1,25 | 4,73E+07 | 4,96E+07 |
| 55 | | 0,0574065 | 0,0431219 | 0,87 | 4,43E+08 | 4,87E+08 |
| 33 | | 0,0575872 | 0,0432747 | 1,42 | 1,44E+07 | 2,22E+07 |
| 14,6 | | 0,0576532 | 0,0432634 | 0,16 | 1,64E+07 | 1,19E+07 |
| 25,6 | | 0,0578825 | 0,0434740 | 0,68 | 6,47E+07 | 4,31E+07 |
| 9,5 | | 0,0579171 | 0,0434625 | 0,25 | 2,54E+07 | 8,81E+06 |
| 8,5 | | 0,0579364 | 0,0434511 | 0,77 | 1,26E+07 | 1,11E+07 |
| 45,1 | | 0,0580601 | 0,0435595 | 0,69 | 4,09E+07 | 6,01E+07 |
| 11,9 | | 0,0581734 | 0,0435985 | 0,73 | 3,95E+07 | 5,63E+07 |
| 7,9 | | 0,0581871 | 0,0435871 | 0,31 | 1,36E+07 | 1,52E+07 |
| 45,1 | | 0,0582906 | 0,0436828 | 0,80 | 2,70E+08 | 2,69E+08 |
| 20,7 | | 0,0583872 | 0,0437039 | 0,73 | 1,28E+08 | 9,05E+07 |
| 13,2 | | 0,0584342 | 0,0436924 | 0,70 | 2,72E+07 | 2,88E+07 |
| 18,1 | | 0,0585066 | 0,0437712 | 0,65 | 3,23E+07 | 2,97E+07 |
| 41 | | 0,0585761 | 0,0437838 | 1,14 | 3,76E+08 | 3,55E+08 |
| 48,5 | | 0,0586470 | 0,0438836 | 1,14 | 4,12E+08 | 4,67E+08 |
| 38,7 | | 0,0587808 | 0,0439046 | 1,41 | 5,26E+07 | 5,64E+07 |
| 22,7 | | 0,0587872 | 0,0438931 | 0,88 | 1,19E+08 | 1,25E+08 |
| 28,8 | | 0,0588463 | 0,0439465 | 0,77 | 2,67E+09 | 2,28E+09 |
| 32,2 | | 0,0588905 | 0,0439602 | 0,84 | 5,70E+08 | 6,29E+08 |
| 2,6 | | 0,0589618 | 0,0440388 | 0,37 | 2,94E+06 | 6,21E+06 |
| 22,1 | | 0,0589699 | 0,0440272 | 0,72 | 3,19E+07 | 4,29E+07 |
| 33,4 | | 0,0590059 | 0,0440764 | 1,12 | 9,75E+07 | 1,01E+08 |
| 28,2 | | 0,0590828 | 0,0440974 | 1,46 | 2,07E+07 | 2,51E+07 |
| 5,4 | | 0,0591037 | 0,0440858 | 0,42 | 1,65E+07 | 9,91E+06 |
| 10,3 | | 0,0593493 | 0,0442072 | 0,58 | 3,19E+07 | 4,47E+07 |
| 27,3 | | 0,0594674 | 0,0443954 | 4,97 | 2,10E+07 | 1,36E+07 |
| 14,4 | | 0,0594947 | 0,0443838 | 2,23 | 4,90E+06 | 5,72E+06 |
| 62,6 | | 0,0594961 | 0,0443722 | 1,08 | 3,09E+08 | 3,09E+08 |
| 28,5 | | 0,0595184 | 0,0443857 | 1,15 | 6,37E+07 | 5,97E+07 |
| 17 | | 0,0595366 | 0,0443741 | 0,07 | 4,19E+07 | 1,38E+08 |
| 22,1 | | 0,0595981 | 0,0443876 | 0,87 | 2,46E+08 | 2,47E+08 |
| 37,6 | | 0,0597177 | 0,0444324 | 0,85 | 1,04E+08 | 9,27E+07 |
| 8 | | 0,0597890 | 0,0444542 | 1,29 | 4,94E+06 | 3,82E+06 |
| 15,5 | | 0,0598009 | 0,0444426 | 0,65 | 2,42E+08 | 1,77E+08 |
| 26,1 | | 0,0598889 | 0,0445447 | 1,25 | 1,41E+08 | 1,77E+08 |
| 48 | | 0,0598973 | 0,0445331 | 0,48 | 4,64E+07 | 5,79E+07 |
| 8,5 | | 0,0599589 | 0,0445695 | 0,80 | 3,82E+07 | 4,18E+07 |
| 28,3 | | 0,0599928 | 0,0445579 | 1,21 | 4,20E+07 | 4,80E+07 |
| 8,4 | | 0,0600042 | 0,0445463 | 2,78 | 4,86E+06 | 1,02E+07 |
| 24,1 | | 0,0601192 | 0,0446076 | 0,87 | 8,70E+07 | 7,77E+07 |
| 76,1 | | 0,0601721 | 0,0446814 | 0,82 | 7,56E+08 | 8,95E+08 |
| 36,4 | | 0,0601756 | 0,0446698 | 1,21 | 3,80E+08 | 3,65E+08 |
| 6,8 | | 0,0601959 | 0,0446582 | 0,74 | 6,80E+06 | 6,68E+06 |
| 46,3 | | 0,0602094 | 0,0446465 | 0,75 | 2,33E+08 | 2,46E+08 |
| 52,6 | | 0,0602481 | 0,0446349 | 1,24 | 1,08E+08 | 1,20E+08 |
| 5 | | 0,0603176 | 0,0446805 | 1,48 | 1,91E+07 | 1,23E+07 |
| 7,4 | | 0,0603535 | 0,0446689 | 0,48 | 4,75E+06 | 5,03E+06 |
| 31,3 | | 0,0603759 | 0,0446573 | 1,37 | 1,32E+08 | 9,89E+07 |

| | | | | | | |
|---|---|---|---|---|---|---|
| 44,2 | | 0,0604003 | 0,0447112 | 0,88 | 1,86E+08 | 1,76E+08 |
| 23,1 | | 0,0605484 | 0,0448077 | 1,93 | 8,27E+06 | 1,48E+07 |
| 49,5 | | 0,0605748 | 0,0448979 | 0,88 | 1,07E+09 | 1,02E+09 |
| 18,5 | | 0,0607557 | 0,0450587 | 1,19 | 3,31E+07 | 2,84E+07 |
| 5,1 | | 0,0607878 | 0,0450719 | 2,48 | 9,85E+06 | 6,03E+06 |
| 25,4 | | 0,0608788 | 0,0450602 | 0,79 | 1,20E+08 | 1,26E+08 |
| 58 | | 0,0610960 | 0,0450818 | 0,86 | 4,63E+09 | 5,33E+09 |
| 45,1 | | 0,0611995 | 0,0451562 | 0,72 | 2,07E+08 | 2,49E+08 |
| 44,7 | | 0,0613152 | 0,0452337 | 0,07 | 1,64E+08 | 6,71E+08 |
| 18,1 | | 0,0613535 | 0,0452220 | 0,79 | 7,39E+07 | 9,62E+07 |
| 20,3 | | 0,0613626 | 0,0452103 | 1,28 | 8,79E+07 | 6,57E+07 |
| 24,6 | | 0,0615263 | 0,0453458 | 0,14 | 2,29E+07 | 3,35E+07 |
| 24,4 | | 0,0615835 | 0,0453340 | 1,18 | 3,56E+08 | 2,95E+08 |
| 14,3 | | 0,0617133 | 0,0453751 | 1,47 | 1,20E+07 | 1,22E+07 |
| 59,9 | | 0,0618048 | 0,0454514 | 0,87 | 8,56E+08 | 8,37E+08 |
| 14,1 | | 0,0618842 | 0,0455173 | 0,22 | 4,30E+07 | 1,40E+07 |
| 17,4 | | 0,0618850 | 0,0455056 | 1,84 | 2,08E+07 | 2,33E+07 |
| 34,7 | | 0,0618893 | 0,0454938 | 0,89 | 3,74E+08 | 3,59E+08 |
| 5,4 | | 0,0618897 | 0,0454820 | 0,58 | 2,55E+06 | 2,88E+06 |
| 15,8 | | 0,0619184 | 0,0454703 | 0,72 | 3,25E+07 | 3,89E+07 |
| 22,1 | | 0,0619506 | 0,0454833 | 0,47 | 5,51E+06 | 4,14E+06 |
| 53,5 | | 0,0620162 | 0,0454716 | 0,80 | 7,67E+08 | 5,89E+08 |
| 21,9 | | 0,0620378 | 0,0454598 | 0,82 | 7,07E+07 | 6,09E+07 |
| 26,5 | | 0,0621111 | 0,0455359 | 1,36 | 9,76E+07 | 1,19E+08 |
| 8,1 | | 0,0621245 | 0,0455521 | 1,81 | 4,65E+06 | 2,73E+06 |
| 10,4 | | 0,0621927 | 0,0455899 | 0,70 | 6,55E+06 | 7,89E+06 |
| 17 | | 0,0626365 | 0,0457155 | 0,36 | 9,03E+06 | 5,28E+06 |
| 18,5 | | 0,0627312 | 0,0457842 | 0,75 | 1,01E+07 | 1,00E+07 |
| 20,5 | | 0,0628007 | 0,0457724 | 0,76 | 3,81E+07 | 4,65E+07 |
| 9,6 | | 0,0628467 | 0,0457884 | 1,55 | 7,97E+06 | 6,50E+06 |
| 60 | | 0,0628757 | 0,0458014 | 1,34 | 1,02E+08 | 9,80E+07 |
| 39,5 | | 0,0629080 | 0,0458216 | 0,92 | 2,96E+07 | 3,09E+07 |
| 5 | | 0,0630831 | 0,0459417 | 1,84 | 3,30E+06 | 4,67E+06 |
| 3,2 | | 0,0632018 | 0,0460670 | 1,59 | 3,04E+07 | 3,29E+07 |
| 12,6 | | 0,0632382 | 0,0460830 | 1,64 | 2,55E+07 | 1,56E+07 |
| 60,1 | | 0,0632433 | 0,0460711 | 1,17 | 1,54E+09 | 1,35E+09 |
| 15,1 | | 0,0632868 | 0,0460912 | 1,39 | 9,95E+06 | 1,52E+07 |
| 19,4 | | 0,0633315 | 0,0461112 | 0,84 | 4,32E+07 | 3,55E+07 |
| 6 | | 0,0634204 | 0,0460994 | 0,71 | 2,14E+07 | 2,54E+07 |
| 37,9 | | 0,0634621 | 0,0461472 | 0,80 | 5,67E+07 | 4,83E+07 |
| 5,7 | | 0,0634936 | 0,0461847 | 0,73 | 1,94E+08 | 1,51E+08 |
| 79,4 | | 0,0636261 | 0,0463117 | 1,30 | 6,33E+07 | 5,85E+07 |
| 11,1 | | 0,0637100 | 0,0464243 | 0,75 | 2,31E+07 | 2,16E+07 |
| 65,1 | | 0,0637740 | 0,0464370 | 0,76 | 5,14E+08 | 6,60E+08 |
| 25,1 | | 0,0637904 | 0,0464570 | 1,58 | 7,80E+07 | 1,16E+08 |
| 8,3 | | 0,0637985 | 0,0464450 | 4,87 | 4,27E+06 | 9,76E+05 |
| 43,3 | | 0,0638282 | 0,0464567 | 1,23 | 1,97E+08 | 2,27E+08 |
| 13,7 | | 0,0638490 | 0,0464448 | 2,24 | 7,68E+06 | 4,48E+06 |
| 3,3 | | 0,0638676 | 0,0464647 | 1,93 | 2,78E+06 | 2,45E+06 |
| 31,6 | | 0,0640325 | 0,0464774 | 0,71 | 5,98E+07 | 4,89E+07 |

| | | | | | | |
|---|---|---|---|---|---|---|
| 18,8 | | 0,0640859 | 0,0464932 | 0,47 | 8,11E+06 | 6,03E+06 |
| 4,9 | | 0,0641148 | 0,0464813 | 0,59 | 2,91E+06 | 3,05E+06 |
| 22,2 | | 0,0642983 | 0,0466540 | 0,83 | 5,99E+07 | 5,64E+07 |
| 39,5 | | 0,0643100 | 0,0466421 | 1,18 | 3,82E+08 | 4,52E+08 |
| 50,5 | | 0,0643935 | 0,0467193 | 1,21 | 9,42E+08 | 1,20E+09 |
| 24,3 | | 0,0644353 | 0,0467073 | 1,25 | 5,55E+07 | 4,75E+07 |
| 26,5 | | 0,0645590 | 0,0468880 | 1,35 | 1,08E+07 | 1,51E+07 |
| 9,7 | | 0,0646064 | 0,0468760 | 0,47 | 7,33E+07 | 9,10E+07 |
| 13,2 | | 0,0646661 | 0,0469275 | 0,78 | 8,25E+06 | 8,21E+06 |
| 31,9 | | 0,0647744 | 0,0470077 | 1,24 | 7,67E+07 | 9,16E+07 |
| 13,4 | | 0,0647831 | 0,0469956 | 2,09 | 3,40E+06 | 4,23E+06 |
| 4,1 | | 0,0648650 | 0,0469836 | 0,50 | 1,40E+07 | 1,41E+07 |
| 32 | | 0,0648691 | 0,0469716 | 1,16 | 1,76E+08 | 1,92E+08 |
| 34,2 | | 0,0649613 | 0,0469872 | 0,88 | 7,15E+08 | 7,79E+08 |
| 19,4 | | 0,0650653 | 0,0470233 | 1,18 | 4,86E+07 | 4,64E+07 |
| 8,3 | | 0,0651165 | 0,0470675 | 3,61 | 3,94E+06 | 8,56E+06 |
| 41,9 | | 0,0652659 | 0,0472691 | 0,69 | 6,37E+08 | 8,59E+08 |
| 11,5 | | 0,0653184 | 0,0473756 | 2,75 | 4,06E+06 | 7,24E+06 |
| 64,8 | | 0,0653247 | 0,0473635 | 0,74 | 1,91E+09 | 2,23E+09 |
| 16,3 | | 0,0653701 | 0,0473994 | 1,21 | 1,19E+07 | 1,30E+07 |
| 45,9 | | 0,0654440 | 0,0474506 | 0,71 | 4,11E+07 | 3,81E+07 |
| 38,7 | | 0,0654978 | 0,0474630 | 1,15 | 4,13E+08 | 4,45E+08 |
| 4,3 | | 0,0655069 | 0,0474835 | 0,85 | 6,85E+06 | 7,80E+06 |
| 13,3 | | 0,0656224 | 0,0476357 | 0,70 | 4,28E+07 | 3,73E+07 |
| 36,9 | | 0,0656758 | 0,0476705 | 0,85 | 9,08E+07 | 1,08E+08 |
| 10,1 | | 0,0658182 | 0,0477144 | 0,37 | 7,36E+06 | 4,31E+06 |
| 51,5 | | 0,0659533 | 0,0477614 | 1,11 | 1,53E+08 | 1,50E+08 |
| 4,2 | | 0,0659803 | 0,0477808 | 0,49 | 7,24E+06 | 4,99E+06 |
| 20,1 | | 0,0660004 | 0,0477687 | 0,68 | 2,86E+07 | 1,73E+07 |
| 16 | | 0,0660582 | 0,0477840 | 0,61 | 2,67E+07 | 2,18E+07 |
| 5,2 | | 0,0660635 | 0,0477718 | 1,92 | 5,57E+06 | 1,08E+07 |
| 31,3 | | 0,0661268 | 0,0478157 | 0,84 | 1,54E+08 | 1,28E+08 |
| 53 | | 0,0661911 | 0,0478666 | 0,90 | 3,12E+08 | 3,33E+08 |
| 15,3 | | 0,0661975 | 0,0478545 | 1,29 | 3,80E+06 | 3,25E+06 |
| 50,2 | | 0,0662496 | 0,0479216 | 0,92 | 3,75E+08 | 3,81E+08 |
| 15,3 | | 0,0663748 | 0,0479664 | 1,42 | 2,45E+07 | 1,88E+07 |
| 2,8 | | 0,0665878 | 0,0481759 | 1,90 | 5,67E+06 | 2,93E+06 |
| 70,7 | | 0,0667053 | 0,0481952 | 0,90 | 2,06E+09 | 2,21E+09 |
| 33,3 | | 0,0667287 | 0,0482460 | 0,41 | 1,15E+07 | 1,71E+07 |
| 8,2 | | 0,0669641 | 0,0483374 | 1,64 | 8,21E+06 | 4,63E+06 |
| 40,9 | | 0,0670689 | 0,0483251 | 1,41 | 4,83E+07 | 3,71E+07 |
| 58,4 | | 0,0670991 | 0,0483128 | 1,31 | 1,02E+08 | 8,84E+07 |
| 70,6 | | 0,0671212 | 0,0483006 | 1,27 | 3,28E+09 | 2,94E+09 |
| 48,3 | | 0,0672157 | 0,0483513 | 0,94 | 2,80E+09 | 2,72E+09 |
| 18 | | 0,0673221 | 0,0485572 | 1,15 | 1,98E+07 | 2,22E+07 |
| 52,5 | | 0,0675357 | 0,0487935 | 1,10 | 5,42E+08 | 5,69E+08 |
| 32,7 | | 0,0675697 | 0,0487811 | 0,90 | 7,35E+08 | 6,49E+08 |
| 48 | | 0,0676016 | 0,0487688 | 1,35 | 1,16E+08 | 8,39E+07 |
| 26,5 | | 0,0677331 | 0,0487807 | 0,89 | 2,55E+08 | 2,78E+08 |
| 38,8 | | 0,0678387 | 0,0488748 | 0,90 | 3,18E+08 | 2,96E+08 |

| | | | | | | |
|---|---|---|---|---|---|---|
| 6,1 | | 0,0678477 | 0,0488857 | 1,37 | 7,67E+06 | 5,95E+06 |
| 31,1 | | 0,0678666 | 0,0488734 | 0,77 | 2,16E+08 | 2,28E+08 |
| 90,4 | | 0,0678728 | 0,0488610 | 1,33 | 4,12E+09 | 4,78E+09 |
| 28 | | 0,0678980 | 0,0488800 | 0,84 | 1,16E+08 | 1,32E+08 |
| 32,2 | | 0,0679169 | 0,0488919 | 1,52 | 5,98E+07 | 3,52E+07 |
| 20,2 | | 0,0679707 | 0,0489615 | 0,55 | 3,35E+07 | 2,16E+07 |
| 21,2 | | 0,0680073 | 0,0489734 | 0,32 | 5,96E+06 | 2,24E+07 |
| 46,2 | | 0,0680868 | 0,0491340 | 0,82 | 3,26E+08 | 3,45E+08 |
| 54,3 | | 0,0680924 | 0,0491216 | 0,68 | 4,01E+08 | 3,72E+08 |
| 19,8 | | 0,0681295 | 0,0491092 | 0,75 | 4,99E+07 | 6,05E+07 |
| 30,6 | | 0,0682113 | 0,0491847 | 0,88 | 5,29E+08 | 4,59E+08 |
| 31,3 | | 0,0682311 | 0,0491723 | 1,57 | 7,81E+07 | 1,11E+08 |
| 17,9 | | 0,0683782 | 0,0492144 | 0,83 | 8,16E+07 | 7,19E+07 |
| 18 | | 0,0684623 | 0,0492566 | 0,61 | 6,74E+06 | 9,97E+06 |
| 36,6 | | 0,0684884 | 0,0492754 | 1,78 | 2,77E+06 | 3,55E+06 |
| 31 | | 0,0685253 | 0,0493216 | 1,28 | 1,09E+08 | 1,39E+08 |
| 28,3 | | 0,0688212 | 0,0496432 | 0,85 | 2,12E+08 | 1,93E+08 |
| 47,9 | | 0,0688237 | 0,0496307 | 1,14 | 4,96E+08 | 5,31E+08 |
| 24,6 | | 0,0689462 | 0,0498068 | 1,25 | 5,12E+07 | 4,58E+07 |
| 53,1 | | 0,0690637 | 0,0498588 | 1,16 | 1,75E+09 | 1,88E+09 |
| 39,2 | | 0,0690875 | 0,0498694 | 1,19 | 2,21E+09 | 2,24E+09 |
| 5 | | 0,0691046 | 0,0498569 | 1,25 | 2,93E+06 | 2,51E+06 |
| 40 | | 0,0692427 | 0,0499380 | 1,14 | 5,99E+08 | 6,22E+08 |
| 45,4 | | 0,0692994 | 0,0499809 | 1,64 | 1,07E+07 | 1,90E+07 |
| 23,9 | | 0,0693924 | 0,0500408 | 0,92 | 9,04E+07 | 8,65E+07 |
| 3,7 | | 0,0694766 | 0,0500282 | 0,40 | 2,19E+07 | 3,25E+07 |
| 36,5 | | 0,0695979 | 0,0500659 | 0,84 | 1,85E+08 | 1,66E+08 |
| 54,5 | | 0,0697285 | 0,0501127 | 0,79 | 1,11E+09 | 1,08E+09 |
| 31,1 | | 0,0697844 | 0,0501001 | 1,15 | 3,80E+07 | 4,38E+07 |
| 32,6 | | 0,0698195 | 0,0500875 | 2,47 | 1,71E+07 | 1,82E+07 |
| 5,9 | | 0,0699114 | 0,0501021 | 1,38 | 5,89E+06 | 8,12E+06 |
| 4,5 | | 0,0700448 | 0,0501438 | 0,41 | 1,45E+07 | 8,15E+06 |
| 67,3 | | 0,0700957 | 0,0502367 | 1,15 | 7,94E+07 | 9,11E+07 |
| 61,8 | | 0,0701172 | 0,0502241 | 1,09 | 2,14E+08 | 2,09E+08 |
| 33,3 | | 0,0701235 | 0,0502115 | 0,84 | 4,92E+07 | 3,95E+07 |
| 3,9 | | 0,0702549 | 0,0503084 | 1,48 | 2,95E+06 | 2,17E+06 |
| 43,9 | | 0,0702807 | 0,0502958 | 1,10 | 3,34E+08 | 3,38E+08 |
| 7,9 | | 0,0703815 | 0,0504147 | 0,83 | 9,11E+07 | 8,55E+07 |
| 8,6 | | 0,0704324 | 0,0504833 | 1,50 | 1,40E+07 | 8,94E+06 |
| 64 | | 0,0704332 | 0,0504706 | 1,12 | 1,73E+09 | 1,79E+09 |
| 35,7 | | 0,0706021 | 0,0505683 | 0,70 | 9,38E+07 | 1,14E+08 |
| 6,5 | | 0,0707377 | 0,0506108 | 0,89 | 5,41E+07 | 5,26E+07 |
| 4,2 | | 0,0708475 | 0,0507315 | 5,00 | 1,62E+07 | 1,71E+06 |
| 49,5 | | 0,0708780 | 0,0507499 | 0,82 | 4,93E+08 | 5,80E+08 |
| 16,6 | | 0,0708810 | 0,0507372 | 0,53 | 5,95E+06 | 1,17E+07 |
| 58,3 | | 0,0709871 | 0,0507756 | 1,20 | 2,69E+08 | 2,32E+08 |
| 7,2 | | 0,0711180 | 0,0508500 | 0,76 | 6,03E+07 | 4,36E+07 |
| 10,6 | | 0,0712124 | 0,0508913 | 1,85 | 5,78E+06 | 3,32E+06 |
| 30,2 | | 0,0713411 | 0,0509637 | 0,76 | 3,57E+08 | 3,63E+08 |
| 13,1 | | 0,0714318 | 0,0511451 | 0,23 | 9,18E+06 | 1,03E+07 |

| | | | | | | |
|---|---|---|---|---|---|---|
| 31,2 | | 0,0715659 | 0,0511634 | 1,25 | 5,14E+07 | 3,75E+07 |
| 50,2 | | 0,0717257 | 0,0513517 | 0,83 | 1,28E+08 | 1,59E+08 |
| 82,3 | | 0,0717774 | 0,0513658 | 0,84 | 7,52E+08 | 9,00E+08 |
| 9,9 | | 0,0717969 | 0,0513530 | 0,35 | 6,68E+06 | 1,89E+06 |
| 47,2 | | 0,0719416 | 0,0514251 | 0,83 | 4,66E+08 | 4,88E+08 |
| 16,6 | | 0,0721099 | 0,0516152 | 0,74 | 1,56E+08 | 1,65E+08 |
| 12,7 | | 0,0721620 | 0,0516333 | 0,63 | 8,81E+07 | 1,16E+08 |
| 56,8 | | 0,0721637 | 0,0516204 | 0,93 | 4,38E+08 | 4,46E+08 |
| 5,2 | | 0,0722771 | 0,0517094 | 1,55 | 3,45E+06 | 2,60E+06 |
| 45,8 | | 0,0723020 | 0,0517234 | 0,84 | 5,59E+08 | 6,79E+08 |
| 1,6 | | 0,0724626 | 0,0519800 | 3,19 | 7,78E+06 | 1,73E+06 |
| 32,3 | | 0,0724994 | 0,0519940 | 0,71 | 1,23E+08 | 1,06E+08 |
| 53,2 | | 0,0725959 | 0,0520629 | 0,73 | 1,50E+09 | 1,90E+09 |
| 40,7 | | 0,0726300 | 0,0521357 | 0,83 | 2,11E+08 | 1,66E+08 |
| 19,4 | | 0,0726787 | 0,0521845 | 0,81 | 1,55E+08 | 1,83E+08 |
| 53,7 | | 0,0727238 | 0,0522612 | 0,89 | 3,71E+08 | 4,14E+08 |
| 18,1 | | 0,0727338 | 0,0522482 | 0,69 | 7,50E+07 | 6,69E+07 |
| 27,5 | | 0,0727549 | 0,0522621 | 0,80 | 1,54E+08 | 1,80E+08 |
| 34,8 | | 0,0728612 | 0,0522949 | 1,16 | 1,76E+08 | 1,83E+08 |
| 7,3 | | 0,0728903 | 0,0523635 | 0,65 | 1,12E+09 | 9,89E+08 |
| 9,6 | | 0,0729023 | 0,0523505 | 2,08 | 1,22E+07 | 1,45E+07 |
| 15 | | 0,0730816 | 0,0525107 | 0,26 | 1,06E+07 | 6,21E+06 |
| 64,9 | | 0,0731309 | 0,0525992 | 1,46 | 1,77E+09 | 2,73E+09 |
| 5,6 | | 0,0731681 | 0,0525861 | 1,51 | 8,18E+06 | 7,34E+06 |
| 18,1 | | 0,0732742 | 0,0526347 | 1,23 | 3,15E+07 | 2,79E+07 |
| 31,2 | | 0,0732878 | 0,0526216 | 0,73 | 1,30E+08 | 1,03E+08 |
| 66,1 | | 0,0733343 | 0,0526085 | 1,42 | 7,72E+07 | 1,26E+08 |
| 14,9 | | 0,0736807 | 0,0527276 | 0,67 | 3,79E+07 | 6,07E+07 |
| 8,9 | | 0,0738133 | 0,0527930 | 1,52 | 1,96E+07 | 2,20E+07 |
| 55 | | 0,0738187 | 0,0527799 | 0,88 | 3,89E+08 | 3,56E+08 |
| 13,3 | | 0,0738571 | 0,0527668 | 1,48 | 7,50E+07 | 6,48E+07 |
| 5,3 | | 0,0739958 | 0,0529384 | 0,20 | 1,57E+07 | 2,01E+07 |
| 28,9 | | 0,0740067 | 0,0529491 | 0,83 | 2,91E+08 | 2,76E+08 |
| 61 | | 0,0741248 | 0,0530333 | 1,15 | 1,30E+09 | 1,17E+09 |
| 2,6 | | 0,0742873 | 0,0531858 | 2,02 | 3,71E+06 | 2,82E+06 |
| 5,5 | | 0,0743057 | 0,0531994 | 0,75 | 9,65E+06 | 8,82E+06 |
| 29,8 | | 0,0743068 | 0,0531862 | 1,14 | 4,38E+08 | 3,81E+08 |
| 29,5 | | 0,0744030 | 0,0531730 | 0,46 | 1,07E+07 | 1,75E+07 |
| 21,4 | | 0,0744793 | 0,0531836 | 0,36 | 3,01E+07 | 2,56E+07 |
| 3 | | 0,0745229 | 0,0532210 | 3,67 | 3,46E+06 | 2,21E+06 |
| 28,2 | | 0,0746455 | 0,0533753 | 0,78 | 2,19E+07 | 1,97E+07 |
| 23,8 | | 0,0746554 | 0,0533621 | 0,73 | 1,48E+08 | 1,29E+08 |
| 31,2 | | 0,0748080 | 0,0535469 | 1,11 | 6,30E+07 | 5,94E+07 |
| 51,5 | | 0,0751867 | 0,0539030 | 0,90 | 1,40E+09 | 1,39E+09 |
| 44,5 | | 0,0752064 | 0,0538896 | 1,21 | 5,52E+07 | 4,86E+07 |
| 11,1 | | 0,0752507 | 0,0539070 | 1,30 | 3,29E+07 | 4,23E+07 |
| 6 | | 0,0753147 | 0,0539510 | 0,54 | 7,25E+06 | 9,12E+06 |
| 12,8 | | 0,0753343 | 0,0539614 | 1,34 | 8,61E+06 | 6,60E+06 |
| 18,1 | | 0,0753547 | 0,0539787 | 1,21 | 9,43E+07 | 1,00E+08 |
| 26,2 | | 0,0757421 | 0,0543203 | 1,21 | 6,51E+07 | 8,42E+07 |

| | | | | | | |
|---|---|---|---|---|---|---|
| 31 | | 0,0757426 | 0,0543069 | 0,84 | 3,51E+08 | 4,02E+08 |
| 21,9 | | 0,0758687 | 0,0543913 | 2,32 | 2,53E+07 | 7,73E+06 |
| 3,9 | | 0,0760743 | 0,0545201 | 3,54 | 1,04E+07 | 1,72E+06 |
| 21,5 | | 0,0761403 | 0,0546380 | 1,42 | 7,79E+07 | 1,20E+08 |
| 44,7 | | 0,0763892 | 0,0548042 | 1,18 | 2,83E+08 | 2,46E+08 |
| 12,2 | | 0,0764109 | 0,0547907 | 0,82 | 1,32E+08 | 1,21E+08 |
| 18,8 | | 0,0765691 | 0,0549667 | 0,39 | 1,64E+07 | 1,53E+07 |
| 6,8 | | 0,0766549 | 0,0550064 | 0,51 | 5,45E+06 | 8,00E+06 |
| 15 | | 0,0768025 | 0,0552256 | 1,84 | 9,29E+06 | 4,54E+06 |
| 32,6 | | 0,0768374 | 0,0552357 | 1,33 | 2,50E+07 | 3,13E+07 |
| 30,8 | | 0,0768397 | 0,0552457 | 1,22 | 6,44E+07 | 6,36E+07 |
| 27,6 | | 0,0772898 | 0,0556747 | 0,82 | 1,87E+08 | 1,41E+08 |
| 39,2 | | 0,0773048 | 0,0556915 | 1,22 | 1,46E+08 | 1,84E+08 |
| 49 | | 0,0774039 | 0,0557557 | 1,14 | 2,22E+08 | 1,94E+08 |
| 67,3 | | 0,0774907 | 0,0558296 | 0,79 | 5,46E+08 | 7,07E+08 |
| 37,6 | | 0,0775588 | 0,0558464 | 0,89 | 1,72E+08 | 1,51E+08 |
| 72,8 | | 0,0775755 | 0,0559104 | 0,82 | 3,68E+09 | 4,33E+09 |
| 15,4 | | 0,0776216 | 0,0559705 | 1,59 | 7,03E+07 | 5,58E+07 |
| 44,2 | | 0,0778218 | 0,0561446 | 1,20 | 9,27E+07 | 1,13E+08 |
| 41,3 | | 0,0779389 | 0,0563424 | 0,76 | 1,91E+08 | 1,45E+08 |
| 5,6 | | 0,0779497 | 0,0563285 | 0,27 | 2,17E+07 | 2,76E+07 |
| 17,8 | | 0,0780194 | 0,0563687 | 0,23 | 1,54E+07 | 1,16E+07 |
| 9 | | 0,0780770 | 0,0564158 | 1,72 | 7,89E+06 | 1,12E+07 |
| 12,1 | | 0,0781046 | 0,0564560 | 0,68 | 1,65E+07 | 2,06E+07 |
| 14,2 | | 0,0781840 | 0,0564422 | 0,88 | 1,06E+08 | 1,16E+08 |
| 4,2 | | 0,0782191 | 0,0564283 | 1,98 | 3,65E+06 | 5,84E+06 |
| 51,6 | | 0,0782770 | 0,0565244 | 1,18 | 1,36E+08 | 1,67E+08 |
| 45,9 | | 0,0784819 | 0,0567698 | 1,17 | 3,61E+08 | 3,48E+08 |
| 35,7 | | 0,0788471 | 0,0570326 | 1,18 | 5,42E+08 | 4,39E+08 |
| 34,1 | | 0,0790009 | 0,0573062 | 1,28 | 4,86E+07 | 3,62E+07 |
| 26,8 | | 0,0790066 | 0,0572921 | 1,13 | 6,55E+07 | 5,83E+07 |
| 25,6 | | 0,0790138 | 0,0572781 | 0,31 | 8,90E+06 | 2,92E+07 |
| 18,6 | | 0,0791326 | 0,0574072 | 0,83 | 5,03E+07 | 4,12E+07 |
| 68,2 | | 0,0791346 | 0,0573931 | 1,31 | 7,73E+08 | 1,09E+09 |
| 16,6 | | 0,0792039 | 0,0574291 | 0,33 | 1,40E+07 | 2,71E+07 |
| 17,1 | | 0,0792647 | 0,0575267 | 1,18 | 7,31E+07 | 5,92E+07 |
| 20,2 | | 0,0793745 | 0,0575890 | 1,76 | 6,18E+06 | 4,79E+06 |
| 38,6 | | 0,0794341 | 0,0575984 | 1,27 | 3,05E+07 | 2,43E+07 |
| 10,1 | | 0,0797654 | 0,0580896 | 1,26 | 8,45E+06 | 9,69E+06 |
| 24,8 | | 0,0798096 | 0,0580989 | 0,84 | 1,45E+08 | 1,59E+08 |
| 44,8 | | 0,0798149 | 0,0580847 | 0,92 | 5,19E+08 | 5,04E+08 |
| 56,3 | | 0,0799592 | 0,0581243 | 0,88 | 2,70E+08 | 3,04E+08 |
| 9,6 | | 0,0799751 | 0,0581404 | 1,25 | 1,21E+07 | 1,17E+07 |
| 21,1 | | 0,0800027 | 0,0581496 | 1,27 | 9,27E+07 | 8,18E+07 |
| 21 | | 0,0800440 | 0,0582224 | 0,84 | 1,22E+08 | 1,27E+08 |
| 70 | | 0,0800683 | 0,0582385 | 1,21 | 7,62E+08 | 5,87E+08 |
| 14,7 | | 0,0800977 | 0,0582849 | 1,13 | 7,77E+06 | 8,25E+06 |
| 11,9 | | 0,0801059 | 0,0582706 | 0,48 | 9,40E+06 | 1,50E+07 |
| 46,1 | | 0,0801467 | 0,0582828 | 0,92 | 3,56E+08 | 3,29E+08 |
| 18,7 | | 0,0802214 | 0,0583721 | 0,82 | 1,85E+07 | 1,89E+07 |

| | | | | | | |
|---|---|---|---|---|---|---|
| 23,1 | | 0,0802706 | 0,0583881 | 0,68 | 8,47E+07 | 8,64E+07 |
| 33,6 | | 0,0802959 | 0,0583973 | 0,84 | 7,72E+07 | 6,61E+07 |
| 23,2 | | 0,0803409 | 0,0583830 | 1,13 | 8,26E+07 | 9,20E+07 |
| 13,2 | | 0,0803866 | 0,0583990 | 1,14 | 9,70E+07 | 9,65E+07 |
| 8 | | 0,0804315 | 0,0584394 | 2,62 | 1,45E+06 | 4,88E+06 |
| 6,4 | | 0,0805062 | 0,0584476 | 0,65 | 1,41E+07 | 8,63E+06 |
| 9,1 | | 0,0805426 | 0,0584333 | 0,83 | 1,33E+08 | 1,67E+08 |
| 4,2 | | 0,0806185 | 0,0584659 | 0,53 | 1,10E+07 | 8,16E+06 |
| 13,1 | | 0,0808410 | 0,0586875 | 0,92 | 8,30E+06 | 7,76E+06 |
| 9,1 | | 0,0808749 | 0,0586732 | 1,36 | 1,28E+07 | 1,26E+07 |
| 57,3 | | 0,0810009 | 0,0587368 | 0,80 | 1,07E+08 | 1,26E+08 |
| 22,5 | | 0,0811965 | 0,0588364 | 1,48 | 2,34E+07 | 3,80E+07 |
| 46,1 | | 0,0812628 | 0,0588445 | 0,87 | 5,84E+08 | 5,14E+08 |
| 44,5 | | 0,0814371 | 0,0590326 | 0,73 | 5,44E+07 | 5,38E+07 |
| 4,8 | | 0,0817274 | 0,0592595 | 1,46 | 3,81E+06 | 4,40E+06 |
| 20,6 | | 0,0818557 | 0,0593521 | 0,74 | 1,34E+08 | 9,71E+07 |
| 24,6 | | 0,0819801 | 0,0593688 | 1,13 | 3,18E+08 | 3,40E+08 |
| 65,9 | | 0,0820059 | 0,0593544 | 1,14 | 1,44E+09 | 1,70E+09 |
| 45,4 | | 0,0820180 | 0,0593400 | 1,10 | 3,39E+08 | 3,65E+08 |
| 40,6 | | 0,0820469 | 0,0593489 | 0,85 | 1,42E+09 | 1,19E+09 |
| 28,6 | | 0,0820780 | 0,0593345 | 0,79 | 2,64E+07 | 3,45E+07 |
| 4,6 | | 0,0821936 | 0,0595056 | 2,29 | 2,86E+06 | 1,43E+06 |
| 68,2 | | 0,0823650 | 0,0595941 | 1,09 | 1,99E+09 | 2,16E+09 |
| 49,7 | | 0,0824923 | 0,0598553 | 0,81 | 1,64E+08 | 2,05E+08 |
| 13,4 | | 0,0826378 | 0,0599544 | 0,82 | 2,68E+07 | 2,14E+07 |
| 20,2 | + | 0,0826690 | 0,0599631 | 0,85 | 1,21E+08 | 1,35E+08 |
| 19,3 | | 0,0827194 | 0,0600252 | 1,30 | 1,25E+08 | 9,86E+07 |
| 16,8 | | 0,0827813 | 0,0600369 | 1,10 | 1,60E+08 | 1,57E+08 |
| 30,1 | | 0,0828550 | 0,0601707 | 0,82 | 6,15E+07 | 6,89E+07 |
| 18,9 | | 0,0828697 | 0,0601561 | 2,79 | 3,29E+06 | 8,74E+06 |
| 59,6 | | 0,0828948 | 0,0601415 | 1,05 | 1,66E+09 | 1,72E+09 |
| 6,7 | | 0,0829920 | 0,0603130 | 1,36 | 1,87E+07 | 2,12E+07 |
| 31,8 | | 0,0830203 | 0,0603207 | 0,89 | 4,08E+08 | 4,01E+08 |
| 3,7 | | 0,0831143 | 0,0604484 | 1,63 | 2,01E+06 | 1,54E+06 |
| 39,3 | | 0,0831427 | 0,0604570 | 1,21 | 5,80E+08 | 5,77E+08 |
| 30,2 | | 0,0831722 | 0,0604424 | 1,10 | 1,62E+08 | 1,52E+08 |
| 47,1 | | 0,0833262 | 0,0606465 | 1,40 | 1,89E+07 | 2,32E+07 |
| 35,8 | | 0,0833692 | 0,0606618 | 1,29 | 3,16E+07 | 2,41E+07 |
| 14 | | 0,0833822 | 0,0606472 | 0,48 | 5,73E+06 | 8,94E+06 |
| 55,8 | | 0,0836192 | 0,0608781 | 0,86 | 8,30E+08 | 1,03E+09 |
| 43,2 | | 0,0836353 | 0,0608866 | 1,34 | 4,09E+08 | 3,30E+08 |
| 39,7 | | 0,0837594 | 0,0609241 | 0,81 | 2,73E+08 | 2,32E+08 |
| 16,4 | | 0,0838047 | 0,0609394 | 0,88 | 2,51E+08 | 2,33E+08 |
| 54 | | 0,0839090 | 0,0611556 | 1,07 | 2,56E+08 | 2,56E+08 |
| 19,5 | | 0,0839201 | 0,0611408 | 0,83 | 3,16E+07 | 3,46E+07 |
| 9,5 | | 0,0839755 | 0,0612361 | 0,56 | 1,73E+07 | 1,89E+07 |
| 39 | | 0,0840507 | 0,0612677 | 1,11 | 1,58E+08 | 1,46E+08 |
| 22,7 | | 0,0842448 | 0,0614517 | 0,78 | 6,65E+07 | 7,20E+07 |
| 5,5 | | 0,0844180 | 0,0616540 | 0,35 | 6,83E+06 | 9,31E+06 |
| 23,5 | | 0,0847245 | 0,0620299 | 0,89 | 7,06E+07 | 7,06E+07 |

| | | | | | | |
|---|---|---|---|---|---|---|
| 9 | | 0,0847628 | 0,0620670 | 0,89 | 1,28E+07 | 1,41E+07 |
| 10,1 | | 0,0848778 | 0,0621051 | 1,06 | 6,38E+07 | 6,32E+07 |
| 28,4 | | 0,0849201 | 0,0620901 | 0,19 | 6,29E+07 | 6,40E+07 |
| 31,3 | | 0,0850384 | 0,0621870 | 0,77 | 6,01E+07 | 7,11E+07 |
| 25,8 | | 0,0850403 | 0,0621720 | 0,77 | 8,73E+07 | 6,31E+07 |
| 29,7 | | 0,0851032 | 0,0622177 | 0,58 | 5,32E+08 | 7,33E+08 |
| 37,2 | | 0,0851524 | 0,0622288 | 0,89 | 1,08E+09 | 9,88E+08 |
| 3,4 | | 0,0852053 | 0,0622667 | 2,29 | 6,46E+06 | 2,74E+06 |
| 16 | | 0,0853381 | 0,0623297 | 1,33 | 2,27E+07 | 1,62E+07 |
| 80,1 | | 0,0853826 | 0,0623754 | 1,06 | 4,97E+09 | 5,12E+09 |
| 15,9 | | 0,0854899 | 0,0624922 | 0,64 | 2,20E+07 | 2,37E+07 |
| 42,6 | | 0,0855473 | 0,0625368 | 0,83 | 7,56E+07 | 6,35E+07 |
| 34,5 | | 0,0855932 | 0,0625939 | 0,79 | 1,79E+08 | 1,96E+08 |
| 53,3 | | 0,0855938 | 0,0625788 | 0,89 | 4,03E+08 | 3,91E+08 |
| 21,5 | | 0,0856749 | 0,0625638 | 0,78 | 3,24E+07 | 3,25E+07 |
| 20,8 | | 0,0857137 | 0,0625786 | 0,66 | 1,08E+08 | 1,90E+08 |
| 10,4 | | 0,0857482 | 0,0626116 | 2,57 | 4,26E+06 | 1,52E+06 |
| 39,4 | | 0,0858160 | 0,0626532 | 0,86 | 5,51E+07 | 5,59E+07 |
| 8,6 | | 0,0858583 | 0,0627208 | 0,43 | 1,91E+07 | 1,86E+07 |
| 26,5 | | 0,0860305 | 0,0628728 | 0,87 | 2,07E+07 | 2,18E+07 |
| 13,1 | | 0,0860514 | 0,0629470 | 1,48 | 1,45E+07 | 1,37E+07 |
| 41,8 | | 0,0860608 | 0,0629319 | 0,71 | 1,16E+08 | 1,30E+08 |
| 63,8 | | 0,0860728 | 0,0629168 | 0,83 | 1,97E+09 | 2,55E+09 |
| 30,6 | | 0,0862655 | 0,0629765 | 0,84 | 2,11E+08 | 2,23E+08 |
| 57,5 | | 0,0864357 | 0,0631494 | 0,80 | 8,93E+07 | 1,22E+08 |
| 23,2 | | 0,0864758 | 0,0631649 | 0,90 | 3,14E+07 | 3,23E+07 |
| 72,4 | | 0,0865399 | 0,0632015 | 1,12 | 1,56E+09 | 1,63E+09 |
| 9 | | 0,0868983 | 0,0635544 | 0,65 | 1,47E+07 | 1,68E+07 |
| 14,3 | | 0,0869983 | 0,0635392 | 1,78 | 1,57E+07 | 1,65E+07 |
| 37 | | 0,0870186 | 0,0635239 | 0,95 | 2,37E+08 | 2,40E+08 |
| 13 | | 0,0871120 | 0,0635384 | 0,67 | 5,30E+06 | 6,31E+06 |
| 17,9 | | 0,0871398 | 0,0635720 | 3,85 | 1,22E+06 | 2,16E+06 |
| 7,7 | | 0,0871537 | 0,0635568 | 1,21 | 7,11E+06 | 6,45E+06 |
| 60,3 | | 0,0872039 | 0,0635416 | 1,24 | 2,39E+09 | 1,98E+09 |
| 23,8 | | 0,0872111 | 0,0635264 | 0,77 | 6,19E+07 | 8,10E+07 |
| 40,9 | | 0,0874248 | 0,0636346 | 1,12 | 1,26E+08 | 1,47E+08 |
| 24,9 | | 0,0874828 | 0,0636950 | 0,90 | 2,12E+07 | 2,13E+07 |
| 19,2 | | 0,0876805 | 0,0639197 | 0,73 | 8,28E+07 | 1,11E+08 |
| 17,5 | | 0,0877324 | 0,0639608 | 0,18 | 7,06E+07 | 5,38E+07 |
| 20,8 | | 0,0878308 | 0,0639752 | 5,09 | 2,39E+06 | 2,62E+07 |
| 45,5 | | 0,0878396 | 0,0639599 | 1,13 | 1,74E+08 | 1,94E+08 |
| 33,5 | | 0,0878537 | 0,0639446 | 0,69 | 2,49E+07 | 2,49E+07 |
| 50,5 | | 0,0880428 | 0,0641241 | 1,08 | 3,43E+08 | 3,44E+08 |
| 32,4 | | 0,0880438 | 0,0641088 | 1,09 | 2,55E+08 | 2,71E+08 |
| 44,2 | | 0,0880972 | 0,0641785 | 1,21 | 2,74E+08 | 2,97E+08 |
| 15,7 | | 0,0881059 | 0,0641632 | 0,72 | 4,39E+07 | 3,01E+07 |
| 75,4 | | 0,0882039 | 0,0642185 | 1,78 | 5,20E+07 | 8,38E+07 |
| 54,1 | | 0,0882637 | 0,0642690 | 0,63 | 3,02E+08 | 2,80E+08 |
| 25,8 | | 0,0884555 | 0,0645197 | 1,20 | 1,37E+07 | 1,68E+07 |
| 28,7 | | 0,0884822 | 0,0645043 | 0,89 | 9,83E+08 | 9,76E+08 |

| | | | | | | |
|---|---|---|---|---|---|---|
| 38,7 | | 0,0884915 | 0,0644889 | 0,40 | 2,41E+07 | 2,28E+07 |
| 24 | | 0,0885002 | 0,0644736 | 0,43 | 8,80E+06 | 9,66E+06 |
| 46,1 | | 0,0886126 | 0,0646316 | 0,93 | 3,14E+08 | 2,98E+08 |
| 5 | | 0,0886433 | 0,0646419 | 1,52 | 1,78E+06 | 2,62E+06 |
| 57,3 | | 0,0886864 | 0,0646265 | 0,72 | 6,72E+07 | 1,02E+08 |
| 21 | | 0,0887025 | 0,0646111 | 1,19 | 1,13E+08 | 1,02E+08 |
| 31,6 | | 0,0887302 | 0,0645958 | 0,91 | 4,39E+08 | 3,78E+08 |
| 29 | | 0,0888123 | 0,0646032 | 0,68 | 1,52E+08 | 1,47E+08 |
| 4,7 | | 0,0888929 | 0,0646183 | 2,35 | 1,38E+06 | 5,12E+06 |
| 12,9 | | 0,0888999 | 0,0646029 | 2,18 | 1,35E+07 | 4,65E+06 |
| 9,5 | | 0,0890344 | 0,0646541 | 0,32 | 2,36E+07 | 2,04E+07 |
| 39 | | 0,0890678 | 0,0646873 | 1,21 | 1,31E+08 | 1,34E+08 |
| 75,2 | | 0,0891641 | 0,0647014 | 1,28 | 1,86E+08 | 2,64E+08 |
| 8,3 | | 0,0892438 | 0,0647943 | 0,69 | 5,78E+06 | 6,40E+06 |
| 12,6 | | 0,0892513 | 0,0647789 | 0,73 | 3,00E+07 | 3,09E+07 |
| 6 | | 0,0894660 | 0,0650047 | 0,68 | 9,71E+06 | 1,24E+07 |
| 29,4 | | 0,0895073 | 0,0649893 | 0,66 | 5,04E+06 | 3,90E+06 |
| 35,3 | | 0,0895116 | 0,0649739 | 0,40 | 5,21E+06 | 6,37E+06 |
| 5,7 | | 0,0895628 | 0,0650353 | 0,40 | 1,25E+07 | 1,76E+07 |
| 21,1 | | 0,0896340 | 0,0650901 | 0,76 | 2,65E+07 | 2,23E+07 |
| 35,8 | | 0,0897212 | 0,0650747 | 0,88 | 1,71E+08 | 1,69E+08 |
| 7,4 | | 0,0897231 | 0,0650593 | 0,47 | 8,20E+06 | 7,74E+06 |
| 26,1 | | 0,0897639 | 0,0650913 | 0,60 | 1,48E+08 | 9,74E+07 |
| 44,3 | | 0,0897667 | 0,0650758 | 0,93 | 5,60E+08 | 6,10E+08 |
| 7,3 | | 0,0898651 | 0,0651429 | 1,91 | 2,36E+06 | 3,92E+06 |
| 46,6 | | 0,0900479 | 0,0653643 | 0,92 | 4,46E+08 | 4,29E+08 |
| 60,1 | | 0,0901641 | 0,0655667 | 0,83 | 1,53E+09 | 1,21E+09 |
| 10,3 | | 0,0904317 | 0,0656723 | 0,43 | 6,54E+06 | 9,60E+06 |
| 49,8 | | 0,0904601 | 0,0657079 | 1,09 | 9,17E+08 | 8,47E+08 |
| 6,1 | | 0,0906496 | 0,0659640 | 2,16 | 2,99E+06 | 3,60E+06 |
| 10 | | 0,0908916 | 0,0660743 | 0,58 | 1,24E+07 | 8,50E+06 |
| 24,9 | | 0,0910195 | 0,0662412 | 0,65 | 2,71E+07 | 1,47E+07 |
| 49,8 | | 0,0911005 | 0,0662965 | 1,14 | 2,76E+08 | 2,82E+08 |
| 7,9 | | 0,0911307 | 0,0663035 | 1,19 | 1,39E+07 | 1,44E+07 |
| 17,4 | | 0,0911790 | 0,0662879 | 1,18 | 4,27E+07 | 4,00E+07 |
| 25,2 | | 0,0912873 | 0,0663752 | 0,78 | 4,12E+08 | 3,94E+08 |
| 7,6 | | 0,0913846 | 0,0664985 | 0,48 | 4,10E+06 | 4,44E+06 |
| 18,4 | | 0,0914416 | 0,0665725 | 0,22 | 2,15E+07 | 2,76E+07 |
| 30,6 | | 0,0914746 | 0,0666050 | 0,87 | 7,06E+07 | 7,28E+07 |
| 31,6 | | 0,0915390 | 0,0666421 | 0,75 | 6,73E+07 | 5,38E+07 |
| 35,7 | | 0,0916383 | 0,0667076 | 0,73 | 5,46E+07 | 6,11E+07 |
| 16 | | 0,0918062 | 0,0667466 | 0,74 | 5,14E+07 | 5,21E+07 |
| 15,6 | | 0,0918607 | 0,0667563 | 1,51 | 1,27E+07 | 2,30E+07 |
| 5,9 | | 0,0918667 | 0,0667698 | 1,16 | 7,76E+06 | 8,71E+06 |
| 10,6 | | 0,0918788 | 0,0667541 | 0,89 | 2,47E+07 | 2,32E+07 |
| 51,7 | | 0,0919205 | 0,0667836 | 1,14 | 1,63E+09 | 1,85E+09 |
| 6,6 | | 0,0919696 | 0,0669375 | 1,23 | 9,10E+06 | 7,48E+06 |
| 8,5 | | 0,0920506 | 0,0670057 | 0,33 | 1,04E+07 | 1,10E+07 |
| 19,8 | | 0,0920627 | 0,0669899 | 0,65 | 2,02E+07 | 1,77E+07 |
| 18,5 | | 0,0922189 | 0,0671032 | 1,58 | 2,69E+07 | 2,90E+07 |

| | | | | | | |
|---|---|---|---|---|---|---|
| 9,6 | | 0,0923564 | 0,0672089 | 2,06 | 4,16E+06 | 9,33E+06 |
| 33,5 | | 0,0923772 | 0,0671930 | 0,88 | 6,48E+08 | 6,23E+08 |
| 43,2 | | 0,0924756 | 0,0672252 | 0,83 | 1,58E+08 | 1,36E+08 |
| 47,8 | | 0,0925131 | 0,0672395 | 1,39 | 1,03E+08 | 1,12E+08 |
| 31,9 | | 0,0925237 | 0,0672463 | 0,84 | 6,19E+07 | 5,76E+07 |
| 20,7 | | 0,0928301 | 0,0675804 | 0,91 | 1,23E+08 | 1,14E+08 |
| 25,7 | | 0,0931172 | 0,0680141 | 0,84 | 1,30E+08 | 1,68E+08 |
| 27,5 | | 0,0931254 | 0,0679981 | 0,83 | 8,69E+07 | 9,16E+07 |
| 26,4 | | 0,0932909 | 0,0681730 | 0,89 | 5,16E+08 | 5,56E+08 |
| 60,6 | | 0,0933043 | 0,0681786 | 0,91 | 1,57E+09 | 1,62E+09 |
| 46,3 | | 0,0935329 | 0,0684238 | 0,81 | 3,15E+08 | 3,25E+08 |
| 14,4 | | 0,0935478 | 0,0684077 | 0,86 | 1,15E+08 | 1,22E+08 |
| 32,7 | | 0,0936552 | 0,0685062 | 1,25 | 1,03E+08 | 1,43E+08 |
| 13 | | 0,0936779 | 0,0685155 | 0,50 | 1,32E+07 | 2,50E+07 |
| 34,7 | | 0,0937783 | 0,0686383 | 1,33 | 4,03E+07 | 6,05E+07 |
| 61,9 | | 0,0937976 | 0,0686222 | 1,17 | 7,52E+08 | 7,86E+08 |
| 27,3 | | 0,0940998 | 0,0687225 | 1,29 | 3,55E+07 | 2,99E+07 |
| 22,3 | | 0,0941505 | 0,0687580 | 0,83 | 1,75E+08 | 2,19E+08 |
| 9,7 | | 0,0941931 | 0,0687672 | 1,24 | 3,42E+07 | 3,88E+07 |
| 31,6 | | 0,0942078 | 0,0688345 | 1,16 | 1,24E+08 | 1,41E+08 |
| 40,3 | | 0,0943298 | 0,0689018 | 0,91 | 3,63E+08 | 3,94E+08 |
| 16 | | 0,0945208 | 0,0690169 | 1,43 | 3,62E+07 | 2,24E+07 |
| 10,4 | | 0,0946374 | 0,0690560 | 0,56 | 2,33E+07 | 5,14E+07 |
| 33 | | 0,0946571 | 0,0690848 | 1,24 | 6,63E+07 | 6,31E+07 |
| 38,5 | | 0,0947152 | 0,0690686 | 0,85 | 3,98E+08 | 4,25E+08 |
| 8,5 | | 0,0947936 | 0,0690815 | 1,64 | 3,80E+06 | 8,22E+06 |
| 44,5 | | 0,0948423 | 0,0690653 | 0,84 | 7,28E+07 | 9,58E+07 |
| 45,4 | | 0,0949669 | 0,0691175 | 1,15 | 2,28E+08 | 2,09E+08 |
| 14,7 | | 0,0950956 | 0,0691771 | 0,75 | 1,92E+07 | 2,25E+07 |
| 18,6 | | 0,0952049 | 0,0693545 | 1,34 | 3,09E+08 | 2,25E+08 |
| 41,6 | | 0,0952504 | 0,0693673 | 1,24 | 1,86E+08 | 1,79E+08 |
| 30,7 | | 0,0954131 | 0,0694530 | 0,90 | 1,48E+08 | 1,62E+08 |
| 11 | | 0,0958243 | 0,0699285 | 0,72 | 1,16E+07 | 1,49E+07 |
| 3,8 | | 0,0960393 | 0,0702879 | 2,65 | 5,54E+06 | 1,20E+06 |
| 41 | | 0,0960733 | 0,0703294 | 1,19 | 2,53E+08 | 2,20E+08 |
| 8,6 | | 0,0961134 | 0,0703858 | 0,71 | 2,31E+07 | 2,12E+07 |
| 23,4 | | 0,0961451 | 0,0704217 | 1,72 | 4,82E+07 | 2,37E+07 |
| 63,9 | | 0,0961623 | 0,0704052 | 1,26 | 1,22E+09 | 1,41E+09 |
| 34,7 | | 0,0962445 | 0,0704756 | 0,50 | 9,74E+07 | 9,94E+07 |
| 46,7 | | 0,0963604 | 0,0705357 | 1,06 | 1,12E+09 | 1,11E+09 |
| 23,3 | + | 0,0964106 | 0,0705930 | 1,43 | 1,21E+07 | 8,82E+06 |
| 33,5 | | 0,0964350 | 0,0706194 | 1,12 | 2,40E+08 | 2,08E+08 |
| 13,8 | | 0,0965280 | 0,0706859 | 0,85 | 2,11E+07 | 2,05E+07 |
| 51,1 | | 0,0966873 | 0,0708131 | 0,87 | 2,09E+08 | 2,32E+08 |
| 6,8 | | 0,0967780 | 0,0708441 | 1,31 | 6,52E+06 | 5,92E+06 |
| 15,6 | | 0,0968358 | 0,0709618 | 2,70 | 1,60E+07 | 4,51E+06 |
| 2,2 | | 0,0968596 | 0,0709453 | 3,46 | 9,41E+06 | 1,29E+06 |
| 41 | | 0,0968949 | 0,0709287 | 1,20 | 1,34E+08 | 1,55E+08 |
| 15,9 | | 0,0970539 | 0,0709346 | 1,62 | 1,67E+07 | 2,21E+07 |
| 11,6 | | 0,0972050 | 0,0711201 | 0,67 | 6,49E+06 | 6,38E+06 |

| | | | | | | |
|---|---|---|---|---|---|---|
| 35,1 | | 0,0972401 | 0,0711287 | 1,10 | 1,75E+08 | 1,51E+08 |
| 51,5 | | 0,0972508 | 0,0711121 | 1,16 | 1,49E+08 | 1,46E+08 |
| 21,9 | | 0,0973878 | 0,0712929 | 1,16 | 4,96E+07 | 4,73E+07 |
| 12 | + | 0,0974317 | 0,0713274 | 6,25 | 6,81E+06 | 7,84E+05 |
| 44,3 | | 0,0975692 | 0,0713806 | 1,41 | 3,52E+07 | 3,43E+07 |
| 45,9 | | 0,0976923 | 0,0714747 | 1,12 | 6,51E+08 | 6,82E+08 |
| 25,6 | | 0,0979814 | 0,0716970 | 1,18 | 5,15E+07 | 4,52E+07 |
| 34,4 | | 0,0980167 | 0,0716803 | 0,87 | 2,32E+08 | 2,27E+08 |
| 45,1 | | 0,0980457 | 0,0716859 | 0,84 | 1,81E+08 | 2,23E+08 |
| 6,5 | | 0,0980995 | 0,0717269 | 0,60 | 4,52E+06 | 3,26E+06 |
| 32,3 | | 0,0982991 | 0,0719127 | 0,65 | 1,99E+07 | 1,46E+07 |
| 31,9 | | 0,0983170 | 0,0719174 | 0,91 | 3,91E+08 | 4,25E+08 |
| 25,7 | | 0,0984000 | 0,0720019 | 1,15 | 9,41E+07 | 8,62E+07 |
| 33,3 | | 0,0984467 | 0,0720074 | 1,12 | 1,60E+08 | 1,53E+08 |
| 4,3 | | 0,0984484 | 0,0719907 | 0,78 | 1,21E+07 | 8,14E+06 |
| 3,6 | | 0,0986466 | 0,0721790 | 0,64 | 5,09E+06 | 3,66E+06 |
| 35,9 | | 0,0986848 | 0,0722448 | 0,65 | 1,58E+08 | 1,38E+08 |
| 43 | | 0,0988359 | 0,0724052 | 0,35 | 8,91E+06 | 4,08E+07 |
| 5,1 | | 0,0988880 | 0,0723884 | 0,19 | 4,40E+07 | 3,04E+07 |
| 23 | | 0,0991058 | 0,0725774 | 0,82 | 8,74E+07 | 7,71E+07 |
| 13,1 | | 0,0991264 | 0,0725606 | 0,67 | 2,38E+07 | 1,73E+07 |
| 30,4 | | 0,0992874 | 0,0726170 | 1,21 | 1,69E+08 | 1,71E+08 |
| 5,4 | | 0,0993468 | 0,0726001 | 1,81 | 1,54E+07 | 6,21E+06 |
| 69 | | 0,0994235 | 0,0727519 | 0,92 | 4,95E+09 | 5,61E+09 |
| 5,7 | | 0,0994283 | 0,0727924 | 1,20 | 5,80E+06 | 5,85E+06 |
| 52,2 | | 0,0996130 | 0,0729190 | 1,12 | 2,50E+08 | 2,54E+08 |
| 12,1 | | 0,0996529 | 0,0729234 | 0,39 | 7,73E+06 | 4,78E+06 |
| 42,3 | | 0,0999110 | 0,0732359 | 0,87 | 4,17E+09 | 4,96E+09 |
| 12,6 | | 0,0999751 | 0,0732698 | 0,78 | 2,17E+07 | 1,68E+07 |
| 12,7 | | 0,0999794 | 0,0732779 | 0,38 | 9,14E+06 | 1,31E+07 |
| 47,4 | | 0,0999905 | 0,0732896 | 1,06 | 4,47E+08 | 4,13E+08 |
| 28,2 | | 0,1000097 | 0,0732726 | 0,60 | 3,21E+07 | 2,76E+07 |
| 34,5 | | 0,1000294 | 0,0732557 | 1,16 | 1,60E+09 | 2,01E+09 |
| 12,5 | | 0,1000577 | 0,0732610 | 0,70 | 3,11E+07 | 2,26E+07 |
| 59,1 | | 0,1001322 | 0,0732690 | 0,91 | 1,20E+09 | 1,16E+09 |
| 55,8 | | 0,1002415 | 0,0733213 | 1,17 | 1,11E+09 | 1,07E+09 |
| 47,7 | | 0,1002689 | 0,0733044 | 1,16 | 1,17E+08 | 1,06E+08 |
| 24,3 | | 0,1002839 | 0,0732875 | 0,88 | 1,07E+08 | 9,72E+07 |
| 27,9 | | 0,1002848 | 0,0732706 | 0,78 | 3,47E+08 | 4,79E+08 |
| 5,4 | | 0,1003795 | 0,0732970 | 0,52 | 5,30E+06 | 5,65E+06 |
| 15,4 | | 0,1004085 | 0,0732801 | 0,51 | 7,23E+06 | 1,81E+07 |
| 30,4 | | 0,1004169 | 0,0732845 | 0,84 | 3,81E+08 | 4,20E+08 |
| 16,6 | | 0,1004726 | 0,0732962 | 1,66 | 1,06E+07 | 1,27E+07 |
| 39,3 | | 0,1005557 | 0,0733263 | 1,13 | 1,09E+08 | 1,02E+08 |
| 10 | | 0,1008179 | 0,0735388 | 1,52 | 3,05E+07 | 2,40E+07 |
| 32,9 | | 0,1008899 | 0,0735735 | 1,14 | 1,30E+08 | 1,52E+08 |
| 26,9 | | 0,1010982 | 0,0737149 | 0,79 | 1,93E+08 | 1,97E+08 |
| 12 | | 0,1011057 | 0,0736980 | 0,79 | 3,15E+07 | 2,40E+07 |
| 20,6 | | 0,1011444 | 0,0736810 | 1,17 | 6,15E+07 | 6,55E+07 |
| 45,7 | | 0,1012007 | 0,0737285 | 1,27 | 9,12E+07 | 7,05E+07 |

| | | | | | | |
|---|---|---|---|---|---|---|
| 15 | | 0,1015139 | 0,0739425 | 0,75 | 2,29E+07 | 1,84E+07 |
| 46,3 | | 0,1019084 | 0,0743128 | 0,84 | 7,40E+07 | 7,97E+07 |
| 20,8 | | 0,1020285 | 0,0743490 | 0,85 | 1,41E+07 | 1,44E+07 |
| 35,4 | | 0,1021072 | 0,0743825 | 1,80 | 2,27E+08 | 1,91E+08 |
| 17,4 | | 0,1021289 | 0,0743939 | 0,55 | 2,07E+07 | 9,65E+06 |
| 19,9 | | 0,1022386 | 0,0743989 | 0,57 | 1,72E+07 | 9,51E+06 |
| 33,4 | | 0,1022481 | 0,0743818 | 0,87 | 3,62E+08 | 3,00E+08 |
| 43,5 | | 0,1023188 | 0,0743895 | 1,17 | 1,77E+08 | 1,87E+08 |
| 44,3 | | 0,1025251 | 0,0745543 | 0,87 | 5,53E+07 | 5,05E+07 |
| 10,9 | | 0,1028250 | 0,0746841 | 0,33 | 1,04E+07 | 1,49E+07 |
| 38,3 | | 0,1028267 | 0,0746670 | 0,90 | 1,27E+08 | 1,31E+08 |
| 1,2 | | 0,1028322 | 0,0746498 | 3,60 | 3,58E+05 | 3,15E+05 |
| 17,5 | | 0,1028810 | 0,0746327 | 0,90 | 2,04E+08 | 2,32E+08 |
| 16,7 | | 0,1030107 | 0,0747404 | 1,71 | 9,45E+06 | 8,44E+06 |
| 35,7 | | 0,1030167 | 0,0747232 | 1,29 | 1,11E+08 | 9,22E+07 |
| 20,6 | | 0,1030276 | 0,0747061 | 0,83 | 6,91E+07 | 8,29E+07 |
| 57,6 | | 0,1030705 | 0,0746890 | 0,88 | 2,62E+08 | 2,87E+08 |
| 15,5 | | 0,1031664 | 0,0746939 | 1,35 | 2,30E+07 | 1,71E+07 |
| 53,4 | | 0,1033134 | 0,0748270 | 1,22 | 6,09E+08 | 6,31E+08 |
| 11,3 | | 0,1033548 | 0,0748099 | 1,11 | 2,81E+07 | 2,38E+07 |
| 62,7 | | 0,1033727 | 0,0748367 | 1,13 | 4,58E+08 | 4,10E+08 |
| 17,1 | | 0,1034536 | 0,0748709 | 1,16 | 7,27E+07 | 6,99E+07 |
| 7,1 | | 0,1037022 | 0,0749792 | 0,45 | 1,10E+07 | 2,67E+07 |
| 9,3 | | 0,1037390 | 0,0749831 | 1,58 | 2,69E+06 | 2,05E+06 |
| 16,2 | | 0,1040497 | 0,0756962 | 0,84 | 3,29E+07 | 3,71E+07 |
| 8,9 | | 0,1042028 | 0,0757969 | 1,20 | 2,24E+07 | 2,58E+07 |
| 27,9 | | 0,1042893 | 0,0758829 | 1,37 | 2,63E+07 | 2,48E+07 |
| 33,4 | | 0,1042940 | 0,0759223 | 0,83 | 7,16E+07 | 8,75E+07 |
| 29,8 | | 0,1046611 | 0,0762368 | 0,73 | 5,47E+07 | 5,57E+07 |
| 58,5 | | 0,1047382 | 0,0763574 | 1,09 | 3,87E+08 | 3,95E+08 |
| 23,2 | | 0,1047521 | 0,0763400 | 0,82 | 3,32E+07 | 2,68E+07 |
| 1,8 | | 0,1048463 | 0,0763444 | 0,75 | 5,19E+07 | 8,28E+07 |
| 14,3 | | 0,1051205 | 0,0764613 | 1,22 | 3,25E+07 | 2,83E+07 |
| 80,3 | | 0,1052221 | 0,0765854 | 1,12 | 9,00E+08 | 1,07E+09 |
| 72,2 | | 0,1053239 | 0,0766391 | 1,83 | 5,75E+08 | 7,79E+08 |
| 16,5 | | 0,1053992 | 0,0767001 | 0,45 | 3,25E+07 | 7,11E+07 |
| 14,3 | | 0,1054203 | 0,0766826 | 0,56 | 7,91E+06 | 4,36E+06 |
| 5,4 | | 0,1054295 | 0,0766861 | 1,36 | 5,29E+06 | 3,46E+06 |
| 30,6 | | 0,1056593 | 0,0768921 | 0,93 | 1,09E+08 | 1,05E+08 |
| 27,7 | | 0,1057675 | 0,0770278 | 1,14 | 5,51E+07 | 5,28E+07 |
| 24 | | 0,1058024 | 0,0770595 | 1,08 | 8,43E+07 | 7,91E+07 |
| 17,9 | | 0,1061748 | 0,0773309 | 0,77 | 8,16E+06 | 6,34E+06 |
| 31,7 | | 0,1062120 | 0,0773661 | 0,77 | 4,72E+07 | 4,94E+07 |
| 10 | | 0,1064150 | 0,0775408 | 2,14 | 2,10E+07 | 2,14E+07 |
| 12,4 | | 0,1064283 | 0,0775523 | 0,27 | 8,79E+06 | 1,90E+06 |
| 7,5 | | 0,1064570 | 0,0775783 | 1,77 | 2,65E+07 | 4,36E+07 |
| 64,2 | | 0,1065430 | 0,0775889 | 1,11 | 1,12E+09 | 9,96E+08 |
| 14,9 | | 0,1065770 | 0,0775712 | 0,61 | 3,05E+08 | 2,86E+08 |
| 11,2 | | 0,1073102 | 0,0783418 | 0,53 | 1,18E+07 | 1,49E+07 |
| 29,3 | | 0,1073544 | 0,0783485 | 0,89 | 1,30E+08 | 1,36E+08 |

| | | | | | | |
|---|---|---|---|---|---|---|
| 12,8 | | 0,1074595 | 0,0785381 | 1,30 | 8,65E+06 | 7,41E+06 |
| 16,9 | | 0,1075930 | 0,0787058 | 1,59 | 6,07E+07 | 3,18E+07 |
| 15,7 | | 0,1078154 | 0,0789270 | 0,65 | 6,66E+06 | 5,51E+06 |
| 29,6 | | 0,1078939 | 0,0790227 | 0,76 | 7,71E+07 | 7,97E+07 |
| 8,8 | | 0,1080746 | 0,0791984 | 0,73 | 2,59E+07 | 3,19E+07 |
| 17,6 | | 0,1081079 | 0,0791804 | 1,59 | 5,56E+06 | 4,43E+06 |
| 42,4 | | 0,1081673 | 0,0791906 | 1,26 | 6,32E+08 | 7,79E+08 |
| 16,9 | | 0,1082652 | 0,0792289 | 0,61 | 1,71E+07 | 1,81E+07 |
| 15,2 | | 0,1083168 | 0,0792390 | 1,45 | 1,05E+07 | 9,98E+06 |
| 28,6 | | 0,1083513 | 0,0792456 | 0,87 | 8,92E+07 | 7,63E+07 |
| 26,4 | | 0,1085478 | 0,0795716 | 0,10 | 1,24E+08 | 1,87E+08 |
| 3,5 | | 0,1086297 | 0,0796515 | 1,64 | 1,37E+07 | 6,04E+06 |
| 12,3 | | 0,1088249 | 0,0797079 | 0,64 | 1,22E+07 | 1,61E+07 |
| 17,8 | | 0,1088510 | 0,0797179 | 1,13 | 6,30E+07 | 7,16E+07 |
| 31,7 | | 0,1088878 | 0,0797715 | 1,25 | 3,31E+07 | 4,07E+07 |
| 15 | | 0,1090252 | 0,0798722 | 0,86 | 1,97E+07 | 1,65E+07 |
| 46 | | 0,1090681 | 0,0799112 | 0,81 | 3,56E+08 | 3,54E+08 |
| 36,2 | | 0,1092888 | 0,0800761 | 0,91 | 2,78E+08 | 2,98E+08 |
| 8,3 | | 0,1093969 | 0,0801894 | 1,63 | 2,89E+07 | 2,64E+07 |
| 29,4 | | 0,1095216 | 0,0803705 | 0,91 | 3,66E+08 | 3,16E+08 |
| 17,5 | | 0,1095292 | 0,0803740 | 0,55 | 4,35E+06 | 4,94E+06 |
| 25,4 | | 0,1095896 | 0,0804047 | 1,09 | 2,19E+08 | 2,38E+08 |
| 12,4 | | 0,1097359 | 0,0805033 | 0,63 | 1,38E+07 | 1,38E+07 |
| 39,7 | | 0,1098531 | 0,0805801 | 1,14 | 1,92E+08 | 2,29E+08 |
| 44,8 | | 0,1102614 | 0,0808632 | 0,86 | 9,11E+07 | 9,87E+07 |
| 10,4 | | 0,1104450 | 0,0809462 | 0,70 | 3,19E+07 | 1,73E+07 |
| 33 | | 0,1104791 | 0,0809523 | 1,35 | 9,70E+07 | 9,95E+07 |
| 38,9 | | 0,1106337 | 0,0809901 | 1,08 | 1,10E+09 | 1,10E+09 |
| 33,5 | | 0,1106606 | 0,0809718 | 1,16 | 1,58E+09 | 1,57E+09 |
| 51,9 | | 0,1106663 | 0,0809535 | 1,27 | 5,09E+07 | 7,46E+07 |
| 52,3 | | 0,1108585 | 0,0810626 | 1,30 | 9,74E+07 | 9,14E+07 |
| 15,2 | | 0,1108893 | 0,0811183 | 0,89 | 3,10E+07 | 3,76E+07 |
| 16,4 | | 0,1109991 | 0,0812210 | 0,73 | 2,70E+06 | 4,53E+06 |
| 28,8 | | 0,1110332 | 0,0812451 | 0,87 | 4,02E+07 | 3,85E+07 |
| 50,9 | | 0,1114042 | 0,0816078 | 1,09 | 1,09E+09 | 1,21E+09 |
| 35,2 | | 0,1115285 | 0,0817202 | 0,59 | 2,01E+07 | 1,28E+07 |
| 6,2 | | 0,1116575 | 0,0819174 | 0,68 | 5,87E+06 | 3,83E+06 |
| 2,6 | | 0,1116692 | 0,0819269 | 0,48 | 8,63E+06 | 4,03E+06 |
| 9,1 | | 0,1118314 | 0,0820555 | 0,77 | 1,02E+07 | 1,05E+07 |
| 32 | | 0,1119127 | 0,0820613 | 0,92 | 5,80E+08 | 6,49E+08 |
| 2,9 | | 0,1121096 | 0,0822592 | 1,55 | 4,56E+06 | 2,93E+06 |
| 29,7 | | 0,1121293 | 0,0822406 | 1,12 | 2,20E+08 | 2,02E+08 |
| 38,9 | | 0,1122516 | 0,0823212 | 0,83 | 4,62E+08 | 4,36E+08 |
| 16,3 | | 0,1123006 | 0,0823243 | 0,85 | 7,58E+07 | 7,92E+07 |
| 18,8 | | 0,1124321 | 0,0823544 | 0,74 | 1,32E+08 | 8,97E+07 |
| 20,8 | | 0,1124555 | 0,0823602 | 0,93 | 3,30E+07 | 3,40E+07 |
| 15,5 | | 0,1124725 | 0,0823633 | 1,19 | 1,51E+07 | 1,21E+07 |
| 8,3 | | 0,1128391 | 0,0825374 | 0,86 | 2,35E+07 | 2,25E+07 |
| 26,7 | | 0,1132591 | 0,0827663 | 0,71 | 8,26E+07 | 5,30E+07 |
| 38,2 | | 0,1133552 | 0,0827998 | 0,78 | 3,22E+08 | 3,91E+08 |

| | | | | | | |
|---|---|---|---|---|---|---|
| 17,5 | | 0,1134553 | 0,0828550 | 1,62 | 6,42E+06 | 1,43E+07 |
| 30,1 | | 0,1134678 | 0,0828363 | 0,94 | 4,52E+07 | 4,12E+07 |
| 51 | | 0,1135026 | 0,0828600 | 0,90 | 2,26E+08 | 2,35E+08 |
| 24,5 | | 0,1137151 | 0,0831182 | 1,16 | 3,96E+08 | 4,27E+08 |
| 21,1 | | 0,1139678 | 0,0832882 | 1,37 | 5,17E+07 | 3,42E+07 |
| 3,6 | | 0,1140730 | 0,0834313 | 2,42 | 4,90E+06 | 1,24E+06 |
| 10,7 | | 0,1141086 | 0,0834646 | 0,82 | 2,88E+07 | 2,47E+07 |
| 34,9 | | 0,1141890 | 0,0835447 | 0,88 | 2,59E+08 | 2,96E+08 |
| 28,6 | | 0,1142880 | 0,0836004 | 0,27 | 1,41E+07 | 1,89E+07 |
| 12,7 | | 0,1142946 | 0,0836023 | 1,33 | 2,37E+06 | 2,77E+06 |
| 46 | | 0,1142984 | 0,0835836 | 1,08 | 5,12E+08 | 4,62E+08 |
| 39,5 | | 0,1144745 | 0,0838277 | 0,81 | 2,95E+08 | 2,68E+08 |
| 29,5 | | 0,1147443 | 0,0839740 | 0,87 | 9,73E+07 | 8,59E+07 |
| 52,3 | | 0,1149550 | 0,0842269 | 1,06 | 5,36E+08 | 4,97E+08 |
| 19,3 | | 0,1151054 | 0,0843085 | 0,83 | 1,20E+08 | 1,21E+08 |
| 19,2 | | 0,1152762 | 0,0845334 | 0,17 | 2,60E+07 | 3,00E+07 |
| 21,7 | | 0,1156188 | 0,0847009 | 0,80 | 2,58E+08 | 3,42E+08 |
| 51,4 | | 0,1156663 | 0,0846819 | 1,09 | 1,65E+09 | 1,50E+09 |
| 18,7 | | 0,1156719 | 0,0847086 | 0,81 | 9,10E+08 | 1,06E+09 |
| 44,1 | | 0,1158875 | 0,0848348 | 0,89 | 2,76E+08 | 2,67E+08 |
| 36,6 | | 0,1160277 | 0,0849940 | 1,16 | 1,54E+08 | 1,86E+08 |
| 62,1 | | 0,1161482 | 0,0852095 | 0,92 | 1,94E+09 | 1,90E+09 |
| 37 | | 0,1162672 | 0,0852397 | 0,89 | 4,49E+08 | 4,88E+08 |
| 14,1 | | 0,1167849 | 0,0857745 | 0,58 | 8,22E+06 | 1,06E+07 |
| 16,9 | | 0,1168759 | 0,0858788 | 4,46 | 1,68E+06 | 1,78E+07 |
| 26,1 | | 0,1169429 | 0,0858873 | 1,12 | 1,14E+08 | 9,51E+07 |
| 8,1 | | 0,1170849 | 0,0860371 | 0,79 | 4,15E+08 | 2,99E+08 |
| 10,3 | | 0,1171527 | 0,0860912 | 2,62 | 1,27E+07 | 4,48E+06 |
| 49,9 | | 0,1173346 | 0,0862177 | 0,92 | 2,56E+09 | 2,81E+09 |
| 31,9 | | 0,1174073 | 0,0863298 | 0,84 | 1,02E+08 | 1,31E+08 |
| 5,1 | | 0,1174534 | 0,0863105 | 0,59 | 3,48E+06 | 7,95E+06 |
| 36 | | 0,1174706 | 0,0863126 | 1,29 | 6,69E+07 | 6,87E+07 |
| 42,5 | | 0,1174987 | 0,0863175 | 1,11 | 3,42E+08 | 3,85E+08 |
| 32,4 | | 0,1175006 | 0,0862982 | 1,07 | 1,86E+08 | 1,88E+08 |
| 31,3 | | 0,1175899 | 0,0863691 | 1,14 | 1,55E+08 | 1,31E+08 |
| 5,4 | | 0,1176068 | 0,0863498 | 0,71 | 9,13E+06 | 1,12E+07 |
| 15,1 | | 0,1176325 | 0,0863823 | 1,07 | 4,24E+07 | 4,25E+07 |
| 11,9 | | 0,1176668 | 0,0863872 | 0,74 | 7,41E+06 | 9,94E+06 |
| 46 | | 0,1177737 | 0,0864901 | 0,91 | 3,49E+08 | 3,50E+08 |
| 74,8 | | 0,1180748 | 0,0868052 | 1,15 | 9,26E+07 | 1,08E+08 |
| 26,6 | | 0,1182661 | 0,0869374 | 0,68 | 5,84E+06 | 4,60E+06 |
| 17,5 | | 0,1183514 | 0,0870187 | 1,16 | 2,92E+07 | 3,53E+07 |
| 28,4 | | 0,1185053 | 0,0870474 | 0,60 | 1,12E+07 | 1,49E+07 |
| 28,1 | | 0,1185430 | 0,0870797 | 1,12 | 3,55E+07 | 3,44E+07 |
| 24,8 | | 0,1186450 | 0,0871387 | 0,89 | 8,60E+07 | 9,19E+07 |
| 14,4 | | 0,1186844 | 0,0871193 | 1,09 | 2,28E+07 | 2,11E+07 |
| 49,8 | | 0,1186887 | 0,0870999 | 0,96 | 1,67E+08 | 1,62E+08 |
| 67,1 | | 0,1186901 | 0,0870806 | 0,87 | 1,71E+09 | 1,50E+09 |
| 7,2 | | 0,1187307 | 0,0871101 | 0,55 | 7,77E+06 | 1,10E+07 |
| 9,4 | | 0,1187414 | 0,0870907 | 0,67 | 1,23E+07 | 1,29E+07 |

| | | | | | | |
|---|---|---|---|---|---|---|
| 24,3 | | 0,1189535 | 0,0873453 | 0,65 | 3,01E+07 | 3,90E+07 |
| 17,4 | | 0,1191050 | 0,0874949 | 0,59 | 2,98E+06 | 3,32E+06 |
| 26,4 | | 0,1191405 | 0,0875315 | 1,21 | 1,40E+08 | 1,20E+08 |
| 16,9 | | 0,1191427 | 0,0875120 | 2,57 | 5,16E+06 | 1,65E+07 |
| 32,2 | | 0,1191908 | 0,0875379 | 0,78 | 2,22E+08 | 1,73E+08 |
| 13,2 | | 0,1192427 | 0,0875424 | 0,84 | 1,68E+07 | 1,64E+07 |
| 11,6 | | 0,1194076 | 0,0875896 | 2,02 | 7,20E+06 | 2,40E+06 |
| 20,9 | | 0,1196376 | 0,0877913 | 0,66 | 2,77E+07 | 2,66E+07 |
| 50 | | 0,1197013 | 0,0877922 | 1,21 | 2,88E+09 | 2,96E+09 |
| 20,9 | | 0,1197736 | 0,0878216 | 2,19 | 6,45E+06 | 2,13E+06 |
| 13,7 | | 0,1197940 | 0,0878305 | 1,31 | 2,40E+07 | 3,48E+07 |
| 4,2 | | 0,1198047 | 0,0878110 | 1,37 | 3,95E+06 | 2,34E+06 |
| 5,4 | | 0,1198998 | 0,0879086 | 1,31 | 7,43E+07 | 8,90E+07 |
| 3 | | 0,1200307 | 0,0880807 | 0,75 | 5,00E+06 | 3,97E+06 |
| 14,3 | | 0,1200714 | 0,0881312 | 0,49 | 4,18E+06 | 6,95E+06 |
| 23 | | 0,1201998 | 0,0881534 | 1,19 | 1,35E+07 | 1,56E+07 |
| 77 | | 0,1204430 | 0,0884582 | 0,80 | 1,10E+09 | 1,07E+09 |
| 25 | | 0,1206000 | 0,0884661 | 0,92 | 1,60E+08 | 1,77E+08 |
| 20,4 | | 0,1206033 | 0,0884465 | 6,69 | 1,60E+06 | 4,28E+07 |
| 55,1 | | 0,1206874 | 0,0885244 | 1,14 | 1,12E+09 | 1,24E+09 |
| 20,5 | | 0,1208817 | 0,0886978 | 0,93 | 3,24E+07 | 3,08E+07 |
| 23,5 | | 0,1208898 | 0,0886782 | 0,60 | 7,58E+07 | 8,97E+07 |
| 35 | | 0,1210925 | 0,0889604 | 0,79 | 4,79E+07 | 3,87E+07 |
| 13,5 | | 0,1211601 | 0,0890673 | 1,47 | 1,44E+07 | 2,79E+07 |
| 38 | | 0,1212406 | 0,0891201 | 0,54 | 4,40E+07 | 1,99E+07 |
| 16,5 | | 0,1217506 | 0,0894463 | 0,67 | 1,48E+07 | 2,55E+07 |
| 65,2 | | 0,1219597 | 0,0895370 | 1,09 | 7,60E+08 | 8,35E+08 |
| 31,2 | | 0,1220126 | 0,0896004 | 1,17 | 1,00E+08 | 1,29E+08 |
| 19,7 | | 0,1220320 | 0,0895806 | 1,23 | 2,24E+07 | 2,27E+07 |
| 13,9 | | 0,1220668 | 0,0896103 | 1,33 | 2,62E+07 | 1,87E+07 |
| 13,5 | | 0,1221880 | 0,0897522 | 0,85 | 4,21E+07 | 4,59E+07 |
| 38,2 | | 0,1223308 | 0,0897977 | 1,89 | 1,09E+08 | 1,42E+08 |
| 6,8 | | 0,1224292 | 0,0898459 | 0,82 | 1,10E+07 | 8,08E+06 |
| 39,3 | | 0,1224648 | 0,0898534 | 0,92 | 5,31E+08 | 5,94E+08 |
| 7,6 | | 0,1224672 | 0,0898336 | 0,70 | 8,06E+06 | 5,39E+06 |
| 42 | | 0,1225770 | 0,0898853 | 0,85 | 2,05E+09 | 2,40E+09 |
| 15,4 | | 0,1227927 | 0,0901205 | 1,19 | 1,38E+07 | 1,49E+07 |
| 63,3 | | 0,1228570 | 0,0901703 | 0,73 | 4,90E+09 | 3,62E+09 |
| 5,7 | | 0,1228864 | 0,0901716 | 0,90 | 1,77E+07 | 1,81E+07 |
| 21,1 | | 0,1229539 | 0,0901799 | 1,21 | 3,14E+07 | 2,87E+07 |
| 6,5 | | 0,1230559 | 0,0902561 | 0,76 | 1,88E+07 | 2,31E+07 |
| 46,9 | | 0,1232992 | 0,0904169 | 1,12 | 4,23E+09 | 4,07E+09 |
| 21,9 | | 0,1235381 | 0,0906878 | 1,27 | 4,89E+07 | 5,09E+07 |
| 56 | | 0,1236431 | 0,0907233 | 1,14 | 7,92E+07 | 7,51E+07 |
| 9,4 | | 0,1239291 | 0,0909923 | 0,51 | 5,98E+06 | 3,94E+06 |
| 15,1 | | 0,1241741 | 0,0911035 | 0,84 | 7,35E+06 | 7,91E+06 |
| 9,6 | | 0,1242000 | 0,0910834 | 0,77 | 1,89E+07 | 1,85E+07 |
| 18,9 | | 0,1244403 | 0,0912641 | 0,54 | 3,32E+06 | 7,42E+06 |
| 48,6 | | 0,1247063 | 0,0914438 | 0,90 | 4,92E+08 | 4,34E+08 |
| 54,5 | | 0,1249586 | 0,0915486 | 1,13 | 7,92E+08 | 7,69E+08 |

| | | | | | | |
|---|---|---|---|---|---|---|
| 22,7 | | 0,1249923 | 0,0915285 | 1,24 | 7,68E+07 | 6,41E+07 |
| 38,3 | | 0,1251083 | 0,0916042 | 0,93 | 5,96E+07 | 5,57E+07 |
| 17,7 | | 0,1253710 | 0,0917239 | 0,92 | 1,50E+08 | 1,46E+08 |
| 35,2 | | 0,1254440 | 0,0917785 | 0,81 | 2,27E+08 | 2,77E+08 |
| 32,9 | | 0,1254775 | 0,0917583 | 1,09 | 2,21E+08 | 2,19E+08 |
| 41,8 | | 0,1257260 | 0,0919895 | 1,23 | 1,23E+08 | 1,34E+08 |
| 8,2 | | 0,1259430 | 0,0921133 | 1,25 | 7,37E+06 | 7,29E+06 |
| 15,7 | | 0,1262565 | 0,0923364 | 0,20 | 4,74E+07 | 3,31E+06 |
| 8,9 | | 0,1263959 | 0,0923785 | 1,43 | 3,54E+07 | 6,33E+07 |
| 76,1 | | 0,1267976 | 0,0926989 | 1,10 | 4,27E+08 | 4,53E+08 |
| 4,6 | | 0,1267985 | 0,0926785 | 1,24 | 5,61E+06 | 5,18E+06 |
| 13,5 | | 0,1268619 | 0,0927214 | 1,43 | 8,03E+06 | 1,30E+07 |
| 19,8 | | 0,1270009 | 0,0928239 | 0,77 | 1,44E+07 | 1,97E+07 |
| 33,4 | | 0,1273064 | 0,0930825 | 1,06 | 2,02E+08 | 1,89E+08 |
| 39,3 | | 0,1274294 | 0,0931682 | 1,79 | 2,74E+07 | 1,30E+07 |
| 26,6 | | 0,1275094 | 0,0932030 | 0,88 | 2,31E+07 | 2,78E+07 |
| 14,9 | | 0,1275169 | 0,0931826 | 0,61 | 5,88E+06 | 9,25E+06 |
| 31,1 | | 0,1275628 | 0,0931858 | 1,08 | 2,70E+08 | 2,38E+08 |
| 88,8 | | 0,1275747 | 0,0931654 | 0,88 | 3,56E+09 | 3,87E+09 |
| 11,5 | | 0,1278770 | 0,0934306 | 1,49 | 1,20E+07 | 1,18E+07 |
| 18,7 | | 0,1279793 | 0,0934793 | 1,12 | 5,87E+07 | 6,93E+07 |
| 40,9 | | 0,1282187 | 0,0936891 | 1,16 | 1,60E+08 | 1,34E+08 |
| 37,9 | | 0,1284563 | 0,0937903 | 0,86 | 2,52E+08 | 2,73E+08 |
| 72,9 | | 0,1285792 | 0,0938862 | 0,84 | 1,98E+09 | 2,65E+09 |
| 17,5 | | 0,1288068 | 0,0941553 | 1,18 | 6,55E+07 | 5,36E+07 |
| 22,7 | | 0,1288624 | 0,0941347 | 0,89 | 7,69E+07 | 6,33E+07 |
| 3,1 | | 0,1289055 | 0,0941378 | 0,61 | 8,27E+06 | 9,29E+06 |
| 0,2 | | 0,1289082 | 0,0941172 | 0,20 | 1,66E+07 | 1,41E+07 |
| 15,1 | | 0,1291485 | 0,0944630 | 1,22 | 1,24E+08 | 1,39E+08 |
| 29 | | 0,1292552 | 0,0945376 | 1,09 | 1,91E+08 | 1,64E+08 |
| 39,7 | | 0,1293387 | 0,0945405 | 0,78 | 7,41E+07 | 6,26E+07 |
| 17,7 | | 0,1294846 | 0,0945671 | 0,84 | 4,34E+07 | 5,11E+07 |
| 28,2 | | 0,1295445 | 0,0946207 | 0,84 | 1,07E+08 | 9,80E+07 |
| 55,9 | | 0,1298546 | 0,0948760 | 0,67 | 4,64E+07 | 3,22E+07 |
| 46,9 | | 0,1299117 | 0,0949094 | 1,41 | 5,80E+07 | 6,31E+07 |
| 44,6 | | 0,1299347 | 0,0948887 | 1,08 | 4,92E+08 | 5,10E+08 |
| 56,6 | | 0,1299876 | 0,0948916 | 0,91 | 3,42E+08 | 3,86E+08 |
| 51,9 | | 0,1300253 | 0,0948709 | 0,65 | 5,86E+07 | 4,92E+07 |
| 55,6 | | 0,1301873 | 0,0949679 | 0,80 | 3,63E+08 | 5,05E+08 |
| 21 | | 0,1302937 | 0,0950397 | 0,81 | 1,33E+08 | 1,51E+08 |
| 67,5 | | 0,1307253 | 0,0954384 | 1,42 | 2,01E+07 | 2,46E+07 |
| 51,9 | | 0,1309151 | 0,0956539 | 1,23 | 3,75E+08 | 3,92E+08 |
| 43,7 | | 0,1310721 | 0,0957368 | 1,12 | 4,37E+08 | 4,46E+08 |
| 21,8 | | 0,1316952 | 0,0962318 | 0,69 | 1,03E+08 | 8,76E+07 |
| 6,1 | | 0,1317004 | 0,0962108 | 1,17 | 6,20E+06 | 7,54E+06 |
| 22,5 | | 0,1318536 | 0,0963267 | 1,19 | 2,60E+07 | 2,35E+07 |
| 39,9 | | 0,1318700 | 0,0963292 | 0,91 | 3,15E+08 | 3,46E+08 |
| 28 | | 0,1319698 | 0,0964728 | 0,81 | 6,16E+07 | 6,32E+07 |
| 34 | | 0,1322227 | 0,0966738 | 1,11 | 3,42E+08 | 3,51E+08 |
| 38,9 | | 0,1322392 | 0,0966527 | 1,15 | 6,93E+07 | 7,72E+07 |

| | | | | | | |
|---|---|---|---|---|---|---|
| 40,2 | | 0,1326595 | 0,0968884 | 0,55 | 5,09E+08 | 1,98E+08 |
| 26,3 | | 0,1327329 | 0,0969578 | 0,24 | 3,90E+07 | 4,35E+07 |
| 19,7 | | 0,1328436 | 0,0970124 | 1,17 | 5,68E+06 | 6,60E+06 |
| 25,2 | | 0,1331214 | 0,0973757 | 1,38 | 2,50E+07 | 2,70E+07 |
| 56,7 | | 0,1332239 | 0,0974510 | 1,06 | 6,13E+08 | 6,81E+08 |
| 14,5 | | 0,1332651 | 0,0974298 | 0,91 | 1,32E+08 | 1,22E+08 |
| 37,7 | | 0,1334033 | 0,0974938 | 1,14 | 3,10E+08 | 3,08E+08 |
| 30,1 | | 0,1335776 | 0,0977715 | 0,92 | 1,09E+08 | 1,10E+08 |
| 9,7 | | 0,1338414 | 0,0980326 | 1,76 | 3,42E+06 | 1,73E+06 |
| 19 | | 0,1338809 | 0,0980851 | 0,92 | 1,03E+08 | 9,21E+07 |
| 4,6 | | 0,1339632 | 0,0980907 | 0,92 | 1,65E+07 | 1,59E+07 |
| 10 | | 0,1340114 | 0,0981745 | 0,86 | 7,36E+06 | 8,74E+06 |
| 42,6 | | 0,1340818 | 0,0981966 | 1,40 | 5,86E+07 | 3,25E+07 |
| 12,4 | | 0,1343770 | 0,0984564 | 1,49 | 3,68E+06 | 2,91E+06 |
| 6,2 | | 0,1344129 | 0,0984350 | 0,59 | 3,83E+06 | 6,80E+06 |
| 3,8 | | 0,1345721 | 0,0985924 | 0,73 | 3,37E+06 | 2,88E+06 |
| 43,2 | | 0,1353715 | 0,0992950 | 0,87 | 5,24E+07 | 5,42E+07 |
| 18,3 | | 0,1354901 | 0,0994417 | 1,44 | 5,26E+07 | 3,94E+07 |
| 37,8 | | 0,1355456 | 0,0994869 | 0,93 | 7,49E+07 | 7,73E+07 |
| 60,9 | | 0,1356080 | 0,0995615 | 1,17 | 1,13E+09 | 9,16E+08 |
| 63,6 | | 0,1357554 | 0,0996023 | 2,80 | 1,20E+07 | 3,24E+06 |
| 34,9 | | 0,1359179 | 0,0997523 | 0,84 | 2,02E+08 | 2,57E+08 |
| 13,1 | | 0,1362110 | 0,1000883 | 1,18 | 1,35E+08 | 1,34E+08 |
| 0,5 | | 0,1362555 | 0,1000900 | 1,56 | 3,17E+07 | 3,86E+07 |
| 5,7 | | 0,1362887 | 0,1000952 | 0,41 | 1,18E+07 | 3,52E+06 |
| 21,7 | | 0,1363318 | 0,1001151 | 1,08 | 8,77E+07 | 8,09E+07 |
| 41,5 | | 0,1364093 | 0,1001583 | 1,08 | 7,97E+07 | 8,77E+07 |
| 69,4 | | 0,1364232 | 0,1001574 | 1,09 | 3,06E+09 | 2,61E+09 |
| 12,6 | | 0,1366418 | 0,1003468 | 1,42 | 4,50E+07 | 4,22E+07 |
| 29,7 | | 0,1367432 | 0,1004609 | 0,86 | 7,61E+07 | 9,17E+07 |
| 5,5 | | 0,1367727 | 0,1004392 | 0,55 | 2,32E+07 | 8,29E+06 |
| 19,6 | | 0,1368049 | 0,1004840 | 0,87 | 4,09E+07 | 3,79E+07 |
| 39,5 | | 0,1371085 | 0,1008270 | 1,24 | 2,58E+08 | 2,82E+08 |
| 4 | | 0,1371179 | 0,1008320 | 1,76 | 1,92E+06 | 4,71E+06 |
| 33,9 | | 0,1372019 | 0,1010270 | 0,89 | 1,22E+08 | 1,19E+08 |
| 27,6 | | 0,1372081 | 0,1010518 | 0,89 | 4,83E+07 | 5,93E+07 |
| 48,5 | | 0,1375262 | 0,1014548 | 0,91 | 3,15E+09 | 3,54E+09 |
| 34,3 | | 0,1375782 | 0,1014562 | 0,90 | 1,15E+08 | 1,38E+08 |
| 76,6 | | 0,1375887 | 0,1014611 | 1,06 | 2,75E+08 | 2,59E+08 |
| 51,5 | | 0,1376301 | 0,1014590 | 1,14 | 4,65E+08 | 3,96E+08 |
| 39,3 | | 0,1377082 | 0,1015079 | 0,91 | 3,05E+08 | 3,12E+08 |
| 61,2 | | 0,1378595 | 0,1016007 | 0,87 | 7,56E+07 | 6,63E+07 |
| 32,2 | | 0,1379374 | 0,1016710 | 1,16 | 4,69E+07 | 3,97E+07 |
| 43,8 | | 0,1380893 | 0,1017569 | 0,86 | 5,42E+08 | 4,73E+08 |
| 25,8 | | 0,1381484 | 0,1017350 | 0,71 | 1,69E+07 | 1,70E+07 |
| 29,7 | | 0,1382075 | 0,1018173 | 1,32 | 1,97E+07 | 2,54E+07 |
| 7,9 | | 0,1382913 | 0,1019203 | 1,48 | 4,73E+06 | 7,69E+06 |
| 46,1 | | 0,1385242 | 0,1022369 | 0,87 | 1,79E+08 | 1,96E+08 |
| 56,9 | | 0,1390864 | 0,1029985 | 0,89 | 4,17E+08 | 4,66E+08 |
| 21,3 | | 0,1391991 | 0,1030771 | 0,86 | 1,96E+08 | 1,80E+08 |

| | | | | | | |
|---|---|---|---|---|---|---|
| 13,7 | | 0,1392483 | 0,1031530 | 0,51 | 6,43E+06 | 5,06E+06 |
| 58,7 | | 0,1393098 | 0,1032014 | 0,93 | 2,56E+09 | 2,67E+09 |
| 25,1 | | 0,1393768 | 0,1032463 | 0,89 | 7,10E+07 | 7,41E+07 |
| 19,5 | | 0,1393817 | 0,1032241 | 0,77 | 1,28E+07 | 1,19E+07 |
| 45,3 | | 0,1394092 | 0,1032019 | 0,87 | 1,69E+08 | 1,69E+08 |
| 23,2 | | 0,1395538 | 0,1033104 | 0,86 | 8,14E+07 | 8,29E+07 |
| 13,4 | | 0,1395889 | 0,1033415 | 1,21 | 2,17E+07 | 2,55E+07 |
| 5,6 | | 0,1400070 | 0,1036906 | 1,90 | 4,50E+06 | 1,48E+06 |
| 38,4 | | 0,1401148 | 0,1038015 | 0,85 | 1,62E+08 | 1,27E+08 |
| 32,3 | | 0,1402984 | 0,1040163 | 1,30 | 1,29E+08 | 1,69E+08 |
| 7,5 | | 0,1403288 | 0,1039940 | 0,91 | 2,06E+07 | 1,99E+07 |
| 51 | | 0,1405220 | 0,1041640 | 1,51 | 7,62E+08 | 6,08E+08 |
| 31 | | 0,1406782 | 0,1043803 | 0,85 | 2,31E+07 | 2,23E+07 |
| 34,8 | | 0,1407716 | 0,1044489 | 1,10 | 9,86E+07 | 8,66E+07 |
| 29,6 | + | 0,1409043 | 0,1046050 | 5,64 | 3,08E+07 | 6,56E+06 |
| 51,1 | | 0,1411907 | 0,1047302 | 0,89 | 2,36E+08 | 2,78E+08 |
| 21,1 | | 0,1412150 | 0,1047077 | 0,88 | 3,99E+07 | 3,86E+07 |
| 7,8 | | 0,1414525 | 0,1049477 | 1,82 | 1,83E+07 | 3,13E+07 |
| 60,5 | | 0,1416491 | 0,1051884 | 0,84 | 1,41E+09 | 1,79E+09 |
| 21,8 | | 0,1417313 | 0,1053811 | 1,11 | 7,40E+07 | 6,75E+07 |
| 6,2 | | 0,1418357 | 0,1055359 | 0,55 | 5,84E+06 | 5,69E+06 |
| 15,8 | | 0,1418447 | 0,1055133 | 0,32 | 4,74E+06 | 1,03E+07 |
| 34,2 | | 0,1423322 | 0,1059045 | 1,17 | 3,04E+08 | 3,23E+08 |
| 41,6 | | 0,1425417 | 0,1060899 | 1,55 | 1,18E+08 | 2,43E+08 |
| 32,9 | | 0,1426438 | 0,1061349 | 0,78 | 5,38E+07 | 4,15E+07 |
| 12,5 | | 0,1427041 | 0,1061652 | 0,85 | 3,46E+07 | 3,82E+07 |
| 25,8 | | 0,1427226 | 0,1061656 | 0,89 | 1,51E+08 | 1,73E+08 |
| 40,8 | | 0,1429937 | 0,1064767 | 1,11 | 4,02E+07 | 3,75E+07 |
| 22,4 | | 0,1432457 | 0,1068338 | 0,86 | 1,71E+08 | 1,36E+08 |
| 16,4 | | 0,1435803 | 0,1071497 | 0,73 | 3,40E+07 | 3,47E+07 |
| 50,9 | | 0,1436203 | 0,1071704 | 1,10 | 2,56E+08 | 2,22E+08 |
| 14,9 | | 0,1438160 | 0,1073442 | 3,46 | 3,16E+06 | 2,47E+07 |
| 20,3 | | 0,1438953 | 0,1074264 | 1,21 | 4,47E+07 | 3,13E+07 |
| 5,7 | | 0,1441199 | 0,1075573 | 0,61 | 3,67E+06 | 2,39E+06 |
| 17,8 | | 0,1441875 | 0,1075539 | 1,40 | 1,09E+07 | 1,18E+07 |
| 12,2 | | 0,1442537 | 0,1075575 | 1,42 | 3,97E+07 | 5,79E+07 |
| 27,5 | | 0,1445631 | 0,1077070 | 1,22 | 1,00E+08 | 6,97E+07 |
| 21,1 | | 0,1446271 | 0,1077319 | 0,91 | 6,54E+07 | 6,04E+07 |
| 29,9 | | 0,1447139 | 0,1077959 | 1,13 | 9,62E+07 | 8,49E+07 |
| 39,9 | | 0,1447858 | 0,1079513 | 0,89 | 1,36E+08 | 1,37E+08 |
| 16,7 | | 0,1447986 | 0,1079283 | 1,86 | 3,92E+07 | 3,28E+07 |
| 15,4 | | 0,1448828 | 0,1079957 | 1,62 | 5,65E+06 | 8,72E+06 |
| 3,9 | | 0,1453913 | 0,1086952 | 0,06 | 4,22E+06 | 8,18E+08 |
| 26,8 | | 0,1454101 | 0,1086721 | 1,17 | 1,48E+08 | 1,31E+08 |
| 46,5 | | 0,1458438 | 0,1091716 | 1,13 | 1,29E+08 | 1,14E+08 |
| 18 | | 0,1458893 | 0,1091918 | 0,94 | 1,65E+08 | 1,67E+08 |
| 38,2 | | 0,1459731 | 0,1092120 | 1,23 | 4,82E+07 | 5,53E+07 |
| 37,6 | | 0,1460155 | 0,1092160 | 1,18 | 2,56E+07 | 3,33E+07 |
| 20,9 | | 0,1461241 | 0,1094288 | 0,78 | 6,12E+07 | 5,69E+07 |
| 53,5 | | 0,1461393 | 0,1094319 | 0,69 | 2,03E+07 | 2,09E+07 |

| | | | | | | |
|---|---|---|---|---|---|---|
| 10,3 | | 0,1461440 | 0,1094086 | 0,68 | 1,96E+07 | 1,33E+07 |
| 27,4 | | 0,1462433 | 0,1094985 | 0,88 | 1,63E+08 | 1,98E+08 |
| 11,4 | | 0,1462804 | 0,1095280 | 3,09 | 7,60E+07 | 7,03E+07 |
| 73,4 | | 0,1464232 | 0,1096298 | 1,14 | 3,03E+08 | 3,42E+08 |
| 28 | | 0,1467121 | 0,1097230 | 0,75 | 3,58E+07 | 3,20E+07 |
| 42,2 | | 0,1469024 | 0,1098605 | 0,78 | 1,82E+08 | 1,35E+08 |
| 9,5 | | 0,1471136 | 0,1099758 | 2,72 | 1,37E+07 | 3,40E+06 |
| 20,7 | | 0,1474956 | 0,1104966 | 0,88 | 1,05E+08 | 9,55E+07 |
| 17,9 | | 0,1475121 | 0,1104731 | 0,85 | 4,34E+07 | 5,56E+07 |
| 21,3 | | 0,1476440 | 0,1106162 | 1,12 | 3,82E+07 | 3,76E+07 |
| 45,3 | | 0,1476526 | 0,1106191 | 0,75 | 9,64E+07 | 1,03E+08 |
| 38,4 | | 0,1480244 | 0,1109048 | 1,12 | 2,32E+08 | 2,11E+08 |
| 24 | | 0,1480460 | 0,1108813 | 1,81 | 1,10E+07 | 8,14E+06 |
| 41,6 | | 0,1480717 | 0,1109036 | 0,87 | 3,89E+08 | 4,45E+08 |
| 31 | | 0,1480847 | 0,1108801 | 2,78 | 2,25E+06 | 7,67E+06 |
| 23,2 | | 0,1481684 | 0,1109100 | 0,68 | 2,54E+07 | 3,49E+07 |
| 14 | | 0,1482792 | 0,1109493 | 1,12 | 2,80E+07 | 3,03E+07 |
| 12,1 | | 0,1483834 | 0,1110836 | 1,26 | 1,37E+07 | 1,12E+07 |
| 7,6 | | 0,1485037 | 0,1110829 | 0,60 | 1,26E+07 | 1,68E+07 |
| 38,6 | | 0,1486046 | 0,1112061 | 1,14 | 6,02E+08 | 6,89E+08 |
| 17,5 | | 0,1487237 | 0,1112512 | 1,13 | 5,59E+07 | 6,16E+07 |
| 22 | | 0,1489593 | 0,1114006 | 1,18 | 1,72E+07 | 1,60E+07 |
| 34,7 | | 0,1492957 | 0,1116982 | 1,27 | 1,69E+08 | 1,17E+08 |
| 9,7 | | 0,1493373 | 0,1116949 | 0,73 | 4,60E+06 | 3,85E+06 |
| 6,4 | | 0,1493404 | 0,1116713 | 0,63 | 6,46E+06 | 3,35E+06 |
| 21,3 | | 0,1494459 | 0,1117916 | 0,78 | 3,24E+07 | 4,22E+07 |
| 30,4 | | 0,1495410 | 0,1117950 | 0,86 | 1,40E+08 | 1,05E+08 |
| 21 | | 0,1495421 | 0,1117714 | 0,87 | 3,87E+07 | 4,17E+07 |
| 56,5 | | 0,1496523 | 0,1119628 | 1,08 | 5,02E+07 | 5,35E+07 |
| 17,1 | | 0,1499094 | 0,1121896 | 0,82 | 2,10E+07 | 1,94E+07 |
| 13,1 | | 0,1504360 | 0,1126575 | 0,67 | 1,20E+07 | 6,82E+06 |
| 59,6 | | 0,1504609 | 0,1126861 | 0,84 | 3,64E+08 | 4,70E+08 |
| 26,1 | | 0,1505930 | 0,1127689 | 0,74 | 1,42E+07 | 1,77E+07 |
| 30,1 | | 0,1506474 | 0,1127915 | 0,72 | 1,10E+07 | 1,08E+07 |
| 15,4 | | 0,1512256 | 0,1133832 | 0,31 | 2,48E+07 | 2,52E+07 |
| 28,1 | | 0,1512496 | 0,1134066 | 0,88 | 6,17E+07 | 5,29E+07 |
| 49,6 | | 0,1515096 | 0,1135703 | 1,06 | 1,18E+09 | 1,23E+09 |
| 48,2 | | 0,1516911 | 0,1137144 | 1,04 | 1,34E+09 | 1,27E+09 |
| 19,3 | | 0,1517563 | 0,1137892 | 0,66 | 2,66E+07 | 3,51E+07 |
| 21,6 | | 0,1522564 | 0,1143750 | 1,12 | 1,76E+07 | 1,55E+07 |
| 4,9 | | 0,1525592 | 0,1147013 | 1,97 | 1,54E+06 | 4,84E+05 |
| 28,5 | | 0,1526519 | 0,1147505 | 0,93 | 7,78E+08 | 7,30E+08 |
| 13 | | 0,1526835 | 0,1147719 | 0,68 | 1,71E+07 | 2,00E+07 |
| 19 | | 0,1527969 | 0,1149114 | 1,37 | 4,31E+06 | 3,25E+06 |
| 14,5 | | 0,1528029 | 0,1148872 | 1,22 | 4,34E+06 | 4,41E+06 |
| 8,9 | | 0,1529143 | 0,1148891 | 1,82 | 2,80E+06 | 2,94E+06 |
| 9,6 | | 0,1530790 | 0,1150361 | 0,80 | 3,84E+06 | 4,71E+06 |
| 20,4 | | 0,1530889 | 0,1150118 | 1,62 | 7,72E+06 | 6,35E+06 |
| 18,4 | | 0,1533582 | 0,1151326 | 0,39 | 2,21E+07 | 2,53E+07 |
| 44,4 | | 0,1537564 | 0,1156022 | 0,85 | 8,92E+07 | 9,07E+07 |

| | | | | | | |
|---|---|---|---|---|---|---|
| 40,5 | | 0,1537706 | 0,1156040 | 1,73 | 2,01E+09 | 2,18E+09 |
| 10 | | 0,1541991 | 0,1159933 | 0,54 | 1,83E+07 | 8,84E+06 |
| 7,4 | | 0,1542408 | 0,1159688 | 1,65 | 3,24E+06 | 3,99E+06 |
| 65,1 | | 0,1544215 | 0,1160160 | 0,85 | 1,21E+08 | 1,31E+08 |
| 24,9 | | 0,1544593 | 0,1160185 | 0,75 | 2,15E+07 | 2,04E+07 |
| 35,7 | | 0,1549352 | 0,1163670 | 0,89 | 1,38E+08 | 1,40E+08 |
| 60,1 | | 0,1549497 | 0,1163425 | 0,85 | 2,30E+08 | 1,95E+08 |
| 9,6 | | 0,1549562 | 0,1163441 | 0,48 | 3,77E+06 | 5,21E+06 |
| 5,2 | | 0,1549995 | 0,1163424 | 2,07 | 2,88E+06 | 1,22E+06 |
| 44,2 | | 0,1553762 | 0,1167267 | 1,11 | 3,08E+08 | 2,87E+08 |
| 27,8 | | 0,1554956 | 0,1168425 | 0,37 | 1,97E+07 | 1,04E+07 |
| 97,8 | | 0,1555314 | 0,1168382 | 2,71 | 4,39E+08 | 4,85E+08 |
| 35,3 | | 0,1555587 | 0,1168397 | 0,85 | 4,82E+07 | 5,28E+07 |
| 8,2 | | 0,1556694 | 0,1169151 | 2,30 | 1,49E+07 | 3,26E+06 |
| 41,1 | | 0,1557204 | 0,1169880 | 0,92 | 9,20E+08 | 8,27E+08 |
| 7,7 | | 0,1557622 | 0,1169903 | 1,54 | 1,71E+07 | 1,02E+07 |
| 39,3 | | 0,1558092 | 0,1169859 | 0,91 | 2,28E+07 | 2,27E+07 |
| 4,1 | | 0,1559488 | 0,1170730 | 0,50 | 2,34E+06 | 8,51E+06 |
| 25,7 | | 0,1561267 | 0,1171307 | 2,35 | 3,94E+06 | 8,75E+06 |
| 4,1 | | 0,1561795 | 0,1171481 | 0,54 | 3,12E+06 | 2,44E+06 |
| 6,9 | | 0,1563861 | 0,1173996 | 2,65 | 5,17E+06 | 1,59E+06 |
| 18 | | 0,1565361 | 0,1175369 | 1,38 | 6,60E+06 | 4,41E+06 |
| 8 | | 0,1570563 | 0,1180440 | 1,22 | 1,63E+07 | 1,25E+07 |
| 1,1 | | 0,1574663 | 0,1184922 | 1,35 | 1,95E+07 | 2,90E+07 |
| 35 | | 0,1576052 | 0,1185077 | 1,34 | 1,33E+07 | 1,07E+07 |
| 49,5 | | 0,1579589 | 0,1188508 | 0,89 | 1,27E+09 | 1,31E+09 |
| 30,5 | | 0,1580280 | 0,1189139 | 1,12 | 8,63E+07 | 7,37E+07 |
| 25,7 | | 0,1583184 | 0,1191579 | 1,13 | 1,77E+09 | 1,63E+09 |
| 50,3 | | 0,1583232 | 0,1191330 | 1,11 | 3,15E+08 | 2,93E+08 |
| 27,7 | | 0,1586897 | 0,1193467 | 2,55 | 4,30E+06 | 2,57E+07 |
| 41,3 | | 0,1587246 | 0,1193418 | 0,90 | 9,00E+08 | 9,09E+08 |
| 25,6 | | 0,1587612 | 0,1193428 | 0,39 | 8,29E+06 | 5,67E+06 |
| 34,2 | | 0,1589805 | 0,1195865 | 1,09 | 1,12E+09 | 1,20E+09 |
| 26,9 | | 0,1589946 | 0,1196142 | 1,15 | 7,28E+07 | 7,97E+07 |
| 3,7 | | 0,1590680 | 0,1196804 | 0,33 | 8,38E+06 | 3,00E+06 |
| 5,3 | | 0,1592700 | 0,1198812 | 1,36 | 2,84E+06 | 3,07E+06 |
| 10,9 | | 0,1596886 | 0,1202275 | 1,20 | 4,80E+07 | 3,49E+07 |
| 27,7 | | 0,1599405 | 0,1205042 | 1,10 | 1,28E+08 | 1,40E+08 |
| 37,4 | | 0,1599872 | 0,1205049 | 1,08 | 1,12E+08 | 1,26E+08 |
| 36 | | 0,1600040 | 0,1204797 | 0,90 | 1,19E+08 | 1,35E+08 |
| 35,4 | | 0,1600041 | 0,1204546 | 1,11 | 1,50E+08 | 1,47E+08 |
| 24,1 | | 0,1600585 | 0,1204553 | 0,84 | 4,41E+07 | 4,79E+07 |
| 2,9 | | 0,1600592 | 0,1204302 | 1,88 | 6,47E+07 | 7,28E+07 |
| 53,1 | | 0,1601107 | 0,1205027 | 1,04 | 1,44E+09 | 1,48E+09 |
| 2,6 | | 0,1601246 | 0,1204776 | 1,46 | 4,36E+06 | 4,60E+06 |
| 45,8 | | 0,1602799 | 0,1207487 | 1,06 | 2,63E+08 | 2,76E+08 |
| 61,9 | | 0,1603142 | 0,1207494 | 1,11 | 7,56E+08 | 8,24E+08 |
| 13,1 | | 0,1605922 | 0,1210163 | 0,80 | 1,29E+07 | 1,28E+07 |
| 84,4 | | 0,1609572 | 0,1213122 | 0,85 | 1,28E+08 | 1,66E+08 |
| 7,1 | | 0,1611126 | 0,1214003 | 0,86 | 1,14E+07 | 1,27E+07 |

| | | | | | | |
|---|---|---|---|---|---|---|
| 35,4 | | 0,1611562 | 0,1214017 | 0,91 | 3,57E+08 | 3,92E+08 |
| 18,5 | | 0,1612251 | 0,1214898 | 0,87 | 3,90E+07 | 3,70E+07 |
| 1,5 | | 0,1614166 | 0,1216212 | 1,86 | 2,96E+08 | 1,21E+08 |
| 40 | | 0,1615366 | 0,1216933 | 0,68 | 5,14E+07 | 5,50E+07 |
| 68,1 | | 0,1617702 | 0,1218804 | 0,90 | 1,11E+08 | 1,01E+08 |
| 39,8 | | 0,1623100 | 0,1223374 | 0,71 | 4,43E+07 | 5,65E+07 |
| 4,4 | | 0,1624627 | 0,1224760 | 1,77 | 2,75E+06 | 2,25E+06 |
| 3,1 | | 0,1626884 | 0,1227252 | 2,85 | 5,35E+07 | 8,56E+06 |
| 19,7 | | 0,1627099 | 0,1227222 | 0,29 | 1,47E+07 | 1,70E+07 |
| 43,3 | | 0,1627566 | 0,1227391 | 1,24 | 1,79E+08 | 2,50E+08 |
| 34,8 | | 0,1628419 | 0,1227818 | 1,14 | 1,33E+08 | 1,31E+08 |
| 5,5 | | 0,1629169 | 0,1228661 | 1,79 | 6,58E+06 | 6,17E+06 |
| 30,2 | | 0,1629266 | 0,1228663 | 0,89 | 2,40E+08 | 2,50E+08 |
| 4,4 | | 0,1630721 | 0,1229605 | 0,51 | 3,02E+06 | 4,06E+06 |
| 2,1 | | 0,1631429 | 0,1229732 | 0,74 | 4,27E+06 | 3,27E+06 |
| 22,5 | | 0,1632694 | 0,1229476 | 1,40 | 1,57E+07 | 2,92E+07 |
| 61,8 | | 0,1633830 | 0,1230260 | 0,92 | 1,15E+08 | 1,23E+08 |
| 13,3 | | 0,1638036 | 0,1232331 | 0,39 | 1,19E+07 | 7,63E+06 |
| 31,7 | | 0,1638349 | 0,1232075 | 5,32 | 2,44E+06 | 4,48E+07 |
| 26,4 | | 0,1638518 | 0,1232334 | 0,93 | 2,78E+09 | 2,76E+09 |
| 1,6 | | 0,1640523 | 0,1234835 | 1,93 | 8,29E+05 | 1,00E+06 |
| 33,7 | | 0,1648188 | 0,1243778 | 1,12 | 9,34E+08 | 8,74E+08 |
| 28,9 | | 0,1648657 | 0,1244142 | 1,25 | 2,01E+08 | 1,67E+08 |
| 5,5 | | 0,1649436 | 0,1244083 | 0,64 | 4,58E+06 | 7,67E+06 |
| 18,9 | | 0,1653774 | 0,1247791 | 2,00 | 1,00E+07 | 6,88E+06 |
| 50,5 | | 0,1655092 | 0,1249390 | 0,91 | 3,79E+08 | 3,29E+08 |
| 39,9 | | 0,1656365 | 0,1250359 | 1,22 | 5,47E+07 | 3,93E+07 |
| 57,3 | | 0,1658232 | 0,1251625 | 0,93 | 5,88E+08 | 6,37E+08 |
| 60,9 | | 0,1660146 | 0,1252336 | 1,16 | 1,71E+09 | 1,48E+09 |
| 11,1 | | 0,1660234 | 0,1252076 | 0,77 | 1,64E+07 | 1,50E+07 |
| 3 | | 0,1661719 | 0,1255380 | 2,07 | 4,48E+06 | 5,38E+06 |
| 52,2 | | 0,1669802 | 0,1262403 | 1,10 | 7,34E+08 | 7,75E+08 |
| 13 | | 0,1670117 | 0,1262340 | 0,91 | 7,59E+06 | 6,79E+06 |
| 69,2 | | 0,1670128 | 0,1262079 | 1,18 | 1,75E+09 | 1,34E+09 |
| 53,1 | | 0,1670909 | 0,1263266 | 1,05 | 6,73E+09 | 6,56E+09 |
| 45,1 | | 0,1672825 | 0,1265389 | 1,13 | 1,74E+08 | 2,05E+08 |
| 31,2 | | 0,1672888 | 0,1265127 | 1,12 | 2,18E+08 | 1,77E+08 |
| 3,2 | | 0,1673213 | 0,1265089 | 1,46 | 1,58E+06 | 2,64E+06 |
| 6,9 | | 0,1673894 | 0,1265084 | 2,02 | 1,89E+06 | 5,75E+05 |
| 15 | | 0,1674113 | 0,1265012 | 1,14 | 1,44E+07 | 1,64E+07 |
| 26,6 | | 0,1674176 | 0,1264751 | 0,88 | 6,42E+07 | 6,83E+07 |
| 4,1 | | 0,1675548 | 0,1265688 | 1,29 | 7,62E+06 | 9,42E+06 |
| 32,1 | | 0,1676009 | 0,1265625 | 0,73 | 5,35E+07 | 3,98E+07 |
| 45,8 | | 0,1676719 | 0,1266223 | 0,88 | 6,10E+08 | 5,94E+08 |
| 8,7 | | 0,1678052 | 0,1267589 | 2,34 | 1,03E+07 | 3,28E+06 |
| 38,5 | | 0,1681316 | 0,1270624 | 0,88 | 2,59E+08 | 2,72E+08 |
| 2,2 | | 0,1681330 | 0,1270361 | 0,79 | 6,14E+06 | 6,19E+06 |
| 53 | | 0,1681829 | 0,1270801 | 0,92 | 4,80E+09 | 4,53E+09 |
| 28,6 | | 0,1683550 | 0,1273172 | 1,21 | 2,17E+07 | 2,48E+07 |
| 13,5 | | 0,1684441 | 0,1273818 | 0,87 | 3,13E+07 | 4,03E+07 |

| | | | | | | |
|---|---|---|---|---|---|---|
| 2,4 | | 0,1687949 | 0,1277277 | 2,17 | 2,83E+06 | 3,41E+06 |
| 21 | | 0,1688552 | 0,1277904 | 1,40 | 7,15E+06 | 9,27E+06 |
| 43 | | 0,1690206 | 0,1278037 | 1,36 | 2,22E+07 | 3,22E+07 |
| 14,3 | | 0,1691741 | 0,1278870 | 1,09 | 2,69E+07 | 2,53E+07 |
| 9,4 | | 0,1693491 | 0,1281113 | 2,48 | 1,17E+07 | 1,18E+07 |
| 10,2 | | 0,1694529 | 0,1281979 | 1,57 | 5,99E+06 | 3,06E+06 |
| 9,1 | | 0,1700426 | 0,1287344 | 0,78 | 8,36E+06 | 1,35E+07 |
| 17 | | 0,1701927 | 0,1288925 | 0,86 | 3,20E+08 | 2,89E+08 |
| 19,4 | | 0,1702427 | 0,1289434 | 0,23 | 3,09E+07 | 3,61E+07 |
| 17,9 | | 0,1703697 | 0,1292850 | 1,13 | 6,30E+07 | 6,76E+07 |
| 51,1 | | 0,1704375 | 0,1293317 | 1,26 | 6,79E+07 | 5,56E+07 |
| 27,3 | | 0,1706691 | 0,1295019 | 1,49 | 5,98E+07 | 3,94E+07 |
| 2,6 | | 0,1707316 | 0,1295402 | 1,39 | 5,39E+06 | 3,53E+06 |
| 10,2 | | 0,1707507 | 0,1295136 | 1,28 | 2,57E+07 | 1,63E+07 |
| 35,8 | | 0,1708437 | 0,1295519 | 0,79 | 1,38E+08 | 1,98E+08 |
| 4,3 | | 0,1710802 | 0,1297474 | 1,69 | 3,08E+06 | 7,60E+06 |
| 30,1 | | 0,1712857 | 0,1299955 | 1,07 | 1,18E+08 | 1,08E+08 |
| 46,2 | | 0,1713293 | 0,1300140 | 1,07 | 1,11E+09 | 1,06E+09 |
| 11,7 | | 0,1713376 | 0,1299873 | 1,31 | 1,39E+07 | 2,20E+07 |
| 23,1 | | 0,1716081 | 0,1303206 | 1,81 | 1,56E+07 | 4,78E+07 |
| 10,3 | | 0,1718066 | 0,1305091 | 1,32 | 4,15E+07 | 2,85E+07 |
| 50,4 | | 0,1718672 | 0,1305078 | 1,17 | 6,54E+07 | 6,75E+07 |
| 30,5 | | 0,1719980 | 0,1306535 | 1,32 | 6,48E+07 | 8,78E+07 |
| 22,8 | | 0,1720324 | 0,1306710 | 0,78 | 2,76E+07 | 1,88E+07 |
| 9,6 | | 0,1720769 | 0,1307148 | 0,80 | 8,47E+06 | 1,17E+07 |
| 8,1 | | 0,1721690 | 0,1307323 | 0,47 | 7,22E+06 | 7,10E+06 |
| 29,3 | | 0,1722119 | 0,1307892 | 1,18 | 8,61E+07 | 7,38E+07 |
| 15,6 | | 0,1724308 | 0,1310242 | 0,85 | 3,28E+07 | 4,04E+07 |
| 21,2 | | 0,1727037 | 0,1312681 | 0,53 | 4,85E+07 | 1,39E+07 |
| 6,4 | | 0,1727311 | 0,1312633 | 0,74 | 8,25E+06 | 8,92E+06 |
| 66,1 | | 0,1729020 | 0,1314349 | 1,10 | 1,20E+09 | 1,21E+09 |
| 25 | | 0,1732189 | 0,1316687 | 1,14 | 5,78E+07 | 5,41E+07 |
| 15,4 | | 0,1734018 | 0,1318180 | 0,25 | 4,14E+07 | 5,02E+07 |
| 11,9 | | 0,1734156 | 0,1318098 | 1,13 | 1,83E+07 | 1,74E+07 |
| 7,4 | | 0,1736641 | 0,1320606 | 0,80 | 6,72E+06 | 7,71E+06 |
| 10,1 | | 0,1737660 | 0,1320967 | 0,85 | 6,23E+06 | 4,71E+06 |
| 31,7 | | 0,1738777 | 0,1321655 | 1,19 | 1,70E+08 | 1,81E+08 |
| 45,1 | | 0,1740541 | 0,1323767 | 1,10 | 1,59E+08 | 1,38E+08 |
| 17,3 | | 0,1741886 | 0,1324397 | 0,82 | 1,30E+07 | 1,44E+07 |
| 63,2 | | 0,1743832 | 0,1325976 | 0,67 | 2,62E+07 | 5,29E+07 |
| 29,8 | | 0,1746316 | 0,1326654 | 1,42 | 9,76E+06 | 1,15E+07 |
| 28,7 | | 0,1746520 | 0,1326579 | 0,83 | 3,66E+07 | 2,87E+07 |
| 23,3 | | 0,1747272 | 0,1326979 | 0,88 | 3,00E+07 | 3,26E+07 |
| 71,5 | | 0,1749070 | 0,1327722 | 0,90 | 4,09E+08 | 3,77E+08 |
| 26,3 | | 0,1749979 | 0,1328955 | 1,30 | 4,19E+07 | 4,41E+07 |
| 18,9 | | 0,1751732 | 0,1331169 | 0,64 | 8,49E+07 | 8,67E+07 |
| 12 | | 0,1756894 | 0,1335026 | 0,85 | 3,71E+07 | 3,39E+07 |
| 10,3 | | 0,1759748 | 0,1339117 | 0,75 | 6,75E+06 | 8,86E+06 |
| 55,4 | | 0,1759814 | 0,1338844 | 0,92 | 2,74E+09 | 2,70E+09 |
| 36,9 | | 0,1764091 | 0,1344665 | 1,20 | 2,11E+08 | 1,77E+08 |

| | | | | | | |
|---|---|---|---|---|---|---|
| 8,4 | | 0,1764194 | 0,1344390 | 1,34 | 7,95E+06 | 1,07E+07 |
| 4,8 | | 0,1768950 | 0,1348363 | 0,67 | 4,06E+06 | 3,00E+06 |
| 23,5 | | 0,1772155 | 0,1351606 | 1,26 | 2,90E+07 | 2,38E+07 |
| 62,1 | | 0,1776272 | 0,1355649 | 0,81 | 1,46E+07 | 1,11E+07 |
| 62,6 | | 0,1780237 | 0,1358817 | 0,93 | 9,06E+08 | 1,03E+09 |
| 21 | | 0,1780732 | 0,1358760 | 0,82 | 6,96E+07 | 8,84E+07 |
| 39,5 | | 0,1781848 | 0,1359853 | 1,11 | 2,23E+08 | 2,16E+08 |
| 10,6 | | 0,1783452 | 0,1361207 | 0,93 | 1,39E+08 | 1,26E+08 |
| 75,5 | | 0,1784367 | 0,1361851 | 1,21 | 7,20E+09 | 6,03E+09 |
| 13,6 | | 0,1786205 | 0,1363612 | 1,12 | 1,97E+07 | 1,96E+07 |
| 33,9 | | 0,1790868 | 0,1367092 | 0,81 | 1,46E+09 | 1,11E+09 |
| 3,8 | | 0,1791734 | 0,1368346 | 0,45 | 5,55E+06 | 3,66E+06 |
| 52,5 | | 0,1793412 | 0,1369534 | 0,90 | 3,55E+08 | 3,94E+08 |
| 22,9 | | 0,1796423 | 0,1374134 | 0,83 | 2,34E+08 | 1,77E+08 |
| 56,2 | | 0,1797039 | 0,1373855 | 1,48 | 2,98E+08 | 5,97E+08 |
| 1,8 | | 0,1797594 | 0,1373966 | 0,54 | 1,18E+07 | 2,47E+07 |
| 28 | | 0,1797833 | 0,1373906 | 1,42 | 1,02E+07 | 2,06E+07 |
| 8,2 | | 0,1800276 | 0,1376142 | 2,00 | 1,92E+06 | 6,17E+06 |
| 58,4 | | 0,1800779 | 0,1376057 | 1,10 | 2,51E+09 | 2,72E+09 |
| 41,6 | | 0,1802300 | 0,1377266 | 1,23 | 6,35E+07 | 7,29E+07 |
| 65,1 | | 0,1805492 | 0,1379687 | 1,11 | 3,75E+09 | 4,65E+09 |
| 19,6 | | 0,1805949 | 0,1380041 | 0,70 | 2,27E+07 | 2,72E+07 |
| 41,3 | | 0,1806731 | 0,1380175 | 1,13 | 3,33E+08 | 2,79E+08 |
| 31,1 | | 0,1807192 | 0,1380333 | 0,96 | 3,04E+08 | 2,82E+08 |
| 6,9 | | 0,1812728 | 0,1385970 | 0,40 | 2,29E+06 | 7,73E+06 |
| 11,6 | | 0,1813883 | 0,1386201 | 2,68 | 4,70E+06 | 3,13E+06 |
| 52,7 | | 0,1814748 | 0,1386334 | 1,16 | 3,69E+08 | 3,84E+08 |
| 2,8 | | 0,1822152 | 0,1393745 | 1,78 | 4,44E+06 | 4,93E+06 |
| 11,5 | | 0,1822224 | 0,1393462 | 0,81 | 3,19E+07 | 3,60E+07 |
| 7,6 | | 0,1822932 | 0,1394064 | 1,19 | 4,93E+07 | 3,49E+07 |
| 44,9 | | 0,1825158 | 0,1396631 | 1,08 | 7,28E+07 | 8,16E+07 |
| 19,8 | | 0,1825878 | 0,1396826 | 1,53 | 1,11E+07 | 9,28E+06 |
| 44,6 | | 0,1826050 | 0,1396794 | 1,54 | 1,60E+08 | 2,21E+08 |
| 25,9 | | 0,1828469 | 0,1398296 | 1,33 | 7,84E+06 | 9,25E+06 |
| 31,2 | | 0,1829831 | 0,1399286 | 0,71 | 4,89E+07 | 3,12E+07 |
| 58,6 | | 0,1830044 | 0,1399830 | 1,05 | 3,13E+09 | 2,97E+09 |
| 16,2 | | 0,1830248 | 0,1399546 | 0,76 | 1,57E+07 | 1,59E+07 |
| 41,5 | | 0,1830364 | 0,1399457 | 0,93 | 8,51E+07 | 9,02E+07 |
| 36,5 | | 0,1831010 | 0,1399903 | 1,07 | 8,75E+07 | 8,28E+07 |
| 22,8 | | 0,1831104 | 0,1399619 | 2,19 | 9,10E+07 | 1,05E+08 |
| 23,5 | | 0,1831139 | 0,1399336 | 1,17 | 1,97E+07 | 1,73E+07 |
| 12,1 | + | 0,1832734 | 0,1402236 | 4,33 | 3,34E+06 | 2,93E+06 |
| 58,9 | | 0,1835702 | 0,1404373 | 0,70 | 7,49E+07 | 7,38E+07 |
| 36,1 | | 0,1836929 | 0,1404753 | 0,90 | 3,34E+08 | 3,82E+08 |
| 19,1 | | 0,1837372 | 0,1405489 | 1,13 | 3,68E+08 | 4,20E+08 |
| 23,3 | | 0,1837561 | 0,1405399 | 0,85 | 2,08E+08 | 1,63E+08 |
| 10,1 | | 0,1837683 | 0,1405114 | 1,12 | 5,64E+07 | 4,43E+07 |
| 5,4 | | 0,1838113 | 0,1405307 | 0,50 | 6,28E+06 | 3,87E+06 |
| 17,9 | | 0,1838459 | 0,1405023 | 0,73 | 4,41E+07 | 4,42E+07 |
| 26,5 | | 0,1843838 | 0,1412867 | 1,16 | 5,44E+08 | 6,55E+08 |

| | | | | | | |
|---|---|---|---|---|---|---|
| 11,6 | | 0,1845969 | 0,1414959 | 1,32 | 1,11E+08 | 1,51E+08 |
| 25,4 | | 0,1846256 | 0,1414673 | 0,77 | 4,66E+07 | 4,64E+07 |
| 66,7 | | 0,1847090 | 0,1415308 | 1,08 | 5,10E+08 | 5,15E+08 |
| 7,4 | | 0,1848373 | 0,1416097 | 0,56 | 5,73E+06 | 7,33E+06 |
| 15,4 | | 0,1851401 | 0,1419269 | 1,09 | 1,02E+08 | 1,10E+08 |
| 3,3 | | 0,1851477 | 0,1418982 | 0,75 | 4,28E+06 | 3,34E+06 |
| 12,3 | | 0,1851698 | 0,1418890 | 0,74 | 6,55E+06 | 8,78E+06 |
| 27,1 | | 0,1852208 | 0,1418603 | 0,91 | 3,92E+08 | 4,40E+08 |
| 29,9 | | 0,1853226 | 0,1419197 | 1,16 | 5,32E+07 | 4,26E+07 |
| 18,8 | | 0,1853539 | 0,1419096 | 0,87 | 1,29E+08 | 1,11E+08 |
| 20,4 | | 0,1854226 | 0,1419407 | 0,67 | 3,83E+07 | 8,26E+07 |
| 26,6 | | 0,1856828 | 0,1421759 | 0,93 | 7,69E+07 | 8,80E+07 |
| 17,7 | | 0,1860691 | 0,1424957 | 0,64 | 3,03E+07 | 2,58E+07 |
| 32,3 | | 0,1862845 | 0,1427113 | 0,84 | 1,19E+09 | 1,39E+09 |
| 33,3 | | 0,1863462 | 0,1428486 | 1,13 | 4,90E+07 | 4,96E+07 |
| 53,4 | | 0,1864693 | 0,1429609 | 0,89 | 1,79E+09 | 2,06E+09 |
| 38 | | 0,1866517 | 0,1431086 | 0,89 | 2,15E+08 | 1,95E+08 |
| 45,5 | | 0,1867678 | 0,1431797 | 0,72 | 2,55E+07 | 3,16E+07 |
| 55,3 | | 0,1867763 | 0,1431758 | 0,90 | 7,05E+08 | 8,38E+08 |
| 22,5 | | 0,1868370 | 0,1432316 | 1,46 | 1,52E+07 | 3,30E+07 |
| 13,3 | | 0,1873575 | 0,1439171 | 0,75 | 1,64E+07 | 1,36E+07 |
| 57,3 | | 0,1875633 | 0,1440902 | 0,85 | 9,26E+08 | 9,95E+08 |
| 15,7 | | 0,1876019 | 0,1440612 | 1,29 | 1,36E+07 | 1,94E+07 |
| 31,2 | | 0,1877801 | 0,1442093 | 0,89 | 6,05E+07 | 6,65E+07 |
| 7,4 | | 0,1879551 | 0,1445287 | 0,79 | 7,45E+06 | 9,97E+06 |
| 9,4 | | 0,1879815 | 0,1444996 | 1,41 | 8,14E+06 | 6,99E+06 |
| 35,8 | | 0,1880347 | 0,1445148 | 1,27 | 3,36E+08 | 4,39E+08 |
| 22,6 | | 0,1883710 | 0,1448420 | 0,83 | 8,35E+07 | 8,78E+07 |
| 7,3 | | 0,1884643 | 0,1449013 | 0,80 | 9,81E+06 | 8,67E+06 |
| 7 | | 0,1886709 | 0,1450579 | 1,29 | 7,23E+06 | 1,03E+07 |
| 7,7 | | 0,1887141 | 0,1451236 | 0,76 | 8,55E+06 | 7,60E+06 |
| 12,3 | | 0,1889313 | 0,1453347 | 1,17 | 1,14E+07 | 1,15E+07 |
| 6,4 | | 0,1891519 | 0,1456132 | 2,12 | 3,73E+06 | 2,43E+06 |
| 13,4 | | 0,1892728 | 0,1457189 | 0,96 | 1,40E+08 | 1,34E+08 |
| 21 | + | 0,1894608 | 0,1458189 | 0,69 | 3,58E+06 | 2,86E+06 |
| 39,9 | | 0,1895741 | 0,1458362 | 0,91 | 4,78E+08 | 4,27E+08 |
| 44,5 | | 0,1897470 | 0,1460470 | 0,82 | 6,82E+07 | 6,23E+07 |
| 12,8 | | 0,1898459 | 0,1460995 | 0,26 | 6,90E+06 | 5,31E+06 |
| 34,1 | | 0,1899269 | 0,1461721 | 0,92 | 2,01E+08 | 2,38E+08 |
| 22,3 | | 0,1899904 | 0,1462551 | 1,06 | 3,41E+07 | 3,10E+07 |
| 15,3 | | 0,1900476 | 0,1462771 | 1,88 | 1,49E+08 | 1,35E+08 |
| 6,7 | | 0,1904086 | 0,1467330 | 1,44 | 8,09E+06 | 4,45E+06 |
| 60,4 | | 0,1905892 | 0,1467781 | 0,95 | 8,72E+08 | 8,71E+08 |
| 58 | | 0,1907600 | 0,1469026 | 1,13 | 2,04E+08 | 1,98E+08 |
| 21,5 | | 0,1910318 | 0,1472114 | 0,83 | 7,19E+07 | 4,89E+07 |
| 40,5 | | 0,1912221 | 0,1474079 | 0,87 | 4,30E+07 | 5,07E+07 |
| 21,8 | | 0,1912579 | 0,1474472 | 1,04 | 7,57E+07 | 6,97E+07 |
| 9,5 | | 0,1919441 | 0,1481386 | 1,31 | 5,59E+06 | 8,24E+06 |
| 45,4 | | 0,1920601 | 0,1483347 | 0,95 | 1,18E+09 | 1,16E+09 |
| 48 | | 0,1923318 | 0,1485204 | 0,91 | 2,11E+08 | 2,05E+08 |

| | | | | | | |
|---|---|---|---|---|---|---|
| 22 | | 0,1923586 | 0,1485347 | 1,55 | 1,32E+08 | 1,32E+08 |
| 54,8 | | 0,1926309 | 0,1486755 | 0,90 | 1,78E+10 | 1,50E+10 |
| 10,3 | | 0,1928649 | 0,1489594 | 1,25 | 3,19E+07 | 3,35E+07 |
| 11,8 | | 0,1929810 | 0,1490312 | 2,45 | 1,86E+06 | 1,78E+06 |
| 12,5 | | 0,1933235 | 0,1494797 | 1,43 | 2,79E+06 | 1,27E+06 |
| 17,3 | | 0,1933908 | 0,1494498 | 1,77 | 1,74E+06 | 1,91E+06 |
| 40,7 | | 0,1933955 | 0,1494199 | 1,09 | 8,94E+07 | 8,27E+07 |
| 28,5 | | 0,1934545 | 0,1494716 | 0,90 | 4,10E+08 | 3,29E+08 |
| 21,8 | | 0,1937123 | 0,1497287 | 0,60 | 2,28E+07 | 3,29E+07 |
| 8,9 | | 0,1938672 | 0,1498490 | 2,39 | 5,81E+06 | 3,28E+06 |
| 6,5 | | 0,1938779 | 0,1498191 | 1,34 | 5,99E+06 | 9,05E+06 |
| 30,4 | | 0,1940150 | 0,1499936 | 0,88 | 1,35E+08 | 1,04E+08 |
| 57,2 | | 0,1940552 | 0,1500100 | 1,16 | 2,89E+08 | 3,77E+08 |
| 25,8 | | 0,1943885 | 0,1502603 | 0,96 | 9,56E+07 | 1,04E+08 |
| 9,1 | | 0,1944532 | 0,1502950 | 0,62 | 1,34E+07 | 1,27E+07 |
| 36,1 | | 0,1944962 | 0,1503240 | 0,93 | 1,05E+08 | 1,09E+08 |
| 17,3 | | 0,1945050 | 0,1502940 | 1,21 | 4,21E+07 | 3,58E+07 |
| 24,6 | | 0,1945920 | 0,1503550 | 0,81 | 2,95E+07 | 4,32E+07 |
| 95,7 | | 0,1948279 | 0,1506106 | 1,10 | 6,50E+09 | 5,71E+09 |
| 3,2 | | 0,1951464 | 0,1508628 | 0,23 | 1,99E+08 | 1,02E+07 |
| 6,2 | | 0,1952232 | 0,1509699 | 0,60 | 5,59E+06 | 1,61E+07 |
| 32,5 | | 0,1955193 | 0,1513041 | 1,08 | 1,24E+08 | 1,13E+08 |
| 19,6 | | 0,1956107 | 0,1513353 | 1,13 | 2,52E+07 | 2,74E+07 |
| 30,9 | | 0,1957894 | 0,1514892 | 1,35 | 3,02E+07 | 3,25E+07 |
| 35,5 | | 0,1959036 | 0,1515499 | 0,86 | 3,93E+08 | 3,10E+08 |
| 38,4 | | 0,1959916 | 0,1516105 | 1,33 | 1,61E+07 | 2,45E+07 |
| 3,6 | | 0,1960439 | 0,1516982 | 1,30 | 6,28E+06 | 5,16E+06 |
| 17,7 | | 0,1961275 | 0,1516895 | 0,84 | 2,13E+07 | 2,30E+07 |
| 44,4 | | 0,1964204 | 0,1518297 | 0,03 | 8,26E+08 | 1,21E+06 |
| 11,4 | | 0,1964696 | 0,1518432 | 0,76 | 1,33E+07 | 1,31E+07 |
| 42,1 | | 0,1965421 | 0,1519029 | 0,93 | 1,14E+09 | 1,21E+09 |
| 6,6 | | 0,1968222 | 0,1521512 | 0,84 | 1,68E+07 | 1,26E+07 |
| 24,8 | | 0,1970925 | 0,1524510 | 0,83 | 3,16E+07 | 4,00E+07 |
| 48,8 | | 0,1971789 | 0,1525137 | 1,05 | 3,65E+08 | 3,60E+08 |
| 14,7 | | 0,1974201 | 0,1527132 | 0,78 | 1,53E+07 | 1,58E+07 |
| 3,5 | | 0,1976505 | 0,1530270 | 0,60 | 2,52E+06 | 6,66E+06 |
| 46,9 | | 0,1979141 | 0,1532772 | 1,19 | 5,12E+07 | 3,60E+07 |
| 21,1 | | 0,1980292 | 0,1534048 | 0,63 | 4,33E+07 | 4,07E+07 |
| 15,4 | | 0,1981177 | 0,1535738 | 1,25 | 2,12E+07 | 3,33E+07 |
| 57,6 | | 0,1981925 | 0,1536251 | 0,92 | 3,90E+09 | 3,53E+09 |
| 46 | | 0,1983315 | 0,1536772 | 0,81 | 3,38E+08 | 2,34E+08 |
| 14,9 | | 0,1986136 | 0,1539595 | 1,27 | 2,17E+07 | 2,84E+07 |
| 12,9 | | 0,1990640 | 0,1542830 | 1,44 | 7,01E+06 | 7,89E+06 |
| 10,6 | | 0,1993362 | 0,1545484 | 0,35 | 1,41E+07 | 4,10E+06 |
| 37,4 | | 0,1993591 | 0,1545178 | 1,16 | 1,65E+07 | 1,73E+07 |
| 9,6 | | 0,1993939 | 0,1545807 | 0,64 | 2,29E+07 | 1,44E+07 |
| 38,9 | | 0,1995927 | 0,1547174 | 0,83 | 2,02E+07 | 2,70E+07 |
| 2,9 | | 0,1997471 | 0,1547462 | 0,81 | 1,89E+07 | 1,33E+07 |
| 5,6 | | 0,1997825 | 0,1547338 | 0,70 | 6,42E+06 | 3,67E+06 |
| 22,3 | | 0,1998662 | 0,1548387 | 0,70 | 1,58E+07 | 8,40E+06 |

| | | | | | | |
|---|---|---|---|---|---|---|
| 8,5 | | 0,1999293 | 0,1548294 | 1,12 | 1,50E+07 | 1,77E+07 |
| 79,2 | | 0,2003719 | 0,1552544 | 0,91 | 3,05E+08 | 3,61E+08 |
| 37,6 | | 0,2004330 | 0,1552886 | 1,12 | 2,95E+08 | 2,70E+08 |
| 32,6 | | 0,2004782 | 0,1553259 | 0,93 | 8,18E+07 | 9,47E+07 |
| 5,7 | | 0,2005426 | 0,1553815 | 1,25 | 5,46E+06 | 7,01E+06 |
| 7 | | 0,2005991 | 0,1554315 | 0,73 | 6,20E+06 | 5,70E+06 |
| 31,5 | | 0,2006238 | 0,1554190 | 2,84 | 1,76E+06 | 1,42E+07 |
| 66,1 | | 0,2007459 | 0,1554127 | 1,07 | 2,68E+09 | 2,78E+09 |
| 78,4 | | 0,2008521 | 0,1554675 | 0,88 | 7,25E+07 | 7,90E+07 |
| 10,9 | + | 0,2012210 | 0,1558932 | 4,20 | 2,70E+07 | 1,10E+06 |
| 40 | | 0,2014582 | 0,1560593 | 0,88 | 1,40E+08 | 1,25E+08 |
| 33,2 | | 0,2015692 | 0,1562001 | 1,29 | 5,13E+07 | 9,10E+07 |
| 20,4 | | 0,2015773 | 0,1561906 | 0,62 | 5,21E+07 | 6,74E+07 |
| 16,7 | | 0,2016883 | 0,1562427 | 0,37 | 1,55E+07 | 1,77E+07 |
| 40,9 | | 0,2019374 | 0,1565153 | 1,06 | 1,91E+08 | 1,80E+08 |
| 50,6 | | 0,2020281 | 0,1565555 | 1,13 | 3,35E+08 | 4,20E+08 |
| 11,5 | | 0,2020399 | 0,1565246 | 0,40 | 1,12E+07 | 1,17E+07 |
| 4,6 | | 0,2023491 | 0,1570047 | 0,40 | 5,69E+06 | 5,92E+06 |
| 49 | | 0,2026491 | 0,1572920 | 0,95 | 7,71E+07 | 8,46E+07 |
| 6,1 | | 0,2028441 | 0,1574765 | 0,63 | 4,07E+06 | 7,93E+06 |
| 52,8 | | 0,2029085 | 0,1574881 | 0,95 | 1,67E+09 | 1,49E+09 |
| 55,7 | | 0,2029507 | 0,1575012 | 0,92 | 7,47E+08 | 7,29E+08 |
| 7,2 | | 0,2031831 | 0,1577282 | 0,77 | 4,03E+09 | 2,75E+09 |
| 44,3 | | 0,2032279 | 0,1577349 | 0,14 | 2,60E+08 | 5,10E+08 |
| 37,4 | | 0,2035810 | 0,1581195 | 1,17 | 1,26E+09 | 1,76E+09 |
| 39,5 | | 0,2036925 | 0,1581696 | 1,65 | 5,45E+07 | 4,61E+07 |
| 64 | | 0,2038699 | 0,1582401 | 1,10 | 2,93E+08 | 2,50E+08 |
| 28 | | 0,2042980 | 0,1586953 | 0,89 | 4,18E+07 | 5,16E+07 |
| 9,2 | | 0,2043926 | 0,1587255 | 0,52 | 4,07E+06 | 5,92E+06 |
| 17,5 | | 0,2044831 | 0,1587628 | 1,47 | 2,65E+07 | 2,55E+07 |
| 9,4 | | 0,2048318 | 0,1591097 | 1,53 | 6,22E+06 | 4,24E+06 |
| 13,3 | | 0,2048952 | 0,1591430 | 0,83 | 1,16E+07 | 1,36E+07 |
| 59 | | 0,2052090 | 0,1593526 | 1,09 | 1,68E+09 | 1,42E+09 |
| 18,6 | | 0,2054197 | 0,1596362 | 0,88 | 8,39E+07 | 9,00E+07 |
| 28,2 | | 0,2055077 | 0,1596733 | 0,37 | 1,33E+07 | 1,93E+07 |
| 23,1 | | 0,2055377 | 0,1596671 | 1,09 | 6,31E+07 | 5,98E+07 |
| 44,8 | | 0,2058902 | 0,1600787 | 0,89 | 4,50E+08 | 4,14E+08 |
| 28,1 | | 0,2063477 | 0,1604697 | 1,30 | 1,15E+07 | 1,72E+07 |
| 34,3 | | 0,2065374 | 0,1607717 | 1,36 | 4,55E+07 | 5,44E+07 |
| 17 | + | 0,2066738 | 0,1608022 | 0,83 | 1,27E+07 | 1,67E+07 |
| 4,5 | | 0,2069377 | 0,1611142 | 1,81 | 5,61E+06 | 1,66E+06 |
| 12,1 | | 0,2071522 | 0,1611926 | 1,64 | 4,16E+06 | 2,72E+06 |
| 18,7 | | 0,2075293 | 0,1616200 | 1,10 | 4,97E+07 | 4,98E+07 |
| 20,3 | | 0,2075598 | 0,1616071 | 0,29 | 3,07E+07 | 5,54E+06 |
| 47,7 | | 0,2076473 | 0,1616460 | 0,89 | 1,50E+09 | 1,40E+09 |
| 54,9 | | 0,2077167 | 0,1617062 | 1,10 | 3,48E+08 | 4,12E+08 |
| 6,6 | | 0,2078089 | 0,1617852 | 0,81 | 3,67E+06 | 5,61E+06 |
| 6 | | 0,2079643 | 0,1617534 | 1,36 | 3,93E+07 | 2,22E+07 |
| 55,1 | | 0,2081352 | 0,1620122 | 1,18 | 9,46E+07 | 6,95E+07 |
| 14 | | 0,2083150 | 0,1620934 | 1,76 | 2,42E+06 | 5,85E+06 |

| | | | | | | |
|---|---|---|---|---|---|---|
| 32 | | 0,2083184 | 0,1620804 | 0,48 | 3,86E+07 | 4,77E+07 |
| 39,3 | | 0,2085474 | 0,1623578 | 1,17 | 2,99E+07 | 2,55E+07 |
| 34,3 | | 0,2088712 | 0,1627378 | 0,70 | 2,67E+08 | 3,22E+08 |
| 39,4 | | 0,2088764 | 0,1627059 | 1,25 | 2,65E+07 | 4,45E+07 |
| 6,9 | | 0,2090059 | 0,1627665 | 0,74 | 1,40E+07 | 8,88E+06 |
| 11 | | 0,2090428 | 0,1627966 | 0,48 | 6,35E+06 | 4,91E+06 |
| 38,2 | | 0,2091519 | 0,1628752 | 0,69 | 1,50E+08 | 1,51E+08 |
| 21,1 | | 0,2091867 | 0,1628809 | 0,35 | 1,55E+07 | 1,55E+07 |
| 41,2 | | 0,2099946 | 0,1638950 | 0,74 | 1,65E+08 | 9,56E+07 |
| 11,8 | | 0,2106626 | 0,1646957 | 0,70 | 8,09E+06 | 1,15E+07 |
| 45,6 | | 0,2108568 | 0,1649454 | 0,96 | 3,27E+08 | 3,54E+08 |
| 52,4 | | 0,2110983 | 0,1650995 | 0,94 | 2,90E+08 | 3,21E+08 |
| 9,4 | | 0,2111564 | 0,1650671 | 0,65 | 1,62E+07 | 1,43E+07 |
| 60,7 | | 0,2114061 | 0,1652556 | 0,93 | 9,58E+07 | 8,55E+07 |
| 11,5 | | 0,2115362 | 0,1653751 | 0,78 | 1,08E+07 | 1,87E+07 |
| 28,2 | | 0,2116063 | 0,1653920 | 1,40 | 7,49E+06 | 9,46E+06 |
| 43,4 | | 0,2117586 | 0,1656038 | 1,15 | 3,61E+08 | 2,67E+08 |
| 50,7 | | 0,2121694 | 0,1659554 | 1,10 | 5,32E+08 | 5,93E+08 |
| 7 | | 0,2121760 | 0,1659230 | 0,66 | 3,25E+06 | 3,01E+06 |
| 26,9 | | 0,2122148 | 0,1659601 | 0,55 | 2,98E+08 | 4,90E+08 |
| 8,8 | | 0,2124806 | 0,1662216 | 0,53 | 9,16E+06 | 1,05E+07 |
| 11,6 | | 0,2126341 | 0,1663009 | 1,16 | 3,57E+07 | 4,59E+07 |
| 0,9 | | 0,2127989 | 0,1664419 | 0,85 | 2,78E+06 | 3,42E+06 |
| 23,9 | | 0,2129164 | 0,1665141 | 1,70 | 3,56E+06 | 3,04E+06 |
| 6,1 | | 0,2130032 | 0,1665026 | 2,06 | 5,75E+06 | 1,53E+06 |
| 15,9 | | 0,2132853 | 0,1666427 | 0,84 | 3,76E+07 | 3,80E+07 |
| 60,2 | | 0,2135208 | 0,1668928 | 0,96 | 9,70E+08 | 9,09E+08 |
| 54 | | 0,2136313 | 0,1669875 | 0,94 | 2,34E+08 | 2,53E+08 |
| 6,1 | | 0,2140659 | 0,1675028 | 0,84 | 6,33E+06 | 6,15E+06 |
| 29,3 | | 0,2140855 | 0,1674951 | 0,95 | 3,65E+07 | 3,42E+07 |
| 15 | | 0,2141338 | 0,1675319 | 1,44 | 1,73E+06 | 3,66E+06 |
| 32,9 | | 0,2142901 | 0,1677176 | 0,82 | 3,39E+08 | 3,03E+08 |
| 56,2 | | 0,2142903 | 0,1676849 | 0,86 | 5,05E+08 | 5,80E+08 |
| 14,3 | | 0,2143339 | 0,1677575 | 1,12 | 2,31E+07 | 1,86E+07 |
| 7,1 | | 0,2144283 | 0,1679439 | 1,46 | 6,29E+06 | 8,45E+06 |
| 27,6 | | 0,2145153 | 0,1679353 | 0,74 | 3,08E+07 | 5,21E+07 |
| 36,7 | | 0,2146160 | 0,1680226 | 0,87 | 8,48E+07 | 9,49E+07 |
| 39,6 | | 0,2146830 | 0,1680109 | 1,18 | 5,79E+07 | 8,28E+07 |
| 17 | | 0,2147399 | 0,1679992 | 1,25 | 3,41E+06 | 3,74E+06 |
| 9,2 | | 0,2149625 | 0,1682679 | 1,76 | 1,23E+06 | 2,76E+06 |
| 57,7 | | 0,2150840 | 0,1683660 | 0,94 | 5,62E+08 | 5,37E+08 |
| 40,7 | | 0,2152319 | 0,1684360 | 0,90 | 5,13E+08 | 4,81E+08 |
| 27,8 | | 0,2154431 | 0,1686468 | 1,18 | 3,09E+07 | 3,98E+07 |
| 9,9 | | 0,2158458 | 0,1689424 | 0,83 | 2,85E+07 | 2,42E+07 |
| 31,4 | | 0,2159454 | 0,1690403 | 1,11 | 8,63E+07 | 9,73E+07 |
| 21,4 | | 0,2159534 | 0,1690074 | 1,14 | 2,50E+07 | 2,68E+07 |
| 14,4 | | 0,2161507 | 0,1692016 | 0,85 | 1,79E+07 | 1,79E+07 |
| 27,7 | | 0,2168487 | 0,1698810 | 0,30 | 6,01E+07 | 4,38E+07 |
| 9,1 | | 0,2170170 | 0,1700393 | 0,33 | 1,62E+07 | 1,70E+07 |
| 4,5 | | 0,2174777 | 0,1703607 | 0,49 | 6,34E+06 | 1,44E+06 |

| | | | | | | |
|---|---|---|---|---|---|---|
| 37,8 | | 0,2175388 | 0,1703882 | 0,94 | 2,01E+08 | 2,10E+08 |
| 19 | | 0,2177251 | 0,1704639 | 0,88 | 1,39E+07 | 1,67E+07 |
| 4,3 | | 0,2180146 | 0,1706281 | 1,23 | 1,19E+07 | 1,19E+07 |
| 27,7 | | 0,2180752 | 0,1706548 | 0,89 | 8,78E+07 | 7,19E+07 |
| 12,5 | | 0,2182856 | 0,1709012 | 0,84 | 7,85E+07 | 8,20E+07 |
| 11,2 | | 0,2182922 | 0,1708890 | 1,24 | 9,24E+07 | 7,52E+07 |
| 31,1 | | 0,2183232 | 0,1708993 | 0,87 | 7,03E+07 | 6,20E+07 |
| 65,6 | | 0,2184992 | 0,1710741 | 0,10 | 6,08E+08 | 4,46E+06 |
| 41,2 | | 0,2188426 | 0,1714111 | 1,15 | 6,79E+07 | 5,71E+07 |
| 43,6 | | 0,2190288 | 0,1715058 | 0,94 | 2,54E+08 | 2,42E+08 |
| 49 | | 0,2190700 | 0,1714935 | 0,95 | 2,37E+09 | 2,33E+09 |
| 52,6 | | 0,2191706 | 0,1716278 | 0,48 | 4,79E+08 | 7,60E+08 |
| 12,4 | | 0,2192722 | 0,1717984 | 1,06 | 3,13E+07 | 2,95E+07 |
| 10,9 | | 0,2193030 | 0,1717891 | 0,50 | 1,19E+07 | 1,32E+07 |
| 19,3 | | 0,2195946 | 0,1720915 | 3,10 | 2,04E+06 | 2,36E+07 |
| 62,9 | | 0,2199869 | 0,1725432 | 1,28 | 1,40E+08 | 1,77E+08 |
| 17,7 | | 0,2201223 | 0,1727252 | 0,73 | 4,55E+07 | 3,58E+07 |
| 11,8 | | 0,2203977 | 0,1730232 | 0,82 | 1,35E+07 | 1,22E+07 |
| 6,3 | | 0,2205274 | 0,1731160 | 0,38 | 1,43E+07 | 7,33E+06 |
| 30,7 | | 0,2208939 | 0,1735091 | 0,88 | 8,45E+07 | 7,04E+07 |
| 11,8 | | 0,2209690 | 0,1736102 | 0,53 | 1,72E+07 | 1,82E+07 |
| 38,2 | | 0,2210306 | 0,1735944 | 1,12 | 6,90E+07 | 7,95E+07 |
| 25,6 | | 0,2212480 | 0,1739841 | 1,13 | 1,38E+07 | 1,15E+07 |
| 39,6 | | 0,2212625 | 0,1739745 | 1,05 | 3,91E+08 | 3,96E+08 |
| 31,6 | | 0,2213087 | 0,1739896 | 2,09 | 1,21E+07 | 1,04E+07 |
| 6,7 | | 0,2216023 | 0,1743782 | 0,48 | 6,40E+06 | 5,65E+06 |
| 22,9 | | 0,2216168 | 0,1744087 | 1,07 | 6,93E+07 | 6,86E+07 |
| 20,8 | | 0,2218575 | 0,1746595 | 1,08 | 5,87E+07 | 6,05E+07 |
| 51,7 | | 0,2219071 | 0,1746643 | 0,98 | 9,07E+09 | 9,08E+09 |
| 15,6 | | 0,2219601 | 0,1746677 | 0,64 | 2,34E+07 | 1,95E+07 |
| 42,2 | | 0,2225702 | 0,1755279 | 0,87 | 1,47E+07 | 1,62E+07 |
| 8,5 | | 0,2229404 | 0,1757451 | 0,48 | 7,20E+06 | 7,87E+06 |
| 5,7 | | 0,2231036 | 0,1759027 | 0,84 | 1,84E+07 | 2,24E+07 |
| 26 | | 0,2233287 | 0,1761876 | 0,86 | 2,53E+07 | 2,57E+07 |
| 43,5 | | 0,2234556 | 0,1763513 | 0,96 | 6,47E+08 | 6,72E+08 |
| 4,8 | | 0,2235750 | 0,1765149 | 0,57 | 1,35E+07 | 4,15E+07 |
| 40,6 | | 0,2236194 | 0,1765016 | 0,91 | 3,33E+08 | 2,68E+08 |
| 14,2 | | 0,2237587 | 0,1766181 | 0,72 | 1,37E+07 | 1,26E+07 |
| 20,6 | | 0,2237767 | 0,1765840 | 1,16 | 3,10E+07 | 3,11E+07 |
| 36,2 | | 0,2239089 | 0,1766741 | 2,06 | 3,33E+06 | 4,97E+06 |
| 28,8 | | 0,2239318 | 0,1766401 | 0,61 | 6,59E+07 | 1,37E+08 |
| 22,1 | | 0,2239335 | 0,1766060 | 0,95 | 3,31E+07 | 3,48E+07 |
| 51,2 | | 0,2240678 | 0,1766784 | 1,21 | 1,20E+08 | 1,82E+08 |
| 28,6 | | 0,2242254 | 0,1767800 | 0,91 | 7,58E+07 | 7,13E+07 |
| 32,9 | | 0,2245377 | 0,1770927 | 1,20 | 3,46E+07 | 2,68E+07 |
| 26,7 | | 0,2248588 | 0,1775547 | 1,23 | 1,77E+08 | 1,07E+08 |
| 43,1 | | 0,2250068 | 0,1777077 | 1,16 | 2,86E+08 | 2,58E+08 |
| 59,6 | | 0,2250091 | 0,1776735 | 1,09 | 2,16E+08 | 1,94E+08 |
| 35,4 | | 0,2250214 | 0,1776393 | 0,91 | 1,73E+08 | 2,14E+08 |
| 15,1 | | 0,2250540 | 0,1776051 | 0,63 | 6,33E+06 | 7,90E+06 |

| | | | | | | |
|---|---|---|---|---|---|---|
| 8,4 | | 0,2252714 | 0,1777764 | 2,90 | 1,70E+06 | 1,80E+07 |
| 6,3 | | 0,2253254 | 0,1778661 | 0,82 | 1,33E+07 | 1,33E+07 |
| 11,1 | | 0,2253362 | 0,1778319 | 1,16 | 2,72E+06 | 2,42E+06 |
| 47,5 | + | 0,2253461 | 0,1777977 | 1,14 | 3,61E+08 | 4,01E+08 |
| 21,8 | | 0,2253964 | 0,1778450 | 0,78 | 4,59E+07 | 6,04E+07 |
| 29,5 | | 0,2255278 | 0,1779815 | 0,79 | 2,64E+07 | 2,85E+07 |
| 12 | | 0,2255474 | 0,1779473 | 1,39 | 2,28E+07 | 1,51E+07 |
| 30 | | 0,2256141 | 0,1780000 | 0,85 | 3,16E+07 | 3,38E+07 |
| 6,2 | | 0,2257009 | 0,1780350 | 0,85 | 3,83E+07 | 3,16E+07 |
| 27 | | 0,2258293 | 0,1781452 | 0,94 | 1,12E+08 | 1,27E+08 |
| 20,5 | | 0,2262545 | 0,1785358 | 1,09 | 8,56E+07 | 8,79E+07 |
| 7,1 | | 0,2265054 | 0,1788257 | 1,82 | 2,44E+06 | 9,71E+06 |
| 26,5 | | 0,2265069 | 0,1787913 | 1,07 | 1,05E+08 | 1,05E+08 |
| 52,1 | | 0,2266444 | 0,1788445 | 1,08 | 3,12E+09 | 2,90E+09 |
| 40,8 | | 0,2266624 | 0,1788463 | 1,14 | 9,31E+08 | 1,07E+09 |
| 29,7 | | 0,2271630 | 0,1793438 | 0,92 | 1,33E+08 | 1,20E+08 |
| 19,6 | | 0,2272452 | 0,1794191 | 0,86 | 1,71E+08 | 1,32E+08 |
| 29,2 | | 0,2273089 | 0,1795129 | 0,65 | 3,28E+08 | 1,48E+08 |
| 5,5 | | 0,2275539 | 0,1797630 | 1,94 | 2,31E+07 | 8,54E+07 |
| 47,9 | | 0,2275687 | 0,1797285 | 0,91 | 2,35E+09 | 2,53E+09 |
| 61,3 | | 0,2280566 | 0,1800920 | 1,06 | 3,92E+08 | 4,45E+08 |
| 32,7 | | 0,2282591 | 0,1803120 | 1,14 | 1,71E+08 | 1,44E+08 |
| 33 | | 0,2284404 | 0,1804775 | 1,03 | 5,26E+08 | 5,08E+08 |
| 9,3 | | 0,2284584 | 0,1804858 | 1,37 | 5,68E+06 | 3,72E+06 |
| 42,4 | | 0,2286478 | 0,1806244 | 0,96 | 6,36E+08 | 6,10E+08 |
| 31,3 | | 0,2287116 | 0,1806741 | 1,10 | 4,79E+07 | 4,52E+07 |
| 7,9 | | 0,2287501 | 0,1806632 | 0,75 | 1,31E+10 | 1,41E+10 |
| 22,5 | | 0,2287907 | 0,1806470 | 1,56 | 8,28E+06 | 3,92E+06 |
| 36 | | 0,2288098 | 0,1806124 | 0,92 | 1,22E+08 | 1,23E+08 |
| 36,4 | | 0,2289759 | 0,1807815 | 0,88 | 2,08E+08 | 1,91E+08 |
| 47,6 | | 0,2290452 | 0,1809068 | 0,90 | 2,72E+08 | 2,12E+08 |
| 33 | | 0,2296614 | 0,1817192 | 1,06 | 6,11E+07 | 6,11E+07 |
| 13,1 | | 0,2296995 | 0,1817671 | 1,21 | 2,64E+07 | 4,02E+07 |
| 22,6 | | 0,2297038 | 0,1817499 | 0,62 | 9,99E+06 | 1,17E+07 |
| 30,1 | | 0,2297069 | 0,1817152 | 0,25 | 2,57E+07 | 2,18E+08 |
| 38,5 | | 0,2298093 | 0,1817599 | 1,43 | 5,14E+06 | 1,03E+07 |
| 14,6 | | 0,2298619 | 0,1817917 | 1,34 | 3,83E+06 | 7,69E+06 |
| 1,2 | | 0,2305790 | 0,1826649 | 0,73 | 3,40E+06 | 6,91E+06 |
| 43,9 | | 0,2307001 | 0,1828562 | 0,94 | 1,85E+09 | 1,91E+09 |
| 49,2 | | 0,2309222 | 0,1829580 | 1,12 | 1,81E+08 | 1,84E+08 |
| 15,5 | | 0,2309485 | 0,1829406 | 0,87 | 3,45E+07 | 2,97E+07 |
| 58,2 | | 0,2313185 | 0,1832791 | 1,07 | 1,92E+08 | 1,96E+08 |
| 15,6 | | 0,2317262 | 0,1837106 | 0,56 | 2,49E+06 | 4,60E+06 |
| 15,5 | | 0,2318761 | 0,1838252 | 2,06 | 6,04E+06 | 1,60E+06 |
| 26,8 | | 0,2322304 | 0,1841061 | 1,39 | 2,49E+07 | 2,52E+07 |
| 16,3 | | 0,2329516 | 0,1850568 | 1,57 | 4,59E+06 | 1,12E+07 |
| 31,2 | | 0,2332448 | 0,1854396 | 0,84 | 2,39E+07 | 1,92E+07 |
| 49,2 | | 0,2337053 | 0,1859375 | 1,13 | 1,08E+08 | 9,29E+07 |
| 10,4 | | 0,2339148 | 0,1861781 | 0,85 | 3,00E+07 | 2,83E+07 |
| 4,9 | | 0,2341380 | 0,1862920 | 1,22 | 4,82E+06 | 4,77E+06 |

| | | | | | | |
|---|---|---|---|---|---|---|
| 40,4 | | 0,2341962 | 0,1863450 | 2,10 | 5,41E+06 | 1,27E+07 |
| 24 | | 0,2345234 | 0,1866136 | 3,07 | 5,69E+06 | 7,18E+07 |
| 2,1 | | 0,2345992 | 0,1867548 | 1,24 | 1,71E+06 | 2,83E+06 |
| 8,3 | | 0,2349861 | 0,1871756 | 0,71 | 4,68E+06 | 5,69E+06 |
| 74,6 | | 0,2353713 | 0,1876092 | 1,05 | 2,69E+09 | 2,53E+09 |
| 6,1 | | 0,2355232 | 0,1878195 | 1,38 | 4,22E+06 | 7,80E+06 |
| 17,3 | | 0,2355537 | 0,1878454 | 1,12 | 4,98E+07 | 4,22E+07 |
| 9,9 | | 0,2356630 | 0,1879604 | 0,65 | 1,31E+07 | 5,87E+06 |
| 14,7 | | 0,2357180 | 0,1879665 | 0,82 | 5,61E+06 | 5,79E+06 |
| 40,9 | | 0,2361204 | 0,1884650 | 1,13 | 4,38E+08 | 5,78E+08 |
| 43,3 | | 0,2367105 | 0,1890729 | 0,89 | 6,61E+07 | 7,74E+07 |
| 22,1 | | 0,2367344 | 0,1890787 | 1,09 | 5,27E+07 | 4,77E+07 |
| 16,3 | | 0,2368587 | 0,1890869 | 0,90 | 2,95E+07 | 3,00E+07 |
| 27,5 | | 0,2372071 | 0,1894768 | 0,85 | 3,84E+07 | 2,94E+07 |
| 41,2 | | 0,2372494 | 0,1894879 | 0,86 | 1,80E+08 | 2,53E+08 |
| 21,3 | | 0,2373210 | 0,1894724 | 0,91 | 1,49E+08 | 1,36E+08 |
| 44,8 | | 0,2374452 | 0,1896188 | 0,82 | 1,62E+09 | 1,12E+09 |
| 62,1 | | 0,2383686 | 0,1908465 | 1,27 | 1,91E+08 | 3,26E+08 |
| 18,3 | | 0,2383994 | 0,1908308 | 1,63 | 1,70E+07 | 2,56E+07 |
| 3,8 | | 0,2384794 | 0,1908621 | 1,98 | 7,58E+05 | 8,60E+05 |
| 57 | | 0,2387719 | 0,1911669 | 0,93 | 2,40E+08 | 2,61E+08 |
| 46,2 | | 0,2390692 | 0,1914897 | 1,13 | 1,31E+08 | 1,04E+08 |
| 19,3 | | 0,2391381 | 0,1915483 | 0,78 | 1,46E+08 | 1,52E+08 |
| 30,3 | | 0,2394243 | 0,1918565 | 0,90 | 3,19E+07 | 3,97E+07 |
| 28,1 | | 0,2399167 | 0,1925547 | 1,88 | 3,99E+06 | 2,10E+06 |
| 27,4 | | 0,2402645 | 0,1929939 | 1,13 | 1,34E+08 | 1,44E+08 |
| 24,1 | | 0,2404730 | 0,1933048 | 0,72 | 1,60E+07 | 1,98E+07 |
| 5,4 | | 0,2406732 | 0,1934736 | 0,18 | 2,71E+07 | 1,42E+07 |
| 54,1 | | 0,2409917 | 0,1937153 | 1,07 | 6,19E+07 | 6,29E+07 |
| 6,8 | | 0,2410773 | 0,1937635 | 0,70 | 2,44E+06 | 2,33E+06 |
| 1,8 | | 0,2411858 | 0,1940148 | 0,59 | 6,84E+06 | 2,50E+06 |
| 17,2 | | 0,2413156 | 0,1942380 | 1,45 | 3,75E+06 | 3,96E+06 |
| 41,5 | | 0,2417217 | 0,1946323 | 1,06 | 4,78E+09 | 4,29E+09 |
| 9,6 | | 0,2418065 | 0,1946136 | 1,06 | 7,44E+06 | 6,87E+06 |
| 36 | | 0,2418083 | 0,1945768 | 0,96 | 7,36E+08 | 7,81E+08 |
| 17,5 | | 0,2421109 | 0,1949216 | 0,90 | 2,07E+07 | 2,11E+07 |
| 25,3 | | 0,2421352 | 0,1949763 | 1,22 | 3,07E+07 | 4,67E+07 |
| 6 | | 0,2428020 | 0,1956768 | 0,73 | 6,26E+06 | 3,73E+06 |
| 70,7 | | 0,2428922 | 0,1956571 | 1,40 | 2,43E+08 | 2,86E+08 |
| 18,5 | | 0,2429259 | 0,1956640 | 1,30 | 2,77E+07 | 2,73E+07 |
| 10,1 | | 0,2434337 | 0,1963329 | 0,83 | 3,81E+07 | 3,24E+07 |
| 41 | | 0,2435655 | 0,1964554 | 0,93 | 2,54E+08 | 2,53E+08 |
| 55,2 | | 0,2437758 | 0,1966436 | 0,90 | 7,26E+09 | 6,43E+09 |
| 18,7 | | 0,2444561 | 0,1972885 | 1,43 | 7,07E+06 | 5,74E+06 |
| 9,4 | | 0,2446346 | 0,1974107 | 0,73 | 6,92E+06 | 6,77E+06 |
| 10,7 | | 0,2453836 | 0,1983908 | 0,81 | 5,16E+07 | 3,75E+07 |
| 32,4 | | 0,2455343 | 0,1985196 | 0,93 | 2,06E+08 | 1,74E+08 |
| 52,5 | | 0,2457798 | 0,1989430 | 1,21 | 5,24E+08 | 8,44E+08 |
| 52,3 | | 0,2458473 | 0,1989258 | 1,16 | 1,18E+08 | 8,32E+07 |
| 9,5 | | 0,2458946 | 0,1989298 | 1,73 | 6,65E+06 | 8,45E+06 |

| | | | | | | |
|---|---|---|---|---|---|---|
| 23,7 | | 0,2460549 | 0,1991995 | 0,44 | 2,41E+07 | 2,27E+07 |
| 51,6 | | 0,2461783 | 0,1993431 | 0,94 | 1,36E+09 | 1,29E+09 |
| 37,5 | | 0,2463711 | 0,1996294 | 0,85 | 1,75E+07 | 1,29E+07 |
| 11,4 | | 0,2465914 | 0,1998672 | 0,83 | 3,92E+06 | 5,82E+06 |
| 10,9 | | 0,2480859 | 0,2016865 | 1,17 | 2,63E+07 | 1,99E+07 |
| 3,3 | | 0,2481751 | 0,2016892 | 0,56 | 3,28E+06 | 3,14E+06 |
| 41,9 | | 0,2481908 | 0,2016745 | 0,96 | 7,93E+09 | 7,33E+09 |
| 44,7 | | 0,2484444 | 0,2018793 | 0,92 | 1,96E+08 | 2,09E+08 |
| 56,1 | | 0,2486682 | 0,2021331 | 0,91 | 1,34E+09 | 1,45E+09 |
| 21,4 | | 0,2488479 | 0,2023211 | 0,89 | 5,66E+07 | 4,64E+07 |
| 45 | | 0,2489714 | 0,2025024 | 0,93 | 5,94E+08 | 6,28E+08 |
| 15,4 | | 0,2491979 | 0,2028561 | 1,10 | 5,52E+07 | 5,71E+07 |
| 9 | | 0,2493468 | 0,2029896 | 0,82 | 4,44E+06 | 6,71E+06 |
| 13,9 | | 0,2493971 | 0,2029755 | 1,21 | 9,19E+06 | 1,39E+07 |
| 73,6 | | 0,2495302 | 0,2030088 | 0,92 | 4,68E+10 | 5,30E+10 |
| 26,6 | | 0,2495740 | 0,2030941 | 0,85 | 1,72E+07 | 1,52E+07 |
| 5,5 | | 0,2496542 | 0,2032471 | 0,72 | 4,02E+06 | 5,68E+06 |
| 16,4 | | 0,2497020 | 0,2032729 | 0,81 | 1,50E+07 | 2,44E+07 |
| 30,3 | | 0,2498117 | 0,2033573 | 0,94 | 5,12E+07 | 5,96E+07 |
| 59,5 | | 0,2500253 | 0,2035297 | 0,97 | 1,28E+09 | 1,36E+09 |
| 25,8 | | 0,2501230 | 0,2036058 | 1,11 | 5,08E+09 | 4,49E+09 |
| 16 | | 0,2503010 | 0,2037638 | 0,60 | 6,13E+07 | 3,57E+07 |
| 64,3 | | 0,2505601 | 0,2038601 | 0,95 | 1,25E+08 | 1,20E+08 |
| 20 | | 0,2506182 | 0,2038579 | 0,88 | 2,38E+07 | 2,73E+07 |
| 3,5 | | 0,2506392 | 0,2038812 | 0,86 | 8,51E+08 | 6,57E+08 |
| 55,1 | | 0,2509124 | 0,2043525 | 0,94 | 2,31E+08 | 2,00E+08 |
| 6,7 | | 0,2509453 | 0,2043141 | 1,41 | 1,56E+06 | 8,49E+05 |
| 18,1 | | 0,2510556 | 0,2044433 | 1,07 | 9,00E+08 | 9,89E+08 |
| 10,6 | | 0,2511138 | 0,2044823 | 0,72 | 7,74E+06 | 1,33E+07 |
| 42,5 | | 0,2512694 | 0,2047187 | 0,85 | 6,38E+07 | 6,70E+07 |
| 45,4 | | 0,2515263 | 0,2051128 | 1,04 | 1,63E+09 | 1,48E+09 |
| 52,1 | | 0,2518491 | 0,2053408 | 0,99 | 3,09E+09 | 3,05E+09 |
| 14,3 | | 0,2518492 | 0,2053023 | 1,24 | 1,27E+07 | 1,39E+07 |
| 32,5 | + | 0,2519118 | 0,2053704 | 2,23 | 1,33E+08 | 5,21E+07 |
| 16,5 | | 0,2519516 | 0,2053784 | 1,16 | 4,19E+07 | 4,12E+07 |
| 8,5 | | 0,2524339 | 0,2058170 | 1,28 | 1,34E+07 | 9,96E+06 |
| 2,4 | | 0,2529861 | 0,2064984 | 1,20 | 1,68E+07 | 2,12E+07 |
| 52,9 | | 0,2534017 | 0,2070956 | 1,11 | 1,92E+07 | 2,40E+07 |
| 50,5 | | 0,2535986 | 0,2072097 | 1,05 | 6,07E+09 | 5,44E+09 |
| 6,8 | | 0,2537836 | 0,2072564 | 0,74 | 3,16E+06 | 3,46E+06 |
| 0,4 | | 0,2538184 | 0,2072610 | 1,88 | 2,88E+06 | 3,91E+06 |
| 49 | | 0,2541317 | 0,2075991 | 1,18 | 3,13E+08 | 3,38E+08 |
| 44,4 | | 0,2541624 | 0,2076037 | 1,23 | 4,07E+07 | 6,34E+07 |
| 7,4 | | 0,2543487 | 0,2078644 | 0,69 | 3,55E+06 | 8,93E+06 |
| 4,4 | | 0,2545925 | 0,2080464 | 1,30 | 2,83E+07 | 2,14E+07 |
| 33 | | 0,2546863 | 0,2081086 | 1,14 | 8,55E+07 | 9,68E+07 |
| 37,7 | | 0,2552210 | 0,2088490 | 0,93 | 7,94E+08 | 9,00E+08 |
| 33 | | 0,2554985 | 0,2091684 | 0,92 | 3,19E+08 | 2,90E+08 |
| 9,6 | | 0,2557792 | 0,2095259 | 1,27 | 5,47E+06 | 5,30E+06 |
| 54,5 | | 0,2558061 | 0,2094867 | 1,13 | 4,53E+09 | 3,60E+09 |

| | | | | | | |
|---|---|---|---|---|---|---|
| 12,9 | | 0,2559257 | 0,2096465 | 0,89 | 1,98E+07 | 1,48E+07 |
| 3,8 | | 0,2562036 | 0,2099491 | 0,75 | 6,81E+06 | 5,43E+06 |
| 8,4 | | 0,2562924 | 0,2100236 | 1,32 | 6,90E+06 | 6,06E+06 |
| 50,9 | | 0,2563801 | 0,2100688 | 0,95 | 8,18E+08 | 7,99E+08 |
| 15,8 | | 0,2569171 | 0,2105730 | 0,96 | 1,41E+08 | 1,43E+08 |
| 62,2 | | 0,2575232 | 0,2115149 | 1,03 | 4,77E+09 | 4,65E+09 |
| 29,7 | | 0,2577298 | 0,2117093 | 0,93 | 5,58E+07 | 5,64E+07 |
| 13,9 | | 0,2578991 | 0,2118297 | 0,54 | 1,31E+07 | 8,65E+06 |
| 39,6 | | 0,2583931 | 0,2124056 | 0,94 | 8,35E+07 | 9,31E+07 |
| 26,4 | | 0,2586424 | 0,2127603 | 1,09 | 1,82E+08 | 1,84E+08 |
| 4,7 | | 0,2591661 | 0,2133336 | 2,90 | 2,85E+06 | 6,15E+06 |
| 28,5 | | 0,2592500 | 0,2134431 | 1,04 | 3,50E+07 | 3,51E+07 |
| 24,4 | | 0,2593087 | 0,2134615 | 0,91 | 9,56E+07 | 8,35E+07 |
| 31,5 | | 0,2602561 | 0,2144642 | 1,17 | 1,07E+08 | 1,11E+08 |
| 14,3 | | 0,2605698 | 0,2146980 | 1,24 | 1,67E+07 | 9,58E+06 |
| 4,7 | | 0,2606817 | 0,2147370 | 0,54 | 6,41E+06 | 5,52E+06 |
| 6,8 | | 0,2607080 | 0,2147380 | 1,12 | 5,85E+07 | 6,18E+07 |
| 52,8 | | 0,2608462 | 0,2147972 | 0,86 | 4,23E+09 | 4,49E+09 |
| 11,9 | | 0,2609127 | 0,2147981 | 1,56 | 1,66E+06 | 4,37E+06 |
| 36,2 | | 0,2611049 | 0,2150376 | 0,83 | 1,55E+08 | 1,55E+08 |
| 54,2 | | 0,2612688 | 0,2151876 | 0,92 | 3,07E+08 | 3,19E+08 |
| 21,6 | | 0,2615352 | 0,2154143 | 0,86 | 2,78E+08 | 2,75E+08 |
| 39,1 | | 0,2617544 | 0,2155806 | 1,31 | 8,52E+07 | 9,27E+07 |
| 11,3 | | 0,2617968 | 0,2155635 | 1,48 | 2,51E+07 | 2,18E+07 |
| 49,2 | | 0,2623430 | 0,2161110 | 0,93 | 8,80E+08 | 1,04E+09 |
| 44,6 | | 0,2624571 | 0,2162100 | 0,44 | 3,22E+07 | 4,04E+07 |
| 48,8 | | 0,2625777 | 0,2163186 | 0,90 | 2,11E+08 | 2,15E+08 |
| 9,4 | | 0,2625842 | 0,2163022 | 0,79 | 1,08E+07 | 1,18E+07 |
| 65,3 | | 0,2634296 | 0,2172807 | 1,05 | 4,72E+08 | 5,11E+08 |
| 5,9 | | 0,2637723 | 0,2177448 | 0,64 | 6,77E+06 | 6,74E+06 |
| 43,9 | | 0,2637957 | 0,2177043 | 0,89 | 1,94E+09 | 2,09E+09 |
| 22,4 | | 0,2639470 | 0,2177449 | 1,42 | 1,08E+07 | 9,05E+06 |
| 51,2 | | 0,2641535 | 0,2179483 | 0,80 | 5,66E+08 | 4,88E+08 |
| 54,6 | | 0,2642125 | 0,2179717 | 1,08 | 7,15E+08 | 7,15E+08 |
| 35,1 | | 0,2642481 | 0,2180613 | 1,12 | 3,77E+07 | 4,70E+07 |
| 3,6 | | 0,2644890 | 0,2184199 | 0,27 | 4,93E+07 | 3,79E+06 |
| 24,7 | | 0,2650896 | 0,2190273 | 1,26 | 6,90E+07 | 5,47E+07 |
| 11,4 | | 0,2653030 | 0,2192474 | 0,73 | 5,09E+06 | 4,97E+06 |
| 64,8 | | 0,2655736 | 0,2196323 | 1,11 | 4,14E+09 | 3,74E+09 |
| 4,9 | | 0,2657323 | 0,2197534 | 0,55 | 1,38E+07 | 1,26E+07 |
| 31,1 | | 0,2659385 | 0,2200512 | 0,95 | 7,60E+08 | 7,24E+08 |
| 4,5 | | 0,2659890 | 0,2200720 | 1,13 | 8,35E+07 | 8,41E+07 |
| 24,8 | | 0,2672085 | 0,2217228 | 0,93 | 8,59E+07 | 7,63E+07 |
| 37,5 | | 0,2674752 | 0,2220312 | 0,92 | 3,82E+08 | 3,66E+08 |
| 19,8 | | 0,2675588 | 0,2221265 | 1,10 | 5,27E+07 | 5,41E+07 |
| 51,7 | | 0,2682835 | 0,2230638 | 0,95 | 9,53E+08 | 9,32E+08 |
| 58,3 | | 0,2684133 | 0,2231048 | 0,94 | 6,49E+08 | 6,51E+08 |
| 2,5 | | 0,2684830 | 0,2230871 | 2,29 | 3,21E+07 | 3,00E+07 |
| 5,1 | | 0,2685992 | 0,2232044 | 0,60 | 4,81E+06 | 3,40E+06 |
| 15 | | 0,2687967 | 0,2233239 | 0,89 | 1,74E+07 | 1,70E+07 |

| | | | | | | |
|---|---|---|---|---|---|---|
| 45,8 | | 0,2692040 | 0,2238629 | 1,13 | 3,58E+08 | 2,65E+08 |
| 15 | | 0,2698196 | 0,2247069 | 1,50 | 9,10E+06 | 1,64E+07 |
| 41,6 | | 0,2699544 | 0,2247809 | 4,35 | 3,11E+06 | 9,50E+07 |
| 14,2 | | 0,2702509 | 0,2251644 | 2,21 | 2,17E+07 | 3,31E+07 |
| 8,1 | | 0,2704624 | 0,2253546 | 0,39 | 1,83E+07 | 2,50E+07 |
| 46,7 | | 0,2707262 | 0,2256653 | 1,09 | 2,65E+08 | 3,00E+08 |
| 62,5 | | 0,2708677 | 0,2257761 | 1,16 | 1,16E+10 | 1,40E+10 |
| 3 | | 0,2713029 | 0,2261858 | 1,99 | 3,37E+06 | 7,78E+06 |
| 4,1 | | 0,2714018 | 0,2262735 | 0,57 | 1,27E+07 | 5,36E+06 |
| 43,6 | | 0,2714934 | 0,2262516 | 1,10 | 5,93E+07 | 6,76E+07 |
| 33,7 | | 0,2715821 | 0,2264257 | 1,26 | 1,48E+08 | 2,16E+08 |
| 36,5 | | 0,2717103 | 0,2265740 | 0,90 | 3,25E+08 | 2,75E+08 |
| 1,9 | | 0,2717793 | 0,2265698 | 1,26 | 3,24E+06 | 1,94E+06 |
| 17,4 | | 0,2737326 | 0,2289050 | 0,80 | 4,05E+06 | 6,17E+06 |
| 46,7 | | 0,2737898 | 0,2289233 | 0,87 | 5,84E+08 | 4,88E+08 |
| 19,8 | | 0,2743791 | 0,2295137 | 0,90 | 1,06E+08 | 1,33E+08 |
| 24,3 | | 0,2747143 | 0,2300454 | 0,91 | 2,01E+08 | 2,02E+08 |
| 36 | | 0,2748457 | 0,2301027 | 1,09 | 1,89E+08 | 2,07E+08 |
| 33,6 | | 0,2748534 | 0,2300838 | 0,90 | 5,43E+07 | 4,35E+07 |
| 36 | | 0,2754321 | 0,2306233 | 1,13 | 2,46E+07 | 3,09E+07 |
| 8,5 | | 0,2758210 | 0,2310090 | 0,51 | 1,99E+07 | 1,13E+07 |
| 4 | | 0,2759153 | 0,2310735 | 1,44 | 2,46E+06 | 4,52E+06 |
| 43,5 | | 0,2760134 | 0,2311578 | 1,06 | 3,00E+08 | 3,27E+08 |
| 8 | | 0,2763510 | 0,2316465 | 0,65 | 2,07E+07 | 1,15E+07 |
| 15,6 | | 0,2763657 | 0,2316443 | 0,82 | 2,62E+07 | 1,61E+07 |
| 29,2 | | 0,2766532 | 0,2318554 | 1,16 | 1,51E+08 | 1,51E+08 |
| 10,5 | | 0,2766917 | 0,2318532 | 1,40 | 2,02E+07 | 3,11E+07 |
| 2,2 | | 0,2769766 | 0,2321681 | 0,65 | 1,17E+08 | 1,36E+08 |
| 14,6 | | 0,2773628 | 0,2325132 | 1,21 | 2,07E+07 | 2,03E+07 |
| 26,6 | | 0,2775857 | 0,2328094 | 1,10 | 3,18E+08 | 2,65E+08 |
| 21,3 | | 0,2783204 | 0,2335375 | 1,14 | 1,05E+08 | 1,44E+08 |
| 4,9 | | 0,2784696 | 0,2335947 | 0,61 | 2,27E+06 | 2,88E+06 |
| 20,5 | | 0,2792251 | 0,2345227 | 0,97 | 4,61E+07 | 4,32E+07 |
| 70,7 | | 0,2795501 | 0,2350011 | 1,08 | 2,53E+09 | 2,10E+09 |
| 58,9 | | 0,2796938 | 0,2350794 | 0,90 | 1,42E+09 | 1,56E+09 |
| 13,9 | | 0,2802552 | 0,2358358 | 1,14 | 3,52E+07 | 3,84E+07 |
| 15,3 | | 0,2810212 | 0,2366045 | 0,81 | 7,70E+07 | 8,12E+07 |
| 25,2 | | 0,2823496 | 0,2383180 | 0,83 | 8,65E+06 | 1,02E+07 |
| 48,8 | | 0,2829514 | 0,2390859 | 0,90 | 1,39E+10 | 1,60E+10 |
| 43,1 | | 0,2829961 | 0,2390817 | 0,90 | 2,12E+08 | 1,61E+08 |
| 4,2 | | 0,2834055 | 0,2394460 | 1,39 | 3,69E+06 | 5,13E+06 |
| 21,1 | | 0,2840454 | 0,2404222 | 1,21 | 3,05E+07 | 3,19E+07 |
| 17,9 | | 0,2840830 | 0,2404192 | 2,11 | 9,77E+06 | 1,80E+07 |
| 30,8 | | 0,2841343 | 0,2404757 | 1,08 | 7,83E+08 | 8,39E+08 |
| 20,4 | | 0,2842440 | 0,2404926 | 0,89 | 3,70E+07 | 3,79E+07 |
| 48,6 | | 0,2844865 | 0,2408328 | 0,88 | 2,59E+08 | 2,41E+08 |
| 68,3 | | 0,2846930 | 0,2411435 | 0,86 | 1,02E+08 | 8,67E+07 |
| 38,2 | | 0,2849357 | 0,2415489 | 0,93 | 2,56E+08 | 2,60E+08 |
| 19,8 | | 0,2849585 | 0,2416022 | 0,68 | 7,87E+06 | 4,03E+06 |
| 10,1 | | 0,2850170 | 0,2416599 | 0,90 | 1,63E+07 | 1,64E+07 |

| | | | | | | |
|---|---|---|---|---|---|---|
| 10 | | 0,2851206 | 0,2418263 | 1,34 | 4,67E+06 | 5,54E+06 |
| 2,6 | | 0,2856866 | 0,2425345 | 1,72 | 1,06E+06 | 1,60E+06 |
| 60,5 | | 0,2858212 | 0,2428475 | 1,09 | 1,68E+08 | 2,00E+08 |
| 11 | | 0,2858556 | 0,2428631 | 2,10 | 7,93E+06 | 1,40E+06 |
| 14,4 | | 0,2859171 | 0,2429565 | 0,67 | 2,35E+07 | 1,33E+07 |
| 59,5 | | 0,2860946 | 0,2431034 | 1,06 | 2,79E+08 | 3,11E+08 |
| 34,3 | | 0,2862211 | 0,2432511 | 1,64 | 3,69E+06 | 1,24E+07 |
| 12,3 | | 0,2863219 | 0,2433231 | 0,90 | 6,88E+06 | 6,99E+06 |
| 7,7 | | 0,2865016 | 0,2435490 | 1,33 | 1,36E+07 | 1,55E+07 |
| 36,8 | | 0,2868879 | 0,2440770 | 0,76 | 7,33E+07 | 6,08E+07 |
| 16,3 | | 0,2869687 | 0,2441268 | 0,79 | 1,96E+07 | 1,22E+07 |
| 37 | | 0,2869784 | 0,2441048 | 0,96 | 1,97E+08 | 1,82E+08 |
| 8,8 | | 0,2871535 | 0,2442447 | 0,85 | 1,77E+07 | 2,02E+07 |
| 27,5 | | 0,2873568 | 0,2444293 | 0,88 | 4,39E+07 | 4,14E+07 |
| 24,7 | | 0,2873744 | 0,2444021 | 0,95 | 8,01E+07 | 9,09E+07 |
| 25,2 | | 0,2883105 | 0,2455577 | 2,19 | 1,15E+07 | 7,37E+06 |
| 26,2 | | 0,2883392 | 0,2455552 | 0,87 | 1,65E+08 | 2,22E+08 |
| 45 | | 0,2883636 | 0,2455476 | 0,88 | 7,34E+07 | 9,31E+07 |
| 27,1 | | 0,2883902 | 0,2455027 | 0,85 | 6,00E+07 | 8,67E+07 |
| 5,7 | | 0,2884166 | 0,2454804 | 0,84 | 2,98E+06 | 3,83E+06 |
| 24,3 | | 0,2884657 | 0,2455577 | 0,85 | 3,14E+07 | 3,03E+07 |
| 71,3 | | 0,2887141 | 0,2457630 | 0,83 | 8,04E+08 | 8,45E+08 |
| 14,8 | | 0,2887772 | 0,2458387 | 0,48 | 2,71E+07 | 2,79E+07 |
| 14,2 | | 0,2889147 | 0,2459817 | 0,42 | 2,40E+06 | 1,51E+07 |
| 8,8 | | 0,2889967 | 0,2460939 | 1,35 | 7,93E+06 | 6,71E+06 |
| 20,7 | | 0,2891242 | 0,2462887 | 0,85 | 4,13E+07 | 4,83E+07 |
| 23,1 | | 0,2898859 | 0,2472772 | 1,25 | 6,85E+07 | 6,75E+07 |
| 9,3 | | 0,2905023 | 0,2480438 | 1,67 | 9,40E+06 | 2,82E+06 |
| 5,6 | | 0,2907622 | 0,2483726 | 1,49 | 2,31E+06 | 6,36E+06 |
| 59,5 | | 0,2908297 | 0,2484127 | 1,48 | 1,60E+08 | 1,65E+08 |
| 4,7 | | 0,2914094 | 0,2490670 | 1,20 | 4,00E+07 | 4,56E+07 |
| 10,4 | | 0,2914561 | 0,2490391 | 0,71 | 6,37E+06 | 8,82E+06 |
| 7,4 | | 0,2915759 | 0,2491542 | 0,74 | 1,28E+07 | 1,58E+07 |
| 39,1 | | 0,2916266 | 0,2492445 | 1,14 | 7,81E+07 | 8,82E+07 |
| 14,1 | | 0,2917382 | 0,2493333 | 0,94 | 5,14E+07 | 5,32E+07 |
| 20,1 | | 0,2917813 | 0,2493331 | 0,80 | 1,17E+07 | 8,23E+06 |
| 3,4 | | 0,2918355 | 0,2493336 | 0,81 | 9,39E+06 | 1,19E+07 |
| 31,8 | | 0,2919970 | 0,2494522 | 0,93 | 2,38E+08 | 2,28E+08 |
| 41 | | 0,2925046 | 0,2500303 | 0,84 | 8,13E+07 | 1,07E+08 |
| 23,5 | | 0,2930232 | 0,2505330 | 0,95 | 3,91E+08 | 3,84E+08 |
| 10,5 | | 0,2930843 | 0,2505471 | 3,08 | 2,65E+06 | 1,89E+07 |
| 46,9 | | 0,2931008 | 0,2505583 | 0,89 | 2,06E+08 | 2,43E+08 |
| 18,3 | | 0,2933011 | 0,2507837 | 1,18 | 4,22E+06 | 3,04E+06 |
| 15,4 | | 0,2933264 | 0,2507381 | 1,59 | 8,22E+06 | 2,69E+07 |
| 12,9 | | 0,2933599 | 0,2506924 | 0,78 | 1,31E+08 | 1,23E+08 |
| 16,2 | | 0,2933745 | 0,2506861 | 0,91 | 1,72E+07 | 1,96E+07 |
| 1,5 | | 0,2934125 | 0,2506405 | 1,52 | 8,53E+06 | 2,84E+06 |
| 43,6 | | 0,2937247 | 0,2508700 | 0,59 | 3,07E+08 | 8,64E+07 |
| 11,6 | | 0,2939854 | 0,2511884 | 1,12 | 4,31E+07 | 3,37E+07 |
| 55 | | 0,2941319 | 0,2513996 | 0,88 | 1,01E+08 | 1,32E+08 |

| | | | | | | |
|---|---|---|---|---|---|---|
| 11,8 | | 0,2941781 | 0,2514128 | 1,31 | 1,55E+07 | 8,66E+06 |
| 43,8 | | 0,2942009 | 0,2514121 | 0,95 | 1,57E+08 | 1,42E+08 |
| 68,2 | | 0,2948525 | 0,2521731 | 1,12 | 1,26E+09 | 1,56E+09 |
| 40,3 | | 0,2951306 | 0,2525789 | 0,94 | 3,60E+08 | 3,26E+08 |
| 17,4 | | 0,2953011 | 0,2527526 | 0,92 | 7,01E+07 | 7,50E+07 |
| 47,2 | | 0,2957926 | 0,2531581 | 0,96 | 3,75E+08 | 4,03E+08 |
| 48,9 | | 0,2958071 | 0,2531121 | 0,92 | 1,00E+08 | 8,35E+07 |
| 3,6 | | 0,2959000 | 0,2531475 | 1,77 | 1,95E+06 | 3,73E+06 |
| 1,6 | | 0,2962236 | 0,2534845 | 0,32 | 3,40E+06 | 3,43E+07 |
| 67,8 | | 0,2965159 | 0,2537537 | 1,04 | 7,36E+09 | 6,82E+09 |
| 7,4 | | 0,2966455 | 0,2538500 | 0,75 | 9,16E+06 | 8,16E+06 |
| 20,2 | | 0,2967140 | 0,2538831 | 0,85 | 1,34E+07 | 9,39E+06 |
| 24,7 | | 0,2973339 | 0,2546277 | 1,06 | 1,28E+08 | 1,26E+08 |
| 51,4 | | 0,2980377 | 0,2557771 | 0,92 | 1,41E+08 | 1,63E+08 |
| 43,1 | | 0,2981501 | 0,2558432 | 0,27 | 5,74E+07 | 5,40E+07 |
| 69 | | 0,2982298 | 0,2559398 | 0,81 | 1,40E+08 | 1,16E+08 |
| 36,7 | | 0,2982980 | 0,2559463 | 1,16 | 1,62E+08 | 1,17E+08 |
| 39,5 | | 0,2984638 | 0,2560783 | 1,12 | 2,05E+09 | 1,59E+09 |
| 5,7 | | 0,2985665 | 0,2561139 | 0,84 | 2,77E+07 | 2,04E+07 |
| 11,4 | | 0,2991113 | 0,2568267 | 1,27 | 3,79E+06 | 5,31E+06 |
| 27,4 | | 0,2993651 | 0,2569976 | 0,91 | 2,63E+07 | 2,94E+07 |
| 7,1 | | 0,2993733 | 0,2569511 | 0,60 | 4,14E+06 | 4,79E+06 |
| 71,4 | | 0,2998029 | 0,2574684 | 0,91 | 6,45E+08 | 7,11E+08 |
| 69,1 | | 0,3000245 | 0,2577906 | 1,04 | 3,06E+09 | 2,96E+09 |
| 40,4 | | 0,3002848 | 0,2582163 | 1,09 | 8,03E+07 | 7,54E+07 |
| 13,3 | | 0,3003681 | 0,2582753 | 1,14 | 2,25E+07 | 1,70E+07 |
| 69,2 | | 0,3012366 | 0,2592214 | 0,89 | 6,29E+08 | 7,02E+08 |
| 53,1 | | 0,3016452 | 0,2595822 | 0,88 | 9,02E+08 | 1,20E+09 |
| 6,6 | | 0,3018523 | 0,2597922 | 0,82 | 8,18E+06 | 7,77E+06 |
| 32,9 | | 0,3021505 | 0,2601896 | 0,92 | 8,95E+07 | 1,02E+08 |
| 37,9 | | 0,3022140 | 0,2602251 | 0,94 | 2,25E+08 | 2,62E+08 |
| 23,1 | | 0,3024450 | 0,2605839 | 1,14 | 3,03E+07 | 2,60E+07 |
| 21,2 | | 0,3027154 | 0,2608168 | 0,86 | 4,54E+07 | 3,67E+07 |
| 39,4 | | 0,3030202 | 0,2612362 | 1,03 | 2,47E+08 | 2,28E+08 |
| 35,1 | | 0,3030920 | 0,2613372 | 1,22 | 8,39E+07 | 9,20E+07 |
| 5,8 | | 0,3032133 | 0,2616327 | 1,30 | 1,55E+07 | 3,27E+07 |
| 46,9 | | 0,3033273 | 0,2616844 | 0,82 | 3,54E+08 | 4,50E+08 |
| 37,8 | | 0,3034641 | 0,2619465 | 0,96 | 1,78E+08 | 1,76E+08 |
| 12 | | 0,3034848 | 0,2619216 | 0,81 | 2,78E+07 | 3,41E+07 |
| 15,1 | | 0,3035799 | 0,2619191 | 3,73 | 8,32E+07 | 6,82E+07 |
| 14,4 | | 0,3036517 | 0,2619332 | 0,58 | 5,28E+06 | 2,42E+07 |
| 37 | | 0,3038681 | 0,2621098 | 1,17 | 7,90E+07 | 5,63E+07 |
| 1,2 | | 0,3041120 | 0,2622964 | 0,66 | 7,15E+06 | 4,17E+06 |
| 76,8 | | 0,3045531 | 0,2631141 | 1,28 | 1,60E+08 | 1,81E+08 |
| 11,8 | | 0,3053297 | 0,2640339 | 1,10 | 5,83E+07 | 5,70E+07 |
| 19 | | 0,3053566 | 0,2640231 | 0,92 | 1,06E+08 | 1,15E+08 |
| 23 | | 0,3056854 | 0,2646134 | 0,42 | 1,08E+07 | 1,64E+07 |
| 5,5 | | 0,3057195 | 0,2646479 | 2,19 | 1,59E+07 | 1,64E+07 |
| 11,1 | | 0,3058267 | 0,2647899 | 1,37 | 2,10E+06 | 3,06E+06 |
| 15,9 | | 0,3058509 | 0,2647811 | 0,91 | 8,36E+07 | 7,22E+07 |

| | | | | | | |
|---|---|---|---|---|---|---|
| 44,5 | | 0,3059801 | 0,2649641 | 0,77 | 1,25E+08 | 1,21E+08 |
| 60 | | 0,3061319 | 0,2650541 | 0,94 | 2,52E+09 | 2,32E+09 |
| 34,9 | | 0,3063156 | 0,2652961 | 0,96 | 3,03E+08 | 2,94E+08 |
| 44,9 | | 0,3065514 | 0,2656519 | 0,89 | 1,98E+08 | 2,71E+08 |
| 21 | | 0,3073460 | 0,2664918 | 1,10 | 3,41E+08 | 3,06E+08 |
| 45,8 | | 0,3079472 | 0,2672709 | 1,14 | 8,35E+08 | 1,06E+09 |
| 23,8 | | 0,3080488 | 0,2674554 | 1,75 | 1,05E+07 | 4,53E+07 |
| 11,1 | | 0,3080769 | 0,2674073 | 0,87 | 1,61E+07 | 1,95E+07 |
| 57,2 | | 0,3084333 | 0,2679546 | 1,17 | 2,71E+08 | 3,25E+08 |
| 11,2 | | 0,3084518 | 0,2679525 | 1,14 | 3,22E+07 | 2,57E+07 |
| 64,1 | | 0,3095412 | 0,2696779 | 1,11 | 8,12E+08 | 9,80E+08 |
| 7,5 | | 0,3097042 | 0,2698172 | 0,86 | 2,53E+08 | 3,11E+08 |
| 16,1 | | 0,3098776 | 0,2699557 | 1,20 | 1,31E+07 | 1,22E+07 |
| 33,6 | | 0,3103316 | 0,2705223 | 0,94 | 3,36E+08 | 3,14E+08 |
| 14,7 | | 0,3104122 | 0,2706240 | 0,87 | 1,33E+07 | 1,28E+07 |
| 36,2 | | 0,3106774 | 0,2709896 | 0,85 | 1,53E+07 | 1,22E+07 |
| 61,3 | | 0,3110740 | 0,2716764 | 0,88 | 1,79E+09 | 1,84E+09 |
| 41,9 | | 0,3111066 | 0,2717017 | 1,21 | 4,42E+07 | 5,69E+07 |
| 5,4 | | 0,3115851 | 0,2724083 | 1,48 | 4,60E+06 | 1,75E+06 |
| 1,3 | | 0,3118919 | 0,2727999 | 0,42 | 2,35E+07 | 7,70E+06 |
| 4,4 | | 0,3124450 | 0,2733652 | 1,16 | 1,64E+06 | 2,13E+06 |
| 24,8 | | 0,3126979 | 0,2737651 | 1,12 | 5,83E+07 | 7,36E+07 |
| 49,3 | | 0,3129201 | 0,2739853 | 1,18 | 1,85E+08 | 2,23E+08 |
| 5,5 | | 0,3130047 | 0,2740338 | 1,48 | 2,87E+06 | 2,83E+06 |
| 30,7 | | 0,3132241 | 0,2742122 | 0,79 | 1,59E+08 | 2,91E+08 |
| 32,7 | | 0,3132551 | 0,2741823 | 1,08 | 3,90E+08 | 4,77E+08 |
| 18,5 | | 0,3136740 | 0,2745868 | 1,70 | 1,45E+07 | 1,32E+07 |
| 17,4 | | 0,3143431 | 0,2754876 | 0,83 | 2,21E+07 | 1,50E+07 |
| 19,1 | | 0,3143531 | 0,2754770 | 0,90 | 1,49E+07 | 1,80E+07 |
| 14 | | 0,3144433 | 0,2755481 | 1,09 | 1,15E+08 | 1,25E+08 |
| 23,1 | | 0,3144941 | 0,2754987 | 0,90 | 6,22E+07 | 7,71E+07 |
| 28,7 | | 0,3146289 | 0,2756651 | 1,11 | 1,28E+08 | 1,31E+08 |
| 12,1 | | 0,3146681 | 0,2756802 | 0,75 | 6,00E+06 | 5,46E+06 |
| 26,3 | | 0,3151329 | 0,2762681 | 1,05 | 3,24E+07 | 3,48E+07 |
| 31,5 | | 0,3152590 | 0,2764752 | 1,14 | 7,43E+07 | 6,31E+07 |
| 5,7 | | 0,3152870 | 0,2764450 | 0,89 | 6,09E+06 | 5,90E+06 |
| 11,7 | | 0,3153487 | 0,2764342 | 0,76 | 1,53E+07 | 1,82E+07 |
| 14 | | 0,3155265 | 0,2766082 | 0,85 | 1,25E+07 | 1,52E+07 |
| 4,5 | | 0,3155681 | 0,2765808 | 0,75 | 5,85E+06 | 4,04E+06 |
| 12,8 | | 0,3155820 | 0,2765507 | 0,87 | 1,06E+07 | 1,45E+07 |
| 31,3 | | 0,3156047 | 0,2765405 | 1,10 | 4,47E+07 | 4,11E+07 |
| 44,3 | | 0,3164488 | 0,2776886 | 1,10 | 1,23E+09 | 1,21E+09 |
| 8,2 | | 0,3165672 | 0,2778264 | 0,53 | 7,06E+06 | 8,12E+06 |
| 34,7 | | 0,3169662 | 0,2783936 | 3,18 | 6,41E+07 | 4,96E+07 |
| 4,5 | | 0,3171542 | 0,2786321 | 1,43 | 3,89E+06 | 1,13E+07 |
| 26,7 | | 0,3172939 | 0,2787732 | 1,32 | 1,66E+07 | 3,68E+07 |
| 70,2 | | 0,3177679 | 0,2793978 | 0,89 | 1,81E+09 | 2,44E+09 |
| 5,7 | | 0,3178157 | 0,2793650 | 0,51 | 1,11E+07 | 8,12E+06 |
| 13,9 | | 0,3181120 | 0,2798177 | 1,08 | 4,49E+07 | 3,87E+07 |
| 68,2 | | 0,3184869 | 0,2803595 | 0,95 | 1,10E+10 | 9,70E+09 |

| | | | | | | |
|---|---|---|---|---|---|---|
| 26,5 | | 0,3185904 | 0,2805467 | 0,86 | 7,27E+07 | 7,25E+07 |
| 28,2 | | 0,3186646 | 0,2805573 | 1,04 | 5,22E+08 | 5,09E+08 |
| 21,8 | | 0,3190061 | 0,2810673 | 0,58 | 1,56E+07 | 2,26E+07 |
| 5,4 | | 0,3190477 | 0,2810871 | 0,71 | 6,74E+06 | 5,22E+06 |
| 43,6 | | 0,3190480 | 0,2810370 | 0,91 | 2,41E+08 | 3,00E+08 |
| 36,4 | | 0,3192785 | 0,2814988 | 0,95 | 7,93E+08 | 8,26E+08 |
| 22,4 | | 0,3194520 | 0,2815877 | 1,23 | 3,10E+06 | 5,07E+06 |
| 51,7 | | 0,3201348 | 0,2824483 | 0,90 | 1,68E+08 | 1,92E+08 |
| 53,1 | | 0,3205363 | 0,2828153 | 0,94 | 4,85E+09 | 5,15E+09 |
| 16,8 | | 0,3206173 | 0,2829033 | 0,94 | 3,86E+07 | 4,35E+07 |
| 1,5 | | 0,3218638 | 0,2845065 | 0,56 | 8,84E+06 | 6,48E+06 |
| 45,5 | | 0,3219100 | 0,2845392 | 1,06 | 3,24E+09 | 3,25E+09 |
| 7,4 | | 0,3219661 | 0,2845056 | 0,40 | 2,32E+07 | 9,36E+06 |
| 19,9 | | 0,3220942 | 0,2847045 | 0,90 | 1,86E+07 | 1,97E+07 |
| 10,2 | | 0,3224079 | 0,2849553 | 0,84 | 5,54E+06 | 5,17E+06 |
| 26,8 | | 0,3224725 | 0,2849979 | 0,92 | 7,83E+07 | 6,33E+07 |
| 16 | | 0,3225867 | 0,2851637 | 0,76 | 6,70E+09 | 3,27E+09 |
| 55,4 | | 0,3227626 | 0,2853281 | 0,81 | 1,48E+08 | 1,36E+08 |
| 60,7 | | 0,3230349 | 0,2856940 | 1,06 | 6,19E+08 | 6,16E+08 |
| 33,5 | | 0,3230735 | 0,2857244 | 0,91 | 1,64E+08 | 1,75E+08 |
| 39 | | 0,3232645 | 0,2858957 | 1,16 | 6,73E+07 | 7,23E+07 |
| 46,5 | | 0,3232914 | 0,2858612 | 1,13 | 3,67E+08 | 4,92E+08 |
| 36,3 | | 0,3235715 | 0,2862723 | 1,07 | 1,53E+09 | 1,56E+09 |
| 39,2 | | 0,3236030 | 0,2862214 | 1,06 | 8,21E+07 | 7,27E+07 |
| 41,2 | | 0,3236377 | 0,2862281 | 1,51 | 2,65E+07 | 7,38E+07 |
| 7,1 | | 0,3236864 | 0,2862170 | 0,72 | 1,09E+07 | 1,05E+07 |
| 63,1 | | 0,3239325 | 0,2864720 | 0,91 | 1,24E+09 | 9,96E+08 |
| 6,9 | | 0,3243254 | 0,2869957 | 1,51 | 8,12E+06 | 9,44E+06 |
| 29,3 | | 0,3246261 | 0,2874020 | 1,09 | 4,71E+07 | 4,29E+07 |
| 11,6 | | 0,3246721 | 0,2874646 | 1,35 | 7,42E+06 | 3,42E+06 |
| 13,9 | | 0,3252671 | 0,2885410 | 1,41 | 5,96E+06 | 1,53E+07 |
| 39,9 | | 0,3261392 | 0,2898074 | 1,12 | 1,64E+08 | 1,99E+08 |
| 26,3 | | 0,3266136 | 0,2905255 | 1,11 | 1,77E+08 | 2,12E+08 |
| 45,6 | | 0,3266190 | 0,2904739 | 0,94 | 5,37E+08 | 5,09E+08 |
| 3,1 | | 0,3267715 | 0,2906539 | 0,33 | 1,27E+07 | 5,84E+07 |
| 2,5 | | 0,3267855 | 0,2906214 | 1,42 | 2,53E+06 | 3,05E+06 |
| 10,7 | | 0,3276022 | 0,2914113 | 1,74 | 2,11E+06 | 1,04E+07 |
| 79,2 | | 0,3282266 | 0,2922939 | 1,06 | 6,09E+09 | 6,71E+09 |
| 54 | | 0,3286157 | 0,2927979 | 1,08 | 4,06E+08 | 4,55E+08 |
| 8,9 | | 0,3288082 | 0,2929972 | 0,73 | 1,27E+07 | 1,74E+07 |
| 13,4 | | 0,3288566 | 0,2929452 | 1,31 | 5,43E+06 | 7,24E+06 |
| 19 | | 0,3290791 | 0,2932423 | 1,50 | 6,55E+06 | 6,51E+06 |
| 7,7 | | 0,3292223 | 0,2933385 | 3,53 | 4,70E+07 | 1,84E+09 |
| 29,1 | | 0,3293604 | 0,2934603 | 1,14 | 1,10E+08 | 1,25E+08 |
| 34,9 | | 0,3302563 | 0,2948605 | 1,22 | 1,69E+08 | 1,79E+08 |
| 17,2 | | 0,3304990 | 0,2951372 | 0,45 | 5,99E+07 | 4,61E+07 |
| 12,3 | | 0,3305026 | 0,2950849 | 1,19 | 4,59E+07 | 4,19E+07 |
| 38,2 | | 0,3308838 | 0,2957165 | 0,92 | 3,27E+07 | 3,62E+07 |
| 5 | | 0,3309139 | 0,2957088 | 0,88 | 3,48E+06 | 2,94E+06 |
| 49,8 | | 0,3310153 | 0,2957549 | 1,10 | 5,10E+09 | 5,15E+09 |

| | | | | | | |
|---|---|---|---|---|---|---|
| 52,9 | | 0,3313880 | 0,2962515 | 0,95 | 3,52E+08 | 3,69E+08 |
| 33,7 | | 0,3314064 | 0,2962535 | 1,12 | 1,57E+08 | 2,00E+08 |
| 15,4 | | 0,3315821 | 0,2965212 | 1,09 | 4,21E+07 | 3,30E+07 |
| 3,2 | | 0,3320152 | 0,2972779 | 1,27 | 8,03E+05 | 1,43E+06 |
| 6 | | 0,3321784 | 0,2975127 | 2,35 | 2,21E+06 | 2,37E+07 |
| 40,4 | | 0,3324377 | 0,2978485 | 1,05 | 1,31E+08 | 1,29E+08 |
| 4,2 | | 0,3329705 | 0,2986499 | 0,71 | 4,19E+06 | 5,68E+06 |
| 48,2 | | 0,3333902 | 0,2993215 | 1,05 | 4,04E+08 | 3,92E+08 |
| 5,8 | | 0,3336691 | 0,2995763 | 0,50 | 7,80E+06 | 7,71E+06 |
| 37,9 | | 0,3336761 | 0,2995424 | 0,98 | 4,21E+08 | 4,33E+08 |
| 27,1 | | 0,3337171 | 0,2995786 | 1,07 | 3,08E+07 | 3,54E+07 |
| 19,8 | | 0,3337181 | 0,2995256 | 0,81 | 2,33E+07 | 1,64E+07 |
| 48,3 | | 0,3346609 | 0,3007542 | 0,87 | 5,13E+08 | 4,26E+08 |
| 16,8 | | 0,3348224 | 0,3009625 | 0,88 | 3,33E+07 | 3,08E+07 |
| 52,5 | | 0,3351472 | 0,3014973 | 0,93 | 6,97E+07 | 8,18E+07 |
| 31,2 | | 0,3351765 | 0,3015161 | 0,99 | 4,96E+08 | 5,01E+08 |
| 19 | | 0,3352784 | 0,3015624 | 1,29 | 2,52E+07 | 3,52E+07 |
| 69,6 | | 0,3356191 | 0,3020318 | 0,95 | 9,72E+09 | 8,87E+09 |
| 34,5 | | 0,3360515 | 0,3025109 | 0,76 | 1,46E+07 | 1,90E+07 |
| 43,1 | | 0,3362379 | 0,3027575 | 1,14 | 1,39E+08 | 1,51E+08 |
| 38,4 | | 0,3364549 | 0,3030895 | 0,51 | 2,06E+07 | 1,03E+07 |
| 11,8 | | 0,3365634 | 0,3032265 | 1,25 | 2,85E+06 | 3,00E+06 |
| 20,9 | | 0,3365907 | 0,3032288 | 1,10 | 4,12E+07 | 3,71E+07 |
| 11,2 | | 0,3366242 | 0,3033101 | 0,82 | 6,07E+07 | 3,31E+07 |
| 22,7 | | 0,3367758 | 0,3035084 | 0,74 | 5,09E+06 | 2,70E+06 |
| 26,1 | | 0,3368425 | 0,3035719 | 1,11 | 9,16E+07 | 1,21E+08 |
| 9,4 | | 0,3368477 | 0,3035184 | 1,28 | 6,18E+06 | 3,49E+06 |
| 64,3 | | 0,3370514 | 0,3036680 | 0,91 | 1,95E+09 | 1,67E+09 |
| 49,2 | | 0,3377616 | 0,3045829 | 1,05 | 1,20E+09 | 1,30E+09 |
| 43,3 | | 0,3381275 | 0,3050085 | 0,95 | 4,42E+08 | 4,80E+08 |
| 19,2 | | 0,3388008 | 0,3057883 | 1,10 | 8,20E+07 | 6,06E+07 |
| 25,2 | | 0,3388905 | 0,3058852 | 0,92 | 3,05E+08 | 3,89E+08 |
| 23,3 | | 0,3390407 | 0,3060884 | 0,92 | 4,60E+07 | 4,73E+07 |
| 53,1 | | 0,3390543 | 0,3060507 | 1,10 | 1,13E+08 | 1,01E+08 |
| 46,2 | | 0,3393245 | 0,3062968 | 0,79 | 9,66E+07 | 8,35E+07 |
| 20,1 | | 0,3402076 | 0,3073939 | 0,81 | 4,62E+07 | 2,50E+07 |
| 15,6 | | 0,3406339 | 0,3081626 | 1,22 | 4,39E+07 | 7,56E+07 |
| 8,3 | | 0,3407039 | 0,3082125 | 1,30 | 1,41E+07 | 1,10E+07 |
| 32 | | 0,3413488 | 0,3089576 | 1,10 | 2,53E+07 | 2,30E+07 |
| 56 | | 0,3415452 | 0,3091706 | 0,76 | 1,58E+09 | 2,57E+09 |
| 46,5 | | 0,3417607 | 0,3095467 | 1,08 | 2,37E+08 | 2,74E+08 |
| 38,2 | | 0,3425150 | 0,3103987 | 1,11 | 4,15E+07 | 5,07E+07 |
| 9,2 | | 0,3428327 | 0,3107400 | 0,70 | 1,17E+07 | 5,84E+06 |
| 32,6 | | 0,3435172 | 0,3116555 | 1,05 | 1,84E+08 | 1,59E+08 |
| 10,1 | | 0,3439143 | 0,3122966 | 0,89 | 1,96E+07 | 1,45E+07 |
| 5,1 | | 0,3443448 | 0,3128285 | 0,76 | 5,84E+06 | 1,05E+07 |
| 3,9 | | 0,3446651 | 0,3132085 | 1,19 | 1,27E+07 | 1,04E+07 |
| 11,4 | | 0,3447515 | 0,3133874 | 0,67 | 4,11E+07 | 8,59E+07 |
| 40,8 | | 0,3448563 | 0,3134104 | 0,86 | 6,84E+08 | 6,39E+08 |
| 13,6 | | 0,3449590 | 0,3134726 | 1,10 | 2,74E+07 | 2,26E+07 |

| | | | | | | |
|---|---|---|---|---|---|---|
| 11,5 | | 0,3456136 | 0,3143999 | 0,78 | 6,64E+06 | 5,96E+06 |
| 12,2 | | 0,3459554 | 0,3148873 | 1,52 | 1,24E+07 | 4,55E+06 |
| 23,9 | | 0,3459761 | 0,3148321 | 1,20 | 1,31E+07 | 1,53E+07 |
| 49,4 | | 0,3462557 | 0,3152105 | 1,05 | 2,09E+08 | 1,97E+08 |
| 36,6 | | 0,3468287 | 0,3160856 | 0,95 | 4,52E+08 | 4,65E+08 |
| 30,8 | | 0,3472317 | 0,3164216 | 0,84 | 1,12E+08 | 1,21E+08 |
| 14,5 | | 0,3474335 | 0,3166186 | 0,29 | 1,60E+08 | 1,23E+08 |
| 49,8 | | 0,3478310 | 0,3172412 | 0,95 | 6,28E+08 | 5,90E+08 |
| 7,9 | | 0,3487573 | 0,3187611 | 0,50 | 3,06E+07 | 6,78E+06 |
| 1,6 | | 0,3487992 | 0,3187403 | 2,89 | 3,92E+07 | 5,26E+07 |
| 11,6 | | 0,3490644 | 0,3190692 | 0,81 | 1,57E+07 | 9,89E+06 |
| 36,1 | | 0,3491187 | 0,3190680 | 1,06 | 1,01E+08 | 9,60E+07 |
| 1,4 | | 0,3491755 | 0,3192118 | 2,40 | 9,26E+05 | 1,69E+05 |
| 17,6 | | 0,3496433 | 0,3197996 | 1,30 | 3,23E+07 | 2,72E+07 |
| 13,9 | | 0,3499223 | 0,3203495 | 0,63 | 8,46E+06 | 7,79E+06 |
| 77,5 | | 0,3500800 | 0,3205098 | 1,03 | 1,08E+10 | 1,03E+10 |
| 6,9 | | 0,3502051 | 0,3206792 | 1,67 | 1,84E+06 | 2,69E+06 |
| 21 | | 0,3504806 | 0,3211047 | 0,89 | 4,97E+07 | 5,04E+07 |
| 20,1 | | 0,3514940 | 0,3223048 | 0,76 | 1,21E+07 | 1,98E+07 |
| 35,7 | | 0,3515967 | 0,3223506 | 0,95 | 1,89E+08 | 1,82E+08 |
| 34,8 | | 0,3518446 | 0,3228288 | 1,07 | 1,35E+08 | 1,22E+08 |
| 16,7 | | 0,3519844 | 0,3229682 | 0,90 | 2,33E+07 | 2,13E+07 |
| 35,6 | | 0,3521489 | 0,3232341 | 0,91 | 5,70E+07 | 5,46E+07 |
| 42,6 | | 0,3522400 | 0,3232937 | 0,89 | 5,59E+08 | 6,90E+08 |
| 5,7 | | 0,3523265 | 0,3234449 | 1,28 | 3,57E+06 | 3,15E+06 |
| 26,3 | | 0,3523572 | 0,3234484 | 1,08 | 7,90E+07 | 7,03E+07 |
| 18,8 | | 0,3524162 | 0,3234877 | 1,06 | 1,17E+08 | 1,10E+08 |
| 14 | | 0,3524748 | 0,3235590 | 0,56 | 1,14E+07 | 1,16E+07 |
| 7,3 | | 0,3526169 | 0,3236521 | 1,28 | 7,78E+06 | 5,34E+06 |
| 32,6 | | 0,3530204 | 0,3242487 | 0,77 | 4,56E+07 | 3,40E+07 |
| 69,9 | | 0,3543155 | 0,3264187 | 0,89 | 2,42E+08 | 2,97E+08 |
| 60,4 | | 0,3547169 | 0,3269176 | 0,96 | 1,14E+09 | 1,18E+09 |
| 40,8 | | 0,3548898 | 0,3270574 | 1,06 | 1,76E+08 | 1,52E+08 |
| 14,4 | | 0,3549356 | 0,3270220 | 0,69 | 2,92E+07 | 2,26E+07 |
| 17,8 | | 0,3549673 | 0,3270208 | 0,42 | 1,07E+07 | 9,80E+06 |
| 7,3 | | 0,3549867 | 0,3270230 | 0,83 | 3,90E+06 | 4,39E+06 |
| 15 | | 0,3553505 | 0,3276532 | 0,76 | 1,98E+08 | 1,98E+08 |
| 9,5 | | 0,3555793 | 0,3278640 | 1,60 | 8,86E+06 | 2,69E+06 |
| 26,8 | | 0,3560107 | 0,3284359 | 0,89 | 1,05E+08 | 1,19E+08 |
| 21,1 | | 0,3561223 | 0,3285446 | 0,58 | 2,04E+07 | 1,75E+07 |
| 74,2 | | 0,3568886 | 0,3297263 | 2,81 | 2,03E+06 | 4,70E+07 |
| 8,4 | | 0,3572446 | 0,3301241 | 0,84 | 6,69E+06 | 1,18E+07 |
| 50,7 | | 0,3574820 | 0,3303614 | 0,91 | 8,77E+08 | 8,45E+08 |
| 54,1 | | 0,3574993 | 0,3303206 | 1,05 | 4,71E+08 | 4,25E+08 |
| 38,1 | | 0,3576140 | 0,3304365 | 0,95 | 2,44E+08 | 2,17E+08 |
| 10,4 | | 0,3578733 | 0,3306318 | 1,21 | 2,49E+07 | 2,04E+07 |
| 56,6 | | 0,3578764 | 0,3305743 | 0,83 | 1,83E+07 | 2,82E+07 |
| 10,1 | | 0,3583645 | 0,3313175 | 1,16 | 3,21E+07 | 3,71E+07 |
| 2,9 | | 0,3584037 | 0,3313142 | 0,70 | 1,84E+06 | 2,17E+06 |
| 7,3 | | 0,3584226 | 0,3313561 | 0,90 | 3,72E+06 | 3,44E+06 |

| | | | | | | |
|---|---|---|---|---|---|---|
| 20,4 | | 0,3586333 | 0,3316993 | 0,90 | 2,06E+07 | 2,11E+07 |
| 7,5 | | 0,3586484 | 0,3316792 | 0,65 | 5,79E+06 | 2,89E+06 |
| 6,1 | | 0,3586861 | 0,3317746 | 0,20 | 2,01E+08 | 1,82E+08 |
| 38,2 | | 0,3588663 | 0,3319965 | 1,03 | 1,51E+08 | 1,48E+08 |
| 36,8 | | 0,3593349 | 0,3326712 | 0,95 | 4,39E+08 | 4,82E+08 |
| 28,8 | | 0,3600306 | 0,3337928 | 1,09 | 9,31E+07 | 1,23E+08 |
| 11,2 | | 0,3601325 | 0,3338696 | 0,85 | 1,00E+07 | 1,05E+07 |
| 8,8 | | 0,3601370 | 0,3338332 | 0,90 | 1,62E+07 | 1,31E+07 |
| 43,2 | | 0,3607377 | 0,3346023 | 0,93 | 7,25E+08 | 7,96E+08 |
| 50,2 | | 0,3611411 | 0,3350042 | 0,91 | 4,79E+08 | 4,32E+08 |
| 18,7 | | 0,3613055 | 0,3351308 | 1,07 | 4,27E+07 | 3,76E+07 |
| 49,5 | | 0,3620267 | 0,3361132 | 1,14 | 1,28E+08 | 1,43E+08 |
| 35,3 | | 0,3620392 | 0,3360764 | 0,93 | 1,48E+08 | 1,80E+08 |
| 10,8 | | 0,3622832 | 0,3366451 | 0,79 | 1,56E+07 | 9,75E+06 |
| 48,9 | | 0,3630136 | 0,3377893 | 0,93 | 2,83E+08 | 2,71E+08 |
| 3,3 | | 0,3635043 | 0,3384748 | 1,16 | 1,08E+07 | 1,10E+07 |
| 38 | | 0,3635418 | 0,3385133 | 0,85 | 7,16E+07 | 4,78E+07 |
| 12,3 | | 0,3635959 | 0,3385524 | 0,61 | 5,80E+06 | 8,12E+06 |
| 5,4 | | 0,3636378 | 0,3385748 | 1,71 | 3,58E+06 | 1,25E+06 |
| 27,5 | | 0,3642272 | 0,3393687 | 1,11 | 8,20E+07 | 1,11E+08 |
| 7 | | 0,3643423 | 0,3394819 | 0,89 | 4,76E+07 | 5,19E+07 |
| 28,5 | | 0,3645057 | 0,3396956 | 1,09 | 1,14E+09 | 9,66E+08 |
| 37,5 | | 0,3646906 | 0,3399549 | 0,93 | 1,66E+08 | 1,45E+08 |
| 4,4 | | 0,3650087 | 0,3402364 | 1,12 | 1,35E+07 | 1,14E+07 |
| 24,7 | | 0,3654854 | 0,3408227 | 1,02 | 3,44E+07 | 3,34E+07 |
| 45,9 | | 0,3662474 | 0,3416847 | 1,08 | 8,23E+08 | 6,77E+08 |
| 9,3 | | 0,3664931 | 0,3421341 | 0,79 | 1,22E+08 | 7,64E+07 |
| 28,2 | | 0,3667078 | 0,3425120 | 1,21 | 7,41E+07 | 1,05E+08 |
| 17 | | 0,3667226 | 0,3424914 | 1,17 | 1,70E+07 | 1,94E+07 |
| 58,6 | | 0,3670365 | 0,3429398 | 1,07 | 6,21E+09 | 7,40E+09 |
| 18,6 | | 0,3675200 | 0,3435513 | 0,94 | 5,88E+07 | 6,67E+07 |
| 22,3 | | 0,3679842 | 0,3442838 | 0,96 | 7,49E+07 | 6,95E+07 |
| 21,9 | | 0,3683264 | 0,3448770 | 1,39 | 2,79E+07 | 2,98E+07 |
| 23,8 | | 0,3683954 | 0,3449377 | 0,88 | 5,88E+06 | 8,03E+06 |
| 7,3 | | 0,3684456 | 0,3449348 | 0,67 | 7,10E+06 | 7,59E+06 |
| 38 | | 0,3685936 | 0,3451290 | 1,04 | 1,13E+08 | 1,18E+08 |
| 16,3 | | 0,3690553 | 0,3456268 | 0,80 | 1,49E+07 | 1,20E+07 |
| 35,7 | | 0,3692204 | 0,3458831 | 0,83 | 4,40E+07 | 3,23E+07 |
| 54,8 | | 0,3697486 | 0,3465978 | 0,96 | 3,17E+08 | 2,85E+08 |
| 63,4 | | 0,3699305 | 0,3467356 | 1,09 | 7,06E+08 | 5,42E+08 |
| 6,2 | | 0,3701092 | 0,3468809 | 0,80 | 8,82E+06 | 1,46E+07 |
| 17,3 | | 0,3705908 | 0,3476524 | 1,15 | 3,92E+07 | 3,48E+07 |
| 5,1 | | 0,3708148 | 0,3479081 | 1,34 | 4,02E+06 | 9,26E+06 |
| 7,2 | | 0,3708851 | 0,3479199 | 0,77 | 4,91E+06 | 4,53E+06 |
| 12 | | 0,3718213 | 0,3495293 | 0,81 | 9,06E+06 | 8,12E+06 |
| 57,3 | | 0,3726741 | 0,3506395 | 0,85 | 3,38E+08 | 2,71E+08 |
| 57,3 | | 0,3734687 | 0,3518163 | 0,95 | 4,29E+10 | 3,70E+10 |
| 24,8 | | 0,3735804 | 0,3519096 | 1,06 | 1,90E+08 | 1,65E+08 |
| 2,1 | | 0,3743076 | 0,3528456 | 1,57 | 1,12E+06 | 1,11E+06 |
| 15,8 | | 0,3744336 | 0,3529268 | 1,13 | 1,62E+07 | 1,12E+07 |

| | | | | | | |
|---|---|---|---|---|---|---|
| 4 | | 0,3749117 | 0,3537637 | 0,53 | 4,42E+06 | 1,19E+06 |
| 25,6 | | 0,3750174 | 0,3539179 | 1,46 | 3,83E+06 | 1,08E+07 |
| 4,5 | | 0,3752088 | 0,3542411 | 0,66 | 5,97E+06 | 7,83E+06 |
| 54,9 | | 0,3752525 | 0,3541986 | 1,08 | 8,15E+09 | 1,01E+10 |
| 28,7 | | 0,3752869 | 0,3541534 | 1,13 | 2,31E+07 | 1,72E+07 |
| 41,6 | | 0,3754298 | 0,3543633 | 1,06 | 2,54E+08 | 2,47E+08 |
| 3,5 | | 0,3754338 | 0,3543023 | 0,67 | 1,21E+06 | 2,45E+06 |
| 59,4 | | 0,3759964 | 0,3552150 | 0,97 | 7,13E+08 | 7,35E+08 |
| 12,8 | | 0,3763062 | 0,3556341 | 1,87 | 6,19E+06 | 5,96E+06 |
| 31,1 | | 0,3763305 | 0,3555894 | 0,63 | 8,19E+06 | 3,65E+06 |
| 20,8 | | 0,3766758 | 0,3559683 | 0,77 | 2,39E+07 | 2,04E+07 |
| 37,5 | | 0,3767203 | 0,3559966 | 0,95 | 8,97E+07 | 9,33E+07 |
| 8,6 | | 0,3767367 | 0,3560207 | 0,68 | 2,19E+06 | 4,57E+06 |
| 18,2 | | 0,3772430 | 0,3567029 | 0,76 | 4,96E+07 | 5,64E+07 |
| 18,6 | + | 0,3777022 | 0,3574834 | 0,46 | 1,74E+09 | 1,25E+09 |
| 18 | | 0,3777048 | 0,3574219 | 0,87 | 9,63E+06 | 1,06E+07 |
| 26,6 | | 0,3778187 | 0,3576178 | 1,04 | 1,55E+08 | 1,40E+08 |
| 2,2 | | 0,3781199 | 0,3580764 | 0,82 | 3,54E+06 | 2,51E+06 |
| 12 | | 0,3781990 | 0,3582205 | 2,11 | 3,45E+06 | 1,99E+07 |
| 8,3 | | 0,3786596 | 0,3586706 | 2,55 | 1,93E+06 | 1,92E+06 |
| 37,3 | | 0,3787500 | 0,3587251 | 1,86 | 2,43E+07 | 2,18E+07 |
| 11,9 | | 0,3788223 | 0,3587625 | 0,57 | 1,10E+07 | 1,72E+06 |
| 25,2 | | 0,3799025 | 0,3602145 | 0,95 | 4,32E+07 | 4,44E+07 |
| 62,1 | | 0,3806049 | 0,3613003 | 1,10 | 2,51E+08 | 2,37E+08 |
| 25,8 | | 0,3807059 | 0,3615193 | 0,94 | 1,21E+08 | 1,23E+08 |
| 37,7 | | 0,3813884 | 0,3628492 | 1,07 | 5,02E+08 | 5,27E+08 |
| 2,3 | | 0,3818776 | 0,3638743 | 0,65 | 5,03E+06 | 3,99E+06 |
| 12,4 | | 0,3819625 | 0,3638942 | 0,93 | 8,84E+06 | 8,01E+06 |
| 73,9 | | 0,3820937 | 0,3640673 | 0,89 | 1,89E+09 | 2,27E+09 |
| 21,3 | | 0,3822608 | 0,3642540 | 1,21 | 1,59E+07 | 2,65E+07 |
| 23 | | 0,3823928 | 0,3643281 | 1,06 | 3,81E+07 | 4,57E+07 |
| 59,5 | | 0,3827836 | 0,3648188 | 0,97 | 4,60E+09 | 5,16E+09 |
| 17,2 | | 0,3830936 | 0,3651741 | 0,95 | 1,23E+08 | 1,17E+08 |
| 20,7 | | 0,3831784 | 0,3652370 | 1,20 | 7,19E+06 | 1,20E+07 |
| 30,4 | | 0,3832263 | 0,3652094 | 2,76 | 3,11E+06 | 5,58E+07 |
| 47,4 | | 0,3833373 | 0,3652771 | 1,05 | 5,06E+09 | 4,60E+09 |
| 5,9 | | 0,3842556 | 0,3666504 | 0,54 | 4,90E+06 | 9,56E+05 |
| 2,5 | | 0,3842907 | 0,3666308 | 1,20 | 6,62E+06 | 9,40E+06 |
| 18 | | 0,3843037 | 0,3665865 | 1,06 | 2,20E+07 | 2,49E+07 |
| 6,9 | | 0,3846712 | 0,3672344 | 1,57 | 4,09E+06 | 1,64E+06 |
| 62,1 | | 0,3847328 | 0,3672489 | 0,92 | 4,25E+08 | 4,96E+08 |
| 60,3 | | 0,3853055 | 0,3679548 | 0,72 | 8,64E+06 | 8,26E+06 |
| 10 | | 0,3854964 | 0,3682076 | 1,31 | 8,91E+06 | 1,23E+07 |
| 33,8 | | 0,3855233 | 0,3681877 | 1,07 | 8,29E+07 | 8,77E+07 |
| 57,2 | | 0,3857766 | 0,3685759 | 1,07 | 2,83E+09 | 2,56E+09 |
| 58,2 | | 0,3860766 | 0,3691414 | 0,95 | 5,02E+09 | 4,36E+09 |
| 8,6 | | 0,3867707 | 0,3704063 | 1,18 | 1,66E+07 | 9,34E+06 |
| 44 | | 0,3868730 | 0,3704641 | 1,08 | 2,20E+08 | 1,72E+08 |
| 62,9 | | 0,3872674 | 0,3711446 | 0,93 | 2,09E+09 | 2,56E+09 |
| 23,9 | | 0,3874437 | 0,3712911 | 1,07 | 1,35E+08 | 1,29E+08 |

| | | | | | | |
|---|---|---|---|---|---|---|
| 24 | | 0,3876388 | 0,3714931 | 1,06 | 4,36E+07 | 4,20E+07 |
| 14,4 | | 0,3877044 | 0,3715226 | 0,77 | 1,26E+07 | 1,83E+07 |
| 57,4 | | 0,3880745 | 0,3722476 | 0,74 | 2,07E+08 | 2,28E+08 |
| 16,3 | | 0,3881379 | 0,3722797 | 0,76 | 2,10E+07 | 8,93E+06 |
| 22,5 | | 0,3888693 | 0,3734555 | 0,75 | 1,04E+07 | 8,35E+06 |
| 10,2 | | 0,3894843 | 0,3744046 | 0,80 | 2,21E+07 | 1,24E+07 |
| 44,3 | | 0,3900024 | 0,3749756 | 1,18 | 2,24E+07 | 2,18E+07 |
| 31,3 | | 0,3900726 | 0,3749621 | 0,92 | 2,08E+08 | 2,15E+08 |
| 42,7 | | 0,3900775 | 0,3748980 | 1,05 | 4,14E+08 | 3,74E+08 |
| 5,4 | | 0,3902157 | 0,3752090 | 1,22 | 8,19E+06 | 5,03E+06 |
| 36,5 | | 0,3908179 | 0,3760779 | 0,80 | 1,91E+08 | 2,55E+08 |
| 31,8 | | 0,3912067 | 0,3767784 | 0,93 | 5,84E+08 | 7,13E+08 |
| 14,9 | | 0,3914815 | 0,3771190 | 0,70 | 5,28E+06 | 4,95E+06 |
| 30,4 | | 0,3915942 | 0,3772096 | 0,91 | 1,57E+08 | 1,79E+08 |
| 38,7 | | 0,3917609 | 0,3773991 | 1,09 | 1,83E+07 | 1,64E+07 |
| 32,5 | | 0,3919586 | 0,3776936 | 1,22 | 5,76E+07 | 4,15E+07 |
| 48,5 | | 0,3920118 | 0,3777268 | 0,89 | 1,94E+08 | 1,72E+08 |
| 31,5 | | 0,3920470 | 0,3777210 | 1,07 | 1,39E+08 | 1,55E+08 |
| 11,7 | | 0,3924296 | 0,3782479 | 0,72 | 8,03E+06 | 7,81E+06 |
| 11,5 | | 0,3924345 | 0,3781834 | 0,90 | 5,92E+06 | 5,14E+06 |
| 24,9 | | 0,3925874 | 0,3782649 | 0,90 | 1,09E+08 | 1,37E+08 |
| 30,3 | | 0,3925967 | 0,3782004 | 1,07 | 3,91E+07 | 4,13E+07 |
| 63,5 | | 0,3928469 | 0,3784699 | 1,06 | 4,65E+08 | 5,57E+08 |
| 40,2 | | 0,3932298 | 0,3790753 | 0,94 | 1,93E+08 | 1,79E+08 |
| 60,6 | | 0,3932410 | 0,3790107 | 0,94 | 1,96E+09 | 1,78E+09 |
| 51,4 | | 0,3937454 | 0,3797268 | 1,22 | 1,28E+08 | 9,56E+07 |
| 45,3 | | 0,3938315 | 0,3798985 | 1,04 | 4,57E+08 | 4,04E+08 |
| 13 | | 0,3940350 | 0,3802445 | 1,06 | 4,89E+07 | 5,24E+07 |
| 17,5 | | 0,3942670 | 0,3805896 | 0,96 | 3,57E+07 | 3,98E+07 |
| 44,9 | | 0,3946192 | 0,3810735 | 1,03 | 1,07E+09 | 1,10E+09 |
| 29,6 | | 0,3946714 | 0,3810849 | 1,05 | 1,40E+08 | 1,53E+08 |
| 19,4 | | 0,3948700 | 0,3813753 | 1,08 | 3,48E+07 | 3,76E+07 |
| 1,5 | | 0,3957675 | 0,3828311 | 1,80 | 1,52E+06 | 6,07E+06 |
| 1,4 | | 0,3958802 | 0,3829776 | 0,53 | 2,25E+07 | 7,78E+06 |
| 30,8 | | 0,3961557 | 0,3832981 | 0,94 | 3,48E+08 | 3,58E+08 |
| 27,4 | | 0,3966136 | 0,3840408 | 0,88 | 4,45E+06 | 6,37E+06 |
| 49,6 | | 0,3972858 | 0,3848479 | 0,92 | 5,47E+08 | 6,89E+08 |
| 23,2 | | 0,3976888 | 0,3855724 | 1,15 | 9,74E+07 | 1,01E+08 |
| 8,3 | | 0,3979627 | 0,3858372 | 0,78 | 6,47E+06 | 5,80E+06 |
| 47,2 | | 0,3980719 | 0,3860020 | 0,80 | 8,75E+08 | 1,35E+09 |
| 66,8 | | 0,3983586 | 0,3863965 | 0,90 | 3,10E+08 | 3,79E+08 |
| 12,9 | | 0,3984882 | 0,3865435 | 1,43 | 2,63E+07 | 7,49E+06 |
| 7,2 | | 0,3987721 | 0,3869852 | 1,21 | 4,38E+06 | 2,75E+06 |
| 18,9 | | 0,3987843 | 0,3869195 | 1,37 | 2,65E+06 | 7,41E+06 |
| 28,3 | | 0,3992332 | 0,3876028 | 0,93 | 7,16E+07 | 5,76E+07 |
| 8,6 | | 0,3992701 | 0,3875947 | 0,41 | 2,19E+07 | 1,65E+07 |
| 18,5 | | 0,3995661 | 0,3880645 | 0,66 | 9,62E+06 | 7,25E+06 |
| 17,1 | | 0,3999141 | 0,3886155 | 2,00 | 1,29E+07 | 1,58E+06 |
| 8,9 | | 0,4016621 | 0,3914504 | 0,80 | 7,65E+06 | 3,90E+06 |
| 5,2 | | 0,4016779 | 0,3914396 | 0,63 | 3,93E+06 | 5,05E+06 |

| | | | | | | |
|---|---|---|---|---|---|---|
| 20,5 | | 0,4017814 | 0,3914628 | 0,98 | 1,70E+07 | 1,70E+07 |
| 9 | | 0,4034521 | 0,3943934 | 0,44 | 3,56E+06 | 2,88E+07 |
| 29 | | 0,4039690 | 0,3950744 | 0,91 | 4,96E+07 | 5,62E+07 |
| 60 | | 0,4040354 | 0,3950888 | 0,93 | 4,33E+08 | 3,36E+08 |
| 13,3 | | 0,4041709 | 0,3952035 | 0,95 | 2,76E+07 | 2,94E+07 |
| 3,9 | | 0,4046025 | 0,3957953 | 0,80 | 7,29E+06 | 1,36E+07 |
| 32,4 | | 0,4047856 | 0,3961010 | 0,78 | 3,28E+07 | 5,99E+07 |
| 15,8 | | 0,4049698 | 0,3963686 | 1,19 | 3,48E+07 | 5,11E+07 |
| 20,2 | | 0,4051081 | 0,3965067 | 1,04 | 8,28E+07 | 7,25E+07 |
| 10,6 | | 0,4059022 | 0,3978842 | 0,89 | 1,49E+07 | 1,26E+07 |
| 6 | | 0,4060398 | 0,3980992 | 1,77 | 4,71E+06 | 1,12E+07 |
| 15,5 | | 0,4060468 | 0,3980318 | 0,59 | 6,59E+06 | 4,70E+06 |
| 12,6 | | 0,4063368 | 0,3984383 | 1,20 | 1,90E+07 | 2,55E+07 |
| 35,6 | | 0,4064975 | 0,3986782 | 0,88 | 1,87E+08 | 1,75E+08 |
| 6,9 | | 0,4068513 | 0,3994613 | 0,54 | 5,04E+06 | 7,22E+05 |
| 19,6 | | 0,4072182 | 0,4001258 | 0,52 | 6,61E+06 | 8,79E+06 |
| 16,4 | | 0,4072202 | 0,4000791 | 1,29 | 2,19E+07 | 4,13E+07 |
| 25 | | 0,4073727 | 0,4002996 | 1,30 | 1,92E+06 | 3,59E+06 |
| 7,7 | | 0,4076545 | 0,4006343 | 0,53 | 6,04E+05 | 4,17E+06 |
| 18,4 | | 0,4078521 | 0,4009520 | 1,10 | 3,76E+07 | 3,70E+07 |
| 11,1 | | 0,4080711 | 0,4012892 | 0,92 | 2,38E+06 | 2,61E+06 |
| 15,1 | | 0,4081174 | 0,4012822 | 1,15 | 6,22E+06 | 4,87E+06 |
| 10,1 | | 0,4088383 | 0,4024923 | 0,71 | 2,58E+06 | 5,87E+06 |
| 63,2 | | 0,4090598 | 0,4027797 | 0,91 | 1,15E+09 | 1,26E+09 |
| 13,2 | | 0,4096419 | 0,4036041 | 0,41 | 2,97E+07 | 2,66E+07 |
| 25,5 | | 0,4096891 | 0,4036285 | 0,62 | 5,84E+06 | 9,44E+06 |
| 34,7 | | 0,4103269 | 0,4045072 | 1,10 | 2,07E+07 | 2,25E+07 |
| 25,1 | | 0,4105798 | 0,4050756 | 1,07 | 2,08E+07 | 2,54E+07 |
| 14,3 | | 0,4106639 | 0,4051362 | 1,07 | 3,08E+07 | 2,75E+07 |
| 28,7 | | 0,4109211 | 0,4053750 | 0,88 | 3,82E+07 | 3,94E+07 |
| 19,6 | | 0,4114979 | 0,4060631 | 0,87 | 2,56E+07 | 1,66E+07 |
| 3,3 | | 0,4115254 | 0,4060493 | 0,70 | 3,91E+06 | 4,92E+06 |
| 57,1 | | 0,4116022 | 0,4060604 | 0,88 | 1,23E+08 | 1,05E+08 |
| 37,2 | | 0,4117602 | 0,4062145 | 1,11 | 1,58E+08 | 1,23E+08 |
| 25,4 | | 0,4123303 | 0,4072453 | 0,77 | 9,65E+07 | 6,40E+07 |
| 46 | | 0,4123503 | 0,4071949 | 0,93 | 5,84E+08 | 7,42E+08 |
| 22,8 | | 0,4123503 | 0,4071262 | 1,16 | 3,04E+07 | 2,29E+07 |
| 9,4 | | 0,4123848 | 0,4071837 | 1,05 | 7,29E+06 | 8,57E+06 |
| 13 | | 0,4127872 | 0,4076664 | 1,16 | 1,26E+07 | 2,17E+07 |
| 31,4 | | 0,4128346 | 0,4076840 | 1,09 | 1,12E+08 | 1,17E+08 |
| 7,9 | | 0,4131132 | 0,4080303 | 1,16 | 1,27E+07 | 1,89E+07 |
| 29,5 | | 0,4133844 | 0,4084406 | 0,92 | 1,15E+09 | 8,57E+08 |
| 54 | | 0,4136174 | 0,4086345 | 0,97 | 1,36E+09 | 1,37E+09 |
| 22,4 | | 0,4136834 | 0,4086646 | 0,88 | 2,22E+07 | 2,49E+07 |
| 26,8 | | 0,4145536 | 0,4099088 | 0,90 | 5,13E+07 | 3,97E+07 |
| 94,1 | | 0,4149951 | 0,4105426 | 1,40 | 3,21E+09 | 5,39E+09 |
| 65,4 | | 0,4150133 | 0,4104735 | 0,86 | 1,93E+09 | 2,69E+09 |
| 9 | | 0,4151605 | 0,4107436 | 0,75 | 4,73E+06 | 2,65E+06 |
| 30,9 | | 0,4155162 | 0,4113985 | 0,90 | 3,36E+08 | 5,03E+08 |
| 5,4 | | 0,4156656 | 0,4114672 | 0,59 | 1,30E+07 | 1,01E+07 |

| | | | | | | |
|---|---|---|---|---|---|---|
| 42,3 | | 0,4158077 | 0,4115641 | 0,91 | 2,51E+08 | 2,87E+08 |
| 8,2 | | 0,4158584 | 0,4115494 | 0,77 | 8,94E+06 | 9,34E+06 |
| 20,4 | | 0,4162924 | 0,4121540 | 0,73 | 2,01E+07 | 2,24E+07 |
| 22 | | 0,4166639 | 0,4128867 | 2,23 | 1,28E+07 | 2,18E+07 |
| 21,6 | | 0,4171004 | 0,4135144 | 0,58 | 1,40E+07 | 2,18E+06 |
| 55,3 | | 0,4173179 | 0,4138501 | 0,87 | 2,24E+09 | 2,24E+09 |
| 43,4 | | 0,4173836 | 0,4140245 | 1,10 | 6,65E+07 | 5,84E+07 |
| 11 | | 0,4179510 | 0,4149157 | 1,54 | 4,36E+06 | 1,27E+07 |
| 5,4 | | 0,4180059 | 0,4149192 | 1,09 | 1,28E+07 | 1,52E+07 |
| 3,2 | | 0,4182279 | 0,4151478 | 1,35 | 3,86E+06 | 2,31E+06 |
| 48,1 | | 0,4189252 | 0,4166816 | 0,92 | 1,27E+08 | 9,52E+07 |
| 22,1 | | 0,4189375 | 0,4166485 | 1,21 | 1,65E+08 | 2,10E+08 |
| 8,4 | | 0,4190863 | 0,4168565 | 1,18 | 5,49E+06 | 3,59E+06 |
| 24,5 | | 0,4196597 | 0,4177436 | 0,80 | 1,85E+07 | 1,59E+07 |
| 16,2 | | 0,4198521 | 0,4179500 | 1,29 | 1,09E+07 | 2,79E+07 |
| 45,5 | | 0,4199164 | 0,4179752 | 0,88 | 6,89E+07 | 6,39E+07 |
| 27,4 | | 0,4206259 | 0,4191856 | 0,91 | 1,18E+07 | 1,25E+07 |
| 6,1 | | 0,4214282 | 0,4201616 | 0,82 | 3,42E+06 | 3,95E+06 |
| 51,2 | | 0,4215288 | 0,4202407 | 1,04 | 2,49E+08 | 2,61E+08 |
| 21,4 | | 0,4221074 | 0,4209507 | 1,12 | 4,95E+07 | 4,82E+07 |
| 12,6 | | 0,4221444 | 0,4209787 | 1,10 | 1,39E+07 | 1,92E+07 |
| 24,1 | | 0,4222973 | 0,4212218 | 0,79 | 2,76E+07 | 1,82E+07 |
| 50,8 | | 0,4226561 | 0,4218683 | 0,95 | 1,09E+10 | 1,05E+10 |
| 49,8 | | 0,4227554 | 0,4219471 | 1,11 | 3,10E+08 | 3,39E+08 |
| 32,8 | | 0,4228119 | 0,4219762 | 1,09 | 3,93E+07 | 5,25E+07 |
| 10,2 | | 0,4235874 | 0,4231440 | 1,13 | 1,70E+07 | 1,51E+07 |
| 28,4 | | 0,4246149 | 0,4248237 | 0,83 | 2,91E+07 | 3,13E+07 |
| 36,7 | | 0,4246525 | 0,4248316 | 1,07 | 2,38E+08 | 2,05E+08 |
| 0,7 | | 0,4250168 | 0,4253115 | 1,08 | 1,11E+08 | 1,21E+08 |
| 23,7 | | 0,4253212 | 0,4256914 | 0,94 | 1,07E+08 | 9,68E+07 |
| 11,3 | | 0,4256035 | 0,4261482 | 0,86 | 1,92E+07 | 1,18E+07 |
| 45,4 | | 0,4258075 | 0,4263620 | 0,87 | 5,40E+07 | 3,52E+07 |
| 3,8 | | 0,4258872 | 0,4263475 | 1,29 | 6,55E+08 | 8,74E+08 |
| 5,6 | | 0,4259399 | 0,4263672 | 1,36 | 6,06E+06 | 2,05E+06 |
| 9,1 | | 0,4265356 | 0,4273600 | 1,16 | 9,60E+06 | 1,39E+07 |
| 15,1 | | 0,4267294 | 0,4275627 | 1,25 | 8,93E+06 | 1,75E+07 |
| 18,2 | | 0,4269338 | 0,4278168 | 0,70 | 5,41E+06 | 9,88E+06 |
| 19,3 | | 0,4269342 | 0,4277453 | 0,79 | 4,95E+07 | 6,02E+07 |
| 46,2 | | 0,4271325 | 0,4281450 | 1,03 | 1,70E+08 | 1,53E+08 |
| 4,3 | | 0,4276141 | 0,4287070 | 1,15 | 5,59E+06 | 9,36E+06 |
| 7,4 | | 0,4278292 | 0,4290102 | 0,78 | 2,92E+07 | 4,30E+07 |
| 60 | | 0,4279459 | 0,4291804 | 1,13 | 8,37E+08 | 8,02E+08 |
| 6,3 | | 0,4279467 | 0,4291087 | 1,77 | 2,92E+06 | 2,05E+07 |
| 12,1 | | 0,4287298 | 0,4303740 | 1,38 | 2,63E+06 | 1,93E+06 |
| 6,5 | | 0,4291240 | 0,4311320 | 1,55 | 1,88E+06 | 2,68E+06 |
| 11,2 | | 0,4291722 | 0,4312577 | 1,11 | 3,78E+07 | 2,89E+07 |
| 5,7 | | 0,4295202 | 0,4319960 | 0,81 | 2,82E+06 | 2,98E+06 |
| 34 | | 0,4301781 | 0,4330497 | 0,89 | 6,89E+07 | 1,03E+08 |
| 24,6 | | 0,4303638 | 0,4332517 | 0,90 | 1,73E+07 | 1,87E+07 |
| 12,9 | | 0,4307758 | 0,4338239 | 0,90 | 4,07E+07 | 3,88E+07 |

| | | | | | | |
|---|---|---|---|---|---|---|
| 10,9 | | 0,4308795 | 0,4339263 | 1,11 | 1,32E+07 | 1,19E+07 |
| 26,7 | | 0,4309497 | 0,4339313 | 1,01 | 9,82E+07 | 9,98E+07 |
| 15,4 | | 0,4317691 | 0,4351472 | 1,02 | 2,71E+07 | 2,57E+07 |
| 0,8 | | 0,4319265 | 0,4353133 | 1,33 | 4,05E+06 | 3,41E+06 |
| 15 | | 0,4326133 | 0,4366092 | 1,23 | 1,76E+07 | 3,40E+07 |
| 6,5 | | 0,4326786 | 0,4365571 | 1,18 | 3,59E+06 | 4,08E+06 |
| 14,3 | | 0,4327645 | 0,4366990 | 0,70 | 5,58E+06 | 7,26E+06 |
| 7,5 | | 0,4327710 | 0,4366622 | 0,90 | 2,89E+07 | 2,18E+07 |
| 26,6 | | 0,4329943 | 0,4369332 | 0,95 | 8,41E+08 | 9,03E+08 |
| 32,9 | | 0,4337431 | 0,4381092 | 0,96 | 5,35E+07 | 5,65E+07 |
| 3,7 | | 0,4339147 | 0,4382554 | 0,83 | 2,67E+06 | 2,90E+06 |
| 37,3 | | 0,4341120 | 0,4384980 | 0,64 | 9,57E+06 | 8,05E+06 |
| 42,4 | | 0,4341250 | 0,4384643 | 1,11 | 9,81E+07 | 8,12E+07 |
| 13,2 | | 0,4341875 | 0,4384819 | 1,05 | 1,73E+07 | 1,86E+07 |
| 31,3 | | 0,4347668 | 0,4393938 | 1,10 | 3,12E+07 | 3,07E+07 |
| 19,3 | | 0,4349998 | 0,4398303 | 0,82 | 2,91E+07 | 1,59E+07 |
| 28,8 | | 0,4352562 | 0,4402102 | 1,08 | 2,67E+08 | 2,12E+08 |
| 9,3 | | 0,4354472 | 0,4404556 | 0,89 | 1,50E+08 | 1,08E+08 |
| 28,1 | | 0,4354570 | 0,4404030 | 0,83 | 2,67E+07 | 3,46E+07 |
| 12,8 | | 0,4359216 | 0,4412088 | 1,13 | 7,59E+06 | 9,21E+06 |
| 11,3 | | 0,4360465 | 0,4413748 | 1,16 | 1,94E+07 | 1,81E+07 |
| 58,5 | | 0,4360629 | 0,4413433 | 0,82 | 3,33E+08 | 2,22E+08 |
| 20,9 | | 0,4363332 | 0,4416322 | 0,97 | 3,20E+08 | 3,12E+08 |
| 27,4 | | 0,4369433 | 0,4425369 | 0,93 | 1,57E+08 | 1,70E+08 |
| 21,4 | | 0,4373332 | 0,4429709 | 0,82 | 1,71E+07 | 1,71E+07 |
| 54,2 | | 0,4373338 | 0,4428974 | 1,04 | 1,97E+09 | 2,27E+09 |
| 14,3 | | 0,4381879 | 0,4441295 | 0,63 | 1,53E+07 | 1,34E+07 |
| 2,2 | | 0,4388865 | 0,4451222 | 1,33 | 1,86E+06 | 3,66E+06 |
| 6,8 | | 0,4390738 | 0,4454858 | 0,66 | 1,22E+07 | 5,47E+06 |
| 18,5 | | 0,4394899 | 0,4462337 | 1,07 | 3,97E+07 | 4,88E+07 |
| 9 | | 0,4400135 | 0,4471379 | 1,38 | 4,98E+06 | 5,39E+06 |
| 26,1 | | 0,4403254 | 0,4477087 | 1,25 | 2,17E+06 | 4,50E+06 |
| 46,3 | | 0,4413298 | 0,4493236 | 0,96 | 3,66E+08 | 3,57E+08 |
| 8,9 | | 0,4415825 | 0,4497519 | 1,44 | 3,78E+06 | 3,01E+06 |
| 9,7 | | 0,4416086 | 0,4497543 | 0,78 | 1,31E+07 | 5,63E+06 |
| 10,3 | | 0,4416835 | 0,4497924 | 1,49 | 3,51E+07 | 1,96E+07 |
| 4,2 | | 0,4418240 | 0,4498982 | 0,66 | 3,88E+06 | 5,51E+06 |
| 48,7 | | 0,4419386 | 0,4500457 | 1,03 | 4,23E+09 | 4,32E+09 |
| 57,8 | | 0,4426299 | 0,4510091 | 1,10 | 7,89E+07 | 6,12E+07 |
| 21,7 | | 0,4426778 | 0,4510921 | 1,12 | 3,79E+07 | 4,84E+07 |
| 15,3 | | 0,4438205 | 0,4529892 | 1,77 | 2,13E+07 | 2,47E+07 |
| 57,3 | | 0,4442585 | 0,4535595 | 0,96 | 2,06E+08 | 2,15E+08 |
| 13 | | 0,4446605 | 0,4541600 | 1,19 | 1,22E+07 | 1,43E+07 |
| 39,7 | | 0,4448800 | 0,4543662 | 0,93 | 4,93E+08 | 5,83E+08 |
| 29,1 | | 0,4449611 | 0,4544228 | 1,03 | 9,12E+07 | 9,23E+07 |
| 25,9 | | 0,4452158 | 0,4547766 | 0,91 | 8,48E+07 | 8,75E+07 |
| 63,8 | | 0,4453077 | 0,4549436 | 0,92 | 1,39E+08 | 1,62E+08 |
| 8,3 | | 0,4459822 | 0,4561747 | 0,71 | 3,09E+07 | 8,33E+06 |
| 13,3 | + | 0,4463448 | 0,4566551 | 1,38 | 2,14E+06 | 1,46E+06 |
| 30,9 | | 0,4471012 | 0,4579140 | 1,12 | 1,33E+08 | 9,13E+07 |

| | | | | | | |
|---|---|---|---|---|---|---|
| 51,1 | | 0,4474282 | 0,4582451 | 0,94 | 9,60E+08 | 8,07E+08 |
| 22,1 | | 0,4476026 | 0,4585774 | 1,18 | 2,64E+07 | 1,93E+07 |
| 6,1 | | 0,4480365 | 0,4592931 | 2,48 | 9,75E+07 | 3,38E+06 |
| 35,8 | | 0,4481975 | 0,4594618 | 1,04 | 1,23E+08 | 1,11E+08 |
| 52,1 | | 0,4486075 | 0,4602367 | 1,03 | 5,61E+08 | 5,86E+08 |
| 7,8 | | 0,4498593 | 0,4621976 | 1,31 | 1,63E+06 | 2,75E+06 |
| 12,9 | | 0,4500708 | 0,4624821 | 0,90 | 1,94E+07 | 2,17E+07 |
| 21,3 | | 0,4501538 | 0,4625378 | 0,93 | 2,82E+07 | 2,37E+07 |
| 20,1 | | 0,4502640 | 0,4626299 | 0,65 | 6,85E+06 | 3,17E+06 |
| 16,6 | | 0,4502643 | 0,4625535 | 1,08 | 5,11E+07 | 4,08E+07 |
| 5,4 | | 0,4505084 | 0,4629612 | 0,57 | 1,94E+06 | 7,79E+06 |
| 56,2 | | 0,4509865 | 0,4635543 | 1,04 | 9,94E+08 | 1,15E+09 |
| 23,6 | | 0,4510765 | 0,4636976 | 1,06 | 4,26E+07 | 3,61E+07 |
| 30,6 | | 0,4511410 | 0,4637512 | 1,06 | 2,41E+08 | 2,97E+08 |
| 40,4 | | 0,4512831 | 0,4640211 | 0,95 | 8,30E+07 | 8,41E+07 |
| 51,8 | | 0,4518549 | 0,4651448 | 0,88 | 1,37E+09 | 1,24E+09 |
| 24,4 | | 0,4519275 | 0,4650681 | 1,15 | 1,06E+07 | 6,27E+06 |
| 20,5 | | 0,4521465 | 0,4653311 | 1,07 | 6,17E+07 | 6,10E+07 |
| 10,8 | | 0,4548521 | 0,4706988 | 0,71 | 6,32E+06 | 8,72E+06 |
| 13,7 | | 0,4553636 | 0,4714210 | 1,06 | 8,31E+07 | 9,42E+07 |
| 12,6 | | 0,4566224 | 0,4737973 | 0,69 | 3,53E+06 | 6,40E+06 |
| 44,6 | | 0,4571350 | 0,4745926 | 0,90 | 4,64E+09 | 3,02E+09 |
| 35,2 | | 0,4575772 | 0,4754859 | 0,78 | 9,54E+07 | 8,04E+07 |
| 19,8 | | 0,4578623 | 0,4759446 | 1,11 | 8,40E+06 | 8,86E+06 |
| 19,5 | | 0,4583418 | 0,4768427 | 0,43 | 1,47E+07 | 1,49E+07 |
| 17,9 | | 0,4583632 | 0,4768155 | 0,92 | 5,58E+07 | 4,28E+07 |
| 48,1 | | 0,4585169 | 0,4771283 | 1,06 | 6,45E+09 | 8,26E+09 |
| 26,6 | | 0,4587016 | 0,4774007 | 0,92 | 3,36E+07 | 2,74E+07 |
| 3,1 | | 0,4591588 | 0,4779845 | 0,55 | 1,54E+08 | 2,18E+07 |
| 14,8 | | 0,4592892 | 0,4781942 | 1,19 | 7,46E+06 | 1,26E+07 |
| 35,2 | | 0,4601227 | 0,4797927 | 0,94 | 5,62E+07 | 5,04E+07 |
| 11 | | 0,4601413 | 0,4797716 | 0,81 | 2,32E+06 | 3,05E+06 |
| 16,7 | | 0,4602904 | 0,4799283 | 1,42 | 7,39E+06 | 7,70E+06 |
| 24,7 | | 0,4603065 | 0,4798855 | 0,93 | 1,81E+08 | 1,62E+08 |
| 5 | | 0,4605530 | 0,4801658 | 0,59 | 6,16E+06 | 5,67E+06 |
| 19,6 | | 0,4606312 | 0,4802479 | 0,91 | 8,79E+08 | 9,22E+08 |
| 7,5 | | 0,4608169 | 0,4803689 | 1,36 | 2,45E+06 | 3,43E+06 |
| 50,8 | | 0,4612231 | 0,4810421 | 0,89 | 1,82E+09 | 1,76E+09 |
| 12,6 | | 0,4613340 | 0,4812904 | 1,14 | 2,49E+07 | 2,83E+07 |
| 16,2 | | 0,4615141 | 0,4814190 | 1,26 | 1,26E+08 | 2,28E+08 |
| 24,6 | | 0,4617988 | 0,4819609 | 0,95 | 8,55E+07 | 9,88E+07 |
| 8,7 | | 0,4622239 | 0,4826419 | 0,68 | 5,72E+06 | 4,12E+06 |
| 28,4 | | 0,4624569 | 0,4829325 | 0,79 | 1,25E+08 | 1,57E+08 |
| 2,2 | | 0,4640809 | 0,4861504 | 1,55 | 9,60E+06 | 4,79E+07 |
| 47,8 | | 0,4641749 | 0,4862407 | 0,93 | 8,97E+07 | 8,09E+07 |
| 6,7 | | 0,4644536 | 0,4865857 | 1,06 | 1,45E+07 | 1,35E+07 |
| 29 | | 0,4645670 | 0,4866299 | 0,86 | 9,16E+07 | 6,79E+07 |
| 33 | | 0,4652133 | 0,4876639 | 0,98 | 1,75E+08 | 1,81E+08 |
| 37,3 | | 0,4660316 | 0,4889575 | 0,92 | 2,34E+08 | 2,68E+08 |
| 13,5 | | 0,4665141 | 0,4897959 | 1,13 | 3,20E+07 | 2,35E+07 |

| | | | | | | |
|---|---|---|---|---|---|---|
| 53,3 | | 0,4667028 | 0,4901604 | 0,94 | 6,82E+07 | 6,69E+07 |
| 13,8 | | 0,4668903 | 0,4903811 | 1,71 | 2,07E+07 | 2,27E+07 |
| 62,8 | | 0,4670587 | 0,4906857 | 1,03 | 1,30E+09 | 1,35E+09 |
| 40,9 | | 0,4673204 | 0,4911148 | 0,84 | 9,04E+06 | 7,01E+06 |
| 15,7 | | 0,4675459 | 0,4915096 | 1,12 | 1,79E+07 | 1,44E+07 |
| 59,8 | | 0,4676103 | 0,4914651 | 1,08 | 1,95E+09 | 1,57E+09 |
| 3,3 | | 0,4681869 | 0,4923434 | 1,30 | 2,43E+06 | 4,91E+06 |
| 46,3 | | 0,4688371 | 0,4935590 | 0,92 | 6,38E+08 | 6,17E+08 |
| 21,9 | | 0,4693868 | 0,4944387 | 1,15 | 5,27E+07 | 3,18E+07 |
| 13,4 | | 0,4695157 | 0,4945837 | 0,49 | 2,26E+07 | 9,02E+06 |
| 53,4 | | 0,4695587 | 0,4946127 | 1,25 | 1,04E+07 | 2,17E+07 |
| 21,6 | | 0,4696778 | 0,4948435 | 0,97 | 1,55E+08 | 1,50E+08 |
| 29,6 | | 0,4702555 | 0,4956359 | 0,92 | 4,44E+08 | 4,25E+08 |
| 19,7 | | 0,4703046 | 0,4956583 | 1,85 | 1,01E+06 | 8,03E+06 |
| 47,8 | | 0,4703574 | 0,4957198 | 1,13 | 2,03E+08 | 1,83E+08 |
| 13,2 | | 0,4704695 | 0,4959012 | 1,30 | 5,42E+06 | 1,45E+07 |
| 4,4 | | 0,4705411 | 0,4958920 | 0,86 | 1,65E+07 | 1,79E+07 |
| 46,3 | | 0,4705505 | 0,4958109 | 1,03 | 8,36E+08 | 7,77E+08 |
| 55,3 | | 0,4714151 | 0,4973050 | 0,98 | 2,41E+09 | 2,52E+09 |
| 23,1 | | 0,4716011 | 0,4975592 | 1,06 | 2,84E+08 | 3,28E+08 |
| 27,9 | | 0,4716869 | 0,4977139 | 0,95 | 5,92E+08 | 5,34E+08 |
| 12,6 | | 0,4720167 | 0,4982452 | 0,89 | 1,30E+07 | 1,27E+07 |
| 69,2 | | 0,4720685 | 0,4981997 | 1,11 | 6,14E+07 | 6,37E+07 |
| 7,5 | | 0,4726150 | 0,4992405 | 1,23 | 3,47E+06 | 4,65E+06 |
| 35,7 | | 0,4727005 | 0,4992681 | 0,88 | 2,50E+07 | 2,76E+07 |
| 8,6 | | 0,4727432 | 0,4992937 | 0,65 | 5,06E+06 | 1,05E+07 |
| 15,3 | | 0,4728041 | 0,4992990 | 0,91 | 3,04E+07 | 2,05E+07 |
| 5,3 | | 0,4730050 | 0,4995722 | 0,71 | 2,69E+06 | 3,55E+06 |
| 7,8 | | 0,4730617 | 0,4996624 | 1,67 | 2,27E+07 | 2,41E+07 |
| 11,9 | | 0,4732030 | 0,4996774 | 1,73 | 9,29E+06 | 6,96E+06 |
| 10,3 | | 0,4740426 | 0,5012208 | 0,58 | 7,63E+07 | 6,29E+07 |
| 27,8 | | 0,4742100 | 0,5013179 | 0,95 | 1,21E+08 | 1,23E+08 |
| 12,1 | | 0,4743563 | 0,5014945 | 1,19 | 3,60E+07 | 2,55E+07 |
| 25,8 | | 0,4743671 | 0,5014532 | 0,96 | 7,50E+07 | 7,36E+07 |
| 40,2 | | 0,4745139 | 0,5015547 | 0,86 | 4,68E+08 | 4,87E+08 |
| 2,3 | | 0,4745157 | 0,5014729 | 0,65 | 2,22E+06 | 2,91E+06 |
| 25,4 | | 0,4748368 | 0,5018738 | 1,07 | 1,42E+07 | 1,46E+07 |
| 35,9 | | 0,4759051 | 0,5037079 | 1,03 | 4,15E+07 | 4,07E+07 |
| 25 | | 0,4765359 | 0,5044616 | 0,96 | 1,21E+08 | 1,42E+08 |
| 14,5 | | 0,4768895 | 0,5050013 | 0,91 | 8,45E+06 | 1,23E+07 |
| 43,4 | | 0,4770790 | 0,5051426 | 1,05 | 6,89E+07 | 7,56E+07 |
| 1,4 | | 0,4772797 | 0,5053920 | 0,31 | 5,80E+07 | 8,74E+07 |
| 12,7 | | 0,4776764 | 0,5059651 | 1,08 | 2,23E+07 | 2,83E+07 |
| 10,3 | | 0,4779701 | 0,5063231 | 1,11 | 1,32E+07 | 1,19E+07 |
| 50 | | 0,4780512 | 0,5063950 | 0,90 | 1,11E+08 | 1,50E+08 |
| 20,8 | | 0,4781641 | 0,5064201 | 1,05 | 5,87E+07 | 6,94E+07 |
| 3,2 | | 0,4791453 | 0,5079147 | 0,88 | 4,13E+06 | 4,11E+06 |
| 20,6 | | 0,4791821 | 0,5079076 | 1,12 | 1,45E+08 | 1,26E+08 |
| 30,6 | | 0,4793382 | 0,5081692 | 0,94 | 9,74E+07 | 1,25E+08 |
| 7 | | 0,4793418 | 0,5081067 | 1,24 | 7,31E+06 | 1,10E+07 |

| | | | | | | |
|---|---|---|---|---|---|---|
| 4,1 | | 0,4798072 | 0,5087496 | 1,12 | 3,08E+07 | 2,78E+07 |
| 14 | | 0,4798973 | 0,5088152 | 0,90 | 3,03E+07 | 2,46E+07 |
| 45,8 | | 0,4805729 | 0,5098995 | 0,80 | 2,42E+08 | 5,51E+08 |
| 35,6 | | 0,4809262 | 0,5107798 | 1,14 | 3,70E+07 | 5,81E+07 |
| 9,8 | | 0,4811642 | 0,5112047 | 1,30 | 1,49E+07 | 1,28E+07 |
| 48,4 | | 0,4823417 | 0,5134779 | 0,82 | 3,64E+08 | 6,67E+08 |
| 54,3 | | 0,4824675 | 0,5135609 | 0,94 | 7,87E+08 | 9,37E+08 |
| 52,5 | | 0,4830161 | 0,5147228 | 0,97 | 9,64E+08 | 1,06E+09 |
| 8,3 | | 0,4831013 | 0,5149407 | 1,13 | 2,54E+07 | 1,59E+07 |
| 30,8 | | 0,4841177 | 0,5168142 | 0,91 | 3,13E+08 | 3,17E+08 |
| 4,2 | | 0,4841606 | 0,5168192 | 0,88 | 2,65E+07 | 2,25E+07 |
| 17,3 | | 0,4842645 | 0,5170692 | 1,18 | 5,76E+06 | 6,87E+06 |
| 6,5 | | 0,4843421 | 0,5170606 | 1,12 | 6,44E+06 | 9,72E+06 |
| 6,8 | | 0,4846406 | 0,5174792 | 1,11 | 7,16E+06 | 5,90E+06 |
| 37,1 | | 0,4846628 | 0,5174303 | 1,17 | 4,30E+06 | 2,45E+06 |
| 44,2 | | 0,4850800 | 0,5179344 | 1,07 | 9,67E+07 | 7,72E+07 |
| 5,5 | | 0,4855737 | 0,5187636 | 0,91 | 7,91E+08 | 5,12E+08 |
| 6,8 | | 0,4876650 | 0,5223108 | 1,88 | 2,05E+07 | 1,74E+07 |
| 13,6 | | 0,4876719 | 0,5222261 | 0,84 | 6,08E+06 | 4,61E+06 |
| 46,6 | | 0,4879126 | 0,5224878 | 0,85 | 1,34E+08 | 1,28E+08 |
| 26,2 | | 0,4879183 | 0,5224031 | 1,17 | 1,60E+08 | 3,11E+08 |
| 41,2 | | 0,4885867 | 0,5237555 | 0,94 | 5,03E+09 | 6,26E+09 |
| 11,6 | | 0,4890036 | 0,5245174 | 0,90 | 3,85E+07 | 4,61E+07 |
| 1,5 | | 0,4892697 | 0,5248966 | 0,55 | 3,06E+06 | 4,95E+06 |
| 7,4 | | 0,4900096 | 0,5261857 | 0,91 | 2,84E+08 | 1,86E+08 |
| 8,2 | | 0,4900862 | 0,5262093 | 1,09 | 1,85E+07 | 2,32E+07 |
| 24,5 | | 0,4905240 | 0,5268207 | 0,94 | 3,37E+07 | 3,02E+07 |
| 9,3 | | 0,4910175 | 0,5273994 | 1,69 | 2,19E+07 | 2,91E+06 |
| 41,7 | | 0,4914930 | 0,5282423 | 1,03 | 1,49E+09 | 1,54E+09 |
| 25,1 | | 0,4915508 | 0,5282299 | 0,95 | 1,02E+08 | 8,51E+07 |
| 0,4 | | 0,4917199 | 0,5284973 | 0,56 | 3,28E+07 | 6,11E+06 |
| 67,1 | | 0,4917238 | 0,5284118 | 1,11 | 3,96E+07 | 5,76E+07 |
| 10,1 | | 0,4922966 | 0,5295187 | 0,95 | 3,57E+07 | 3,96E+07 |
| 36,6 | | 0,4930622 | 0,5307890 | 1,06 | 3,13E+08 | 2,77E+08 |
| 45,6 | | 0,4931007 | 0,5308274 | 1,56 | 1,04E+07 | 5,13E+07 |
| 42,4 | | 0,4936562 | 0,5318421 | 0,86 | 3,08E+07 | 4,46E+07 |
| 56 | | 0,4936943 | 0,5318519 | 1,04 | 4,59E+08 | 5,31E+08 |
| 11,7 | + | 0,4938582 | 0,5320731 | 1,21 | 5,75E+06 | 7,14E+06 |
| 13,7 | | 0,4941888 | 0,5326519 | 1,55 | 1,85E+06 | 3,56E+06 |
| 66,1 | | 0,4947566 | 0,5334284 | 0,92 | 2,03E+09 | 2,67E+09 |
| 1,8 | | 0,4948799 | 0,5335303 | 1,32 | 2,55E+06 | 2,59E+06 |
| 5,2 | | 0,4950411 | 0,5337304 | 1,32 | 2,50E+06 | 7,50E+06 |
| 36,8 | | 0,4952524 | 0,5340307 | 1,06 | 2,17E+08 | 2,62E+08 |
| 31,3 | | 0,4959232 | 0,5352174 | 1,04 | 7,44E+07 | 8,62E+07 |
| 28,7 | | 0,4960005 | 0,5353184 | 0,79 | 5,10E+06 | 1,12E+07 |
| 6,5 | | 0,4960143 | 0,5352319 | 0,67 | 4,21E+06 | 8,50E+06 |
| 11,9 | | 0,4960968 | 0,5352475 | 0,75 | 2,79E+07 | 2,23E+07 |
| 3 | | 0,4964820 | 0,5356564 | 0,84 | 4,77E+07 | 3,01E+07 |
| 8 | | 0,4965283 | 0,5356100 | 0,68 | 1,32E+07 | 2,84E+06 |
| 6,4 | | 0,4967217 | 0,5358476 | 0,73 | 9,64E+06 | 2,38E+06 |

| | | | | | | |
|---|---|---|---|---|---|---|
| 18,8 | | 0,4969651 | 0,5361459 | 0,87 | 8,94E+06 | 7,57E+06 |
| 30,9 | | 0,4987339 | 0,5392301 | 0,92 | 1,88E+07 | 2,03E+07 |
| 11 | | 0,4988214 | 0,5393044 | 1,53 | 9,59E+06 | 2,09E+06 |
| 9,5 | | 0,4992683 | 0,5399677 | 0,91 | 5,42E+06 | 5,22E+06 |
| 28,9 | | 0,4993723 | 0,5400826 | 0,86 | 8,89E+07 | 7,60E+07 |
| 36 | | 0,4995406 | 0,5402702 | 1,08 | 1,81E+07 | 1,79E+07 |
| 52,9 | | 0,4997037 | 0,5404275 | 0,87 | 4,82E+07 | 7,39E+07 |
| 3,4 | | 0,5001892 | 0,5412166 | 1,11 | 6,95E+07 | 4,37E+07 |
| 62,1 | | 0,5004790 | 0,5415407 | 0,90 | 9,09E+07 | 9,69E+07 |
| 50 | | 0,5005931 | 0,5417042 | 2,25 | 1,35E+07 | 1,73E+06 |
| 10,5 | | 0,5021099 | 0,5444150 | 0,66 | 8,94E+06 | 1,18E+07 |
| 27 | | 0,5021277 | 0,5444188 | 0,95 | 7,99E+07 | 8,17E+07 |
| 12 | | 0,5029474 | 0,5458579 | 1,12 | 1,91E+07 | 2,16E+07 |
| 13,7 | | 0,5031577 | 0,5462164 | 1,08 | 3,94E+07 | 3,90E+07 |
| 21,4 | | 0,5032551 | 0,5463648 | 0,89 | 5,32E+07 | 3,26E+07 |
| 22,9 | | 0,5035210 | 0,5468268 | 0,93 | 1,09E+07 | 1,49E+07 |
| 9,5 | | 0,5036933 | 0,5469004 | 1,39 | 4,77E+06 | 3,41E+06 |
| 25,2 | | 0,5038397 | 0,5470647 | 0,89 | 1,61E+07 | 1,55E+07 |
| 11,6 | | 0,5042586 | 0,5478352 | 0,91 | 5,67E+07 | 4,66E+07 |
| 51,9 | | 0,5050734 | 0,5492992 | 0,96 | 2,04E+08 | 2,44E+08 |
| 19,4 | | 0,5063055 | 0,5514242 | 0,92 | 1,88E+07 | 1,71E+07 |
| 22,1 | | 0,5069197 | 0,5523390 | 0,90 | 4,96E+07 | 3,44E+07 |
| 53,3 | | 0,5071506 | 0,5527249 | 1,06 | 2,14E+08 | 2,73E+08 |
| 50,3 | | 0,5073843 | 0,5530328 | 0,97 | 8,03E+07 | 8,90E+07 |
| 21,2 | | 0,5077340 | 0,5536113 | 0,94 | 5,14E+07 | 6,55E+07 |
| 31,3 | | 0,5082500 | 0,5544950 | 0,97 | 2,64E+07 | 2,71E+07 |
| 37,3 | | 0,5086338 | 0,5552974 | 0,98 | 1,62E+08 | 1,76E+08 |
| 9 | | 0,5087782 | 0,5555206 | 0,78 | 6,09E+06 | 1,38E+07 |
| 12,2 | | 0,5093189 | 0,5562814 | 0,88 | 6,64E+06 | 1,09E+07 |
| 7,1 | | 0,5094546 | 0,5563945 | 1,37 | 2,99E+06 | 6,78E+06 |
| 3,3 | | 0,5101587 | 0,5577986 | 1,11 | 1,02E+07 | 1,33E+07 |
| 36,6 | | 0,5105982 | 0,5587405 | 1,02 | 2,08E+08 | 2,17E+08 |
| 35,3 | | 0,5117514 | 0,5609119 | 0,85 | 7,03E+06 | 5,01E+06 |
| 50,9 | | 0,5118104 | 0,5609291 | 1,04 | 3,41E+08 | 3,54E+08 |
| 13,9 | | 0,5120339 | 0,5611838 | 1,07 | 4,68E+07 | 4,77E+07 |
| 19,3 | | 0,5124604 | 0,5618177 | 1,06 | 1,92E+07 | 1,57E+07 |
| 55,4 | | 0,5129088 | 0,5625843 | 0,95 | 9,81E+07 | 9,54E+07 |
| 22,2 | | 0,5129772 | 0,5625768 | 1,06 | 1,91E+07 | 1,80E+07 |
| 12,9 | | 0,5133023 | 0,5630897 | 0,93 | 2,93E+07 | 2,21E+07 |
| 20,1 | | 0,5135911 | 0,5636684 | 1,09 | 3,60E+08 | 2,53E+08 |
| 32,3 | | 0,5136258 | 0,5636562 | 0,76 | 1,14E+08 | 1,30E+08 |
| 4,9 | | 0,5139741 | 0,5642956 | 0,82 | 4,01E+06 | 4,24E+06 |
| 8,9 | | 0,5139850 | 0,5642052 | 1,58 | 3,05E+06 | 1,57E+07 |
| 12,4 | | 0,5141491 | 0,5643237 | 0,95 | 3,07E+07 | 3,24E+07 |
| 27,5 | | 0,5141701 | 0,5642333 | 0,78 | 2,89E+07 | 3,18E+07 |
| 7,4 | | 0,5151359 | 0,5660423 | 1,11 | 6,66E+06 | 6,98E+06 |
| 14,2 | | 0,5155933 | 0,5666167 | 1,75 | 1,31E+07 | 1,34E+06 |
| 41,9 | | 0,5156381 | 0,5666470 | 1,07 | 2,04E+08 | 2,34E+08 |
| 9,7 | | 0,5162650 | 0,5675837 | 1,08 | 2,10E+07 | 1,64E+07 |
| 56,2 | | 0,5163373 | 0,5676375 | 1,12 | 3,23E+07 | 4,69E+07 |

| | | | | | | |
|---|---|---|---|---|---|---|
| 6,3 | | 0,5164500 | 0,5677490 | 0,77 | 7,85E+06 | 3,06E+06 |
| 5,9 | | 0,5172182 | 0,5691383 | 0,77 | 9,50E+06 | 6,38E+06 |
| 31,8 | | 0,5175648 | 0,5695913 | 0,95 | 6,78E+07 | 5,62E+07 |
| 38,9 | | 0,5177722 | 0,5699347 | 0,97 | 1,61E+08 | 1,45E+08 |
| 22,6 | | 0,5180417 | 0,5703817 | 0,94 | 3,33E+07 | 3,12E+07 |
| 35,4 | | 0,5195779 | 0,5727217 | 0,94 | 1,22E+08 | 1,21E+08 |
| 49,1 | | 0,5196084 | 0,5727702 | 0,92 | 4,50E+07 | 4,98E+07 |
| 37,2 | | 0,5200646 | 0,5735248 | 0,97 | 2,93E+08 | 2,97E+08 |
| 10,9 | | 0,5203250 | 0,5739946 | 1,23 | 5,91E+06 | 1,40E+07 |
| 15,4 | | 0,5206537 | 0,5745512 | 0,57 | 4,49E+07 | 2,92E+07 |
| 4,9 | | 0,5207469 | 0,5746019 | 1,20 | 4,52E+06 | 2,18E+06 |
| 51,2 | | 0,5208284 | 0,5745996 | 0,96 | 4,78E+08 | 5,21E+08 |
| 21,8 | + | 0,5213419 | 0,5753711 | 0,39 | 2,08E+06 | 3,38E+06 |
| 15,3 | | 0,5215592 | 0,5756741 | 0,96 | 9,51E+07 | 9,62E+07 |
| 41,7 | | 0,5218141 | 0,5760511 | 1,11 | 6,69E+07 | 8,87E+07 |
| 18,8 | | 0,5219397 | 0,5760696 | 1,29 | 2,09E+08 | 7,92E+07 |
| 3,7 | | 0,5221586 | 0,5764126 | 1,17 | 2,01E+06 | 2,39E+06 |
| 17 | | 0,5225779 | 0,5771897 | 0,96 | 4,75E+07 | 4,26E+07 |
| 76,3 | | 0,5226339 | 0,5771927 | 1,02 | 1,36E+09 | 1,32E+09 |
| 21,7 | | 0,5237578 | 0,5789486 | 0,86 | 3,55E+07 | 2,11E+07 |
| 55,7 | | 0,5239421 | 0,5791932 | 0,93 | 4,92E+08 | 3,71E+08 |
| 42,1 | | 0,5245208 | 0,5801678 | 0,97 | 7,12E+08 | 7,39E+08 |
| 7,7 | | 0,5248340 | 0,5807778 | 0,68 | 3,49E+06 | 3,24E+06 |
| 19,5 | | 0,5249283 | 0,5808338 | 1,12 | 2,49E+07 | 2,59E+07 |
| 22,2 | | 0,5250600 | 0,5808866 | 1,07 | 7,47E+07 | 8,89E+07 |
| 20,2 | | 0,5251097 | 0,5809407 | 1,22 | 5,61E+06 | 1,05E+07 |
| 15,3 | | 0,5255703 | 0,5816273 | 0,90 | 2,88E+07 | 2,64E+07 |
| 54,1 | | 0,5257925 | 0,5820013 | 1,10 | 7,57E+08 | 8,63E+08 |
| 37,5 | | 0,5263416 | 0,5830986 | 1,03 | 1,79E+09 | 1,54E+09 |
| 49,1 | | 0,5264722 | 0,5832294 | 0,92 | 3,51E+08 | 4,45E+08 |
| 1,9 | | 0,5276785 | 0,5851693 | 1,16 | 7,56E+06 | 1,36E+07 |
| 27 | | 0,5277904 | 0,5852889 | 1,07 | 4,01E+07 | 3,93E+07 |
| 33,9 | | 0,5278361 | 0,5853601 | 1,07 | 6,74E+07 | 7,13E+07 |
| 28 | | 0,5282453 | 0,5858650 | 0,88 | 5,15E+07 | 6,99E+07 |
| 15,3 | | 0,5285006 | 0,5860659 | 1,45 | 2,10E+06 | 1,15E+07 |
| 30,9 | | 0,5285889 | 0,5861624 | 1,06 | 1,48E+08 | 1,47E+08 |
| 17,3 | | 0,5286379 | 0,5861716 | 1,07 | 7,21E+07 | 9,17E+07 |
| 17,9 | | 0,5286638 | 0,5861509 | 0,90 | 3,78E+07 | 5,61E+07 |
| 4,3 | | 0,5286891 | 0,5861302 | 1,15 | 3,12E+06 | 3,99E+06 |
| 31,7 | | 0,5296239 | 0,5878715 | 1,07 | 2,46E+07 | 2,78E+07 |
| 33,9 | | 0,5306022 | 0,5894856 | 0,94 | 6,12E+07 | 6,39E+07 |
| 50,6 | | 0,5318164 | 0,5913111 | 0,94 | 5,31E+08 | 6,73E+08 |
| 70,5 | | 0,5319032 | 0,5914034 | 1,03 | 1,09E+09 | 1,23E+09 |
| 21,4 | | 0,5321110 | 0,5916623 | 1,08 | 3,00E+07 | 3,30E+07 |
| 44,1 | | 0,5326814 | 0,5923974 | 1,03 | 2,32E+08 | 2,46E+08 |
| 26,6 | | 0,5336500 | 0,5939517 | 1,03 | 2,64E+08 | 2,37E+08 |
| 44,4 | | 0,5338380 | 0,5942800 | 1,04 | 6,20E+08 | 5,93E+08 |
| 56,3 | | 0,5341519 | 0,5947645 | 0,86 | 7,56E+08 | 6,00E+08 |
| 30,8 | | 0,5342922 | 0,5948359 | 1,11 | 5,21E+07 | 3,83E+07 |
| 36,3 | | 0,5345768 | 0,5954612 | 0,86 | 4,44E+07 | 5,94E+07 |

| | | | | | | |
|---|---|---|---|---|---|---|
| 8,7 | | 0,5349138 | 0,5960680 | 1,07 | 3,52E+07 | 2,67E+07 |
| 10 | | 0,5352457 | 0,5965151 | 0,93 | 1,81E+07 | 2,51E+07 |
| 38,4 | | 0,5354444 | 0,5967658 | 1,04 | 7,79E+07 | 7,78E+07 |
| 17,7 | | 0,5360299 | 0,5977765 | 1,04 | 3,39E+08 | 3,71E+08 |
| 26,9 | | 0,5361533 | 0,5978353 | 1,02 | 5,43E+07 | 5,87E+07 |
| 13 | | 0,5369141 | 0,5992072 | 1,40 | 6,69E+06 | 1,57E+06 |
| 39,6 | | 0,5370948 | 0,5995526 | 0,89 | 1,65E+07 | 1,48E+07 |
| 6,6 | | 0,5383109 | 0,6018623 | 0,92 | 9,26E+06 | 1,18E+07 |
| 11,7 | | 0,5383906 | 0,6018544 | 1,18 | 5,90E+06 | 7,55E+06 |
| 9,8 | | 0,5389765 | 0,6028950 | 0,92 | 2,74E+06 | 3,47E+06 |
| 21,8 | | 0,5394648 | 0,6036886 | 1,07 | 2,38E+07 | 2,01E+07 |
| 41,7 | | 0,5395378 | 0,6036779 | 0,92 | 2,83E+08 | 2,84E+08 |
| 38,9 | | 0,5400157 | 0,6042847 | 0,95 | 4,03E+08 | 4,12E+08 |
| 25,7 | | 0,5405084 | 0,6051487 | 0,91 | 5,53E+07 | 8,60E+07 |
| 46,7 | | 0,5405918 | 0,6052645 | 1,09 | 1,75E+08 | 2,08E+08 |
| 50 | | 0,5410929 | 0,6060520 | 0,93 | 7,80E+07 | 7,99E+07 |
| 30,7 | | 0,5411870 | 0,6060985 | 0,88 | 2,61E+07 | 3,67E+07 |
| 7,9 | | 0,5418245 | 0,6070282 | 1,30 | 5,21E+06 | 5,37E+06 |
| 58,3 | | 0,5420667 | 0,6075572 | 0,97 | 1,89E+09 | 2,15E+09 |
| 20,7 | | 0,5423471 | 0,6078619 | 1,05 | 2,20E+07 | 2,72E+07 |
| 19 | | 0,5426324 | 0,6082875 | 1,31 | 3,22E+06 | 2,44E+06 |
| 35,9 | | 0,5446417 | 0,6114701 | 0,72 | 5,90E+06 | 4,01E+06 |
| 44,1 | | 0,5446842 | 0,6114721 | 0,95 | 6,90E+08 | 7,54E+08 |
| 67,8 | | 0,5451653 | 0,6122405 | 1,19 | 3,70E+09 | 4,21E+09 |
| 5 | | 0,5456448 | 0,6129075 | 0,61 | 4,98E+06 | 1,19E+06 |
| 5,8 | | 0,5458340 | 0,6133724 | 1,44 | 7,33E+06 | 7,31E+06 |
| 28,5 | | 0,5458865 | 0,6133936 | 1,12 | 3,98E+08 | 2,85E+08 |
| 17,1 | | 0,5462679 | 0,6140683 | 1,60 | 1,44E+07 | 2,52E+06 |
| 16,1 | | 0,5465549 | 0,6144108 | 0,94 | 6,44E+07 | 6,13E+07 |
| 38,1 | | 0,5467022 | 0,6145147 | 1,06 | 2,78E+08 | 2,30E+08 |
| 2,8 | | 0,5472579 | 0,6152534 | 2,09 | 1,58E+06 | 3,47E+07 |
| 29,9 | | 0,5477472 | 0,6158875 | 1,16 | 4,86E+07 | 9,77E+07 |
| 59,7 | | 0,5480275 | 0,6162553 | 0,94 | 1,21E+09 | 9,28E+08 |
| 30,2 | | 0,5481749 | 0,6164303 | 1,05 | 4,81E+09 | 5,32E+09 |
| 36,1 | | 0,5483766 | 0,6167449 | 1,07 | 3,08E+07 | 3,88E+07 |
| 6,8 | | 0,5484575 | 0,6167530 | 0,63 | 1,13E+07 | 1,63E+06 |
| 37,3 | | 0,5489566 | 0,6177603 | 1,03 | 1,22E+08 | 1,16E+08 |
| 26,7 | | 0,5493896 | 0,6185696 | 1,07 | 2,26E+07 | 2,89E+07 |
| 24,3 | | 0,5494118 | 0,6185219 | 1,05 | 1,65E+07 | 1,58E+07 |
| 7,5 | | 0,5494281 | 0,6185139 | 0,90 | 1,44E+08 | 2,10E+08 |
| 31,6 | | 0,5494468 | 0,6184794 | 1,02 | 1,06E+08 | 1,11E+08 |
| 57,7 | | 0,5506233 | 0,6203099 | 0,97 | 6,82E+08 | 6,89E+08 |
| 4,1 | | 0,5508141 | 0,6204203 | 1,17 | 6,18E+06 | 7,13E+06 |
| 38,5 | | 0,5508405 | 0,6203565 | 1,04 | 4,08E+08 | 3,32E+08 |
| 1,9 | | 0,5510822 | 0,6207456 | 0,25 | 5,25E+06 | 1,79E+08 |
| 74,7 | | 0,5510942 | 0,6206477 | 0,97 | 4,41E+09 | 4,34E+09 |
| 14,3 | | 0,5511158 | 0,6205802 | 1,18 | 4,98E+07 | 2,88E+07 |
| 59,1 | | 0,5511770 | 0,6205788 | 1,08 | 3,18E+09 | 3,03E+09 |
| 24,9 | | 0,5513153 | 0,6206859 | 1,12 | 1,61E+07 | 1,49E+07 |
| 33,9 | | 0,5517025 | 0,6212789 | 1,05 | 1,79E+08 | 1,49E+08 |

| | | | | | | |
|---|---|---|---|---|---|---|
| 32,4 | | 0,5520627 | 0,6217388 | 1,03 | 1,83E+08 | 1,68E+08 |
| 43,9 | | 0,5528115 | 0,6227940 | 1,11 | 2,06E+08 | 2,20E+08 |
| 5,2 | | 0,5538992 | 0,6246678 | 1,10 | 6,18E+07 | 4,22E+07 |
| 53,3 | | 0,5539723 | 0,6246236 | 1,09 | 2,92E+09 | 3,56E+09 |
| 28 | | 0,5543908 | 0,6249907 | 0,89 | 7,55E+06 | 4,58E+06 |
| 42,7 | | 0,5555257 | 0,6268879 | 1,06 | 1,03E+08 | 1,23E+08 |
| 18 | | 0,5561428 | 0,6276128 | 1,46 | 3,22E+06 | 4,67E+06 |
| 51,3 | | 0,5563066 | 0,6277350 | 0,93 | 5,39E+08 | 7,41E+08 |
| 45,5 | | 0,5565177 | 0,6279710 | 1,02 | 2,24E+08 | 2,28E+08 |
| 16,4 | | 0,5568516 | 0,6283870 | 0,93 | 3,26E+07 | 3,33E+07 |
| 17,6 | | 0,5576641 | 0,6296756 | 1,03 | 2,61E+07 | 2,24E+07 |
| 29,5 | | 0,5580942 | 0,6301265 | 1,06 | 1,52E+08 | 1,45E+08 |
| 67,6 | | 0,5581263 | 0,6300613 | 1,04 | 4,00E+09 | 4,29E+09 |
| 27,2 | | 0,5584538 | 0,6305145 | 0,96 | 1,04E+08 | 9,43E+07 |
| 33,8 | | 0,5586224 | 0,6307499 | 0,97 | 3,25E+08 | 3,11E+08 |
| 16,8 | | 0,5588946 | 0,6311374 | 0,77 | 3,32E+06 | 3,13E+06 |
| 16,7 | | 0,5598005 | 0,6327518 | 0,94 | 3,30E+07 | 2,51E+07 |
| 29,2 | | 0,5605855 | 0,6339786 | 1,03 | 9,35E+07 | 8,60E+07 |
| 8,1 | | 0,5613099 | 0,6351126 | 0,85 | 5,24E+06 | 6,82E+06 |
| 34,8 | | 0,5615988 | 0,6355281 | 0,88 | 3,67E+08 | 4,30E+08 |
| 23,6 | | 0,5617486 | 0,6356193 | 1,21 | 4,50E+07 | 2,50E+07 |
| 10,6 | | 0,5620834 | 0,6360628 | 0,96 | 3,10E+09 | 3,07E+09 |
| 5,1 | | 0,5621062 | 0,6359824 | 1,35 | 1,11E+07 | 1,27E+07 |
| 11,9 | | 0,5621092 | 0,6358826 | 0,84 | 4,51E+06 | 4,27E+06 |
| 5,6 | | 0,5626096 | 0,6363032 | 1,30 | 6,95E+06 | 5,21E+06 |
| 4,2 | | 0,5630297 | 0,6370609 | 1,41 | 2,99E+06 | 5,23E+06 |
| 52 | | 0,5634568 | 0,6376118 | 1,07 | 1,18E+08 | 1,13E+08 |
| 28,3 | | 0,5637921 | 0,6380709 | 1,01 | 2,29E+08 | 2,42E+08 |
| 18,2 | | 0,5642242 | 0,6387608 | 0,98 | 1,02E+08 | 9,03E+07 |
| 8,4 | | 0,5650637 | 0,6401982 | 1,33 | 7,10E+06 | 2,08E+06 |
| 45,6 | | 0,5651914 | 0,6403174 | 0,96 | 8,35E+07 | 9,83E+07 |
| 27,4 | | 0,5653942 | 0,6405695 | 1,06 | 8,82E+07 | 7,59E+07 |
| 56,5 | | 0,5655284 | 0,6406572 | 1,07 | 7,64E+08 | 9,76E+08 |
| 34,3 | | 0,5656311 | 0,6407480 | 1,06 | 1,74E+08 | 2,37E+08 |
| 59,7 | | 0,5658180 | 0,6409378 | 0,97 | 5,93E+08 | 6,79E+08 |
| 16,1 | | 0,5660484 | 0,6413106 | 1,17 | 3,42E+06 | 6,43E+06 |
| 3,8 | | 0,5664912 | 0,6418330 | 0,81 | 5,86E+06 | 2,19E+06 |
| 7,3 | | 0,5667732 | 0,6422393 | 0,78 | 1,08E+07 | 8,38E+06 |
| 4,6 | | 0,5669197 | 0,6423248 | 0,74 | 1,23E+07 | 7,50E+06 |
| 8,9 | | 0,5675060 | 0,6433298 | 1,08 | 7,37E+06 | 8,23E+06 |
| 13,2 | | 0,5675747 | 0,6433124 | 0,87 | 2,49E+07 | 2,73E+07 |
| 22,5 | | 0,5676637 | 0,6433588 | 0,88 | 2,28E+08 | 3,04E+08 |
| 30,6 | | 0,5682424 | 0,6441058 | 1,12 | 5,60E+07 | 7,70E+07 |
| 8,3 | | 0,5684422 | 0,6443055 | 1,45 | 5,78E+06 | 1,46E+06 |
| 4 | | 0,5695724 | 0,6463996 | 0,89 | 3,22E+06 | 3,59E+06 |
| 29 | | 0,5697377 | 0,6464581 | 0,95 | 9,27E+07 | 1,09E+08 |
| 14,7 | | 0,5699562 | 0,6468932 | 0,96 | 2,98E+07 | 2,96E+07 |
| 23,4 | | 0,5700758 | 0,6469434 | 0,96 | 3,02E+09 | 2,70E+09 |
| 6,7 | | 0,5708930 | 0,6482884 | 0,79 | 7,18E+06 | 4,40E+06 |
| 41,1 | | 0,5709177 | 0,6482208 | 0,96 | 2,72E+07 | 2,91E+07 |

| | | | | | | |
|---|---|---|---|---|---|---|
| 7,3 | | 0,5718108 | 0,6496295 | 1,03 | 4,03E+07 | 3,48E+07 |
| 6,7 | | 0,5718295 | 0,6495811 | 1,14 | 5,62E+06 | 5,16E+06 |
| 16,7 | | 0,5720781 | 0,6500547 | 0,91 | 3,10E+07 | 3,18E+07 |
| 26,3 | | 0,5723375 | 0,6503713 | 0,95 | 1,01E+08 | 9,80E+07 |
| 36,7 | | 0,5724830 | 0,6505477 | 0,97 | 1,15E+08 | 1,22E+08 |
| 38,8 | | 0,5731579 | 0,6513583 | 0,86 | 2,01E+08 | 3,86E+08 |
| 30 | | 0,5731837 | 0,6513060 | 0,93 | 6,71E+07 | 6,92E+07 |
| 51,9 | | 0,5743366 | 0,6528907 | 0,96 | 2,05E+08 | 2,60E+08 |
| 23,6 | | 0,5744516 | 0,6529636 | 1,11 | 1,55E+08 | 1,16E+08 |
| 44,4 | | 0,5756293 | 0,6546987 | 1,13 | 3,46E+07 | 5,54E+07 |
| 25,6 | | 0,5758866 | 0,6550036 | 0,97 | 2,14E+08 | 1,88E+08 |
| 22,9 | | 0,5768269 | 0,6565119 | 0,93 | 7,51E+07 | 6,94E+07 |
| 43,7 | | 0,5774901 | 0,6577850 | 0,83 | 6,48E+07 | 4,61E+07 |
| 4,1 | | 0,5780809 | 0,6586228 | 1,17 | 5,98E+06 | 4,16E+06 |
| 21,2 | | 0,5781631 | 0,6586898 | 0,89 | 3,14E+07 | 5,27E+07 |
| 35,5 | | 0,5781885 | 0,6586351 | 1,04 | 1,24E+08 | 1,02E+08 |
| 52,1 | | 0,5790050 | 0,6598179 | 0,92 | 3,25E+08 | 3,34E+08 |
| 27,7 | | 0,5804997 | 0,6623087 | 1,51 | 1,59E+06 | 1,17E+07 |
| 32,1 | | 0,5815076 | 0,6640100 | 0,88 | 7,10E+06 | 1,20E+07 |
| 43,2 | | 0,5817003 | 0,6642307 | 0,98 | 2,51E+08 | 2,48E+08 |
| 32,5 | | 0,5822969 | 0,6651120 | 0,98 | 1,82E+08 | 1,69E+08 |
| 4,7 | | 0,5823795 | 0,6650726 | 0,79 | 3,77E+06 | 1,00E+07 |
| 24,2 | | 0,5827378 | 0,6656501 | 1,02 | 7,49E+07 | 8,14E+07 |
| 5,1 | | 0,5830860 | 0,6661371 | 0,82 | 5,67E+06 | 7,92E+06 |
| 7,9 | | 0,5834225 | 0,6665242 | 0,75 | 4,55E+06 | 4,06E+06 |
| 5,5 | | 0,5835956 | 0,6666889 | 0,86 | 1,21E+07 | 1,15E+07 |
| 55,6 | | 0,5836268 | 0,6666405 | 1,09 | 1,62E+09 | 1,22E+09 |
| 3,9 | | 0,5839284 | 0,6670741 | 0,78 | 1,79E+06 | 2,90E+06 |
| 6,6 | | 0,5841543 | 0,6672928 | 1,16 | 1,33E+07 | 8,45E+06 |
| 91,5 | | 0,5844373 | 0,6676564 | 1,04 | 1,24E+09 | 1,16E+09 |
| 16,9 | | 0,5846165 | 0,6680510 | 1,05 | 2,07E+07 | 1,77E+07 |
| 6,5 | | 0,5846311 | 0,6679664 | 0,73 | 7,69E+06 | 3,03E+06 |
| 11,3 | | 0,5854654 | 0,6692761 | 1,08 | 3,93E+07 | 5,42E+07 |
| 5,1 | | 0,5855900 | 0,6693518 | 1,37 | 3,02E+06 | 6,34E+06 |
| 61,9 | | 0,5857010 | 0,6695214 | 1,06 | 1,51E+09 | 1,92E+09 |
| 31,4 | | 0,5859181 | 0,6696841 | 0,94 | 1,12E+08 | 1,45E+08 |
| 46,7 | | 0,5873602 | 0,6715019 | 0,92 | 6,97E+07 | 1,09E+08 |
| 39,7 | | 0,5878069 | 0,6721113 | 0,96 | 3,18E+08 | 2,75E+08 |
| 16,4 | | 0,5899461 | 0,6748264 | 1,08 | 3,22E+07 | 3,85E+07 |
| 39,5 | | 0,5900780 | 0,6749323 | 0,91 | 3,10E+08 | 3,29E+08 |
| 60,3 | | 0,5905724 | 0,6756632 | 0,94 | 5,28E+08 | 4,94E+08 |
| 12,4 | | 0,5916167 | 0,6770419 | 1,05 | 2,64E+07 | 3,22E+07 |
| 3,8 | | 0,5920236 | 0,6776357 | 0,89 | 3,66E+06 | 3,40E+06 |
| 21,6 | | 0,5921244 | 0,6777752 | 0,88 | 1,69E+07 | 1,53E+07 |
| 20,6 | | 0,5922362 | 0,6777426 | 1,08 | 3,30E+07 | 3,86E+07 |
| 36,5 | | 0,5923266 | 0,6777585 | 1,08 | 2,15E+07 | 2,75E+07 |
| 57,8 | | 0,5923902 | 0,6777278 | 0,98 | 4,36E+08 | 4,36E+08 |
| 11,2 | | 0,5929884 | 0,6784525 | 1,22 | 1,35E+06 | 3,47E+06 |
| 17,4 | | 0,5936227 | 0,6794501 | 0,88 | 1,10E+07 | 9,69E+06 |
| 11,8 | | 0,5936449 | 0,6794123 | 1,17 | 1,39E+07 | 1,36E+07 |

| | | | | | | |
|---|---|---|---|---|---|---|
| 17,6 | | 0,5939153 | 0,6796091 | 0,89 | 1,38E+08 | 8,56E+07 |
| 3,9 | | 0,5950164 | 0,6812128 | 0,74 | 7,09E+06 | 6,57E+06 |
| 40,3 | | 0,5955878 | 0,6822323 | 1,01 | 3,46E+08 | 3,21E+08 |
| 12,1 | + | 0,5959800 | 0,6827696 | 0,63 | 9,49E+08 | 1,85E+08 |
| 14,7 | | 0,5962791 | 0,6829868 | 0,92 | 2,53E+07 | 1,85E+07 |
| 21,7 | | 0,5964612 | 0,6830942 | 0,96 | 1,97E+07 | 2,33E+07 |
| 8 | | 0,5964937 | 0,6830727 | 1,12 | 2,37E+06 | 2,99E+06 |
| 18,8 | | 0,5966413 | 0,6831999 | 0,80 | 6,13E+06 | 6,60E+06 |
| 38,5 | | 0,5967023 | 0,6832074 | 1,04 | 3,31E+08 | 3,31E+08 |
| 7,1 | | 0,5971604 | 0,6837230 | 1,09 | 1,23E+07 | 1,44E+07 |
| 37,6 | | 0,5993980 | 0,6866223 | 0,90 | 2,45E+08 | 2,71E+08 |
| 13,8 | | 0,5995375 | 0,6866634 | 0,96 | 1,11E+08 | 1,40E+08 |
| 49,1 | | 0,6008966 | 0,6883183 | 1,10 | 5,38E+07 | 7,62E+07 |
| 16,3 | | 0,6013808 | 0,6889406 | 1,07 | 2,06E+07 | 1,89E+07 |
| 30,8 | | 0,6018610 | 0,6896982 | 0,95 | 3,73E+08 | 4,39E+08 |
| 22,4 | | 0,6024122 | 0,6903677 | 0,66 | 1,68E+06 | 1,94E+07 |
| 33,8 | | 0,6026810 | 0,6908061 | 1,07 | 6,70E+07 | 8,49E+07 |
| 12,6 | | 0,6037450 | 0,6925514 | 1,48 | 3,72E+07 | 2,02E+07 |
| 52,9 | | 0,6045768 | 0,6938419 | 0,97 | 5,50E+09 | 4,75E+09 |
| 34 | | 0,6048452 | 0,6941565 | 0,98 | 1,01E+09 | 1,03E+09 |
| 62,4 | | 0,6055494 | 0,6949918 | 1,03 | 1,88E+09 | 2,02E+09 |
| 23,7 | | 0,6060458 | 0,6957143 | 0,94 | 2,44E+07 | 2,05E+07 |
| 8,2 | | 0,6061548 | 0,6957354 | 1,18 | 5,71E+06 | 6,43E+06 |
| 34,4 | | 0,6066176 | 0,6963858 | 1,04 | 8,23E+07 | 6,54E+07 |
| 19,9 | | 0,6066262 | 0,6962782 | 0,58 | 1,95E+07 | 1,82E+06 |
| 28,9 | | 0,6066986 | 0,6962695 | 0,98 | 1,09E+08 | 1,14E+08 |
| 3,4 | | 0,6067658 | 0,6962750 | 1,25 | 8,60E+06 | 2,22E+07 |
| 52,4 | | 0,6075667 | 0,6973061 | 0,96 | 2,61E+09 | 3,28E+09 |
| 14,9 | | 0,6101064 | 0,7007358 | 1,07 | 1,33E+07 | 1,35E+07 |
| 34,4 | | 0,6104837 | 0,7011069 | 1,05 | 5,22E+07 | 5,75E+07 |
| 9,1 | | 0,6109296 | 0,7014951 | 0,88 | 1,88E+06 | 2,44E+06 |
| 21,3 | | 0,6125757 | 0,7036641 | 0,89 | 7,42E+07 | 1,41E+08 |
| 6,5 | | 0,6131243 | 0,7044840 | 1,26 | 2,11E+06 | 1,95E+06 |
| 35,9 | | 0,6142283 | 0,7061895 | 0,92 | 3,72E+08 | 2,90E+08 |
| 23,3 | | 0,6144056 | 0,7062750 | 0,30 | 8,79E+08 | 2,48E+06 |
| 24,3 | | 0,6147932 | 0,7067911 | 1,21 | 6,55E+06 | 2,03E+07 |
| 19,2 | | 0,6149315 | 0,7068524 | 1,09 | 1,19E+07 | 1,63E+07 |
| 3,3 | | 0,6150037 | 0,7069500 | 1,35 | 1,44E+07 | 2,53E+06 |
| 13,5 | | 0,6157627 | 0,7080944 | 1,06 | 2,02E+07 | 2,31E+07 |
| 22,3 | | 0,6163255 | 0,7089954 | 1,08 | 7,02E+07 | 9,68E+07 |
| 10,8 | | 0,6164527 | 0,7091333 | 1,40 | 6,20E+06 | 4,55E+06 |
| 2,9 | | 0,6164623 | 0,7090240 | 1,08 | 1,13E+07 | 7,58E+06 |
| 3,9 | | 0,6167428 | 0,7094135 | 1,05 | 1,42E+08 | 1,12E+08 |
| 17,1 | | 0,6170897 | 0,7097621 | 1,07 | 5,24E+07 | 6,08E+07 |
| 46,7 | | 0,6173829 | 0,7099029 | 0,98 | 1,76E+08 | 1,81E+08 |
| 6,7 | | 0,6201173 | 0,7139181 | 0,78 | 2,07E+06 | 2,49E+06 |
| 53,5 | | 0,6201241 | 0,7138278 | 1,02 | 1,56E+09 | 1,40E+09 |
| 3,9 | | 0,6209088 | 0,7147385 | 1,29 | 4,42E+06 | 1,37E+07 |
| 19 | | 0,6216130 | 0,7155048 | 1,26 | 6,27E+06 | 2,11E+06 |
| 7,9 | | 0,6221860 | 0,7160991 | 1,08 | 9,43E+06 | 1,01E+07 |

| | | | | | | |
|---|---|---|---|---|---|---|
| 69,3 | | 0,6222520 | 0,7161237 | 0,98 | 2,41E+09 | 2,37E+09 |
| 7,1 | | 0,6229637 | 0,7170409 | 0,82 | 1,83E+07 | 6,64E+06 |
| 37 | | 0,6231658 | 0,7172605 | 0,95 | 7,05E+09 | 5,40E+09 |
| 70,5 | | 0,6239686 | 0,7183588 | 0,97 | 8,60E+09 | 8,14E+09 |
| 12,7 | | 0,6243191 | 0,7187688 | 1,09 | 9,20E+06 | 5,53E+06 |
| 17,6 | | 0,6245605 | 0,7190218 | 0,96 | 1,77E+07 | 1,53E+07 |
| 52,5 | | 0,6250666 | 0,7194095 | 1,06 | 9,92E+07 | 8,58E+07 |
| 14,2 | | 0,6251309 | 0,7194883 | 0,94 | 1,75E+08 | 1,69E+08 |
| 5,4 | | 0,6254205 | 0,7198198 | 0,91 | 4,26E+06 | 6,69E+06 |
| 3,6 | | 0,6255690 | 0,7198186 | 1,25 | 8,49E+06 | 8,07E+06 |
| 6,3 | | 0,6259874 | 0,7201746 | 0,93 | 9,12E+06 | 7,07E+06 |
| 21,3 | | 0,6261084 | 0,7202754 | 1,06 | 3,15E+07 | 3,39E+07 |
| 53,2 | | 0,6268961 | 0,7213630 | 1,09 | 1,48E+08 | 1,32E+08 |
| 35,7 | | 0,6284679 | 0,7234562 | 1,04 | 3,73E+07 | 3,78E+07 |
| 31 | | 0,6291116 | 0,7242691 | 0,96 | 6,30E+07 | 5,92E+07 |
| 3,6 | | 0,6292021 | 0,7242881 | 0,84 | 1,26E+07 | 2,05E+07 |
| 10 | | 0,6293448 | 0,7243771 | 1,07 | 4,97E+06 | 5,02E+06 |
| 12 | | 0,6296050 | 0,7244581 | 1,19 | 4,20E+06 | 1,87E+06 |
| 21,8 | | 0,6304880 | 0,7253937 | 1,07 | 2,58E+07 | 2,46E+07 |
| 41,1 | | 0,6318297 | 0,7273008 | 1,54 | 1,64E+07 | 1,29E+06 |
| 22,2 | | 0,6326924 | 0,7285064 | 0,92 | 4,11E+07 | 3,58E+07 |
| 14,4 | | 0,6332765 | 0,7291445 | 1,16 | 7,98E+06 | 4,81E+06 |
| 34,2 | | 0,6337327 | 0,7296426 | 0,96 | 1,43E+08 | 1,57E+08 |
| 35,3 | | 0,6338892 | 0,7298067 | 1,02 | 1,04E+08 | 9,79E+07 |
| 5 | | 0,6349408 | 0,7315424 | 0,91 | 7,31E+06 | 7,51E+06 |
| 7,1 | | 0,6359728 | 0,7326740 | 1,23 | 1,04E+07 | 3,63E+06 |
| 10,4 | | 0,6362832 | 0,7331804 | 0,95 | 3,52E+07 | 3,99E+07 |
| 25,4 | | 0,6379525 | 0,7356021 | 0,91 | 7,36E+06 | 4,87E+06 |
| 65,5 | | 0,6383659 | 0,7360398 | 0,90 | 2,04E+08 | 1,88E+08 |
| 38,7 | | 0,6386657 | 0,7363733 | 1,13 | 1,39E+08 | 1,18E+08 |
| 10,1 | | 0,6390294 | 0,7368617 | 0,89 | 1,06E+07 | 6,51E+06 |
| 17,9 | | 0,6393016 | 0,7370325 | 0,89 | 1,30E+07 | 2,64E+07 |
| 33,8 | | 0,6409814 | 0,7393316 | 0,97 | 6,32E+08 | 6,57E+08 |
| 49,8 | | 0,6410681 | 0,7393495 | 0,98 | 1,73E+09 | 1,78E+09 |
| 18,3 | | 0,6412284 | 0,7394623 | 1,04 | 1,17E+08 | 1,16E+08 |
| 6 | + | 0,6415185 | 0,7396999 | 0,89 | 4,10E+06 | 4,81E+06 |
| 26,8 | | 0,6416190 | 0,7398028 | 1,04 | 1,89E+08 | 2,37E+08 |
| 40,1 | | 0,6417929 | 0,7398212 | 0,97 | 2,73E+08 | 2,56E+08 |
| 4,4 | | 0,6425876 | 0,7408918 | 1,09 | 1,10E+07 | 9,00E+06 |
| 54,8 | | 0,6435839 | 0,7421255 | 0,95 | 5,99E+07 | 5,71E+07 |
| 31,3 | | 0,6444028 | 0,7430252 | 1,06 | 4,54E+07 | 4,38E+07 |
| 31,2 | | 0,6445407 | 0,7430627 | 0,88 | 6,49E+06 | 6,97E+06 |
| 10 | | 0,6450323 | 0,7437039 | 1,11 | 2,82E+06 | 3,99E+06 |
| 6,6 | | 0,6453957 | 0,7439957 | 1,11 | 1,94E+06 | 3,53E+06 |
| 12,4 | | 0,6461244 | 0,7447876 | 1,04 | 5,34E+07 | 6,23E+07 |
| 35,5 | | 0,6464209 | 0,7451428 | 1,04 | 1,33E+08 | 1,22E+08 |
| 4 | | 0,6468019 | 0,7455167 | 0,93 | 4,59E+06 | 5,37E+06 |
| 2,7 | | 0,6471966 | 0,7461314 | 1,19 | 2,29E+06 | 2,06E+06 |
| 72,3 | | 0,6473473 | 0,7462283 | 1,01 | 3,06E+09 | 3,16E+09 |
| 44,1 | | 0,6474608 | 0,7462841 | 1,04 | 1,41E+08 | 1,28E+08 |

| | | | | | | |
|---|---|---|---|---|---|---|
| 6,2 | | 0,6477910 | 0,7467090 | 1,05 | 1,36E+07 | 1,51E+07 |
| 61,3 | | 0,6483620 | 0,7474020 | 0,94 | 1,68E+09 | 1,92E+09 |
| 9,6 | | 0,6486114 | 0,7476708 | 0,85 | 2,40E+06 | 6,41E+06 |
| 32,5 | | 0,6495109 | 0,7489166 | 0,96 | 9,25E+08 | 1,03E+09 |
| 42,6 | | 0,6497265 | 0,7490808 | 1,06 | 6,17E+07 | 6,13E+07 |
| 26,4 | | 0,6509592 | 0,7504176 | 1,72 | 4,96E+06 | 1,20E+08 |
| 14,8 | | 0,6510540 | 0,7504508 | 1,06 | 1,51E+07 | 1,20E+07 |
| 35,2 | | 0,6512158 | 0,7505127 | 1,02 | 2,29E+08 | 2,30E+08 |
| 29,9 | | 0,6516446 | 0,7510566 | 1,05 | 8,17E+07 | 7,03E+07 |
| 42,1 | | 0,6522985 | 0,7517346 | 1,04 | 1,14E+09 | 9,03E+08 |
| 38 | | 0,6526163 | 0,7520744 | 0,97 | 3,30E+08 | 3,50E+08 |
| 16,8 | | 0,6526841 | 0,7520634 | 0,94 | 1,56E+08 | 2,11E+08 |
| 22,8 | | 0,6537352 | 0,7536807 | 0,89 | 3,29E+07 | 1,47E+07 |
| 55,3 | | 0,6538404 | 0,7536817 | 0,97 | 2,68E+08 | 2,77E+08 |
| 34,2 | | 0,6550097 | 0,7551233 | 0,97 | 8,28E+07 | 7,30E+07 |
| 62,3 | | 0,6564576 | 0,7564084 | 0,98 | 7,21E+08 | 7,81E+08 |
| 19,3 | | 0,6565156 | 0,7563974 | 0,97 | 4,88E+07 | 4,06E+07 |
| 43,8 | | 0,6571276 | 0,7570439 | 0,97 | 2,94E+08 | 3,68E+08 |
| 65,6 | | 0,6575327 | 0,7576335 | 0,97 | 5,80E+09 | 6,68E+09 |
| 10,7 | | 0,6579000 | 0,7579446 | 0,86 | 1,64E+06 | 3,58E+06 |
| 12,5 | | 0,6579343 | 0,7578870 | 0,94 | 8,71E+06 | 1,07E+07 |
| 13,4 | | 0,6582418 | 0,7582132 | 0,94 | 2,61E+07 | 2,15E+07 |
| 48,1 | | 0,6584733 | 0,7583401 | 0,99 | 4,32E+08 | 4,24E+08 |
| 13,1 | | 0,6591011 | 0,7591342 | 1,08 | 1,03E+07 | 1,16E+07 |
| 68,6 | | 0,6605576 | 0,7613167 | 0,96 | 5,68E+09 | 5,61E+09 |
| 44,1 | + | 0,6606788 | 0,7613475 | 0,85 | 1,04E+08 | 5,89E+07 |
| 46,7 | | 0,6609168 | 0,7614112 | 1,02 | 8,01E+07 | 7,62E+07 |
| 31 | | 0,6615159 | 0,7622117 | 0,94 | 1,15E+08 | 8,42E+07 |
| 19,6 | | 0,6617050 | 0,7622917 | 0,94 | 3,97E+07 | 4,77E+07 |
| 24 | | 0,6619795 | 0,7627129 | 0,93 | 5,41E+07 | 7,12E+07 |
| 8,1 | | 0,6621348 | 0,7628280 | 1,16 | 8,33E+06 | 1,18E+07 |
| 43,8 | | 0,6624345 | 0,7631444 | 1,06 | 7,86E+08 | 7,04E+08 |
| 23,9 | | 0,6626323 | 0,7632947 | 0,95 | 1,16E+07 | 1,48E+07 |
| 40 | | 0,6628031 | 0,7633477 | 0,98 | 2,07E+08 | 2,34E+08 |
| 27,5 | | 0,6639314 | 0,7649117 | 1,05 | 2,52E+07 | 2,34E+07 |
| 24,4 | | 0,6644210 | 0,7654810 | 1,10 | 2,35E+07 | 1,48E+07 |
| 5,6 | | 0,6644384 | 0,7653793 | 0,88 | 9,08E+06 | 3,81E+06 |
| 6 | | 0,6645787 | 0,7654739 | 1,61 | 6,10E+06 | 3,90E+05 |
| 19,3 | | 0,6648035 | 0,7657865 | 1,03 | 1,59E+08 | 1,54E+08 |
| 1,8 | | 0,6657932 | 0,7672451 | 1,26 | 3,07E+06 | 2,30E+06 |
| 7,4 | | 0,6658262 | 0,7671911 | 1,14 | 2,66E+06 | 2,38E+06 |
| 6,7 | | 0,6658309 | 0,7670747 | 1,11 | 5,49E+06 | 3,41E+06 |
| 5 | | 0,6663719 | 0,7677171 | 0,90 | 2,55E+06 | 1,79E+06 |
| 25,8 | | 0,6668962 | 0,7682489 | 0,93 | 9,31E+07 | 9,23E+07 |
| 3,3 | | 0,6670332 | 0,7683520 | 1,24 | 4,70E+06 | 4,47E+06 |
| 7,4 | | 0,6676740 | 0,7691135 | 0,69 | 1,54E+07 | 1,55E+07 |
| 2,3 | | 0,6686962 | 0,7703443 | 0,79 | 5,39E+06 | 3,62E+06 |
| 7,1 | | 0,6693980 | 0,7710955 | 1,61 | 3,25E+07 | 1,49E+06 |
| 43,6 | | 0,6696005 | 0,7712194 | 1,03 | 2,96E+08 | 3,51E+08 |
| 45,6 | | 0,6711509 | 0,7731904 | 0,97 | 5,81E+08 | 5,31E+08 |

| | | | | | | |
|---|---|---|---|---|---|---|
| 17 | | 0,6714601 | 0,7734473 | 1,14 | 8,18E+06 | 5,54E+06 |
| 7,4 | | 0,6727733 | 0,7750403 | 0,63 | 8,88E+06 | 6,86E+06 |
| 18,7 | | 0,6736479 | 0,7759921 | 0,51 | 2,97E+07 | 9,96E+06 |
| 23,2 | | 0,6743649 | 0,7768152 | 0,85 | 2,45E+07 | 1,81E+07 |
| 52,2 | | 0,6745315 | 0,7769029 | 1,02 | 1,21E+09 | 1,35E+09 |
| 25,7 | | 0,6748292 | 0,7771051 | 0,51 | 6,40E+07 | 5,65E+07 |
| 11 | | 0,6748827 | 0,7771086 | 1,20 | 2,01E+06 | 5,77E+06 |
| 2,9 | + | 0,6756322 | 0,7778219 | 0,73 | 2,65E+06 | 1,97E+06 |
| 26,9 | | 0,6756371 | 0,7777229 | 0,92 | 1,24E+07 | 1,54E+07 |
| 38,8 | | 0,6756577 | 0,7776434 | 1,06 | 9,51E+07 | 1,23E+08 |
| 26,6 | | 0,6756725 | 0,7775257 | 0,94 | 5,48E+07 | 5,65E+07 |
| 1,4 | | 0,6756808 | 0,7774268 | 0,79 | 3,04E+06 | 1,19E+07 |
| 53,8 | | 0,6759257 | 0,7778429 | 1,03 | 5,85E+08 | 4,92E+08 |
| 14,6 | | 0,6762575 | 0,7780744 | 1,05 | 1,95E+07 | 2,78E+07 |
| 10,8 | | 0,6769701 | 0,7789660 | 0,83 | 2,15E+06 | 5,66E+06 |
| 21,9 | | 0,6773916 | 0,7793551 | 0,78 | 6,73E+06 | 1,86E+06 |
| 7,9 | | 0,6779026 | 0,7798107 | 0,88 | 3,46E+06 | 4,79E+06 |
| 47,5 | | 0,6781427 | 0,7799389 | 1,04 | 1,60E+09 | 1,65E+09 |
| 12,6 | | 0,6789467 | 0,7807051 | 0,80 | 6,50E+06 | 5,66E+06 |
| 46,8 | | 0,6790155 | 0,7806747 | 1,05 | 6,22E+08 | 6,57E+08 |
| 2 | | 0,6792748 | 0,7809225 | 0,77 | 4,06E+06 | 1,91E+06 |
| 35 | | 0,6793547 | 0,7808710 | 1,21 | 3,38E+07 | 3,84E+07 |
| 70,1 | | 0,6795664 | 0,7811730 | 0,94 | 8,57E+08 | 6,46E+08 |
| 58,4 | | 0,6799495 | 0,7816949 | 1,05 | 2,36E+08 | 3,30E+08 |
| 14,2 | | 0,6802272 | 0,7819254 | 1,04 | 9,46E+06 | 1,22E+07 |
| 24,9 | | 0,6804618 | 0,7820242 | 0,89 | 2,09E+07 | 1,00E+07 |
| 44,1 | | 0,6810133 | 0,7824436 | 0,89 | 2,83E+07 | 2,73E+07 |
| 17,9 | | 0,6811773 | 0,7826999 | 1,28 | 1,06E+07 | 6,66E+06 |
| 1 | | 0,6816765 | 0,7829989 | 0,81 | 8,50E+05 | 1,11E+06 |
| 4,8 | | 0,6821090 | 0,7835225 | 0,68 | 1,00E+06 | 1,47E+06 |
| 36,5 | | 0,6822758 | 0,7835624 | 1,06 | 6,24E+07 | 9,13E+07 |
| 13,2 | | 0,6826607 | 0,7839221 | 1,14 | 3,48E+06 | 7,62E+06 |
| 10,5 | | 0,6844888 | 0,7860202 | 0,93 | 1,28E+07 | 1,58E+07 |
| 75,8 | | 0,6849143 | 0,7864507 | 0,96 | 4,29E+08 | 5,06E+08 |
| 10,1 | | 0,6855473 | 0,7873533 | 1,13 | 4,84E+06 | 1,04E+07 |
| 8,1 | | 0,6856555 | 0,7873010 | 1,15 | 1,05E+07 | 4,68E+06 |
| 10,6 | | 0,6856834 | 0,7872552 | 1,08 | 8,07E+06 | 7,08E+06 |
| 21,8 | | 0,6867819 | 0,7887700 | 1,26 | 5,86E+06 | 7,11E+06 |
| 45,7 | | 0,6873535 | 0,7892757 | 1,07 | 6,66E+07 | 8,93E+07 |
| 18,1 | | 0,6873720 | 0,7892086 | 0,65 | 1,15E+08 | 6,50E+06 |
| 9,2 | | 0,6875460 | 0,7891999 | 1,09 | 3,03E+07 | 1,62E+07 |
| 60 | | 0,6876185 | 0,7892160 | 1,02 | 5,14E+08 | 5,49E+08 |
| 18,6 | | 0,6878478 | 0,7894948 | 0,95 | 3,15E+07 | 2,51E+07 |
| 9,8 | | 0,6886232 | 0,7902054 | 1,11 | 2,30E+07 | 1,82E+07 |
| 49,7 | | 0,6886283 | 0,7900864 | 0,97 | 2,04E+08 | 2,37E+08 |
| 21,2 | | 0,6887915 | 0,7903162 | 0,97 | 3,97E+08 | 4,28E+08 |
| 33,3 | | 0,6895555 | 0,7910811 | 0,96 | 1,09E+08 | 1,30E+08 |
| 22,3 | | 0,6902930 | 0,7919266 | 0,80 | 1,47E+07 | 2,56E+06 |
| 40,3 | | 0,6903977 | 0,7920265 | 0,94 | 5,46E+08 | 4,22E+08 |
| 42,8 | | 0,6904657 | 0,7919236 | 1,08 | 5,46E+07 | 4,48E+07 |

| | | | | | | |
|---|---|---|---|---|---|---|
| 2,8 | | 0,6906578 | 0,7920427 | 1,04 | 7,98E+07 | 9,24E+07 |
| 33,6 | | 0,6907883 | 0,7920836 | 0,95 | 1,11E+08 | 1,03E+08 |
| 51,8 | | 0,6909039 | 0,7921311 | 1,01 | 1,06E+09 | 1,03E+09 |
| 15,8 | | 0,6909102 | 0,7920120 | 0,98 | 6,33E+07 | 5,87E+07 |
| 5,1 | | 0,6909442 | 0,7919393 | 0,95 | 3,12E+06 | 3,37E+06 |
| 20,4 | | 0,6914566 | 0,7924360 | 1,05 | 8,59E+07 | 1,28E+08 |
| 19,4 | | 0,6933090 | 0,7943580 | 1,04 | 3,17E+07 | 2,70E+07 |
| 13,2 | | 0,6933286 | 0,7942759 | 1,07 | 4,71E+06 | 6,20E+06 |
| 39,8 | | 0,6940429 | 0,7951128 | 0,97 | 4,35E+08 | 3,46E+08 |
| 30,6 | | 0,6952057 | 0,7964724 | 1,26 | 1,05E+07 | 2,39E+06 |
| 9,6 | | 0,6959513 | 0,7972456 | 0,83 | 1,65E+07 | 1,97E+07 |
| 13,6 | | 0,6968508 | 0,7983178 | 0,92 | 6,49E+07 | 5,64E+07 |
| 8,6 | | 0,6973390 | 0,7987241 | 0,69 | 9,27E+05 | 5,86E+06 |
| 15,2 | | 0,6973549 | 0,7986042 | 0,97 | 3,12E+07 | 3,06E+07 |
| 26,3 | | 0,6974499 | 0,7985960 | 0,88 | 3,35E+06 | 7,39E+06 |
| 16,7 | | 0,6978616 | 0,7988640 | 1,22 | 5,27E+07 | 4,29E+07 |
| 43,4 | | 0,6995694 | 0,8005655 | 0,95 | 2,59E+08 | 3,65E+08 |
| 53,2 | | 0,6997838 | 0,8006819 | 0,96 | 2,94E+08 | 2,60E+08 |
| 69,5 | | 0,7000273 | 0,8007730 | 0,98 | 1,18E+09 | 1,17E+09 |
| 31,6 | | 0,7003768 | 0,8010681 | 0,98 | 1,16E+08 | 1,41E+08 |
| 43,9 | | 0,7005986 | 0,8011897 | 0,99 | 1,00E+08 | 1,01E+08 |
| 8,2 | | 0,7006137 | 0,8010984 | 1,16 | 1,43E+06 | 2,85E+06 |
| 11,5 | | 0,7009778 | 0,8014605 | 1,03 | 1,08E+08 | 9,85E+07 |
| 26,4 | | 0,7014483 | 0,8019262 | 0,96 | 4,01E+07 | 4,74E+07 |
| 10,5 | | 0,7030674 | 0,8040210 | 0,83 | 6,27E+06 | 2,29E+06 |
| 11,1 | + | 0,7036063 | 0,8044886 | 0,89 | 1,92E+07 | 3,71E+07 |
| 3,9 | | 0,7039493 | 0,8048356 | 0,89 | 5,56E+06 | 3,53E+06 |
| 77,8 | | 0,7040758 | 0,8048169 | 1,08 | 3,38E+08 | 4,91E+08 |
| 12,8 | | 0,7049654 | 0,8056431 | 1,13 | 4,98E+06 | 3,06E+06 |
| 39,9 | | 0,7051082 | 0,8056777 | 1,06 | 5,20E+07 | 5,35E+07 |
| 9,5 | | 0,7052268 | 0,8057487 | 0,83 | 6,16E+06 | 8,62E+06 |
| 27,9 | | 0,7053764 | 0,8057394 | 1,06 | 8,40E+07 | 1,03E+08 |
| 9,8 | | 0,7055008 | 0,8058146 | 0,95 | 6,93E+07 | 8,47E+07 |
| 34,5 | | 0,7061659 | 0,8065874 | 0,96 | 7,90E+07 | 9,51E+07 |
| 36,1 | | 0,7067120 | 0,8071073 | 1,03 | 2,67E+08 | 2,22E+08 |
| 17,9 | | 0,7070954 | 0,8074726 | 1,07 | 1,22E+07 | 1,88E+07 |
| 18 | | 0,7076289 | 0,8081227 | 0,86 | 8,09E+06 | 2,77E+06 |
| 5,2 | | 0,7081407 | 0,8084841 | 0,91 | 2,49E+06 | 3,54E+06 |
| 34,2 | | 0,7084641 | 0,8088138 | 1,05 | 3,38E+07 | 4,33E+07 |
| 10,1 | | 0,7086415 | 0,8088950 | 0,95 | 4,43E+07 | 6,18E+07 |
| 49,9 | | 0,7089207 | 0,8090947 | 1,02 | 2,72E+08 | 3,05E+08 |
| 36,2 | | 0,7092955 | 0,8094270 | 1,05 | 2,14E+07 | 1,82E+07 |
| 31,6 | | 0,7095712 | 0,8098514 | 1,04 | 1,67E+08 | 1,81E+08 |
| 37,9 | | 0,7097547 | 0,8099659 | 0,97 | 4,16E+08 | 3,73E+08 |
| 12,6 | | 0,7113962 | 0,8118392 | 0,92 | 5,27E+06 | 7,98E+06 |
| 47,1 | | 0,7115918 | 0,8119611 | 0,98 | 9,42E+08 | 9,78E+08 |
| 24,6 | | 0,7123592 | 0,8126305 | 0,97 | 1,52E+08 | 1,28E+08 |
| 73,8 | | 0,7128802 | 0,8130481 | 1,03 | 5,85E+08 | 6,81E+08 |
| 52,2 | | 0,7148942 | 0,8151773 | 0,95 | 1,73E+09 | 1,77E+09 |
| 5,4 | | 0,7150547 | 0,8152599 | 0,95 | 4,58E+06 | 5,99E+06 |

| | | | | | | |
|---|---|---|---|---|---|---|
| 9,1 | | 0,7153595 | 0,8155341 | 1,30 | 3,39E+06 | 2,45E+06 |
| 14,8 | | 0,7153602 | 0,8154124 | 0,88 | 2,96E+07 | 1,21E+07 |
| 41,3 | | 0,7157146 | 0,8157206 | 0,95 | 1,46E+09 | 1,59E+09 |
| 36,8 | | 0,7160384 | 0,8159952 | 0,98 | 1,67E+08 | 1,70E+08 |
| 9 | | 0,7166288 | 0,8165539 | 0,93 | 9,41E+06 | 6,01E+06 |
| 31,7 | | 0,7166523 | 0,8164321 | 1,15 | 1,18E+08 | 4,54E+07 |
| 43 | | 0,7171266 | 0,8169023 | 1,11 | 2,66E+07 | 1,46E+07 |
| 6,4 | | 0,7174365 | 0,8172267 | 0,87 | 2,92E+06 | 7,66E+06 |
| 29,3 | + | 0,7177339 | 0,8175111 | 1,21 | 2,34E+08 | 1,08E+08 |
| 6,2 | | 0,7183780 | 0,8179332 | 1,07 | 2,74E+06 | 2,35E+06 |
| 3,2 | | 0,7189229 | 0,8183390 | 1,43 | 1,92E+07 | 1,65E+06 |
| 3,9 | | 0,7191761 | 0,8184735 | 0,85 | 3,36E+06 | 5,59E+06 |
| 12,3 | | 0,7194374 | 0,8185613 | 1,07 | 9,27E+06 | 6,55E+06 |
| 9,9 | | 0,7197260 | 0,8188131 | 0,79 | 2,28E+06 | 5,47E+06 |
| 49,6 | | 0,7197547 | 0,8187072 | 0,97 | 3,46E+09 | 3,93E+09 |
| 5,7 | | 0,7203634 | 0,8194285 | 0,91 | 6,24E+06 | 6,46E+06 |
| 30 | | 0,7208221 | 0,8198099 | 0,89 | 5,66E+06 | 3,27E+06 |
| 14,2 | | 0,7210815 | 0,8200244 | 0,95 | 6,59E+07 | 6,83E+07 |
| 2,7 | | 0,7212526 | 0,8200042 | 1,13 | 2,70E+06 | 1,88E+06 |
| 2,7 | | 0,7213587 | 0,8200655 | 0,75 | 1,59E+07 | 1,16E+07 |
| 16,7 | | 0,7217696 | 0,8202841 | 0,94 | 1,50E+08 | 1,39E+08 |
| 5,5 | | 0,7222147 | 0,8206258 | 1,60 | 2,64E+06 | 6,84E+07 |
| 27,8 | | 0,7225877 | 0,8208323 | 1,09 | 4,71E+07 | 4,17E+07 |
| 12,7 | | 0,7231560 | 0,8213250 | 0,74 | 5,20E+06 | 1,54E+07 |
| 2,7 | | 0,7242121 | 0,8223776 | 0,91 | 6,91E+06 | 6,69E+06 |
| 2,7 | | 0,7244623 | 0,8226677 | 0,87 | 6,07E+06 | 2,80E+06 |
| 22,7 | | 0,7255968 | 0,8236972 | 0,94 | 6,10E+08 | 5,25E+08 |
| 53,3 | | 0,7257930 | 0,8238186 | 0,97 | 2,53E+08 | 3,17E+08 |
| 10,5 | | 0,7259162 | 0,8238537 | 1,05 | 1,77E+07 | 1,41E+07 |
| 11,5 | | 0,7261271 | 0,8239548 | 0,96 | 8,96E+07 | 7,27E+07 |
| 54,7 | | 0,7269276 | 0,8246886 | 0,96 | 1,88E+08 | 1,95E+08 |
| 38,2 | | 0,7280196 | 0,8257206 | 0,98 | 2,95E+08 | 2,80E+08 |
| 5 | | 0,7286103 | 0,8260895 | 0,87 | 9,23E+06 | 2,65E+06 |
| 14,7 | | 0,7286126 | 0,8259851 | 0,92 | 1,22E+07 | 1,06E+07 |
| 72,6 | | 0,7289337 | 0,8261780 | 1,09 | 5,18E+08 | 8,83E+08 |
| 13,8 | | 0,7290167 | 0,8261462 | 0,93 | 2,16E+07 | 3,60E+07 |
| 17,7 | | 0,7291885 | 0,8261352 | 0,91 | 1,33E+07 | 1,89E+07 |
| 26,1 | | 0,7300447 | 0,8268571 | 0,87 | 1,86E+08 | 5,79E+08 |
| 17,1 | | 0,7305079 | 0,8272552 | 0,97 | 2,17E+07 | 2,81E+07 |
| 41,5 | | 0,7308479 | 0,8274323 | 0,98 | 3,74E+08 | 4,39E+08 |
| 7,9 | | 0,7321467 | 0,8288241 | 1,11 | 5,92E+06 | 9,81E+06 |
| 31,6 | | 0,7325101 | 0,8291517 | 1,04 | 6,60E+07 | 5,86E+07 |
| 18,7 | | 0,7328785 | 0,8294334 | 1,03 | 9,78E+07 | 1,17E+08 |
| 3,8 | | 0,7330731 | 0,8295270 | 0,85 | 6,49E+06 | 6,08E+06 |
| 49,1 | | 0,7332978 | 0,8296508 | 1,05 | 4,39E+07 | 3,08E+07 |
| 49,6 | | 0,7342751 | 0,8305731 | 0,98 | 6,22E+08 | 7,32E+08 |
| 27,5 | | 0,7343613 | 0,8305668 | 1,04 | 1,24E+07 | 1,21E+07 |
| 5,8 | | 0,7344341 | 0,8305890 | 0,96 | 3,87E+06 | 4,30E+06 |
| 19,7 | | 0,7360827 | 0,8325130 | 1,19 | 1,32E+07 | 5,76E+07 |
| 2 | | 0,7361998 | 0,8325491 | 0,66 | 3,22E+06 | 3,90E+07 |

| | | | | | | |
|---|---|---|---|---|---|---|
| 51,9 | | 0,7365244 | 0,8328513 | 0,98 | 7,87E+08 | 8,41E+08 |
| 8,4 | | 0,7366768 | 0,8328743 | 1,06 | 6,02E+07 | 6,18E+07 |
| 15,1 | | 0,7377179 | 0,8339659 | 1,11 | 1,42E+07 | 5,33E+06 |
| 7,4 | | 0,7380515 | 0,8340587 | 0,96 | 4,25E+06 | 3,68E+06 |
| 22,9 | | 0,7395803 | 0,8354650 | 1,05 | 6,70E+07 | 1,03E+08 |
| 17,7 | | 0,7401327 | 0,8358211 | 0,91 | 1,25E+07 | 2,93E+07 |
| 3,5 | | 0,7404259 | 0,8359343 | 1,10 | 5,35E+06 | 2,45E+06 |
| 66,5 | | 0,7408371 | 0,8362576 | 0,97 | 1,07E+09 | 1,21E+09 |
| 36,5 | | 0,7410145 | 0,8363044 | 1,03 | 1,68E+08 | 1,94E+08 |
| 46,8 | | 0,7419118 | 0,8370977 | 1,02 | 9,58E+08 | 1,02E+09 |
| 11,7 | | 0,7438416 | 0,8390647 | 0,93 | 5,53E+06 | 8,48E+06 |
| 16 | | 0,7441359 | 0,8393779 | 0,76 | 7,99E+07 | 2,87E+08 |
| 8,9 | | 0,7446148 | 0,8397893 | 1,22 | 2,07E+06 | 1,22E+06 |
| 31,3 | | 0,7447050 | 0,8397994 | 1,07 | 1,71E+07 | 2,17E+07 |
| 6,9 | | 0,7448860 | 0,8398391 | 0,95 | 2,27E+07 | 1,59E+07 |
| 10,7 | | 0,7450094 | 0,8399071 | 1,11 | 2,75E+06 | 4,92E+06 |
| 5,4 | | 0,7457840 | 0,8407287 | 0,98 | 1,19E+07 | 1,27E+07 |
| 46,3 | | 0,7462990 | 0,8409486 | 1,03 | 6,55E+09 | 6,67E+09 |
| 7,1 | | 0,7467970 | 0,8413647 | 0,84 | 1,44E+07 | 3,05E+06 |
| 8,9 | | 0,7468088 | 0,8412539 | 1,02 | 1,42E+07 | 1,33E+07 |
| 2,3 | | 0,7468695 | 0,8411296 | 1,21 | 1,06E+07 | 1,28E+07 |
| 6 | | 0,7468923 | 0,8410195 | 1,06 | 5,01E+06 | 8,47E+06 |
| 6,4 | | 0,7483568 | 0,8427307 | 1,12 | 2,18E+06 | 3,45E+06 |
| 3,9 | | 0,7485212 | 0,8428721 | 1,22 | 6,74E+06 | 2,03E+06 |
| 11,8 | | 0,7488218 | 0,8431091 | 0,97 | 1,93E+07 | 2,44E+07 |
| 10,7 | | 0,7492145 | 0,8433349 | 1,18 | 3,21E+06 | 7,52E+05 |
| 5,9 | | 0,7493915 | 0,8433722 | 1,06 | 6,45E+06 | 5,53E+06 |
| 20,9 | | 0,7494461 | 0,8433310 | 1,08 | 2,92E+07 | 1,55E+07 |
| 6,2 | | 0,7495393 | 0,8434125 | 1,05 | 1,74E+07 | 1,38E+07 |
| 15,4 | | 0,7500966 | 0,8438678 | 0,92 | 1,53E+07 | 3,29E+07 |
| 19,5 | | 0,7503339 | 0,8440655 | 0,98 | 7,58E+07 | 8,59E+07 |
| 55,3 | | 0,7517745 | 0,8453396 | 1,03 | 1,08E+08 | 8,38E+07 |
| 75,1 | | 0,7517840 | 0,8452338 | 1,03 | 6,21E+09 | 6,72E+09 |
| 51,7 | | 0,7521782 | 0,8456177 | 0,97 | 2,18E+09 | 2,64E+09 |
| 19 | | 0,7527041 | 0,8459932 | 1,06 | 3,56E+07 | 2,08E+07 |
| 13,6 | | 0,7535846 | 0,8467243 | 1,06 | 6,23E+06 | 5,26E+06 |
| 8,3 | | 0,7538264 | 0,8468176 | 1,09 | 2,52E+06 | 2,66E+06 |
| 12,8 | | 0,7541233 | 0,8471403 | 0,95 | 1,95E+07 | 1,45E+07 |
| 22,7 | | 0,7544145 | 0,8474877 | 1,03 | 3,03E+07 | 2,68E+07 |
| 13,1 | | 0,7548407 | 0,8478350 | 1,08 | 1,21E+07 | 2,37E+07 |
| 31,7 | | 0,7553350 | 0,8481244 | 1,03 | 6,99E+07 | 5,94E+07 |
| 3,6 | | 0,7560515 | 0,8489364 | 0,91 | 7,19E+06 | 5,46E+06 |
| 5,8 | | 0,7564603 | 0,8492349 | 1,06 | 1,44E+07 | 8,99E+06 |
| 50,2 | | 0,7566497 | 0,8492890 | 1,02 | 4,38E+09 | 3,92E+09 |
| 7,7 | | 0,7569990 | 0,8496858 | 1,12 | 4,56E+06 | 5,51E+06 |
| 16,2 | | 0,7572737 | 0,8499794 | 0,82 | 6,28E+06 | 2,94E+06 |
| 9,3 | | 0,7577650 | 0,8504449 | 0,92 | 1,48E+07 | 1,57E+07 |
| 33,3 | | 0,7583551 | 0,8510810 | 1,04 | 8,47E+08 | 1,17E+09 |
| 31 | | 0,7586970 | 0,8513542 | 0,89 | 1,26E+08 | 1,11E+08 |
| 10,4 | | 0,7597063 | 0,8523355 | 1,22 | 6,87E+06 | 7,31E+06 |

| | | | | | | |
|---|---|---|---|---|---|---|
| 37,1 | | 0,7598399 | 0,8523813 | 1,01 | 2,92E+08 | 3,16E+08 |
| 20,6 | | 0,7598901 | 0,8522742 | 1,02 | 3,57E+07 | 3,06E+07 |
| 66,7 | | 0,7599784 | 0,8521965 | 0,97 | 3,17E+08 | 2,58E+08 |
| 20,8 | | 0,7607058 | 0,8527812 | 1,09 | 9,26E+06 | 6,35E+06 |
| 27,8 | | 0,7608919 | 0,8529016 | 1,06 | 4,10E+08 | 4,52E+08 |
| 73,2 | | 0,7617609 | 0,8537818 | 0,98 | 2,71E+09 | 2,85E+09 |
| 19 | | 0,7620306 | 0,8540156 | 0,99 | 1,65E+08 | 1,53E+08 |
| 4 | | 0,7628401 | 0,8546855 | 0,94 | 4,94E+06 | 6,37E+06 |
| 21,3 | | 0,7629321 | 0,8546193 | 0,83 | 2,56E+06 | 6,26E+06 |
| 50,7 | | 0,7630020 | 0,8545754 | 0,94 | 5,81E+07 | 1,13E+08 |
| 27,9 | | 0,7630085 | 0,8544680 | 0,98 | 6,78E+07 | 7,82E+07 |
| 3,1 | | 0,7630406 | 0,8543566 | 0,72 | 2,06E+06 | 1,15E+06 |
| 18,8 | | 0,7639439 | 0,8553432 | 0,91 | 2,33E+06 | 3,86E+06 |
| 44,8 | | 0,7642440 | 0,8556687 | 1,02 | 5,57E+08 | 4,85E+08 |
| 8,2 | | 0,7643407 | 0,8556864 | 0,97 | 1,80E+07 | 1,29E+07 |
| 31 | | 0,7644239 | 0,8556483 | 0,95 | 1,50E+07 | 1,67E+07 |
| 58 | | 0,7644270 | 0,8555227 | 1,02 | 3,03E+08 | 3,63E+08 |
| 40,4 | | 0,7653219 | 0,8564555 | 0,99 | 1,43E+08 | 1,57E+08 |
| 49 | | 0,7656963 | 0,8567994 | 1,01 | 2,07E+08 | 1,99E+08 |
| 36,9 | | 0,7660521 | 0,8571015 | 1,02 | 2,16E+08 | 2,37E+08 |
| 15 | | 0,7660609 | 0,8569758 | 1,08 | 8,99E+07 | 8,37E+07 |
| 72,9 | | 0,7664640 | 0,8572379 | 1,03 | 3,77E+08 | 4,09E+08 |
| 44,8 | | 0,7666799 | 0,8572365 | 0,95 | 1,39E+08 | 2,21E+08 |
| 71,8 | | 0,7669215 | 0,8574328 | 1,05 | 3,13E+08 | 3,52E+08 |
| 7,2 | | 0,7672299 | 0,8575804 | 0,89 | 3,17E+06 | 9,83E+05 |
| 18,5 | | 0,7677863 | 0,8580340 | 1,05 | 8,66E+06 | 9,64E+06 |
| 19,8 | | 0,7678593 | 0,8579405 | 1,06 | 1,16E+07 | 1,73E+07 |
| 18,6 | | 0,7679646 | 0,8579555 | 0,94 | 5,99E+07 | 3,26E+07 |
| 4 | | 0,7684790 | 0,8586104 | 0,90 | 4,47E+06 | 4,09E+06 |
| 20,3 | | 0,7685765 | 0,8586487 | 0,96 | 3,37E+07 | 4,21E+07 |
| 44,7 | | 0,7685858 | 0,8585387 | 0,97 | 4,23E+08 | 3,66E+08 |
| 20,9 | | 0,7701060 | 0,8599813 | 1,02 | 1,47E+08 | 1,37E+08 |
| 41,2 | | 0,7701260 | 0,8598893 | 1,03 | 1,19E+08 | 9,45E+07 |
| 22,9 | | 0,7702809 | 0,8599777 | 0,96 | 2,14E+08 | 1,79E+08 |
| 66,6 | | 0,7702988 | 0,8598519 | 1,02 | 5,82E+08 | 6,98E+08 |
| 39,9 | | 0,7705049 | 0,8599321 | 0,98 | 1,10E+08 | 1,29E+08 |
| 19,8 | | 0,7736839 | 0,8631371 | 1,03 | 1,99E+07 | 2,31E+07 |
| 6,1 | | 0,7742097 | 0,8636213 | 0,87 | 3,04E+06 | 3,28E+06 |
| 52,9 | | 0,7742958 | 0,8635429 | 0,97 | 6,52E+08 | 8,54E+08 |
| 65 | | 0,7745033 | 0,8635740 | 1,02 | 2,36E+08 | 2,81E+08 |
| 5,1 | | 0,7745435 | 0,8634635 | 1,06 | 6,40E+06 | 5,61E+06 |
| 10,9 | | 0,7753383 | 0,8640234 | 0,91 | 1,90E+07 | 1,18E+07 |
| 35,1 | | 0,7753933 | 0,8639503 | 0,97 | 3,50E+08 | 3,86E+08 |
| 36 | | 0,7756452 | 0,8640567 | 1,01 | 3,13E+07 | 2,99E+07 |
| 40 | | 0,7757162 | 0,8639924 | 0,83 | 5,13E+06 | 5,15E+06 |
| 36,2 | | 0,7770443 | 0,8650330 | 1,01 | 5,15E+08 | 4,86E+08 |
| 6,7 | | 0,7770520 | 0,8649066 | 0,98 | 5,40E+06 | 5,22E+06 |
| 15 | | 0,7779877 | 0,8657387 | 0,97 | 4,21E+07 | 3,93E+07 |
| 43,6 | | 0,7784668 | 0,8660359 | 1,05 | 1,37E+09 | 9,25E+08 |
| 4,1 | + | 0,7813922 | 0,8690850 | 0,89 | 4,49E+06 | 4,32E+06 |

| | | | | | | |
|---|---|---|---|---|---|---|
| 15,4 | | 0,7813940 | 0,8689581 | 0,95 | 3,14E+07 | 1,91E+07 |
| 9,8 | | 0,7815918 | 0,8689784 | 1,27 | 5,33E+06 | 6,06E+05 |
| 17,7 | | 0,7818099 | 0,8691032 | 1,03 | 2,87E+07 | 3,74E+07 |
| 4,2 | | 0,7818844 | 0,8690091 | 1,14 | 6,16E+06 | 1,46E+06 |
| 69,3 | | 0,7826060 | 0,8697627 | 0,97 | 4,32E+08 | 5,64E+08 |
| 62,3 | | 0,7827091 | 0,8697525 | 0,99 | 9,30E+08 | 8,21E+08 |
| 36,1 | | 0,7834255 | 0,8705169 | 1,02 | 4,01E+08 | 4,62E+08 |
| 33,9 | | 0,7847230 | 0,8717240 | 0,96 | 4,78E+07 | 5,99E+07 |
| 4,8 | | 0,7852065 | 0,8719247 | 1,02 | 4,97E+06 | 5,34E+06 |
| 2,9 | | 0,7856921 | 0,8722795 | 0,97 | 1,02E+07 | 1,19E+07 |
| 26,3 | | 0,7858227 | 0,8722339 | 0,98 | 3,72E+08 | 4,01E+08 |
| 41,8 | | 0,7859186 | 0,8721546 | 1,04 | 7,49E+07 | 9,52E+07 |
| 12 | | 0,7865212 | 0,8727803 | 1,04 | 2,99E+07 | 4,62E+07 |
| 7,8 | | 0,7867718 | 0,8727738 | 1,03 | 1,36E+07 | 1,79E+07 |
| 22,1 | | 0,7869797 | 0,8729200 | 0,78 | 4,23E+06 | 9,60E+05 |
| 27,3 | | 0,7870071 | 0,8728108 | 1,01 | 1,69E+07 | 1,69E+07 |
| 16,6 | | 0,7873458 | 0,8730188 | 0,98 | 6,06E+07 | 4,72E+07 |
| 39,4 | | 0,7884265 | 0,8739732 | 1,02 | 7,57E+07 | 7,94E+07 |
| 9 | | 0,7887927 | 0,8741990 | 0,94 | 1,70E+07 | 1,92E+07 |
| 7,8 | | 0,7897916 | 0,8749304 | 0,84 | 1,64E+08 | 7,93E+07 |
| 18,4 | | 0,7900353 | 0,8749614 | 0,90 | 1,03E+07 | 8,73E+06 |
| 8,8 | | 0,7907899 | 0,8756191 | 0,95 | 3,35E+07 | 2,96E+07 |
| 44,1 | | 0,7916313 | 0,8764024 | 0,94 | 1,50E+07 | 2,87E+07 |
| 11,1 | | 0,7917640 | 0,8764291 | 1,04 | 9,68E+06 | 1,21E+07 |
| 3,9 | | 0,7919788 | 0,8765367 | 0,95 | 2,43E+07 | 3,40E+07 |
| 51,2 | | 0,7926049 | 0,8768987 | 0,98 | 2,46E+08 | 2,83E+08 |
| 11,9 | | 0,7927378 | 0,8768625 | 1,15 | 9,49E+06 | 4,35E+06 |
| 51,2 | | 0,7929500 | 0,8769281 | 1,01 | 4,99E+08 | 4,60E+08 |
| 38,1 | | 0,7929768 | 0,8768163 | 1,03 | 1,99E+07 | 2,46E+07 |
| 20,6 | | 0,7931291 | 0,8767568 | 1,11 | 2,38E+06 | 5,73E+06 |
| 20,7 | | 0,7936769 | 0,8772203 | 1,04 | 6,79E+07 | 5,16E+07 |
| 55,6 | | 0,7937263 | 0,8771085 | 0,95 | 1,61E+07 | 1,23E+07 |
| 6,3 | | 0,7938495 | 0,8770990 | 0,88 | 4,17E+06 | 5,01E+06 |
| 16 | | 0,7939603 | 0,8770355 | 1,07 | 6,78E+06 | 7,44E+06 |
| 25,6 | | 0,7940264 | 0,8769400 | 0,98 | 3,04E+08 | 3,44E+08 |
| 4,5 | | 0,7941246 | 0,8768986 | 1,10 | 2,51E+06 | 3,85E+06 |
| 34,1 | | 0,7943309 | 0,8770478 | 0,91 | 3,98E+07 | 6,63E+07 |
| 10 | | 0,7944678 | 0,8770383 | 1,04 | 1,00E+07 | 8,09E+06 |
| 1,2 | | 0,7948627 | 0,8773688 | 0,91 | 6,57E+06 | 7,14E+06 |
| 39 | | 0,7951395 | 0,8774981 | 0,98 | 3,08E+08 | 3,37E+08 |
| 9,1 | | 0,7968683 | 0,8791224 | 1,18 | 2,85E+06 | 8,83E+06 |
| 7,1 | | 0,7969826 | 0,8791580 | 1,22 | 1,05E+07 | 7,82E+06 |
| 44,8 | | 0,7969921 | 0,8790489 | 0,97 | 2,24E+09 | 1,79E+09 |
| 67,8 | | 0,7971339 | 0,8790328 | 1,02 | 1,46E+10 | 1,86E+10 |
| 2,5 | | 0,7985094 | 0,8802682 | 1,33 | 1,80E+06 | 1,60E+07 |
| 14,4 | | 0,7991325 | 0,8806907 | 1,11 | 5,61E+06 | 1,94E+06 |
| 12,9 | | 0,8005561 | 0,8821291 | 0,90 | 9,28E+06 | 3,04E+07 |
| 20,8 | | 0,8006625 | 0,8821665 | 0,83 | 2,25E+07 | 1,25E+07 |
| 3,4 | | 0,8007034 | 0,8820525 | 0,94 | 5,51E+06 | 2,79E+06 |
| 30,6 | | 0,8007188 | 0,8819432 | 0,97 | 2,57E+08 | 2,19E+08 |

| | | | | | | |
|---|---|---|---|---|---|---|
| 70,8 | | 0,8010735 | 0,8820937 | 1,29 | 1,95E+06 | 7,93E+06 |
| 7,5 | | 0,8011243 | 0,8820313 | 1,11 | 2,01E+06 | 3,49E+06 |
| 5 | | 0,8028966 | 0,8836550 | 1,10 | 8,09E+06 | 1,38E+07 |
| 29,3 | | 0,8030757 | 0,8836365 | 0,98 | 3,65E+08 | 4,27E+08 |
| 25,7 | | 0,8034162 | 0,8838713 | 1,09 | 3,47E+07 | 4,67E+07 |
| 65,4 | | 0,8037325 | 0,8840052 | 1,02 | 5,24E+08 | 5,91E+08 |
| 4,7 | | 0,8037472 | 0,8838772 | 0,93 | 3,22E+06 | 4,91E+06 |
| 65,9 | | 0,8044768 | 0,8847410 | 1,03 | 5,79E+08 | 6,17E+08 |
| 14,1 | | 0,8045489 | 0,8847305 | 0,99 | 8,08E+07 | 7,89E+07 |
| 15,5 | | 0,8051227 | 0,8849574 | 0,97 | 1,45E+07 | 2,06E+07 |
| 32,3 | | 0,8053184 | 0,8850094 | 1,04 | 5,86E+06 | 7,68E+06 |
| 67,4 | | 0,8054138 | 0,8850214 | 0,98 | 5,50E+08 | 6,09E+08 |
| 43,6 | | 0,8066299 | 0,8860466 | 1,01 | 9,77E+07 | 8,67E+07 |
| 25 | | 0,8083559 | 0,8874848 | 1,04 | 1,42E+07 | 1,61E+07 |
| 4,3 | | 0,8094375 | 0,8886639 | 0,96 | 5,73E+06 | 4,05E+06 |
| 23,1 | | 0,8100208 | 0,8892008 | 0,71 | 5,70E+07 | 6,55E+05 |
| 62,9 | | 0,8100815 | 0,8891781 | 1,00 | 3,72E+09 | 3,88E+09 |
| 38,6 | | 0,8105334 | 0,8893798 | 0,98 | 2,18E+08 | 2,00E+08 |
| 10,6 | | 0,8106073 | 0,8893432 | 1,08 | 4,24E+06 | 8,67E+06 |
| 28,4 | | 0,8112384 | 0,8898918 | 1,05 | 7,24E+07 | 7,89E+07 |
| 6,4 | + | 0,8112497 | 0,8897632 | 0,81 | 4,92E+06 | 2,59E+06 |
| 52,9 | | 0,8115390 | 0,8898300 | 0,99 | 2,20E+08 | 2,12E+08 |
| 5,5 | + | 0,8131337 | 0,8909970 | 1,07 | 6,34E+06 | 9,51E+06 |
| 11,2 | | 0,8132906 | 0,8909312 | 0,96 | 2,00E+06 | 2,57E+06 |
| 38,2 | | 0,8142510 | 0,8916365 | 1,01 | 3,63E+07 | 4,14E+07 |
| 2,6 | | 0,8143064 | 0,8915891 | 1,19 | 1,50E+06 | 4,85E+06 |
| 14,2 | | 0,8148634 | 0,8920127 | 0,95 | 9,36E+06 | 5,23E+06 |
| 9,1 | | 0,8153748 | 0,8923888 | 0,94 | 1,93E+07 | 1,30E+07 |
| 3,2 | | 0,8159359 | 0,8927047 | 1,14 | 2,99E+06 | 1,26E+07 |
| 23,7 | | 0,8159972 | 0,8926122 | 0,99 | 1,23E+08 | 1,35E+08 |
| 5,6 | | 0,8173190 | 0,8937936 | 0,94 | 5,67E+06 | 4,70E+06 |
| 78,6 | | 0,8180127 | 0,8943557 | 0,99 | 1,01E+10 | 9,38E+09 |
| 35,4 | | 0,8184738 | 0,8947744 | 0,97 | 3,65E+07 | 2,52E+07 |
| 3,3 | | 0,8187903 | 0,8949934 | 0,92 | 1,21E+07 | 9,94E+06 |
| 49,5 | | 0,8196858 | 0,8955741 | 1,03 | 4,31E+09 | 5,95E+09 |
| 30,3 | | 0,8200333 | 0,8957871 | 0,98 | 5,33E+07 | 5,49E+07 |
| 15,5 | | 0,8214513 | 0,8971199 | 1,01 | 4,91E+07 | 5,45E+07 |
| 10,5 | | 0,8228111 | 0,8982798 | 1,04 | 1,25E+07 | 1,49E+07 |
| 59,5 | | 0,8233526 | 0,8987351 | 1,03 | 2,02E+08 | 2,40E+08 |
| 31,2 | | 0,8233734 | 0,8986234 | 0,98 | 1,88E+08 | 1,91E+08 |
| 5,7 | | 0,8239039 | 0,8990336 | 0,92 | 5,99E+06 | 7,63E+06 |
| 42 | | 0,8240248 | 0,8990297 | 1,02 | 1,97E+08 | 2,26E+08 |
| 24,5 | | 0,8242193 | 0,8990621 | 0,96 | 3,98E+07 | 2,42E+07 |
| 29,7 | | 0,8242386 | 0,8989464 | 0,99 | 1,90E+08 | 2,10E+08 |
| 5,7 | | 0,8247474 | 0,8993926 | 1,07 | 4,45E+06 | 3,56E+06 |
| 29,6 | | 0,8257000 | 0,9001354 | 0,96 | 5,89E+07 | 6,45E+07 |
| 34,6 | | 0,8274226 | 0,9013850 | 1,01 | 2,68E+08 | 2,61E+08 |
| 8,1 | | 0,8276834 | 0,9015685 | 1,08 | 7,39E+06 | 9,07E+06 |
| 8,3 | | 0,8278899 | 0,9016075 | 1,04 | 7,72E+06 | 9,09E+06 |
| 37,4 | | 0,8280188 | 0,9016222 | 1,05 | 1,04E+08 | 7,28E+07 |

| | | | | | | |
|---|---|---|---|---|---|---|
| 12,8 | | 0,8281546 | 0,9016709 | 0,97 | 7,47E+06 | 7,96E+06 |
| 21,1 | | 0,8283954 | 0,9017824 | 1,05 | 6,40E+07 | 7,23E+07 |
| 21,2 | | 0,8285623 | 0,9017781 | 0,90 | 2,91E+07 | 7,42E+06 |
| 26,8 | | 0,8286141 | 0,9016996 | 0,97 | 2,20E+07 | 2,06E+07 |
| 51,5 | | 0,8292402 | 0,9022011 | 1,02 | 3,88E+09 | 3,40E+09 |
| 16,9 | | 0,8295427 | 0,9024563 | 1,05 | 3,73E+06 | 2,67E+06 |
| 6,1 | | 0,8325856 | 0,9053968 | 0,95 | 3,56E+06 | 3,47E+06 |
| 57,7 | | 0,8328593 | 0,9054622 | 0,97 | 1,26E+08 | 1,41E+08 |
| 31,2 | | 0,8329328 | 0,9054206 | 0,96 | 5,86E+09 | 4,15E+09 |
| 55,6 | | 0,8330876 | 0,9053784 | 0,98 | 4,93E+08 | 4,08E+08 |
| 67,6 | | 0,8331241 | 0,9052850 | 1,04 | 5,91E+07 | 1,05E+08 |
| 15,7 | | 0,8333844 | 0,9053671 | 1,06 | 3,01E+06 | 3,21E+06 |
| 10,2 | | 0,8335911 | 0,9053876 | 0,94 | 2,13E+06 | 3,51E+06 |
| 9 | | 0,8336731 | 0,9053046 | 0,90 | 2,45E+06 | 1,33E+06 |
| 9,1 | | 0,8350002 | 0,9062974 | 0,87 | 1,21E+06 | 4,55E+06 |
| 27,5 | | 0,8359123 | 0,9070032 | 1,15 | 1,57E+06 | 1,34E+06 |
| 2,3 | | 0,8360929 | 0,9070906 | 0,96 | 3,28E+06 | 2,32E+06 |
| 50,6 | | 0,8363568 | 0,9071671 | 1,01 | 1,57E+08 | 1,82E+08 |
| 42 | | 0,8363941 | 0,9070685 | 0,98 | 7,71E+07 | 6,51E+07 |
| 55 | | 0,8380776 | 0,9083664 | 1,01 | 1,29E+09 | 1,42E+09 |
| 6,3 | | 0,8381713 | 0,9082853 | 1,20 | 5,38E+06 | 1,02E+07 |
| 16,8 | | 0,8394284 | 0,9092755 | 1,09 | 2,61E+06 | 2,41E+06 |
| 17,6 | | 0,8395850 | 0,9092960 | 0,91 | 3,81E+07 | 1,38E+07 |
| 9 | | 0,8396941 | 0,9092844 | 0,95 | 2,45E+07 | 2,48E+07 |
| 61,7 | | 0,8409495 | 0,9102155 | 0,96 | 2,70E+08 | 3,78E+08 |
| 21,7 | | 0,8411722 | 0,9102123 | 0,92 | 6,41E+07 | 2,71E+07 |
| 19,6 | | 0,8415580 | 0,9104908 | 0,96 | 1,09E+07 | 1,96E+07 |
| 12,4 | | 0,8421594 | 0,9107221 | 0,96 | 7,93E+06 | 1,39E+07 |
| 55,6 | | 0,8428969 | 0,9111994 | 1,03 | 3,04E+09 | 2,20E+09 |
| 8,4 | | 0,8432262 | 0,9114386 | 1,04 | 1,09E+07 | 8,05E+06 |
| 22,9 | | 0,8433223 | 0,9113883 | 1,03 | 2,68E+07 | 3,99E+07 |
| 12,7 | | 0,8437721 | 0,9116486 | 1,02 | 1,33E+08 | 1,29E+08 |
| 25,9 | | 0,8439325 | 0,9116194 | 0,98 | 1,82E+08 | 1,60E+08 |
| 54,2 | | 0,8441436 | 0,9116940 | 0,99 | 5,71E+08 | 6,32E+08 |
| 50,9 | | 0,8445983 | 0,9120642 | 1,02 | 2,84E+08 | 3,59E+08 |
| 30,1 | | 0,8451993 | 0,9125294 | 0,98 | 2,68E+08 | 2,57E+08 |
| 14,2 | | 0,8473495 | 0,9143537 | 1,04 | 7,30E+06 | 6,00E+06 |
| 6,9 | | 0,8474551 | 0,9142675 | 0,92 | 4,21E+06 | 4,84E+06 |
| 8 | | 0,8474792 | 0,9141503 | 0,95 | 6,84E+06 | 5,68E+06 |
| 30,4 | | 0,8479082 | 0,9144953 | 1,01 | 5,23E+07 | 5,58E+07 |
| 12,8 | | 0,8480319 | 0,9144898 | 0,94 | 3,20E+06 | 7,44E+06 |
| 8,5 | | 0,8483520 | 0,9147298 | 0,92 | 7,98E+06 | 2,46E+07 |
| 9,4 | | 0,8491209 | 0,9154798 | 0,94 | 2,62E+06 | 2,72E+06 |
| 38,2 | | 0,8493186 | 0,9154753 | 0,98 | 1,14E+08 | 1,32E+08 |
| 10,8 | | 0,8494454 | 0,9154358 | 0,98 | 1,21E+07 | 1,25E+07 |
| 12,8 | | 0,8495523 | 0,9154828 | 1,08 | 5,85E+06 | 4,20E+06 |
| 72,2 | | 0,8505347 | 0,9163518 | 0,99 | 3,22E+08 | 2,97E+08 |
| 21,7 | | 0,8506765 | 0,9162991 | 0,95 | 1,10E+07 | 8,42E+06 |
| 49,1 | | 0,8508060 | 0,9163179 | 0,96 | 1,41E+07 | 1,89E+07 |
| 21,3 | | 0,8510527 | 0,9163917 | 0,98 | 4,86E+07 | 5,23E+07 |

| | | | | | | |
|---|---|---|---|---|---|---|
| 17,8 | | 0,8514118 | 0,9165002 | 1,03 | 2,70E+07 | 2,46E+07 |
| 45,2 | | 0,8520370 | 0,9167711 | 0,96 | 4,34E+07 | 5,54E+07 |
| 16,3 | | 0,8522323 | 0,9168584 | 1,01 | 6,99E+07 | 7,78E+07 |
| 29,7 | | 0,8527655 | 0,9172383 | 1,02 | 1,20E+08 | 1,31E+08 |
| 24,1 | | 0,8533077 | 0,9177363 | 0,98 | 2,48E+08 | 2,31E+08 |
| 3 | | 0,8535191 | 0,9176647 | 0,92 | 7,38E+06 | 3,13E+06 |
| 25,8 | | 0,8537537 | 0,9176924 | 1,02 | 1,64E+08 | 1,78E+08 |
| 14,1 | | 0,8541928 | 0,9178869 | 0,89 | 3,59E+06 | 1,72E+07 |
| 2,8 | | 0,8559149 | 0,9192440 | 0,97 | 6,44E+06 | 4,92E+06 |
| 12,5 | | 0,8561483 | 0,9193880 | 0,97 | 1,60E+07 | 1,64E+07 |
| 40,4 | | 0,8571738 | 0,9201353 | 1,01 | 1,55E+08 | 1,46E+08 |
| 7,9 | | 0,8581646 | 0,9208744 | 1,07 | 1,14E+07 | 5,23E+06 |
| 18,3 | | 0,8581810 | 0,9207744 | 1,08 | 3,43E+06 | 7,04E+06 |
| 49,5 | | 0,8582897 | 0,9207207 | 1,01 | 4,53E+08 | 5,17E+08 |
| 58,8 | | 0,8585876 | 0,9209334 | 0,98 | 3,11E+09 | 2,33E+09 |
| 35,4 | | 0,8588464 | 0,9211483 | 0,98 | 1,00E+08 | 8,80E+07 |
| 58,9 | | 0,8590229 | 0,9211989 | 1,02 | 7,17E+08 | 6,50E+08 |
| 15,2 | | 0,8590446 | 0,9211275 | 0,95 | 6,26E+06 | 6,15E+06 |
| 32,9 | | 0,8601040 | 0,9220105 | 0,99 | 2,94E+08 | 2,42E+08 |
| 16,2 | | 0,8601117 | 0,9218968 | 0,96 | 9,15E+06 | 1,08E+07 |
| 66,9 | | 0,8601241 | 0,9217654 | 0,99 | 2,36E+08 | 2,35E+08 |
| 13 | | 0,8609148 | 0,9223619 | 1,01 | 5,04E+07 | 4,58E+07 |
| 18,3 | | 0,8613480 | 0,9225274 | 0,97 | 2,63E+07 | 4,06E+07 |
| 31,8 | | 0,8615786 | 0,9226137 | 1,01 | 1,21E+08 | 9,64E+07 |
| 32,9 | | 0,8623769 | 0,9233443 | 1,04 | 8,71E+06 | 5,64E+06 |
| 47,7 | | 0,8626331 | 0,9233461 | 1,02 | 6,40E+07 | 6,65E+07 |
| 28,1 | | 0,8640575 | 0,9245473 | 0,99 | 2,84E+07 | 2,91E+07 |
| 12,5 | | 0,8646266 | 0,9250376 | 1,04 | 1,14E+07 | 7,18E+06 |
| 55 | | 0,8647172 | 0,9250038 | 0,99 | 1,70E+09 | 1,88E+09 |
| 2 | | 0,8655596 | 0,9255827 | 1,32 | 2,54E+07 | 1,85E+05 |
| 18,6 | | 0,8659358 | 0,9257823 | 1,02 | 2,59E+07 | 2,95E+07 |
| 1,3 | | 0,8667592 | 0,9264445 | 0,94 | 2,97E+06 | 5,31E+06 |
| 23,6 | | 0,8675557 | 0,9269267 | 0,98 | 1,38E+08 | 1,47E+08 |
| 25,1 | | 0,8682588 | 0,9275494 | 1,02 | 5,21E+07 | 7,14E+07 |
| 65,3 | | 0,8685984 | 0,9279306 | 0,99 | 4,18E+08 | 4,87E+08 |
| 9,2 | | 0,8689738 | 0,9281530 | 0,98 | 7,92E+06 | 1,00E+07 |
| 5,2 | | 0,8694194 | 0,9284152 | 1,04 | 2,01E+07 | 2,37E+07 |
| 5,1 | | 0,8695038 | 0,9284077 | 0,94 | 8,72E+06 | 2,54E+06 |
| 83,7 | | 0,8698407 | 0,9284765 | 0,94 | 1,24E+08 | 2,18E+08 |
| 18,8 | + | 0,8701482 | 0,9285492 | 1,14 | 3,58E+08 | 1,99E+08 |
| 55,4 | | 0,8702217 | 0,9284928 | 0,98 | 1,18E+08 | 1,41E+08 |
| 23,1 | | 0,8702370 | 0,9283785 | 0,93 | 6,77E+06 | 3,34E+06 |
| 12,7 | | 0,8704047 | 0,9283597 | 0,94 | 3,68E+06 | 2,05E+06 |
| 50,4 | | 0,8707121 | 0,9284767 | 1,02 | 9,77E+07 | 1,12E+08 |
| 31,7 | | 0,8711357 | 0,9286715 | 1,06 | 5,11E+07 | 6,59E+07 |
| 15,2 | | 0,8718023 | 0,9290281 | 1,02 | 6,01E+07 | 5,11E+07 |
| 11,9 | | 0,8721538 | 0,9291773 | 0,96 | 1,34E+07 | 2,66E+07 |
| 15,6 | | 0,8724493 | 0,9293055 | 1,03 | 2,27E+07 | 1,53E+07 |
| 17,2 | | 0,8740480 | 0,9303262 | 0,97 | 7,47E+07 | 4,31E+07 |
| 27,8 | | 0,8740499 | 0,9301942 | 0,98 | 1,49E+10 | 1,09E+10 |

| | | | | | | |
|---|---|---|---|---|---|---|
| 10,6 | | 0,8745416 | 0,9303789 | 1,02 | 4,13E+07 | 4,91E+07 |
| 17,6 | | 0,8751165 | 0,9308263 | 1,03 | 8,83E+07 | 7,60E+07 |
| 33,7 | | 0,8751676 | 0,9307300 | 1,02 | 4,85E+08 | 3,33E+08 |
| 39,6 | | 0,8753618 | 0,9308000 | 0,99 | 1,60E+09 | 1,96E+09 |
| 23,2 | | 0,8777121 | 0,9326490 | 0,99 | 8,18E+07 | 7,67E+07 |
| 35,7 | | 0,8783101 | 0,9329002 | 0,99 | 1,15E+08 | 1,18E+08 |
| 5,1 | | 0,8783907 | 0,9328348 | 0,94 | 6,08E+06 | 6,57E+06 |
| 4,6 | | 0,8788603 | 0,9330944 | 1,03 | 4,56E+06 | 6,44E+06 |
| 18 | | 0,8791742 | 0,9332179 | 0,95 | 1,93E+07 | 1,29E+07 |
| 25,7 | | 0,8794721 | 0,9333753 | 1,02 | 1,43E+07 | 2,21E+07 |
| 43,2 | | 0,8794986 | 0,9332719 | 1,02 | 5,97E+08 | 6,98E+08 |
| 31,9 | | 0,8795514 | 0,9331884 | 0,99 | 4,47E+07 | 4,43E+07 |
| 6 | | 0,8796447 | 0,9331645 | 0,97 | 9,47E+06 | 7,77E+06 |
| 8,2 | | 0,8808258 | 0,9341201 | 0,96 | 4,53E+06 | 3,06E+06 |
| 17,1 | | 0,8817311 | 0,9349577 | 0,97 | 4,66E+07 | 5,24E+07 |
| 6,4 | | 0,8820353 | 0,9352518 | 0,98 | 3,90E+06 | 3,94E+06 |
| 39,9 | | 0,8824030 | 0,9354818 | 0,98 | 3,10E+08 | 3,19E+08 |
| 9,2 | | 0,8833277 | 0,9360883 | 1,06 | 7,68E+06 | 3,54E+06 |
| 8,4 | | 0,8837466 | 0,9363227 | 0,91 | 7,83E+06 | 2,97E+06 |
| 1,5 | | 0,8843453 | 0,9364704 | 1,03 | 4,38E+06 | 2,52E+06 |
| 42,7 | | 0,8844601 | 0,9364341 | 0,99 | 1,35E+08 | 1,31E+08 |
| 41 | | 0,8855023 | 0,9371720 | 1,01 | 6,08E+08 | 7,67E+08 |
| 24,5 | | 0,8856328 | 0,9371838 | 1,02 | 3,81E+07 | 3,86E+07 |
| 31,7 | | 0,8856352 | 0,9370687 | 0,99 | 1,82E+08 | 1,61E+08 |
| 37,7 | | 0,8859283 | 0,9371003 | 1,02 | 4,22E+07 | 5,21E+07 |
| 12,9 | | 0,8859779 | 0,9370419 | 1,14 | 1,69E+07 | 1,51E+06 |
| 63,2 | | 0,8864582 | 0,9373448 | 1,00 | 2,63E+08 | 2,82E+08 |
| 16,1 | | 0,8866263 | 0,9374436 | 1,02 | 4,66E+07 | 4,19E+07 |
| 38,3 | | 0,8872999 | 0,9378736 | 0,97 | 2,07E+09 | 1,19E+09 |
| 33,3 | | 0,8878559 | 0,9382159 | 0,99 | 1,32E+09 | 1,27E+09 |
| 25,5 | | 0,8884263 | 0,9386768 | 0,99 | 5,13E+07 | 5,47E+07 |
| 29,1 | | 0,8886523 | 0,9388251 | 1,02 | 3,35E+07 | 3,86E+07 |
| 16,9 | | 0,8886644 | 0,9387106 | 1,02 | 2,46E+07 | 2,64E+07 |
| 22,9 | | 0,8887303 | 0,9385780 | 0,97 | 1,45E+07 | 1,01E+07 |
| 35,4 | | 0,8890866 | 0,9388561 | 1,03 | 9,75E+07 | 8,95E+07 |
| 28,4 | | 0,8901201 | 0,9396973 | 1,02 | 2,74E+07 | 2,32E+07 |
| 15,7 | | 0,8904018 | 0,9397735 | 1,01 | 3,03E+07 | 2,61E+07 |
| 1,9 | | 0,8909603 | 0,9400678 | 0,90 | 1,62E+06 | 2,03E+06 |
| 5,4 | | 0,8914699 | 0,9403743 | 0,97 | 5,23E+06 | 4,70E+06 |
| 54,5 | | 0,8916634 | 0,9404273 | 0,98 | 1,33E+09 | 1,13E+09 |
| 5,2 | | 0,8927181 | 0,9411548 | 1,07 | 1,87E+06 | 4,84E+06 |
| 4,3 | | 0,8931449 | 0,9413256 | 0,97 | 4,22E+06 | 3,73E+06 |
| 20,8 | | 0,8935569 | 0,9416081 | 1,01 | 7,88E+07 | 9,01E+07 |
| 23,1 | | 0,8944638 | 0,9424192 | 1,01 | 5,78E+07 | 4,72E+07 |
| 39,2 | | 0,8946051 | 0,9424555 | 0,99 | 1,96E+08 | 1,67E+08 |
| 7,2 | | 0,8947305 | 0,9424614 | 0,98 | 4,40E+06 | 6,38E+06 |
| 14,5 | | 0,8949619 | 0,9424836 | 0,96 | 8,96E+06 | 8,75E+06 |
| 31 | | 0,8950161 | 0,9423868 | 0,98 | 1,34E+08 | 9,74E+07 |
| 2,6 | | 0,8951237 | 0,9423414 | 0,98 | 3,54E+06 | 5,81E+06 |
| 29,3 | | 0,8955971 | 0,9426669 | 1,02 | 1,09E+08 | 1,42E+08 |

| | | | | | | |
|---|---|---|---|---|---|---|
| 48,6 | | 0,8956320 | 0,9425476 | 1,01 | 2,57E+08 | 2,16E+08 |
| 16 | | 0,8958700 | 0,9426808 | 1,01 | 8,52E+07 | 8,31E+07 |
| 30,9 | | 0,8964045 | 0,9429170 | 0,96 | 4,56E+07 | 4,02E+07 |
| 14,8 | | 0,8967788 | 0,9430180 | 1,02 | 2,37E+07 | 2,77E+07 |
| 2,8 | | 0,8981770 | 0,9441752 | 0,97 | 2,90E+06 | 2,76E+06 |
| 17,2 | | 0,8985067 | 0,9442354 | 1,01 | 7,23E+07 | 8,48E+07 |
| 7,4 | | 0,8985292 | 0,9441335 | 0,94 | 1,05E+07 | 2,88E+06 |
| 44,1 | | 0,8985642 | 0,9440135 | 1,03 | 2,80E+07 | 5,05E+07 |
| 85,3 | | 0,8987532 | 0,9439916 | 1,01 | 8,10E+09 | 7,26E+09 |
| 5 | | 0,8988130 | 0,9438936 | 1,10 | 7,74E+06 | 9,51E+06 |
| 49,2 | | 0,8990776 | 0,9439398 | 0,99 | 1,62E+09 | 1,60E+09 |
| 7,4 | | 0,8991870 | 0,9438683 | 1,05 | 2,96E+06 | 1,88E+06 |
| 20 | | 0,8995790 | 0,9440641 | 1,02 | 1,37E+07 | 1,47E+07 |
| 4,5 | | 0,9006437 | 0,9448467 | 0,98 | 7,70E+06 | 1,16E+07 |
| 10,2 | | 0,9006870 | 0,9447791 | 0,98 | 2,03E+07 | 1,17E+07 |
| 25,5 | | 0,9007485 | 0,9446946 | 1,01 | 3,10E+08 | 2,90E+08 |
| 40,7 | | 0,9013716 | 0,9450437 | 1,02 | 5,17E+07 | 6,06E+07 |
| 13,8 | | 0,9014368 | 0,9449958 | 0,98 | 1,14E+07 | 1,55E+07 |
| 21,1 | | 0,9024787 | 0,9456202 | 1,01 | 5,07E+07 | 6,06E+07 |
| 39,7 | | 0,9029728 | 0,9459657 | 0,99 | 3,71E+08 | 3,39E+08 |
| 15,9 | | 0,9030199 | 0,9458328 | 1,04 | 1,17E+07 | 2,14E+07 |
| 22,1 | | 0,9030344 | 0,9456999 | 0,99 | 9,07E+07 | 9,76E+07 |
| 16 | | 0,9031044 | 0,9456154 | 1,01 | 4,31E+08 | 5,26E+08 |
| 28 | | 0,9033204 | 0,9456393 | 0,99 | 7,20E+07 | 8,57E+07 |
| 7,7 | | 0,9036498 | 0,9457582 | 1,21 | 4,02E+07 | 1,55E+06 |
| 19,3 | | 0,9040155 | 0,9458495 | 0,99 | 6,88E+07 | 6,77E+07 |
| 5,2 | | 0,9042996 | 0,9459267 | 0,91 | 4,88E+06 | 6,35E+06 |
| 58,7 | | 0,9045487 | 0,9460331 | 0,99 | 8,58E+08 | 8,77E+08 |
| 43,4 | | 0,9056669 | 0,9467217 | 1,02 | 5,93E+07 | 6,04E+07 |
| 34,9 | | 0,9057889 | 0,9466506 | 0,99 | 1,58E+08 | 1,98E+08 |
| 44,5 | | 0,9066810 | 0,9473018 | 1,00 | 1,95E+08 | 2,23E+08 |
| 22,1 | | 0,9072837 | 0,9476880 | 0,98 | 4,07E+07 | 2,73E+07 |
| 11,1 | | 0,9077814 | 0,9481077 | 0,98 | 3,26E+07 | 4,33E+07 |
| 29,9 | | 0,9096117 | 0,9495192 | 0,98 | 2,13E+08 | 1,59E+08 |
| 4,8 | | 0,9099175 | 0,9495151 | 0,97 | 5,82E+06 | 5,80E+06 |
| 17,7 | | 0,9101208 | 0,9495827 | 1,03 | 1,91E+07 | 3,63E+07 |
| 18,3 | | 0,9110547 | 0,9503524 | 0,98 | 1,33E+07 | 2,00E+07 |
| 52 | | 0,9121747 | 0,9512033 | 1,01 | 1,97E+08 | 2,53E+08 |
| 27,2 | | 0,9135735 | 0,9521929 | 0,99 | 1,32E+08 | 1,82E+08 |
| 32,5 | | 0,9143524 | 0,9526732 | 1,01 | 5,72E+08 | 6,25E+08 |
| 25,5 | | 0,9146043 | 0,9527779 | 0,98 | 7,01E+07 | 1,16E+08 |
| 9,2 | | 0,9151332 | 0,9531443 | 1,11 | 1,99E+07 | 5,69E+07 |
| 4,1 | | 0,9153939 | 0,9531750 | 0,94 | 7,58E+06 | 8,33E+06 |
| 45,7 | | 0,9165089 | 0,9541188 | 1,01 | 1,90E+08 | 2,26E+08 |
| 5,1 | | 0,9165396 | 0,9540031 | 1,01 | 5,75E+06 | 4,11E+06 |
| 16,8 | | 0,9169012 | 0,9541658 | 1,00 | 1,77E+08 | 1,68E+08 |
| 22,1 | | 0,9176450 | 0,9547742 | 1,00 | 5,59E+07 | 6,16E+07 |
| 48,8 | | 0,9183156 | 0,9552923 | 0,98 | 2,20E+08 | 2,71E+08 |
| 11,7 | | 0,9184341 | 0,9552924 | 1,02 | 7,66E+07 | 9,42E+07 |
| 5,5 | | 0,9192231 | 0,9559121 | 1,04 | 5,98E+06 | 6,99E+06 |

| | | | | | | |
|---|---|---|---|---|---|---|
| 21,3 | | 0,9198998 | 0,9562211 | 1,16 | 5,66E+08 | 7,32E+06 |
| 13,5 | | 0,9205869 | 0,9566491 | 0,98 | 3,87E+06 | 7,85E+06 |
| 23,1 | | 0,9205997 | 0,9565461 | 0,99 | 1,21E+08 | 1,14E+08 |
| 13,2 | | 0,9210400 | 0,9568263 | 1,03 | 7,40E+06 | 1,23E+07 |
| 13,9 | | 0,9213424 | 0,9570376 | 1,04 | 5,33E+06 | 1,74E+06 |
| 27,7 | | 0,9219614 | 0,9574827 | 1,01 | 9,70E+07 | 1,01E+08 |
| 46 | | 0,9220461 | 0,9574137 | 1,02 | 1,06E+08 | 1,22E+08 |
| 14 | | 0,9229345 | 0,9581375 | 1,04 | 2,52E+06 | 2,52E+06 |
| 14,3 | | 0,9230968 | 0,9580841 | 0,94 | 8,17E+06 | 1,55E+07 |
| 24,3 | | 0,9231495 | 0,9579905 | 1,02 | 8,99E+06 | 1,48E+07 |
| 4,9 | | 0,9236204 | 0,9581154 | 0,99 | 3,79E+06 | 3,44E+06 |
| 17,7 | | 0,9237173 | 0,9580447 | 0,99 | 4,04E+07 | 4,10E+07 |
| 24,6 | | 0,9242714 | 0,9584227 | 0,99 | 3,43E+08 | 4,75E+08 |
| 51,9 | | 0,9243080 | 0,9583061 | 0,98 | 1,92E+07 | 3,20E+07 |
| 2,9 | | 0,9246457 | 0,9585348 | 1,04 | 4,98E+06 | 2,75E+06 |
| 0,8 | | 0,9270581 | 0,9605339 | 1,02 | 1,56E+06 | 1,62E+06 |
| 6,3 | | 0,9271353 | 0,9604741 | 1,04 | 2,68E+06 | 7,59E+06 |
| 37,2 | | 0,9272955 | 0,9604729 | 0,99 | 5,00E+07 | 5,75E+07 |
| 53,1 | | 0,9282139 | 0,9610300 | 0,99 | 4,26E+08 | 4,09E+08 |
| 20,1 | | 0,9282394 | 0,9608959 | 0,99 | 1,83E+07 | 2,48E+07 |
| 68,1 | | 0,9282979 | 0,9607769 | 0,99 | 5,75E+07 | 7,07E+07 |
| 42,9 | | 0,9288346 | 0,9610848 | 1,00 | 3,51E+08 | 3,86E+08 |
| 16,6 | + | 0,9293430 | 0,9613804 | 1,06 | 3,20E+07 | 2,34E+07 |
| 36 | | 0,9295466 | 0,9614550 | 1,01 | 8,44E+07 | 1,12E+08 |
| 49,1 | | 0,9299949 | 0,9617805 | 0,99 | 4,41E+08 | 4,02E+08 |
| 2 | | 0,9304801 | 0,9620502 | 0,94 | 2,89E+06 | 3,42E+06 |
| 29,3 | | 0,9315898 | 0,9628730 | 0,99 | 1,22E+08 | 1,66E+08 |
| 8,5 | | 0,9320164 | 0,9631291 | 1,08 | 2,13E+07 | 2,41E+07 |
| 34,4 | | 0,9320800 | 0,9630584 | 1,00 | 2,32E+08 | 2,35E+08 |
| 11,7 | | 0,9322434 | 0,9631605 | 1,01 | 5,29E+06 | 5,71E+06 |
| 40,5 | | 0,9322495 | 0,9630263 | 0,98 | 1,59E+07 | 3,04E+07 |
| 35,8 | | 0,9322625 | 0,9629401 | 0,99 | 4,58E+07 | 4,06E+07 |
| 32 | | 0,9328504 | 0,9633698 | 1,01 | 1,84E+08 | 2,48E+08 |
| 55,7 | | 0,9330833 | 0,9633833 | 1,00 | 2,82E+09 | 2,88E+09 |
| 63,6 | | 0,9332104 | 0,9633583 | 1,01 | 4,00E+09 | 3,91E+09 |
| 19,6 | | 0,9340180 | 0,9638340 | 0,99 | 2,17E+07 | 1,63E+07 |
| 7,9 | | 0,9340340 | 0,9636998 | 1,00 | 3,98E+06 | 4,36E+06 |
| 7,4 | | 0,9344325 | 0,9639660 | 0,98 | 5,31E+06 | 7,32E+06 |
| 30,8 | | 0,9347972 | 0,9641253 | 1,01 | 4,47E+08 | 5,76E+08 |
| 28,5 | | 0,9350322 | 0,9642037 | 0,99 | 1,69E+07 | 2,77E+07 |
| 7,6 | | 0,9352752 | 0,9643306 | 0,99 | 2,12E+07 | 2,13E+07 |
| 3,2 | | 0,9355484 | 0,9643868 | 0,97 | 3,20E+06 | 5,57E+06 |
| 16,8 | | 0,9359822 | 0,9646638 | 1,01 | 5,08E+07 | 5,78E+07 |
| 48,4 | | 0,9371896 | 0,9655855 | 0,99 | 2,38E+08 | 3,10E+08 |
| 9,8 | | 0,9372359 | 0,9655035 | 0,99 | 1,52E+07 | 1,81E+07 |
| 40,8 | | 0,9379515 | 0,9659894 | 1,00 | 1,81E+08 | 1,92E+08 |
| 2,3 | | 0,9388852 | 0,9667532 | 0,94 | 3,68E+06 | 1,46E+06 |
| 37,1 | | 0,9401457 | 0,9675891 | 1,00 | 2,66E+08 | 3,17E+08 |
| 20,1 | | 0,9410353 | 0,9681462 | 0,99 | 4,32E+07 | 5,59E+07 |
| 51,3 | | 0,9425003 | 0,9692262 | 0,99 | 1,55E+08 | 1,76E+08 |

| | | | | | | |
|---|---|---|---|---|---|---|
| 44,4 | | 0,9425990 | 0,9691583 | 1,00 | 5,28E+08 | 5,23E+08 |
| 10 | | 0,9426083 | 0,9690236 | 1,04 | 7,16E+06 | 2,09E+06 |
| 24,4 | | 0,9427790 | 0,9690190 | 0,99 | 1,34E+08 | 1,18E+08 |
| 35,1 | | 0,9429715 | 0,9689739 | 1,00 | 1,44E+08 | 1,36E+08 |
| 24,9 | | 0,9430132 | 0,9688393 | 0,98 | 1,99E+08 | 9,63E+07 |
| 22,7 | | 0,9430698 | 0,9687398 | 1,01 | 4,39E+07 | 3,93E+07 |
| 52,9 | | 0,9432163 | 0,9687602 | 0,99 | 9,58E+07 | 9,57E+07 |
| 26,8 | | 0,9432246 | 0,9686258 | 1,01 | 4,73E+07 | 6,34E+07 |
| 27,7 | | 0,9433361 | 0,9685785 | 0,98 | 1,10E+07 | 1,46E+07 |
| 27,8 | | 0,9442321 | 0,9691551 | 0,98 | 9,83E+06 | 2,05E+07 |
| 12,5 | | 0,9460744 | 0,9706286 | 1,02 | 1,06E+08 | 9,74E+07 |
| 14,2 | | 0,9464855 | 0,9708768 | 0,96 | 1,01E+07 | 9,09E+06 |
| 61,4 | | 0,9466797 | 0,9708636 | 1,00 | 2,59E+09 | 2,44E+09 |
| 29,6 | | 0,9469521 | 0,9709908 | 1,01 | 9,94E+07 | 1,16E+08 |
| 7,3 | | 0,9469761 | 0,9708695 | 0,97 | 3,65E+06 | 2,96E+06 |
| 21,6 | | 0,9472273 | 0,9709817 | 1,00 | 2,02E+08 | 2,33E+08 |
| 4,6 | | 0,9475354 | 0,9711565 | 0,96 | 1,08E+07 | 4,54E+06 |
| 24,1 | | 0,9477677 | 0,9711993 | 1,01 | 8,82E+07 | 1,08E+08 |
| 10,8 | | 0,9479331 | 0,9711789 | 1,00 | 1,13E+07 | 9,81E+06 |
| 18,5 | | 0,9485278 | 0,9715820 | 1,01 | 3,14E+07 | 3,92E+07 |
| 42,9 | | 0,9489116 | 0,9717872 | 1,01 | 2,94E+07 | 2,59E+07 |
| 1,5 | | 0,9498107 | 0,9721784 | 1,02 | 2,68E+06 | 2,00E+06 |
| 58,4 | | 0,9498795 | 0,9720610 | 1,00 | 3,01E+08 | 2,92E+08 |
| 31,6 | | 0,9499382 | 0,9719679 | 1,01 | 1,41E+08 | 1,20E+08 |
| 17,2 | | 0,9513091 | 0,9727246 | 1,00 | 8,37E+07 | 9,05E+07 |
| 5,3 | | 0,9515672 | 0,9727499 | 0,99 | 7,45E+06 | 6,23E+06 |
| 29,3 | | 0,9519046 | 0,9730129 | 1,01 | 1,49E+08 | 1,34E+08 |
| 16,1 | | 0,9520633 | 0,9729734 | 1,02 | 2,40E+06 | 2,80E+06 |
| 48,6 | | 0,9532157 | 0,9738298 | 0,99 | 1,66E+09 | 2,11E+09 |
| 28,8 | | 0,9536532 | 0,9739900 | 1,00 | 1,79E+08 | 1,74E+08 |
| 61,5 | | 0,9539182 | 0,9740872 | 0,99 | 2,14E+08 | 3,00E+08 |
| 4,9 | | 0,9543275 | 0,9743055 | 1,03 | 1,01E+07 | 3,27E+06 |
| 49,7 | | 0,9544352 | 0,9742327 | 0,99 | 9,16E+08 | 9,95E+08 |
| 4,2 | | 0,9544409 | 0,9740979 | 1,02 | 1,18E+07 | 8,95E+06 |
| 55,1 | | 0,9548711 | 0,9743421 | 1,00 | 4,67E+08 | 4,22E+08 |
| 55,2 | | 0,9549040 | 0,9742074 | 1,01 | 7,75E+09 | 6,74E+09 |
| 14,2 | | 0,9552850 | 0,9742845 | 0,99 | 1,19E+07 | 1,20E+07 |
| 17,7 | | 0,9554364 | 0,9742068 | 0,99 | 1,03E+07 | 1,25E+07 |
| 7,1 | | 0,9557546 | 0,9743021 | 0,98 | 5,50E+06 | 6,80E+06 |
| 27,1 | | 0,9563214 | 0,9746943 | 1,01 | 1,13E+07 | 1,34E+07 |
| 18,5 | | 0,9563534 | 0,9745745 | 1,01 | 9,10E+07 | 7,70E+07 |
| 42,4 | | 0,9568640 | 0,9749831 | 1,01 | 9,52E+08 | 1,10E+09 |
| 47,7 | | 0,9583796 | 0,9758431 | 1,00 | 1,32E+08 | 1,57E+08 |
| 27,5 | | 0,9584137 | 0,9757602 | 1,01 | 5,83E+07 | 4,51E+07 |
| 25,7 | | 0,9584690 | 0,9757067 | 0,99 | 7,10E+06 | 7,72E+06 |
| 47,1 | | 0,9591859 | 0,9762010 | 1,00 | 1,72E+08 | 1,51E+08 |
| 8 | | 0,9600728 | 0,9767351 | 1,03 | 7,08E+06 | 1,00E+07 |
| 65,6 | | 0,9612282 | 0,9773379 | 1,00 | 4,34E+09 | 3,76E+09 |
| 2,2 | | 0,9627830 | 0,9786286 | 0,99 | 4,44E+06 | 4,97E+06 |
| 49,4 | | 0,9628270 | 0,9785404 | 1,01 | 1,02E+09 | 1,41E+09 |

| | | | | | | |
|---|---|---|---|---|---|---|
| 20,4 | | 0,9637667 | 0,9791080 | 1,00 | 3,18E+07 | 3,44E+07 |
| 27,1 | | 0,9642399 | 0,9793339 | 0,99 | 7,85E+08 | 1,10E+09 |
| 11,9 | | 0,9643693 | 0,9792573 | 1,01 | 6,15E+06 | 8,79E+06 |
| 18,4 | | 0,9649527 | 0,9795790 | 0,99 | 2,03E+08 | 1,08E+08 |
| 24,9 | | 0,9657012 | 0,9799239 | 1,02 | 1,88E+08 | 8,19E+07 |
| 26,1 | | 0,9667509 | 0,9806757 | 1,00 | 8,64E+07 | 8,79E+07 |
| 12,6 | | 0,9672183 | 0,9809017 | 1,00 | 1,99E+07 | 1,84E+07 |
| 23,3 | | 0,9674390 | 0,9810245 | 1,00 | 3,97E+07 | 3,92E+07 |
| 45,5 | | 0,9676595 | 0,9811578 | 1,00 | 9,30E+08 | 8,51E+08 |
| 8,5 | | 0,9698273 | 0,9828137 | 1,01 | 6,28E+07 | 4,05E+07 |
| 10,5 | | 0,9708135 | 0,9834086 | 1,04 | 1,76E+06 | 3,50E+07 |
| 16,2 | | 0,9711675 | 0,9834505 | 1,00 | 5,18E+07 | 5,31E+07 |
| 29,5 | | 0,9717979 | 0,9836473 | 1,00 | 4,40E+07 | 4,89E+07 |
| 17,8 | | 0,9718819 | 0,9835763 | 1,01 | 6,16E+06 | 3,81E+06 |
| 53,6 | | 0,9724771 | 0,9839267 | 1,00 | 2,56E+08 | 2,39E+08 |
| 8,7 | | 0,9730784 | 0,9843624 | 1,03 | 9,96E+05 | 8,63E+06 |
| 3,4 | | 0,9731280 | 0,9842588 | 0,99 | 1,23E+07 | 6,31E+06 |
| 38,8 | | 0,9735363 | 0,9844163 | 1,00 | 6,93E+08 | 7,98E+08 |
| 16,8 | | 0,9739592 | 0,9845500 | 1,00 | 2,71E+07 | 2,76E+07 |
| 1,1 | | 0,9739880 | 0,9844410 | 1,03 | 1,13E+06 | 2,25E+06 |
| 51,9 | | 0,9743677 | 0,9845912 | 1,01 | 1,23E+08 | 1,06E+08 |
| 10,2 | | 0,9746523 | 0,9845916 | 1,01 | 2,05E+07 | 1,18E+07 |
| 9,6 | | 0,9748323 | 0,9846433 | 0,99 | 1,00E+07 | 1,14E+07 |
| 28,3 | | 0,9755725 | 0,9851120 | 1,01 | 2,11E+07 | 2,33E+07 |
| 6,7 | | 0,9757760 | 0,9852229 | 1,01 | 6,37E+06 | 7,40E+06 |
| 60,9 | | 0,9762487 | 0,9855149 | 1,00 | 4,67E+08 | 4,92E+08 |
| 3,5 | | 0,9764067 | 0,9854871 | 1,01 | 1,06E+07 | 1,28E+07 |
| 5,8 | + | 0,9768152 | 0,9856673 | 0,98 | 4,64E+07 | 3,33E+07 |
| 12,7 | | 0,9772472 | 0,9859200 | 1,00 | 1,34E+07 | 1,67E+07 |
| 60,4 | | 0,9776200 | 0,9860836 | 1,00 | 2,02E+09 | 1,69E+09 |
| 20,1 | | 0,9777802 | 0,9860998 | 1,00 | 3,08E+07 | 2,38E+07 |
| 36,8 | | 0,9783562 | 0,9863237 | 1,00 | 3,74E+07 | 4,79E+07 |
| 48,7 | | 0,9793885 | 0,9870609 | 1,00 | 7,84E+08 | 7,30E+08 |
| 21,3 | | 0,9809365 | 0,9883176 | 1,00 | 8,23E+07 | 6,36E+07 |
| 11,7 | | 0,9810878 | 0,9883066 | 0,99 | 1,02E+07 | 7,39E+06 |
| 31,9 | | 0,9811347 | 0,9882010 | 1,00 | 9,58E+07 | 1,16E+08 |
| 23,6 | | 0,9814976 | 0,9883724 | 1,00 | 1,67E+07 | 2,20E+07 |
| 39,4 | | 0,9825182 | 0,9891301 | 1,00 | 4,13E+09 | 3,87E+09 |
| 30,9 | | 0,9828104 | 0,9892541 | 1,00 | 7,56E+07 | 5,55E+07 |
| 29,4 | | 0,9834889 | 0,9896470 | 1,00 | 1,50E+07 | 1,70E+07 |
| 50,5 | | 0,9836843 | 0,9897439 | 1,00 | 1,68E+08 | 1,51E+08 |
| 43,5 | | 0,9846911 | 0,9906120 | 1,00 | 2,79E+08 | 3,25E+08 |
| 24 | | 0,9848729 | 0,9906006 | 1,00 | 2,09E+07 | 2,21E+07 |
| 13,1 | | 0,9854033 | 0,9910283 | 1,01 | 2,46E+06 | 1,13E+07 |
| 11,9 | | 0,9861667 | 0,9915392 | 0,99 | 4,12E+06 | 1,32E+06 |
| 31,4 | | 0,9863406 | 0,9915930 | 1,00 | 2,11E+08 | 1,90E+08 |
| 50,2 | | 0,9863430 | 0,9914570 | 1,00 | 7,88E+08 | 1,11E+09 |
| 35 | | 0,9882811 | 0,9928336 | 1,00 | 3,07E+09 | 2,75E+09 |
| 7,9 | | 0,9887236 | 0,9930040 | 1,00 | 4,28E+06 | 3,05E+06 |
| 30,1 | | 0,9910892 | 0,9947385 | 1,00 | 8,14E+07 | 9,03E+07 |

| | | | | | | |
|---|---|---|---|---|---|---|
| 10,7 | | 0,9911722 | 0,9946975 | 1,01 | 2,18E+07 | 4,58E+06 |
| 54,4 | | 0,9923289 | 0,9956361 | 1,00 | 4,28E+08 | 4,33E+08 |
| 26,4 | | 0,9929323 | 0,9959912 | 1,00 | 4,22E+07 | 4,61E+07 |
| 12,6 | | 0,9942124 | 0,9968296 | 1,00 | 2,09E+06 | 3,72E+06 |
| 35,8 | | 0,9944759 | 0,9970141 | 1,00 | 1,34E+08 | 1,34E+08 |
| 4,5 | | 0,9947229 | 0,9970819 | 1,00 | 2,57E+07 | 6,70E+06 |
| 33,9 | | 0,9953579 | 0,9975555 | 1,00 | 6,88E+07 | 6,20E+07 |
| 11,8 | | 0,9959397 | 0,9977864 | 1,00 | 1,65E+06 | 4,63E+06 |
| 28,6 | | 0,9966225 | 0,9981689 | 1,00 | 3,94E+07 | 3,09E+07 |
| 30 | | 0,9976538 | 0,9987331 | 1,00 | 6,32E+07 | 5,05E+07 |
| 19,1 | | 0,9985173 | 0,9992095 | 1,00 | 1,91E+06 | 2,25E+06 |
| 22,6 | | 0,9990260 | 0,9994499 | 1,00 | 1,48E+07 | 2,30E+06 |
| 49 | | 0,9993075 | 0,9995638 | 1,00 | 2,95E+08 | 3,19E+07 |
| 21,1 | | 0,9998000 | 0,9998331 | 1,00 | 5,83E+07 | 6,75E+07 |

| | OE | | | | | | |
|---|---|---|---|---|---|---|---|
| LFQ intensity C3 | LFQ intensity OE1 | LFQ intensity OE2 | LFQ intensity OE3 | Mol. weight [kDa] | Q-value | Score | MS/MS count |
| 1,32E+09 | 1,75E+09 | 1,74E+09 | 1,73E+09 | 88,79 | 0 | 323,31 | 332 |
| 1,32E+08 | 5,62E+06 | 5,95E+06 | 6,02E+06 | 47,117 | 0 | 101,18 | 31 |
| 1,20E+08 | 1,50E+07 | 1,49E+07 | 1,56E+07 | 43,567 | 0 | 53,291 | 49 |
| 1,40E+09 | 7,69E+06 | 9,40E+06 | 8,27E+06 | 128,95 | 0 | 323,31 | 159 |
| 4,69E+09 | 1,77E+09 | 1,83E+09 | 1,83E+09 | 75,884 | 0 | 323,31 | 492 |
| 7,98E+09 | 1,91E+09 | 1,99E+09 | 2,03E+09 | 24,87 | 0 | 323,31 | 351 |
| 8,11E+08 | 2,08E+08 | 1,96E+08 | 1,94E+08 | 131,28 | 0 | 323,31 | 176 |
| 1,88E+09 | 1,07E+09 | 1,06E+09 | 1,05E+09 | 72,585 | 0 | 323,31 | 265 |
| 4,22E+08 | 2,01E+08 | 2,02E+08 | 2,00E+08 | 175,01 | 0 | 323,31 | 182 |
| 3,59E+07 | 1,02E+08 | 9,79E+07 | 9,97E+07 | 42,443 | 0 | 29,914 | 38 |
| 8,41E+07 | 2,30E+08 | 2,34E+08 | 2,44E+08 | 33,466 | 0 | 46,608 | 51 |
| 1,38E+09 | 3,44E+08 | 3,60E+08 | 3,74E+08 | 16,442 | 0 | 103,6 | 54 |
| 7,88E+07 | 2,93E+07 | 2,89E+07 | 2,87E+07 | 115,38 | 0 | 37,562 | 58 |
| 7,45E+07 | 1,44E+09 | 1,40E+09 | 1,49E+09 | 25,97 | 0 | 323,31 | 122 |
| 5,17E+08 | 1,14E+08 | 1,07E+08 | 1,05E+08 | 235,31 | 0 | 323,31 | 199 |
| 3,37E+08 | 9,46E+08 | 9,31E+08 | 9,46E+08 | 184,67 | 0 | 323,31 | 256 |
| 2,73E+08 | 9,27E+07 | 9,10E+07 | 9,42E+07 | 64,254 | 0 | 68,498 | 95 |
| 1,35E+08 | 4,41E+07 | 4,38E+07 | 4,64E+07 | 14,379 | 0 | 59,99 | 42 |
| 4,87E+08 | 2,14E+09 | 2,17E+09 | 2,07E+09 | 61,174 | 0 | 323,31 | 251 |
| 1,67E+09 | 3,13E+06 | 4,23E+06 | 2,85E+06 | 70,148 | 0 | 323,31 | 180 |
| 1,24E+09 | 9,51E+05 | 1,16E+06 | 8,64E+05 | 404,05 | 0 | 323,31 | 249 |
| 5,38E+08 | 7,83E+08 | 7,83E+08 | 7,97E+08 | 167,34 | 0 | 323,31 | 288 |
| 2,58E+09 | 1,59E+09 | 1,56E+09 | 1,58E+09 | 34,106 | 0 | 323,31 | 343 |
| 8,75E+08 | 1,87E+08 | 1,76E+08 | 1,73E+08 | 55,262 | 0 | 141,55 | 107 |
| 3,30E+08 | 2,32E+08 | 2,33E+08 | 2,37E+08 | 57,241 | 0 | 323,31 | 122 |
| 2,13E+08 | 3,42E+06 | 4,30E+06 | 4,59E+06 | 32,926 | 0 | 74,935 | 53 |
| 6,78E+07 | 1,46E+07 | 1,57E+07 | 1,41E+07 | 42,164 | 0 | 32,551 | 34 |
| 6,05E+08 | 1,42E+09 | 1,49E+09 | 1,44E+09 | 94,381 | 0 | 323,31 | 258 |
| 3,65E+07 | 1,60E+07 | 1,62E+07 | 1,67E+07 | 23,603 | 0 | 28,22 | 13 |
| 4,53E+08 | 8,17E+07 | 7,51E+07 | 7,54E+07 | 115,2 | 0 | 192,16 | 123 |
| 1,33E+07 | 2,37E+07 | 2,42E+07 | 2,41E+07 | 16,399 | 0 | 17,177 | 10 |
| 9,30E+08 | 2,17E+08 | 2,07E+08 | 2,22E+08 | 69,406 | 0 | 259,01 | 122 |
| 1,51E+09 | 3,36E+09 | 3,35E+09 | 3,31E+09 | 121,94 | 0 | 323,31 | 444 |
| 1,53E+09 | 3,11E+08 | 2,71E+08 | 2,90E+08 | 31,474 | 0 | 323,31 | 188 |
| 3,81E+06 | 5,98E+07 | 5,28E+07 | 4,92E+07 | 48,377 | 0 | 36,535 | 34 |
| 4,12E+08 | 9,40E+07 | 8,94E+07 | 9,17E+07 | 18,904 | 0 | 249,53 | 56 |
| 7,86E+08 | 2,82E+09 | 2,75E+09 | 2,95E+09 | 164,34 | 0 | 323,31 | 388 |
| 5,82E+07 | 4,48E+09 | 3,90E+09 | 3,78E+09 | 52,576 | 0 | 166,64 | 265 |
| 1,69E+08 | 3,43E+08 | 3,31E+08 | 3,33E+08 | 52,921 | 0 | 323,31 | 84 |
| 5,53E+09 | 1,01E+10 | 1,05E+10 | 1,02E+10 | 57,058 | 0 | 323,31 | 815 |
| 7,06E+08 | 2,15E+09 | 2,00E+09 | 2,03E+09 | 177,3 | 0 | 323,31 | 369 |
| 8,15E+07 | 1,53E+08 | 1,54E+08 | 1,56E+08 | 35,242 | 0 | 34,574 | 49 |
| 3,87E+06 | 1,65E+08 | 1,46E+08 | 1,53E+08 | 167,07 | 0 | 39,724 | 65 |
| 2,71E+08 | 1,11E+07 | 9,10E+06 | 9,69E+06 | 48,343 | 0 | 174,71 | 68 |
| 5,27E+08 | 2,66E+08 | 2,61E+08 | 2,61E+08 | 1009,9 | 0 | 233,35 | 246 |
| 1,99E+08 | 6,49E+08 | 7,02E+08 | 6,93E+08 | 34,45 | 0 | 85,232 | 112 |

| | | | | | | | |
|---|---|---|---|---|---|---|---|
| 9,17E+08 | 1,66E+09 | 1,57E+09 | 1,62E+09 | 102,29 | 0 | 323,31 | 375 |
| 2,19E+08 | 1,19E+08 | 1,21E+08 | 1,24E+08 | 147,59 | 0 | 175,32 | 131 |
| 1,13E+07 | 8,08E+07 | 7,50E+07 | 7,65E+07 | 72,484 | 0 | 15,286 | 39 |
| 1,40E+09 | 2,40E+09 | 2,41E+09 | 2,37E+09 | 103,32 | 0 | 323,31 | 362 |
| 8,22E+08 | 3,27E+08 | 3,06E+08 | 3,23E+08 | 102,8 | 0 | 323,31 | 249 |
| 6,80E+08 | 1,09E+09 | 1,13E+09 | 1,10E+09 | 39,547 | 0 | 323,31 | 152 |
| 1,93E+08 | 2,57E+08 | 2,55E+08 | 2,57E+08 | 40,691 | 0 | 219,31 | 102 |
| 5,07E+07 | 1,39E+08 | 1,44E+08 | 1,32E+08 | 127,42 | 0 | 105,19 | 85 |
| 5,95E+07 | 2,61E+08 | 2,87E+08 | 2,82E+08 | 180,25 | 0 | 51,7 | 84 |
| 2,75E+08 | 1,14E+08 | 1,08E+08 | 1,09E+08 | 102,04 | 0 | 64,24 | 117 |
| 1,40E+08 | 6,84E+07 | 7,29E+07 | 7,01E+07 | 131,11 | 0 | 37,035 | 63 |
| 7,02E+07 | 1,60E+08 | 1,69E+08 | 1,66E+08 | 97,666 | 0 | 77,641 | 70 |
| 2,19E+06 | 1,31E+08 | 1,36E+08 | 1,45E+08 | 22,43 | 0 | 124,24 | 17 |
| 6,49E+08 | 4,39E+06 | 4,11E+06 | 2,69E+06 | 60,931 | 0 | 238,93 | 79 |
| 2,60E+08 | 6,25E+08 | 6,05E+08 | 5,96E+08 | 95,986 | 0 | 211,45 | 217 |
| 2,52E+08 | 4,60E+08 | 4,69E+08 | 4,76E+08 | 153,97 | 0 | 201,71 | 172 |
| 1,92E+08 | 6,47E+08 | 6,24E+08 | 6,61E+08 | 66,365 | 0 | 310,51 | 132 |
| 2,41E+08 | 7,25E+08 | 7,54E+08 | 7,58E+08 | 61,359 | 0 | 218,3 | 112 |
| 7,58E+07 | 3,96E+08 | 4,32E+08 | 4,03E+08 | 50,48 | 0 | 60,553 | 77 |
| 1,23E+08 | 2,67E+08 | 2,49E+08 | 2,54E+08 | 40,068 | 0 | 175,16 | 107 |
| 1,53E+09 | 4,53E+08 | 4,85E+08 | 4,60E+08 | 43,953 | 0 | 323,31 | 209 |
| 1,18E+08 | 3,15E+08 | 3,32E+08 | 3,08E+08 | 50,282 | 0 | 166,4 | 100 |
| 3,99E+08 | 3,93E+06 | 4,01E+06 | 6,05E+06 | 132,35 | 0 | 110,53 | 69 |
| 1,77E+08 | 2,37E+06 | 1,76E+06 | 2,65E+06 | 164,7 | 0,000155 | 4,4206 | 12 |
| 3,40E+07 | 3,46E+06 | 3,12E+06 | 4,01E+06 | 42,586 | 0,000148 | 3,3515 | 7 |
| 1,90E+08 | 2,60E+07 | 2,98E+07 | 2,49E+07 | 51,599 | 0 | 77,254 | 52 |
| 1,01E+08 | 2,48E+06 | 1,80E+06 | 2,68E+06 | 61,001 | 0 | 38,739 | 27 |
| 2,82E+08 | 8,04E+07 | 7,43E+07 | 7,36E+07 | 135,63 | 0 | 91,585 | 120 |
| 4,40E+08 | 1,05E+09 | 1,02E+09 | 1,01E+09 | 35,406 | 0 | 195,75 | 117 |
| 6,56E+07 | 2,29E+08 | 2,38E+08 | 2,28E+08 | 89,679 | 0 | 86,865 | 68 |
| 8,17E+06 | 5,69E+07 | 6,98E+07 | 6,81E+07 | 326,74 | 0 | 28,079 | 59 |
| 1,80E+06 | 2,45E+07 | 2,51E+07 | 2,42E+07 | 73,947 | 0 | 10,98 | 18 |
| 1,03E+08 | 2,68E+08 | 2,92E+08 | 2,87E+08 | 149,32 | 0 | 70,636 | 125 |
| 1,26E+08 | 5,16E+07 | 4,90E+07 | 5,42E+07 | 54,381 | 0 | 23,607 | 48 |
| 1,12E+08 | 8,50E+06 | 8,58E+06 | 6,69E+06 | 154,64 | 0 | 59,981 | 70 |
| 1,41E+08 | 4,65E+08 | 4,32E+08 | 4,68E+08 | 72,571 | 0 | 107,26 | 153 |
| 2,46E+08 | 4,76E+08 | 4,74E+08 | 4,58E+08 | 21,606 | 0 | 68,951 | 99 |
| 8,12E+08 | 1,65E+08 | 1,77E+08 | 1,61E+08 | 53,514 | 0 | 264,65 | 132 |
| 2,57E+09 | 1,20E+09 | 1,30E+09 | 1,21E+09 | 44,816 | 0 | 323,31 | 212 |
| 9,35E+08 | 3,50E+08 | 3,39E+08 | 3,41E+08 | 33,155 | 0 | 323,31 | 98 |
| 2,73E+09 | 8,36E+09 | 8,54E+09 | 8,09E+09 | 34,127 | 0 | 323,31 | 507 |
| 1,64E+08 | 4,95E+08 | 5,47E+08 | 5,59E+08 | 79,327 | 0 | 323,31 | 140 |
| 4,51E+08 | 2,68E+08 | 2,64E+08 | 2,72E+08 | 25,661 | 0 | 32,97 | 117 |
| 6,00E+07 | 1,26E+08 | 1,29E+08 | 1,34E+08 | 84,755 | 0 | 20,903 | 36 |
| 5,08E+08 | 5,27E+07 | 4,71E+07 | 5,99E+07 | 210,8 | 0 | 74,023 | 96 |
| 3,10E+07 | 1,56E+08 | 1,39E+08 | 1,42E+08 | 25,172 | 0 | 93,014 | 40 |
| 4,59E+06 | 3,43E+07 | 3,85E+07 | 3,88E+07 | 56,84 | 0,000158 | 5,0652 | 17 |
| 2,40E+08 | 7,77E+05 | 1,18E+06 | 1,47E+06 | 46,158 | 0 | 52,742 | 47 |
| 3,92E+08 | 1,42E+08 | 1,36E+08 | 1,50E+08 | 50,118 | 0 | 77,406 | 87 |
| 3,23E+06 | 4,49E+08 | 4,15E+08 | 5,68E+08 | 28,793 | 0 | 272,79 | 16 |

| | | | | | | | |
|---|---|---|---|---|---|---|---|
| 9,66E+07 | 4,29E+06 | 3,28E+06 | 3,69E+06 | 174,89 | 0 | 96,576 | 47 |
| 2,15E+08 | 4,50E+07 | 5,06E+07 | 4,42E+07 | 131,83 | 0 | 105,52 | 90 |
| 3,13E+08 | 4,14E+07 | 4,01E+07 | 3,82E+07 | 36,724 | 0 | 180,99 | 74 |
| 5,94E+08 | 2,16E+08 | 2,14E+08 | 2,34E+08 | 44,889 | 0 | 160,57 | 141 |
| 2,01E+08 | 5,36E+08 | 4,97E+08 | 4,80E+08 | 22,329 | 0 | 124,55 | 78 |
| 1,75E+08 | 3,55E+06 | 3,12E+06 | 2,29E+06 | 25,006 | 0 | 323,31 | 9 |
| 3,20E+06 | 2,81E+08 | 2,93E+08 | 2,92E+08 | 49,609 | 0 | 53,721 | 59 |
| 3,24E+07 | 5,16E+07 | 5,27E+07 | 5,00E+07 | 115,69 | 0 | 70,271 | 59 |
| 3,67E+08 | 5,51E+08 | 5,36E+08 | 5,48E+08 | 54,317 | 0 | 94,328 | 120 |
| 5,62E+09 | 3,36E+09 | 3,35E+09 | 3,36E+09 | 274,22 | 0 | 323,31 | 1073 |
| 8,64E+08 | 3,60E+08 | 3,62E+08 | 3,83E+08 | 35,649 | 0 | 294,55 | 95 |
| 7,69E+07 | 2,14E+08 | 1,95E+08 | 2,12E+08 | 78,101 | 0 | 147,7 | 94 |
| 7,84E+07 | 5,09E+07 | 5,26E+07 | 5,20E+07 | 126,64 | 0 | 30,635 | 74 |
| 1,75E+09 | 4,06E+09 | 4,30E+09 | 4,35E+09 | 62,766 | 0 | 323,31 | 592 |
| 1,60E+09 | 2,22E+09 | 2,28E+09 | 2,22E+09 | 119,49 | 0 | 323,31 | 340 |
| 3,05E+07 | 6,40E+06 | 6,27E+06 | 6,88E+06 | 46,3 | 0 | 10,463 | 13 |
| 3,57E+08 | 4,80E+05 | 9,85E+05 | 5,42E+05 | 24,394 | 0 | 28,31 | 46 |
| 1,12E+08 | 3,14E+06 | 2,77E+06 | 1,94E+06 | 37,859 | 0 | 72,442 | 26 |
| 1,04E+08 | 4,06E+06 | 5,66E+06 | 4,01E+06 | 37,548 | 0 | 55,75 | 24 |
| 7,64E+07 | 3,48E+06 | 2,53E+06 | 3,75E+06 | 123,12 | 0 | 78,89 | 43 |
| 1,65E+07 | 1,71E+06 | 2,13E+06 | 1,82E+06 | 280,81 | 0 | 8,7785 | 16 |
| 3,75E+08 | 2,60E+08 | 2,51E+08 | 2,49E+08 | 76,293 | 0 | 280,59 | 147 |
| 1,71E+08 | 4,53E+08 | 4,51E+08 | 4,95E+08 | 20,802 | 0 | 268,45 | 65 |
| 8,55E+07 | 1,72E+08 | 1,64E+08 | 1,58E+08 | 70,292 | 0 | 60,904 | 81 |
| 1,43E+08 | 2,66E+06 | 1,72E+06 | 2,01E+06 | 71,062 | 0 | 209,61 | 29 |
| 6,44E+08 | 8,66E+08 | 8,95E+08 | 8,97E+08 | 29,435 | 0 | 285,56 | 192 |
| 1,12E+09 | 6,22E+08 | 6,16E+08 | 5,83E+08 | 74,18 | 0 | 323,31 | 208 |
| 1,48E+08 | 2,89E+06 | 4,74E+06 | 3,94E+06 | 55,939 | 0 | 162,63 | 33 |
| 1,82E+09 | 3,83E+09 | 3,81E+09 | 3,70E+09 | 49,816 | 0 | 323,31 | 298 |
| 1,14E+09 | 2,75E+09 | 2,82E+09 | 2,91E+09 | 37,387 | 0 | 180,98 | 247 |
| 3,00E+08 | 2,07E+08 | 2,12E+08 | 2,10E+08 | 96,304 | 0 | 64,345 | 132 |
| 1,35E+08 | 2,04E+06 | 3,49E+06 | 2,37E+06 | 81,8 | 0 | 55,811 | 41 |
| 2,46E+10 | 4,09E+10 | 4,12E+10 | 4,05E+10 | 534,18 | 0 | 323,31 | 4055 |
| 1,84E+08 | 1,21E+08 | 1,19E+08 | 1,14E+08 | 108,19 | 0 | 92,052 | 121 |
| 7,60E+08 | 2,20E+09 | 2,27E+09 | 2,08E+09 | 44,953 | 0 | 323,31 | 198 |
| 1,43E+09 | 3,91E+08 | 3,97E+08 | 4,21E+08 | 122,16 | 0 | 323,31 | 253 |
| 2,20E+08 | 3,57E+07 | 4,39E+07 | 3,85E+07 | 65,918 | 0 | 75,127 | 61 |
| 4,36E+07 | 2,48E+06 | 1,87E+06 | 2,40E+06 | 181,39 | 0 | 28,832 | 29 |
| 5,89E+08 | 2,97E+08 | 3,06E+08 | 3,06E+08 | 175,05 | 0 | 266,63 | 229 |
| 8,90E+07 | 1,56E+08 | 1,59E+08 | 1,56E+08 | 87,196 | 0 | 76,476 | 87 |
| 1,48E+08 | 8,35E+06 | 9,08E+06 | 7,78E+06 | 70,683 | 0 | 134,29 | 50 |
| 2,36E+08 | 4,54E+08 | 4,17E+08 | 4,42E+08 | 39,254 | 0 | 208,06 | 128 |
| 9,82E+07 | 2,97E+06 | 1,77E+06 | 2,01E+06 | 35,866 | 0 | 31,026 | 32 |
| 3,13E+06 | 3,29E+08 | 2,25E+08 | 2,18E+08 | 23,045 | 0 | 88,544 | 56 |
| 9,86E+08 | 4,95E+08 | 4,86E+08 | 4,83E+08 | 206,23 | 0 | 323,31 | 311 |
| 2,98E+08 | 1,16E+08 | 1,12E+08 | 1,24E+08 | 35,428 | 0 | 323,31 | 38 |
| 3,46E+08 | 6,62E+08 | 6,87E+08 | 7,06E+08 | 27,61 | 0 | 323,31 | 129 |
| 6,69E+07 | 1,32E+07 | 1,29E+07 | 1,10E+07 | 44,814 | 0 | 8,1694 | 16 |
| 8,66E+07 | 3,14E+08 | 2,99E+08 | 2,99E+08 | 22,45 | 0 | 119,46 | 57 |
| 5,03E+07 | 3,04E+08 | 3,36E+08 | 2,99E+08 | 253,62 | 0 | 180,75 | 192 |

| | | | | | | | |
|---|---|---|---|---|---|---|---|
| 2,46E+08 | 8,58E+08 | 8,67E+08 | 8,28E+08 | 146,56 | 0 | 323,31 | 228 |
| 2,07E+07 | 3,68E+07 | 3,47E+07 | 3,66E+07 | 12,047 | 0 | 7,0778 | 16 |
| 4,07E+07 | 1,14E+08 | 1,30E+08 | 1,24E+08 | 45,126 | 0 | 44,061 | 35 |
| 4,44E+07 | 9,54E+07 | 1,08E+08 | 1,02E+08 | 136,61 | 0 | 32,604 | 50 |
| 1,62E+08 | 1,13E+08 | 1,11E+08 | 1,17E+08 | 93,39 | 0 | 88,446 | 94 |
| 1,90E+09 | 5,88E+09 | 6,77E+09 | 6,23E+09 | 27,515 | 0 | 323,31 | 143 |
| 8,65E+07 | 7,91E+08 | 6,48E+08 | 8,49E+08 | 22,197 | 0 | 258,24 | 88 |
| 2,18E+08 | 3,40E+08 | 3,53E+08 | 3,39E+08 | 125,78 | 0 | 323,31 | 153 |
| 1,05E+09 | 1,52E+09 | 1,60E+09 | 1,55E+09 | 69,05 | 0 | 323,31 | 258 |
| 1,18E+09 | 2,02E+09 | 2,02E+09 | 2,16E+09 | 36,586 | 0 | 323,31 | 182 |
| 1,43E+08 | 8,37E+07 | 8,47E+07 | 8,47E+07 | 55,792 | 0 | 83,939 | 77 |
| 3,92E+07 | 4,98E+07 | 4,98E+07 | 5,01E+07 | 156,68 | 0 | 14,402 | 36 |
| 5,73E+08 | 2,50E+06 | 5,20E+06 | 3,88E+06 | 128,56 | 0 | 323,31 | 110 |
| 3,32E+08 | 3,17E+06 | 2,96E+06 | 1,66E+06 | 74,621 | 0 | 141,4 | 55 |
| 8,03E+08 | 4,58E+08 | 4,53E+08 | 4,38E+08 | 304,27 | 0 | 323,31 | 433 |
| 6,89E+08 | 4,16E+08 | 4,28E+08 | 4,04E+08 | 398,29 | 0 | 323,31 | 363 |
| 3,92E+08 | 1,89E+08 | 2,02E+08 | 1,93E+08 | 99,875 | 0 | 323,31 | 159 |
| 3,80E+06 | 2,93E+07 | 2,53E+07 | 2,79E+07 | 164,08 | 0 | 11,861 | 20 |
| 1,84E+08 | 6,74E+08 | 6,29E+08 | 6,29E+08 | 115,04 | 0 | 234,56 | 122 |
| 1,14E+09 | 6,11E+08 | 5,99E+08 | 5,82E+08 | 273,81 | 0 | 323,31 | 380 |
| 2,13E+08 | 1,49E+08 | 1,51E+08 | 1,56E+08 | 41,287 | 0 | 80,275 | 86 |
| 1,00E+09 | 2,81E+06 | 1,26E+06 | 2,07E+06 | 77,286 | 0 | 323,31 | 91 |
| 6,42E+07 | 2,00E+07 | 2,03E+07 | 2,21E+07 | 29,69 | 0 | 6,764 | 19 |
| 1,05E+08 | 2,46E+08 | 2,23E+08 | 2,33E+08 | 46,646 | 0 | 105,99 | 92 |
| 9,02E+08 | 2,58E+08 | 2,34E+08 | 2,51E+08 | 61,48 | 0 | 323,31 | 194 |
| 8,11E+08 | 2,39E+09 | 2,42E+09 | 2,62E+09 | 61,417 | 0 | 323,31 | 315 |
| 7,15E+07 | 2,88E+06 | 1,69E+06 | 2,28E+06 | 136,45 | 0 | 86,602 | 33 |
| 4,27E+08 | 1,58E+08 | 1,66E+08 | 1,44E+08 | 41,813 | 0 | 214,78 | 63 |
| 2,86E+08 | 1,32E+08 | 1,27E+08 | 1,26E+08 | 64,989 | 0 | 107,64 | 104 |
| 1,30E+09 | 5,45E+08 | 4,95E+08 | 5,38E+08 | 28,673 | 0 | 323,31 | 147 |
| 1,12E+09 | 6,42E+08 | 6,32E+08 | 6,02E+08 | 117,37 | 0 | 290,49 | 294 |
| 1,38E+08 | 2,30E+08 | 2,37E+08 | 2,44E+08 | 56,431 | 0 | 101,09 | 83 |
| 1,51E+08 | 8,39E+07 | 8,80E+07 | 8,51E+07 | 31,875 | 0 | 283,05 | 47 |
| 8,56E+07 | 1,80E+08 | 1,74E+08 | 1,93E+08 | 18,02 | 0 | 50,043 | 28 |
| 1,00E+08 | 1,52E+07 | 1,32E+07 | 1,48E+07 | 59,211 | 0 | 22,616 | 24 |
| 1,52E+06 | 1,53E+08 | 1,61E+08 | 1,45E+08 | 30,953 | 0 | 58,744 | 35 |
| 4,23E+09 | 1,46E+09 | 1,39E+09 | 1,30E+09 | 123,59 | 0 | 323,31 | 504 |
| 3,82E+09 | 9,25E+09 | 8,99E+09 | 8,57E+09 | 117,81 | 0 | 323,31 | 668 |
| 3,06E+08 | 2,30E+08 | 2,35E+08 | 2,30E+08 | 168,93 | 0 | 218,01 | 193 |
| 3,32E+09 | 2,27E+09 | 2,24E+09 | 2,16E+09 | 18,749 | 0 | 323,31 | 234 |
| 3,07E+06 | 2,40E+07 | 2,56E+07 | 2,82E+07 | 75,294 | 0 | 10,811 | 22 |
| 3,32E+06 | 1,26E+07 | 1,58E+07 | 1,45E+07 | 53,435 | 0 | 5,9026 | 8 |
| 8,36E+07 | 2,05E+07 | 1,95E+07 | 1,82E+07 | 38,058 | 0 | 17,023 | 36 |
| 8,56E+06 | 1,61E+07 | 1,68E+07 | 1,79E+07 | 52,995 | 0,000158 | 5,1027 | 15 |
| 2,66E+08 | 3,12E+07 | 2,36E+07 | 2,89E+07 | 80,109 | 0 | 65,635 | 64 |
| 3,04E+08 | 4,42E+06 | 2,07E+06 | 3,19E+06 | 71,342 | 0 | 323,31 | 74 |
| 2,43E+06 | 4,57E+07 | 4,69E+07 | 4,71E+07 | 90,029 | 0 | 10,338 | 15 |
| 1,03E+09 | 3,56E+08 | 4,06E+08 | 3,49E+08 | 58,758 | 0 | 323,31 | 133 |
| 1,20E+08 | 2,87E+08 | 2,55E+08 | 2,69E+08 | 52,295 | 0 | 37,677 | 76 |
| 1,96E+08 | 3,48E+08 | 3,55E+08 | 3,36E+08 | 294 | 0 | 125,13 | 212 |

| | | | | | | | |
|---|---|---|---|---|---|---|---|
| 2,16E+08 | 3,69E+08 | 3,61E+08 | 3,93E+08 | 67,513 | 0 | 144,9 | 117 |
| 4,32E+08 | 1,93E+08 | 1,87E+08 | 1,84E+08 | 46,674 | 0 | 173,22 | 151 |
| 4,46E+08 | 6,91E+08 | 6,95E+08 | 7,04E+08 | 88,947 | 0 | 323,31 | 233 |
| 5,80E+08 | 8,78E+08 | 8,72E+08 | 9,17E+08 | 532,16 | 0 | 323,31 | 467 |
| 5,43E+08 | 9,71E+08 | 1,05E+09 | 1,06E+09 | 103,99 | 0 | 218,64 | 243 |
| 9,48E+08 | 1,35E+09 | 1,34E+09 | 1,33E+09 | 290,08 | 0 | 323,31 | 496 |
| 1,44E+07 | 2,70E+08 | 2,04E+08 | 2,20E+08 | 21,298 | 0 | 165,31 | 18 |
| 8,17E+08 | 1,50E+09 | 1,45E+09 | 1,45E+09 | 87,22 | 0 | 323,31 | 276 |
| 1,34E+09 | 5,33E+08 | 5,02E+08 | 4,74E+08 | 121,55 | 0 | 323,31 | 265 |
| 3,78E+08 | 1,36E+08 | 1,57E+08 | 1,45E+08 | 38,6 | 0 | 68,44 | 51 |
| 3,09E+08 | 4,59E+06 | 2,78E+06 | 2,10E+06 | 30,26 | 0 | 97,788 | 41 |
| 3,30E+09 | 5,15E+09 | 5,15E+09 | 5,20E+09 | 110,45 | 0 | 323,31 | 394 |
| 3,38E+08 | 2,45E+08 | 2,55E+08 | 2,48E+08 | 51,737 | 0 | 108,59 | 127 |
| 1,18E+08 | 4,33E+07 | 4,38E+07 | 3,80E+07 | 143,07 | 0 | 53,264 | 82 |
| 3,24E+08 | 8,63E+07 | 8,01E+07 | 8,38E+07 | 131,96 | 0 | 141,12 | 121 |
| 9,46E+07 | 1,35E+08 | 1,40E+08 | 1,39E+08 | 233,03 | 0 | 85,056 | 121 |
| 9,39E+07 | 3,13E+08 | 2,98E+08 | 3,00E+08 | 16,131 | 0 | 97,74 | 58 |
| 1,06E+09 | 2,08E+09 | 2,15E+09 | 2,35E+09 | 77,738 | 0 | 323,31 | 324 |
| 9,98E+09 | 2,27E+10 | 2,55E+10 | 2,61E+10 | 44,55 | 0 | 323,31 | 1145 |
| 7,42E+07 | 1,32E+06 | 2,47E+06 | 1,53E+06 | 70,733 | 0 | 19,726 | 33 |
| 8,66E+08 | 1,54E+09 | 1,56E+09 | 1,52E+09 | 132,89 | 0 | 323,31 | 288 |
| 1,83E+07 | 3,30E+06 | 3,66E+06 | 2,74E+06 | 79,081 | 0,000157 | 4,9835 | 13 |
| 3,73E+08 | 1,55E+08 | 1,65E+08 | 1,55E+08 | 37,392 | 0 | 118,32 | 77 |
| 4,40E+08 | 2,08E+08 | 2,09E+08 | 2,16E+08 | 41,291 | 0 | 323,31 | 106 |
| 3,27E+09 | 5,28E+09 | 5,06E+09 | 5,29E+09 | 136,33 | 0 | 323,31 | 517 |
| 2,31E+08 | 8,84E+06 | 6,23E+06 | 9,22E+06 | 38,555 | 0 | 125,79 | 30 |
| 1,93E+08 | 9,14E+07 | 9,37E+07 | 1,00E+08 | 130,45 | 0 | 299,04 | 103 |
| 5,34E+09 | 8,25E+09 | 8,59E+09 | 8,43E+09 | 188,74 | 0 | 323,31 | 952 |
| 1,10E+09 | 8,13E+08 | 8,32E+08 | 8,11E+08 | 59,236 | 0 | 323,31 | 227 |
| 1,08E+09 | 1,45E+09 | 1,51E+09 | 1,47E+09 | 782,72 | 0 | 323,31 | 712 |
| 7,62E+07 | 3,40E+06 | 2,01E+06 | 2,64E+06 | 66,107 | 0 | 14,03 | 15 |
| 2,77E+09 | 3,51E+09 | 3,56E+09 | 3,56E+09 | 102,09 | 0 | 323,31 | 517 |
| 5,13E+08 | 7,76E+08 | 8,19E+08 | 7,81E+08 | 83,041 | 0 | 323,31 | 199 |
| 2,98E+08 | 6,72E+08 | 7,51E+08 | 6,66E+08 | 59,601 | 0 | 235,54 | 125 |
| 3,14E+06 | 4,72E+07 | 5,16E+07 | 5,82E+07 | 20,437 | 0 | 9,2876 | 12 |
| 9,46E+07 | 1,98E+08 | 1,86E+08 | 1,87E+08 | 91,836 | 0 | 59,082 | 79 |
| 7,64E+08 | 7,07E+06 | 4,06E+06 | 3,51E+06 | 72,107 | 0 | 6,4523 | 14 |
| 1,12E+09 | 5,76E+08 | 5,96E+08 | 5,94E+08 | 72,399 | 0 | 291,6 | 156 |
| 2,02E+06 | 6,37E+09 | 6,75E+09 | 6,37E+09 | 21,785 | 0 | 323,31 | 143 |
| 9,90E+07 | 5,15E+07 | 5,27E+07 | 5,02E+07 | 133,51 | 0 | 115,96 | 60 |
| 2,75E+08 | 4,34E+06 | 2,03E+06 | 2,22E+06 | 25,492 | 0 | 74,756 | 29 |
| 1,11E+08 | 6,94E+07 | 7,01E+07 | 6,95E+07 | 73,162 | 0 | 138,94 | 70 |
| 3,72E+06 | 9,12E+08 | 9,81E+08 | 9,26E+08 | 103,34 | 0 | 323,31 | 154 |
| 8,29E+07 | 9,59E+06 | 7,33E+06 | 1,07E+07 | 80,371 | 0 | 179,92 | 50 |
| 9,79E+07 | 1,88E+08 | 1,84E+08 | 1,92E+08 | 170,39 | 0 | 72,891 | 108 |
| 1,45E+08 | 3,11E+08 | 3,34E+08 | 3,29E+08 | 28,944 | 0 | 43,141 | 56 |
| 1,39E+08 | 3,35E+07 | 4,15E+07 | 3,60E+07 | 32,45 | 0 | 36,228 | 48 |
| 2,03E+08 | 9,79E+07 | 1,02E+08 | 1,09E+08 | 260,64 | 0 | 92,637 | 126 |
| 3,92E+08 | 2,25E+08 | 2,31E+08 | 2,18E+08 | 87,334 | 0 | 135,78 | 154 |
| 6,15E+08 | 1,41E+09 | 1,25E+09 | 1,30E+09 | 21,066 | 0 | 323,31 | 172 |

| | | | | | | | |
|---|---|---|---|---|---|---|---|
| 1,53E+08 | 2,47E+07 | 2,58E+07 | 3,03E+07 | 21,66 | 0 | 58,508 | 41 |
| 9,12E+07 | 5,60E+08 | 5,00E+08 | 5,01E+08 | 27,853 | 0 | 94,2 | 77 |
| 2,07E+08 | 1,37E+08 | 1,42E+08 | 1,45E+08 | 71,011 | 0 | 180,56 | 102 |
| 1,09E+09 | 4,28E+08 | 4,26E+08 | 4,21E+08 | 63,294 | 0 | 281,57 | 211 |
| 3,40E+09 | 2,35E+09 | 2,41E+09 | 2,38E+09 | 96,023 | 0 | 323,31 | 419 |
| 1,03E+08 | 2,14E+08 | 2,19E+08 | 2,10E+08 | 23,121 | 0 | 119,57 | 74 |
| 1,72E+07 | 1,12E+07 | 1,10E+07 | 1,05E+07 | 43,452 | 0,00015 | 3,611 | 17 |
| 8,40E+08 | 1,60E+09 | 1,55E+09 | 1,48E+09 | 160,91 | 0 | 323,31 | 391 |
| 2,57E+07 | 1,48E+07 | 1,61E+07 | 1,48E+07 | 43,835 | 0 | 6,2256 | 16 |
| 4,55E+08 | 8,91E+08 | 9,01E+08 | 8,34E+08 | 290,71 | 0 | 323,31 | 292 |
| 7,19E+08 | 1,18E+09 | 1,17E+09 | 1,11E+09 | 53,706 | 0 | 293,79 | 213 |
| 2,38E+08 | 1,50E+08 | 1,49E+08 | 1,58E+08 | 142 | 0 | 168,82 | 125 |
| 2,78E+08 | 1,75E+08 | 1,64E+08 | 1,71E+08 | 75,1 | 0 | 158,34 | 114 |
| 4,88E+08 | 1,03E+09 | 1,04E+09 | 1,01E+09 | 61,859 | 0 | 323,31 | 272 |
| 2,57E+08 | 1,47E+08 | 1,54E+08 | 1,59E+08 | 85,247 | 0 | 56,274 | 61 |
| 4,54E+07 | 2,34E+06 | 3,21E+06 | 2,90E+06 | 169,42 | 0 | 55,777 | 24 |
| 8,22E+07 | 5,15E+07 | 4,97E+07 | 4,84E+07 | 95,974 | 0 | 60,754 | 73 |
| 9,53E+07 | 1,35E+07 | 1,68E+07 | 1,25E+07 | 34,737 | 0 | 28,147 | 34 |
| 3,48E+08 | 1,01E+08 | 9,53E+07 | 9,95E+07 | 38,114 | 0 | 271,95 | 95 |
| 1,62E+08 | 2,33E+08 | 2,40E+08 | 2,31E+08 | 84,186 | 0 | 35,16 | 105 |
| 1,61E+09 | 2,71E+09 | 2,71E+09 | 2,76E+09 | 70,741 | 0 | 323,31 | 322 |
| 3,37E+08 | 6,78E+08 | 6,23E+08 | 6,43E+08 | 103,16 | 0 | 240,23 | 199 |
| 3,09E+07 | 5,52E+07 | 5,75E+07 | 5,66E+07 | 137,51 | 0 | 54,497 | 51 |
| 6,46E+08 | 3,36E+08 | 3,06E+08 | 3,19E+08 | 82,228 | 0 | 323,31 | 182 |
| 7,74E+07 | 1,45E+06 | 2,52E+06 | 2,55E+06 | 37,402 | 0 | 65,025 | 32 |
| 4,40E+07 | 7,67E+07 | 7,07E+07 | 7,35E+07 | 80,376 | 0 | 23,593 | 57 |
| 8,47E+08 | 1,53E+09 | 1,43E+09 | 1,54E+09 | 106,98 | 0 | 323,31 | 379 |
| 1,16E+08 | 4,33E+07 | 4,65E+07 | 4,79E+07 | 99,66 | 0 | 153,2 | 84 |
| 7,35E+07 | 5,02E+06 | 3,32E+06 | 3,37E+06 | 76,941 | 0 | 38,412 | 31 |
| 1,65E+10 | 1,12E+10 | 1,12E+10 | 1,11E+10 | 56,3 | 0 | 323,31 | 969 |
| 6,57E+08 | 5,00E+08 | 5,11E+08 | 5,04E+08 | 54,614 | 0 | 323,31 | 145 |
| 6,06E+09 | 1,95E+06 | 2,46E+06 | 7,15E+06 | 54,565 | 0 | 323,31 | 256 |
| 3,30E+09 | 4,61E+09 | 4,56E+09 | 4,47E+09 | 235,63 | 0 | 323,31 | 876 |
| 5,22E+07 | 1,96E+07 | 2,25E+07 | 2,01E+07 | 38,522 | 0 | 94,148 | 26 |
| 7,76E+08 | 2,40E+08 | 2,56E+08 | 2,45E+08 | 74,466 | 0 | 323,31 | 185 |
| 4,26E+08 | 8,47E+08 | 8,11E+08 | 8,13E+08 | 831,87 | 0 | 323,31 | 521 |
| 5,26E+07 | 2,96E+06 | 2,14E+06 | 1,78E+06 | 81,635 | 0 | 20,806 | 20 |
| 3,07E+07 | 2,00E+06 | 1,83E+06 | 1,18E+06 | 44,059 | 0 | 6,3813 | 7 |
| 8,20E+06 | 6,48E+07 | 5,33E+07 | 5,63E+07 | 61,554 | 0 | 38,898 | 31 |
| 4,38E+07 | 2,12E+06 | 3,10E+06 | 1,90E+06 | 25,814 | 0 | 323,31 | 11 |
| 1,50E+08 | 3,14E+06 | 3,82E+06 | 6,23E+06 | 53,844 | 0 | 24,909 | 38 |
| 8,82E+08 | 1,35E+09 | 1,30E+09 | 1,32E+09 | 54,357 | 0 | 323,31 | 236 |
| 1,68E+08 | 6,47E+07 | 6,70E+07 | 7,01E+07 | 58,408 | 0 | 152,1 | 82 |
| 1,66E+09 | 7,89E+08 | 7,79E+08 | 7,16E+08 | 80,208 | 0 | 323,31 | 292 |
| 1,24E+07 | 2,81E+06 | 2,82E+06 | 2,69E+06 | 69,598 | 0,000148 | 3,3911 | 6 |
| 8,82E+08 | 1,65E+09 | 1,51E+09 | 1,55E+09 | 35,49 | 0 | 323,31 | 184 |
| 3,75E+06 | 4,36E+07 | 4,42E+07 | 4,88E+07 | 82,348 | 0 | 15,068 | 29 |
| 6,79E+07 | 3,26E+07 | 3,30E+07 | 2,92E+07 | 65,978 | 0 | 34,996 | 31 |
| 1,47E+08 | 1,10E+07 | 7,99E+06 | 1,33E+07 | 151,86 | 0 | 62,661 | 73 |
| 1,30E+08 | 2,88E+07 | 2,38E+07 | 2,06E+07 | 25,958 | 0 | 65,597 | 29 |

| | | | | | | | |
|---|---|---|---|---|---|---|---|
| 3,73E+08 | 9,83E+08 | 1,06E+09 | 1,16E+09 | 27,738 | 0 | 192,96 | 108 |
| 2,40E+06 | 1,16E+08 | 1,72E+08 | 1,88E+08 | 35,286 | 0 | 56,341 | 50 |
| 1,23E+08 | 1,88E+08 | 1,85E+08 | 1,96E+08 | 37,817 | 0 | 323,31 | 58 |
| 1,40E+09 | 1,89E+09 | 2,00E+09 | 1,97E+09 | 584,78 | 0 | 323,31 | 866 |
| 1,28E+08 | 7,58E+06 | 9,33E+06 | 6,09E+06 | 27,914 | 0 | 15,313 | 19 |
| 5,16E+07 | 3,10E+07 | 3,30E+07 | 3,13E+07 | 111,77 | 0 | 42,475 | 52 |
| 3,62E+08 | 3,80E+06 | 1,69E+06 | 3,97E+06 | 122,06 | 0 | 135,54 | 99 |
| 8,48E+07 | 5,39E+07 | 5,40E+07 | 5,38E+07 | 42,136 | 0 | 61,835 | 38 |
| 4,46E+07 | 1,36E+07 | 1,10E+07 | 1,38E+07 | 40,156 | 0 | 53,382 | 17 |
| 8,64E+07 | 1,26E+08 | 1,19E+08 | 1,22E+08 | 172,11 | 0 | 59,618 | 105 |
| 9,30E+08 | 1,31E+09 | 1,31E+09 | 1,27E+09 | 203,7 | 0 | 323,31 | 432 |
| 4,37E+06 | 1,76E+08 | 1,70E+08 | 1,60E+08 | 81,303 | 0 | 85,708 | 66 |
| 3,54E+07 | 1,19E+07 | 1,04E+07 | 1,08E+07 | 109,9 | 0 | 54,364 | 27 |
| 3,42E+08 | 4,04E+08 | 4,07E+08 | 4,08E+08 | 40,744 | 0 | 323,31 | 133 |
| 5,03E+08 | 1,74E+08 | 1,67E+08 | 1,87E+08 | 92,371 | 0 | 181,62 | 157 |
| 4,56E+08 | 1,60E+08 | 1,56E+08 | 1,62E+08 | 80,581 | 0 | 159,93 | 123 |
| 1,32E+09 | 3,70E+06 | 1,52E+06 | 5,15E+06 | 201,65 | 0 | 323,31 | 197 |
| 5,02E+09 | 3,04E+09 | 2,88E+09 | 3,16E+09 | 67,63 | 0 | 323,31 | 579 |
| 3,85E+07 | 1,64E+07 | 1,82E+07 | 1,54E+07 | 125,24 | 0 | 20,512 | 38 |
| 3,08E+08 | 1,86E+08 | 1,81E+08 | 1,96E+08 | 106,86 | 0 | 310,52 | 126 |
| 2,58E+07 | 9,28E+06 | 9,11E+06 | 7,96E+06 | 55,546 | 0 | 8,0411 | 16 |
| 1,09E+09 | 2,08E+09 | 2,01E+09 | 2,00E+09 | 101,2 | 0 | 323,31 | 408 |
| 1,67E+09 | 2,32E+09 | 2,48E+09 | 2,39E+09 | 105,84 | 0 | 323,31 | 478 |
| 5,86E+08 | 3,43E+08 | 3,37E+08 | 3,52E+08 | 72,415 | 0 | 291,51 | 154 |
| 6,58E+07 | 1,23E+08 | 1,21E+08 | 1,27E+08 | 48,064 | 0 | 126,03 | 70 |
| 2,96E+08 | 6,55E+05 | 1,86E+06 | 8,34E+05 | 44,541 | 0 | 323,31 | 60 |
| 6,82E+08 | 1,06E+09 | 9,89E+08 | 1,03E+09 | 55,446 | 0 | 323,31 | 193 |
| 1,33E+08 | 3,39E+08 | 3,30E+08 | 3,73E+08 | 33,428 | 0 | 323,31 | 115 |
| 5,22E+07 | 2,49E+06 | 1,65E+06 | 1,24E+06 | 40,141 | 0 | 23,43 | 25 |
| 1,60E+07 | 3,49E+06 | 2,75E+06 | 3,05E+06 | 14,458 | 0 | 5,8697 | 10 |
| 1,01E+08 | 6,80E+07 | 6,59E+07 | 6,64E+07 | 53,721 | 0 | 149,44 | 45 |
| 3,62E+08 | 1,02E+09 | 9,94E+08 | 1,11E+09 | 81,604 | 0 | 323,31 | 223 |
| 3,64E+08 | 4,35E+06 | 1,64E+06 | 3,40E+06 | 35,692 | 0 | 120,73 | 24 |
| 8,70E+08 | 5,00E+08 | 4,81E+08 | 4,92E+08 | 38,158 | 0 | 323,31 | 119 |
| 1,70E+09 | 5,14E+09 | 4,93E+09 | 4,92E+09 | 207,12 | 0 | 323,31 | 741 |
| 1,22E+08 | 7,59E+07 | 6,89E+07 | 6,86E+07 | 167,32 | 0 | 32,183 | 70 |
| 3,35E+08 | 7,73E+08 | 7,36E+08 | 7,03E+08 | 69,042 | 0 | 226,88 | 141 |
| 5,54E+06 | 2,36E+09 | 2,03E+09 | 2,37E+09 | 24,57 | 0 | 138,85 | 148 |
| 2,00E+07 | 6,50E+07 | 5,43E+07 | 5,33E+07 | 81,163 | 0 | 127,95 | 42 |
| 1,38E+08 | 6,48E+07 | 6,82E+07 | 7,05E+07 | 47,466 | 0 | 35,173 | 80 |
| 2,38E+08 | 3,63E+06 | 5,52E+06 | 3,66E+06 | 18,032 | 0 | 288,16 | 21 |
| 7,63E+07 | 4,39E+07 | 4,74E+07 | 4,61E+07 | 41,486 | 0 | 22,177 | 42 |
| 5,76E+09 | 1,18E+10 | 1,28E+10 | 1,32E+10 | 32,931 | 0 | 138,57 | 330 |
| 5,99E+07 | 3,49E+07 | 3,24E+07 | 3,12E+07 | 33,07 | 0 | 10,87 | 23 |
| 3,57E+08 | 7,97E+08 | 9,42E+08 | 8,16E+08 | 70,175 | 0 | 127,53 | 165 |
| 4,87E+07 | 4,51E+06 | 3,87E+06 | 6,03E+06 | 73,142 | 0 | 21,787 | 16 |
| 5,61E+08 | 3,44E+08 | 3,35E+08 | 3,37E+08 | 111,34 | 0 | 257,92 | 202 |
| 2,30E+08 | 3,67E+08 | 3,58E+08 | 3,59E+08 | 103,49 | 0 | 211,89 | 123 |
| 6,30E+08 | 2,58E+08 | 2,70E+08 | 2,47E+08 | 82,105 | 0 | 323,31 | 214 |
| 1,82E+09 | 2,78E+09 | 2,62E+09 | 2,81E+09 | 82,669 | 0 | 323,31 | 422 |

| | | | | | | | |
|---|---|---|---|---|---|---|---|
| 7,08E+06 | 6,60E+07 | 7,68E+07 | 6,62E+07 | 69,136 | 0 | 33,264 | 22 |
| 1,48E+09 | 2,12E+09 | 2,25E+09 | 2,13E+09 | 51,386 | 0 | 259,14 | 262 |
| 5,33E+07 | 1,49E+06 | 2,00E+06 | 2,87E+06 | 56,59 | 0 | 15,03 | 17 |
| 5,27E+07 | 1,04E+08 | 1,04E+08 | 1,03E+08 | 62,132 | 0 | 36,062 | 62 |
| 3,95E+08 | 2,51E+08 | 2,41E+08 | 2,41E+08 | 245,16 | 0 | 249,79 | 239 |
| 3,98E+08 | 8,98E+05 | 1,03E+06 | 7,18E+05 | 48,064 | 0 | 98,396 | 59 |
| 4,00E+08 | 5,13E+08 | 5,11E+08 | 5,00E+08 | 112,32 | 0 | 272,5 | 253 |
| 2,16E+08 | 1,90E+06 | 1,20E+06 | 2,46E+06 | 35,908 | 0 | 141,64 | 39 |
| 7,30E+06 | 1,76E+08 | 1,73E+08 | 1,65E+08 | 25,485 | 0 | 40,667 | 18 |
| 2,94E+08 | 1,15E+08 | 1,27E+08 | 1,12E+08 | 59,084 | 0 | 169,58 | 90 |
| 9,18E+08 | 1,83E+09 | 1,73E+09 | 1,67E+09 | 95,923 | 0 | 323,31 | 346 |
| 3,60E+07 | 1,63E+07 | 1,64E+07 | 1,76E+07 | 63,301 | 0 | 50,383 | 25 |
| 3,40E+08 | 2,94E+06 | 4,96E+06 | 1,79E+06 | 81,317 | 0 | 226,21 | 70 |
| 6,22E+08 | 2,77E+08 | 2,82E+08 | 2,87E+08 | 119,27 | 0 | 209,15 | 176 |
| 8,68E+07 | 2,38E+06 | 1,70E+06 | 9,97E+05 | 170,82 | 0 | 105,76 | 60 |
| 1,80E+08 | 7,78E+07 | 7,57E+07 | 8,59E+07 | 24,715 | 0 | 92,783 | 51 |
| 1,69E+06 | 6,08E+07 | 5,65E+07 | 5,35E+07 | 122,46 | 0 | 42,119 | 28 |
| 8,98E+08 | 1,45E+09 | 1,35E+09 | 1,34E+09 | 54,866 | 0 | 323,31 | 231 |
| 2,58E+06 | 4,24E+08 | 5,37E+08 | 4,89E+08 | 21,539 | 0 | 92,827 | 52 |
| 1,54E+08 | 7,32E+07 | 6,87E+07 | 7,12E+07 | 60,629 | 0 | 72,837 | 59 |
| 3,46E+08 | 8,86E+07 | 8,60E+07 | 9,79E+07 | 33,996 | 0 | 192,66 | 62 |
| 3,78E+09 | 2,15E+09 | 2,12E+09 | 2,24E+09 | 55,814 | 0 | 323,31 | 409 |
| 1,37E+07 | 1,25E+08 | 1,18E+08 | 1,44E+08 | 46,897 | 0 | 34,091 | 25 |
| 1,37E+08 | 9,05E+07 | 8,62E+07 | 8,68E+07 | 285,66 | 0 | 58,064 | 88 |
| 1,24E+09 | 4,60E+08 | 4,68E+08 | 4,88E+08 | 69,012 | 0 | 323,31 | 164 |
| 1,91E+08 | 1,52E+07 | 1,54E+07 | 2,33E+07 | 119,48 | 0 | 103,13 | 61 |
| 3,22E+08 | 1,42E+08 | 1,39E+08 | 1,45E+08 | 171,18 | 0 | 161,21 | 170 |
| 7,21E+07 | 3,63E+07 | 3,52E+07 | 3,22E+07 | 46,451 | 0 | 28,967 | 53 |
| 1,98E+08 | 6,34E+07 | 7,98E+07 | 7,12E+07 | 79,921 | 0 | 70,2 | 104 |
| 2,45E+08 | 5,10E+08 | 4,74E+08 | 4,98E+08 | 83,594 | 0 | 237,59 | 159 |
| 5,42E+08 | 4,22E+06 | 5,02E+06 | 1,71E+06 | 63,864 | 0 | 323,31 | 92 |
| 1,57E+09 | 2,07E+09 | 2,18E+09 | 2,07E+09 | 55,249 | 0 | 323,31 | 283 |
| 4,29E+08 | 2,27E+08 | 2,15E+08 | 2,04E+08 | 139,49 | 0 | 323,31 | 179 |
| 2,97E+07 | 8,76E+07 | 8,49E+07 | 8,10E+07 | 66,654 | 0 | 27,745 | 47 |
| 7,36E+08 | 5,69E+08 | 5,61E+08 | 5,36E+08 | 199,08 | 0 | 323,31 | 361 |
| 1,45E+08 | 9,51E+07 | 9,36E+07 | 1,00E+08 | 84,877 | 0 | 124,7 | 100 |
| 1,02E+08 | 1,40E+06 | 3,25E+06 | 2,29E+06 | 48,267 | 0 | 47,989 | 25 |
| 1,53E+10 | 2,77E+10 | 2,91E+10 | 2,84E+10 | 36,498 | 0 | 323,31 | 1405 |
| 5,05E+08 | 7,40E+08 | 6,92E+08 | 6,96E+08 | 83,056 | 0 | 323,31 | 192 |
| 2,83E+07 | 4,72E+08 | 3,97E+08 | 4,00E+08 | 37,057 | 0 | 74,7 | 75 |
| 5,13E+07 | 2,94E+07 | 2,72E+07 | 3,00E+07 | 61,361 | 0 | 26,041 | 42 |
| 9,83E+08 | 1,48E+09 | 1,57E+09 | 1,64E+09 | 66,682 | 0 | 323,31 | 231 |
| 6,71E+08 | 1,11E+09 | 1,13E+09 | 1,14E+09 | 194,74 | 0 | 323,31 | 364 |
| 1,88E+08 | 3,02E+08 | 2,91E+08 | 2,82E+08 | 59,795 | 0 | 246,79 | 101 |
| 3,84E+08 | 2,45E+08 | 2,45E+08 | 2,60E+08 | 98,72 | 0 | 226,83 | 159 |
| 2,79E+07 | 2,29E+08 | 2,18E+08 | 2,29E+08 | 18,227 | 0 | 40,757 | 48 |
| 1,84E+07 | 2,86E+06 | 4,10E+06 | 3,55E+06 | 18,862 | 0,000154 | 4,2664 | 8 |
| 2,60E+07 | 1,28E+07 | 1,10E+07 | 1,12E+07 | 48,375 | 0 | 10,446 | 21 |
| 6,81E+07 | 2,07E+08 | 1,82E+08 | 1,98E+08 | 153,74 | 0 | 165,59 | 95 |
| 6,39E+08 | 4,46E+08 | 4,65E+08 | 4,36E+08 | 79,381 | 0 | 323,31 | 203 |

| | | | | | | | |
|---|---|---|---|---|---|---|---|
| 4,43E+07 | 2,68E+07 | 2,75E+07 | 2,93E+07 | 48,745 | 0 | 5,8988 | 21 |
| 3,10E+07 | 7,10E+07 | 6,98E+07 | 6,79E+07 | 81,444 | 0 | 47,821 | 48 |
| 3,41E+08 | 1,07E+08 | 1,05E+08 | 8,59E+07 | 78,025 | 0 | 323,31 | 108 |
| 1,56E+08 | 2,59E+08 | 2,64E+08 | 2,53E+08 | 101,27 | 0 | 133,61 | 102 |
| 9,12E+08 | 6,39E+08 | 6,63E+08 | 6,28E+08 | 118,26 | 0 | 323,31 | 273 |
| 3,22E+08 | 1,71E+08 | 1,83E+08 | 1,78E+08 | 33,913 | 0 | 323,31 | 64 |
| 1,43E+09 | 9,53E+08 | 9,88E+08 | 1,01E+09 | 102,53 | 0 | 323,31 | 288 |
| 9,03E+07 | 1,70E+08 | 1,71E+08 | 1,79E+08 | 57,105 | 0 | 181,17 | 82 |
| 1,70E+08 | 5,53E+08 | 4,96E+08 | 4,95E+08 | 30,276 | 0 | 109,87 | 78 |
| 2,32E+07 | 8,10E+06 | 8,96E+06 | 7,10E+06 | 65,076 | 0 | 17,745 | 16 |
| 1,47E+09 | 9,85E+08 | 9,86E+08 | 1,03E+09 | 90,875 | 0 | 323,31 | 297 |
| 9,38E+08 | 5,57E+08 | 5,58E+08 | 5,28E+08 | 36,265 | 0 | 323,31 | 150 |
| 9,48E+08 | 1,24E+09 | 1,33E+09 | 1,27E+09 | 266,8 | 0 | 323,31 | 488 |
| 5,60E+08 | 3,49E+08 | 3,74E+08 | 3,51E+08 | 87,021 | 0 | 323,31 | 133 |
| 1,97E+09 | 2,42E+09 | 2,42E+09 | 2,43E+09 | 102,71 | 0 | 323,31 | 343 |
| 1,25E+10 | 6,66E+09 | 6,37E+09 | 6,21E+09 | 36,384 | 0 | 323,31 | 480 |
| 5,27E+08 | 1,78E+08 | 1,66E+08 | 1,36E+08 | 94,946 | 0 | 323,31 | 112 |
| 1,79E+08 | 8,15E+07 | 7,75E+07 | 8,23E+07 | 28,263 | 0 | 229,75 | 22 |
| 2,77E+07 | 2,20E+06 | 1,35E+06 | 2,31E+06 | 110,58 | 0 | 8,1952 | 16 |
| 9,30E+07 | 1,86E+08 | 1,74E+08 | 1,62E+08 | 45,343 | 0 | 18,346 | 65 |
| 4,29E+09 | 6,77E+09 | 6,94E+09 | 7,39E+09 | 28,832 | 0 | 323,31 | 286 |
| 2,55E+08 | 1,28E+08 | 1,38E+08 | 1,40E+08 | 44,93 | 0 | 301,8 | 117 |
| 2,51E+08 | 2,16E+08 | 2,14E+08 | 2,16E+08 | 195,06 | 0 | 166,36 | 177 |
| 6,09E+08 | 3,26E+08 | 3,06E+08 | 3,17E+08 | 78,538 | 0 | 235,62 | 204 |
| 1,20E+07 | 5,51E+06 | 5,08E+06 | 5,03E+06 | 71,275 | 0 | 8,7117 | 16 |
| 2,91E+08 | 1,02E+08 | 1,23E+08 | 1,11E+08 | 61,378 | 0 | 125,94 | 109 |
| 2,45E+08 | 1,13E+08 | 1,16E+08 | 1,20E+08 | 91,215 | 0 | 194,38 | 125 |
| 1,02E+09 | 5,33E+08 | 5,30E+08 | 5,14E+08 | 94,138 | 0 | 323,31 | 188 |
| 1,85E+08 | 8,70E+07 | 8,99E+07 | 8,39E+07 | 30,148 | 0 | 64,359 | 45 |
| 7,38E+07 | 2,45E+08 | 2,54E+08 | 2,53E+08 | 121,47 | 0 | 120,08 | 91 |
| 3,54E+08 | 1,66E+08 | 1,40E+08 | 1,58E+08 | 152,13 | 0 | 192,09 | 115 |
| 1,62E+08 | 1,11E+08 | 1,14E+08 | 1,12E+08 | 56,985 | 0 | 102,43 | 84 |
| 5,41E+07 | 1,06E+08 | 1,04E+08 | 1,17E+08 | 221,67 | 0 | 43,934 | 89 |
| 1,43E+10 | 9,07E+09 | 8,09E+09 | 8,30E+09 | 38,734 | 0 | 323,31 | 725 |
| 1,23E+08 | 8,63E+07 | 8,28E+07 | 8,78E+07 | 68,203 | 0 | 70,429 | 62 |
| 5,72E+08 | 3,32E+08 | 2,98E+08 | 3,33E+08 | 31,381 | 0 | 175,76 | 144 |
| 7,72E+07 | 1,65E+07 | 1,71E+07 | 1,54E+07 | 32,503 | 0 | 7,0372 | 23 |
| 5,74E+07 | 4,54E+06 | 6,12E+06 | 3,39E+06 | 71,545 | 0 | 24,863 | 24 |
| 2,58E+08 | 4,82E+08 | 4,39E+08 | 4,65E+08 | 98,19 | 0 | 176,59 | 206 |
| 2,62E+08 | 2,04E+08 | 2,02E+08 | 1,97E+08 | 88,817 | 0 | 150,68 | 121 |
| 3,05E+08 | 4,41E+08 | 4,59E+08 | 4,45E+08 | 22,223 | 0 | 191,77 | 77 |
| 1,20E+07 | 6,46E+07 | 5,24E+07 | 4,99E+07 | 161,67 | 0 | 24,219 | 41 |
| 7,66E+08 | 1,02E+09 | 9,88E+08 | 1,02E+09 | 46,839 | 0 | 323,31 | 174 |
| 8,85E+07 | 6,22E+06 | 4,64E+06 | 6,79E+06 | 295,23 | 0 | 30,061 | 52 |
| 8,05E+07 | 2,51E+06 | 1,97E+06 | 4,36E+06 | 26,588 | 0 | 46,923 | 24 |
| 7,33E+08 | 9,55E+08 | 9,90E+08 | 9,76E+08 | 84,921 | 0 | 323,31 | 231 |
| 2,96E+08 | 1,29E+08 | 1,13E+08 | 1,30E+08 | 243,16 | 0 | 228,84 | 171 |
| 2,23E+08 | 4,84E+08 | 4,72E+08 | 4,94E+08 | 50,521 | 0 | 323,31 | 152 |
| 7,64E+07 | 1,70E+08 | 1,64E+08 | 1,45E+08 | 35,943 | 0 | 310,18 | 47 |
| 7,62E+08 | 4,11E+08 | 3,71E+08 | 4,25E+08 | 65,583 | 0 | 323,31 | 190 |

| | | | | | | | |
|---|---|---|---|---|---|---|---|
| 4,59E+08 | 5,97E+08 | 6,02E+08 | 5,87E+08 | 85,47 | 0 | 323,31 | 204 |
| 1,13E+07 | 2,53E+07 | 2,76E+07 | 2,50E+07 | 108,01 | 0 | 10,016 | 28 |
| 5,09E+08 | 3,31E+08 | 3,11E+08 | 3,05E+08 | 66,78 | 0 | 323,31 | 159 |
| 3,42E+08 | 1,35E+08 | 1,43E+08 | 1,34E+08 | 113,07 | 0 | 175,96 | 106 |
| 2,78E+08 | 9,12E+07 | 8,98E+07 | 1,06E+08 | 57,568 | 0 | 165,45 | 81 |
| 3,50E+08 | 4,81E+08 | 4,72E+08 | 4,94E+08 | 70,289 | 0 | 295,44 | 173 |
| 2,38E+07 | 3,87E+07 | 3,54E+07 | 3,64E+07 | 36,426 | 0 | 10,169 | 35 |
| 3,71E+08 | 5,49E+08 | 5,46E+08 | 5,27E+08 | 113,88 | 0 | 264,2 | 214 |
| 4,42E+08 | 4,53E+07 | 7,32E+07 | 5,26E+07 | 25,096 | 0 | 37,849 | 58 |
| 1,72E+09 | 1,07E+09 | 1,13E+09 | 1,13E+09 | 43,688 | 0 | 323,31 | 172 |
| 7,36E+06 | 2,13E+07 | 2,41E+07 | 1,95E+07 | 128,5 | 0,00015 | 3,6939 | 5 |
| 5,73E+07 | 2,66E+06 | 4,40E+06 | 2,60E+06 | 34,668 | 0 | 16,798 | 23 |
| 6,02E+07 | 3,28E+07 | 3,43E+07 | 3,20E+07 | 37,316 | 0 | 113,8 | 34 |
| 7,50E+07 | 1,38E+08 | 1,25E+08 | 1,38E+08 | 43,544 | 0 | 168,53 | 38 |
| 8,41E+08 | 1,36E+09 | 1,35E+09 | 1,44E+09 | 38,62 | 0 | 323,31 | 262 |
| 2,52E+06 | 1,12E+08 | 1,29E+08 | 1,34E+08 | 27,269 | 0 | 72,163 | 17 |
| 2,67E+09 | 3,97E+09 | 4,16E+09 | 4,19E+09 | 31,319 | 0 | 323,31 | 231 |
| 1,87E+09 | 1,18E+09 | 1,10E+09 | 1,08E+09 | 87,676 | 0 | 323,31 | 381 |
| 1,51E+09 | 6,42E+08 | 6,48E+08 | 5,81E+08 | 85,73 | 0 | 323,31 | 247 |
| 2,88E+08 | 6,00E+08 | 5,67E+08 | 5,51E+08 | 98,241 | 0 | 323,31 | 172 |
| 2,95E+08 | 1,85E+08 | 1,88E+08 | 1,89E+08 | 82,408 | 0 | 323,31 | 146 |
| 3,29E+07 | 1,54E+07 | 1,52E+07 | 1,35E+07 | 43,98 | 0 | 25,347 | 32 |
| 1,30E+09 | 9,31E+08 | 8,80E+08 | 8,81E+08 | 117,45 | 0 | 323,31 | 373 |
| 7,06E+07 | 1,65E+08 | 1,66E+08 | 1,63E+08 | 69,912 | 0 | 74,165 | 82 |
| 7,83E+08 | 6,02E+08 | 6,00E+08 | 6,23E+08 | 79,776 | 0 | 323,31 | 241 |
| 1,45E+08 | 2,85E+08 | 2,62E+08 | 2,83E+08 | 40,251 | 0 | 50,049 | 73 |
| 1,35E+08 | 2,00E+08 | 1,97E+08 | 1,99E+08 | 409,76 | 0 | 112,78 | 170 |
| 1,71E+08 | 3,22E+08 | 3,32E+08 | 3,10E+08 | 88,651 | 0 | 147,52 | 154 |
| 4,11E+07 | 1,64E+07 | 1,99E+07 | 1,89E+07 | 64,955 | 0 | 27,877 | 35 |
| 2,93E+07 | 3,90E+06 | 4,74E+06 | 3,41E+06 | 45,464 | 0 | 8,0767 | 10 |
| 2,32E+08 | 4,31E+07 | 2,71E+07 | 3,75E+07 | 40,1 | 0 | 56,972 | 33 |
| 2,52E+08 | 1,63E+08 | 1,51E+08 | 1,48E+08 | 132,68 | 0 | 128,65 | 112 |
| 2,98E+08 | 5,12E+08 | 5,61E+08 | 5,95E+08 | 62,63 | 0 | 162,08 | 174 |
| 4,20E+07 | 1,97E+06 | 1,15E+06 | 2,47E+06 | 59,088 | 0 | 11,857 | 12 |
| 1,28E+08 | 2,27E+08 | 2,26E+08 | 2,34E+08 | 112,41 | 0 | 62,456 | 115 |
| 5,07E+07 | 1,43E+08 | 1,41E+08 | 1,31E+08 | 78,751 | 0 | 102,89 | 63 |
| 7,90E+07 | 4,84E+07 | 4,98E+07 | 4,68E+07 | 32,032 | 0 | 30,106 | 52 |
| 3,04E+06 | 1,05E+07 | 1,46E+07 | 1,14E+07 | 25,746 | 0,002099 | 2,169 | 4 |
| 6,24E+07 | 1,30E+08 | 1,20E+08 | 1,27E+08 | 57,455 | 0 | 43,343 | 58 |
| 9,57E+06 | 1,75E+07 | 1,82E+07 | 1,69E+07 | 21,491 | 0 | 8,895 | 24 |
| 2,25E+08 | 2,20E+06 | 6,05E+06 | 2,63E+06 | 32,423 | 0 | 247,29 | 31 |
| 4,96E+07 | 1,31E+06 | 1,23E+06 | 1,85E+06 | 163,24 | 0 | 12,733 | 23 |
| 9,94E+07 | 2,62E+08 | 2,49E+08 | 2,63E+08 | 77,738 | 0 | 83,294 | 106 |
| 8,98E+07 | 6,50E+07 | 6,34E+07 | 6,36E+07 | 50,239 | 0 | 83,525 | 57 |
| 1,34E+08 | 6,33E+07 | 6,58E+07 | 5,68E+07 | 69,819 | 0 | 43,729 | 80 |
| 2,15E+08 | 2,54E+06 | 6,91E+06 | 4,08E+06 | 53,389 | 0 | 302,26 | 35 |
| 5,51E+07 | 1,24E+08 | 1,18E+08 | 1,24E+08 | 131,55 | 0 | 40,401 | 77 |
| 1,14E+08 | 1,73E+08 | 1,60E+08 | 1,61E+08 | 54,985 | 0 | 308,47 | 61 |
| 2,13E+08 | 5,71E+08 | 5,40E+08 | 5,71E+08 | 51,24 | 0 | 240,63 | 119 |
| 8,13E+08 | 1,22E+09 | 1,18E+09 | 1,23E+09 | 65,696 | 0 | 251,2 | 189 |

| | | | | | | | |
|---|---|---|---|---|---|---|---|
| 7,29E+07 | 4,64E+07 | 4,59E+07 | 4,51E+07 | 129,05 | 0 | 88,354 | 55 |
| 4,54E+07 | 6,89E+07 | 7,54E+07 | 7,24E+07 | 31,157 | 0 | 28,227 | 37 |
| 7,26E+09 | 4,41E+09 | 4,22E+09 | 4,12E+09 | 284,59 | 0 | 323,31 | 1309 |
| 1,96E+08 | 1,44E+08 | 1,46E+08 | 1,46E+08 | 62,384 | 0 | 66,83 | 118 |
| 2,43E+08 | 4,51E+08 | 4,26E+08 | 4,07E+08 | 23,782 | 0 | 109,61 | 72 |
| 3,18E+07 | 2,56E+06 | 1,60E+06 | 1,32E+06 | 54,386 | 0 | 11,651 | 13 |
| 1,04E+09 | 1,98E+09 | 1,86E+09 | 1,94E+09 | 66,765 | 0 | 323,31 | 267 |
| 1,50E+09 | 6,54E+08 | 5,56E+08 | 6,06E+08 | 39,735 | 0 | 323,31 | 162 |
| 5,50E+07 | 1,16E+08 | 1,18E+08 | 1,10E+08 | 101,55 | 0 | 47,991 | 79 |
| 3,83E+08 | 1,88E+08 | 1,83E+08 | 1,61E+08 | 41,297 | 0 | 128,91 | 112 |
| 5,02E+08 | 3,16E+08 | 3,10E+08 | 3,16E+08 | 108,85 | 0 | 323,31 | 199 |
| 2,65E+08 | 1,65E+08 | 1,65E+08 | 1,67E+08 | 26,925 | 0 | 32,851 | 44 |
| 9,94E+07 | 1,98E+08 | 1,85E+08 | 1,76E+08 | 77,92 | 0 | 32,024 | 88 |
| 4,51E+08 | 5,69E+08 | 5,76E+08 | 5,75E+08 | 66,704 | 0 | 247,66 | 193 |
| 1,16E+08 | 6,15E+07 | 6,67E+07 | 5,87E+07 | 74,198 | 0 | 70,107 | 64 |
| 2,18E+08 | 3,74E+08 | 4,32E+08 | 3,83E+08 | 70,264 | 0 | 165,39 | 118 |
| 4,45E+08 | 6,85E+08 | 6,92E+08 | 7,22E+08 | 69,483 | 0 | 133,64 | 200 |
| 1,28E+08 | 3,18E+06 | 1,06E+06 | 2,08E+06 | 39,394 | 0 | 72,652 | 28 |
| 7,61E+07 | 1,62E+06 | 3,53E+06 | 3,38E+06 | 94,654 | 0 | 29,083 | 35 |
| 4,73E+08 | 2,58E+08 | 2,26E+08 | 2,61E+08 | 212,73 | 0 | 323,31 | 219 |
| 2,60E+07 | 7,82E+07 | 7,63E+07 | 6,82E+07 | 45,464 | 0 | 43,857 | 25 |
| 1,49E+08 | 7,91E+07 | 8,11E+07 | 8,98E+07 | 89,25 | 0 | 76,176 | 64 |
| 2,64E+07 | 8,41E+06 | 7,39E+06 | 8,55E+06 | 184,15 | 0 | 16,894 | 23 |
| 4,73E+07 | 3,45E+05 | 5,78E+05 | 1,09E+06 | 109,15 | 0 | 67,686 | 32 |
| 1,89E+06 | 1,56E+07 | 1,50E+07 | 1,99E+07 | 66,801 | 0,000153 | 4,153 | 11 |
| 1,59E+10 | 9,46E+09 | 1,07E+10 | 1,00E+10 | 59,752 | 0 | 323,31 | 687 |
| 2,60E+08 | 1,15E+08 | 1,06E+08 | 1,15E+08 | 118,87 | 0 | 149,12 | 110 |
| 5,37E+08 | 3,17E+08 | 3,53E+08 | 3,47E+08 | 269,8 | 0 | 196,84 | 248 |
| 1,05E+08 | 6,58E+07 | 7,06E+07 | 7,21E+07 | 38,604 | 0 | 79,709 | 36 |
| 3,22E+09 | 1,43E+09 | 1,40E+09 | 1,36E+09 | 116,74 | 0 | 323,31 | 402 |
| 7,43E+08 | 4,42E+08 | 4,81E+08 | 4,83E+08 | 80,428 | 0 | 277,15 | 200 |
| 8,42E+07 | 1,09E+08 | 1,10E+08 | 1,16E+08 | 103,96 | 0 | 27,88 | 78 |
| 6,18E+08 | 3,41E+08 | 3,40E+08 | 3,55E+08 | 76,205 | 0 | 180,74 | 159 |
| 4,44E+08 | 2,53E+08 | 2,75E+08 | 2,77E+08 | 89,413 | 0 | 276,77 | 154 |
| 5,46E+07 | 8,77E+07 | 8,16E+07 | 8,05E+07 | 42,189 | 0 | 37,817 | 68 |
| 2,89E+08 | 1,95E+08 | 1,78E+08 | 1,95E+08 | 65,853 | 0 | 223,7 | 99 |
| 5,10E+07 | 1,07E+08 | 1,19E+08 | 1,10E+08 | 59,735 | 0 | 41,817 | 46 |
| 7,66E+07 | 4,37E+07 | 4,44E+07 | 4,07E+07 | 57,143 | 0 | 40,824 | 49 |
| 1,91E+08 | 1,15E+08 | 1,23E+08 | 1,22E+08 | 214,43 | 0 | 89,848 | 171 |
| 7,22E+07 | 1,21E+08 | 1,20E+08 | 1,20E+08 | 47,369 | 0 | 24,768 | 26 |
| 1,59E+08 | 5,56E+07 | 5,82E+07 | 5,42E+07 | 62,586 | 0 | 89,953 | 69 |
| 8,87E+07 | 3,72E+06 | 5,79E+06 | 2,90E+06 | 35,019 | 0 | 51,041 | 8 |
| 6,01E+07 | 1,85E+07 | 2,39E+07 | 2,14E+07 | 29,063 | 0 | 68,925 | 31 |
| 1,46E+08 | 1,29E+06 | 2,37E+06 | 3,81E+06 | 57,187 | 0 | 64,618 | 58 |
| 2,63E+08 | 4,17E+08 | 4,25E+08 | 4,13E+08 | 211,19 | 0 | 235,03 | 181 |
| 9,87E+08 | 7,97E+08 | 7,78E+08 | 7,59E+08 | 112,8 | 0 | 323,31 | 320 |
| 5,08E+08 | 1,09E+09 | 1,15E+09 | 1,14E+09 | 104,02 | 0 | 323,31 | 232 |
| 2,58E+08 | 8,37E+07 | 9,48E+07 | 9,34E+07 | 80,614 | 0 | 73,816 | 80 |
| 9,46E+08 | 4,12E+08 | 4,77E+08 | 4,74E+08 | 57,93 | 0 | 323,31 | 204 |
| 1,50E+08 | 2,20E+08 | 2,40E+08 | 2,25E+08 | 359,4 | 0 | 103,09 | 188 |

| | | | | | | | |
|---|---|---|---|---|---|---|---|
| 1,08E+09 | 1,38E+09 | 1,41E+09 | 1,44E+09 | 102,1 | 0 | 323,31 | 352 |
| 1,34E+08 | 1,90E+08 | 1,85E+08 | 1,94E+08 | 86,454 | 0 | 67,016 | 102 |
| 3,42E+07 | 1,10E+06 | 1,56E+06 | 7,35E+05 | 60,163 | 0 | 18,991 | 26 |
| 2,49E+09 | 3,05E+09 | 3,17E+09 | 3,17E+09 | 102,08 | 0 | 323,31 | 556 |
| 9,57E+08 | 5,81E+08 | 5,85E+08 | 5,57E+08 | 59,795 | 0 | 323,31 | 195 |
| 7,44E+08 | 1,85E+06 | 2,32E+06 | 7,22E+06 | 80,253 | 0 | 323,31 | 111 |
| 3,02E+09 | 8,87E+08 | 7,15E+08 | 6,24E+08 | 62,747 | 0 | 323,31 | 215 |
| 1,31E+08 | 1,15E+07 | 1,34E+07 | 6,92E+06 | 64,061 | 0 | 61,605 | 46 |
| 1,09E+08 | 1,58E+08 | 1,70E+08 | 1,56E+08 | 133,37 | 0 | 108,66 | 113 |
| 9,11E+07 | 2,24E+08 | 1,98E+08 | 2,14E+08 | 30,534 | 0 | 34,327 | 38 |
| 6,60E+07 | 4,16E+07 | 3,72E+07 | 3,83E+07 | 77,594 | 0 | 31,368 | 51 |
| 5,25E+07 | 1,12E+08 | 1,13E+08 | 1,01E+08 | 198,95 | 0 | 36,25 | 94 |
| 4,69E+06 | 2,17E+08 | 2,24E+08 | 2,28E+08 | 128,36 | 0 | 143,62 | 69 |
| 4,54E+07 | 4,37E+06 | 2,14E+06 | 2,61E+06 | 93,481 | 0 | 25,993 | 19 |
| 1,97E+09 | 1,51E+09 | 1,50E+09 | 1,55E+09 | 341,12 | 0 | 323,31 | 651 |
| 6,13E+08 | 7,30E+08 | 7,33E+08 | 7,53E+08 | 66,741 | 0 | 323,31 | 202 |
| 2,10E+09 | 3,22E+09 | 3,05E+09 | 2,98E+09 | 49,508 | 0 | 323,31 | 381 |
| 7,44E+07 | 3,20E+07 | 3,04E+07 | 3,25E+07 | 42,407 | 0 | 31,195 | 38 |
| 5,35E+07 | 3,71E+07 | 3,64E+07 | 3,82E+07 | 38,018 | 0 | 11,803 | 27 |
| 7,42E+08 | 4,51E+08 | 5,00E+08 | 4,97E+08 | 72,679 | 0 | 125,77 | 191 |
| 2,92E+07 | 6,08E+07 | 6,11E+07 | 5,78E+07 | 20,11 | 0,000155 | 4,4623 | 19 |
| 2,95E+06 | 2,56E+07 | 2,25E+07 | 2,25E+07 | 24,33 | 0 | 11,866 | 11 |
| 6,24E+06 | 1,51E+08 | 1,65E+08 | 1,83E+08 | 520,68 | 0 | 60,46 | 119 |
| 1,23E+08 | 1,91E+08 | 1,86E+08 | 1,87E+08 | 54,043 | 0 | 111,71 | 53 |
| 3,92E+06 | 3,37E+07 | 3,50E+07 | 4,48E+07 | 101,68 | 0 | 9,568 | 20 |
| 1,68E+08 | 9,11E+07 | 9,53E+07 | 9,59E+07 | 71,558 | 0 | 42,347 | 91 |
| 2,10E+08 | 4,74E+08 | 4,37E+08 | 4,11E+08 | 46,167 | 0 | 323,31 | 91 |
| 7,25E+08 | 4,63E+08 | 4,81E+08 | 4,97E+08 | 106 | 0 | 121,52 | 191 |
| 1,52E+08 | 8,53E+07 | 9,17E+07 | 8,93E+07 | 73,704 | 0 | 27,581 | 76 |
| 3,39E+08 | 5,09E+06 | 4,01E+06 | 5,92E+06 | 35,194 | 0 | 49,685 | 32 |
| 1,05E+08 | 1,54E+08 | 1,59E+08 | 1,62E+08 | 105,97 | 0 | 81,908 | 106 |
| 1,61E+07 | 4,37E+08 | 4,26E+08 | 4,81E+08 | 68,587 | 0 | 168,52 | 94 |
| 1,05E+09 | 1,52E+09 | 1,53E+09 | 1,55E+09 | 70,568 | 0 | 323,31 | 295 |
| 4,53E+07 | 3,61E+06 | 2,64E+06 | 3,92E+06 | 71,194 | 0 | 57,122 | 26 |
| 4,81E+07 | 1,34E+08 | 1,43E+08 | 1,53E+08 | 24,275 | 0 | 177,77 | 42 |
| 2,57E+09 | 2,06E+09 | 1,94E+09 | 2,04E+09 | 183,19 | 0 | 323,31 | 621 |
| 3,59E+08 | 9,93E+08 | 9,72E+08 | 9,70E+08 | 21,394 | 0 | 323,31 | 178 |
| 8,50E+08 | 6,51E+08 | 6,16E+08 | 6,06E+08 | 94,475 | 0 | 294,96 | 274 |
| 6,46E+08 | 4,66E+08 | 4,90E+08 | 4,66E+08 | 52,471 | 0 | 277,17 | 160 |
| 1,06E+07 | 1,53E+08 | 1,20E+08 | 1,38E+08 | 214,06 | 0 | 81,573 | 68 |
| 1,51E+08 | 2,43E+08 | 2,46E+08 | 2,74E+08 | 55,055 | 0 | 128,68 | 46 |
| 6,74E+07 | 3,28E+07 | 3,05E+07 | 3,65E+07 | 67,843 | 0 | 19,535 | 32 |
| 1,44E+08 | 2,86E+08 | 3,48E+08 | 3,01E+08 | 57,488 | 0 | 323,31 | 79 |
| 9,17E+08 | 4,24E+08 | 4,56E+08 | 5,02E+08 | 40,581 | 0 | 323,31 | 102 |
| 3,13E+07 | 1,05E+07 | 1,03E+07 | 1,30E+07 | 23,062 | 0 | 47,188 | 30 |
| 8,00E+07 | 1,43E+08 | 1,34E+08 | 1,27E+08 | 59,846 | 0 | 93,684 | 80 |
| 1,91E+06 | 9,84E+07 | 1,13E+08 | 9,98E+07 | 42,791 | 0 | 36,058 | 30 |
| 4,94E+08 | 6,32E+08 | 6,30E+08 | 6,38E+08 | 52,511 | 0 | 323,31 | 165 |
| 6,88E+07 | 3,94E+07 | 4,04E+07 | 4,44E+07 | 63,393 | 0 | 13,254 | 37 |
| 1,98E+09 | 1,80E+09 | 1,83E+09 | 1,83E+09 | 26,372 | 0 | 323,31 | 123 |

| | | | | | | | |
|---|---|---|---|---|---|---|---|
| 1,82E+08 | 1,42E+07 | 2,43E+07 | 1,57E+07 | 202,37 | 0 | 112,76 | 94 |
| 3,14E+09 | 1,94E+09 | 2,01E+09 | 2,02E+09 | 273,99 | 0 | 323,31 | 722 |
| 4,73E+06 | 1,11E+08 | 8,36E+07 | 1,10E+08 | 46,203 | 0 | 63,275 | 25 |
| 1,71E+08 | 1,39E+08 | 1,42E+08 | 1,38E+08 | 79,512 | 0 | 113,79 | 103 |
| 3,55E+07 | 6,45E+07 | 7,36E+07 | 6,59E+07 | 33,574 | 0 | 42,822 | 35 |
| 3,07E+08 | 1,51E+08 | 1,54E+08 | 1,62E+08 | 160,12 | 0 | 323,31 | 189 |
| 2,39E+08 | 6,74E+08 | 7,16E+08 | 7,15E+08 | 25,716 | 0 | 188,27 | 90 |
| 4,66E+07 | 6,66E+07 | 7,14E+07 | 6,92E+07 | 45,521 | 0 | 22,987 | 64 |
| 6,11E+07 | 9,46E+07 | 1,05E+08 | 1,05E+08 | 186,72 | 0 | 62,662 | 65 |
| 2,03E+06 | 4,12E+07 | 4,50E+07 | 4,62E+07 | 24,727 | 0 | 27,404 | 11 |
| 3,21E+08 | 6,99E+07 | 7,77E+07 | 8,15E+07 | 57,463 | 0 | 287,5 | 65 |
| 1,65E+08 | 2,34E+08 | 2,29E+08 | 2,26E+08 | 56,876 | 0 | 170,61 | 101 |
| 4,34E+08 | 2,47E+09 | 2,11E+09 | 2,24E+09 | 46,103 | 0 | 323,31 | 124 |
| 2,11E+08 | 6,53E+06 | 2,09E+06 | 3,49E+06 | 85,025 | 0 | 134,66 | 59 |
| 1,72E+07 | 3,25E+07 | 3,20E+07 | 3,11E+07 | 75,072 | 0 | 11,138 | 36 |
| 6,58E+07 | 4,47E+06 | 8,12E+06 | 6,76E+06 | 15,944 | 0 | 49,986 | 25 |
| 9,12E+07 | 5,64E+07 | 6,10E+07 | 6,05E+07 | 21,597 | 0 | 117,65 | 60 |
| 9,83E+06 | 2,79E+07 | 2,73E+07 | 2,99E+07 | 55,325 | 0,000159 | 5,2934 | 14 |
| 1,22E+09 | 7,20E+08 | 7,22E+08 | 7,01E+08 | 113,1 | 0 | 227,61 | 279 |
| 7,36E+08 | 1,44E+09 | 1,47E+09 | 1,46E+09 | 75,167 | 0 | 323,31 | 199 |
| 1,07E+08 | 2,67E+07 | 2,08E+07 | 1,83E+07 | 96,804 | 0 | 43,488 | 54 |
| 2,48E+08 | 5,96E+08 | 6,33E+08 | 6,21E+08 | 137,38 | 0 | 265,3 | 217 |
| 3,36E+07 | 1,71E+07 | 1,51E+07 | 1,48E+07 | 46,329 | 0,000156 | 4,7254 | 21 |
| 2,05E+08 | 1,41E+08 | 1,42E+08 | 1,40E+08 | 51,709 | 0 | 35,917 | 75 |
| 9,30E+08 | 1,64E+09 | 1,58E+09 | 1,60E+09 | 27,497 | 0 | 323,31 | 175 |
| 1,16E+07 | 3,33E+07 | 3,15E+07 | 3,49E+07 | 35,396 | 0 | 10,629 | 25 |
| 1,19E+09 | 8,21E+08 | 8,66E+08 | 8,58E+08 | 247,64 | 0 | 323,31 | 510 |
| 1,32E+08 | 6,34E+07 | 7,23E+07 | 7,31E+07 | 59,452 | 0 | 79,589 | 76 |
| 1,08E+09 | 8,32E+08 | 8,06E+08 | 8,63E+08 | 123,79 | 0 | 81,909 | 236 |
| 3,14E+07 | 5,04E+06 | 5,98E+06 | 3,75E+06 | 331,56 | 0 | 14,059 | 26 |
| 7,51E+06 | 8,71E+07 | 8,65E+07 | 8,44E+07 | 149,57 | 0 | 35,398 | 51 |
| 1,90E+08 | 1,54E+08 | 1,47E+08 | 1,56E+08 | 78,685 | 0 | 53,699 | 116 |
| 6,55E+09 | 4,92E+09 | 5,19E+09 | 5,19E+09 | 93,55 | 0 | 323,31 | 686 |
| 2,92E+08 | 1,12E+08 | 1,25E+08 | 1,22E+08 | 33,097 | 0 | 81,706 | 39 |
| 9,22E+07 | 2,03E+08 | 2,15E+08 | 2,17E+08 | 28,308 | 0 | 218,14 | 43 |
| 4,33E+07 | 6,09E+06 | 7,73E+06 | 5,21E+06 | 97,047 | 0 | 19,465 | 25 |
| 5,12E+08 | 1,48E+08 | 1,36E+08 | 1,25E+08 | 85,454 | 0 | 323,31 | 173 |
| 3,87E+07 | 6,15E+07 | 5,89E+07 | 6,26E+07 | 18,352 | 0 | 32,327 | 16 |
| 3,01E+09 | 1,65E+09 | 1,77E+09 | 1,92E+09 | 88,231 | 0 | 323,31 | 314 |
| 1,76E+08 | 1,09E+08 | 1,02E+08 | 9,96E+07 | 67,558 | 0 | 63,09 | 73 |
| 5,39E+07 | 3,84E+06 | 5,80E+06 | 5,57E+06 | 19,099 | 0 | 24,954 | 13 |
| 4,63E+06 | 1,93E+07 | 2,57E+07 | 2,76E+07 | 147,94 | 0 | 7,2077 | 9 |
| 3,86E+07 | 2,31E+06 | 1,53E+06 | 3,50E+06 | 62,623 | 0 | 29,729 | 22 |
| 1,29E+08 | 1,69E+08 | 1,74E+08 | 1,81E+08 | 208,69 | 0 | 125,52 | 166 |
| 5,22E+08 | 2,84E+08 | 3,18E+08 | 3,26E+08 | 197,35 | 0 | 232,48 | 219 |
| 1,53E+08 | 8,77E+07 | 8,46E+07 | 7,52E+07 | 93,85 | 0 | 65,862 | 81 |
| 3,61E+08 | 1,60E+08 | 1,92E+08 | 1,83E+08 | 79,343 | 0 | 85,521 | 110 |
| 1,45E+08 | 1,03E+08 | 1,09E+08 | 1,03E+08 | 79,864 | 0 | 98,797 | 83 |
| 4,63E+08 | 5,78E+08 | 5,73E+08 | 5,75E+08 | 274,21 | 0 | 323,31 | 266 |
| 3,26E+08 | 1,10E+08 | 1,09E+08 | 1,09E+08 | 102,7 | 0 | 261,12 | 124 |

| | | | | | | | |
|---|---|---|---|---|---|---|---|
| 4,38E+07 | 2,54E+07 | 2,30E+07 | 2,35E+07 | 43,803 | 0 | 6,9614 | 15 |
| 2,21E+09 | 1,73E+06 | 4,99E+06 | 6,13E+05 | 228,99 | 0 | 323,31 | 358 |
| 1,25E+09 | 7,09E+08 | 6,97E+08 | 7,65E+08 | 56,001 | 0 | 323,31 | 314 |
| 2,65E+07 | 5,02E+07 | 5,85E+07 | 5,69E+07 | 57,908 | 0 | 14,531 | 37 |
| 3,50E+08 | 4,72E+08 | 4,49E+08 | 4,37E+08 | 103,93 | 0 | 240,56 | 131 |
| 1,29E+09 | 2,70E+09 | 2,35E+09 | 2,53E+09 | 43,953 | 0 | 323,31 | 351 |
| 3,37E+08 | 7,38E+07 | 7,95E+07 | 6,52E+07 | 226,94 | 0,006081 | 1,6021 | 13 |
| 7,24E+08 | 5,89E+08 | 6,16E+08 | 5,88E+08 | 132,06 | 0 | 245,04 | 249 |
| 6,32E+09 | 5,06E+05 | 3,39E+06 | 4,49E+06 | 28,106 | 0 | 323,31 | 141 |
| 1,17E+09 | 8,19E+08 | 8,05E+08 | 7,62E+08 | 67,941 | 0 | 322,83 | 269 |
| 4,46E+07 | 8,99E+07 | 1,04E+08 | 1,01E+08 | 31,055 | 0 | 89,383 | 44 |
| 6,32E+07 | 2,26E+06 | 1,82E+06 | 1,71E+06 | 70,407 | 0 | 68,99 | 20 |
| 4,23E+07 | 1,20E+07 | 9,42E+06 | 9,73E+06 | 37,268 | 0,000158 | 5,0785 | 14 |
| 1,20E+09 | 7,84E+08 | 7,85E+08 | 7,88E+08 | 112,26 | 0 | 323,31 | 307 |
| 8,04E+07 | 4,47E+06 | 5,86E+06 | 5,52E+06 | 67,121 | 0 | 75,396 | 30 |
| 1,08E+08 | 1,53E+08 | 1,55E+08 | 1,61E+08 | 22,909 | 0 | 24,602 | 57 |
| 2,10E+07 | 4,88E+07 | 4,96E+07 | 5,45E+07 | 51,881 | 0,000151 | 3,8362 | 18 |
| 1,91E+08 | 3,18E+08 | 3,29E+08 | 2,92E+08 | 92,201 | 0 | 183,82 | 133 |
| 5,74E+07 | 9,95E+07 | 9,98E+07 | 8,81E+07 | 24,245 | 0 | 61,725 | 48 |
| 2,85E+07 | 4,82E+07 | 4,54E+07 | 5,23E+07 | 26,494 | 0 | 23,595 | 23 |
| 6,94E+07 | 7,56E+07 | 7,49E+07 | 7,53E+07 | 160,81 | 0 | 72,264 | 71 |
| 1,76E+09 | 1,46E+09 | 1,43E+09 | 1,45E+09 | 49,925 | 0 | 323,31 | 245 |
| 3,07E+08 | 1,80E+08 | 1,72E+08 | 1,67E+08 | 141,03 | 0 | 88,392 | 133 |
| 1,20E+08 | 2,60E+08 | 2,61E+08 | 2,28E+08 | 13,815 | 0 | 51,964 | 56 |
| 1,15E+07 | 3,92E+06 | 3,99E+06 | 4,15E+06 | 82,784 | 0,000429 | 2,6895 | 5 |
| 2,20E+08 | 1,54E+08 | 1,47E+08 | 1,44E+08 | 71,051 | 0 | 133,79 | 130 |
| 1,44E+08 | 6,87E+07 | 6,78E+07 | 7,01E+07 | 52,164 | 0 | 186,73 | 51 |
| 3,26E+09 | 4,25E+09 | 4,13E+09 | 4,18E+09 | 156,61 | 0 | 323,31 | 705 |
| 9,77E+07 | 5,21E+07 | 4,86E+07 | 5,60E+07 | 60,526 | 0 | 84,981 | 43 |
| 1,49E+08 | 5,79E+06 | 5,84E+06 | 5,33E+06 | 207,54 | 0 | 67,883 | 62 |
| 2,92E+07 | 5,69E+07 | 6,74E+07 | 6,21E+07 | 94,128 | 0 | 18,97 | 52 |
| 5,38E+08 | 7,52E+08 | 7,70E+08 | 7,27E+08 | 44,783 | 0 | 318,64 | 166 |
| 1,55E+08 | 2,52E+08 | 2,42E+08 | 2,63E+08 | 70,678 | 0 | 323,31 | 100 |
| 6,03E+07 | 2,50E+07 | 2,65E+07 | 2,31E+07 | 46,58 | 0 | 23,279 | 39 |
| 2,73E+09 | 3,93E+09 | 3,86E+09 | 3,96E+09 | 116,72 | 0 | 323,31 | 754 |
| 3,93E+07 | 1,29E+06 | 1,37E+06 | 2,94E+06 | 31,083 | 0 | 12,563 | 16 |
| 2,03E+09 | 2,79E+09 | 2,82E+09 | 2,72E+09 | 48,354 | 0 | 323,31 | 288 |
| 2,15E+09 | 5,15E+09 | 5,46E+09 | 4,74E+09 | 54,831 | 0 | 323,31 | 322 |
| 1,30E+08 | 6,43E+07 | 6,31E+07 | 6,60E+07 | 17,09 | 0 | 174,27 | 59 |
| 3,11E+08 | 1,61E+08 | 1,62E+08 | 1,64E+08 | 74,318 | 0 | 223,09 | 116 |
| 7,53E+08 | 6,18E+08 | 5,97E+08 | 6,29E+08 | 308,35 | 0 | 323,31 | 397 |
| 1,74E+08 | 8,44E+07 | 8,44E+07 | 8,17E+07 | 124,49 | 0 | 142,85 | 90 |
| 1,67E+08 | 1,31E+08 | 1,33E+08 | 1,34E+08 | 194,44 | 0 | 95,943 | 142 |
| 6,79E+07 | 1,53E+08 | 1,54E+08 | 1,29E+08 | 38,167 | 0 | 61,486 | 60 |
| 7,26E+07 | 1,30E+08 | 1,26E+08 | 1,37E+08 | 41,669 | 0 | 81,665 | 51 |
| 7,16E+07 | 9,13E+07 | 9,75E+07 | 9,22E+07 | 152,01 | 0 | 91,767 | 76 |
| 2,59E+06 | 1,25E+07 | 8,31E+06 | 1,09E+07 | 27,902 | 0,000148 | 3,3459 | 6 |
| 8,98E+08 | 4,35E+08 | 4,71E+08 | 5,03E+08 | 80,597 | 0 | 57,765 | 103 |
| 1,35E+09 | 8,33E+08 | 8,08E+08 | 7,98E+08 | 38,988 | 0 | 323,31 | 236 |
| 6,46E+07 | 1,46E+08 | 1,28E+08 | 1,49E+08 | 34,975 | 0 | 85,589 | 59 |

| | | | | | | | |
|---|---|---|---|---|---|---|---|
| 1,25E+09 | 8,94E+08 | 9,02E+08 | 8,99E+08 | 117,64 | 0 | 323,31 | 368 |
| 2,05E+07 | 3,21E+06 | 2,57E+06 | 2,72E+06 | 56,331 | 0 | 14,45 | 17 |
| 1,75E+07 | 1,32E+08 | 1,01E+08 | 1,10E+08 | 303,97 | 0 | 80,899 | 88 |
| 1,25E+09 | 7,66E+08 | 7,60E+08 | 8,22E+08 | 50,113 | 0 | 252,46 | 182 |
| 4,22E+08 | 3,31E+08 | 3,28E+08 | 3,48E+08 | 100,31 | 0 | 308,77 | 172 |
| 2,35E+08 | 1,19E+08 | 1,16E+08 | 1,09E+08 | 73,231 | 0 | 307,48 | 107 |
| 1,63E+09 | 2,84E+09 | 2,47E+09 | 2,57E+09 | 85,941 | 0 | 323,31 | 266 |
| 6,80E+08 | 3,46E+08 | 3,15E+08 | 3,38E+08 | 69,622 | 0 | 314,79 | 193 |
| 5,45E+08 | 3,81E+08 | 3,83E+08 | 3,72E+08 | 49,74 | 0 | 323,31 | 121 |
| 5,09E+07 | 2,22E+07 | 2,55E+07 | 2,24E+07 | 112,06 | 0 | 41,427 | 47 |
| 3,27E+10 | 4,87E+10 | 5,07E+10 | 5,24E+10 | 57,844 | 0 | 323,31 | 2171 |
| 5,54E+07 | 1,59E+06 | 1,37E+06 | 3,46E+06 | 62,535 | 0 | 20,241 | 22 |
| 6,54E+07 | 3,77E+07 | 4,10E+07 | 4,00E+07 | 31,716 | 0 | 13,725 | 31 |
| 1,76E+08 | 1,27E+08 | 1,26E+08 | 1,35E+08 | 46,486 | 0 | 20,984 | 77 |
| 1,56E+08 | 2,08E+08 | 2,17E+08 | 2,20E+08 | 70,641 | 0 | 244,57 | 104 |
| 1,49E+08 | 3,12E+08 | 2,71E+08 | 2,84E+08 | 34,54 | 0 | 90,878 | 83 |
| 2,44E+08 | 1,27E+08 | 1,28E+08 | 1,19E+08 | 29,82 | 0 | 308,79 | 46 |
| 2,30E+09 | 1,77E+10 | 1,70E+10 | 1,40E+10 | 11,721 | 0 | 287,68 | 635 |
| 9,07E+07 | 4,69E+06 | 3,26E+06 | 3,22E+06 | 55,357 | 0 | 28,688 | 26 |
| 1,17E+08 | 9,20E+07 | 9,70E+07 | 9,36E+07 | 114,32 | 0 | 44,254 | 111 |
| 8,24E+07 | 1,09E+08 | 1,09E+08 | 1,04E+08 | 163,16 | 0 | 78,149 | 99 |
| 5,04E+06 | 7,86E+06 | 7,05E+06 | 7,41E+06 | 96,754 | 0,001272 | 2,4295 | 10 |
| 7,76E+07 | 1,36E+08 | 1,50E+08 | 1,65E+08 | 60,112 | 0 | 42,434 | 32 |
| 2,69E+07 | 1,10E+08 | 9,30E+07 | 1,07E+08 | 36,276 | 0 | 53,03 | 45 |
| 5,55E+07 | 2,00E+07 | 1,82E+07 | 2,46E+07 | 13,667 | 0 | 9,8463 | 22 |
| 3,78E+08 | 2,13E+08 | 1,84E+08 | 2,03E+08 | 36,09 | 0 | 60,476 | 98 |
| 3,49E+08 | 5,89E+08 | 6,08E+08 | 5,70E+08 | 140,83 | 0 | 313,9 | 197 |
| 2,48E+06 | 3,73E+08 | 3,90E+08 | 3,54E+08 | 99,795 | 0 | 323,31 | 66 |
| 5,04E+08 | 7,11E+08 | 6,78E+08 | 6,59E+08 | 53,97 | 0 | 156,06 | 170 |
| 1,60E+08 | 3,96E+06 | 1,41E+06 | 4,25E+06 | 25,061 | 0 | 68,13 | 33 |
| 1,46E+08 | 3,01E+06 | 6,97E+06 | 2,22E+06 | 58,812 | 0 | 87,996 | 51 |
| 4,18E+08 | 2,47E+08 | 2,45E+08 | 2,60E+08 | 116,97 | 0 | 130,79 | 191 |
| 7,25E+06 | 1,83E+06 | 1,85E+06 | 1,78E+06 | 86,075 | 0 | 8,4588 | 4 |
| 2,02E+08 | 3,60E+08 | 3,44E+08 | 3,73E+08 | 17,008 | 0 | 106,23 | 45 |
| 2,28E+08 | 1,58E+08 | 1,43E+08 | 1,43E+08 | 63,928 | 0 | 104,38 | 146 |
| 2,03E+08 | 3,31E+08 | 3,47E+08 | 3,74E+08 | 420,08 | 0 | 216,37 | 271 |
| 4,53E+07 | 2,35E+07 | 2,49E+07 | 2,64E+07 | 219,88 | 0 | 33,764 | 55 |
| 8,59E+07 | 4,14E+07 | 4,75E+07 | 4,69E+07 | 42,071 | 0 | 23,857 | 43 |
| 1,23E+08 | 1,94E+08 | 1,85E+08 | 1,72E+08 | 49,935 | 0 | 29,776 | 77 |
| 1,04E+07 | 3,91E+07 | 5,20E+07 | 4,19E+07 | 74,274 | 0 | 17,582 | 25 |
| 1,49E+08 | 9,37E+07 | 9,16E+07 | 8,78E+07 | 14,098 | 0 | 60,629 | 52 |
| 2,86E+09 | 1,86E+09 | 1,82E+09 | 1,78E+09 | 294,84 | 0 | 323,31 | 580 |
| 6,48E+07 | 1,96E+07 | 2,47E+07 | 2,17E+07 | 40,178 | 0 | 20,23 | 29 |
| 4,04E+08 | 8,84E+08 | 9,06E+08 | 9,12E+08 | 17,274 | 0 | 52,563 | 82 |
| 8,80E+07 | 1,56E+08 | 1,56E+08 | 1,56E+08 | 347,86 | 0 | 81,912 | 128 |
| 2,08E+07 | 6,57E+06 | 7,96E+06 | 5,90E+06 | 93,572 | 0 | 9,4964 | 17 |
| 2,22E+08 | 3,34E+08 | 3,14E+08 | 3,01E+08 | 49,295 | 0 | 46,79 | 121 |
| 3,23E+08 | 1,12E+08 | 1,13E+08 | 9,30E+07 | 43,004 | 0 | 313,05 | 81 |
| 3,10E+07 | 1,57E+07 | 1,65E+07 | 1,86E+07 | 147,64 | 0 | 20,05 | 31 |
| 3,30E+08 | 1,87E+08 | 1,98E+08 | 1,92E+08 | 33,196 | 0 | 78,841 | 93 |

| | | | | | | | |
|---|---|---|---|---|---|---|---|
| 6,69E+08 | 2,27E+07 | 4,42E+07 | 5,33E+07 | 113,58 | 0 | 243,78 | 125 |
| 1,11E+07 | 4,32E+06 | 3,83E+06 | 3,82E+06 | 92,555 | 0 | 9,9809 | 15 |
| 1,90E+08 | 2,30E+08 | 2,25E+08 | 2,27E+08 | 92,023 | 0 | 50,144 | 112 |
| 1,47E+08 | 2,35E+08 | 2,18E+08 | 2,44E+08 | 27,385 | 0 | 152,46 | 46 |
| 9,87E+07 | 1,93E+08 | 1,94E+08 | 1,70E+08 | 21,536 | 0 | 200,52 | 51 |
| 1,23E+09 | 5,67E+08 | 5,32E+08 | 6,06E+08 | 47,024 | 0 | 323,31 | 190 |
| 9,76E+09 | 1,41E+10 | 1,49E+10 | 1,50E+10 | 22,176 | 0 | 305,8 | 618 |
| 1,10E+08 | 6,84E+07 | 7,72E+07 | 7,10E+07 | 36,3 | 0 | 50,512 | 58 |
| 1,20E+08 | 1,96E+08 | 1,77E+08 | 1,73E+08 | 50,192 | 0 | 41,954 | 41 |
| 4,65E+07 | 2,71E+07 | 2,61E+07 | 2,83E+07 | 59,854 | 0 | 15,333 | 35 |
| 5,27E+07 | 2,20E+07 | 1,85E+07 | 1,79E+07 | 43,144 | 0 | 14,378 | 35 |
| 6,01E+07 | 3,72E+06 | 7,94E+06 | 4,16E+06 | 118,61 | 0 | 65,615 | 23 |
| 5,27E+07 | 8,14E+06 | 1,08E+07 | 6,07E+06 | 33,676 | 0 | 21,383 | 25 |
| 1,24E+09 | 1,99E+09 | 2,36E+09 | 2,18E+09 | 20,861 | 0 | 67,318 | 141 |
| 2,31E+09 | 2,95E+09 | 2,79E+09 | 3,00E+09 | 53,247 | 0 | 323,31 | 277 |
| 4,38E+07 | 4,62E+08 | 3,51E+08 | 4,10E+08 | 105,62 | 0 | 168,83 | 95 |
| 3,55E+09 | 6,00E+09 | 6,10E+09 | 5,94E+09 | 291,12 | 0 | 323,31 | 885 |
| 3,12E+08 | 1,49E+08 | 1,66E+08 | 1,76E+08 | 23,473 | 0 | 40,818 | 79 |
| 5,52E+08 | 3,78E+08 | 4,17E+08 | 3,84E+08 | 64,469 | 0 | 323,31 | 123 |
| 4,05E+08 | 2,45E+08 | 2,50E+08 | 2,30E+08 | 16,665 | 0 | 234,3 | 100 |
| 3,02E+08 | 2,55E+08 | 2,48E+08 | 2,54E+08 | 315,45 | 0 | 199,28 | 208 |
| 1,55E+09 | 2,49E+09 | 2,34E+09 | 2,45E+09 | 82,499 | 0 | 323,31 | 417 |
| 5,30E+07 | 2,93E+07 | 3,05E+07 | 2,83E+07 | 83,94 | 0 | 37,661 | 38 |
| 5,52E+07 | 9,70E+07 | 8,54E+07 | 8,89E+07 | 59,05 | 0 | 33,978 | 46 |
| 1,42E+07 | 2,52E+07 | 2,39E+07 | 2,55E+07 | 94,578 | 0 | 10,445 | 25 |
| 7,10E+06 | 2,59E+07 | 3,61E+07 | 3,47E+07 | 192,79 | 0 | 6,0393 | 13 |
| 2,16E+07 | 4,41E+06 | 2,37E+06 | 3,29E+06 | 198,51 | 0 | 13,613 | 19 |
| 2,13E+08 | 1,19E+08 | 1,27E+08 | 1,09E+08 | 18,382 | 0 | 11,958 | 46 |
| 1,90E+09 | 2,95E+09 | 2,67E+09 | 2,64E+09 | 197,09 | 0 | 323,31 | 463 |
| 5,50E+08 | 7,76E+08 | 7,79E+08 | 7,97E+08 | 139,52 | 0 | 323,31 | 280 |
| 1,35E+08 | 3,07E+07 | 2,25E+07 | 2,10E+07 | 52,284 | 0 | 20,726 | 31 |
| 1,61E+08 | 8,57E+08 | 8,74E+08 | 6,47E+08 | 192,99 | 0 | 323,31 | 83 |
| 5,61E+08 | 3,39E+08 | 3,25E+08 | 3,08E+08 | 31,814 | 0 | 110 | 101 |
| 5,08E+07 | 8,45E+07 | 9,47E+07 | 9,46E+07 | 63,931 | 0,000993 | 2,5045 | 18 |
| 2,10E+06 | 3,25E+07 | 2,15E+07 | 2,25E+07 | 55,913 | 0 | 6,5606 | 18 |
| 4,50E+08 | 2,99E+08 | 3,08E+08 | 2,80E+08 | 68,489 | 0 | 269,46 | 142 |
| 3,28E+08 | 2,30E+08 | 2,36E+08 | 2,39E+08 | 67,473 | 0 | 281,14 | 108 |
| 1,75E+08 | 1,27E+08 | 1,23E+08 | 1,32E+08 | 290,84 | 0 | 100,88 | 114 |
| 6,68E+07 | 2,26E+06 | 1,57E+06 | 4,42E+06 | 37,824 | 0,000156 | 4,7982 | 11 |
| 2,33E+08 | 2,08E+06 | 5,54E+06 | 6,70E+06 | 22,851 | 0 | 61,241 | 34 |
| 1,03E+08 | 1,33E+06 | 4,65E+06 | 1,93E+06 | 73,823 | 0 | 69,389 | 42 |
| 5,14E+08 | 7,67E+08 | 8,35E+08 | 7,46E+08 | 46,975 | 0 | 73,572 | 84 |
| 1,68E+08 | 8,79E+07 | 8,90E+07 | 7,64E+07 | 129,98 | 0 | 119,46 | 103 |
| 1,50E+08 | 4,99E+07 | 5,99E+07 | 6,51E+07 | 47,953 | 0 | 159,39 | 52 |
| 1,53E+08 | 4,43E+07 | 2,95E+07 | 3,03E+07 | 70,146 | 0 | 14,766 | 46 |
| 9,62E+07 | 1,70E+08 | 1,65E+08 | 1,60E+08 | 211,1 | 0 | 85,774 | 133 |
| 1,34E+09 | 2,22E+09 | 2,14E+09 | 2,18E+09 | 27,623 | 0 | 323,31 | 224 |
| 2,27E+09 | 1,85E+09 | 1,78E+09 | 1,84E+09 | 54,079 | 0 | 323,31 | 307 |
| 4,50E+08 | 7,14E+08 | 6,95E+08 | 6,73E+08 | 110,78 | 0 | 323,31 | 184 |
| 1,15E+09 | 7,24E+08 | 7,35E+08 | 7,68E+08 | 265,65 | 0 | 323,31 | 348 |

| | | | | | | | |
|---|---|---|---|---|---|---|---|
| 6,52E+07 | 1,52E+06 | 3,47E+06 | 3,46E+06 | 301,92 | 0 | 74,494 | 56 |
| 2,12E+08 | 4,13E+06 | 1,27E+06 | 4,76E+06 | 81,594 | 0 | 48,461 | 51 |
| 2,65E+09 | 1,72E+10 | 1,77E+10 | 2,04E+10 | 69,293 | 0 | 323,31 | 765 |
| 4,12E+06 | 2,56E+07 | 1,96E+07 | 2,23E+07 | 15,394 | 0 | 6,3287 | 9 |
| 3,18E+09 | 1,89E+09 | 2,17E+09 | 2,10E+09 | 50,421 | 0 | 323,31 | 333 |
| 1,23E+08 | 9,25E+07 | 9,16E+07 | 9,11E+07 | 24,944 | 0 | 24,756 | 57 |
| 7,30E+09 | 4,92E+09 | 5,20E+09 | 4,87E+09 | 56,537 | 0 | 323,31 | 544 |
| 3,17E+06 | 1,51E+07 | 1,62E+07 | 1,62E+07 | 196,58 | 0 | 8,3573 | 18 |
| 8,54E+07 | 2,32E+07 | 2,21E+07 | 1,64E+07 | 100,81 | 0 | 51,645 | 42 |
| 2,74E+08 | 3,12E+08 | 3,26E+08 | 3,26E+08 | 72,241 | 0 | 248,77 | 145 |
| 6,20E+07 | 7,83E+07 | 7,82E+07 | 8,28E+07 | 56,026 | 0 | 25,108 | 45 |
| 4,35E+06 | 1,47E+07 | 1,73E+07 | 1,57E+07 | 69,561 | 0,000434 | 2,9063 | 11 |
| 3,11E+08 | 1,88E+08 | 1,63E+08 | 1,90E+08 | 93,275 | 0 | 168,25 | 156 |
| 1,37E+09 | 1,16E+09 | 1,18E+09 | 1,21E+09 | 122,79 | 0 | 323,31 | 322 |
| 2,91E+08 | 2,42E+08 | 2,28E+08 | 2,42E+08 | 24,634 | 0 | 323,31 | 121 |
| 7,20E+06 | 3,00E+07 | 2,51E+07 | 3,02E+07 | 32,001 | 0 | 20,319 | 19 |
| 1,04E+10 | 1,24E+10 | 1,27E+10 | 1,21E+10 | 233,32 | 0 | 323,31 | 1758 |
| 6,73E+07 | 1,10E+08 | 1,04E+08 | 1,03E+08 | 39,414 | 0 | 105,9 | 41 |
| 3,18E+07 | 4,75E+06 | 2,43E+06 | 3,62E+06 | 44,901 | 0,000157 | 4,8238 | 8 |
| 1,36E+08 | 9,60E+07 | 8,96E+07 | 8,35E+07 | 121,02 | 0 | 213,84 | 104 |
| 3,17E+08 | 2,25E+08 | 2,34E+08 | 2,39E+08 | 49,593 | 0 | 149,28 | 121 |
| 4,51E+08 | 7,25E+08 | 6,92E+08 | 6,76E+08 | 37,67 | 0 | 65,494 | 130 |
| 2,20E+08 | 1,23E+08 | 1,34E+08 | 1,19E+08 | 46,299 | 0 | 292,17 | 83 |
| 5,23E+08 | 2,98E+08 | 3,17E+08 | 3,14E+08 | 19,023 | 0 | 323,31 | 61 |
| 3,83E+08 | 2,66E+08 | 2,67E+08 | 2,53E+08 | 60,545 | 0 | 158,45 | 120 |
| 3,77E+08 | 1,57E+08 | 1,71E+08 | 1,70E+08 | 35,765 | 0 | 323,31 | 84 |
| 1,53E+08 | 2,31E+08 | 2,21E+08 | 2,30E+08 | 32,928 | 0 | 102,46 | 95 |
| 3,49E+08 | 7,26E+08 | 7,05E+08 | 7,43E+08 | 109,73 | 0 | 272,3 | 208 |
| 8,58E+06 | 3,70E+07 | 3,39E+07 | 4,23E+07 | 123,89 | 0 | 21,379 | 26 |
| 1,25E+08 | 2,40E+08 | 2,72E+08 | 2,82E+08 | 42,432 | 0 | 184,34 | 62 |
| 9,02E+07 | 1,03E+07 | 6,89E+06 | 5,16E+06 | 58,843 | 0 | 30,267 | 43 |
| 5,95E+07 | 1,63E+08 | 1,46E+08 | 1,53E+08 | 58,244 | 0 | 254,05 | 90 |
| 7,91E+07 | 1,32E+08 | 1,37E+08 | 1,30E+08 | 11,241 | 0 | 41,049 | 37 |
| 6,92E+07 | 9,71E+07 | 9,41E+07 | 9,88E+07 | 60,28 | 0 | 90,668 | 53 |
| 2,68E+06 | 1,70E+07 | 1,69E+07 | 1,81E+07 | 73,148 | 0 | 9,4386 | 18 |
| 1,96E+07 | 4,96E+07 | 4,39E+07 | 4,00E+07 | 53,656 | 0 | 22,158 | 29 |
| 1,85E+08 | 9,49E+07 | 8,18E+07 | 8,18E+07 | 115,91 | 0 | 72,714 | 102 |
| 5,81E+08 | 6,50E+08 | 6,62E+08 | 6,54E+08 | 191,09 | 0 | 323,31 | 279 |
| 1,53E+09 | 7,33E+08 | 7,32E+08 | 7,02E+08 | 95,531 | 0 | 323,31 | 263 |
| 6,68E+09 | 9,55E+05 | 3,90E+06 | 4,21E+06 | 50,708 | 0 | 323,31 | 188 |
| 5,75E+07 | 9,65E+07 | 9,40E+07 | 9,29E+07 | 124,2 | 0 | 98,551 | 64 |
| 5,06E+08 | 4,10E+08 | 3,88E+08 | 4,06E+08 | 40,965 | 0 | 323,31 | 177 |
| 5,62E+08 | 4,18E+08 | 4,16E+08 | 4,21E+08 | 68,075 | 0 | 145,02 | 190 |
| 6,12E+07 | 4,23E+07 | 3,89E+07 | 3,71E+07 | 99,55 | 0 | 63,65 | 56 |
| 4,80E+08 | 4,11E+08 | 3,97E+08 | 4,18E+08 | 266,23 | 0 | 323,31 | 351 |
| 3,83E+08 | 2,37E+08 | 2,44E+08 | 2,58E+08 | 106,25 | 0 | 237,69 | 101 |
| 5,76E+07 | 8,90E+07 | 9,15E+07 | 9,13E+07 | 87,365 | 0 | 61,602 | 60 |
| 2,42E+08 | 1,24E+08 | 1,32E+08 | 1,47E+08 | 92,273 | 0 | 192,14 | 130 |
| 1,35E+08 | 2,28E+08 | 2,12E+08 | 2,30E+08 | 138,97 | 0 | 139,7 | 133 |
| 4,99E+06 | 1,66E+07 | 1,95E+07 | 1,63E+07 | 40,092 | 0,000149 | 3,5263 | 9 |

| | | | | | | | |
|---|---|---|---|---|---|---|---|
| 6,84E+07 | 1,83E+07 | 2,64E+07 | 2,46E+07 | 63,012 | 0 | 93,618 | 45 |
| 2,62E+07 | 9,37E+06 | 8,63E+06 | 8,85E+06 | 27,977 | 0 | 21,582 | 22 |
| 3,06E+08 | 1,93E+08 | 2,15E+08 | 2,14E+08 | 40,84 | 0 | 222,13 | 98 |
| 7,23E+07 | 9,16E+07 | 8,88E+07 | 8,73E+07 | 39,79 | 0 | 20,628 | 40 |
| 3,70E+08 | 1,97E+08 | 1,91E+08 | 1,90E+08 | 74,194 | 0 | 145,22 | 103 |
| 4,54E+07 | 3,35E+07 | 3,45E+07 | 3,57E+07 | 41,119 | 0 | 31,648 | 32 |
| 2,82E+06 | 7,34E+07 | 7,80E+07 | 9,97E+07 | 17,785 | 0 | 19,334 | 19 |
| 1,61E+08 | 1,21E+08 | 1,23E+08 | 1,30E+08 | 23,653 | 0 | 69,135 | 82 |
| 8,11E+07 | 3,60E+06 | 1,24E+06 | 3,25E+06 | 49,656 | 0 | 17,08 | 22 |
| 6,60E+08 | 1,01E+09 | 8,90E+08 | 9,61E+08 | 59,105 | 0 | 208,36 | 236 |
| 2,93E+07 | 5,38E+07 | 6,01E+07 | 5,13E+07 | 96,582 | 0 | 24,179 | 37 |
| 1,45E+07 | 2,41E+07 | 2,40E+07 | 2,59E+07 | 68,954 | 0 | 12,242 | 11 |
| 6,50E+07 | 3,49E+07 | 4,11E+07 | 3,93E+07 | 42,223 | 0 | 65,491 | 43 |
| 8,82E+07 | 1,13E+08 | 1,08E+08 | 1,07E+08 | 80,646 | 0 | 103,64 | 67 |
| 1,56E+09 | 1,29E+09 | 1,29E+09 | 1,31E+09 | 101,43 | 0 | 323,31 | 329 |
| 2,44E+08 | 1,27E+08 | 1,35E+08 | 1,21E+08 | 97,747 | 0 | 300,11 | 122 |
| 1,55E+07 | 5,53E+07 | 5,34E+07 | 5,76E+07 | 77,999 | 0 | 12,497 | 36 |
| 4,63E+08 | 3,05E+08 | 3,31E+08 | 3,31E+08 | 87,142 | 0 | 323,31 | 186 |
| 1,58E+09 | 2,00E+09 | 1,90E+09 | 1,91E+09 | 37,497 | 0 | 323,31 | 214 |
| 4,04E+07 | 2,60E+07 | 2,85E+07 | 2,85E+07 | 52,639 | 0 | 88,615 | 27 |
| 1,22E+09 | 8,62E+08 | 8,93E+08 | 9,18E+08 | 150,72 | 0 | 323,31 | 247 |
| 5,36E+07 | 5,04E+06 | 1,90E+06 | 2,20E+06 | 123,37 | 0 | 38,501 | 44 |
| 2,46E+08 | 1,30E+08 | 1,24E+08 | 1,23E+08 | 64,476 | 0 | 203,24 | 102 |
| 2,56E+08 | 3,13E+08 | 3,20E+08 | 3,04E+08 | 199,15 | 0 | 297,48 | 214 |
| 1,25E+08 | 5,64E+07 | 5,52E+07 | 6,58E+07 | 104,83 | 0 | 27,021 | 66 |
| 2,30E+08 | 3,08E+08 | 2,99E+08 | 2,86E+08 | 44,113 | 0 | 116,51 | 102 |
| 9,16E+08 | 1,21E+09 | 1,24E+09 | 1,22E+09 | 88,303 | 0 | 323,31 | 293 |
| 1,88E+08 | 2,66E+08 | 2,44E+08 | 2,70E+08 | 97,303 | 0 | 228,86 | 139 |
| 9,45E+07 | 6,51E+07 | 6,78E+07 | 6,92E+07 | 113,35 | 0 | 119,22 | 66 |
| 6,07E+08 | 8,27E+08 | 7,82E+08 | 7,76E+08 | 39,47 | 0 | 323,31 | 131 |
| 7,68E+07 | 6,63E+07 | 6,68E+07 | 6,48E+07 | 158,17 | 0 | 31,973 | 76 |
| 2,54E+07 | 5,18E+06 | 2,82E+06 | 2,95E+06 | 46,474 | 0 | 12,103 | 11 |
| 1,89E+09 | 1,40E+09 | 1,31E+09 | 1,35E+09 | 144,27 | 0 | 323,31 | 403 |
| 4,24E+07 | 1,90E+08 | 2,09E+08 | 2,20E+08 | 60,648 | 0 | 109,68 | 74 |
| 3,58E+08 | 2,56E+08 | 2,46E+08 | 2,66E+08 | 75,418 | 0 | 143,2 | 126 |
| 1,68E+06 | 2,65E+08 | 2,78E+08 | 2,60E+08 | 49,788 | 0 | 111,16 | 58 |
| 4,11E+08 | 2,72E+08 | 2,66E+08 | 2,56E+08 | 152,4 | 0 | 190,54 | 189 |
| 9,72E+06 | 5,14E+07 | 4,62E+07 | 6,93E+07 | 71,318 | 0 | 21,26 | 38 |
| 6,59E+07 | 2,54E+07 | 3,22E+07 | 3,18E+07 | 129,24 | 0 | 42,065 | 48 |
| 5,45E+07 | 2,69E+07 | 2,81E+07 | 3,01E+07 | 52,224 | 0 | 54,829 | 28 |
| 1,65E+08 | 7,20E+07 | 7,14E+07 | 7,87E+07 | 116,51 | 0 | 119,12 | 83 |
| 1,03E+09 | 1,55E+09 | 1,60E+09 | 1,65E+09 | 94,859 | 0 | 323,31 | 323 |
| 6,98E+07 | 1,06E+08 | 1,02E+08 | 9,60E+07 | 74,75 | 0 | 31,97 | 83 |
| 2,17E+08 | 1,26E+08 | 1,14E+08 | 1,13E+08 | 43,095 | 0 | 176 | 92 |
| 1,53E+08 | 1,62E+06 | 6,45E+06 | 3,11E+06 | 29,645 | 0 | 53,807 | 38 |
| 7,47E+07 | 5,30E+07 | 5,35E+07 | 5,85E+07 | 143,07 | 0 | 87,527 | 68 |
| 9,37E+06 | 2,68E+06 | 3,16E+06 | 3,57E+06 | 94,475 | 0 | 9,1977 | 13 |
| 8,07E+08 | 6,17E+08 | 6,13E+08 | 6,37E+08 | 136,67 | 0 | 323,31 | 288 |
| 3,59E+09 | 4,05E+09 | 4,12E+09 | 4,23E+09 | 51,065 | 0 | 323,31 | 304 |
| 1,39E+08 | 8,44E+07 | 8,95E+07 | 7,70E+07 | 40,036 | 0 | 100,53 | 73 |

| | | | | | | | |
|---|---|---|---|---|---|---|---|
| 5,61E+07 | 1,05E+07 | 5,45E+06 | 8,52E+06 | 42,481 | 0 | 23,164 | 16 |
| 6,78E+08 | 5,34E+08 | 4,99E+08 | 5,19E+08 | 85,938 | 0 | 222,88 | 220 |
| 1,32E+08 | 9,08E+07 | 7,77E+07 | 8,09E+07 | 238,98 | 0 | 47,794 | 103 |
| 5,24E+07 | 4,73E+06 | 1,88E+06 | 3,73E+06 | 110,91 | 0 | 12,384 | 20 |
| 1,95E+08 | 1,61E+08 | 1,51E+08 | 1,49E+08 | 30,641 | 0 | 103,6 | 91 |
| 2,97E+08 | 2,14E+08 | 2,09E+08 | 2,30E+08 | 51,691 | 0 | 323,31 | 89 |
| 1,77E+07 | 6,23E+07 | 6,02E+07 | 6,80E+07 | 25,535 | 0 | 76,114 | 27 |
| 5,80E+07 | 2,39E+06 | 6,34E+06 | 3,59E+06 | 29,237 | 0 | 11,517 | 20 |
| 2,20E+06 | 1,55E+08 | 9,18E+07 | 9,40E+07 | 50,167 | 0 | 30,221 | 38 |
| 2,74E+08 | 1,76E+08 | 1,77E+08 | 1,64E+08 | 24,72 | 0 | 33,647 | 59 |
| 2,60E+07 | 1,54E+07 | 1,60E+07 | 1,54E+07 | 84,55 | 0 | 15,086 | 26 |
| 4,07E+06 | 1,11E+07 | 1,15E+07 | 1,26E+07 | 26,047 | 0,000156 | 4,6334 | 11 |
| 4,57E+06 | 4,71E+07 | 2,74E+07 | 2,98E+07 | 149,29 | 0,000151 | 3,7965 | 14 |
| 2,55E+08 | 1,52E+08 | 1,57E+08 | 1,39E+08 | 62,995 | 0 | 102,33 | 99 |
| 2,19E+09 | 1,45E+09 | 1,44E+09 | 1,33E+09 | 77,043 | 0 | 323,31 | 369 |
| 1,25E+08 | 6,37E+07 | 5,47E+07 | 5,32E+07 | 29,572 | 0 | 30,812 | 49 |
| 1,08E+09 | 1,41E+09 | 1,47E+09 | 1,57E+09 | 134,75 | 0 | 323,31 | 277 |
| 1,27E+08 | 2,84E+08 | 2,90E+08 | 3,04E+08 | 59,902 | 0 | 42,685 | 95 |
| 1,13E+07 | 7,05E+06 | 7,42E+06 | 7,84E+06 | 155,17 | 0 | 6,1788 | 20 |
| 1,94E+08 | 1,58E+08 | 1,51E+08 | 1,63E+08 | 203,51 | 0 | 240,16 | 144 |
| 7,26E+07 | 8,91E+07 | 8,97E+07 | 9,61E+07 | 61,012 | 0 | 79,312 | 63 |
| 5,18E+08 | 4,10E+08 | 4,31E+08 | 4,25E+08 | 51,091 | 0 | 323,31 | 149 |
| 3,45E+07 | 1,19E+08 | 1,18E+08 | 1,13E+08 | 58,193 | 0 | 44,815 | 74 |
| 2,30E+08 | 3,67E+08 | 3,48E+08 | 3,41E+08 | 46,567 | 0 | 81,053 | 92 |
| 7,99E+06 | 4,02E+07 | 3,89E+07 | 4,10E+07 | 21,509 | 0 | 11,35 | 24 |
| 3,93E+06 | 1,03E+07 | 9,33E+06 | 1,25E+07 | 68,455 | 0,000153 | 4,194 | 10 |
| 5,17E+07 | 1,13E+08 | 1,09E+08 | 9,85E+07 | 70,088 | 0 | 52,845 | 55 |
| 3,82E+08 | 6,55E+08 | 6,63E+08 | 6,85E+08 | 52,227 | 0 | 323,31 | 155 |
| 1,57E+08 | 1,22E+08 | 1,17E+08 | 1,13E+08 | 74,101 | 0 | 41,455 | 67 |
| 2,13E+07 | 6,32E+07 | 5,55E+07 | 6,58E+07 | 92,258 | 0 | 41,977 | 62 |
| 5,05E+08 | 3,18E+08 | 2,99E+08 | 3,16E+08 | 103 | 0 | 323,31 | 192 |
| 1,84E+07 | 2,79E+07 | 2,79E+07 | 2,55E+07 | 56,268 | 0 | 11,051 | 30 |
| 1,40E+08 | 1,39E+07 | 1,41E+07 | 1,52E+07 | 22,687 | 0 | 234,74 | 29 |
| 1,10E+08 | 1,40E+08 | 1,52E+08 | 1,47E+08 | 91,982 | 0 | 225,81 | 86 |
| 8,45E+07 | 1,37E+08 | 1,34E+08 | 1,23E+08 | 50,964 | 0 | 32,084 | 48 |
| 5,22E+07 | 3,38E+06 | 2,41E+06 | 6,35E+06 | 36,213 | 0 | 15,518 | 12 |
| 1,93E+08 | 1,16E+08 | 1,09E+08 | 1,29E+08 | 67,188 | 0 | 95,348 | 62 |
| 1,54E+08 | 2,66E+08 | 2,93E+08 | 2,74E+08 | 60,736 | 0 | 259,78 | 79 |
| 2,04E+09 | 1,46E+09 | 1,48E+09 | 1,56E+09 | 96,02 | 0 | 323,31 | 275 |
| 1,71E+08 | 1,21E+08 | 1,07E+08 | 1,03E+08 | 94,284 | 0 | 118,02 | 115 |
| 9,26E+05 | 1,49E+08 | 1,78E+08 | 1,57E+08 | 129,68 | 0 | 89,603 | 57 |
| 7,25E+07 | 1,42E+08 | 1,37E+08 | 1,46E+08 | 50,777 | 0 | 96,464 | 77 |
| 5,03E+08 | 6,44E+08 | 6,35E+08 | 6,11E+08 | 20,082 | 0 | 123,93 | 89 |
| 1,29E+08 | 1,94E+08 | 2,06E+08 | 2,07E+08 | 10,99 | 0 | 90,875 | 61 |
| 1,44E+07 | 2,98E+06 | 3,29E+06 | 1,91E+06 | 24,663 | 0,000154 | 4,3225 | 9 |
| 6,06E+07 | 8,34E+07 | 7,91E+07 | 8,00E+07 | 39,292 | 0 | 10,566 | 34 |
| 2,42E+07 | 3,45E+06 | 4,59E+06 | 2,39E+06 | 31,458 | 0 | 9,4679 | 16 |
| 1,13E+08 | 2,11E+07 | 3,20E+07 | 3,28E+07 | 47,97 | 0 | 47,666 | 27 |
| 5,04E+08 | 2,71E+08 | 2,13E+08 | 2,20E+08 | 41,551 | 0 | 315,46 | 121 |
| 2,47E+06 | 3,12E+07 | 3,09E+07 | 2,88E+07 | 8,941 | 0 | 87,884 | 6 |

| | | | | | | | |
|---|---|---|---|---|---|---|---|
| 1,78E+08 | 4,03E+08 | 4,48E+08 | 4,59E+08 | 35,877 | 0 | 323,31 | 80 |
| 9,01E+07 | 5,93E+07 | 5,70E+07 | 6,03E+07 | 78 | 0 | 175,1 | 64 |
| 5,31E+08 | 2,21E+08 | 2,35E+08 | 2,31E+08 | 35,363 | 0 | 75,783 | 91 |
| 1,67E+07 | 2,12E+06 | 1,95E+06 | 1,14E+06 | 135,59 | 0 | 19,422 | 25 |
| 1,09E+08 | 1,95E+08 | 1,83E+08 | 1,71E+08 | 82,58 | 0 | 165,9 | 87 |
| 3,47E+07 | 2,02E+07 | 2,08E+07 | 1,95E+07 | 39,978 | 0 | 37,232 | 37 |
| 6,09E+08 | 9,06E+08 | 1,02E+09 | 1,04E+09 | 33,75 | 0 | 227,15 | 138 |
| 1,30E+08 | 1,09E+08 | 1,15E+08 | 1,11E+08 | 65,218 | 0 | 70,341 | 72 |
| 2,39E+07 | 3,26E+07 | 3,01E+07 | 3,16E+07 | 331,33 | 0 | 12,38 | 42 |
| 1,40E+08 | 3,50E+08 | 2,89E+08 | 2,85E+08 | 78,169 | 0 | 165,42 | 104 |
| 7,46E+07 | 5,35E+07 | 5,19E+07 | 5,57E+07 | 33,56 | 0 | 17,455 | 35 |
| 9,67E+07 | 7,24E+08 | 7,00E+08 | 7,91E+08 | 39,904 | 0 | 323,31 | 94 |
| 7,53E+08 | 6,20E+08 | 5,68E+08 | 5,83E+08 | 117,3 | 0 | 323,31 | 212 |
| 1,30E+08 | 7,38E+07 | 7,52E+07 | 8,13E+07 | 164,18 | 0 | 72,957 | 112 |
| 7,48E+07 | 9,83E+07 | 1,02E+08 | 1,01E+08 | 26,949 | 0 | 70,433 | 73 |
| 8,68E+07 | 1,47E+08 | 1,65E+08 | 1,51E+08 | 119,46 | 0 | 62,131 | 113 |
| 1,12E+08 | 3,41E+07 | 4,12E+07 | 4,33E+07 | 106,36 | 0 | 82,882 | 54 |
| 1,25E+08 | 8,85E+07 | 8,17E+07 | 7,98E+07 | 25,25 | 0 | 16,455 | 20 |
| 3,96E+08 | 7,68E+08 | 8,18E+08 | 8,02E+08 | 91,83 | 0 | 323,31 | 200 |
| 6,12E+07 | 1,14E+08 | 1,11E+08 | 1,02E+08 | 28,803 | 0 | 18,264 | 50 |
| 2,59E+08 | 4,08E+08 | 4,02E+08 | 3,90E+08 | 102,94 | 0 | 134,57 | 119 |
| 4,31E+08 | 3,53E+08 | 3,59E+08 | 3,40E+08 | 58,951 | 0 | 155,48 | 138 |
| 5,57E+07 | 9,92E+05 | 1,72E+06 | 3,59E+06 | 44,326 | 0 | 21,724 | 22 |
| 2,51E+08 | 3,21E+08 | 3,12E+08 | 3,29E+08 | 133,32 | 0 | 151,24 | 173 |
| 1,50E+08 | 2,21E+08 | 2,02E+08 | 2,00E+08 | 59,419 | 0 | 67,878 | 91 |
| 2,93E+07 | 4,22E+07 | 4,55E+07 | 4,65E+07 | 101,97 | 0 | 22,584 | 47 |
| 1,96E+07 | 5,11E+07 | 4,74E+07 | 4,66E+07 | 67,581 | 0 | 20,773 | 37 |
| 9,32E+07 | 5,71E+07 | 6,11E+07 | 5,41E+07 | 300,22 | 0 | 65,838 | 117 |
| 2,42E+08 | 1,98E+08 | 1,83E+08 | 1,82E+08 | 118,41 | 0 | 99,238 | 154 |
| 3,10E+06 | 5,69E+07 | 5,21E+07 | 4,02E+07 | 118,73 | 0 | 18,865 | 39 |
| 4,53E+08 | 3,25E+08 | 3,03E+08 | 2,79E+08 | 69,047 | 0 | 132,83 | 149 |
| 4,50E+08 | 3,73E+08 | 3,86E+08 | 3,60E+08 | 23,984 | 0 | 201,56 | 83 |
| 1,50E+07 | 7,52E+06 | 6,31E+06 | 6,57E+06 | 17,551 | 0,000155 | 4,53 | 10 |
| 1,11E+09 | 8,35E+08 | 8,75E+08 | 9,01E+08 | 834,21 | 0 | 323,31 | 753 |
| 9,76E+07 | 7,49E+07 | 7,47E+07 | 7,23E+07 | 164,55 | 0 | 116,24 | 94 |
| 2,24E+07 | 1,03E+07 | 1,27E+07 | 1,03E+07 | 149,73 | 0 | 6,2197 | 20 |
| 9,75E+07 | 6,47E+07 | 6,91E+07 | 6,19E+07 | 48,384 | 0 | 40,639 | 66 |
| 2,08E+06 | 1,51E+07 | 1,32E+07 | 1,52E+07 | 57,433 | 0 | 7,3797 | 13 |
| 6,69E+07 | 1,00E+08 | 1,02E+08 | 1,07E+08 | 55,662 | 0 | 60,201 | 73 |
| 3,13E+07 | 1,50E+07 | 1,61E+07 | 1,67E+07 | 54,716 | 0 | 25,872 | 24 |
| 7,85E+07 | 4,73E+06 | 1,46E+06 | 3,65E+06 | 127,88 | 0 | 37,701 | 32 |
| 5,89E+07 | 7,97E+06 | 5,22E+06 | 3,14E+06 | 35,56 | 0 | 36,872 | 27 |
| 4,23E+07 | 2,87E+07 | 2,75E+07 | 2,91E+07 | 72,318 | 0 | 31,689 | 65 |
| 1,50E+07 | 3,45E+07 | 4,03E+07 | 3,93E+07 | 73,049 | 0 | 32,345 | 23 |
| 3,60E+06 | 1,74E+07 | 1,25E+07 | 1,13E+07 | 377,62 | 0 | 6,2975 | 10 |
| 2,87E+08 | 2,11E+08 | 2,26E+08 | 2,16E+08 | 84,213 | 0 | 159,18 | 133 |
| 7,12E+08 | 1,29E+09 | 1,21E+09 | 1,13E+09 | 138,56 | 0 | 323,31 | 231 |
| 5,54E+08 | 2,60E+08 | 2,26E+08 | 2,42E+08 | 33,363 | 0 | 97,225 | 87 |
| 2,79E+07 | 1,29E+07 | 1,27E+07 | 9,83E+06 | 64,49 | 0 | 12,712 | 17 |
| 1,41E+09 | 8,25E+08 | 9,34E+08 | 9,53E+08 | 19,53 | 0 | 120,87 | 144 |

| | | | | | | | |
|---|---|---|---|---|---|---|---|
| 2,53E+06 | 1,60E+07 | 2,36E+07 | 3,09E+07 | 284,73 | 0 | 12,717 | 25 |
| 9,95E+06 | 3,17E+07 | 4,28E+07 | 3,21E+07 | 157,01 | 0 | 12,015 | 26 |
| 1,28E+08 | 9,99E+07 | 1,01E+08 | 9,51E+07 | 56,763 | 0 | 105,33 | 36 |
| 2,77E+08 | 3,79E+08 | 3,84E+08 | 3,75E+08 | 152,25 | 0 | 297,47 | 180 |
| 1,93E+07 | 1,21E+08 | 1,14E+08 | 1,42E+08 | 106,19 | 0 | 44,2 | 38 |
| 2,39E+09 | 1,85E+09 | 1,94E+09 | 1,95E+09 | 50,372 | 0 | 323,31 | 191 |
| 2,96E+08 | 2,07E+08 | 1,87E+08 | 1,96E+08 | 34,376 | 0 | 118,14 | 101 |
| 4,58E+07 | 2,95E+07 | 2,65E+07 | 2,88E+07 | 265,49 | 0 | 96,877 | 40 |
| 4,41E+10 | 6,36E+10 | 6,54E+10 | 6,40E+10 | 53,687 | 0 | 323,31 | 2784 |
| 6,93E+09 | 9,11E+09 | 9,83E+09 | 9,23E+09 | 152,04 | 0 | 323,31 | 889 |
| 1,05E+07 | 2,89E+06 | 3,24E+06 | 2,00E+06 | 66,54 | 0,000428 | 2,6579 | 4 |
| 1,17E+09 | 7,38E+08 | 7,43E+08 | 8,10E+08 | 50,02 | 0 | 323,31 | 199 |
| 3,60E+07 | 3,01E+06 | 1,64E+06 | 3,86E+06 | 70,022 | 0 | 21,65 | 14 |
| 4,14E+06 | 1,49E+08 | 1,36E+08 | 1,39E+08 | 67,35 | 0 | 245 | 38 |
| 1,94E+08 | 2,78E+08 | 2,80E+08 | 2,73E+08 | 20,451 | 0 | 21,414 | 43 |
| 3,16E+08 | 1,65E+08 | 1,62E+08 | 1,88E+08 | 23,349 | 0 | 217,78 | 44 |
| 1,62E+08 | 5,42E+06 | 3,28E+06 | 1,23E+06 | 49,906 | 0 | 49,364 | 4 |
| 1,35E+08 | 3,36E+07 | 3,73E+07 | 4,72E+07 | 46,402 | 0 | 29,074 | 54 |
| 1,35E+08 | 6,07E+07 | 7,37E+07 | 5,86E+07 | 46,379 | 0 | 119,85 | 46 |
| 6,43E+08 | 2,26E+09 | 1,91E+09 | 1,67E+09 | 45,681 | 0 | 32,638 | 56 |
| 7,83E+07 | 2,37E+06 | 7,07E+06 | 2,60E+06 | 36,572 | 0 | 85,006 | 19 |
| 2,00E+08 | 1,35E+08 | 1,20E+08 | 1,28E+08 | 56,775 | 0 | 83,646 | 92 |
| 1,94E+09 | 3,39E+09 | 3,19E+09 | 3,04E+09 | 58,388 | 0 | 323,31 | 357 |
| 1,07E+08 | 1,42E+08 | 1,41E+08 | 1,46E+08 | 67,95 | 0 | 51,117 | 79 |
| 2,42E+07 | 1,20E+07 | 8,53E+06 | 9,61E+06 | 42,248 | 0 | 46,515 | 19 |
| 1,28E+08 | 2,35E+08 | 2,46E+08 | 2,59E+08 | 38,124 | 0 | 103,96 | 96 |
| 5,91E+07 | 1,65E+06 | 2,83E+06 | 5,32E+06 | 72,28 | 0 | 73,567 | 32 |
| 1,26E+07 | 2,27E+07 | 2,00E+07 | 2,51E+07 | 68,474 | 0 | 8,5307 | 18 |
| 7,72E+07 | 3,77E+07 | 3,47E+07 | 3,90E+07 | 61,445 | 0 | 11,418 | 40 |
| 3,39E+07 | 5,16E+07 | 4,96E+07 | 5,45E+07 | 46,49 | 0 | 7,96 | 30 |
| 2,59E+06 | 3,44E+07 | 3,28E+07 | 2,63E+07 | 127,69 | 0 | 10,04 | 11 |
| 1,61E+10 | 1,23E+10 | 1,13E+10 | 1,08E+10 | 35,752 | 0 | 323,31 | 697 |
| 9,89E+06 | 4,80E+07 | 5,22E+07 | 3,67E+07 | 113,18 | 0 | 15,814 | 33 |
| 3,46E+07 | 4,87E+06 | 5,81E+06 | 2,78E+06 | 34,001 | 0,000429 | 2,6896 | 11 |
| 1,74E+08 | 3,15E+06 | 3,62E+06 | 2,48E+06 | 117,53 | 0 | 215,47 | 117 |
| 4,57E+08 | 6,05E+08 | 6,71E+08 | 6,27E+08 | 49,278 | 0 | 323,31 | 181 |
| 1,40E+08 | 6,24E+06 | 1,49E+06 | 1,87E+06 | 68,035 | 0 | 35,277 | 48 |
| 5,81E+07 | 1,04E+08 | 9,83E+07 | 9,90E+07 | 15,743 | 0 | 132,09 | 44 |
| 7,79E+06 | 3,13E+06 | 2,07E+06 | 2,17E+06 | 32,88 | 0,000154 | 4,4044 | 6 |
| 4,11E+07 | 7,59E+07 | 7,09E+07 | 7,07E+07 | 45,341 | 0 | 17,404 | 32 |
| 2,21E+08 | 3,79E+07 | 2,58E+07 | 1,58E+07 | 121,9 | 0 | 130,29 | 60 |
| 1,82E+08 | 1,09E+08 | 1,06E+08 | 1,19E+08 | 74,305 | 0 | 46,212 | 72 |
| 8,22E+07 | 5,64E+07 | 4,90E+07 | 5,27E+07 | 282,71 | 0 | 66,856 | 65 |
| 3,27E+08 | 2,56E+08 | 2,36E+08 | 2,53E+08 | 215,99 | 0 | 167,96 | 192 |
| 3,07E+08 | 2,73E+08 | 2,64E+08 | 2,69E+08 | 154,91 | 0 | 236,28 | 165 |
| 2,83E+08 | 2,20E+08 | 2,18E+08 | 2,19E+08 | 31,994 | 0 | 71,618 | 95 |
| 3,59E+08 | 4,84E+08 | 5,08E+08 | 5,09E+08 | 10,871 | 0 | 32,822 | 29 |
| 5,71E+07 | 2,12E+06 | 2,84E+06 | 1,63E+06 | 80,623 | 0 | 19,303 | 20 |
| 2,70E+08 | 5,53E+08 | 5,05E+08 | 5,34E+08 | 61,035 | 0 | 279,18 | 113 |
| 1,89E+07 | 3,41E+06 | 4,97E+06 | 2,58E+06 | 84,438 | 0 | 11,845 | 17 |

| | | | | | | | |
|---|---|---|---|---|---|---|---|
| 4,10E+07 | 6,77E+07 | 5,90E+07 | 6,58E+07 | 65,239 | 0 | 37,166 | 53 |
| 7,63E+07 | 1,87E+06 | 1,25E+06 | 4,66E+06 | 37,804 | 0 | 14,761 | 24 |
| 1,67E+08 | 8,75E+07 | 9,11E+07 | 8,25E+07 | 32,402 | 0 | 71,664 | 88 |
| 1,68E+09 | 1,20E+09 | 1,22E+09 | 1,20E+09 | 80,953 | 0 | 323,31 | 403 |
| 4,46E+08 | 6,94E+08 | 7,07E+08 | 6,95E+08 | 33,692 | 0 | 323,31 | 179 |
| 6,11E+08 | 7,96E+08 | 8,11E+08 | 8,16E+08 | 52,767 | 0 | 214,62 | 190 |
| 1,28E+08 | 1,88E+08 | 1,71E+08 | 1,68E+08 | 113,09 | 0 | 299,11 | 130 |
| 1,75E+07 | 4,02E+06 | 5,10E+06 | 2,86E+06 | 38,347 | 0 | 8,2978 | 10 |
| 6,21E+07 | 1,95E+06 | 4,32E+06 | 1,28E+06 | 87,653 | 0 | 45,169 | 59 |
| 5,76E+07 | 1,06E+07 | 1,65E+07 | 9,49E+06 | 81,1 | 0 | 107,16 | 22 |
| 2,23E+08 | 1,60E+08 | 1,79E+08 | 1,67E+08 | 64,514 | 0 | 106,85 | 129 |
| 4,05E+08 | 2,45E+08 | 2,14E+08 | 2,28E+08 | 37,863 | 0 | 207,68 | 87 |
| 7,47E+06 | 1,99E+08 | 1,94E+08 | 2,42E+08 | 23,182 | 0 | 24,906 | 31 |
| 1,46E+07 | 4,11E+07 | 4,20E+07 | 3,70E+07 | 17,444 | 0 | 33,944 | 22 |
| 2,46E+08 | 1,78E+08 | 1,96E+08 | 1,83E+08 | 100,74 | 0 | 101,75 | 115 |
| 1,02E+09 | 1,67E+09 | 1,50E+09 | 1,66E+09 | 38,442 | 0 | 323,31 | 252 |
| 1,93E+08 | 1,24E+08 | 1,10E+08 | 1,14E+08 | 88,606 | 0 | 178,5 | 117 |
| 5,10E+07 | 4,31E+07 | 4,18E+07 | 4,32E+07 | 115,03 | 0 | 26,962 | 59 |
| 4,39E+08 | 6,44E+08 | 6,72E+08 | 6,79E+08 | 123,79 | 0 | 323,31 | 245 |
| 1,34E+08 | 3,35E+07 | 3,41E+07 | 2,14E+07 | 38,224 | 0 | 88,833 | 39 |
| 3,78E+07 | 1,72E+07 | 1,70E+07 | 1,41E+07 | 49,753 | 0 | 9,3498 | 36 |
| 5,75E+07 | 1,23E+07 | 1,93E+07 | 1,26E+07 | 30,482 | 0 | 27,204 | 30 |
| 3,69E+06 | 3,00E+07 | 2,30E+07 | 2,39E+07 | 63,056 | 0 | 8,5491 | 12 |
| 5,71E+07 | 3,56E+07 | 3,76E+07 | 3,65E+07 | 25,356 | 0 | 77,003 | 32 |
| 9,60E+07 | 6,33E+07 | 6,78E+07 | 5,96E+07 | 59,514 | 0 | 58,996 | 66 |
| 4,04E+07 | 1,91E+07 | 1,72E+07 | 1,63E+07 | 66,22 | 0 | 33,059 | 24 |
| 1,56E+09 | 1,94E+09 | 1,89E+09 | 1,91E+09 | 152,59 | 0 | 323,31 | 529 |
| 3,56E+08 | 5,27E+08 | 5,33E+08 | 5,55E+08 | 104,2 | 0 | 270,93 | 186 |
| 2,02E+07 | 4,38E+07 | 4,03E+07 | 4,38E+07 | 89,713 | 0 | 72,359 | 39 |
| 8,80E+07 | 8,29E+06 | 1,50E+07 | 8,04E+06 | 24,794 | 0 | 34,329 | 25 |
| 6,64E+09 | 8,74E+09 | 8,82E+09 | 8,53E+09 | 67,277 | 0 | 323,31 | 1162 |
| 1,69E+07 | 3,77E+06 | 4,73E+06 | 3,85E+06 | 30,308 | 0 | 9,3718 | 12 |
| 4,10E+07 | 1,36E+07 | 1,71E+07 | 1,94E+07 | 40,045 | 0 | 228,51 | 12 |
| 1,49E+08 | 7,23E+07 | 7,94E+07 | 7,39E+07 | 54,776 | 0 | 154,26 | 90 |
| 4,12E+08 | 2,36E+08 | 2,22E+08 | 1,93E+08 | 55,731 | 0 | 111,39 | 115 |
| 1,38E+08 | 9,11E+07 | 9,15E+07 | 1,03E+08 | 53,998 | 0 | 120,04 | 57 |
| 1,80E+08 | 1,15E+08 | 1,14E+08 | 1,02E+08 | 48,385 | 0 | 74,665 | 104 |
| 1,20E+08 | 2,69E+06 | 8,52E+06 | 7,10E+06 | 98,026 | 0 | 60,764 | 50 |
| 2,53E+08 | 3,40E+06 | 2,20E+06 | 5,22E+06 | 72,931 | 0 | 50,741 | 47 |
| 2,47E+09 | 2,16E+09 | 2,08E+09 | 2,16E+09 | 105,73 | 0 | 323,31 | 418 |
| 6,35E+06 | 5,22E+07 | 5,23E+07 | 4,25E+07 | 9,5842 | 0 | 8,7457 | 15 |
| 1,29E+08 | 1,78E+08 | 1,90E+08 | 1,69E+08 | 31,639 | 0 | 147,29 | 56 |
| 2,39E+09 | 1,76E+09 | 1,86E+09 | 1,80E+09 | 80,751 | 0 | 323,31 | 455 |
| 5,36E+07 | 3,07E+07 | 2,51E+07 | 3,10E+07 | 85,995 | 0 | 45,411 | 44 |
| 3,44E+08 | 2,30E+08 | 1,99E+08 | 1,98E+08 | 78,491 | 0 | 196,77 | 95 |
| 1,43E+08 | 2,61E+08 | 2,42E+08 | 2,69E+08 | 22,604 | 0 | 29,907 | 44 |
| 3,86E+07 | 4,34E+06 | 1,53E+06 | 2,85E+06 | 24,241 | 0,000436 | 2,9966 | 16 |
| 3,65E+08 | 2,84E+08 | 2,78E+08 | 2,64E+08 | 61,341 | 0 | 155,17 | 76 |
| 3,03E+08 | 2,49E+08 | 2,59E+08 | 2,63E+08 | 28,062 | 0 | 142,03 | 71 |
| 4,69E+07 | 3,29E+07 | 2,91E+07 | 3,19E+07 | 108,43 | 0 | 24,76 | 53 |

| | | | | | | | |
|---|---|---|---|---|---|---|---|
| 1,98E+08 | 1,53E+08 | 1,41E+08 | 1,37E+08 | 63,131 | 0 | 134,44 | 96 |
| 3,97E+06 | 2,08E+08 | 1,70E+08 | 7,25E+07 | 40,982 | 0 | 33,667 | 19 |
| 9,54E+08 | 1,30E+09 | 1,31E+09 | 1,26E+09 | 55,971 | 0 | 323,31 | 224 |
| 6,10E+08 | 4,21E+08 | 3,73E+08 | 3,46E+08 | 37,872 | 0 | 323,31 | 142 |
| 1,03E+07 | 1,50E+06 | 2,08E+06 | 2,19E+06 | 72,692 | 0,000157 | 4,8938 | 6 |
| 5,82E+07 | 9,51E+07 | 9,91E+07 | 9,70E+07 | 48,513 | 0 | 27,382 | 72 |
| 6,36E+08 | 7,29E+08 | 7,40E+08 | 7,06E+08 | 74,569 | 0 | 323,31 | 217 |
| 1,96E+08 | 3,85E+07 | 5,27E+07 | 5,88E+07 | 27,706 | 0 | 18,05 | 47 |
| 1,96E+08 | 1,50E+08 | 1,47E+08 | 1,47E+08 | 72,349 | 0 | 58,131 | 106 |
| 6,53E+06 | 2,98E+06 | 3,26E+06 | 2,38E+06 | 42,407 | 0,002095 | 2,135 | 5 |
| 2,11E+07 | 2,71E+07 | 2,67E+07 | 2,62E+07 | 57,673 | 0 | 10,456 | 21 |
| 5,74E+07 | 2,84E+07 | 2,96E+07 | 2,97E+07 | 74,812 | 0 | 78,239 | 41 |
| 5,65E+07 | 1,85E+06 | 4,52E+06 | 1,27E+06 | 12,255 | 0 | 12,947 | 17 |
| 4,33E+08 | 3,30E+08 | 3,02E+08 | 3,39E+08 | 148,07 | 0 | 181,35 | 228 |
| 1,11E+08 | 1,69E+08 | 1,48E+08 | 1,56E+08 | 98 | 0 | 55,547 | 124 |
| 2,62E+08 | 8,60E+06 | 1,80E+06 | 1,77E+06 | 148,23 | 0 | 101,32 | 94 |
| 6,06E+07 | 2,23E+07 | 1,83E+07 | 1,58E+07 | 21,166 | 0 | 37,34 | 18 |
| 1,16E+08 | 1,97E+08 | 1,80E+08 | 1,66E+08 | 61,75 | 0 | 90,582 | 87 |
| 3,19E+08 | 2,49E+08 | 2,48E+08 | 2,49E+08 | 106,08 | 0 | 192,84 | 151 |
| 4,61E+07 | 1,68E+06 | 3,85E+06 | 4,96E+06 | 36,354 | 0 | 35,004 | 25 |
| 5,75E+08 | 2,90E+08 | 2,64E+08 | 2,56E+08 | 25,976 | 0 | 42,676 | 131 |
| 5,57E+08 | 3,81E+08 | 3,81E+08 | 3,60E+08 | 84,276 | 0 | 323,31 | 168 |
| 1,24E+08 | 3,77E+06 | 7,61E+05 | 6,84E+05 | 75,916 | 0 | 70,877 | 46 |
| 1,77E+09 | 2,12E+09 | 2,15E+09 | 2,10E+09 | 107,06 | 0 | 323,31 | 475 |
| 1,16E+09 | 1,45E+09 | 1,45E+09 | 1,44E+09 | 144,63 | 0 | 323,31 | 385 |
| 7,58E+07 | 1,44E+08 | 1,55E+08 | 1,61E+08 | 58,861 | 0 | 242,25 | 79 |
| 5,92E+08 | 8,27E+08 | 8,63E+08 | 8,36E+08 | 28,152 | 0 | 323,31 | 167 |
| 4,05E+07 | 5,70E+07 | 5,93E+07 | 6,53E+07 | 57,648 | 0 | 40,535 | 36 |
| 6,36E+08 | 4,10E+08 | 4,36E+08 | 4,24E+08 | 9,3267 | 0 | 8,7925 | 41 |
| 2,86E+08 | 1,96E+08 | 1,88E+08 | 1,67E+08 | 36,717 | 0 | 112,94 | 96 |
| 1,79E+08 | 2,60E+08 | 2,60E+08 | 2,75E+08 | 43,5 | 0 | 160,26 | 71 |
| 8,33E+07 | 9,78E+05 | 4,36E+06 | 1,46E+06 | 63,221 | 0 | 36,626 | 28 |
| 7,62E+07 | 1,02E+08 | 9,76E+07 | 1,06E+08 | 32,549 | 0 | 24,725 | 46 |
| 2,77E+07 | 5,88E+07 | 5,56E+07 | 6,62E+07 | 47,744 | 0 | 24,182 | 29 |
| 3,35E+08 | 2,12E+08 | 2,28E+08 | 1,97E+08 | 45,78 | 0 | 81,846 | 108 |
| 1,71E+08 | 2,41E+08 | 2,60E+08 | 2,51E+08 | 90,244 | 0 | 251,74 | 114 |
| 3,63E+08 | 5,99E+08 | 6,36E+08 | 6,46E+08 | 68,988 | 0 | 265,75 | 147 |
| 3,84E+07 | 1,05E+08 | 1,06E+08 | 1,13E+08 | 60,336 | 0 | 57,218 | 63 |
| 1,31E+07 | 4,39E+06 | 2,66E+06 | 3,68E+06 | 59,959 | 0 | 6,3433 | 8 |
| 1,30E+09 | 1,90E+09 | 1,96E+09 | 1,94E+09 | 225,65 | 0 | 323,31 | 606 |
| 2,50E+09 | 1,57E+09 | 1,53E+09 | 1,50E+09 | 28,467 | 0 | 323,31 | 307 |
| 1,87E+07 | 2,31E+06 | 2,90E+06 | 4,45E+06 | 28,531 | 0,000151 | 3,7669 | 12 |
| 2,22E+08 | 4,32E+08 | 4,01E+08 | 4,04E+08 | 63,953 | 0 | 323,31 | 144 |
| 4,97E+07 | 3,35E+07 | 2,80E+07 | 3,03E+07 | 49,61 | 0 | 28,677 | 29 |
| 4,95E+07 | 9,63E+07 | 9,69E+07 | 9,41E+07 | 82,975 | 0 | 22,968 | 51 |
| 3,50E+08 | 3,11E+08 | 3,10E+08 | 2,96E+08 | 69,874 | 0 | 91,183 | 107 |
| 1,03E+08 | 7,68E+07 | 8,26E+07 | 8,40E+07 | 74,919 | 0 | 31,559 | 64 |
| 1,45E+08 | 1,08E+08 | 1,11E+08 | 1,08E+08 | 75,596 | 0 | 155,51 | 51 |
| 1,24E+08 | 1,80E+08 | 1,79E+08 | 1,78E+08 | 36,946 | 0 | 158,95 | 53 |
| 7,51E+07 | 1,04E+07 | 7,62E+06 | 5,20E+06 | 56,782 | 0 | 53,715 | 41 |

| | | | | | | | |
|---|---|---|---|---|---|---|---|
| 1,66E+08 | 1,07E+08 | 1,09E+08 | 1,16E+08 | 52,812 | 0 | 29,383 | 55 |
| 1,12E+07 | 2,31E+07 | 2,13E+07 | 1,97E+07 | 40,446 | 0 | 10,988 | 25 |
| 4,83E+07 | 9,51E+07 | 1,00E+08 | 9,54E+07 | 40,035 | 0 | 58,622 | 34 |
| 1,13E+08 | 5,68E+07 | 6,34E+07 | 6,76E+07 | 34,425 | 0 | 138,08 | 65 |
| 2,39E+08 | 1,43E+07 | 8,77E+06 | 3,44E+06 | 69,422 | 0 | 133 | 48 |
| 3,07E+07 | 1,25E+07 | 1,29E+07 | 1,12E+07 | 48,937 | 0 | 12,998 | 24 |
| 3,63E+07 | 1,41E+08 | 1,33E+08 | 1,37E+08 | 218,71 | 0 | 91,986 | 76 |
| 3,68E+07 | 5,61E+07 | 5,09E+07 | 5,56E+07 | 34,46 | 0 | 22,072 | 33 |
| 2,46E+08 | 1,78E+08 | 1,63E+08 | 1,68E+08 | 196,38 | 0 | 233,44 | 145 |
| 8,46E+06 | 4,84E+06 | 5,72E+06 | 4,72E+06 | 17,871 | 0 | 5,7743 | 6 |
| 2,60E+07 | 4,12E+07 | 3,86E+07 | 3,58E+07 | 28,231 | 0 | 13,276 | 26 |
| 1,40E+08 | 3,64E+08 | 3,38E+08 | 2,95E+08 | 15,184 | 0 | 55,862 | 62 |
| 3,23E+08 | 4,25E+08 | 4,57E+08 | 4,49E+08 | 45,463 | 0 | 105,49 | 112 |
| 6,16E+07 | 1,32E+07 | 8,95E+06 | 1,26E+07 | 56,602 | 0 | 46,694 | 28 |
| 9,27E+07 | 1,67E+08 | 1,65E+08 | 1,79E+08 | 21,834 | 0 | 50,301 | 47 |
| 5,14E+08 | 7,22E+08 | 7,32E+08 | 6,50E+08 | 241,44 | 0 | 323,31 | 327 |
| 2,35E+07 | 5,13E+07 | 4,69E+07 | 5,78E+07 | 22,607 | 0 | 22,452 | 21 |
| 9,20E+09 | 4,16E+09 | 4,21E+09 | 4,30E+09 | 23,713 | 0 | 323,31 | 392 |
| 1,77E+08 | 4,78E+08 | 4,94E+08 | 4,48E+08 | 21,024 | 0 | 106,37 | 89 |
| 4,99E+08 | 6,39E+08 | 7,05E+08 | 6,47E+08 | 70,449 | 0 | 97,297 | 171 |
| 1,00E+09 | 1,54E+09 | 1,50E+09 | 1,44E+09 | 121,12 | 0 | 323,31 | 355 |
| 7,31E+07 | 5,33E+07 | 5,90E+07 | 5,41E+07 | 120,81 | 0 | 30,818 | 47 |
| 2,72E+07 | 8,23E+06 | 5,10E+06 | 5,14E+06 | 27,294 | 0,000151 | 3,886 | 7 |
| 1,12E+08 | 7,38E+07 | 7,90E+07 | 8,23E+07 | 59,71 | 0 | 25,119 | 55 |
| 4,21E+06 | 4,29E+07 | 3,43E+07 | 4,18E+07 | 16,782 | 0,000155 | 4,4445 | 3 |
| 1,12E+08 | 2,25E+08 | 2,08E+08 | 2,41E+08 | 49,736 | 0 | 62,12 | 66 |
| 8,66E+08 | 1,16E+09 | 1,15E+09 | 1,16E+09 | 96,001 | 0 | 323,31 | 273 |
| 7,00E+06 | 8,95E+06 | 9,95E+06 | 9,29E+06 | 119,17 | 0,00015 | 3,6407 | 10 |
| 3,65E+07 | 6,94E+07 | 6,44E+07 | 5,55E+07 | 69,703 | 0 | 31,711 | 57 |
| 4,12E+07 | 7,66E+06 | 3,19E+06 | 4,33E+06 | 36,807 | 0 | 16,459 | 17 |
| 2,75E+07 | 3,26E+06 | 3,20E+06 | 2,30E+06 | 66,048 | 0 | 21,802 | 16 |
| 3,15E+06 | 1,24E+08 | 1,12E+08 | 1,18E+08 | 53,553 | 0 | 94,281 | 43 |
| 2,08E+09 | 1,68E+09 | 1,69E+09 | 1,57E+09 | 107,5 | 0 | 323,31 | 398 |
| 1,02E+09 | 1,12E+09 | 1,12E+09 | 1,12E+09 | 120,66 | 0 | 323,31 | 374 |
| 7,33E+07 | 1,25E+08 | 1,20E+08 | 1,17E+08 | 34,406 | 0 | 74,559 | 44 |
| 1,03E+09 | 4,84E+08 | 4,75E+08 | 4,56E+08 | 37,616 | 0 | 323,31 | 144 |
| 1,78E+08 | 2,42E+08 | 2,17E+08 | 2,35E+08 | 37,781 | 0 | 121,65 | 92 |
| 1,41E+08 | 2,25E+08 | 2,52E+08 | 2,66E+08 | 39,799 | 0 | 128,7 | 79 |
| 3,94E+06 | 4,69E+07 | 5,84E+07 | 7,48E+07 | 139,07 | 0 | 7,0858 | 30 |
| 3,99E+08 | 3,41E+08 | 3,57E+08 | 3,57E+08 | 86,876 | 0 | 113,14 | 222 |
| 1,69E+07 | 2,93E+06 | 2,65E+06 | 2,85E+06 | 36,977 | 0,003183 | 1,9833 | 7 |
| 5,15E+07 | 1,02E+08 | 9,26E+07 | 9,27E+07 | 50,683 | 0 | 123,76 | 74 |
| 2,48E+07 | 1,61E+07 | 1,62E+07 | 1,38E+07 | 49,908 | 0 | 52,421 | 28 |
| 1,01E+08 | 1,60E+08 | 1,59E+08 | 1,43E+08 | 223,37 | 0 | 124,01 | 129 |
| 3,87E+09 | 2,53E+09 | 2,69E+09 | 2,53E+09 | 98,124 | 0 | 96,931 | 193 |
| 1,03E+10 | 1,25E+10 | 1,28E+10 | 1,25E+10 | 89,321 | 0 | 323,31 | 1061 |
| 7,17E+08 | 4,29E+08 | 4,38E+08 | 4,99E+08 | 58,47 | 0 | 206,38 | 156 |
| 4,51E+08 | 5,42E+08 | 5,12E+08 | 5,13E+08 | 19,992 | 0 | 27,743 | 82 |
| 1,20E+08 | 7,80E+07 | 8,89E+07 | 8,11E+07 | 161,97 | 0 | 127,22 | 93 |
| 2,99E+08 | 4,13E+08 | 3,89E+08 | 3,77E+08 | 57,223 | 0 | 107,01 | 127 |

| | | | | | | | |
|---|---|---|---|---|---|---|---|
| 2,31E+07 | 5,27E+06 | 6,15E+06 | 6,70E+06 | 31,524 | 0 | 6,2374 | 19 |
| 1,59E+07 | 3,77E+07 | 3,53E+07 | 3,57E+07 | 210,46 | 0 | 17,533 | 39 |
| 8,26E+07 | 5,04E+07 | 5,06E+07 | 4,08E+07 | 78,765 | 0 | 45,194 | 69 |
| 2,26E+07 | 7,34E+07 | 7,63E+07 | 6,98E+07 | 73,606 | 0 | 15,798 | 39 |
| 2,01E+08 | 2,74E+08 | 2,80E+08 | 3,15E+08 | 74,452 | 0 | 323,31 | 41 |
| 3,75E+07 | 8,96E+07 | 8,28E+07 | 7,76E+07 | 26,132 | 0 | 85,498 | 28 |
| 2,27E+08 | 1,49E+08 | 1,62E+08 | 1,73E+08 | 49,116 | 0 | 121,7 | 88 |
| 2,43E+06 | 2,09E+07 | 2,56E+07 | 2,41E+07 | 81,035 | 0 | 15,395 | 12 |
| 2,13E+08 | 1,79E+08 | 1,75E+08 | 1,80E+08 | 30,501 | 0 | 158,2 | 90 |
| 6,35E+06 | 2,43E+07 | 2,66E+07 | 3,94E+07 | 53,771 | 0 | 13,169 | 18 |
| 1,05E+08 | 1,87E+08 | 2,08E+08 | 2,12E+08 | 54,537 | 0 | 102,25 | 74 |
| 2,99E+08 | 2,20E+06 | 1,18E+07 | 3,78E+06 | 180,69 | 0 | 323,31 | 110 |
| 6,50E+08 | 5,02E+08 | 4,93E+08 | 5,12E+08 | 24,043 | 0 | 323,31 | 83 |
| 3,91E+08 | 3,06E+08 | 2,99E+08 | 2,84E+08 | 96,729 | 0 | 139,93 | 183 |
| 1,04E+09 | 1,60E+09 | 1,71E+09 | 1,72E+09 | 113,84 | 0 | 319,87 | 312 |
| 1,03E+08 | 6,83E+07 | 6,24E+07 | 6,93E+07 | 119,91 | 0 | 77,831 | 77 |
| 2,68E+06 | 1,27E+07 | 1,48E+07 | 1,46E+07 | 21,949 | 0,000157 | 4,9033 | 10 |
| 2,89E+08 | 2,18E+08 | 2,21E+08 | 1,99E+08 | 43,267 | 0 | 323,31 | 110 |
| 1,17E+08 | 8,28E+07 | 8,31E+07 | 8,78E+07 | 47,13 | 0 | 19,42 | 64 |
| 3,40E+09 | 3,03E+09 | 2,98E+09 | 3,12E+09 | 27,855 | 0 | 323,31 | 255 |
| 2,26E+08 | 1,19E+08 | 1,29E+08 | 1,38E+08 | 55,76 | 0 | 32,121 | 73 |
| 6,76E+08 | 9,01E+08 | 9,55E+08 | 9,39E+08 | 74,042 | 0 | 294,87 | 252 |
| 9,91E+06 | 1,85E+07 | 1,79E+07 | 1,81E+07 | 99,146 | 0 | 19,044 | 24 |
| 2,00E+07 | 3,44E+07 | 3,92E+07 | 3,33E+07 | 45,908 | 0 | 15,054 | 30 |
| 1,56E+07 | 8,47E+06 | 6,80E+06 | 7,04E+06 | 82,262 | 0 | 48,083 | 14 |
| 2,54E+08 | 2,16E+08 | 2,05E+08 | 2,01E+08 | 32,214 | 0 | 73,947 | 84 |
| 4,65E+08 | 6,20E+08 | 6,51E+08 | 6,58E+08 | 15,344 | 0 | 179,52 | 54 |
| 4,88E+07 | 3,66E+06 | 1,91E+06 | 1,82E+06 | 38,204 | 0 | 96,31 | 45 |
| 1,58E+08 | 2,10E+08 | 2,03E+08 | 2,04E+08 | 135,94 | 0 | 80,974 | 140 |
| 2,12E+09 | 1,67E+09 | 1,51E+09 | 1,51E+09 | 23,363 | 0 | 282,59 | 123 |
| 1,32E+08 | 2,05E+08 | 1,92E+08 | 2,03E+08 | 55,518 | 0 | 140,48 | 115 |
| 1,15E+07 | 1,73E+06 | 3,26E+06 | 1,69E+06 | 115,97 | 0 | 5,5669 | 15 |
| 1,08E+08 | 6,92E+07 | 5,80E+07 | 6,96E+07 | 30,658 | 0 | 24,633 | 48 |
| 1,96E+07 | 6,30E+07 | 5,01E+07 | 5,50E+07 | 33,061 | 0 | 30,037 | 33 |
| 2,72E+08 | 4,19E+08 | 3,70E+08 | 3,66E+08 | 66,223 | 0 | 221,24 | 95 |
| 1,61E+07 | 3,44E+07 | 3,60E+07 | 3,68E+07 | 17,768 | 0 | 13,047 | 7 |
| 4,71E+07 | 8,49E+07 | 7,52E+07 | 8,12E+07 | 20,791 | 0 | 139,7 | 53 |
| 1,08E+09 | 1,77E+09 | 1,75E+09 | 1,79E+09 | 54,191 | 0 | 323,31 | 199 |
| 4,98E+07 | 2,07E+07 | 2,17E+07 | 2,20E+07 | 165,25 | 0 | 41,834 | 53 |
| 7,88E+07 | 6,02E+07 | 5,33E+07 | 5,77E+07 | 118,46 | 0 | 24,005 | 59 |
| 8,28E+06 | 3,85E+07 | 4,34E+07 | 4,93E+07 | 24,612 | 0 | 7,9975 | 12 |
| 1,55E+09 | 1,08E+09 | 1,17E+09 | 1,12E+09 | 33,659 | 0 | 323,31 | 180 |
| 1,34E+09 | 1,61E+09 | 1,54E+09 | 1,61E+09 | 108,3 | 0 | 323,31 | 352 |
| 8,98E+06 | 1,51E+06 | 2,92E+06 | 2,10E+06 | 175,55 | 0,000156 | 4,7201 | 7 |
| 1,10E+07 | 3,71E+06 | 2,79E+06 | 2,14E+06 | 55,747 | 0 | 20,641 | 14 |
| 2,45E+07 | 1,14E+07 | 1,38E+07 | 9,94E+06 | 27,01 | 0 | 10,465 | 17 |
| 2,28E+07 | 5,50E+07 | 7,07E+07 | 7,11E+07 | 121,6 | 0 | 62,44 | 56 |
| 1,11E+07 | 5,11E+06 | 3,60E+06 | 3,90E+06 | 177,53 | 0 | 11,181 | 12 |
| 8,04E+07 | 1,95E+06 | 3,77E+06 | 7,62E+06 | 53,246 | 0 | 18,866 | 24 |
| 5,84E+07 | 4,37E+06 | 4,40E+06 | 1,08E+07 | 39,203 | 0 | 16,18 | 28 |

| | | | | | | | |
|---|---|---|---|---|---|---|---|
| 9,99E+07 | 6,49E+07 | 5,26E+07 | 5,54E+07 | 134,36 | 0 | 67,745 | 78 |
| 2,85E+08 | 1,89E+08 | 2,04E+08 | 1,99E+08 | 32,379 | 0 | 323,31 | 67 |
| 5,77E+07 | 5,32E+06 | 2,10E+06 | 6,34E+06 | 92,555 | 0 | 13,598 | 33 |
| 7,90E+08 | 3,50E+08 | 3,99E+08 | 3,91E+08 | 86,041 | 0,008982 | 1,4614 | 19 |
| 4,43E+07 | 2,42E+07 | 2,72E+07 | 2,09E+07 | 102,73 | 0 | 12,86 | 18 |
| 4,67E+09 | 3,69E+09 | 3,29E+09 | 3,43E+09 | 31,062 | 0 | 323,31 | 448 |
| 5,85E+07 | 3,73E+07 | 4,29E+07 | 4,10E+07 | 103,67 | 0 | 21,959 | 54 |
| 1,44E+08 | 2,57E+06 | 6,79E+05 | 4,25E+06 | 37,402 | 0 | 225,96 | 39 |
| 1,36E+08 | 2,00E+08 | 2,04E+08 | 2,12E+08 | 50,545 | 0 | 159,48 | 69 |
| 1,46E+08 | 9,59E+07 | 9,74E+07 | 8,54E+07 | 54,495 | 0 | 75,108 | 66 |
| 4,29E+07 | 1,47E+06 | 1,67E+06 | 4,60E+06 | 20,123 | 0,000156 | 4,6591 | 16 |
| 5,55E+07 | 1,17E+08 | 1,08E+08 | 1,09E+08 | 21,618 | 0 | 52,285 | 40 |
| 9,61E+07 | 1,59E+08 | 1,63E+08 | 1,72E+08 | 80,363 | 0 | 100,63 | 76 |
| 1,71E+08 | 1,44E+08 | 1,42E+08 | 1,50E+08 | 190,76 | 0 | 48,285 | 144 |
| 2,64E+07 | 2,20E+07 | 2,19E+07 | 2,22E+07 | 115,31 | 0 | 15,968 | 24 |
| 2,77E+08 | 1,61E+08 | 1,87E+08 | 1,78E+08 | 52,755 | 0 | 133,37 | 69 |
| 1,05E+09 | 1,61E+09 | 1,48E+09 | 1,49E+09 | 108,85 | 0 | 323,31 | 333 |
| 2,74E+07 | 2,42E+08 | 1,93E+08 | 2,42E+08 | 42,663 | 0 | 203,55 | 28 |
| 3,93E+06 | 7,31E+06 | 6,76E+06 | 7,51E+06 | 23,766 | 0,002096 | 2,155 | 4 |
| 2,32E+08 | 1,68E+08 | 1,62E+08 | 1,56E+08 | 44,127 | 0 | 106,48 | 74 |
| 7,85E+07 | 4,55E+07 | 4,61E+07 | 5,17E+07 | 60,775 | 0 | 65,37 | 34 |
| 7,66E+07 | 6,08E+07 | 5,78E+07 | 6,04E+07 | 47,357 | 0 | 19,161 | 44 |
| 9,39E+07 | 1,35E+06 | 3,24E+06 | 5,14E+06 | 55,446 | 0 | 50,822 | 25 |
| 3,34E+07 | 1,56E+07 | 1,60E+07 | 1,13E+07 | 39,508 | 0 | 11,454 | 22 |
| 3,95E+07 | 3,43E+07 | 3,31E+07 | 3,20E+07 | 41,983 | 0 | 20,386 | 31 |
| 2,04E+07 | 9,74E+05 | 2,62E+06 | 2,01E+06 | 67,953 | 0 | 14,524 | 17 |
| 1,62E+08 | 1,29E+08 | 1,15E+08 | 1,17E+08 | 46,201 | 0 | 31,918 | 50 |
| 1,03E+09 | 8,54E+08 | 8,39E+08 | 8,81E+08 | 197,24 | 0 | 323,31 | 379 |
| 4,49E+08 | 3,17E+08 | 3,16E+08 | 2,96E+08 | 28,543 | 0 | 57,68 | 82 |
| 4,89E+08 | 6,11E+08 | 6,33E+08 | 6,22E+08 | 102,36 | 0 | 323,31 | 186 |
| 7,77E+06 | 1,62E+06 | 2,11E+06 | 2,94E+06 | 49,198 | 0,000149 | 3,551 | 5 |
| 6,68E+07 | 3,55E+07 | 2,86E+07 | 2,62E+07 | 40,751 | 0 | 38,323 | 36 |
| 3,69E+07 | 6,01E+07 | 6,64E+07 | 7,26E+07 | 42,63 | 0 | 58,169 | 38 |
| 6,25E+08 | 4,38E+08 | 4,92E+08 | 4,70E+08 | 181,43 | 0 | 323,31 | 205 |
| 1,41E+08 | 2,16E+08 | 2,23E+08 | 1,88E+08 | 189,08 | 0 | 61,528 | 145 |
| 1,29E+07 | 5,44E+06 | 6,91E+06 | 6,21E+06 | 92,185 | 0 | 6,5029 | 12 |
| 2,67E+08 | 3,89E+08 | 3,79E+08 | 3,56E+08 | 123,97 | 0 | 131,14 | 183 |
| 7,36E+08 | 1,25E+09 | 1,14E+09 | 1,18E+09 | 69,036 | 0 | 314,95 | 203 |
| 6,07E+07 | 1,09E+08 | 9,95E+07 | 1,17E+08 | 40,143 | 0 | 71,261 | 56 |
| 2,67E+07 | 6,84E+07 | 6,54E+07 | 6,01E+07 | 65,804 | 0 | 26,335 | 56 |
| 4,14E+06 | 3,34E+07 | 3,75E+07 | 4,37E+07 | 18,136 | 0 | 6,9938 | 7 |
| 1,13E+08 | 8,76E+07 | 7,51E+07 | 8,09E+07 | 126,4 | 0 | 66,136 | 96 |
| 1,04E+09 | 1,25E+09 | 1,21E+09 | 1,23E+09 | 572,28 | 0 | 323,31 | 618 |
| 1,87E+08 | 2,40E+08 | 2,27E+08 | 2,39E+08 | 121,19 | 0 | 114,64 | 143 |
| 6,04E+07 | 1,32E+08 | 1,47E+08 | 1,46E+08 | 191,96 | 0 | 109,66 | 96 |
| 8,86E+09 | 2,29E+06 | 4,19E+07 | 1,81E+07 | 47,538 | 0 | 323,31 | 369 |
| 1,40E+07 | 2,54E+06 | 3,13E+06 | 2,09E+06 | 41,9 | 0 | 9,1054 | 8 |
| 1,24E+08 | 2,42E+08 | 2,16E+08 | 2,54E+08 | 74,028 | 0 | 119,05 | 91 |
| 3,59E+08 | 2,50E+08 | 2,70E+08 | 2,53E+08 | 124,31 | 0 | 192,97 | 186 |
| 2,11E+07 | 2,08E+06 | 4,09E+06 | 4,57E+06 | 44,411 | 0 | 14,182 | 15 |

| | | | | | | | |
|---|---|---|---|---|---|---|---|
| 1,24E+07 | 6,03E+06 | 6,48E+06 | 4,54E+06 | 64,609 | 0 | 18,366 | 11 |
| 5,85E+07 | 6,77E+07 | 6,77E+07 | 6,89E+07 | 134,59 | 0 | 43,934 | 86 |
| 3,96E+07 | 7,31E+07 | 8,26E+07 | 8,06E+07 | 142,88 | 0 | 24,082 | 36 |
| 3,53E+08 | 4,81E+08 | 4,85E+08 | 5,01E+08 | 72,89 | 0 | 323,31 | 187 |
| 2,33E+07 | 6,12E+07 | 6,52E+07 | 5,77E+07 | 31,966 | 0 | 18,258 | 29 |
| 2,59E+07 | 4,10E+07 | 3,99E+07 | 4,44E+07 | 87,274 | 0 | 24,651 | 45 |
| 7,07E+08 | 5,41E+08 | 5,35E+08 | 5,26E+08 | 106,72 | 0 | 323,31 | 284 |
| 2,35E+09 | 1,50E+09 | 1,44E+09 | 1,37E+09 | 39,632 | 0 | 323,31 | 310 |
| 3,76E+08 | 5,06E+08 | 5,37E+08 | 5,23E+08 | 40,243 | 0 | 323,31 | 105 |
| 7,15E+08 | 8,80E+08 | 8,22E+08 | 8,61E+08 | 56,613 | 0 | 323,31 | 166 |
| 2,18E+08 | 3,08E+08 | 3,34E+08 | 2,90E+08 | 85,268 | 0 | 242,09 | 154 |
| 8,54E+08 | 7,14E+08 | 7,33E+08 | 7,12E+08 | 43,231 | 0 | 198,25 | 131 |
| 1,71E+07 | 8,78E+07 | 8,85E+07 | 9,83E+07 | 41,704 | 0 | 18,505 | 40 |
| 2,65E+08 | 3,80E+08 | 3,60E+08 | 3,74E+08 | 34,909 | 0 | 323,31 | 112 |
| 7,45E+06 | 3,24E+06 | 2,00E+06 | 2,31E+06 | 49,967 | 0 | 5,4708 | 12 |
| 2,70E+07 | 1,24E+08 | 1,54E+08 | 1,54E+08 | 78,359 | 0 | 97,443 | 75 |
| 1,01E+08 | 6,53E+07 | 5,74E+07 | 6,82E+07 | 97,075 | 0 | 50,451 | 60 |
| 6,44E+07 | 9,07E+07 | 8,32E+07 | 8,05E+07 | 101,44 | 0 | 36,318 | 78 |
| 4,21E+06 | 6,60E+06 | 7,86E+06 | 8,66E+06 | 23,705 | 0,000432 | 2,8239 | 8 |
| 6,05E+08 | 4,14E+08 | 3,86E+08 | 3,52E+08 | 106,86 | 0 | 154,5 | 84 |
| 4,17E+06 | 2,40E+07 | 1,85E+07 | 2,03E+07 | 85,411 | 0 | 11,191 | 14 |
| 2,90E+08 | 2,09E+08 | 1,95E+08 | 2,18E+08 | 70,367 | 0 | 196,63 | 132 |
| 1,24E+08 | 1,90E+08 | 2,25E+08 | 2,33E+08 | 19,116 | 0 | 42,19 | 58 |
| 9,74E+06 | 1,44E+07 | 1,58E+07 | 1,71E+07 | 77,279 | 0 | 7,8314 | 20 |
| 3,51E+07 | 9,36E+07 | 1,03E+08 | 1,06E+08 | 71,651 | 0 | 29,497 | 50 |
| 1,75E+07 | 2,03E+07 | 1,95E+07 | 2,06E+07 | 38,883 | 0 | 5,568 | 20 |
| 6,17E+07 | 1,01E+08 | 1,02E+08 | 1,04E+08 | 36,464 | 0 | 46,308 | 44 |
| 1,41E+08 | 8,68E+07 | 9,89E+07 | 8,69E+07 | 62,607 | 0 | 89,502 | 79 |
| 1,80E+08 | 2,85E+08 | 3,21E+08 | 2,81E+08 | 67,568 | 0 | 105,67 | 156 |
| 4,98E+08 | 3,28E+08 | 3,25E+08 | 3,53E+08 | 135 | 0 | 276,26 | 151 |
| 5,46E+08 | 4,54E+08 | 4,28E+08 | 4,14E+08 | 40,718 | 0 | 323,31 | 153 |
| 5,87E+07 | 8,19E+07 | 8,06E+07 | 7,30E+07 | 38,452 | 0 | 127,39 | 46 |
| 9,52E+06 | 2,02E+07 | 2,63E+07 | 2,79E+07 | 27,952 | 0 | 6,4289 | 13 |
| 2,96E+06 | 1,01E+07 | 8,68E+06 | 9,30E+06 | 57,031 | 0,000155 | 4,5913 | 11 |
| 1,13E+08 | 5,91E+07 | 4,69E+07 | 5,81E+07 | 31,595 | 0 | 19,134 | 65 |
| 1,14E+08 | 1,56E+08 | 1,39E+08 | 1,51E+08 | 30,367 | 0 | 88,133 | 67 |
| 1,88E+08 | 1,28E+08 | 1,17E+08 | 1,10E+08 | 42,215 | 0 | 31,013 | 83 |
| 1,10E+08 | 1,51E+08 | 1,60E+08 | 1,51E+08 | 215,92 | 0 | 150,83 | 135 |
| 5,18E+07 | 7,75E+06 | 2,28E+06 | 3,59E+06 | 53,617 | 0 | 51,508 | 16 |
| 1,78E+07 | 4,45E+06 | 1,96E+06 | 2,75E+06 | 45,043 | 0 | 17,553 | 10 |
| 2,00E+07 | 2,29E+06 | 4,33E+06 | 3,38E+06 | 20,548 | 0 | 6,2935 | 11 |
| 1,90E+09 | 1,45E+09 | 1,53E+09 | 1,45E+09 | 85,359 | 0 | 323,31 | 331 |
| 1,03E+08 | 2,79E+08 | 3,23E+08 | 2,72E+08 | 21,71 | 0 | 15,952 | 54 |
| 1,32E+07 | 5,17E+07 | 4,86E+07 | 4,82E+07 | 47,833 | 0 | 20,685 | 30 |
| 3,39E+08 | 2,69E+08 | 2,74E+08 | 2,74E+08 | 67,333 | 0 | 191,88 | 157 |
| 6,78E+07 | 3,49E+06 | 1,50E+06 | 6,28E+06 | 121,15 | 0 | 30,727 | 41 |
| 1,82E+08 | 2,87E+08 | 2,80E+08 | 3,24E+08 | 85,651 | 0 | 181,36 | 110 |
| 3,08E+07 | 1,91E+07 | 2,21E+07 | 2,01E+07 | 41,398 | 0 | 58,639 | 13 |
| 1,65E+08 | 3,07E+08 | 3,42E+08 | 3,78E+08 | 28,721 | 0 | 47,417 | 56 |
| 2,87E+07 | 7,35E+07 | 6,84E+07 | 7,85E+07 | 43,81 | 0 | 12,742 | 47 |

| | | | | | | | |
|---|---|---|---|---|---|---|---|
| 1,50E+08 | 9,57E+07 | 8,95E+07 | 9,08E+07 | 59,445 | 0 | 89,304 | 60 |
| 1,73E+09 | 3,70E+09 | 3,79E+09 | 3,94E+09 | 40,472 | 0 | 300,38 | 286 |
| 4,91E+07 | 9,47E+05 | 2,77E+06 | 4,05E+06 | 70,02 | 0 | 14,395 | 17 |
| 7,58E+08 | 1,55E+09 | 1,28E+09 | 1,41E+09 | 36,511 | 0 | 323,31 | 117 |
| 3,92E+08 | 2,66E+08 | 2,79E+08 | 2,64E+08 | 37,436 | 0 | 51,234 | 118 |
| 8,26E+09 | 6,74E+09 | 7,11E+09 | 6,86E+09 | 272,43 | 0 | 323,31 | 1243 |
| 3,31E+08 | 2,31E+06 | 8,84E+06 | 3,60E+06 | 39,947 | 0 | 80,649 | 39 |
| 3,43E+07 | 4,81E+06 | 4,95E+06 | 9,88E+06 | 59,573 | 0 | 7,8904 | 15 |
| 1,64E+07 | 4,62E+06 | 2,37E+06 | 3,61E+06 | 61,987 | 0,000159 | 5,2713 | 10 |
| 8,31E+07 | 4,69E+07 | 4,30E+07 | 5,25E+07 | 41,709 | 0 | 21,831 | 43 |
| 2,64E+08 | 4,63E+08 | 3,95E+08 | 4,31E+08 | 62,289 | 0 | 36,807 | 72 |
| 3,90E+07 | 4,89E+07 | 5,28E+07 | 5,05E+07 | 55,865 | 0 | 41,654 | 35 |
| 5,93E+07 | 4,32E+06 | 3,81E+06 | 1,78E+06 | 15,404 | 0 | 71,328 | 5 |
| 7,95E+07 | 3,60E+06 | 3,38E+06 | 1,11E+07 | 46,169 | 0 | 15,226 | 29 |
| 2,38E+07 | 1,55E+07 | 1,59E+07 | 1,49E+07 | 77,774 | 0 | 15,98 | 33 |
| 1,75E+07 | 4,74E+06 | 5,64E+06 | 5,94E+06 | 107,83 | 0 | 8,3346 | 14 |
| 1,95E+07 | 3,82E+07 | 3,62E+07 | 3,68E+07 | 14,83 | 0 | 6,9973 | 28 |
| 1,59E+09 | 1,89E+09 | 1,91E+09 | 1,83E+09 | 85,451 | 0 | 323,31 | 471 |
| 2,70E+08 | 4,90E+08 | 4,51E+08 | 4,97E+08 | 160,58 | 0 | 323,31 | 210 |
| 1,64E+08 | 2,30E+08 | 2,59E+08 | 2,26E+08 | 72,05 | 0 | 163,58 | 119 |
| 3,69E+08 | 3,96E+08 | 4,08E+08 | 3,98E+08 | 71,721 | 0 | 237,92 | 129 |
| 2,03E+10 | 2,94E+10 | 3,30E+10 | 2,99E+10 | 47,14 | 0 | 323,31 | 1113 |
| 2,80E+09 | 1,55E+09 | 1,61E+09 | 1,43E+09 | 22,395 | 0 | 323,31 | 547 |
| 7,49E+07 | 4,38E+07 | 5,12E+07 | 4,53E+07 | 38,725 | 0 | 75,79 | 46 |
| 2,64E+08 | 7,94E+07 | 1,10E+08 | 1,11E+08 | 135,5 | 0 | 18,529 | 31 |
| 6,51E+07 | 3,75E+06 | 4,83E+06 | 1,21E+06 | 79,916 | 0 | 13,939 | 30 |
| 7,46E+06 | 1,74E+07 | 1,80E+07 | 1,62E+07 | 38,399 | 0 | 10,075 | 17 |
| 7,30E+07 | 1,30E+08 | 1,64E+08 | 1,60E+08 | 30,959 | 0 | 24,578 | 40 |
| 1,03E+08 | 1,24E+08 | 1,29E+08 | 1,20E+08 | 103,7 | 0 | 45,225 | 74 |
| 3,28E+08 | 1,26E+08 | 1,14E+08 | 1,16E+08 | 100,29 | 0 | 323,31 | 109 |
| 6,08E+07 | 9,89E+07 | 9,26E+07 | 8,48E+07 | 75,023 | 0 | 72,122 | 53 |
| 2,53E+06 | 6,86E+06 | 1,04E+07 | 1,09E+07 | 37,574 | 0,000156 | 4,6327 | 13 |
| 8,50E+07 | 1,21E+08 | 1,16E+08 | 1,11E+08 | 108,14 | 0 | 112,81 | 95 |
| 1,68E+08 | 1,13E+08 | 1,15E+08 | 1,19E+08 | 217,36 | 0 | 171,32 | 125 |
| 5,87E+07 | 4,73E+06 | 4,02E+06 | 1,30E+06 | 47,123 | 0 | 17,151 | 27 |
| 1,37E+07 | 1,30E+06 | 2,73E+06 | 1,32E+06 | 72,227 | 0 | 10,785 | 20 |
| 3,51E+08 | 6,76E+08 | 6,06E+08 | 5,33E+08 | 27,883 | 0 | 323,31 | 143 |
| 1,83E+08 | 2,34E+08 | 2,43E+08 | 2,26E+08 | 64,016 | 0 | 245,51 | 132 |
| 6,31E+07 | 1,24E+08 | 1,38E+08 | 1,20E+08 | 77,519 | 0 | 46,252 | 82 |
| 9,15E+07 | 1,58E+08 | 1,70E+08 | 1,76E+08 | 42,88 | 0 | 92,032 | 57 |
| 1,66E+06 | 2,70E+07 | 2,23E+07 | 2,84E+07 | 58,688 | 0 | 12,744 | 14 |
| 5,91E+07 | 1,14E+08 | 1,15E+08 | 9,25E+07 | 104,2 | 0 | 105,92 | 52 |
| 2,03E+06 | 1,53E+07 | 1,15E+07 | 1,63E+07 | 80,118 | 0 | 7,01 | 10 |
| 1,45E+08 | 1,92E+08 | 2,11E+08 | 1,83E+08 | 217,29 | 0 | 55,303 | 159 |
| 2,95E+08 | 2,34E+08 | 2,40E+08 | 2,46E+08 | 62,981 | 0 | 142,43 | 91 |
| 7,61E+07 | 3,44E+07 | 4,25E+07 | 4,54E+07 | 128,43 | 0 | 37,19 | 57 |
| 2,25E+07 | 1,66E+07 | 1,65E+07 | 1,59E+07 | 76,824 | 0 | 16,82 | 24 |
| 8,90E+09 | 1,05E+10 | 1,04E+10 | 1,06E+10 | 191,55 | 0 | 323,31 | 1291 |
| 8,73E+08 | 6,16E+08 | 6,12E+08 | 6,18E+08 | 109,62 | 0 | 323,31 | 288 |
| 1,16E+08 | 1,79E+08 | 1,71E+08 | 1,95E+08 | 56,794 | 0 | 67,473 | 79 |

| | | | | | | | |
|---|---|---|---|---|---|---|---|
| 7,93E+07 | 4,46E+07 | 3,87E+07 | 3,93E+07 | 120,31 | 0 | 73,55 | 61 |
| 7,54E+07 | 5,11E+07 | 5,69E+07 | 5,07E+07 | 77,256 | 0 | 29,717 | 41 |
| 4,48E+08 | 8,18E+08 | 8,42E+08 | 9,02E+08 | 33,189 | 0 | 323,31 | 168 |
| 3,99E+06 | 2,57E+06 | 2,28E+06 | 2,52E+06 | 26,923 | 0,000854 | 2,5848 | 9 |
| 2,66E+07 | 1,57E+07 | 1,29E+07 | 1,44E+07 | 97,721 | 0 | 42,105 | 25 |
| 1,71E+07 | 3,43E+06 | 2,30E+06 | 3,87E+06 | 23,934 | 0,000147 | 3,2901 | 12 |
| 1,21E+08 | 7,70E+07 | 5,73E+07 | 6,59E+07 | 86,721 | 0 | 45,955 | 70 |
| 9,88E+07 | 1,65E+08 | 1,53E+08 | 1,68E+08 | 26,248 | 0 | 75,516 | 50 |
| 2,18E+08 | 1,69E+08 | 1,48E+08 | 1,47E+08 | 81,084 | 0 | 154,29 | 91 |
| 2,70E+07 | 6,90E+07 | 7,03E+07 | 6,56E+07 | 22,08 | 0 | 106,2 | 32 |
| 1,15E+07 | 2,43E+07 | 1,92E+07 | 2,01E+07 | 39,154 | 0 | 11,478 | 21 |
| 1,25E+08 | 8,48E+07 | 8,24E+07 | 9,32E+07 | 44,092 | 0 | 105,14 | 60 |
| 3,53E+07 | 5,23E+07 | 4,87E+07 | 4,84E+07 | 51,126 | 0 | 14,339 | 48 |
| 2,76E+09 | 1,71E+09 | 1,91E+09 | 1,63E+09 | 112,98 | 0 | 323,31 | 415 |
| 5,24E+08 | 7,14E+08 | 7,72E+08 | 7,09E+08 | 11,101 | 0 | 88,844 | 64 |
| 3,06E+07 | 4,72E+07 | 4,80E+07 | 5,30E+07 | 291,46 | 0 | 21,768 | 50 |
| 9,75E+07 | 1,73E+08 | 1,70E+08 | 1,69E+08 | 10,425 | 0 | 229,29 | 9 |
| 8,18E+07 | 6,86E+07 | 6,87E+07 | 6,61E+07 | 155,79 | 0 | 34,96 | 75 |
| 6,93E+07 | 5,09E+07 | 5,21E+07 | 5,68E+07 | 86,518 | 0 | 33,811 | 64 |
| 2,51E+07 | 1,36E+07 | 1,02E+07 | 1,05E+07 | 328,8 | 0 | 29,461 | 28 |
| 6,45E+07 | 4,70E+07 | 5,03E+07 | 4,30E+07 | 79,573 | 0 | 68,067 | 55 |
| 6,89E+07 | 4,73E+07 | 5,05E+07 | 4,88E+07 | 123,1 | 0 | 24,37 | 56 |
| 2,30E+08 | 3,33E+08 | 3,26E+08 | 3,56E+08 | 29,174 | 0 | 113,53 | 75 |
| 6,83E+07 | 1,05E+08 | 9,54E+07 | 1,03E+08 | 36,793 | 0 | 131,7 | 56 |
| 3,48E+08 | 3,14E+08 | 2,95E+08 | 3,08E+08 | 149,58 | 0 | 223,69 | 194 |
| 6,88E+07 | 1,01E+07 | 5,70E+06 | 1,51E+07 | 47,379 | 0 | 29,905 | 34 |
| 6,45E+06 | 4,26E+06 | 4,64E+06 | 4,70E+06 | 38,966 | 0,000149 | 3,5088 | 11 |
| 5,73E+07 | 1,26E+08 | 1,10E+08 | 1,10E+08 | 38,961 | 0 | 27,444 | 68 |
| 7,87E+08 | 5,60E+08 | 6,23E+08 | 6,18E+08 | 71,06 | 0 | 323,31 | 205 |
| 2,73E+08 | 3,53E+08 | 3,97E+08 | 3,87E+08 | 29,718 | 0 | 113,22 | 99 |
| 2,22E+08 | 1,54E+08 | 1,51E+08 | 1,39E+08 | 82,175 | 0 | 194,8 | 132 |
| 3,89E+07 | 3,18E+06 | 8,81E+06 | 4,62E+06 | 44,357 | 0 | 19,764 | 27 |
| 1,44E+08 | 4,99E+07 | 4,79E+07 | 2,91E+07 | 153,8 | 0 | 91,172 | 69 |
| 9,63E+07 | 5,97E+07 | 6,08E+07 | 6,05E+07 | 37,335 | 0 | 38,093 | 69 |
| 6,64E+07 | 3,12E+07 | 3,88E+07 | 3,69E+07 | 96,683 | 0 | 50,501 | 54 |
| 1,25E+07 | 2,03E+07 | 1,91E+07 | 2,07E+07 | 18,187 | 0,000158 | 5,0544 | 9 |
| 1,61E+07 | 4,02E+07 | 4,53E+07 | 4,65E+07 | 51,996 | 0 | 32,024 | 36 |
| 1,79E+08 | 2,44E+08 | 2,33E+08 | 2,24E+08 | 103,22 | 0 | 191,48 | 158 |
| 5,32E+08 | 3,85E+08 | 4,18E+08 | 4,18E+08 | 32,562 | 0 | 124,23 | 54 |
| 1,67E+09 | 2,66E+09 | 2,58E+09 | 2,49E+09 | 41,655 | 0 | 229,5 | 210 |
| 3,31E+08 | 5,02E+08 | 5,04E+08 | 4,68E+08 | 58,408 | 0 | 160,23 | 125 |
| 3,93E+07 | 2,40E+07 | 2,64E+07 | 2,16E+07 | 162,54 | 0 | 24,077 | 45 |
| 6,84E+09 | 5,08E+09 | 5,57E+09 | 5,30E+09 | 50,976 | 0 | 323,31 | 922 |
| 5,69E+06 | 2,61E+07 | 3,15E+07 | 3,09E+07 | 60,305 | 0 | 13,07 | 30 |
| 3,18E+07 | 7,18E+07 | 6,48E+07 | 7,84E+07 | 52,347 | 0 | 21,567 | 52 |
| 4,15E+07 | 3,39E+07 | 3,29E+07 | 3,52E+07 | 406,57 | 0 | 15,124 | 32 |
| 6,99E+06 | 3,24E+06 | 3,94E+06 | 4,00E+06 | 26,035 | 0,000429 | 2,7253 | 6 |
| 8,72E+06 | 5,39E+07 | 5,73E+07 | 6,29E+07 | 138,82 | 0 | 17,81 | 30 |
| 1,53E+09 | 1,15E+09 | 1,12E+09 | 1,12E+09 | 207,78 | 0 | 323,31 | 505 |
| 4,73E+07 | 4,42E+06 | 2,60E+06 | 7,59E+06 | 110,24 | 0 | 36,678 | 31 |

| | | | | | | | |
|---|---|---|---|---|---|---|---|
| 1,38E+07 | 9,11E+06 | 8,97E+06 | 9,07E+06 | 104,58 | 0 | 6,2601 | 22 |
| 8,25E+08 | 5,24E+08 | 4,71E+08 | 4,61E+08 | 87,869 | 0 | 323,31 | 241 |
| 1,40E+08 | 1,22E+08 | 1,25E+08 | 1,27E+08 | 87,046 | 0 | 278,36 | 89 |
| 1,87E+08 | 2,24E+08 | 2,20E+08 | 2,41E+08 | 57,611 | 0 | 58,13 | 126 |
| 8,86E+06 | 3,35E+07 | 3,66E+07 | 4,51E+07 | 37,798 | 0 | 7,6203 | 13 |
| 2,59E+09 | 3,01E+09 | 3,10E+09 | 2,94E+09 | 81,21 | 0 | 323,31 | 437 |
| 1,26E+07 | 2,17E+07 | 2,21E+07 | 2,57E+07 | 22,241 | 0 | 21,272 | 6 |
| 1,43E+08 | 1,06E+08 | 9,16E+07 | 9,97E+07 | 32,851 | 0 | 34,848 | 52 |
| 9,76E+08 | 8,79E+08 | 8,63E+08 | 8,60E+08 | 46,284 | 0 | 323,31 | 235 |
| 1,06E+08 | 6,46E+07 | 6,96E+07 | 7,65E+07 | 78,346 | 0 | 26,807 | 78 |
| 4,46E+08 | 2,76E+08 | 3,15E+08 | 3,26E+08 | 34,785 | 0 | 250,44 | 111 |
| 4,45E+07 | 7,59E+07 | 7,51E+07 | 7,33E+07 | 46,551 | 0 | 16,472 | 39 |
| 8,66E+07 | 5,97E+06 | 5,29E+06 | 1,48E+06 | 33,931 | 0 | 12,72 | 20 |
| 3,60E+08 | 7,06E+08 | 6,57E+08 | 6,37E+08 | 24,603 | 0 | 216,1 | 61 |
| 1,30E+07 | 6,86E+06 | 6,16E+06 | 5,48E+06 | 51,719 | 0 | 7,28 | 14 |
| 1,58E+08 | 1,14E+08 | 1,02E+08 | 1,02E+08 | 168,08 | 0 | 77,079 | 133 |
| 2,57E+07 | 1,12E+07 | 1,07E+07 | 1,39E+07 | 53,517 | 0 | 95,237 | 23 |
| 1,36E+08 | 6,46E+06 | 7,90E+06 | 3,08E+06 | 39,351 | 0 | 125,74 | 34 |
| 1,76E+07 | 1,03E+07 | 1,17E+07 | 1,13E+07 | 35,024 | 0 | 9,5821 | 17 |
| 6,87E+08 | 2,18E+08 | 2,38E+08 | 2,19E+08 | 27,057 | 0 | 323,31 | 105 |
| 2,79E+07 | 1,08E+07 | 8,55E+06 | 1,34E+07 | 80,088 | 0 | 31,627 | 30 |
| 3,80E+07 | 2,30E+07 | 2,33E+07 | 1,83E+07 | 52,603 | 0 | 70,221 | 55 |
| 9,15E+07 | 5,56E+07 | 5,77E+07 | 5,52E+07 | 44,721 | 0 | 28,475 | 45 |
| 1,22E+08 | 4,09E+06 | 1,20E+07 | 5,90E+06 | 27,189 | 0 | 110,82 | 13 |
| 3,33E+07 | 1,81E+07 | 2,07E+07 | 2,11E+07 | 32,542 | 0 | 52,741 | 26 |
| 1,33E+08 | 2,08E+08 | 1,94E+08 | 1,98E+08 | 110,54 | 0 | 163,03 | 139 |
| 3,40E+07 | 4,80E+07 | 4,99E+07 | 5,05E+07 | 58,326 | 0 | 26,128 | 63 |
| 2,00E+08 | 2,21E+08 | 2,16E+08 | 2,26E+08 | 119,24 | 0 | 134,2 | 132 |
| 2,58E+08 | 1,84E+08 | 1,94E+08 | 1,76E+08 | 139,34 | 0 | 67,678 | 142 |
| 2,68E+08 | 4,39E+08 | 3,88E+08 | 4,22E+08 | 98,577 | 0 | 323,31 | 136 |
| 5,58E+06 | 3,19E+07 | 2,81E+07 | 2,77E+07 | 75,732 | 0 | 7,4013 | 20 |
| 4,03E+08 | 2,54E+08 | 2,50E+08 | 2,74E+08 | 81,58 | 0 | 150,5 | 115 |
| 3,46E+08 | 2,85E+08 | 2,54E+08 | 2,72E+08 | 123,05 | 0 | 231,22 | 144 |
| 9,22E+07 | 3,69E+07 | 2,92E+07 | 3,43E+07 | 27,807 | 0 | 19,028 | 27 |
| 4,03E+08 | 2,46E+08 | 2,29E+08 | 2,41E+08 | 10,071 | 0 | 54,954 | 31 |
| 2,93E+08 | 3,53E+08 | 3,74E+08 | 3,68E+08 | 65,89 | 0 | 323,31 | 164 |
| 1,50E+08 | 2,71E+08 | 2,70E+08 | 2,56E+08 | 33,601 | 0 | 84,89 | 98 |
| 9,38E+07 | 3,91E+06 | 2,07E+06 | 3,26E+06 | 204 | 0 | 323,31 | 177 |
| 2,13E+09 | 1,29E+09 | 1,43E+09 | 1,52E+09 | 15,137 | 0 | 323,31 | 175 |
| 8,86E+07 | 1,21E+08 | 1,07E+08 | 1,17E+08 | 133,93 | 0 | 51,899 | 76 |
| 7,83E+07 | 6,41E+06 | 5,43E+06 | 1,71E+06 | 74,172 | 0 | 115,14 | 28 |
| 1,11E+08 | 6,84E+07 | 6,98E+07 | 6,34E+07 | 41,383 | 0 | 43,664 | 67 |
| 4,94E+07 | 9,23E+06 | 3,88E+06 | 4,46E+06 | 111,67 | 0 | 22,104 | 28 |
| 1,73E+09 | 1,37E+09 | 1,32E+09 | 1,31E+09 | 63,691 | 0 | 323,31 | 305 |
| 4,01E+08 | 1,32E+08 | 1,59E+08 | 1,39E+08 | 8,4689 | 0 | 15,866 | 87 |
| 1,30E+09 | 1,71E+09 | 1,66E+09 | 1,57E+09 | 76,221 | 0 | 323,31 | 322 |
| 4,29E+08 | 5,19E+08 | 5,64E+08 | 5,29E+08 | 65,934 | 0 | 300,33 | 141 |
| 3,31E+08 | 2,09E+08 | 2,35E+08 | 2,50E+08 | 45,437 | 0 | 323,31 | 109 |
| 1,48E+07 | 4,46E+06 | 6,13E+06 | 5,47E+06 | 63,036 | 0 | 27,235 | 13 |
| 7,08E+06 | 2,78E+08 | 2,82E+08 | 2,99E+08 | 23,802 | 0 | 14,831 | 45 |

| | | | | | | | |
|---|---|---|---|---|---|---|---|
| 3,08E+06 | 3,15E+07 | 2,81E+07 | 2,54E+07 | 29,774 | 0 | 11,558 | 23 |
| 9,11E+07 | 5,06E+07 | 5,67E+07 | 5,92E+07 | 15,531 | 0 | 23,617 | 46 |
| 3,62E+06 | 1,97E+07 | 1,95E+07 | 1,60E+07 | 13,719 | 0,000435 | 2,9803 | 4 |
| 3,92E+07 | 3,09E+06 | 1,45E+06 | 5,37E+06 | 50,254 | 0 | 21,242 | 21 |
| 2,73E+06 | 3,72E+07 | 3,64E+07 | 3,50E+07 | 20,814 | 0 | 5,348 | 17 |
| 4,42E+07 | 2,88E+07 | 3,36E+07 | 3,03E+07 | 101,69 | 0 | 26,993 | 55 |
| 1,81E+08 | 1,51E+08 | 1,51E+08 | 1,37E+08 | 39,344 | 0 | 55,865 | 75 |
| 5,50E+06 | 2,76E+06 | 2,70E+06 | 2,53E+06 | 79,006 | 0,000157 | 4,9415 | 8 |
| 2,74E+07 | 3,98E+06 | 3,62E+06 | 1,59E+06 | 57,713 | 0 | 8,5727 | 11 |
| 4,10E+08 | 7,02E+08 | 7,44E+08 | 8,21E+08 | 27,855 | 0 | 183,16 | 86 |
| 3,81E+08 | 5,54E+08 | 4,96E+08 | 5,58E+08 | 40,148 | 0 | 173,65 | 103 |
| 4,44E+07 | 8,39E+07 | 8,09E+07 | 8,56E+07 | 30,708 | 0 | 63,887 | 46 |
| 2,71E+07 | 1,16E+07 | 8,78E+06 | 1,30E+07 | 131,72 | 0 | 18,087 | 36 |
| 3,51E+07 | 2,71E+06 | 6,46E+06 | 3,72E+06 | 33,026 | 0 | 26,76 | 29 |
| 5,62E+06 | 9,37E+06 | 1,01E+07 | 9,79E+06 | 24,286 | 0,000148 | 3,3648 | 8 |
| 2,02E+07 | 9,22E+06 | 1,20E+07 | 9,99E+06 | 23,384 | 0 | 5,7727 | 8 |
| 6,70E+08 | 8,50E+08 | 9,34E+08 | 8,45E+08 | 36,428 | 0 | 323,31 | 202 |
| 1,08E+08 | 7,62E+07 | 7,25E+07 | 7,10E+07 | 100,46 | 0 | 104,51 | 80 |
| 2,05E+07 | 7,21E+07 | 5,04E+07 | 5,25E+07 | 54,532 | 0 | 16,444 | 29 |
| 2,87E+07 | 5,38E+06 | 2,37E+06 | 4,99E+06 | 123,87 | 0 | 16,589 | 24 |
| 3,17E+08 | 1,41E+06 | 6,26E+06 | 4,86E+05 | 22,753 | 0 | 288,05 | 56 |
| 1,68E+08 | 1,08E+08 | 1,12E+08 | 1,18E+08 | 104,84 | 0 | 122,07 | 118 |
| 9,27E+07 | 1,12E+08 | 1,24E+08 | 1,27E+08 | 96,932 | 0 | 57,724 | 111 |
| 9,37E+07 | 1,76E+08 | 1,56E+08 | 1,47E+08 | 96,481 | 0 | 103,47 | 92 |
| 4,31E+08 | 5,23E+08 | 5,06E+08 | 4,98E+08 | 128,78 | 0 | 209,45 | 279 |
| 2,77E+07 | 4,80E+07 | 4,14E+07 | 4,01E+07 | 19,947 | 0 | 27,421 | 18 |
| 1,91E+09 | 2,61E+07 | 1,56E+06 | 2,41E+06 | 74,414 | 0 | 323,31 | 155 |
| 1,72E+08 | 9,70E+06 | 1,37E+06 | 4,28E+06 | 92,725 | 0 | 91,895 | 47 |
| 8,43E+08 | 9,69E+08 | 9,46E+08 | 9,79E+08 | 59,434 | 0 | 234,41 | 228 |
| 5,24E+07 | 1,15E+07 | 1,97E+07 | 1,61E+07 | 116,34 | 0 | 37,869 | 43 |
| 1,47E+07 | 1,20E+08 | 1,84E+08 | 1,52E+08 | 80,774 | 0 | 117,29 | 57 |
| 1,65E+08 | 1,38E+08 | 1,30E+08 | 1,33E+08 | 128,21 | 0 | 111,91 | 148 |
| 2,28E+08 | 2,39E+06 | 4,09E+06 | 4,26E+06 | 75,329 | 0 | 323,31 | 76 |
| 5,99E+08 | 4,53E+08 | 4,32E+08 | 4,24E+08 | 158,23 | 0 | 323,31 | 263 |
| 1,11E+09 | 8,12E+08 | 9,13E+08 | 8,79E+08 | 69,448 | 0 | 187,21 | 227 |
| 1,23E+08 | 2,74E+08 | 2,45E+08 | 2,78E+08 | 88,25 | 0 | 124,31 | 118 |
| 7,73E+07 | 5,10E+07 | 5,29E+07 | 5,82E+07 | 119,31 | 0 | 38,035 | 74 |
| 3,96E+07 | 2,38E+07 | 2,68E+07 | 2,04E+07 | 125,22 | 0 | 13,264 | 35 |
| 3,44E+07 | 1,83E+07 | 1,37E+07 | 1,73E+07 | 55,045 | 0 | 26,567 | 28 |
| 7,28E+07 | 9,31E+07 | 8,71E+07 | 8,68E+07 | 46,172 | 0 | 142,96 | 55 |
| 8,59E+07 | 4,59E+07 | 5,18E+07 | 5,12E+07 | 71,58 | 0 | 93,311 | 75 |
| 2,16E+07 | 3,12E+07 | 3,11E+07 | 3,51E+07 | 136,95 | 0 | 23,439 | 33 |
| 5,55E+08 | 7,90E+08 | 7,90E+08 | 8,32E+08 | 48,956 | 0 | 323,31 | 83 |
| 1,39E+08 | 7,34E+07 | 6,99E+07 | 6,94E+07 | 74,929 | 0 | 56,344 | 56 |
| 2,19E+08 | 1,47E+08 | 1,54E+08 | 1,39E+08 | 29,404 | 0 | 58,472 | 43 |
| 1,60E+08 | 2,46E+08 | 2,66E+08 | 2,44E+08 | 50,858 | 0 | 142,39 | 117 |
| 2,48E+07 | 3,21E+06 | 4,60E+06 | 1,86E+06 | 84,185 | 0 | 18,116 | 8 |
| 2,20E+09 | 1,94E+09 | 1,87E+09 | 1,83E+09 | 49,184 | 0 | 323,31 | 321 |
| 4,91E+07 | 9,66E+07 | 9,01E+07 | 1,04E+08 | 24,918 | 0 | 9,0014 | 32 |
| 1,55E+08 | 2,58E+08 | 2,35E+08 | 2,71E+08 | 38,167 | 0 | 25,111 | 84 |

| | | | | | | | |
|---|---|---|---|---|---|---|---|
| 5,35E+08 | 2,98E+08 | 2,51E+08 | 2,53E+08 | 14,427 | 0 | 103,23 | 115 |
| 3,98E+08 | 2,78E+08 | 2,96E+08 | 2,93E+08 | 45,303 | 0 | 200,95 | 117 |
| 5,62E+07 | 2,62E+07 | 2,56E+07 | 2,14E+07 | 22,336 | 0 | 25,953 | 32 |
| 3,34E+09 | 2,25E+09 | 2,31E+09 | 2,18E+09 | 62,277 | 0 | 323,31 | 359 |
| 1,72E+08 | 1,01E+08 | 8,93E+07 | 7,84E+07 | 64,402 | 0 | 176,74 | 75 |
| 1,01E+08 | 1,20E+08 | 1,23E+08 | 1,22E+08 | 54,174 | 0 | 51,669 | 48 |
| 3,48E+07 | 5,45E+06 | 4,99E+06 | 1,07E+07 | 47,585 | 0 | 22,472 | 25 |
| 7,20E+09 | 5,74E+09 | 5,61E+09 | 6,11E+09 | 50,06 | 0 | 323,31 | 605 |
| 1,04E+09 | 6,57E+08 | 6,14E+08 | 5,71E+08 | 723,3 | 0 | 323,31 | 277 |
| 7,64E+07 | 7,93E+06 | 6,40E+06 | 2,09E+06 | 56,005 | 0 | 49,487 | 35 |
| 1,89E+08 | 3,67E+07 | 7,05E+07 | 4,07E+07 | 106,8 | 0 | 153,73 | 67 |
| 1,22E+07 | 4,40E+06 | 2,64E+06 | 4,31E+06 | 24,162 | 0,00291 | 2,0073 | 6 |
| 1,58E+07 | 3,56E+07 | 3,98E+07 | 3,34E+07 | 135,52 | 0 | 19,352 | 39 |
| 3,21E+08 | 2,76E+08 | 2,82E+08 | 2,75E+08 | 70,158 | 0 | 323,31 | 131 |
| 4,08E+07 | 5,46E+06 | 1,22E+06 | 2,08E+06 | 42,043 | 0 | 20,108 | 28 |
| 1,27E+07 | 2,13E+07 | 2,47E+07 | 2,90E+07 | 54,926 | 0 | 7,0387 | 22 |
| 8,08E+07 | 6,49E+07 | 5,92E+07 | 6,24E+07 | 213,86 | 0 | 61,037 | 70 |
| 1,05E+09 | 5,11E+08 | 5,15E+08 | 5,81E+08 | 20,021 | 0 | 131,13 | 154 |
| 1,16E+09 | 6,99E+08 | 6,77E+08 | 6,50E+08 | 40,429 | 0 | 147,68 | 128 |
| 1,48E+07 | 1,93E+07 | 1,75E+07 | 1,87E+07 | 62,693 | 0,000292 | 3,0746 | 8 |
| 3,28E+07 | 2,06E+06 | 4,13E+06 | 9,62E+05 | 22,875 | 0 | 5,3299 | 11 |
| 4,23E+07 | 1,26E+07 | 1,44E+07 | 1,93E+07 | 79,582 | 0 | 23,73 | 28 |
| 5,38E+06 | 2,97E+07 | 2,47E+07 | 2,18E+07 | 48,031 | 0 | 9,5185 | 17 |
| 8,67E+08 | 5,62E+08 | 5,52E+08 | 5,21E+08 | 152,48 | 0 | 323,31 | 301 |
| 9,62E+07 | 1,70E+08 | 1,79E+08 | 1,61E+08 | 26,336 | 0 | 51,551 | 75 |
| 4,25E+07 | 1,05E+07 | 1,68E+07 | 1,14E+07 | 48,605 | 0 | 24,589 | 33 |
| 6,30E+06 | 3,98E+07 | 5,78E+07 | 3,25E+07 | 54,71 | 0 | 22,644 | 20 |
| 6,01E+07 | 3,77E+07 | 4,44E+07 | 4,08E+07 | 117,55 | 0 | 44,143 | 57 |
| 1,75E+09 | 2,06E+09 | 1,95E+09 | 2,12E+09 | 79,871 | 0 | 323,31 | 297 |
| 2,85E+07 | 6,02E+07 | 5,53E+07 | 5,12E+07 | 59,323 | 0 | 13,644 | 34 |
| 2,49E+08 | 3,65E+08 | 3,35E+08 | 3,89E+08 | 11,045 | 0 | 209,88 | 99 |
| 1,57E+08 | 1,04E+08 | 9,23E+07 | 9,58E+07 | 39,291 | 0 | 29,942 | 69 |
| 3,99E+06 | 6,21E+08 | 6,25E+08 | 6,29E+08 | 101,59 | 0 | 191,68 | 121 |
| 1,85E+09 | 1,05E+09 | 1,21E+09 | 1,09E+09 | 19,724 | 0 | 43,114 | 98 |
| 9,16E+08 | 7,85E+08 | 7,97E+08 | 8,02E+08 | 101,96 | 0 | 323,31 | 251 |
| 1,52E+09 | 1,37E+09 | 1,39E+09 | 1,31E+09 | 15,423 | 0 | 58,416 | 241 |
| 1,35E+07 | 1,90E+07 | 1,65E+07 | 1,75E+07 | 77,957 | 0 | 65,692 | 20 |
| 3,13E+06 | 1,43E+07 | 9,87E+06 | 1,05E+07 | 45,571 | 0 | 9,83 | 5 |
| 7,06E+07 | 8,67E+07 | 8,27E+07 | 8,09E+07 | 59,386 | 0 | 33,532 | 76 |
| 7,79E+08 | 1,42E+09 | 1,25E+09 | 1,30E+09 | 97,255 | 0 | 87,793 | 172 |
| 1,72E+08 | 6,81E+07 | 7,44E+07 | 8,08E+07 | 23,596 | 0 | 266,88 | 48 |
| 5,12E+08 | 5,90E+08 | 6,31E+08 | 5,92E+08 | 482,63 | 0 | 323,31 | 395 |
| 1,58E+09 | 1,74E+09 | 1,71E+09 | 1,68E+09 | 60,602 | 0 | 323,31 | 376 |
| 1,67E+08 | 1,11E+08 | 8,94E+07 | 9,96E+07 | 88,75 | 0 | 116,46 | 95 |
| 4,12E+08 | 3,03E+08 | 3,00E+08 | 3,06E+08 | 82,311 | 0 | 250,77 | 143 |
| 2,92E+06 | 6,22E+07 | 6,60E+07 | 6,24E+07 | 60,438 | 0 | 12,172 | 25 |
| 3,09E+09 | 2,05E+09 | 2,19E+09 | 2,10E+09 | 176,43 | 0 | 323,31 | 636 |
| 4,54E+08 | 5,77E+08 | 5,52E+08 | 5,22E+08 | 132,6 | 0 | 323,31 | 208 |
| 1,01E+09 | 1,46E+09 | 1,38E+09 | 1,46E+09 | 33,995 | 0 | 323,31 | 233 |
| 1,37E+08 | 7,64E+07 | 7,70E+07 | 6,78E+07 | 15,572 | 0 | 69,68 | 60 |

| | | | | | | | |
|---|---|---|---|---|---|---|---|
| 2,72E+09 | 3,75E+09 | 3,80E+09 | 3,69E+09 | 61,336 | 0 | 323,31 | 391 |
| 1,68E+08 | 3,16E+08 | 3,37E+08 | 3,30E+08 | 67,942 | 0 | 178,87 | 175 |
| 9,94E+07 | 4,56E+07 | 5,10E+07 | 5,76E+07 | 50,503 | 0 | 59,694 | 38 |
| 2,30E+07 | 6,43E+06 | 2,68E+06 | 3,09E+06 | 67,017 | 0 | 24,114 | 23 |
| 1,76E+08 | 1,01E+08 | 8,51E+07 | 1,09E+08 | 95,959 | 0 | 190,63 | 84 |
| 4,16E+08 | 3,42E+08 | 3,48E+08 | 3,51E+08 | 194,11 | 0 | 323,31 | 269 |
| 6,40E+07 | 3,71E+06 | 5,78E+06 | 1,18E+07 | 95,351 | 0 | 15,212 | 30 |
| 1,73E+08 | 8,09E+06 | 1,34E+06 | 8,98E+05 | 65,091 | 0 | 38,943 | 54 |
| 9,78E+07 | 4,84E+07 | 5,05E+07 | 3,84E+07 | 80,851 | 0 | 33,783 | 64 |
| 1,40E+08 | 1,81E+08 | 1,88E+08 | 1,83E+08 | 107,14 | 0 | 136,62 | 89 |
| 4,63E+08 | 3,24E+08 | 3,34E+08 | 3,55E+08 | 53,984 | 0 | 275,84 | 68 |
| 2,67E+09 | 3,35E+09 | 3,58E+09 | 3,65E+09 | 123,09 | 0 | 323,31 | 557 |
| 7,48E+07 | 8,96E+07 | 9,42E+07 | 9,04E+07 | 81,288 | 0 | 60,049 | 59 |
| 1,74E+07 | 2,71E+06 | 4,50E+06 | 2,67E+06 | 49,64 | 0 | 6,9704 | 10 |
| 1,54E+08 | 2,06E+08 | 2,04E+08 | 1,95E+08 | 127,2 | 0 | 116,18 | 119 |
| 2,15E+07 | 1,77E+06 | 1,25E+06 | 4,02E+06 | 31,512 | 0 | 8,3307 | 11 |
| 3,21E+08 | 2,32E+08 | 2,36E+08 | 2,15E+08 | 70,046 | 0 | 207,95 | 128 |
| 5,38E+07 | 2,86E+06 | 7,30E+06 | 5,07E+06 | 54,313 | 0 | 17,9 | 20 |
| 1,36E+08 | 6,99E+07 | 8,87E+07 | 7,75E+07 | 106,25 | 0 | 105,4 | 86 |
| 2,04E+09 | 2,45E+09 | 2,36E+09 | 2,34E+09 | 83,899 | 0 | 323,31 | 304 |
| 3,24E+07 | 2,34E+07 | 2,04E+07 | 2,11E+07 | 28,847 | 0 | 60,572 | 17 |
| 2,28E+07 | 3,23E+06 | 6,17E+06 | 5,82E+06 | 43,208 | 0 | 5,6038 | 10 |
| 9,78E+07 | 8,46E+06 | 1,55E+06 | 1,95E+06 | 49,429 | 0 | 72,305 | 45 |
| 1,66E+07 | 6,81E+06 | 7,47E+06 | 6,07E+06 | 38,272 | 0 | 12,058 | 14 |
| 2,86E+08 | 2,61E+08 | 2,65E+08 | 2,69E+08 | 60,576 | 0 | 68,495 | 103 |
| 2,06E+08 | 2,29E+08 | 2,41E+08 | 2,42E+08 | 89,159 | 0 | 251,44 | 131 |
| 2,42E+07 | 3,58E+07 | 3,20E+07 | 3,35E+07 | 143,23 | 0 | 12,74 | 41 |
| 8,13E+06 | 1,32E+06 | 2,89E+06 | 1,77E+06 | 74,564 | 0 | 8,5439 | 12 |
| 2,38E+08 | 2,09E+08 | 2,12E+08 | 2,08E+08 | 199,59 | 0 | 194,71 | 122 |
| 4,48E+08 | 5,90E+08 | 6,12E+08 | 6,16E+08 | 157,45 | 0 | 323,31 | 275 |
| 7,55E+08 | 5,05E+08 | 4,85E+08 | 4,47E+08 | 108,48 | 0 | 323,31 | 189 |
| 7,63E+06 | 1,96E+06 | 3,41E+06 | 2,76E+06 | 13,643 | 0 | 18,242 | 6 |
| 4,69E+08 | 2,18E+08 | 2,74E+08 | 2,56E+08 | 59,584 | 0 | 323,31 | 98 |
| 5,55E+07 | 1,78E+06 | 7,21E+06 | 2,49E+06 | 122,38 | 0 | 26,715 | 27 |
| 1,16E+09 | 6,60E+08 | 7,72E+08 | 8,33E+08 | 60,786 | 0 | 323,31 | 199 |
| 7,73E+06 | 2,10E+07 | 2,73E+07 | 3,76E+07 | 12,974 | 0,000151 | 3,8106 | 13 |
| 8,07E+07 | 9,98E+07 | 1,02E+08 | 9,85E+07 | 42,064 | 0 | 40,107 | 63 |
| 1,79E+08 | 2,38E+08 | 2,21E+08 | 2,44E+08 | 56,99 | 0 | 213,78 | 90 |
| 1,79E+08 | 1,28E+08 | 1,10E+08 | 1,31E+08 | 38,816 | 0 | 45,261 | 60 |
| 2,97E+07 | 4,16E+07 | 3,82E+07 | 3,65E+07 | 63,473 | 0 | 17,454 | 45 |
| 1,60E+08 | 2,59E+08 | 2,77E+08 | 2,80E+08 | 50,942 | 0 | 127,97 | 133 |
| 3,01E+07 | 1,67E+07 | 1,65E+07 | 1,85E+07 | 49,994 | 0 | 44,516 | 21 |
| 2,55E+08 | 1,10E+07 | 2,58E+07 | 8,05E+06 | 63,45 | 0 | 74,541 | 42 |
| 2,05E+07 | 1,01E+07 | 9,07E+06 | 9,99E+06 | 28,507 | 0 | 5,9758 | 20 |
| 2,02E+08 | 1,23E+08 | 1,30E+08 | 1,46E+08 | 59,743 | 0 | 52,836 | 100 |
| 9,56E+06 | 2,17E+06 | 2,28E+06 | 2,22E+06 | 39,294 | 0,000435 | 2,9563 | 4 |
| 1,84E+09 | 2,31E+09 | 2,69E+09 | 2,52E+09 | 36,108 | 0 | 323,31 | 411 |
| 7,31E+08 | 5,87E+08 | 5,78E+08 | 5,61E+08 | 73,697 | 0 | 36,608 | 149 |
| 1,67E+08 | 2,82E+08 | 3,67E+08 | 3,92E+08 | 140,99 | 0 | 323,31 | 84 |
| 1,50E+09 | 1,76E+09 | 1,82E+09 | 1,74E+09 | 97,512 | 0 | 323,31 | 410 |

| | | | | | | | |
|---|---|---|---|---|---|---|---|
| 5,70E+07 | 3,55E+07 | 3,60E+07 | 4,22E+07 | 118,39 | 0 | 103 | 68 |
| 2,37E+08 | 1,54E+08 | 1,61E+08 | 1,68E+08 | 61,976 | 0 | 323,31 | 113 |
| 3,21E+07 | 5,93E+07 | 5,01E+07 | 6,38E+07 | 11,692 | 0,003568 | 1,8692 | 14 |
| 1,11E+07 | 3,25E+07 | 2,99E+07 | 2,42E+07 | 54,968 | 0 | 17,714 | 33 |
| 2,93E+08 | 4,96E+08 | 5,97E+08 | 6,01E+08 | 12,811 | 0 | 306,28 | 74 |
| 6,40E+07 | 4,59E+07 | 4,83E+07 | 4,49E+07 | 43,126 | 0 | 24,06 | 51 |
| 3,62E+08 | 3,02E+08 | 3,00E+08 | 3,04E+08 | 167,22 | 0 | 323,31 | 239 |
| 1,19E+07 | 3,02E+06 | 2,40E+06 | 4,68E+06 | 192,63 | 0 | 9,2926 | 10 |
| 3,34E+08 | 2,29E+08 | 2,28E+08 | 2,39E+08 | 53,932 | 0 | 323,31 | 144 |
| 8,05E+07 | 2,36E+07 | 2,10E+07 | 1,13E+07 | 23,578 | 0 | 30,836 | 23 |
| 2,13E+07 | 7,92E+07 | 6,12E+07 | 5,98E+07 | 99,536 | 0 | 23,536 | 29 |
| 5,74E+08 | 3,06E+08 | 2,81E+08 | 2,83E+08 | 108,29 | 0 | 323,31 | 204 |
| 4,67E+07 | 3,56E+07 | 2,89E+07 | 3,05E+07 | 40,977 | 0 | 85,227 | 36 |
| 4,97E+07 | 3,10E+07 | 2,16E+07 | 2,41E+07 | 57,943 | 0 | 23,505 | 37 |
| 1,78E+08 | 1,49E+08 | 1,44E+08 | 1,43E+08 | 51,029 | 0 | 72,213 | 108 |
| 4,43E+07 | 8,85E+07 | 7,93E+07 | 6,72E+07 | 43,776 | 0 | 33,928 | 60 |
| 2,42E+08 | 2,04E+08 | 1,91E+08 | 2,02E+08 | 145,27 | 0 | 99,033 | 148 |
| 7,07E+07 | 4,75E+07 | 4,01E+07 | 4,82E+07 | 48,994 | 0 | 17,639 | 37 |
| 4,83E+08 | 3,08E+08 | 3,04E+08 | 3,06E+08 | 129,6 | 0 | 323,31 | 236 |
| 6,52E+07 | 1,05E+08 | 9,89E+07 | 9,79E+07 | 39,182 | 0 | 18,732 | 84 |
| 1,93E+08 | 2,58E+08 | 2,99E+08 | 2,79E+08 | 47,624 | 0 | 81,604 | 108 |
| 1,94E+08 | 1,09E+08 | 1,23E+08 | 1,29E+08 | 32,932 | 0 | 47,114 | 47 |
| 2,93E+09 | 6,41E+09 | 6,11E+09 | 5,52E+09 | 11,675 | 0 | 98,249 | 365 |
| 1,31E+08 | 2,72E+08 | 2,84E+08 | 2,93E+08 | 92,194 | 0 | 158,68 | 87 |
| 5,70E+07 | 5,20E+06 | 4,33E+06 | 5,05E+06 | 13,559 | 0 | 75,128 | 8 |
| 2,17E+08 | 1,42E+08 | 1,34E+08 | 1,51E+08 | 23,553 | 0 | 55,669 | 15 |
| 6,29E+06 | 6,01E+07 | 3,24E+07 | 6,95E+07 | 39,234 | 0 | 14,841 | 20 |
| 1,94E+08 | 1,42E+08 | 1,29E+08 | 1,26E+08 | 26,624 | 0 | 88,491 | 66 |
| 3,36E+08 | 5,09E+08 | 5,05E+08 | 4,69E+08 | 52,6 | 0 | 316 | 143 |
| 1,91E+08 | 7,44E+07 | 6,35E+07 | 8,12E+07 | 14,467 | 0 | 14,949 | 25 |
| 2,35E+08 | 1,96E+08 | 2,03E+08 | 2,10E+08 | 83,54 | 0 | 152,24 | 171 |
| 2,40E+08 | 1,63E+08 | 1,61E+08 | 1,40E+08 | 70,095 | 0 | 195,99 | 98 |
| 5,97E+07 | 3,95E+07 | 3,04E+07 | 2,96E+07 | 67,556 | 0 | 24,366 | 63 |
| 1,36E+09 | 1,15E+09 | 1,12E+09 | 1,13E+09 | 65,574 | 0 | 323,31 | 247 |
| 4,40E+07 | 2,54E+07 | 2,75E+07 | 2,69E+07 | 86,174 | 0 | 12,601 | 37 |
| 5,83E+07 | 8,68E+06 | 1,36E+07 | 1,88E+07 | 44,881 | 0 | 116,42 | 13 |
| 9,53E+07 | 8,20E+07 | 7,41E+07 | 7,57E+07 | 26,175 | 0 | 28,608 | 53 |
| 6,64E+08 | 5,97E+08 | 5,88E+08 | 5,95E+08 | 178,46 | 0 | 323,31 | 298 |
| 9,98E+08 | 8,87E+08 | 8,32E+08 | 8,73E+08 | 108,84 | 0 | 323,31 | 350 |
| 2,29E+07 | 6,81E+07 | 6,31E+07 | 6,27E+07 | 126,34 | 0 | 25,29 | 39 |
| 5,89E+07 | 3,92E+07 | 3,73E+07 | 3,12E+07 | 58,478 | 0 | 18,286 | 45 |
| 3,57E+08 | 2,19E+08 | 2,15E+08 | 2,35E+08 | 15,241 | 0 | 50,307 | 82 |
| 1,25E+07 | 3,67E+07 | 4,32E+07 | 4,64E+07 | 65,361 | 0 | 21,849 | 33 |
| 4,74E+08 | 1,15E+09 | 9,26E+08 | 9,89E+08 | 35,732 | 0 | 269,66 | 125 |
| 5,06E+08 | 4,82E+08 | 4,74E+08 | 4,76E+08 | 112,03 | 0 | 323,31 | 263 |
| 1,57E+07 | 4,76E+07 | 3,70E+07 | 4,88E+07 | 43,9 | 0 | 12,036 | 24 |
| 8,94E+07 | 1,07E+08 | 1,13E+08 | 1,06E+08 | 137,32 | 0 | 117,34 | 73 |
| 4,30E+08 | 3,67E+08 | 3,87E+08 | 3,85E+08 | 213,67 | 0 | 323,31 | 231 |
| 4,84E+07 | 1,25E+06 | 3,46E+06 | 5,87E+06 | 86,738 | 0 | 15,317 | 24 |
| 1,60E+07 | 2,05E+07 | 2,11E+07 | 2,17E+07 | 16,182 | 0 | 18,344 | 16 |

| | | | | | | | |
|---|---|---|---|---|---|---|---|
| 1,24E+08 | 1,80E+08 | 1,73E+08 | 1,53E+08 | 83,281 | 0 | 130,16 | 89 |
| 9,65E+06 | 5,59E+06 | 5,12E+06 | 3,98E+06 | 66,899 | 0 | 23,679 | 12 |
| 1,61E+07 | 4,92E+06 | 2,53E+06 | 4,45E+06 | 42,428 | 0 | 6,2985 | 11 |
| 2,03E+08 | 1,23E+07 | 9,93E+06 | 2,07E+06 | 42,829 | 0 | 53,031 | 47 |
| 1,99E+07 | 4,41E+07 | 3,64E+07 | 4,13E+07 | 37,742 | 0 | 42,174 | 25 |
| 2,97E+07 | 4,70E+07 | 4,07E+07 | 4,45E+07 | 49,387 | 0 | 10,389 | 26 |
| 3,33E+09 | 2,74E+09 | 2,81E+09 | 2,84E+09 | 145,81 | 0 | 323,31 | 728 |
| 8,54E+07 | 4,57E+07 | 4,55E+07 | 5,44E+07 | 176,66 | 0 | 55,472 | 94 |
| 1,17E+08 | 1,63E+08 | 1,70E+08 | 1,63E+08 | 21,984 | 0 | 100,11 | 76 |
| 7,96E+08 | 1,09E+09 | 1,13E+09 | 1,03E+09 | 55,988 | 0 | 323,31 | 151 |
| 3,28E+09 | 2,44E+09 | 2,49E+09 | 2,49E+09 | 172,79 | 0 | 323,31 | 739 |
| 9,81E+07 | 1,76E+08 | 1,58E+08 | 1,73E+08 | 32,931 | 0 | 118,16 | 84 |
| 1,07E+08 | 8,08E+07 | 8,41E+07 | 8,07E+07 | 67,755 | 0 | 154,49 | 93 |
| 6,90E+06 | 4,09E+07 | 5,05E+07 | 5,05E+07 | 53,341 | 0 | 147,16 | 12 |
| 3,06E+07 | 1,86E+07 | 2,20E+07 | 1,74E+07 | 90,093 | 0 | 8,6147 | 27 |
| 2,75E+08 | 5,64E+08 | 4,95E+08 | 4,19E+08 | 43,008 | 0 | 253,39 | 124 |
| 1,10E+08 | 1,84E+08 | 1,85E+08 | 1,94E+08 | 41,984 | 0 | 49,87 | 73 |
| 7,27E+07 | 4,25E+07 | 4,61E+07 | 5,19E+07 | 16,502 | 0,003445 | 1,9242 | 16 |
| 5,54E+07 | 1,30E+08 | 1,10E+08 | 1,08E+08 | 26,635 | 0 | 178,78 | 74 |
| 8,87E+07 | 5,39E+07 | 5,39E+07 | 5,53E+07 | 76,869 | 0 | 76,74 | 59 |
| 1,47E+08 | 2,03E+06 | 1,23E+07 | 7,10E+06 | 116,08 | 0 | 45,862 | 50 |
| 7,03E+06 | 3,01E+06 | 3,12E+06 | 2,72E+06 | 33,638 | 0,000436 | 3,0161 | 9 |
| 2,07E+07 | 1,36E+07 | 1,50E+07 | 1,31E+07 | 85,872 | 0 | 28,919 | 36 |
| 5,37E+07 | 4,85E+07 | 4,78E+07 | 4,62E+07 | 34,08 | 0 | 30,898 | 18 |
| 2,84E+07 | 3,17E+07 | 3,03E+07 | 3,13E+07 | 52,573 | 0 | 56,493 | 26 |
| 2,50E+08 | 2,06E+08 | 2,03E+08 | 2,12E+08 | 47,983 | 0 | 211,83 | 124 |
| 1,34E+09 | 1,23E+09 | 1,21E+09 | 1,16E+09 | 122,89 | 0 | 323,31 | 281 |
| 7,80E+06 | 3,56E+07 | 7,52E+07 | 3,62E+07 | 24,636 | 0 | 14,311 | 19 |
| 1,78E+07 | 1,00E+06 | 3,16E+06 | 1,78E+06 | 43,975 | 0,000149 | 3,4292 | 7 |
| 3,05E+08 | 2,38E+08 | 2,30E+08 | 2,50E+08 | 114,14 | 0 | 219,99 | 132 |
| 3,26E+08 | 2,49E+08 | 2,13E+08 | 2,42E+08 | 257,56 | 0 | 225,5 | 198 |
| 3,51E+08 | 2,20E+08 | 2,08E+08 | 2,20E+08 | 33,83 | 0 | 85,178 | 100 |
| 6,21E+07 | 9,56E+07 | 1,07E+08 | 1,09E+08 | 48,906 | 0 | 16,099 | 51 |
| 2,56E+08 | 1,78E+08 | 2,00E+08 | 1,64E+08 | 26,131 | 0 | 210,18 | 145 |
| 1,32E+07 | 2,39E+06 | 4,19E+06 | 4,87E+06 | 83,511 | 0 | 18,982 | 10 |
| 4,71E+08 | 1,32E+09 | 1,30E+09 | 1,13E+09 | 34,155 | 0 | 278,75 | 135 |
| 1,17E+08 | 1,46E+08 | 1,58E+08 | 1,43E+08 | 31,231 | 0 | 16,397 | 47 |
| 6,29E+08 | 1,17E+09 | 1,11E+09 | 1,24E+09 | 19,048 | 0 | 157,15 | 96 |
| 1,94E+07 | 8,99E+06 | 6,10E+06 | 9,08E+06 | 79,029 | 0 | 13,956 | 21 |
| 1,74E+07 | 4,53E+07 | 4,20E+07 | 3,50E+07 | 19,587 | 0 | 8,4484 | 20 |
| 5,00E+07 | 2,54E+06 | 4,87E+06 | 2,00E+06 | 14,237 | 0 | 55,166 | 9 |
| 1,39E+07 | 4,69E+06 | 2,33E+06 | 4,01E+06 | 42,935 | 0,000153 | 4,2608 | 9 |
| 2,00E+08 | 1,13E+08 | 1,23E+08 | 1,14E+08 | 41,207 | 0 | 174,06 | 66 |
| 6,79E+08 | 5,06E+08 | 4,96E+08 | 5,06E+08 | 34,702 | 0 | 210,81 | 141 |
| 8,23E+07 | 5,54E+07 | 5,94E+07 | 5,32E+07 | 53,961 | 0 | 35,644 | 50 |
| 4,70E+06 | 8,54E+07 | 7,58E+07 | 8,42E+07 | 96,449 | 0 | 13,541 | 35 |
| 1,90E+09 | 2,90E+09 | 2,69E+09 | 2,56E+09 | 106,91 | 0 | 323,31 | 458 |
| 7,91E+07 | 5,96E+07 | 5,10E+07 | 5,53E+07 | 111,23 | 0 | 20,297 | 43 |
| 1,82E+08 | 2,20E+08 | 2,37E+08 | 2,54E+08 | 15,081 | 0 | 15,291 | 41 |
| 3,55E+08 | 2,34E+08 | 2,17E+08 | 1,98E+08 | 102,91 | 0 | 187,74 | 138 |

| | | | | | | | |
|---|---|---|---|---|---|---|---|
| 1,46E+08 | 1,00E+08 | 9,17E+07 | 8,06E+07 | 18,891 | 0 | 58,676 | 37 |
| 4,17E+07 | 2,60E+07 | 2,98E+07 | 2,28E+07 | 186,44 | 0 | 36,626 | 43 |
| 3,42E+07 | 2,25E+07 | 2,11E+07 | 2,59E+07 | 27,784 | 0 | 9,0445 | 23 |
| 6,69E+07 | 8,79E+07 | 8,33E+07 | 8,18E+07 | 23,277 | 0 | 67,28 | 36 |
| 6,71E+07 | 1,16E+08 | 1,22E+08 | 1,26E+08 | 50,37 | 0 | 82,305 | 62 |
| 1,34E+08 | 9,35E+06 | 2,07E+06 | 1,19E+06 | 90,105 | 0 | 85,269 | 52 |
| 8,53E+07 | 1,77E+08 | 1,58E+08 | 2,07E+08 | 20,504 | 0 | 60,724 | 38 |
| 2,95E+06 | 4,22E+06 | 4,56E+06 | 3,74E+06 | 69,115 | 0,000151 | 3,8848 | 6 |
| 1,18E+08 | 2,33E+07 | 2,30E+07 | 2,63E+07 | 34,904 | 0 | 86,005 | 19 |
| 2,32E+07 | 3,41E+06 | 2,75E+06 | 7,30E+06 | 122,71 | 0 | 43,605 | 18 |
| 3,70E+07 | 2,73E+07 | 2,56E+07 | 2,47E+07 | 33,221 | 0 | 9,7368 | 25 |
| 2,66E+08 | 2,05E+08 | 2,06E+08 | 2,12E+08 | 118,55 | 0 | 189,06 | 131 |
| 3,44E+07 | 1,62E+06 | 4,57E+06 | 2,42E+06 | 102,53 | 0 | 12,009 | 16 |
| 3,16E+08 | 4,25E+08 | 4,49E+08 | 4,70E+08 | 106,69 | 0 | 269,91 | 169 |
| 3,90E+09 | 5,48E+09 | 4,85E+09 | 5,28E+09 | 32,351 | 0 | 323,31 | 285 |
| 1,67E+07 | 2,51E+06 | 3,92E+06 | 5,38E+06 | 23,284 | 0,000147 | 3,2278 | 4 |
| 5,45E+06 | 3,27E+06 | 2,70E+06 | 3,25E+06 | 22,133 | 0,001409 | 2,357 | 3 |
| 6,58E+07 | 8,88E+07 | 1,02E+08 | 9,91E+07 | 165,8 | 0 | 142,94 | 66 |
| 5,37E+06 | 1,64E+07 | 1,48E+07 | 2,16E+07 | 12,691 | 0 | 12,718 | 10 |
| 2,29E+07 | 5,45E+06 | 2,71E+06 | 1,71E+06 | 38,201 | 0 | 29,509 | 8 |
| 6,76E+08 | 3,91E+08 | 3,67E+08 | 4,22E+08 | 63,798 | 0 | 323,31 | 109 |
| 2,53E+07 | 4,10E+07 | 4,87E+07 | 4,84E+07 | 57,034 | 0 | 48,667 | 25 |
| 6,47E+08 | 8,93E+08 | 8,87E+08 | 8,82E+08 | 57,788 | 0 | 323,31 | 158 |
| 4,54E+07 | 2,27E+07 | 2,36E+07 | 2,43E+07 | 164,33 | 0 | 24,175 | 45 |
| 1,37E+10 | 1,02E+10 | 1,08E+10 | 1,09E+10 | 34,196 | 0 | 323,31 | 711 |
| 1,72E+07 | 3,80E+06 | 2,30E+06 | 1,35E+06 | 74,331 | 0 | 7,8884 | 15 |
| 1,20E+07 | 2,64E+06 | 4,52E+06 | 2,65E+06 | 186,53 | 0 | 12,756 | 7 |
| 9,66E+06 | 3,33E+06 | 2,07E+06 | 2,08E+06 | 36,728 | 0,000148 | 3,3075 | 5 |
| 9,39E+06 | 2,23E+07 | 1,90E+07 | 2,77E+07 | 14,916 | 0 | 11,144 | 20 |
| 4,38E+08 | 6,84E+08 | 6,50E+08 | 7,07E+08 | 32,967 | 0 | 323,31 | 108 |
| 3,89E+06 | 2,18E+07 | 1,72E+07 | 1,94E+07 | 27,392 | 0 | 5,7751 | 12 |
| 9,38E+06 | 3,37E+06 | 2,90E+06 | 4,22E+06 | 23,693 | 0 | 6,7781 | 7 |
| 7,70E+07 | 4,41E+07 | 5,36E+07 | 5,41E+07 | 138,59 | 0 | 36,824 | 63 |
| 3,57E+07 | 5,98E+07 | 5,37E+07 | 6,16E+07 | 25,068 | 0 | 127,31 | 51 |
| 4,98E+07 | 4,21E+07 | 3,93E+07 | 4,02E+07 | 72,407 | 0 | 31,24 | 45 |
| 3,70E+08 | 4,14E+08 | 4,34E+08 | 4,12E+08 | 133,32 | 0 | 268,8 | 255 |
| 4,14E+07 | 4,64E+06 | 8,25E+06 | 2,44E+06 | 172,14 | 0 | 25,501 | 30 |
| 4,89E+07 | 8,75E+07 | 1,08E+08 | 1,14E+08 | 69,948 | 0 | 83,155 | 48 |
| 9,86E+07 | 8,14E+07 | 7,77E+07 | 8,54E+07 | 72,673 | 0 | 34,396 | 88 |
| 3,08E+08 | 5,19E+08 | 5,54E+08 | 5,37E+08 | 22,338 | 0 | 89,334 | 97 |
| 2,34E+08 | 1,51E+08 | 1,53E+08 | 1,44E+08 | 138,04 | 0 | 95,838 | 119 |
| 3,91E+08 | 3,32E+08 | 3,26E+08 | 3,08E+08 | 70,429 | 0 | 105,31 | 187 |
| 8,97E+07 | 4,47E+08 | 3,11E+08 | 3,25E+08 | 18,841 | 0,000146 | 3,0953 | 25 |
| 1,70E+08 | 2,38E+08 | 2,11E+08 | 2,45E+08 | 211,46 | 0 | 71,074 | 154 |
| 4,19E+07 | 4,73E+06 | 1,17E+06 | 3,75E+06 | 79,102 | 0 | 7,5739 | 11 |
| 1,45E+07 | 4,27E+07 | 3,80E+07 | 3,91E+07 | 15,382 | 0 | 6,9433 | 20 |
| 3,10E+08 | 2,42E+08 | 2,44E+08 | 2,51E+08 | 81,699 | 0 | 88,104 | 86 |
| 2,70E+08 | 2,40E+08 | 2,43E+08 | 2,51E+08 | 44,176 | 0 | 323,31 | 107 |
| 1,83E+08 | 9,85E+07 | 1,04E+08 | 1,18E+08 | 149,67 | 0 | 59,787 | 114 |
| 6,51E+07 | 1,34E+07 | 7,21E+06 | 4,51E+06 | 129,65 | 0 | 33,649 | 28 |

| | | | | | | | |
|---|---|---|---|---|---|---|---|
| 1,77E+10 | 2,40E+10 | 2,40E+10 | 2,36E+10 | 281,22 | 0 | 323,31 | 1573 |
| 9,03E+08 | 1,75E+07 | 2,41E+07 | 8,77E+07 | 129,77 | 0 | 18,617 | 35 |
| 1,24E+09 | 7,65E+08 | 6,64E+08 | 6,40E+08 | 63,676 | 0 | 197,28 | 174 |
| 5,16E+07 | 1,98E+07 | 1,79E+07 | 1,98E+07 | 43,975 | 0 | 11,966 | 13 |
| 1,12E+07 | 1,77E+07 | 1,63E+07 | 1,45E+07 | 104,81 | 0,000149 | 3,4733 | 18 |
| 4,54E+07 | 2,59E+06 | 2,96E+06 | 8,99E+06 | 26,87 | 0 | 7,72 | 11 |
| 1,86E+08 | 1,49E+08 | 1,51E+08 | 1,63E+08 | 235,39 | 0 | 72,146 | 148 |
| 1,09E+08 | 1,41E+08 | 1,46E+08 | 1,37E+08 | 93,629 | 0 | 126,56 | 73 |
| 1,08E+08 | 1,52E+08 | 1,36E+08 | 1,40E+08 | 224,32 | 0 | 177,07 | 135 |
| 2,85E+06 | 9,03E+06 | 1,05E+07 | 7,55E+06 | 46,972 | 0 | 7,7603 | 10 |
| 2,37E+07 | 2,07E+06 | 4,88E+06 | 3,02E+06 | 20,748 | 0,003704 | 1,8614 | 4 |
| 5,11E+07 | 4,00E+07 | 4,25E+07 | 4,46E+07 | 53,831 | 0 | 20,115 | 37 |
| 8,15E+07 | 1,15E+08 | 1,08E+08 | 1,26E+08 | 25,961 | 0 | 96,457 | 48 |
| 2,17E+08 | 1,78E+08 | 1,68E+08 | 1,65E+08 | 36,79 | 0 | 56,321 | 71 |
| 5,65E+09 | 7,47E+09 | 7,79E+09 | 7,07E+09 | 84,787 | 0 | 323,31 | 986 |
| 3,82E+06 | 3,25E+08 | 2,92E+08 | 7,58E+08 | 84,152 | 0 | 295,47 | 54 |
| 2,73E+09 | 3,78E+09 | 3,46E+09 | 3,32E+09 | 103,07 | 0 | 323,31 | 556 |
| 3,04E+07 | 6,27E+06 | 5,65E+06 | 3,75E+06 | 42,248 | 0 | 16,407 | 14 |
| 1,13E+08 | 8,21E+07 | 7,82E+07 | 7,17E+07 | 43,295 | 0 | 28,377 | 56 |
| 7,24E+07 | 6,18E+07 | 6,03E+07 | 6,54E+07 | 34,773 | 0 | 96,606 | 56 |
| 5,91E+08 | 7,05E+08 | 6,84E+08 | 7,12E+08 | 114,99 | 0 | 323,31 | 223 |
| 3,70E+07 | 1,16E+07 | 9,25E+06 | 5,02E+06 | 35,701 | 0 | 14,012 | 23 |
| 1,86E+07 | 3,29E+08 | 2,77E+08 | 2,98E+08 | 56,63 | 0 | 95,937 | 63 |
| 2,10E+09 | 3,41E+09 | 3,90E+09 | 3,99E+09 | 13,777 | 0 | 301,25 | 236 |
| 3,30E+08 | 3,74E+08 | 3,82E+08 | 3,98E+08 | 79,763 | 0 | 222,12 | 152 |
| 2,88E+08 | 1,57E+08 | 1,69E+08 | 1,49E+08 | 65,46 | 0 | 90,444 | 126 |
| 1,93E+07 | 3,09E+07 | 2,58E+07 | 2,61E+07 | 135,25 | 0 | 9,6957 | 37 |
| 2,04E+08 | 1,25E+08 | 1,03E+08 | 1,08E+08 | 209,62 | 0 | 40,887 | 65 |
| 4,61E+06 | 1,49E+07 | 2,01E+07 | 2,02E+07 | 102,04 | 0 | 17,221 | 14 |
| 2,95E+08 | 1,79E+08 | 1,87E+08 | 1,58E+08 | 40,342 | 0 | 167,08 | 104 |
| 2,16E+09 | 1,82E+09 | 1,74E+09 | 1,61E+09 | 46,415 | 0 | 323,31 | 314 |
| 1,89E+07 | 2,57E+07 | 2,63E+07 | 3,06E+07 | 127,61 | 0 | 27,622 | 23 |
| 4,80E+07 | 3,36E+07 | 3,34E+07 | 3,05E+07 | 32,281 | 0 | 27,083 | 32 |
| 1,69E+07 | 2,84E+07 | 2,93E+07 | 4,03E+07 | 20,936 | 0 | 8,7945 | 16 |
| 2,56E+07 | 1,95E+07 | 1,78E+07 | 1,56E+07 | 41,546 | 0 | 17,735 | 18 |
| 2,97E+07 | 6,02E+07 | 5,91E+07 | 5,21E+07 | 59,083 | 0 | 41,725 | 44 |
| 3,71E+08 | 5,72E+08 | 7,99E+08 | 7,48E+08 | 7,3384 | 0 | 323,31 | 50 |
| 4,16E+06 | 2,65E+07 | 1,52E+07 | 1,94E+07 | 24,784 | 0 | 11,036 | 7 |
| 2,38E+07 | 6,66E+07 | 7,88E+07 | 6,72E+07 | 63,375 | 0 | 39,585 | 23 |
| 1,40E+07 | 1,72E+08 | 2,09E+08 | 1,83E+08 | 22,727 | 0 | 219,65 | 29 |
| 2,31E+07 | 7,13E+06 | 1,05E+07 | 7,41E+06 | 47,967 | 0 | 12,201 | 18 |
| 6,63E+06 | 1,12E+07 | 1,13E+07 | 1,07E+07 | 55,144 | 0,000291 | 3,0362 | 9 |
| 2,90E+08 | 3,40E+08 | 3,62E+08 | 3,43E+08 | 34,958 | 0 | 323,31 | 111 |
| 3,71E+08 | 4,78E+08 | 5,54E+08 | 5,37E+08 | 27,956 | 0 | 323,31 | 197 |
| 1,12E+08 | 7,45E+07 | 7,77E+07 | 8,56E+07 | 54,27 | 0 | 36,389 | 52 |
| 7,57E+07 | 6,57E+07 | 6,23E+07 | 6,40E+07 | 23,477 | 0 | 8,3102 | 27 |
| 2,16E+07 | 1,44E+07 | 1,33E+07 | 1,62E+07 | 52,904 | 0 | 10,702 | 29 |
| 5,38E+06 | 1,19E+07 | 1,43E+07 | 1,45E+07 | 35,823 | 0,000157 | 4,9583 | 8 |
| 7,00E+08 | 4,11E+08 | 3,79E+08 | 3,95E+08 | 20,583 | 0 | 323,31 | 83 |
| 2,22E+08 | 1,10E+08 | 8,15E+07 | 8,70E+07 | 83,14 | 0 | 54,246 | 80 |

| | | | | | | | |
|---|---|---|---|---|---|---|---|
| 1,22E+07 | 3,58E+06 | 4,19E+06 | 5,96E+06 | 15,032 | 0,000156 | 4,6349 | 5 |
| 2,02E+08 | 1,60E+08 | 1,68E+08 | 1,68E+08 | 72,648 | 0 | 85,458 | 108 |
| 7,42E+07 | 1,47E+08 | 1,43E+08 | 1,70E+08 | 11,101 | 0 | 89,619 | 28 |
| 3,34E+09 | 5,17E+09 | 5,39E+09 | 5,56E+09 | 88,846 | 0 | 323,31 | 536 |
| 8,06E+06 | 1,27E+07 | 1,18E+07 | 1,06E+07 | 92,714 | 0 | 12,995 | 23 |
| 3,34E+08 | 3,90E+08 | 3,99E+08 | 3,72E+08 | 14,454 | 0 | 71,71 | 69 |
| 7,26E+07 | 3,13E+07 | 3,99E+07 | 3,69E+07 | 92,632 | 0 | 84,684 | 35 |
| 5,87E+09 | 3,92E+09 | 3,68E+09 | 4,00E+09 | 67,766 | 0 | 323,31 | 680 |
| 2,54E+08 | 3,31E+08 | 3,36E+08 | 3,23E+08 | 30,048 | 0 | 186,46 | 100 |
| 9,00E+05 | 3,46E+07 | 2,28E+07 | 2,75E+07 | 75,795 | 0 | 14,79 | 24 |
| 1,70E+07 | 1,41E+07 | 1,29E+07 | 1,24E+07 | 22,436 | 0,003182 | 1,9721 | 7 |
| 7,16E+08 | 6,53E+08 | 6,78E+08 | 6,71E+08 | 265,25 | 0 | 323,31 | 322 |
| 3,97E+07 | 5,73E+06 | 1,29E+07 | 1,00E+07 | 86,625 | 0 | 60,189 | 27 |
| 2,82E+07 | 2,13E+06 | 4,82E+06 | 1,73E+06 | 83,043 | 0 | 10,463 | 18 |
| 4,14E+06 | 5,87E+07 | 3,03E+07 | 2,87E+07 | 51,595 | 0 | 17,114 | 29 |
| 2,52E+08 | 3,15E+08 | 3,06E+08 | 2,94E+08 | 232,75 | 0 | 271,27 | 240 |
| 5,95E+08 | 7,00E+08 | 6,96E+08 | 7,44E+08 | 76,526 | 0 | 323,31 | 190 |
| 1,71E+08 | 1,10E+07 | 6,67E+06 | 1,45E+06 | 37,17 | 0 | 71,221 | 24 |
| 1,36E+07 | 5,99E+07 | 6,68E+07 | 6,55E+07 | 57,915 | 0 | 35,266 | 40 |
| 3,05E+07 | 1,92E+07 | 2,33E+07 | 2,26E+07 | 62,255 | 0 | 7,0203 | 19 |
| 6,22E+07 | 3,07E+06 | 7,28E+06 | 8,42E+06 | 100,83 | 0 | 20,691 | 30 |
| 1,01E+09 | 7,91E+08 | 8,26E+08 | 8,35E+08 | 115,42 | 0 | 323,31 | 325 |
| 1,47E+08 | 1,19E+08 | 1,12E+08 | 1,22E+08 | 71,11 | 0 | 96,036 | 92 |
| 5,01E+07 | 1,95E+07 | 1,82E+07 | 1,80E+07 | 49,566 | 0 | 73,839 | 24 |
| 3,84E+08 | 3,23E+08 | 3,07E+08 | 3,02E+08 | 38,822 | 0 | 70,577 | 90 |
| 3,83E+08 | 4,67E+08 | 5,33E+08 | 5,03E+08 | 25,604 | 0 | 8,5827 | 52 |
| 4,26E+09 | 2,32E+09 | 2,38E+09 | 2,39E+09 | 19,462 | 0 | 265,5 | 439 |
| 5,63E+06 | 1,07E+07 | 9,02E+06 | 8,73E+06 | 16,256 | 0,000154 | 4,2992 | 11 |
| 2,17E+09 | 1,90E+09 | 1,80E+09 | 1,88E+09 | 47,357 | 0 | 323,31 | 357 |
| 1,30E+07 | 6,98E+06 | 5,25E+06 | 5,96E+06 | 43,016 | 0 | 7,0364 | 8 |
| 2,44E+07 | 1,48E+07 | 1,11E+07 | 1,44E+07 | 121,42 | 0 | 22,85 | 35 |
| 2,56E+07 | 5,10E+06 | 4,43E+06 | 6,13E+06 | 43,371 | 0 | 5,345 | 17 |
| 1,83E+07 | 6,15E+07 | 5,48E+07 | 4,96E+07 | 53,872 | 0 | 114,63 | 30 |
| 4,89E+07 | 1,11E+08 | 1,27E+08 | 1,17E+08 | 33,365 | 0 | 30,029 | 53 |
| 8,10E+06 | 1,67E+07 | 1,42E+07 | 1,44E+07 | 19,017 | 0 | 9,6316 | 14 |
| 6,15E+07 | 8,75E+05 | 5,54E+06 | 2,27E+06 | 8,5497 | 0 | 5,9312 | 17 |
| 3,00E+07 | 6,55E+06 | 8,04E+06 | 3,17E+06 | 231,37 | 0 | 20,608 | 27 |
| 1,27E+09 | 7,92E+08 | 8,64E+08 | 8,67E+08 | 10,344 | 0 | 9,9715 | 94 |
| 1,54E+06 | 1,97E+07 | 1,57E+07 | 2,98E+07 | 32,887 | 0,0021 | 2,1901 | 8 |
| 1,95E+07 | 1,13E+08 | 1,15E+08 | 9,91E+07 | 162,33 | 0 | 74,414 | 91 |
| 2,13E+08 | 2,93E+08 | 2,90E+08 | 3,35E+08 | 34,834 | 0 | 323,31 | 65 |
| 3,19E+07 | 4,91E+07 | 5,25E+07 | 4,77E+07 | 73,631 | 0 | 19,479 | 39 |
| 1,69E+08 | 9,06E+07 | 1,08E+08 | 1,02E+08 | 54,097 | 0 | 6,3956 | 18 |
| 1,07E+08 | 6,57E+07 | 6,90E+07 | 7,19E+07 | 28,6 | 0 | 5,9635 | 22 |
| 3,97E+07 | 2,35E+07 | 1,73E+07 | 1,43E+07 | 141,48 | 0 | 32,611 | 37 |
| 1,25E+06 | 1,18E+08 | 7,87E+07 | 9,08E+07 | 169,04 | 0 | 22,756 | 53 |
| 2,18E+07 | 1,24E+07 | 1,18E+07 | 1,57E+07 | 76,992 | 0 | 11,708 | 33 |
| 1,76E+07 | 3,66E+06 | 2,57E+06 | 6,44E+06 | 34,623 | 0 | 9,3473 | 13 |
| 2,34E+08 | 1,64E+08 | 1,72E+08 | 1,67E+08 | 103,32 | 0 | 66,49 | 121 |
| 6,84E+07 | 4,52E+07 | 4,35E+07 | 4,54E+07 | 44,609 | 0 | 49,573 | 38 |

| | | | | | | | |
|---|---|---|---|---|---|---|---|
| 4,95E+07 | 3,47E+07 | 3,86E+07 | 3,39E+07 | 48,219 | 0 | 7,5452 | 26 |
| 1,76E+09 | 2,10E+09 | 2,41E+09 | 2,37E+09 | 43,096 | 0 | 323,31 | 271 |
| 5,31E+08 | 6,72E+08 | 6,59E+08 | 6,86E+08 | 48,162 | 0 | 323,31 | 150 |
| 2,18E+08 | 1,83E+08 | 1,79E+08 | 1,84E+08 | 91,841 | 0 | 289,39 | 110 |
| 5,65E+08 | 4,65E+08 | 4,95E+08 | 4,54E+08 | 27,127 | 0 | 263,26 | 100 |
| 8,37E+07 | 5,97E+07 | 6,01E+07 | 5,46E+07 | 32,402 | 0 | 28,222 | 46 |
| 3,61E+07 | 4,44E+07 | 4,21E+07 | 4,15E+07 | 29,374 | 0 | 17,135 | 41 |
| 1,22E+07 | 8,82E+06 | 7,43E+06 | 8,87E+06 | 34,06 | 0 | 37,021 | 14 |
| 1,57E+09 | 2,12E+09 | 1,90E+09 | 1,90E+09 | 64,282 | 0 | 323,31 | 348 |
| 2,37E+08 | 2,95E+08 | 3,00E+08 | 3,01E+08 | 46,524 | 0 | 191,5 | 77 |
| 2,36E+08 | 1,54E+08 | 1,67E+08 | 1,75E+08 | 79,076 | 0 | 177,44 | 106 |
| 1,32E+07 | 1,54E+06 | 1,56E+06 | 1,85E+06 | 50,939 | 0 | 6,8665 | 5 |
| 2,33E+08 | 3,40E+08 | 2,91E+08 | 3,27E+08 | 39,431 | 0 | 274,47 | 100 |
| 1,66E+07 | 4,18E+06 | 1,62E+06 | 3,05E+06 | 67,703 | 0 | 8,2368 | 13 |
| 2,66E+07 | 1,68E+07 | 1,12E+07 | 1,36E+07 | 108,04 | 0 | 13,147 | 21 |
| 1,09E+09 | 8,62E+08 | 8,34E+08 | 7,67E+08 | 121,5 | 0 | 202,8 | 176 |
| 2,59E+06 | 1,24E+07 | 1,33E+07 | 1,41E+07 | 27,895 | 0,000151 | 3,8164 | 15 |
| 1,30E+07 | 1,34E+08 | 1,64E+08 | 1,37E+08 | 35,469 | 0 | 17,283 | 32 |
| 1,25E+07 | 3,19E+06 | 1,10E+06 | 2,58E+06 | 113,51 | 0,000152 | 3,9948 | 11 |
| 5,47E+07 | 2,20E+07 | 2,55E+07 | 1,74E+07 | 46,325 | 0 | 142,73 | 27 |
| 1,48E+08 | 1,20E+08 | 1,15E+08 | 1,29E+08 | 28,381 | 0 | 67,839 | 70 |
| 1,09E+08 | 9,31E+07 | 9,43E+07 | 9,53E+07 | 49,593 | 0 | 78,641 | 58 |
| 1,53E+08 | 2,70E+08 | 2,77E+08 | 3,02E+08 | 63,538 | 0 | 125,56 | 126 |
| 3,56E+08 | 5,04E+08 | 4,41E+08 | 4,35E+08 | 49,812 | 0 | 323,31 | 137 |
| 9,36E+08 | 7,60E+08 | 8,24E+08 | 8,22E+08 | 82,657 | 0 | 323,31 | 284 |
| 3,43E+07 | 1,30E+07 | 1,90E+07 | 1,66E+07 | 83,365 | 0 | 16,225 | 33 |
| 7,36E+09 | 3,60E+09 | 3,27E+09 | 2,55E+09 | 32,56 | 0 | 323,31 | 662 |
| 3,62E+07 | 5,35E+07 | 4,91E+07 | 5,64E+07 | 102,45 | 0 | 31,043 | 41 |
| 2,49E+08 | 3,45E+08 | 3,29E+08 | 3,54E+08 | 51,1 | 0 | 323,31 | 162 |
| 6,83E+08 | 4,81E+08 | 5,40E+08 | 5,09E+08 | 46,481 | 0 | 102,96 | 131 |
| 1,50E+07 | 2,62E+06 | 8,95E+05 | 2,72E+06 | 29,494 | 0,004631 | 1,7315 | 4 |
| 2,88E+08 | 5,37E+08 | 4,37E+08 | 4,26E+08 | 28,729 | 0 | 323,31 | 160 |
| 4,68E+07 | 3,45E+07 | 3,42E+07 | 3,10E+07 | 108,85 | 0 | 69,552 | 44 |
| 2,24E+07 | 4,64E+06 | 1,08E+06 | 2,28E+06 | 96,541 | 0 | 8,5518 | 17 |
| 1,02E+08 | 1,66E+08 | 1,52E+08 | 1,52E+08 | 51,681 | 0 | 51,051 | 96 |
| 5,06E+08 | 7,14E+08 | 7,25E+08 | 7,15E+08 | 49,654 | 0 | 323,31 | 209 |
| 7,75E+07 | 9,71E+07 | 9,36E+07 | 9,45E+07 | 109,67 | 0 | 100,61 | 57 |
| 3,18E+06 | 6,48E+07 | 5,82E+07 | 6,58E+07 | 183,67 | 0 | 33,615 | 24 |
| 1,59E+08 | 9,22E+07 | 9,76E+07 | 1,03E+08 | 34,371 | 0 | 97,524 | 51 |
| 1,45E+07 | 9,08E+06 | 7,52E+06 | 7,85E+06 | 94,774 | 0,000431 | 2,7974 | 12 |
| 3,50E+08 | 3,95E+08 | 3,92E+08 | 3,98E+08 | 162,4 | 0 | 254,08 | 217 |
| 6,59E+08 | 8,49E+08 | 9,20E+08 | 8,66E+08 | 110,65 | 0 | 323,31 | 226 |
| 1,07E+07 | 3,85E+06 | 1,49E+06 | 1,81E+06 | 89,448 | 0 | 6,1684 | 9 |
| 6,12E+07 | 9,47E+07 | 1,04E+08 | 9,76E+07 | 215,08 | 0 | 30,378 | 81 |
| 9,53E+06 | 2,80E+06 | 1,84E+06 | 3,78E+06 | 147,94 | 0,000148 | 3,4019 | 9 |
| 1,13E+10 | 9,54E+09 | 1,01E+10 | 9,37E+09 | 87,917 | 0 | 323,31 | 789 |
| 8,39E+06 | 1,56E+07 | 1,44E+07 | 1,61E+07 | 62,216 | 0 | 5,9475 | 17 |
| 3,33E+08 | 2,82E+08 | 2,85E+08 | 2,85E+08 | 38,978 | 0 | 117,16 | 123 |
| 7,98E+08 | 1,09E+09 | 1,11E+09 | 1,06E+09 | 150,33 | 0 | 323,31 | 384 |
| 1,43E+08 | 2,94E+08 | 2,66E+08 | 2,64E+08 | 57,621 | 0 | 74,61 | 91 |

| | | | | | | | |
|---|---|---|---|---|---|---|---|
| 4,63E+08 | 6,08E+08 | 7,20E+08 | 6,54E+08 | 89,519 | 0 | 211,8 | 168 |
| 5,85E+08 | 8,85E+08 | 8,50E+08 | 8,30E+08 | 18,721 | 0 | 176,81 | 29 |
| 8,74E+06 | 5,86E+06 | 5,51E+06 | 5,66E+06 | 24,074 | 0,00015 | 3,6384 | 11 |
| 3,00E+07 | 4,76E+07 | 5,24E+07 | 5,11E+07 | 141,78 | 0 | 32,244 | 47 |
| 3,20E+08 | 2,09E+08 | 1,76E+08 | 2,02E+08 | 39,666 | 0 | 179,79 | 137 |
| 3,68E+08 | 4,24E+08 | 4,66E+08 | 4,49E+08 | 35,284 | 0 | 119,17 | 64 |
| 9,47E+08 | 7,94E+08 | 7,26E+08 | 7,42E+08 | 79,481 | 0 | 244,83 | 197 |
| 5,28E+07 | 1,08E+06 | 2,45E+06 | 3,58E+06 | 122,42 | 0 | 16,435 | 18 |
| 5,28E+08 | 3,54E+08 | 3,50E+08 | 3,35E+08 | 108,77 | 0 | 323,31 | 216 |
| 2,02E+08 | 1,20E+08 | 1,24E+08 | 1,28E+08 | 52,534 | 0 | 189,22 | 45 |
| 1,02E+08 | 3,74E+07 | 3,72E+07 | 5,59E+07 | 36,129 | 0 | 92,8 | 65 |
| 1,36E+07 | 7,26E+06 | 7,13E+06 | 4,56E+06 | 93,076 | 0 | 7,2217 | 13 |
| 1,54E+08 | 9,55E+07 | 9,47E+07 | 8,68E+07 | 104,83 | 0 | 85,341 | 85 |
| 5,28E+09 | 4,23E+09 | 4,56E+09 | 4,59E+09 | 69,289 | 0 | 323,31 | 360 |
| 8,55E+07 | 5,42E+06 | 2,03E+06 | 4,72E+06 | 5,6794 | 0 | 16,263 | 9 |
| 3,20E+07 | 2,82E+07 | 2,64E+07 | 2,72E+07 | 38,39 | 0 | 23,001 | 15 |
| 4,49E+08 | 3,51E+08 | 3,41E+08 | 3,28E+08 | 247,17 | 0 | 323,31 | 237 |
| 6,85E+07 | 4,93E+07 | 4,42E+07 | 5,25E+07 | 18,098 | 0 | 21,529 | 26 |
| 2,11E+07 | 1,37E+07 | 1,58E+07 | 1,55E+07 | 119,56 | 0 | 17,932 | 16 |
| 1,26E+06 | 2,38E+07 | 1,68E+07 | 1,64E+07 | 72,954 | 0 | 9,016 | 9 |
| 3,26E+08 | 1,78E+08 | 1,88E+08 | 1,78E+08 | 21,064 | 0 | 77,41 | 28 |
| 5,05E+07 | 9,12E+06 | 2,02E+06 | 4,65E+06 | 215,74 | 0 | 49,971 | 38 |
| 1,09E+08 | 6,53E+07 | 7,40E+07 | 6,02E+07 | 52,106 | 0 | 46,513 | 58 |
| 2,72E+06 | 4,70E+07 | 4,42E+07 | 4,48E+07 | 143,86 | 0 | 23,915 | 29 |
| 7,29E+07 | 1,81E+07 | 8,03E+06 | 7,68E+06 | 46,291 | 0 | 125,63 | 38 |
| 2,42E+08 | 2,68E+08 | 2,65E+08 | 2,75E+08 | 41,518 | 0 | 181,89 | 120 |
| 1,12E+08 | 7,57E+07 | 7,88E+07 | 7,43E+07 | 67,076 | 0 | 119,31 | 49 |
| 8,95E+07 | 1,22E+08 | 1,30E+08 | 1,22E+08 | 34,952 | 0 | 59,99 | 74 |
| 8,62E+08 | 1,29E+09 | 1,24E+09 | 1,26E+09 | 71,781 | 0 | 185,57 | 341 |
| 1,52E+07 | 5,95E+06 | 3,75E+06 | 2,41E+06 | 15,676 | 0,002917 | 2,0364 | 4 |
| 9,56E+06 | 4,38E+06 | 6,20E+06 | 5,19E+06 | 68,889 | 0,000435 | 2,9697 | 12 |
| 2,86E+08 | 4,43E+08 | 4,11E+08 | 4,25E+08 | 52,206 | 0 | 229,66 | 117 |
| 6,38E+07 | 3,41E+07 | 3,51E+07 | 4,61E+07 | 17,996 | 0 | 13,802 | 31 |
| 2,21E+09 | 1,59E+09 | 1,60E+09 | 1,41E+09 | 49,027 | 0 | 323,31 | 250 |
| 6,83E+06 | 1,21E+07 | 1,33E+07 | 1,23E+07 | 77,573 | 0 | 6,2004 | 12 |
| 1,42E+07 | 3,57E+07 | 3,94E+07 | 3,45E+07 | 50,006 | 0 | 13,571 | 22 |
| 4,99E+06 | 1,05E+07 | 1,04E+07 | 1,47E+07 | 91,98 | 0,00015 | 3,6881 | 6 |
| 1,04E+08 | 1,41E+08 | 1,42E+08 | 1,52E+08 | 122,17 | 0 | 73,973 | 114 |
| 5,92E+06 | 1,16E+07 | 1,05E+07 | 1,23E+07 | 21,825 | 0,000148 | 3,3231 | 7 |
| 2,07E+08 | 1,63E+08 | 1,53E+08 | 1,68E+08 | 87,752 | 0 | 57,15 | 111 |
| 1,62E+08 | 2,15E+08 | 2,49E+08 | 2,26E+08 | 65,214 | 0 | 292,47 | 73 |
| 1,76E+07 | 3,42E+07 | 2,93E+07 | 3,82E+07 | 33,701 | 0 | 13,998 | 34 |
| 3,12E+08 | 4,07E+08 | 4,19E+08 | 4,71E+08 | 250,55 | 0 | 93,906 | 138 |
| 2,70E+08 | 6,00E+08 | 5,21E+08 | 5,45E+08 | 38,113 | 0 | 323,31 | 178 |
| 1,34E+07 | 7,61E+06 | 7,22E+06 | 6,17E+06 | 41,967 | 0 | 8,8937 | 12 |
| 7,66E+08 | 3,88E+08 | 3,80E+08 | 4,18E+08 | 20,02 | 0 | 112,08 | 106 |
| 2,08E+08 | 3,31E+08 | 3,32E+08 | 3,21E+08 | 197,03 | 0 | 164,49 | 185 |
| 2,06E+07 | 3,98E+07 | 3,39E+07 | 3,49E+07 | 22,749 | 0 | 11,497 | 12 |
| 9,64E+08 | 1,15E+09 | 1,28E+09 | 1,17E+09 | 272,53 | 0 | 323,31 | 442 |
| 7,59E+06 | 4,33E+06 | 3,00E+06 | 3,53E+06 | 31,133 | 0,003571 | 1,8856 | 8 |

| | | | | | | | |
|---|---|---|---|---|---|---|---|
| 1,14E+08 | 1,04E+08 | 9,74E+07 | 1,02E+08 | 20,368 | 0 | 12,551 | 38 |
| 7,45E+08 | 4,97E+08 | 4,75E+08 | 4,76E+08 | 105,77 | 0 | 323,31 | 306 |
| 3,86E+08 | 3,40E+08 | 3,43E+08 | 3,42E+08 | 196,69 | 0 | 323,31 | 235 |
| 3,76E+07 | 7,38E+07 | 7,60E+07 | 6,57E+07 | 29,415 | 0 | 62,537 | 33 |
| 7,07E+07 | 8,48E+07 | 8,89E+07 | 8,62E+07 | 35,705 | 0 | 62,374 | 48 |
| 1,05E+08 | 1,71E+08 | 1,45E+08 | 1,71E+08 | 51,66 | 0 | 67,217 | 77 |
| 7,10E+08 | 1,31E+09 | 1,35E+09 | 1,19E+09 | 19,526 | 0 | 323,31 | 108 |
| 6,60E+07 | 9,73E+07 | 1,19E+08 | 1,06E+08 | 211,87 | 0 | 87,744 | 68 |
| 4,20E+06 | 3,29E+07 | 1,91E+07 | 2,56E+07 | 65,19 | 0 | 9,3061 | 16 |
| 3,61E+07 | 7,46E+07 | 6,71E+07 | 6,45E+07 | 20,114 | 0 | 104,91 | 20 |
| 2,05E+07 | 2,65E+07 | 2,66E+07 | 2,79E+07 | 208,96 | 0 | 38,436 | 41 |
| 1,57E+09 | 1,97E+09 | 1,87E+09 | 1,92E+09 | 270,25 | 0 | 323,31 | 541 |
| 5,10E+07 | 2,87E+07 | 2,97E+07 | 2,95E+07 | 81,488 | 0 | 42,126 | 28 |
| 2,57E+08 | 2,29E+08 | 2,17E+08 | 2,19E+08 | 106,2 | 0 | 195,57 | 197 |
| 4,78E+06 | 8,68E+06 | 9,85E+06 | 1,15E+07 | 161,14 | 0 | 5,9695 | 13 |
| 2,72E+08 | 4,22E+08 | 4,41E+08 | 4,61E+08 | 72,3 | 0 | 252,91 | 129 |
| 6,85E+07 | 1,00E+08 | 9,62E+07 | 1,10E+08 | 237,04 | 0 | 38,595 | 94 |
| 1,93E+07 | 1,39E+07 | 1,55E+07 | 1,61E+07 | 72,037 | 0,002099 | 2,1758 | 7 |
| 7,74E+07 | 1,21E+08 | 1,13E+08 | 1,19E+08 | 66,839 | 0 | 319,34 | 55 |
| 9,84E+07 | 1,15E+08 | 1,17E+08 | 1,28E+08 | 109,08 | 0 | 39,766 | 94 |
| 4,36E+08 | 5,06E+08 | 5,28E+08 | 5,19E+08 | 55,832 | 0 | 174,09 | 146 |
| 8,02E+07 | 1,24E+08 | 1,22E+08 | 1,34E+08 | 57,985 | 0 | 100,07 | 73 |
| 3,96E+06 | 2,03E+07 | 1,98E+07 | 3,06E+07 | 22,628 | 0 | 13,582 | 15 |
| 2,43E+07 | 4,42E+07 | 3,82E+07 | 4,34E+07 | 54,906 | 0 | 14,132 | 29 |
| 2,32E+08 | 2,85E+08 | 2,82E+08 | 2,95E+08 | 51,45 | 0 | 323,31 | 99 |
| 2,67E+10 | 3,40E+10 | 3,06E+10 | 3,08E+10 | 38,676 | 0 | 323,31 | 1229 |
| 9,70E+06 | 2,30E+06 | 3,82E+06 | 2,78E+06 | 43,437 | 0,000429 | 2,717 | 10 |
| 3,41E+08 | 2,39E+08 | 2,58E+08 | 2,77E+08 | 51,954 | 0 | 186,61 | 69 |
| 7,19E+07 | 9,61E+07 | 1,10E+08 | 9,97E+07 | 36,365 | 0 | 164,57 | 47 |
| 9,05E+06 | 1,35E+06 | 3,39E+06 | 2,02E+06 | 151,62 | 0,000852 | 2,5238 | 8 |
| 3,43E+06 | 9,34E+06 | 9,84E+06 | 1,27E+07 | 37,432 | 0 | 5,862 | 13 |
| 5,06E+07 | 1,77E+08 | 2,03E+08 | 2,44E+08 | 250,36 | 0 | 62,606 | 91 |
| 1,18E+08 | 6,98E+07 | 6,67E+07 | 8,67E+07 | 63,758 | 0 | 44,314 | 66 |
| 7,44E+08 | 5,68E+08 | 5,96E+08 | 6,22E+08 | 56,614 | 0 | 211,36 | 172 |
| 1,01E+08 | 1,46E+08 | 1,93E+08 | 1,89E+08 | 21,533 | 0 | 13,043 | 41 |
| 9,26E+07 | 7,03E+07 | 5,61E+07 | 6,37E+07 | 56,061 | 0 | 121,98 | 29 |
| 3,47E+08 | 5,05E+08 | 4,55E+08 | 4,59E+08 | 46,132 | 0 | 302,54 | 165 |
| 3,11E+07 | 2,34E+07 | 1,88E+07 | 2,06E+07 | 51,611 | 0 | 6,6535 | 19 |
| 1,44E+08 | 1,76E+08 | 1,67E+08 | 1,67E+08 | 40,612 | 0 | 21,116 | 64 |
| 1,07E+08 | 1,91E+08 | 1,60E+08 | 1,86E+08 | 87,16 | 0 | 132,91 | 71 |
| 6,00E+06 | 4,24E+06 | 3,04E+06 | 3,46E+06 | 49,306 | 0,000152 | 3,9628 | 10 |
| 3,69E+07 | 2,63E+06 | 2,62E+06 | 3,23E+06 | 83,65 | 0 | 10,305 | 15 |
| 7,92E+08 | 9,03E+08 | 9,66E+08 | 1,01E+09 | 31,893 | 0 | 323,31 | 180 |
| 1,53E+07 | 2,20E+07 | 2,19E+07 | 2,65E+07 | 52,685 | 0,000154 | 4,2878 | 11 |
| 4,27E+07 | 1,80E+08 | 1,39E+08 | 1,63E+08 | 67,811 | 0 | 30,97 | 46 |
| 3,30E+08 | 5,03E+08 | 4,15E+08 | 4,56E+08 | 21,694 | 0 | 323,31 | 103 |
| 2,90E+09 | 2,00E+09 | 2,31E+09 | 2,17E+09 | 100,2 | 0 | 323,31 | 496 |
| 2,24E+08 | 1,20E+08 | 1,40E+08 | 1,50E+08 | 19,606 | 0 | 60,109 | 71 |
| 9,42E+05 | 1,30E+08 | 1,26E+08 | 1,02E+08 | 22,794 | 0 | 37,426 | 27 |
| 8,85E+07 | 1,78E+07 | 5,03E+06 | 1,52E+07 | 142,02 | 0 | 53,056 | 73 |

| | | | | | | | |
|---|---|---|---|---|---|---|---|
| 7,22E+07 | 5,07E+07 | 5,36E+07 | 4,97E+07 | 49,619 | 0 | 202,36 | 35 |
| 1,83E+08 | 1,27E+08 | 1,17E+08 | 1,27E+08 | 33,31 | 0 | 300,89 | 31 |
| 5,93E+07 | 8,02E+07 | 8,17E+07 | 8,94E+07 | 29,227 | 0 | 25,169 | 48 |
| 1,16E+07 | 4,54E+07 | 3,87E+07 | 3,67E+07 | 136,77 | 0 | 8,2111 | 26 |
| 3,08E+09 | 2,41E+09 | 2,54E+09 | 2,48E+09 | 141,74 | 0 | 323,31 | 522 |
| 1,18E+09 | 1,35E+09 | 1,36E+09 | 1,30E+09 | 67,083 | 0 | 323,31 | 226 |
| 2,15E+08 | 2,83E+08 | 2,89E+08 | 2,68E+08 | 36,12 | 0 | 323,31 | 60 |
| 1,17E+07 | 8,25E+06 | 8,43E+06 | 7,62E+06 | 34,366 | 0,002644 | 2,0679 | 7 |
| 7,91E+08 | 6,15E+08 | 5,94E+08 | 5,76E+08 | 310,89 | 0 | 323,31 | 323 |
| 2,00E+08 | 2,26E+08 | 2,26E+08 | 2,36E+08 | 103,35 | 0 | 187,87 | 115 |
| 9,21E+08 | 1,48E+09 | 1,63E+09 | 1,82E+09 | 29,622 | 0 | 323,31 | 141 |
| 2,59E+09 | 2,26E+09 | 2,26E+09 | 2,41E+09 | 114,86 | 0 | 323,31 | 491 |
| 3,25E+07 | 1,64E+07 | 1,46E+07 | 1,69E+07 | 40,168 | 0 | 9,3945 | 35 |
| 4,49E+08 | 6,55E+08 | 6,17E+08 | 6,15E+08 | 58,727 | 0 | 323,31 | 142 |
| 2,02E+07 | 4,24E+07 | 4,21E+07 | 5,94E+07 | 40,978 | 0 | 25,58 | 11 |
| 3,60E+08 | 2,47E+08 | 2,26E+08 | 2,20E+08 | 46,797 | 0 | 125,71 | 140 |
| 2,84E+08 | 2,05E+08 | 1,70E+08 | 2,01E+08 | 28,09 | 0 | 71,776 | 34 |
| 1,39E+08 | 2,40E+08 | 2,27E+08 | 2,53E+08 | 76,187 | 0 | 323,31 | 97 |
| 1,03E+08 | 8,49E+07 | 9,28E+07 | 8,64E+07 | 69,971 | 0 | 66,676 | 58 |
| 1,00E+09 | 8,74E+08 | 7,99E+08 | 8,47E+08 | 41,275 | 0 | 238,68 | 156 |
| 3,17E+08 | 2,24E+08 | 2,32E+08 | 2,48E+08 | 85,682 | 0 | 75,598 | 178 |
| 3,30E+07 | 6,03E+07 | 6,20E+07 | 6,04E+07 | 100,78 | 0 | 58,621 | 48 |
| 2,85E+06 | 1,01E+07 | 1,12E+07 | 1,21E+07 | 61,867 | 0,000151 | 3,8901 | 9 |
| 2,45E+08 | 1,89E+08 | 2,06E+08 | 2,01E+08 | 93,092 | 0 | 265,26 | 113 |
| 5,32E+07 | 4,19E+07 | 4,50E+07 | 4,20E+07 | 87,441 | 0 | 24,311 | 60 |
| 4,76E+07 | 6,49E+07 | 7,17E+07 | 5,95E+07 | 197,06 | 0 | 17,716 | 58 |
| 1,70E+08 | 1,03E+08 | 1,31E+08 | 1,24E+08 | 54,093 | 0 | 136,09 | 85 |
| 2,89E+07 | 3,53E+07 | 4,21E+07 | 4,06E+07 | 25,493 | 0 | 8,3582 | 22 |
| 4,86E+08 | 6,29E+08 | 6,97E+08 | 6,62E+08 | 75,29 | 0 | 323,31 | 137 |
| 8,72E+07 | 1,51E+08 | 1,39E+08 | 1,26E+08 | 81,945 | 0 | 215,13 | 63 |
| 1,45E+08 | 9,80E+07 | 9,10E+07 | 1,13E+08 | 86,34 | 0 | 70,302 | 83 |
| 1,76E+07 | 7,79E+06 | 1,13E+07 | 1,03E+07 | 25,261 | 0,000571 | 2,6407 | 13 |
| 7,66E+06 | 3,10E+07 | 2,93E+07 | 3,27E+07 | 21,151 | 0 | 13,67 | 19 |
| 1,22E+07 | 1,76E+07 | 1,79E+07 | 1,73E+07 | 161,83 | 0 | 10,208 | 31 |
| 3,27E+07 | 4,19E+07 | 3,93E+07 | 4,44E+07 | 133,84 | 0 | 23,16 | 50 |
| 2,20E+08 | 2,67E+06 | 3,82E+06 | 1,97E+07 | 130,16 | 0 | 167,17 | 81 |
| 4,34E+08 | 5,69E+08 | 5,57E+08 | 5,29E+08 | 22,165 | 0 | 75,674 | 89 |
| 1,44E+08 | 1,15E+08 | 1,14E+08 | 1,12E+08 | 101,21 | 0 | 258,83 | 87 |
| 1,02E+08 | 6,37E+07 | 4,30E+07 | 5,87E+07 | 35,235 | 0 | 29,711 | 45 |
| 3,25E+08 | 5,72E+08 | 4,89E+08 | 5,08E+08 | 63,458 | 0 | 154,55 | 128 |
| 5,54E+06 | 4,80E+07 | 5,09E+07 | 4,67E+07 | 75,313 | 0 | 14,769 | 44 |
| 1,50E+09 | 9,37E+08 | 9,41E+08 | 1,00E+09 | 155,12 | 0 | 323,31 | 398 |
| 4,67E+07 | 7,11E+07 | 9,87E+07 | 8,86E+07 | 6,954 | 0,000154 | 4,295 | 28 |
| 8,39E+08 | 5,04E+08 | 5,44E+08 | 5,01E+08 | 32,296 | 0 | 144,22 | 88 |
| 8,10E+08 | 6,39E+08 | 6,18E+08 | 6,18E+08 | 72,322 | 0 | 323,31 | 262 |
| 5,02E+07 | 4,40E+07 | 4,29E+07 | 4,30E+07 | 37,408 | 0 | 13,713 | 38 |
| 2,43E+06 | 9,50E+06 | 6,88E+06 | 6,24E+06 | 32,625 | 0,000151 | 3,7612 | 10 |
| 2,45E+08 | 2,01E+08 | 1,91E+08 | 2,08E+08 | 25,776 | 0 | 136,9 | 113 |
| 8,95E+07 | 1,43E+08 | 1,44E+08 | 1,40E+08 | 100,58 | 0 | 75,804 | 84 |
| 1,47E+08 | 1,02E+08 | 1,02E+08 | 9,52E+07 | 50,569 | 0 | 81,2 | 80 |

| | | | | | | | |
|---|---|---|---|---|---|---|---|
| 2,20E+09 | 2,71E+09 | 2,57E+09 | 2,73E+09 | 92,866 | 0 | 323,31 | 551 |
| 1,15E+08 | 5,40E+07 | 6,75E+07 | 5,39E+07 | 148,52 | 0 | 77,267 | 88 |
| 2,10E+07 | 5,80E+07 | 4,56E+07 | 4,55E+07 | 41,937 | 0 | 58,339 | 27 |
| 1,35E+07 | 7,90E+07 | 8,04E+07 | 8,17E+07 | 121,28 | 0 | 21,779 | 41 |
| 3,11E+07 | 1,43E+07 | 1,41E+07 | 1,50E+07 | 86,379 | 0,000148 | 3,3168 | 12 |
| 1,19E+07 | 1,79E+07 | 1,85E+07 | 2,36E+07 | 12,73 | 0 | 5,7426 | 9 |
| 3,45E+07 | 4,23E+06 | 1,05E+07 | 3,61E+06 | 127,65 | 0 | 11,432 | 21 |
| 2,87E+07 | 6,93E+06 | 1,58E+06 | 1,79E+06 | 77,444 | 0 | 22,566 | 18 |
| 1,10E+08 | 7,40E+07 | 8,16E+07 | 7,18E+07 | 58,867 | 0 | 69,173 | 70 |
| 1,77E+07 | 2,25E+06 | 6,31E+06 | 2,73E+06 | 42,132 | 0,000158 | 5,1145 | 12 |
| 1,02E+08 | 5,61E+07 | 5,13E+07 | 4,67E+07 | 28,302 | 0 | 56,725 | 43 |
| 3,33E+09 | 2,76E+09 | 2,57E+09 | 2,82E+09 | 27,013 | 0 | 323,31 | 236 |
| 1,74E+08 | 1,39E+08 | 1,17E+08 | 1,15E+08 | 52,714 | 0 | 187,39 | 76 |
| 8,31E+07 | 1,50E+08 | 1,42E+08 | 1,53E+08 | 31,726 | 0 | 24,669 | 54 |
| 2,12E+07 | 4,91E+06 | 1,34E+06 | 2,96E+06 | 38,383 | 0 | 6,5056 | 13 |
| 5,36E+08 | 4,44E+08 | 4,78E+08 | 4,32E+08 | 118,07 | 0 | 103,12 | 114 |
| 2,85E+08 | 2,43E+08 | 2,25E+08 | 2,29E+08 | 61,217 | 0 | 323,31 | 127 |
| 3,09E+08 | 3,88E+08 | 3,54E+08 | 3,57E+08 | 68,77 | 0 | 119,22 | 150 |
| 1,27E+08 | 1,74E+08 | 1,66E+08 | 1,68E+08 | 82,341 | 0 | 79,967 | 108 |
| 2,63E+07 | 4,03E+07 | 4,37E+07 | 4,24E+07 | 30,608 | 0,000434 | 2,9112 | 12 |
| 1,13E+09 | 1,01E+09 | 9,68E+08 | 9,28E+08 | 217,17 | 0 | 323,31 | 448 |
| 1,77E+08 | 1,24E+08 | 1,23E+08 | 1,37E+08 | 74,077 | 0 | 36,44 | 92 |
| 2,19E+06 | 5,17E+07 | 4,53E+07 | 5,27E+07 | 28,504 | 0 | 15,455 | 28 |
| 3,39E+08 | 2,95E+08 | 2,61E+08 | 2,58E+08 | 71,078 | 0 | 214,42 | 149 |
| 5,79E+06 | 2,39E+07 | 1,95E+07 | 2,82E+07 | 210,94 | 0 | 12,016 | 26 |
| 6,20E+09 | 7,20E+09 | 7,14E+09 | 6,94E+09 | 269,82 | 0 | 323,31 | 1161 |
| 1,34E+08 | 7,86E+07 | 8,91E+07 | 8,93E+07 | 58,955 | 0 | 80,481 | 84 |
| 1,38E+08 | 9,83E+07 | 9,59E+07 | 8,02E+07 | 23,414 | 0 | 19,85 | 50 |
| 1,45E+09 | 2,10E+09 | 2,08E+09 | 1,98E+09 | 35,009 | 0 | 323,31 | 167 |
| 2,15E+07 | 4,12E+07 | 3,75E+07 | 3,47E+07 | 68,733 | 0 | 8,1309 | 24 |
| 3,12E+07 | 6,68E+07 | 7,80E+07 | 8,80E+07 | 59,534 | 0 | 23,088 | 26 |
| 9,88E+08 | 7,26E+08 | 7,46E+08 | 7,72E+08 | 74,112 | 0 | 198,87 | 213 |
| 2,55E+08 | 2,08E+08 | 2,11E+08 | 1,95E+08 | 42,699 | 0 | 87,017 | 108 |
| 7,68E+08 | 9,61E+08 | 9,50E+08 | 9,49E+08 | 117,96 | 0 | 323,31 | 334 |
| 3,20E+06 | 6,58E+06 | 6,90E+06 | 6,10E+06 | 80,214 | 0,000149 | 3,5112 | 12 |
| 9,67E+07 | 7,82E+07 | 7,82E+07 | 8,23E+07 | 51,827 | 0 | 147,37 | 73 |
| 3,99E+07 | 6,25E+06 | 2,03E+06 | 2,55E+06 | 27,585 | 0 | 7,1325 | 16 |
| 2,86E+08 | 3,46E+08 | 3,84E+08 | 4,06E+08 | 43,474 | 0 | 323,31 | 95 |
| 2,87E+09 | 3,91E+09 | 4,32E+09 | 3,79E+09 | 45,345 | 0 | 323,31 | 389 |
| 1,28E+08 | 1,68E+08 | 1,63E+08 | 1,65E+08 | 12,605 | 0 | 35,679 | 44 |
| 2,03E+07 | 2,14E+06 | 6,42E+06 | 2,67E+06 | 108,14 | 0 | 13,288 | 16 |
| 9,69E+07 | 1,36E+08 | 1,48E+08 | 1,48E+08 | 44,013 | 0 | 323,31 | 72 |
| 1,94E+07 | 1,52E+06 | 5,62E+06 | 1,95E+06 | 43,463 | 0 | 40,276 | 14 |
| 4,35E+07 | 7,02E+07 | 6,73E+07 | 5,55E+07 | 20,852 | 0 | 100,72 | 28 |
| 3,24E+07 | 3,34E+06 | 6,81E+06 | 1,45E+06 | 34,69 | 0 | 40,963 | 16 |
| 4,01E+08 | 6,05E+08 | 5,55E+08 | 6,34E+08 | 30,972 | 0 | 119,04 | 74 |
| 1,07E+08 | 8,22E+07 | 8,40E+07 | 9,44E+07 | 153,55 | 0 | 51,898 | 94 |
| 7,83E+08 | 6,25E+08 | 5,81E+08 | 5,58E+08 | 88,265 | 0 | 315,88 | 240 |
| 2,15E+08 | 2,50E+08 | 2,50E+08 | 2,38E+08 | 80,597 | 0 | 143,96 | 125 |
| 6,25E+06 | 1,66E+07 | 1,37E+07 | 1,41E+07 | 110,23 | 0,000152 | 4,046 | 9 |

| | | | | | | | |
|---|---|---|---|---|---|---|---|
| 5,79E+08 | 6,81E+08 | 7,09E+08 | 7,30E+08 | 121,64 | 0 | 323,31 | 227 |
| 9,09E+08 | 6,56E+08 | 7,18E+08 | 7,19E+08 | 45,133 | 0 | 323,31 | 148 |
| 4,67E+07 | 8,30E+07 | 7,15E+07 | 7,18E+07 | 61,18 | 0 | 81,23 | 67 |
| 1,30E+08 | 1,44E+08 | 1,42E+08 | 1,41E+08 | 38,97 | 0 | 171,89 | 103 |
| 2,57E+08 | 2,17E+08 | 2,08E+08 | 2,20E+08 | 60,123 | 0 | 119,45 | 126 |
| 1,44E+08 | 1,91E+08 | 2,11E+08 | 1,82E+08 | 35,159 | 0 | 71,568 | 72 |
| 4,10E+07 | 1,91E+07 | 2,19E+07 | 1,41E+07 | 45,37 | 0 | 18,72 | 33 |
| 2,41E+08 | 2,96E+08 | 3,21E+08 | 3,29E+08 | 18,977 | 0 | 102,38 | 65 |
| 5,71E+08 | 4,11E+08 | 3,78E+08 | 3,87E+08 | 75,435 | 0 | 74,205 | 126 |
| 1,35E+08 | 2,17E+08 | 2,12E+08 | 2,21E+08 | 138,85 | 0 | 74,511 | 89 |
| 1,18E+08 | 7,80E+07 | 5,65E+07 | 7,64E+07 | 11,987 | 0 | 13,465 | 28 |
| 4,53E+07 | 1,88E+07 | 1,56E+07 | 8,41E+06 | 16,718 | 0 | 37,757 | 21 |
| 3,46E+07 | 9,40E+06 | 1,55E+07 | 6,51E+06 | 124,79 | 0 | 23,427 | 36 |
| 1,32E+08 | 1,13E+08 | 1,18E+08 | 1,09E+08 | 39,713 | 0 | 160,45 | 62 |
| 6,29E+07 | 9,02E+07 | 8,60E+07 | 7,99E+07 | 113,32 | 0 | 37,162 | 78 |
| 2,82E+07 | 4,27E+06 | 4,06E+06 | 8,10E+06 | 67,031 | 0 | 6,3809 | 11 |
| 2,41E+07 | 1,41E+07 | 1,50E+07 | 1,09E+07 | 60,468 | 0 | 9,0058 | 20 |
| 1,62E+08 | 2,85E+08 | 3,08E+08 | 3,09E+08 | 31,715 | 0 | 43,453 | 91 |
| 1,56E+08 | 1,07E+08 | 1,15E+08 | 1,26E+08 | 26,678 | 0 | 28,757 | 53 |
| 1,28E+09 | 1,50E+09 | 1,68E+09 | 1,72E+09 | 48,968 | 0 | 191 | 187 |
| 3,75E+06 | 4,81E+07 | 4,66E+07 | 3,79E+07 | 164,54 | 0 | 18,745 | 25 |
| 1,38E+07 | 9,75E+06 | 9,72E+06 | 8,02E+06 | 20,642 | 0 | 5,4005 | 15 |
| 1,09E+09 | 8,35E+08 | 8,03E+08 | 8,65E+08 | 29,579 | 0 | 323,31 | 133 |
| 3,52E+08 | 3,13E+08 | 2,99E+08 | 2,82E+08 | 129,23 | 0 | 81,575 | 163 |
| 3,75E+08 | 5,02E+08 | 4,93E+08 | 4,42E+08 | 61,23 | 0 | 322,13 | 167 |
| 1,38E+07 | 2,53E+06 | 5,35E+06 | 4,49E+06 | 56,939 | 0,000434 | 2,8894 | 10 |
| 3,80E+07 | 6,05E+07 | 5,56E+07 | 5,80E+07 | 211,9 | 0 | 64,784 | 65 |
| 1,30E+07 | 9,14E+06 | 7,53E+06 | 9,14E+06 | 101,12 | 0 | 10,643 | 11 |
| 2,98E+08 | 3,71E+08 | 3,82E+08 | 3,87E+08 | 43,887 | 0 | 146,98 | 128 |
| 2,10E+08 | 1,58E+08 | 1,53E+08 | 1,54E+08 | 59,771 | 0 | 163,76 | 78 |
| 4,94E+06 | 1,22E+07 | 1,08E+07 | 1,27E+07 | 111,66 | 0,000159 | 5,2155 | 16 |
| 2,87E+06 | 4,39E+06 | 4,04E+06 | 4,59E+06 | 47,544 | 0,000432 | 2,8252 | 2 |
| 1,86E+06 | 1,05E+07 | 1,71E+07 | 1,71E+07 | 30,209 | 0 | 6,237 | 11 |
| 4,38E+07 | 5,31E+07 | 6,03E+07 | 5,55E+07 | 91,467 | 0 | 27,908 | 47 |
| 1,06E+07 | 2,75E+06 | 2,61E+06 | 9,65E+05 | 92,022 | 0 | 7,2022 | 9 |
| 2,09E+09 | 1,42E+09 | 1,43E+09 | 1,41E+09 | 67,839 | 0 | 323,31 | 327 |
| 1,16E+08 | 1,89E+08 | 1,66E+08 | 1,50E+08 | 65,862 | 0 | 202,62 | 70 |
| 1,20E+09 | 1,47E+09 | 1,40E+09 | 1,43E+09 | 95,832 | 0 | 323,31 | 273 |
| 7,65E+07 | 9,16E+07 | 8,90E+07 | 9,76E+07 | 46,587 | 0 | 55,526 | 52 |
| 8,52E+07 | 6,67E+07 | 7,31E+07 | 6,77E+07 | 50,069 | 0 | 145,81 | 42 |
| 1,41E+08 | 1,26E+08 | 1,16E+08 | 1,12E+08 | 82,056 | 0 | 66,335 | 64 |
| 2,12E+08 | 2,60E+08 | 2,68E+08 | 3,13E+08 | 72,654 | 0 | 165,36 | 96 |
| 2,84E+06 | 3,90E+08 | 3,13E+08 | 3,00E+08 | 20,538 | 0 | 56,312 | 75 |
| 4,89E+07 | 1,09E+08 | 9,69E+07 | 9,54E+07 | 35,31 | 0 | 44,438 | 52 |
| 3,62E+07 | 5,44E+07 | 6,07E+07 | 5,62E+07 | 26,306 | 0 | 6,2962 | 17 |
| 7,80E+07 | 1,13E+08 | 1,13E+08 | 1,06E+08 | 74,854 | 0 | 23,967 | 63 |
| 1,28E+06 | 7,63E+06 | 9,69E+06 | 1,18E+07 | 46,228 | 0,000155 | 4,5299 | 9 |
| 2,12E+09 | 1,79E+09 | 1,83E+09 | 1,69E+09 | 54,356 | 0 | 323,31 | 238 |
| 1,67E+08 | 1,30E+08 | 1,36E+08 | 1,39E+08 | 19,628 | 0 | 33,865 | 47 |
| 7,34E+08 | 5,65E+08 | 6,38E+08 | 6,25E+08 | 120,26 | 0 | 323,31 | 273 |

| | | | | | | | |
|---|---|---|---|---|---|---|---|
| 2,47E+07 | 3,21E+06 | 1,45E+06 | 6,25E+06 | 72,878 | 0,000159 | 5,1807 | 12 |
| 4,95E+08 | 4,09E+08 | 3,84E+08 | 3,44E+08 | 98,768 | 0 | 237,72 | 166 |
| 2,87E+07 | 2,24E+07 | 2,23E+07 | 2,22E+07 | 107,79 | 0 | 8,2613 | 20 |
| 5,96E+07 | 7,65E+07 | 8,03E+07 | 8,97E+07 | 66,455 | 0 | 96,013 | 51 |
| 1,50E+08 | 1,04E+08 | 1,18E+08 | 1,26E+08 | 27,328 | 0 | 31,758 | 65 |
| 5,71E+08 | 7,36E+08 | 7,82E+08 | 8,42E+08 | 14,275 | 0 | 109,54 | 65 |
| 8,13E+09 | 1,17E+10 | 1,28E+10 | 1,19E+10 | 39,355 | 0 | 323,31 | 351 |
| 1,21E+09 | 9,34E+08 | 9,42E+08 | 1,06E+09 | 93,28 | 0 | 253,11 | 322 |
| 7,44E+07 | 9,37E+07 | 1,01E+08 | 8,73E+07 | 41,349 | 0 | 73,125 | 70 |
| 2,44E+07 | 1,34E+07 | 1,53E+07 | 1,38E+07 | 66,924 | 0 | 50,412 | 23 |
| 3,30E+07 | 6,26E+07 | 6,57E+07 | 6,28E+07 | 139,95 | 0 | 36,933 | 53 |
| 1,28E+07 | 3,40E+06 | 4,28E+06 | 5,76E+06 | 28,616 | 0,000855 | 2,6098 | 8 |
| 1,58E+08 | 1,33E+08 | 1,39E+08 | 1,37E+08 | 35,848 | 0 | 148,02 | 76 |
| 5,14E+07 | 1,93E+07 | 2,66E+07 | 2,98E+07 | 41,06 | 0 | 226,76 | 24 |
| 1,21E+07 | 7,56E+06 | 9,40E+06 | 8,61E+06 | 48,732 | 0 | 5,6076 | 13 |
| 1,19E+08 | 6,93E+07 | 6,68E+07 | 6,30E+07 | 9,6601 | 0 | 8,0995 | 28 |
| 3,48E+08 | 4,97E+08 | 5,49E+08 | 4,59E+08 | 19,414 | 0 | 126,77 | 110 |
| 7,48E+07 | 3,86E+07 | 4,54E+07 | 5,46E+07 | 29,642 | 0 | 21,981 | 47 |
| 1,17E+08 | 9,72E+07 | 8,49E+07 | 9,81E+07 | 18,761 | 0 | 27,529 | 51 |
| 7,02E+08 | 8,34E+08 | 9,62E+08 | 9,64E+08 | 18,192 | 0 | 181,94 | 71 |
| 4,39E+07 | 2,55E+07 | 2,68E+07 | 3,21E+07 | 21,199 | 0 | 19,614 | 6 |
| 2,46E+08 | 3,71E+08 | 4,44E+08 | 3,56E+08 | 13,584 | 0 | 82,829 | 76 |
| 9,27E+07 | 1,12E+08 | 1,09E+08 | 1,11E+08 | 28,598 | 0 | 70,927 | 46 |
| 6,25E+08 | 6,84E+08 | 7,06E+08 | 7,09E+08 | 57,229 | 0 | 159,01 | 197 |
| 2,59E+07 | 3,68E+07 | 4,32E+07 | 4,43E+07 | 37,195 | 0 | 31,569 | 22 |
| 4,53E+08 | 6,67E+08 | 6,01E+08 | 5,82E+08 | 84,487 | 0 | 276,7 | 194 |
| 9,08E+07 | 7,90E+07 | 7,47E+07 | 7,77E+07 | 28,969 | 0 | 17,664 | 36 |
| 9,56E+07 | 6,64E+07 | 7,81E+07 | 7,39E+07 | 100,56 | 0 | 50,282 | 64 |
| 1,26E+08 | 1,02E+08 | 9,78E+07 | 9,30E+07 | 62,022 | 0 | 68,087 | 55 |
| 4,72E+08 | 5,50E+08 | 5,74E+08 | 5,47E+08 | 82,843 | 0 | 277,18 | 136 |
| 8,23E+07 | 1,89E+08 | 1,71E+08 | 1,65E+08 | 32,754 | 0 | 65,756 | 70 |
| 2,16E+08 | 2,53E+08 | 2,51E+08 | 2,73E+08 | 60,484 | 0 | 147,24 | 73 |
| 1,08E+08 | 8,21E+07 | 8,26E+07 | 9,00E+07 | 70,889 | 0 | 51,143 | 56 |
| 6,81E+07 | 3,90E+07 | 4,12E+07 | 5,19E+07 | 80,466 | 0 | 49,941 | 41 |
| 4,60E+06 | 6,66E+07 | 2,35E+07 | 5,78E+07 | 160,68 | 0 | 13,38 | 27 |
| 3,17E+07 | 3,99E+07 | 4,55E+07 | 5,14E+07 | 13,557 | 0 | 36,869 | 25 |
| 4,29E+07 | 1,14E+08 | 9,05E+07 | 1,03E+08 | 11,655 | 0 | 73,019 | 15 |
| 9,49E+07 | 1,34E+07 | 3,01E+07 | 2,34E+07 | 52,664 | 0 | 52,269 | 45 |
| 9,12E+08 | 5,96E+08 | 6,17E+08 | 5,41E+08 | 109,94 | 0 | 323,31 | 231 |
| 5,17E+06 | 9,91E+06 | 1,01E+07 | 7,13E+06 | 39,955 | 0 | 7,0458 | 11 |
| 8,00E+07 | 6,40E+07 | 6,82E+07 | 7,07E+07 | 31,344 | 0 | 67,34 | 69 |
| 7,44E+07 | 5,97E+07 | 5,72E+07 | 5,90E+07 | 63,237 | 0 | 23,2 | 42 |
| 8,79E+08 | 6,48E+08 | 6,23E+08 | 6,23E+08 | 128,62 | 0 | 323,31 | 297 |
| 9,38E+07 | 4,08E+08 | 3,17E+08 | 3,73E+08 | 58,643 | 0 | 78,982 | 49 |
| 6,44E+06 | 2,70E+07 | 3,73E+07 | 3,90E+07 | 20,262 | 0 | 35,865 | 16 |
| 2,81E+07 | 3,88E+07 | 4,93E+07 | 4,59E+07 | 37,772 | 0 | 143,6 | 23 |
| 3,14E+06 | 1,43E+07 | 1,99E+07 | 1,81E+07 | 53,663 | 0 | 6,9629 | 16 |
| 1,83E+06 | 5,99E+07 | 5,05E+07 | 3,88E+07 | 167,28 | 0 | 19,38 | 36 |
| 1,45E+08 | 1,30E+08 | 1,24E+08 | 1,22E+08 | 146,41 | 0 | 97,647 | 120 |
| 2,56E+08 | 2,26E+08 | 2,27E+08 | 2,05E+08 | 254,61 | 0 | 157,88 | 228 |

| | | | | | | | |
|---|---|---|---|---|---|---|---|
| 3,03E+09 | 2,40E+09 | 2,32E+09 | 2,44E+09 | 69,062 | 0 | 323,31 | 363 |
| 8,91E+06 | 2,17E+06 | 4,59E+06 | 2,90E+06 | 90,83 | 0,003174 | 1,9458 | 10 |
| 1,80E+07 | 2,85E+06 | 5,13E+06 | 2,27E+06 | 97,802 | 0 | 19,793 | 15 |
| 5,02E+09 | 3,93E+09 | 3,95E+09 | 4,07E+09 | 21,778 | 0 | 323,31 | 249 |
| 1,05E+07 | 3,42E+07 | 2,67E+07 | 2,89E+07 | 19,454 | 0 | 21,672 | 20 |
| 3,09E+07 | 6,03E+06 | 2,22E+06 | 6,22E+06 | 21,487 | 0 | 22,607 | 12 |
| 6,52E+06 | 3,14E+06 | 2,11E+06 | 3,60E+06 | 14,51 | 0,002099 | 2,1692 | 2 |
| 7,43E+07 | 3,95E+07 | 3,32E+07 | 4,34E+07 | 25,049 | 0 | 101,71 | 30 |
| 3,38E+07 | 4,40E+07 | 4,06E+07 | 4,32E+07 | 106,58 | 0 | 18,635 | 36 |
| 1,24E+08 | 9,00E+07 | 1,01E+08 | 9,82E+07 | 36,867 | 0 | 29,02 | 43 |
| 5,66E+07 | 9,90E+07 | 1,09E+08 | 1,18E+08 | 78,776 | 0 | 32,33 | 47 |
| 8,64E+07 | 7,42E+07 | 7,85E+07 | 7,47E+07 | 23,952 | 0 | 5,3427 | 17 |
| 2,23E+07 | 1,50E+06 | 4,51E+06 | 4,63E+06 | 47,766 | 0 | 60,768 | 34 |
| 9,53E+08 | 7,50E+08 | 7,85E+08 | 8,28E+08 | 57,543 | 0 | 323,31 | 197 |
| 4,08E+07 | 5,41E+07 | 5,09E+07 | 5,08E+07 | 61,445 | 0 | 10,114 | 33 |
| 4,69E+07 | 4,35E+06 | 4,18E+06 | 1,54E+07 | 49,408 | 0 | 16,851 | 34 |
| 2,04E+06 | 2,41E+07 | 2,04E+07 | 2,41E+07 | 41,226 | 0 | 5,4704 | 15 |
| 8,48E+08 | 5,37E+08 | 4,15E+08 | 5,60E+08 | 12,496 | 0 | 309,48 | 101 |
| 1,20E+08 | 9,98E+07 | 1,01E+08 | 9,08E+07 | 71,793 | 0 | 74,344 | 86 |
| 6,75E+08 | 5,89E+08 | 6,06E+08 | 6,14E+08 | 75,892 | 0 | 207,69 | 185 |
| 6,38E+06 | 1,65E+07 | 1,44E+07 | 1,74E+07 | 233,26 | 0,000153 | 4,1485 | 10 |
| 7,73E+06 | 3,06E+06 | 3,92E+06 | 4,95E+06 | 119,1 | 0,000158 | 5,1236 | 9 |
| 8,76E+07 | 6,40E+07 | 6,86E+07 | 5,73E+07 | 47,335 | 0 | 106,79 | 40 |
| 2,09E+08 | 1,63E+08 | 1,68E+08 | 1,54E+08 | 28,489 | 0 | 79,081 | 82 |
| 3,61E+08 | 2,70E+08 | 2,64E+08 | 2,38E+08 | 60,18 | 0 | 323,31 | 153 |
| 1,19E+08 | 1,63E+08 | 1,61E+08 | 1,74E+08 | 11,154 | 0 | 83,729 | 134 |
| 1,07E+07 | 2,47E+07 | 2,54E+07 | 2,48E+07 | 40,674 | 0 | 33,407 | 21 |
| 2,02E+07 | 1,57E+07 | 1,35E+07 | 1,39E+07 | 66,109 | 0 | 16,044 | 28 |
| 2,60E+09 | 2,74E+09 | 2,73E+09 | 2,84E+09 | 105,73 | 0 | 323,31 | 520 |
| 3,24E+07 | 2,91E+07 | 2,92E+07 | 3,05E+07 | 49,094 | 0 | 101,67 | 25 |
| 1,53E+07 | 2,78E+06 | 4,42E+06 | 2,11E+06 | 20,818 | 0 | 6,7308 | 8 |
| 3,41E+07 | 2,60E+07 | 2,09E+07 | 1,89E+07 | 73,275 | 0 | 20,072 | 35 |
| 6,50E+07 | 4,32E+07 | 4,45E+07 | 3,85E+07 | 43,876 | 0 | 84,226 | 49 |
| 2,26E+08 | 1,68E+08 | 1,95E+08 | 1,73E+08 | 59,744 | 0 | 55,414 | 89 |
| 2,34E+08 | 1,86E+08 | 1,64E+08 | 1,62E+08 | 36,942 | 0 | 148,51 | 63 |
| 5,46E+07 | 2,83E+07 | 3,00E+07 | 3,54E+07 | 152,17 | 0 | 38,534 | 38 |
| 4,85E+08 | 4,21E+08 | 4,03E+08 | 4,04E+08 | 41,192 | 0 | 96,103 | 137 |
| 1,23E+07 | 6,28E+06 | 3,62E+06 | 5,90E+06 | 247,08 | 0 | 6,7624 | 12 |
| 1,62E+08 | 2,37E+08 | 2,19E+08 | 2,20E+08 | 54,201 | 0 | 83,596 | 106 |
| 2,42E+09 | 2,22E+09 | 2,24E+09 | 2,25E+09 | 504,74 | 0 | 323,31 | 742 |
| 1,91E+09 | 1,48E+09 | 1,50E+09 | 1,47E+09 | 123,97 | 0 | 323,31 | 401 |
| 3,58E+07 | 7,82E+06 | 4,14E+06 | 3,65E+06 | 38,075 | 0 | 8,414 | 12 |
| 2,50E+07 | 4,49E+07 | 4,10E+07 | 4,30E+07 | 30,047 | 0 | 12,01 | 22 |
| 4,46E+07 | 3,45E+07 | 3,60E+07 | 3,92E+07 | 69,573 | 0 | 73,97 | 49 |
| 1,26E+07 | 8,38E+06 | 6,65E+06 | 7,62E+06 | 97,774 | 0 | 6,8482 | 14 |
| 1,46E+08 | 1,24E+08 | 1,15E+08 | 1,06E+08 | 71,459 | 0 | 274,13 | 93 |
| 3,86E+07 | 4,89E+07 | 5,34E+07 | 5,61E+07 | 51,984 | 0 | 48,73 | 41 |
| 1,46E+08 | 1,71E+08 | 1,60E+08 | 1,66E+08 | 59,964 | 0 | 266,69 | 66 |
| 4,65E+07 | 3,27E+06 | 1,49E+06 | 5,99E+06 | 96,658 | 0 | 19,591 | 22 |
| 1,57E+10 | 1,85E+10 | 2,12E+10 | 1,99E+10 | 72,421 | 0 | 323,31 | 1226 |

| | | | | | | | |
|---|---|---|---|---|---|---|---|
| 3,23E+08 | 3,55E+08 | 3,90E+08 | 3,62E+08 | 65,318 | 0 | 167,79 | 157 |
| 6,84E+08 | 8,44E+08 | 8,52E+08 | 7,81E+08 | 57,598 | 0 | 323,31 | 183 |
| 4,03E+07 | 6,86E+07 | 7,11E+07 | 6,72E+07 | 70,525 | 0 | 47,896 | 58 |
| 7,33E+07 | 1,01E+08 | 1,38E+08 | 1,23E+08 | 34,492 | 0 | 35,54 | 38 |
| 3,40E+08 | 5,05E+08 | 5,73E+08 | 5,75E+08 | 21,308 | 0 | 213,45 | 98 |
| 1,45E+07 | 6,39E+06 | 4,69E+06 | 2,55E+06 | 24,443 | 0 | 5,8415 | 9 |
| 5,09E+07 | 8,65E+07 | 8,58E+07 | 8,68E+07 | 328,31 | 0 | 51,981 | 83 |
| 3,14E+07 | 4,65E+07 | 5,25E+07 | 4,94E+07 | 27,166 | 0,000159 | 5,3034 | 14 |
| 6,61E+07 | 3,79E+07 | 4,04E+07 | 4,35E+07 | 95,694 | 0 | 99,233 | 54 |
| 3,46E+07 | 7,04E+06 | 1,86E+06 | 9,61E+05 | 20,165 | 0 | 323,31 | 42 |
| 2,73E+08 | 4,82E+08 | 3,98E+08 | 3,87E+08 | 49,035 | 0 | 323,31 | 197 |
| 4,36E+07 | 1,93E+07 | 2,56E+07 | 2,56E+07 | 11,993 | 0 | 15,928 | 15 |
| 2,34E+08 | 1,57E+08 | 1,89E+08 | 1,63E+08 | 35,147 | 0 | 146,94 | 77 |
| 5,49E+07 | 9,73E+06 | 2,09E+06 | 1,30E+06 | 94,154 | 0 | 28,402 | 43 |
| 3,43E+07 | 5,19E+07 | 4,57E+07 | 5,56E+07 | 34,971 | 0 | 17,102 | 34 |
| 2,94E+08 | 2,10E+08 | 2,33E+08 | 2,41E+08 | 18,757 | 0 | 162,42 | 87 |
| 3,95E+07 | 1,34E+07 | 1,73E+07 | 2,33E+07 | 72,201 | 0 | 16,11 | 28 |
| 1,31E+08 | 1,10E+08 | 1,08E+08 | 1,05E+08 | 87,004 | 0 | 36,306 | 104 |
| 1,66E+08 | 2,15E+08 | 2,34E+08 | 2,18E+08 | 55,028 | 0 | 204,16 | 100 |
| 5,16E+07 | 8,78E+05 | 7,63E+06 | 2,37E+06 | 159,18 | 0 | 43,014 | 47 |
| 2,10E+08 | 1,66E+08 | 1,60E+08 | 1,77E+08 | 51,456 | 0 | 148,03 | 125 |
| 2,39E+07 | 9,78E+06 | 8,66E+06 | 7,00E+06 | 62,118 | 0 | 102,4 | 18 |
| 7,85E+08 | 9,09E+08 | 9,85E+08 | 8,78E+08 | 89,508 | 0 | 191,5 | 232 |
| 1,48E+08 | 1,78E+08 | 1,80E+08 | 1,99E+08 | 54,713 | 0 | 90,714 | 73 |
| 8,03E+06 | 5,24E+06 | 3,99E+06 | 5,05E+06 | 42,527 | 0,004628 | 1,7221 | 6 |
| 2,62E+07 | 3,27E+07 | 2,96E+07 | 3,44E+07 | 51,588 | 0 | 8,9696 | 32 |
| 1,01E+09 | 5,69E+08 | 5,31E+08 | 5,83E+08 | 33,752 | 0 | 90,83 | 32 |
| 1,10E+09 | 1,60E+09 | 1,46E+09 | 1,41E+09 | 28,024 | 0 | 323,31 | 205 |
| 1,11E+08 | 2,25E+08 | 1,82E+08 | 2,00E+08 | 52,768 | 0 | 164,56 | 95 |
| 3,96E+08 | 2,42E+08 | 2,27E+08 | 2,30E+08 | 23,443 | 0 | 115,65 | 37 |
| 3,01E+08 | 2,15E+08 | 2,42E+08 | 2,37E+08 | 93,21 | 0 | 129 | 125 |
| 4,22E+07 | 3,11E+07 | 2,65E+07 | 3,38E+07 | 35,332 | 0 | 39,693 | 22 |
| 3,66E+08 | 2,86E+08 | 3,01E+08 | 2,85E+08 | 117,94 | 0 | 237,3 | 170 |
| 3,53E+06 | 4,42E+07 | 3,33E+07 | 3,89E+07 | 132,35 | 0 | 21,7 | 48 |
| 8,21E+07 | 5,32E+07 | 5,51E+07 | 6,72E+07 | 44,433 | 0 | 27,414 | 54 |
| 3,73E+09 | 3,29E+09 | 3,25E+09 | 3,36E+09 | 350,86 | 0 | 323,31 | 1008 |
| 2,68E+09 | 3,43E+09 | 3,24E+09 | 3,44E+09 | 24,423 | 0 | 169,48 | 134 |
| 9,56E+06 | 8,93E+05 | 2,46E+06 | 1,88E+06 | 100,72 | 0,000159 | 5,3009 | 7 |
| 5,28E+07 | 2,09E+07 | 2,13E+07 | 9,81E+06 | 97,466 | 0 | 23,63 | 30 |
| 3,10E+07 | 1,53E+07 | 1,52E+07 | 1,28E+07 | 38,677 | 0 | 11,046 | 13 |
| 5,39E+07 | 4,49E+07 | 4,15E+07 | 4,11E+07 | 87,986 | 0 | 65,607 | 58 |
| 1,78E+07 | 4,13E+06 | 5,86E+06 | 1,84E+06 | 36,764 | 0 | 5,7588 | 10 |
| 7,49E+08 | 1,01E+09 | 1,03E+09 | 9,83E+08 | 65,798 | 0 | 323,31 | 241 |
| 7,78E+08 | 5,04E+08 | 4,38E+08 | 4,09E+08 | 71,439 | 0 | 152,05 | 115 |
| 2,66E+07 | 4,17E+07 | 4,22E+07 | 4,50E+07 | 24,776 | 0 | 63,37 | 20 |
| 3,92E+07 | 2,86E+07 | 2,83E+07 | 2,82E+07 | 77,268 | 0 | 86,97 | 25 |
| 9,64E+06 | 2,25E+07 | 2,12E+07 | 2,78E+07 | 104,54 | 0 | 12,743 | 23 |
| 4,31E+06 | 3,70E+07 | 4,65E+07 | 4,28E+07 | 35,216 | 0 | 9,3853 | 14 |
| 1,55E+07 | 8,91E+06 | 8,20E+06 | 9,24E+06 | 39,056 | 0 | 6,7676 | 22 |
| 2,54E+07 | 5,93E+07 | 5,79E+07 | 5,82E+07 | 37,963 | 0 | 21,303 | 41 |

| | | | | | | | |
|---|---|---|---|---|---|---|---|
| 1,52E+09 | 1,87E+09 | 1,86E+09 | 1,85E+09 | 172,88 | 0 | 323,31 | 418 |
| 2,12E+08 | 3,30E+08 | 3,40E+08 | 3,53E+08 | 34,986 | 0 | 82,785 | 120 |
| 8,02E+07 | 1,05E+08 | 1,14E+08 | 9,36E+07 | 46,338 | 0 | 15,096 | 33 |
| 1,22E+09 | 9,73E+08 | 1,09E+09 | 9,64E+08 | 23,656 | 0 | 44,797 | 220 |
| 1,70E+09 | 2,06E+09 | 1,94E+09 | 1,89E+09 | 119,73 | 0 | 323,31 | 553 |
| 6,08E+07 | 9,33E+07 | 9,48E+07 | 9,69E+07 | 68,683 | 0 | 28,03 | 73 |
| 4,64E+06 | 1,35E+07 | 1,03E+07 | 2,47E+07 | 38,152 | 0 | 8,7205 | 18 |
| 7,68E+06 | 1,19E+07 | 1,00E+07 | 1,17E+07 | 35,411 | 0 | 9,5795 | 16 |
| 1,20E+08 | 9,96E+07 | 1,01E+08 | 1,08E+08 | 82,039 | 0 | 83,829 | 107 |
| 7,75E+06 | 4,09E+07 | 4,36E+07 | 5,34E+07 | 45,456 | 0 | 25,393 | 28 |
| 7,98E+07 | 4,34E+07 | 4,22E+07 | 4,54E+07 | 22,168 | 0 | 127,13 | 15 |
| 1,37E+08 | 1,69E+08 | 1,85E+08 | 2,09E+08 | 32,072 | 0 | 139,69 | 78 |
| 5,26E+07 | 6,48E+07 | 5,79E+07 | 6,22E+07 | 24,493 | 0 | 60,127 | 43 |
| 3,68E+08 | 4,48E+08 | 4,51E+08 | 4,64E+08 | 45,349 | 0 | 110,45 | 109 |
| 1,19E+08 | 1,47E+08 | 1,35E+08 | 1,55E+08 | 88,337 | 0 | 145,57 | 87 |
| 9,10E+08 | 5,67E+08 | 5,89E+08 | 5,40E+08 | 39,502 | 0 | 323,31 | 144 |
| 1,13E+08 | 7,25E+07 | 8,61E+07 | 9,08E+07 | 19,478 | 0 | 10,591 | 30 |
| 2,97E+06 | 1,04E+08 | 8,72E+07 | 9,99E+07 | 161,66 | 0 | 33,342 | 46 |
| 4,46E+06 | 2,16E+06 | 3,05E+06 | 2,04E+06 | 53,571 | 0,000159 | 5,2226 | 11 |
| 4,86E+08 | 3,45E+08 | 3,32E+08 | 3,13E+08 | 65,615 | 0 | 323,31 | 139 |
| 1,45E+07 | 8,36E+06 | 6,18E+06 | 9,01E+06 | 59,231 | 0,000159 | 5,2249 | 20 |
| 2,57E+07 | 1,28E+07 | 1,11E+07 | 7,90E+06 | 10,101 | 0,00015 | 3,6481 | 5 |
| 6,31E+07 | 1,09E+08 | 1,16E+08 | 1,11E+08 | 47,406 | 0 | 35,77 | 81 |
| 7,51E+07 | 4,92E+07 | 3,38E+07 | 3,49E+07 | 38,098 | 0 | 9,0294 | 21 |
| 4,79E+07 | 1,12E+06 | 8,67E+06 | 2,55E+06 | 24,35 | 0,000158 | 5,1559 | 11 |
| 4,40E+07 | 1,95E+07 | 2,93E+07 | 2,77E+07 | 81,462 | 0 | 24,446 | 47 |
| 9,07E+07 | 1,84E+07 | 2,99E+06 | 2,43E+06 | 42,713 | 0 | 186,28 | 48 |
| 4,22E+07 | 7,60E+06 | 1,09E+06 | 1,49E+06 | 126,74 | 0 | 10,587 | 27 |
| 8,99E+08 | 9,87E+08 | 9,38E+08 | 9,60E+08 | 116,8 | 0 | 323,31 | 188 |
| 1,03E+09 | 1,17E+09 | 1,24E+09 | 1,20E+09 | 28,532 | 0 | 323,31 | 205 |
| 6,49E+08 | 3,35E+06 | 2,73E+07 | 5,02E+06 | 182,17 | 0 | 310,14 | 81 |
| 5,74E+07 | 1,96E+07 | 9,61E+06 | 1,08E+07 | 21,482 | 0 | 33,437 | 24 |
| 1,14E+07 | 6,80E+06 | 6,86E+06 | 8,37E+06 | 47,538 | 0 | 8,2941 | 10 |
| 3,30E+08 | 2,81E+08 | 2,80E+08 | 2,72E+08 | 33,541 | 0 | 128,98 | 96 |
| 1,07E+08 | 8,36E+07 | 7,98E+07 | 9,17E+07 | 94,118 | 0 | 25,311 | 80 |
| 1,42E+08 | 1,18E+08 | 1,20E+08 | 1,19E+08 | 62,981 | 0 | 120,83 | 92 |
| 3,85E+07 | 4,55E+07 | 5,19E+07 | 4,59E+07 | 121,71 | 0 | 10,723 | 30 |
| 1,94E+08 | 1,30E+08 | 1,19E+08 | 1,18E+08 | 127,74 | 0 | 135,5 | 115 |
| 9,83E+08 | 8,64E+08 | 8,35E+08 | 8,73E+08 | 153,49 | 0 | 323,31 | 335 |
| 7,28E+06 | 3,88E+06 | 3,67E+06 | 2,92E+06 | 31,155 | 0,000147 | 3,1809 | 6 |
| 1,14E+08 | 9,50E+07 | 1,02E+08 | 1,00E+08 | 54,033 | 0 | 51,306 | 69 |
| 4,14E+07 | 5,00E+07 | 5,15E+07 | 4,86E+07 | 205,94 | 0 | 28,176 | 65 |
| 1,49E+08 | 2,22E+08 | 2,27E+08 | 2,26E+08 | 21,311 | 0 | 10,69 | 28 |
| 1,46E+07 | 5,31E+06 | 5,51E+06 | 2,79E+06 | 31,193 | 0 | 9,1427 | 16 |
| 1,95E+06 | 1,25E+07 | 1,21E+07 | 1,40E+07 | 46,311 | 0 | 15,398 | 3 |
| 1,42E+07 | 1,21E+07 | 9,97E+06 | 1,05E+07 | 42,304 | 0 | 6,3556 | 14 |
| 2,95E+07 | 2,31E+07 | 2,23E+07 | 2,61E+07 | 26,76 | 0 | 6,2534 | 20 |
| 2,20E+07 | 2,78E+07 | 3,12E+07 | 3,01E+07 | 151,25 | 0 | 16,043 | 32 |
| 1,40E+08 | 1,66E+08 | 1,77E+08 | 1,64E+08 | 408,21 | 0 | 123,59 | 163 |
| 1,52E+08 | 1,05E+08 | 1,07E+08 | 1,10E+08 | 52,597 | 0 | 185,02 | 62 |

| | | | | | | | |
|---|---|---|---|---|---|---|---|
| 9,69E+07 | 1,30E+08 | 1,34E+08 | 1,25E+08 | 117,08 | 0 | 54,235 | 93 |
| 7,81E+08 | 1,26E+09 | 1,25E+09 | 1,35E+09 | 9,4771 | 0 | 12,412 | 37 |
| 1,49E+09 | 1,25E+09 | 1,26E+09 | 1,18E+09 | 68,539 | 0 | 323,31 | 284 |
| 3,34E+08 | 3,02E+08 | 2,87E+08 | 2,92E+08 | 68,46 | 0 | 126,52 | 162 |
| 3,15E+08 | 2,54E+08 | 2,49E+08 | 2,63E+08 | 85,423 | 0 | 296,97 | 112 |
| 1,32E+08 | 1,66E+08 | 1,70E+08 | 1,62E+08 | 126,62 | 0 | 160,43 | 124 |
| 9,35E+09 | 8,63E+09 | 8,76E+09 | 8,59E+09 | 47,153 | 0 | 323,31 | 607 |
| 3,47E+07 | 2,78E+07 | 2,93E+07 | 2,51E+07 | 46,04 | 0,000155 | 4,4238 | 7 |
| 3,78E+07 | 5,86E+07 | 5,96E+07 | 5,73E+07 | 40,32 | 0 | 23,23 | 42 |
| 2,13E+08 | 2,51E+08 | 2,40E+08 | 2,52E+08 | 51,354 | 0 | 151,19 | 136 |
| 6,80E+06 | 1,44E+07 | 1,13E+07 | 1,10E+07 | 22,514 | 0 | 9,8949 | 23 |
| 3,00E+07 | 1,85E+07 | 2,01E+07 | 1,72E+07 | 15,437 | 0 | 7,0421 | 16 |
| 3,47E+07 | 4,71E+07 | 5,14E+07 | 5,00E+07 | 66,204 | 0 | 21,949 | 30 |
| 1,15E+07 | 5,15E+06 | 2,41E+06 | 4,17E+06 | 58,005 | 0 | 8,9235 | 10 |
| 2,25E+07 | 2,91E+07 | 3,58E+07 | 3,84E+07 | 42,896 | 0 | 20,277 | 26 |
| 2,49E+08 | 1,40E+08 | 1,40E+08 | 1,48E+08 | 52,383 | 0 | 323,31 | 102 |
| 8,47E+08 | 1,05E+09 | 9,46E+08 | 9,69E+08 | 59,262 | 0 | 323,31 | 259 |
| 3,14E+08 | 2,87E+06 | 3,90E+06 | 3,54E+07 | 128,9 | 0 | 323,31 | 139 |
| 2,30E+07 | 8,07E+06 | 2,99E+06 | 6,14E+06 | 82,407 | 0 | 11,462 | 16 |
| 6,86E+07 | 1,09E+08 | 9,53E+07 | 1,09E+08 | 33,34 | 0 | 49,445 | 58 |
| 2,13E+08 | 1,55E+06 | 1,21E+06 | 1,17E+07 | 87,331 | 0 | 81,29 | 47 |
| 1,59E+06 | 2,43E+07 | 2,64E+07 | 2,45E+07 | 70,775 | 0 | 9,1404 | 21 |
| 6,70E+07 | 9,01E+07 | 8,86E+07 | 9,63E+07 | 81,737 | 0 | 24,968 | 72 |
| 5,09E+07 | 3,24E+07 | 2,90E+07 | 3,45E+07 | 45,382 | 0 | 32,197 | 35 |
| 3,33E+07 | 3,15E+06 | 1,67E+06 | 8,51E+06 | 34,134 | 0 | 16,689 | 17 |
| 4,61E+07 | 7,21E+07 | 6,61E+07 | 6,77E+07 | 56,078 | 0 | 78,546 | 45 |
| 2,94E+08 | 2,44E+08 | 2,51E+08 | 2,55E+08 | 67,175 | 0 | 117,71 | 111 |
| 6,33E+08 | 4,38E+08 | 4,73E+08 | 4,98E+08 | 41,11 | 0 | 153,73 | 96 |
| 2,16E+09 | 1,73E+09 | 1,77E+09 | 1,82E+09 | 39,638 | 0 | 193,97 | 220 |
| 7,65E+07 | 6,17E+07 | 5,38E+07 | 5,47E+07 | 60,522 | 0 | 41,298 | 64 |
| 5,31E+09 | 4,60E+09 | 4,28E+09 | 3,86E+09 | 77,648 | 0 | 323,31 | 935 |
| 2,62E+08 | 2,41E+08 | 2,30E+08 | 2,28E+08 | 80,86 | 0 | 323,31 | 128 |
| 1,04E+08 | 1,32E+08 | 1,34E+08 | 1,22E+08 | 20,747 | 0 | 74,301 | 107 |
| 5,93E+09 | 1,01E+10 | 1,10E+10 | 1,09E+10 | 32,191 | 0 | 323,31 | 537 |
| 6,28E+07 | 4,27E+07 | 4,28E+07 | 3,57E+07 | 38,287 | 0 | 14,816 | 43 |
| 4,40E+08 | 4,09E+08 | 3,78E+08 | 3,70E+08 | 82,435 | 0 | 296,66 | 145 |
| 7,61E+07 | 6,61E+07 | 6,15E+07 | 6,86E+07 | 107,58 | 0 | 29,927 | 76 |
| 4,97E+06 | 1,88E+07 | 1,78E+07 | 1,94E+07 | 49,75 | 0 | 7,6268 | 17 |
| 5,90E+08 | 7,26E+08 | 7,75E+08 | 7,05E+08 | 50,56 | 0 | 136,21 | 200 |
| 2,99E+07 | 1,99E+07 | 2,01E+07 | 2,35E+07 | 102,31 | 0 | 10,588 | 31 |
| 1,44E+08 | 1,64E+08 | 1,60E+08 | 1,55E+08 | 83,725 | 0 | 82,823 | 115 |
| 2,63E+08 | 1,15E+08 | 1,31E+08 | 1,19E+08 | 93,275 | 0 | 45,455 | 57 |
| 1,05E+07 | 7,51E+06 | 5,40E+06 | 7,07E+06 | 24,441 | 0,000156 | 4,6116 | 13 |
| 1,66E+08 | 4,99E+07 | 6,05E+07 | 5,31E+07 | 66,357 | 0 | 323,31 | 89 |
| 1,13E+08 | 7,33E+06 | 1,99E+07 | 2,52E+06 | 114,65 | 0 | 31,318 | 50 |
| 7,37E+08 | 6,43E+08 | 6,02E+08 | 6,60E+08 | 143,13 | 0 | 323,31 | 274 |
| 4,15E+07 | 2,40E+07 | 2,98E+07 | 2,37E+07 | 42,799 | 0 | 7,6861 | 26 |
| 6,09E+07 | 3,92E+07 | 4,56E+07 | 4,94E+07 | 31,058 | 0 | 12,153 | 26 |
| 1,86E+07 | 1,59E+07 | 1,50E+07 | 1,65E+07 | 48,883 | 0 | 16,572 | 18 |
| 1,02E+07 | 8,40E+06 | 6,64E+06 | 7,41E+06 | 73,853 | 0 | 11,615 | 21 |

| | | | | | | | |
|---|---|---|---|---|---|---|---|
| 2,06E+08 | 2,73E+08 | 2,91E+08 | 3,21E+08 | 17,018 | 0 | 49,426 | 78 |
| 8,49E+06 | 3,93E+07 | 2,18E+07 | 1,89E+07 | 76,487 | 0 | 8,3118 | 22 |
| 9,80E+08 | 8,38E+08 | 8,56E+08 | 8,77E+08 | 133,91 | 0 | 323,31 | 350 |
| 2,85E+07 | 4,81E+07 | 5,04E+07 | 5,84E+07 | 127,71 | 0 | 10,695 | 44 |
| 2,11E+07 | 3,02E+07 | 3,01E+07 | 3,69E+07 | 11,877 | 0 | 5,5766 | 20 |
| 3,37E+06 | 5,68E+06 | 5,00E+06 | 7,13E+06 | 63,362 | 0,008979 | 1,4561 | 6 |
| 5,21E+07 | 9,21E+07 | 9,28E+07 | 1,01E+08 | 42,747 | 0 | 13,876 | 56 |
| 2,52E+09 | 2,07E+09 | 2,15E+09 | 2,04E+09 | 34,384 | 0 | 208,43 | 301 |
| 2,78E+08 | 1,99E+08 | 2,28E+08 | 2,18E+08 | 18,607 | 0 | 323,31 | 35 |
| 1,96E+08 | 1,26E+08 | 1,01E+08 | 1,24E+08 | 48,477 | 0 | 257,5 | 57 |
| 2,08E+08 | 2,54E+08 | 2,59E+08 | 2,65E+08 | 23,914 | 0 | 29,501 | 79 |
| 3,56E+07 | 8,13E+07 | 7,75E+07 | 8,26E+07 | 20,619 | 0 | 14,912 | 18 |
| 8,30E+06 | 1,25E+08 | 1,78E+08 | 1,17E+08 | 22,417 | 0 | 212,61 | 26 |
| 2,40E+08 | 2,71E+08 | 2,90E+08 | 2,94E+08 | 88,053 | 0 | 246,89 | 148 |
| 1,51E+08 | 2,22E+08 | 1,97E+08 | 2,13E+08 | 25,452 | 0 | 142,62 | 79 |
| 1,59E+08 | 2,39E+08 | 2,40E+08 | 2,06E+08 | 99,523 | 0 | 55,414 | 71 |
| 2,42E+08 | 1,90E+08 | 1,85E+08 | 2,08E+08 | 41,747 | 0 | 193,88 | 93 |
| 3,30E+07 | 2,52E+07 | 2,11E+07 | 2,63E+07 | 32,941 | 0 | 14,764 | 19 |
| 4,46E+07 | 7,82E+07 | 8,35E+07 | 8,26E+07 | 48,774 | 0 | 37,003 | 54 |
| 2,99E+06 | 1,07E+06 | 1,37E+06 | 1,45E+06 | 81,783 | 0,00237 | 2,097 | 4 |
| 7,79E+06 | 3,26E+06 | 1,89E+06 | 3,95E+06 | 47,115 | 0,000157 | 4,9473 | 9 |
| 7,04E+07 | 1,22E+08 | 1,10E+08 | 9,21E+07 | 41,555 | 0 | 36,01 | 44 |
| 2,93E+08 | 2,12E+08 | 2,23E+08 | 2,03E+08 | 83,345 | 0 | 44,422 | 102 |
| 1,27E+07 | 2,96E+07 | 2,75E+07 | 2,27E+07 | 43,746 | 0 | 6,7934 | 16 |
| 1,43E+08 | 1,72E+08 | 1,74E+08 | 1,72E+08 | 6,6478 | 0 | 6,4886 | 34 |
| 8,95E+08 | 1,10E+09 | 1,00E+09 | 1,05E+09 | 133,73 | 0 | 323,31 | 352 |
| 1,40E+07 | 3,09E+07 | 2,75E+07 | 3,47E+07 | 101,02 | 0 | 20,299 | 30 |
| 8,63E+08 | 1,14E+09 | 1,05E+09 | 1,15E+09 | 46,67 | 0 | 323,31 | 202 |
| 1,22E+08 | 9,42E+07 | 9,44E+07 | 8,70E+07 | 41,624 | 0 | 54,156 | 55 |
| 1,45E+08 | 1,78E+08 | 1,77E+08 | 2,02E+08 | 147,85 | 0 | 55,834 | 130 |
| 9,88E+07 | 7,92E+07 | 7,76E+07 | 8,87E+07 | 13,331 | 0 | 39,919 | 48 |
| 1,51E+08 | 1,24E+08 | 1,17E+08 | 1,21E+08 | 66,275 | 0 | 126,4 | 85 |
| 5,14E+07 | 8,45E+07 | 9,03E+07 | 7,35E+07 | 98,921 | 0 | 39,204 | 70 |
| 2,15E+08 | 1,67E+08 | 1,59E+08 | 1,75E+08 | 55,707 | 0 | 113,53 | 100 |
| 9,20E+07 | 1,25E+08 | 1,27E+08 | 1,36E+08 | 88,196 | 0 | 53,897 | 100 |
| 2,05E+08 | 1,29E+08 | 1,22E+08 | 1,24E+08 | 27,656 | 0 | 56,124 | 57 |
| 2,17E+07 | 2,68E+07 | 2,59E+07 | 2,69E+07 | 103,68 | 0 | 18,346 | 35 |
| 4,05E+07 | 4,82E+07 | 4,51E+07 | 4,61E+07 | 131,84 | 0 | 20,683 | 43 |
| 7,19E+06 | 2,69E+06 | 3,28E+06 | 1,35E+06 | 75,109 | 0,000148 | 3,3619 | 8 |
| 1,87E+08 | 9,40E+07 | 1,02E+08 | 1,41E+08 | 17,6 | 0 | 133,08 | 49 |
| 4,01E+07 | 2,16E+07 | 1,31E+07 | 2,15E+07 | 39,881 | 0 | 9,1617 | 22 |
| 9,23E+07 | 1,19E+08 | 1,21E+08 | 1,23E+08 | 77,378 | 0 | 49,593 | 70 |
| 9,09E+08 | 8,10E+08 | 7,88E+08 | 7,16E+08 | 106,72 | 0 | 299,22 | 297 |
| 5,73E+08 | 4,90E+08 | 4,61E+08 | 4,58E+08 | 69,228 | 0 | 87,596 | 197 |
| 5,88E+08 | 7,53E+08 | 7,53E+08 | 7,42E+08 | 224,13 | 0 | 318,31 | 291 |
| 1,44E+08 | 1,67E+08 | 1,74E+08 | 1,91E+08 | 26,738 | 0 | 178,19 | 68 |
| 4,29E+08 | 5,87E+08 | 6,09E+08 | 6,35E+08 | 44,373 | 0 | 33,129 | 59 |
| 3,30E+08 | 2,27E+08 | 2,44E+08 | 2,57E+08 | 46,648 | 0 | 323,31 | 168 |
| 8,68E+07 | 7,44E+07 | 7,59E+07 | 7,41E+07 | 30,887 | 0 | 45,924 | 59 |
| 1,09E+08 | 9,20E+07 | 7,73E+07 | 8,63E+07 | 101,48 | 0 | 32,869 | 98 |

| | | | | | | | |
|---|---|---|---|---|---|---|---|
| 5,66E+07 | 4,68E+07 | 4,71E+07 | 5,07E+07 | 38,789 | 0 | 12,497 | 37 |
| 6,23E+08 | 5,24E+08 | 5,35E+08 | 5,09E+08 | 212,98 | 0 | 323,31 | 300 |
| 4,35E+07 | 3,35E+07 | 3,46E+07 | 3,68E+07 | 60,399 | 0 | 16,035 | 37 |
| 8,65E+07 | 3,98E+07 | 4,13E+07 | 5,14E+07 | 41,313 | 0 | 47,27 | 40 |
| 1,40E+08 | 1,83E+08 | 1,87E+08 | 1,68E+08 | 56,87 | 0 | 115,25 | 67 |
| 5,05E+06 | 1,83E+07 | 1,52E+07 | 1,66E+07 | 26,133 | 0,000151 | 3,7907 | 7 |
| 5,63E+08 | 5,27E+08 | 5,27E+08 | 5,46E+08 | 194,75 | 0 | 307,83 | 280 |
| 1,88E+07 | 1,39E+06 | 3,23E+06 | 6,37E+06 | 17,897 | 0 | 7,9921 | 10 |
| 4,36E+07 | 6,38E+07 | 6,81E+07 | 5,78E+07 | 46,852 | 0 | 24,563 | 37 |
| 5,92E+07 | 1,39E+07 | 2,37E+07 | 2,97E+07 | 53,094 | 0 | 17,617 | 37 |
| 1,22E+09 | 8,70E+08 | 8,91E+08 | 7,78E+08 | 20,809 | 0 | 250,36 | 155 |
| 3,54E+07 | 4,39E+07 | 4,62E+07 | 4,64E+07 | 56,708 | 0 | 26,546 | 34 |
| 1,15E+08 | 8,40E+07 | 8,74E+07 | 8,61E+07 | 90,66 | 0 | 67,04 | 77 |
| 2,52E+08 | 1,46E+08 | 1,41E+08 | 1,31E+08 | 35,658 | 0 | 194,27 | 69 |
| 5,52E+08 | 5,17E+08 | 4,93E+08 | 4,93E+08 | 85,893 | 0 | 138,44 | 205 |
| 4,79E+07 | 8,98E+07 | 6,88E+07 | 6,47E+07 | 26,519 | 0 | 86,495 | 51 |
| 2,17E+06 | 4,09E+07 | 9,87E+07 | 1,03E+08 | 187,37 | 0 | 25,579 | 51 |
| 2,71E+08 | 2,30E+08 | 2,45E+08 | 2,45E+08 | 32,168 | 0 | 323,31 | 97 |
| 3,24E+06 | 1,64E+06 | 1,24E+06 | 1,98E+06 | 180,25 | 0,001546 | 2,2938 | 3 |
| 5,37E+07 | 7,36E+07 | 6,41E+07 | 7,15E+07 | 28,705 | 0 | 16,703 | 39 |
| 4,09E+07 | 1,04E+07 | 3,80E+06 | 9,98E+06 | 119,64 | 0 | 15,178 | 18 |
| 6,91E+07 | 1,07E+08 | 1,01E+08 | 1,07E+08 | 172,44 | 0 | 44,695 | 79 |
| 1,54E+08 | 1,35E+08 | 1,27E+08 | 1,38E+08 | 45,755 | 0 | 124,09 | 96 |
| 5,86E+07 | 5,35E+07 | 5,07E+07 | 5,16E+07 | 155,27 | 0 | 33,382 | 77 |
| 8,24E+07 | 1,05E+08 | 9,72E+07 | 1,03E+08 | 72,602 | 0 | 25,82 | 84 |
| 1,22E+07 | 1,68E+07 | 1,64E+07 | 1,54E+07 | 24,908 | 0 | 10,517 | 14 |
| 3,14E+07 | 4,81E+07 | 4,26E+07 | 4,20E+07 | 45,785 | 0 | 55,313 | 30 |
| 1,51E+07 | 8,96E+06 | 1,12E+07 | 1,12E+07 | 86,355 | 0 | 18,873 | 25 |
| 2,48E+08 | 1,76E+08 | 1,98E+08 | 1,77E+08 | 87,56 | 0 | 155,03 | 132 |
| 9,89E+07 | 1,18E+08 | 1,15E+08 | 1,22E+08 | 30,977 | 0 | 33,11 | 71 |
| 3,04E+07 | 3,73E+07 | 4,03E+07 | 4,43E+07 | 105,33 | 0 | 9,9551 | 35 |
| 1,50E+08 | 1,93E+08 | 2,03E+08 | 2,08E+08 | 62,829 | 0 | 291,82 | 118 |
| 1,14E+09 | 6,27E+08 | 7,26E+08 | 6,61E+08 | 29,994 | 0 | 278,75 | 138 |
| 1,51E+08 | 1,60E+08 | 1,57E+08 | 1,56E+08 | 168,86 | 0 | 133,6 | 148 |
| 5,61E+07 | 6,60E+07 | 6,85E+07 | 6,06E+07 | 10,261 | 0 | 12,643 | 27 |
| 7,64E+07 | 1,32E+08 | 1,27E+08 | 1,32E+08 | 29,435 | 0 | 11,902 | 44 |
| 1,17E+08 | 8,90E+07 | 1,03E+08 | 9,49E+07 | 35,217 | 0 | 92,397 | 42 |
| 3,42E+08 | 5,00E+08 | 5,23E+08 | 5,25E+08 | 110,7 | 0 | 322,53 | 155 |
| 4,36E+07 | 5,24E+07 | 5,26E+07 | 4,84E+07 | 125,53 | 0 | 35,48 | 50 |
| 6,95E+06 | 2,86E+07 | 3,13E+07 | 3,60E+07 | 24,429 | 0 | 13,018 | 18 |
| 1,54E+07 | 2,64E+07 | 2,51E+07 | 2,13E+07 | 19,1 | 0 | 14,484 | 15 |
| 5,83E+07 | 3,76E+07 | 4,56E+07 | 4,68E+07 | 20,677 | 0 | 22,348 | 27 |
| 1,07E+07 | 1,41E+07 | 1,33E+07 | 1,58E+07 | 106,54 | 0 | 9,2546 | 13 |
| 9,53E+06 | 3,64E+06 | 3,17E+06 | 5,18E+06 | 104,53 | 0 | 5,4491 | 12 |
| 1,07E+08 | 5,95E+07 | 7,17E+07 | 8,24E+07 | 50,933 | 0 | 41,077 | 66 |
| 4,56E+07 | 2,16E+06 | 1,08E+07 | 3,15E+06 | 148,97 | 0,000147 | 3,2921 | 8 |
| 4,18E+07 | 2,03E+07 | 2,14E+07 | 1,93E+07 | 52,297 | 0 | 77,042 | 38 |
| 1,23E+07 | 1,86E+07 | 1,85E+07 | 1,90E+07 | 31,497 | 0,000157 | 4,9748 | 14 |
| 7,58E+07 | 3,01E+07 | 1,23E+07 | 2,86E+07 | 213,03 | 0 | 85,552 | 73 |
| 6,59E+07 | 5,30E+07 | 4,30E+07 | 4,06E+07 | 42,067 | 0 | 11,874 | 32 |

| | | | | | | | |
|---|---|---|---|---|---|---|---|
| 8,99E+06 | 2,87E+07 | 3,10E+07 | 2,02E+07 | 47,016 | 0 | 12,841 | 19 |
| 8,71E+06 | 4,31E+06 | 4,71E+06 | 4,60E+06 | 81,858 | 0,000151 | 3,8813 | 13 |
| 7,69E+06 | 4,11E+06 | 5,70E+06 | 3,65E+06 | 17,913 | 0,000152 | 4,0553 | 14 |
| 1,04E+09 | 9,39E+08 | 8,92E+08 | 9,69E+08 | 68,721 | 0 | 232,88 | 219 |
| 3,00E+08 | 2,47E+08 | 2,41E+08 | 2,48E+08 | 42,116 | 0 | 163,43 | 111 |
| 5,07E+07 | 1,55E+08 | 1,53E+08 | 1,46E+08 | 42,532 | 0 | 37,153 | 38 |
| 2,43E+07 | 2,78E+06 | 8,61E+06 | 3,13E+06 | 107,83 | 0 | 51,04 | 14 |
| 6,89E+07 | 1,26E+07 | 4,34E+06 | 1,59E+06 | 31,01 | 0 | 25,662 | 23 |
| 5,39E+07 | 4,19E+07 | 4,71E+07 | 3,94E+07 | 94,393 | 0 | 24,129 | 42 |
| 7,77E+06 | 2,00E+07 | 2,16E+07 | 1,89E+07 | 58,507 | 0 | 43,346 | 20 |
| 1,65E+07 | 9,20E+06 | 9,08E+06 | 8,56E+06 | 93,433 | 0 | 11,805 | 9 |
| 1,61E+08 | 2,24E+08 | 2,88E+08 | 2,77E+08 | 44,41 | 0 | 154,84 | 59 |
| 2,34E+08 | 3,49E+08 | 3,86E+08 | 4,11E+08 | 21,627 | 0 | 34,817 | 118 |
| 2,30E+07 | 4,10E+06 | 2,69E+06 | 4,54E+06 | 37,379 | 0,000156 | 4,6184 | 5 |
| 5,69E+07 | 8,58E+06 | 1,61E+06 | 7,07E+06 | 211,49 | 0 | 52,051 | 53 |
| 8,31E+07 | 1,91E+06 | 1,38E+07 | 1,08E+07 | 72,108 | 0,000149 | 3,534 | 13 |
| 9,95E+08 | 1,25E+09 | 1,35E+09 | 1,15E+09 | 22,086 | 0 | 323,31 | 324 |
| 2,99E+07 | 4,18E+07 | 4,62E+07 | 4,06E+07 | 29,371 | 0 | 126,85 | 25 |
| 9,44E+07 | 7,27E+07 | 8,44E+07 | 7,82E+07 | 91,785 | 0 | 40,786 | 67 |
| 2,28E+08 | 1,98E+08 | 1,90E+08 | 1,77E+08 | 142,57 | 0 | 251,26 | 169 |
| 1,29E+08 | 6,30E+07 | 7,03E+07 | 5,81E+07 | 24,752 | 0 | 91,978 | 68 |
| 2,78E+07 | 2,00E+07 | 2,29E+07 | 1,73E+07 | 9,7608 | 0 | 6,4648 | 6 |
| 1,55E+09 | 4,55E+09 | 4,71E+09 | 4,77E+09 | 38,418 | 0 | 323,31 | 141 |
| 1,63E+08 | 1,31E+08 | 1,40E+08 | 1,39E+08 | 38,458 | 0 | 323,31 | 50 |
| 1,30E+08 | 1,82E+08 | 1,74E+08 | 1,73E+08 | 157,25 | 0 | 94,929 | 120 |
| 1,41E+07 | 6,05E+06 | 1,61E+06 | 2,76E+06 | 23,461 | 0 | 7,079 | 12 |
| 3,93E+07 | 3,16E+07 | 2,63E+07 | 2,71E+07 | 68,464 | 0 | 58,043 | 38 |
| 2,16E+07 | 1,34E+06 | 3,80E+06 | 1,38E+06 | 106,17 | 0 | 9,6038 | 20 |
| 3,53E+08 | 2,67E+08 | 2,96E+08 | 3,05E+08 | 89,518 | 0 | 210,45 | 127 |
| 1,09E+09 | 1,27E+09 | 1,20E+09 | 1,20E+09 | 100,96 | 0 | 323,31 | 309 |
| 1,01E+09 | 1,48E+09 | 1,44E+09 | 1,44E+09 | 24,307 | 0 | 274,39 | 160 |
| 4,07E+07 | 6,73E+07 | 6,55E+07 | 6,74E+07 | 344,69 | 0 | 42,592 | 90 |
| 4,06E+08 | 3,02E+08 | 3,37E+08 | 3,39E+08 | 102,8 | 0 | 112,23 | 163 |
| 6,88E+07 | 5,01E+07 | 4,03E+07 | 3,70E+07 | 59,085 | 0 | 19,202 | 52 |
| 4,54E+07 | 3,16E+07 | 2,89E+07 | 3,11E+07 | 56,146 | 0 | 9,6473 | 29 |
| 1,08E+08 | 3,58E+06 | 1,48E+07 | 1,68E+07 | 135,76 | 0 | 19,889 | 25 |
| 2,49E+07 | 3,48E+07 | 4,54E+07 | 4,03E+07 | 26,645 | 0 | 23,854 | 23 |
| 2,18E+07 | 1,23E+06 | 3,45E+06 | 5,33E+06 | 8,3585 | 0 | 27,741 | 9 |
| 1,39E+08 | 1,04E+08 | 1,17E+08 | 1,09E+08 | 30,55 | 0 | 42,313 | 53 |
| 3,49E+08 | 6,04E+08 | 5,27E+08 | 5,40E+08 | 56,888 | 0 | 323,31 | 100 |
| 2,48E+07 | 1,24E+07 | 1,03E+07 | 1,63E+07 | 11,475 | 0 | 204,41 | 7 |
| 2,15E+08 | 3,05E+08 | 2,92E+08 | 3,12E+08 | 128,81 | 0 | 197,23 | 165 |
| 3,47E+07 | 4,37E+07 | 5,00E+07 | 4,38E+07 | 97,346 | 0 | 20,256 | 44 |
| 7,94E+06 | 1,12E+08 | 1,03E+08 | 1,01E+08 | 319,59 | 0 | 50,571 | 83 |
| 4,58E+08 | 7,07E+08 | 6,59E+08 | 7,57E+08 | 29,506 | 0 | 323,31 | 109 |
| 2,84E+06 | 1,24E+07 | 1,08E+07 | 1,23E+07 | 20,978 | 0,000149 | 3,456 | 13 |
| 2,79E+07 | 6,37E+06 | 4,76E+06 | 1,92E+06 | 45,801 | 0 | 10,044 | 16 |
| 6,70E+07 | 5,32E+07 | 5,48E+07 | 5,92E+07 | 59,064 | 0 | 45,631 | 57 |
| 7,06E+07 | 1,01E+08 | 1,07E+08 | 1,11E+08 | 47,238 | 0 | 33,241 | 53 |
| 9,34E+08 | 1,09E+09 | 1,01E+09 | 1,03E+09 | 73,963 | 0 | 323,31 | 246 |

| | | | | | | | |
|---|---|---|---|---|---|---|---|
| 1,92E+07 | 1,10E+07 | 1,21E+07 | 7,14E+06 | 52,175 | 0 | 14,331 | 19 |
| 4,60E+07 | 1,57E+08 | 1,25E+08 | 1,79E+08 | 25,134 | 0 | 105,13 | 65 |
| 4,08E+07 | 8,37E+06 | 1,37E+07 | 9,02E+06 | 104,68 | 0,009746 | 1,4076 | 7 |
| 4,47E+06 | 2,95E+06 | 1,58E+06 | 1,94E+06 | 81,7 | 0,000436 | 3,0072 | 6 |
| 3,01E+08 | 2,87E+08 | 2,72E+08 | 2,83E+08 | 50,41 | 0 | 69,94 | 115 |
| 6,63E+07 | 1,10E+08 | 9,80E+07 | 1,09E+08 | 65,893 | 0 | 47,721 | 76 |
| 5,07E+08 | 4,33E+08 | 4,02E+08 | 3,98E+08 | 70,573 | 0 | 323,31 | 194 |
| 7,73E+07 | 1,10E+08 | 1,16E+08 | 1,20E+08 | 50,898 | 0 | 44,071 | 69 |
| 4,97E+06 | 1,88E+07 | 2,46E+07 | 1,48E+07 | 23,42 | 0 | 8,0582 | 14 |
| 3,13E+07 | 4,50E+07 | 4,38E+07 | 4,16E+07 | 204,08 | 0 | 118,3 | 44 |
| 3,55E+06 | 3,90E+07 | 3,91E+07 | 4,91E+07 | 49,462 | 0 | 22,432 | 22 |
| 2,43E+09 | 2,19E+09 | 2,22E+09 | 2,14E+09 | 57,927 | 0 | 323,31 | 268 |
| 1,92E+08 | 1,35E+08 | 1,39E+08 | 1,44E+08 | 28,038 | 0 | 85,595 | 36 |
| 7,73E+06 | 2,11E+07 | 2,84E+07 | 3,45E+07 | 57,873 | 0,001274 | 2,4503 | 6 |
| 2,91E+08 | 2,35E+08 | 2,37E+08 | 2,59E+08 | 36,731 | 0 | 179,87 | 91 |
| 3,63E+07 | 4,17E+07 | 4,39E+07 | 4,11E+07 | 51,637 | 0 | 30,56 | 39 |
| 3,01E+06 | 1,13E+07 | 8,67E+06 | 2,26E+07 | 17,153 | 0,005027 | 1,6855 | 6 |
| 3,83E+07 | 4,83E+07 | 4,69E+07 | 4,49E+07 | 76,756 | 0 | 175,56 | 43 |
| 8,55E+07 | 1,09E+08 | 9,90E+07 | 1,05E+08 | 32,314 | 0 | 323,31 | 53 |
| 4,38E+08 | 2,86E+08 | 2,49E+08 | 2,50E+08 | 34,005 | 0 | 116,88 | 110 |
| 9,70E+06 | 2,78E+07 | 2,83E+07 | 3,79E+07 | 9,7463 | 0 | 6,1991 | 7 |
| 2,70E+08 | 2,18E+08 | 1,84E+08 | 1,95E+08 | 63,301 | 0 | 166,47 | 114 |
| 5,45E+07 | 2,89E+07 | 2,99E+07 | 3,76E+07 | 18,236 | 0 | 18,652 | 35 |
| 4,36E+07 | 8,52E+07 | 7,05E+07 | 7,32E+07 | 48,819 | 0 | 32,131 | 41 |
| 2,48E+09 | 2,14E+09 | 2,29E+09 | 2,31E+09 | 24,146 | 0 | 111,83 | 270 |
| 8,33E+06 | 3,19E+07 | 4,37E+07 | 4,84E+07 | 96,002 | 0 | 25,425 | 31 |
| 1,75E+07 | 3,25E+07 | 3,57E+07 | 3,77E+07 | 19,727 | 0,000158 | 5,1254 | 9 |
| 1,08E+09 | 8,30E+08 | 8,67E+08 | 8,50E+08 | 38,937 | 0 | 191,25 | 232 |
| 4,71E+08 | 3,94E+08 | 4,04E+08 | 4,21E+08 | 26,201 | 0 | 177,79 | 95 |
| 2,06E+07 | 2,43E+07 | 2,61E+07 | 2,50E+07 | 10,139 | 0,000433 | 2,8731 | 8 |
| 1,10E+08 | 8,37E+07 | 9,60E+07 | 9,25E+07 | 29,199 | 0 | 21,503 | 49 |
| 5,38E+06 | 3,48E+06 | 2,71E+06 | 3,69E+06 | 40,789 | 0 | 11,855 | 14 |
| 2,00E+09 | 1,86E+09 | 1,86E+09 | 1,87E+09 | 88,544 | 0 | 323,31 | 367 |
| 1,39E+10 | 1,48E+10 | 1,61E+10 | 1,60E+10 | 18,559 | 0 | 323,31 | 643 |
| 2,89E+07 | 9,11E+06 | 2,24E+06 | 6,43E+06 | 20,059 | 0 | 16,915 | 19 |
| 2,16E+07 | 1,69E+07 | 1,64E+07 | 1,33E+07 | 17,307 | 0 | 18,728 | 13 |
| 2,97E+08 | 1,81E+08 | 1,57E+08 | 1,62E+08 | 97,055 | 0 | 323,31 | 108 |
| 6,75E+07 | 8,41E+07 | 8,44E+07 | 7,90E+07 | 34,989 | 0 | 31,727 | 23 |
| 7,74E+07 | 5,88E+07 | 5,62E+07 | 5,66E+07 | 26,621 | 0 | 11,885 | 29 |
| 6,82E+08 | 1,03E+09 | 9,97E+08 | 9,47E+08 | 52,572 | 0 | 323,31 | 203 |
| 1,33E+08 | 8,75E+07 | 1,02E+08 | 9,47E+07 | 53,184 | 0 | 69,988 | 56 |
| 2,76E+07 | 1,60E+07 | 9,28E+06 | 8,89E+06 | 63,119 | 0 | 12,578 | 22 |
| 1,30E+09 | 1,64E+09 | 1,55E+09 | 1,69E+09 | 81,877 | 0 | 323,31 | 282 |
| 1,15E+07 | 6,67E+06 | 6,31E+06 | 8,98E+06 | 50,632 | 0 | 5,6583 | 8 |
| 1,22E+07 | 3,96E+07 | 4,10E+07 | 3,48E+07 | 72,577 | 0 | 41,659 | 25 |
| 1,55E+07 | 2,34E+07 | 1,96E+07 | 1,87E+07 | 79,673 | 0 | 18,735 | 18 |
| 3,40E+06 | 1,18E+07 | 6,30E+06 | 8,17E+06 | 57,791 | 0,003829 | 1,8036 | 7 |
| 7,08E+07 | 1,33E+08 | 1,46E+08 | 1,36E+08 | 33,942 | 0 | 14,805 | 41 |
| 1,04E+09 | 1,13E+09 | 1,22E+09 | 1,16E+09 | 143,23 | 0 | 323,31 | 431 |
| 5,80E+07 | 4,46E+07 | 4,96E+07 | 4,84E+07 | 77,187 | 0 | 106,73 | 53 |

| | | | | | | | |
|---|---|---|---|---|---|---|---|
| 7,51E+06 | 1,21E+07 | 1,75E+07 | 1,63E+07 | 29,657 | 0 | 10,501 | 18 |
| 5,43E+07 | 1,01E+08 | 8,27E+07 | 7,96E+07 | 27,483 | 0 | 16,855 | 36 |
| 5,72E+09 | 7,16E+09 | 8,34E+09 | 9,29E+09 | 14,957 | 0 | 323,31 | 590 |
| 3,75E+06 | 5,64E+06 | 5,43E+06 | 4,39E+06 | 28,239 | 0,000152 | 4,0176 | 11 |
| 7,32E+07 | 9,65E+07 | 9,64E+07 | 9,55E+07 | 60,263 | 0 | 31,592 | 57 |
| 1,62E+08 | 1,02E+08 | 9,47E+07 | 8,15E+07 | 27,581 | 0 | 106,44 | 39 |
| 5,31E+07 | 8,95E+07 | 9,04E+07 | 9,58E+07 | 21,539 | 0 | 8,9894 | 40 |
| 3,62E+07 | 2,20E+07 | 2,44E+07 | 2,72E+07 | 78,631 | 0 | 35,846 | 35 |
| 3,16E+07 | 2,45E+07 | 2,03E+07 | 2,56E+07 | 81,332 | 0 | 19,292 | 36 |
| 6,10E+09 | 4,94E+09 | 5,11E+09 | 5,04E+09 | 140,21 | 0 | 323,31 | 750 |
| 7,08E+06 | 3,69E+06 | 2,76E+06 | 2,29E+06 | 35,742 | 0,003175 | 1,9505 | 6 |
| 3,74E+08 | 3,18E+08 | 3,03E+08 | 3,29E+08 | 25,826 | 0 | 126,91 | 107 |
| 7,74E+08 | 1,03E+09 | 9,52E+08 | 9,92E+08 | 42,516 | 0 | 323,31 | 194 |
| 3,42E+08 | 2,65E+08 | 2,51E+08 | 2,58E+08 | 99,307 | 0 | 323,31 | 174 |
| 1,26E+07 | 2,20E+07 | 2,16E+07 | 2,18E+07 | 51,532 | 0 | 16,105 | 8 |
| 1,67E+08 | 1,34E+08 | 1,36E+08 | 1,50E+08 | 31,246 | 0 | 101,44 | 73 |
| 5,05E+07 | 3,85E+07 | 3,99E+07 | 3,84E+07 | 125,38 | 0 | 32,74 | 42 |
| 3,11E+08 | 2,12E+08 | 2,54E+08 | 2,59E+08 | 55,503 | 0 | 214,09 | 50 |
| 8,35E+08 | 1,09E+09 | 1,07E+09 | 1,16E+09 | 30,436 | 0 | 139,69 | 138 |
| 2,34E+08 | 1,93E+08 | 1,91E+08 | 2,00E+08 | 110,2 | 0 | 177,23 | 130 |
| 2,78E+08 | 3,60E+08 | 3,64E+08 | 3,75E+08 | 134,03 | 0 | 241,6 | 151 |
| 1,63E+07 | 9,99E+06 | 1,19E+07 | 9,85E+06 | 44,325 | 0 | 5,7673 | 16 |
| 2,32E+08 | 1,95E+08 | 1,92E+08 | 2,03E+08 | 95,306 | 0 | 68,113 | 131 |
| 6,37E+08 | 4,61E+08 | 4,32E+08 | 4,12E+08 | 24,942 | 0 | 117,53 | 81 |
| 4,27E+09 | 5,14E+09 | 5,22E+09 | 5,08E+09 | 104,98 | 0 | 323,31 | 494 |
| 5,87E+08 | 9,53E+08 | 9,49E+08 | 9,76E+08 | 131,63 | 0 | 247,68 | 281 |
| 1,19E+08 | 1,34E+08 | 1,52E+08 | 1,48E+08 | 53,451 | 0 | 66,396 | 76 |
| 6,22E+06 | 5,26E+07 | 5,47E+07 | 6,07E+07 | 93,769 | 0 | 26,516 | 31 |
| 3,94E+07 | 6,01E+07 | 5,37E+07 | 5,85E+07 | 77,229 | 0 | 35,417 | 46 |
| 3,33E+07 | 2,21E+07 | 2,26E+07 | 2,61E+07 | 19,147 | 0,000147 | 3,2963 | 7 |
| 3,67E+08 | 2,99E+08 | 3,23E+08 | 3,21E+08 | 76,429 | 0 | 71,349 | 136 |
| 1,71E+08 | 4,00E+06 | 5,01E+06 | 4,89E+06 | 37,367 | 0 | 78,684 | 21 |
| 1,53E+07 | 2,59E+07 | 2,38E+07 | 2,32E+07 | 37,935 | 0 | 8,2424 | 10 |
| 1,18E+07 | 2,29E+07 | 2,29E+07 | 2,52E+07 | 76,956 | 0,000153 | 4,1648 | 15 |
| 1,99E+08 | 1,65E+08 | 1,79E+08 | 1,68E+08 | 88,324 | 0 | 41,285 | 63 |
| 7,60E+07 | 5,56E+07 | 5,60E+07 | 6,00E+07 | 82,687 | 0 | 21,969 | 49 |
| 8,48E+07 | 1,31E+08 | 1,37E+08 | 1,08E+08 | 38,225 | 0 | 42,122 | 65 |
| 2,04E+08 | 2,76E+08 | 2,63E+08 | 2,99E+08 | 54,218 | 0 | 233,45 | 124 |
| 2,09E+07 | 1,54E+06 | 2,04E+06 | 3,58E+06 | 55,968 | 0 | 52,003 | 29 |
| 3,97E+08 | 2,35E+08 | 3,20E+08 | 2,96E+08 | 12,497 | 0 | 12,602 | 43 |
| 1,79E+08 | 1,15E+08 | 1,51E+08 | 1,24E+08 | 105,04 | 0 | 46,786 | 49 |
| 1,56E+08 | 1,68E+08 | 1,69E+08 | 1,67E+08 | 195,99 | 0 | 87,159 | 144 |
| 3,00E+09 | 4,15E+09 | 4,28E+09 | 4,86E+09 | 42,435 | 0 | 323,31 | 352 |
| 2,66E+08 | 4,22E+08 | 4,28E+08 | 4,35E+08 | 83,967 | 0 | 117,11 | 151 |
| 1,32E+08 | 2,23E+08 | 1,85E+08 | 1,83E+08 | 22,166 | 0 | 118,23 | 73 |
| 2,18E+07 | 7,83E+06 | 2,09E+06 | 2,81E+06 | 204,1 | 0 | 9,6987 | 11 |
| 2,27E+08 | 1,83E+08 | 2,03E+08 | 2,05E+08 | 118,78 | 0 | 144,77 | 110 |
| 2,58E+08 | 2,03E+08 | 2,28E+08 | 2,08E+08 | 68,249 | 0 | 241,96 | 136 |
| 2,43E+07 | 3,21E+07 | 3,26E+07 | 2,95E+07 | 48,9 | 0 | 17,97 | 33 |
| 8,59E+08 | 7,93E+08 | 7,73E+08 | 7,51E+08 | 29,116 | 0 | 278,86 | 127 |

| | | | | | | | |
|---|---|---|---|---|---|---|---|
| 1,05E+08 | 1,50E+08 | 1,61E+08 | 1,62E+08 | 152,6 | 0 | 55,631 | 109 |
| 1,09E+08 | 1,59E+08 | 1,59E+08 | 1,57E+08 | 88,927 | 0 | 117,22 | 114 |
| 1,24E+07 | 2,28E+07 | 2,07E+07 | 2,59E+07 | 99,611 | 0 | 6,8248 | 25 |
| 1,27E+10 | 9,37E+09 | 1,11E+10 | 1,04E+10 | 37,402 | 0 | 323,31 | 1079 |
| 2,92E+08 | 3,73E+08 | 3,41E+08 | 3,26E+08 | 146,22 | 0 | 274,53 | 173 |
| 1,85E+08 | 1,23E+08 | 1,41E+08 | 1,45E+08 | 30,169 | 0 | 20,314 | 49 |
| 5,57E+08 | 6,93E+08 | 7,06E+08 | 7,19E+08 | 23,01 | 0 | 323,31 | 77 |
| 1,64E+08 | 2,08E+08 | 2,19E+08 | 2,38E+08 | 86,239 | 0 | 323,31 | 97 |
| 7,49E+07 | 8,72E+07 | 1,03E+08 | 9,49E+07 | 72,859 | 0 | 98,195 | 48 |
| 1,45E+09 | 1,36E+09 | 1,25E+09 | 1,31E+09 | 92,188 | 0 | 323,31 | 323 |
| 7,82E+05 | 8,65E+07 | 9,17E+07 | 8,01E+07 | 60,612 | 0 | 22,224 | 38 |
| 3,07E+08 | 2,23E+08 | 2,50E+08 | 2,20E+08 | 39,468 | 0 | 63,635 | 51 |
| 2,72E+08 | 2,19E+08 | 2,11E+08 | 2,28E+08 | 59,715 | 0 | 268,26 | 105 |
| 5,95E+07 | 7,53E+07 | 7,19E+07 | 7,15E+07 | 37,374 | 0 | 22,78 | 49 |
| 9,11E+06 | 2,87E+07 | 2,71E+07 | 2,88E+07 | 77,383 | 0 | 10,923 | 21 |
| 3,98E+07 | 3,10E+07 | 3,01E+07 | 2,63E+07 | 48,364 | 0 | 11,9 | 27 |
| 9,37E+07 | 5,48E+07 | 7,01E+07 | 6,75E+07 | 48,568 | 0 | 43,457 | 57 |
| 2,10E+08 | 1,50E+08 | 1,34E+08 | 1,39E+08 | 36,744 | 0 | 132,97 | 86 |
| 1,71E+07 | 2,26E+07 | 3,14E+07 | 2,65E+07 | 26,529 | 0 | 11,208 | 28 |
| 3,17E+07 | 2,43E+07 | 2,41E+07 | 2,02E+07 | 46,795 | 0 | 14,243 | 30 |
| 3,58E+07 | 6,01E+06 | 6,37E+06 | 1,09E+06 | 88,529 | 0 | 17,452 | 20 |
| 1,20E+08 | 8,91E+07 | 9,44E+07 | 7,87E+07 | 25,82 | 0 | 20,823 | 30 |
| 7,84E+08 | 5,32E+08 | 6,18E+08 | 6,12E+08 | 33,665 | 0 | 168,44 | 103 |
| 2,42E+08 | 3,15E+06 | 3,66E+07 | 2,20E+06 | 98,255 | 0 | 310,84 | 74 |
| 8,46E+08 | 7,01E+08 | 7,72E+08 | 7,62E+08 | 41,489 | 0 | 230,37 | 134 |
| 1,29E+07 | 6,02E+06 | 8,34E+06 | 8,58E+06 | 26,407 | 0,000159 | 5,3135 | 14 |
| 2,70E+07 | 8,06E+06 | 8,90E+06 | 4,03E+06 | 248,74 | 0 | 40,758 | 23 |
| 1,94E+08 | 3,60E+08 | 3,65E+08 | 3,59E+08 | 9,1175 | 0 | 115,66 | 49 |
| 2,86E+08 | 2,50E+08 | 2,50E+08 | 2,50E+08 | 27,285 | 0 | 118,11 | 96 |
| 2,66E+08 | 2,36E+08 | 2,12E+08 | 2,08E+08 | 61,211 | 0 | 49,159 | 124 |
| 1,08E+08 | 1,44E+08 | 1,36E+08 | 1,20E+08 | 14,164 | 0 | 9,6861 | 40 |
| 1,62E+07 | 2,53E+07 | 2,09E+07 | 2,00E+07 | 24,863 | 0 | 7,1563 | 18 |
| 1,30E+08 | 2,16E+08 | 1,97E+08 | 2,03E+08 | 73,12 | 0 | 127,84 | 78 |
| 8,90E+07 | 3,64E+07 | 4,18E+07 | 4,42E+07 | 46,288 | 0 | 64,699 | 38 |
| 1,68E+07 | 7,15E+06 | 2,62E+06 | 2,05E+06 | 33,431 | 0,000155 | 4,495 | 10 |
| 1,02E+08 | 8,51E+07 | 9,19E+07 | 9,42E+07 | 29,725 | 0 | 19,791 | 43 |
| 4,46E+08 | 3,68E+08 | 3,40E+08 | 3,68E+08 | 74,775 | 0 | 83,72 | 192 |
| 2,75E+07 | 4,61E+07 | 4,21E+07 | 4,30E+07 | 87,02 | 0 | 62,475 | 45 |
| 2,86E+07 | 4,63E+07 | 6,45E+07 | 4,94E+07 | 51,702 | 0 | 16,11 | 35 |
| 1,68E+08 | 1,98E+08 | 2,18E+08 | 2,18E+08 | 25,193 | 0 | 160,31 | 67 |
| 2,66E+09 | 3,58E+09 | 3,26E+09 | 3,55E+09 | 43,698 | 0 | 323,31 | 517 |
| 5,30E+07 | 8,53E+07 | 7,89E+07 | 7,43E+07 | 126,18 | 0 | 112,52 | 79 |
| 7,61E+07 | 6,17E+07 | 5,16E+07 | 6,15E+07 | 79,546 | 0 | 101,49 | 53 |
| 3,64E+08 | 3,02E+08 | 2,49E+08 | 2,64E+08 | 116,41 | 0 | 323,31 | 221 |
| 6,35E+08 | 5,66E+08 | 5,74E+08 | 5,78E+08 | 77,795 | 0 | 306,56 | 206 |
| 1,08E+07 | 5,27E+06 | 3,98E+06 | 1,87E+06 | 43,837 | 0 | 30,037 | 13 |
| 5,30E+07 | 3,13E+07 | 2,53E+07 | 1,49E+07 | 67,464 | 0 | 41,53 | 31 |
| 1,85E+08 | 1,39E+08 | 1,56E+08 | 1,57E+08 | 49,714 | 0 | 45,674 | 116 |
| 7,80E+08 | 1,25E+09 | 1,23E+09 | 1,33E+09 | 82,613 | 0 | 296,48 | 228 |
| 1,36E+08 | 1,58E+08 | 1,47E+08 | 1,48E+08 | 20,677 | 0 | 53,456 | 70 |

| | | | | | | | |
|---|---|---|---|---|---|---|---|
| 2,79E+08 | 1,99E+08 | 2,08E+08 | 2,44E+08 | 52,01 | 0 | 141,84 | 59 |
| 2,15E+07 | 3,02E+07 | 2,96E+07 | 2,93E+07 | 23,522 | 0 | 12,432 | 26 |
| 2,95E+08 | 3,19E+08 | 3,20E+08 | 3,47E+08 | 132,76 | 0 | 258,18 | 148 |
| 2,62E+07 | 1,73E+07 | 2,24E+07 | 1,80E+07 | 50,201 | 0,000149 | 3,5857 | 16 |
| 6,32E+07 | 1,11E+08 | 1,05E+08 | 1,22E+08 | 92,612 | 0 | 64,815 | 71 |
| 9,30E+07 | 1,09E+08 | 1,12E+08 | 1,31E+08 | 172,35 | 0 | 48,351 | 114 |
| 3,92E+07 | 5,22E+07 | 6,45E+07 | 6,37E+07 | 34,278 | 0 | 24,088 | 36 |
| 4,67E+06 | 3,12E+06 | 2,89E+06 | 2,70E+06 | 39,453 | 0,001548 | 2,3034 | 6 |
| 6,92E+07 | 4,43E+07 | 3,87E+07 | 5,16E+07 | 12,111 | 0 | 40,144 | 38 |
| 1,54E+08 | 1,92E+08 | 1,71E+08 | 1,77E+08 | 17,319 | 0 | 203,25 | 55 |
| 4,27E+08 | 3,33E+08 | 3,01E+08 | 3,12E+08 | 43,528 | 0 | 323,31 | 100 |
| 4,48E+08 | 2,75E+08 | 3,11E+08 | 2,79E+08 | 64,979 | 0 | 279,23 | 201 |
| 6,18E+08 | 4,37E+08 | 4,42E+08 | 4,58E+08 | 229,2 | 0 | 297,62 | 270 |
| 2,39E+09 | 1,99E+09 | 1,87E+09 | 2,02E+09 | 30,53 | 0 | 323,31 | 236 |
| 5,23E+08 | 8,09E+08 | 7,35E+08 | 7,75E+08 | 25,206 | 0 | 323,31 | 113 |
| 1,37E+08 | 9,56E+07 | 1,14E+08 | 1,17E+08 | 11,97 | 0 | 25,446 | 45 |
| 2,75E+08 | 2,47E+08 | 2,12E+08 | 2,09E+08 | 73,088 | 0 | 206,47 | 108 |
| 7,51E+08 | 1,26E+09 | 1,09E+09 | 9,27E+08 | 37,778 | 0 | 323,31 | 237 |
| 1,29E+08 | 8,51E+07 | 1,06E+08 | 1,06E+08 | 31,375 | 0 | 88,887 | 56 |
| 6,14E+08 | 5,25E+08 | 5,39E+08 | 5,06E+08 | 170,29 | 0 | 323,31 | 301 |
| 1,73E+07 | 3,20E+06 | 2,51E+06 | 8,01E+06 | 33,243 | 0 | 5,6184 | 10 |
| 6,68E+08 | 5,61E+08 | 5,94E+08 | 5,35E+08 | 38,384 | 0 | 287,31 | 115 |
| 3,63E+07 | 5,21E+07 | 6,62E+07 | 7,88E+07 | 39,376 | 0 | 31,699 | 27 |
| 5,31E+07 | 4,09E+07 | 4,38E+07 | 4,36E+07 | 25,766 | 0 | 9,7938 | 33 |
| 6,55E+07 | 4,10E+07 | 5,14E+07 | 4,49E+07 | 84,778 | 0 | 91,772 | 47 |
| 3,12E+07 | 2,09E+07 | 1,71E+07 | 1,34E+07 | 39,105 | 0 | 9,3591 | 19 |
| 7,91E+06 | 3,44E+07 | 3,48E+07 | 4,15E+07 | 55,142 | 0 | 15,093 | 14 |
| 2,29E+07 | 5,99E+06 | 4,83E+06 | 1,62E+06 | 32,951 | 0 | 7,3471 | 11 |
| 6,80E+07 | 9,81E+07 | 8,41E+07 | 7,96E+07 | 69,885 | 0 | 30,727 | 54 |
| 6,05E+09 | 4,81E+09 | 4,78E+09 | 4,86E+09 | 69,632 | 0 | 323,31 | 522 |
| 1,13E+09 | 1,34E+09 | 1,26E+09 | 1,32E+09 | 11,922 | 0 | 204,15 | 203 |
| 4,36E+07 | 2,58E+07 | 3,01E+07 | 3,29E+07 | 105,35 | 0 | 18,008 | 17 |
| 6,39E+07 | 5,55E+07 | 4,81E+07 | 5,30E+07 | 104,41 | 0 | 37,947 | 72 |
| 6,11E+07 | 1,05E+08 | 9,99E+07 | 1,01E+08 | 265,63 | 0 | 121,71 | 85 |
| 4,63E+08 | 3,02E+08 | 3,07E+08 | 2,86E+08 | 11,424 | 0 | 15,675 | 61 |
| 1,42E+07 | 1,06E+07 | 7,79E+06 | 1,02E+07 | 62,208 | 0 | 30,85 | 20 |
| 2,86E+06 | 4,59E+07 | 3,72E+07 | 4,99E+07 | 43,349 | 0 | 44,19 | 25 |
| 2,94E+08 | 2,12E+08 | 2,11E+08 | 2,22E+08 | 34,322 | 0 | 59,15 | 68 |
| 7,46E+07 | 6,38E+07 | 6,72E+07 | 7,00E+07 | 60,878 | 0 | 323,31 | 33 |
| 3,72E+09 | 4,50E+09 | 4,37E+09 | 4,18E+09 | 64,217 | 0 | 323,31 | 527 |
| 2,52E+07 | 1,87E+07 | 1,74E+07 | 1,59E+07 | 85,98 | 0 | 24,25 | 17 |
| 1,58E+07 | 5,27E+06 | 3,93E+06 | 9,15E+06 | 289,07 | 0,001823 | 2,2363 | 10 |
| 1,10E+09 | 9,06E+08 | 8,80E+08 | 9,77E+08 | 206,5 | 0 | 323,31 | 393 |
| 1,56E+07 | 1,18E+07 | 9,35E+06 | 1,19E+07 | 25,029 | 0 | 5,8232 | 15 |
| 9,96E+08 | 1,24E+09 | 1,24E+09 | 1,36E+09 | 38,117 | 0 | 323,31 | 129 |
| 1,03E+09 | 8,74E+08 | 8,60E+08 | 8,45E+08 | 317,74 | 0 | 323,31 | 507 |
| 1,96E+08 | 1,63E+08 | 1,38E+08 | 1,54E+08 | 43,88 | 0 | 142,84 | 82 |
| 8,14E+07 | 1,67E+08 | 1,80E+08 | 1,84E+08 | 33,569 | 0 | 68,023 | 46 |
| 2,93E+07 | 4,24E+07 | 4,17E+07 | 4,65E+07 | 55,064 | 0 | 13,693 | 15 |
| 3,94E+08 | 3,41E+08 | 3,17E+08 | 3,04E+08 | 59,481 | 0 | 318,12 | 162 |

| | | | | | | | |
|---|---|---|---|---|---|---|---|
| 3,93E+07 | 5,63E+07 | 5,33E+07 | 5,40E+07 | 25,262 | 0,000156 | 4,7355 | 18 |
| 1,05E+07 | 2,31E+07 | 1,42E+07 | 1,97E+07 | 47,152 | 0 | 8,2614 | 15 |
| 1,05E+08 | 1,32E+08 | 1,20E+08 | 1,39E+08 | 41,614 | 0 | 323,31 | 67 |
| 2,11E+08 | 1,73E+08 | 1,42E+08 | 1,36E+08 | 47,404 | 0 | 71,37 | 104 |
| 6,13E+08 | 7,22E+08 | 7,26E+08 | 6,54E+08 | 17,725 | 0 | 107,82 | 157 |
| 5,22E+08 | 6,15E+08 | 8,02E+08 | 7,72E+08 | 26,334 | 0 | 187,6 | 114 |
| 1,75E+07 | 7,51E+07 | 1,06E+08 | 1,21E+08 | 19,3 | 0 | 56,269 | 25 |
| 4,38E+08 | 3,50E+08 | 3,72E+08 | 3,79E+08 | 35,328 | 0 | 192,78 | 132 |
| 2,36E+06 | 5,83E+07 | 4,66E+07 | 4,22E+07 | 29,596 | 0 | 20,773 | 22 |
| 5,04E+07 | 3,97E+07 | 4,26E+07 | 4,00E+07 | 44,415 | 0 | 20,888 | 27 |
| 3,65E+08 | 3,21E+08 | 3,09E+08 | 3,08E+08 | 47,598 | 0 | 22,868 | 79 |
| 3,37E+08 | 4,05E+08 | 4,23E+08 | 4,57E+08 | 25,426 | 0 | 239,92 | 104 |
| 2,29E+08 | 1,80E+08 | 1,97E+08 | 1,95E+08 | 52,515 | 0 | 79,588 | 129 |
| 3,23E+07 | 6,24E+07 | 5,90E+07 | 7,26E+07 | 93,696 | 0 | 43,817 | 39 |
| 4,29E+08 | 5,65E+08 | 5,21E+08 | 5,54E+08 | 36,023 | 0 | 323,31 | 150 |
| 9,87E+07 | 7,92E+07 | 7,62E+07 | 8,72E+07 | 96,21 | 0 | 53,236 | 104 |
| 5,36E+09 | 6,13E+09 | 6,49E+09 | 6,75E+09 | 94,132 | 0 | 323,31 | 572 |
| 2,87E+08 | 3,24E+08 | 3,52E+08 | 3,60E+08 | 73,98 | 0 | 178,76 | 110 |
| 6,24E+09 | 5,14E+09 | 5,24E+09 | 4,97E+09 | 273,61 | 0 | 323,31 | 1114 |
| 6,62E+06 | 3,74E+06 | 5,22E+06 | 4,88E+06 | 26,864 | 0,000292 | 3,077 | 9 |
| 5,94E+07 | 4,94E+07 | 5,36E+07 | 4,74E+07 | 28,702 | 0 | 120,65 | 39 |
| 2,41E+08 | 3,27E+08 | 2,91E+08 | 3,23E+08 | 51,344 | 0 | 226,4 | 104 |
| 2,76E+06 | 1,59E+07 | 1,31E+07 | 8,92E+06 | 23,73 | 0 | 5,3673 | 6 |
| 2,15E+07 | 1,06E+07 | 9,70E+06 | 1,54E+07 | 56,178 | 0 | 6,2853 | 16 |
| 3,57E+08 | 3,04E+08 | 3,10E+08 | 2,82E+08 | 115,36 | 0 | 221,47 | 183 |
| 8,18E+06 | 2,30E+06 | 9,08E+05 | 3,46E+06 | 28,644 | 0 | 8,5185 | 7 |
| 1,30E+07 | 4,13E+06 | 8,29E+06 | 5,11E+06 | 50,943 | 0 | 6,2669 | 14 |
| 1,92E+07 | 1,13E+07 | 1,43E+07 | 1,51E+07 | 67,67 | 0 | 17,853 | 26 |
| 2,87E+07 | 2,54E+07 | 2,22E+07 | 2,56E+07 | 86,075 | 0 | 20,807 | 39 |
| 1,85E+08 | 6,50E+08 | 5,30E+08 | 9,68E+08 | 99,136 | 0 | 59,021 | 44 |
| 1,32E+06 | 6,26E+06 | 5,88E+06 | 5,71E+06 | 46,017 | 0,003702 | 1,8479 | 5 |
| 1,18E+09 | 7,42E+08 | 7,45E+08 | 6,64E+08 | 63,845 | 0 | 323,31 | 234 |
| 4,46E+07 | 3,64E+07 | 4,08E+07 | 3,79E+07 | 35,827 | 0 | 20,805 | 29 |
| 8,90E+07 | 7,30E+07 | 7,26E+07 | 5,86E+07 | 98,992 | 0 | 87,922 | 67 |
| 4,55E+07 | 4,06E+07 | 3,60E+07 | 3,41E+07 | 33,04 | 0 | 40,991 | 25 |
| 3,37E+07 | 4,47E+07 | 5,10E+07 | 5,40E+07 | 151,11 | 0 | 37,604 | 51 |
| 6,65E+06 | 1,50E+06 | 2,09E+06 | 1,91E+06 | 104,9 | 0,003831 | 1,8104 | 2 |
| 2,55E+09 | 2,76E+09 | 2,79E+09 | 2,63E+09 | 96,735 | 0 | 323,31 | 454 |
| 2,83E+09 | 3,63E+09 | 3,41E+09 | 3,49E+09 | 75,673 | 0 | 323,31 | 475 |
| 1,15E+08 | 1,76E+06 | 3,09E+06 | 2,45E+07 | 30,448 | 0 | 18,112 | 18 |
| 7,93E+07 | 5,04E+07 | 6,04E+07 | 6,50E+07 | 80,687 | 0 | 26,719 | 77 |
| 8,66E+06 | 2,34E+07 | 1,43E+07 | 1,51E+07 | 121,87 | 0,000157 | 4,8192 | 11 |
| 3,54E+08 | 4,77E+08 | 4,18E+08 | 4,04E+08 | 47,744 | 0 | 232,33 | 65 |
| 4,60E+06 | 6,84E+06 | 6,72E+06 | 6,78E+06 | 30,014 | 0,000851 | 2,5148 | 7 |
| 8,58E+06 | 4,52E+06 | 4,87E+06 | 6,02E+06 | 42,145 | 0 | 6,0573 | 9 |
| 1,21E+08 | 1,03E+08 | 8,84E+07 | 8,18E+07 | 94,602 | 0 | 162,7 | 68 |
| 5,30E+09 | 1,07E+10 | 1,00E+10 | 1,11E+10 | 24,409 | 0 | 323,31 | 187 |
| 8,27E+08 | 1,24E+09 | 9,81E+08 | 1,07E+09 | 10,102 | 0 | 178,96 | 116 |
| 1,55E+08 | 1,47E+08 | 1,33E+08 | 1,37E+08 | 86,186 | 0 | 58,38 | 83 |
| 4,05E+06 | 7,06E+06 | 7,59E+06 | 5,99E+06 | 81,758 | 0,000157 | 4,9549 | 9 |

| | | | | | | | |
|---|---|---|---|---|---|---|---|
| 5,83E+08 | 6,59E+08 | 7,41E+08 | 7,05E+08 | 18,709 | 0 | 202,92 | 76 |
| 1,00E+07 | 2,71E+06 | 4,22E+06 | 4,47E+06 | 26,995 | 0 | 11,045 | 6 |
| 1,84E+08 | 2,23E+08 | 2,31E+08 | 2,04E+08 | 47,874 | 0 | 107,41 | 86 |
| 3,03E+08 | 2,21E+08 | 2,19E+08 | 2,20E+08 | 39,774 | 0 | 138,3 | 126 |
| 2,19E+07 | 1,75E+07 | 1,95E+07 | 1,90E+07 | 57,971 | 0 | 8,7775 | 26 |
| 9,89E+06 | 1,92E+06 | 1,17E+06 | 4,25E+06 | 52,28 | 0,00846 | 1,4816 | 4 |
| 1,67E+07 | 1,10E+07 | 1,22E+07 | 1,42E+07 | 17,067 | 0 | 6,8187 | 23 |
| 2,25E+08 | 1,31E+08 | 1,34E+08 | 1,06E+08 | 52,938 | 0 | 200,17 | 85 |
| 2,70E+08 | 2,26E+08 | 2,19E+08 | 2,15E+08 | 184,29 | 0 | 59,727 | 146 |
| 7,51E+07 | 6,42E+06 | 1,50E+07 | 1,45E+06 | 18,955 | 0 | 43,517 | 24 |
| 2,28E+07 | 1,62E+07 | 1,96E+07 | 1,86E+07 | 51,589 | 0,000156 | 4,714 | 13 |
| 9,46E+06 | 5,36E+06 | 5,78E+06 | 7,84E+06 | 46,202 | 0 | 14,616 | 16 |
| 6,06E+06 | 1,38E+07 | 1,80E+07 | 1,60E+07 | 87,411 | 0,000152 | 3,9449 | 9 |
| 4,06E+08 | 3,04E+08 | 3,45E+08 | 3,58E+08 | 15,661 | 0 | 160,42 | 41 |
| 5,66E+07 | 3,77E+07 | 4,12E+07 | 4,00E+07 | 67,948 | 0 | 19,326 | 32 |
| 1,22E+09 | 1,50E+09 | 1,59E+09 | 1,49E+09 | 117,43 | 0 | 323,31 | 331 |
| 2,33E+07 | 2,73E+07 | 2,66E+07 | 3,11E+07 | 56,076 | 0,000152 | 3,9861 | 9 |
| 1,00E+08 | 1,21E+08 | 1,18E+08 | 1,13E+08 | 64,615 | 0 | 98,368 | 106 |
| 7,96E+08 | 1,00E+09 | 9,68E+08 | 1,07E+09 | 68,839 | 0 | 323,31 | 153 |
| 6,13E+06 | 9,55E+06 | 8,95E+06 | 8,71E+06 | 208,33 | 0 | 8,4518 | 21 |
| 2,80E+08 | 1,94E+08 | 2,07E+08 | 2,45E+08 | 27,541 | 0 | 323,31 | 56 |
| 1,31E+09 | 1,72E+09 | 1,93E+09 | 1,84E+09 | 47,907 | 0 | 323,31 | 204 |
| 6,83E+08 | 7,33E+08 | 7,66E+08 | 7,62E+08 | 141,67 | 0 | 323,31 | 323 |
| 6,46E+07 | 4,32E+07 | 5,45E+07 | 4,96E+07 | 47,307 | 0 | 35,813 | 44 |
| 1,58E+09 | 2,78E+08 | 1,36E+08 | 1,55E+08 | 34,595 | 0 | 17,024 | 32 |
| 6,58E+07 | 5,29E+07 | 4,37E+07 | 5,58E+07 | 89,125 | 0 | 55,103 | 77 |
| 3,12E+07 | 1,93E+06 | 1,25E+07 | 2,61E+06 | 45,234 | 0 | 8,3703 | 17 |
| 7,88E+07 | 6,38E+07 | 5,79E+07 | 4,97E+07 | 121,8 | 0 | 36,324 | 68 |
| 8,12E+06 | 4,73E+06 | 5,01E+06 | 3,47E+06 | 58,999 | 0 | 6,9769 | 9 |
| 8,74E+07 | 9,35E+07 | 9,83E+07 | 9,95E+07 | 91,935 | 0 | 36,584 | 70 |
| 7,05E+06 | 9,49E+06 | 9,05E+06 | 8,11E+06 | 23,13 | 0 | 5,5131 | 12 |
| 2,61E+10 | 1,85E+10 | 1,96E+10 | 1,90E+10 | 95,313 | 0 | 323,31 | 1613 |
| 2,47E+08 | 1,77E+08 | 1,31E+08 | 1,40E+08 | 87,072 | 0 | 77,339 | 73 |
| 1,24E+06 | 1,54E+07 | 1,76E+07 | 1,23E+07 | 67,711 | 0 | 6,058 | 11 |
| 1,96E+07 | 3,84E+06 | 5,26E+06 | 3,12E+06 | 41,397 | 0 | 10,554 | 13 |
| 4,38E+07 | 7,28E+06 | 6,99E+06 | 1,15E+06 | 32,19 | 0 | 12,547 | 14 |
| 3,78E+08 | 2,86E+08 | 3,37E+08 | 2,72E+08 | 92,849 | 0 | 111,25 | 169 |
| 5,84E+07 | 8,02E+07 | 8,04E+07 | 9,02E+07 | 116,66 | 0 | 43,926 | 61 |
| 8,60E+08 | 7,24E+08 | 7,13E+08 | 8,10E+08 | 57,432 | 0 | 288,95 | 155 |
| 2,68E+07 | 3,57E+07 | 4,54E+07 | 4,08E+07 | 35,957 | 0 | 25,163 | 37 |
| 1,11E+09 | 1,00E+09 | 9,77E+08 | 1,02E+09 | 36,954 | 0 | 323,31 | 145 |
| 1,14E+08 | 6,51E+07 | 6,16E+07 | 7,22E+07 | 13,996 | 0 | 19,956 | 39 |
| 8,93E+06 | 1,95E+07 | 2,19E+07 | 1,75E+07 | 45,296 | 0 | 39,849 | 20 |
| 2,20E+08 | 3,02E+08 | 2,63E+08 | 2,74E+08 | 250,93 | 0 | 192,84 | 161 |
| 4,71E+07 | 4,11E+07 | 3,51E+07 | 3,14E+07 | 47,173 | 0 | 22,375 | 44 |
| 4,25E+08 | 5,73E+08 | 5,47E+08 | 5,57E+08 | 48,043 | 0 | 59,026 | 107 |
| 7,23E+06 | 2,13E+06 | 1,42E+06 | 4,24E+06 | 16,667 | 0,000433 | 2,8436 | 5 |
| 1,17E+09 | 1,10E+09 | 1,00E+09 | 9,78E+08 | 32,886 | 0 | 66,768 | 145 |
| 3,93E+08 | 1,53E+08 | 2,11E+08 | 1,77E+08 | 49,924 | 0 | 74,035 | 42 |
| 1,45E+07 | 3,97E+06 | 3,39E+06 | 2,86E+06 | 50,075 | 0 | 7,5216 | 9 |

| | | | | | | | |
|---|---|---|---|---|---|---|---|
| 6,38E+08 | 5,94E+08 | 5,65E+08 | 5,65E+08 | 160,82 | 0 | 323,31 | 242 |
| 9,75E+07 | 1,34E+08 | 1,28E+08 | 1,42E+08 | 77,951 | 0 | 46,809 | 95 |
| 5,77E+07 | 8,12E+07 | 7,69E+07 | 6,93E+07 | 43,256 | 0 | 88,501 | 49 |
| 6,22E+07 | 7,03E+07 | 7,67E+07 | 7,78E+07 | 79,83 | 0 | 21,757 | 50 |
| 1,12E+09 | 9,03E+08 | 1,02E+09 | 8,60E+08 | 39,275 | 0 | 323,31 | 170 |
| 1,27E+07 | 2,88E+06 | 7,06E+06 | 2,42E+06 | 85,547 | 0 | 8,1992 | 15 |
| 1,08E+08 | 1,64E+08 | 1,60E+08 | 1,41E+08 | 61,341 | 0 | 42,525 | 30 |
| 1,29E+08 | 1,02E+08 | 1,10E+08 | 1,06E+08 | 26,289 | 0 | 33,089 | 49 |
| 1,38E+08 | 1,13E+08 | 1,09E+08 | 1,09E+08 | 48,609 | 0 | 29,809 | 68 |
| 2,82E+06 | 4,60E+06 | 5,34E+06 | 3,42E+06 | 77,776 | 0 | 5,9642 | 6 |
| 2,62E+07 | 6,48E+06 | 5,07E+06 | 6,14E+06 | 323,86 | 0 | 10,341 | 17 |
| 1,90E+09 | 2,52E+09 | 2,74E+09 | 2,34E+09 | 28,785 | 0 | 323,31 | 308 |
| 7,19E+07 | 6,05E+07 | 6,41E+07 | 5,90E+07 | 86,187 | 0 | 81,731 | 75 |
| 3,08E+06 | 1,82E+06 | 1,82E+06 | 1,87E+06 | 124,11 | 0,000571 | 2,6328 | 5 |
| 4,75E+06 | 8,84E+06 | 9,74E+06 | 9,93E+06 | 40,627 | 0,000149 | 3,4792 | 13 |
| 8,30E+07 | 1,39E+08 | 1,31E+08 | 1,52E+08 | 14,286 | 0 | 9,3819 | 63 |
| 1,51E+09 | 1,84E+09 | 1,91E+09 | 1,89E+09 | 17,138 | 0 | 129,46 | 149 |
| 4,08E+07 | 3,05E+07 | 3,12E+07 | 2,31E+07 | 32,2 | 0 | 30,422 | 26 |
| 3,89E+09 | 4,82E+09 | 5,40E+09 | 5,78E+09 | 23,407 | 0 | 323,31 | 354 |
| 4,64E+07 | 4,03E+07 | 3,91E+07 | 3,76E+07 | 39,889 | 0 | 21,958 | 35 |
| 3,28E+08 | 2,99E+08 | 2,61E+08 | 2,94E+08 | 30,097 | 0 | 106,55 | 52 |
| 4,23E+07 | 2,28E+07 | 1,06E+07 | 2,01E+07 | 39,317 | 0 | 21,231 | 18 |
| 3,15E+08 | 1,97E+08 | 1,83E+08 | 1,69E+08 | 26,778 | 0 | 323,31 | 115 |
| 7,65E+08 | 6,11E+08 | 6,83E+08 | 7,02E+08 | 43,221 | 0 | 323,31 | 134 |
| 4,42E+07 | 2,70E+07 | 1,94E+07 | 2,96E+07 | 40,642 | 0 | 14,946 | 37 |
| 4,12E+06 | 9,65E+06 | 8,83E+06 | 8,73E+06 | 57,919 | 0,000158 | 5,1086 | 14 |
| 3,82E+09 | 3,35E+09 | 3,48E+09 | 3,48E+09 | 94,629 | 0 | 323,31 | 408 |
| 2,23E+08 | 1,66E+08 | 1,70E+08 | 1,63E+08 | 286,52 | 0 | 88,819 | 150 |
| 3,90E+06 | 1,02E+06 | 1,04E+06 | 1,69E+06 | 46,826 | 0,000291 | 3,0297 | 3 |
| 4,05E+08 | 5,00E+08 | 5,39E+08 | 4,84E+08 | 128,65 | 0 | 323,31 | 216 |
| 5,94E+09 | 6,41E+09 | 6,62E+09 | 6,18E+09 | 17,222 | 0 | 71,034 | 344 |
| 8,75E+07 | 1,18E+08 | 1,06E+08 | 1,12E+08 | 52,246 | 0 | 49,279 | 73 |
| 1,27E+08 | 1,18E+08 | 1,19E+08 | 1,16E+08 | 71,686 | 0 | 89,414 | 93 |
| 2,67E+08 | 1,36E+06 | 1,06E+07 | 2,16E+07 | 332,91 | 0 | 95,496 | 100 |
| 2,13E+09 | 1,93E+09 | 1,79E+09 | 1,94E+09 | 51,112 | 0 | 323,31 | 299 |
| 2,21E+08 | 1,86E+08 | 1,62E+08 | 1,92E+08 | 69,84 | 0 | 129,81 | 106 |
| 3,59E+07 | 4,00E+07 | 3,85E+07 | 3,98E+07 | 35,041 | 0 | 11,651 | 21 |
| 6,52E+09 | 5,86E+09 | 6,01E+09 | 5,95E+09 | 59,623 | 0 | 323,31 | 614 |
| 2,42E+08 | 3,09E+08 | 3,04E+08 | 2,80E+08 | 38,219 | 0 | 277,32 | 99 |
| 8,78E+07 | 1,09E+08 | 1,08E+08 | 9,91E+07 | 49,795 | 0 | 60,784 | 92 |
| 2,52E+09 | 2,88E+09 | 2,94E+09 | 2,80E+09 | 138,43 | 0 | 323,31 | 505 |
| 2,19E+07 | 1,76E+07 | 2,03E+07 | 1,93E+07 | 60,841 | 0 | 17,226 | 27 |
| 1,03E+08 | 1,50E+08 | 1,38E+08 | 1,47E+08 | 79,198 | 0 | 66,923 | 124 |
| 8,09E+08 | 9,69E+08 | 8,83E+08 | 9,36E+08 | 46,247 | 0 | 266,86 | 142 |
| 3,01E+07 | 5,66E+06 | 1,67E+06 | 9,68E+06 | 17,024 | 0 | 19,67 | 17 |
| 2,06E+08 | 1,84E+08 | 1,47E+08 | 1,46E+08 | 30,036 | 0 | 24,545 | 50 |
| 2,00E+07 | 6,78E+06 | 1,12E+07 | 1,07E+07 | 17,525 | 0 | 23,933 | 11 |
| 2,68E+06 | 1,86E+07 | 1,89E+07 | 1,31E+07 | 40,474 | 0,000157 | 4,8727 | 17 |
| 2,89E+07 | 2,27E+07 | 2,35E+07 | 1,72E+07 | 65,877 | 0 | 21,679 | 21 |
| 1,23E+08 | 1,15E+08 | 9,97E+07 | 1,04E+08 | 63,02 | 0 | 115,46 | 100 |

| | | | | | | | |
|---|---|---|---|---|---|---|---|
| 2,35E+07 | 3,40E+07 | 3,30E+07 | 3,79E+07 | 33,471 | 0 | 29,411 | 23 |
| 2,82E+09 | 2,59E+09 | 2,49E+09 | 2,49E+09 | 48,647 | 0 | 323,31 | 496 |
| 2,36E+06 | 3,78E+06 | 5,54E+06 | 4,75E+06 | 46,986 | 0,001409 | 2,3568 | 5 |
| 1,22E+08 | 2,77E+06 | 7,93E+06 | 2,60E+07 | 55,569 | 0 | 12,191 | 15 |
| 5,25E+07 | 7,21E+07 | 7,15E+07 | 7,80E+07 | 35,796 | 0 | 19,998 | 53 |
| 1,17E+08 | 7,73E+07 | 7,56E+07 | 7,21E+07 | 128,63 | 0 | 204,97 | 72 |
| 7,89E+07 | 1,04E+08 | 1,08E+08 | 1,20E+08 | 52,797 | 0 | 44,127 | 72 |
| 1,05E+09 | 1,60E+09 | 1,60E+09 | 1,77E+09 | 31,078 | 0 | 323,31 | 83 |
| 2,74E+07 | 1,64E+06 | 7,37E+06 | 2,00E+06 | 36,438 | 0 | 8,9601 | 20 |
| 1,66E+08 | 1,15E+08 | 1,37E+08 | 1,09E+08 | 63,752 | 0 | 24,858 | 74 |
| 1,41E+08 | 2,55E+08 | 2,35E+08 | 2,51E+08 | 86,052 | 0 | 215,35 | 106 |
| 1,12E+08 | 1,47E+08 | 1,35E+08 | 1,48E+08 | 56,677 | 0 | 97,259 | 79 |
| 4,28E+08 | 6,55E+08 | 5,74E+08 | 6,51E+08 | 145,24 | 0 | 159,31 | 184 |
| 1,08E+08 | 5,47E+07 | 5,69E+07 | 5,88E+07 | 37,22 | 0 | 23,629 | 22 |
| 2,89E+08 | 1,78E+08 | 2,05E+08 | 1,83E+08 | 24,57 | 0 | 137,04 | 64 |
| 1,49E+08 | 1,31E+08 | 1,21E+08 | 1,18E+08 | 165,59 | 0 | 256,95 | 116 |
| 3,32E+07 | 1,43E+06 | 9,47E+06 | 4,32E+06 | 63,634 | 0 | 8,5891 | 12 |
| 4,90E+07 | 4,21E+07 | 3,96E+07 | 3,30E+07 | 25,414 | 0 | 10,931 | 29 |
| 4,75E+07 | 7,01E+07 | 6,45E+07 | 6,75E+07 | 79,434 | 0 | 22,598 | 63 |
| 7,54E+08 | 6,33E+08 | 6,39E+08 | 6,32E+08 | 33,558 | 0 | 323,31 | 137 |
| 1,54E+07 | 4,07E+07 | 2,93E+07 | 2,74E+07 | 27,808 | 0 | 15,556 | 32 |
| 5,52E+07 | 4,94E+07 | 4,02E+07 | 3,78E+07 | 46,6 | 0 | 135,74 | 38 |
| 9,61E+07 | 6,81E+07 | 7,26E+07 | 7,61E+07 | 122,61 | 0 | 28,463 | 65 |
| 1,24E+07 | 2,87E+07 | 2,10E+07 | 2,26E+07 | 44,356 | 0 | 19,812 | 24 |
| 1,63E+08 | 2,52E+08 | 1,95E+08 | 1,97E+08 | 17,66 | 0 | 109,07 | 50 |
| 6,44E+08 | 6,23E+08 | 6,16E+08 | 5,89E+08 | 206,72 | 0 | 323,31 | 261 |
| 4,21E+08 | 3,01E+08 | 3,78E+08 | 3,42E+08 | 46,175 | 0 | 173,95 | 100 |
| 2,20E+08 | 1,62E+08 | 1,76E+08 | 1,38E+08 | 31,443 | 0 | 121,4 | 60 |
| 1,14E+08 | 1,52E+08 | 1,71E+08 | 1,41E+08 | 81,644 | 0 | 103,6 | 71 |
| 1,12E+08 | 1,05E+08 | 1,03E+08 | 1,01E+08 | 28,979 | 0 | 25,386 | 72 |
| 7,24E+06 | 9,98E+06 | 1,10E+07 | 1,14E+07 | 51,226 | 0,000156 | 4,5998 | 16 |
| 3,24E+07 | 2,67E+07 | 2,77E+07 | 3,04E+07 | 35,905 | 0 | 11,385 | 30 |
| 2,68E+08 | 2,15E+08 | 2,11E+08 | 1,94E+08 | 12,293 | 0 | 71,78 | 49 |
| 3,24E+08 | 4,01E+08 | 3,55E+08 | 3,57E+08 | 48,327 | 0 | 261,85 | 116 |
| 2,30E+09 | 2,82E+09 | 2,86E+09 | 2,62E+09 | 26,268 | 0 | 323,31 | 458 |
| 3,02E+07 | 2,34E+07 | 2,54E+07 | 2,58E+07 | 290,8 | 0 | 7,5848 | 25 |
| 6,67E+07 | 9,63E+07 | 9,88E+07 | 7,82E+07 | 66,289 | 0 | 98,007 | 54 |
| 4,08E+07 | 5,21E+07 | 4,41E+07 | 5,01E+07 | 140,58 | 0 | 22,05 | 46 |
| 5,82E+08 | 6,38E+08 | 6,29E+08 | 6,03E+08 | 66,545 | 0 | 322,19 | 171 |
| 2,81E+06 | 5,30E+06 | 5,28E+06 | 6,04E+06 | 30,469 | 0,00291 | 2,009 | 5 |
| 9,91E+07 | 1,14E+08 | 1,49E+08 | 1,44E+08 | 18,582 | 0 | 171,98 | 56 |
| 2,90E+07 | 4,60E+07 | 4,28E+07 | 3,94E+07 | 56,405 | 0 | 26,58 | 31 |
| 2,96E+07 | 2,58E+07 | 2,71E+07 | 2,54E+07 | 39,394 | 0 | 34,902 | 33 |
| 7,66E+08 | 7,23E+08 | 6,72E+08 | 7,13E+08 | 145,09 | 0 | 323,31 | 276 |
| 9,46E+07 | 1,37E+08 | 1,28E+08 | 1,40E+08 | 54,044 | 0 | 38,095 | 75 |
| 5,91E+07 | 4,58E+07 | 4,40E+07 | 5,24E+07 | 63,396 | 0 | 37,25 | 54 |
| 4,22E+06 | 6,96E+06 | 6,42E+06 | 7,88E+06 | 64,13 | 0,00141 | 2,3593 | 9 |
| 9,38E+06 | 1,62E+07 | 1,31E+07 | 1,44E+07 | 43,182 | 0,000432 | 2,8257 | 11 |
| 2,05E+08 | 1,85E+08 | 1,46E+08 | 1,58E+08 | 32,905 | 0 | 161,83 | 97 |
| 1,59E+08 | 1,23E+08 | 1,02E+08 | 9,61E+07 | 65,636 | 0 | 60,909 | 73 |

| | | | | | | | |
|---|---|---|---|---|---|---|---|
| 3,72E+08 | 3,40E+08 | 3,20E+08 | 3,25E+08 | 88,062 | 0 | 323,31 | 151 |
| 8,97E+07 | 1,64E+08 | 1,66E+08 | 1,44E+08 | 29,884 | 0 | 24,184 | 58 |
| 1,25E+08 | 1,08E+08 | 9,26E+07 | 1,02E+08 | 48,206 | 0 | 79,973 | 76 |
| 4,12E+07 | 3,41E+07 | 2,86E+07 | 3,40E+07 | 96,4 | 0 | 17,6 | 38 |
| 4,52E+07 | 5,67E+07 | 6,37E+07 | 5,59E+07 | 24,464 | 0 | 62,497 | 23 |
| 5,38E+05 | 2,62E+06 | 2,18E+06 | 1,69E+06 | 36,384 | 0 | 56,088 | 5 |
| 2,60E+07 | 6,74E+06 | 1,55E+07 | 8,15E+06 | 18,226 | 0 | 12,609 | 14 |
| 5,08E+07 | 3,45E+07 | 4,07E+07 | 4,43E+07 | 31,739 | 0 | 12,736 | 19 |
| 4,35E+06 | 1,01E+07 | 1,01E+07 | 7,68E+06 | 69,601 | 0 | 7,3993 | 14 |
| 5,26E+07 | 7,54E+07 | 8,65E+07 | 7,78E+07 | 57,62 | 0 | 30,627 | 60 |
| 2,93E+07 | 3,63E+07 | 3,24E+07 | 3,88E+07 | 122,43 | 0 | 20,511 | 43 |
| 3,44E+08 | 2,52E+08 | 2,60E+08 | 3,08E+08 | 91,719 | 0 | 125,87 | 122 |
| 1,47E+07 | 2,68E+07 | 1,90E+07 | 1,91E+07 | 166,41 | 0 | 14,598 | 37 |
| 2,69E+07 | 1,44E+07 | 1,73E+07 | 1,74E+07 | 23,612 | 0 | 16,576 | 15 |
| 4,45E+07 | 1,60E+07 | 2,52E+07 | 2,20E+07 | 55,948 | 0 | 21,321 | 45 |
| 3,43E+08 | 2,96E+08 | 2,97E+08 | 3,26E+08 | 84,937 | 0 | 177,51 | 120 |
| 5,90E+07 | 4,30E+07 | 4,41E+07 | 5,03E+07 | 34,772 | 0 | 13,381 | 35 |
| 1,46E+07 | 3,43E+07 | 2,58E+07 | 2,54E+07 | 30,654 | 0 | 13,663 | 24 |
| 1,72E+07 | 1,43E+07 | 1,07E+07 | 1,27E+07 | 62,907 | 0 | 9,0857 | 27 |
| 1,15E+07 | 4,77E+06 | 3,92E+06 | 1,69E+06 | 43,136 | 0,000155 | 4,4833 | 6 |
| 2,16E+07 | 1,35E+08 | 1,21E+08 | 1,21E+08 | 115,61 | 0 | 23,853 | 55 |
| 2,48E+07 | 1,66E+07 | 2,05E+07 | 1,68E+07 | 43,231 | 0 | 6,2976 | 20 |
| 4,45E+07 | 3,44E+06 | 1,66E+06 | 1,52E+07 | 32,523 | 0 | 16,423 | 13 |
| 1,87E+07 | 2,28E+06 | 9,78E+06 | 4,06E+06 | 31,022 | 0 | 11,5 | 15 |
| 2,12E+07 | 4,41E+08 | 5,81E+08 | 6,49E+08 | 54,748 | 0 | 17,678 | 40 |
| 5,07E+07 | 2,26E+07 | 7,40E+06 | 4,22E+06 | 38,932 | 0 | 27,927 | 15 |
| 1,74E+07 | 3,22E+07 | 3,54E+07 | 3,48E+07 | 25,444 | 0 | 11,826 | 11 |
| 2,28E+07 | 4,29E+06 | 1,04E+07 | 1,16E+07 | 31,3 | 0 | 12,644 | 11 |
| 1,73E+07 | 1,49E+07 | 1,29E+07 | 1,35E+07 | 91,521 | 0 | 20,267 | 37 |
| 2,11E+09 | 2,53E+09 | 2,64E+09 | 2,77E+09 | 66,612 | 0 | 323,31 | 368 |
| 4,14E+07 | 5,11E+07 | 5,80E+07 | 6,18E+07 | 55,227 | 0 | 231,32 | 45 |
| 3,58E+06 | 2,54E+06 | 2,80E+06 | 2,56E+06 | 50,201 | 0,000855 | 2,5995 | 2 |
| 4,55E+06 | 1,22E+07 | 1,28E+07 | 5,93E+06 | 140,97 | 0 | 8,5532 | 6 |
| 4,22E+07 | 2,99E+07 | 2,16E+07 | 2,99E+07 | 27,631 | 0 | 52,548 | 22 |
| 2,98E+07 | 1,75E+07 | 2,04E+07 | 2,47E+07 | 162,34 | 0 | 24,336 | 36 |
| 2,05E+07 | 2,66E+07 | 2,30E+07 | 2,74E+07 | 37,825 | 0 | 13,377 | 23 |
| 2,70E+08 | 1,71E+08 | 1,97E+08 | 2,07E+08 | 46,895 | 0 | 84,951 | 100 |
| 3,62E+06 | 9,64E+06 | 1,11E+07 | 1,80E+07 | 124,59 | 0 | 7,4627 | 13 |
| 8,67E+06 | 2,01E+07 | 1,97E+07 | 1,97E+07 | 18,473 | 0,004633 | 1,7339 | 6 |
| 6,79E+07 | 1,58E+06 | 3,11E+06 | 1,03E+06 | 44,091 | 0,003174 | 1,9436 | 4 |
| 3,69E+06 | 7,44E+06 | 6,45E+06 | 7,84E+06 | 57,669 | 0,000155 | 4,4347 | 10 |
| 1,23E+08 | 1,59E+08 | 1,39E+08 | 1,46E+08 | 60,211 | 0 | 121,82 | 83 |
| 1,51E+08 | 1,32E+08 | 1,05E+08 | 1,02E+08 | 93,055 | 0 | 121,59 | 90 |
| 2,86E+08 | 2,15E+08 | 1,82E+08 | 1,91E+08 | 22,371 | 0 | 265,03 | 72 |
| 5,87E+07 | 4,66E+07 | 4,86E+07 | 5,30E+07 | 148,95 | 0 | 48,69 | 61 |
| 6,32E+06 | 2,00E+07 | 3,69E+07 | 3,27E+07 | 15,766 | 0 | 7,2436 | 19 |
| 9,62E+06 | 1,94E+07 | 1,45E+07 | 2,08E+07 | 53,941 | 0 | 6,291 | 16 |
| 4,08E+06 | 8,14E+06 | 7,86E+06 | 1,04E+07 | 155,8 | 0 | 11,33 | 10 |
| 9,02E+06 | 3,86E+07 | 2,59E+07 | 3,33E+07 | 35,294 | 0 | 21,35 | 12 |
| 2,09E+07 | 2,83E+06 | 1,33E+06 | 7,89E+06 | 33,479 | 0,002098 | 2,168 | 5 |

| | | | | | | | |
|---|---|---|---|---|---|---|---|
| 1,36E+07 | 2,15E+07 | 1,78E+07 | 2,41E+07 | 136,1 | 0,008201 | 1,5041 | 5 |
| 7,98E+07 | 8,70E+07 | 8,71E+07 | 8,17E+07 | 18,127 | 0 | 198,73 | 46 |
| 1,25E+07 | 6,89E+06 | 7,31E+06 | 6,14E+06 | 28,572 | 0,0021 | 2,1931 | 5 |
| 1,09E+08 | 9,04E+07 | 8,70E+07 | 7,32E+07 | 108,35 | 0 | 22,29 | 79 |
| 3,63E+08 | 3,91E+08 | 3,91E+08 | 3,76E+08 | 57,624 | 0 | 263,29 | 169 |
| 3,05E+08 | 2,79E+08 | 2,62E+08 | 2,62E+08 | 31,474 | 0 | 173,89 | 77 |
| 3,22E+07 | 1,55E+07 | 1,55E+07 | 1,45E+07 | 126,84 | 0 | 20,398 | 35 |
| 2,60E+07 | 1,79E+07 | 1,42E+07 | 1,60E+07 | 24,914 | 0 | 17,82 | 21 |
| 1,87E+08 | 1,58E+08 | 1,70E+08 | 1,54E+08 | 35,079 | 0 | 86,952 | 98 |
| 2,74E+06 | 4,44E+06 | 4,58E+06 | 4,22E+06 | 22,144 | 0,005813 | 1,6077 | 5 |
| 5,18E+09 | 4,25E+09 | 4,21E+09 | 4,61E+09 | 59,555 | 0 | 323,31 | 714 |
| 6,91E+06 | 9,46E+06 | 9,37E+06 | 8,62E+06 | 69,775 | 0,000156 | 4,7835 | 12 |
| 1,94E+07 | 1,82E+06 | 9,04E+06 | 2,86E+06 | 32,904 | 0 | 13,502 | 9 |
| 6,26E+07 | 1,55E+08 | 1,38E+08 | 1,54E+08 | 34,165 | 0 | 31,165 | 41 |
| 1,63E+08 | 2,43E+08 | 2,30E+08 | 2,38E+08 | 137,5 | 0 | 169,49 | 139 |
| 5,13E+06 | 9,53E+06 | 1,20E+07 | 7,92E+06 | 98,573 | 0,004891 | 1,6871 | 7 |
| 6,31E+06 | 1,92E+07 | 9,55E+06 | 1,01E+07 | 80,121 | 0 | 9,3092 | 8 |
| 9,59E+06 | 6,42E+06 | 5,07E+06 | 2,82E+06 | 36,807 | 0,000433 | 2,8344 | 12 |
| 2,21E+07 | 1,15E+07 | 1,04E+07 | 1,55E+07 | 72,138 | 0 | 17,231 | 18 |
| 1,45E+07 | 8,59E+06 | 4,03E+06 | 4,31E+06 | 79,075 | 0 | 9,8459 | 15 |
| 1,17E+07 | 5,88E+06 | 6,39E+06 | 4,70E+06 | 46,475 | 0 | 6,102 | 10 |
| 2,88E+09 | 3,33E+09 | 3,17E+09 | 3,17E+09 | 82,406 | 0 | 323,31 | 532 |
| 2,24E+07 | 1,44E+07 | 1,91E+07 | 1,82E+07 | 68,171 | 0 | 138,07 | 18 |
| 4,69E+06 | 1,42E+07 | 1,47E+07 | 1,48E+07 | 63,98 | 0 | 9,5558 | 11 |
| 5,77E+07 | 7,79E+07 | 7,43E+07 | 7,43E+07 | 58,458 | 0 | 45,143 | 46 |
| 3,77E+07 | 6,40E+07 | 4,86E+07 | 6,96E+07 | 99,229 | 0 | 19,032 | 55 |
| 7,85E+07 | 6,64E+06 | 1,56E+07 | 1,76E+06 | 53,516 | 0 | 19,778 | 38 |
| 2,69E+07 | 4,89E+07 | 5,13E+07 | 4,65E+07 | 68,689 | 0 | 13,834 | 54 |
| 1,58E+07 | 1,07E+07 | 7,03E+06 | 5,54E+06 | 17,499 | 0 | 14,648 | 7 |
| 6,14E+08 | 5,58E+08 | 5,36E+08 | 5,60E+08 | 58,278 | 0 | 244,15 | 212 |
| 3,50E+06 | 2,09E+07 | 2,95E+07 | 5,67E+07 | 47,323 | 0,001547 | 2,2985 | 7 |
| 7,31E+08 | 3,24E+08 | 4,00E+08 | 2,47E+08 | 10,803 | 0 | 135,18 | 45 |
| 2,13E+07 | 2,72E+07 | 3,26E+07 | 2,58E+07 | 24,748 | 0 | 45,518 | 19 |
| 7,31E+07 | 8,09E+07 | 7,91E+07 | 8,41E+07 | 139,61 | 0 | 41,601 | 77 |
| 6,65E+07 | 5,97E+07 | 5,08E+07 | 5,64E+07 | 30,593 | 0 | 28,723 | 51 |
| 1,16E+08 | 6,94E+07 | 8,40E+07 | 8,53E+07 | 51,313 | 0 | 88,252 | 25 |
| 2,75E+08 | 2,08E+08 | 2,05E+08 | 1,75E+08 | 56,7 | 0 | 323,31 | 92 |
| 2,29E+07 | 3,69E+07 | 3,03E+07 | 3,60E+07 | 37,202 | 0 | 7,0501 | 21 |
| 1,26E+08 | 1,08E+08 | 1,15E+08 | 1,15E+08 | 53,255 | 0 | 25,311 | 64 |
| 2,42E+08 | 2,12E+08 | 2,18E+08 | 2,25E+08 | 169,23 | 0 | 203,43 | 172 |
| 1,70E+06 | 5,36E+06 | 8,29E+06 | 6,90E+06 | 52,612 | 0,000148 | 3,402 | 5 |
| 6,08E+07 | 4,41E+07 | 4,07E+07 | 4,21E+07 | 85,843 | 0 | 34,523 | 53 |
| 6,52E+07 | 8,78E+07 | 9,30E+07 | 7,73E+07 | 349,41 | 0 | 41,648 | 112 |
| 5,11E+06 | 2,01E+06 | 2,87E+06 | 2,12E+06 | 28,954 | 0 | 14,907 | 5 |
| 1,78E+07 | 1,13E+07 | 1,03E+07 | 1,47E+07 | 41,246 | 0 | 9,0603 | 15 |
| 6,35E+07 | 8,08E+07 | 7,03E+07 | 6,96E+07 | 76,125 | 0 | 31,658 | 56 |
| 2,71E+07 | 2,06E+07 | 2,33E+07 | 1,73E+07 | 109,96 | 0 | 28,865 | 28 |
| 1,87E+09 | 2,38E+09 | 2,50E+09 | 2,45E+09 | 55,983 | 0 | 323,31 | 322 |
| 6,34E+07 | 7,57E+07 | 8,37E+07 | 9,91E+07 | 55,026 | 0 | 79,12 | 41 |
| 3,95E+08 | 4,30E+08 | 4,17E+08 | 4,21E+08 | 74,095 | 0 | 246,55 | 171 |

| | | | | | | | |
|---|---|---|---|---|---|---|---|
| 6,16E+08 | 5,16E+08 | 5,34E+08 | 5,54E+08 | 85,221 | 0 | 77,114 | 190 |
| 2,82E+08 | 2,55E+08 | 2,52E+08 | 2,72E+08 | 26,022 | 0 | 172,9 | 69 |
| 2,22E+07 | 1,39E+07 | 1,57E+07 | 8,00E+06 | 119,32 | 0 | 14,343 | 29 |
| 6,62E+09 | 8,58E+09 | 8,35E+09 | 8,34E+09 | 47,994 | 0 | 323,31 | 829 |
| 2,73E+08 | 2,27E+08 | 2,13E+08 | 2,18E+08 | 32,399 | 0 | 58,109 | 96 |
| 8,07E+07 | 1,35E+08 | 1,24E+08 | 1,38E+08 | 83,145 | 0 | 35,381 | 66 |
| 8,58E+08 | 1,13E+09 | 1,07E+09 | 1,09E+09 | 181,91 | 0 | 323,31 | 438 |
| 6,62E+07 | 1,05E+08 | 1,00E+08 | 1,18E+08 | 145,78 | 0 | 39,74 | 67 |
| 3,04E+06 | 1,08E+07 | 1,30E+07 | 1,30E+07 | 94,678 | 0,000151 | 3,767 | 6 |
| 3,36E+08 | 2,02E+08 | 2,12E+08 | 2,13E+08 | 35,185 | 0 | 104,42 | 78 |
| 1,42E+07 | 2,54E+07 | 3,34E+07 | 3,49E+07 | 41,433 | 0 | 12,489 | 15 |
| 1,29E+07 | 2,04E+07 | 1,70E+07 | 2,33E+07 | 70,048 | 0,002779 | 2,0465 | 16 |
| 7,44E+07 | 1,05E+08 | 1,06E+08 | 1,18E+08 | 38,724 | 0 | 87,108 | 57 |
| 5,50E+07 | 3,78E+07 | 3,55E+07 | 4,38E+07 | 127 | 0 | 34,955 | 53 |
| 8,50E+06 | 6,40E+06 | 7,75E+06 | 5,76E+06 | 117,08 | 0,00015 | 3,6262 | 12 |
| 3,96E+08 | 4,26E+08 | 4,66E+08 | 4,58E+08 | 166,59 | 0 | 323,31 | 240 |
| 6,49E+06 | 1,86E+08 | 1,01E+08 | 1,99E+08 | 101,51 | 0 | 13,835 | 28 |
| 1,41E+09 | 9,89E+08 | 1,00E+09 | 1,05E+09 | 26,51 | 0 | 302,74 | 111 |
| 4,36E+08 | 5,85E+08 | 5,50E+08 | 6,49E+08 | 34,086 | 0 | 145,11 | 106 |
| 3,38E+07 | 1,70E+07 | 1,66E+07 | 2,64E+07 | 42,354 | 0 | 9,5853 | 23 |
| 1,26E+08 | 1,06E+08 | 1,05E+08 | 9,34E+07 | 88,66 | 0 | 164,68 | 103 |
| 3,19E+07 | 2,70E+07 | 2,32E+07 | 1,98E+07 | 72,638 | 0 | 60,809 | 41 |
| 1,39E+08 | 1,06E+08 | 1,24E+08 | 1,27E+08 | 70,68 | 0 | 180,34 | 71 |
| 2,08E+06 | 8,95E+06 | 1,09E+07 | 1,18E+07 | 105,37 | 0 | 8,607 | 9 |
| 9,38E+08 | 1,02E+09 | 9,84E+08 | 1,03E+09 | 52,22 | 0 | 323,31 | 321 |
| 3,14E+08 | 4,08E+08 | 3,92E+08 | 4,06E+08 | 36,71 | 0 | 123,46 | 77 |
| 3,60E+07 | 4,52E+07 | 5,14E+07 | 4,04E+07 | 162,27 | 0 | 22,945 | 44 |
| 9,21E+06 | 1,54E+07 | 2,19E+07 | 2,00E+07 | 40,477 | 0 | 10,752 | 25 |
| 4,68E+07 | 5,20E+07 | 4,85E+07 | 5,19E+07 | 102,54 | 0 | 47,7 | 53 |
| 3,12E+07 | 2,77E+07 | 2,06E+07 | 2,33E+07 | 50,548 | 0 | 8,8475 | 28 |
| 1,54E+09 | 1,27E+09 | 1,37E+09 | 1,31E+09 | 121,63 | 0 | 323,31 | 477 |
| 1,17E+08 | 7,57E+07 | 7,02E+07 | 7,53E+07 | 164,99 | 0 | 116,42 | 68 |
| 3,21E+06 | 9,23E+06 | 1,25E+07 | 8,40E+06 | 57,342 | 0,000152 | 4,0053 | 11 |
| 1,42E+08 | 7,32E+07 | 8,45E+07 | 7,92E+07 | 38,637 | 0 | 53,915 | 41 |
| 2,13E+08 | 1,89E+08 | 1,87E+08 | 1,95E+08 | 140,74 | 0 | 130,05 | 159 |
| 1,08E+08 | 1,68E+08 | 1,82E+08 | 1,70E+08 | 65,052 | 0 | 77,489 | 99 |
| 6,75E+07 | 2,33E+06 | 1,35E+07 | 1,46E+07 | 25,608 | 0,000159 | 5,2568 | 14 |
| 5,35E+06 | 2,73E+07 | 2,50E+07 | 9,16E+06 | 154,35 | 0 | 17,421 | 24 |
| 2,88E+08 | 4,37E+08 | 3,61E+08 | 5,04E+08 | 43,865 | 0 | 44,755 | 55 |
| 1,39E+09 | 1,06E+09 | 1,10E+09 | 1,09E+09 | 33,188 | 0 | 323,31 | 133 |
| 2,12E+07 | 3,26E+07 | 3,29E+07 | 4,08E+07 | 144,8 | 0 | 17,089 | 34 |
| 2,77E+08 | 3,60E+08 | 2,95E+08 | 3,32E+08 | 44,123 | 0 | 110,84 | 104 |
| 2,11E+09 | 1,79E+09 | 1,97E+09 | 1,99E+09 | 46,839 | 0 | 323,31 | 358 |
| 1,14E+08 | 7,91E+07 | 7,01E+07 | 7,31E+07 | 34,577 | 0 | 204,85 | 65 |
| 1,26E+07 | 7,16E+06 | 7,58E+06 | 6,00E+06 | 76,865 | 0 | 7,172 | 13 |
| 1,14E+08 | 8,14E+07 | 8,70E+07 | 7,85E+07 | 70,755 | 0 | 102,17 | 71 |
| 2,12E+07 | 1,16E+07 | 1,46E+07 | 1,71E+07 | 47,227 | 0 | 7,5126 | 17 |
| 2,98E+07 | 1,98E+07 | 1,67E+07 | 2,37E+07 | 85,515 | 0 | 28,002 | 46 |
| 8,41E+08 | 1,03E+09 | 1,10E+09 | 1,09E+09 | 58,537 | 0 | 304,19 | 182 |
| 1,02E+07 | 8,70E+06 | 9,47E+06 | 8,44E+06 | 65,314 | 0 | 11,2 | 15 |

| | | | | | | | |
|---|---|---|---|---|---|---|---|
| 1,67E+07 | 9,65E+06 | 9,29E+06 | 6,01E+06 | 68,529 | 0 | 10,448 | 18 |
| 9,66E+07 | 7,89E+07 | 7,51E+07 | 6,86E+07 | 270,64 | 0 | 43,965 | 98 |
| 2,80E+07 | 3,20E+07 | 3,69E+07 | 3,90E+07 | 212,4 | 0 | 43,397 | 35 |
| 3,31E+07 | 1,80E+07 | 1,96E+07 | 2,04E+07 | 48,154 | 0 | 24,079 | 36 |
| 9,88E+07 | 8,51E+07 | 8,22E+07 | 9,47E+07 | 109,05 | 0 | 47,934 | 100 |
| 4,49E+08 | 4,28E+08 | 3,94E+08 | 4,21E+08 | 36,588 | 0 | 105,37 | 83 |
| 1,53E+09 | 8,80E+08 | 1,04E+09 | 9,61E+08 | 37,331 | 0 | 323,31 | 114 |
| 3,30E+07 | 1,59E+07 | 1,81E+07 | 8,14E+06 | 98,76 | 0 | 48,272 | 18 |
| 1,16E+08 | 9,76E+07 | 1,04E+08 | 1,03E+08 | 109,98 | 0 | 57,634 | 75 |
| 1,25E+09 | 1,45E+09 | 1,55E+09 | 1,57E+09 | 105,53 | 0 | 323,31 | 330 |
| 9,74E+06 | 2,32E+06 | 6,43E+06 | 3,64E+06 | 48,972 | 0 | 7,3864 | 8 |
| 1,16E+07 | 7,09E+06 | 3,41E+06 | 2,41E+06 | 23,252 | 0 | 5,3269 | 17 |
| 8,55E+08 | 6,77E+08 | 6,47E+08 | 6,84E+08 | 70,527 | 0 | 323,31 | 205 |
| 5,62E+08 | 4,80E+08 | 4,82E+08 | 4,95E+08 | 53,254 | 0 | 323,31 | 164 |
| 1,75E+07 | 1,88E+07 | 2,32E+07 | 2,16E+07 | 71,577 | 0,000156 | 4,7574 | 12 |
| 1,52E+07 | 2,46E+06 | 9,19E+06 | 3,21E+06 | 37,941 | 0,000159 | 5,1659 | 11 |
| 1,29E+08 | 1,88E+08 | 1,93E+08 | 1,74E+08 | 31,845 | 0 | 25,9 | 47 |
| 7,30E+06 | 1,02E+07 | 1,46E+07 | 1,04E+07 | 34,381 | 0,000153 | 4,2046 | 7 |
| 7,03E+07 | 1,43E+08 | 9,52E+07 | 9,27E+07 | 31,46 | 0 | 30,04 | 49 |
| 6,23E+09 | 7,58E+09 | 6,78E+09 | 6,79E+09 | 29,597 | 0 | 323,31 | 391 |
| 1,06E+07 | 3,59E+06 | 7,13E+06 | 5,69E+06 | 119,17 | 0 | 11,726 | 7 |
| 2,09E+06 | 5,62E+06 | 6,67E+06 | 5,72E+06 | 126,03 | 0,000433 | 2,8511 | 6 |
| 2,89E+09 | 3,45E+09 | 3,29E+09 | 3,40E+09 | 47,436 | 0 | 323,31 | 370 |
| 5,12E+06 | 1,37E+07 | 8,04E+06 | 8,61E+06 | 27,686 | 0 | 7,5972 | 7 |
| 7,30E+06 | 2,81E+07 | 1,75E+07 | 1,89E+07 | 93,474 | 0 | 11,409 | 13 |
| 1,56E+08 | 1,89E+08 | 2,63E+08 | 3,01E+08 | 22,06 | 0 | 27,095 | 78 |
| 1,27E+09 | 9,35E+08 | 9,49E+08 | 9,15E+08 | 59,406 | 0 | 323,31 | 166 |
| 8,28E+06 | 1,29E+07 | 1,81E+07 | 1,87E+07 | 37,021 | 0 | 35,754 | 12 |
| 2,17E+08 | 1,98E+08 | 1,77E+08 | 1,85E+08 | 43,934 | 0 | 164,14 | 55 |
| 2,07E+08 | 2,48E+08 | 2,31E+08 | 2,45E+08 | 83,013 | 0 | 142,93 | 88 |
| 1,50E+09 | 1,77E+09 | 1,90E+09 | 1,88E+09 | 83,355 | 0 | 323,31 | 295 |
| 2,16E+07 | 1,59E+07 | 1,55E+07 | 1,71E+07 | 22,066 | 0 | 7,923 | 21 |
| 2,46E+08 | 2,11E+08 | 2,22E+08 | 2,27E+08 | 70,543 | 0 | 109,8 | 119 |
| 1,70E+07 | 1,04E+07 | 5,83E+06 | 6,59E+06 | 24,97 | 0 | 18,404 | 19 |
| 5,04E+06 | 7,32E+06 | 7,87E+06 | 6,82E+06 | 77,39 | 0,000146 | 3,1119 | 15 |
| 3,26E+06 | 8,46E+06 | 5,28E+06 | 5,27E+06 | 16,018 | 0,000291 | 3,0625 | 6 |
| 1,22E+08 | 1,53E+08 | 1,51E+08 | 1,35E+08 | 78,756 | 0 | 101,26 | 113 |
| 7,93E+08 | 6,00E+08 | 6,87E+08 | 6,93E+08 | 27,254 | 0 | 323,31 | 125 |
| 2,18E+07 | 2,04E+07 | 1,74E+07 | 1,86E+07 | 56,631 | 0 | 7,605 | 14 |
| 7,10E+06 | 1,72E+07 | 2,21E+07 | 1,81E+07 | 93,896 | 0 | 13,125 | 24 |
| 2,02E+08 | 3,05E+08 | 2,92E+08 | 2,51E+08 | 141,28 | 0 | 195,21 | 176 |
| 1,18E+08 | 1,32E+08 | 1,61E+08 | 1,48E+08 | 41,879 | 0 | 72,547 | 70 |
| 3,00E+08 | 2,32E+08 | 2,70E+08 | 2,37E+08 | 123,81 | 0 | 234,22 | 161 |
| 1,03E+08 | 1,46E+08 | 1,80E+08 | 1,73E+08 | 24,148 | 0 | 22,528 | 33 |
| 8,60E+07 | 1,17E+08 | 1,38E+08 | 1,27E+08 | 77,546 | 0 | 42,437 | 76 |
| 5,73E+08 | 4,82E+08 | 4,90E+08 | 5,08E+08 | 41,215 | 0 | 313,75 | 167 |
| 2,45E+07 | 1,24E+07 | 6,65E+06 | 1,42E+07 | 45,894 | 0 | 11,765 | 16 |
| 1,69E+07 | 5,06E+07 | 4,58E+07 | 5,35E+07 | 55,464 | 0 | 7,272 | 16 |
| 2,77E+06 | 1,21E+07 | 1,54E+07 | 1,35E+07 | 96,155 | 0 | 5,945 | 17 |
| 3,18E+08 | 3,86E+08 | 3,80E+08 | 3,47E+08 | 33,772 | 0 | 323,31 | 134 |

| | | | | | | | |
|---|---|---|---|---|---|---|---|
| 2,16E+07 | 3,35E+07 | 2,90E+07 | 3,39E+07 | 147,45 | 0 | 12,538 | 31 |
| 2,69E+09 | 2,03E+09 | 2,18E+09 | 2,04E+09 | 31,052 | 0 | 323,31 | 154 |
| 3,87E+07 | 3,01E+07 | 3,12E+07 | 3,58E+07 | 36,161 | 0 | 16,607 | 25 |
| 2,03E+06 | 2,17E+07 | 5,37E+07 | 7,18E+06 | 40,084 | 0,000291 | 3,0443 | 4 |
| 1,50E+07 | 6,66E+06 | 8,91E+06 | 5,02E+06 | 24,319 | 0 | 145,46 | 12 |
| 3,55E+08 | 3,33E+08 | 3,22E+08 | 3,28E+08 | 50,304 | 0 | 124,14 | 134 |
| 2,52E+07 | 9,24E+07 | 8,57E+07 | 6,18E+07 | 237,77 | 0 | 32,206 | 42 |
| 3,19E+08 | 2,42E+08 | 2,31E+08 | 2,13E+08 | 117,96 | 0 | 323,31 | 122 |
| 4,43E+08 | 3,08E+08 | 2,61E+08 | 2,93E+08 | 49,067 | 0 | 286,87 | 139 |
| 4,59E+07 | 1,17E+08 | 9,29E+07 | 9,18E+07 | 47,316 | 0 | 30,235 | 34 |
| 5,96E+06 | 1,36E+06 | 3,67E+06 | 2,00E+06 | 91,382 | 0 | 5,5668 | 10 |
| 1,24E+07 | 2,00E+07 | 1,66E+07 | 1,48E+07 | 33,298 | 0 | 18,522 | 13 |
| 9,63E+07 | 1,99E+08 | 1,25E+08 | 1,41E+08 | 38,884 | 0 | 156,78 | 84 |
| 3,87E+07 | 6,81E+07 | 6,02E+07 | 6,31E+07 | 130,35 | 0 | 30,022 | 40 |
| 6,22E+08 | 4,41E+08 | 3,60E+08 | 4,47E+08 | 27,418 | 0 | 286,07 | 86 |
| 5,21E+07 | 8,21E+07 | 7,80E+07 | 7,95E+07 | 63,869 | 0 | 18,242 | 50 |
| 3,39E+06 | 1,01E+07 | 8,82E+06 | 8,95E+06 | 46,001 | 0,000149 | 3,5129 | 3 |
| 4,72E+08 | 3,40E+08 | 3,27E+08 | 3,62E+08 | 32,921 | 0 | 263,98 | 50 |
| 2,81E+07 | 1,81E+07 | 1,77E+07 | 2,06E+07 | 52,336 | 0 | 21,552 | 30 |
| 6,08E+06 | 3,76E+06 | 4,69E+06 | 3,66E+06 | 31,49 | 0 | 6,4517 | 13 |
| 1,89E+07 | 7,61E+06 | 7,63E+06 | 5,99E+06 | 154,51 | 0 | 6,4545 | 18 |
| 2,14E+07 | 2,97E+07 | 5,73E+07 | 3,92E+07 | 16,168 | 0 | 37,363 | 12 |
| 1,22E+08 | 1,15E+08 | 1,16E+08 | 1,07E+08 | 77,566 | 0 | 49,511 | 64 |
| 5,63E+08 | 9,42E+08 | 9,47E+08 | 9,22E+08 | 60,574 | 0 | 323,31 | 204 |
| 2,09E+07 | 1,20E+07 | 1,14E+07 | 1,51E+07 | 22,311 | 0,000432 | 2,8182 | 9 |
| 1,07E+07 | 2,85E+07 | 2,90E+07 | 2,52E+07 | 100,31 | 0 | 15,397 | 35 |
| 1,41E+09 | 1,69E+09 | 1,61E+09 | 1,67E+09 | 52,895 | 0 | 323,31 | 344 |
| 1,27E+08 | 1,17E+08 | 1,19E+08 | 1,21E+08 | 95,102 | 0 | 70,409 | 106 |
| 1,99E+06 | 4,94E+06 | 4,54E+06 | 3,65E+06 | 20,039 | 0,002916 | 2,0309 | 4 |
| 3,56E+07 | 4,84E+07 | 6,38E+07 | 5,06E+07 | 17,675 | 0 | 10,884 | 37 |
| 9,19E+07 | 6,63E+07 | 7,62E+07 | 6,10E+07 | 208,5 | 0 | 60,815 | 72 |
| 2,18E+07 | 5,75E+07 | 4,84E+07 | 6,53E+07 | 49,988 | 0 | 53,081 | 17 |
| 4,85E+07 | 5,27E+07 | 5,62E+07 | 6,52E+07 | 21,725 | 0 | 13,649 | 29 |
| 2,83E+06 | 2,84E+07 | 1,89E+07 | 2,21E+07 | 47,101 | 0 | 7,0552 | 18 |
| 6,71E+06 | 2,12E+06 | 9,71E+05 | 3,90E+06 | 35,604 | 0,002909 | 2,0044 | 5 |
| 4,53E+07 | 2,06E+07 | 2,41E+07 | 1,21E+07 | 119,27 | 0 | 23,009 | 37 |
| 4,41E+08 | 2,97E+08 | 3,23E+08 | 3,16E+08 | 86,161 | 0 | 323,31 | 152 |
| 5,84E+08 | 6,10E+08 | 5,98E+08 | 6,35E+08 | 94,944 | 0 | 323,31 | 227 |
| 8,54E+08 | 7,28E+08 | 6,69E+08 | 6,31E+08 | 41,241 | 0 | 221,63 | 199 |
| 3,62E+07 | 6,93E+07 | 5,68E+07 | 5,85E+07 | 72,499 | 0 | 14,009 | 39 |
| 7,71E+06 | 5,53E+06 | 4,40E+06 | 5,51E+06 | 12,266 | 0,001822 | 2,2316 | 9 |
| 2,49E+07 | 1,24E+07 | 1,14E+07 | 1,56E+07 | 29,188 | 0,000148 | 3,4082 | 12 |
| 1,36E+08 | 1,26E+08 | 1,20E+08 | 1,27E+08 | 73,156 | 0 | 96,179 | 79 |
| 4,07E+09 | 4,23E+09 | 4,21E+09 | 4,11E+09 | 161,93 | 0 | 323,31 | 611 |
| 5,09E+08 | 6,15E+08 | 6,33E+08 | 5,76E+08 | 34,567 | 0 | 265,39 | 130 |
| 4,15E+08 | 3,52E+08 | 2,58E+08 | 3,19E+08 | 53,267 | 0 | 60,176 | 71 |
| 3,59E+06 | 3,34E+06 | 2,44E+06 | 2,65E+06 | 46,585 | 0,002782 | 2,0611 | 6 |
| 1,18E+07 | 6,30E+06 | 6,93E+06 | 6,71E+06 | 11,999 | 0,00015 | 3,7596 | 7 |
| 3,29E+07 | 6,20E+06 | 1,44E+07 | 4,49E+06 | 78,863 | 0 | 18,977 | 16 |
| 1,31E+09 | 1,12E+09 | 1,23E+09 | 1,18E+09 | 170,65 | 0 | 323,31 | 487 |

| | | | | | | | |
|---|---|---|---|---|---|---|---|
| 1,77E+06 | 4,27E+06 | 6,59E+06 | 5,73E+06 | 100,46 | 0,001409 | 2,3328 | 8 |
| 1,86E+07 | 1,25E+07 | 1,51E+07 | 1,34E+07 | 67,7 | 0 | 8,2718 | 16 |
| 8,35E+07 | 4,44E+07 | 6,43E+07 | 5,23E+07 | 37,52 | 0 | 36,385 | 36 |
| 7,58E+08 | 7,24E+08 | 6,46E+08 | 6,11E+08 | 72,799 | 0 | 323,31 | 211 |
| 2,94E+08 | 2,50E+08 | 2,63E+08 | 2,58E+08 | 150,07 | 0 | 147,45 | 197 |
| 2,20E+07 | 1,68E+07 | 1,90E+07 | 1,95E+07 | 101,67 | 0 | 7,6086 | 21 |
| 6,35E+06 | 1,64E+06 | 2,27E+06 | 4,18E+06 | 25,338 | 0,00833 | 1,4946 | 3 |
| 4,28E+07 | 6,51E+07 | 1,08E+08 | 8,32E+07 | 21,876 | 0 | 13,463 | 38 |
| 3,63E+07 | 6,00E+07 | 5,03E+07 | 5,61E+07 | 39,74 | 0 | 15,682 | 28 |
| 2,71E+08 | 2,03E+08 | 2,23E+08 | 1,96E+08 | 73,118 | 0 | 143,04 | 125 |
| 1,72E+06 | 4,51E+06 | 5,23E+06 | 4,71E+06 | 52,055 | 0,000434 | 2,9053 | 7 |
| 3,15E+08 | 4,15E+08 | 3,60E+08 | 4,15E+08 | 111,61 | 0 | 168,46 | 226 |
| 9,82E+08 | 8,78E+08 | 8,44E+08 | 7,72E+08 | 128,47 | 0 | 303,47 | 371 |
| 5,58E+06 | 2,88E+06 | 2,90E+06 | 4,17E+06 | 85,764 | 0,000151 | 3,8014 | 5 |
| 4,49E+08 | 3,54E+08 | 3,54E+08 | 3,52E+08 | 35,268 | 0 | 153,69 | 102 |
| 2,70E+07 | 3,40E+07 | 3,12E+07 | 3,39E+07 | 136,71 | 0 | 11,405 | 33 |
| 8,84E+06 | 3,94E+07 | 1,54E+07 | 2,15E+07 | 47,853 | 0 | 5,6256 | 15 |
| 1,12E+07 | 7,92E+06 | 5,50E+06 | 5,04E+06 | 105,96 | 0 | 6,6509 | 15 |
| 1,49E+07 | 1,72E+07 | 2,00E+07 | 2,00E+07 | 63,643 | 0 | 37,415 | 16 |
| 7,94E+06 | 5,66E+06 | 3,95E+06 | 3,71E+06 | 186,48 | 0 | 7,4656 | 14 |
| 1,92E+07 | 2,00E+07 | 2,26E+07 | 2,31E+07 | 175,31 | 0 | 24,862 | 38 |
| 1,71E+07 | 1,42E+07 | 1,30E+07 | 1,61E+07 | 163,45 | 0 | 6,7156 | 21 |
| 2,86E+09 | 1,97E+09 | 1,77E+09 | 1,74E+09 | 54,507 | 0 | 323,31 | 258 |
| 3,11E+07 | 6,11E+07 | 6,14E+07 | 6,26E+07 | 87,34 | 0 | 50,141 | 63 |
| 1,42E+07 | 1,11E+07 | 5,25E+06 | 7,73E+06 | 59,925 | 0 | 10,671 | 21 |
| 9,82E+07 | 1,64E+08 | 1,85E+08 | 1,64E+08 | 59,71 | 0 | 64,833 | 67 |
| 7,42E+06 | 4,08E+06 | 5,23E+06 | 2,17E+06 | 32,5 | 0,000148 | 3,4195 | 5 |
| 3,22E+08 | 5,74E+08 | 4,88E+08 | 5,36E+08 | 10,344 | 0 | 134,08 | 51 |
| 8,96E+06 | 8,22E+06 | 7,02E+06 | 7,64E+06 | 38,525 | 0,002368 | 2,0849 | 13 |
| 7,87E+06 | 8,58E+06 | 1,02E+07 | 1,15E+07 | 99,46 | 0 | 7,4532 | 14 |
| 1,11E+07 | 8,65E+06 | 2,99E+06 | 3,27E+06 | 51,469 | 0,00015 | 3,7195 | 6 |
| 6,22E+08 | 9,04E+08 | 8,29E+08 | 8,57E+08 | 48,37 | 0 | 323,31 | 98 |
| 1,77E+08 | 1,68E+08 | 1,64E+08 | 1,75E+08 | 62,942 | 0 | 54,4 | 96 |
| 1,99E+08 | 2,24E+08 | 2,52E+08 | 2,72E+08 | 11,557 | 0 | 19,251 | 35 |
| 3,25E+08 | 3,63E+08 | 3,46E+08 | 3,41E+08 | 46,325 | 0 | 214,56 | 84 |
| 3,46E+07 | 4,61E+07 | 5,33E+07 | 5,02E+07 | 42,879 | 0 | 18,562 | 38 |
| 6,70E+06 | 2,52E+06 | 3,62E+06 | 3,41E+06 | 99,403 | 0,00015 | 3,666 | 8 |
| 3,27E+08 | 3,90E+08 | 3,92E+08 | 4,00E+08 | 87,264 | 0 | 178,41 | 197 |
| 5,67E+08 | 5,03E+08 | 4,68E+08 | 5,23E+08 | 67,544 | 0 | 122,48 | 121 |
| 1,32E+08 | 2,92E+06 | 1,48E+06 | 2,83E+07 | 134,71 | 0 | 41,156 | 50 |
| 1,43E+09 | 1,05E+09 | 9,06E+08 | 1,03E+09 | 56,477 | 0 | 323,31 | 123 |
| 1,46E+08 | 1,27E+08 | 1,30E+08 | 1,27E+08 | 156,09 | 0 | 97,712 | 148 |
| 1,25E+08 | 1,38E+08 | 1,59E+08 | 1,57E+08 | 237,79 | 0 | 83,419 | 124 |
| 4,56E+08 | 3,49E+08 | 3,53E+08 | 3,40E+08 | 79,248 | 0 | 323,31 | 177 |
| 1,01E+08 | 1,17E+08 | 1,31E+08 | 1,37E+08 | 101,89 | 0 | 62,103 | 96 |
| 8,36E+07 | 7,83E+07 | 6,69E+07 | 6,82E+07 | 43,546 | 0 | 26,49 | 34 |
| 1,12E+08 | 1,56E+08 | 1,34E+08 | 1,77E+08 | 39,124 | 0 | 33,988 | 64 |
| 1,96E+08 | 2,59E+08 | 2,32E+08 | 2,64E+08 | 115,42 | 0 | 24,709 | 86 |
| 2,20E+08 | 1,89E+08 | 1,88E+08 | 2,00E+08 | 59,865 | 0 | 201,65 | 94 |
| 4,99E+07 | 1,75E+07 | 4,02E+06 | 1,25E+07 | 155,61 | 0 | 60,986 | 22 |

| | | | | | | | |
|---|---|---|---|---|---|---|---|
| 2,70E+07 | 2,04E+07 | 1,99E+07 | 1,29E+07 | 46,879 | 0,000158 | 5,0194 | 20 |
| 6,70E+07 | 5,89E+07 | 6,31E+07 | 5,92E+07 | 45,132 | 0 | 16,372 | 55 |
| 2,87E+07 | 3,13E+07 | 4,29E+07 | 4,48E+07 | 7,8648 | 0 | 10,131 | 18 |
| 1,45E+09 | 1,15E+09 | 1,03E+09 | 1,05E+09 | 13,527 | 0 | 323,31 | 201 |
| 4,75E+07 | 4,15E+07 | 3,92E+07 | 4,18E+07 | 69,96 | 0,003696 | 1,8195 | 8 |
| 5,25E+08 | 4,58E+08 | 4,25E+08 | 4,91E+08 | 38,113 | 0 | 216,15 | 126 |
| 6,23E+06 | 1,08E+07 | 1,39E+07 | 1,26E+07 | 58,877 | 0,000153 | 4,1701 | 9 |
| 7,04E+08 | 8,65E+08 | 9,65E+08 | 8,51E+08 | 73,101 | 0 | 323,31 | 127 |
| 3,86E+07 | 6,27E+07 | 6,42E+07 | 6,87E+07 | 258,13 | 0 | 56,682 | 57 |
| 3,10E+08 | 2,58E+08 | 2,85E+08 | 2,69E+08 | 95,675 | 0 | 134,86 | 169 |
| 1,75E+08 | 1,31E+08 | 1,34E+08 | 1,41E+08 | 65,229 | 0 | 96,44 | 92 |
| 1,22E+07 | 7,74E+06 | 1,00E+07 | 6,37E+06 | 64,679 | 0 | 5,9873 | 12 |
| 3,63E+07 | 5,13E+07 | 4,23E+07 | 4,55E+07 | 20,67 | 0 | 9,8354 | 24 |
| 3,62E+07 | 4,26E+07 | 4,02E+07 | 4,01E+07 | 20,082 | 0 | 8,4152 | 31 |
| 1,63E+08 | 1,99E+08 | 2,25E+08 | 2,58E+08 | 21,701 | 0 | 10,656 | 27 |
| 3,91E+07 | 5,29E+07 | 4,40E+07 | 4,66E+07 | 35,652 | 0 | 8,5023 | 26 |
| 6,84E+08 | 5,53E+08 | 6,28E+08 | 6,38E+08 | 59,374 | 0 | 268,65 | 147 |
| 4,98E+08 | 4,51E+08 | 4,65E+08 | 4,14E+08 | 80,222 | 0 | 242,06 | 171 |
| 1,59E+09 | 9,37E+08 | 1,06E+09 | 9,80E+08 | 44,593 | 0 | 323,31 | 184 |
| 5,78E+07 | 7,61E+07 | 6,25E+07 | 7,72E+07 | 35,942 | 0 | 177,88 | 49 |
| 4,20E+06 | 4,68E+06 | 5,91E+06 | 7,02E+06 | 69,855 | 0,000156 | 4,7744 | 7 |
| 9,47E+06 | 1,33E+07 | 1,52E+07 | 1,88E+07 | 165,87 | 0 | 6,8853 | 12 |
| 1,43E+08 | 1,31E+08 | 1,18E+08 | 1,24E+08 | 51,739 | 0 | 91,185 | 86 |
| 1,40E+07 | 9,89E+06 | 1,20E+07 | 1,03E+07 | 44,413 | 0,000436 | 2,9851 | 11 |
| 8,35E+06 | 1,72E+07 | 1,58E+07 | 1,72E+07 | 44,852 | 0 | 26,45 | 18 |
| 4,75E+09 | 3,88E+09 | 4,38E+09 | 4,15E+09 | 59,652 | 0 | 323,31 | 750 |
| 1,41E+07 | 2,14E+07 | 1,75E+07 | 1,67E+07 | 74,049 | 0 | 9,241 | 25 |
| 9,85E+06 | 1,29E+07 | 1,26E+07 | 1,91E+07 | 24,945 | 0 | 17,058 | 11 |
| 2,53E+07 | 4,32E+07 | 4,10E+07 | 4,10E+07 | 79,624 | 0 | 16,419 | 40 |
| 5,92E+06 | 7,65E+06 | 1,07E+07 | 1,10E+07 | 14,588 | 0,000435 | 2,952 | 7 |
| 1,78E+07 | 1,68E+07 | 1,47E+07 | 1,46E+07 | 94,5 | 0 | 22,311 | 23 |
| 1,24E+07 | 2,35E+06 | 7,04E+06 | 5,05E+06 | 76,461 | 0 | 10,769 | 7 |
| 3,03E+07 | 2,06E+07 | 2,02E+07 | 2,25E+07 | 46,217 | 0 | 19,806 | 25 |
| 1,51E+07 | 1,22E+07 | 1,25E+07 | 1,46E+07 | 22,983 | 0 | 6,662 | 17 |
| 2,95E+07 | 1,36E+07 | 1,52E+07 | 1,62E+07 | 34,465 | 0 | 9,5156 | 16 |
| 1,58E+07 | 5,41E+06 | 9,23E+06 | 9,24E+06 | 46,279 | 0 | 15,065 | 18 |
| 3,05E+06 | 5,72E+06 | 5,14E+06 | 5,09E+06 | 85,233 | 0,001685 | 2,271 | 5 |
| 4,50E+08 | 5,09E+08 | 5,01E+08 | 5,08E+08 | 126,82 | 0 | 215,49 | 227 |
| 1,80E+08 | 1,72E+08 | 1,56E+08 | 1,60E+08 | 19,294 | 0 | 40,793 | 60 |
| 2,70E+06 | 6,26E+06 | 9,09E+06 | 1,06E+07 | 270,51 | 0 | 5,4608 | 9 |
| 2,75E+07 | 3,34E+07 | 3,37E+07 | 2,86E+07 | 35,979 | 0 | 34,735 | 29 |
| 1,95E+08 | 2,33E+08 | 2,10E+08 | 2,10E+08 | 58,042 | 0 | 146,24 | 77 |
| 2,46E+08 | 2,77E+08 | 2,83E+08 | 2,69E+08 | 22,037 | 0 | 101,39 | 74 |
| 1,36E+08 | 1,73E+08 | 1,57E+08 | 1,85E+08 | 54,073 | 0 | 112,92 | 45 |
| 1,16E+07 | 2,34E+07 | 1,43E+07 | 1,98E+07 | 339,51 | 0,000153 | 4,0924 | 9 |
| 2,27E+08 | 1,80E+08 | 1,71E+08 | 1,49E+08 | 37,827 | 0 | 105,46 | 64 |
| 2,69E+08 | 3,40E+08 | 3,30E+08 | 3,74E+08 | 92,718 | 0 | 180,11 | 146 |
| 1,27E+08 | 1,13E+08 | 1,15E+08 | 1,22E+08 | 92,31 | 0 | 29,59 | 69 |
| 7,37E+07 | 5,41E+07 | 6,35E+07 | 6,89E+07 | 25,637 | 0 | 11,431 | 40 |
| 4,98E+07 | 9,06E+07 | 8,90E+07 | 8,60E+07 | 25,988 | 0 | 16,943 | 42 |

| | | | | | | | |
|---|---|---|---|---|---|---|---|
| 9,56E+06 | 2,26E+07 | 2,20E+07 | 2,48E+07 | 46,512 | 0 | 14,864 | 17 |
| 2,13E+07 | 4,37E+06 | 9,80E+06 | 9,04E+06 | 60,211 | 0 | 30,647 | 13 |
| 8,36E+06 | 4,46E+07 | 1,36E+07 | 1,89E+07 | 81,033 | 0 | 71,279 | 21 |
| 7,94E+07 | 6,65E+07 | 5,52E+07 | 6,15E+07 | 83,2 | 0 | 52,483 | 53 |
| 1,73E+08 | 1,46E+08 | 1,24E+08 | 1,50E+08 | 80,839 | 0 | 100,6 | 101 |
| 5,94E+07 | 6,90E+07 | 6,53E+07 | 7,55E+07 | 46,575 | 0 | 31,327 | 52 |
| 1,12E+10 | 1,17E+10 | 1,17E+10 | 1,25E+10 | 31,231 | 0 | 323,31 | 559 |
| 4,65E+08 | 3,59E+08 | 4,17E+08 | 4,31E+08 | 31,835 | 0 | 323,31 | 157 |
| 2,45E+07 | 1,01E+07 | 8,56E+06 | 1,52E+07 | 62,948 | 0 | 21,202 | 17 |
| 1,07E+08 | 1,30E+08 | 1,47E+08 | 1,37E+08 | 14,127 | 0 | 235,23 | 46 |
| 1,72E+07 | 9,40E+06 | 1,29E+07 | 1,41E+07 | 72,185 | 0,000147 | 3,1903 | 15 |
| 1,00E+07 | 1,26E+07 | 1,18E+07 | 1,24E+07 | 55,324 | 0,000154 | 4,4105 | 17 |
| 3,93E+07 | 5,29E+07 | 5,72E+07 | 5,69E+07 | 313,16 | 0 | 45,698 | 56 |
| 1,16E+10 | 8,27E+09 | 8,66E+09 | 1,00E+10 | 21,977 | 0 | 26,751 | 181 |
| 3,38E+06 | 2,98E+06 | 1,88E+06 | 2,13E+06 | 267,46 | 0,004366 | 1,7491 | 3 |
| 2,37E+08 | 1,94E+08 | 2,27E+08 | 2,08E+08 | 37,314 | 0 | 70,174 | 94 |
| 5,33E+09 | 4,64E+09 | 4,74E+09 | 4,96E+09 | 60,448 | 0 | 323,31 | 598 |
| 5,67E+07 | 2,30E+07 | 2,30E+07 | 1,08E+07 | 26,163 | 0 | 117,97 | 33 |
| 4,84E+07 | 2,91E+07 | 4,11E+07 | 3,79E+07 | 19,476 | 0 | 16,635 | 15 |
| 2,43E+09 | 3,29E+09 | 3,12E+09 | 3,21E+09 | 23,489 | 0 | 323,31 | 307 |
| 4,34E+06 | 2,30E+07 | 2,43E+07 | 2,72E+07 | 22,502 | 0 | 10,265 | 13 |
| 9,78E+07 | 7,86E+07 | 7,34E+07 | 8,97E+07 | 50,662 | 0 | 68,493 | 58 |
| 5,07E+07 | 7,49E+07 | 8,30E+07 | 7,49E+07 | 93,462 | 0 | 128,44 | 58 |
| 6,11E+07 | 3,70E+07 | 5,01E+07 | 4,21E+07 | 69,424 | 0 | 32,063 | 39 |
| 7,28E+07 | 8,57E+07 | 9,76E+07 | 1,02E+08 | 57,412 | 0 | 19,454 | 60 |
| 3,16E+08 | 3,89E+08 | 3,93E+08 | 4,24E+08 | 28,78 | 0 | 293,54 | 98 |
| 4,72E+06 | 2,05E+06 | 3,33E+06 | 2,22E+06 | 136,81 | 0,000153 | 4,1477 | 8 |
| 2,05E+08 | 3,35E+08 | 3,10E+08 | 3,67E+08 | 22,228 | 0 | 37,586 | 59 |
| 1,21E+07 | 1,93E+07 | 1,50E+07 | 1,48E+07 | 20,746 | 0,000152 | 3,9973 | 10 |
| 4,23E+08 | 3,77E+08 | 3,90E+08 | 3,73E+08 | 80,076 | 0 | 257,91 | 202 |
| 7,89E+06 | 4,31E+06 | 3,54E+06 | 6,98E+06 | 26,273 | 0 | 5,3283 | 11 |
| 1,13E+08 | 9,10E+07 | 9,38E+07 | 1,09E+08 | 32,989 | 0 | 27,243 | 46 |
| 2,50E+08 | 2,00E+08 | 2,25E+08 | 2,26E+08 | 76,912 | 0 | 103,41 | 130 |
| 1,35E+07 | 3,33E+07 | 3,33E+07 | 3,95E+07 | 44,812 | 0 | 36,636 | 21 |
| 2,39E+07 | 3,07E+07 | 4,38E+07 | 3,20E+07 | 17,107 | 0 | 25,901 | 17 |
| 1,13E+09 | 1,22E+09 | 1,16E+09 | 1,19E+09 | 117,77 | 0 | 249,17 | 250 |
| 1,70E+07 | 2,52E+07 | 2,85E+07 | 2,88E+07 | 57,225 | 0 | 7,2105 | 17 |
| 1,68E+08 | 1,21E+08 | 1,24E+08 | 1,35E+08 | 40,724 | 0 | 55,045 | 46 |
| 1,49E+06 | 5,17E+07 | 4,00E+07 | 2,50E+07 | 38,249 | 0 | 7,7474 | 20 |
| 1,07E+07 | 9,48E+06 | 6,19E+06 | 7,59E+06 | 98,405 | 0 | 7,5979 | 13 |
| 2,64E+08 | 3,89E+08 | 3,85E+08 | 4,17E+08 | 29,522 | 0 | 123,3 | 62 |
| 1,53E+06 | 2,24E+07 | 2,19E+07 | 2,33E+07 | 36,622 | 0 | 5,5673 | 9 |
| 1,21E+07 | 9,14E+06 | 7,99E+06 | 1,10E+07 | 37,066 | 0 | 16,084 | 9 |
| 8,17E+06 | 3,13E+07 | 1,94E+07 | 2,42E+07 | 8,0355 | 0,001548 | 2,3035 | 10 |
| 1,48E+09 | 1,43E+09 | 1,44E+09 | 1,34E+09 | 60,248 | 0 | 323,31 | 279 |
| 2,20E+07 | 1,25E+07 | 9,68E+06 | 1,35E+07 | 101,22 | 0 | 7,358 | 17 |
| 1,43E+08 | 1,28E+08 | 1,05E+08 | 1,21E+08 | 92,079 | 0 | 61,084 | 72 |
| 3,29E+06 | 4,25E+06 | 5,17E+06 | 3,91E+06 | 24,955 | 0,001822 | 2,2336 | 4 |
| 5,97E+07 | 4,02E+07 | 4,31E+07 | 5,27E+07 | 14,674 | 0 | 20,445 | 23 |
| 3,42E+07 | 5,73E+07 | 5,03E+07 | 5,46E+07 | 51,69 | 0 | 5,3285 | 13 |

| | | | | | | | |
|---|---|---|---|---|---|---|---|
| 2,28E+09 | 1,68E+09 | 1,77E+09 | 1,70E+09 | 134,19 | 0 | 323,31 | 441 |
| 1,88E+08 | 1,18E+08 | 1,25E+08 | 1,13E+08 | 32,733 | 0 | 323,31 | 67 |
| 4,59E+08 | 5,69E+08 | 4,88E+08 | 5,26E+08 | 28,463 | 0 | 104,53 | 153 |
| 2,09E+07 | 1,46E+07 | 1,24E+07 | 1,24E+07 | 19,285 | 0 | 10,15 | 15 |
| 3,76E+08 | 4,44E+08 | 4,08E+08 | 4,17E+08 | 187,19 | 0 | 117,49 | 243 |
| 1,31E+08 | 1,15E+08 | 1,18E+08 | 9,59E+07 | 43,088 | 0 | 58,689 | 54 |
| 6,25E+07 | 6,85E+07 | 6,70E+07 | 7,47E+07 | 20,156 | 0 | 63,756 | 39 |
| 5,54E+07 | 3,38E+07 | 3,40E+07 | 1,47E+07 | 45,23 | 0 | 43,632 | 33 |
| 4,82E+07 | 4,17E+07 | 4,11E+07 | 3,67E+07 | 87,337 | 0 | 45,412 | 53 |
| 4,87E+07 | 5,20E+07 | 5,49E+07 | 5,06E+07 | 88,047 | 0 | 31,776 | 72 |
| 4,35E+06 | 3,43E+06 | 3,67E+06 | 3,04E+06 | 122,76 | 0,009492 | 1,4221 | 11 |
| 6,87E+07 | 8,02E+07 | 8,27E+07 | 8,33E+07 | 79,375 | 0 | 126,16 | 65 |
| 4,40E+06 | 2,75E+06 | 1,11E+06 | 2,93E+06 | 14,283 | 0,00015 | 3,7033 | 4 |
| 1,47E+07 | 1,57E+06 | 7,12E+06 | 3,76E+06 | 23,396 | 0 | 7,314 | 11 |
| 4,57E+07 | 2,79E+07 | 3,51E+07 | 3,01E+07 | 38,107 | 0 | 149,54 | 36 |
| 2,39E+07 | 1,21E+07 | 5,27E+06 | 1,07E+07 | 158,46 | 0 | 17,275 | 19 |
| 8,61E+06 | 9,54E+06 | 1,54E+07 | 1,53E+07 | 17,813 | 0 | 40,54 | 9 |
| 6,58E+09 | 6,39E+09 | 6,10E+09 | 6,14E+09 | 60,629 | 0 | 323,31 | 728 |
| 7,39E+06 | 9,15E+06 | 8,66E+06 | 8,43E+06 | 36,632 | 0,00043 | 2,7566 | 12 |
| 9,74E+07 | 1,06E+08 | 1,24E+08 | 1,29E+08 | 37,309 | 0 | 21,308 | 55 |
| 9,75E+07 | 7,45E+07 | 7,51E+07 | 9,16E+07 | 73,104 | 0 | 37,186 | 48 |
| 9,57E+08 | 8,99E+08 | 8,97E+08 | 9,32E+08 | 26,24 | 0 | 162,21 | 130 |
| 3,65E+08 | 3,91E+08 | 4,24E+08 | 4,39E+08 | 52,287 | 0 | 125,59 | 112 |
| 6,64E+07 | 4,49E+07 | 4,77E+07 | 4,41E+07 | 56,613 | 0 | 55,815 | 42 |
| 7,80E+07 | 9,49E+07 | 8,16E+07 | 9,42E+07 | 76,684 | 0 | 25,625 | 56 |
| 2,04E+08 | 2,74E+08 | 2,93E+08 | 2,78E+08 | 46,434 | 0 | 124,98 | 82 |
| 2,30E+08 | 3,20E+08 | 3,31E+08 | 3,21E+08 | 49,279 | 0 | 323,31 | 70 |
| 1,49E+07 | 4,38E+06 | 4,18E+06 | 8,09E+06 | 27,397 | 0 | 121,51 | 11 |
| 1,36E+07 | 1,30E+07 | 1,03E+07 | 1,04E+07 | 87,676 | 0 | 11,424 | 30 |
| 3,52E+06 | 2,36E+07 | 1,78E+07 | 1,27E+07 | 155,27 | 0 | 9,3619 | 22 |
| 1,97E+09 | 1,32E+09 | 1,32E+09 | 1,03E+09 | 19,329 | 0 | 238,54 | 98 |
| 5,08E+08 | 5,53E+08 | 5,86E+08 | 5,42E+08 | 58,951 | 0 | 67,206 | 52 |
| 4,43E+07 | 9,71E+07 | 9,16E+07 | 1,01E+08 | 7,9972 | 0 | 54,502 | 27 |
| 6,34E+07 | 7,76E+07 | 6,70E+07 | 7,83E+07 | 85,652 | 0 | 23,22 | 61 |
| 5,73E+07 | 4,70E+07 | 5,14E+07 | 5,29E+07 | 44,772 | 0 | 15,579 | 38 |
| 1,33E+08 | 8,38E+07 | 7,64E+07 | 9,49E+07 | 103,17 | 0 | 18,935 | 42 |
| 1,06E+08 | 8,60E+07 | 9,88E+07 | 8,98E+07 | 46,132 | 0 | 66,705 | 78 |
| 3,67E+08 | 2,80E+08 | 2,99E+08 | 2,84E+08 | 26,151 | 0 | 148,29 | 115 |
| 3,40E+09 | 3,88E+09 | 4,28E+09 | 4,17E+09 | 48,1 | 0 | 323,31 | 322 |
| 5,94E+08 | 5,58E+08 | 5,19E+08 | 5,65E+08 | 106,17 | 0 | 323,31 | 194 |
| 2,23E+08 | 2,19E+08 | 2,13E+08 | 2,19E+08 | 167,14 | 0 | 106,32 | 155 |
| 5,20E+08 | 3,59E+08 | 4,87E+08 | 3,70E+08 | 55,302 | 0 | 302,45 | 106 |
| 1,05E+07 | 1,25E+07 | 1,20E+07 | 1,40E+07 | 68,966 | 0 | 6,986 | 19 |
| 3,40E+08 | 4,34E+08 | 4,82E+08 | 4,57E+08 | 12,473 | 0 | 285,95 | 65 |
| 1,30E+07 | 6,76E+06 | 4,87E+06 | 7,75E+06 | 21,656 | 0 | 11,812 | 14 |
| 1,27E+07 | 2,81E+06 | 4,97E+06 | 4,67E+06 | 178,88 | 0,000147 | 3,1468 | 6 |
| 9,37E+07 | 9,91E+07 | 1,11E+08 | 1,11E+08 | 46,553 | 0 | 47,696 | 53 |
| 1,99E+07 | 2,71E+07 | 2,54E+07 | 2,63E+07 | 163,45 | 0 | 16,867 | 36 |
| 5,20E+07 | 8,52E+07 | 8,18E+07 | 8,30E+07 | 47,972 | 0 | 39,134 | 54 |
| 1,70E+07 | 2,29E+07 | 2,64E+07 | 2,89E+07 | 42,119 | 0 | 8,283 | 16 |

| | | | | | | | |
|---|---|---|---|---|---|---|---|
| 8,52E+08 | 4,14E+08 | 3,93E+08 | 3,59E+08 | 37,27 | 0 | 323,31 | 119 |
| 3,72E+07 | 3,15E+07 | 2,91E+07 | 3,40E+07 | 35,764 | 0 | 20,111 | 46 |
| 2,99E+07 | 3,23E+07 | 3,35E+07 | 3,90E+07 | 19,003 | 0 | 24,425 | 31 |
| 4,12E+08 | 6,60E+08 | 6,58E+08 | 5,37E+08 | 58,886 | 0 | 323,31 | 123 |
| 1,37E+08 | 1,15E+08 | 1,26E+08 | 1,29E+08 | 201,57 | 0 | 323,31 | 134 |
| 5,34E+08 | 4,93E+08 | 4,92E+08 | 5,25E+08 | 132,39 | 0 | 190,19 | 237 |
| 1,06E+08 | 1,02E+08 | 8,67E+07 | 9,66E+07 | 161 | 0 | 69,877 | 94 |
| 3,11E+08 | 4,99E+08 | 4,65E+08 | 4,46E+08 | 71,513 | 0 | 136,85 | 179 |
| 9,26E+08 | 9,54E+08 | 9,92E+08 | 1,00E+09 | 133,57 | 0 | 323,31 | 275 |
| 1,21E+07 | 3,40E+06 | 3,18E+06 | 1,00E+07 | 39,992 | 0 | 5,7242 | 9 |
| 1,29E+07 | 9,83E+06 | 2,62E+06 | 3,78E+06 | 87,87 | 0 | 9,7367 | 14 |
| 4,85E+08 | 7,42E+08 | 7,88E+08 | 7,09E+08 | 21,557 | 0 | 310,38 | 95 |
| 5,22E+07 | 6,10E+07 | 5,76E+07 | 7,83E+07 | 41,866 | 0 | 23,197 | 28 |
| 3,41E+07 | 2,18E+07 | 2,64E+07 | 2,58E+07 | 103,73 | 0 | 13,59 | 36 |
| 6,39E+06 | 3,43E+06 | 1,76E+06 | 2,74E+06 | 50,255 | 0 | 6,3744 | 6 |
| 4,43E+07 | 5,36E+07 | 5,79E+07 | 5,37E+07 | 63,769 | 0 | 14,034 | 39 |
| 6,29E+08 | 8,67E+08 | 8,16E+08 | 8,03E+08 | 44,116 | 0 | 323,31 | 206 |
| 1,32E+07 | 9,18E+06 | 6,62E+06 | 8,19E+06 | 90,726 | 0 | 12,576 | 13 |
| 9,43E+06 | 8,30E+06 | 6,06E+06 | 7,58E+06 | 68,582 | 0,000154 | 4,3671 | 14 |
| 5,97E+07 | 4,26E+07 | 4,45E+07 | 5,54E+07 | 48,58 | 0 | 58,98 | 33 |
| 2,83E+08 | 1,88E+08 | 2,37E+08 | 2,62E+08 | 38,241 | 0 | 193,5 | 61 |
| 4,39E+06 | 2,12E+06 | 2,64E+06 | 3,64E+06 | 59,77 | 0,000431 | 2,7661 | 6 |
| 3,22E+08 | 2,93E+08 | 3,14E+08 | 3,06E+08 | 105,21 | 0 | 250,37 | 181 |
| 9,87E+06 | 1,74E+07 | 1,78E+07 | 1,25E+07 | 47,416 | 0 | 6,6588 | 19 |
| 2,40E+08 | 2,07E+08 | 2,05E+08 | 2,08E+08 | 42,636 | 0 | 158,35 | 104 |
| 1,35E+08 | 1,21E+08 | 1,17E+08 | 1,12E+08 | 55,66 | 0 | 218,15 | 72 |
| 2,41E+07 | 1,25E+07 | 1,15E+07 | 6,26E+06 | 113,55 | 0 | 7,5721 | 25 |
| 3,78E+07 | 5,62E+07 | 6,63E+07 | 5,33E+07 | 55,741 | 0 | 10,7 | 33 |
| 3,32E+09 | 2,71E+09 | 2,66E+09 | 2,65E+09 | 71,982 | 0 | 323,31 | 505 |
| 3,58E+08 | 2,96E+08 | 2,90E+08 | 2,91E+08 | 165,13 | 0 | 288,91 | 191 |
| 1,84E+07 | 9,07E+06 | 8,87E+06 | 1,95E+06 | 60,521 | 0,000152 | 4,0663 | 18 |
| 2,65E+10 | 1,86E+10 | 2,00E+10 | 1,77E+10 | 83,28 | 0 | 323,31 | 1226 |
| 9,60E+06 | 1,32E+07 | 1,08E+07 | 1,59E+07 | 108,22 | 0 | 7,9171 | 17 |
| 3,82E+08 | 2,21E+08 | 2,18E+08 | 2,18E+08 | 60,273 | 0 | 228,64 | 104 |
| 2,20E+09 | 2,02E+09 | 2,00E+09 | 2,02E+09 | 50,213 | 0 | 323,31 | 369 |
| 1,09E+09 | 1,18E+09 | 1,25E+09 | 1,31E+09 | 155,66 | 0 | 323,31 | 441 |
| 5,18E+07 | 4,38E+07 | 4,98E+07 | 4,56E+07 | 31,739 | 0 | 50,763 | 43 |
| 1,89E+07 | 1,37E+07 | 1,16E+07 | 1,23E+07 | 72,62 | 0 | 20,086 | 18 |
| 1,61E+07 | 2,86E+07 | 2,82E+07 | 2,72E+07 | 9,3308 | 0 | 6,8126 | 13 |
| 3,03E+07 | 2,60E+06 | 2,73E+06 | 3,45E+06 | 85,841 | 0 | 7,388 | 15 |
| 8,29E+08 | 1,02E+09 | 9,32E+08 | 9,32E+08 | 39,832 | 0 | 323,31 | 247 |
| 3,19E+08 | 3,25E+08 | 3,97E+08 | 3,80E+08 | 61,422 | 0 | 57,502 | 109 |
| 2,17E+07 | 9,74E+06 | 1,81E+07 | 1,58E+07 | 101,28 | 0 | 8,6363 | 23 |
| 3,26E+08 | 2,98E+08 | 2,61E+08 | 2,66E+08 | 29,473 | 0 | 191,79 | 86 |
| 3,21E+07 | 2,95E+07 | 2,58E+07 | 2,11E+07 | 20,667 | 0 | 21,507 | 49 |
| 6,18E+06 | 3,52E+06 | 3,03E+06 | 4,37E+06 | 49,299 | 0,000433 | 2,8821 | 7 |
| 1,89E+08 | 1,41E+08 | 1,44E+08 | 1,49E+08 | 143,64 | 0 | 88,608 | 115 |
| 3,84E+07 | 2,60E+07 | 1,74E+07 | 2,67E+07 | 126,26 | 0 | 10,429 | 36 |
| 2,44E+07 | 2,68E+07 | 3,26E+07 | 3,49E+07 | 43,823 | 0 | 7,4431 | 21 |
| 3,88E+09 | 3,64E+09 | 3,72E+09 | 3,70E+09 | 50,537 | 0 | 323,31 | 535 |

| | | | | | | | |
|---|---|---|---|---|---|---|---|
| 1,18E+09 | 1,09E+09 | 1,14E+09 | 1,06E+09 | 108,69 | 0 | 323,31 | 338 |
| 9,75E+07 | 1,42E+08 | 1,16E+08 | 1,30E+08 | 54,257 | 0 | 50,995 | 86 |
| 3,13E+08 | 3,02E+08 | 2,94E+08 | 2,66E+08 | 113,28 | 0 | 153,55 | 124 |
| 3,43E+06 | 1,66E+07 | 1,24E+07 | 1,53E+07 | 36,588 | 0 | 9,0318 | 17 |
| 1,48E+07 | 2,45E+07 | 2,39E+07 | 3,08E+07 | 30,016 | 0 | 6,4305 | 11 |
| 8,84E+07 | 6,33E+07 | 7,12E+07 | 4,82E+07 | 15,674 | 0 | 35,078 | 49 |
| 1,84E+07 | 2,37E+07 | 1,90E+07 | 2,11E+07 | 146,71 | 0 | 14,649 | 30 |
| 1,76E+08 | 1,38E+08 | 1,19E+08 | 1,25E+08 | 53,542 | 0 | 250,72 | 87 |
| 8,60E+08 | 6,47E+08 | 6,12E+08 | 6,61E+08 | 38,358 | 0 | 112,97 | 178 |
| 2,31E+08 | 1,65E+08 | 1,86E+08 | 2,20E+08 | 22,101 | 0 | 323,31 | 78 |
| 3,82E+08 | 2,97E+08 | 3,37E+08 | 3,20E+08 | 13,077 | 0 | 210,11 | 70 |
| 3,40E+08 | 4,70E+08 | 4,95E+08 | 5,06E+08 | 100,15 | 0 | 323,31 | 108 |
| 3,12E+10 | 3,59E+10 | 3,53E+10 | 3,70E+10 | 74,237 | 0 | 323,31 | 1789 |
| 1,73E+08 | 2,12E+08 | 2,21E+08 | 2,07E+08 | 44,688 | 0 | 66,926 | 107 |
| 4,39E+07 | 6,84E+07 | 7,06E+07 | 6,49E+07 | 128,66 | 0 | 36,128 | 56 |
| 4,64E+08 | 6,16E+08 | 6,41E+08 | 5,67E+08 | 22,137 | 0 | 41,648 | 99 |
| 7,16E+08 | 9,20E+08 | 8,61E+08 | 7,62E+08 | 42,679 | 0 | 323,31 | 170 |
| 2,79E+07 | 4,08E+07 | 5,27E+07 | 5,74E+07 | 30,202 | 0 | 8,2592 | 19 |
| 3,08E+08 | 4,14E+08 | 3,72E+08 | 3,96E+08 | 35,08 | 0 | 127,99 | 127 |
| 1,77E+07 | 1,60E+07 | 1,35E+07 | 1,20E+07 | 87,142 | 0 | 9,4962 | 28 |
| 1,19E+07 | 8,25E+06 | 1,05E+07 | 1,02E+07 | 75,042 | 0 | 21,04 | 19 |
| 1,08E+07 | 4,35E+06 | 6,59E+06 | 1,37E+06 | 63,732 | 0,001275 | 2,4691 | 5 |
| 1,73E+08 | 1,68E+08 | 1,55E+08 | 1,58E+08 | 28,093 | 0 | 50,787 | 74 |
| 4,45E+08 | 4,73E+08 | 4,76E+08 | 4,47E+08 | 218,13 | 0 | 323,31 | 235 |
| 4,35E+08 | 3,64E+08 | 3,63E+08 | 4,30E+08 | 80,817 | 0 | 200,45 | 129 |
| 2,61E+08 | 2,25E+08 | 2,07E+08 | 2,17E+08 | 39,106 | 0 | 167,86 | 59 |
| 3,35E+08 | 3,47E+08 | 3,79E+08 | 3,50E+08 | 46,478 | 0 | 323,31 | 111 |
| 4,94E+07 | 6,42E+07 | 5,94E+07 | 5,58E+07 | 74,331 | 0 | 16,757 | 44 |
| 6,55E+08 | 7,55E+08 | 7,50E+08 | 7,73E+08 | 116,86 | 0 | 323,31 | 289 |
| 8,03E+07 | 7,54E+07 | 6,10E+07 | 6,19E+07 | 68,927 | 0 | 16,625 | 45 |
| 4,73E+06 | 3,02E+07 | 2,83E+07 | 2,56E+07 | 189,41 | 0 | 7,6026 | 21 |
| 1,54E+08 | 1,45E+08 | 1,47E+08 | 1,49E+08 | 52,443 | 0 | 31,352 | 68 |
| 7,42E+07 | 2,28E+07 | 1,62E+07 | 3,44E+06 | 194,29 | 0 | 39,218 | 39 |
| 7,76E+06 | 6,70E+06 | 4,75E+06 | 2,76E+06 | 24,677 | 0,00015 | 3,7382 | 9 |
| 3,88E+07 | 2,59E+07 | 2,90E+07 | 2,07E+07 | 16,32 | 0 | 16,072 | 31 |
| 5,33E+06 | 1,02E+07 | 1,03E+07 | 5,52E+06 | 33,607 | 0 | 5,6849 | 5 |
| 2,41E+07 | 2,90E+07 | 2,74E+07 | 2,86E+07 | 292,12 | 0,000147 | 3,1697 | 15 |
| 8,65E+06 | 5,72E+06 | 6,69E+06 | 7,06E+06 | 13,641 | 0 | 17,836 | 9 |
| 8,90E+08 | 6,24E+08 | 6,44E+08 | 6,16E+08 | 77,454 | 0 | 186,35 | 230 |
| 6,08E+07 | 1,08E+08 | 8,71E+07 | 9,05E+07 | 19,431 | 0 | 13,158 | 48 |
| 1,11E+08 | 1,28E+08 | 1,26E+08 | 1,31E+08 | 72,057 | 0 | 115,77 | 78 |
| 2,95E+06 | 9,73E+06 | 7,78E+06 | 6,27E+06 | 37,437 | 0,000151 | 3,7783 | 8 |
| 8,72E+07 | 9,47E+07 | 1,05E+08 | 9,56E+07 | 77,785 | 0 | 44,72 | 69 |
| 1,18E+09 | 1,79E+09 | 1,82E+09 | 1,90E+09 | 34,741 | 0 | 134,46 | 150 |
| 8,84E+08 | 8,05E+08 | 7,95E+08 | 7,92E+08 | 61,475 | 0 | 323,31 | 198 |
| 1,69E+07 | 1,66E+06 | 9,00E+05 | 5,26E+06 | 182,36 | 0 | 30,13 | 10 |
| 3,81E+09 | 4,72E+09 | 4,68E+09 | 4,72E+09 | 85,462 | 0 | 323,31 | 411 |
| 4,49E+08 | 3,29E+08 | 3,18E+08 | 3,59E+08 | 77,06 | 0 | 274,58 | 120 |
| 2,32E+07 | 6,12E+07 | 4,58E+07 | 5,44E+07 | 84,684 | 0 | 18,45 | 41 |
| 1,70E+07 | 1,23E+07 | 1,57E+07 | 1,19E+07 | 254,47 | 0,000154 | 4,3462 | 15 |

| | | | | | | | |
|---|---|---|---|---|---|---|---|
| 2,59E+07 | 2,32E+07 | 2,28E+07 | 2,17E+07 | 186,48 | 0 | 7,7357 | 28 |
| 5,71E+06 | 1,53E+07 | 1,46E+07 | 1,42E+07 | 66,062 | 0 | 9,8804 | 18 |
| 4,85E+06 | 8,42E+06 | 6,96E+06 | 7,87E+06 | 27,333 | 0,000992 | 2,4794 | 6 |
| 3,89E+07 | 5,02E+07 | 5,38E+07 | 4,78E+07 | 80,504 | 0 | 17,711 | 49 |
| 1,63E+07 | 2,79E+07 | 2,10E+07 | 2,27E+07 | 66,739 | 0 | 9,6782 | 22 |
| 5,55E+06 | 3,44E+07 | 2,89E+07 | 3,42E+07 | 77,795 | 0 | 35,374 | 31 |
| 2,57E+07 | 3,19E+07 | 2,89E+07 | 2,63E+07 | 51,957 | 0 | 10,379 | 20 |
| 4,74E+06 | 4,22E+06 | 2,49E+06 | 2,74E+06 | 39,532 | 0 | 7,7062 | 10 |
| 2,10E+06 | 4,08E+06 | 3,05E+06 | 4,60E+06 | 70,799 | 0,000146 | 3,1076 | 5 |
| 5,17E+07 | 4,53E+07 | 4,58E+07 | 3,52E+07 | 34,798 | 0 | 40,829 | 23 |
| 9,82E+07 | 8,19E+07 | 8,21E+07 | 8,36E+07 | 60,501 | 0 | 25,949 | 58 |
| 1,24E+07 | 1,67E+07 | 1,84E+07 | 1,86E+07 | 26,738 | 0 | 17,139 | 7 |
| 1,58E+08 | 1,23E+08 | 1,20E+08 | 1,09E+08 | 56,536 | 0 | 64,65 | 81 |
| 6,25E+06 | 5,26E+06 | 1,70E+06 | 3,25E+06 | 129,72 | 0 | 5,7359 | 11 |
| 1,44E+07 | 1,80E+07 | 1,76E+07 | 2,15E+07 | 135,12 | 0 | 11,834 | 24 |
| 4,87E+09 | 5,27E+09 | 5,30E+09 | 5,32E+09 | 97,183 | 0 | 323,31 | 720 |
| 1,90E+08 | 1,77E+08 | 1,56E+08 | 1,49E+08 | 85,291 | 0 | 136,4 | 122 |
| 2,94E+09 | 2,27E+09 | 2,49E+09 | 2,44E+09 | 27,744 | 0 | 118,52 | 419 |
| 2,20E+08 | 4,05E+08 | 3,20E+08 | 3,57E+08 | 39,842 | 0 | 26,636 | 89 |
| 1,79E+08 | 1,63E+08 | 1,69E+08 | 1,76E+08 | 429,64 | 0 | 121,97 | 140 |
| 1,19E+07 | 2,28E+07 | 1,76E+07 | 3,28E+07 | 70,505 | 0 | 5,7577 | 11 |
| 6,07E+07 | 4,34E+07 | 5,61E+07 | 5,35E+07 | 119,27 | 0 | 31,159 | 56 |
| 2,22E+07 | 3,81E+06 | 6,82E+05 | 8,42E+06 | 11,402 | 0,000157 | 4,8959 | 6 |
| 5,38E+07 | 3,93E+07 | 3,83E+07 | 3,76E+07 | 263,3 | 0 | 185,13 | 54 |
| 3,43E+08 | 3,05E+08 | 2,76E+08 | 3,18E+08 | 41,87 | 0 | 106,72 | 93 |
| 6,57E+06 | 2,13E+07 | 2,07E+07 | 2,14E+07 | 180,2 | 0 | 9,6886 | 28 |
| 2,89E+07 | 2,47E+07 | 2,69E+07 | 2,63E+07 | 77,972 | 0 | 23,939 | 40 |
| 2,47E+08 | 2,71E+08 | 2,58E+08 | 2,55E+08 | 238,83 | 0 | 172,04 | 210 |
| 2,61E+06 | 3,72E+06 | 3,04E+06 | 5,15E+06 | 63,898 | 0,003962 | 1,7819 | 4 |
| 6,79E+07 | 8,72E+07 | 8,12E+07 | 9,41E+07 | 59,146 | 0 | 43,449 | 47 |
| 5,83E+07 | 4,18E+07 | 2,72E+07 | 3,33E+07 | 29,628 | 0 | 14,157 | 25 |
| 1,59E+07 | 1,54E+06 | 4,28E+06 | 8,91E+06 | 54,324 | 0 | 8,6652 | 10 |
| 1,02E+07 | 3,11E+07 | 1,92E+07 | 2,10E+07 | 146,75 | 0 | 53,288 | 9 |
| 7,87E+07 | 1,15E+08 | 1,14E+08 | 9,28E+07 | 39,493 | 0 | 74,8 | 47 |
| 3,74E+07 | 4,16E+07 | 4,36E+07 | 3,87E+07 | 46,276 | 0 | 15,145 | 35 |
| 2,74E+07 | 4,57E+07 | 3,64E+07 | 3,55E+07 | 47,24 | 0 | 39,893 | 47 |
| 1,38E+08 | 9,75E+07 | 9,90E+07 | 1,06E+08 | 86,305 | 0 | 80,879 | 80 |
| 3,57E+07 | 2,62E+06 | 1,49E+07 | 1,31E+07 | 65,475 | 0 | 28,105 | 27 |
| 5,20E+07 | 8,48E+07 | 9,29E+07 | 9,05E+07 | 204,65 | 0 | 29,203 | 77 |
| 1,41E+08 | 1,25E+08 | 1,24E+08 | 1,31E+08 | 64,667 | 0 | 95,547 | 97 |
| 9,40E+08 | 7,28E+08 | 8,72E+08 | 7,79E+08 | 53,12 | 0 | 323,31 | 176 |
| 8,97E+06 | 1,48E+06 | 1,24E+06 | 5,35E+06 | 29,218 | 0 | 24,514 | 6 |
| 1,65E+08 | 9,18E+07 | 6,42E+07 | 1,18E+08 | 53,762 | 0 | 37,086 | 18 |
| 3,89E+06 | 8,85E+06 | 1,26E+07 | 1,61E+07 | 140,37 | 0 | 9,7383 | 13 |
| 6,36E+07 | 9,10E+07 | 1,06E+08 | 1,04E+08 | 103,01 | 0 | 63,259 | 89 |
| 1,39E+07 | 1,12E+07 | 1,31E+07 | 1,14E+07 | 70,932 | 0 | 9,5014 | 24 |
| 1,21E+07 | 3,08E+07 | 4,03E+07 | 7,12E+07 | 26,851 | 0 | 15,621 | 19 |
| 6,11E+07 | 8,10E+07 | 8,26E+07 | 8,88E+07 | 15,525 | 0 | 57,167 | 25 |
| 1,64E+08 | 1,89E+08 | 1,85E+08 | 1,69E+08 | 53,398 | 0 | 316,61 | 106 |
| 4,44E+07 | 9,83E+06 | 2,32E+07 | 2,98E+07 | 191,49 | 0 | 24,073 | 41 |

| | | | | | | | |
|---|---|---|---|---|---|---|---|
| 1,56E+08 | 8,07E+07 | 8,42E+07 | 7,99E+07 | 25,42 | 0 | 222,4 | 48 |
| 2,72E+07 | 3,26E+07 | 3,84E+07 | 4,09E+07 | 12,228 | 0 | 61,963 | 25 |
| 2,48E+08 | 1,95E+08 | 1,97E+08 | 2,20E+08 | 56,471 | 0 | 323,31 | 71 |
| 1,12E+09 | 1,20E+09 | 1,28E+09 | 1,28E+09 | 82,545 | 0 | 323,31 | 283 |
| 2,22E+07 | 1,83E+07 | 2,07E+07 | 1,98E+07 | 59,505 | 0 | 20,838 | 26 |
| 4,04E+08 | 4,85E+08 | 5,75E+08 | 5,64E+08 | 12,097 | 0 | 94,647 | 88 |
| 5,17E+08 | 4,90E+08 | 4,85E+08 | 4,62E+08 | 94,036 | 0 | 323,31 | 223 |
| 2,65E+08 | 4,77E+08 | 4,45E+08 | 4,90E+08 | 104,17 | 0 | 164,32 | 157 |
| 1,40E+08 | 1,29E+08 | 1,37E+08 | 1,25E+08 | 24,575 | 0 | 75,064 | 53 |
| 1,45E+10 | 1,56E+10 | 1,69E+10 | 1,67E+10 | 56,678 | 0 | 323,31 | 1058 |
| 1,32E+08 | 9,23E+07 | 8,34E+07 | 9,48E+07 | 96,134 | 0 | 26,083 | 47 |
| 1,98E+07 | 1,72E+07 | 1,54E+07 | 1,92E+07 | 37,516 | 0 | 12,922 | 20 |
| 9,69E+07 | 1,08E+08 | 1,29E+08 | 1,33E+08 | 74,29 | 0 | 48,009 | 74 |
| 3,53E+08 | 2,61E+08 | 2,53E+08 | 3,39E+08 | 16,08 | 0 | 186,11 | 72 |
| 1,69E+07 | 2,45E+07 | 3,01E+07 | 2,11E+07 | 36,127 | 0 | 8,8233 | 15 |
| 5,20E+07 | 5,93E+07 | 5,80E+07 | 5,25E+07 | 54,815 | 0 | 136,76 | 62 |
| 2,45E+09 | 2,85E+09 | 2,76E+09 | 2,83E+09 | 47,756 | 0 | 323,31 | 335 |
| 5,11E+07 | 4,03E+07 | 3,78E+07 | 3,44E+07 | 14,92 | 0 | 33,248 | 37 |
| 1,29E+07 | 8,27E+06 | 8,23E+06 | 3,97E+06 | 89,559 | 0 | 9,7733 | 12 |
| 5,15E+07 | 7,99E+07 | 7,59E+07 | 7,45E+07 | 14,617 | 0,000149 | 3,4922 | 17 |
| 2,45E+07 | 2,60E+07 | 2,77E+07 | 2,62E+07 | 203,15 | 0 | 37,437 | 36 |
| 1,44E+07 | 2,05E+07 | 1,99E+07 | 2,56E+07 | 49,979 | 0 | 12,584 | 20 |
| 3,12E+07 | 4,11E+07 | 3,94E+07 | 3,29E+07 | 22,812 | 0 | 13,01 | 33 |
| 3,25E+07 | 4,75E+07 | 5,53E+07 | 6,29E+07 | 20,046 | 0 | 323,31 | 60 |
| 4,58E+06 | 6,84E+06 | 9,35E+06 | 1,09E+07 | 20,923 | 0,000148 | 3,3611 | 6 |
| 5,95E+07 | 7,24E+07 | 1,20E+08 | 1,01E+08 | 9,6418 | 0 | 85,117 | 28 |
| 2,49E+08 | 2,69E+08 | 2,83E+08 | 2,75E+08 | 14,679 | 0 | 323,31 | 95 |
| 6,22E+06 | 1,27E+07 | 1,37E+07 | 8,94E+06 | 19,209 | 0 | 5,3438 | 13 |
| 1,24E+09 | 1,03E+09 | 1,17E+09 | 1,14E+09 | 30,817 | 0 | 120,08 | 123 |
| 1,68E+08 | 1,98E+08 | 1,71E+08 | 1,91E+08 | 435,87 | 0 | 114,77 | 213 |
| 2,83E+07 | 2,69E+07 | 2,58E+07 | 2,46E+07 | 96,789 | 0 | 31,32 | 29 |
| 1,73E+07 | 7,14E+06 | 1,10E+07 | 1,14E+07 | 118,56 | 0,000153 | 4,2211 | 10 |
| 1,49E+07 | 2,92E+07 | 3,57E+07 | 4,68E+07 | 25,465 | 0 | 9,6791 | 20 |
| 4,01E+06 | 2,15E+06 | 1,79E+06 | 2,84E+06 | 45,094 | 0,000146 | 3,1121 | 6 |
| 1,10E+09 | 1,22E+09 | 1,33E+09 | 1,30E+09 | 42,536 | 0 | 259,93 | 215 |
| 5,90E+07 | 6,14E+07 | 7,44E+07 | 7,47E+07 | 80,731 | 0 | 323,31 | 21 |
| 1,06E+09 | 9,00E+08 | 9,95E+08 | 1,00E+09 | 99,551 | 0 | 321,81 | 171 |
| 1,52E+09 | 1,64E+09 | 1,76E+09 | 1,55E+09 | 42,809 | 0 | 323,31 | 268 |
| 4,36E+07 | 2,80E+07 | 3,09E+07 | 3,76E+07 | 22,659 | 0 | 31,354 | 40 |
| 1,86E+06 | 6,92E+06 | 8,77E+06 | 5,35E+06 | 48,769 | 0,002092 | 2,1148 | 5 |
| 3,11E+08 | 2,57E+08 | 2,36E+08 | 2,40E+08 | 101,93 | 0 | 323,31 | 138 |
| 2,51E+08 | 2,22E+08 | 2,02E+08 | 2,24E+08 | 43,537 | 0 | 122,3 | 93 |
| 6,95E+07 | 9,73E+07 | 9,46E+07 | 1,10E+08 | 51,69 | 0 | 74,425 | 58 |
| 5,26E+07 | 6,77E+07 | 5,98E+07 | 6,34E+07 | 134,94 | 0 | 20,068 | 64 |
| 2,87E+07 | 4,29E+07 | 4,31E+07 | 3,62E+07 | 35,612 | 0 | 44,33 | 39 |
| 9,34E+06 | 2,53E+06 | 2,23E+06 | 4,94E+06 | 66,487 | 0 | 5,5917 | 7 |
| 6,36E+07 | 5,60E+07 | 6,21E+07 | 5,56E+07 | 24,725 | 0 | 7,9143 | 23 |
| 7,55E+07 | 1,16E+08 | 1,19E+08 | 1,13E+08 | 50,216 | 0 | 90,325 | 77 |
| 9,43E+07 | 4,15E+07 | 4,06E+07 | 3,20E+07 | 85,669 | 0 | 198,06 | 31 |
| 2,82E+09 | 2,50E+09 | 2,48E+09 | 2,76E+09 | 32,667 | 0 | 323,31 | 339 |

| | | | | | | | |
|---|---|---|---|---|---|---|---|
| 6,21E+07 | 4,46E+07 | 4,44E+07 | 4,63E+07 | 94,525 | 0 | 35,511 | 64 |
| 1,67E+08 | 1,35E+08 | 1,54E+08 | 1,52E+08 | 117,33 | 0 | 124,04 | 130 |
| 4,16E+06 | 9,16E+06 | 9,41E+06 | 6,71E+06 | 9,2858 | 0,000146 | 3,1199 | 5 |
| 1,26E+08 | 1,69E+08 | 1,48E+08 | 1,40E+08 | 43,323 | 0 | 125,25 | 94 |
| 8,02E+07 | 7,81E+07 | 7,86E+07 | 7,69E+07 | 238,59 | 0 | 55,802 | 101 |
| 7,99E+06 | 1,16E+07 | 1,57E+07 | 1,46E+07 | 79,853 | 0 | 13,217 | 14 |
| 3,68E+07 | 4,12E+07 | 3,71E+07 | 4,05E+07 | 24,001 | 0 | 46,531 | 21 |
| 1,32E+09 | 1,81E+09 | 2,00E+09 | 1,91E+09 | 23,609 | 0 | 323,31 | 179 |
| 1,96E+07 | 1,81E+07 | 1,52E+07 | 1,77E+07 | 199,13 | 0 | 7,8475 | 20 |
| 2,30E+08 | 2,09E+08 | 1,85E+08 | 2,23E+08 | 26,753 | 0 | 240,42 | 38 |
| 4,18E+07 | 8,98E+07 | 8,62E+07 | 8,28E+07 | 46,244 | 0 | 29,269 | 46 |
| 5,46E+09 | 5,45E+09 | 4,87E+09 | 4,94E+09 | 46,533 | 0 | 323,31 | 632 |
| 2,37E+08 | 1,63E+08 | 1,50E+08 | 1,78E+08 | 11,158 | 0 | 18,652 | 32 |
| 3,66E+07 | 4,48E+07 | 4,04E+07 | 3,81E+07 | 38,817 | 0 | 18,31 | 45 |
| 3,08E+08 | 1,83E+08 | 1,41E+08 | 1,54E+08 | 105,69 | 0 | 66,678 | 73 |
| 1,75E+08 | 1,65E+08 | 1,47E+08 | 1,36E+08 | 17 | 0 | 54,928 | 69 |
| 8,23E+07 | 9,57E+07 | 1,04E+08 | 1,02E+08 | 36,938 | 0 | 42,165 | 50 |
| 2,23E+07 | 3,18E+07 | 2,50E+07 | 3,05E+07 | 41,626 | 0 | 10,867 | 31 |
| 1,06E+08 | 8,89E+07 | 9,89E+07 | 9,49E+07 | 33,288 | 0 | 28,144 | 66 |
| 1,01E+07 | 5,36E+06 | 5,28E+06 | 1,00E+07 | 45,17 | 0 | 11,967 | 17 |
| 2,28E+07 | 1,17E+08 | 1,03E+08 | 1,06E+08 | 99,098 | 0 | 50,486 | 64 |
| 2,48E+06 | 1,63E+07 | 1,16E+07 | 1,11E+07 | 18,232 | 0 | 5,8912 | 9 |
| 2,90E+07 | 3,96E+07 | 4,27E+07 | 3,84E+07 | 127,89 | 0 | 19,631 | 49 |
| 1,12E+07 | 1,28E+07 | 1,21E+07 | 1,49E+07 | 12,325 | 0 | 8,6509 | 7 |
| 3,78E+06 | 1,36E+07 | 1,60E+07 | 1,64E+07 | 31,992 | 0,001548 | 2,3154 | 6 |
| 1,47E+07 | 1,45E+07 | 1,25E+07 | 1,11E+07 | 21,494 | 0,000151 | 3,8041 | 12 |
| 1,99E+07 | 1,56E+07 | 1,43E+07 | 1,14E+07 | 56,242 | 0 | 8,7241 | 16 |
| 9,35E+06 | 2,89E+06 | 6,67E+06 | 5,63E+06 | 136,54 | 0,000155 | 4,4913 | 5 |
| 1,09E+08 | 9,24E+07 | 1,06E+08 | 1,01E+08 | 68,97 | 0 | 105,81 | 62 |
| 2,36E+09 | 2,12E+09 | 2,06E+09 | 2,03E+09 | 176,07 | 0 | 323,31 | 393 |
| 7,95E+06 | 8,33E+06 | 9,72E+06 | 8,76E+06 | 54,174 | 0 | 5,9534 | 13 |
| 2,13E+08 | 2,00E+08 | 1,70E+08 | 1,88E+08 | 155,89 | 0 | 109,73 | 150 |
| 3,72E+07 | 2,76E+07 | 2,69E+07 | 3,15E+07 | 24,416 | 0,000158 | 5,1308 | 21 |
| 3,09E+07 | 1,57E+07 | 2,55E+07 | 2,58E+07 | 83,628 | 0 | 44,687 | 30 |
| 9,47E+07 | 1,23E+08 | 1,42E+08 | 1,69E+08 | 14,982 | 0 | 9,901 | 25 |
| 3,27E+06 | 5,40E+06 | 5,38E+06 | 8,34E+06 | 130,12 | 0 | 11,615 | 13 |
| 5,31E+07 | 5,98E+07 | 6,23E+07 | 6,33E+07 | 86,841 | 0 | 64,243 | 47 |
| 2,10E+08 | 1,30E+08 | 1,68E+08 | 1,90E+08 | 16,292 | 0 | 39,275 | 17 |
| 7,28E+08 | 6,35E+08 | 6,72E+08 | 6,97E+08 | 26,946 | 0 | 89,775 | 133 |
| 1,27E+07 | 4,89E+07 | 4,23E+07 | 3,96E+07 | 118,98 | 0 | 14,879 | 28 |
| 8,12E+06 | 2,20E+07 | 2,04E+07 | 1,92E+07 | 60,755 | 0 | 6,6869 | 19 |
| 1,59E+08 | 1,14E+08 | 1,50E+08 | 1,14E+08 | 42,161 | 0 | 112,79 | 123 |
| 2,96E+08 | 1,95E+08 | 2,43E+08 | 2,42E+08 | 86,014 | 0 | 68,859 | 120 |
| 8,45E+07 | 8,49E+07 | 9,84E+07 | 9,71E+07 | 55,979 | 0 | 23,63 | 37 |
| 4,78E+07 | 3,68E+07 | 4,20E+07 | 3,98E+07 | 163,68 | 0 | 49,256 | 51 |
| 4,20E+07 | 6,89E+07 | 5,57E+07 | 5,90E+07 | 41,204 | 0 | 27,228 | 52 |
| 1,28E+07 | 1,04E+07 | 1,06E+07 | 8,91E+06 | 72,172 | 0 | 6,9106 | 10 |
| 6,15E+07 | 9,71E+07 | 7,83E+07 | 8,55E+07 | 25,171 | 0 | 8,1671 | 41 |
| 7,68E+08 | 7,00E+08 | 6,31E+08 | 6,56E+08 | 17,163 | 0 | 250,74 | 122 |
| 8,30E+07 | 7,84E+07 | 7,81E+07 | 7,32E+07 | 98,314 | 0 | 117,71 | 58 |

| | | | | | | | |
|---|---|---|---|---|---|---|---|
| 2,10E+08 | 2,82E+08 | 2,94E+08 | 2,58E+08 | 40,88 | 0 | 94,354 | 125 |
| 2,42E+07 | 3,01E+07 | 3,48E+07 | 3,64E+07 | 125,79 | 0 | 15,401 | 37 |
| 8,56E+08 | 7,74E+08 | 7,27E+08 | 7,35E+08 | 28,823 | 0 | 196,64 | 126 |
| 7,17E+07 | 9,46E+07 | 8,78E+07 | 7,84E+07 | 44,375 | 0 | 56,719 | 52 |
| 3,81E+07 | 1,38E+06 | 2,23E+07 | 1,83E+06 | 60,253 | 0 | 17,712 | 31 |
| 5,69E+06 | 7,60E+06 | 8,65E+06 | 9,34E+06 | 45,996 | 0 | 7,8194 | 10 |
| 6,56E+07 | 3,39E+07 | 3,99E+07 | 4,77E+07 | 138,58 | 0 | 15,351 | 48 |
| 4,67E+08 | 6,47E+08 | 6,46E+08 | 6,52E+08 | 38,221 | 0 | 323,31 | 154 |
| 2,71E+08 | 4,28E+08 | 3,40E+08 | 3,49E+08 | 57,922 | 0 | 323,31 | 99 |
| 1,65E+08 | 1,26E+08 | 1,58E+08 | 1,41E+08 | 36,118 | 0 | 305,92 | 66 |
| 2,01E+06 | 3,83E+06 | 4,12E+06 | 3,00E+06 | 54,369 | 0,00043 | 2,7544 | 2 |
| 1,07E+08 | 1,05E+08 | 1,05E+08 | 9,86E+07 | 100,27 | 0 | 49,861 | 75 |
| 1,67E+10 | 2,01E+10 | 1,76E+10 | 1,98E+10 | 17,971 | 0 | 323,31 | 934 |
| 9,13E+06 | 2,91E+07 | 1,72E+07 | 2,07E+07 | 41,295 | 0 | 8,3559 | 8 |
| 3,15E+08 | 4,37E+08 | 4,99E+08 | 4,21E+08 | 20,198 | 0 | 193,07 | 69 |
| 5,09E+08 | 4,52E+08 | 4,47E+08 | 4,59E+08 | 26,974 | 0 | 162,64 | 136 |
| 1,26E+08 | 5,83E+06 | 3,27E+06 | 7,61E+07 | 31,852 | 0 | 168,64 | 38 |
| 2,72E+06 | 5,63E+06 | 4,87E+06 | 6,32E+06 | 74,492 | 0,00015 | 3,5968 | 7 |
| 1,88E+06 | 2,44E+06 | 2,87E+06 | 4,35E+06 | 27,129 | 0,000853 | 2,5688 | 4 |
| 1,71E+07 | 1,27E+07 | 1,38E+07 | 1,28E+07 | 33,413 | 0 | 6,7101 | 17 |
| 5,29E+08 | 2,52E+08 | 3,74E+08 | 3,27E+08 | 16,288 | 0 | 323,31 | 45 |
| 5,78E+06 | 7,12E+06 | 7,23E+06 | 6,34E+06 | 48,476 | 0,000151 | 3,8285 | 9 |
| 1,78E+07 | 1,35E+07 | 1,37E+07 | 1,38E+07 | 43,992 | 0 | 8,6961 | 22 |
| 1,27E+07 | 1,42E+07 | 1,37E+07 | 1,41E+07 | 46,297 | 0 | 6,8484 | 17 |
| 8,62E+07 | 5,49E+07 | 5,37E+07 | 6,53E+07 | 21,523 | 0 | 13,798 | 41 |
| 5,72E+08 | 5,33E+08 | 4,95E+08 | 5,03E+08 | 28,948 | 0 | 69,568 | 145 |
| 3,07E+08 | 3,39E+08 | 3,96E+08 | 3,79E+08 | 66,213 | 0 | 165,06 | 142 |
| 5,99E+07 | 8,29E+07 | 7,61E+07 | 8,60E+07 | 22,292 | 0 | 59,732 | 30 |
| 3,25E+07 | 2,46E+07 | 2,75E+07 | 3,13E+07 | 17,014 | 0,002779 | 2,0486 | 18 |
| 3,54E+07 | 3,08E+07 | 3,21E+07 | 3,25E+07 | 15,792 | 0 | 7,1401 | 22 |
| 9,55E+06 | 1,07E+07 | 1,40E+07 | 1,24E+07 | 35,512 | 0 | 17,41 | 14 |
| 5,99E+06 | 5,87E+06 | 4,75E+06 | 4,80E+06 | 24,822 | 0,000436 | 2,9999 | 7 |
| 5,99E+06 | 2,22E+06 | 2,31E+06 | 3,56E+06 | 108,24 | 0,000149 | 3,4273 | 6 |
| 5,11E+06 | 2,76E+06 | 4,36E+06 | 2,64E+06 | 39,909 | 0,000436 | 2,9897 | 8 |
| 2,79E+06 | 1,56E+07 | 1,70E+07 | 1,81E+07 | 143,05 | 0,000153 | 4,1906 | 10 |
| 5,84E+08 | 3,64E+08 | 3,80E+08 | 3,95E+08 | 31,3 | 0 | 85,233 | 100 |
| 1,05E+07 | 2,17E+07 | 1,55E+07 | 1,74E+07 | 53,006 | 0,000158 | 5,0093 | 15 |
| 3,63E+07 | 3,15E+07 | 2,82E+07 | 3,06E+07 | 42,853 | 0 | 14,46 | 35 |
| 1,00E+08 | 8,40E+07 | 8,46E+07 | 9,00E+07 | 99,191 | 0 | 48,335 | 70 |
| 8,94E+07 | 7,84E+07 | 6,71E+07 | 7,53E+07 | 83,424 | 0 | 83,038 | 63 |
| 4,73E+06 | 1,22E+07 | 1,68E+07 | 1,45E+07 | 63,617 | 0,000158 | 5,0941 | 15 |
| 1,69E+07 | 1,16E+07 | 8,89E+06 | 6,68E+06 | 13,922 | 0 | 12,501 | 12 |
| 2,05E+07 | 2,41E+07 | 2,79E+07 | 2,81E+07 | 45,121 | 0 | 9,9291 | 25 |
| 6,30E+07 | 7,25E+07 | 8,35E+07 | 7,22E+07 | 21,85 | 0 | 222,81 | 35 |
| 5,99E+07 | 3,39E+07 | 5,04E+07 | 4,78E+07 | 151,12 | 0 | 42,222 | 54 |
| 7,55E+06 | 1,20E+07 | 1,11E+07 | 1,14E+07 | 63,438 | 0,000152 | 4,0237 | 13 |
| 2,62E+09 | 3,25E+09 | 3,13E+09 | 3,18E+09 | 91,006 | 0 | 323,31 | 647 |
| 4,00E+07 | 6,75E+07 | 5,50E+07 | 5,52E+07 | 22,193 | 0 | 17,401 | 33 |
| 6,25E+07 | 6,46E+07 | 7,56E+07 | 7,39E+07 | 21,236 | 0 | 12,54 | 34 |
| 3,91E+07 | 2,48E+07 | 2,98E+07 | 3,27E+07 | 120,81 | 0 | 6,7213 | 14 |

| | | | | | | | |
|---|---|---|---|---|---|---|---|
| 2,30E+07 | 2,06E+07 | 1,52E+07 | 2,05E+07 | 47,133 | 0 | 12,438 | 22 |
| 2,91E+09 | 2,10E+09 | 2,26E+09 | 2,34E+09 | 33,377 | 0 | 323,31 | 176 |
| 1,11E+08 | 1,27E+08 | 1,37E+08 | 1,41E+08 | 88,733 | 0 | 58,525 | 82 |
| 1,12E+08 | 1,28E+08 | 1,28E+08 | 1,32E+08 | 56,346 | 0 | 92,226 | 52 |
| 3,95E+08 | 4,13E+08 | 4,57E+08 | 4,13E+08 | 36,539 | 0 | 323,31 | 96 |
| 7,62E+08 | 9,32E+08 | 8,80E+08 | 9,13E+08 | 40,709 | 0 | 323,31 | 195 |
| 2,17E+08 | 1,57E+08 | 1,50E+08 | 1,62E+08 | 45,288 | 0 | 202,85 | 81 |
| 1,98E+08 | 2,25E+08 | 2,40E+08 | 2,23E+08 | 194,31 | 0 | 54,271 | 131 |
| 3,03E+07 | 5,96E+07 | 7,65E+07 | 7,79E+07 | 100,84 | 0 | 17,676 | 42 |
| 1,27E+07 | 6,20E+06 | 4,18E+06 | 6,38E+06 | 67,177 | 0 | 9,2961 | 18 |
| 3,11E+09 | 2,22E+09 | 2,45E+09 | 2,07E+09 | 65,322 | 0 | 323,31 | 380 |
| 4,21E+06 | 6,97E+06 | 6,28E+06 | 8,48E+06 | 95,895 | 0,000154 | 4,3216 | 3 |
| 1,46E+08 | 1,18E+08 | 1,16E+08 | 7,82E+07 | 15,261 | 0 | 15,447 | 41 |
| 4,23E+08 | 3,71E+08 | 3,59E+08 | 3,18E+08 | 24,977 | 0 | 172,44 | 108 |
| 1,07E+09 | 1,11E+09 | 1,17E+09 | 1,08E+09 | 33,997 | 0 | 323,31 | 191 |
| 1,46E+07 | 2,18E+07 | 1,91E+07 | 2,24E+07 | 38,835 | 0 | 7,7593 | 29 |
| 4,09E+08 | 4,66E+08 | 4,75E+08 | 5,22E+08 | 39,776 | 0 | 323,31 | 88 |
| 1,23E+08 | 1,15E+08 | 1,12E+08 | 9,30E+07 | 20,921 | 0 | 15,04 | 48 |
| 1,41E+07 | 2,93E+06 | 3,68E+06 | 7,06E+06 | 42,098 | 0 | 8,7545 | 3 |
| 2,56E+07 | 1,33E+07 | 1,47E+07 | 1,14E+07 | 15,348 | 0 | 41,126 | 8 |
| 2,41E+07 | 3,35E+07 | 4,44E+07 | 3,82E+07 | 163,62 | 0 | 16,356 | 35 |
| 1,66E+06 | 1,21E+07 | 1,56E+07 | 1,40E+07 | 62,641 | 0 | 8,9291 | 11 |
| 5,44E+08 | 3,78E+08 | 4,11E+08 | 4,07E+08 | 37,186 | 0 | 96,208 | 110 |
| 6,58E+09 | 5,15E+09 | 5,86E+09 | 5,53E+09 | 75,441 | 0 | 323,31 | 650 |
| 4,30E+06 | 1,21E+07 | 1,59E+07 | 1,46E+07 | 77,456 | 0,001824 | 2,2386 | 5 |
| 2,89E+08 | 4,11E+08 | 3,48E+08 | 3,71E+08 | 26,31 | 0 | 58,599 | 72 |
| 1,17E+08 | 1,34E+08 | 1,25E+08 | 1,47E+08 | 36,836 | 0 | 32,956 | 58 |
| 4,38E+07 | 3,06E+07 | 2,88E+07 | 2,90E+07 | 43,346 | 0 | 38,157 | 25 |
| 1,35E+07 | 1,95E+06 | 1,12E+07 | 3,36E+06 | 15,953 | 0,000151 | 3,9169 | 7 |
| 8,39E+07 | 7,23E+07 | 6,51E+07 | 8,43E+07 | 20,472 | 0 | 22,883 | 23 |
| 2,20E+08 | 1,99E+08 | 2,02E+08 | 1,98E+08 | 165,1 | 0 | 302,32 | 158 |
| 3,50E+09 | 3,16E+09 | 3,06E+09 | 3,08E+09 | 66,785 | 0 | 323,31 | 418 |
| 1,14E+09 | 9,25E+08 | 9,40E+08 | 9,89E+08 | 22,906 | 0 | 180,2 | 234 |
| 2,54E+08 | 2,38E+08 | 2,36E+08 | 2,47E+08 | 38,914 | 0 | 47,392 | 111 |
| 2,10E+07 | 2,52E+07 | 3,15E+07 | 2,51E+07 | 28,928 | 0,000148 | 3,4065 | 12 |
| 2,02E+08 | 2,28E+08 | 2,30E+08 | 2,43E+08 | 42,256 | 0 | 293,21 | 86 |
| 1,01E+08 | 1,19E+08 | 1,05E+08 | 1,08E+08 | 42,774 | 0 | 21,069 | 54 |
| 1,06E+08 | 1,07E+08 | 1,13E+08 | 1,11E+08 | 58,552 | 0 | 16,223 | 33 |
| 1,63E+08 | 1,95E+08 | 2,03E+08 | 1,63E+08 | 49,372 | 0 | 231,66 | 40 |
| 4,23E+08 | 3,87E+08 | 3,95E+08 | 3,94E+08 | 122,74 | 0 | 122,99 | 156 |
| 1,27E+07 | 2,42E+07 | 2,57E+07 | 2,49E+07 | 17,271 | 0 | 13,286 | 19 |
| 2,35E+08 | 2,37E+08 | 2,62E+08 | 2,58E+08 | 149,19 | 0 | 309,57 | 138 |
| 1,72E+09 | 2,38E+09 | 2,13E+09 | 2,15E+09 | 11,475 | 0 | 323,31 | 65 |
| 7,64E+06 | 1,43E+07 | 1,19E+07 | 1,29E+07 | 75,683 | 0 | 6,3347 | 9 |
| 2,80E+07 | 1,76E+07 | 2,39E+07 | 2,60E+07 | 155,76 | 0 | 17,109 | 47 |
| 1,20E+08 | 1,69E+08 | 1,53E+08 | 1,47E+08 | 43,691 | 0 | 86,2 | 76 |
| 9,89E+08 | 4,27E+08 | 1,20E+08 | 6,42E+08 | 5,0256 | 0 | 31,043 | 46 |
| 1,24E+08 | 1,40E+08 | 1,42E+08 | 1,36E+08 | 96,948 | 0 | 75,167 | 100 |
| 2,61E+07 | 2,46E+07 | 2,13E+07 | 2,37E+07 | 32,281 | 0 | 6,8073 | 21 |
| 4,45E+08 | 4,11E+08 | 4,24E+08 | 4,19E+08 | 44,699 | 0 | 52,315 | 132 |

| | | | | | | | |
|---|---|---|---|---|---|---|---|
| 3,03E+09 | 1,85E+09 | 1,67E+09 | 1,94E+09 | 21,266 | 0 | 104,86 | 72 |
| 4,17E+07 | 5,38E+07 | 5,38E+07 | 6,53E+07 | 31,445 | 0 | 24,946 | 36 |
| 5,08E+08 | 4,44E+08 | 4,09E+08 | 3,95E+08 | 56,494 | 0 | 323,31 | 198 |
| 1,82E+07 | 2,55E+07 | 2,36E+07 | 2,74E+07 | 45,652 | 0 | 10,803 | 33 |
| 1,24E+07 | 7,03E+06 | 6,63E+05 | 2,22E+06 | 44,472 | 0 | 7,51 | 9 |
| 4,32E+07 | 3,72E+07 | 3,29E+07 | 3,16E+07 | 29,852 | 0 | 10,363 | 20 |
| 4,35E+06 | 2,35E+06 | 3,16E+06 | 1,96E+06 | 74,517 | 0 | 5,44 | 16 |
| 9,71E+06 | 8,24E+06 | 7,72E+06 | 9,66E+06 | 24,087 | 0 | 6,195 | 15 |
| 5,79E+07 | 3,11E+07 | 4,01E+07 | 3,68E+07 | 28,996 | 0 | 52,301 | 30 |
| 3,91E+07 | 3,17E+07 | 3,26E+07 | 3,25E+07 | 40,956 | 0 | 8,8072 | 22 |
| 8,85E+06 | 7,21E+06 | 1,72E+06 | 4,22E+06 | 52,734 | 0,000158 | 5,1429 | 8 |
| 3,36E+08 | 2,21E+08 | 2,27E+08 | 2,52E+08 | 53,076 | 0 | 83,457 | 140 |
| 1,03E+08 | 7,37E+07 | 7,13E+07 | 8,80E+07 | 125,3 | 0 | 38,303 | 48 |
| 2,30E+07 | 1,79E+07 | 1,52E+07 | 2,27E+07 | 38,362 | 0 | 6,6964 | 12 |
| 4,26E+07 | 1,86E+07 | 2,18E+07 | 2,80E+07 | 48,223 | 0 | 7,7989 | 25 |
| 4,08E+08 | 4,10E+08 | 4,56E+08 | 4,34E+08 | 91,709 | 0 | 146,63 | 133 |
| 4,00E+08 | 4,93E+08 | 4,68E+08 | 4,96E+08 | 72,082 | 0 | 244,91 | 207 |
| 7,47E+07 | 9,86E+07 | 8,11E+07 | 7,73E+07 | 38,015 | 0 | 34,753 | 48 |
| 1,18E+08 | 1,15E+08 | 1,06E+08 | 9,75E+07 | 79,761 | 0 | 41,671 | 85 |
| 1,97E+09 | 1,60E+09 | 1,89E+09 | 1,80E+09 | 32,994 | 0 | 208,7 | 185 |
| 5,98E+08 | 5,66E+08 | 4,68E+08 | 4,71E+08 | 57,772 | 0 | 323,31 | 109 |
| 1,08E+07 | 2,29E+06 | 1,96E+06 | 2,23E+06 | 73,023 | 0,000434 | 2,8889 | 3 |
| 4,69E+07 | 3,05E+07 | 3,00E+07 | 2,59E+07 | 35,897 | 0 | 10,285 | 22 |
| 8,82E+07 | 1,08E+08 | 1,05E+08 | 1,06E+08 | 33,832 | 0 | 63,853 | 48 |
| 3,10E+07 | 4,23E+07 | 3,73E+07 | 3,16E+07 | 32,459 | 0 | 17,553 | 38 |
| 1,19E+07 | 4,19E+06 | 9,44E+06 | 3,68E+06 | 104,26 | 0 | 5,9908 | 8 |
| 2,64E+07 | 2,33E+07 | 2,15E+07 | 1,49E+07 | 28,367 | 0 | 10,646 | 17 |
| 2,78E+06 | 4,47E+07 | 4,58E+07 | 4,78E+07 | 46,313 | 0 | 13,911 | 32 |
| 2,13E+06 | 8,40E+06 | 8,73E+06 | 9,00E+06 | 20,892 | 0,00127 | 2,3935 | 4 |
| 2,86E+08 | 3,33E+08 | 3,18E+08 | 3,24E+08 | 22,314 | 0 | 151,44 | 108 |
| 7,11E+07 | 7,19E+07 | 7,51E+07 | 7,64E+07 | 37,06 | 0 | 15,318 | 41 |
| 2,21E+07 | 1,56E+07 | 3,89E+06 | 8,40E+05 | 65,01 | 0 | 39,679 | 20 |
| 2,12E+08 | 2,03E+08 | 1,95E+08 | 2,10E+08 | 133,41 | 0 | 125,85 | 135 |
| 8,85E+07 | 7,94E+07 | 8,68E+07 | 7,51E+07 | 32,918 | 0 | 28,324 | 75 |
| 4,82E+06 | 5,64E+06 | 5,39E+06 | 6,40E+06 | 34,169 | 0,00383 | 1,8053 | 7 |
| 1,87E+08 | 1,67E+08 | 1,25E+08 | 1,05E+08 | 9,461 | 0 | 39,382 | 130 |
| 1,35E+08 | 1,95E+08 | 1,85E+08 | 1,85E+08 | 34,786 | 0 | 139,57 | 51 |
| 7,30E+07 | 1,87E+07 | 4,40E+07 | 2,67E+07 | 22,464 | 0 | 87,104 | 20 |
| 4,60E+07 | 3,83E+07 | 3,17E+07 | 3,12E+07 | 20,811 | 0 | 10,843 | 29 |
| 4,17E+07 | 5,62E+07 | 4,73E+07 | 5,53E+07 | 53,237 | 0 | 18,301 | 60 |
| 2,64E+06 | 1,49E+07 | 1,38E+07 | 1,37E+07 | 50,469 | 0,000148 | 3,3459 | 11 |
| 7,27E+07 | 6,74E+07 | 6,87E+07 | 6,99E+07 | 81,792 | 0 | 19,589 | 56 |
| 8,02E+08 | 6,00E+08 | 7,04E+08 | 7,16E+08 | 52,292 | 0 | 323,31 | 187 |
| 3,16E+08 | 3,92E+08 | 4,61E+08 | 4,28E+08 | 40,855 | 0 | 275,14 | 93 |
| 7,70E+06 | 4,35E+06 | 6,24E+06 | 5,31E+06 | 46,914 | 0,00015 | 3,7578 | 12 |
| 1,76E+08 | 1,62E+08 | 1,55E+08 | 1,71E+08 | 29,1 | 0 | 29,303 | 68 |
| 9,54E+07 | 1,45E+08 | 1,22E+08 | 1,34E+08 | 43,735 | 0 | 81,552 | 83 |
| 1,52E+07 | 2,32E+07 | 1,94E+07 | 2,58E+07 | 43,209 | 0,009742 | 1,4059 | 7 |
| 9,78E+06 | 4,10E+06 | 2,49E+06 | 2,46E+06 | 30,546 | 0,004498 | 1,7362 | 5 |
| 1,47E+08 | 1,78E+08 | 1,72E+08 | 1,62E+08 | 44,789 | 0 | 113,2 | 69 |

| | | | | | | | |
|---|---|---|---|---|---|---|---|
| 1,82E+08 | 1,70E+08 | 1,47E+08 | 1,64E+08 | 63,471 | 0 | 141,29 | 97 |
| 1,61E+07 | 2,33E+07 | 1,93E+07 | 3,16E+07 | 37,189 | 0 | 7,38 | 24 |
| 9,21E+08 | 8,97E+08 | 8,52E+08 | 9,06E+08 | 58,841 | 0 | 323,31 | 245 |
| 2,72E+07 | 3,40E+07 | 3,40E+07 | 3,70E+07 | 58,664 | 0 | 10,323 | 29 |
| 3,06E+06 | 1,26E+07 | 1,68E+07 | 1,31E+07 | 305,39 | 0 | 11,285 | 19 |
| 1,32E+08 | 1,11E+08 | 1,06E+08 | 8,45E+07 | 45,389 | 0 | 158,73 | 65 |
| 4,40E+09 | 4,05E+09 | 4,19E+09 | 4,06E+09 | 244,54 | 0 | 323,31 | 1057 |
| 1,61E+08 | 1,49E+08 | 1,46E+08 | 1,43E+08 | 68,561 | 0 | 178,67 | 87 |
| 7,13E+08 | 4,90E+06 | 5,95E+07 | 9,20E+07 | 58,863 | 0 | 303,18 | 60 |
| 8,04E+07 | 6,45E+07 | 7,18E+07 | 6,10E+07 | 195,53 | 0 | 48,975 | 82 |
| 7,62E+07 | 1,03E+08 | 1,02E+08 | 8,88E+07 | 48,768 | 0 | 110,63 | 37 |
| 2,35E+07 | 1,12E+07 | 4,96E+06 | 8,64E+05 | 93,433 | 0 | 29,656 | 22 |
| 3,07E+08 | 3,57E+08 | 3,77E+08 | 3,92E+08 | 87,149 | 0 | 129,71 | 81 |
| 7,87E+06 | 1,51E+07 | 1,44E+07 | 1,68E+07 | 53,201 | 0 | 8,0365 | 18 |
| 7,92E+08 | 7,94E+08 | 6,96E+08 | 6,87E+08 | 53,408 | 0 | 225,94 | 230 |
| 1,10E+07 | 2,81E+06 | 9,45E+06 | 2,75E+06 | 29,19 | 0 | 18,37 | 7 |
| 4,35E+07 | 4,86E+07 | 5,67E+07 | 4,74E+07 | 44,607 | 0 | 75,137 | 28 |
| 3,97E+08 | 3,59E+08 | 3,24E+08 | 3,19E+08 | 58,213 | 0 | 321,29 | 70 |
| 1,46E+06 | 1,34E+06 | 1,17E+06 | 1,31E+06 | 48,974 | 0,000153 | 4,0902 | 5 |
| 2,59E+07 | 2,17E+07 | 2,53E+07 | 2,24E+07 | 19,824 | 0 | 6,3841 | 6 |
| 9,85E+06 | 3,70E+06 | 2,25E+06 | 2,84E+06 | 57,971 | 0 | 11,717 | 15 |
| 6,66E+08 | 5,01E+08 | 5,47E+08 | 5,69E+08 | 37,35 | 0 | 323,31 | 163 |
| 6,97E+07 | 4,92E+07 | 5,81E+07 | 5,90E+07 | 141 | 0 | 23,526 | 66 |
| 7,95E+07 | 1,26E+08 | 1,33E+08 | 1,39E+08 | 41,565 | 0 | 28,005 | 61 |
| 5,25E+06 | 9,27E+06 | 6,50E+06 | 6,60E+06 | 70,279 | 0,000155 | 4,4631 | 7 |
| 6,47E+06 | 4,90E+06 | 5,97E+06 | 3,87E+06 | 33,339 | 0,000151 | 3,834 | 9 |
| 1,41E+07 | 5,23E+06 | 3,05E+06 | 2,03E+06 | 21,434 | 0 | 22,149 | 8 |
| 1,06E+07 | 9,42E+06 | 6,45E+06 | 7,54E+06 | 46,541 | 0 | 10,24 | 19 |
| 3,23E+07 | 2,91E+07 | 2,79E+07 | 3,06E+07 | 33,975 | 0 | 12,193 | 29 |
| 8,28E+06 | 9,27E+06 | 1,10E+07 | 1,56E+07 | 34,067 | 0,000429 | 2,7146 | 9 |
| 8,09E+07 | 1,44E+08 | 1,06E+08 | 1,28E+08 | 13,48 | 0 | 55,186 | 43 |
| 2,92E+07 | 2,87E+07 | 2,77E+07 | 2,60E+07 | 31,139 | 0 | 24,04 | 27 |
| 7,09E+06 | 7,89E+06 | 8,19E+06 | 1,06E+07 | 124,68 | 0,000159 | 5,2183 | 12 |
| 5,20E+07 | 6,69E+07 | 5,19E+07 | 6,05E+07 | 36,975 | 0 | 73,212 | 18 |
| 2,15E+07 | 2,61E+07 | 3,59E+07 | 4,00E+07 | 80,428 | 0 | 9,2984 | 25 |
| 1,29E+09 | 1,58E+09 | 1,57E+09 | 1,74E+09 | 102,45 | 0 | 323,31 | 284 |
| 1,25E+07 | 1,69E+07 | 1,85E+07 | 1,63E+07 | 71,617 | 0 | 26,75 | 26 |
| 3,57E+07 | 3,07E+07 | 3,12E+07 | 3,35E+07 | 40,152 | 0 | 53,56 | 26 |
| 1,94E+07 | 1,39E+07 | 1,40E+07 | 1,94E+07 | 276,81 | 0 | 12,825 | 31 |
| 6,25E+07 | 4,40E+07 | 4,82E+07 | 4,17E+07 | 49,798 | 0 | 36,535 | 42 |
| 1,81E+08 | 1,42E+08 | 1,40E+08 | 1,06E+08 | 124,52 | 0,00015 | 3,6384 | 17 |
| 5,17E+07 | 6,43E+07 | 7,63E+07 | 8,66E+07 | 11,042 | 0 | 105,71 | 29 |
| 1,60E+07 | 1,44E+07 | 1,55E+07 | 1,50E+07 | 65,937 | 0 | 5,5651 | 15 |
| 4,60E+08 | 4,17E+08 | 4,09E+08 | 4,06E+08 | 35,738 | 0 | 323,31 | 142 |
| 9,28E+07 | 1,16E+08 | 1,86E+08 | 1,54E+08 | 22,924 | 0 | 17,758 | 40 |
| 7,73E+06 | 1,30E+07 | 1,59E+07 | 1,80E+07 | 93,454 | 0 | 5,9249 | 12 |
| 1,81E+08 | 2,56E+08 | 2,65E+08 | 2,25E+08 | 40,301 | 0 | 83,929 | 85 |
| 2,60E+06 | 9,27E+06 | 1,12E+07 | 9,73E+06 | 34,644 | 0,000147 | 3,228 | 6 |
| 1,56E+06 | 3,05E+06 | 5,90E+06 | 4,24E+06 | 54,543 | 0,003444 | 1,9206 | 3 |
| 5,55E+07 | 4,28E+07 | 4,49E+07 | 3,07E+07 | 25,222 | 0 | 9,9544 | 16 |

| | | | | | | | |
|---|---|---|---|---|---|---|---|
| 5,27E+06 | 1,78E+06 | 3,77E+06 | 4,10E+06 | 26,354 | 0,001961 | 2,2061 | 5 |
| 3,80E+06 | 2,68E+06 | 1,39E+06 | 1,89E+06 | 28,986 | 0,003445 | 1,9275 | 5 |
| 4,72E+07 | 4,47E+07 | 4,55E+07 | 4,53E+07 | 79,565 | 0 | 18,19 | 62 |
| 3,94E+08 | 5,19E+08 | 4,65E+08 | 4,61E+08 | 53,442 | 0 | 102,24 | 159 |
| 1,13E+09 | 1,37E+09 | 1,32E+09 | 1,27E+09 | 84,818 | 0 | 323,31 | 223 |
| 5,92E+07 | 6,35E+07 | 7,57E+07 | 6,34E+07 | 41,762 | 0 | 44,022 | 42 |
| 1,19E+07 | 1,84E+07 | 1,76E+07 | 1,49E+07 | 30,509 | 0 | 18,76 | 13 |
| 1,86E+08 | 4,14E+07 | 5,78E+07 | 5,37E+07 | 36,464 | 0 | 145,75 | 27 |
| 9,60E+06 | 5,86E+06 | 6,86E+06 | 7,77E+06 | 32,423 | 0 | 5,8803 | 9 |
| 6,98E+07 | 1,00E+08 | 1,00E+08 | 9,23E+07 | 40,42 | 0 | 128,61 | 49 |
| 8,16E+06 | 1,21E+07 | 1,10E+07 | 8,03E+06 | 40,489 | 0 | 5,6481 | 8 |
| 9,94E+06 | 4,67E+06 | 5,20E+06 | 1,03E+07 | 80,251 | 0 | 9,9923 | 9 |
| 1,66E+08 | 2,22E+08 | 1,98E+08 | 1,98E+08 | 96,43 | 0 | 88,596 | 102 |
| 6,71E+08 | 6,62E+08 | 6,36E+08 | 5,98E+08 | 67,795 | 0 | 235,44 | 185 |
| 4,96E+07 | 6,14E+07 | 5,92E+07 | 5,03E+07 | 72,51 | 0 | 36,958 | 49 |
| 1,50E+06 | 1,31E+07 | 1,21E+07 | 1,49E+07 | 51,88 | 0,000159 | 5,252 | 6 |
| 9,82E+08 | 6,44E+08 | 5,45E+08 | 5,07E+08 | 20,373 | 0 | 97,751 | 112 |
| 1,99E+06 | 7,95E+06 | 1,21E+07 | 1,26E+07 | 43,5 | 0,000152 | 3,9601 | 7 |
| 2,89E+09 | 1,71E+09 | 1,72E+09 | 1,68E+09 | 42,598 | 0 | 323,31 | 334 |
| 1,50E+07 | 1,60E+07 | 1,69E+07 | 1,50E+07 | 59,716 | 0 | 6,3018 | 19 |
| 3,65E+07 | 2,54E+07 | 3,53E+07 | 2,27E+07 | 15,661 | 0 | 46,227 | 10 |
| 3,92E+08 | 4,47E+08 | 5,18E+08 | 4,74E+08 | 117,94 | 0 | 259,86 | 250 |
| 6,99E+06 | 6,77E+06 | 5,88E+06 | 5,82E+06 | 54,937 | 0,00383 | 1,8102 | 7 |
| 4,51E+07 | 2,39E+07 | 3,73E+07 | 2,73E+07 | 133,49 | 0 | 14,021 | 34 |
| 1,10E+08 | 8,95E+07 | 8,23E+07 | 8,86E+07 | 30,603 | 0 | 63,49 | 44 |
| 3,54E+06 | 2,85E+06 | 9,67E+05 | 2,15E+06 | 25,277 | 0,000149 | 3,532 | 3 |
| 1,62E+08 | 1,63E+08 | 1,84E+08 | 1,69E+08 | 70,843 | 0 | 142,27 | 138 |
| 1,07E+07 | 4,51E+06 | 2,49E+06 | 4,01E+06 | 117,46 | 0,000855 | 2,6039 | 10 |
| 2,09E+07 | 1,57E+07 | 1,53E+07 | 1,34E+07 | 32,698 | 0 | 29,995 | 19 |
| 1,71E+07 | 1,74E+07 | 1,24E+07 | 1,06E+07 | 66,25 | 0 | 7,1409 | 15 |
| 7,40E+06 | 2,08E+07 | 1,22E+07 | 1,25E+07 | 41,357 | 0 | 21,763 | 12 |
| 1,58E+08 | 1,18E+08 | 1,25E+08 | 1,26E+08 | 55,854 | 0 | 167,5 | 67 |
| 3,19E+08 | 3,08E+08 | 2,90E+08 | 2,72E+08 | 38,527 | 0 | 323,31 | 112 |
| 3,25E+06 | 3,93E+06 | 4,19E+06 | 5,25E+06 | 45,924 | 0 | 7,5732 | 9 |
| 3,89E+08 | 3,72E+08 | 3,36E+08 | 3,49E+08 | 34,134 | 0 | 323,31 | 115 |
| 1,52E+07 | 2,71E+07 | 2,58E+07 | 2,90E+07 | 88,49 | 0 | 12,411 | 24 |
| 5,80E+06 | 6,79E+06 | 1,02E+07 | 9,63E+06 | 54,707 | 0,000434 | 2,89 | 7 |
| 1,96E+09 | 1,80E+09 | 1,91E+09 | 1,93E+09 | 27,372 | 0 | 323,31 | 307 |
| 1,32E+07 | 1,05E+07 | 5,27E+06 | 3,36E+06 | 13,18 | 0 | 5,617 | 11 |
| 7,14E+06 | 9,82E+06 | 9,46E+06 | 1,30E+07 | 69,301 | 0,000153 | 4,1994 | 11 |
| 3,39E+07 | 5,73E+07 | 4,69E+07 | 6,28E+07 | 20,926 | 0 | 12,753 | 37 |
| 8,10E+07 | 9,92E+07 | 1,29E+08 | 1,28E+08 | 16,331 | 0 | 37,901 | 40 |
| 4,04E+09 | 4,23E+09 | 4,45E+09 | 4,21E+09 | 29,174 | 0 | 323,31 | 736 |
| 2,76E+09 | 2,56E+09 | 2,72E+09 | 2,54E+09 | 76,775 | 0 | 323,31 | 346 |
| 2,36E+07 | 2,42E+07 | 2,57E+07 | 2,51E+07 | 110,18 | 0 | 14,446 | 32 |
| 5,21E+08 | 5,66E+08 | 6,07E+08 | 6,18E+08 | 86,491 | 0 | 323,31 | 245 |
| 7,16E+08 | 6,34E+08 | 5,99E+08 | 6,45E+08 | 42,295 | 0 | 229,67 | 101 |
| 9,15E+07 | 1,17E+08 | 1,26E+08 | 1,50E+08 | 16,998 | 0 | 31,261 | 28 |
| 2,79E+08 | 2,58E+08 | 2,38E+08 | 2,29E+08 | 201,44 | 0 | 227,22 | 171 |
| 3,18E+08 | 2,65E+08 | 2,74E+08 | 2,99E+08 | 118,03 | 0 | 152,1 | 170 |

| | | | | | | | |
|---|---|---|---|---|---|---|---|
| 6,43E+06 | 8,70E+06 | 1,10E+07 | 7,83E+06 | 130,71 | 0 | 9,5463 | 15 |
| 1,71E+08 | 1,74E+08 | 1,53E+08 | 1,46E+08 | 28,97 | 0 | 146,34 | 92 |
| 4,12E+09 | 4,72E+09 | 6,68E+09 | 6,03E+09 | 12,504 | 0 | 323,31 | 149 |
| 1,07E+08 | 1,02E+08 | 1,03E+08 | 9,35E+07 | 49,997 | 0 | 84,693 | 64 |
| 4,69E+07 | 7,38E+07 | 6,11E+07 | 7,65E+07 | 10,333 | 0 | 52,276 | 33 |
| 1,76E+07 | 9,54E+06 | 1,48E+07 | 1,50E+07 | 48,795 | 0 | 6,9329 | 19 |
| 2,08E+07 | 3,19E+06 | 5,24E+06 | 5,35E+06 | 40,634 | 0 | 7,9284 | 15 |
| 4,08E+08 | 2,74E+08 | 2,87E+08 | 3,19E+08 | 64,808 | 0 | 57,25 | 128 |
| 2,85E+08 | 1,88E+08 | 2,54E+08 | 2,78E+08 | 46,763 | 0 | 309,93 | 86 |
| 4,90E+07 | 4,77E+07 | 3,52E+07 | 3,80E+07 | 60,484 | 0 | 67,303 | 58 |
| 4,51E+08 | 4,31E+08 | 4,12E+08 | 4,17E+08 | 53,912 | 0 | 323,31 | 103 |
| 1,37E+08 | 1,90E+08 | 1,68E+08 | 1,45E+08 | 21,951 | 0 | 42,937 | 25 |
| 6,83E+07 | 5,55E+07 | 6,38E+07 | 6,55E+07 | 117,17 | 0 | 19,628 | 53 |
| 6,52E+06 | 4,03E+06 | 6,20E+06 | 3,96E+06 | 21,778 | 0,000151 | 3,9009 | 12 |
| 2,28E+06 | 3,87E+06 | 7,36E+06 | 4,45E+06 | 11,323 | 0,000432 | 2,8132 | 6 |
| 1,03E+08 | 1,52E+08 | 1,38E+08 | 1,54E+08 | 65,335 | 0 | 79,874 | 78 |
| 1,81E+08 | 1,62E+08 | 1,55E+08 | 1,81E+08 | 93,933 | 0 | 60,363 | 89 |
| 4,55E+08 | 5,83E+08 | 5,32E+08 | 5,72E+08 | 90,973 | 0 | 323,31 | 153 |
| 3,89E+07 | 5,77E+07 | 5,18E+07 | 5,95E+07 | 13,274 | 0 | 87,376 | 31 |
| 1,58E+09 | 1,88E+09 | 2,08E+09 | 2,06E+09 | 14,173 | 0 | 323,31 | 132 |
| 2,30E+09 | 2,46E+09 | 2,54E+09 | 3,08E+09 | 20,732 | 0 | 119,83 | 144 |
| 2,64E+06 | 3,84E+06 | 2,93E+06 | 3,38E+06 | 64,778 | 0,00318 | 1,9701 | 8 |
| 7,02E+08 | 6,99E+08 | 7,37E+08 | 7,46E+08 | 136,97 | 0 | 323,31 | 273 |
| 1,30E+07 | 1,84E+07 | 2,46E+07 | 2,55E+07 | 24,739 | 0 | 11,935 | 19 |
| 9,65E+07 | 8,31E+07 | 8,58E+07 | 8,17E+07 | 52,212 | 0 | 21,257 | 32 |
| 2,62E+07 | 1,57E+07 | 1,46E+07 | 5,25E+06 | 252,3 | 0 | 6,5117 | 20 |
| 2,07E+08 | 1,65E+08 | 1,54E+08 | 1,50E+08 | 120,8 | 0 | 101,48 | 140 |
| 1,45E+09 | 9,90E+08 | 9,30E+08 | 9,41E+08 | 104,92 | 0 | 323,31 | 302 |
| 4,06E+07 | 4,75E+07 | 4,34E+07 | 4,94E+07 | 35 | 0 | 20,564 | 35 |
| 5,96E+06 | 3,23E+07 | 2,67E+07 | 3,26E+07 | 24,728 | 0 | 10,952 | 23 |
| 5,44E+06 | 8,21E+06 | 9,65E+06 | 8,67E+06 | 45,53 | 0,001275 | 2,4745 | 12 |
| 1,71E+07 | 3,16E+06 | 8,53E+06 | 5,13E+06 | 296,18 | 0 | 7,571 | 20 |
| 8,31E+07 | 1,04E+08 | 9,89E+07 | 8,99E+07 | 24,183 | 0 | 60,734 | 60 |
| 2,23E+08 | 2,41E+08 | 2,21E+08 | 2,45E+08 | 12,405 | 0 | 61,912 | 80 |
| 4,90E+07 | 3,87E+07 | 3,79E+07 | 3,81E+07 | 54,325 | 0 | 24,878 | 46 |
| 3,59E+06 | 4,52E+06 | 4,44E+06 | 3,71E+06 | 147,56 | 0,000854 | 2,5882 | 6 |
| 3,59E+08 | 3,73E+08 | 3,60E+08 | 4,00E+08 | 82,302 | 0 | 323,31 | 143 |
| 7,43E+07 | 7,22E+07 | 7,29E+07 | 6,36E+07 | 192,06 | 0 | 12,634 | 43 |
| 8,35E+06 | 1,49E+07 | 1,60E+07 | 1,46E+07 | 116,24 | 0 | 7,9425 | 18 |
| 1,54E+09 | 1,93E+09 | 1,85E+09 | 1,90E+09 | 60,718 | 0 | 323,31 | 327 |
| 1,39E+08 | 8,74E+07 | 8,78E+07 | 6,73E+07 | 23,648 | 0 | 159,96 | 36 |
| 5,11E+07 | 4,58E+07 | 4,41E+07 | 5,11E+07 | 364,21 | 0 | 25,632 | 57 |
| 7,11E+06 | 2,89E+07 | 3,07E+07 | 2,77E+07 | 93,827 | 0,00015 | 3,617 | 12 |
| 6,50E+08 | 4,64E+08 | 4,82E+08 | 4,57E+08 | 70,793 | 0 | 169,66 | 170 |
| 1,40E+07 | 5,54E+06 | 5,25E+06 | 4,86E+06 | 33,19 | 0 | 24,528 | 10 |
| 2,52E+08 | 2,95E+08 | 2,74E+08 | 3,39E+08 | 44,97 | 0 | 128,78 | 109 |
| 5,71E+07 | 4,39E+07 | 4,07E+07 | 3,63E+07 | 78,112 | 0 | 13,895 | 25 |
| 3,12E+06 | 7,41E+06 | 9,44E+06 | 5,44E+06 | 34,843 | 0,005412 | 1,6279 | 5 |
| 5,01E+08 | 3,08E+08 | 3,02E+08 | 3,11E+08 | 17,364 | 0 | 55,043 | 43 |
| 1,05E+07 | 6,57E+06 | 2,19E+06 | 7,98E+05 | 41,412 | 0 | 6,1585 | 9 |

| | | | | | | | |
|---|---|---|---|---|---|---|---|
| 4,53E+07 | 5,53E+07 | 5,69E+07 | 5,46E+07 | 66,197 | 0 | 26,11 | 44 |
| 1,24E+08 | 1,11E+08 | 1,12E+08 | 1,16E+08 | 24,814 | 0 | 64,245 | 56 |
| 7,68E+08 | 6,11E+08 | 7,11E+08 | 6,97E+08 | 23,897 | 0 | 323,31 | 125 |
| 3,07E+06 | 1,80E+06 | 8,07E+05 | 1,10E+06 | 14,414 | 0,002094 | 2,1327 | 6 |
| 5,49E+08 | 4,67E+08 | 3,92E+08 | 3,85E+08 | 55,72 | 0 | 323,31 | 208 |
| 1,20E+08 | 9,67E+07 | 1,07E+08 | 1,23E+08 | 62,188 | 0 | 13,805 | 50 |
| 7,06E+07 | 4,94E+07 | 5,10E+07 | 7,24E+07 | 66,047 | 0 | 51,833 | 46 |
| 4,17E+08 | 4,00E+08 | 4,20E+08 | 3,89E+08 | 47,037 | 0 | 177,08 | 197 |
| 3,60E+06 | 5,32E+06 | 6,13E+06 | 3,70E+06 | 93,557 | 0,000147 | 3,2447 | 6 |
| 6,20E+08 | 5,36E+08 | 5,47E+08 | 4,71E+08 | 51,847 | 0 | 243,73 | 155 |
| 6,54E+06 | 1,37E+07 | 1,32E+07 | 1,57E+07 | 339,09 | 0,000154 | 4,379 | 11 |
| 7,77E+07 | 6,83E+07 | 6,88E+07 | 7,74E+07 | 17,437 | 0 | 45,944 | 28 |
| 1,33E+09 | 1,03E+09 | 1,08E+09 | 1,33E+09 | 13,916 | 0 | 323,31 | 209 |
| 1,84E+08 | 1,59E+08 | 1,43E+08 | 1,60E+08 | 46,71 | 0 | 87,214 | 78 |
| 1,37E+08 | 1,24E+08 | 1,32E+08 | 1,28E+08 | 74,692 | 0 | 103,8 | 70 |
| 4,20E+08 | 3,39E+08 | 3,58E+08 | 3,74E+08 | 37,028 | 0 | 227,41 | 132 |
| 1,09E+08 | 6,04E+07 | 5,61E+07 | 5,35E+07 | 28,702 | 0 | 166,79 | 29 |
| 1,34E+08 | 1,32E+08 | 1,19E+08 | 1,22E+08 | 52,387 | 0 | 23,884 | 74 |
| 2,11E+08 | 2,16E+08 | 2,34E+08 | 2,11E+08 | 37,24 | 0 | 57,979 | 78 |
| 8,00E+08 | 5,24E+08 | 5,55E+08 | 8,39E+08 | 103,88 | 0 | 175,38 | 73 |
| 1,00E+07 | 1,43E+07 | 3,45E+07 | 3,23E+07 | 17,37 | 0 | 6,659 | 24 |
| 7,54E+06 | 2,33E+06 | 7,70E+05 | 4,86E+06 | 12,823 | 0 | 15,185 | 4 |
| 2,52E+09 | 3,93E+09 | 3,00E+09 | 3,21E+09 | 22,127 | 0 | 323,31 | 182 |
| 1,26E+07 | 1,28E+07 | 1,50E+07 | 1,37E+07 | 59,362 | 0 | 12,881 | 10 |
| 3,72E+07 | 4,08E+07 | 3,90E+07 | 3,82E+07 | 104,9 | 0 | 33,231 | 52 |
| 1,58E+08 | 1,02E+08 | 9,30E+07 | 8,80E+07 | 36,525 | 0 | 108,03 | 99 |
| 1,08E+08 | 1,41E+08 | 1,45E+08 | 1,47E+08 | 12,737 | 0 | 18,365 | 51 |
| 5,62E+07 | 2,91E+07 | 3,62E+07 | 3,71E+07 | 35,384 | 0 | 20,338 | 23 |
| 1,48E+07 | 2,41E+07 | 2,55E+07 | 3,61E+07 | 96,449 | 0 | 33,39 | 28 |
| 4,20E+08 | 3,57E+08 | 3,32E+08 | 3,35E+08 | 31,487 | 0 | 69,509 | 122 |
| 6,09E+07 | 7,35E+07 | 1,21E+08 | 1,08E+08 | 41,963 | 0 | 10,143 | 31 |
| 1,49E+07 | 9,98E+06 | 3,69E+06 | 1,00E+06 | 52,694 | 0,000433 | 2,8447 | 5 |
| 2,52E+08 | 2,49E+08 | 2,33E+08 | 2,01E+08 | 69,437 | 0 | 323,31 | 95 |
| 1,35E+09 | 1,45E+09 | 1,37E+09 | 1,58E+09 | 31,345 | 0 | 323,31 | 168 |
| 7,32E+06 | 9,77E+06 | 7,63E+06 | 8,45E+06 | 63 | 0,000429 | 2,6719 | 8 |
| 9,94E+06 | 8,36E+06 | 7,52E+06 | 5,71E+06 | 97,276 | 0 | 7,9172 | 12 |
| 3,90E+08 | 4,47E+08 | 4,40E+08 | 4,88E+08 | 42,329 | 0 | 323,31 | 74 |
| 1,89E+07 | 8,83E+06 | 9,45E+06 | 4,03E+06 | 24,041 | 0 | 6,641 | 16 |
| 2,71E+07 | 6,69E+06 | 2,31E+07 | 6,43E+06 | 56,909 | 0 | 19,918 | 11 |
| 4,24E+06 | 1,76E+07 | 4,27E+06 | 2,13E+07 | 123,48 | 0,003438 | 1,9032 | 3 |
| 2,11E+07 | 1,35E+07 | 1,85E+07 | 1,75E+07 | 20,703 | 0 | 27,876 | 13 |
| 1,03E+08 | 7,73E+07 | 1,02E+08 | 9,49E+07 | 86,606 | 0 | 30,682 | 69 |
| 6,71E+07 | 6,81E+07 | 6,86E+07 | 7,42E+07 | 27,578 | 0 | 9,3015 | 36 |
| 1,25E+09 | 1,17E+09 | 1,25E+09 | 1,24E+09 | 75,029 | 0 | 268,83 | 215 |
| 5,22E+07 | 6,44E+07 | 7,02E+07 | 5,51E+07 | 22,934 | 0 | 54,646 | 38 |
| 3,38E+07 | 4,31E+07 | 5,46E+07 | 4,40E+07 | 62,367 | 0 | 15,991 | 29 |
| 4,94E+06 | 4,26E+06 | 2,62E+06 | 4,72E+06 | 56,569 | 0 | 6,9239 | 5 |
| 5,79E+06 | 9,54E+06 | 8,60E+06 | 9,66E+06 | 32,786 | 0,003703 | 1,8558 | 17 |
| 8,05E+07 | 1,20E+08 | 1,11E+08 | 1,02E+08 | 47,947 | 0 | 26,708 | 45 |
| 8,02E+07 | 9,26E+07 | 9,07E+07 | 9,23E+07 | 104,68 | 0 | 45,536 | 88 |

| | | | | | | | |
|---|---|---|---|---|---|---|---|
| 4,36E+08 | 3,48E+08 | 3,08E+08 | 3,39E+08 | 118,1 | 0 | 302,93 | 142 |
| 1,73E+07 | 3,55E+07 | 3,14E+07 | 3,77E+07 | 25,948 | 0 | 12,57 | 19 |
| 3,05E+06 | 1,35E+07 | 1,32E+07 | 1,37E+07 | 120,94 | 0 | 10,429 | 8 |
| 8,03E+07 | 1,17E+08 | 1,32E+08 | 1,38E+08 | 8,1828 | 0 | 129,79 | 18 |
| 3,03E+08 | 3,44E+08 | 3,23E+08 | 3,09E+08 | 98,47 | 0 | 323,31 | 160 |
| 1,01E+08 | 9,96E+07 | 9,32E+07 | 9,61E+07 | 225,47 | 0 | 43,74 | 82 |
| 2,20E+07 | 1,04E+07 | 3,24E+06 | 9,84E+06 | 28,033 | 0,000153 | 4,0976 | 11 |
| 6,28E+06 | 3,74E+06 | 4,80E+06 | 1,97E+06 | 51,466 | 0,000159 | 5,2137 | 6 |
| 4,14E+06 | 1,10E+07 | 9,64E+06 | 1,04E+07 | 35,071 | 0,000155 | 4,423 | 8 |
| 2,14E+07 | 3,08E+07 | 3,57E+07 | 3,62E+07 | 34,51 | 0 | 20,334 | 29 |
| 5,47E+07 | 6,75E+07 | 8,35E+07 | 7,15E+07 | 35,977 | 0 | 224,61 | 33 |
| 1,61E+08 | 1,27E+08 | 1,36E+08 | 1,35E+08 | 83,108 | 0 | 68,243 | 78 |
| 1,47E+08 | 2,05E+08 | 1,79E+08 | 1,98E+08 | 44,12 | 0 | 90,11 | 94 |
| 2,04E+08 | 2,21E+08 | 2,34E+08 | 2,53E+08 | 30,059 | 0 | 116,26 | 80 |
| 7,46E+08 | 5,31E+08 | 5,34E+08 | 5,09E+08 | 29,729 | 0 | 323,31 | 133 |
| 1,75E+08 | 1,45E+08 | 1,53E+08 | 1,42E+08 | 45,029 | 0 | 79,778 | 76 |
| 3,77E+09 | 3,65E+09 | 3,12E+09 | 2,93E+09 | 29,82 | 0 | 323,31 | 223 |
| 4,28E+07 | 6,71E+07 | 9,79E+07 | 1,03E+08 | 59,646 | 0 | 23,221 | 33 |
| 9,53E+07 | 1,27E+08 | 1,23E+08 | 1,10E+08 | 34,732 | 0 | 52,985 | 62 |
| 1,33E+08 | 1,09E+08 | 1,21E+08 | 1,23E+08 | 15,977 | 0 | 134,48 | 49 |
| 1,71E+07 | 2,23E+06 | 1,39E+07 | 6,80E+06 | 334,06 | 0 | 14,582 | 22 |
| 1,76E+07 | 1,02E+06 | 6,12E+06 | 6,45E+06 | 8,4807 | 0,005411 | 1,6234 | 8 |
| 1,47E+07 | 2,09E+07 | 1,41E+07 | 2,26E+07 | 116,1 | 0 | 6,5873 | 20 |
| 2,35E+07 | 1,66E+07 | 1,41E+07 | 1,07E+07 | 52,181 | 0 | 8,3563 | 20 |
| 9,71E+07 | 9,51E+07 | 9,49E+07 | 9,13E+07 | 97,964 | 0 | 98,112 | 57 |
| 8,69E+06 | 8,95E+06 | 1,48E+07 | 1,08E+07 | 76,543 | 0,001548 | 2,3122 | 6 |
| 1,56E+08 | 1,81E+08 | 1,90E+08 | 1,69E+08 | 23,707 | 0 | 50,762 | 56 |
| 3,98E+08 | 4,25E+08 | 4,75E+08 | 3,96E+08 | 77,8 | 0 | 190,46 | 181 |
| 4,90E+08 | 6,14E+08 | 5,49E+08 | 5,63E+08 | 42,314 | 0 | 250,6 | 141 |
| 3,93E+07 | 5,43E+07 | 5,64E+07 | 4,69E+07 | 19,127 | 0 | 19,421 | 24 |
| 6,94E+07 | 7,09E+07 | 7,36E+07 | 7,36E+07 | 82,252 | 0 | 63,169 | 55 |
| 2,02E+07 | 8,61E+06 | 2,66E+06 | 6,74E+06 | 43,833 | 0 | 9,2128 | 15 |
| 4,75E+07 | 3,67E+07 | 3,60E+07 | 4,26E+07 | 65,444 | 0 | 26,478 | 45 |
| 8,29E+08 | 1,07E+09 | 1,19E+09 | 1,24E+09 | 44,729 | 0 | 249,79 | 223 |
| 1,96E+07 | 9,18E+06 | 1,07E+07 | 2,71E+06 | 54,177 | 0 | 15,957 | 17 |
| 6,55E+07 | 8,17E+07 | 7,81E+07 | 7,26E+07 | 131,36 | 0 | 22,978 | 71 |
| 2,79E+06 | 8,12E+06 | 6,80E+06 | 8,20E+06 | 22,375 | 0 | 5,6485 | 7 |
| 2,33E+07 | 3,42E+07 | 2,93E+07 | 3,52E+07 | 35,182 | 0 | 17,352 | 27 |
| 7,56E+06 | 9,57E+06 | 1,20E+07 | 1,07E+07 | 56,474 | 0 | 8,2377 | 21 |
| 1,87E+08 | 1,40E+08 | 1,31E+08 | 1,36E+08 | 63,505 | 0 | 68,681 | 73 |
| 5,57E+08 | 4,71E+08 | 4,86E+08 | 5,00E+08 | 43,735 | 0 | 318,78 | 148 |
| 3,05E+08 | 2,64E+08 | 2,37E+08 | 2,70E+08 | 34,819 | 0 | 313,64 | 122 |
| 1,06E+07 | 1,22E+07 | 1,43E+07 | 1,65E+07 | 53,578 | 0 | 6,9647 | 12 |
| 6,79E+07 | 9,92E+07 | 1,11E+08 | 9,52E+07 | 132,96 | 0 | 146,3 | 84 |
| 1,03E+08 | 9,46E+07 | 1,05E+08 | 9,58E+07 | 73,066 | 0 | 27,509 | 70 |
| 7,02E+08 | 8,02E+08 | 8,76E+08 | 7,98E+08 | 47,262 | 0 | 323,31 | 208 |
| 7,36E+06 | 9,52E+06 | 8,69E+06 | 8,26E+06 | 34,747 | 0,000151 | 3,7953 | 14 |
| 1,64E+07 | 3,84E+06 | 9,65E+06 | 6,69E+06 | 44,838 | 0 | 7,3983 | 7 |
| 3,62E+08 | 3,08E+08 | 3,19E+08 | 3,33E+08 | 49,895 | 0 | 155,04 | 110 |
| 1,54E+07 | 1,36E+07 | 1,37E+07 | 1,60E+07 | 43,619 | 0 | 8,0901 | 15 |

| | | | | | | | |
|---|---|---|---|---|---|---|---|
| 9,72E+07 | 4,38E+07 | 7,13E+07 | 7,09E+07 | 78,86 | 0 | 83,634 | 41 |
| 7,67E+07 | 5,62E+07 | 6,10E+07 | 6,78E+07 | 44,337 | 0 | 58,223 | 52 |
| 8,25E+07 | 9,15E+07 | 1,04E+08 | 9,50E+07 | 128,92 | 0 | 62,635 | 83 |
| 9,63E+07 | 1,22E+08 | 1,10E+08 | 1,00E+08 | 47,66 | 0 | 23,33 | 28 |
| 4,84E+06 | 1,02E+07 | 7,59E+06 | 7,99E+06 | 53,423 | 0,000151 | 3,8916 | 8 |
| 1,43E+07 | 8,95E+06 | 7,93E+06 | 6,60E+06 | 119,43 | 0 | 6,2455 | 21 |
| 1,68E+08 | 1,32E+08 | 1,30E+08 | 1,25E+08 | 40,711 | 0 | 69,507 | 44 |
| 1,13E+07 | 3,21E+06 | 6,31E+06 | 7,34E+06 | 139,17 | 0,000151 | 3,8702 | 10 |
| 8,29E+06 | 7,84E+06 | 7,14E+06 | 7,47E+06 | 74,361 | 0 | 6,7778 | 16 |
| 9,23E+06 | 1,81E+07 | 1,44E+07 | 1,43E+07 | 102,31 | 0 | 11,423 | 23 |
| 1,48E+08 | 1,05E+08 | 9,90E+07 | 9,75E+07 | 11,636 | 0 | 115,17 | 39 |
| 3,83E+07 | 4,42E+07 | 4,90E+07 | 5,08E+07 | 65,23 | 0 | 53,031 | 34 |
| 5,93E+08 | 5,27E+08 | 4,56E+08 | 4,81E+08 | 68,625 | 0 | 224,17 | 254 |
| 4,57E+07 | 4,59E+07 | 3,79E+07 | 2,99E+07 | 38,146 | 0 | 23,426 | 36 |
| 3,55E+06 | 5,55E+06 | 7,50E+06 | 4,45E+06 | 66,825 | 0,003573 | 1,8961 | 6 |
| 9,06E+07 | 7,46E+07 | 8,60E+07 | 7,43E+07 | 37,131 | 0 | 90,621 | 42 |
| 2,97E+08 | 3,83E+08 | 3,63E+08 | 3,37E+08 | 278,58 | 0 | 323,31 | 207 |
| 1,66E+09 | 1,86E+09 | 1,83E+09 | 1,77E+09 | 47,006 | 0 | 323,31 | 260 |
| 3,26E+08 | 3,75E+08 | 3,62E+08 | 3,90E+08 | 63,17 | 0 | 323,31 | 131 |
| 1,15E+09 | 1,07E+09 | 9,91E+08 | 1,11E+09 | 86,751 | 0 | 323,31 | 182 |
| 2,99E+07 | 2,59E+07 | 2,49E+07 | 2,07E+07 | 61,534 | 0 | 13,875 | 40 |
| 1,17E+06 | 2,52E+06 | 5,68E+06 | 4,01E+06 | 31,978 | 0,000713 | 2,6231 | 2 |
| 1,95E+09 | 2,12E+09 | 2,26E+09 | 2,26E+09 | 45,626 | 0 | 323,31 | 553 |
| 2,12E+08 | 1,46E+08 | 1,56E+08 | 1,69E+08 | 33,578 | 0 | 35,085 | 106 |
| 2,29E+07 | 1,82E+07 | 1,89E+07 | 2,15E+07 | 63,273 | 0 | 9,0388 | 24 |
| 1,21E+08 | 9,37E+07 | 1,14E+08 | 1,12E+08 | 54,099 | 0 | 53,837 | 49 |
| 8,65E+07 | 1,39E+08 | 1,25E+08 | 1,33E+08 | 61,075 | 0 | 25,004 | 49 |
| 1,63E+08 | 1,79E+08 | 1,63E+08 | 1,86E+08 | 200,24 | 0 | 67,204 | 119 |
| 6,22E+07 | 5,54E+07 | 4,49E+07 | 5,81E+07 | 72,462 | 0 | 39,501 | 47 |
| 1,95E+06 | 1,04E+07 | 1,22E+07 | 9,61E+06 | 36,141 | 0 | 5,5071 | 11 |
| 1,68E+09 | 1,81E+09 | 1,77E+09 | 1,72E+09 | 44,172 | 0 | 323,31 | 424 |
| 1,36E+07 | 2,48E+07 | 2,44E+07 | 2,25E+07 | 41,653 | 0 | 10,798 | 11 |
| 3,82E+08 | 3,26E+08 | 3,67E+08 | 3,75E+08 | 40,528 | 0 | 84,353 | 95 |
| 1,84E+06 | 3,30E+06 | 3,81E+06 | 1,98E+06 | 103,89 | 0,002094 | 2,132 | 5 |
| 4,68E+08 | 7,06E+08 | 6,30E+08 | 6,18E+08 | 53,902 | 0 | 148,51 | 142 |
| 1,67E+08 | 1,82E+08 | 1,65E+08 | 1,81E+08 | 69,571 | 0 | 31,668 | 78 |
| 1,63E+07 | 2,21E+07 | 2,84E+07 | 3,15E+07 | 9,7951 | 0 | 12,464 | 27 |
| 2,47E+07 | 3,35E+07 | 3,88E+07 | 3,08E+07 | 22,948 | 0 | 87,643 | 28 |
| 4,93E+06 | 1,76E+06 | 3,90E+06 | 3,98E+06 | 35,482 | 0,000155 | 4,5991 | 6 |
| 8,62E+08 | 7,75E+08 | 7,71E+08 | 7,78E+08 | 56,937 | 0 | 323,31 | 204 |
| 4,33E+08 | 6,24E+08 | 4,52E+08 | 4,96E+08 | 20,9 | 0 | 23,861 | 52 |
| 3,20E+08 | 2,22E+08 | 2,20E+08 | 2,19E+08 | 84,059 | 0 | 229,99 | 124 |
| 2,31E+08 | 1,96E+08 | 2,05E+08 | 2,30E+08 | 89,798 | 0 | 30,948 | 66 |
| 2,72E+08 | 2,77E+08 | 2,71E+08 | 2,93E+08 | 44,229 | 0 | 154,49 | 87 |
| 3,80E+07 | 2,74E+07 | 2,63E+07 | 3,25E+07 | 46,585 | 0 | 25,107 | 20 |
| 3,21E+07 | 1,02E+07 | 1,70E+07 | 1,06E+07 | 38,781 | 0 | 230,16 | 11 |
| 1,34E+08 | 1,61E+08 | 1,62E+08 | 1,64E+08 | 43,037 | 0 | 31,387 | 74 |
| 7,22E+07 | 5,47E+07 | 6,60E+07 | 4,61E+07 | 15,103 | 0 | 75,246 | 30 |
| 8,96E+06 | 2,20E+06 | 7,00E+06 | 1,62E+06 | 52,926 | 0,004632 | 1,7324 | 4 |
| 6,47E+07 | 6,14E+07 | 6,55E+07 | 5,67E+07 | 31,709 | 0 | 48,93 | 30 |

| | | | | | | | |
|---|---|---|---|---|---|---|---|
| 1,40E+07 | 1,11E+07 | 1,29E+07 | 1,23E+07 | 131,56 | 0,000156 | 4,6504 | 24 |
| 5,91E+07 | 6,72E+07 | 6,54E+07 | 6,49E+07 | 600,24 | 0 | 85,573 | 93 |
| 6,75E+07 | 2,93E+06 | 2,86E+07 | 2,28E+07 | 86,689 | 0 | 21,156 | 45 |
| 5,69E+07 | 4,15E+07 | 4,65E+07 | 5,73E+07 | 23,34 | 0 | 11,7 | 38 |
| 7,36E+07 | 5,09E+07 | 6,44E+07 | 5,59E+07 | 76,166 | 0 | 32,023 | 56 |
| 1,14E+09 | 4,13E+08 | 5,29E+08 | 4,03E+08 | 19,779 | 0 | 323,31 | 55 |
| 1,11E+09 | 9,10E+08 | 9,11E+08 | 1,01E+09 | 28,537 | 0 | 71,721 | 142 |
| 2,48E+06 | 5,41E+06 | 9,95E+06 | 9,80E+06 | 282,96 | 0,000147 | 3,2556 | 8 |
| 1,62E+07 | 2,35E+07 | 2,69E+07 | 2,23E+07 | 37,08 | 0 | 8,9578 | 26 |
| 5,38E+09 | 5,45E+09 | 5,52E+09 | 5,37E+09 | 58,066 | 0 | 323,31 | 453 |
| 3,43E+07 | 1,93E+07 | 1,26E+07 | 1,92E+07 | 37,848 | 0 | 9,7532 | 15 |
| 7,88E+07 | 6,05E+07 | 5,59E+07 | 6,51E+07 | 37,648 | 0 | 18,303 | 44 |
| 1,43E+08 | 1,22E+08 | 1,40E+08 | 1,43E+08 | 82,019 | 0 | 38,418 | 84 |
| 4,57E+08 | 3,62E+08 | 3,70E+08 | 3,81E+08 | 43,185 | 0 | 323,31 | 143 |
| 3,88E+07 | 2,23E+07 | 2,88E+07 | 3,00E+07 | 83,193 | 0 | 15,384 | 37 |
| 1,48E+08 | 1,09E+08 | 9,67E+07 | 8,10E+07 | 37,629 | 0 | 8,263 | 25 |
| 5,94E+06 | 9,12E+06 | 9,54E+06 | 7,51E+06 | 54,376 | 0,000152 | 4,0278 | 6 |
| 6,64E+07 | 4,80E+07 | 5,14E+07 | 5,28E+07 | 39,599 | 0 | 71,158 | 57 |
| 1,27E+07 | 3,71E+06 | 8,41E+06 | 1,18E+07 | 52,247 | 0,001825 | 2,2598 | 8 |
| 2,03E+07 | 1,89E+07 | 1,96E+07 | 1,61E+07 | 41,12 | 0 | 19,101 | 37 |
| 8,46E+06 | 1,69E+07 | 1,83E+07 | 1,78E+07 | 55,591 | 0 | 6,1887 | 21 |
| 8,88E+07 | 6,57E+07 | 7,88E+07 | 9,28E+07 | 22,517 | 0 | 94,026 | 27 |
| 2,05E+09 | 1,77E+09 | 1,89E+09 | 1,73E+09 | 43,062 | 0 | 323,31 | 242 |
| 1,80E+08 | 1,58E+08 | 1,83E+08 | 1,74E+08 | 105,72 | 0 | 323,31 | 122 |
| 1,16E+08 | 8,93E+07 | 8,72E+07 | 8,29E+07 | 39,41 | 0 | 71,069 | 64 |
| 3,08E+07 | 2,62E+07 | 2,85E+07 | 3,05E+07 | 30,488 | 0 | 7,1791 | 14 |
| 1,68E+09 | 1,73E+09 | 1,74E+09 | 1,97E+09 | 40,742 | 0 | 323,31 | 280 |
| 9,07E+06 | 9,05E+06 | 8,25E+06 | 8,32E+06 | 91,317 | 0 | 6,1413 | 10 |
| 7,48E+06 | 2,28E+07 | 2,21E+07 | 2,18E+07 | 20,14 | 0,000159 | 5,1621 | 12 |
| 2,37E+08 | 2,34E+08 | 2,27E+08 | 2,15E+08 | 80,085 | 0 | 108,77 | 118 |
| 5,79E+06 | 2,81E+06 | 4,27E+06 | 4,92E+06 | 36,347 | 0,000157 | 4,8786 | 12 |
| 8,87E+06 | 9,63E+06 | 1,35E+07 | 1,03E+07 | 19,928 | 0,00043 | 2,729 | 6 |
| 5,52E+06 | 7,06E+06 | 8,21E+06 | 7,82E+06 | 43,907 | 0 | 5,3794 | 7 |
| 2,56E+09 | 2,56E+09 | 3,13E+09 | 2,91E+09 | 17,951 | 0 | 323,31 | 525 |
| 7,36E+07 | 6,38E+07 | 5,58E+07 | 4,81E+07 | 50,466 | 0 | 30,791 | 49 |
| 1,33E+08 | 1,56E+08 | 1,44E+08 | 1,56E+08 | 81,817 | 0 | 92,623 | 98 |
| 2,43E+07 | 1,98E+07 | 1,96E+07 | 2,06E+07 | 60,063 | 0 | 74,101 | 32 |
| 8,58E+07 | 6,02E+07 | 6,24E+07 | 8,31E+07 | 53,435 | 0 | 25,185 | 32 |
| 4,01E+07 | 2,10E+06 | 1,93E+07 | 2,08E+07 | 41,067 | 0 | 8,2334 | 12 |
| 4,11E+06 | 3,16E+07 | 3,27E+07 | 3,29E+07 | 22,739 | 0 | 7,8755 | 8 |
| 1,89E+08 | 2,29E+08 | 2,07E+08 | 1,96E+08 | 50,714 | 0 | 323,31 | 103 |
| 3,03E+07 | 2,36E+07 | 1,88E+07 | 1,41E+07 | 21,308 | 0 | 8,9327 | 22 |
| 3,66E+08 | 3,93E+08 | 3,89E+08 | 3,59E+08 | 33,572 | 0 | 323,31 | 149 |
| 2,38E+08 | 2,75E+08 | 2,76E+08 | 2,82E+08 | 213,42 | 0 | 238,51 | 197 |
| 2,24E+08 | 3,30E+08 | 3,19E+08 | 3,10E+08 | 61,227 | 0 | 84,759 | 124 |
| 3,15E+07 | 2,56E+07 | 2,85E+07 | 2,14E+07 | 169,1 | 0 | 33,229 | 41 |
| 1,16E+08 | 1,34E+08 | 1,74E+08 | 1,21E+08 | 12,861 | 0 | 323,31 | 35 |
| 2,08E+08 | 1,25E+08 | 2,23E+08 | 1,61E+08 | 16,101 | 0 | 86,213 | 54 |
| 1,61E+07 | 1,95E+07 | 1,68E+07 | 1,95E+07 | 33,95 | 0 | 7,4802 | 16 |
| 8,51E+08 | 7,92E+08 | 8,48E+08 | 8,53E+08 | 131,03 | 0 | 323,31 | 233 |

| | | | | | | | |
|---|---|---|---|---|---|---|---|
| 2,99E+07 | 1,07E+07 | 4,97E+06 | 1,98E+07 | 41,021 | 0 | 20,435 | 17 |
| 9,52E+06 | 5,65E+06 | 5,95E+06 | 1,86E+06 | 42,723 | 0 | 14,817 | 9 |
| 3,35E+08 | 2,87E+08 | 2,96E+08 | 2,93E+08 | 106,52 | 0 | 323,31 | 184 |
| 2,23E+06 | 3,02E+06 | 2,71E+06 | 4,43E+06 | 71,443 | 0,000154 | 4,2894 | 4 |
| 1,01E+08 | 6,29E+07 | 5,99E+07 | 6,96E+07 | 17,143 | 0 | 170,93 | 49 |
| 9,78E+07 | 1,36E+08 | 1,27E+08 | 1,10E+08 | 219,89 | 0 | 132,82 | 130 |
| 3,96E+08 | 3,62E+08 | 3,68E+08 | 3,65E+08 | 113,97 | 0 | 290,67 | 175 |
| 1,25E+08 | 7,02E+07 | 1,10E+08 | 1,16E+08 | 23,668 | 0 | 106,77 | 47 |
| 2,70E+06 | 5,86E+06 | 6,02E+06 | 7,07E+06 | 60,674 | 0,000147 | 3,2464 | 6 |
| 1,26E+07 | 2,22E+07 | 1,99E+07 | 1,85E+07 | 35,466 | 0 | 30,388 | 13 |
| 2,59E+07 | 1,53E+07 | 9,41E+06 | 2,79E+06 | 61,822 | 0,000154 | 4,3226 | 16 |
| 1,14E+08 | 1,68E+08 | 1,60E+08 | 1,33E+08 | 44,503 | 0 | 61,461 | 76 |
| 2,20E+08 | 2,99E+08 | 2,98E+08 | 2,58E+08 | 17,749 | 0 | 287,66 | 99 |
| 9,22E+06 | 4,14E+06 | 4,97E+06 | 5,50E+06 | 59,065 | 0,000156 | 4,6804 | 13 |
| 4,23E+07 | 2,36E+07 | 2,24E+07 | 2,94E+07 | 91,29 | 0 | 64,622 | 28 |
| 1,34E+07 | 5,99E+06 | 9,09E+06 | 9,31E+06 | 59,986 | 0,002231 | 2,1089 | 7 |
| 5,27E+06 | 2,24E+06 | 3,64E+06 | 3,64E+06 | 12,473 | 0,002093 | 2,1215 | 7 |
| 3,81E+06 | 2,01E+06 | 1,06E+06 | 3,92E+06 | 13,209 | 0,000155 | 4,5809 | 5 |
| 1,16E+07 | 7,00E+06 | 2,50E+06 | 9,10E+06 | 73,904 | 0 | 5,6967 | 6 |
| 3,15E+07 | 2,05E+07 | 2,26E+07 | 1,78E+07 | 40,533 | 0 | 20,67 | 21 |
| 1,43E+08 | 1,42E+08 | 1,40E+08 | 1,42E+08 | 23,764 | 0 | 207,55 | 52 |
| 8,05E+06 | 6,66E+06 | 3,85E+06 | 2,06E+06 | 57,464 | 0,001547 | 2,3021 | 8 |
| 1,24E+08 | 4,92E+07 | 8,03E+07 | 9,55E+07 | 60,319 | 0 | 101,87 | 29 |
| 5,63E+08 | 5,46E+08 | 5,26E+08 | 5,47E+08 | 77,504 | 0 | 303,6 | 153 |
| 4,90E+06 | 4,73E+06 | 9,18E+06 | 7,31E+06 | 93,515 | 0,000154 | 4,3927 | 8 |
| 4,85E+08 | 4,09E+08 | 4,11E+08 | 4,26E+08 | 42,722 | 0 | 131,8 | 139 |
| 1,37E+09 | 1,04E+09 | 1,20E+09 | 1,18E+09 | 30,016 | 0 | 323,31 | 101 |
| 1,11E+07 | 7,56E+06 | 2,98E+06 | 2,53E+06 | 20,247 | 0,000435 | 2,945 | 6 |
| 9,61E+08 | 9,92E+08 | 9,66E+08 | 1,02E+09 | 181,82 | 0 | 323,31 | 348 |
| 1,25E+06 | 5,82E+06 | 5,73E+06 | 4,07E+06 | 45,899 | 0,005947 | 1,6049 | 3 |
| 9,12E+06 | 4,04E+06 | 5,53E+06 | 8,48E+06 | 66,554 | 0,000148 | 3,3723 | 9 |
| 1,77E+07 | 1,40E+07 | 1,25E+07 | 1,09E+07 | 44,418 | 0 | 53,029 | 20 |
| 2,34E+08 | 3,03E+08 | 3,01E+08 | 2,97E+08 | 29,98 | 0 | 178,49 | 80 |
| 1,22E+07 | 1,68E+07 | 1,70E+07 | 1,42E+07 | 78,97 | 0 | 8,6888 | 23 |
| 3,60E+07 | 4,33E+07 | 4,48E+07 | 5,15E+07 | 62,183 | 0 | 22,811 | 20 |
| 2,94E+08 | 2,94E+08 | 2,95E+08 | 2,57E+08 | 31,172 | 0 | 323,31 | 57 |
| 4,96E+06 | 3,86E+06 | 2,09E+06 | 1,25E+06 | 36,134 | 0,000432 | 2,8061 | 2 |
| 1,92E+07 | 3,72E+06 | 1,78E+06 | 1,79E+07 | 39,221 | 0,003181 | 1,9703 | 8 |
| 5,98E+07 | 5,86E+07 | 5,69E+07 | 6,05E+07 | 70,236 | 0 | 68,735 | 64 |
| 8,35E+07 | 4,84E+07 | 5,31E+07 | 4,97E+07 | 84,389 | 0 | 133,31 | 49 |
| 3,97E+07 | 3,34E+07 | 3,67E+07 | 4,16E+07 | 21,681 | 0 | 16,24 | 25 |
| 3,47E+07 | 3,51E+07 | 3,34E+07 | 3,25E+07 | 53,477 | 0 | 13,263 | 45 |
| 2,02E+07 | 2,86E+07 | 2,84E+07 | 2,51E+07 | 44,363 | 0 | 7,2633 | 25 |
| 7,59E+06 | 8,74E+06 | 1,02E+07 | 8,87E+06 | 70,999 | 0,000152 | 4,0167 | 19 |
| 2,33E+07 | 2,31E+07 | 1,95E+07 | 2,06E+07 | 61,851 | 0 | 21,054 | 30 |
| 1,93E+09 | 1,94E+09 | 2,16E+09 | 2,08E+09 | 16,93 | 0 | 323,31 | 229 |
| 6,83E+06 | 8,87E+06 | 1,01E+07 | 9,58E+06 | 40,803 | 0,00237 | 2,1 | 7 |
| 1,06E+07 | 9,53E+06 | 2,33E+06 | 1,90E+06 | 106,36 | 0 | 5,817 | 11 |
| 1,82E+07 | 1,47E+07 | 8,38E+06 | 1,48E+07 | 59,083 | 0 | 9,6247 | 31 |
| 3,55E+07 | 6,79E+07 | 4,52E+07 | 3,54E+07 | 54,173 | 0 | 9,5411 | 25 |

| | | | | | | | |
|---|---|---|---|---|---|---|---|
| 7,27E+06 | 8,62E+06 | 1,68E+07 | 1,71E+07 | 124,42 | 0 | 10,164 | 12 |
| 6,43E+08 | 5,12E+08 | 5,46E+08 | 6,25E+08 | 40,555 | 0 | 303,64 | 115 |
| 1,83E+08 | 1,34E+08 | 1,29E+08 | 1,29E+08 | 42,554 | 0 | 113,6 | 85 |
| 7,10E+07 | 1,39E+08 | 1,17E+08 | 1,35E+08 | 28,027 | 0 | 25,34 | 59 |
| 4,74E+07 | 4,59E+07 | 4,74E+07 | 4,55E+07 | 35,887 | 0 | 25,898 | 41 |
| 1,28E+08 | 1,06E+08 | 1,16E+08 | 1,11E+08 | 254,2 | 0 | 63,566 | 126 |
| 1,39E+08 | 1,22E+08 | 1,20E+08 | 1,25E+08 | 45,852 | 0 | 40,982 | 55 |
| 7,39E+07 | 6,28E+07 | 7,33E+07 | 7,37E+07 | 73,039 | 0 | 59,243 | 64 |
| 4,68E+08 | 4,41E+08 | 4,71E+08 | 4,54E+08 | 182,77 | 0 | 179,51 | 199 |
| 1,43E+09 | 1,37E+09 | 1,39E+09 | 1,45E+09 | 49,03 | 0 | 323,31 | 224 |
| 2,60E+08 | 2,14E+08 | 2,48E+08 | 2,68E+08 | 62,488 | 0 | 126,81 | 132 |
| 1,17E+08 | 8,90E+07 | 1,05E+08 | 1,11E+08 | 17,285 | 0,000157 | 4,9738 | 23 |
| 1,21E+08 | 1,47E+08 | 1,45E+08 | 1,64E+08 | 59,266 | 0 | 39,364 | 59 |
| 9,25E+06 | 7,80E+06 | 8,57E+06 | 5,80E+06 | 83,099 | 0 | 8,0851 | 20 |
| 5,28E+07 | 7,47E+07 | 6,57E+07 | 6,22E+07 | 28,263 | 0 | 26,488 | 41 |
| 8,83E+08 | 1,01E+09 | 8,48E+08 | 9,78E+08 | 25,086 | 0 | 143,26 | 148 |
| 4,01E+07 | 5,07E+07 | 3,86E+07 | 4,71E+07 | 13,335 | 0,000151 | 3,8155 | 17 |
| 1,68E+08 | 1,50E+08 | 1,61E+08 | 1,53E+08 | 120,62 | 0 | 124,76 | 114 |
| 2,75E+07 | 4,12E+07 | 4,17E+07 | 4,10E+07 | 29,237 | 0 | 19,104 | 14 |
| 1,17E+08 | 1,38E+08 | 1,59E+08 | 1,47E+08 | 50,235 | 0 | 130,15 | 62 |
| 3,94E+08 | 3,23E+08 | 3,63E+08 | 3,57E+08 | 174,06 | 0 | 321,19 | 212 |
| 2,49E+07 | 3,52E+07 | 3,72E+07 | 4,53E+07 | 94,219 | 0 | 12,945 | 36 |
| 4,61E+07 | 1,91E+07 | 2,61E+07 | 1,91E+07 | 137,47 | 0 | 9,981 | 26 |
| 8,59E+07 | 9,04E+07 | 8,76E+07 | 8,55E+07 | 33,34 | 0 | 107,35 | 51 |
| 4,66E+08 | 3,44E+08 | 3,45E+08 | 4,10E+08 | 45,739 | 0 | 99,722 | 108 |
| 5,76E+06 | 8,53E+06 | 9,65E+06 | 9,62E+06 | 128,51 | 0 | 7,842 | 8 |
| 8,59E+07 | 7,17E+07 | 7,16E+07 | 6,85E+07 | 30,345 | 0 | 29,538 | 47 |
| 2,30E+08 | 2,78E+08 | 2,29E+08 | 2,63E+08 | 37,279 | 0 | 323,31 | 95 |
| 2,85E+07 | 1,85E+07 | 1,84E+07 | 1,51E+07 | 83,825 | 0 | 15,905 | 36 |
| 3,15E+08 | 4,40E+08 | 3,56E+08 | 3,39E+08 | 46,582 | 0 | 199,72 | 142 |
| 2,45E+08 | 2,52E+08 | 2,45E+08 | 2,53E+08 | 58,608 | 0 | 212,79 | 125 |
| 1,41E+08 | 1,33E+08 | 1,31E+08 | 1,42E+08 | 79,558 | 0 | 172,98 | 85 |
| 1,26E+07 | 1,18E+07 | 7,56E+06 | 9,19E+06 | 34,468 | 0,000149 | 3,5572 | 9 |
| 3,14E+06 | 8,13E+06 | 7,17E+06 | 6,67E+06 | 123,72 | 0,000432 | 2,8205 | 7 |
| 2,76E+08 | 3,00E+08 | 3,17E+08 | 2,72E+08 | 38,897 | 0 | 65,213 | 82 |
| 3,30E+07 | 1,52E+07 | 2,04E+07 | 1,83E+07 | 76,007 | 0 | 8,4657 | 16 |
| 5,14E+07 | 6,81E+07 | 6,92E+07 | 6,32E+07 | 40,433 | 0 | 15,459 | 40 |
| 1,74E+09 | 1,89E+09 | 1,78E+09 | 1,76E+09 | 47,408 | 0 | 323,31 | 308 |
| 2,29E+08 | 8,86E+07 | 5,66E+07 | 5,45E+07 | 18,637 | 0 | 179,27 | 51 |
| 1,21E+09 | 1,21E+09 | 1,22E+09 | 1,22E+09 | 146,89 | 0 | 323,31 | 393 |
| 8,71E+06 | 1,14E+07 | 1,36E+07 | 1,76E+07 | 26,59 | 0 | 9,1209 | 14 |
| 2,03E+08 | 2,39E+08 | 2,43E+08 | 2,45E+08 | 41,987 | 0 | 110,84 | 75 |
| 2,28E+07 | 1,61E+07 | 1,90E+07 | 2,01E+07 | 61,508 | 0 | 8,7431 | 17 |
| 2,06E+08 | 1,98E+08 | 1,97E+08 | 1,67E+08 | 83,025 | 0 | 239,84 | 125 |
| 4,51E+06 | 8,21E+06 | 6,71E+06 | 7,11E+06 | 67,195 | 0,000154 | 4,3486 | 12 |
| 1,95E+07 | 2,99E+07 | 2,90E+07 | 3,22E+07 | 51,965 | 0 | 27,205 | 24 |
| 3,55E+06 | 1,31E+07 | 1,11E+07 | 1,23E+07 | 128,66 | 0,000156 | 4,6768 | 7 |
| 1,54E+08 | 1,62E+08 | 2,02E+08 | 1,68E+08 | 70,306 | 0 | 181,8 | 151 |
| 2,55E+07 | 2,45E+07 | 4,65E+07 | 3,53E+07 | 76,851 | 0 | 50,014 | 24 |
| 6,97E+06 | 3,15E+06 | 5,95E+06 | 4,60E+06 | 40,249 | 0,000852 | 2,5369 | 13 |

| | | | | | | | |
|---|---|---|---|---|---|---|---|
| 1,63E+08 | 1,74E+08 | 1,80E+08 | 1,83E+08 | 33,816 | 0 | 75,653 | 65 |
| 1,39E+08 | 1,75E+08 | 1,48E+08 | 1,85E+08 | 33,209 | 0 | 80,829 | 73 |
| 4,24E+07 | 4,97E+07 | 5,29E+07 | 5,88E+07 | 86,962 | 0 | 36,098 | 44 |
| 1,10E+07 | 3,18E+07 | 3,07E+07 | 1,47E+07 | 51,723 | 0 | 9,5555 | 17 |
| 2,88E+07 | 3,49E+07 | 5,71E+07 | 4,86E+07 | 10,731 | 0 | 25,197 | 19 |
| 5,90E+08 | 6,78E+08 | 7,57E+08 | 7,20E+08 | 47,346 | 0 | 313,84 | 152 |
| 4,23E+07 | 5,85E+07 | 4,95E+07 | 5,55E+07 | 65,514 | 0 | 19,238 | 47 |
| 2,67E+08 | 2,20E+08 | 1,96E+08 | 2,17E+08 | 97,679 | 0 | 189,24 | 143 |
| 2,34E+08 | 1,73E+08 | 1,82E+08 | 1,78E+08 | 57,23 | 0 | 196,03 | 119 |
| 7,25E+06 | 2,60E+06 | 3,20E+06 | 2,74E+06 | 92,663 | 0,000156 | 4,7741 | 9 |
| 2,58E+07 | 1,40E+07 | 1,01E+07 | 1,46E+07 | 35,316 | 0 | 16,828 | 21 |
| 4,51E+08 | 3,77E+08 | 3,94E+08 | 3,79E+08 | 237,47 | 0 | 230,34 | 272 |
| 1,04E+08 | 1,05E+08 | 1,18E+08 | 1,04E+08 | 94,526 | 0 | 64,936 | 82 |
| 1,82E+08 | 1,79E+08 | 1,85E+08 | 1,89E+08 | 48,057 | 0 | 94,743 | 73 |
| 9,36E+06 | 7,48E+06 | 7,74E+06 | 7,51E+06 | 29,445 | 0,004893 | 1,6921 | 7 |
| 5,20E+06 | 3,36E+06 | 3,65E+06 | 2,10E+06 | 100,8 | 0,000146 | 3,0827 | 9 |
| 1,11E+08 | 6,28E+07 | 1,12E+08 | 9,53E+07 | 13,813 | 0 | 10,848 | 61 |
| 9,89E+06 | 5,19E+06 | 4,88E+06 | 5,92E+06 | 25,393 | 0 | 8,2649 | 17 |
| 5,72E+06 | 9,15E+06 | 2,32E+06 | 2,65E+06 | 25,652 | 0,000146 | 3,0866 | 2 |
| 6,53E+07 | 5,79E+07 | 6,26E+07 | 6,66E+07 | 21,733 | 0 | 42,148 | 32 |
| 2,14E+07 | 1,31E+07 | 1,09E+07 | 1,88E+07 | 28,588 | 0,000151 | 3,8061 | 16 |
| 1,32E+08 | 1,77E+08 | 1,97E+08 | 1,94E+08 | 25,345 | 0 | 72,542 | 73 |
| 8,59E+06 | 1,69E+07 | 1,47E+07 | 1,96E+07 | 147,29 | 0,000151 | 3,8158 | 15 |
| 5,50E+09 | 4,97E+09 | 4,95E+09 | 4,77E+09 | 109,36 | 0 | 323,31 | 604 |
| 5,66E+06 | 7,23E+06 | 7,77E+06 | 5,88E+06 | 110,06 | 0,000432 | 2,8308 | 14 |
| 2,16E+08 | 2,71E+08 | 2,70E+08 | 2,64E+08 | 46,936 | 0 | 105,47 | 110 |
| 7,26E+06 | 3,94E+06 | 3,71E+06 | 1,12E+06 | 19,642 | 0,001273 | 2,447 | 2 |
| 4,94E+09 | 4,32E+09 | 3,81E+09 | 4,04E+09 | 38,354 | 0 | 161,65 | 342 |
| 2,49E+07 | 1,70E+07 | 1,56E+07 | 1,61E+07 | 85,287 | 0 | 20,107 | 20 |
| 1,21E+07 | 9,32E+06 | 2,01E+06 | 4,16E+06 | 58,217 | 0,000158 | 5,0951 | 9 |
| 4,18E+08 | 4,49E+08 | 4,65E+08 | 4,42E+08 | 124,25 | 0 | 323,31 | 226 |
| 5,98E+07 | 2,36E+07 | 2,34E+07 | 2,07E+07 | 19,611 | 0 | 39,808 | 28 |
| 1,76E+09 | 2,01E+09 | 2,05E+09 | 2,14E+09 | 31,013 | 0 | 323,31 | 153 |
| 2,61E+07 | 2,14E+07 | 1,39E+07 | 2,09E+07 | 119,31 | 0 | 12,666 | 27 |
| 1,32E+09 | 1,07E+09 | 1,12E+09 | 1,15E+09 | 63,299 | 0 | 323,31 | 274 |
| 9,25E+08 | 1,11E+09 | 1,22E+09 | 1,31E+09 | 33,499 | 0 | 323,31 | 171 |
| 1,01E+08 | 1,20E+08 | 1,17E+08 | 1,39E+08 | 30,26 | 0 | 71,168 | 55 |
| 1,05E+08 | 9,64E+07 | 8,13E+07 | 9,46E+07 | 52,028 | 0 | 23,678 | 44 |
| 4,91E+08 | 3,53E+08 | 3,42E+08 | 3,24E+08 | 34,208 | 0 | 104,76 | 138 |
| 7,85E+06 | 1,93E+06 | 3,23E+06 | 5,20E+06 | 49,425 | 0,000149 | 3,4939 | 6 |
| 1,26E+07 | 5,00E+06 | 8,41E+06 | 5,29E+06 | 45,882 | 0,000152 | 4,0776 | 9 |
| 4,26E+08 | 3,80E+08 | 3,58E+08 | 2,98E+08 | 78,734 | 0 | 190,27 | 159 |
| 6,75E+06 | 2,16E+07 | 1,36E+07 | 1,42E+07 | 58,282 | 0,000159 | 5,2652 | 19 |
| 1,12E+08 | 1,35E+08 | 1,15E+08 | 1,16E+08 | 53,129 | 0 | 98,654 | 72 |
| 1,81E+07 | 2,96E+07 | 4,60E+07 | 3,45E+07 | 63,388 | 0 | 8,1649 | 29 |
| 1,55E+08 | 1,65E+08 | 1,76E+08 | 1,58E+08 | 83,926 | 0 | 57,803 | 96 |
| 1,49E+08 | 1,24E+08 | 1,45E+08 | 1,55E+08 | 61,363 | 0 | 46,277 | 58 |
| 3,48E+07 | 2,26E+07 | 2,32E+07 | 2,43E+07 | 52,264 | 0 | 20,834 | 21 |
| 7,17E+07 | 7,04E+07 | 7,60E+07 | 8,56E+07 | 37,253 | 0 | 19,677 | 40 |
| 8,36E+07 | 1,21E+08 | 9,39E+07 | 9,59E+07 | 20,52 | 0 | 120,23 | 42 |

| | | | | | | | |
|---|---|---|---|---|---|---|---|
| 1,58E+07 | 1,55E+07 | 1,20E+07 | 1,52E+07 | 64,91 | 0 | 5,8597 | 22 |
| 9,04E+07 | 7,06E+07 | 7,38E+07 | 6,14E+07 | 15,147 | 0 | 33,384 | 26 |
| 1,55E+07 | 1,31E+07 | 1,36E+07 | 1,08E+07 | 53,107 | 0 | 19,421 | 26 |
| 1,00E+08 | 3,52E+08 | 3,06E+08 | 2,33E+08 | 19,592 | 0 | 94,961 | 39 |
| 1,99E+07 | 7,08E+06 | 1,10E+07 | 8,71E+06 | 16,699 | 0 | 42,595 | 15 |
| 9,83E+06 | 6,38E+06 | 5,02E+06 | 9,37E+06 | 55,269 | 0 | 10,014 | 11 |
| 2,94E+08 | 2,78E+08 | 2,70E+08 | 2,79E+08 | 38,194 | 0 | 133,27 | 80 |
| 2,14E+08 | 2,43E+08 | 2,29E+08 | 2,05E+08 | 106,3 | 0 | 144,44 | 136 |
| 6,12E+07 | 5,17E+07 | 4,75E+07 | 4,63E+07 | 34,888 | 0 | 60,312 | 58 |
| 1,20E+07 | 3,27E+06 | 1,07E+07 | 1,81E+06 | 51,842 | 0 | 5,8878 | 12 |
| 1,38E+08 | 1,23E+08 | 1,08E+08 | 1,25E+08 | 40,89 | 0 | 74,434 | 69 |
| 1,49E+06 | 2,36E+06 | 3,26E+06 | 1,01E+06 | 540,18 | 0,000429 | 2,6762 | 5 |
| 2,38E+08 | 1,93E+08 | 2,09E+08 | 2,00E+08 | 86,555 | 0 | 249,23 | 111 |
| 4,63E+06 | 9,94E+06 | 1,21E+07 | 1,53E+07 | 46,713 | 0,000149 | 3,4792 | 26 |
| 1,36E+08 | 1,31E+08 | 1,49E+08 | 1,54E+08 | 25,921 | 0 | 149,79 | 58 |
| 6,29E+07 | 6,32E+07 | 5,60E+07 | 5,82E+07 | 86,32 | 0 | 20,497 | 59 |
| 2,70E+08 | 2,67E+08 | 2,23E+08 | 2,33E+08 | 31,253 | 0 | 70,6 | 98 |
| 2,12E+07 | 2,64E+07 | 3,38E+07 | 2,31E+07 | 123,97 | 0 | 12,277 | 30 |
| 4,79E+08 | 6,75E+08 | 6,62E+08 | 7,50E+08 | 40,079 | 0 | 323,31 | 106 |
| 2,72E+07 | 2,97E+07 | 2,93E+07 | 2,89E+07 | 108,9 | 0 | 19,615 | 25 |
| 5,03E+08 | 5,25E+08 | 5,15E+08 | 5,08E+08 | 23,334 | 0 | 323,31 | 95 |
| 8,71E+07 | 9,29E+07 | 8,47E+07 | 8,89E+07 | 124,13 | 0 | 77,677 | 55 |
| 2,32E+07 | 4,91E+06 | 1,12E+07 | 1,10E+07 | 115,71 | 0 | 8,0006 | 21 |
| 2,82E+06 | 5,47E+06 | 2,80E+06 | 4,00E+06 | 37,701 | 0,002912 | 2,0126 | 3 |
| 2,88E+07 | 2,83E+07 | 2,56E+07 | 2,90E+07 | 44,072 | 0 | 5,9155 | 13 |
| 2,01E+07 | 2,96E+07 | 2,70E+07 | 2,51E+07 | 87,194 | 0 | 7,502 | 26 |
| 3,91E+07 | 3,81E+07 | 3,90E+07 | 4,43E+07 | 36,679 | 0 | 11,797 | 28 |
| 6,44E+07 | 6,20E+07 | 6,11E+07 | 6,05E+07 | 37,506 | 0 | 37,073 | 48 |
| 7,66E+07 | 5,39E+07 | 4,46E+07 | 3,73E+07 | 68,945 | 0 | 99,545 | 60 |
| 4,40E+08 | 4,41E+08 | 4,49E+08 | 4,37E+08 | 15,957 | 0 | 281,27 | 79 |
| 3,48E+07 | 2,44E+07 | 2,46E+07 | 2,85E+07 | 24,328 | 0 | 11,339 | 28 |
| 6,59E+07 | 5,25E+07 | 4,79E+07 | 4,73E+07 | 509,43 | 0 | 6,6375 | 28 |
| 3,77E+07 | 4,01E+07 | 3,66E+07 | 4,26E+07 | 158,48 | 0 | 25,503 | 50 |
| 9,88E+08 | 1,07E+09 | 1,08E+09 | 1,14E+09 | 36,46 | 0 | 323,31 | 209 |
| 3,26E+08 | 1,02E+09 | 1,17E+09 | 7,42E+08 | 11,186 | 0 | 299,9 | 289 |
| 9,97E+07 | 2,34E+07 | 2,24E+07 | 4,04E+07 | 98,334 | 0 | 11,744 | 22 |
| 3,17E+06 | 2,53E+06 | 2,46E+06 | 3,10E+06 | 26,031 | 0,000993 | 2,4918 | 10 |
| 4,48E+06 | 5,06E+06 | 6,14E+06 | 6,66E+06 | 105,98 | 0,000148 | 3,3196 | 5 |
| 1,17E+08 | 1,05E+08 | 1,03E+08 | 1,01E+08 | 64,892 | 0 | 62,664 | 54 |
| 6,05E+07 | 6,99E+07 | 5,86E+07 | 6,43E+07 | 107,98 | 0 | 45,132 | 69 |
| 7,76E+07 | 8,26E+07 | 8,70E+07 | 9,08E+07 | 68,446 | 0 | 42,112 | 60 |
| 9,63E+06 | 6,42E+06 | 6,17E+06 | 5,75E+06 | 9,8175 | 0,000146 | 3,0911 | 10 |
| 3,58E+07 | 3,60E+07 | 3,66E+07 | 2,93E+07 | 13,649 | 0 | 75,377 | 27 |
| 1,82E+07 | 8,76E+07 | 3,42E+07 | 2,67E+07 | 89,406 | 0 | 5,5309 | 20 |
| 1,27E+07 | 2,40E+06 | 1,12E+06 | 1,63E+06 | 22,694 | 0 | 10,132 | 11 |
| 1,96E+07 | 3,85E+07 | 5,13E+07 | 6,39E+07 | 52,265 | 0 | 15,179 | 22 |
| 9,58E+08 | 1,15E+09 | 1,13E+09 | 1,11E+09 | 64,403 | 0 | 323,31 | 236 |
| 5,30E+08 | 1,79E+08 | 2,82E+08 | 2,08E+08 | 20,697 | 0 | 320,73 | 45 |
| 1,18E+07 | 4,35E+06 | 5,98E+06 | 1,19E+07 | 74,647 | 0 | 5,6005 | 9 |
| 1,17E+08 | 1,09E+08 | 1,10E+08 | 1,21E+08 | 41,234 | 0 | 13,581 | 43 |

| | | | | | | | |
|---|---|---|---|---|---|---|---|
| 1,12E+07 | 1,13E+07 | 1,12E+07 | 1,25E+07 | 26,839 | 0,005422 | 1,6497 | 9 |
| 3,92E+07 | 8,10E+07 | 5,43E+07 | 6,85E+07 | 28,032 | 0 | 7,8677 | 29 |
| 4,96E+06 | 2,69E+06 | 5,20E+06 | 3,54E+06 | 9,2908 | 0,003449 | 1,9371 | 4 |
| 8,02E+07 | 7,43E+07 | 6,21E+07 | 4,74E+07 | 66,031 | 0 | 11,441 | 42 |
| 1,96E+07 | 1,67E+07 | 1,85E+07 | 2,04E+07 | 42,957 | 0 | 6,8412 | 11 |
| 7,07E+06 | 1,06E+07 | 6,20E+06 | 1,06E+07 | 30,269 | 0 | 5,9517 | 5 |
| 9,20E+08 | 9,50E+08 | 1,03E+09 | 9,31E+08 | 37,895 | 0 | 323,31 | 214 |
| 1,15E+07 | 6,45E+06 | 1,18E+07 | 1,08E+07 | 54,052 | 0 | 8,5289 | 22 |
| 6,27E+06 | 1,34E+07 | 1,36E+07 | 1,09E+07 | 45,64 | 0 | 5,5985 | 20 |
| 7,27E+07 | 6,69E+07 | 7,20E+07 | 6,88E+07 | 41,873 | 0 | 27,42 | 41 |
| 2,44E+08 | 4,76E+07 | 6,12E+07 | 2,05E+06 | 166,58 | 0 | 104,39 | 66 |
| 8,62E+06 | 1,56E+07 | 1,35E+07 | 1,50E+07 | 111,81 | 0,000432 | 2,8232 | 5 |
| 1,27E+07 | 6,04E+06 | 9,06E+06 | 1,20E+07 | 51,51 | 0 | 6,3729 | 13 |
| 6,25E+07 | 7,06E+07 | 8,04E+07 | 7,19E+07 | 17,315 | 0 | 7,5026 | 20 |
| 3,39E+07 | 5,30E+07 | 4,28E+07 | 3,95E+07 | 34,728 | 0 | 22,879 | 41 |
| 1,56E+07 | 1,45E+07 | 1,51E+07 | 1,51E+07 | 47,642 | 0 | 6,0785 | 17 |
| 2,66E+08 | 2,62E+08 | 2,52E+08 | 2,74E+08 | 47,922 | 0 | 59,191 | 112 |
| 2,55E+08 | 2,47E+08 | 2,56E+08 | 2,50E+08 | 91,707 | 0 | 184,78 | 141 |
| 1,39E+07 | 3,37E+07 | 3,37E+07 | 4,07E+07 | 184,65 | 0 | 14,408 | 37 |
| 3,27E+08 | 2,94E+08 | 3,09E+08 | 3,12E+08 | 28,567 | 0 | 172,95 | 52 |
| 8,74E+06 | 2,10E+06 | 3,86E+06 | 3,81E+06 | 23,333 | 0,008986 | 1,4664 | 2 |
| 2,11E+08 | 2,41E+08 | 2,35E+08 | 2,50E+08 | 244,56 | 0 | 107,37 | 169 |
| 7,07E+06 | 7,29E+06 | 7,06E+06 | 6,56E+06 | 46,584 | 0,000149 | 3,4959 | 13 |
| 2,13E+08 | 2,25E+08 | 2,40E+08 | 2,54E+08 | 46,297 | 0 | 205,51 | 73 |
| 9,07E+07 | 8,20E+07 | 7,08E+07 | 8,95E+07 | 25,782 | 0 | 158,01 | 40 |
| 2,25E+07 | 1,65E+07 | 1,57E+07 | 1,62E+07 | 128,19 | 0 | 14,306 | 27 |
| 1,24E+08 | 1,64E+08 | 1,12E+08 | 1,62E+08 | 24,687 | 0 | 39,763 | 36 |
| 9,94E+08 | 1,16E+09 | 1,11E+09 | 1,17E+09 | 84,939 | 0 | 323,31 | 233 |
| 1,28E+09 | 1,78E+09 | 1,72E+09 | 1,63E+09 | 37,984 | 0 | 155,89 | 230 |
| 6,88E+07 | 8,31E+07 | 8,19E+07 | 7,87E+07 | 40,139 | 0 | 38,181 | 64 |
| 7,29E+07 | 1,35E+08 | 1,03E+08 | 1,02E+08 | 7,4158 | 0 | 9,1307 | 35 |
| 3,29E+07 | 2,97E+07 | 2,92E+07 | 3,08E+07 | 101,29 | 0 | 11,859 | 36 |
| 3,89E+06 | 2,78E+06 | 2,54E+06 | 2,58E+06 | 36,62 | 0,000159 | 5,2295 | 8 |
| 4,49E+07 | 3,91E+07 | 3,56E+07 | 3,25E+07 | 24,48 | 0 | 12,244 | 32 |
| 1,24E+09 | 1,24E+09 | 1,30E+09 | 1,31E+09 | 57,401 | 0 | 323,31 | 179 |
| 2,49E+07 | 8,10E+06 | 1,48E+07 | 1,08E+07 | 16,367 | 0 | 85,247 | 14 |
| 3,17E+06 | 3,09E+06 | 2,51E+06 | 2,85E+06 | 44,501 | 0,008204 | 1,5108 | 2 |
| 4,87E+06 | 1,74E+06 | 2,44E+06 | 4,47E+06 | 44,095 | 0,009619 | 1,4161 | 2 |
| 1,44E+07 | 9,14E+06 | 9,81E+06 | 7,76E+06 | 136,23 | 0 | 7,6399 | 23 |
| 5,88E+08 | 5,69E+08 | 5,68E+08 | 5,36E+08 | 62,907 | 0 | 197,83 | 175 |
| 4,77E+06 | 4,59E+06 | 7,18E+06 | 7,25E+06 | 61,531 | 0,001274 | 2,4548 | 5 |
| 1,92E+08 | 2,32E+08 | 2,42E+08 | 2,12E+08 | 94,371 | 0 | 158,7 | 101 |
| 3,49E+08 | 3,37E+08 | 3,65E+08 | 3,28E+08 | 28,031 | 0 | 42,955 | 63 |
| 6,79E+07 | 6,85E+07 | 5,53E+07 | 6,61E+07 | 67,903 | 0 | 23,801 | 37 |
| 1,03E+08 | 9,54E+07 | 6,90E+07 | 7,44E+07 | 26,172 | 0 | 15,315 | 33 |
| 3,30E+07 | 2,89E+07 | 3,09E+07 | 3,28E+07 | 73,215 | 0 | 17,4 | 39 |
| 1,18E+07 | 1,56E+07 | 1,63E+07 | 1,44E+07 | 35,607 | 0 | 16,826 | 17 |
| 2,82E+07 | 2,18E+07 | 2,18E+07 | 1,96E+07 | 197,19 | 0 | 14,219 | 38 |
| 7,23E+07 | 4,15E+07 | 4,75E+07 | 5,87E+07 | 28,055 | 0 | 13,177 | 36 |
| 2,65E+08 | 2,70E+08 | 2,29E+08 | 2,57E+08 | 58,675 | 0 | 147,43 | 115 |

| | | | | | | | |
|---|---|---|---|---|---|---|---|
| 9,24E+06 | 1,38E+07 | 1,62E+07 | 1,62E+07 | 16,069 | 0,000159 | 5,2224 | 5 |
| 4,14E+07 | 4,01E+07 | 4,02E+07 | 4,01E+07 | 44,687 | 0 | 11,861 | 39 |
| 2,17E+08 | 2,14E+08 | 2,12E+08 | 1,87E+08 | 34,537 | 0 | 42,393 | 112 |
| 3,37E+08 | 4,42E+08 | 4,31E+08 | 4,67E+08 | 30,381 | 0 | 323,31 | 74 |
| 4,03E+07 | 6,80E+07 | 4,79E+07 | 5,66E+07 | 57,321 | 0 | 9,8836 | 34 |
| 2,72E+06 | 6,10E+06 | 4,50E+06 | 8,49E+06 | 55,086 | 0,000157 | 4,8932 | 5 |
| 3,09E+07 | 2,30E+07 | 2,05E+07 | 2,61E+07 | 45,318 | 0,000158 | 5,0513 | 15 |
| 2,42E+08 | 2,22E+08 | 2,36E+08 | 2,40E+08 | 104,63 | 0 | 185,52 | 132 |
| 1,61E+07 | 1,45E+07 | 3,72E+06 | 1,64E+06 | 37,202 | 0 | 12,808 | 17 |
| 2,86E+06 | 4,47E+06 | 2,90E+06 | 3,37E+06 | 63,137 | 0,003442 | 1,9136 | 10 |
| 4,55E+08 | 5,25E+08 | 5,17E+08 | 5,02E+08 | 65,41 | 0 | 212,19 | 143 |
| 2,22E+08 | 2,00E+08 | 1,99E+08 | 2,37E+08 | 13,722 | 0 | 37,606 | 21 |
| 8,92E+07 | 8,74E+07 | 7,15E+07 | 7,89E+07 | 20,824 | 0 | 19,741 | 39 |
| 5,38E+08 | 5,41E+08 | 5,60E+08 | 5,60E+08 | 86,436 | 0 | 323,31 | 236 |
| 9,25E+07 | 9,41E+07 | 8,74E+07 | 9,40E+07 | 27,785 | 0 | 15,163 | 34 |
| 1,86E+07 | 7,57E+05 | 7,99E+06 | 1,22E+07 | 18,105 | 0 | 8,8086 | 11 |
| 3,19E+08 | 2,58E+08 | 2,64E+08 | 2,09E+08 | 49,911 | 0 | 164,68 | 78 |
| 1,55E+09 | 1,61E+09 | 1,71E+09 | 1,82E+09 | 67,314 | 0 | 323,31 | 235 |
| 9,95E+08 | 6,79E+08 | 9,44E+08 | 7,99E+08 | 14,552 | 0 | 95,199 | 31 |
| 2,63E+08 | 2,67E+08 | 2,30E+08 | 2,24E+08 | 60,518 | 0 | 100,97 | 129 |
| 1,78E+08 | 1,82E+08 | 2,05E+08 | 2,14E+08 | 56,802 | 0 | 323,31 | 66 |
| 1,70E+09 | 1,74E+09 | 1,66E+09 | 1,67E+09 | 87,265 | 0 | 323,31 | 362 |
| 4,80E+08 | 4,25E+08 | 3,77E+08 | 4,56E+08 | 125,98 | 0 | 252,78 | 179 |
| 1,93E+07 | 6,24E+06 | 8,42E+06 | 6,28E+06 | 37,986 | 0 | 152,3 | 10 |
| 2,26E+06 | 1,51E+07 | 1,87E+07 | 2,11E+07 | 34,004 | 0,000153 | 4,0943 | 11 |
| 1,04E+08 | 1,19E+08 | 1,13E+08 | 1,17E+08 | 83,693 | 0 | 51,726 | 96 |
| 3,20E+08 | 2,35E+08 | 2,82E+08 | 2,93E+08 | 18,186 | 0 | 8,925 | 27 |
| 2,48E+06 | 1,42E+07 | 1,20E+07 | 1,48E+07 | 17,478 | 0,003964 | 1,7846 | 6 |
| 2,58E+09 | 2,48E+09 | 2,54E+09 | 2,28E+09 | 54,54 | 0 | 323,31 | 540 |
| 1,09E+08 | 1,02E+08 | 8,92E+07 | 9,60E+07 | 38,737 | 0 | 74,223 | 62 |
| 5,40E+06 | 2,69E+06 | 3,97E+06 | 2,80E+06 | 54,851 | 0,000154 | 4,3634 | 8 |
| 4,69E+07 | 8,29E+07 | 7,50E+07 | 7,44E+07 | 47,552 | 0 | 41,285 | 69 |
| 3,34E+08 | 3,73E+08 | 3,95E+08 | 4,04E+08 | 101,98 | 0 | 217,6 | 179 |
| 1,71E+08 | 1,99E+08 | 1,93E+08 | 1,89E+08 | 134,11 | 0 | 104,1 | 181 |
| 1,35E+08 | 1,73E+08 | 1,53E+08 | 1,54E+08 | 76,003 | 0 | 53,414 | 82 |
| 1,42E+07 | 6,59E+06 | 7,82E+06 | 9,95E+06 | 35,776 | 0 | 53,182 | 8 |
| 4,53E+07 | 4,41E+07 | 4,84E+07 | 4,71E+07 | 158,36 | 0 | 24,486 | 41 |
| 8,05E+06 | 5,36E+06 | 5,54E+06 | 7,96E+06 | 50,735 | 0 | 6,7461 | 5 |
| 3,52E+08 | 2,93E+08 | 3,21E+08 | 3,44E+08 | 29,26 | 0 | 121,26 | 82 |
| 1,11E+08 | 1,27E+08 | 1,18E+08 | 1,11E+08 | 16,989 | 0 | 100,61 | 49 |
| 5,74E+06 | 3,78E+06 | 2,60E+06 | 4,85E+06 | 33,177 | 0,000157 | 4,8548 | 15 |
| 2,80E+07 | 3,40E+07 | 3,59E+07 | 3,70E+07 | 89,135 | 0 | 12,308 | 35 |
| 2,45E+07 | 8,02E+06 | 9,46E+06 | 1,17E+07 | 41,236 | 0 | 16,244 | 11 |
| 3,05E+07 | 3,82E+07 | 3,90E+07 | 3,51E+07 | 48,627 | 0 | 35,868 | 33 |
| 8,01E+07 | 7,20E+07 | 8,28E+07 | 7,58E+07 | 59,145 | 0 | 14,569 | 28 |
| 2,14E+07 | 2,41E+07 | 2,51E+07 | 2,21E+07 | 70,092 | 0 | 24,753 | 30 |
| 1,55E+08 | 1,56E+08 | 1,55E+08 | 1,53E+08 | 50,716 | 0 | 51,711 | 104 |
| 1,89E+09 | 1,51E+09 | 1,43E+09 | 1,49E+09 | 32,237 | 0 | 323,31 | 295 |
| 4,58E+06 | 4,47E+06 | 4,96E+06 | 2,91E+06 | 31,235 | 0,002783 | 2,0626 | 6 |
| 1,31E+07 | 5,77E+06 | 1,04E+07 | 1,06E+07 | 40,32 | 0,000852 | 2,5452 | 6 |

| | | | | | | | |
|---|---|---|---|---|---|---|---|
| 3,79E+07 | 3,10E+07 | 2,50E+07 | 1,56E+07 | 60,211 | 0 | 16,954 | 27 |
| 3,02E+06 | 2,43E+06 | 1,08E+06 | 2,34E+06 | 34,035 | 0 | 5,9492 | 6 |
| 1,14E+08 | 1,74E+08 | 1,43E+08 | 1,37E+08 | 31,593 | 0 | 56,908 | 43 |
| 2,14E+07 | 3,71E+07 | 2,14E+07 | 3,88E+07 | 64,327 | 0 | 7,157 | 20 |
| 1,53E+08 | 1,48E+08 | 1,24E+08 | 1,52E+08 | 60,644 | 0 | 95,275 | 57 |
| 2,05E+07 | 1,43E+07 | 1,66E+07 | 1,43E+07 | 97,291 | 0 | 6,9196 | 19 |
| 5,52E+06 | 7,79E+06 | 1,20E+07 | 8,44E+06 | 27,662 | 0,000149 | 3,525 | 8 |
| 1,54E+07 | 1,30E+07 | 1,44E+07 | 1,75E+07 | 69,207 | 0 | 29,623 | 25 |
| 2,19E+09 | 3,28E+09 | 3,18E+09 | 3,16E+09 | 8,2038 | 0 | 168,66 | 245 |
| 6,64E+06 | 1,22E+07 | 7,77E+06 | 1,01E+07 | 38,362 | 0,000153 | 4,1085 | 9 |
| 2,98E+07 | 4,42E+07 | 3,84E+07 | 3,29E+07 | 83,739 | 0 | 15,303 | 42 |
| 2,70E+06 | 3,77E+06 | 4,01E+06 | 4,25E+06 | 144,63 | 0,000155 | 4,5899 | 7 |
| 5,56E+07 | 9,41E+07 | 9,34E+07 | 9,40E+07 | 97,356 | 0,000156 | 4,6734 | 20 |
| 4,67E+06 | 2,76E+06 | 4,27E+06 | 3,37E+06 | 95,397 | 0,003176 | 1,9509 | 7 |
| 5,58E+06 | 1,51E+06 | 4,73E+06 | 2,68E+06 | 55,532 | 0,000435 | 2,9294 | 8 |
| 1,35E+07 | 1,90E+07 | 1,68E+07 | 1,48E+07 | 24,583 | 0 | 9,0248 | 23 |
| 1,39E+09 | 8,06E+08 | 9,95E+08 | 1,03E+09 | 34,357 | 0 | 323,31 | 317 |
| 1,61E+08 | 1,48E+08 | 1,60E+08 | 1,52E+08 | 61,271 | 0 | 35,296 | 64 |
| 5,54E+06 | 3,79E+07 | 5,83E+07 | 5,15E+07 | 35,856 | 0 | 40,194 | 15 |
| 1,41E+09 | 1,48E+09 | 1,42E+09 | 1,40E+09 | 63,988 | 0 | 323,31 | 277 |
| 2,86E+07 | 2,82E+07 | 2,86E+07 | 2,87E+07 | 50,829 | 0 | 29,958 | 23 |
| 4,28E+07 | 3,94E+07 | 3,22E+07 | 5,01E+07 | 25,766 | 0 | 23,634 | 32 |
| 3,69E+07 | 3,19E+07 | 2,78E+07 | 3,79E+07 | 20,101 | 0 | 18,489 | 23 |
| 2,20E+07 | 3,31E+07 | 2,94E+07 | 2,92E+07 | 66,99 | 0 | 10,577 | 28 |
| 5,46E+07 | 2,12E+07 | 2,03E+07 | 1,78E+07 | 59,411 | 0 | 17,389 | 26 |
| 1,35E+07 | 1,06E+07 | 1,28E+07 | 1,10E+07 | 32,108 | 0 | 9,0321 | 13 |
| 8,01E+08 | 8,58E+08 | 9,19E+08 | 8,24E+08 | 90,209 | 0 | 323,31 | 232 |
| 1,12E+08 | 1,26E+08 | 1,41E+08 | 1,30E+08 | 45,269 | 0 | 22,886 | 84 |
| 1,78E+07 | 2,23E+07 | 2,68E+07 | 2,84E+07 | 41,279 | 0 | 8,0158 | 24 |
| 2,99E+07 | 3,10E+07 | 3,14E+07 | 3,54E+07 | 83,42 | 0 | 6,9752 | 33 |
| 5,60E+07 | 3,91E+07 | 4,10E+07 | 4,09E+07 | 150,88 | 0 | 23,926 | 51 |
| 3,08E+08 | 3,62E+08 | 3,32E+08 | 2,66E+08 | 10,35 | 0 | 323,31 | 68 |
| 1,03E+07 | 7,59E+06 | 7,82E+06 | 8,55E+06 | 28,708 | 0 | 32,877 | 15 |
| 5,95E+08 | 5,01E+08 | 5,25E+08 | 5,47E+08 | 111,71 | 0 | 187,56 | 220 |
| 7,54E+06 | 4,56E+06 | 6,22E+06 | 3,98E+06 | 57,345 | 0,000852 | 2,5444 | 6 |
| 2,70E+09 | 2,10E+09 | 2,00E+09 | 1,95E+09 | 39,24 | 0 | 323,31 | 213 |
| 1,19E+07 | 1,46E+07 | 1,58E+07 | 1,79E+07 | 46,942 | 0,000147 | 3,2928 | 12 |
| 6,09E+09 | 3,51E+09 | 3,80E+09 | 3,21E+09 | 27,771 | 0 | 323,31 | 302 |
| 1,72E+07 | 1,69E+07 | 1,42E+07 | 1,65E+07 | 214,38 | 0 | 7,1619 | 24 |
| 2,33E+07 | 3,63E+07 | 3,19E+07 | 3,22E+07 | 18,75 | 0 | 74,287 | 16 |
| 2,39E+07 | 1,70E+07 | 2,01E+07 | 1,34E+07 | 196,53 | 0 | 14,867 | 23 |
| 4,38E+09 | 4,28E+09 | 5,18E+09 | 4,80E+09 | 13,509 | 0 | 176,82 | 177 |
| 6,95E+07 | 6,83E+07 | 7,83E+07 | 6,63E+07 | 46,942 | 0 | 78,04 | 30 |
| 6,47E+07 | 8,78E+07 | 7,86E+07 | 8,32E+07 | 24,576 | 0 | 136,24 | 56 |
| 1,29E+07 | 3,58E+06 | 3,40E+06 | 3,27E+06 | 46,597 | 0,000158 | 5,0545 | 5 |
| 8,68E+06 | 5,75E+06 | 6,84E+06 | 7,51E+06 | 34,912 | 0 | 9,5907 | 11 |
| 1,86E+07 | 1,83E+07 | 1,41E+07 | 1,13E+07 | 45,864 | 0 | 6,749 | 14 |
| 5,02E+06 | 1,77E+06 | 3,53E+06 | 3,15E+06 | 19,157 | 0,000571 | 2,6358 | 4 |
| 4,52E+08 | 4,21E+08 | 4,38E+08 | 3,87E+08 | 39,866 | 0 | 323,31 | 130 |
| 9,37E+08 | 9,35E+08 | 9,50E+08 | 9,22E+08 | 40,512 | 0 | 323,31 | 153 |

| | | | | | | | |
|---|---|---|---|---|---|---|---|
| 5,75E+07 | 7,15E+07 | 9,17E+07 | 8,22E+07 | 58,704 | 0 | 13,742 | 51 |
| 5,36E+07 | 5,27E+07 | 5,08E+07 | 5,39E+07 | 21,732 | 0 | 16,125 | 36 |
| 1,42E+08 | 1,25E+08 | 1,33E+08 | 1,44E+08 | 292 | 0 | 147,41 | 117 |
| 1,93E+08 | 1,80E+08 | 1,78E+08 | 2,00E+08 | 50,638 | 0 | 42,04 | 94 |
| 2,29E+08 | 2,56E+08 | 2,50E+08 | 2,24E+08 | 155,13 | 0 | 248,85 | 173 |
| 9,46E+07 | 1,38E+08 | 1,41E+08 | 1,51E+08 | 23,025 | 0 | 129,24 | 59 |
| 9,61E+06 | 1,15E+07 | 9,27E+06 | 9,39E+06 | 117,78 | 0 | 11,498 | 11 |
| 2,40E+07 | 2,05E+06 | 4,35E+06 | 3,49E+06 | 42,111 | 0,001272 | 2,4277 | 6 |
| 5,49E+07 | 8,12E+07 | 6,82E+07 | 6,55E+07 | 26,477 | 0,001686 | 2,28 | 13 |
| 4,33E+08 | 4,56E+08 | 5,32E+08 | 4,65E+08 | 18,672 | 0 | 96,851 | 55 |
| 5,43E+06 | 8,25E+06 | 6,30E+06 | 5,75E+06 | 63,391 | 0,000291 | 3,0344 | 5 |
| 1,15E+07 | 1,85E+07 | 1,56E+07 | 1,22E+07 | 55,603 | 0 | 6,2004 | 19 |
| 1,59E+07 | 1,55E+07 | 1,17E+07 | 1,14E+07 | 46,306 | 0 | 10,934 | 27 |
| 1,96E+08 | 2,12E+08 | 2,09E+08 | 1,98E+08 | 73,862 | 0 | 233,67 | 97 |
| 2,89E+07 | 4,52E+07 | 2,83E+07 | 4,58E+07 | 24,192 | 0 | 11,362 | 20 |
| 2,38E+07 | 2,08E+07 | 2,24E+07 | 2,27E+07 | 24,499 | 0 | 11,438 | 28 |
| 7,49E+06 | 4,74E+06 | 2,99E+06 | 6,46E+06 | 24,829 | 0,005812 | 1,6064 | 8 |
| 2,46E+08 | 2,64E+08 | 2,70E+08 | 2,77E+08 | 117,19 | 0 | 264,45 | 162 |
| 4,42E+09 | 3,48E+09 | 3,58E+09 | 3,33E+09 | 20,83 | 0 | 323,31 | 343 |
| 6,45E+06 | 1,51E+07 | 1,45E+07 | 1,37E+07 | 55,714 | 0,000151 | 3,7959 | 18 |
| 6,03E+07 | 6,69E+07 | 6,99E+07 | 7,27E+07 | 54,417 | 0 | 27,079 | 35 |
| 1,29E+08 | 1,73E+08 | 1,53E+08 | 1,61E+08 | 51,066 | 0 | 74,867 | 65 |
| 3,25E+08 | 2,46E+08 | 2,37E+08 | 2,48E+08 | 53,394 | 0 | 235,01 | 96 |
| 2,56E+09 | 1,99E+09 | 2,04E+09 | 1,96E+09 | 33,296 | 0 | 323,31 | 292 |
| 6,88E+07 | 7,19E+07 | 7,85E+07 | 6,95E+07 | 61,734 | 0 | 93,868 | 44 |
| 7,25E+07 | 6,53E+07 | 6,18E+07 | 6,19E+07 | 149,07 | 0 | 40,438 | 83 |
| 9,83E+06 | 7,57E+06 | 3,33E+06 | 6,72E+06 | 51,909 | 0 | 12,527 | 3 |
| 1,21E+06 | 1,52E+06 | 1,13E+06 | 1,26E+06 | 442,36 | 0,004633 | 1,7339 | 6 |
| 1,00E+08 | 1,42E+08 | 1,61E+08 | 1,39E+08 | 135,29 | 0 | 20,263 | 67 |
| 1,82E+08 | 1,91E+08 | 1,93E+08 | 2,01E+08 | 83,217 | 0 | 125,72 | 93 |
| 8,44E+07 | 4,69E+07 | 5,96E+07 | 6,56E+07 | 41,989 | 0 | 89,467 | 37 |
| 5,09E+07 | 4,15E+07 | 4,57E+07 | 3,56E+07 | 87,908 | 0 | 32,269 | 47 |
| 1,33E+08 | 9,62E+07 | 9,13E+07 | 9,33E+07 | 23,119 | 0 | 26,403 | 32 |
| 4,13E+07 | 2,00E+07 | 2,51E+07 | 3,71E+07 | 12,24 | 0 | 15,294 | 28 |
| 3,65E+07 | 6,51E+07 | 7,20E+07 | 7,99E+07 | 16,881 | 0 | 110,14 | 36 |
| 5,61E+08 | 5,56E+08 | 5,77E+08 | 5,55E+08 | 92,104 | 0 | 323,31 | 191 |
| 3,51E+08 | 3,09E+08 | 3,36E+08 | 3,39E+08 | 61,71 | 0 | 277 | 133 |
| 8,24E+07 | 4,34E+07 | 2,87E+07 | 5,14E+07 | 11,819 | 0 | 113,19 | 43 |
| 3,90E+08 | 3,60E+08 | 3,46E+08 | 2,93E+08 | 32,939 | 0 | 323,31 | 95 |
| 1,04E+08 | 1,03E+08 | 1,08E+08 | 9,94E+07 | 160,96 | 0 | 71,859 | 97 |
| 1,42E+07 | 2,27E+07 | 3,16E+07 | 2,84E+07 | 17,667 | 0 | 64,484 | 18 |
| 3,02E+08 | 3,78E+08 | 4,51E+08 | 4,89E+08 | 52,149 | 0 | 244,5 | 151 |
| 3,72E+08 | 4,71E+08 | 4,69E+08 | 4,58E+08 | 82,876 | 0 | 162,4 | 152 |
| 1,44E+08 | 6,66E+07 | 6,67E+07 | 9,74E+07 | 26,313 | 0 | 323,31 | 35 |
| 5,76E+06 | 7,38E+06 | 7,42E+06 | 7,89E+06 | 52,465 | 0,00043 | 2,7447 | 10 |
| 3,07E+07 | 2,88E+07 | 3,35E+07 | 3,25E+07 | 44,41 | 0 | 21,914 | 31 |
| 3,72E+08 | 3,10E+08 | 3,22E+08 | 3,04E+08 | 112,62 | 0 | 174,06 | 174 |
| 6,03E+07 | 4,02E+07 | 5,91E+07 | 5,18E+07 | 87,044 | 0 | 51,554 | 42 |
| 2,94E+08 | 3,71E+08 | 3,71E+08 | 3,55E+08 | 60,824 | 0 | 108,86 | 98 |
| 6,41E+07 | 8,78E+07 | 8,12E+07 | 7,34E+07 | 43,771 | 0 | 49,645 | 59 |

| | | | | | | | |
|---|---|---|---|---|---|---|---|
| 2,68E+08 | 1,76E+08 | 2,06E+08 | 1,24E+08 | 11,845 | 0 | 143,94 | 26 |
| 1,19E+08 | 6,41E+06 | 5,34E+07 | 8,11E+06 | 55,278 | 0 | 23,277 | 22 |
| 6,07E+06 | 7,68E+06 | 7,67E+06 | 6,20E+06 | 42,514 | 0,000146 | 3,1191 | 17 |
| 1,87E+07 | 2,48E+07 | 3,78E+07 | 3,55E+07 | 30,335 | 0 | 10,7 | 25 |
| 6,57E+08 | 6,82E+08 | 7,04E+08 | 6,85E+08 | 64,124 | 0 | 296,36 | 184 |
| 1,24E+08 | 1,10E+08 | 1,08E+08 | 1,25E+08 | 47,147 | 0 | 51,888 | 36 |
| 3,39E+08 | 4,11E+08 | 3,44E+08 | 3,38E+08 | 155,59 | 0 | 323,31 | 174 |
| 1,01E+08 | 9,16E+07 | 1,01E+08 | 1,02E+08 | 83,794 | 0 | 29,395 | 73 |
| 3,07E+06 | 7,09E+06 | 3,82E+06 | 3,65E+06 | 94,315 | 0 | 6,5111 | 12 |
| 9,02E+07 | 8,82E+07 | 8,89E+07 | 8,54E+07 | 133,47 | 0 | 53,089 | 71 |
| 1,63E+07 | 1,54E+07 | 1,57E+07 | 1,37E+07 | 161,35 | 0 | 6,394 | 15 |
| 7,42E+06 | 5,98E+06 | 7,36E+06 | 6,81E+06 | 65,505 | 0 | 14,503 | 10 |
| 3,99E+07 | 6,00E+07 | 5,45E+07 | 6,35E+07 | 39,991 | 0 | 15,849 | 47 |
| 2,60E+06 | 5,43E+06 | 3,11E+06 | 5,46E+06 | 59,651 | 0,000158 | 5,0664 | 5 |
| 4,39E+06 | 1,90E+06 | 4,02E+06 | 3,11E+06 | 56,043 | 0,001275 | 2,459 | 3 |
| 4,68E+06 | 2,65E+06 | 2,26E+06 | 3,01E+06 | 74,303 | 0 | 5,7616 | 7 |
| 5,73E+07 | 5,34E+07 | 4,75E+07 | 4,16E+07 | 28,991 | 0 | 30,24 | 45 |
| 2,71E+07 | 5,43E+07 | 5,16E+07 | 6,06E+07 | 33,794 | 0 | 46,42 | 29 |
| 8,55E+07 | 7,35E+07 | 7,33E+07 | 7,36E+07 | 37,809 | 0 | 31,733 | 55 |
| 9,44E+08 | 1,30E+09 | 1,10E+09 | 1,10E+09 | 25,378 | 0 | 311,73 | 147 |
| 2,03E+07 | 2,80E+07 | 2,19E+07 | 2,82E+07 | 12,403 | 0 | 13,383 | 13 |
| 1,91E+08 | 1,86E+08 | 1,74E+08 | 1,83E+08 | 55,586 | 0 | 109,97 | 78 |
| 1,04E+08 | 1,53E+08 | 1,41E+08 | 1,44E+08 | 28,834 | 0 | 9,8514 | 41 |
| 1,97E+07 | 4,33E+07 | 3,67E+07 | 5,81E+07 | 169,12 | 0,009487 | 1,4184 | 10 |
| 7,28E+06 | 3,13E+06 | 1,44E+06 | 4,47E+06 | 59,267 | 0,000154 | 4,3286 | 4 |
| 9,24E+07 | 9,09E+07 | 9,46E+07 | 9,71E+07 | 95,91 | 0 | 37,121 | 68 |
| 8,60E+07 | 9,62E+07 | 8,83E+07 | 8,86E+07 | 46,325 | 0 | 43,339 | 96 |
| 2,77E+09 | 3,00E+09 | 3,07E+09 | 3,14E+09 | 47,688 | 0 | 323,31 | 323 |
| 2,64E+07 | 4,42E+07 | 5,61E+07 | 5,77E+07 | 28,787 | 0,000292 | 3,0705 | 14 |
| 9,39E+07 | 8,21E+07 | 7,02E+07 | 7,17E+07 | 38,341 | 0 | 25,482 | 52 |
| 1,01E+07 | 8,03E+06 | 6,45E+06 | 6,15E+06 | 115,02 | 0,000433 | 2,8339 | 10 |
| 3,51E+07 | 2,89E+07 | 3,56E+07 | 3,42E+07 | 82,063 | 0 | 23,609 | 33 |
| 1,99E+08 | 2,99E+08 | 2,78E+08 | 3,36E+08 | 27,453 | 0 | 179,64 | 87 |
| 1,97E+06 | 5,04E+06 | 4,81E+06 | 3,98E+06 | 107,37 | 0,003179 | 1,9619 | 3 |
| 1,27E+08 | 9,76E+07 | 1,18E+08 | 1,13E+08 | 55,957 | 0 | 55,602 | 74 |
| 5,52E+07 | 4,93E+07 | 4,69E+07 | 4,86E+07 | 67,554 | 0 | 23,806 | 66 |
| 2,98E+09 | 2,82E+09 | 2,94E+09 | 2,97E+09 | 135,55 | 0 | 323,31 | 511 |
| 1,24E+08 | 1,13E+08 | 1,16E+08 | 1,09E+08 | 38,028 | 0 | 115,32 | 41 |
| 2,72E+08 | 2,95E+08 | 2,87E+08 | 2,71E+08 | 29,546 | 0 | 323,31 | 102 |
| 4,46E+08 | 5,54E+08 | 4,72E+08 | 4,69E+08 | 33,88 | 0 | 163,96 | 213 |
| 2,64E+08 | 2,69E+08 | 2,66E+08 | 2,61E+08 | 130,89 | 0 | 323,31 | 184 |
| 7,37E+07 | 6,10E+07 | 7,04E+07 | 5,75E+07 | 23,416 | 0 | 87,262 | 51 |
| 4,03E+07 | 5,19E+07 | 5,24E+07 | 4,33E+07 | 83,924 | 0 | 41,394 | 61 |
| 4,40E+08 | 4,00E+08 | 3,85E+08 | 4,63E+08 | 22,187 | 0 | 215,15 | 82 |
| 2,20E+07 | 1,06E+07 | 1,15E+07 | 1,81E+07 | 14,235 | 0,000156 | 4,6157 | 14 |
| 2,99E+07 | 3,75E+07 | 3,32E+07 | 2,77E+07 | 41,049 | 0 | 10,525 | 25 |
| 4,26E+06 | 9,82E+06 | 7,40E+06 | 6,90E+06 | 92,37 | 0,001686 | 2,2878 | 10 |
| 2,08E+08 | 1,87E+08 | 1,71E+08 | 1,50E+08 | 44,59 | 0 | 123,9 | 97 |
| 5,02E+08 | 4,24E+08 | 4,22E+08 | 3,79E+08 | 88,382 | 0 | 323,31 | 209 |
| 1,63E+08 | 1,75E+08 | 1,46E+08 | 1,43E+08 | 198,92 | 0 | 55,11 | 128 |

| | | | | | | | |
|---|---|---|---|---|---|---|---|
| 4,63E+06 | 1,49E+06 | 2,68E+06 | 5,03E+06 | 44,591 | 0 | 12,09 | 6 |
| 2,34E+09 | 2,37E+09 | 2,35E+09 | 2,29E+09 | 55,238 | 0 | 323,31 | 279 |
| 6,35E+07 | 5,80E+07 | 6,71E+07 | 6,07E+07 | 99,104 | 0 | 61,224 | 66 |
| 1,25E+07 | 1,11E+07 | 1,09E+07 | 7,22E+06 | 46,794 | 0 | 12,501 | 15 |
| 1,56E+08 | 1,46E+08 | 1,25E+08 | 1,60E+08 | 34,278 | 0 | 91,292 | 85 |
| 6,39E+07 | 6,52E+07 | 6,42E+07 | 6,48E+07 | 61,916 | 0 | 60,938 | 45 |
| 1,88E+07 | 2,40E+07 | 2,90E+07 | 2,66E+07 | 27,283 | 0,001825 | 2,2568 | 9 |
| 3,39E+06 | 6,22E+06 | 4,42E+06 | 5,63E+06 | 83,56 | 0,009752 | 1,4146 | 4 |
| 1,67E+08 | 1,29E+08 | 1,30E+08 | 1,25E+08 | 29,127 | 0 | 56,643 | 41 |
| 1,03E+08 | 1,73E+08 | 1,72E+08 | 1,68E+08 | 49,981 | 0 | 66,664 | 92 |
| 1,87E+07 | 1,91E+07 | 1,82E+07 | 1,65E+07 | 242,25 | 0 | 60,329 | 25 |
| 3,99E+08 | 7,28E+08 | 1,10E+09 | 8,03E+08 | 11,083 | 0 | 169,66 | 229 |
| 2,02E+07 | 2,11E+07 | 1,59E+07 | 1,89E+07 | 18,766 | 0 | 9,1779 | 17 |
| 8,53E+07 | 9,61E+07 | 9,72E+07 | 1,03E+08 | 66,556 | 0 | 40,428 | 62 |
| 2,49E+06 | 1,52E+07 | 5,02E+07 | 1,18E+08 | 65,432 | 0 | 24,299 | 27 |
| 2,25E+08 | 2,14E+08 | 2,17E+08 | 2,22E+08 | 74,648 | 0 | 264,03 | 143 |
| 4,55E+07 | 3,99E+07 | 3,46E+07 | 3,48E+07 | 33,543 | 0 | 27,869 | 25 |
| 1,11E+07 | 3,60E+07 | 2,61E+07 | 4,05E+07 | 392,33 | 0 | 24,821 | 49 |
| 1,50E+09 | 1,45E+09 | 1,33E+09 | 1,17E+09 | 47,655 | 0 | 323,31 | 202 |
| 6,39E+07 | 7,91E+07 | 7,82E+07 | 7,04E+07 | 17,261 | 0 | 6,521 | 15 |
| 6,42E+06 | 6,28E+06 | 2,51E+06 | 2,20E+06 | 49,192 | 0,004366 | 1,7535 | 4 |
| 9,39E+06 | 3,96E+06 | 4,80E+06 | 7,83E+05 | 57,037 | 0 | 14,434 | 7 |
| 2,44E+08 | 3,51E+08 | 3,34E+08 | 3,28E+08 | 106,6 | 0 | 84,931 | 185 |
| 2,34E+08 | 3,02E+08 | 3,13E+08 | 2,66E+08 | 18,255 | 0 | 193,31 | 53 |
| 6,55E+07 | 3,97E+07 | 4,07E+07 | 4,34E+07 | 55,504 | 0 | 15,523 | 43 |
| 4,62E+07 | 3,34E+07 | 3,54E+07 | 3,13E+07 | 179,33 | 0 | 25,055 | 50 |
| 1,73E+08 | 1,52E+08 | 1,56E+08 | 1,36E+08 | 144,44 | 0 | 122,79 | 122 |
| 3,37E+07 | 3,96E+07 | 4,26E+07 | 4,08E+07 | 35,988 | 0 | 32,913 | 32 |
| 1,52E+08 | 1,19E+08 | 1,35E+08 | 1,40E+08 | 110,03 | 0 | 113,8 | 86 |
| 3,42E+07 | 1,84E+07 | 3,34E+07 | 2,55E+07 | 28,391 | 0,000153 | 4,2023 | 20 |
| 2,38E+08 | 2,63E+08 | 2,47E+08 | 2,77E+08 | 37,605 | 0 | 214,75 | 91 |
| 2,41E+07 | 4,74E+07 | 4,00E+07 | 4,11E+07 | 67,334 | 0 | 21,436 | 39 |
| 4,03E+07 | 4,63E+07 | 4,64E+07 | 4,67E+07 | 45,658 | 0 | 66,013 | 31 |
| 5,21E+06 | 2,92E+06 | 1,79E+06 | 2,03E+06 | 43,585 | 0,004627 | 1,7218 | 2 |
| 7,19E+06 | 1,26E+07 | 1,20E+07 | 1,67E+07 | 20,891 | 0 | 25,804 | 14 |
| 7,32E+07 | 8,69E+07 | 6,83E+07 | 8,05E+07 | 14,544 | 0 | 20,915 | 29 |
| 8,03E+07 | 1,06E+08 | 9,67E+07 | 9,94E+07 | 50,835 | 0 | 17,893 | 33 |
| 6,79E+07 | 6,30E+07 | 5,84E+07 | 5,55E+07 | 36,674 | 0 | 60,68 | 23 |
| 1,00E+08 | 9,72E+07 | 1,13E+08 | 1,06E+08 | 72,728 | 0 | 114,76 | 68 |
| 1,62E+08 | 1,22E+08 | 1,36E+08 | 1,27E+08 | 30,597 | 0 | 201,32 | 66 |
| 1,34E+07 | 4,19E+07 | 4,76E+07 | 5,51E+07 | 120,28 | 0 | 9,9614 | 31 |
| 1,01E+07 | 8,67E+06 | 1,68E+07 | 1,46E+07 | 35,593 | 0 | 7,0797 | 22 |
| 5,32E+07 | 5,01E+06 | 3,60E+06 | 2,51E+06 | 82,978 | 0,003701 | 1,8435 | 7 |
| 1,13E+08 | 1,47E+08 | 1,46E+08 | 1,62E+08 | 88,179 | 0 | 92,467 | 99 |
| 1,08E+08 | 1,23E+08 | 1,31E+08 | 1,44E+08 | 71,292 | 0 | 53,05 | 80 |
| 1,49E+08 | 1,48E+08 | 1,49E+08 | 1,52E+08 | 111,84 | 0 | 81,363 | 86 |
| 7,06E+07 | 7,04E+07 | 7,28E+07 | 6,77E+07 | 32,373 | 0 | 42,177 | 47 |
| 3,42E+07 | 3,68E+07 | 3,75E+07 | 3,51E+07 | 21,802 | 0 | 13,034 | 29 |
| 4,98E+07 | 5,41E+07 | 3,56E+07 | 4,35E+07 | 22,321 | 0 | 46,386 | 34 |
| 2,49E+07 | 2,10E+07 | 1,55E+07 | 1,07E+07 | 19,796 | 0 | 115,24 | 37 |

| | | | | | | | |
|---|---|---|---|---|---|---|---|
| 2,74E+07 | 1,22E+07 | 1,31E+07 | 1,41E+07 | 82,998 | 0 | 15,658 | 22 |
| 1,83E+08 | 1,52E+08 | 1,73E+08 | 1,56E+08 | 21,39 | 0 | 232,67 | 69 |
| 1,16E+07 | 9,50E+07 | 1,46E+08 | 1,32E+08 | 49,909 | 0 | 323,31 | 30 |
| 3,71E+08 | 3,54E+08 | 4,15E+08 | 3,93E+08 | 17,356 | 0 | 323,31 | 89 |
| 5,24E+07 | 2,93E+07 | 2,66E+07 | 3,24E+07 | 21,491 | 0 | 19,653 | 20 |
| 1,19E+08 | 1,13E+08 | 1,22E+08 | 9,96E+07 | 38,968 | 0 | 28,085 | 59 |
| 2,15E+06 | 1,25E+07 | 1,22E+07 | 1,33E+07 | 83,693 | 0,000149 | 3,4606 | 7 |
| 1,21E+08 | 9,91E+07 | 9,00E+07 | 9,17E+07 | 200,8 | 0 | 62,441 | 93 |
| 5,28E+07 | 4,62E+07 | 4,35E+07 | 3,85E+07 | 129,99 | 0 | 25,181 | 53 |
| 3,44E+07 | 3,97E+07 | 4,56E+07 | 3,82E+07 | 62,875 | 0 | 24,291 | 41 |
| 7,02E+07 | 5,45E+07 | 7,82E+07 | 6,95E+07 | 13,251 | 0 | 34,541 | 44 |
| 1,88E+08 | 2,37E+08 | 2,42E+08 | 2,25E+08 | 105,61 | 0 | 160,88 | 128 |
| 3,67E+06 | 1,31E+07 | 1,14E+07 | 1,31E+07 | 17,504 | 0 | 11,682 | 11 |
| 3,81E+08 | 3,18E+08 | 3,89E+08 | 3,54E+08 | 32,342 | 0 | 60,251 | 74 |
| 1,57E+07 | 1,56E+07 | 1,93E+07 | 1,96E+07 | 23,275 | 0,000151 | 3,8034 | 12 |
| 2,58E+07 | 2,50E+07 | 1,35E+07 | 2,16E+07 | 51,421 | 0 | 13,366 | 28 |
| 2,69E+07 | 3,39E+07 | 3,34E+07 | 2,86E+07 | 78,832 | 0 | 40,332 | 23 |
| 1,21E+07 | 1,30E+07 | 1,48E+07 | 1,92E+07 | 74,091 | 0 | 8,9123 | 22 |
| 7,76E+06 | 5,22E+06 | 7,43E+06 | 9,35E+06 | 52,743 | 0,000148 | 3,3609 | 10 |
| 6,37E+08 | 6,52E+08 | 8,14E+08 | 7,45E+08 | 17,982 | 0 | 190,48 | 57 |
| 6,37E+07 | 6,10E+07 | 7,19E+07 | 7,16E+07 | 34,085 | 0 | 77,077 | 27 |
| 1,60E+07 | 1,62E+07 | 2,06E+07 | 2,17E+07 | 23,848 | 0 | 6,3063 | 14 |
| 1,25E+08 | 1,92E+08 | 1,51E+08 | 1,76E+08 | 60,621 | 0 | 190,59 | 75 |
| 6,70E+06 | 3,43E+06 | 4,09E+06 | 3,29E+06 | 35,502 | 0,000147 | 3,1922 | 3 |
| 5,36E+06 | 3,95E+06 | 2,23E+06 | 3,31E+06 | 88,057 | 0,000155 | 4,4487 | 6 |
| 4,81E+07 | 2,87E+07 | 3,69E+07 | 2,92E+07 | 41,666 | 0 | 50,194 | 28 |
| 1,27E+08 | 1,09E+08 | 1,07E+08 | 1,01E+08 | 101,52 | 0 | 48,426 | 97 |
| 3,34E+07 | 3,50E+07 | 3,06E+07 | 3,35E+07 | 59,173 | 0 | 16,113 | 29 |
| 5,25E+07 | 5,93E+07 | 5,21E+07 | 5,76E+07 | 26,03 | 0 | 16,782 | 33 |
| 2,38E+07 | 1,58E+07 | 1,58E+07 | 2,12E+07 | 53,693 | 0 | 7,6439 | 21 |
| 8,80E+06 | 4,73E+06 | 5,83E+06 | 7,97E+06 | 81,75 | 0 | 7,1692 | 10 |
| 4,11E+08 | 3,72E+08 | 3,68E+08 | 3,04E+08 | 52,625 | 0 | 80,887 | 139 |
| 1,99E+07 | 1,67E+07 | 1,07E+07 | 1,13E+07 | 41,877 | 0 | 18,016 | 22 |
| 1,23E+07 | 7,54E+06 | 6,35E+06 | 1,16E+07 | 26,297 | 0 | 8,1131 | 11 |
| 1,56E+07 | 1,22E+07 | 1,81E+06 | 1,29E+07 | 48,379 | 0,000153 | 4,2098 | 11 |
| 6,22E+07 | 5,16E+07 | 4,77E+07 | 5,64E+07 | 82,043 | 0 | 39,101 | 54 |
| 1,11E+09 | 1,27E+09 | 1,22E+09 | 1,24E+09 | 151,08 | 0 | 323,31 | 366 |
| 1,34E+09 | 1,36E+09 | 1,34E+09 | 1,38E+09 | 225,63 | 0 | 323,31 | 541 |
| 1,75E+07 | 1,61E+07 | 2,08E+07 | 1,42E+07 | 19,686 | 0 | 8,2958 | 13 |
| 1,52E+07 | 1,94E+07 | 1,79E+07 | 1,67E+07 | 50,88 | 0 | 13,445 | 20 |
| 1,45E+06 | 1,82E+06 | 2,37E+06 | 1,91E+06 | 94,202 | 0,001685 | 2,275 | 8 |
| 8,41E+08 | 7,38E+08 | 7,23E+08 | 7,19E+08 | 59,152 | 0 | 323,31 | 172 |
| 1,36E+07 | 1,33E+07 | 7,81E+06 | 1,41E+07 | 47,234 | 0 | 7,9911 | 23 |
| 4,49E+06 | 7,31E+06 | 4,55E+06 | 4,89E+06 | 40,56 | 0 | 6,5197 | 7 |
| 3,12E+06 | 4,89E+06 | 4,81E+06 | 4,62E+06 | 40,633 | 0,000156 | 4,6527 | 12 |
| 6,99E+06 | 8,32E+06 | 8,25E+06 | 5,06E+06 | 118,53 | 0,00015 | 3,7379 | 10 |
| 3,24E+06 | 2,78E+06 | 3,26E+06 | 3,37E+06 | 32,64 | 0,001274 | 2,4526 | 9 |
| 1,56E+07 | 1,58E+07 | 1,48E+07 | 1,39E+07 | 54,26 | 0 | 14,191 | 15 |
| 1,14E+07 | 3,82E+06 | 1,85E+07 | 5,17E+06 | 47,38 | 0,003444 | 1,9234 | 8 |
| 7,30E+07 | 8,01E+07 | 6,63E+07 | 6,89E+07 | 42,758 | 0 | 85,45 | 54 |

| | | | | | | | |
|---|---|---|---|---|---|---|---|
| 1,91E+09 | 2,18E+09 | 6,30E+09 | 3,17E+09 | 12,254 | 0 | 323,31 | 452 |
| 1,00E+07 | 4,87E+06 | 4,90E+06 | 1,09E+07 | 111,86 | 0 | 9,7091 | 19 |
| 1,57E+06 | 4,19E+06 | 4,82E+06 | 4,51E+06 | 51,385 | 0,005814 | 1,61 | 9 |
| 1,48E+08 | 1,13E+08 | 1,28E+08 | 9,84E+07 | 10,214 | 0 | 25,926 | 55 |
| 3,32E+07 | 1,66E+07 | 2,02E+07 | 1,85E+07 | 21,11 | 0 | 49,453 | 15 |
| 1,24E+08 | 1,32E+08 | 1,12E+08 | 1,16E+08 | 27,403 | 0 | 47,123 | 54 |
| 1,76E+08 | 1,81E+08 | 1,54E+08 | 1,75E+08 | 20,463 | 0 | 92,308 | 40 |
| 4,53E+06 | 9,58E+05 | 2,61E+06 | 3,84E+06 | 44,207 | 0,000148 | 3,3142 | 5 |
| 3,07E+06 | 6,87E+06 | 5,29E+06 | 2,62E+06 | 57,401 | 0,007279 | 1,5423 | 3 |
| 2,54E+08 | 3,30E+08 | 3,02E+08 | 3,13E+08 | 25,238 | 0 | 113,25 | 81 |
| 5,56E+06 | 1,83E+06 | 3,78E+06 | 8,30E+06 | 41,99 | 0 | 9,5032 | 10 |
| 1,42E+09 | 2,08E+09 | 3,53E+09 | 8,19E+08 | 10,051 | 0 | 323,31 | 458 |
| 5,00E+07 | 4,70E+07 | 4,66E+07 | 3,61E+07 | 38,381 | 0 | 34,666 | 53 |
| 5,98E+06 | 2,18E+07 | 1,24E+07 | 1,32E+07 | 58,699 | 0,000148 | 3,3693 | 6 |
| 8,17E+08 | 8,36E+08 | 7,63E+08 | 7,49E+08 | 144,8 | 0 | 323,31 | 301 |
| 8,32E+06 | 1,82E+07 | 1,38E+07 | 2,13E+07 | 77,524 | 0 | 6,4778 | 16 |
| 2,64E+07 | 2,17E+07 | 2,11E+07 | 2,26E+07 | 21,737 | 0 | 80,487 | 16 |
| 6,12E+06 | 2,50E+06 | 2,02E+06 | 2,94E+06 | 39,106 | 0,003046 | 1,9964 | 2 |
| 1,86E+07 | 1,77E+07 | 3,01E+07 | 1,57E+07 | 31,852 | 0 | 5,5092 | 17 |
| 3,48E+06 | 8,84E+05 | 2,85E+06 | 1,65E+06 | 136,76 | 0,003441 | 1,9115 | 6 |
| 1,86E+06 | 2,84E+06 | 1,03E+07 | 9,83E+06 | 79,159 | 0 | 16,073 | 6 |
| 7,19E+06 | 1,02E+07 | 7,04E+06 | 7,75E+06 | 26,596 | 0,00015 | 3,6673 | 12 |
| 1,11E+07 | 1,54E+07 | 1,58E+07 | 1,70E+07 | 139,3 | 0 | 9,3922 | 22 |
| 2,50E+07 | 4,20E+07 | 2,87E+07 | 2,87E+07 | 145,66 | 0,000157 | 4,8989 | 13 |
| 1,59E+07 | 1,80E+07 | 1,41E+07 | 2,15E+07 | 48,758 | 0 | 120,66 | 21 |
| 1,08E+09 | 1,13E+09 | 1,02E+09 | 1,09E+09 | 41,063 | 0 | 323,31 | 131 |
| 6,98E+07 | 8,44E+07 | 8,36E+07 | 8,89E+07 | 60,132 | 0 | 199,32 | 51 |
| 1,50E+09 | 1,99E+09 | 1,67E+09 | 1,85E+09 | 42,444 | 0 | 154,52 | 161 |
| 2,61E+08 | 3,35E+08 | 3,12E+08 | 3,13E+08 | 50,197 | 0 | 70,718 | 141 |
| 1,08E+07 | 1,96E+07 | 3,08E+07 | 3,28E+07 | 44,905 | 0 | 13,397 | 35 |
| 7,71E+08 | 7,97E+08 | 7,93E+08 | 7,29E+08 | 60,866 | 0 | 89,822 | 148 |
| 2,04E+07 | 7,37E+06 | 1,97E+06 | 4,04E+06 | 27,167 | 0 | 123,19 | 3 |
| 1,03E+09 | 1,27E+09 | 1,24E+09 | 1,15E+09 | 104,58 | 0 | 323,31 | 254 |
| 9,18E+07 | 9,95E+07 | 9,49E+07 | 8,57E+07 | 46,119 | 0 | 28,118 | 47 |
| 7,56E+06 | 5,71E+06 | 1,13E+06 | 1,06E+06 | 74,152 | 0,003699 | 1,8333 | 3 |
| 3,80E+06 | 3,20E+06 | 5,03E+06 | 5,22E+06 | 25,081 | 0,000147 | 3,1468 | 3 |
| 3,76E+07 | 4,64E+07 | 4,49E+07 | 5,26E+07 | 108,23 | 0 | 23,415 | 32 |
| 1,47E+08 | 1,63E+08 | 1,52E+08 | 1,43E+08 | 95,584 | 0 | 72,061 | 98 |
| 1,21E+08 | 1,35E+08 | 1,27E+08 | 1,24E+08 | 57,506 | 0 | 84,122 | 79 |
| 1,44E+08 | 1,16E+08 | 1,26E+08 | 1,15E+08 | 23,159 | 0 | 242 | 53 |
| 1,23E+08 | 1,57E+08 | 1,57E+08 | 1,52E+08 | 32,313 | 0 | 116,39 | 53 |
| 4,85E+07 | 3,45E+07 | 3,68E+07 | 4,76E+07 | 28,135 | 0 | 7,0357 | 17 |
| 3,05E+07 | 1,49E+08 | 6,43E+07 | 9,89E+07 | 83,355 | 0,000432 | 2,8196 | 7 |
| 1,42E+09 | 1,51E+09 | 1,45E+09 | 1,56E+09 | 139,88 | 0 | 323,31 | 342 |
| 2,48E+06 | 4,52E+06 | 6,31E+06 | 5,44E+06 | 152,47 | 0,000853 | 2,5489 | 5 |
| 2,62E+08 | 2,96E+08 | 2,70E+08 | 2,80E+08 | 53,133 | 0 | 297,2 | 126 |
| 6,67E+08 | 8,48E+08 | 8,17E+08 | 8,28E+08 | 31,775 | 0 | 274,83 | 227 |
| 1,29E+07 | 1,12E+07 | 1,22E+07 | 8,05E+06 | 41,245 | 0,000157 | 4,8203 | 11 |
| 1,60E+08 | 1,32E+08 | 1,18E+08 | 1,37E+08 | 14,255 | 0 | 20,688 | 50 |
| 1,00E+07 | 1,06E+07 | 9,97E+06 | 8,77E+06 | 61,521 | 0 | 5,8111 | 17 |

| | | | | | | | |
|---|---|---|---|---|---|---|---|
| 3,27E+08 | 3,40E+08 | 3,17E+08 | 3,16E+08 | 84,921 | 0 | 161,32 | 177 |
| 3,88E+07 | 3,52E+07 | 3,70E+07 | 2,88E+07 | 100,3 | 0 | 30,086 | 41 |
| 9,18E+07 | 2,60E+08 | 2,54E+08 | 3,20E+08 | 73,427 | 0,000854 | 2,5765 | 38 |
| 3,05E+07 | 2,36E+07 | 3,43E+07 | 3,39E+07 | 14,267 | 0 | 14,473 | 29 |
| 1,21E+08 | 9,80E+07 | 9,45E+07 | 1,06E+08 | 23,261 | 0 | 161,95 | 59 |
| 4,49E+07 | 2,44E+07 | 3,74E+07 | 4,48E+07 | 36,513 | 0 | 62,108 | 37 |
| 1,93E+06 | 6,94E+06 | 4,09E+06 | 2,32E+06 | 42,918 | 0,000433 | 2,8474 | 3 |
| 9,05E+06 | 4,26E+07 | 5,70E+07 | 3,93E+07 | 161,12 | 0,000431 | 2,7662 | 9 |
| 1,01E+07 | 4,81E+06 | 1,22E+07 | 1,09E+06 | 22,807 | 0 | 6,2583 | 6 |
| 1,70E+08 | 2,55E+08 | 2,46E+08 | 2,29E+08 | 35,44 | 0 | 96,814 | 92 |
| 1,58E+08 | 1,69E+08 | 1,63E+08 | 1,47E+08 | 24,936 | 0 | 132,55 | 27 |
| 2,37E+06 | 8,36E+06 | 6,98E+06 | 9,50E+06 | 88,344 | 0,000149 | 3,4709 | 12 |
| 2,02E+08 | 2,01E+08 | 2,02E+08 | 2,12E+08 | 96,428 | 0 | 84,679 | 149 |
| 6,83E+06 | 4,05E+06 | 1,40E+06 | 1,98E+06 | 56,141 | 0,002911 | 2,0109 | 2 |
| 2,96E+06 | 2,70E+06 | 1,96E+06 | 3,15E+06 | 132,87 | 0,002782 | 2,0611 | 7 |
| 2,50E+07 | 3,05E+07 | 3,51E+07 | 2,91E+07 | 53,324 | 0 | 21,395 | 34 |
| 1,12E+08 | 1,12E+08 | 9,80E+07 | 1,11E+08 | 31,129 | 0 | 152,63 | 83 |
| 1,88E+07 | 7,19E+06 | 7,97E+06 | 1,73E+06 | 46,625 | 0 | 63,461 | 6 |
| 4,74E+07 | 8,09E+07 | 1,01E+08 | 9,58E+07 | 59,948 | 0 | 131,96 | 45 |
| 2,61E+09 | 2,46E+09 | 2,72E+09 | 2,42E+09 | 136,7 | 0 | 50,94 | 113 |
| 7,89E+05 | 1,83E+06 | 3,10E+06 | 8,35E+05 | 76,255 | 0,009489 | 1,4218 | 4 |
| 7,52E+08 | 9,70E+08 | 9,55E+08 | 9,25E+08 | 39,93 | 0 | 141,67 | 115 |
| 2,58E+08 | 2,68E+08 | 2,68E+08 | 2,37E+08 | 26,585 | 0 | 272,16 | 96 |
| 8,69E+06 | 3,44E+06 | 3,94E+06 | 6,03E+06 | 71,017 | 0,000154 | 4,3931 | 8 |
| 2,63E+06 | 1,37E+07 | 1,02E+07 | 1,04E+07 | 47,661 | 0 | 9,5067 | 10 |
| 3,29E+08 | 2,96E+08 | 3,33E+08 | 3,10E+08 | 59,291 | 0 | 323,31 | 92 |
| 5,70E+07 | 5,89E+07 | 6,31E+07 | 6,05E+07 | 21,137 | 0 | 15,465 | 28 |
| 5,85E+08 | 5,60E+08 | 5,34E+08 | 5,94E+08 | 27,341 | 0 | 323,31 | 111 |
| 1,36E+09 | 1,94E+09 | 1,67E+09 | 1,64E+09 | 21,881 | 0 | 323,31 | 249 |
| 1,15E+07 | 1,33E+07 | 1,07E+07 | 8,91E+06 | 59,656 | 0,000155 | 4,4302 | 15 |
| 1,41E+06 | 7,02E+06 | 7,72E+06 | 5,57E+06 | 123,07 | 0,002783 | 2,0633 | 11 |
| 8,65E+08 | 9,06E+08 | 8,98E+08 | 8,15E+08 | 43,512 | 0 | 323,31 | 129 |
| 6,35E+06 | 6,28E+06 | 6,42E+06 | 6,20E+06 | 68,425 | 0 | 7,4795 | 10 |
| 1,80E+09 | 1,84E+09 | 1,88E+09 | 1,98E+09 | 38,092 | 0 | 323,31 | 362 |
| 6,39E+09 | 6,95E+09 | 6,57E+09 | 7,05E+09 | 56,585 | 0 | 323,31 | 416 |
| 1,64E+08 | 2,13E+08 | 1,98E+08 | 1,98E+08 | 71,593 | 0 | 191,82 | 118 |
| 1,97E+08 | 2,10E+08 | 2,23E+08 | 2,26E+08 | 90,195 | 0 | 73,965 | 117 |
| 1,61E+06 | 3,55E+06 | 2,83E+06 | 2,10E+06 | 45,952 | 0,003697 | 1,8223 | 4 |
| 1,27E+06 | 3,28E+06 | 2,30E+06 | 1,50E+06 | 35,767 | 0,005023 | 1,6735 | 8 |
| 1,78E+07 | 1,78E+07 | 1,73E+07 | 2,03E+07 | 54,158 | 0 | 10,796 | 22 |
| 7,99E+07 | 6,51E+07 | 6,36E+07 | 5,86E+07 | 57,685 | 0 | 21,707 | 55 |
| 8,36E+06 | 1,06E+07 | 8,69E+06 | 1,41E+07 | 241,76 | 0 | 6,3918 | 11 |
| 5,86E+07 | 2,85E+07 | 3,62E+07 | 4,72E+07 | 32,446 | 0 | 29,537 | 30 |
| 7,34E+08 | 5,90E+08 | 5,36E+08 | 5,83E+08 | 55,843 | 0 | 323,31 | 208 |
| 1,77E+07 | 1,67E+07 | 2,36E+07 | 1,95E+07 | 128,74 | 0,000152 | 4,0535 | 13 |
| 2,11E+08 | 2,19E+08 | 2,17E+08 | 2,11E+08 | 47,211 | 0 | 79,324 | 105 |
| 6,50E+06 | 4,35E+06 | 6,55E+06 | 4,26E+06 | 116,81 | 0,001685 | 2,2703 | 9 |
| 4,35E+09 | 3,88E+09 | 4,25E+09 | 4,46E+09 | 24,916 | 0 | 323,31 | 210 |
| 2,23E+07 | 2,84E+07 | 3,28E+07 | 2,28E+07 | 53,517 | 0 | 18,744 | 42 |
| 3,80E+07 | 3,36E+07 | 3,08E+07 | 3,07E+07 | 114,07 | 0 | 12,214 | 45 |

| | | | | | | | |
|---|---|---|---|---|---|---|---|
| 2,03E+06 | 3,59E+06 | 1,40E+07 | 4,00E+06 | 300,14 | 0,000853 | 2,5507 | 9 |
| 4,66E+06 | 9,85E+06 | 9,39E+06 | 9,19E+06 | 25,524 | 0,000148 | 3,3951 | 8 |
| 3,93E+07 | 3,72E+07 | 4,72E+07 | 4,00E+07 | 12,354 | 0 | 13,898 | 20 |
| 2,38E+07 | 2,98E+07 | 2,60E+07 | 2,74E+07 | 120,33 | 0 | 13,566 | 45 |
| 2,38E+06 | 1,53E+07 | 1,59E+07 | 2,07E+07 | 83,347 | 0,000155 | 4,4983 | 18 |
| 2,58E+06 | 5,46E+06 | 6,61E+06 | 5,06E+06 | 57,568 | 0 | 5,9831 | 12 |
| 1,02E+07 | 7,63E+06 | 7,84E+06 | 9,11E+06 | 107,95 | 0 | 10,746 | 13 |
| 3,60E+08 | 2,65E+08 | 3,13E+08 | 2,58E+08 | 52,792 | 0 | 41,687 | 51 |
| 5,12E+07 | 3,51E+07 | 1,13E+07 | 1,86E+06 | 52,966 | 0 | 32,304 | 13 |
| 6,06E+07 | 6,32E+07 | 7,58E+07 | 7,73E+07 | 101,49 | 0 | 25,355 | 67 |
| 8,62E+07 | 8,33E+07 | 9,62E+07 | 8,09E+07 | 10,275 | 0 | 80,699 | 40 |
| 2,68E+07 | 5,38E+07 | 6,65E+07 | 5,87E+07 | 15,515 | 0 | 9,4725 | 28 |
| 6,75E+06 | 7,76E+06 | 6,29E+06 | 7,13E+06 | 273,93 | 0,000158 | 5,162 | 12 |
| 1,89E+07 | 2,92E+07 | 2,38E+07 | 2,41E+07 | 86,258 | 0 | 9,813 | 24 |
| 1,38E+08 | 1,21E+08 | 1,11E+08 | 1,41E+08 | 83,097 | 0 | 141,84 | 123 |
| 4,54E+06 | 1,01E+07 | 5,68E+06 | 8,96E+06 | 81,794 | 0 | 7,7944 | 6 |
| 1,05E+08 | 1,20E+08 | 1,19E+08 | 1,15E+08 | 89,519 | 0 | 59,042 | 72 |
| 1,16E+09 | 1,15E+09 | 1,14E+09 | 1,27E+09 | 56,784 | 0 | 323,31 | 194 |
| 2,11E+07 | 2,35E+07 | 2,22E+07 | 2,77E+07 | 76,088 | 0 | 8,3759 | 25 |
| 4,35E+07 | 5,75E+07 | 5,97E+07 | 5,67E+07 | 108,98 | 0 | 46,012 | 57 |
| 2,44E+07 | 3,83E+07 | 3,83E+07 | 4,55E+07 | 216,13 | 0 | 20,266 | 41 |
| 7,65E+07 | 7,13E+07 | 8,02E+07 | 9,42E+07 | 13,194 | 0 | 79,003 | 36 |
| 5,03E+07 | 9,27E+07 | 8,06E+07 | 8,77E+07 | 16,564 | 0 | 88,15 | 19 |
| 2,16E+07 | 1,75E+07 | 1,46E+07 | 2,07E+07 | 42,779 | 0 | 31,838 | 19 |
| 1,05E+07 | 7,15E+06 | 9,71E+06 | 7,75E+06 | 37,525 | 0,000291 | 3,0433 | 10 |
| 6,84E+06 | 1,90E+06 | 2,30E+06 | 8,18E+06 | 69,558 | 0,000149 | 3,4984 | 6 |
| 1,01E+08 | 1,07E+08 | 9,43E+07 | 1,05E+08 | 43,99 | 0 | 168,15 | 36 |
| 4,56E+07 | 3,15E+07 | 3,47E+07 | 3,34E+07 | 135,04 | 0 | 44,826 | 36 |
| 1,89E+07 | 1,26E+07 | 1,04E+07 | 1,42E+07 | 64,227 | 0 | 33,826 | 31 |
| 1,16E+07 | 7,69E+06 | 8,60E+06 | 5,25E+06 | 119,4 | 0 | 9,8575 | 15 |
| 1,11E+09 | 1,17E+09 | 1,30E+09 | 1,41E+09 | 45,536 | 0 | 272,67 | 382 |
| 5,44E+07 | 7,03E+07 | 6,53E+07 | 5,46E+07 | 101,13 | 0 | 46,253 | 62 |
| 4,52E+07 | 2,93E+07 | 2,23E+06 | 2,37E+07 | 27,847 | 0 | 107,7 | 22 |
| 1,50E+07 | 1,76E+07 | 2,02E+07 | 1,92E+07 | 84,48 | 0 | 12,267 | 31 |
| 6,46E+06 | 5,27E+06 | 4,62E+06 | 7,03E+06 | 69,734 | 0,000153 | 4,1323 | 12 |
| 4,98E+06 | 4,82E+06 | 4,03E+06 | 4,56E+06 | 13,089 | 0,003448 | 1,936 | 4 |
| 1,35E+08 | 1,87E+08 | 2,11E+08 | 1,76E+08 | 51,029 | 0 | 107,47 | 82 |
| 1,65E+08 | 1,65E+08 | 1,69E+08 | 1,71E+08 | 62,832 | 0 | 199,58 | 117 |
| 1,36E+07 | 1,25E+07 | 8,86E+06 | 1,27E+07 | 84,253 | 0 | 8,1753 | 21 |
| 3,08E+07 | 1,85E+07 | 2,54E+07 | 2,72E+07 | 7,3184 | 0 | 50,678 | 19 |
| 6,51E+06 | 9,95E+06 | 1,47E+07 | 1,42E+07 | 11,411 | 0 | 9,8118 | 13 |
| 2,61E+07 | 2,32E+07 | 2,72E+07 | 2,51E+07 | 31,172 | 0 | 19,839 | 31 |
| 3,15E+07 | 3,15E+07 | 2,43E+07 | 2,72E+07 | 74,922 | 0 | 14,403 | 30 |
| 3,69E+08 | 3,39E+08 | 3,17E+08 | 3,85E+08 | 18,37 | 0 | 323,31 | 111 |
| 6,31E+07 | 6,39E+07 | 7,46E+07 | 5,38E+07 | 23,445 | 0 | 17,938 | 19 |
| 4,71E+07 | 3,44E+07 | 6,42E+07 | 4,08E+07 | 51,392 | 0 | 8,4717 | 25 |
| 2,77E+07 | 2,52E+07 | 2,84E+07 | 2,97E+07 | 34,244 | 0,000154 | 4,3587 | 18 |
| 1,09E+07 | 6,47E+06 | 5,32E+06 | 7,85E+06 | 52,292 | 0,003178 | 1,9573 | 7 |
| 3,06E+09 | 2,58E+09 | 2,76E+09 | 2,45E+09 | 45,729 | 0 | 323,31 | 639 |
| 2,47E+08 | 2,79E+08 | 2,34E+08 | 2,45E+08 | 35,247 | 0 | 61,619 | 60 |

| | | | | | | | |
|---|---|---|---|---|---|---|---|
| 1,41E+07 | 1,40E+07 | 1,28E+07 | 1,62E+07 | 150,49 | 0 | 5,6842 | 18 |
| 3,12E+06 | 1,42E+06 | 2,72E+06 | 2,91E+06 | 30,666 | 0,004369 | 1,7582 | 4 |
| 3,17E+07 | 3,21E+07 | 3,09E+07 | 4,40E+07 | 34,139 | 0 | 49,597 | 25 |
| 1,29E+07 | 8,71E+06 | 1,05E+07 | 1,23E+07 | 15,236 | 0 | 11,235 | 18 |
| 9,53E+08 | 8,71E+08 | 9,33E+08 | 8,89E+08 | 71,683 | 0 | 323,31 | 185 |
| 6,82E+07 | 7,30E+07 | 5,57E+07 | 5,75E+07 | 54,874 | 0 | 67,741 | 50 |
| 2,59E+08 | 2,43E+08 | 2,61E+08 | 2,72E+08 | 91,784 | 0 | 229,48 | 141 |
| 1,23E+08 | 1,16E+08 | 1,17E+08 | 1,27E+08 | 60,182 | 0 | 99,003 | 41 |
| 4,88E+09 | 7,32E+09 | 6,74E+09 | 7,68E+09 | 10,963 | 0 | 57,838 | 396 |
| 1,65E+07 | 1,91E+07 | 2,20E+07 | 2,15E+07 | 73,798 | 0 | 17,185 | 32 |
| 1,58E+09 | 1,00E+09 | 1,08E+09 | 1,26E+09 | 25,476 | 0 | 312,06 | 389 |
| 5,49E+06 | 5,39E+06 | 1,69E+06 | 1,10E+06 | 192,29 | 0,000156 | 4,7931 | 5 |
| 4,35E+08 | 3,59E+08 | 3,69E+08 | 3,36E+08 | 29,949 | 0 | 159,26 | 83 |
| 1,63E+08 | 1,63E+08 | 1,51E+08 | 1,59E+08 | 44,399 | 0 | 17,629 | 56 |
| 6,06E+08 | 6,36E+08 | 7,59E+08 | 7,19E+08 | 11,605 | 0 | 287,31 | 134 |
| 2,28E+07 | 6,64E+06 | 1,81E+07 | 8,65E+06 | 172,12 | 0,006074 | 1,5884 | 8 |
| 1,16E+07 | 2,05E+07 | 1,88E+07 | 1,82E+07 | 28,575 | 0 | 8,9882 | 24 |
| 7,39E+06 | 9,64E+06 | 9,00E+06 | 8,02E+06 | 40,753 | 0,004098 | 1,7735 | 5 |
| 2,62E+09 | 3,07E+09 | 3,02E+09 | 2,58E+09 | 31,293 | 0 | 323,31 | 189 |
| 4,90E+07 | 8,30E+07 | 6,79E+07 | 7,57E+07 | 31,218 | 0 | 40,643 | 48 |
| 4,30E+09 | 4,80E+09 | 4,74E+09 | 4,52E+09 | 55,758 | 0 | 323,31 | 445 |
| 3,09E+07 | 2,76E+07 | 1,68E+07 | 1,40E+07 | 90,372 | 0 | 11,603 | 31 |
| 3,17E+08 | 3,14E+08 | 3,78E+08 | 3,59E+08 | 56,979 | 0 | 233,49 | 95 |
| 3,05E+08 | 2,90E+08 | 2,82E+08 | 2,81E+08 | 228,84 | 0 | 211,34 | 225 |
| 1,57E+07 | 2,60E+06 | 2,18E+06 | 3,09E+06 | 41,7 | 0,000429 | 2,6748 | 4 |
| 2,02E+07 | 2,22E+07 | 2,27E+07 | 1,14E+07 | 74,993 | 0,000157 | 4,8944 | 11 |
| 2,92E+08 | 3,74E+08 | 4,09E+08 | 4,25E+08 | 48,381 | 0 | 323,31 | 122 |
| 2,97E+06 | 3,81E+06 | 8,73E+06 | 1,10E+07 | 139,1 | 0,000431 | 2,78 | 6 |
| 2,31E+07 | 2,35E+07 | 2,43E+07 | 2,44E+07 | 90,868 | 0 | 32,995 | 28 |
| 4,01E+07 | 5,09E+07 | 4,55E+07 | 4,99E+07 | 72,503 | 0 | 13,028 | 26 |
| 7,39E+07 | 8,54E+07 | 8,33E+07 | 7,74E+07 | 44,716 | 0 | 45,547 | 84 |
| 1,05E+07 | 9,79E+06 | 1,64E+07 | 2,41E+07 | 54,066 | 0 | 6,4927 | 13 |
| 9,08E+07 | 2,57E+08 | 2,00E+08 | 2,29E+08 | 15,054 | 0 | 43,154 | 96 |
| 1,32E+07 | 1,53E+07 | 1,33E+07 | 1,12E+07 | 23,849 | 0 | 6,9884 | 14 |
| 2,56E+07 | 2,07E+07 | 2,29E+07 | 2,91E+07 | 18,245 | 0 | 45,983 | 31 |
| 3,27E+09 | 3,25E+09 | 3,22E+09 | 3,41E+09 | 170,08 | 0 | 323,31 | 802 |
| 1,45E+07 | 9,77E+06 | 1,63E+07 | 1,01E+07 | 26,02 | 0,000147 | 3,258 | 8 |
| 8,68E+07 | 7,53E+07 | 8,14E+07 | 8,72E+07 | 71,421 | 0 | 58,043 | 79 |
| 7,65E+07 | 8,54E+07 | 9,07E+07 | 8,81E+07 | 39,859 | 0 | 29,248 | 64 |
| 8,88E+07 | 1,85E+08 | 5,06E+08 | 9,57E+07 | 11,43 | 0 | 5,3838 | 47 |
| 1,82E+07 | 1,80E+07 | 2,45E+07 | 2,26E+07 | 18,795 | 0 | 7,1675 | 17 |
| 4,52E+07 | 4,00E+07 | 2,25E+07 | 3,99E+07 | 55,207 | 0 | 6,301 | 12 |
| 1,38E+08 | 5,63E+07 | 7,33E+07 | 6,43E+07 | 6,6767 | 0 | 6,1957 | 86 |
| 3,89E+08 | 3,52E+08 | 3,39E+08 | 3,08E+08 | 91,349 | 0 | 107,96 | 162 |
| 4,28E+08 | 5,07E+08 | 4,49E+08 | 4,17E+08 | 105,1 | 0 | 323,31 | 145 |
| 2,04E+08 | 1,41E+08 | 1,69E+08 | 1,76E+08 | 52,107 | 0 | 116,63 | 104 |
| 5,09E+07 | 5,65E+07 | 5,58E+07 | 5,65E+07 | 10,678 | 0 | 6,9551 | 7 |
| 2,42E+06 | 2,46E+06 | 2,97E+06 | 1,02E+06 | 34,118 | 0,0037 | 1,8374 | 3 |
| 5,20E+07 | 4,58E+07 | 3,59E+07 | 2,41E+07 | 37,998 | 0 | 16,321 | 29 |
| 5,34E+08 | 5,86E+08 | 7,22E+08 | 7,01E+08 | 25,258 | 0 | 217,02 | 99 |

| | | | | | | | |
|---|---|---|---|---|---|---|---|
| 1,81E+08 | 2,31E+08 | 1,76E+08 | 1,71E+08 | 15,808 | 0 | 63,956 | 71 |
| 2,85E+07 | 3,22E+07 | 3,03E+07 | 2,83E+07 | 58,477 | 0 | 23,608 | 35 |
| 4,91E+08 | 4,99E+08 | 5,60E+08 | 5,78E+08 | 56,55 | 0 | 256,13 | 153 |
| 6,80E+06 | 1,83E+06 | 5,23E+06 | 5,30E+06 | 58,439 | 0,002782 | 2,0609 | 4 |
| 1,00E+08 | 1,20E+08 | 1,16E+08 | 1,04E+08 | 67,091 | 0 | 142,07 | 50 |
| 2,96E+06 | 3,47E+06 | 2,33E+06 | 2,25E+06 | 100,93 | 0,000435 | 2,9522 | 5 |
| 4,64E+06 | 4,65E+06 | 4,51E+06 | 5,15E+06 | 20,724 | 0,001823 | 2,2386 | 10 |
| 4,29E+08 | 4,11E+08 | 3,88E+08 | 3,49E+08 | 30,891 | 0 | 111,22 | 70 |
| 3,99E+07 | 5,61E+07 | 4,95E+07 | 5,12E+07 | 47,384 | 0 | 42,449 | 44 |
| 9,70E+07 | 9,64E+07 | 9,22E+07 | 1,03E+08 | 37,969 | 0 | 40,603 | 69 |
| 7,27E+07 | 4,81E+07 | 3,79E+07 | 3,81E+07 | 12,694 | 0,001271 | 2,4205 | 7 |
| 7,88E+07 | 7,80E+07 | 7,45E+07 | 7,42E+07 | 84,669 | 0 | 33,361 | 70 |
| 2,29E+07 | 2,71E+07 | 1,60E+07 | 1,06E+07 | 55,541 | 0 | 10,557 | 25 |
| 1,61E+09 | 1,00E+09 | 1,23E+09 | 1,26E+09 | 49,199 | 0 | 323,31 | 373 |
| 4,97E+07 | 5,75E+07 | 6,25E+07 | 4,83E+07 | 17,348 | 0 | 11,245 | 21 |
| 1,64E+09 | 1,50E+09 | 1,68E+09 | 1,67E+09 | 60,342 | 0 | 323,31 | 381 |
| 1,75E+08 | 1,64E+08 | 1,88E+08 | 1,69E+08 | 93,282 | 0 | 137,85 | 85 |
| 3,15E+07 | 2,95E+07 | 2,13E+07 | 1,50E+07 | 13,915 | 0 | 8,1971 | 17 |
| 6,93E+08 | 6,28E+08 | 7,05E+08 | 6,66E+08 | 67,589 | 0 | 323,31 | 192 |
| 2,85E+07 | 3,58E+07 | 3,67E+07 | 3,40E+07 | 18,342 | 0 | 33,925 | 20 |
| 1,35E+07 | 1,45E+07 | 8,06E+06 | 1,09E+07 | 22,123 | 0,000146 | 3,0958 | 7 |
| 1,25E+09 | 9,85E+08 | 8,51E+08 | 8,34E+08 | 26,468 | 0 | 200,66 | 161 |
| 2,34E+07 | 2,51E+07 | 2,34E+07 | 2,28E+07 | 57,541 | 0 | 19,77 | 24 |
| 7,58E+07 | 5,55E+07 | 6,05E+07 | 6,32E+07 | 76,068 | 0 | 58,805 | 47 |
| 8,91E+06 | 6,43E+06 | 5,83E+06 | 8,73E+06 | 76,441 | 0 | 5,3384 | 7 |
| 1,16E+07 | 1,66E+07 | 1,12E+07 | 9,91E+06 | 71,021 | 0,000153 | 4,1907 | 9 |
| 2,81E+08 | 3,76E+08 | 4,63E+08 | 4,91E+08 | 20,626 | 0 | 17,913 | 85 |
| 7,97E+07 | 7,05E+07 | 5,71E+07 | 8,40E+07 | 73,976 | 0 | 21,482 | 27 |
| 7,11E+06 | 7,24E+06 | 7,76E+06 | 5,41E+06 | 41,982 | 0,000852 | 2,5229 | 8 |
| 6,56E+06 | 8,53E+06 | 1,12E+07 | 1,10E+07 | 40,421 | 0 | 8,5404 | 7 |
| 8,08E+06 | 5,42E+06 | 8,58E+06 | 4,93E+06 | 69,858 | 0,000155 | 4,4702 | 11 |
| 1,06E+07 | 1,08E+07 | 1,39E+07 | 1,48E+07 | 29,266 | 0 | 6,1481 | 13 |
| 2,49E+06 | 7,19E+06 | 1,19E+07 | 2,53E+06 | 45,544 | 0,001961 | 2,2033 | 2 |
| 1,31E+08 | 1,25E+08 | 1,31E+08 | 1,32E+08 | 104,77 | 0 | 21,139 | 70 |
| 6,14E+06 | 2,76E+06 | 2,48E+06 | 3,03E+06 | 18,281 | 0,003177 | 1,9554 | 3 |
| 5,03E+08 | 4,49E+08 | 4,31E+08 | 4,06E+08 | 80,027 | 0 | 323,31 | 159 |
| 8,79E+07 | 6,75E+07 | 5,83E+07 | 5,16E+07 | 14,555 | 0 | 29,047 | 45 |
| 2,56E+07 | 6,59E+05 | 3,59E+06 | 6,97E+06 | 88,712 | 0,000433 | 2,8672 | 5 |
| 2,28E+08 | 2,11E+08 | 2,00E+08 | 2,01E+08 | 119,15 | 0 | 140,12 | 139 |
| 3,47E+07 | 3,50E+07 | 3,48E+07 | 3,55E+07 | 50,66 | 0 | 14,371 | 36 |
| 1,78E+08 | 1,44E+08 | 5,57E+08 | 2,95E+08 | 28,911 | 0 | 12,056 | 27 |
| 3,88E+06 | 7,34E+06 | 8,02E+06 | 7,00E+06 | 122,68 | 0 | 6,0331 | 12 |
| 8,20E+08 | 8,31E+08 | 8,35E+08 | 7,77E+08 | 100,94 | 0 | 323,31 | 351 |
| 2,08E+08 | 2,63E+08 | 2,31E+08 | 2,01E+08 | 17,8 | 0 | 161,28 | 61 |
| 6,40E+07 | 5,01E+07 | 4,72E+07 | 5,39E+07 | 107,72 | 0 | 18,113 | 57 |
| 3,90E+07 | 3,95E+07 | 4,06E+07 | 3,45E+07 | 12,399 | 0 | 26,789 | 20 |
| 7,28E+07 | 7,46E+07 | 7,77E+07 | 7,52E+07 | 48,572 | 0 | 78,032 | 36 |
| 4,97E+06 | 6,95E+06 | 8,98E+06 | 8,16E+06 | 46,846 | 0,00043 | 2,7313 | 12 |
| 1,12E+09 | 1,16E+09 | 1,07E+09 | 1,06E+09 | 119,82 | 0 | 323,31 | 300 |
| 1,86E+08 | 1,70E+08 | 1,80E+08 | 1,99E+08 | 44,054 | 0 | 53,475 | 94 |

| | | | | | | | |
|---|---|---|---|---|---|---|---|
| 5,70E+07 | 1,64E+08 | 1,46E+08 | 1,56E+08 | 23,689 | 0 | 7,455 | 18 |
| 1,80E+10 | 1,50E+10 | 1,61E+10 | 1,46E+10 | 70,87 | 0 | 323,31 | 1362 |
| 2,16E+07 | 3,46E+07 | 3,44E+07 | 3,77E+07 | 90,001 | 0 | 35,752 | 34 |
| 1,01E+07 | 8,67E+06 | 7,30E+06 | 7,83E+06 | 44,792 | 0,000153 | 4,1111 | 6 |
| 1,94E+06 | 2,54E+06 | 2,76E+06 | 2,88E+06 | 31,669 | 0,000147 | 3,1596 | 10 |
| 5,38E+06 | 4,20E+06 | 4,88E+06 | 4,82E+06 | 18,666 | 0,000155 | 4,5204 | 2 |
| 9,11E+07 | 9,25E+07 | 9,00E+07 | 1,05E+08 | 40,506 | 0 | 100,46 | 69 |
| 3,77E+08 | 3,40E+08 | 3,27E+08 | 3,38E+08 | 68,512 | 0 | 199,07 | 134 |
| 3,52E+07 | 2,05E+07 | 2,76E+07 | 1,02E+07 | 49,612 | 0 | 24,649 | 28 |
| 8,86E+05 | 5,88E+06 | 5,73E+06 | 6,85E+06 | 52,058 | 0,000156 | 4,6875 | 12 |
| 7,14E+06 | 9,18E+06 | 7,88E+06 | 1,28E+07 | 115,84 | 0 | 7,379 | 14 |
| 1,07E+08 | 1,00E+08 | 1,04E+08 | 9,79E+07 | 24,683 | 0 | 241,88 | 37 |
| 2,90E+08 | 3,96E+08 | 3,59E+08 | 3,48E+08 | 33,097 | 0 | 191,66 | 107 |
| 1,05E+08 | 9,63E+07 | 9,80E+07 | 9,58E+07 | 83,745 | 0 | 186,57 | 72 |
| 1,21E+07 | 5,61E+06 | 1,45E+07 | 6,17E+06 | 40,129 | 0,002916 | 2,031 | 4 |
| 1,09E+08 | 1,08E+08 | 1,02E+08 | 9,32E+07 | 57,236 | 0 | 95,908 | 77 |
| 2,94E+07 | 4,32E+07 | 3,81E+07 | 4,75E+07 | 70,839 | 0 | 17,54 | 35 |
| 4,00E+07 | 3,16E+07 | 3,27E+07 | 2,59E+07 | 39,091 | 0 | 59,469 | 52 |
| 5,32E+09 | 6,21E+09 | 6,42E+09 | 6,58E+09 | 11,651 | 0 | 323,31 | 170 |
| 1,33E+08 | 1,54E+07 | 1,37E+07 | 1,66E+07 | 76,86 | 0,004632 | 1,7334 | 4 |
| 7,73E+06 | 6,40E+06 | 4,61E+06 | 5,13E+06 | 56,909 | 0 | 8,6873 | 4 |
| 1,31E+08 | 1,28E+08 | 1,36E+08 | 1,31E+08 | 103,45 | 0 | 101,67 | 105 |
| 2,62E+07 | 2,72E+07 | 2,79E+07 | 3,44E+07 | 31,68 | 0 | 8,0773 | 22 |
| 1,82E+07 | 3,95E+07 | 3,48E+07 | 3,19E+07 | 22,975 | 0 | 10,773 | 25 |
| 3,49E+08 | 3,05E+08 | 3,36E+08 | 2,67E+08 | 36,154 | 0 | 132,36 | 57 |
| 2,93E+07 | 3,32E+07 | 2,76E+07 | 3,00E+07 | 23,201 | 0 | 8,0958 | 26 |
| 4,08E+06 | 5,31E+06 | 7,10E+06 | 7,63E+06 | 97,302 | 0,000151 | 3,7981 | 7 |
| 2,78E+07 | 2,25E+07 | 2,08E+07 | 1,73E+07 | 104,55 | 0 | 94,96 | 39 |
| 3,45E+06 | 2,85E+05 | 1,25E+05 | 2,98E+06 | 68,692 | 0 | 249,12 | 34 |
| 8,65E+06 | 7,97E+06 | 1,08E+07 | 7,82E+06 | 65,74 | 0 | 6,3818 | 16 |
| 1,26E+09 | 1,04E+09 | 1,15E+09 | 1,17E+09 | 91,712 | 0 | 323,31 | 164 |
| 1,68E+07 | 1,42E+07 | 1,29E+07 | 1,14E+07 | 202,07 | 0 | 19,297 | 20 |
| 4,45E+07 | 3,44E+07 | 2,85E+07 | 3,33E+07 | 29,213 | 0 | 10,044 | 25 |
| 3,93E+08 | 3,96E+08 | 3,82E+08 | 3,90E+08 | 33,524 | 0 | 141,29 | 101 |
| 1,29E+07 | 8,68E+06 | 1,40E+07 | 1,23E+07 | 18,754 | 0,00015 | 3,5914 | 11 |
| 4,00E+06 | 2,01E+06 | 3,48E+06 | 2,05E+06 | 86,307 | 0,000429 | 2,6864 | 11 |
| 4,74E+07 | 5,29E+07 | 4,95E+07 | 5,62E+07 | 22,893 | 0 | 14,315 | 30 |
| 3,44E+07 | 1,41E+07 | 2,89E+07 | 3,73E+07 | 97,967 | 0 | 33,973 | 37 |
| 2,21E+07 | 3,00E+07 | 3,20E+07 | 3,20E+07 | 62,412 | 0 | 17,006 | 26 |
| 3,45E+09 | 3,07E+09 | 3,57E+09 | 3,33E+09 | 21,897 | 0 | 323,31 | 202 |
| 2,54E+08 | 2,38E+08 | 1,90E+08 | 2,38E+08 | 15,076 | 0 | 14,547 | 63 |
| 1,75E+07 | 2,43E+07 | 3,04E+07 | 3,00E+07 | 83,981 | 0 | 11,945 | 33 |
| 4,50E+06 | 6,61E+06 | 1,13E+07 | 9,84E+06 | 17,985 | 0,000571 | 2,6413 | 5 |
| 1,39E+07 | 8,74E+06 | 1,33E+06 | 2,99E+06 | 65,329 | 0 | 8,4874 | 7 |
| 2,23E+07 | 2,11E+07 | 2,19E+07 | 2,12E+07 | 40,656 | 0 | 15,757 | 32 |
| 3,84E+07 | 1,69E+07 | 1,50E+07 | 1,30E+07 | 229,39 | 0 | 11,302 | 29 |
| 2,82E+07 | 2,31E+07 | 1,87E+07 | 2,03E+07 | 23,144 | 0 | 12,448 | 27 |
| 1,85E+07 | 1,16E+07 | 1,45E+07 | 1,46E+07 | 346,45 | 0,000155 | 4,4435 | 17 |
| 7,18E+06 | 4,48E+06 | 4,08E+06 | 3,22E+06 | 175,77 | 0,000431 | 2,7926 | 9 |
| 7,94E+06 | 7,37E+06 | 6,04E+06 | 8,00E+06 | 35,208 | 0 | 11,525 | 20 |

| | | | | | | | |
|---|---|---|---|---|---|---|---|
| 1,72E+07 | 2,03E+07 | 1,88E+07 | 1,68E+07 | 211,94 | 0 | 16,596 | 32 |
| 2,98E+08 | 2,87E+08 | 2,83E+08 | 3,01E+08 | 10,403 | 0 | 101,58 | 68 |
| 2,32E+08 | 3,12E+08 | 2,92E+08 | 2,86E+08 | 43,066 | 0 | 63,964 | 116 |
| 8,99E+07 | 8,09E+07 | 8,17E+07 | 8,54E+07 | 91,39 | 0 | 59,838 | 83 |
| 8,55E+06 | 9,68E+06 | 8,45E+06 | 7,77E+06 | 65,547 | 0,000146 | 3,1 | 9 |
| 7,77E+06 | 6,83E+06 | 3,87E+06 | 4,06E+06 | 25,548 | 0,000159 | 5,2664 | 6 |
| 1,27E+07 | 1,96E+07 | 2,15E+07 | 1,71E+07 | 37,417 | 0 | 7,3858 | 15 |
| 3,10E+09 | 3,05E+09 | 3,07E+09 | 3,02E+09 | 49,548 | 0 | 323,31 | 562 |
| 9,34E+07 | 6,67E+07 | 7,75E+07 | 6,97E+07 | 14,523 | 0 | 113,3 | 58 |
| 4,01E+06 | 2,64E+07 | 1,70E+07 | 1,96E+07 | 32,836 | 0,000158 | 5,0097 | 12 |
| 1,07E+08 | 1,08E+08 | 1,14E+08 | 1,07E+08 | 36,721 | 0 | 25,316 | 48 |
| 7,09E+07 | 9,10E+07 | 8,81E+07 | 8,83E+07 | 68,155 | 0 | 92,934 | 70 |
| 1,32E+08 | 4,39E+07 | 6,45E+07 | 3,81E+07 | 35,914 | 0 | 323,31 | 33 |
| 1,31E+07 | 1,60E+06 | 9,26E+06 | 1,25E+07 | 22,781 | 0 | 9,4032 | 6 |
| 1,95E+08 | 1,88E+08 | 2,05E+08 | 2,07E+08 | 53,056 | 0 | 161,37 | 102 |
| 3,40E+08 | 4,20E+08 | 4,17E+08 | 3,89E+08 | 26,249 | 0 | 197,32 | 110 |
| 5,14E+06 | 9,80E+06 | 2,76E+06 | 1,60E+06 | 28,88 | 0,002506 | 2,0713 | 2 |
| 3,20E+06 | 8,74E+05 | 1,34E+06 | 5,78E+06 | 50,185 | 0,008203 | 1,5096 | 5 |
| 8,38E+07 | 7,38E+07 | 7,89E+07 | 7,92E+07 | 40,658 | 0 | 99,107 | 57 |
| 4,63E+06 | 2,16E+06 | 4,28E+06 | 4,10E+06 | 67,579 | 0,00015 | 3,6108 | 7 |
| 1,53E+09 | 1,49E+09 | 1,47E+09 | 1,46E+09 | 60,909 | 0 | 323,31 | 216 |
| 6,35E+08 | 6,50E+08 | 6,20E+08 | 6,67E+08 | 28,357 | 0 | 323,31 | 99 |
| 4,73E+09 | 3,05E+09 | 2,62E+09 | 3,04E+09 | 22,099 | 0 | 34,597 | 59 |
| 1,42E+08 | 9,69E+07 | 3,16E+06 | 1,67E+08 | 23,263 | 0 | 63,811 | 16 |
| 1,34E+09 | 1,67E+09 | 1,75E+09 | 1,64E+09 | 13,015 | 0 | 162,89 | 92 |
| 1,94E+07 | 5,17E+07 | 6,10E+07 | 7,00E+07 | 26,832 | 0 | 13,235 | 32 |
| 2,93E+08 | 2,91E+08 | 3,02E+08 | 3,21E+08 | 14,141 | 0 | 323,31 | 48 |
| 5,05E+07 | 4,37E+07 | 4,44E+07 | 4,00E+07 | 37,965 | 0 | 48,307 | 33 |
| 7,11E+06 | 2,60E+06 | 1,53E+06 | 6,05E+06 | 64,322 | 0 | 5,7889 | 6 |
| 1,26E+07 | 2,69E+07 | 3,55E+07 | 2,84E+07 | 27,873 | 0 | 5,3267 | 8 |
| 7,57E+06 | 1,36E+07 | 8,17E+06 | 6,40E+06 | 119,96 | 0 | 9,6305 | 11 |
| 1,00E+07 | 9,57E+06 | 1,13E+07 | 8,38E+06 | 36,067 | 0 | 37,051 | 16 |
| 1,70E+09 | 1,72E+09 | 1,76E+09 | 1,77E+09 | 9,1413 | 0 | 323,31 | 224 |
| 7,00E+07 | 6,53E+07 | 7,35E+07 | 7,46E+07 | 158,83 | 0 | 54,135 | 70 |
| 4,12E+06 | 2,51E+06 | 2,14E+06 | 9,63E+06 | 15,085 | 0,001823 | 2,2386 | 5 |
| 5,61E+07 | 7,11E+07 | 6,34E+07 | 6,09E+07 | 109,32 | 0 | 34,818 | 78 |
| 5,16E+08 | 3,89E+08 | 4,02E+08 | 4,41E+08 | 6,3814 | 0 | 191,45 | 14 |
| 1,91E+07 | 1,86E+07 | 1,88E+07 | 2,39E+07 | 25,083 | 0 | 54,253 | 14 |
| 6,72E+07 | 8,25E+07 | 9,47E+07 | 5,37E+07 | 22,16 | 0 | 71,591 | 20 |
| 1,11E+07 | 1,02E+07 | 1,14E+07 | 1,14E+07 | 68,886 | 0 | 7,5018 | 13 |
| 2,22E+06 | 4,99E+06 | 3,87E+06 | 6,33E+06 | 46,717 | 0 | 12,809 | 3 |
| 1,36E+06 | 3,67E+06 | 4,32E+06 | 4,32E+06 | 29,003 | 0,000159 | 5,3034 | 6 |
| 4,27E+07 | 4,92E+07 | 5,62E+07 | 5,11E+07 | 102,61 | 0 | 57,094 | 47 |
| 4,90E+07 | 1,53E+07 | 2,96E+06 | 4,46E+06 | 31,802 | 0 | 161,95 | 13 |
| 1,54E+09 | 1,17E+09 | 1,50E+09 | 1,31E+09 | 29,976 | 0 | 323,31 | 97 |
| 4,24E+08 | 4,41E+08 | 4,40E+08 | 4,18E+08 | 33,814 | 0 | 323,31 | 87 |
| 5,49E+06 | 4,09E+06 | 3,64E+06 | 4,00E+06 | 55,448 | 0,001409 | 2,3412 | 7 |
| 3,50E+07 | 3,49E+07 | 4,97E+07 | 4,41E+07 | 67,296 | 0,000147 | 3,2813 | 18 |
| 7,25E+07 | 9,53E+07 | 8,33E+07 | 9,79E+07 | 17,27 | 0 | 290,27 | 48 |
| 7,86E+06 | 1,09E+07 | 8,00E+06 | 6,99E+06 | 25,057 | 0 | 5,5002 | 16 |

| | | | | | | | |
|---|---|---|---|---|---|---|---|
| 5,61E+07 | 9,12E+06 | 4,59E+07 | 2,74E+07 | 13,784 | 0 | 142,42 | 17 |
| 2,21E+07 | 3,26E+07 | 2,72E+07 | 3,00E+07 | 13,131 | 0 | 116,81 | 18 |
| 2,24E+08 | 1,21E+08 | 2,45E+08 | 2,19E+08 | 7,7662 | 0 | 6,2046 | 22 |
| 3,28E+07 | 4,19E+07 | 4,30E+07 | 4,22E+07 | 23,629 | 0 | 20,58 | 39 |
| 1,25E+07 | 6,55E+06 | 9,34E+06 | 1,05E+07 | 85,277 | 0 | 12,492 | 20 |
| 6,05E+06 | 7,15E+06 | 1,97E+06 | 1,51E+06 | 33,949 | 0,004622 | 1,6965 | 4 |
| 7,34E+07 | 9,00E+07 | 7,05E+07 | 8,51E+07 | 18,963 | 0 | 12,558 | 53 |
| 1,37E+07 | 2,06E+07 | 3,22E+06 | 2,13E+06 | 59,807 | 0 | 11,122 | 14 |
| 9,21E+07 | 8,31E+07 | 7,57E+07 | 9,46E+07 | 22,03 | 0 | 178,22 | 67 |
| 5,84E+06 | 6,43E+06 | 6,68E+06 | 4,40E+06 | 48,757 | 0,000158 | 5,0679 | 15 |
| 3,54E+08 | 3,31E+08 | 3,25E+08 | 3,37E+08 | 68,995 | 0 | 170,9 | 167 |
| 3,11E+08 | 3,06E+08 | 2,85E+08 | 2,74E+08 | 38,549 | 0 | 67,694 | 115 |
| 2,57E+07 | 7,78E+06 | 1,65E+07 | 1,30E+07 | 13,918 | 0,000156 | 4,6904 | 8 |
| 9,60E+07 | 9,08E+07 | 8,52E+07 | 8,18E+07 | 17,086 | 0 | 63,136 | 52 |
| 1,44E+07 | 1,24E+07 | 1,09E+07 | 1,00E+07 | 64,324 | 0,000155 | 4,4716 | 14 |
| 4,52E+06 | 1,08E+07 | 9,34E+06 | 8,74E+06 | 23,662 | 0,000157 | 4,9465 | 15 |
| 2,99E+08 | 3,72E+08 | 3,49E+08 | 3,35E+08 | 14,254 | 0 | 88,476 | 46 |
| 4,72E+08 | 5,77E+08 | 5,99E+08 | 5,80E+08 | 51,863 | 0 | 169,19 | 168 |
| 5,44E+06 | 3,75E+06 | 2,13E+06 | 1,95E+06 | 50,319 | 0,000149 | 3,5677 | 7 |
| 6,33E+08 | 4,86E+08 | 1,85E+08 | 1,73E+08 | 32,065 | 0 | 47,245 | 49 |
| 3,83E+06 | 5,42E+06 | 2,10E+06 | 4,72E+06 | 24,631 | 0,000993 | 2,491 | 5 |
| 3,70E+07 | 4,56E+07 | 5,04E+07 | 4,10E+07 | 45,96 | 0 | 38,097 | 11 |
| 2,41E+06 | 2,59E+06 | 2,33E+06 | 2,38E+06 | 307,63 | 0,005024 | 1,6765 | 8 |
| 1,50E+06 | 6,98E+06 | 3,30E+06 | 3,48E+06 | 22,645 | 0 | 6,7891 | 13 |
| 5,33E+06 | 1,19E+07 | 5,75E+06 | 6,01E+06 | 60,015 | 0,000154 | 4,3963 | 10 |
| 3,43E+07 | 2,43E+07 | 3,41E+07 | 3,47E+07 | 58,965 | 0 | 11,553 | 26 |
| 9,43E+08 | 9,28E+08 | 8,62E+08 | 9,13E+08 | 74,792 | 0 | 323,31 | 239 |
| 2,26E+08 | 2,14E+08 | 2,17E+08 | 2,36E+08 | 33,104 | 0 | 162,58 | 68 |
| 6,21E+06 | 4,55E+06 | 4,78E+06 | 6,61E+06 | 52,278 | 0,003177 | 1,9565 | 8 |
| 3,38E+07 | 3,13E+07 | 3,45E+07 | 3,31E+07 | 54,968 | 0 | 42,758 | 34 |
| 3,25E+06 | 3,43E+06 | 3,88E+06 | 4,63E+06 | 18,363 | 0,000155 | 4,5623 | 3 |
| 2,21E+08 | 2,19E+08 | 2,27E+08 | 2,52E+08 | 69,552 | 0 | 30,929 | 83 |
| 7,07E+08 | 4,91E+08 | 5,06E+08 | 5,34E+08 | 29,367 | 0 | 122,46 | 113 |
| 1,98E+07 | 2,37E+07 | 2,11E+07 | 2,37E+07 | 90,025 | 0 | 12,192 | 17 |
| 1,42E+07 | 1,21E+07 | 1,63E+07 | 1,21E+07 | 124,87 | 0 | 5,8283 | 22 |
| 4,93E+07 | 3,83E+07 | 2,53E+07 | 3,27E+07 | 47,318 | 0 | 118,54 | 41 |
| 8,79E+07 | 7,19E+07 | 9,25E+07 | 7,03E+07 | 55,196 | 0 | 77,512 | 81 |
| 7,85E+07 | 8,35E+07 | 8,42E+07 | 8,82E+07 | 46,719 | 0 | 85,708 | 57 |
| 2,48E+06 | 4,76E+06 | 3,71E+06 | 3,55E+06 | 17,43 | 0,001271 | 2,422 | 11 |
| 4,38E+06 | 3,76E+06 | 5,18E+06 | 4,14E+06 | 26,986 | 0,00318 | 1,97 | 4 |
| 6,07E+08 | 5,09E+08 | 5,39E+08 | 5,53E+08 | 44,868 | 0 | 228,8 | 111 |
| 4,10E+08 | 4,04E+08 | 4,21E+08 | 4,37E+08 | 75,409 | 0 | 301,46 | 178 |
| 3,03E+07 | 4,19E+07 | 4,25E+07 | 3,43E+07 | 79,576 | 0 | 25,011 | 40 |
| 1,89E+07 | 2,12E+07 | 1,82E+07 | 1,94E+07 | 31,79 | 0,000158 | 5,0596 | 16 |
| 8,35E+07 | 1,08E+08 | 9,11E+07 | 9,63E+07 | 19,784 | 0 | 11,235 | 41 |
| 2,01E+07 | 2,87E+07 | 2,62E+07 | 2,67E+07 | 16,037 | 0,000159 | 5,1661 | 15 |
| 1,32E+07 | 1,24E+07 | 1,38E+07 | 1,49E+07 | 33,689 | 0,00015 | 3,5931 | 18 |
| 5,56E+07 | 2,62E+07 | 4,72E+07 | 3,21E+06 | 52,96 | 0 | 16,248 | 22 |
| 1,42E+07 | 1,05E+07 | 1,18E+06 | 1,17E+07 | 28,048 | 0,000431 | 2,7685 | 11 |
| 5,02E+06 | 1,95E+06 | 1,34E+06 | 2,12E+06 | 130,79 | 0 | 7,7051 | 8 |

| | | | | | | | |
|---|---|---|---|---|---|---|---|
| 1,88E+08 | 1,91E+08 | 1,80E+08 | 1,93E+08 | 33,656 | 0 | 120,67 | 60 |
| 1,64E+07 | 1,42E+07 | 1,51E+07 | 1,23E+07 | 20,809 | 0,001273 | 2,4428 | 16 |
| 8,07E+06 | 1,17E+07 | 1,42E+07 | 1,29E+07 | 114,58 | 0 | 7,4483 | 9 |
| 7,35E+07 | 6,45E+07 | 6,55E+07 | 7,63E+07 | 32,41 | 0 | 10,443 | 17 |
| 7,86E+07 | 7,36E+07 | 7,68E+07 | 5,21E+07 | 46,434 | 0 | 6,027 | 28 |
| 8,31E+07 | 1,30E+08 | 8,21E+07 | 1,02E+08 | 527,45 | 0 | 36,041 | 78 |
| 8,22E+07 | 6,75E+07 | 6,06E+07 | 5,83E+07 | 29,689 | 0 | 103,81 | 43 |
| 7,30E+06 | 1,83E+06 | 2,78E+06 | 3,91E+06 | 14,486 | 0 | 96,805 | 12 |
| 7,21E+07 | 8,39E+07 | 7,64E+07 | 6,66E+07 | 33,512 | 0 | 13,214 | 52 |
| 2,55E+08 | 2,40E+08 | 2,48E+08 | 2,17E+08 | 47,269 | 0 | 147,96 | 61 |
| 2,63E+09 | 2,32E+09 | 2,27E+09 | 2,33E+09 | 293,02 | 0 | 323,31 | 824 |
| 2,50E+09 | 4,61E+08 | 5,55E+08 | 3,99E+08 | 28,302 | 0 | 323,31 | 95 |
| 2,86E+07 | 3,29E+07 | 2,97E+07 | 3,22E+07 | 104,96 | 0 | 15,036 | 43 |
| 1,14E+07 | 8,20E+06 | 1,14E+07 | 2,40E+06 | 48,235 | 0,000151 | 3,8886 | 11 |
| 1,77E+07 | 3,48E+07 | 2,54E+07 | 2,87E+07 | 32,355 | 0 | 14,502 | 22 |
| 1,10E+08 | 2,18E+08 | 1,70E+08 | 1,54E+08 | 13,553 | 0 | 127,82 | 48 |
| 2,25E+07 | 2,70E+07 | 2,49E+07 | 2,12E+07 | 39,906 | 0 | 25,026 | 19 |
| 1,03E+07 | 9,82E+06 | 1,18E+07 | 8,07E+06 | 63,303 | 0 | 6,5851 | 17 |
| 2,04E+06 | 2,96E+06 | 1,10E+06 | 3,48E+06 | 164,74 | 0 | 5,6674 | 12 |
| 6,69E+07 | 5,86E+07 | 6,55E+07 | 7,04E+07 | 21,84 | 0 | 185,77 | 31 |
| 2,67E+07 | 1,37E+07 | 1,90E+07 | 4,72E+06 | 157,2 | 0 | 12,059 | 22 |
| 6,93E+07 | 7,34E+07 | 8,00E+07 | 9,22E+07 | 23,192 | 0 | 32,413 | 31 |
| 1,32E+07 | 1,29E+07 | 1,61E+07 | 1,47E+07 | 14,281 | 0,000157 | 4,899 | 15 |
| 4,02E+08 | 3,90E+08 | 4,29E+08 | 4,27E+08 | 75,656 | 0 | 323,31 | 143 |
| 2,54E+06 | 1,57E+07 | 1,15E+07 | 1,61E+07 | 35,333 | 0 | 14,414 | 17 |
| 6,05E+06 | 6,62E+06 | 3,17E+06 | 1,16E+06 | 102,65 | 0,000292 | 3,0702 | 3 |
| 6,12E+07 | 7,32E+07 | 6,78E+07 | 7,15E+07 | 78,894 | 0 | 32,922 | 51 |
| 5,15E+07 | 6,17E+07 | 6,34E+07 | 5,90E+07 | 138,76 | 0 | 43,04 | 77 |
| 9,24E+09 | 8,69E+09 | 9,12E+09 | 8,97E+09 | 532,04 | 0 | 323,31 | 2037 |
| 1,16E+07 | 7,36E+06 | 1,23E+07 | 1,56E+07 | 104,52 | 0 | 11,776 | 21 |
| 1,31E+07 | 1,32E+07 | 1,42E+07 | 1,12E+07 | 22,905 | 0,000158 | 5,1107 | 12 |
| 7,07E+06 | 1,47E+06 | 3,44E+06 | 8,64E+06 | 54,726 | 0,003443 | 1,9189 | 7 |
| 1,47E+07 | 1,50E+07 | 1,53E+07 | 1,55E+07 | 56,753 | 0,000431 | 2,802 | 14 |
| 2,59E+07 | 2,72E+07 | 1,95E+07 | 2,00E+07 | 53,956 | 0 | 12,69 | 29 |
| 6,45E+08 | 6,29E+08 | 6,56E+08 | 6,02E+08 | 116,95 | 0 | 323,31 | 261 |
| 4,49E+07 | 1,55E+07 | 1,57E+07 | 1,90E+07 | 25,342 | 0 | 6,0071 | 7 |
| 2,94E+08 | 2,80E+08 | 2,72E+08 | 2,56E+08 | 104,55 | 0 | 275,37 | 159 |
| 1,38E+07 | 1,23E+07 | 1,18E+07 | 6,04E+06 | 95,644 | 0 | 8,5002 | 19 |
| 3,35E+07 | 3,06E+07 | 4,32E+07 | 3,83E+07 | 49,126 | 0 | 9,7794 | 21 |
| 9,67E+05 | 6,42E+06 | 4,53E+06 | 4,84E+06 | 22,656 | 0 | 15,547 | 5 |
| 7,59E+07 | 3,27E+07 | 7,62E+07 | 6,33E+07 | 5,8378 | 0,008456 | 1,4743 | 8 |
| 3,68E+07 | 3,27E+07 | 3,25E+07 | 3,43E+07 | 74,85 | 0 | 20,942 | 42 |
| 1,60E+08 | 1,98E+08 | 1,84E+08 | 1,69E+08 | 37,547 | 0 | 323,31 | 41 |
| 7,30E+07 | 6,03E+07 | 6,70E+07 | 7,44E+07 | 56,41 | 0 | 52,218 | 60 |
| 3,49E+07 | 3,72E+07 | 3,31E+07 | 4,58E+07 | 19,52 | 0 | 11,888 | 33 |
| 1,38E+08 | 1,61E+08 | 1,76E+08 | 1,73E+08 | 49,374 | 0 | 17,709 | 43 |
| 3,09E+08 | 3,95E+08 | 3,00E+08 | 3,02E+08 | 51,373 | 0 | 33,265 | 183 |
| 2,02E+08 | 2,27E+08 | 2,01E+08 | 2,36E+08 | 22,689 | 0 | 120,49 | 65 |
| 1,84E+08 | 1,78E+08 | 1,68E+08 | 1,73E+08 | 41,745 | 0 | 30,01 | 82 |
| 3,61E+06 | 2,25E+06 | 4,49E+06 | 4,40E+06 | 48,791 | 0,000431 | 2,7851 | 6 |

| | | | | | | | |
|---|---|---|---|---|---|---|---|
| 1,32E+07 | 1,93E+07 | 2,21E+07 | 2,30E+07 | 39,548 | 0,000852 | 2,5304 | 7 |
| 1,02E+07 | 1,18E+07 | 1,04E+07 | 8,19E+06 | 39,485 | 0,000152 | 4,0001 | 9 |
| 2,98E+06 | 3,61E+06 | 3,13E+06 | 2,70E+06 | 39,469 | 0,001961 | 2,2019 | 8 |
| 3,73E+08 | 3,66E+08 | 4,47E+08 | 4,89E+08 | 38,252 | 0 | 323,31 | 102 |
| 3,37E+07 | 3,62E+07 | 3,75E+07 | 3,26E+07 | 103,69 | 0 | 18,043 | 55 |
| 1,77E+07 | 2,14E+07 | 1,75E+07 | 1,79E+07 | 32,909 | 0 | 12,076 | 23 |
| 1,21E+07 | 2,35E+07 | 1,73E+07 | 2,73E+07 | 128,23 | 0 | 10,601 | 32 |
| 3,57E+07 | 3,14E+07 | 3,22E+07 | 2,29E+07 | 35,082 | 0 | 20,888 | 39 |
| 3,98E+07 | 3,55E+07 | 2,64E+07 | 3,18E+07 | 336 | 0 | 12,651 | 35 |
| 1,18E+08 | 1,05E+08 | 1,13E+08 | 1,16E+08 | 106,78 | 0 | 94,533 | 96 |
| 7,23E+07 | 8,67E+07 | 9,05E+07 | 9,10E+07 | 27,052 | 0 | 116,93 | 16 |
| 3,60E+06 | 8,34E+06 | 8,88E+06 | 6,91E+06 | 70,801 | 0,000158 | 5,088 | 8 |
| 1,12E+08 | 1,10E+08 | 1,24E+08 | 1,10E+08 | 29,518 | 0 | 35,636 | 47 |
| 2,68E+09 | 3,16E+09 | 2,95E+09 | 3,25E+09 | 23,384 | 0 | 323,31 | 171 |
| 1,28E+09 | 1,23E+09 | 1,27E+09 | 1,21E+09 | 22,186 | 0 | 182,24 | 160 |
| 1,10E+08 | 1,14E+08 | 1,08E+08 | 1,13E+08 | 83,305 | 0 | 59,817 | 93 |
| 1,20E+08 | 1,18E+08 | 1,21E+08 | 1,20E+08 | 63,379 | 0 | 117,28 | 56 |
| 1,35E+08 | 1,52E+08 | 1,04E+08 | 1,13E+08 | 15,391 | 0 | 48,152 | 50 |
| 5,45E+07 | 6,72E+07 | 7,50E+07 | 1,55E+08 | 97,213 | 0,000153 | 4,2388 | 12 |
| 2,95E+09 | 2,39E+09 | 2,28E+09 | 2,40E+09 | 64,279 | 0 | 323,31 | 283 |
| 4,11E+08 | 4,41E+08 | 4,32E+08 | 4,43E+08 | 21,331 | 0 | 138,94 | 113 |
| 1,51E+08 | 1,52E+08 | 1,95E+08 | 1,87E+08 | 17,806 | 0 | 56,495 | 28 |
| 5,06E+08 | 5,47E+08 | 5,29E+08 | 5,13E+08 | 185,21 | 0 | 303,18 | 267 |
| 6,75E+06 | 9,23E+06 | 6,79E+06 | 5,86E+06 | 88,993 | 0 | 6,0659 | 13 |
| 6,54E+08 | 6,24E+08 | 6,15E+08 | 5,92E+08 | 96,831 | 0 | 290,82 | 207 |
| 5,07E+07 | 4,71E+07 | 5,40E+07 | 5,71E+07 | 62,377 | 0 | 36,877 | 45 |
| 1,93E+10 | 8,57E+09 | 1,53E+10 | 1,13E+10 | 13,992 | 0 | 323,31 | 206 |
| 1,06E+07 | 9,06E+06 | 1,24E+07 | 1,16E+07 | 25,001 | 0 | 8,8763 | 10 |
| 1,06E+08 | 1,00E+08 | 1,08E+08 | 1,13E+08 | 34,238 | 0 | 88,459 | 72 |
| 1,64E+08 | 1,79E+08 | 1,48E+08 | 1,68E+08 | 49,583 | 0 | 127,09 | 84 |
| 2,29E+08 | 2,17E+08 | 2,15E+08 | 2,05E+08 | 53,924 | 0 | 160,18 | 111 |
| 6,71E+07 | 7,00E+07 | 6,40E+07 | 6,65E+07 | 63,085 | 0 | 38,461 | 59 |
| 3,71E+07 | 4,22E+07 | 4,29E+07 | 3,84E+07 | 129,33 | 0 | 25,005 | 54 |
| 1,02E+07 | 9,01E+06 | 9,41E+06 | 3,41E+06 | 9,2234 | 0,000149 | 3,5525 | 5 |
| 4,79E+08 | 3,68E+07 | 7,17E+07 | 1,62E+07 | 15,328 | 0 | 253,27 | 17 |
| 6,10E+06 | 7,70E+06 | 1,24E+07 | 9,80E+06 | 18,066 | 0 | 16,959 | 13 |
| 6,16E+06 | 7,69E+06 | 7,40E+06 | 7,65E+06 | 27,798 | 0,000154 | 4,339 | 11 |
| 4,28E+06 | 3,85E+06 | 3,06E+06 | 3,37E+06 | 114,07 | 0,00127 | 2,4028 | 5 |
| 1,85E+09 | 1,61E+09 | 1,79E+09 | 1,86E+09 | 40,489 | 0 | 323,31 | 204 |
| 2,22E+08 | 2,36E+08 | 2,04E+08 | 2,14E+08 | 47,502 | 0 | 96,973 | 79 |
| 2,82E+07 | 2,81E+07 | 2,32E+07 | 2,96E+07 | 39,552 | 0 | 6,7369 | 24 |
| 1,75E+08 | 2,13E+08 | 1,91E+08 | 1,97E+08 | 16,98 | 0 | 95,191 | 35 |
| 3,12E+06 | 1,81E+06 | 9,69E+05 | 3,54E+06 | 37,231 | 0,000157 | 4,8794 | 5 |
| 5,12E+06 | 1,12E+07 | 4,22E+06 | 9,12E+06 | 48,103 | 0 | 6,3117 | 11 |
| 4,99E+07 | 4,64E+07 | 3,97E+07 | 4,58E+07 | 24,97 | 0 | 20,855 | 24 |
| 1,22E+07 | 1,41E+07 | 1,51E+07 | 1,14E+07 | 51,51 | 0 | 11,139 | 17 |
| 2,66E+07 | 2,24E+07 | 1,67E+07 | 1,91E+07 | 31,254 | 0 | 67,934 | 22 |
| 9,90E+07 | 1,00E+08 | 1,10E+08 | 1,31E+08 | 43,485 | 0 | 163,81 | 34 |
| 2,77E+07 | 2,01E+07 | 2,38E+07 | 3,00E+07 | 126,77 | 0 | 33,172 | 31 |
| 3,42E+06 | 6,21E+06 | 4,98E+06 | 4,64E+06 | 43,286 | 0,000146 | 3,1124 | 5 |

| | | | | | | | |
|---|---|---|---|---|---|---|---|
| 1,92E+07 | 4,91E+07 | 1,50E+07 | 1,67E+07 | 10,505 | 0 | 23,841 | 11 |
| 5,14E+07 | 6,91E+07 | 1,01E+08 | 8,73E+07 | 19,997 | 0 | 5,4608 | 21 |
| 2,44E+06 | 2,73E+06 | 3,02E+06 | 2,72E+06 | 115,37 | 0,00357 | 1,8781 | 5 |
| 4,04E+06 | 2,23E+06 | 4,88E+06 | 3,51E+06 | 46,37 | 0,000428 | 2,6663 | 5 |
| 2,80E+09 | 2,78E+09 | 2,90E+09 | 2,73E+09 | 91,545 | 0 | 323,31 | 503 |
| 8,55E+06 | 1,00E+07 | 9,47E+06 | 7,87E+06 | 117,98 | 0,00043 | 2,7583 | 8 |
| 4,05E+07 | 4,45E+07 | 5,05E+07 | 5,35E+07 | 78,861 | 0 | 24,031 | 43 |
| 5,01E+06 | 4,36E+06 | 4,37E+06 | 5,57E+06 | 68,612 | 0 | 19,929 | 12 |
| 8,57E+06 | 5,11E+06 | 5,71E+06 | 5,30E+06 | 33,934 | 0,000148 | 3,3895 | 15 |
| 5,20E+08 | 5,59E+08 | 6,19E+08 | 5,52E+08 | 15,733 | 0 | 17,641 | 115 |
| 5,98E+07 | 6,52E+07 | 5,74E+07 | 5,68E+07 | 19,92 | 0 | 103,69 | 49 |
| 4,27E+07 | 5,05E+07 | 5,38E+07 | 5,16E+07 | 63,56 | 0 | 36,685 | 45 |
| 3,29E+07 | 2,56E+07 | 2,63E+07 | 3,16E+07 | 61,088 | 0 | 12,452 | 27 |
| 3,55E+07 | 3,31E+07 | 3,02E+07 | 2,48E+07 | 61,758 | 0 | 12,755 | 30 |
| 2,40E+08 | 1,97E+08 | 1,95E+08 | 1,80E+08 | 55,356 | 0 | 132,78 | 83 |
| 1,51E+08 | 1,30E+08 | 1,22E+08 | 1,47E+08 | 80,796 | 0 | 83,018 | 73 |
| 1,35E+09 | 1,03E+09 | 1,34E+09 | 9,96E+08 | 17,258 | 0 | 8,392 | 171 |
| 2,74E+08 | 3,71E+08 | 3,16E+08 | 2,96E+08 | 25,194 | 0 | 95,315 | 76 |
| 7,81E+06 | 2,64E+07 | 2,47E+07 | 2,23E+07 | 67,844 | 0 | 14,663 | 22 |
| 3,19E+06 | 3,60E+06 | 2,16E+06 | 2,07E+06 | 105,28 | 0,000152 | 4,0792 | 4 |
| 2,54E+08 | 2,40E+08 | 2,49E+08 | 2,11E+08 | 23,348 | 0 | 193,84 | 67 |
| 1,18E+08 | 1,46E+08 | 1,32E+08 | 1,21E+08 | 32,208 | 0 | 38,652 | 50 |
| 8,92E+07 | 9,02E+07 | 1,07E+08 | 9,84E+07 | 43,039 | 0 | 56,027 | 45 |
| 3,87E+07 | 3,48E+07 | 3,28E+07 | 3,13E+07 | 42,196 | 0 | 28,75 | 31 |
| 5,88E+06 | 5,87E+06 | 1,34E+07 | 4,17E+06 | 27,723 | 0 | 86,15 | 6 |
| 1,70E+08 | 1,77E+08 | 1,80E+08 | 1,50E+08 | 120,8 | 0 | 62,66 | 118 |
| 9,03E+06 | 9,82E+06 | 1,06E+07 | 1,04E+07 | 23,034 | 0,000155 | 4,5001 | 23 |
| 6,26E+08 | 1,01E+07 | 6,25E+06 | 2,35E+07 | 14,135 | 0 | 323,31 | 38 |
| 6,03E+07 | 7,19E+07 | 6,37E+07 | 6,21E+07 | 19,509 | 0 | 68,97 | 37 |
| 3,95E+06 | 1,51E+06 | 1,74E+06 | 2,91E+06 | 71,642 | 0,000154 | 4,37 | 4 |
| 2,04E+06 | 2,20E+06 | 1,71E+06 | 1,93E+06 | 62,185 | 0,003698 | 1,8243 | 2 |
| 2,50E+06 | 7,61E+06 | 3,69E+06 | 4,04E+06 | 10,135 | 0,000434 | 2,917 | 6 |
| 4,17E+09 | 4,82E+09 | 4,59E+09 | 4,65E+09 | 21,644 | 0 | 107,64 | 182 |
| 7,01E+06 | 7,02E+06 | 7,96E+06 | 7,69E+06 | 109,88 | 0 | 8,9169 | 18 |
| 8,00E+08 | 7,33E+08 | 7,48E+08 | 7,54E+08 | 100,45 | 0 | 221,22 | 251 |
| 2,09E+07 | 1,63E+07 | 2,04E+07 | 2,03E+07 | 48,811 | 0 | 6,4541 | 18 |
| 3,20E+07 | 4,78E+07 | 3,89E+07 | 4,50E+07 | 38,02 | 0 | 9,9575 | 22 |
| 4,35E+06 | 3,99E+06 | 2,43E+06 | 4,08E+06 | 50,459 | 0,000149 | 3,4657 | 4 |
| 3,25E+08 | 6,12E+08 | 3,64E+08 | 2,78E+08 | 9,7681 | 0 | 18,077 | 30 |
| 4,35E+07 | 3,88E+07 | 5,28E+07 | 3,57E+07 | 54,022 | 0 | 288,58 | 22 |
| 3,36E+07 | 3,14E+07 | 3,38E+07 | 2,25E+07 | 195,83 | 0 | 11,441 | 35 |
| 2,29E+08 | 2,41E+08 | 2,33E+08 | 2,08E+08 | 35,334 | 0 | 206,53 | 95 |
| 6,07E+09 | 5,40E+09 | 6,44E+09 | 6,01E+09 | 34,4 | 0 | 215,49 | 548 |
| 3,31E+06 | 9,48E+06 | 6,23E+06 | 6,62E+06 | 28,378 | 0,000853 | 2,5629 | 10 |
| 1,33E+07 | 5,79E+06 | 5,94E+06 | 7,07E+06 | 75,358 | 0 | 7,3901 | 15 |
| 3,20E+07 | 3,40E+07 | 2,87E+07 | 3,42E+07 | 214,42 | 0 | 16,821 | 46 |
| 1,92E+08 | 1,69E+08 | 1,85E+08 | 1,75E+08 | 134,5 | 0 | 166,46 | 166 |
| 6,87E+08 | 7,85E+08 | 8,41E+08 | 8,11E+08 | 47,789 | 0 | 323,31 | 207 |
| 1,11E+08 | 1,15E+08 | 1,23E+08 | 1,21E+08 | 28,372 | 0 | 34,93 | 65 |
| 2,27E+06 | 8,25E+06 | 9,00E+06 | 8,97E+06 | 93,393 | 0 | 7,8946 | 11 |

| | | | | | | | |
|---|---|---|---|---|---|---|---|
| 1,91E+07 | 2,87E+06 | 1,73E+07 | 1,78E+07 | 44,459 | 0,000156 | 4,6786 | 14 |
| 1,46E+09 | 1,36E+09 | 1,28E+09 | 1,23E+09 | 58,792 | 0 | 323,31 | 199 |
| 1,35E+07 | 1,11E+07 | 1,17E+07 | 1,43E+07 | 23,099 | 0 | 38,659 | 16 |
| 4,92E+06 | 3,65E+06 | 3,76E+06 | 4,66E+06 | 42,552 | 0,000146 | 3,0898 | 8 |
| 1,93E+07 | 2,61E+07 | 2,71E+07 | 2,25E+07 | 64,907 | 0 | 19,647 | 25 |
| 1,29E+06 | 8,37E+05 | 1,15E+06 | 2,38E+06 | 50,899 | 0,003047 | 1,9965 | 1 |
| 7,76E+09 | 7,57E+09 | 7,43E+09 | 7,04E+09 | 46,153 | 0 | 323,31 | 759 |
| 2,01E+08 | 2,07E+08 | 1,70E+08 | 1,82E+08 | 99,986 | 0 | 156,1 | 146 |
| 1,63E+09 | 1,22E+09 | 1,43E+09 | 1,36E+09 | 34,577 | 0 | 317,86 | 210 |
| 4,30E+07 | 4,02E+07 | 4,43E+07 | 4,45E+07 | 55,973 | 0 | 40,906 | 43 |
| 5,25E+08 | 5,23E+08 | 5,51E+08 | 5,43E+08 | 80,797 | 0 | 309,31 | 139 |
| 6,72E+07 | 6,13E+07 | 6,55E+07 | 7,10E+07 | 161,94 | 0 | 51,585 | 62 |
| 4,41E+06 | 4,40E+06 | 4,40E+06 | 3,69E+06 | 43,814 | 0,000152 | 4,08 | 6 |
| 1,37E+07 | 1,57E+07 | 1,35E+07 | 1,47E+07 | 26,053 | 0 | 16,96 | 11 |
| 4,33E+10 | 4,25E+10 | 4,38E+10 | 4,50E+10 | 50,113 | 0 | 323,31 | 1406 |
| 1,51E+07 | 1,09E+07 | 1,57E+07 | 1,44E+07 | 24,408 | 0 | 5,8158 | 24 |
| 4,47E+06 | 2,75E+06 | 5,25E+06 | 2,59E+06 | 87,81 | 0,0082 | 1,5027 | 5 |
| 1,98E+07 | 1,78E+07 | 1,53E+07 | 1,40E+07 | 34,041 | 0,000149 | 3,4326 | 8 |
| 5,43E+07 | 5,01E+07 | 5,22E+07 | 5,25E+07 | 19,538 | 0 | 11,941 | 26 |
| 1,38E+09 | 1,31E+09 | 1,28E+09 | 1,30E+09 | 70,875 | 0 | 323,31 | 333 |
| 3,92E+09 | 5,08E+09 | 4,84E+09 | 4,95E+09 | 23,564 | 0 | 46,532 | 238 |
| 1,65E+07 | 2,08E+07 | 1,75E+07 | 2,09E+07 | 54,388 | 0 | 8,9754 | 16 |
| 1,32E+08 | 1,20E+08 | 1,24E+08 | 1,14E+08 | 21,336 | 0 | 236,04 | 53 |
| 2,70E+07 | 2,02E+07 | 2,69E+07 | 2,20E+07 | 50,469 | 0 | 16,311 | 27 |
| 7,09E+08 | 7,46E+08 | 5,65E+08 | 5,91E+08 | 33,338 | 0,000148 | 3,3086 | 35 |
| 2,11E+08 | 2,05E+08 | 2,03E+08 | 1,95E+08 | 47,513 | 0 | 156,1 | 137 |
| 1,68E+06 | 2,02E+06 | 2,19E+06 | 1,40E+06 | 89,075 | 0,000147 | 3,1787 | 12 |
| 9,16E+08 | 9,41E+08 | 9,81E+08 | 1,07E+09 | 12,215 | 0 | 6,9606 | 124 |
| 6,68E+06 | 7,52E+06 | 6,86E+06 | 5,03E+06 | 78,572 | 0 | 6,7522 | 21 |
| 5,25E+07 | 6,16E+07 | 4,45E+07 | 5,04E+07 | 36,971 | 0 | 35,226 | 47 |
| 1,55E+09 | 1,61E+09 | 1,59E+09 | 1,66E+09 | 141,55 | 0 | 323,31 | 497 |
| 3,01E+09 | 3,01E+09 | 3,01E+09 | 3,04E+09 | 243,24 | 0 | 323,31 | 799 |
| 8,81E+06 | 1,23E+07 | 1,55E+07 | 1,56E+07 | 34,168 | 0 | 5,7166 | 20 |
| 7,25E+07 | 6,11E+07 | 2,88E+08 | 3,14E+08 | 59,51 | 0 | 76,858 | 67 |
| 3,21E+07 | 4,64E+07 | 4,88E+07 | 3,84E+07 | 89,744 | 0 | 16,911 | 44 |
| 7,63E+06 | 1,53E+07 | 1,26E+07 | 1,11E+07 | 35,095 | 0 | 35,04 | 7 |
| 1,83E+07 | 1,85E+07 | 2,19E+07 | 2,78E+07 | 97,189 | 0,000151 | 3,8348 | 18 |
| 2,20E+07 | 2,51E+07 | 2,22E+07 | 2,47E+07 | 20,405 | 0 | 10,215 | 25 |
| 5,79E+09 | 6,04E+09 | 5,85E+09 | 6,28E+09 | 34,216 | 0 | 323,31 | 418 |
| 3,37E+06 | 3,85E+06 | 2,01E+06 | 1,92E+06 | 25,616 | 0 | 5,4941 | 4 |
| 9,44E+05 | 3,60E+06 | 3,23E+06 | 6,05E+06 | 337 | 0,000434 | 2,9137 | 3 |
| 2,27E+08 | 3,21E+08 | 3,55E+08 | 3,47E+08 | 17,247 | 0 | 277,14 | 79 |
| 6,24E+07 | 6,24E+07 | 7,32E+07 | 6,49E+07 | 29,639 | 0 | 20,445 | 46 |
| 6,47E+06 | 3,62E+06 | 4,11E+06 | 4,55E+06 | 62,289 | 0,000152 | 3,946 | 9 |
| 4,10E+07 | 3,31E+07 | 4,02E+07 | 4,15E+07 | 250,63 | 0 | 12,119 | 25 |
| 1,09E+08 | 1,23E+08 | 9,71E+07 | 1,12E+08 | 19,43 | 0 | 40,805 | 16 |
| 7,66E+08 | 7,13E+08 | 7,95E+08 | 7,65E+08 | 116,05 | 0 | 323,31 | 241 |
| 3,28E+08 | 2,64E+08 | 2,87E+08 | 3,13E+08 | 63,506 | 0 | 133,6 | 85 |
| 8,35E+06 | 7,14E+06 | 7,12E+06 | 9,74E+06 | 88,326 | 0 | 9,3039 | 10 |
| 3,57E+09 | 4,01E+09 | 4,46E+09 | 4,66E+09 | 15,942 | 0 | 205,41 | 185 |

| | | | | | | | |
|---|---|---|---|---|---|---|---|
| 1,81E+07 | 1,54E+07 | 1,48E+07 | 1,62E+07 | 93,275 | 0 | 38,535 | 26 |
| 8,63E+06 | 3,69E+06 | 5,25E+06 | 6,81E+06 | 94,495 | 0,000854 | 2,5849 | 17 |
| 1,08E+07 | 1,20E+07 | 8,07E+06 | 1,09E+07 | 29,659 | 0,003443 | 1,9178 | 8 |
| 7,41E+08 | 7,81E+08 | 7,33E+08 | 7,32E+08 | 90,884 | 0 | 323,31 | 281 |
| 1,34E+08 | 1,36E+08 | 1,28E+08 | 1,38E+08 | 317,06 | 0 | 93,79 | 113 |
| 4,43E+09 | 4,67E+09 | 4,80E+09 | 4,83E+09 | 66,406 | 0 | 323,31 | 453 |
| 4,79E+07 | 4,92E+07 | 5,13E+07 | 4,76E+07 | 34,706 | 0 | 7,1287 | 32 |
| 1,07E+07 | 2,37E+06 | 1,07E+07 | 7,43E+06 | 34,105 | 0 | 10,521 | 7 |
| 8,85E+07 | 8,93E+07 | 7,86E+07 | 8,03E+07 | 51,777 | 0 | 76,386 | 64 |
| 2,00E+08 | 1,83E+08 | 2,12E+08 | 2,22E+08 | 139,99 | 0 | 122,03 | 100 |
| 6,77E+06 | 6,36E+07 | 8,64E+06 | 5,25E+06 | 110,58 | 0,000431 | 2,8038 | 6 |
| 3,57E+07 | 3,48E+07 | 3,69E+07 | 3,82E+07 | 51,23 | 0 | 13,557 | 31 |
| 7,98E+07 | 8,59E+07 | 7,65E+07 | 7,24E+07 | 94,927 | 0 | 32,158 | 59 |
| 7,68E+07 | 1,06E+08 | 1,18E+08 | 1,18E+08 | 47,095 | 0 | 21,486 | 62 |
| 1,35E+07 | 1,52E+07 | 1,65E+07 | 1,62E+07 | 68,464 | 0 | 9,7104 | 23 |
| 8,76E+06 | 4,72E+06 | 6,81E+06 | 1,55E+06 | 62,898 | 0 | 33,818 | 5 |
| 4,80E+07 | 6,41E+07 | 6,49E+07 | 5,81E+07 | 521,23 | 0 | 28,452 | 61 |
| 3,23E+09 | 3,18E+09 | 3,31E+09 | 3,75E+09 | 20,252 | 0 | 226,49 | 224 |
| 4,57E+06 | 5,95E+06 | 4,76E+06 | 4,47E+06 | 57,329 | 0 | 5,5585 | 7 |
| 1,02E+08 | 1,09E+08 | 1,14E+08 | 1,15E+08 | 72,461 | 0 | 98,288 | 69 |
| 3,28E+08 | 2,64E+08 | 3,20E+08 | 3,00E+08 | 37,703 | 0 | 278,1 | 92 |
| 2,07E+08 | 1,93E+08 | 2,15E+08 | 2,41E+08 | 78,947 | 0 | 34,332 | 51 |
| 5,78E+07 | 7,59E+07 | 1,08E+08 | 1,26E+08 | 32,97 | 0 | 97,919 | 42 |
| 9,96E+06 | 2,84E+07 | 2,20E+07 | 2,82E+07 | 73,432 | 0 | 29,929 | 25 |
| 8,96E+08 | 9,01E+08 | 8,65E+08 | 8,44E+08 | 82,897 | 0 | 323,31 | 254 |
| 4,87E+07 | 5,05E+06 | 3,61E+07 | 2,91E+07 | 7,4043 | 0 | 23,049 | 16 |
| 1,72E+08 | 1,82E+08 | 1,67E+08 | 1,87E+08 | 19,133 | 0 | 204,09 | 53 |
| 8,63E+06 | 6,26E+06 | 1,08E+07 | 7,90E+06 | 39,651 | 0,000154 | 4,4126 | 10 |
| 5,12E+08 | 5,41E+08 | 5,03E+08 | 5,19E+08 | 23,412 | 0 | 323,31 | 109 |
| 2,45E+06 | 3,37E+06 | 2,94E+06 | 3,00E+06 | 103,53 | 0,000149 | 3,5895 | 4 |
| 2,35E+09 | 1,87E+09 | 2,14E+09 | 1,66E+09 | 17,779 | 0 | 18,535 | 130 |
| 1,89E+07 | 1,60E+07 | 2,35E+07 | 1,41E+07 | 56,624 | 0 | 19,22 | 20 |
| 8,28E+08 | 5,30E+08 | 5,14E+08 | 4,33E+08 | 28,086 | 0 | 323,31 | 144 |
| 6,11E+08 | 6,94E+08 | 7,42E+08 | 7,66E+08 | 13,281 | 0 | 256,15 | 64 |
| 5,07E+07 | 5,10E+07 | 4,98E+07 | 5,02E+07 | 25,594 | 0 | 50,376 | 28 |
| 2,01E+07 | 1,24E+06 | 1,35E+07 | 4,23E+06 | 28,274 | 0,004626 | 1,705 | 2 |
| 3,86E+07 | 7,50E+07 | 6,29E+07 | 6,20E+07 | 105,11 | 0 | 40,882 | 58 |
| 3,34E+06 | 4,76E+06 | 2,66E+06 | 2,65E+06 | 51,427 | 0,000433 | 2,8315 | 7 |
| 4,89E+09 | 4,55E+09 | 4,83E+09 | 4,70E+09 | 62,581 | 0 | 323,31 | 536 |
| 3,76E+07 | 6,12E+06 | 1,01E+07 | 1,74E+07 | 53,525 | 0,001547 | 2,2985 | 11 |
| 7,54E+08 | 7,25E+08 | 6,65E+08 | 7,45E+08 | 49,694 | 0 | 323,31 | 154 |
| 9,16E+07 | 8,16E+07 | 1,06E+08 | 1,06E+08 | 45,345 | 0 | 60,312 | 18 |
| 7,08E+07 | 6,90E+07 | 7,18E+07 | 7,43E+07 | 52,307 | 0 | 55,909 | 49 |
| 4,35E+08 | 3,66E+08 | 3,32E+08 | 3,85E+08 | 9,9583 | 0 | 6,9889 | 25 |
| 4,79E+07 | 5,31E+07 | 6,40E+07 | 5,32E+07 | 22,787 | 0 | 14,028 | 10 |
| 1,01E+09 | 9,22E+08 | 8,75E+08 | 9,64E+08 | 78,025 | 0 | 323,31 | 252 |
| 5,86E+08 | 6,21E+08 | 5,97E+08 | 5,59E+08 | 35,88 | 0 | 144,51 | 126 |
| 4,51E+06 | 4,07E+07 | 3,39E+07 | 3,78E+07 | 147,71 | 0,002915 | 2,0251 | 6 |
| 3,00E+06 | 1,08E+06 | 2,77E+06 | 3,61E+06 | 60,916 | 0,000436 | 3,0254 | 4 |
| 1,55E+07 | 1,71E+07 | 1,49E+07 | 1,27E+07 | 36,919 | 0 | 7,6877 | 12 |

| | | | | | | | |
|---|---|---|---|---|---|---|---|
| 3,17E+08 | 3,35E+08 | 3,40E+08 | 3,82E+08 | 29,812 | 0 | 238,84 | 103 |
| 5,58E+06 | 1,57E+07 | 1,28E+07 | 1,40E+07 | 46,039 | 0 | 16,012 | 21 |
| 7,25E+07 | 2,09E+08 | 6,43E+07 | 1,31E+08 | 12,746 | 0 | 323,31 | 13 |
| 4,37E+06 | 2,85E+07 | 3,74E+07 | 3,20E+07 | 24,56 | 0 | 74,742 | 19 |
| 8,23E+06 | 3,75E+06 | 2,22E+07 | 2,74E+06 | 29,215 | 0,003965 | 1,7879 | 3 |
| 2,43E+08 | 2,79E+08 | 2,93E+08 | 3,03E+08 | 73,384 | 0 | 57,873 | 151 |
| 1,71E+10 | 1,63E+10 | 1,66E+10 | 1,59E+10 | 92,475 | 0 | 323,31 | 1436 |
| 1,55E+06 | 8,98E+06 | 3,95E+06 | 9,09E+06 | 111,67 | 0,001549 | 2,3228 | 6 |
| 1,49E+07 | 4,44E+06 | 1,05E+07 | 4,01E+06 | 70,751 | 0,005022 | 1,6727 | 5 |
| 5,24E+07 | 6,59E+07 | 6,32E+07 | 6,75E+07 | 16,954 | 0 | 15,061 | 35 |
| 1,29E+08 | 2,17E+08 | 1,67E+08 | 2,29E+08 | 35,955 | 0 | 38,406 | 77 |
| 2,57E+08 | 2,33E+08 | 2,66E+08 | 2,68E+08 | 74,152 | 0 | 124,56 | 146 |
| 2,49E+06 | 2,85E+06 | 3,88E+06 | 2,82E+06 | 131,29 | 0,008198 | 1,5005 | 5 |
| 7,12E+06 | 4,70E+06 | 4,13E+06 | 4,72E+06 | 46,145 | 0 | 5,9667 | 10 |
| 6,42E+08 | 4,48E+08 | 5,70E+08 | 4,74E+08 | 24,604 | 0 | 167,35 | 224 |
| 1,24E+08 | 1,17E+08 | 1,09E+08 | 1,01E+08 | 65,704 | 0 | 14,986 | 71 |
| 1,73E+08 | 1,54E+08 | 1,80E+08 | 1,88E+08 | 37,022 | 0 | 197,87 | 55 |
| 2,29E+08 | 2,28E+08 | 2,45E+08 | 2,11E+08 | 34,934 | 0 | 53,686 | 60 |
| 5,38E+07 | 4,96E+07 | 4,33E+07 | 4,24E+07 | 41,598 | 0 | 20,253 | 30 |
| 2,68E+07 | 3,37E+07 | 3,22E+07 | 2,69E+07 | 31,379 | 0 | 9,4604 | 39 |
| 1,01E+07 | 1,17E+07 | 1,03E+07 | 2,48E+06 | 12,159 | 0,009242 | 1,4426 | 9 |
| 4,46E+06 | 7,37E+06 | 3,63E+06 | 5,49E+06 | 78,946 | 0,009109 | 1,4428 | 3 |
| 3,21E+08 | 3,10E+08 | 3,52E+08 | 3,43E+08 | 21,258 | 0 | 176,45 | 147 |
| 3,20E+07 | 9,33E+06 | 1,69E+07 | 1,32E+07 | 107,54 | 0 | 7,7843 | 14 |
| 2,51E+07 | 1,82E+07 | 1,92E+07 | 1,64E+07 | 52,244 | 0 | 14,749 | 15 |
| 1,24E+08 | 1,97E+08 | 1,49E+08 | 1,48E+08 | 26,157 | 0 | 32,527 | 90 |
| 1,34E+07 | 2,35E+07 | 3,50E+07 | 2,83E+07 | 301,05 | 0 | 38,567 | 41 |
| 2,10E+08 | 1,27E+08 | 1,34E+08 | 5,42E+07 | 107,25 | 0,008976 | 1,4511 | 13 |
| 2,10E+07 | 1,85E+07 | 2,82E+07 | 3,00E+07 | 37,869 | 0 | 30,92 | 16 |
| 3,30E+08 | 3,24E+08 | 3,23E+08 | 3,50E+08 | 327,45 | 0 | 323,31 | 250 |
| 1,13E+08 | 1,44E+08 | 1,40E+08 | 1,26E+08 | 96,466 | 0 | 39,79 | 89 |
| 5,62E+06 | 1,13E+06 | 2,96E+06 | 2,45E+06 | 30,25 | 0,000428 | 2,6692 | 5 |
| 4,36E+07 | 4,29E+07 | 4,19E+07 | 4,40E+07 | 112,76 | 0 | 13,826 | 37 |
| 2,13E+09 | 2,44E+09 | 2,34E+09 | 2,53E+09 | 14,865 | 0 | 208,69 | 253 |
| 1,81E+09 | 1,40E+09 | 1,56E+09 | 1,35E+09 | 18,007 | 0 | 239,61 | 169 |
| 4,88E+07 | 4,30E+07 | 5,03E+07 | 4,58E+07 | 191,57 | 0 | 22,286 | 59 |
| 5,00E+07 | 4,99E+07 | 5,34E+07 | 6,26E+07 | 19,778 | 0,00015 | 3,6772 | 22 |
| 9,08E+06 | 9,76E+06 | 7,87E+06 | 6,04E+06 | 32,436 | 0 | 6,341 | 22 |
| 1,85E+10 | 1,37E+10 | 1,42E+10 | 1,53E+10 | 76,722 | 0 | 323,31 | 961 |
| 1,80E+08 | 1,69E+08 | 1,55E+08 | 1,70E+08 | 39,916 | 0 | 54,543 | 86 |
| 2,56E+06 | 5,29E+06 | 6,77E+06 | 3,66E+06 | 52,614 | 0,000436 | 3,0238 | 2 |
| 4,72E+07 | 4,88E+07 | 3,90E+07 | 4,28E+07 | 56,576 | 0 | 13,246 | 30 |
| 2,36E+06 | 1,64E+07 | 1,73E+07 | 1,37E+07 | 43,458 | 0 | 7,7944 | 11 |
| 9,33E+08 | 9,69E+08 | 8,87E+08 | 8,88E+08 | 137,61 | 0 | 281,43 | 326 |
| 4,00E+07 | 2,84E+07 | 3,55E+07 | 3,89E+07 | 72,917 | 0 | 22,63 | 35 |
| 2,00E+08 | 2,32E+08 | 1,98E+08 | 1,88E+08 | 40,685 | 0 | 59,453 | 93 |
| 6,65E+07 | 7,37E+07 | 7,24E+07 | 6,91E+07 | 20,959 | 0 | 31,119 | 50 |
| 2,84E+08 | 2,59E+08 | 2,62E+08 | 2,25E+08 | 29,8 | 0 | 136,3 | 114 |
| 1,14E+07 | 5,63E+06 | 4,83E+06 | 4,11E+06 | 38,076 | 0 | 6,287 | 10 |
| 1,87E+07 | 1,31E+07 | 1,63E+07 | 1,68E+07 | 110,09 | 0 | 8,0341 | 16 |

| | | | | | | | |
|---|---|---|---|---|---|---|---|
| 2,81E+06 | 7,11E+06 | 4,79E+06 | 5,14E+06 | 47,8 | 0,000151 | 3,7725 | 6 |
| 1,44E+06 | 5,28E+06 | 1,82E+06 | 1,30E+06 | 72,088 | 0,000434 | 2,8927 | 2 |
| 2,05E+08 | 2,05E+08 | 2,00E+08 | 2,15E+08 | 33,448 | 0 | 39,829 | 73 |
| 9,20E+06 | 9,43E+06 | 9,63E+06 | 1,05E+07 | 43,144 | 0,000152 | 3,9833 | 8 |
| 8,09E+06 | 9,89E+06 | 7,68E+06 | 1,02E+07 | 30,673 | 0,000154 | 4,3159 | 11 |
| 2,70E+08 | 3,13E+08 | 2,99E+08 | 2,96E+08 | 24,106 | 0 | 254,38 | 131 |
| 1,17E+07 | 1,56E+07 | 1,20E+07 | 1,26E+07 | 23,562 | 0 | 23,331 | 14 |
| 8,39E+06 | 6,55E+06 | 6,08E+06 | 7,39E+06 | 39,991 | 0 | 67,219 | 9 |
| 7,78E+06 | 1,81E+07 | 1,60E+07 | 1,32E+07 | 158,83 | 0 | 9,4888 | 24 |
| 8,30E+07 | 7,97E+07 | 5,16E+07 | 3,88E+07 | 7,7412 | 0 | 12,474 | 61 |
| 1,25E+07 | 9,14E+06 | 1,17E+07 | 1,37E+07 | 11,813 | 0,00015 | 3,5987 | 8 |
| 1,98E+08 | 1,82E+08 | 1,85E+08 | 1,89E+08 | 85,299 | 0 | 131,84 | 102 |
| 2,62E+07 | 1,66E+07 | 1,73E+07 | 2,02E+07 | 155,98 | 0 | 14,601 | 22 |
| 5,07E+07 | 4,62E+07 | 3,43E+07 | 3,94E+07 | 95,911 | 0 | 61,94 | 41 |
| 8,60E+07 | 7,98E+07 | 7,97E+07 | 8,45E+07 | 184,83 | 0 | 155,67 | 101 |
| 1,41E+06 | 1,05E+07 | 1,03E+07 | 1,15E+07 | 28,103 | 0,000148 | 3,351 | 17 |
| 1,82E+08 | 1,44E+08 | 1,61E+08 | 1,87E+08 | 42,024 | 0 | 162,55 | 46 |
| 6,53E+07 | 6,52E+07 | 6,94E+07 | 6,69E+07 | 35,297 | 0 | 32,043 | 48 |
| 5,62E+07 | 5,45E+07 | 5,65E+07 | 5,77E+07 | 69,579 | 0 | 29,478 | 48 |
| 4,52E+06 | 3,02E+06 | 2,84E+06 | 3,61E+06 | 53,406 | 0,008074 | 1,5153 | 6 |
| 3,58E+07 | 3,51E+07 | 2,38E+07 | 2,52E+07 | 76,425 | 0 | 20,959 | 32 |
| 5,63E+08 | 7,02E+08 | 5,27E+08 | 5,90E+08 | 19,889 | 0 | 323,31 | 122 |
| 5,19E+06 | 7,72E+06 | 4,95E+06 | 1,12E+07 | 52,717 | 0 | 9,1916 | 16 |
| 1,59E+07 | 6,84E+06 | 2,04E+06 | 2,99E+06 | 51,847 | 0,002917 | 2,0406 | 7 |
| 5,62E+06 | 5,76E+06 | 1,18E+07 | 1,09E+07 | 110,07 | 0 | 5,6729 | 10 |
| 3,26E+07 | 3,21E+07 | 3,87E+07 | 3,27E+07 | 45,863 | 0 | 12,411 | 37 |
| 6,94E+07 | 5,94E+07 | 1,03E+08 | 1,03E+08 | 13,163 | 0 | 9,8923 | 18 |
| 9,66E+06 | 8,67E+06 | 1,23E+07 | 1,11E+07 | 125,06 | 0,000156 | 4,6213 | 10 |
| 5,31E+06 | 6,56E+06 | 5,23E+06 | 7,58E+06 | 65,825 | 0,000155 | 4,5161 | 9 |
| 6,64E+07 | 2,23E+08 | 1,47E+08 | 1,74E+08 | 29,661 | 0 | 195,46 | 130 |
| 4,80E+07 | 6,75E+07 | 4,21E+07 | 5,26E+07 | 180,38 | 0,000291 | 3,0671 | 14 |
| 1,38E+07 | 6,66E+06 | 8,70E+06 | 4,74E+06 | 89,312 | 0 | 10,153 | 18 |
| 2,47E+07 | 1,06E+07 | 1,13E+07 | 1,71E+07 | 146,22 | 0 | 9,1308 | 18 |
| 1,06E+08 | 1,17E+08 | 9,63E+07 | 9,70E+07 | 46,207 | 0 | 166,97 | 75 |
| 4,59E+07 | 4,95E+07 | 4,51E+07 | 4,65E+07 | 79,638 | 0 | 58,218 | 45 |
| 6,82E+06 | 7,33E+06 | 7,96E+06 | 5,68E+06 | 61,509 | 0 | 6,5025 | 14 |
| 1,11E+07 | 9,21E+06 | 1,11E+07 | 6,49E+06 | 100,36 | 0,001549 | 2,3209 | 9 |
| 2,05E+08 | 2,19E+08 | 2,13E+08 | 1,90E+08 | 31,195 | 0 | 197,36 | 60 |
| 1,32E+08 | 9,51E+07 | 8,25E+07 | 8,60E+07 | 21,139 | 0 | 126,47 | 42 |
| 3,69E+08 | 3,88E+08 | 3,61E+08 | 3,39E+08 | 242,09 | 0 | 323,31 | 242 |
| 1,46E+06 | 3,63E+07 | 8,55E+06 | 6,92E+06 | 56,253 | 0,000148 | 3,3777 | 14 |
| 2,48E+08 | 1,79E+08 | 2,29E+08 | 2,14E+08 | 18,698 | 0 | 50,916 | 77 |
| 4,27E+06 | 4,17E+06 | 5,31E+06 | 4,09E+06 | 31,205 | 0,000148 | 3,4224 | 10 |
| 2,16E+07 | 2,74E+07 | 3,24E+07 | 2,16E+07 | 104,28 | 0 | 11,239 | 27 |
| 8,78E+07 | 6,38E+07 | 9,88E+07 | 1,07E+08 | 24,778 | 0 | 77,29 | 16 |
| 1,85E+07 | 1,49E+07 | 1,91E+07 | 1,64E+07 | 58,151 | 0 | 8,3596 | 28 |
| 4,04E+06 | 6,74E+06 | 8,89E+06 | 5,77E+06 | 444,01 | 0,000147 | 3,29 | 7 |
| 2,01E+08 | 1,46E+08 | 6,51E+07 | 1,13E+08 | 11,451 | 0 | 17,205 | 32 |
| 3,79E+07 | 4,68E+07 | 4,29E+07 | 3,82E+07 | 36,953 | 0 | 13,68 | 22 |
| 1,26E+08 | 9,37E+07 | 1,16E+08 | 1,06E+08 | 9,1275 | 0 | 7,157 | 27 |

| | | | | | | | |
|---|---|---|---|---|---|---|---|
| 1,29E+07 | 2,06E+07 | 1,30E+07 | 1,45E+07 | 62,781 | 0 | 10,201 | 24 |
| 1,39E+08 | 1,39E+08 | 1,40E+08 | 1,38E+08 | 50,87 | 0 | 117,01 | 67 |
| 1,15E+09 | 1,41E+09 | 1,55E+09 | 1,44E+09 | 29,47 | 0 | 323,31 | 110 |
| 3,82E+08 | 3,36E+08 | 3,43E+08 | 3,27E+08 | 67,317 | 0 | 238,82 | 147 |
| 6,15E+07 | 6,12E+07 | 6,81E+07 | 6,03E+07 | 192,21 | 0 | 44,934 | 86 |
| 4,22E+08 | 3,80E+08 | 3,88E+08 | 3,81E+08 | 49,783 | 0 | 85,783 | 111 |
| 9,47E+07 | 8,91E+07 | 7,93E+07 | 8,85E+07 | 19,742 | 0 | 27,582 | 43 |
| 3,84E+06 | 8,66E+06 | 7,77E+06 | 2,32E+06 | 70,251 | 0,000147 | 3,2429 | 4 |
| 2,75E+07 | 2,51E+06 | 1,60E+07 | 2,53E+06 | 81,733 | 0,006075 | 1,5904 | 5 |
| 7,56E+09 | 7,34E+09 | 7,66E+09 | 7,61E+09 | 70,67 | 0 | 323,31 | 569 |
| 4,46E+06 | 5,92E+06 | 5,34E+06 | 4,48E+06 | 141,74 | 0,00015 | 3,635 | 8 |
| 9,06E+06 | 8,99E+06 | 8,13E+06 | 9,62E+06 | 44,713 | 0 | 18,564 | 13 |
| 1,30E+08 | 1,24E+08 | 1,39E+08 | 1,45E+08 | 70,042 | 0 | 39,081 | 66 |
| 1,30E+08 | 1,38E+08 | 1,33E+08 | 1,24E+08 | 17,121 | 0 | 197,23 | 70 |
| 5,29E+07 | 3,81E+07 | 5,07E+07 | 1,59E+06 | 12,436 | 0 | 37,466 | 14 |
| 1,76E+08 | 1,39E+08 | 1,20E+08 | 9,06E+07 | 16,311 | 0 | 240,97 | 46 |
| 1,38E+08 | 1,40E+08 | 1,56E+08 | 1,85E+08 | 26,178 | 0 | 20,608 | 65 |
| 1,72E+09 | 1,79E+09 | 2,02E+09 | 2,17E+09 | 23,532 | 0 | 323,31 | 99 |
| 2,75E+07 | 2,60E+07 | 1,85E+07 | 1,91E+07 | 215,35 | 0 | 6,7461 | 11 |
| 5,09E+06 | 4,52E+06 | 5,71E+06 | 8,10E+06 | 15,838 | 0 | 28,833 | 6 |
| 2,37E+07 | 2,21E+07 | 2,60E+07 | 2,41E+07 | 33,753 | 0 | 12,43 | 25 |
| 5,55E+06 | 4,69E+06 | 1,22E+06 | 4,06E+06 | 41,234 | 0,000852 | 2,5166 | 5 |
| 8,15E+08 | 6,17E+08 | 6,94E+08 | 6,65E+08 | 23,035 | 0 | 285,73 | 98 |
| 2,79E+09 | 3,00E+09 | 3,00E+09 | 3,13E+09 | 57,147 | 0 | 323,31 | 446 |
| 9,05E+07 | 8,69E+07 | 8,34E+07 | 9,68E+07 | 37,704 | 0 | 90,104 | 68 |
| 2,36E+07 | 2,31E+07 | 2,61E+07 | 2,24E+07 | 21,607 | 0,000153 | 4,1154 | 22 |
| 7,37E+08 | 7,20E+08 | 5,95E+08 | 5,28E+08 | 32,277 | 0 | 245,74 | 218 |
| 8,70E+08 | 8,39E+08 | 9,02E+08 | 8,60E+08 | 31,27 | 0 | 323,31 | 153 |
| 5,17E+06 | 4,74E+06 | 5,92E+06 | 6,38E+06 | 63,377 | 0,000153 | 4,112 | 10 |
| 1,09E+08 | 8,35E+07 | 9,36E+07 | 9,76E+07 | 98,112 | 0 | 74,522 | 98 |
| 2,30E+08 | 2,32E+08 | 2,17E+08 | 2,24E+08 | 24,301 | 0 | 141,46 | 53 |
| 2,27E+07 | 3,40E+07 | 2,78E+07 | 2,77E+07 | 20,14 | 0 | 6,7747 | 19 |
| 4,90E+07 | 3,10E+07 | 4,09E+07 | 4,10E+07 | 130,28 | 0 | 40,938 | 40 |
| 2,43E+08 | 2,55E+08 | 2,47E+08 | 2,42E+08 | 59,69 | 0 | 71,409 | 101 |
| 1,40E+08 | 1,14E+08 | 1,42E+08 | 1,22E+08 | 63,893 | 0 | 74,285 | 56 |
| 2,11E+07 | 3,14E+07 | 2,63E+07 | 2,86E+07 | 88,216 | 0,000152 | 3,9444 | 16 |
| 2,67E+08 | 2,74E+08 | 3,26E+08 | 2,67E+08 | 9,0303 | 0 | 86,09 | 47 |
| 1,75E+08 | 1,60E+08 | 1,75E+08 | 1,74E+08 | 43,082 | 0 | 195,85 | 71 |
| 3,18E+07 | 1,81E+07 | 2,95E+07 | 3,01E+07 | 36,707 | 0 | 6,0397 | 9 |
| 2,68E+06 | 8,11E+07 | 9,01E+07 | 1,08E+08 | 87,455 | 0 | 10,131 | 25 |
| 1,62E+07 | 6,38E+06 | 8,49E+06 | 7,43E+06 | 104,92 | 0 | 23,368 | 18 |
| 5,46E+07 | 8,06E+07 | 7,18E+07 | 6,66E+07 | 42,469 | 0 | 39,552 | 50 |
| 3,05E+06 | 1,88E+06 | 2,96E+06 | 4,60E+06 | 130,4 | 0,008455 | 1,474 | 4 |
| 1,24E+08 | 1,60E+08 | 2,77E+08 | 1,69E+08 | 15,537 | 0 | 269,44 | 101 |
| 4,76E+07 | 5,76E+07 | 5,58E+07 | 6,55E+07 | 231,4 | 0 | 24,521 | 65 |
| 1,30E+08 | 1,08E+08 | 1,14E+08 | 1,00E+08 | 71,453 | 0 | 143,26 | 69 |
| 3,14E+06 | 2,67E+06 | 9,87E+06 | 1,55E+06 | 30,902 | 0 | 6,5598 | 5 |
| 2,20E+06 | 1,59E+07 | 1,90E+07 | 1,98E+07 | 42,691 | 0,000149 | 3,5808 | 9 |
| 2,82E+06 | 5,69E+06 | 2,52E+06 | 3,24E+06 | 38,636 | 0,000433 | 2,8437 | 4 |
| 6,30E+07 | 6,52E+07 | 6,41E+07 | 6,79E+07 | 41,789 | 0 | 33,506 | 31 |

| | | | | | | | |
|---|---|---|---|---|---|---|---|
| 2,33E+08 | 1,11E+08 | 1,33E+08 | 1,10E+08 | 33,046 | 0 | 116,14 | 66 |
| 2,76E+09 | 2,42E+09 | 2,39E+09 | 2,31E+09 | 15,524 | 0 | 323,31 | 214 |
| 2,95E+08 | 2,81E+08 | 2,76E+08 | 3,03E+08 | 134,07 | 0 | 304,71 | 153 |
| 2,08E+08 | 1,93E+08 | 2,02E+08 | 2,02E+08 | 39,384 | 0 | 38,05 | 102 |
| 3,06E+08 | 3,02E+08 | 3,68E+08 | 3,83E+08 | 116,92 | 0 | 215,27 | 128 |
| 1,17E+09 | 1,11E+09 | 1,29E+09 | 1,08E+09 | 48,994 | 0 | 323,31 | 176 |
| 4,29E+07 | 4,38E+07 | 4,78E+07 | 5,26E+07 | 54,594 | 0 | 16,721 | 49 |
| 2,03E+07 | 1,37E+07 | 1,59E+07 | 1,91E+07 | 74,234 | 0 | 15,024 | 26 |
| 4,26E+08 | 4,24E+08 | 3,70E+08 | 3,86E+08 | 23,598 | 0 | 199,95 | 92 |
| 2,24E+07 | 2,81E+07 | 3,22E+07 | 3,04E+07 | 115,85 | 0 | 22,013 | 36 |
| 1,05E+09 | 9,73E+08 | 1,04E+09 | 1,11E+09 | 44,93 | 0 | 323,31 | 181 |
| 2,18E+08 | 2,22E+08 | 1,95E+08 | 2,53E+08 | 142,88 | 0 | 13,252 | 24 |
| 1,36E+07 | 1,65E+07 | 1,98E+07 | 1,16E+07 | 46,596 | 0,000159 | 5,1754 | 16 |
| 2,85E+08 | 2,95E+08 | 3,01E+08 | 2,83E+08 | 123,84 | 0 | 219,44 | 144 |
| 9,10E+06 | 1,00E+07 | 1,01E+07 | 1,01E+07 | 24,959 | 0,000158 | 5,1506 | 14 |
| 1,06E+07 | 1,14E+07 | 1,19E+07 | 8,84E+06 | 21,283 | 0 | 7,8561 | 21 |
| 2,45E+09 | 1,67E+09 | 1,94E+09 | 1,69E+09 | 63,963 | 0 | 323,31 | 278 |
| 3,35E+07 | 4,76E+07 | 5,32E+07 | 5,92E+07 | 14,904 | 0 | 199,15 | 30 |
| 4,07E+06 | 3,50E+06 | 5,24E+06 | 5,83E+06 | 68,001 | 0,000148 | 3,3647 | 6 |
| 3,66E+06 | 5,90E+06 | 6,43E+06 | 1,27E+06 | 370,66 | 0,006076 | 1,5952 | 4 |
| 1,60E+06 | 2,40E+06 | 2,04E+06 | 1,77E+06 | 40,308 | 0,006075 | 1,5939 | 6 |
| 5,61E+07 | 7,34E+07 | 6,36E+07 | 7,19E+07 | 24,8 | 0 | 5,4331 | 16 |
| 1,79E+08 | 1,80E+08 | 2,51E+08 | 2,69E+08 | 26,032 | 0 | 80,212 | 57 |
| 4,02E+06 | 2,53E+06 | 5,82E+06 | 7,20E+06 | 46,846 | 0,002507 | 2,0747 | 4 |
| 2,81E+08 | 2,05E+08 | 1,90E+08 | 1,63E+08 | 43,131 | 0 | 33,459 | 67 |
| 4,54E+08 | 4,73E+08 | 4,89E+08 | 4,56E+08 | 42,613 | 0 | 323,31 | 96 |
| 3,56E+06 | 1,55E+07 | 1,66E+07 | 1,30E+07 | 77,832 | 0 | 12,68 | 23 |
| 1,69E+07 | 1,29E+07 | 1,85E+07 | 1,35E+07 | 20,398 | 0 | 13,11 | 11 |
| 1,36E+07 | 1,30E+07 | 1,43E+07 | 1,43E+07 | 70,617 | 0 | 12,685 | 28 |
| 1,04E+08 | 1,22E+08 | 1,15E+08 | 1,36E+08 | 53,918 | 0 | 56,323 | 40 |
| 5,86E+07 | 5,66E+07 | 5,83E+07 | 6,27E+07 | 35,048 | 0 | 24,538 | 44 |
| 1,46E+08 | 1,42E+08 | 1,34E+08 | 1,74E+08 | 58,279 | 0 | 241,85 | 72 |
| 4,84E+06 | 2,72E+06 | 6,38E+06 | 3,77E+06 | 46,572 | 0,000152 | 4,0171 | 9 |
| 3,38E+07 | 3,56E+07 | 3,77E+07 | 3,29E+07 | 29,326 | 0 | 10,757 | 33 |
| 5,05E+07 | 6,64E+07 | 7,26E+07 | 7,32E+07 | 26,713 | 0 | 48,682 | 29 |
| 5,98E+06 | 6,55E+06 | 4,81E+06 | 4,73E+06 | 160,85 | 0,000149 | 3,4449 | 15 |
| 1,06E+07 | 1,33E+07 | 7,59E+06 | 1,29E+07 | 98,601 | 0,000158 | 5,068 | 17 |
| 1,55E+07 | 1,12E+07 | 1,03E+07 | 1,55E+07 | 48,967 | 0 | 13,751 | 16 |
| 6,87E+06 | 3,66E+06 | 5,96E+06 | 3,17E+06 | 127,01 | 0,000436 | 3,023 | 5 |
| 1,54E+07 | 1,23E+07 | 1,16E+07 | 1,09E+07 | 46,151 | 0 | 38,49 | 11 |
| 3,71E+07 | 5,00E+07 | 4,10E+07 | 4,36E+07 | 22,876 | 0 | 12,042 | 31 |
| 9,89E+08 | 1,21E+09 | 1,19E+09 | 1,37E+09 | 21,45 | 0 | 125,1 | 188 |
| 6,30E+06 | 2,35E+06 | 1,14E+07 | 1,96E+06 | 53,645 | 0 | 7,4819 | 6 |
| 2,74E+06 | 5,79E+07 | 6,65E+07 | 7,30E+07 | 11,365 | 0 | 6,4844 | 18 |
| 7,46E+06 | 1,07E+07 | 9,02E+06 | 9,90E+06 | 133,74 | 0 | 9,935 | 9 |
| 2,95E+07 | 3,14E+07 | 3,76E+07 | 3,51E+07 | 35,467 | 0 | 13,09 | 28 |
| 1,97E+09 | 2,00E+09 | 1,78E+09 | 1,73E+09 | 35,915 | 0 | 323,31 | 209 |
| 1,76E+06 | 2,23E+06 | 2,57E+06 | 3,68E+06 | 36,684 | 0,002918 | 2,0419 | 6 |
| 3,71E+07 | 4,06E+07 | 4,37E+07 | 4,56E+07 | 218,83 | 0 | 45,375 | 76 |
| 1,08E+10 | 1,02E+10 | 1,02E+10 | 9,53E+09 | 73,46 | 0 | 323,31 | 880 |

| | | | | | | | |
|---|---|---|---|---|---|---|---|
| 6,59E+07 | 4,63E+07 | 6,60E+07 | 7,12E+07 | 65,194 | 0 | 45,114 | 54 |
| 5,71E+08 | 5,62E+08 | 5,49E+08 | 5,55E+08 | 20,605 | 0 | 155,38 | 57 |
| 1,77E+07 | 4,11E+06 | 1,72E+07 | 1,68E+07 | 30,849 | 0 | 10,143 | 11 |
| 6,65E+06 | 3,39E+06 | 7,78E+06 | 3,20E+06 | 109,81 | 0 | 6,1778 | 6 |
| 2,88E+08 | 2,34E+08 | 2,51E+08 | 2,70E+08 | 36,883 | 0 | 229,25 | 81 |
| 8,44E+08 | 8,23E+08 | 8,00E+08 | 7,22E+08 | 70,402 | 0 | 323,31 | 306 |
| 5,29E+06 | 4,91E+06 | 6,14E+06 | 5,17E+06 | 32,485 | 0 | 25,092 | 8 |
| 2,02E+08 | 1,49E+08 | 1,67E+08 | 1,91E+08 | 23,294 | 0 | 272,65 | 40 |
| 5,77E+09 | 5,20E+09 | 4,76E+09 | 4,77E+09 | 58,004 | 0 | 323,31 | 388 |
| 4,47E+07 | 4,10E+07 | 4,08E+07 | 3,74E+07 | 93,703 | 0 | 18,146 | 47 |
| 5,99E+06 | 1,66E+06 | 4,03E+06 | 9,18E+06 | 214,89 | 0,000147 | 3,3062 | 4 |
| 3,48E+09 | 3,77E+09 | 3,63E+09 | 3,21E+09 | 111,18 | 0 | 323,31 | 516 |
| 1,29E+07 | 7,05E+06 | 1,34E+06 | 1,88E+07 | 36,296 | 0,000148 | 3,3621 | 3 |
| 1,73E+07 | 1,58E+07 | 1,97E+07 | 1,51E+07 | 37,851 | 0 | 15,342 | 23 |
| 6,86E+06 | 5,67E+06 | 5,44E+06 | 3,77E+06 | 49,429 | 0,000151 | 3,908 | 7 |
| 6,48E+07 | 6,10E+07 | 6,62E+07 | 6,24E+07 | 65,91 | 0 | 38,541 | 48 |
| 3,71E+09 | 3,91E+09 | 3,28E+09 | 2,81E+09 | 49,67 | 0 | 323,31 | 217 |
| 8,20E+07 | 9,29E+07 | 1,01E+08 | 9,41E+07 | 22,748 | 0 | 30,499 | 42 |
| 7,23E+08 | 6,90E+08 | 6,88E+08 | 6,94E+08 | 33,316 | 0 | 142,9 | 135 |
| 1,34E+08 | 1,43E+08 | 1,39E+08 | 1,45E+08 | 59,097 | 0 | 41,073 | 76 |
| 9,64E+07 | 9,37E+07 | 7,82E+07 | 1,00E+08 | 25,617 | 0 | 54,788 | 40 |
| 3,55E+08 | 4,65E+08 | 4,55E+08 | 4,33E+08 | 55,429 | 0 | 133,56 | 140 |
| 1,31E+09 | 1,51E+09 | 1,60E+09 | 1,57E+09 | 210,43 | 0 | 323,31 | 373 |
| 8,12E+07 | 8,32E+07 | 7,87E+07 | 8,75E+07 | 20,978 | 0 | 76,049 | 29 |
| 7,97E+07 | 9,28E+07 | 8,60E+07 | 6,76E+07 | 13,373 | 0 | 30,638 | 22 |
| 7,43E+06 | 4,06E+06 | 7,62E+06 | 1,01E+07 | 70,274 | 0,000155 | 4,5392 | 10 |
| 1,04E+09 | 1,09E+09 | 9,67E+08 | 9,12E+08 | 19,667 | 0 | 158,41 | 90 |
| 4,17E+06 | 8,92E+06 | 1,75E+07 | 7,10E+06 | 82,997 | 0 | 5,7802 | 8 |
| 5,17E+07 | 5,50E+07 | 4,65E+07 | 5,22E+07 | 20,139 | 0 | 84,242 | 27 |
| 5,03E+06 | 5,56E+06 | 9,09E+06 | 6,23E+06 | 27,562 | 0,000154 | 4,3664 | 7 |
| 7,13E+06 | 1,39E+07 | 1,30E+07 | 1,00E+07 | 45,145 | 0 | 7,8221 | 18 |
| 1,54E+08 | 1,71E+08 | 2,16E+08 | 1,91E+08 | 17,32 | 0 | 22,063 | 35 |
| 1,59E+08 | 2,20E+08 | 1,99E+08 | 1,87E+08 | 21,978 | 0 | 126,13 | 62 |
| 6,03E+08 | 5,06E+08 | 4,91E+08 | 5,45E+08 | 68,747 | 0 | 148,82 | 168 |
| 4,76E+07 | 3,45E+07 | 1,87E+07 | 2,04E+06 | 68,24 | 0,000291 | 3,0415 | 9 |
| 6,00E+06 | 4,95E+06 | 3,84E+06 | 6,96E+06 | 136,97 | 0,004364 | 1,7438 | 6 |
| 3,43E+06 | 9,50E+06 | 7,35E+06 | 5,68E+06 | 58,128 | 0 | 5,5583 | 9 |
| 5,64E+09 | 6,55E+09 | 6,40E+09 | 6,52E+09 | 35,076 | 0 | 323,31 | 350 |
| 4,13E+08 | 5,15E+08 | 4,58E+08 | 4,13E+08 | 38,096 | 0 | 311,9 | 173 |
| 2,05E+07 | 1,91E+07 | 8,18E+06 | 1,12E+07 | 135,88 | 0 | 8,7155 | 26 |
| 9,82E+06 | 1,35E+07 | 7,60E+06 | 8,51E+06 | 54,009 | 0 | 19,374 | 7 |
| 1,28E+07 | 1,32E+07 | 1,93E+07 | 7,20E+06 | 23,305 | 0 | 18,965 | 12 |
| 8,71E+08 | 1,06E+09 | 1,46E+09 | 2,16E+09 | 60 | 0,000429 | 2,6931 | 32 |
| 8,78E+07 | 1,09E+08 | 1,22E+08 | 1,36E+08 | 34,49 | 0 | 78,835 | 39 |
| 1,04E+08 | 1,78E+08 | 1,63E+08 | 1,97E+08 | 23,547 | 0 | 323,31 | 29 |
| 2,41E+07 | 3,12E+07 | 4,02E+07 | 4,73E+06 | 26,857 | 0 | 11,54 | 8 |
| 2,74E+07 | 4,39E+07 | 4,42E+07 | 4,62E+07 | 161,96 | 0 | 15,246 | 37 |
| 3,82E+07 | 3,70E+07 | 3,08E+07 | 3,10E+07 | 54,766 | 0 | 37,165 | 46 |
| 2,84E+06 | 2,61E+06 | 2,38E+06 | 3,24E+06 | 47,508 | 0,008197 | 1,499 | 7 |
| 4,23E+09 | 4,92E+09 | 5,08E+09 | 5,94E+09 | 24,305 | 0 | 236,12 | 324 |

| | | | | | | | |
|---|---|---|---|---|---|---|---|
| 4,00E+08 | 3,54E+08 | 3,70E+08 | 3,42E+08 | 31,825 | 0 | 323,31 | 122 |
| 2,08E+08 | 2,24E+08 | 1,91E+08 | 2,13E+08 | 44,402 | 0 | 323,31 | 82 |
| 3,77E+07 | 4,27E+07 | 4,26E+07 | 3,77E+07 | 39,432 | 0 | 13,431 | 27 |
| 1,28E+06 | 1,37E+06 | 1,20E+06 | 1,84E+06 | 113,5 | 0,003703 | 1,8495 | 8 |
| 2,10E+07 | 2,53E+07 | 2,12E+07 | 2,64E+07 | 224,92 | 0 | 9,8023 | 19 |
| 1,45E+08 | 1,46E+08 | 1,42E+08 | 1,36E+08 | 78,429 | 0 | 76,745 | 95 |
| 1,96E+06 | 2,47E+06 | 2,69E+06 | 2,47E+06 | 130,65 | 0,000156 | 4,6335 | 7 |
| 3,77E+08 | 4,30E+08 | 3,80E+08 | 4,21E+08 | 34,481 | 0 | 170,45 | 144 |
| 7,86E+06 | 3,49E+06 | 1,40E+06 | 1,22E+07 | 55,838 | 0,003572 | 1,8892 | 4 |
| 4,02E+08 | 4,03E+08 | 4,09E+08 | 4,12E+08 | 135,08 | 0 | 226,82 | 209 |
| 2,96E+07 | 3,23E+07 | 3,40E+07 | 3,61E+07 | 21,86 | 0 | 32,231 | 20 |
| 2,08E+07 | 2,10E+07 | 1,21E+07 | 1,67E+07 | 55,354 | 0 | 25,272 | 21 |
| 6,18E+08 | 5,00E+08 | 4,42E+08 | 4,07E+08 | 29,343 | 0 | 242,91 | 112 |
| 4,16E+07 | 3,16E+07 | 2,71E+07 | 3,42E+07 | 113,95 | 0 | 17,759 | 40 |
| 7,46E+07 | 6,44E+07 | 7,50E+07 | 7,12E+07 | 31,654 | 0 | 47,238 | 53 |
| 5,14E+08 | 5,00E+08 | 5,01E+08 | 4,91E+08 | 215,54 | 0 | 323,31 | 307 |
| 2,24E+07 | 5,09E+07 | 2,98E+07 | 2,81E+07 | 22,282 | 0 | 29,93 | 28 |
| 1,03E+10 | 9,15E+09 | 9,24E+09 | 9,09E+09 | 277,82 | 0 | 323,31 | 1344 |
| 3,18E+07 | 1,65E+07 | 1,85E+07 | 1,25E+07 | 32,79 | 0 | 23,998 | 26 |
| 1,80E+08 | 2,04E+08 | 1,79E+08 | 1,51E+08 | 16,859 | 0 | 18,734 | 85 |
| 2,20E+07 | 2,77E+06 | 1,46E+07 | 1,55E+07 | 11,213 | 0 | 81,669 | 7 |
| 5,39E+06 | 4,64E+06 | 4,51E+06 | 4,30E+06 | 13,3 | 0,004757 | 1,6944 | 3 |
| 3,25E+07 | 3,71E+07 | 4,00E+07 | 4,45E+07 | 12,017 | 0,000147 | 3,2731 | 16 |
| 3,93E+07 | 3,41E+07 | 3,52E+07 | 3,64E+07 | 46,577 | 0 | 8,5294 | 24 |
| 2,70E+06 | 2,68E+06 | 1,76E+06 | 3,20E+06 | 7,8359 | 0,00015 | 3,6339 | 2 |
| 1,15E+08 | 1,31E+08 | 1,17E+08 | 1,15E+08 | 46,121 | 0 | 111,88 | 50 |
| 3,98E+06 | 7,22E+06 | 5,64E+06 | 4,39E+06 | 91,264 | 0 | 6,735 | 12 |
| 2,14E+09 | 1,88E+09 | 1,76E+09 | 1,58E+09 | 51,564 | 0 | 323,31 | 246 |
| 1,14E+09 | 1,32E+09 | 1,21E+09 | 1,31E+09 | 48,234 | 0 | 323,31 | 152 |
| 4,57E+08 | 4,09E+08 | 4,36E+08 | 4,68E+08 | 65,146 | 0 | 148,03 | 166 |
| 7,06E+07 | 7,50E+07 | 7,70E+07 | 8,15E+07 | 34,67 | 0 | 56,281 | 32 |
| 3,44E+08 | 3,12E+08 | 2,98E+08 | 3,40E+08 | 31,395 | 0 | 74,294 | 69 |
| 3,94E+07 | 4,53E+07 | 3,79E+07 | 3,83E+07 | 30,084 | 0 | 11,105 | 36 |
| 1,28E+08 | 1,16E+08 | 1,37E+08 | 1,21E+08 | 33,758 | 0 | 246,45 | 75 |
| 6,99E+07 | 9,50E+07 | 5,98E+07 | 4,91E+07 | 29,395 | 0 | 106,78 | 37 |
| 3,68E+07 | 2,50E+07 | 3,03E+07 | 3,02E+07 | 31,617 | 0 | 7,847 | 16 |
| 7,07E+07 | 8,28E+07 | 6,68E+07 | 7,66E+07 | 58,567 | 0 | 10,369 | 23 |
| 1,15E+07 | 2,30E+07 | 1,05E+07 | 1,61E+07 | 63,023 | 0 | 5,3251 | 11 |
| 1,89E+07 | 2,59E+07 | 2,31E+07 | 2,47E+07 | 8,7608 | 0 | 5,5031 | 22 |
| 3,73E+09 | 1,83E+09 | 2,08E+09 | 1,75E+09 | 44,547 | 0 | 323,31 | 183 |
| 2,37E+08 | 2,71E+08 | 2,90E+08 | 2,45E+08 | 11,528 | 0 | 31,008 | 50 |
| 4,62E+07 | 5,30E+07 | 4,38E+07 | 5,79E+07 | 34,084 | 0 | 31,065 | 42 |
| 1,04E+07 | 5,29E+06 | 4,59E+06 | 1,02E+07 | 13,725 | 0,000154 | 4,3593 | 8 |
| 1,72E+08 | 1,83E+08 | 1,73E+08 | 1,84E+08 | 28,37 | 0 | 173,06 | 54 |
| 2,07E+07 | 1,69E+07 | 1,54E+07 | 1,57E+07 | 118,56 | 0 | 8,8104 | 18 |
| 1,02E+07 | 7,95E+06 | 7,43E+06 | 4,72E+06 | 83,1 | 0,003441 | 1,908 | 11 |
| 7,95E+06 | 1,33E+07 | 1,29E+07 | 1,02E+07 | 140,98 | 0,000157 | 4,9301 | 19 |
| 5,73E+07 | 5,45E+07 | 5,23E+07 | 2,18E+07 | 85,533 | 0 | 7,5008 | 23 |
| 8,46E+08 | 5,38E+08 | 7,73E+08 | 5,66E+08 | 16,687 | 0 | 170,31 | 29 |
| 2,22E+07 | 2,84E+07 | 2,35E+07 | 2,74E+07 | 56,868 | 0 | 6,6494 | 20 |

| | | | | | | | |
|---|---|---|---|---|---|---|---|
| 9,00E+06 | 3,72E+06 | 6,46E+06 | 7,00E+06 | 41,791 | 0 | 75,038 | 17 |
| 4,25E+06 | 1,00E+07 | 6,70E+06 | 1,27E+07 | 57,093 | 0 | 13,864 | 9 |
| 2,20E+07 | 2,21E+07 | 2,07E+07 | 1,68E+07 | 17,73 | 0 | 27,254 | 11 |
| 1,99E+08 | 2,27E+08 | 1,98E+08 | 2,09E+08 | 42,309 | 0 | 323,31 | 105 |
| 3,98E+08 | 4,10E+08 | 4,00E+08 | 4,31E+08 | 93,841 | 0 | 323,31 | 160 |
| 1,82E+08 | 1,15E+08 | 1,25E+08 | 1,02E+08 | 49,558 | 0 | 239,67 | 50 |
| 8,28E+07 | 3,33E+06 | 1,25E+08 | 9,23E+07 | 116,97 | 0 | 18,21 | 28 |
| 5,48E+08 | 5,25E+08 | 5,55E+08 | 5,90E+08 | 71,726 | 0 | 323,31 | 170 |
| 4,09E+06 | 4,06E+06 | 7,55E+06 | 3,52E+06 | 164,62 | 0 | 5,5842 | 8 |
| 2,34E+06 | 5,00E+07 | 5,93E+07 | 3,91E+07 | 366,22 | 0,003439 | 1,9061 | 10 |
| 1,09E+07 | 1,02E+07 | 7,55E+06 | 1,19E+07 | 48,189 | 0 | 11,192 | 15 |
| 8,76E+07 | 9,56E+07 | 1,08E+08 | 9,81E+07 | 87,175 | 0 | 121,39 | 82 |
| 2,59E+06 | 1,78E+06 | 1,21E+06 | 2,63E+06 | 205,73 | 0,007676 | 1,5271 | 6 |
| 1,48E+07 | 3,34E+07 | 3,22E+07 | 2,66E+07 | 59,008 | 0 | 23,053 | 21 |
| 3,48E+06 | 5,51E+06 | 5,23E+06 | 2,03E+06 | 80,698 | 0 | 7,6027 | 6 |
| 1,09E+10 | 1,08E+10 | 1,17E+10 | 1,07E+10 | 60,955 | 0 | 323,31 | 1142 |
| 4,20E+06 | 5,82E+06 | 2,02E+06 | 8,20E+06 | 41,771 | 0,001272 | 2,4322 | 2 |
| 4,77E+07 | 3,58E+07 | 4,35E+07 | 5,32E+07 | 81,215 | 0 | 20,161 | 45 |
| 2,88E+07 | 1,48E+07 | 1,56E+07 | 1,33E+07 | 38,607 | 0 | 20,652 | 19 |
| 1,72E+08 | 1,75E+08 | 1,58E+08 | 1,83E+08 | 37,527 | 0 | 44,122 | 72 |
| 1,34E+08 | 1,34E+08 | 1,53E+08 | 1,30E+08 | 76,006 | 0 | 233,43 | 92 |
| 1,78E+07 | 1,68E+07 | 1,83E+07 | 2,09E+07 | 54,297 | 0 | 11,134 | 23 |
| 6,51E+07 | 4,74E+07 | 5,97E+07 | 5,46E+07 | 43,353 | 0 | 193,37 | 39 |
| 4,69E+08 | 4,90E+08 | 5,24E+08 | 4,92E+08 | 76,908 | 0 | 206,89 | 186 |
| 5,56E+06 | 3,83E+06 | 5,15E+06 | 6,62E+06 | 53,193 | 0,00127 | 2,3875 | 3 |
| 6,48E+07 | 7,12E+07 | 8,23E+07 | 7,69E+07 | 64,622 | 0 | 22,997 | 49 |
| 1,31E+08 | 1,22E+08 | 1,32E+08 | 1,26E+08 | 76,784 | 0 | 323,31 | 54 |
| 9,90E+06 | 2,08E+06 | 1,27E+07 | 8,66E+06 | 37,262 | 0,001409 | 2,3569 | 11 |
| 3,77E+06 | 7,51E+06 | 5,57E+06 | 7,87E+06 | 29,978 | 0,008207 | 1,5124 | 6 |
| 3,01E+07 | 2,87E+07 | 1,86E+07 | 3,96E+07 | 57,317 | 0 | 51,286 | 28 |
| 3,57E+08 | 2,58E+08 | 2,77E+08 | 2,49E+08 | 33,429 | 0 | 323,31 | 88 |
| 1,30E+09 | 1,19E+09 | 1,15E+09 | 1,12E+09 | 96,406 | 0 | 323,31 | 395 |
| 1,70E+08 | 1,71E+08 | 1,69E+08 | 1,86E+08 | 93,09 | 0 | 88,311 | 117 |
| 1,78E+07 | 8,22E+06 | 2,26E+07 | 2,08E+07 | 14,155 | 0 | 5,9187 | 8 |
| 1,18E+08 | 7,23E+06 | 1,28E+07 | 9,89E+06 | 16,607 | 0 | 12,166 | 23 |
| 2,52E+06 | 3,10E+06 | 2,99E+06 | 2,71E+06 | 70,955 | 0,005422 | 1,6489 | 5 |
| 4,23E+08 | 2,17E+08 | 1,75E+08 | 1,93E+08 | 42,06 | 0 | 239,41 | 79 |
| 2,09E+06 | 5,76E+06 | 6,50E+06 | 5,43E+06 | 79,284 | 0 | 6,8504 | 11 |
| 8,73E+07 | 7,83E+07 | 9,79E+07 | 9,85E+07 | 20,655 | 0 | 42,658 | 31 |
| 7,17E+06 | 1,02E+07 | 3,65E+06 | 1,37E+07 | 24,913 | 0,00043 | 2,7588 | 8 |
| 3,09E+07 | 3,11E+07 | 4,45E+07 | 4,74E+07 | 20,911 | 0 | 20,751 | 22 |
| 9,20E+06 | 7,28E+06 | 8,33E+06 | 6,98E+06 | 68,234 | 0,000157 | 4,9657 | 14 |
| 1,08E+09 | 7,75E+08 | 8,75E+08 | 8,96E+08 | 109,55 | 0 | 323,31 | 364 |
| 4,15E+08 | 4,61E+08 | 4,71E+08 | 4,39E+08 | 49,485 | 0 | 205,5 | 104 |
| 2,24E+08 | 2,20E+08 | 2,26E+08 | 2,07E+08 | 30,244 | 0 | 110,3 | 60 |
| 2,03E+07 | 1,89E+07 | 2,93E+07 | 3,29E+07 | 56,789 | 0 | 30,802 | 17 |
| 3,04E+07 | 2,36E+07 | 2,14E+07 | 1,78E+07 | 21,918 | 0 | 11,943 | 27 |
| 2,48E+07 | 3,10E+07 | 4,02E+07 | 3,67E+07 | 9,0788 | 0,00872 | 1,4716 | 17 |
| 5,37E+06 | 2,14E+06 | 2,09E+06 | 1,64E+06 | 69,24 | 0,007809 | 1,5238 | 6 |
| 2,90E+06 | 3,39E+06 | 3,01E+06 | 2,64E+06 | 67,031 | 0,000158 | 5,0113 | 10 |

| | | | | | | | |
|---|---|---|---|---|---|---|---|
| 2,47E+07 | 2,18E+07 | 2,13E+07 | 1,71E+07 | 81,265 | 0 | 18,685 | 28 |
| 1,59E+06 | 2,72E+06 | 1,84E+06 | 1,45E+06 | 38,752 | 0,000147 | 3,2047 | 3 |
| 1,89E+08 | 1,74E+06 | 1,74E+08 | 1,86E+08 | 116,23 | 0 | 5,6143 | 6 |
| 1,56E+08 | 1,55E+08 | 1,65E+08 | 1,50E+08 | 47,36 | 0 | 221,26 | 93 |
| 5,11E+08 | 4,35E+08 | 4,46E+08 | 4,75E+08 | 64,802 | 0 | 300,04 | 110 |
| 1,04E+08 | 1,16E+08 | 1,19E+08 | 1,11E+08 | 58,118 | 0 | 46,203 | 56 |
| 7,02E+06 | 7,45E+06 | 9,06E+06 | 6,86E+06 | 113,03 | 0 | 15,273 | 15 |
| 1,44E+07 | 1,19E+07 | 1,20E+07 | 1,54E+07 | 103 | 0 | 8,0755 | 15 |
| 8,46E+08 | 7,75E+08 | 7,70E+08 | 6,68E+08 | 50,09 | 0 | 323,31 | 141 |
| 3,62E+08 | 4,05E+08 | 3,67E+08 | 3,85E+08 | 26,476 | 0 | 61,967 | 92 |
| 3,70E+07 | 4,57E+07 | 3,83E+07 | 4,20E+07 | 59,519 | 0 | 65,428 | 36 |
| 1,40E+08 | 1,83E+08 | 1,68E+08 | 1,23E+08 | 22,382 | 0 | 211,25 | 87 |
| 1,46E+08 | 1,56E+08 | 1,41E+08 | 1,41E+08 | 44,669 | 0 | 45,402 | 95 |
| 1,55E+07 | 1,05E+07 | 8,02E+06 | 1,40E+07 | 107,22 | 0 | 15,196 | 18 |
| 2,35E+08 | 2,21E+08 | 2,52E+08 | 2,58E+08 | 49,939 | 0 | 95,817 | 129 |
| 7,61E+06 | 1,24E+07 | 9,50E+06 | 1,21E+07 | 60,303 | 0,002915 | 2,0296 | 8 |
| 4,37E+07 | 4,97E+07 | 4,41E+07 | 4,15E+07 | 9,2983 | 0 | 66,614 | 16 |
| 6,35E+06 | 6,51E+06 | 6,57E+06 | 1,60E+06 | 61,852 | 0 | 9,1911 | 14 |
| 7,23E+06 | 4,55E+06 | 5,28E+06 | 6,69E+06 | 65,722 | 0,000156 | 4,662 | 6 |
| 1,04E+08 | 1,16E+08 | 1,10E+08 | 1,00E+08 | 72,871 | 0 | 41,158 | 78 |
| 6,50E+07 | 4,36E+07 | 4,75E+07 | 5,47E+07 | 223,93 | 0 | 8,4029 | 29 |
| 9,78E+08 | 9,98E+08 | 1,11E+09 | 1,26E+09 | 32,68 | 0 | 323,31 | 171 |
| 1,40E+08 | 1,49E+08 | 1,27E+08 | 1,44E+08 | 50,966 | 0 | 68,54 | 90 |
| 9,41E+06 | 1,35E+07 | 1,28E+07 | 1,18E+07 | 163,58 | 0 | 5,5502 | 23 |
| 3,25E+07 | 3,43E+07 | 3,36E+07 | 3,43E+07 | 40,659 | 0 | 16,03 | 28 |
| 6,39E+08 | 7,91E+08 | 7,33E+08 | 7,84E+08 | 15,86 | 0 | 135,33 | 254 |
| 5,89E+07 | 6,03E+07 | 5,86E+07 | 7,70E+07 | 43,092 | 0,000149 | 3,5122 | 9 |
| 1,37E+08 | 1,34E+08 | 1,25E+08 | 1,12E+08 | 40,57 | 0 | 242,92 | 35 |
| 2,52E+07 | 2,41E+07 | 2,79E+07 | 1,98E+07 | 46,224 | 0 | 7,8063 | 21 |
| 6,08E+09 | 7,17E+09 | 6,67E+09 | 7,16E+09 | 32,838 | 0 | 323,31 | 410 |
| 7,11E+07 | 6,05E+07 | 6,02E+07 | 6,39E+07 | 110,22 | 0 | 41,465 | 59 |
| 7,04E+07 | 6,45E+07 | 7,36E+07 | 6,74E+07 | 83,446 | 0 | 41,837 | 62 |
| 4,99E+07 | 6,59E+07 | 6,01E+07 | 2,82E+07 | 44,229 | 0 | 10,242 | 24 |
| 5,56E+06 | 5,20E+06 | 5,62E+06 | 6,15E+06 | 48,3 | 0 | 9,8358 | 15 |
| 5,11E+06 | 5,64E+06 | 2,04E+06 | 7,07E+06 | 49,912 | 0,00344 | 1,9077 | 6 |
| 1,12E+08 | 1,12E+08 | 1,26E+08 | 1,19E+08 | 36,566 | 0 | 196,36 | 52 |
| 7,40E+06 | 1,01E+07 | 8,71E+06 | 7,76E+06 | 83,773 | 0 | 26,33 | 23 |
| 5,98E+07 | 3,85E+07 | 3,20E+07 | 3,89E+07 | 27,974 | 0 | 82,621 | 25 |
| 2,90E+08 | 2,93E+08 | 2,77E+08 | 2,89E+08 | 48,34 | 0 | 139,64 | 128 |
| 7,02E+08 | 7,02E+08 | 7,18E+08 | 6,95E+08 | 24,313 | 0 | 253,6 | 118 |
| 6,97E+06 | 8,04E+06 | 7,95E+06 | 7,20E+06 | 49,316 | 0,001269 | 2,3848 | 8 |
| 2,62E+07 | 4,06E+07 | 4,05E+07 | 3,29E+07 | 22,77 | 0 | 8,5765 | 21 |
| 9,79E+06 | 1,01E+07 | 9,90E+06 | 8,80E+06 | 130,32 | 0 | 11,971 | 13 |
| 5,11E+06 | 4,09E+06 | 5,51E+06 | 2,28E+06 | 44,576 | 0,005423 | 1,6501 | 8 |
| 9,87E+06 | 4,86E+06 | 9,30E+06 | 8,51E+06 | 19,25 | 0,000147 | 3,2779 | 7 |
| 4,19E+08 | 2,51E+08 | 3,47E+08 | 2,75E+08 | 10,435 | 0 | 177,15 | 62 |
| 4,30E+10 | 3,93E+10 | 3,78E+10 | 3,94E+10 | 11,367 | 0 | 318,64 | 2366 |
| 1,73E+08 | 1,79E+08 | 1,79E+08 | 1,99E+08 | 63,961 | 0 | 206,48 | 60 |
| 1,72E+06 | 1,63E+06 | 1,10E+06 | 4,65E+06 | 73,871 | 0,006079 | 1,5984 | 4 |
| 1,47E+07 | 1,68E+07 | 1,44E+07 | 1,58E+07 | 61,057 | 0 | 11,443 | 28 |

| | | | | | | | |
|---|---|---|---|---|---|---|---|
| 7,30E+06 | 9,30E+05 | 2,58E+06 | 2,40E+06 | 47,514 | 0 | 38,358 | 6 |
| 3,26E+06 | 7,84E+06 | 8,00E+06 | 6,66E+06 | 17,537 | 0,000148 | 3,3095 | 11 |
| 5,08E+06 | 1,82E+06 | 6,03E+06 | 6,14E+06 | 29,37 | 0,005416 | 1,639 | 5 |
| 9,40E+09 | 9,08E+09 | 1,05E+10 | 1,02E+10 | 39,652 | 0 | 323,31 | 899 |
| 1,86E+07 | 1,86E+07 | 2,52E+07 | 2,31E+07 | 14,251 | 0 | 7,8034 | 22 |
| 2,96E+08 | 2,89E+08 | 2,75E+08 | 2,79E+08 | 141,46 | 0 | 323,31 | 170 |
| 1,74E+06 | 5,96E+05 | 1,27E+06 | 2,01E+06 | 42,913 | 0,003184 | 1,9879 | 5 |
| 6,65E+08 | 6,85E+08 | 6,87E+08 | 6,76E+08 | 51,596 | 0 | 248,18 | 198 |
| 2,22E+06 | 1,15E+07 | 1,62E+07 | 2,86E+06 | 26,433 | 0,001271 | 2,4125 | 3 |
| 5,97E+06 | 6,62E+06 | 1,72E+06 | 4,00E+06 | 11,48 | 0 | 323,31 | 11 |
| 4,57E+07 | 1,90E+07 | 2,17E+07 | 2,50E+07 | 27,964 | 0 | 8,0202 | 27 |
| 1,00E+08 | 8,81E+07 | 8,53E+07 | 9,65E+07 | 58,006 | 0 | 67,371 | 78 |
| 5,51E+06 | 4,41E+06 | 2,15E+06 | 1,79E+06 | 58,73 | 0,000159 | 5,3064 | 2 |
| 6,87E+07 | 6,69E+07 | 4,62E+07 | 2,74E+07 | 96,739 | 0 | 22,127 | 46 |
| 2,49E+08 | 9,45E+08 | 3,26E+08 | 1,75E+08 | 22,975 | 0 | 9,4427 | 60 |
| 7,15E+06 | 7,06E+06 | 7,84E+06 | 8,83E+06 | 18,305 | 0,00741 | 1,5378 | 5 |
| 1,38E+08 | 1,49E+08 | 1,52E+08 | 1,48E+08 | 81,626 | 0 | 323,31 | 80 |
| 2,57E+06 | 3,02E+06 | 1,73E+06 | 2,45E+06 | 221,57 | 0,000154 | 4,2985 | 10 |
| 4,05E+06 | 5,14E+06 | 1,84E+07 | 2,76E+07 | 49,679 | 0 | 6,36 | 19 |
| 2,44E+05 | 9,97E+05 | 8,61E+06 | 1,73E+06 | 19,837 | 0,000855 | 2,6005 | 6 |
| 4,17E+06 | 1,80E+07 | 1,92E+07 | 4,15E+07 | 9,2864 | 0 | 14,618 | 14 |
| 6,41E+06 | 2,95E+06 | 2,05E+06 | 3,62E+06 | 56,413 | 0 | 8,035 | 8 |
| 4,65E+07 | 4,32E+07 | 4,55E+07 | 3,91E+07 | 37,829 | 0 | 11,06 | 32 |
| 1,92E+08 | 2,34E+08 | 2,35E+08 | 2,78E+08 | 10,458 | 0 | 117,72 | 54 |
| 1,16E+08 | 1,02E+08 | 1,14E+08 | 1,24E+08 | 203,91 | 0 | 65,178 | 121 |
| 6,12E+08 | 6,26E+08 | 5,92E+08 | 5,42E+08 | 59,125 | 0 | 153,9 | 176 |
| 1,83E+06 | 4,09E+06 | 1,69E+06 | 1,43E+06 | 60,291 | 0,005411 | 1,6235 | 2 |
| 7,34E+06 | 6,97E+06 | 8,10E+06 | 7,52E+06 | 62,529 | 0 | 10,246 | 13 |
| 2,70E+09 | 1,86E+09 | 2,24E+09 | 1,97E+09 | 18,521 | 0 | 217,56 | 291 |
| 2,93E+07 | 2,77E+07 | 3,00E+07 | 2,62E+07 | 48,396 | 0 | 15,309 | 21 |
| 3,81E+07 | 4,50E+07 | 4,26E+07 | 4,19E+07 | 83,99 | 0 | 34,503 | 57 |
| 4,78E+09 | 4,58E+09 | 4,80E+09 | 4,63E+09 | 106,91 | 0 | 323,31 | 554 |
| 1,03E+08 | 1,09E+08 | 1,08E+08 | 1,08E+08 | 186,06 | 0 | 53,092 | 118 |
| 9,46E+06 | 1,11E+07 | 9,36E+06 | 1,35E+07 | 25,985 | 0,000431 | 2,8026 | 10 |
| 8,25E+07 | 6,61E+07 | 7,87E+07 | 5,79E+07 | 16,369 | 0 | 117,62 | 13 |
| 4,39E+09 | 4,66E+09 | 5,21E+09 | 4,92E+09 | 51,736 | 0 | 322,53 | 357 |
| 5,05E+06 | 8,37E+05 | 2,26E+06 | 1,97E+06 | 41,89 | 0,000152 | 4,0498 | 4 |
| 5,42E+06 | 9,42E+06 | 9,14E+06 | 6,84E+06 | 52,622 | 0,000429 | 2,6872 | 7 |
| 2,63E+07 | 2,69E+07 | 2,49E+07 | 2,55E+07 | 54,208 | 0 | 11,492 | 32 |
| 6,59E+06 | 4,69E+06 | 8,52E+06 | 4,28E+06 | 35,891 | 0,00015 | 3,5995 | 8 |
| 3,89E+08 | 3,72E+08 | 4,02E+08 | 4,31E+08 | 21,897 | 0 | 181,11 | 90 |
| 2,24E+07 | 7,09E+06 | 9,99E+06 | 8,36E+06 | 14,149 | 0,000157 | 4,8281 | 12 |
| 5,38E+06 | 8,30E+06 | 1,30E+07 | 1,23E+07 | 81,951 | 0,000153 | 4,2286 | 16 |
| 7,35E+07 | 8,26E+07 | 8,34E+07 | 9,44E+07 | 103,74 | 0 | 160,87 | 88 |
| 3,21E+09 | 3,00E+09 | 3,24E+09 | 2,95E+09 | 16,085 | 0 | 47,725 | 127 |
| 4,46E+09 | 4,49E+09 | 4,23E+09 | 4,47E+09 | 24,205 | 0 | 323,31 | 215 |
| 1,36E+07 | 1,47E+07 | 1,54E+07 | 1,51E+07 | 133,99 | 0 | 6,5373 | 21 |
| 2,13E+08 | 2,23E+08 | 2,17E+08 | 2,10E+08 | 48,699 | 0 | 323,31 | 85 |
| 2,10E+09 | 2,15E+09 | 1,99E+09 | 2,14E+09 | 36,169 | 0 | 323,31 | 168 |
| 1,09E+08 | 1,29E+08 | 1,35E+08 | 1,34E+08 | 20,529 | 0 | 89,552 | 47 |

| | | | | | | | |
|---|---|---|---|---|---|---|---|
| 4,29E+07 | 4,05E+07 | 4,92E+07 | 4,68E+07 | 34,729 | 0 | 8,8149 | 29 |
| 1,50E+07 | 1,62E+07 | 7,25E+06 | 1,35E+07 | 83,516 | 0 | 8,3521 | 14 |
| 4,68E+08 | 2,86E+08 | 2,00E+08 | 1,58E+08 | 22,031 | 0 | 323,31 | 68 |
| 1,47E+07 | 1,26E+07 | 1,18E+07 | 8,20E+06 | 23,112 | 0,000433 | 2,8623 | 9 |
| 1,48E+07 | 5,12E+06 | 1,13E+07 | 9,61E+06 | 30,501 | 0 | 7,7816 | 16 |
| 2,45E+07 | 1,65E+07 | 1,26E+07 | 1,66E+07 | 172,62 | 0 | 9,0807 | 27 |
| 1,71E+07 | 1,85E+07 | 2,40E+07 | 3,06E+07 | 12,441 | 0 | 5,8807 | 29 |
| 2,15E+08 | 2,13E+08 | 2,13E+08 | 1,63E+08 | 27,987 | 0 | 55,804 | 62 |
| 3,50E+08 | 4,13E+08 | 3,87E+08 | 3,94E+08 | 48,063 | 0 | 128,55 | 139 |
| 4,37E+06 | 5,82E+06 | 7,93E+06 | 7,17E+06 | 133,34 | 0,000431 | 2,8014 | 9 |
| 3,98E+08 | 2,62E+08 | 1,92E+08 | 1,94E+08 | 20,524 | 0 | 50,335 | 67 |
| 7,06E+08 | 5,94E+08 | 6,58E+08 | 6,13E+08 | 121,04 | 0 | 323,31 | 272 |
| 6,52E+06 | 7,30E+06 | 3,86E+06 | 2,07E+06 | 40,447 | 0,000152 | 3,9427 | 11 |
| 2,18E+08 | 1,75E+08 | 1,58E+08 | 1,67E+08 | 23,399 | 0 | 20,204 | 55 |
| 1,81E+07 | 2,19E+07 | 1,65E+07 | 1,94E+07 | 21,092 | 0 | 13,062 | 20 |
| 2,88E+07 | 5,46E+07 | 5,02E+07 | 4,56E+07 | 29,454 | 0 | 38,23 | 36 |
| 2,43E+08 | 1,59E+08 | 1,82E+08 | 1,98E+08 | 45,745 | 0 | 302,41 | 98 |
| 1,64E+08 | 1,76E+08 | 1,62E+08 | 1,50E+08 | 42,761 | 0 | 79,962 | 56 |
| 8,31E+06 | 1,14E+07 | 4,36E+06 | 3,93E+06 | 42,241 | 0,004627 | 1,713 | 8 |
| 5,22E+06 | 5,23E+06 | 5,55E+06 | 3,94E+06 | 25,552 | 0 | 7,2133 | 9 |
| 9,49E+07 | 1,01E+08 | 9,74E+07 | 1,05E+08 | 73,797 | 0 | 19,545 | 45 |
| 4,02E+07 | 4,87E+07 | 3,99E+07 | 4,04E+07 | 37,657 | 0 | 17,574 | 48 |
| 5,11E+08 | 5,66E+08 | 5,27E+08 | 5,24E+08 | 22,406 | 0 | 323,31 | 101 |
| 2,20E+08 | 1,88E+08 | 1,91E+08 | 1,76E+08 | 57,454 | 0 | 103,32 | 94 |
| 1,97E+09 | 1,73E+09 | 1,66E+09 | 2,00E+09 | 35,73 | 0 | 323,31 | 182 |
| 6,45E+07 | 1,05E+08 | 1,06E+08 | 1,31E+08 | 20,116 | 0 | 132,33 | 29 |
| 4,43E+08 | 4,43E+08 | 4,56E+08 | 4,53E+08 | 91,736 | 0 | 323,31 | 215 |
| 4,80E+07 | 4,70E+07 | 5,62E+07 | 5,61E+07 | 202,24 | 0 | 31,855 | 67 |
| 3,79E+07 | 3,85E+07 | 3,55E+07 | 3,48E+07 | 38,842 | 0 | 12,571 | 29 |
| 9,86E+08 | 1,07E+09 | 1,08E+09 | 1,12E+09 | 44,76 | 0 | 323,31 | 221 |
| 1,33E+08 | 1,48E+08 | 1,42E+08 | 1,54E+08 | 82,779 | 0 | 49,41 | 94 |
| 3,46E+07 | 4,42E+07 | 3,83E+07 | 3,38E+07 | 39,694 | 0 | 9,4969 | 34 |
| 6,31E+06 | 3,10E+06 | 1,08E+07 | 1,02E+07 | 94,316 | 0,009748 | 1,4103 | 2 |
| 4,36E+06 | 2,21E+06 | 1,12E+07 | 4,58E+06 | 173,27 | 0,00463 | 1,7262 | 7 |
| 3,01E+08 | 3,36E+08 | 3,05E+08 | 3,06E+08 | 80,564 | 0 | 116,59 | 144 |
| 4,49E+06 | 5,05E+06 | 4,15E+06 | 4,14E+06 | 22,628 | 0 | 5,7112 | 8 |
| 5,21E+08 | 5,36E+08 | 5,46E+08 | 5,23E+08 | 66,943 | 0 | 273,21 | 148 |
| 6,37E+07 | 9,36E+07 | 9,91E+07 | 1,04E+08 | 33,395 | 0,000159 | 5,1788 | 24 |
| 1,26E+07 | 6,60E+06 | 6,80E+06 | 5,02E+06 | 33,735 | 0,000151 | 3,7811 | 4 |
| 1,76E+09 | 1,16E+09 | 8,17E+08 | 1,14E+09 | 14,463 | 0 | 116,47 | 117 |
| 2,63E+08 | 2,86E+08 | 2,89E+08 | 2,78E+08 | 21,777 | 0 | 57,144 | 106 |
| 1,32E+07 | 1,70E+07 | 1,86E+07 | 2,42E+07 | 85,339 | 0 | 8,8644 | 20 |
| 2,35E+06 | 3,33E+06 | 3,46E+06 | 4,39E+06 | 51,346 | 0,000435 | 2,9366 | 5 |
| 6,28E+06 | 6,80E+06 | 8,26E+06 | 5,69E+06 | 24,986 | 0 | 8,4293 | 11 |
| 6,76E+07 | 5,76E+07 | 6,52E+07 | 6,02E+07 | 18,376 | 0 | 21,212 | 47 |
| 1,80E+06 | 1,82E+06 | 1,02E+07 | 2,33E+06 | 63,235 | 0,00043 | 2,739 | 6 |
| 8,98E+06 | 7,43E+06 | 1,00E+07 | 2,40E+06 | 50,104 | 0 | 7,3145 | 8 |
| 1,53E+07 | 1,18E+07 | 1,29E+07 | 1,64E+07 | 34,516 | 0 | 7,7832 | 12 |
| 5,38E+06 | 4,60E+06 | 3,33E+06 | 5,35E+06 | 39,246 | 0,000151 | 3,8499 | 11 |
| 1,29E+07 | 6,48E+06 | 4,73E+06 | 2,07E+06 | 37,489 | 0,000147 | 3,1493 | 2 |

| | | | | | | | |
|---|---|---|---|---|---|---|---|
| 1,71E+07 | 1,74E+07 | 1,61E+07 | 1,65E+07 | 43,561 | 0 | 6,3406 | 24 |
| 2,71E+07 | 5,36E+06 | 2,50E+06 | 1,73E+07 | 21,18 | 0 | 37,494 | 4 |
| 4,91E+07 | 5,38E+07 | 3,95E+07 | 4,86E+07 | 67,5 | 0 | 39,939 | 39 |
| 4,08E+08 | 3,59E+08 | 3,59E+08 | 3,70E+08 | 26,791 | 0 | 115,9 | 97 |
| 2,87E+07 | 2,47E+07 | 2,93E+07 | 2,77E+07 | 52,364 | 0 | 7,2029 | 34 |
| 9,78E+06 | 6,36E+06 | 1,08E+07 | 7,13E+06 | 132,8 | 0,004499 | 1,7372 | 6 |
| 8,12E+07 | 4,37E+07 | 4,30E+07 | 4,02E+07 | 15,815 | 0 | 11,102 | 12 |
| 2,82E+07 | 4,83E+07 | 4,44E+07 | 3,91E+07 | 21,102 | 0 | 10,047 | 14 |
| 7,75E+07 | 7,96E+07 | 7,93E+07 | 8,24E+07 | 215,68 | 0 | 34,274 | 86 |
| 1,13E+07 | 1,30E+07 | 1,21E+07 | 9,55E+06 | 65,235 | 0,000147 | 3,2881 | 11 |
| 1,36E+06 | 6,62E+06 | 8,23E+06 | 7,32E+06 | 82,455 | 0,000435 | 2,9725 | 11 |
| 7,24E+06 | 1,24E+06 | 7,32E+06 | 5,21E+06 | 16,335 | 0,000436 | 2,9892 | 13 |
| 2,60E+07 | 2,53E+07 | 2,22E+07 | 3,82E+07 | 37,768 | 0 | 5,4932 | 27 |
| 2,57E+08 | 2,07E+08 | 1,67E+08 | 1,65E+08 | 53,398 | 0 | 274,19 | 72 |
| 5,27E+06 | 1,91E+06 | 1,27E+06 | 1,22E+06 | 22,344 | 0,003572 | 1,8866 | 2 |
| 1,16E+07 | 9,10E+06 | 1,15E+06 | 9,04E+06 | 16,299 | 0 | 6,5791 | 10 |
| 4,68E+07 | 5,66E+07 | 4,80E+07 | 3,38E+07 | 53,245 | 0 | 19,818 | 26 |
| 1,77E+06 | 2,83E+06 | 4,19E+06 | 2,27E+06 | 11,797 | 0,000291 | 3,0626 | 5 |
| 2,09E+06 | 5,31E+05 | 7,23E+05 | 2,01E+06 | 98,579 | 0,003569 | 1,8718 | 5 |
| 2,87E+07 | 4,18E+07 | 3,41E+07 | 3,75E+07 | 39,1 | 0 | 10,023 | 25 |
| 2,02E+06 | 2,26E+06 | 1,89E+06 | 2,25E+06 | 31,582 | 0,003829 | 1,7992 | 8 |
| 7,48E+06 | 5,86E+06 | 7,52E+06 | 7,90E+06 | 36,572 | 0 | 9,5167 | 15 |
| 6,96E+06 | 4,29E+06 | 4,02E+06 | 2,15E+06 | 160,51 | 0 | 10,808 | 11 |
| 1,52E+09 | 1,19E+09 | 1,31E+09 | 1,08E+09 | 52,901 | 0 | 323,31 | 255 |
| 2,00E+06 | 2,19E+06 | 5,26E+06 | 9,23E+06 | 65,21 | 0 | 16,035 | 13 |
| 6,50E+06 | 1,76E+06 | 9,44E+06 | 5,24E+06 | 46,952 | 0 | 6,8554 | 11 |
| 2,44E+07 | 2,63E+07 | 2,06E+07 | 2,79E+07 | 31,068 | 0 | 7,9713 | 20 |
| 2,43E+07 | 2,52E+07 | 2,66E+07 | 2,33E+07 | 49 | 0 | 13,717 | 32 |
| 2,58E+07 | 2,80E+07 | 2,88E+07 | 3,32E+07 | 51,026 | 0 | 7,5737 | 20 |
| 5,55E+07 | 4,51E+07 | 3,49E+07 | 3,56E+07 | 76,107 | 0 | 59,345 | 47 |
| 1,96E+07 | 1,47E+07 | 1,82E+07 | 2,02E+07 | 32,905 | 0 | 22,764 | 15 |
| 9,40E+06 | 5,75E+06 | 2,27E+06 | 4,78E+06 | 143,89 | 0,000147 | 3,161 | 6 |
| 8,65E+07 | 7,96E+07 | 1,08E+08 | 8,96E+07 | 6,784 | 0 | 63,824 | 29 |
| 1,13E+08 | 1,56E+08 | 1,28E+08 | 1,52E+08 | 15,874 | 0 | 42,886 | 25 |
| 3,98E+07 | 3,64E+07 | 5,29E+07 | 5,75E+07 | 26,263 | 0 | 43,658 | 22 |
| 6,22E+08 | 6,27E+08 | 5,46E+08 | 6,23E+08 | 53,662 | 0 | 323,31 | 138 |
| 3,86E+07 | 3,95E+07 | 3,48E+07 | 3,09E+07 | 41,135 | 0 | 15,478 | 33 |
| 7,56E+06 | 8,16E+06 | 7,81E+06 | 8,67E+06 | 59,653 | 0 | 6,9068 | 13 |
| 1,47E+07 | 1,87E+07 | 1,83E+07 | 1,84E+07 | 10,531 | 0,003448 | 1,9362 | 9 |
| 8,62E+07 | 1,13E+08 | 1,13E+08 | 1,15E+08 | 29,524 | 0 | 11,492 | 43 |
| 1,68E+07 | 1,52E+07 | 2,21E+07 | 1,86E+07 | 85,402 | 0 | 8,0689 | 18 |
| 1,14E+09 | 9,68E+08 | 9,41E+08 | 9,46E+08 | 43,158 | 0 | 107,64 | 129 |
| 1,24E+09 | 1,30E+09 | 1,29E+09 | 1,24E+09 | 52,851 | 0 | 323,31 | 240 |
| 3,14E+07 | 2,60E+07 | 1,88E+07 | 2,41E+07 | 50,871 | 0 | 47,044 | 30 |
| 3,76E+07 | 3,42E+07 | 3,97E+07 | 4,15E+07 | 47,737 | 0 | 9,5152 | 34 |
| 1,07E+10 | 6,28E+09 | 9,12E+09 | 8,84E+09 | 14,866 | 0 | 323,31 | 308 |
| 3,17E+09 | 1,85E+09 | 2,21E+09 | 2,53E+09 | 14,839 | 0 | 258,14 | 375 |
| 4,95E+06 | 3,10E+06 | 1,87E+06 | 4,45E+06 | 47,203 | 0 | 11,693 | 5 |
| 3,88E+08 | 3,73E+08 | 3,65E+08 | 3,49E+08 | 55,074 | 0 | 323,31 | 93 |
| 9,69E+06 | 2,41E+06 | 1,72E+07 | 6,45E+06 | 104,05 | 0,000853 | 2,5548 | 6 |

| | | | | | | | |
|---|---|---|---|---|---|---|---|
| 3,46E+08 | 2,77E+08 | 2,78E+08 | 2,46E+08 | 14,954 | 0 | 170,03 | 67 |
| 7,15E+06 | 6,78E+06 | 4,01E+06 | 1,02E+07 | 118,53 | 0,000157 | 4,8835 | 14 |
| 5,97E+07 | 2,28E+07 | 1,88E+07 | 2,39E+07 | 24,198 | 0 | 11,014 | 12 |
| 1,23E+06 | 1,52E+07 | 1,68E+07 | 1,48E+07 | 41,441 | 0 | 12,095 | 13 |
| 4,02E+06 | 3,46E+06 | 1,73E+06 | 3,97E+06 | 26,279 | 0,000157 | 4,9211 | 5 |
| 3,17E+09 | 1,84E+09 | 2,54E+09 | 2,27E+09 | 17,208 | 0 | 323,31 | 183 |
| 4,77E+07 | 6,11E+07 | 6,78E+07 | 5,94E+07 | 58,316 | 0 | 65,282 | 58 |
| 2,08E+07 | 1,40E+07 | 2,09E+07 | 1,43E+07 | 17,158 | 0 | 28 | 9 |
| 1,65E+07 | 1,46E+07 | 1,73E+07 | 1,62E+07 | 248,49 | 0 | 9,8725 | 25 |
| 1,53E+06 | 2,78E+06 | 2,51E+06 | 4,82E+06 | 85,489 | 0,009375 | 1,44 | 3 |
| 1,04E+08 | 9,27E+07 | 9,91E+07 | 1,06E+08 | 12,648 | 0 | 22,928 | 47 |
| 1,17E+08 | 1,62E+08 | 1,80E+08 | 2,43E+08 | 28,246 | 0 | 14,402 | 35 |
| 3,05E+06 | 5,18E+06 | 4,49E+06 | 4,25E+06 | 42,498 | 0,000151 | 3,9129 | 5 |
| 1,40E+07 | 1,68E+07 | 1,57E+07 | 8,18E+06 | 24,393 | 0 | 7,5987 | 17 |
| 2,05E+07 | 2,62E+07 | 2,34E+07 | 2,17E+07 | 37,158 | 0 | 6,4622 | 23 |
| 5,27E+07 | 4,30E+07 | 5,53E+07 | 6,57E+07 | 13,974 | 0 | 26,093 | 25 |
| 1,56E+07 | 1,34E+07 | 1,18E+07 | 1,11E+07 | 23,18 | 0,000158 | 5,1608 | 14 |
| 5,48E+06 | 3,21E+06 | 2,67E+06 | 4,78E+06 | 47,754 | 0,000147 | 3,1437 | 9 |
| 2,60E+08 | 2,71E+08 | 2,47E+08 | 2,84E+08 | 108,54 | 0 | 212,57 | 190 |
| 6,27E+07 | 7,07E+07 | 5,70E+07 | 5,19E+07 | 31,586 | 0 | 104,82 | 33 |
| 1,42E+07 | 1,83E+07 | 1,72E+07 | 1,61E+07 | 28,433 | 0 | 10,891 | 21 |
| 3,03E+07 | 2,55E+07 | 1,29E+07 | 2,29E+07 | 9,8742 | 0,000147 | 3,2628 | 16 |
| 1,27E+10 | 1,11E+10 | 1,09E+10 | 1,02E+10 | 226,37 | 0 | 323,31 | 2235 |
| 4,42E+08 | 3,77E+08 | 4,25E+08 | 3,91E+08 | 25,581 | 0 | 323,31 | 96 |
| 4,33E+07 | 5,21E+07 | 4,40E+07 | 5,08E+07 | 27,393 | 0 | 49,765 | 42 |
| 1,15E+07 | 1,58E+07 | 1,84E+07 | 1,46E+07 | 108,07 | 0 | 13,021 | 28 |
| 4,69E+07 | 3,49E+07 | 3,22E+07 | 2,22E+07 | 16,353 | 0 | 69,05 | 19 |
| 2,39E+08 | 2,68E+08 | 2,27E+08 | 2,33E+08 | 42,281 | 0 | 139,01 | 61 |
| 9,22E+07 | 1,13E+08 | 1,25E+08 | 1,12E+08 | 3906,4 | 0 | 5,687 | 39 |
| 8,92E+07 | 8,24E+07 | 9,64E+07 | 9,56E+07 | 120,63 | 0 | 212,26 | 81 |
| 1,26E+07 | 1,24E+07 | 1,09E+07 | 1,37E+07 | 36,227 | 0 | 5,7833 | 15 |
| 5,13E+07 | 3,67E+07 | 4,73E+07 | 3,67E+07 | 21,418 | 0 | 158,33 | 40 |
| 3,29E+08 | 7,64E+08 | 7,32E+08 | 7,29E+08 | 210,29 | 0,000434 | 2,906 | 13 |
| 4,04E+06 | 6,38E+06 | 3,94E+06 | 4,98E+06 | 20,828 | 0,001272 | 2,4232 | 4 |
| 1,66E+07 | 1,66E+07 | 1,41E+07 | 1,48E+07 | 23,528 | 0 | 8,9453 | 14 |
| 7,85E+06 | 1,47E+07 | 1,24E+07 | 1,31E+07 | 35,757 | 0 | 11,797 | 16 |
| 3,63E+06 | 6,15E+06 | 2,35E+06 | 4,55E+06 | 32,081 | 0,002095 | 2,1405 | 4 |
| 5,66E+07 | 6,66E+07 | 4,61E+07 | 2,70E+07 | 34,709 | 0 | 36,631 | 15 |
| 1,59E+08 | 1,64E+08 | 1,62E+08 | 1,72E+08 | 35,7 | 0 | 115,65 | 47 |
| 8,28E+06 | 8,66E+06 | 8,08E+06 | 9,51E+06 | 219,09 | 0,000154 | 4,4021 | 13 |
| 2,65E+07 | 4,01E+07 | 1,89E+07 | 2,10E+07 | 55,872 | 0 | 80,03 | 29 |
| 8,59E+08 | 1,21E+09 | 7,46E+08 | 9,30E+08 | 26,511 | 0 | 323,31 | 59 |
| 1,89E+07 | 2,28E+07 | 1,61E+07 | 1,71E+07 | 85,187 | 0 | 42,058 | 10 |
| 2,66E+06 | 1,64E+06 | 4,47E+06 | 4,85E+06 | 29,749 | 0,003569 | 1,8732 | 4 |
| 1,64E+06 | 2,71E+06 | 1,49E+06 | 7,65E+06 | 22,204 | 0,008459 | 1,4788 | 3 |
| 2,81E+07 | 3,61E+07 | 3,01E+07 | 3,88E+07 | 57,223 | 0 | 5,5064 | 29 |
| 5,96E+06 | 2,88E+06 | 3,31E+06 | 2,75E+06 | 89,542 | 0,000433 | 2,839 | 4 |
| 7,35E+07 | 7,64E+07 | 6,67E+07 | 7,24E+07 | 42,61 | 0 | 67,654 | 58 |
| 1,55E+07 | 1,91E+07 | 1,37E+07 | 1,40E+07 | 39,125 | 0 | 5,6443 | 19 |
| 4,97E+07 | 3,35E+07 | 3,62E+07 | 4,66E+07 | 59,856 | 0 | 7,2469 | 23 |

| | | | | | | | |
|---|---|---|---|---|---|---|---|
| 1,07E+07 | 1,54E+07 | 1,40E+07 | 1,07E+07 | 35,866 | 0,000158 | 5,1447 | 16 |
| 9,82E+07 | 9,92E+07 | 1,03E+08 | 9,82E+07 | 88,339 | 0 | 152,26 | 65 |
| 2,71E+07 | 2,79E+07 | 2,64E+07 | 2,74E+07 | 109,51 | 0 | 17,461 | 38 |
| 3,19E+06 | 8,69E+06 | 2,91E+06 | 4,13E+06 | 143,97 | 0,002914 | 2,0238 | 3 |
| 3,48E+07 | 3,75E+07 | 2,91E+07 | 3,53E+07 | 42,732 | 0 | 36,607 | 25 |
| 2,93E+06 | 5,66E+06 | 3,46E+06 | 3,56E+06 | 46,273 | 0,000146 | 3,0917 | 9 |
| 2,46E+06 | 5,09E+06 | 2,29E+06 | 2,99E+06 | 36,112 | 0,003696 | 1,8206 | 6 |
| 2,14E+07 | 2,46E+07 | 2,01E+07 | 1,99E+07 | 158,06 | 0 | 27,381 | 34 |
| 7,31E+08 | 7,59E+08 | 7,93E+08 | 7,81E+08 | 108,18 | 0 | 249,1 | 262 |
| 5,07E+07 | 5,48E+07 | 5,04E+07 | 4,92E+07 | 54,616 | 0 | 105,59 | 57 |
| 3,25E+06 | 1,75E+06 | 2,28E+06 | 3,58E+06 | 80,383 | 0,000291 | 3,0657 | 4 |
| 6,19E+06 | 1,87E+06 | 8,19E+06 | 8,25E+06 | 13,194 | 0 | 6,7002 | 11 |
| 1,14E+08 | 1,20E+08 | 1,02E+08 | 9,97E+07 | 14,194 | 0 | 33,679 | 44 |
| 1,60E+07 | 1,71E+07 | 1,80E+07 | 1,96E+07 | 75,596 | 0 | 18,515 | 21 |
| 2,79E+07 | 3,84E+07 | 3,36E+07 | 2,73E+07 | 56,104 | 0 | 32,015 | 46 |
| 3,13E+07 | 2,24E+07 | 1,67E+07 | 2,12E+07 | 58,221 | 0 | 42,403 | 19 |
| 2,11E+08 | 2,56E+08 | 2,26E+08 | 2,60E+08 | 37,35 | 0 | 27,459 | 66 |
| 1,12E+08 | 1,25E+08 | 9,57E+07 | 1,09E+08 | 203,75 | 0 | 67,707 | 81 |
| 3,08E+07 | 2,62E+07 | 1,74E+07 | 3,54E+07 | 20,733 | 0 | 9,969 | 22 |
| 6,61E+06 | 8,28E+06 | 7,57E+06 | 1,06E+07 | 39,787 | 0,000151 | 3,8289 | 13 |
| 2,67E+07 | 2,61E+07 | 2,90E+07 | 1,93E+07 | 124,41 | 0 | 11,218 | 27 |
| 4,77E+08 | 2,62E+08 | 2,87E+08 | 2,63E+08 | 28,211 | 0 | 323,31 | 105 |
| 3,07E+08 | 3,22E+08 | 3,01E+08 | 2,92E+08 | 337,18 | 0 | 178,2 | 187 |
| 1,31E+08 | 1,49E+08 | 1,34E+08 | 1,41E+08 | 51,007 | 0 | 74,994 | 73 |
| 1,42E+07 | 1,33E+07 | 1,92E+07 | 9,03E+06 | 18,53 | 0,000158 | 5,0683 | 13 |
| 2,00E+09 | 2,20E+09 | 2,08E+09 | 2,22E+09 | 50,906 | 0 | 323,31 | 330 |
| 1,04E+07 | 2,84E+06 | 1,18E+07 | 1,58E+07 | 37,723 | 0 | 13,65 | 10 |
| 1,99E+06 | 5,19E+06 | 2,41E+06 | 2,52E+06 | 127,53 | 0,000157 | 4,8549 | 4 |
| 2,81E+06 | 5,21E+06 | 2,38E+06 | 4,34E+06 | 119,1 | 0 | 8,2502 | 11 |
| 3,96E+07 | 4,59E+07 | 4,80E+07 | 4,25E+07 | 62,49 | 0 | 40,706 | 48 |
| 1,67E+06 | 4,83E+06 | 5,00E+06 | 4,90E+06 | 55,577 | 0,000855 | 2,6072 | 4 |
| 3,17E+06 | 3,10E+06 | 5,32E+06 | 3,70E+06 | 17,994 | 0,000146 | 3,0794 | 5 |
| 3,19E+08 | 3,35E+08 | 3,28E+08 | 3,39E+08 | 35,274 | 0 | 64,485 | 115 |
| 1,20E+07 | 7,33E+06 | 7,50E+06 | 7,44E+06 | 76,578 | 0,000431 | 2,7663 | 10 |
| 1,27E+07 | 7,60E+06 | 6,76E+06 | 8,77E+06 | 61,688 | 0,000148 | 3,3079 | 12 |
| 4,87E+07 | 8,86E+07 | 2,35E+07 | 5,34E+07 | 72,442 | 0,000154 | 4,2947 | 17 |
| 3,96E+06 | 4,33E+06 | 1,14E+06 | 4,98E+06 | 62,537 | 0,005816 | 1,6161 | 4 |
| 3,87E+09 | 4,26E+09 | 4,38E+09 | 4,16E+09 | 17,718 | 0 | 105,93 | 359 |
| 6,54E+07 | 6,46E+07 | 7,57E+07 | 8,62E+07 | 7,7059 | 0 | 61,986 | 38 |
| 5,87E+07 | 5,62E+07 | 5,53E+07 | 4,89E+07 | 161,09 | 0 | 37,856 | 69 |
| 3,08E+06 | 2,19E+07 | 1,88E+07 | 2,17E+07 | 19,069 | 0 | 33,246 | 6 |
| 1,92E+08 | 1,90E+08 | 1,92E+08 | 2,09E+08 | 25,57 | 0 | 286,4 | 83 |
| 8,22E+06 | 1,17E+07 | 1,21E+07 | 1,68E+07 | 13,219 | 0 | 16,864 | 6 |
| 4,99E+08 | 5,48E+08 | 4,28E+08 | 4,87E+08 | 43,214 | 0 | 196,34 | 135 |
| 1,01E+08 | 9,97E+07 | 9,71E+07 | 9,59E+07 | 22,848 | 0 | 75,261 | 36 |
| 6,45E+07 | 6,78E+07 | 6,90E+07 | 7,82E+07 | 45,926 | 0 | 32,789 | 59 |
| 1,37E+08 | 1,32E+08 | 1,56E+08 | 1,16E+08 | 13,36 | 0 | 42,44 | 41 |
| 1,29E+07 | 1,19E+07 | 1,24E+07 | 7,98E+06 | 141,43 | 0 | 13,691 | 18 |
| 4,77E+06 | 2,57E+06 | 4,41E+06 | 3,45E+06 | 11,756 | 0,001271 | 2,421 | 3 |
| 9,58E+07 | 1,10E+08 | 1,13E+08 | 1,30E+08 | 48,072 | 0 | 20,241 | 42 |

| | | | | | | | |
|---|---|---|---|---|---|---|---|
| 1,03E+09 | 9,21E+08 | 8,19E+08 | 8,61E+08 | 50,549 | 0 | 323,31 | 151 |
| 1,36E+07 | 2,56E+07 | 2,06E+07 | 2,19E+07 | 40,856 | 0 | 8,325 | 23 |
| 7,45E+07 | 7,84E+07 | 7,00E+07 | 6,81E+07 | 231,8 | 0 | 7,0292 | 18 |
| 1,26E+08 | 1,26E+08 | 1,26E+08 | 1,20E+08 | 73,999 | 0 | 183,95 | 55 |
| 5,71E+08 | 6,15E+08 | 5,61E+08 | 5,85E+08 | 45,095 | 0 | 317,78 | 165 |
| 2,71E+06 | 5,19E+06 | 2,57E+06 | 2,07E+06 | 36,219 | 0,001271 | 2,4172 | 6 |
| 1,54E+07 | 1,57E+07 | 1,55E+07 | 1,94E+07 | 69,378 | 0 | 5,8299 | 24 |
| 3,19E+07 | 2,45E+07 | 2,63E+07 | 2,65E+07 | 77,5 | 0 | 19,452 | 45 |
| 4,80E+06 | 6,04E+06 | 1,24E+06 | 3,78E+06 | 20,441 | 0,000433 | 2,8698 | 6 |
| 4,04E+07 | 5,21E+07 | 4,27E+07 | 4,82E+07 | 67,146 | 0 | 96,266 | 45 |
| 9,98E+06 | 2,89E+06 | 6,73E+06 | 1,40E+06 | 73,453 | 0,000432 | 2,8097 | 7 |
| 1,04E+09 | 1,10E+09 | 1,10E+09 | 1,09E+09 | 54,084 | 0 | 323,31 | 241 |
| 3,36E+07 | 3,98E+07 | 4,13E+07 | 3,79E+07 | 40,783 | 0 | 31,604 | 28 |
| 2,93E+08 | 2,99E+08 | 2,82E+08 | 2,95E+08 | 39,681 | 0 | 232,86 | 100 |
| 7,24E+07 | 7,32E+07 | 8,04E+07 | 7,45E+07 | 45,019 | 0 | 72,019 | 69 |
| 1,85E+09 | 1,57E+09 | 1,14E+09 | 1,18E+09 | 25,147 | 0 | 296,41 | 168 |
| 7,19E+06 | 8,11E+06 | 8,79E+06 | 1,03E+07 | 32,715 | 0,000155 | 4,4936 | 14 |
| 4,98E+07 | 5,73E+07 | 6,71E+07 | 6,02E+07 | 27,639 | 0 | 112,01 | 49 |
| 4,85E+06 | 7,10E+06 | 2,13E+06 | 6,27E+06 | 43,272 | 0,000148 | 3,4022 | 6 |
| 7,43E+07 | 8,99E+07 | 8,57E+07 | 8,99E+07 | 184,08 | 0 | 32,861 | 58 |
| 7,54E+06 | 1,82E+06 | 6,11E+06 | 5,12E+06 | 34,379 | 0,000151 | 3,8965 | 3 |
| 3,71E+09 | 3,20E+09 | 3,65E+09 | 3,22E+09 | 16,605 | 0 | 35,588 | 85 |
| 1,93E+08 | 1,17E+08 | 8,10E+07 | 7,42E+07 | 29,799 | 0 | 198,14 | 42 |
| 1,11E+07 | 1,18E+07 | 1,13E+07 | 8,61E+06 | 26,82 | 0 | 13,908 | 18 |
| 5,62E+07 | 5,14E+06 | 3,01E+06 | 6,25E+07 | 25,017 | 0,000435 | 2,9502 | 8 |
| 4,39E+07 | 4,21E+07 | 4,77E+07 | 4,12E+07 | 75,149 | 0 | 16,452 | 44 |
| 7,87E+09 | 7,85E+09 | 8,14E+09 | 7,91E+09 | 30,831 | 0 | 323,31 | 482 |
| 3,31E+07 | 2,63E+07 | 2,72E+07 | 3,33E+07 | 25,745 | 0 | 9,8751 | 29 |
| 1,40E+07 | 1,90E+07 | 2,28E+07 | 1,76E+07 | 35,882 | 0,003436 | 1,8999 | 7 |
| 1,39E+07 | 1,17E+07 | 1,57E+07 | 1,20E+07 | 54,914 | 0 | 6,5347 | 15 |
| 5,40E+07 | 4,94E+07 | 4,59E+07 | 5,68E+07 | 62,734 | 0 | 51,923 | 60 |
| 4,13E+06 | 3,39E+06 | 2,65E+06 | 1,74E+06 | 55,933 | 0 | 6,5066 | 11 |
| 7,52E+06 | 4,89E+06 | 2,15E+07 | 1,17E+07 | 15,421 | 0 | 11,373 | 9 |
| 2,12E+08 | 1,58E+08 | 1,81E+08 | 1,74E+08 | 139,57 | 0 | 191,97 | 78 |
| 3,87E+06 | 4,57E+06 | 6,94E+06 | 8,61E+05 | 90,41 | 0,000148 | 3,3535 | 7 |
| 7,77E+08 | 7,06E+08 | 9,58E+08 | 7,04E+08 | 6,4066 | 0 | 24,671 | 39 |
| 2,25E+06 | 2,15E+06 | 3,10E+06 | 7,08E+06 | 109,95 | 0,00015 | 3,6782 | 5 |
| 2,57E+09 | 1,62E+09 | 2,04E+09 | 1,77E+09 | 14,759 | 0 | 235,06 | 57 |
| 2,28E+07 | 2,20E+07 | 2,95E+07 | 3,61E+07 | 86,37 | 0 | 10,783 | 35 |
| 9,68E+07 | 1,41E+08 | 2,11E+08 | 1,86E+08 | 79,996 | 0 | 97,427 | 61 |
| 8,43E+07 | 8,83E+07 | 8,06E+07 | 8,68E+07 | 103,71 | 0 | 79,505 | 53 |
| 6,44E+06 | 1,48E+06 | 4,93E+06 | 6,55E+06 | 38,516 | 0 | 5,4468 | 5 |
| 1,40E+08 | 6,36E+07 | 1,39E+08 | 1,55E+08 | 8,9723 | 0 | 68,381 | 22 |
| 4,90E+07 | 4,16E+07 | 4,31E+07 | 4,67E+07 | 102,01 | 0,005154 | 1,6597 | 5 |
| 6,79E+07 | 7,15E+07 | 7,24E+07 | 7,75E+07 | 18,213 | 0 | 53,591 | 31 |
| 1,19E+07 | 1,38E+07 | 1,32E+07 | 1,52E+07 | 139,61 | 0 | 11,028 | 26 |
| 5,17E+07 | 6,99E+07 | 5,00E+07 | 5,78E+07 | 28,709 | 0 | 15,934 | 54 |
| 1,78E+08 | 1,70E+08 | 1,82E+08 | 1,72E+08 | 110,28 | 0 | 169,02 | 131 |
| 3,11E+08 | 2,60E+08 | 2,65E+08 | 2,23E+08 | 18,237 | 0 | 114,44 | 61 |
| 1,97E+07 | 2,59E+07 | 2,97E+07 | 2,75E+07 | 102,18 | 0 | 49,012 | 29 |

| | | | | | | | |
|---|---|---|---|---|---|---|---|
| 6,48E+07 | 7,32E+07 | 5,82E+07 | 5,72E+07 | 29,781 | 0 | 45,493 | 53 |
| 2,95E+06 | 1,61E+07 | 2,04E+07 | 2,12E+07 | 47,527 | 0 | 11,799 | 16 |
| 1,46E+09 | 1,46E+09 | 1,39E+09 | 1,38E+09 | 56,141 | 0 | 323,31 | 364 |
| 4,69E+06 | 6,34E+06 | 6,09E+06 | 4,54E+06 | 11,667 | 0,000146 | 3,0858 | 9 |
| 1,96E+07 | 2,38E+07 | 1,86E+07 | 1,63E+07 | 87,38 | 0 | 11,567 | 27 |
| 1,49E+09 | 1,69E+09 | 1,74E+09 | 1,95E+09 | 14,525 | 0 | 323,31 | 80 |
| 4,80E+06 | 8,03E+06 | 4,11E+06 | 3,83E+06 | 175,77 | 0 | 5,8297 | 9 |
| 8,11E+08 | 6,96E+08 | 6,18E+08 | 5,83E+08 | 22,677 | 0 | 323,31 | 79 |
| 3,87E+07 | 3,93E+07 | 4,55E+07 | 5,53E+07 | 48,597 | 0 | 20,214 | 29 |
| 8,64E+06 | 1,08E+06 | 1,46E+07 | 1,28E+07 | 16,815 | 0 | 81,25 | 4 |
| 1,08E+07 | 1,27E+07 | 1,86E+07 | 2,03E+07 | 16,683 | 0 | 17,893 | 21 |
| 1,67E+08 | 1,48E+08 | 1,61E+08 | 1,48E+08 | 156,46 | 0 | 99,826 | 115 |
| 4,77E+08 | 3,78E+08 | 3,71E+08 | 5,02E+08 | 22,988 | 0 | 75,747 | 77 |
| 1,01E+07 | 1,26E+07 | 7,45E+06 | 5,51E+06 | 34,734 | 0,000156 | 4,6779 | 10 |
| 1,30E+08 | 2,02E+08 | 1,63E+08 | 2,14E+08 | 20,495 | 0 | 323,31 | 57 |
| 1,45E+07 | 1,37E+07 | 1,38E+07 | 1,33E+07 | 71,442 | 0 | 5,6249 | 13 |
| 2,89E+07 | 1,48E+07 | 1,96E+07 | 1,86E+07 | 120,89 | 0,00015 | 3,7336 | 10 |
| 7,59E+08 | 8,65E+08 | 7,84E+08 | 8,01E+08 | 83,549 | 0 | 225,49 | 224 |
| 2,55E+09 | 2,40E+09 | 2,52E+09 | 2,42E+09 | 51,828 | 0 | 323,31 | 392 |
| 2,67E+08 | 2,97E+08 | 3,01E+08 | 3,27E+08 | 288,79 | 0 | 83,614 | 180 |
| 4,75E+08 | 5,06E+08 | 4,98E+08 | 5,12E+08 | 37,54 | 0 | 323,31 | 60 |
| 9,01E+06 | 8,61E+06 | 1,05E+07 | 1,16E+07 | 78,403 | 0 | 12,701 | 17 |
| 4,59E+07 | 6,43E+07 | 7,27E+07 | 5,33E+07 | 18,706 | 0 | 205,05 | 58 |
| 8,19E+06 | 5,51E+06 | 7,05E+06 | 6,33E+06 | 18,584 | 0,001824 | 2,2511 | 9 |
| 2,87E+07 | 2,81E+07 | 1,78E+07 | 2,75E+07 | 13,215 | 0,000159 | 5,1946 | 15 |
| 9,53E+06 | 4,31E+06 | 2,37E+06 | 1,33E+07 | 87,862 | 0 | 5,7365 | 8 |
| 2,54E+07 | 2,36E+07 | 2,11E+07 | 2,40E+07 | 75,81 | 0 | 20,67 | 28 |
| 5,52E+06 | 1,38E+06 | 2,77E+06 | 5,02E+06 | 95,083 | 0 | 5,7225 | 7 |
| 3,68E+06 | 1,73E+07 | 3,12E+07 | 1,74E+07 | 177,27 | 0 | 6,6164 | 22 |
| 1,04E+06 | 6,16E+06 | 8,02E+06 | 7,01E+06 | 53,182 | 0,000434 | 2,9188 | 7 |
| 1,73E+08 | 1,62E+07 | 1,02E+08 | 9,91E+07 | 21,05 | 0,000156 | 4,6458 | 60 |
| 1,39E+08 | 1,13E+08 | 1,34E+08 | 1,16E+08 | 72,263 | 0 | 57,621 | 74 |
| 5,28E+07 | 4,92E+07 | 4,44E+07 | 3,78E+07 | 36,778 | 0 | 69,747 | 36 |
| 7,31E+07 | 7,34E+07 | 6,38E+07 | 7,58E+07 | 34,299 | 0 | 16,994 | 39 |
| 8,17E+08 | 4,99E+08 | 5,41E+08 | 4,41E+08 | 34,883 | 0 | 323,31 | 86 |
| 1,25E+07 | 2,78E+06 | 2,74E+06 | 2,97E+06 | 117,86 | 0,008985 | 1,4653 | 4 |
| 1,53E+07 | 1,39E+07 | 1,85E+07 | 1,51E+07 | 16,493 | 0 | 13,416 | 12 |
| 4,36E+07 | 4,05E+07 | 4,36E+07 | 4,61E+07 | 24,131 | 0 | 63,143 | 20 |
| 1,32E+08 | 1,25E+08 | 1,27E+08 | 1,28E+08 | 18,017 | 0 | 40,392 | 36 |
| 1,05E+07 | 8,66E+06 | 1,03E+07 | 9,21E+06 | 43,374 | 0 | 8,8614 | 10 |
| 7,44E+07 | 7,24E+07 | 8,67E+07 | 7,23E+07 | 37,632 | 0 | 64,419 | 51 |
| 3,18E+06 | 1,67E+06 | 6,29E+07 | 4,49E+06 | 487,06 | 0,008208 | 1,5128 | 5 |
| 2,59E+07 | 2,49E+07 | 2,66E+07 | 3,07E+07 | 102,27 | 0 | 12,998 | 33 |
| 1,12E+07 | 1,06E+07 | 1,58E+07 | 1,42E+07 | 87,732 | 0 | 7,0023 | 18 |
| 1,17E+08 | 1,10E+08 | 9,52E+07 | 1,35E+08 | 7,4454 | 0 | 244,06 | 12 |
| 6,53E+07 | 7,28E+07 | 6,77E+07 | 6,28E+07 | 81,034 | 0 | 61,672 | 58 |
| 2,78E+06 | 3,80E+06 | 2,90E+06 | 2,98E+06 | 53,574 | 0,000852 | 2,5428 | 7 |
| 9,13E+07 | 1,33E+08 | 1,23E+08 | 1,43E+08 | 33,982 | 0 | 97,329 | 72 |
| 1,04E+08 | 1,02E+08 | 1,04E+08 | 9,98E+07 | 44,895 | 0 | 77,442 | 51 |
| 4,46E+06 | 7,54E+06 | 1,05E+07 | 8,63E+06 | 76,81 | 0 | 6,9123 | 9 |

| | | | | | | | |
|---|---|---|---|---|---|---|---|
| 2,03E+07 | 2,92E+07 | 3,38E+07 | 2,51E+07 | 219,66 | 0 | 10,503 | 26 |
| 2,03E+07 | 2,05E+07 | 2,18E+07 | 2,50E+07 | 32,677 | 0 | 28,217 | 33 |
| 4,04E+08 | 2,25E+08 | 3,67E+08 | 3,36E+08 | 17,595 | 0 | 81,569 | 62 |
| 4,23E+07 | 6,09E+07 | 4,27E+07 | 5,18E+07 | 17,595 | 0 | 11,168 | 29 |
| 5,24E+06 | 1,24E+07 | 1,59E+07 | 1,12E+07 | 17,476 | 0 | 36,329 | 7 |
| 7,20E+08 | 4,15E+08 | 5,96E+08 | 3,94E+08 | 34,463 | 0 | 323,31 | 107 |
| 8,98E+08 | 7,15E+08 | 8,85E+08 | 8,59E+08 | 32,25 | 0 | 192,34 | 125 |
| 9,63E+08 | 9,69E+08 | 9,87E+08 | 9,54E+08 | 133,42 | 0 | 323,31 | 325 |
| 1,87E+07 | 2,30E+07 | 1,92E+07 | 2,46E+07 | 85,493 | 0 | 14,216 | 20 |
| 3,02E+08 | 2,52E+08 | 2,44E+08 | 3,62E+08 | 18,936 | 0 | 57,346 | 25 |
| 2,32E+07 | 1,56E+07 | 2,19E+07 | 2,74E+07 | 35,575 | 0 | 323,31 | 14 |
| 5,22E+06 | 7,29E+06 | 9,50E+06 | 4,86E+06 | 39,835 | 0 | 5,8338 | 13 |
| 8,86E+06 | 8,85E+06 | 8,14E+06 | 1,10E+07 | 96,81 | 0 | 6,4951 | 15 |
| 5,29E+06 | 5,68E+06 | 6,63E+06 | 8,11E+06 | 45,325 | 0,0021 | 2,1818 | 11 |
| 2,49E+06 | 2,95E+06 | 3,91E+06 | 3,62E+06 | 10,366 | 0 | 323,31 | 3 |
| 7,32E+07 | 8,14E+07 | 9,19E+07 | 9,06E+07 | 52,451 | 0 | 92,713 | 66 |
| 6,27E+08 | 5,48E+08 | 5,65E+08 | 6,09E+08 | 228,58 | 0 | 8,975 | 32 |
| 1,59E+06 | 1,64E+07 | 1,65E+07 | 1,39E+07 | 46,412 | 0 | 42,912 | 6 |
| 4,74E+06 | 5,43E+06 | 5,12E+06 | 2,87E+06 | 22,47 | 0,000154 | 4,3937 | 5 |
| 7,41E+07 | 9,39E+07 | 7,48E+07 | 1,09E+08 | 13,724 | 0 | 268,24 | 18 |
| 2,14E+08 | 2,31E+08 | 2,62E+08 | 2,79E+08 | 40,707 | 0 | 64,59 | 58 |
| 5,06E+09 | 4,95E+09 | 5,38E+09 | 5,02E+09 | 149,47 | 0 | 323,31 | 700 |
| 3,13E+07 | 2,85E+07 | 3,80E+07 | 3,69E+07 | 89,972 | 0 | 7,6998 | 22 |
| 1,61E+06 | 3,75E+05 | 3,52E+06 | 3,03E+06 | 222,51 | 0,00015 | 3,6509 | 2 |
| 2,47E+08 | 2,21E+08 | 2,12E+08 | 2,10E+08 | 36,983 | 0 | 40,899 | 32 |
| 1,65E+07 | 2,10E+07 | 2,24E+07 | 1,94E+07 | 33,275 | 0,000435 | 2,9837 | 15 |
| 3,80E+07 | 2,87E+07 | 3,26E+07 | 3,41E+07 | 20,996 | 0 | 17,232 | 21 |
| 2,41E+07 | 1,78E+07 | 1,91E+07 | 2,18E+07 | 21,333 | 0,000159 | 5,2726 | 8 |
| 1,62E+09 | 1,66E+09 | 1,55E+09 | 1,55E+09 | 80,859 | 0 | 323,31 | 192 |
| 8,76E+07 | 9,31E+07 | 8,49E+07 | 8,29E+07 | 102,15 | 0 | 25,908 | 75 |
| 3,91E+06 | 2,91E+06 | 4,20E+06 | 1,14E+07 | 304,47 | 0,002507 | 2,0759 | 6 |
| 4,49E+07 | 5,93E+07 | 5,19E+07 | 4,52E+07 | 16,297 | 0 | 17,751 | 42 |
| 4,24E+07 | 3,49E+07 | 3,72E+07 | 3,99E+07 | 297,59 | 0 | 25,61 | 48 |
| 3,55E+08 | 3,29E+08 | 3,25E+08 | 3,41E+08 | 61,449 | 0 | 230,89 | 112 |
| 6,84E+07 | 6,25E+07 | 5,15E+07 | 4,35E+07 | 18,111 | 0 | 162 | 26 |
| 5,61E+07 | 3,27E+07 | 4,36E+07 | 3,47E+07 | 18,566 | 0 | 11,434 | 39 |
| 4,62E+08 | 5,23E+08 | 5,14E+08 | 4,75E+08 | 20,767 | 0 | 235,51 | 80 |
| 6,10E+06 | 4,77E+06 | 8,62E+06 | 1,07E+07 | 85,686 | 0 | 8,8374 | 14 |
| 1,15E+07 | 7,30E+06 | 9,38E+06 | 4,17E+06 | 33,125 | 0 | 50,966 | 5 |
| 1,91E+09 | 2,07E+09 | 1,93E+09 | 2,05E+09 | 12,784 | 0 | 165,02 | 133 |
| 2,81E+06 | 1,92E+06 | 3,27E+06 | 6,81E+06 | 92,07 | 0,000149 | 3,5735 | 2 |
| 5,32E+06 | 8,67E+06 | 5,10E+06 | 5,17E+06 | 66,095 | 0,00043 | 2,7303 | 9 |
| 1,99E+08 | 2,44E+08 | 2,41E+08 | 2,32E+08 | 71,543 | 0 | 227,86 | 94 |
| 8,47E+07 | 8,15E+07 | 9,10E+07 | 8,34E+07 | 74,398 | 0 | 56,353 | 54 |
| 5,26E+06 | 7,33E+06 | 3,98E+06 | 5,11E+06 | 13,881 | 0,000853 | 2,5643 | 9 |
| 2,67E+07 | 6,15E+06 | 6,92E+06 | 6,68E+06 | 34,052 | 0 | 49,713 | 7 |
| 5,12E+07 | 4,24E+07 | 1,62E+07 | 1,95E+07 | 18,929 | 0,002094 | 2,1298 | 10 |
| 2,20E+07 | 2,46E+07 | 2,57E+07 | 2,96E+07 | 109,05 | 0 | 9,8131 | 10 |
| 1,29E+07 | 5,36E+06 | 5,30E+06 | 5,47E+06 | 22,853 | 0,000148 | 3,4075 | 5 |
| 3,46E+06 | 3,19E+06 | 3,15E+06 | 3,09E+06 | 104,2 | 0,002095 | 2,1499 | 5 |

| | | | | | | | |
|---|---|---|---|---|---|---|---|
| 6,61E+06 | 5,81E+06 | 9,15E+06 | 5,63E+06 | 17,458 | 0,000147 | 3,2662 | 7 |
| 2,59E+07 | 2,14E+07 | 1,84E+07 | 1,98E+07 | 27,205 | 0 | 14,044 | 12 |
| 1,18E+07 | 1,26E+07 | 7,07E+06 | 9,45E+06 | 65,617 | 0,000156 | 4,611 | 9 |
| 6,05E+06 | 6,42E+06 | 4,43E+06 | 4,53E+06 | 31,805 | 0,001269 | 2,3739 | 7 |
| 1,04E+08 | 1,06E+08 | 7,55E+07 | 5,57E+07 | 15,192 | 0 | 11,775 | 17 |
| 1,80E+07 | 2,34E+07 | 1,66E+07 | 1,88E+07 | 13,43 | 0 | 11,275 | 15 |
| 4,01E+07 | 4,31E+07 | 4,32E+07 | 5,04E+07 | 13,841 | 0 | 19,322 | 35 |
| 5,34E+07 | 6,59E+07 | 6,27E+07 | 5,42E+07 | 85,873 | 0,002781 | 2,0549 | 5 |
| 6,94E+07 | 9,43E+07 | 6,70E+07 | 6,94E+07 | 13,02 | 0 | 69,557 | 42 |
| 1,91E+07 | 3,69E+07 | 3,94E+07 | 3,48E+06 | 9,584 | 0 | 9,8297 | 5 |
| 6,53E+06 | 2,96E+06 | 3,89E+06 | 1,70E+07 | 31,61 | 0 | 14,167 | 2 |
| 9,33E+07 | 7,74E+07 | 7,82E+07 | 8,76E+07 | 53,846 | 0 | 98,468 | 60 |
| 3,05E+07 | 2,25E+07 | 2,82E+07 | 2,82E+07 | 126,78 | 0 | 12,314 | 34 |
| 4,47E+07 | 5,33E+07 | 4,22E+07 | 3,86E+07 | 33,497 | 0 | 19,219 | 22 |
| 4,50E+07 | 4,12E+07 | 4,00E+07 | 3,34E+07 | 67,235 | 0 | 24,52 | 51 |
| 1,20E+07 | 1,20E+07 | 1,14E+07 | 1,16E+07 | 28,535 | 0 | 9,5517 | 21 |
| 4,83E+06 | 1,28E+07 | 5,65E+06 | 2,89E+06 | 110,82 | 0 | 6,6666 | 17 |
| 2,28E+07 | 1,69E+07 | 1,84E+07 | 1,28E+07 | 64,617 | 0 | 21,89 | 20 |
| 4,59E+07 | 5,18E+07 | 3,69E+07 | 4,83E+07 | 105,06 | 0 | 8,6373 | 15 |
| 2,35E+08 | 2,21E+08 | 2,08E+08 | 2,24E+08 | 39,218 | 0 | 148,19 | 81 |
| 1,61E+07 | 1,92E+07 | 1,32E+07 | 1,58E+07 | 31,136 | 0 | 41,548 | 7 |
| 3,66E+07 | 4,22E+07 | 3,38E+07 | 3,23E+07 | 31,791 | 0 | 9,9842 | 27 |
| 2,38E+08 | 2,69E+08 | 2,47E+08 | 2,47E+08 | 44,929 | 0 | 114,66 | 120 |
| 8,09E+07 | 7,93E+07 | 7,94E+07 | 8,43E+07 | 40,744 | 0 | 61,42 | 48 |
| 5,98E+07 | 5,14E+07 | 5,40E+07 | 6,03E+07 | 78,3 | 0 | 32,293 | 44 |
| 2,46E+07 | 2,67E+07 | 2,53E+07 | 2,36E+07 | 44,049 | 0 | 17,483 | 22 |
| 1,76E+08 | 1,73E+08 | 1,60E+08 | 1,68E+08 | 100,12 | 0 | 87,969 | 101 |
| 1,13E+07 | 8,66E+06 | 1,07E+07 | 4,93E+06 | 74,053 | 0 | 7,4603 | 12 |
| 9,63E+06 | 9,20E+06 | 7,44E+06 | 7,06E+06 | 56,793 | 0 | 6,7979 | 18 |
| 3,65E+06 | 1,03E+07 | 6,02E+06 | 3,04E+06 | 46,502 | 0 | 6,4814 | 6 |
| 8,58E+06 | 1,08E+07 | 1,11E+07 | 1,32E+07 | 47,32 | 0,003704 | 1,8593 | 6 |
| 2,10E+08 | 2,06E+08 | 2,26E+08 | 2,17E+08 | 73,742 | 0 | 154,52 | 113 |
| 6,37E+06 | 3,64E+06 | 7,37E+06 | 5,12E+06 | 22,576 | 0 | 12,289 | 16 |
| 3,50E+08 | 3,32E+08 | 3,57E+08 | 4,07E+08 | 31,55 | 0 | 125,22 | 72 |
| 3,59E+07 | 4,50E+07 | 4,47E+07 | 4,90E+07 | 142,05 | 0 | 23,574 | 41 |
| 1,66E+07 | 1,77E+07 | 1,68E+07 | 2,00E+07 | 39,072 | 0 | 6,7993 | 23 |
| 1,13E+08 | 9,11E+07 | 1,03E+08 | 9,78E+07 | 28,712 | 0 | 78,62 | 56 |
| 1,79E+07 | 2,15E+07 | 2,09E+07 | 1,65E+07 | 41,144 | 0 | 12,421 | 31 |
| 2,61E+07 | 2,61E+07 | 2,23E+07 | 2,37E+07 | 160,49 | 0 | 19,612 | 39 |
| 3,37E+08 | 3,37E+08 | 3,13E+08 | 3,75E+08 | 58,482 | 0 | 229,08 | 64 |
| 4,55E+07 | 4,73E+07 | 6,85E+07 | 9,17E+07 | 10,916 | 0 | 31,148 | 22 |
| 3,37E+06 | 1,82E+06 | 3,95E+06 | 4,33E+06 | 60,705 | 0,002094 | 2,1319 | 7 |
| 2,47E+07 | 1,77E+07 | 1,49E+07 | 1,75E+07 | 43,268 | 0,004629 | 1,7245 | 8 |
| 2,69E+07 | 2,90E+07 | 3,03E+07 | 2,61E+07 | 91,391 | 0 | 17,388 | 26 |
| 1,49E+07 | 1,14E+07 | 2,30E+07 | 2,50E+07 | 15,771 | 0 | 25,724 | 28 |
| 7,94E+06 | 1,01E+07 | 8,16E+06 | 6,19E+06 | 123,81 | 0 | 30,359 | 12 |
| 1,50E+07 | 1,18E+07 | 1,12E+07 | 1,07E+07 | 22 | 0,000434 | 2,9052 | 5 |
| 2,78E+08 | 2,54E+08 | 2,75E+08 | 2,35E+08 | 26,021 | 0 | 218,65 | 79 |
| 1,71E+07 | 1,70E+07 | 2,01E+07 | 2,17E+07 | 101,78 | 0 | 8,2007 | 20 |
| 2,79E+07 | 4,38E+07 | 3,86E+07 | 3,57E+07 | 20,833 | 0 | 13,568 | 37 |

| | | | | | | | |
|---|---|---|---|---|---|---|---|
| 6,86E+06 | 3,21E+06 | 3,74E+06 | 6,38E+06 | 132,4 | 0,000153 | 4,2254 | 2 |
| 4,59E+06 | 6,99E+06 | 2,74E+06 | 6,62E+06 | 37,849 | 0,003438 | 1,9036 | 7 |
| 6,66E+07 | 5,58E+07 | 6,39E+07 | 6,11E+07 | 51,658 | 0 | 85,802 | 42 |
| 1,55E+08 | 1,57E+08 | 1,41E+08 | 1,49E+08 | 26,259 | 0 | 14,335 | 40 |
| 2,62E+07 | 2,61E+07 | 2,92E+07 | 3,00E+07 | 20,886 | 0 | 133,28 | 27 |
| 1,55E+08 | 1,20E+08 | 1,19E+08 | 1,33E+08 | 48,584 | 0 | 116,99 | 92 |
| 4,01E+07 | 4,10E+07 | 4,90E+07 | 3,52E+07 | 19,556 | 0 | 47,718 | 29 |
| 3,05E+08 | 3,05E+08 | 2,72E+08 | 2,95E+08 | 170,87 | 0 | 234,74 | 154 |
| 6,29E+06 | 9,97E+06 | 8,35E+06 | 1,17E+07 | 90,959 | 0 | 5,5937 | 19 |
| 2,55E+07 | 8,47E+07 | 1,11E+07 | 6,62E+06 | 36,302 | 0,000153 | 4,1325 | 11 |
| 2,20E+06 | 2,78E+06 | 3,72E+06 | 3,63E+06 | 136,69 | 0,000149 | 3,4231 | 8 |
| 5,17E+08 | 5,10E+08 | 5,09E+08 | 4,43E+08 | 89,69 | 0 | 268,13 | 210 |
| 1,42E+08 | 2,82E+06 | 5,52E+06 | 3,74E+06 | 51,267 | 0 | 110,44 | 8 |
| 8,12E+07 | 8,26E+07 | 8,53E+07 | 9,22E+07 | 149,47 | 0 | 61,819 | 72 |
| 8,12E+07 | 1,01E+08 | 9,58E+07 | 6,81E+07 | 28,73 | 0 | 42,543 | 52 |
| 1,47E+08 | 2,54E+08 | 1,16E+08 | 1,76E+08 | 17,587 | 0 | 9,2024 | 63 |
| 3,96E+06 | 3,78E+06 | 3,06E+06 | 2,66E+06 | 122,52 | 0,001962 | 2,2215 | 3 |
| 4,27E+07 | 3,92E+07 | 4,15E+07 | 4,67E+07 | 175,89 | 0 | 30,641 | 46 |
| 1,41E+09 | 1,41E+09 | 1,33E+09 | 1,44E+09 | 41,829 | 0 | 323,31 | 267 |
| 3,87E+07 | 3,17E+07 | 2,57E+07 | 2,29E+07 | 9,4618 | 0,000155 | 4,425 | 14 |
| 5,17E+08 | 4,30E+08 | 4,05E+08 | 4,35E+08 | 38,813 | 0 | 323,31 | 60 |
| 6,76E+08 | 6,51E+08 | 6,73E+08 | 7,38E+08 | 82,988 | 0 | 323,31 | 220 |
| 3,65E+06 | 5,59E+06 | 8,26E+05 | 2,78E+06 | 80,534 | 0,000435 | 2,9405 | 5 |
| 2,72E+07 | 2,11E+07 | 3,50E+07 | 3,35E+07 | 23,415 | 0 | 156,34 | 29 |
| 1,07E+08 | 9,43E+07 | 9,46E+07 | 9,86E+07 | 116,05 | 0 | 60,346 | 62 |
| 9,94E+06 | 1,47E+07 | 7,21E+06 | 1,01E+07 | 43,11 | 0 | 6,4243 | 21 |
| 3,01E+07 | 2,10E+07 | 2,31E+07 | 3,41E+07 | 32,494 | 0 | 60,023 | 32 |
| 1,21E+09 | 1,03E+09 | 9,76E+08 | 1,05E+09 | 83,953 | 0 | 323,31 | 229 |
| 1,72E+09 | 1,71E+09 | 1,79E+09 | 1,72E+09 | 59,696 | 0 | 323,31 | 227 |
| 3,05E+08 | 3,57E+08 | 3,29E+08 | 3,19E+08 | 42,525 | 0 | 198,06 | 138 |
| 7,08E+06 | 9,45E+06 | 1,02E+07 | 1,19E+07 | 275,75 | 0,009371 | 1,4381 | 7 |
| 5,23E+07 | 4,48E+07 | 4,76E+07 | 4,70E+07 | 56,202 | 0 | 15,347 | 35 |
| 9,08E+07 | 8,79E+07 | 7,66E+07 | 7,98E+07 | 41,026 | 0 | 38,384 | 70 |
| 3,79E+07 | 4,43E+07 | 4,61E+07 | 4,63E+07 | 45,336 | 0 | 51,804 | 43 |
| 8,44E+06 | 8,58E+06 | 6,63E+06 | 1,11E+07 | 21,961 | 0 | 5,635 | 8 |
| 1,42E+08 | 1,83E+08 | 1,38E+08 | 1,46E+08 | 140,97 | 0 | 187,62 | 88 |
| 7,56E+07 | 9,52E+07 | 8,40E+07 | 7,64E+07 | 48,508 | 0 | 9,9127 | 46 |
| 3,59E+07 | 3,38E+07 | 4,08E+07 | 4,02E+07 | 38,195 | 0 | 14,369 | 33 |
| 6,07E+06 | 5,15E+06 | 4,78E+06 | 4,61E+06 | 68,317 | 0,00015 | 3,6698 | 7 |
| 3,02E+07 | 3,32E+07 | 2,88E+07 | 2,61E+07 | 34,443 | 0 | 10,577 | 32 |
| 4,76E+07 | 5,35E+07 | 5,61E+07 | 5,10E+07 | 20,745 | 0 | 323,31 | 39 |
| 6,94E+08 | 6,28E+08 | 5,56E+08 | 5,89E+08 | 42,785 | 0 | 323,31 | 120 |
| 1,18E+09 | 1,21E+09 | 1,21E+09 | 1,16E+09 | 51,572 | 0 | 323,31 | 402 |
| 2,63E+07 | 2,70E+07 | 3,47E+07 | 3,46E+07 | 51,691 | 0 | 20,283 | 29 |
| 2,44E+08 | 2,30E+08 | 2,50E+08 | 2,63E+08 | 28,369 | 0 | 35,577 | 52 |
| 2,61E+08 | 2,54E+08 | 2,52E+08 | 2,82E+08 | 88,796 | 0 | 184,64 | 120 |
| 5,40E+08 | 6,39E+08 | 6,29E+08 | 5,58E+08 | 54,087 | 0 | 323,31 | 175 |
| 1,22E+09 | 8,32E+08 | 6,58E+08 | 6,38E+08 | 27,778 | 0 | 323,31 | 116 |
| 3,09E+07 | 4,17E+07 | 4,63E+07 | 4,38E+07 | 75,59 | 0 | 42,926 | 42 |
| 2,78E+07 | 3,43E+07 | 3,80E+07 | 3,58E+07 | 11,753 | 0 | 9,1597 | 26 |

| | | | | | | | |
|---|---|---|---|---|---|---|---|
| 2,66E+07 | 3,01E+07 | 3,17E+07 | 3,17E+07 | 284 | 0 | 27,218 | 48 |
| 1,86E+07 | 2,00E+07 | 1,91E+07 | 1,76E+07 | 150,79 | 0 | 20,679 | 26 |
| 6,70E+07 | 7,63E+07 | 7,38E+07 | 8,09E+07 | 24,385 | 0 | 29,58 | 50 |
| 3,22E+08 | 3,35E+08 | 3,85E+08 | 3,53E+08 | 93,245 | 0 | 73,892 | 92 |
| 5,79E+07 | 5,79E+07 | 5,81E+07 | 5,76E+07 | 45 | 0 | 19,904 | 36 |
| 1,90E+06 | 2,71E+06 | 5,31E+06 | 3,78E+06 | 8,873 | 0,003699 | 1,825 | 3 |
| 2,36E+07 | 1,68E+07 | 1,32E+07 | 1,84E+07 | 22,238 | 0 | 24,156 | 26 |
| 8,47E+06 | 7,73E+06 | 9,10E+06 | 1,02E+07 | 141,59 | 0,000291 | 3,0585 | 9 |
| 3,48E+06 | 7,03E+06 | 5,39E+06 | 6,66E+06 | 31,331 | 0 | 6,121 | 8 |
| 2,43E+06 | 2,93E+06 | 2,23E+06 | 2,70E+06 | 33,925 | 0 | 7,3474 | 8 |
| 2,26E+07 | 2,55E+07 | 2,67E+07 | 1,97E+07 | 42,106 | 0 | 18,721 | 23 |
| 2,00E+08 | 2,38E+08 | 2,24E+08 | 2,38E+08 | 37,596 | 0 | 146,99 | 76 |
| 4,88E+08 | 4,24E+08 | 4,33E+08 | 3,83E+08 | 46,638 | 0 | 323,31 | 121 |
| 7,81E+07 | 5,82E+07 | 6,68E+07 | 7,12E+07 | 15,847 | 0 | 68,734 | 32 |
| 1,37E+08 | 1,97E+08 | 1,94E+08 | 1,69E+08 | 68,393 | 0 | 183,5 | 193 |
| 7,77E+07 | 5,99E+07 | 8,29E+07 | 7,95E+07 | 15,944 | 0 | 20,545 | 40 |
| 3,79E+07 | 3,94E+07 | 2,52E+07 | 2,48E+07 | 15,121 | 0 | 7,6286 | 42 |
| 3,83E+06 | 8,08E+06 | 2,94E+06 | 9,81E+06 | 69,451 | 0,000436 | 2,9957 | 10 |
| 1,88E+09 | 1,96E+09 | 1,93E+09 | 1,85E+09 | 70,299 | 0 | 323,31 | 261 |
| 2,29E+07 | 2,39E+07 | 2,72E+07 | 2,45E+07 | 19,655 | 0 | 33,967 | 15 |
| 8,93E+06 | 5,22E+06 | 6,68E+06 | 4,56E+06 | 11,314 | 0,000154 | 4,2759 | 4 |
| 5,06E+06 | 7,11E+06 | 1,40E+06 | 4,50E+06 | 15,801 | 0 | 10,832 | 4 |
| 8,79E+08 | 6,73E+08 | 7,39E+08 | 7,80E+08 | 98,144 | 0 | 323,31 | 221 |
| 8,47E+09 | 5,70E+09 | 6,34E+09 | 6,09E+09 | 16,837 | 0 | 323,31 | 400 |
| 1,06E+07 | 4,48E+06 | 2,58E+06 | 1,25E+06 | 51,247 | 0 | 8,6496 | 3 |
| 1,66E+06 | 8,74E+06 | 3,96E+06 | 7,66E+06 | 114,35 | 0,000152 | 4,0122 | 8 |
| 2,73E+08 | 4,02E+08 | 2,78E+08 | 3,84E+08 | 40,538 | 0 | 323,31 | 62 |
| 1,65E+06 | 9,61E+06 | 6,52E+06 | 3,91E+06 | 15,806 | 0 | 5,5298 | 6 |
| 4,79E+07 | 5,55E+07 | 5,36E+07 | 5,30E+07 | 104,19 | 0 | 24,277 | 56 |
| 2,86E+08 | 2,53E+08 | 2,90E+08 | 2,94E+08 | 49,419 | 0 | 152,04 | 79 |
| 5,81E+06 | 3,68E+06 | 2,71E+07 | 2,91E+07 | 46,977 | 0,003702 | 1,847 | 2 |
| 9,09E+07 | 8,16E+07 | 8,74E+07 | 9,44E+07 | 7,645 | 0,000147 | 3,2486 | 14 |
| 1,17E+09 | 9,85E+08 | 1,06E+09 | 1,06E+09 | 35,638 | 0 | 323,31 | 182 |
| 4,54E+09 | 4,62E+09 | 5,20E+09 | 5,52E+09 | 32,904 | 0 | 282,7 | 237 |
| 3,43E+07 | 4,36E+07 | 3,31E+07 | 3,50E+07 | 18,885 | 0 | 7,5024 | 23 |
| 7,04E+06 | 2,46E+06 | 1,94E+06 | 6,87E+06 | 28,287 | 0,002092 | 2,1159 | 2 |
| 1,26E+08 | 1,28E+08 | 1,16E+08 | 1,33E+08 | 51,213 | 0 | 30,822 | 56 |
| 2,08E+07 | 2,51E+07 | 2,63E+07 | 2,51E+07 | 14,677 | 0 | 323,31 | 16 |
| 1,58E+07 | 1,88E+07 | 1,46E+07 | 1,74E+07 | 41,033 | 0 | 7,7728 | 15 |
| 1,57E+08 | 1,20E+08 | 1,68E+08 | 1,71E+08 | 38,439 | 0 | 8,2832 | 12 |
| 1,06E+08 | 1,04E+08 | 1,12E+08 | 1,14E+08 | 86,072 | 0 | 32,873 | 90 |
| 6,33E+08 | 6,24E+08 | 6,27E+08 | 6,97E+08 | 27,545 | 0 | 73,923 | 172 |
| 3,44E+06 | 7,35E+06 | 5,22E+06 | 6,36E+06 | 166,67 | 0 | 5,6809 | 11 |
| 3,70E+08 | 3,77E+08 | 3,70E+08 | 4,10E+08 | 94,652 | 0 | 226,51 | 184 |
| 1,01E+08 | 7,73E+05 | 5,34E+06 | 3,67E+08 | 129,69 | 0,008588 | 1,4728 | 5 |
| 3,90E+09 | 4,04E+09 | 3,90E+09 | 4,31E+09 | 64,464 | 0 | 323,31 | 493 |
| 2,14E+07 | 3,62E+07 | 3,81E+07 | 3,60E+07 | 49,509 | 0 | 11,895 | 19 |
| 3,32E+09 | 2,82E+09 | 3,41E+09 | 4,17E+09 | 16,832 | 0 | 323,31 | 187 |
| 2,30E+07 | 2,44E+07 | 1,66E+07 | 1,92E+07 | 40,241 | 0 | 42,028 | 22 |
| 1,50E+08 | 1,74E+08 | 1,52E+08 | 1,75E+08 | 94,422 | 0 | 169,02 | 90 |

| | | | | | | | |
|---|---|---|---|---|---|---|---|
| 1,99E+08 | 1,84E+08 | 1,92E+08 | 1,92E+08 | 25,564 | 0 | 23,63 | 61 |
| 1,33E+08 | 1,92E+08 | 1,93E+08 | 2,25E+08 | 23,598 | 0 | 67,99 | 49 |
| 6,12E+07 | 5,10E+07 | 6,90E+07 | 6,11E+07 | 237,32 | 0 | 9,8697 | 39 |
| 4,38E+09 | 3,47E+09 | 4,06E+09 | 4,14E+09 | 17,363 | 0 | 180,85 | 245 |
| 6,48E+06 | 6,36E+06 | 5,70E+06 | 4,29E+06 | 14,468 | 0,001275 | 2,4686 | 6 |
| 1,38E+08 | 1,32E+08 | 1,21E+08 | 1,31E+08 | 12,51 | 0 | 97,859 | 55 |
| 2,74E+06 | 1,66E+06 | 7,12E+06 | 1,09E+07 | 33,667 | 0,000428 | 2,6646 | 8 |
| 5,97E+08 | 6,49E+08 | 5,59E+08 | 5,30E+08 | 75,38 | 0 | 323,31 | 147 |
| 2,03E+08 | 2,21E+08 | 2,21E+08 | 2,28E+08 | 103,47 | 0 | 323,31 | 182 |
| 2,97E+07 | 2,57E+07 | 2,80E+07 | 3,63E+07 | 74,817 | 0 | 36,461 | 33 |
| 2,25E+07 | 2,39E+07 | 2,44E+07 | 2,48E+07 | 48,004 | 0 | 11,267 | 37 |
| 1,87E+08 | 1,82E+08 | 1,72E+08 | 1,56E+08 | 64,86 | 0 | 142,61 | 109 |
| 3,51E+09 | 3,99E+09 | 4,05E+09 | 4,18E+09 | 68,527 | 0 | 323,31 | 377 |
| 9,04E+07 | 8,63E+07 | 1,01E+08 | 8,95E+07 | 83,886 | 0 | 136,43 | 64 |
| 3,43E+08 | 2,89E+08 | 3,31E+08 | 3,28E+08 | 48,936 | 0 | 81,724 | 71 |
| 4,18E+06 | 4,22E+06 | 1,19E+06 | 3,89E+06 | 32,167 | 0,000146 | 3,1025 | 5 |
| 2,59E+07 | 2,43E+07 | 2,87E+07 | 2,55E+07 | 89,085 | 0 | 13,177 | 40 |
| 9,35E+07 | 9,65E+07 | 8,71E+07 | 9,75E+07 | 66,662 | 0 | 25,632 | 53 |
| 4,45E+06 | 4,97E+06 | 6,46E+06 | 3,08E+06 | 36,03 | 0,002098 | 2,1659 | 6 |
| 2,27E+08 | 2,65E+08 | 3,16E+08 | 2,94E+08 | 37,623 | 0 | 28,791 | 47 |
| 5,35E+07 | 6,85E+07 | 4,21E+07 | 3,67E+07 | 57,696 | 0 | 35,034 | 31 |
| 3,09E+09 | 3,34E+09 | 2,91E+09 | 2,66E+09 | 42,004 | 0 | 21,907 | 51 |
| 2,95E+06 | 1,26E+07 | 8,27E+06 | 9,91E+06 | 182,56 | 0,000146 | 3,0912 | 8 |
| 3,13E+06 | 5,18E+06 | 2,19E+06 | 3,15E+06 | 25,704 | 0,001546 | 2,2942 | 3 |
| 2,41E+06 | 6,75E+06 | 3,44E+06 | 8,17E+06 | 43,165 | 0,001272 | 2,4353 | 3 |
| 3,91E+06 | 8,81E+06 | 9,90E+06 | 1,96E+06 | 114,26 | 0,000149 | 3,5301 | 5 |
| 8,89E+07 | 1,08E+08 | 1,06E+08 | 1,25E+08 | 37,252 | 0 | 51,104 | 69 |
| 2,32E+08 | 2,37E+08 | 2,38E+08 | 2,35E+08 | 152,53 | 0 | 134,72 | 175 |
| 9,04E+07 | 9,14E+07 | 9,29E+07 | 9,05E+07 | 171,12 | 0 | 75,067 | 85 |
| 3,41E+06 | 3,14E+06 | 4,51E+06 | 8,38E+06 | 36,897 | 0,003697 | 1,8236 | 2 |
| 1,00E+08 | 9,16E+07 | 9,28E+07 | 8,61E+07 | 32,718 | 0 | 159,93 | 59 |
| 6,52E+07 | 8,12E+07 | 7,95E+07 | 7,97E+07 | 128,39 | 0 | 25,289 | 76 |
| 7,75E+08 | 7,82E+08 | 1,00E+09 | 9,00E+08 | 22,541 | 0 | 323,31 | 150 |
| 1,89E+08 | 2,11E+08 | 2,10E+08 | 2,09E+08 | 63,232 | 0 | 87,002 | 82 |
| 6,80E+08 | 6,60E+08 | 6,35E+08 | 5,88E+08 | 58,369 | 0 | 156,72 | 207 |
| 7,76E+06 | 6,92E+06 | 5,94E+06 | 6,66E+06 | 14,324 | 0,000153 | 4,194 | 4 |
| 5,17E+06 | 3,54E+06 | 4,08E+06 | 2,41E+06 | 31,564 | 0,00502 | 1,6645 | 2 |
| 5,75E+06 | 3,24E+06 | 7,58E+06 | 1,02E+07 | 26,093 | 0,000158 | 5,0739 | 5 |
| 2,62E+06 | 5,74E+06 | 4,98E+06 | 3,43E+06 | 20,178 | 0,005414 | 1,6333 | 7 |
| 8,26E+06 | 7,24E+06 | 1,08E+07 | 8,10E+06 | 151,57 | 0 | 8,4906 | 15 |
| 1,69E+07 | 2,75E+07 | 1,62E+07 | 1,70E+07 | 44,338 | 0 | 6,7473 | 22 |
| 4,53E+08 | 2,97E+08 | 2,95E+08 | 2,42E+08 | 85,616 | 0 | 323,31 | 84 |
| 9,05E+07 | 1,02E+08 | 7,08E+07 | 7,56E+07 | 20,302 | 0 | 8,3158 | 29 |
| 1,10E+07 | 6,20E+06 | 6,38E+06 | 7,12E+06 | 74,767 | 0,000435 | 2,9232 | 11 |
| 5,23E+06 | 2,80E+06 | 4,12E+06 | 3,71E+06 | 115,79 | 0,000155 | 4,5628 | 6 |
| 8,16E+07 | 8,42E+07 | 8,81E+07 | 9,39E+07 | 17,876 | 0 | 12,879 | 30 |
| 3,22E+07 | 2,67E+07 | 2,86E+07 | 3,28E+07 | 32,014 | 0 | 5,9798 | 17 |
| 3,21E+09 | 2,87E+09 | 3,04E+09 | 2,66E+09 | 52,673 | 0 | 323,31 | 268 |
| 1,30E+07 | 4,07E+06 | 5,60E+06 | 8,82E+06 | 74,22 | 0 | 5,6719 | 11 |
| 2,77E+07 | 3,07E+07 | 2,55E+07 | 2,46E+07 | 21,266 | 0 | 11,536 | 19 |

| | | | | | | | |
|---|---|---|---|---|---|---|---|
| 3,84E+07 | 4,04E+07 | 3,69E+07 | 3,96E+07 | 171,75 | 0 | 13,003 | 33 |
| 3,38E+06 | 6,05E+06 | 3,89E+06 | 6,26E+06 | 52,172 | 0,000149 | 3,5035 | 11 |
| 2,19E+07 | 2,28E+07 | 2,43E+07 | 2,98E+07 | 22,96 | 0 | 54,8 | 20 |
| 1,23E+08 | 1,11E+08 | 9,77E+07 | 9,68E+07 | 112,9 | 0 | 48,561 | 95 |
| 1,03E+08 | 1,06E+08 | 1,11E+08 | 1,13E+08 | 60,341 | 0 | 162,59 | 76 |
| 4,28E+08 | 2,93E+08 | 3,00E+08 | 2,40E+08 | 22,705 | 0 | 323,31 | 134 |
| 9,48E+07 | 6,90E+07 | 6,62E+07 | 7,73E+07 | 54,395 | 0 | 51,989 | 55 |
| 2,47E+08 | 2,33E+08 | 2,27E+08 | 2,18E+08 | 35,975 | 0 | 235,91 | 93 |
| 8,57E+07 | 1,18E+08 | 1,31E+08 | 1,38E+08 | 30,537 | 0 | 59,547 | 57 |
| 3,02E+07 | 3,75E+07 | 4,79E+07 | 4,62E+07 | 15,236 | 0 | 7,1696 | 24 |
| 2,14E+08 | 1,93E+08 | 2,04E+08 | 2,00E+08 | 188,91 | 0 | 139,93 | 148 |
| 6,23E+07 | 6,14E+07 | 5,56E+07 | 7,76E+07 | 13,608 | 0 | 89,756 | 21 |
| 6,41E+07 | 2,86E+07 | 4,77E+07 | 7,92E+07 | 15,422 | 0 | 86,554 | 30 |
| 9,05E+06 | 5,47E+06 | 7,86E+06 | 8,40E+06 | 53,212 | 0,000147 | 3,2774 | 8 |
| 4,19E+07 | 4,70E+07 | 3,35E+07 | 3,07E+07 | 24,038 | 0 | 29,552 | 30 |
| 1,13E+08 | 1,16E+08 | 1,11E+08 | 1,25E+08 | 54,274 | 0 | 71,214 | 56 |
| 2,35E+08 | 3,12E+08 | 2,53E+08 | 2,55E+08 | 8,2355 | 0 | 27,179 | 121 |
| 1,31E+07 | 9,78E+06 | 9,73E+06 | 8,84E+06 | 12,858 | 0,000434 | 2,891 | 10 |
| 1,42E+07 | 9,04E+06 | 1,01E+07 | 9,10E+06 | 15,278 | 0,000156 | 4,6939 | 12 |
| 2,34E+08 | 2,42E+08 | 2,31E+08 | 2,47E+08 | 91,612 | 0 | 139,51 | 173 |
| 1,88E+08 | 1,83E+08 | 1,71E+08 | 1,73E+08 | 35,623 | 0 | 174,27 | 74 |
| 5,41E+06 | 7,76E+06 | 4,05E+06 | 3,25E+06 | 121 | 0,000147 | 3,2748 | 5 |
| 8,42E+07 | 8,22E+07 | 8,07E+07 | 8,25E+07 | 89,365 | 0 | 21,177 | 68 |
| 7,03E+06 | 1,06E+07 | 4,18E+06 | 3,90E+06 | 53,376 | 0,004369 | 1,7616 | 6 |
| 5,83E+06 | 1,42E+06 | 7,26E+06 | 4,33E+06 | 42,325 | 0,003176 | 1,951 | 6 |
| 5,72E+06 | 7,75E+06 | 9,10E+06 | 7,22E+06 | 34,817 | 0,009372 | 1,4382 | 9 |
| 2,02E+09 | 1,77E+09 | 1,69E+09 | 1,73E+09 | 16,604 | 0 | 323,31 | 180 |
| 2,35E+06 | 3,92E+06 | 1,31E+06 | 1,14E+06 | 138,1 | 0 | 8,1432 | 4 |
| 1,10E+07 | 1,93E+07 | 9,73E+06 | 1,03E+07 | 70,812 | 0 | 5,3794 | 18 |
| 1,38E+09 | 1,22E+09 | 1,40E+09 | 1,31E+09 | 13,17 | 0 | 255,85 | 166 |
| 1,70E+07 | 1,72E+07 | 2,00E+07 | 2,11E+07 | 81,477 | 0 | 15,265 | 27 |
| 2,17E+06 | 1,92E+06 | 1,74E+06 | 5,81E+06 | 36,947 | 0,005022 | 1,6729 | 2 |
| 5,61E+07 | 5,84E+07 | 5,42E+07 | 4,74E+07 | 56,77 | 0 | 56,212 | 36 |
| 2,49E+06 | 2,02E+06 | 7,79E+06 | 7,83E+06 | 103,3 | 0,000431 | 2,7756 | 7 |
| 1,98E+09 | 1,92E+09 | 1,78E+09 | 1,99E+09 | 37,063 | 0 | 323,31 | 286 |
| 1,31E+08 | 1,34E+08 | 1,24E+08 | 1,08E+08 | 116,45 | 0 | 79,251 | 105 |
| 9,41E+07 | 8,82E+07 | 8,54E+07 | 7,36E+07 | 33,289 | 0 | 110,55 | 46 |
| 2,69E+08 | 2,56E+08 | 2,88E+08 | 2,86E+08 | 90,059 | 0 | 90,356 | 147 |
| 2,78E+07 | 3,10E+07 | 3,37E+07 | 4,10E+07 | 61,115 | 0 | 8,6573 | 32 |
| 2,00E+08 | 2,35E+08 | 2,56E+08 | 2,56E+08 | 41,492 | 0 | 144,75 | 68 |
| 4,16E+08 | 5,04E+08 | 3,94E+08 | 4,57E+08 | 25,853 | 0 | 132,11 | 80 |
| 2,84E+07 | 3,28E+07 | 3,11E+07 | 2,72E+07 | 176,52 | 0 | 14,557 | 48 |
| 2,72E+06 | 3,45E+06 | 3,41E+06 | 2,07E+06 | 76,862 | 0,00141 | 2,3587 | 4 |
| 9,02E+06 | 1,35E+07 | 1,24E+07 | 9,38E+06 | 17,389 | 0 | 7,9275 | 9 |
| 2,61E+07 | 3,80E+07 | 3,02E+07 | 3,65E+07 | 37,39 | 0 | 10,964 | 24 |
| 2,14E+07 | 2,07E+07 | 2,79E+07 | 2,72E+07 | 36,313 | 0 | 6,258 | 22 |
| 4,83E+08 | 4,47E+08 | 4,34E+08 | 4,45E+08 | 49,223 | 0 | 323,31 | 154 |
| 3,28E+06 | 3,91E+06 | 2,27E+06 | 3,17E+06 | 9,5839 | 0,002911 | 2,0096 | 4 |
| 1,95E+07 | 1,05E+07 | 1,09E+07 | 1,24E+07 | 24,264 | 0 | 5,7051 | 16 |
| 2,92E+07 | 2,28E+07 | 2,35E+07 | 1,66E+07 | 94,41 | 0 | 10,332 | 21 |

| | | | | | | | |
|---|---|---|---|---|---|---|---|
| 8,27E+07 | 8,34E+07 | 7,70E+07 | 1,09E+08 | 45,58 | 0 | 10,085 | 22 |
| 4,33E+06 | 5,28E+06 | 1,70E+06 | 9,15E+06 | 107,72 | 0,000153 | 4,1634 | 5 |
| 3,35E+08 | 3,33E+08 | 3,37E+08 | 3,46E+08 | 95,125 | 0 | 323,31 | 147 |
| 8,85E+08 | 2,23E+08 | 1,58E+08 | 1,10E+09 | 26,558 | 0 | 12,801 | 14 |
| 2,29E+07 | 1,97E+07 | 1,71E+07 | 2,49E+07 | 57,368 | 0 | 8,6297 | 20 |
| 2,39E+07 | 2,15E+07 | 2,25E+07 | 2,01E+07 | 32,978 | 0 | 14,494 | 18 |
| 2,67E+06 | 3,35E+06 | 3,81E+06 | 2,08E+06 | 42,03 | 0,004367 | 1,7564 | 6 |
| 3,65E+06 | 8,21E+06 | 3,63E+06 | 2,50E+06 | 25,61 | 0,000151 | 3,7623 | 8 |
| 2,77E+08 | 3,31E+08 | 3,19E+08 | 3,19E+08 | 56,381 | 0 | 70,673 | 104 |
| 9,09E+06 | 1,13E+07 | 1,31E+07 | 1,39E+07 | 42,442 | 0 | 18,24 | 13 |
| 1,45E+08 | 1,94E+08 | 1,89E+08 | 1,89E+08 | 39,617 | 0 | 56,388 | 74 |
| 1,30E+08 | 1,20E+08 | 1,23E+08 | 1,21E+08 | 26,497 | 0,000157 | 4,9648 | 39 |
| 8,88E+07 | 8,85E+07 | 8,19E+07 | 6,70E+07 | 12,758 | 0 | 106,35 | 81 |
| 1,44E+07 | 1,86E+07 | 1,75E+07 | 2,11E+07 | 23,644 | 0 | 6,8245 | 16 |
| 3,27E+08 | 3,28E+08 | 3,69E+08 | 3,76E+08 | 30,149 | 0 | 100,46 | 79 |
| 1,12E+07 | 4,70E+06 | 4,13E+06 | 5,32E+06 | 20,633 | 0,000153 | 4,2025 | 3 |
| 5,50E+07 | 7,05E+07 | 7,34E+07 | 7,49E+07 | 44,422 | 0 | 84,464 | 42 |
| 1,43E+08 | 3,35E+07 | 1,25E+08 | 8,37E+07 | 15,142 | 0 | 221,47 | 19 |
| 5,23E+09 | 5,02E+09 | 5,23E+09 | 4,79E+09 | 46,109 | 0 | 323,31 | 485 |
| 9,32E+08 | 9,73E+08 | 9,94E+08 | 9,53E+08 | 164,42 | 0 | 323,31 | 322 |
| 1,78E+09 | 1,96E+09 | 2,02E+09 | 1,84E+09 | 29,546 | 0 | 233,43 | 248 |
| 2,38E+07 | 2,45E+07 | 2,23E+07 | 1,82E+07 | 33,988 | 0 | 7,7411 | 20 |
| 2,94E+06 | 7,92E+06 | 4,36E+06 | 5,20E+06 | 69,147 | 0,000158 | 5,0385 | 10 |
| 6,70E+07 | 7,27E+07 | 7,77E+07 | 7,26E+07 | 63,202 | 0 | 95,42 | 71 |
| 7,21E+06 | 2,20E+06 | 1,46E+07 | 1,53E+06 | 17,554 | 0,000434 | 2,9129 | 4 |
| 1,02E+08 | 1,03E+08 | 1,02E+08 | 1,12E+08 | 59,606 | 0 | 129,18 | 44 |
| 3,11E+07 | 2,34E+07 | 2,56E+07 | 1,92E+07 | 61,062 | 0 | 42,767 | 9 |
| 2,79E+09 | 2,88E+09 | 2,63E+09 | 2,79E+09 | 34,877 | 0 | 323,31 | 252 |
| 1,75E+07 | 1,59E+07 | 1,85E+07 | 1,33E+07 | 82,975 | 0 | 7,7501 | 20 |
| 5,19E+07 | 6,45E+07 | 4,79E+07 | 5,90E+07 | 43,408 | 0 | 37,694 | 47 |
| 3,40E+06 | 2,95E+06 | 1,74E+06 | 2,06E+06 | 56,44 | 0,002911 | 2,0109 | 7 |
| 9,21E+07 | 8,33E+07 | 1,07E+08 | 7,58E+07 | 48,166 | 0 | 40,431 | 21 |
| 1,98E+06 | 4,05E+06 | 3,61E+06 | 1,10E+06 | 30,939 | 0,003447 | 1,9319 | 2 |
| 4,71E+08 | 3,13E+08 | 3,63E+08 | 3,52E+08 | 16,284 | 0 | 9,9257 | 18 |
| 2,40E+06 | 1,34E+06 | 3,68E+07 | 2,89E+06 | 227,03 | 0 | 249,28 | 60 |
| 1,55E+07 | 1,62E+07 | 1,33E+07 | 1,69E+07 | 24,56 | 0 | 6,2514 | 14 |
| 1,91E+07 | 1,94E+07 | 1,47E+07 | 1,69E+07 | 19,519 | 0 | 28,289 | 15 |
| 1,11E+07 | 9,66E+06 | 1,13E+07 | 8,94E+06 | 69,926 | 0,000429 | 2,7227 | 8 |
| 1,99E+07 | 2,58E+07 | 1,89E+07 | 2,26E+07 | 67,272 | 0 | 7,2868 | 25 |
| 6,25E+07 | 7,61E+07 | 7,86E+07 | 8,92E+07 | 86,766 | 0 | 59,381 | 62 |
| 8,91E+05 | 5,11E+06 | 3,97E+06 | 3,37E+06 | 32,739 | 0 | 5,8602 | 3 |
| 9,74E+06 | 1,11E+07 | 1,10E+07 | 8,64E+06 | 41,614 | 0,005815 | 1,6129 | 6 |
| 1,33E+08 | 1,22E+08 | 1,41E+08 | 1,42E+08 | 107,58 | 0,003962 | 1,779 | 10 |
| 4,35E+07 | 4,97E+07 | 5,43E+07 | 6,21E+07 | 55,839 | 0 | 48,906 | 48 |
| 1,66E+08 | 1,73E+08 | 1,77E+08 | 1,62E+08 | 34,995 | 0 | 65,982 | 93 |
| 5,51E+06 | 3,89E+06 | 2,92E+06 | 1,18E+06 | 86,889 | 0,000432 | 2,8193 | 6 |
| 1,52E+09 | 1,55E+09 | 1,47E+09 | 1,54E+09 | 63,53 | 0 | 323,31 | 250 |
| 3,98E+06 | 1,11E+07 | 4,80E+06 | 9,61E+06 | 40,619 | 0,00043 | 2,7658 | 3 |
| 2,51E+06 | 2,44E+06 | 4,38E+06 | 6,21E+06 | 17,355 | 0,002096 | 2,1555 | 4 |
| 8,11E+06 | 7,77E+06 | 1,03E+07 | 1,23E+07 | 51,216 | 0,000157 | 4,9012 | 11 |

| | | | | | | | |
|---|---|---|---|---|---|---|---|
| 2,29E+09 | 2,25E+09 | 2,45E+09 | 2,26E+09 | 26,411 | 0 | 323,31 | 272 |
| 1,01E+07 | 1,02E+07 | 5,52E+06 | 1,20E+07 | 54,014 | 0,000149 | 3,5776 | 16 |
| 5,85E+09 | 5,33E+09 | 5,94E+09 | 6,11E+09 | 13,373 | 0 | 71,545 | 165 |
| 7,36E+09 | 7,56E+09 | 7,39E+09 | 8,38E+09 | 16,445 | 0 | 63,607 | 346 |
| 6,39E+06 | 6,85E+06 | 7,09E+06 | 8,76E+06 | 20,769 | 0,000151 | 3,8833 | 12 |
| 1,53E+07 | 1,71E+07 | 1,57E+07 | 1,36E+07 | 124,84 | 0 | 19,052 | 31 |
| 7,20E+07 | 9,79E+07 | 9,52E+07 | 7,89E+07 | 41,928 | 0 | 54,743 | 78 |
| 1,21E+08 | 1,49E+08 | 1,43E+08 | 1,38E+08 | 111,24 | 0 | 55,248 | 99 |
| 4,91E+06 | 5,35E+06 | 5,22E+06 | 3,80E+06 | 58,154 | 0,000147 | 3,1617 | 7 |
| 2,41E+06 | 6,82E+06 | 5,68E+06 | 8,37E+06 | 120,83 | 0,00015 | 3,746 | 8 |
| 8,56E+06 | 9,34E+06 | 6,17E+06 | 7,66E+06 | 53,511 | 0,003963 | 1,7844 | 7 |
| 2,64E+07 | 2,88E+07 | 3,72E+07 | 3,11E+07 | 58,575 | 0 | 19,586 | 26 |
| 2,17E+08 | 1,90E+08 | 1,79E+08 | 1,60E+08 | 37,694 | 0 | 195,29 | 73 |
| 4,52E+07 | 4,37E+07 | 3,88E+07 | 4,21E+07 | 29,814 | 0 | 71,505 | 25 |
| 5,87E+07 | 6,09E+07 | 5,09E+07 | 6,32E+07 | 28,886 | 0 | 44,41 | 25 |
| 3,57E+07 | 1,32E+07 | 2,29E+07 | 1,79E+07 | 82,361 | 0 | 29,611 | 8 |
| 3,80E+06 | 4,07E+06 | 5,43E+06 | 5,25E+06 | 104,47 | 0,000154 | 4,3563 | 10 |
| 4,37E+06 | 2,70E+06 | 4,06E+06 | 5,33E+06 | 35,694 | 0,002097 | 2,1604 | 3 |
| 3,46E+07 | 3,26E+07 | 3,16E+07 | 2,61E+07 | 87,21 | 0 | 36,378 | 44 |
| 5,26E+06 | 1,25E+07 | 3,51E+06 | 9,19E+06 | 28,78 | 0 | 6,2065 | 6 |
| 2,88E+07 | 3,40E+07 | 2,56E+07 | 3,82E+07 | 45,932 | 0 | 7,9454 | 26 |
| 3,83E+06 | 8,22E+06 | 4,34E+06 | 6,39E+06 | 38,301 | 0,000432 | 2,8063 | 16 |
| 1,75E+08 | 1,57E+08 | 1,66E+08 | 1,33E+08 | 45,701 | 0 | 90,084 | 65 |
| 1,02E+08 | 1,08E+08 | 9,87E+07 | 1,03E+08 | 76,631 | 0 | 56,468 | 64 |
| 8,70E+06 | 5,29E+06 | 9,85E+06 | 6,86E+06 | 63,66 | 0 | 12,938 | 6 |
| 6,52E+06 | 4,52E+06 | 9,66E+06 | 1,06E+07 | 26,96 | 0,000436 | 3,0104 | 12 |
| 3,81E+07 | 3,07E+07 | 3,70E+07 | 4,09E+07 | 129,02 | 0 | 11,868 | 38 |
| 7,23E+06 | 6,64E+06 | 4,57E+06 | 6,47E+06 | 23,058 | 0 | 7,5245 | 15 |
| 3,56E+08 | 2,30E+08 | 2,11E+08 | 2,06E+08 | 16,723 | 0 | 194,92 | 38 |
| 6,97E+07 | 9,37E+07 | 1,38E+08 | 1,27E+08 | 21,649 | 0 | 51,259 | 48 |
| 8,58E+06 | 1,02E+07 | 5,23E+06 | 7,75E+06 | 57,026 | 0 | 6,5286 | 13 |
| 2,31E+07 | 1,85E+07 | 1,90E+07 | 1,61E+07 | 66,855 | 0 | 10,309 | 25 |
| 5,76E+08 | 6,30E+08 | 5,62E+08 | 6,23E+08 | 91,926 | 0 | 288,42 | 226 |
| 1,55E+09 | 1,58E+09 | 1,65E+09 | 1,71E+09 | 58,336 | 0 | 323,31 | 213 |
| 9,43E+07 | 1,11E+08 | 1,08E+08 | 1,20E+08 | 85,648 | 0 | 16,064 | 27 |
| 6,42E+06 | 3,82E+06 | 3,68E+06 | 6,44E+06 | 50,66 | 0,006078 | 1,5983 | 3 |
| 2,07E+08 | 2,10E+08 | 2,26E+08 | 2,17E+08 | 169,43 | 0 | 262,79 | 141 |
| 2,85E+08 | 2,89E+08 | 2,39E+08 | 2,63E+08 | 62,368 | 0 | 107,02 | 111 |
| 1,63E+07 | 1,26E+07 | 1,23E+07 | 1,35E+07 | 33,228 | 0,001686 | 2,2884 | 10 |
| 4,60E+07 | 5,64E+07 | 5,27E+07 | 4,55E+07 | 11,549 | 0 | 17,539 | 37 |
| 4,91E+07 | 3,86E+07 | 5,53E+07 | 5,51E+07 | 73,624 | 0 | 35,636 | 56 |
| 3,68E+06 | 4,70E+06 | 3,64E+06 | 6,55E+06 | 21,011 | 0 | 10,717 | 7 |
| 4,08E+06 | 4,73E+06 | 4,71E+06 | 2,82E+06 | 32,517 | 0,006878 | 1,5662 | 1 |
| 1,77E+06 | 2,54E+06 | 2,47E+06 | 2,68E+06 | 35,063 | 0,000152 | 4,0702 | 5 |
| 6,06E+07 | 6,55E+07 | 6,39E+07 | 5,41E+07 | 264,28 | 0 | 35,228 | 56 |
| 1,57E+08 | 1,40E+08 | 1,49E+08 | 1,37E+08 | 86,525 | 0 | 254,13 | 96 |
| 3,67E+06 | 3,46E+06 | 4,95E+06 | 4,23E+06 | 172,41 | 0,006745 | 1,5733 | 8 |
| 7,89E+05 | 1,63E+06 | 2,02E+06 | 1,88E+06 | 49,484 | 0,003178 | 1,9613 | 5 |
| 3,16E+09 | 3,08E+09 | 3,23E+09 | 3,16E+09 | 76,723 | 0 | 323,31 | 444 |
| 1,61E+08 | 1,46E+08 | 1,42E+08 | 1,55E+08 | 42,795 | 0 | 140,54 | 71 |

| | | | | | | | |
|---|---|---|---|---|---|---|---|
| 1,08E+07 | 1,26E+07 | 1,44E+07 | 1,44E+07 | 212,75 | 0 | 15,882 | 26 |
| 2,55E+09 | 2,03E+09 | 1,83E+09 | 1,82E+09 | 24,693 | 0 | 323,31 | 136 |
| 6,64E+06 | 4,15E+06 | 4,21E+06 | 3,57E+06 | 92,262 | 0 | 5,6959 | 10 |
| 1,11E+09 | 9,40E+08 | 1,10E+09 | 9,15E+08 | 18,562 | 0 | 226,03 | 89 |
| 4,32E+07 | 6,23E+07 | 5,74E+07 | 5,48E+07 | 66,308 | 0 | 39,36 | 60 |
| 1,56E+08 | 8,68E+07 | 7,43E+07 | 7,35E+07 | 12,459 | 0 | 8,716 | 9 |
| 1,29E+07 | 1,68E+07 | 1,16E+07 | 1,45E+07 | 94,078 | 0 | 7,2308 | 24 |
| 2,36E+08 | 2,48E+08 | 2,41E+08 | 2,20E+08 | 95,044 | 0 | 323,31 | 128 |
| 5,89E+07 | 6,86E+07 | 7,20E+07 | 7,99E+07 | 87,693 | 0 | 51,202 | 58 |
| 9,71E+08 | 9,83E+08 | 1,06E+09 | 1,07E+09 | 61,249 | 0 | 323,31 | 125 |
| 2,92E+08 | 3,15E+08 | 3,05E+08 | 3,23E+08 | 48,376 | 0 | 60,796 | 118 |
| 1,50E+08 | 1,44E+08 | 1,64E+08 | 1,75E+08 | 39,917 | 0 | 69,689 | 49 |
| 1,99E+07 | 2,17E+07 | 1,75E+07 | 1,79E+07 | 25,557 | 0 | 7,2266 | 10 |
| 2,29E+08 | 2,51E+08 | 2,61E+08 | 2,36E+08 | 14,015 | 0 | 18,674 | 50 |
| 8,58E+07 | 7,42E+07 | 8,64E+07 | 7,29E+07 | 29,298 | 0 | 18,215 | 40 |
| 7,31E+08 | 7,04E+08 | 7,31E+08 | 7,63E+08 | 46,852 | 0 | 323,31 | 189 |
| 4,38E+07 | 3,94E+07 | 4,23E+07 | 4,68E+07 | 84,728 | 0 | 15,798 | 44 |
| 3,13E+08 | 3,20E+08 | 3,04E+08 | 3,14E+08 | 61,174 | 0 | 85,596 | 124 |
| 5,48E+09 | 5,57E+09 | 5,90E+09 | 5,91E+09 | 47,411 | 0 | 323,31 | 552 |
| 4,76E+06 | 2,42E+06 | 3,02E+06 | 2,40E+06 | 15,777 | 0,000156 | 4,6204 | 7 |
| 1,14E+07 | 9,44E+06 | 1,15E+07 | 8,17E+06 | 52,156 | 0 | 24,77 | 16 |
| 1,91E+07 | 2,03E+07 | 1,81E+07 | 2,43E+07 | 33,24 | 0 | 5,9951 | 16 |
| 4,14E+08 | 4,22E+08 | 4,14E+08 | 4,26E+08 | 108,34 | 0 | 204,87 | 242 |
| 8,37E+06 | 9,30E+06 | 9,70E+06 | 1,41E+07 | 28,05 | 0 | 7,4846 | 15 |
| 5,43E+09 | 4,67E+09 | 6,11E+09 | 5,43E+09 | 29,885 | 0 | 323,31 | 570 |
| 1,37E+08 | 5,47E+07 | 1,29E+08 | 7,18E+07 | 62,129 | 0 | 235,51 | 57 |
| 7,67E+07 | 8,58E+07 | 7,40E+07 | 7,87E+07 | 38,444 | 0 | 51,935 | 69 |
| 1,18E+08 | 9,97E+07 | 8,68E+07 | 1,10E+08 | 41,139 | 0 | 89,848 | 48 |
| 3,11E+07 | 3,53E+07 | 4,08E+07 | 3,37E+07 | 60,575 | 0 | 18,704 | 30 |
| 4,47E+07 | 5,80E+07 | 4,73E+07 | 5,11E+07 | 18,901 | 0 | 8,7675 | 24 |
| 4,57E+06 | 6,57E+06 | 1,03E+07 | 1,04E+07 | 161,21 | 0 | 9,1252 | 24 |
| 9,66E+08 | 9,65E+08 | 7,63E+08 | 8,54E+08 | 25,053 | 0 | 285,3 | 146 |
| 1,01E+07 | 1,05E+07 | 1,14E+07 | 1,22E+07 | 31,872 | 0 | 9,0681 | 20 |
| 2,07E+08 | 2,04E+08 | 2,15E+08 | 2,14E+08 | 21,423 | 0 | 69,281 | 60 |
| 1,88E+07 | 2,23E+07 | 2,14E+07 | 2,71E+07 | 44,845 | 0 | 29,239 | 32 |
| 1,25E+07 | 1,98E+07 | 1,85E+07 | 1,59E+07 | 22,113 | 0 | 6,8336 | 22 |
| 4,88E+06 | 5,30E+06 | 4,57E+06 | 4,83E+06 | 80,44 | 0 | 5,7253 | 7 |
| 4,30E+06 | 5,66E+06 | 6,43E+06 | 1,18E+06 | 30,337 | 0 | 51,756 | 3 |
| 1,34E+08 | 1,48E+08 | 1,58E+08 | 1,52E+08 | 52,213 | 0 | 59,452 | 72 |
| 1,44E+06 | 1,15E+06 | 4,89E+06 | 3,61E+06 | 76,433 | 0,001824 | 2,2438 | 3 |
| 2,33E+06 | 3,61E+06 | 3,84E+06 | 1,59E+06 | 83,508 | 0,002914 | 2,0223 | 7 |
| 5,04E+06 | 4,21E+06 | 4,38E+06 | 6,96E+06 | 40,96 | 0 | 5,6534 | 7 |
| 2,24E+06 | 2,73E+06 | 1,97E+06 | 1,40E+06 | 35,693 | 0 | 26,23 | 6 |
| 6,32E+07 | 7,65E+07 | 8,74E+07 | 6,57E+07 | 34,155 | 0 | 10,828 | 45 |
| 4,75E+06 | 1,33E+07 | 2,77E+06 | 5,11E+06 | 149,64 | 0,008853 | 1,4669 | 8 |
| 2,00E+07 | 2,29E+06 | 2,55E+07 | 2,65E+07 | 42,993 | 0,009749 | 1,4107 | 8 |
| 4,39E+06 | 1,45E+06 | 8,49E+06 | 3,38E+06 | 132,53 | 0,004364 | 1,7452 | 4 |
| 3,24E+07 | 1,39E+07 | 1,99E+07 | 2,38E+07 | 37,41 | 0,000436 | 2,9941 | 5 |
| 3,28E+08 | 3,66E+08 | 3,33E+08 | 3,09E+08 | 47,778 | 0 | 190,19 | 144 |
| 4,89E+08 | 5,31E+08 | 4,84E+08 | 5,40E+08 | 53,291 | 0 | 323,31 | 158 |

| | | | | | | | |
|---|---|---|---|---|---|---|---|
| 1,49E+07 | 1,03E+07 | 1,06E+07 | 9,21E+06 | 17,922 | 0 | 16,673 | 6 |
| 1,62E+08 | 1,34E+07 | 1,29E+07 | 1,43E+07 | 112,73 | 0 | 10,371 | 20 |
| 4,82E+08 | 3,96E+06 | 8,79E+07 | 5,30E+07 | 22,006 | 0 | 16,785 | 8 |
| 3,51E+07 | 2,95E+07 | 1,16E+07 | 2,80E+07 | 66,376 | 0 | 32,5 | 34 |
| 1,36E+09 | 1,37E+09 | 1,35E+09 | 1,28E+09 | 104,15 | 0 | 323,31 | 327 |
| 7,86E+06 | 5,06E+07 | 6,72E+07 | 1,12E+06 | 24,489 | 0 | 323,31 | 5 |
| 7,22E+06 | 6,06E+06 | 5,70E+06 | 4,22E+06 | 52,122 | 0,000156 | 4,7552 | 9 |
| 1,77E+07 | 4,36E+06 | 2,88E+06 | 2,85E+06 | 62,378 | 0 | 11,689 | 4 |
| 2,12E+07 | 1,79E+07 | 1,34E+07 | 1,31E+07 | 58,597 | 0 | 98,841 | 25 |
| 9,59E+07 | 1,04E+08 | 1,29E+08 | 9,80E+07 | 38,616 | 0 | 35,804 | 64 |
| 3,99E+07 | 4,36E+07 | 4,33E+07 | 5,46E+07 | 49,639 | 0 | 15,075 | 49 |
| 1,13E+07 | 5,20E+06 | 1,00E+07 | 3,84E+06 | 80,471 | 0,00608 | 1,6007 | 3 |
| 5,66E+08 | 6,03E+08 | 5,41E+08 | 5,45E+08 | 40,024 | 0 | 323,31 | 133 |
| 2,27E+07 | 2,72E+07 | 2,26E+07 | 2,35E+07 | 38,722 | 0 | 21,894 | 21 |
| 4,33E+06 | 3,81E+06 | 1,72E+06 | 4,60E+06 | 44,309 | 0,000149 | 3,4678 | 5 |
| 8,26E+06 | 4,53E+06 | 1,98E+06 | 5,42E+06 | 19,652 | 0,000429 | 2,6759 | 7 |
| 6,25E+06 | 5,18E+06 | 2,64E+06 | 5,18E+06 | 60,761 | 0,000158 | 5,0531 | 7 |
| 1,98E+09 | 1,84E+09 | 1,90E+09 | 1,65E+09 | 22,889 | 0 | 323,31 | 148 |
| 7,12E+06 | 1,91E+06 | 8,57E+06 | 8,21E+06 | 109,46 | 0,000152 | 4,0833 | 15 |
| 8,59E+08 | 8,28E+08 | 7,21E+08 | 6,85E+08 | 21,708 | 0 | 63,838 | 84 |
| 6,72E+06 | 7,38E+06 | 1,66E+06 | 1,91E+06 | 203,8 | 0,000435 | 2,9431 | 4 |
| 2,53E+07 | 8,67E+07 | 2,64E+07 | 2,53E+07 | 33,675 | 0 | 21,419 | 28 |
| 5,75E+08 | 6,87E+08 | 6,72E+08 | 5,80E+08 | 10 | 0 | 323,31 | 98 |
| 3,29E+08 | 3,34E+08 | 2,96E+08 | 3,03E+08 | 35,565 | 0 | 323,31 | 120 |
| 1,08E+07 | 1,07E+07 | 1,05E+07 | 1,26E+07 | 54,008 | 0 | 9,6391 | 12 |
| 2,24E+07 | 1,53E+07 | 1,66E+07 | 1,30E+07 | 19,967 | 0 | 10,242 | 15 |
| 4,00E+07 | 4,33E+07 | 1,96E+07 | 2,57E+07 | 22,942 | 0 | 252,17 | 17 |
| 1,77E+06 | 7,87E+06 | 4,63E+06 | 7,22E+06 | 17,908 | 0,001272 | 2,4324 | 7 |
| 2,12E+06 | 2,11E+06 | 8,64E+05 | 5,95E+05 | 176,66 | 0,000854 | 2,5794 | 5 |
| 5,97E+06 | 5,76E+05 | 8,55E+05 | 5,54E+06 | 75,128 | 0,000436 | 3,0008 | 3 |
| 7,32E+07 | 8,58E+07 | 8,66E+07 | 6,73E+07 | 47,33 | 0 | 75,304 | 36 |
| 6,20E+06 | 8,43E+06 | 6,08E+06 | 4,70E+06 | 24,452 | 0,00015 | 3,6245 | 10 |
| 1,31E+07 | 1,60E+07 | 1,36E+07 | 9,94E+06 | 95,224 | 0,000156 | 4,8061 | 20 |
| 3,97E+08 | 4,09E+08 | 3,91E+08 | 4,76E+08 | 10,834 | 0 | 183,03 | 60 |
| 1,23E+07 | 8,98E+06 | 1,02E+07 | 9,85E+06 | 105,11 | 0 | 5,724 | 19 |
| 7,90E+06 | 1,10E+07 | 9,64E+06 | 5,54E+06 | 133,77 | 0 | 8,2187 | 18 |
| 4,83E+06 | 6,18E+06 | 8,09E+06 | 6,92E+06 | 79,532 | 0 | 9,247 | 11 |
| 3,30E+06 | 2,45E+06 | 1,07E+07 | 1,06E+07 | 19,282 | 0,000993 | 2,4964 | 5 |
| 5,73E+07 | 6,61E+07 | 7,25E+07 | 8,67E+07 | 43,838 | 0 | 30,982 | 56 |
| 1,02E+07 | 5,58E+06 | 2,48E+07 | 1,51E+07 | 92,774 | 0 | 20,695 | 14 |
| 2,09E+07 | 2,31E+07 | 2,61E+07 | 2,17E+07 | 137,31 | 0 | 10,454 | 22 |
| 4,89E+08 | 5,33E+08 | 5,11E+08 | 5,31E+08 | 68,016 | 0 | 323,31 | 200 |
| 2,08E+07 | 2,29E+07 | 2,58E+07 | 2,37E+07 | 69,007 | 0 | 11,007 | 33 |
| 4,02E+07 | 3,28E+07 | 2,61E+07 | 2,71E+07 | 31,603 | 0,000158 | 5,1488 | 16 |
| 2,38E+08 | 2,21E+08 | 2,04E+08 | 2,35E+08 | 62,34 | 0 | 104,66 | 120 |
| 3,28E+08 | 3,57E+08 | 3,71E+08 | 3,79E+08 | 91,512 | 0 | 53,189 | 126 |
| 1,46E+08 | 1,30E+08 | 1,24E+08 | 1,15E+08 | 49,475 | 0 | 223,29 | 84 |
| 3,95E+06 | 4,62E+06 | 4,18E+06 | 3,90E+06 | 15,824 | 0 | 14,087 | 9 |
| 6,62E+08 | 4,42E+08 | 5,04E+08 | 5,65E+08 | 37,377 | 0 | 323,31 | 70 |
| 7,16E+07 | 6,81E+07 | 6,50E+07 | 4,92E+07 | 30,44 | 0 | 21,346 | 58 |

| | | | | | | | |
|---|---|---|---|---|---|---|---|
| 9,52E+07 | 1,01E+08 | 1,01E+08 | 7,86E+07 | 58,731 | 0,005289 | 1,6534 | 18 |
| 1,46E+08 | 1,06E+08 | 1,20E+08 | 1,13E+08 | 39,706 | 0 | 187,97 | 44 |
| 1,02E+09 | 9,90E+08 | 1,07E+09 | 1,09E+09 | 51,991 | 0 | 281,12 | 188 |
| 6,99E+07 | 5,88E+07 | 6,36E+07 | 6,45E+07 | 65,961 | 0 | 37,937 | 48 |
| 2,81E+06 | 2,70E+06 | 3,68E+06 | 2,53E+06 | 29,573 | 0,000158 | 5,1596 | 8 |
| 1,15E+08 | 1,14E+08 | 1,15E+08 | 1,13E+08 | 54,959 | 0 | 171,83 | 46 |
| 2,62E+07 | 2,51E+07 | 3,06E+07 | 3,32E+07 | 65,288 | 0 | 10,548 | 33 |
| 7,19E+06 | 5,50E+06 | 7,99E+06 | 5,93E+06 | 39,967 | 0,000159 | 5,1862 | 8 |
| 3,79E+08 | 3,75E+08 | 3,63E+08 | 3,84E+08 | 53,93 | 0 | 136,45 | 124 |
| 5,45E+06 | 3,39E+06 | 1,05E+07 | 7,72E+06 | 12,274 | 0 | 239,63 | 6 |
| 2,80E+07 | 7,53E+06 | 2,40E+07 | 2,87E+07 | 191,97 | 0 | 24,041 | 27 |
| 3,41E+07 | 4,46E+07 | 5,01E+07 | 4,35E+07 | 23,231 | 0 | 8,5085 | 27 |
| 7,08E+06 | 8,55E+05 | 1,96E+06 | 7,40E+06 | 39,968 | 0,003573 | 1,8919 | 4 |
| 2,77E+07 | 2,88E+07 | 2,64E+07 | 3,20E+07 | 132,62 | 0 | 25,03 | 45 |
| 7,16E+06 | 3,55E+06 | 6,33E+06 | 5,34E+06 | 8,496 | 0 | 14,182 | 4 |
| 3,81E+07 | 8,17E+07 | 8,86E+07 | 2,14E+07 | 16,46 | 0 | 192,81 | 23 |
| 3,30E+08 | 3,37E+08 | 2,92E+08 | 2,72E+08 | 64,958 | 0 | 323,31 | 67 |
| 3,23E+08 | 3,10E+08 | 2,87E+08 | 2,48E+08 | 47,832 | 0 | 311,01 | 126 |
| 1,08E+09 | 1,11E+09 | 1,18E+09 | 1,08E+09 | 17,861 | 0 | 323,31 | 141 |
| 1,23E+08 | 1,25E+08 | 1,21E+08 | 1,23E+08 | 75,363 | 0 | 101,96 | 92 |
| 9,20E+07 | 9,75E+07 | 9,88E+07 | 9,21E+07 | 29,05 | 0 | 103,39 | 59 |
| 4,41E+06 | 3,65E+06 | 3,18E+06 | 2,39E+06 | 50,175 | 0,0037 | 1,8394 | 3 |
| 8,88E+07 | 1,03E+08 | 9,57E+07 | 1,03E+08 | 133,84 | 0 | 27,995 | 50 |
| 4,61E+07 | 5,08E+07 | 3,79E+07 | 4,00E+07 | 55,384 | 0 | 12,94 | 37 |
| 1,01E+07 | 5,26E+06 | 3,45E+06 | 4,57E+06 | 38,56 | 0,00015 | 3,6964 | 6 |
| 3,80E+07 | 1,93E+07 | 3,36E+07 | 2,95E+07 | 65,056 | 0 | 6,8646 | 20 |
| 4,65E+06 | 4,42E+06 | 5,80E+06 | 2,54E+06 | 41,635 | 0,002096 | 2,1562 | 2 |
| 2,61E+08 | 3,71E+08 | 3,72E+08 | 3,92E+08 | 20,451 | 0 | 254,33 | 116 |
| 1,89E+06 | 3,08E+06 | 3,35E+06 | 3,99E+06 | 10,642 | 0,002781 | 2,0583 | 6 |
| 3,43E+07 | 4,95E+07 | 4,61E+07 | 5,00E+07 | 20,358 | 0 | 27,223 | 26 |
| 4,29E+06 | 1,05E+07 | 2,57E+06 | 4,86E+06 | 39,452 | 0,000157 | 4,9154 | 7 |
| 1,21E+08 | 1,12E+08 | 1,28E+08 | 8,85E+07 | 42,158 | 0 | 95,631 | 28 |
| 6,24E+07 | 7,49E+07 | 7,43E+07 | 5,61E+07 | 83,603 | 0 | 9,0545 | 32 |
| 1,01E+08 | 9,23E+07 | 9,41E+07 | 7,79E+07 | 68,079 | 0 | 148,94 | 63 |
| 2,21E+08 | 2,60E+08 | 2,39E+08 | 2,30E+08 | 88,066 | 0 | 85,739 | 161 |
| 2,02E+07 | 1,78E+07 | 1,82E+07 | 1,74E+07 | 31,61 | 0,000158 | 5,0676 | 19 |
| 3,17E+06 | 4,68E+06 | 3,34E+06 | 2,92E+06 | 17,875 | 0,000429 | 2,6835 | 8 |
| 3,19E+06 | 2,58E+06 | 4,03E+06 | 2,08E+06 | 67,733 | 0,000147 | 3,1986 | 5 |
| 5,19E+07 | 4,58E+07 | 4,45E+07 | 4,33E+07 | 25,263 | 0 | 33,084 | 27 |
| 5,68E+07 | 5,90E+07 | 5,27E+07 | 4,23E+07 | 179,83 | 0 | 20,281 | 46 |
| 2,81E+08 | 3,09E+08 | 2,99E+08 | 2,69E+08 | 40,603 | 0 | 136,18 | 83 |
| 1,60E+07 | 2,12E+07 | 1,60E+07 | 2,15E+07 | 28,967 | 0 | 8,7752 | 12 |
| 1,32E+08 | 1,66E+08 | 1,61E+08 | 1,67E+08 | 23,366 | 0 | 13,072 | 38 |
| 4,11E+08 | 3,50E+08 | 4,20E+08 | 4,03E+08 | 75,199 | 0 | 49,196 | 109 |
| 4,36E+06 | 4,05E+06 | 5,46E+06 | 6,35E+06 | 45,28 | 0,000147 | 3,1966 | 13 |
| 8,53E+08 | 8,77E+08 | 9,12E+08 | 9,33E+08 | 107,66 | 0 | 323,31 | 256 |
| 1,14E+08 | 1,23E+08 | 1,33E+08 | 1,21E+08 | 11,642 | 0 | 10,389 | 18 |
| 5,40E+08 | 6,76E+08 | 6,03E+08 | 5,82E+08 | 21,867 | 0 | 141,28 | 70 |
| 2,59E+09 | 1,80E+09 | 2,00E+09 | 1,89E+09 | 7,8409 | 0 | 107,37 | 217 |
| 5,02E+06 | 5,16E+06 | 3,90E+06 | 5,79E+06 | 24,187 | 0,000151 | 3,8235 | 9 |

| | | | | | | | |
|---|---|---|---|---|---|---|---|
| 8,19E+06 | 9,57E+06 | 9,28E+06 | 1,70E+06 | 18,787 | 0,006079 | 1,5996 | 4 |
| 1,72E+07 | 2,02E+07 | 1,14E+07 | 1,85E+07 | 47,342 | 0 | 28,691 | 19 |
| 1,86E+09 | 1,28E+09 | 1,67E+09 | 1,75E+09 | 12,259 | 0 | 323,31 | 39 |
| 1,68E+08 | 1,80E+08 | 1,67E+08 | 1,49E+08 | 103,13 | 0 | 102,7 | 116 |
| 5,88E+06 | 7,56E+06 | 5,45E+06 | 6,53E+06 | 33,57 | 0,000151 | 3,7983 | 6 |
| 1,15E+08 | 1,42E+08 | 7,42E+07 | 8,98E+07 | 28,713 | 0 | 171 | 42 |
| 3,51E+07 | 2,95E+07 | 2,35E+07 | 2,69E+07 | 24,774 | 0 | 148,96 | 14 |
| 7,49E+06 | 4,22E+06 | 3,67E+06 | 7,00E+06 | 40,693 | 0,005026 | 1,6854 | 7 |
| 2,10E+08 | 9,74E+07 | 4,06E+08 | 2,36E+08 | 66,017 | 0 | 323,31 | 84 |
| 3,18E+06 | 2,55E+06 | 2,44E+06 | 4,09E+06 | 45,54 | 0,000153 | 4,0924 | 7 |
| 1,00E+07 | 3,28E+06 | 1,35E+07 | 2,08E+07 | 25,079 | 0 | 24,726 | 6 |
| 4,82E+06 | 1,81E+06 | 5,80E+06 | 5,39E+06 | 34,257 | 0,008202 | 1,5077 | 4 |
| 1,05E+07 | 1,05E+07 | 9,65E+06 | 7,70E+06 | 25,761 | 0,000153 | 4,2297 | 7 |
| 7,31E+06 | 1,36E+06 | 5,40E+06 | 6,21E+06 | 43,371 | 0,005025 | 1,6826 | 6 |
| 2,98E+09 | 3,06E+09 | 3,34E+09 | 3,57E+09 | 126,85 | 0 | 323,31 | 557 |
| 3,47E+06 | 3,76E+06 | 4,81E+06 | 5,87E+06 | 63,823 | 0,008462 | 1,485 | 8 |
| 8,61E+06 | 3,83E+06 | 6,16E+06 | 4,71E+06 | 9,4226 | 0 | 9,3671 | 13 |
| 4,78E+07 | 4,86E+07 | 6,09E+07 | 6,20E+07 | 111,9 | 0 | 27,333 | 65 |
| 3,08E+06 | 3,95E+06 | 1,59E+06 | 3,61E+06 | 147,41 | 0,001686 | 2,2762 | 3 |
| 3,26E+06 | 1,07E+07 | 1,16E+07 | 2,08E+06 | 89,526 | 0,007146 | 1,5523 | 6 |
| 9,03E+07 | 1,02E+08 | 1,15E+08 | 1,32E+08 | 49,369 | 0 | 38,461 | 54 |
| 3,93E+07 | 9,65E+07 | 8,07E+06 | 3,76E+07 | 69,612 | 0,000157 | 4,8566 | 9 |
| 8,23E+07 | 7,04E+07 | 5,74E+07 | 5,19E+07 | 18,733 | 0 | 28,861 | 59 |
| 1,61E+07 | 2,11E+06 | 1,19E+07 | 2,10E+07 | 62,953 | 0,006611 | 1,5765 | 8 |
| 3,42E+06 | 3,99E+06 | 5,64E+06 | 5,28E+06 | 45,588 | 0,009488 | 1,4199 | 5 |
| 4,93E+06 | 2,89E+06 | 2,89E+06 | 6,66E+06 | 253,91 | 0,00043 | 2,7525 | 7 |
| 3,80E+08 | 4,27E+08 | 4,53E+08 | 5,27E+08 | 39,025 | 0 | 165,7 | 37 |
| 2,44E+08 | 2,75E+08 | 2,57E+08 | 2,50E+08 | 15,242 | 0 | 44,187 | 45 |
| 1,15E+07 | 1,62E+07 | 1,43E+07 | 1,42E+07 | 16,275 | 0,002912 | 2,0158 | 19 |
| 6,37E+07 | 6,93E+07 | 7,44E+07 | 7,19E+07 | 53,57 | 0 | 9,2777 | 42 |
| 1,41E+08 | 1,74E+08 | 1,56E+08 | 1,70E+08 | 20,68 | 0 | 44,509 | 77 |
| 2,48E+08 | 2,78E+08 | 2,63E+08 | 2,65E+08 | 113,52 | 0 | 305,21 | 168 |
| 5,15E+06 | 4,12E+06 | 4,27E+06 | 4,77E+06 | 43,081 | 0,005416 | 1,6387 | 3 |
| 7,10E+06 | 6,83E+06 | 8,57E+06 | 1,21E+07 | 38,509 | 0 | 5,8535 | 15 |
| 9,00E+08 | 1,08E+09 | 6,65E+08 | 7,42E+08 | 25,926 | 0 | 323,31 | 148 |
| 3,91E+07 | 3,25E+07 | 2,98E+07 | 2,52E+07 | 35,298 | 0 | 17,311 | 26 |
| 9,25E+06 | 9,46E+06 | 1,52E+07 | 1,23E+07 | 67,83 | 0 | 9,6348 | 21 |
| 4,79E+08 | 3,08E+08 | 4,05E+08 | 2,75E+08 | 45,024 | 0 | 323,31 | 47 |
| 2,46E+07 | 2,42E+07 | 2,37E+07 | 2,41E+07 | 70,222 | 0 | 10,436 | 36 |
| 4,37E+08 | 4,07E+08 | 4,10E+08 | 4,05E+08 | 90,582 | 0 | 323,31 | 131 |
| 5,69E+06 | 4,97E+06 | 8,16E+06 | 1,13E+07 | 85,927 | 0 | 7,4755 | 15 |
| 5,89E+07 | 5,12E+07 | 7,28E+07 | 6,96E+07 | 23,466 | 0 | 180,32 | 169 |
| 1,32E+08 | 1,21E+08 | 1,16E+08 | 1,19E+08 | 128,33 | 0 | 152,58 | 84 |
| 2,14E+06 | 4,05E+06 | 5,41E+06 | 2,40E+06 | 144,51 | 0,00502 | 1,6634 | 5 |
| 4,70E+07 | 4,48E+07 | 4,08E+07 | 4,03E+07 | 12,326 | 0 | 10,202 | 25 |
| 6,08E+08 | 6,12E+08 | 6,68E+08 | 6,31E+08 | 59,43 | 0 | 230,64 | 144 |
| 1,52E+07 | 1,38E+07 | 1,55E+07 | 1,19E+07 | 55,028 | 0 | 18,186 | 24 |
| 4,60E+06 | 4,84E+06 | 3,34E+06 | 4,17E+06 | 115,45 | 0,000149 | 3,4764 | 9 |
| 5,33E+07 | 4,43E+07 | 4,43E+07 | 3,49E+07 | 17,312 | 0,000292 | 3,0671 | 15 |
| 4,13E+07 | 5,06E+07 | 4,13E+06 | 7,28E+06 | 50,575 | 0,008206 | 1,5118 | 3 |

| | | | | | | | |
|---|---|---|---|---|---|---|---|
| 9,23E+08 | 7,90E+08 | 8,68E+08 | 8,41E+08 | 100,11 | 0 | 323,31 | 284 |
| 9,70E+07 | 7,29E+07 | 7,93E+07 | 7,40E+07 | 294,82 | 0 | 19,417 | 41 |
| 9,78E+06 | 1,14E+07 | 8,37E+06 | 1,08E+07 | 79,13 | 0 | 9,3853 | 13 |
| 4,17E+06 | 3,84E+06 | 3,16E+06 | 4,71E+06 | 70,268 | 0,000431 | 2,8005 | 6 |
| 9,38E+07 | 8,20E+07 | 9,42E+07 | 9,72E+07 | 86,016 | 0 | 32,133 | 56 |
| 2,51E+07 | 1,94E+07 | 1,88E+07 | 1,91E+07 | 54,917 | 0 | 6,1478 | 26 |
| 4,20E+06 | 3,74E+06 | 5,57E+06 | 3,53E+06 | 80,491 | 0,001549 | 2,3218 | 4 |
| 1,44E+09 | 1,24E+09 | 1,20E+09 | 1,14E+09 | 41,525 | 0 | 323,31 | 227 |
| 2,18E+08 | 1,97E+08 | 1,95E+08 | 1,99E+08 | 38,368 | 0 | 69 | 105 |
| 1,00E+09 | 9,11E+08 | 1,05E+09 | 1,09E+09 | 35,327 | 0 | 323,31 | 93 |
| 7,22E+06 | 8,84E+06 | 4,96E+06 | 6,21E+06 | 13,188 | 0,000149 | 3,4957 | 7 |
| 3,98E+08 | 3,16E+08 | 2,81E+08 | 4,44E+07 | 13,779 | 0,000155 | 4,516 | 54 |
| 7,96E+06 | 4,43E+06 | 2,70E+06 | 3,08E+06 | 46,864 | 0,000148 | 3,4004 | 8 |
| 2,66E+07 | 1,77E+07 | 2,71E+07 | 2,50E+07 | 17,823 | 0 | 6,4095 | 13 |
| 1,53E+07 | 1,66E+07 | 1,52E+07 | 1,88E+07 | 49,816 | 0 | 5,7307 | 21 |
| 6,73E+06 | 6,29E+06 | 5,33E+06 | 3,73E+06 | 46,886 | 0,00015 | 3,7268 | 9 |
| 1,33E+07 | 1,25E+07 | 1,14E+07 | 1,33E+07 | 75,769 | 0 | 8,2643 | 12 |
| 5,89E+09 | 5,85E+09 | 7,53E+09 | 6,38E+09 | 33,509 | 0 | 323,31 | 492 |
| 3,29E+06 | 3,90E+06 | 5,44E+06 | 3,99E+06 | 78,918 | 0,00463 | 1,7275 | 8 |
| 1,35E+07 | 1,23E+07 | 1,47E+07 | 1,51E+07 | 165,96 | 0 | 10,513 | 24 |
| 2,38E+06 | 9,39E+06 | 6,39E+06 | 9,57E+06 | 91,977 | 0,000153 | 4,1482 | 12 |
| 8,05E+06 | 8,27E+06 | 6,64E+06 | 7,49E+06 | 94,64 | 0,000157 | 4,8924 | 13 |
| 5,62E+06 | 4,20E+06 | 4,97E+06 | 2,82E+06 | 49,011 | 0,006207 | 1,5811 | 3 |
| 1,01E+06 | 3,34E+06 | 2,08E+06 | 3,62E+06 | 77,269 | 0,005413 | 1,6292 | 3 |
| 2,61E+07 | 2,15E+07 | 2,22E+07 | 2,34E+07 | 50,08 | 0,000157 | 4,8709 | 29 |
| 3,02E+06 | 1,97E+06 | 2,53E+06 | 2,39E+06 | 28,691 | 0,000147 | 3,148 | 7 |
| 9,78E+06 | 6,92E+06 | 7,76E+06 | 7,77E+06 | 33,859 | 0 | 65,016 | 6 |
| 2,44E+07 | 2,97E+07 | 2,36E+07 | 1,99E+07 | 71,343 | 0 | 19,335 | 39 |
| 1,08E+07 | 1,58E+07 | 1,39E+07 | 1,36E+07 | 42,058 | 0,000159 | 5,1983 | 13 |
| 2,12E+07 | 2,55E+07 | 1,80E+07 | 1,80E+07 | 65,384 | 0 | 12,583 | 32 |
| 7,63E+07 | 8,28E+07 | 7,75E+07 | 7,33E+07 | 118,2 | 0 | 32,817 | 66 |
| 8,06E+07 | 9,65E+07 | 9,66E+07 | 8,64E+07 | 21,878 | 0 | 37,936 | 52 |
| 7,71E+09 | 7,52E+09 | 7,16E+09 | 6,47E+09 | 57,477 | 0 | 323,31 | 869 |
| 2,97E+09 | 2,44E+09 | 2,58E+09 | 2,47E+09 | 34,306 | 0 | 323,31 | 168 |
| 3,53E+07 | 2,94E+07 | 3,10E+07 | 3,46E+07 | 16,355 | 0 | 104,72 | 6 |
| 8,75E+06 | 7,36E+06 | 7,65E+06 | 6,03E+06 | 42,132 | 0,000152 | 3,9685 | 17 |
| 1,87E+06 | 2,95E+06 | 3,35E+06 | 1,63E+06 | 58,779 | 0,000853 | 2,5609 | 3 |
| 2,39E+07 | 1,84E+07 | 1,68E+07 | 1,87E+07 | 171,65 | 0 | 38,54 | 28 |
| 2,49E+07 | 2,96E+07 | 3,09E+07 | 2,43E+07 | 91,514 | 0 | 16,229 | 37 |
| 1,98E+07 | 1,58E+07 | 2,36E+07 | 1,92E+07 | 108,96 | 0 | 10,372 | 20 |
| 6,01E+07 | 5,98E+07 | 6,22E+07 | 7,28E+07 | 67,969 | 0 | 28,568 | 50 |
| 2,99E+06 | 4,21E+06 | 5,38E+06 | 3,94E+06 | 67,411 | 0,007278 | 1,5422 | 6 |
| 7,89E+06 | 1,12E+07 | 9,83E+06 | 1,12E+07 | 112,36 | 0,000152 | 4,075 | 14 |
| 3,71E+09 | 4,12E+09 | 3,88E+09 | 4,19E+09 | 24,893 | 0 | 323,31 | 253 |
| 8,98E+06 | 1,13E+07 | 4,22E+06 | 6,70E+06 | 62,57 | 0,000154 | 4,417 | 11 |
| 8,99E+06 | 8,69E+06 | 6,30E+06 | 1,67E+06 | 21,748 | 0 | 5,8879 | 1 |
| 7,69E+06 | 1,30E+07 | 1,02E+07 | 1,05E+07 | 50,508 | 0 | 16,634 | 7 |
| 1,05E+09 | 1,17E+09 | 9,82E+08 | 1,01E+09 | 24,911 | 0 | 323,31 | 105 |
| 5,60E+07 | 9,61E+07 | 1,12E+08 | 5,17E+07 | 10,561 | 0 | 27,799 | 15 |
| 8,40E+06 | 3,11E+07 | 5,59E+06 | 4,45E+06 | 45,125 | 0 | 5,9909 | 10 |

| | | | | | | | |
|---|---|---|---|---|---|---|---|
| 3,16E+08 | 3,14E+08 | 3,08E+08 | 3,08E+08 | 99,818 | 0 | 295,13 | 193 |
| 3,45E+07 | 3,22E+07 | 3,68E+07 | 3,38E+07 | 93,312 | 0 | 22,176 | 57 |
| 3,38E+08 | 2,76E+08 | 3,11E+08 | 2,95E+08 | 22,432 | 0 | 82,398 | 92 |
| 4,57E+06 | 7,74E+06 | 5,07E+06 | 8,87E+06 | 26,445 | 0,000158 | 5,149 | 10 |
| 7,19E+08 | 5,66E+08 | 5,26E+08 | 5,32E+08 | 20,396 | 0 | 108,99 | 39 |
| 2,68E+09 | 2,34E+09 | 2,93E+09 | 2,81E+09 | 31,732 | 0 | 323,31 | 252 |
| 1,45E+08 | 1,49E+08 | 1,58E+08 | 1,50E+08 | 96,904 | 0 | 80,61 | 55 |
| 4,50E+06 | 4,27E+06 | 4,28E+06 | 6,55E+06 | 169,64 | 0,00015 | 3,6628 | 2 |
| 9,00E+05 | 2,37E+06 | 2,09E+06 | 1,68E+06 | 25,433 | 0,000149 | 3,4605 | 7 |
| 8,69E+07 | 7,62E+07 | 8,57E+07 | 7,20E+07 | 23,275 | 0 | 37,877 | 36 |
| 6,86E+07 | 7,61E+07 | 7,14E+07 | 6,23E+07 | 39,109 | 0 | 17,366 | 60 |
| 2,33E+07 | 2,01E+06 | 6,37E+06 | 1,63E+06 | 50,135 | 0,007409 | 1,5373 | 3 |
| 6,09E+06 | 3,56E+06 | 3,65E+06 | 3,21E+06 | 24,2 | 0,004625 | 1,7043 | 8 |
| 4,97E+08 | 4,86E+08 | 5,19E+08 | 5,64E+08 | 16,534 | 0 | 208,68 | 84 |
| 1,73E+07 | 1,56E+07 | 1,60E+07 | 1,46E+07 | 118,3 | 0,000157 | 4,8863 | 18 |
| 1,08E+07 | 1,47E+07 | 1,07E+07 | 1,46E+07 | 23,058 | 0 | 13,529 | 17 |
| 3,12E+08 | 3,29E+08 | 3,37E+08 | 3,27E+08 | 39,166 | 0 | 304,01 | 148 |
| 1,51E+08 | 1,55E+08 | 1,50E+08 | 1,41E+08 | 60,193 | 0 | 88,849 | 83 |
| 1,87E+08 | 2,04E+08 | 1,95E+08 | 1,99E+08 | 69,552 | 0 | 288,15 | 115 |
| 1,98E+08 | 2,22E+08 | 2,21E+08 | 2,16E+08 | 80,491 | 0 | 137,93 | 111 |
| 1,38E+08 | 1,31E+08 | 1,31E+08 | 7,57E+07 | 182,35 | 0 | 17,707 | 51 |
| 3,19E+08 | 3,49E+08 | 3,66E+08 | 4,20E+08 | 20,411 | 0 | 100,19 | 103 |
| 2,16E+08 | 1,61E+08 | 1,88E+08 | 1,87E+08 | 14,308 | 0 | 7,7315 | 39 |
| 5,35E+08 | 4,12E+08 | 4,04E+08 | 4,13E+08 | 29,891 | 0 | 323,31 | 140 |
| 2,26E+06 | 1,99E+06 | 1,31E+06 | 1,90E+06 | 32,883 | 0,000148 | 3,3221 | 7 |
| 6,12E+06 | 7,60E+06 | 8,30E+06 | 9,33E+06 | 20,153 | 0 | 12,321 | 10 |
| 1,79E+07 | 1,97E+07 | 1,63E+07 | 1,33E+07 | 66,839 | 0 | 6,3863 | 23 |
| 4,78E+07 | 3,86E+07 | 4,85E+07 | 4,17E+07 | 20,203 | 0 | 8,5766 | 6 |
| 1,77E+06 | 2,29E+06 | 3,32E+06 | 3,14E+06 | 55,837 | 0,00015 | 3,6665 | 4 |
| 3,05E+07 | 3,45E+07 | 3,98E+07 | 2,76E+07 | 42,202 | 0 | 9,9455 | 17 |
| 5,09E+08 | 4,18E+08 | 4,47E+08 | 3,82E+08 | 35,377 | 0 | 323,31 | 78 |
| 1,51E+08 | 1,60E+08 | 1,34E+08 | 1,49E+08 | 62,279 | 0 | 314,66 | 51 |
| 8,92E+07 | 1,04E+08 | 1,13E+08 | 9,38E+07 | 42,916 | 0 | 76,895 | 54 |
| 1,91E+08 | 1,62E+08 | 1,67E+08 | 2,38E+08 | 34,79 | 0 | 222,66 | 58 |
| 6,44E+08 | 6,32E+08 | 6,78E+08 | 6,44E+08 | 68,325 | 0 | 323,31 | 223 |
| 1,16E+08 | 1,19E+08 | 1,17E+08 | 1,13E+08 | 41,019 | 0 | 62,106 | 88 |
| 2,31E+07 | 2,63E+07 | 2,23E+07 | 1,98E+07 | 52,962 | 0 | 37,203 | 30 |
| 2,85E+06 | 3,69E+06 | 4,61E+06 | 1,11E+06 | 52,081 | 0,000153 | 4,0979 | 5 |
| 8,04E+08 | 7,67E+08 | 7,45E+08 | 7,27E+08 | 47,447 | 0 | 323,31 | 146 |
| 2,80E+08 | 2,63E+08 | 2,79E+08 | 2,67E+08 | 20,348 | 0 | 289,65 | 79 |
| 9,77E+06 | 8,10E+06 | 8,30E+06 | 6,21E+06 | 51,837 | 0,000431 | 2,7807 | 6 |
| 6,73E+06 | 1,16E+07 | 8,43E+06 | 1,15E+07 | 62,055 | 0 | 6,2717 | 15 |
| 2,77E+08 | 3,07E+08 | 3,18E+08 | 3,49E+08 | 75,474 | 0 | 312,98 | 103 |
| 3,32E+07 | 3,05E+07 | 3,17E+07 | 3,34E+07 | 36,852 | 0 | 32,48 | 34 |
| 4,80E+06 | 1,98E+06 | 1,35E+07 | 2,77E+06 | 9,3398 | 0 | 34,95 | 9 |
| 5,36E+08 | 5,46E+08 | 5,06E+08 | 5,03E+08 | 68,542 | 0 | 237,86 | 127 |
| 5,18E+06 | 4,46E+06 | 5,93E+06 | 5,12E+06 | 55,357 | 0 | 5,5597 | 7 |
| 3,94E+07 | 3,84E+07 | 3,39E+07 | 4,60E+07 | 165,2 | 0 | 96,526 | 38 |
| 8,13E+08 | 1,08E+09 | 1,05E+09 | 1,04E+09 | 13,293 | 0 | 9,6381 | 54 |
| 7,72E+06 | 9,37E+06 | 3,44E+06 | 3,29E+06 | 101,24 | 0,00015 | 3,7148 | 3 |

| | | | | | | | |
|---|---|---|---|---|---|---|---|
| 1,92E+07 | 2,21E+07 | 1,83E+07 | 2,43E+07 | 20,795 | 0 | 6,1303 | 12 |
| 2,88E+06 | 1,85E+06 | 1,52E+06 | 6,76E+06 | 46,384 | 0,001274 | 2,4559 | 3 |
| 3,88E+07 | 3,58E+07 | 3,41E+07 | 3,72E+07 | 67,319 | 0 | 22,948 | 30 |
| 3,23E+06 | 3,83E+06 | 3,98E+06 | 2,80E+06 | 138,83 | 0,000146 | 3,0983 | 7 |
| 4,75E+08 | 4,44E+08 | 5,27E+08 | 4,54E+08 | 14,478 | 0 | 235,63 | 126 |
| 9,09E+08 | 8,40E+08 | 9,02E+08 | 8,81E+08 | 69,59 | 0 | 323,31 | 190 |
| 3,77E+08 | 4,05E+08 | 4,23E+08 | 4,31E+08 | 291,55 | 0 | 253,8 | 349 |
| 3,60E+07 | 4,56E+07 | 4,21E+07 | 4,70E+07 | 34,412 | 0 | 14,38 | 52 |
| 5,74E+06 | 5,50E+06 | 5,11E+06 | 5,67E+06 | 86,939 | 0,001273 | 2,4375 | 11 |
| 1,26E+07 | 1,05E+07 | 1,32E+07 | 1,01E+07 | 120,07 | 0,005417 | 1,6395 | 13 |
| 3,94E+08 | 3,44E+08 | 4,01E+08 | 4,05E+08 | 32,316 | 0 | 65,106 | 70 |
| 9,39E+07 | 7,51E+07 | 9,86E+07 | 1,01E+08 | 17,936 | 0 | 82,284 | 49 |
| 3,60E+07 | 4,34E+07 | 3,52E+07 | 3,69E+07 | 32,985 | 0 | 6,3648 | 22 |
| 1,38E+07 | 1,40E+07 | 1,54E+07 | 1,71E+07 | 82,962 | 0 | 5,834 | 8 |
| 1,57E+07 | 3,67E+06 | 4,38E+06 | 1,91E+06 | 13,565 | 0,000149 | 3,437 | 6 |
| 1,54E+07 | 1,66E+07 | 1,61E+07 | 1,69E+07 | 29,922 | 0 | 7,5404 | 20 |
| 5,35E+07 | 5,24E+07 | 4,85E+07 | 5,60E+07 | 69,04 | 0 | 29,391 | 37 |
| 9,04E+07 | 8,49E+07 | 9,02E+07 | 7,56E+07 | 33,263 | 0 | 25,953 | 40 |
| 2,25E+07 | 1,29E+07 | 2,34E+07 | 2,05E+07 | 46,12 | 0,00015 | 3,6741 | 17 |
| 2,11E+07 | 5,50E+07 | 5,42E+07 | 5,50E+07 | 212,92 | 0 | 11,224 | 40 |
| 3,29E+06 | 6,86E+06 | 4,61E+06 | 6,74E+06 | 8,7558 | 0,000992 | 2,4843 | 4 |
| 2,51E+07 | 2,10E+07 | 3,14E+07 | 3,28E+07 | 37,57 | 0,000432 | 2,8115 | 26 |
| 1,76E+07 | 1,78E+07 | 1,77E+07 | 2,03E+07 | 12,428 | 0 | 16,032 | 23 |
| 1,37E+07 | 9,96E+06 | 1,40E+07 | 1,29E+07 | 68,346 | 0,000148 | 3,3954 | 13 |
| 1,83E+07 | 2,16E+07 | 2,54E+07 | 2,35E+07 | 75,847 | 0,004368 | 1,7577 | 11 |
| 3,19E+08 | 2,69E+08 | 2,72E+08 | 2,84E+08 | 44,141 | 0 | 156,25 | 101 |
| 1,47E+07 | 1,64E+07 | 4,82E+06 | 1,19E+07 | 50,664 | 0,000157 | 4,9076 | 6 |
| 5,03E+08 | 5,00E+08 | 4,89E+08 | 4,85E+08 | 85,965 | 0 | 323,31 | 210 |
| 2,04E+07 | 2,01E+07 | 2,47E+07 | 2,16E+07 | 12,707 | 0 | 27,691 | 16 |
| 6,17E+06 | 3,25E+06 | 5,44E+06 | 6,49E+06 | 7,743 | 0,009493 | 1,4241 | 3 |
| 6,82E+07 | 5,44E+07 | 6,32E+07 | 7,82E+07 | 29,664 | 0 | 120,2 | 25 |
| 2,18E+07 | 1,43E+07 | 1,63E+07 | 1,62E+07 | 12,33 | 0 | 6,8235 | 21 |
| 1,30E+07 | 3,24E+06 | 7,20E+06 | 7,97E+06 | 55,011 | 0,00357 | 1,8826 | 4 |
| 3,50E+06 | 6,21E+06 | 5,74E+06 | 6,05E+06 | 39,24 | 0 | 5,3609 | 11 |
| 2,83E+08 | 3,03E+08 | 2,97E+08 | 3,13E+08 | 262,1 | 0 | 128,83 | 262 |
| 3,10E+06 | 1,91E+06 | 3,66E+06 | 5,69E+06 | 33,288 | 0,003046 | 1,9962 | 3 |
| 3,71E+07 | 2,46E+07 | 5,95E+07 | 5,09E+07 | 9,0167 | 0 | 9,5607 | 17 |
| 1,02E+07 | 1,19E+07 | 9,35E+06 | 8,26E+06 | 83,976 | 0,000155 | 4,5668 | 18 |
| 5,74E+06 | 1,08E+07 | 5,79E+06 | 3,19E+06 | 143,37 | 0,001269 | 2,3775 | 8 |
| 2,86E+08 | 3,43E+08 | 2,95E+08 | 2,74E+08 | 42,906 | 0 | 52,223 | 97 |
| 1,34E+07 | 1,04E+07 | 3,88E+06 | 1,38E+07 | 60,582 | 0 | 8,4543 | 8 |
| 1,15E+06 | 4,85E+06 | 4,36E+06 | 8,04E+06 | 71,817 | 0,000158 | 5,0395 | 7 |
| 2,37E+09 | 1,89E+09 | 2,14E+09 | 2,16E+09 | 24,162 | 0 | 323,31 | 281 |
| 1,52E+10 | 1,62E+10 | 1,58E+10 | 1,73E+10 | 35,611 | 0 | 323,31 | 742 |
| 6,26E+07 | 1,40E+07 | 1,98E+07 | 1,52E+07 | 245,25 | 0,002368 | 2,0789 | 6 |
| 5,79E+06 | 5,05E+06 | 3,22E+06 | 5,34E+06 | 23,901 | 0,000292 | 3,0675 | 4 |
| 3,12E+07 | 1,74E+07 | 2,02E+07 | 1,81E+07 | 144,55 | 0 | 17,938 | 30 |
| 5,55E+06 | 1,24E+07 | 3,49E+06 | 2,09E+07 | 38,333 | 0 | 8,1854 | 16 |
| 3,26E+06 | 3,86E+06 | 3,43E+06 | 3,18E+06 | 47,321 | 0,000147 | 3,2924 | 6 |
| 3,01E+08 | 2,18E+08 | 2,67E+08 | 2,64E+08 | 79,755 | 0 | 95,886 | 98 |

| | | | | | | | |
|---|---|---|---|---|---|---|---|
| 9,21E+06 | 2,96E+07 | 1,72E+06 | 6,04E+06 | 12,483 | 0 | 157,2 | 5 |
| 5,20E+06 | 6,52E+06 | 2,64E+06 | 2,93E+06 | 148,65 | 0,000153 | 4,2323 | 9 |
| 2,60E+07 | 1,41E+07 | 1,49E+07 | 1,83E+07 | 297,48 | 0 | 16,941 | 30 |
| 3,48E+08 | 3,84E+08 | 3,69E+08 | 3,64E+08 | 93,496 | 0 | 214 | 116 |
| 1,68E+07 | 3,68E+07 | 2,76E+07 | 3,45E+07 | 66,856 | 0 | 11,332 | 36 |
| 5,67E+08 | 6,17E+08 | 5,69E+08 | 5,24E+08 | 36,776 | 0 | 323,31 | 141 |
| 3,66E+06 | 2,15E+06 | 4,55E+06 | 4,71E+06 | 34,053 | 0,005019 | 1,6605 | 6 |
| 7,98E+08 | 6,50E+08 | 7,38E+08 | 6,47E+08 | 38,724 | 0 | 233,63 | 143 |
| 9,04E+07 | 7,91E+07 | 8,29E+07 | 8,47E+07 | 48,08 | 0 | 34,135 | 28 |
| 1,68E+07 | 1,67E+07 | 1,54E+07 | 1,78E+07 | 22,017 | 0,000433 | 2,8418 | 12 |
| 7,21E+06 | 5,79E+06 | 8,49E+06 | 7,36E+06 | 10,254 | 0,000435 | 2,9785 | 8 |
| 4,60E+08 | 5,27E+08 | 5,11E+08 | 5,35E+08 | 40,526 | 0 | 323,31 | 141 |
| 9,40E+07 | 9,08E+07 | 9,96E+07 | 9,12E+07 | 15,183 | 0 | 11,497 | 23 |
| 1,86E+07 | 1,81E+07 | 1,34E+07 | 1,95E+07 | 16,188 | 0,000154 | 4,2864 | 22 |
| 5,93E+06 | 4,86E+06 | 4,40E+06 | 5,75E+06 | 148,06 | 0,000154 | 4,2972 | 7 |
| 1,79E+06 | 2,51E+06 | 3,81E+06 | 2,45E+06 | 76,723 | 0 | 55,674 | 15 |
| 3,77E+09 | 3,72E+09 | 3,89E+09 | 3,82E+09 | 126,61 | 0 | 323,31 | 612 |
| 1,77E+08 | 1,92E+08 | 1,82E+08 | 2,10E+08 | 68,3 | 0 | 202,98 | 106 |
| 9,25E+06 | 9,04E+06 | 8,94E+06 | 5,32E+06 | 48,743 | 0 | 6,5001 | 16 |
| 4,34E+07 | 6,38E+07 | 6,57E+07 | 6,84E+07 | 20,249 | 0,000152 | 4,0689 | 26 |
| 3,89E+07 | 5,68E+06 | 7,59E+06 | 6,13E+06 | 51,621 | 0 | 38,061 | 6 |
| 2,40E+08 | 2,18E+08 | 2,11E+08 | 2,35E+08 | 47,499 | 0 | 70,431 | 104 |
| 3,81E+06 | 6,67E+06 | 6,96E+06 | 6,07E+06 | 51,113 | 0,002101 | 2,1955 | 4 |
| 1,99E+06 | 1,62E+06 | 2,02E+06 | 2,73E+06 | 40,933 | 0,00043 | 2,7488 | 5 |
| 4,13E+07 | 4,29E+07 | 4,01E+07 | 3,76E+07 | 28,954 | 0 | 26,571 | 36 |
| 1,65E+06 | 2,58E+06 | 7,91E+06 | 1,00E+06 | 69,792 | 0,001409 | 2,3444 | 1 |
| 1,05E+07 | 8,02E+06 | 7,57E+06 | 7,20E+06 | 70,862 | 0 | 11,231 | 16 |
| 7,81E+06 | 1,27E+07 | 1,08E+07 | 1,17E+07 | 57,344 | 0,000154 | 4,2811 | 9 |
| 6,68E+06 | 3,89E+06 | 8,37E+06 | 1,14E+07 | 96,275 | 0,000433 | 2,8848 | 4 |
| 1,47E+08 | 1,34E+08 | 1,33E+08 | 1,32E+08 | 329,46 | 0 | 84,673 | 162 |
| 3,57E+06 | 5,01E+06 | 2,68E+06 | 5,79E+06 | 57,569 | 0,000155 | 4,4308 | 2 |
| 9,65E+09 | 9,28E+09 | 9,65E+09 | 1,00E+10 | 26,674 | 0 | 286,37 | 620 |
| 2,57E+07 | 2,86E+07 | 3,13E+07 | 2,38E+07 | 30,355 | 0 | 7,0828 | 16 |
| 4,23E+06 | 8,24E+06 | 6,52E+06 | 7,45E+06 | 202,97 | 0 | 6,7 | 15 |
| 5,93E+09 | 5,84E+09 | 5,37E+09 | 5,25E+09 | 21,423 | 0 | 298,85 | 409 |
| 6,36E+07 | 5,10E+07 | 5,98E+07 | 5,79E+07 | 8,4977 | 0 | 6,1597 | 18 |
| 5,75E+07 | 5,40E+07 | 5,37E+07 | 5,49E+07 | 69,56 | 0 | 11,838 | 37 |
| 8,35E+06 | 1,25E+07 | 1,16E+07 | 1,22E+07 | 21,626 | 0,000431 | 2,7675 | 13 |
| 2,75E+08 | 2,30E+08 | 2,78E+08 | 2,25E+08 | 23,076 | 0 | 210,72 | 80 |
| 1,96E+08 | 1,69E+08 | 1,86E+08 | 2,14E+08 | 49,348 | 0 | 57,687 | 67 |
| 6,29E+06 | 2,99E+06 | 9,08E+06 | 8,17E+06 | 82,198 | 0,000154 | 4,286 | 11 |
| 2,24E+08 | 2,41E+08 | 2,23E+08 | 1,96E+08 | 52,422 | 0 | 314,6 | 98 |
| 3,77E+07 | 3,14E+07 | 2,64E+07 | 3,81E+07 | 28,899 | 0 | 97,022 | 22 |
| 2,10E+08 | 1,90E+08 | 2,07E+08 | 2,05E+08 | 98,047 | 0 | 74,019 | 125 |
| 1,97E+06 | 4,24E+06 | 2,91E+06 | 3,06E+06 | 104,12 | 0,000147 | 3,2862 | 10 |
| 3,86E+07 | 4,63E+07 | 5,09E+07 | 5,52E+07 | 34,727 | 0 | 13,386 | 56 |
| 2,24E+08 | 2,47E+08 | 2,54E+08 | 2,62E+08 | 56,855 | 0 | 104,37 | 105 |
| 3,17E+06 | 6,42E+06 | 7,05E+06 | 5,90E+06 | 66,264 | 0,000158 | 5,0164 | 8 |
| 1,15E+07 | 1,12E+07 | 8,06E+06 | 9,93E+06 | 54,688 | 0 | 12,088 | 18 |
| 1,18E+08 | 9,18E+07 | 8,09E+07 | 1,40E+08 | 31,794 | 0 | 19,776 | 53 |

| | | | | | | | |
|---|---|---|---|---|---|---|---|
| 9,76E+06 | 7,37E+06 | 7,67E+06 | 9,49E+06 | 80,491 | 0 | 9,5217 | 17 |
| 4,52E+07 | 8,77E+07 | 4,74E+07 | 5,90E+07 | 23,599 | 0 | 8,8039 | 42 |
| 8,12E+06 | 1,23E+07 | 1,07E+07 | 9,75E+06 | 12,575 | 0,000155 | 4,4517 | 9 |
| 2,93E+07 | 2,15E+07 | 2,40E+07 | 2,39E+07 | 25,686 | 0 | 19,128 | 22 |
| 3,55E+09 | 3,30E+09 | 3,73E+09 | 3,99E+09 | 15,798 | 0 | 144,7 | 115 |
| 5,29E+06 | 3,89E+06 | 3,64E+06 | 4,27E+06 | 17,829 | 0,000151 | 3,9094 | 8 |
| 5,72E+06 | 4,75E+06 | 4,27E+06 | 3,01E+06 | 91,921 | 0 | 14,145 | 9 |
| 9,94E+07 | 1,29E+08 | 1,14E+08 | 1,11E+08 | 24,967 | 0 | 41,212 | 58 |
| 3,59E+09 | 3,68E+09 | 4,43E+09 | 4,80E+09 | 13,742 | 0 | 43,104 | 82 |
| 5,04E+08 | 4,61E+08 | 4,52E+08 | 4,65E+08 | 63,589 | 0 | 265,23 | 184 |
| 6,96E+07 | 6,68E+07 | 8,74E+07 | 8,38E+07 | 16,786 | 0 | 61,476 | 54 |
| 6,56E+06 | 3,88E+06 | 3,80E+06 | 5,16E+06 | 25,947 | 0,000156 | 4,6046 | 7 |
| 4,04E+06 | 4,38E+06 | 2,77E+06 | 2,05E+06 | 37,899 | 0,001273 | 2,4372 | 6 |
| 5,49E+06 | 1,61E+06 | 2,30E+06 | 3,53E+06 | 33,394 | 0,009368 | 1,4358 | 5 |
| 5,13E+06 | 4,09E+06 | 3,85E+06 | 1,20E+06 | 66,365 | 0,000156 | 4,6357 | 6 |
| 6,53E+06 | 1,66E+06 | 5,40E+06 | 2,31E+06 | 17,08 | 0,000429 | 2,6925 | 3 |
| 2,01E+06 | 2,08E+06 | 2,80E+06 | 2,35E+06 | 78,748 | 0,006077 | 1,5965 | 3 |
| 1,56E+08 | 1,65E+08 | 1,60E+08 | 1,75E+08 | 48,186 | 0 | 95,518 | 74 |
| 9,38E+07 | 7,67E+07 | 7,75E+07 | 7,38E+07 | 32,183 | 0 | 20,343 | 37 |
| 1,25E+09 | 1,33E+09 | 1,30E+09 | 1,37E+09 | 54,272 | 0 | 323,31 | 301 |
| 2,80E+06 | 1,17E+07 | 1,41E+06 | 1,62E+07 | 122,67 | 0,000152 | 4,0786 | 5 |
| 5,13E+06 | 2,08E+06 | 3,34E+06 | 5,96E+06 | 22,921 | 0 | 12,963 | 6 |
| 6,18E+07 | 2,74E+07 | 3,53E+07 | 2,51E+07 | 22,96 | 0 | 50,843 | 14 |
| 1,31E+07 | 1,45E+07 | 2,00E+07 | 2,33E+07 | 55,676 | 0 | 23,861 | 7 |
| 2,13E+08 | 2,51E+08 | 2,77E+08 | 2,80E+08 | 13,382 | 0 | 115,57 | 49 |
| 3,63E+07 | 6,24E+07 | 2,66E+07 | 3,01E+07 | 20,313 | 0 | 71,154 | 10 |
| 1,96E+07 | 1,77E+07 | 1,33E+07 | 1,55E+07 | 49,917 | 0 | 12,37 | 21 |
| 1,23E+07 | 1,17E+07 | 8,93E+06 | 1,15E+07 | 113,18 | 0 | 6,4929 | 26 |
| 3,47E+09 | 3,38E+09 | 2,57E+09 | 2,95E+09 | 31,419 | 0 | 323,31 | 360 |
| 6,37E+06 | 7,80E+06 | 1,12E+07 | 7,28E+06 | 92,942 | 0,00015 | 3,6426 | 16 |
| 2,77E+07 | 3,28E+07 | 3,15E+07 | 3,12E+07 | 20,201 | 0 | 104,11 | 23 |
| 1,04E+08 | 1,21E+08 | 1,25E+08 | 1,24E+08 | 398,75 | 0 | 45,536 | 130 |
| 1,59E+08 | 1,45E+08 | 1,81E+08 | 1,68E+08 | 38,404 | 0 | 9,7239 | 38 |
| 5,77E+08 | 6,00E+08 | 5,96E+08 | 5,70E+08 | 54,65 | 0 | 323,31 | 147 |
| 2,86E+08 | 3,15E+08 | 3,21E+08 | 3,04E+08 | 50,834 | 0 | 285,13 | 121 |
| 2,11E+08 | 2,67E+08 | 2,43E+08 | 2,11E+08 | 56,369 | 0 | 290,04 | 82 |
| 5,89E+06 | 9,68E+06 | 5,81E+06 | 5,20E+06 | 22,949 | 0,000155 | 4,4254 | 7 |
| 1,68E+06 | 4,56E+06 | 3,01E+06 | 1,93E+06 | 68,592 | 0,001547 | 2,3022 | 3 |
| 2,84E+06 | 4,30E+06 | 4,33E+06 | 5,01E+06 | 52,214 | 0,002913 | 2,0188 | 6 |
| 6,15E+07 | 5,20E+07 | 5,83E+07 | 6,17E+07 | 19,029 | 0 | 101,93 | 41 |
| 7,85E+06 | 5,72E+06 | 5,66E+06 | 4,81E+06 | 46,084 | 0 | 63,456 | 13 |
| 1,48E+07 | 1,48E+07 | 1,96E+07 | 7,72E+06 | 78,668 | 0,000154 | 4,3649 | 19 |
| 6,20E+06 | 3,47E+06 | 3,44E+06 | 3,11E+06 | 29,128 | 0 | 5,4362 | 5 |
| 1,01E+08 | 1,17E+08 | 1,02E+08 | 1,21E+08 | 20,208 | 0 | 74,086 | 40 |
| 1,16E+07 | 1,23E+07 | 9,82E+06 | 1,37E+07 | 86,622 | 0 | 8,7467 | 21 |
| 5,23E+06 | 2,75E+06 | 6,69E+06 | 8,70E+06 | 33,485 | 0 | 11,585 | 6 |
| 2,99E+08 | 2,66E+08 | 3,17E+08 | 3,26E+08 | 23,255 | 0 | 143 | 78 |
| 7,62E+06 | 1,19E+07 | 5,89E+06 | 8,78E+06 | 27,754 | 0 | 6,6757 | 8 |
| 2,30E+07 | 1,88E+07 | 2,26E+07 | 1,27E+07 | 12,957 | 0 | 47,812 | 18 |
| 4,94E+07 | 5,37E+07 | 4,14E+07 | 5,37E+07 | 28,38 | 0 | 11,176 | 32 |

| | | | | | | | |
|---|---|---|---|---|---|---|---|
| 2,46E+07 | 2,07E+07 | 3,12E+07 | 2,71E+07 | 61,558 | 0 | 8,924 | 25 |
| 3,01E+07 | 4,90E+07 | 3,21E+07 | 4,06E+07 | 10,477 | 0 | 52,397 | 21 |
| 6,35E+07 | 7,19E+07 | 7,18E+07 | 6,94E+07 | 198,79 | 0 | 42,163 | 79 |
| 1,61E+08 | 1,53E+08 | 1,51E+08 | 1,18E+08 | 47,038 | 0 | 148,07 | 57 |
| 2,37E+08 | 2,85E+08 | 2,20E+08 | 2,04E+08 | 44,355 | 0 | 155,75 | 59 |
| 2,17E+06 | 5,63E+06 | 2,76E+06 | 2,48E+06 | 151,46 | 0,003568 | 1,8694 | 4 |
| 1,70E+08 | 2,16E+08 | 1,68E+08 | 1,46E+08 | 43,388 | 0 | 123,25 | 74 |
| 2,95E+06 | 4,10E+06 | 6,39E+06 | 4,97E+06 | 53,11 | 0 | 5,4544 | 4 |
| 4,13E+06 | 4,11E+06 | 5,04E+06 | 5,75E+06 | 58,732 | 0,003175 | 1,9465 | 5 |
| 1,12E+07 | 1,23E+07 | 1,41E+07 | 1,56E+07 | 38,502 | 0 | 31,294 | 6 |
| 1,50E+08 | 1,71E+08 | 1,32E+08 | 1,57E+08 | 57,584 | 0 | 69,067 | 58 |
| 1,12E+07 | 1,02E+07 | 1,34E+07 | 5,97E+06 | 83,096 | 0 | 5,4202 | 14 |
| 1,62E+06 | 3,52E+06 | 3,70E+06 | 3,82E+06 | 16,875 | 0,000149 | 3,426 | 6 |
| 4,37E+08 | 4,39E+08 | 4,89E+08 | 4,94E+08 | 13,527 | 0 | 80,457 | 91 |
| 3,06E+09 | 2,77E+09 | 2,56E+09 | 2,95E+09 | 18,916 | 0 | 178,02 | 327 |
| 1,14E+08 | 1,06E+08 | 9,65E+07 | 9,29E+07 | 73,961 | 0 | 58,681 | 84 |
| 8,82E+08 | 6,74E+08 | 8,60E+08 | 7,58E+08 | 16,318 | 0 | 323,31 | 87 |
| 5,35E+06 | 9,76E+06 | 4,65E+06 | 3,85E+06 | 27,229 | 0,000157 | 5,0001 | 15 |
| 2,85E+08 | 2,73E+08 | 2,61E+08 | 2,74E+08 | 155,81 | 0 | 158,86 | 166 |
| 1,70E+07 | 1,02E+07 | 1,24E+07 | 1,19E+07 | 71,709 | 0 | 26,144 | 17 |
| 2,10E+08 | 2,26E+08 | 2,42E+08 | 2,06E+08 | 16,062 | 0 | 109,39 | 76 |
| 4,80E+07 | 4,56E+07 | 4,97E+07 | 5,00E+07 | 70,865 | 0 | 59,722 | 21 |
| 2,73E+07 | 3,00E+07 | 3,21E+07 | 2,79E+07 | 50,248 | 0 | 8,938 | 26 |
| 1,00E+08 | 1,01E+08 | 1,10E+08 | 1,09E+08 | 44,202 | 0 | 66,625 | 47 |
| 4,12E+06 | 5,92E+06 | 5,69E+06 | 6,80E+06 | 25,323 | 0 | 56,169 | 26 |
| 5,48E+07 | 6,08E+07 | 5,50E+07 | 7,37E+07 | 42,89 | 0 | 51,452 | 59 |
| 3,07E+07 | 3,14E+07 | 3,04E+07 | 2,57E+07 | 21,916 | 0 | 13,136 | 27 |
| 8,45E+06 | 7,05E+06 | 9,34E+06 | 1,17E+07 | 31,398 | 0 | 19,377 | 12 |
| 1,50E+09 | 1,71E+09 | 1,62E+09 | 1,67E+09 | 116,45 | 0 | 323,31 | 416 |
| 1,86E+06 | 4,06E+06 | 5,28E+06 | 9,25E+05 | 268,88 | 0,002909 | 2,0062 | 7 |
| 2,39E+07 | 2,21E+07 | 3,16E+07 | 2,78E+07 | 64,959 | 0 | 10,784 | 36 |
| 6,38E+06 | 3,46E+06 | 6,81E+06 | 3,60E+06 | 192,56 | 0,000152 | 3,9565 | 4 |
| 1,93E+08 | 1,71E+08 | 1,49E+08 | 1,45E+08 | 23,59 | 0 | 156,27 | 41 |
| 7,19E+07 | 6,19E+07 | 7,53E+07 | 6,13E+07 | 53,748 | 0 | 97,313 | 40 |
| 4,63E+08 | 4,60E+08 | 4,63E+08 | 4,31E+08 | 37,548 | 0 | 323,31 | 173 |
| 8,32E+06 | 7,46E+06 | 9,93E+06 | 8,43E+06 | 92,563 | 0 | 5,7606 | 18 |
| 1,18E+07 | 1,90E+07 | 2,04E+07 | 1,63E+07 | 47,783 | 0,000146 | 3,0815 | 10 |
| 6,02E+06 | 4,24E+06 | 4,35E+06 | 5,91E+06 | 53,771 | 0,002097 | 2,1564 | 5 |
| 9,02E+07 | 9,44E+07 | 1,99E+08 | 1,09E+08 | 10,119 | 0 | 68,701 | 62 |
| 3,15E+07 | 1,52E+08 | 1,14E+08 | 1,90E+08 | 24,348 | 0 | 8,3748 | 47 |
| 1,71E+08 | 1,51E+08 | 1,34E+08 | 1,33E+08 | 20,864 | 0 | 90,507 | 67 |
| 6,29E+06 | 2,42E+06 | 5,88E+06 | 8,00E+06 | 19,815 | 0 | 6,2095 | 9 |
| 1,73E+06 | 3,67E+06 | 1,98E+06 | 1,50E+06 | 33,375 | 0,004623 | 1,7012 | 3 |
| 8,76E+07 | 9,08E+07 | 1,04E+08 | 1,06E+08 | 15,072 | 0 | 76,196 | 44 |
| 1,20E+08 | 7,81E+07 | 1,09E+08 | 5,60E+07 | 16,629 | 0 | 115,15 | 21 |
| 7,52E+07 | 5,32E+07 | 6,56E+07 | 7,12E+07 | 26,215 | 0 | 13,292 | 27 |
| 1,73E+07 | 1,98E+07 | 1,67E+07 | 1,66E+07 | 36,059 | 0 | 10,016 | 24 |
| 1,42E+07 | 1,64E+07 | 1,81E+07 | 1,79E+07 | 86,791 | 0 | 9,4327 | 26 |
| 6,84E+07 | 6,26E+07 | 5,68E+07 | 5,67E+07 | 63,798 | 0 | 74,104 | 44 |
| 1,21E+10 | 1,30E+10 | 1,12E+10 | 1,27E+10 | 22,576 | 0 | 84,722 | 261 |

| | | | | | | | |
|---|---|---|---|---|---|---|---|
| 3,45E+07 | 4,15E+07 | 3,95E+07 | 4,51E+07 | 83,772 | 0 | 16,935 | 26 |
| 5,24E+07 | 7,31E+07 | 7,34E+07 | 7,09E+07 | 43,843 | 0 | 7,2912 | 34 |
| 3,44E+08 | 3,95E+08 | 4,01E+08 | 3,73E+08 | 20,825 | 0 | 323,31 | 38 |
| 1,92E+09 | 1,81E+09 | 1,79E+09 | 1,80E+09 | 217,75 | 0 | 323,31 | 532 |
| 7,60E+07 | 6,94E+07 | 7,44E+07 | 8,90E+07 | 113,88 | 0 | 157 | 69 |
| 1,05E+08 | 1,11E+08 | 1,09E+08 | 1,16E+08 | 96,965 | 0 | 59,284 | 88 |
| 5,83E+06 | 9,88E+06 | 2,81E+06 | 6,97E+06 | 28,507 | 0,001547 | 2,3015 | 4 |
| 3,51E+06 | 5,31E+06 | 4,57E+06 | 4,64E+06 | 136,76 | 0,000147 | 3,1634 | 13 |
| 8,51E+06 | 1,26E+07 | 9,16E+06 | 1,60E+07 | 30,114 | 0,000155 | 4,4782 | 14 |
| 1,93E+07 | 1,98E+07 | 1,71E+07 | 1,92E+07 | 31,01 | 0 | 11,744 | 22 |
| 6,18E+08 | 5,54E+08 | 6,52E+08 | 7,49E+08 | 28,127 | 0 | 82,43 | 65 |
| 3,84E+07 | 4,15E+07 | 4,41E+07 | 4,05E+07 | 26,457 | 0 | 18,59 | 38 |
| 5,94E+06 | 5,61E+06 | 7,63E+06 | 9,27E+06 | 61,84 | 0 | 16,708 | 9 |
| 6,62E+06 | 3,35E+06 | 5,33E+06 | 4,54E+06 | 54,152 | 0,002913 | 2,0216 | 6 |
| 3,00E+07 | 4,31E+07 | 3,77E+07 | 4,14E+07 | 37,803 | 0 | 8,7154 | 28 |
| 5,36E+06 | 5,04E+06 | 3,55E+06 | 4,28E+06 | 38,954 | 0,000992 | 2,4842 | 9 |
| 2,47E+08 | 2,63E+08 | 3,20E+08 | 2,77E+08 | 47,129 | 0 | 93,383 | 66 |
| 3,90E+06 | 6,94E+06 | 6,66E+06 | 2,76E+06 | 37,275 | 0,00043 | 2,7355 | 7 |
| 1,64E+07 | 1,23E+07 | 6,37E+06 | 3,69E+06 | 90,11 | 0,000156 | 4,8158 | 7 |
| 3,69E+06 | 2,79E+06 | 3,48E+06 | 4,65E+06 | 175,18 | 0,002097 | 2,1595 | 5 |
| 1,56E+08 | 1,43E+08 | 1,32E+08 | 1,42E+08 | 23,805 | 0 | 32,554 | 36 |
| 6,23E+08 | 6,76E+08 | 6,66E+08 | 6,68E+08 | 142,28 | 0 | 323,31 | 318 |
| 2,71E+07 | 3,30E+07 | 3,81E+07 | 3,35E+07 | 45,724 | 0 | 34,819 | 28 |
| 1,52E+08 | 1,40E+08 | 1,72E+08 | 1,78E+08 | 35,015 | 0 | 50,484 | 57 |
| 4,21E+07 | 3,87E+07 | 4,78E+07 | 5,28E+07 | 16,524 | 0 | 301,43 | 27 |
| 2,36E+07 | 1,01E+07 | 9,50E+06 | 9,34E+06 | 53,961 | 0 | 17,839 | 11 |
| 2,75E+08 | 2,62E+08 | 2,81E+08 | 2,82E+08 | 20,486 | 0 | 94,734 | 54 |
| 4,61E+07 | 5,55E+07 | 4,67E+07 | 3,68E+07 | 25,219 | 0 | 5,5775 | 25 |
| 1,98E+09 | 1,55E+09 | 1,63E+09 | 1,77E+09 | 11,614 | 0 | 110,8 | 207 |
| 1,46E+09 | 1,40E+09 | 1,40E+09 | 1,20E+09 | 20,855 | 0 | 323,31 | 253 |
| 4,71E+07 | 4,83E+07 | 5,33E+07 | 5,02E+07 | 64,562 | 0 | 18,338 | 42 |
| 4,61E+07 | 3,96E+07 | 3,70E+07 | 4,26E+07 | 35,851 | 0 | 16,98 | 31 |
| 1,82E+07 | 2,01E+07 | 2,30E+07 | 2,71E+07 | 82,26 | 0 | 25,907 | 34 |
| 8,92E+06 | 1,13E+07 | 1,31E+07 | 8,09E+06 | 41,032 | 0 | 44,527 | 9 |
| 1,80E+08 | 1,21E+08 | 1,25E+08 | 1,14E+08 | 16,191 | 0 | 117,03 | 39 |
| 2,92E+07 | 3,32E+07 | 2,23E+07 | 2,66E+07 | 35,152 | 0 | 27,303 | 24 |
| 2,36E+07 | 2,94E+07 | 2,58E+07 | 2,56E+07 | 79,196 | 0 | 17,663 | 33 |
| 8,78E+06 | 6,10E+06 | 1,23E+06 | 2,83E+06 | 66,126 | 0,004367 | 1,7572 | 3 |
| 2,96E+06 | 4,08E+06 | 3,83E+06 | 4,32E+06 | 67,416 | 0,000148 | 3,3853 | 11 |
| 1,05E+09 | 9,64E+08 | 1,23E+09 | 1,27E+09 | 12,554 | 0 | 7,9039 | 122 |
| 4,26E+06 | 2,64E+06 | 2,42E+06 | 7,37E+06 | 88,548 | 0,000146 | 3,1263 | 6 |
| 2,33E+06 | 3,13E+06 | 3,37E+06 | 3,21E+06 | 67,182 | 0,001271 | 2,4203 | 8 |
| 7,41E+07 | 8,75E+07 | 7,81E+07 | 7,92E+07 | 54,352 | 0 | 17,934 | 47 |
| 5,81E+07 | 5,11E+07 | 5,71E+07 | 5,61E+07 | 82,956 | 0 | 24,877 | 54 |
| 1,85E+08 | 2,11E+08 | 1,64E+08 | 1,68E+08 | 43,247 | 0 | 64,256 | 74 |
| 6,78E+06 | 5,54E+06 | 5,35E+06 | 6,06E+06 | 55,164 | 0,005424 | 1,6503 | 6 |
| 3,44E+06 | 5,94E+06 | 6,60E+06 | 6,01E+06 | 38,642 | 0 | 34,963 | 8 |
| 1,53E+08 | 1,26E+08 | 1,18E+08 | 1,27E+08 | 56,598 | 0 | 101,21 | 74 |
| 4,05E+06 | 3,93E+06 | 4,94E+06 | 4,01E+06 | 147,11 | 0,000146 | 3,1114 | 12 |
| 1,85E+08 | 1,40E+08 | 1,41E+08 | 1,54E+08 | 22,278 | 0 | 136,15 | 54 |

| | | | | | | | |
|---|---|---|---|---|---|---|---|
| 2,55E+08 | 2,61E+08 | 2,27E+08 | 2,47E+08 | 145,31 | 0 | 206,67 | 162 |
| 7,93E+07 | 9,19E+07 | 8,24E+07 | 7,59E+07 | 140,6 | 0 | 34,719 | 59 |
| 8,11E+07 | 3,85E+07 | 6,78E+07 | 5,08E+07 | 6,1773 | 0,000156 | 4,6444 | 11 |
| 3,45E+07 | 2,84E+07 | 2,98E+07 | 2,80E+07 | 86,771 | 0 | 23,984 | 30 |
| 5,12E+06 | 3,70E+06 | 3,49E+06 | 2,91E+06 | 145 | 0,002916 | 2,0326 | 7 |
| 8,42E+07 | 7,97E+07 | 7,89E+07 | 8,39E+07 | 21,539 | 0 | 24,932 | 19 |
| 4,26E+06 | 3,05E+06 | 7,07E+06 | 4,98E+06 | 100,08 | 0,000152 | 3,9935 | 9 |
| 4,11E+07 | 3,60E+07 | 3,69E+07 | 4,75E+07 | 16,28 | 0 | 69,205 | 27 |
| 8,69E+09 | 6,88E+09 | 8,36E+09 | 9,26E+09 | 35,81 | 0 | 323,31 | 1539 |
| 2,75E+06 | 1,93E+06 | 1,02E+07 | 1,38E+07 | 190,48 | 0,005419 | 1,6457 | 6 |
| 1,50E+09 | 1,47E+09 | 1,56E+09 | 1,67E+09 | 134,24 | 0 | 323,31 | 393 |
| 4,44E+06 | 4,08E+06 | 1,89E+06 | 3,70E+06 | 80,952 | 0,007147 | 1,5571 | 4 |
| 1,97E+07 | 1,52E+07 | 1,70E+07 | 1,61E+07 | 46,452 | 0 | 6,9048 | 19 |
| 1,04E+07 | 9,03E+06 | 9,36E+06 | 1,05E+07 | 48,157 | 0 | 6,1354 | 12 |
| 1,24E+07 | 1,30E+07 | 1,55E+07 | 1,37E+07 | 65,381 | 0 | 22,842 | 14 |
| 3,64E+08 | 3,10E+08 | 3,32E+08 | 3,28E+08 | 136,48 | 0 | 246,8 | 159 |
| 4,56E+07 | 4,36E+07 | 5,63E+07 | 6,14E+07 | 10,854 | 0 | 17,83 | 37 |
| 9,53E+06 | 1,07E+07 | 1,04E+07 | 1,42E+07 | 37,535 | 0 | 9,0678 | 18 |
| 4,18E+07 | 5,16E+07 | 4,85E+07 | 5,35E+07 | 19,086 | 0 | 6,5433 | 37 |
| 3,84E+08 | 3,75E+08 | 3,60E+08 | 3,52E+08 | 156,58 | 0 | 323,31 | 185 |
| 2,20E+07 | 2,83E+07 | 1,33E+07 | 1,64E+07 | 70,837 | 0 | 15,278 | 19 |
| 9,60E+07 | 8,93E+07 | 8,95E+07 | 1,04E+08 | 136,37 | 0 | 49,636 | 91 |
| 6,14E+08 | 5,52E+08 | 5,17E+08 | 5,08E+08 | 68,461 | 0 | 323,31 | 96 |
| 7,40E+07 | 7,65E+07 | 7,47E+07 | 7,81E+07 | 52,322 | 0 | 15,872 | 50 |
| 3,89E+07 | 2,11E+06 | 4,19E+07 | 4,88E+07 | 36,453 | 0,000292 | 3,0724 | 6 |
| 5,39E+07 | 5,76E+07 | 7,04E+07 | 5,96E+07 | 29,608 | 0 | 14,047 | 26 |
| 3,27E+06 | 8,92E+06 | 1,08E+06 | 8,01E+06 | 64,282 | 0,003311 | 1,9384 | 7 |
| 1,09E+09 | 9,69E+08 | 9,83E+08 | 8,30E+08 | 30,752 | 0 | 323,31 | 85 |
| 9,44E+07 | 7,44E+07 | 6,56E+07 | 7,35E+07 | 12,552 | 0 | 7,4214 | 12 |
| 1,49E+08 | 1,64E+08 | 1,67E+08 | 1,65E+08 | 10,194 | 0 | 74,052 | 27 |
| 2,03E+08 | 2,07E+08 | 2,06E+08 | 2,04E+08 | 63,908 | 0 | 135,75 | 101 |
| 2,36E+07 | 3,34E+07 | 2,97E+07 | 2,46E+07 | 62,062 | 0 | 11,255 | 25 |
| 3,58E+07 | 2,83E+07 | 4,18E+07 | 4,04E+07 | 49,804 | 0 | 7,1352 | 24 |
| 1,35E+08 | 1,48E+08 | 1,77E+08 | 1,65E+08 | 20,987 | 0 | 253,41 | 38 |
| 3,73E+06 | 6,91E+06 | 3,88E+06 | 4,33E+06 | 71,037 | 0,009373 | 1,439 | 8 |
| 3,44E+07 | 3,66E+07 | 2,09E+07 | 3,44E+07 | 20,555 | 0 | 247,65 | 21 |
| 1,11E+07 | 1,45E+07 | 1,43E+07 | 1,34E+07 | 32,823 | 0 | 8,9284 | 29 |
| 2,07E+08 | 2,21E+08 | 1,98E+08 | 2,43E+08 | 22,435 | 0 | 48,027 | 63 |
| 1,42E+08 | 1,46E+08 | 1,46E+08 | 1,54E+08 | 53,844 | 0 | 145,76 | 52 |
| 5,03E+08 | 5,82E+08 | 5,44E+08 | 5,81E+08 | 23,474 | 0 | 114,31 | 71 |
| 9,82E+07 | 7,56E+07 | 1,10E+08 | 9,04E+07 | 16,958 | 0 | 17,421 | 17 |
| 4,30E+07 | 1,05E+08 | 7,69E+06 | 8,17E+07 | 35,988 | 0,000148 | 3,3237 | 10 |
| 2,50E+06 | 7,39E+06 | 7,46E+06 | 2,38E+06 | 59,145 | 0,000157 | 4,9777 | 7 |
| 2,07E+08 | 2,22E+08 | 2,16E+08 | 1,90E+08 | 32,631 | 0 | 168,59 | 80 |
| 5,08E+06 | 4,75E+06 | 5,04E+06 | 5,18E+06 | 61,344 | 0,002098 | 2,1657 | 9 |
| 1,72E+08 | 1,78E+08 | 1,79E+08 | 1,59E+08 | 55,598 | 0 | 9,9047 | 34 |
| 5,72E+07 | 5,81E+07 | 6,06E+07 | 5,53E+07 | 81,598 | 0 | 18,164 | 48 |
| 1,63E+08 | 2,00E+08 | 2,09E+08 | 2,21E+08 | 9,7251 | 0 | 191,71 | 147 |
| 5,07E+07 | 6,77E+07 | 7,43E+07 | 7,73E+07 | 21,175 | 0,000146 | 3,1336 | 14 |
| 2,56E+06 | 3,52E+06 | 4,64E+06 | 7,40E+06 | 111,72 | 0 | 5,3948 | 11 |

| | | | | | | | |
|---|---|---|---|---|---|---|---|
| 3,83E+08 | 1,25E+08 | 1,18E+08 | 1,67E+08 | 51,813 | 0 | 67,517 | 31 |
| 7,17E+06 | 5,93E+06 | 5,27E+06 | 6,47E+06 | 49,266 | 0,000152 | 4,0329 | 9 |
| 1,41E+08 | 1,24E+08 | 1,27E+08 | 1,22E+08 | 216,82 | 0 | 57,315 | 133 |
| 1,86E+07 | 1,28E+07 | 1,38E+07 | 1,05E+07 | 58,917 | 0 | 37,472 | 14 |
| 5,71E+06 | 5,16E+06 | 3,75E+06 | 3,11E+06 | 24,787 | 0,000153 | 4,2085 | 4 |
| 1,32E+08 | 1,16E+08 | 1,05E+08 | 1,11E+08 | 48,051 | 0 | 60,495 | 62 |
| 1,33E+08 | 1,62E+08 | 1,02E+08 | 1,10E+08 | 29,758 | 0 | 147,02 | 48 |
| 7,31E+06 | 3,05E+06 | 3,87E+06 | 4,40E+06 | 19,06 | 0 | 6,8862 | 4 |
| 1,68E+07 | 1,92E+07 | 2,43E+07 | 3,74E+06 | 8,1384 | 0,003695 | 1,8109 | 9 |
| 1,44E+07 | 1,23E+07 | 1,27E+07 | 1,29E+07 | 12,295 | 0,000153 | 4,1165 | 11 |
| 3,80E+06 | 3,88E+06 | 2,74E+06 | 4,45E+06 | 71,064 | 0,000855 | 2,6015 | 6 |
| 4,80E+07 | 4,33E+07 | 4,30E+07 | 4,20E+07 | 51,225 | 0 | 13,167 | 22 |
| 3,10E+08 | 3,72E+08 | 3,54E+08 | 3,68E+08 | 54,734 | 0 | 323,31 | 109 |
| 2,73E+07 | 2,44E+07 | 2,29E+07 | 2,85E+07 | 14,188 | 0 | 12,5 | 23 |
| 2,52E+06 | 4,67E+06 | 1,73E+06 | 4,82E+06 | 63,662 | 0,000432 | 2,8175 | 2 |
| 1,14E+06 | 1,14E+06 | 1,76E+06 | 1,51E+06 | 140,94 | 0,009367 | 1,4354 | 7 |
| 8,20E+06 | 3,63E+06 | 7,40E+06 | 7,03E+06 | 119,16 | 0,000152 | 4,0124 | 11 |
| 6,04E+07 | 5,29E+07 | 4,80E+07 | 6,61E+07 | 27,815 | 0 | 151,16 | 37 |
| 4,71E+08 | 4,40E+08 | 4,52E+08 | 4,06E+08 | 30,224 | 0 | 323,31 | 83 |
| 1,79E+07 | 1,79E+07 | 1,97E+07 | 2,22E+07 | 25,335 | 0 | 6,5819 | 15 |
| 8,70E+07 | 6,24E+07 | 8,52E+07 | 6,36E+07 | 13,37 | 0 | 31,968 | 35 |
| 3,56E+08 | 3,49E+08 | 3,65E+08 | 3,83E+08 | 82,308 | 0 | 173,17 | 148 |
| 5,03E+06 | 2,84E+07 | 1,33E+07 | 1,19E+07 | 18,974 | 0,00015 | 3,7531 | 20 |
| 1,01E+08 | 1,04E+08 | 9,47E+07 | 1,00E+08 | 29,463 | 0 | 19,569 | 59 |
| 2,94E+08 | 4,06E+08 | 3,58E+08 | 3,46E+08 | 33,901 | 0 | 323,31 | 82 |
| 7,63E+06 | 7,43E+06 | 1,15E+06 | 7,29E+06 | 79,513 | 0,003965 | 1,7851 | 2 |
| 1,32E+08 | 1,64E+08 | 1,13E+08 | 1,40E+08 | 6,6979 | 0,000148 | 3,3329 | 34 |
| 3,73E+06 | 2,45E+07 | 2,53E+07 | 3,91E+06 | 40,849 | 0,009744 | 1,4066 | 6 |
| 2,32E+08 | 2,39E+08 | 2,42E+08 | 2,21E+08 | 68,09 | 0 | 49,602 | 103 |
| 4,18E+06 | 4,72E+06 | 4,62E+06 | 6,01E+06 | 59,71 | 0,000152 | 3,9693 | 9 |
| 2,15E+07 | 2,14E+07 | 2,04E+07 | 2,26E+07 | 13,658 | 0 | 200,14 | 39 |
| 6,34E+07 | 4,60E+07 | 5,46E+07 | 4,52E+07 | 21,942 | 0 | 32,195 | 42 |
| 1,96E+08 | 2,26E+08 | 2,04E+08 | 2,00E+08 | 33,073 | 0 | 19,659 | 61 |
| 2,59E+09 | 3,00E+09 | 2,63E+09 | 2,63E+09 | 18,431 | 0 | 322,61 | 204 |
| 3,09E+09 | 3,41E+09 | 3,73E+09 | 3,90E+09 | 17,804 | 0 | 323,31 | 99 |
| 1,53E+07 | 2,11E+07 | 1,40E+07 | 1,76E+07 | 35,869 | 0 | 10,244 | 17 |
| 4,10E+06 | 3,96E+06 | 3,93E+06 | 4,52E+06 | 19,664 | 0 | 9,3495 | 3 |
| 9,07E+06 | 6,17E+06 | 5,92E+06 | 9,15E+06 | 21,501 | 0,003571 | 1,8844 | 6 |
| 6,42E+08 | 6,65E+08 | 5,39E+08 | 4,79E+08 | 78,168 | 0 | 323,31 | 155 |
| 2,10E+07 | 2,04E+07 | 1,82E+07 | 2,54E+07 | 22,652 | 0 | 6,2398 | 14 |
| 1,60E+07 | 1,89E+07 | 1,86E+07 | 2,00E+07 | 92,403 | 0 | 9,4388 | 23 |
| 3,21E+06 | 5,40E+06 | 2,09E+06 | 4,64E+06 | 122,33 | 0,008461 | 1,482 | 3 |
| 3,98E+07 | 4,83E+07 | 4,99E+07 | 4,98E+07 | 149,54 | 0 | 62,767 | 58 |
| 2,95E+08 | 2,41E+08 | 2,97E+08 | 2,95E+08 | 39,296 | 0 | 323,31 | 85 |
| 1,49E+07 | 1,40E+07 | 1,56E+07 | 1,83E+07 | 26,81 | 0,000151 | 3,8281 | 10 |
| 1,70E+08 | 1,80E+08 | 1,80E+08 | 1,84E+08 | 34,196 | 0 | 62,172 | 83 |
| 1,82E+06 | 8,34E+05 | 8,68E+06 | 1,11E+06 | 118,72 | 0,000432 | 2,8158 | 4 |
| 2,69E+08 | 2,85E+08 | 2,89E+08 | 2,72E+08 | 150,55 | 0 | 269,15 | 183 |
| 5,39E+07 | 4,86E+07 | 4,99E+07 | 5,26E+07 | 80,169 | 0 | 56,996 | 42 |
| 1,14E+08 | 1,60E+08 | 1,24E+08 | 1,52E+08 | 35,41 | 0 | 54,55 | 82 |

| | | | | | | | |
|---|---|---|---|---|---|---|---|
| 5,82E+08 | 5,22E+08 | 5,83E+08 | 5,22E+08 | 65,629 | 0 | 323,31 | 140 |
| 8,23E+06 | 4,39E+06 | 3,18E+06 | 1,00E+07 | 60,063 | 0,000153 | 4,1237 | 6 |
| 1,47E+08 | 1,39E+08 | 1,22E+08 | 1,33E+08 | 160,91 | 0 | 154,47 | 119 |
| 1,41E+08 | 1,39E+08 | 1,39E+08 | 1,43E+08 | 38,926 | 0 | 86,389 | 66 |
| 1,56E+08 | 1,80E+08 | 1,53E+08 | 1,02E+08 | 52,429 | 0 | 27,798 | 54 |
| 4,04E+07 | 4,93E+07 | 3,42E+07 | 4,24E+07 | 86,599 | 0 | 21,233 | 47 |
| 8,04E+07 | 6,95E+07 | 9,41E+07 | 1,09E+08 | 19,859 | 0 | 15,867 | 34 |
| 4,45E+07 | 4,42E+07 | 5,68E+07 | 5,49E+07 | 15,709 | 0 | 29,506 | 27 |
| 2,59E+07 | 2,00E+07 | 1,42E+07 | 1,38E+07 | 23,299 | 0 | 19,981 | 14 |
| 1,10E+07 | 1,38E+07 | 1,12E+07 | 1,36E+07 | 36,94 | 0 | 7,391 | 13 |
| 5,10E+07 | 1,09E+08 | 5,15E+07 | 1,00E+08 | 67,955 | 0 | 6,8604 | 17 |
| 7,03E+06 | 2,74E+06 | 1,26E+07 | 1,65E+07 | 17,872 | 0,000152 | 3,9556 | 5 |
| 2,47E+09 | 2,53E+09 | 2,53E+09 | 2,42E+09 | 91,369 | 0 | 323,31 | 505 |
| 1,38E+08 | 1,25E+08 | 1,25E+08 | 1,03E+08 | 70,626 | 0 | 67,051 | 63 |
| 4,06E+06 | 1,82E+06 | 6,11E+06 | 3,66E+06 | 43,371 | 0,001275 | 2,4687 | 5 |
| 2,23E+08 | 2,29E+08 | 2,10E+08 | 2,20E+08 | 114,31 | 0 | 77,838 | 101 |
| 2,62E+06 | 9,98E+06 | 1,84E+06 | 6,13E+06 | 63,218 | 0,002369 | 2,0897 | 4 |
| 8,11E+07 | 8,97E+07 | 8,82E+07 | 9,92E+07 | 86,108 | 0 | 29,85 | 57 |
| 9,29E+06 | 1,03E+07 | 1,01E+07 | 1,00E+07 | 106,32 | 0 | 23,846 | 11 |
| 4,65E+07 | 4,07E+07 | 3,38E+07 | 4,27E+07 | 54,444 | 0 | 31,62 | 26 |
| 2,69E+07 | 2,20E+07 | 2,60E+07 | 3,68E+07 | 18,066 | 0 | 9,7152 | 18 |
| 1,78E+06 | 1,44E+06 | 3,25E+06 | 2,16E+06 | 102,25 | 0,00898 | 1,4578 | 4 |
| 2,63E+08 | 2,89E+08 | 2,87E+08 | 2,82E+08 | 48,542 | 0 | 69,261 | 150 |
| 1,37E+08 | 1,19E+08 | 1,43E+08 | 1,39E+08 | 133,68 | 0 | 72,183 | 122 |
| 8,12E+07 | 7,82E+07 | 8,28E+07 | 9,39E+07 | 91,859 | 0 | 67,948 | 51 |
| 8,47E+06 | 7,53E+06 | 6,61E+06 | 7,75E+06 | 62,808 | 0,002913 | 2,0166 | 6 |
| 1,02E+08 | 1,27E+08 | 1,34E+08 | 1,21E+08 | 19,481 | 0 | 6,1418 | 35 |
| 6,21E+06 | 3,71E+06 | 4,51E+06 | 2,65E+06 | 22,715 | 0,001824 | 2,2556 | 5 |
| 1,59E+09 | 1,64E+09 | 1,76E+09 | 1,89E+09 | 53,516 | 0 | 323,31 | 296 |
| 1,48E+08 | 1,58E+08 | 1,62E+08 | 1,83E+08 | 39,807 | 0 | 96,838 | 51 |
| 2,59E+08 | 3,31E+08 | 2,15E+08 | 2,26E+08 | 29,486 | 0 | 255,37 | 112 |
| 1,66E+07 | 5,31E+06 | 9,38E+06 | 1,21E+07 | 195,81 | 0 | 6,8788 | 16 |
| 1,11E+09 | 7,57E+08 | 1,03E+09 | 1,26E+09 | 15,807 | 0 | 70,768 | 87 |
| 6,90E+06 | 7,92E+06 | 7,01E+06 | 1,37E+07 | 152,88 | 0 | 7,2419 | 11 |
| 5,45E+08 | 5,08E+08 | 4,60E+08 | 4,67E+08 | 56,475 | 0 | 323,31 | 217 |
| 5,76E+09 | 6,84E+09 | 6,41E+09 | 6,97E+09 | 22,591 | 0 | 85,611 | 305 |
| 1,51E+07 | 1,58E+07 | 1,11E+07 | 1,21E+07 | 63,905 | 0 | 7,9185 | 14 |
| 1,48E+07 | 1,46E+07 | 9,51E+06 | 1,34E+07 | 35,321 | 0 | 8,2328 | 12 |
| 2,90E+06 | 5,04E+06 | 4,08E+06 | 5,03E+06 | 15,207 | 0,003698 | 1,8243 | 6 |
| 1,64E+07 | 1,30E+07 | 1,33E+07 | 1,46E+07 | 30,427 | 0 | 11,866 | 15 |
| 6,73E+07 | 7,73E+07 | 7,60E+07 | 8,16E+07 | 96,329 | 0 | 147,18 | 63 |
| 8,08E+08 | 8,20E+08 | 1,04E+09 | 1,02E+09 | 28,405 | 0 | 323,31 | 157 |
| 1,46E+08 | 1,50E+08 | 1,46E+08 | 1,40E+08 | 32,131 | 0 | 57,017 | 65 |
| 6,92E+07 | 7,80E+07 | 4,59E+07 | 5,25E+07 | 28,96 | 0 | 72,374 | 41 |
| 1,33E+07 | 9,77E+06 | 8,94E+06 | 8,07E+06 | 63,457 | 0 | 15,349 | 17 |
| 1,71E+08 | 1,67E+08 | 1,56E+08 | 1,70E+08 | 60,988 | 0 | 91,605 | 64 |
| 1,12E+07 | 3,06E+06 | 1,56E+07 | 1,82E+07 | 21,709 | 0,003181 | 1,971 | 6 |
| 4,82E+09 | 4,08E+09 | 4,23E+09 | 4,63E+09 | 16,273 | 0 | 323,31 | 265 |
| 6,55E+06 | 4,38E+06 | 4,08E+06 | 7,81E+06 | 246,32 | 0,00043 | 2,749 | 11 |
| 1,45E+09 | 1,24E+09 | 1,33E+09 | 1,28E+09 | 17,695 | 0 | 56,629 | 154 |

| | | | | | | | |
|---|---|---|---|---|---|---|---|
| 2,85E+07 | 3,46E+07 | 3,08E+07 | 2,95E+07 | 84,18 | 0 | 19,867 | 49 |
| 8,24E+08 | 9,76E+08 | 9,28E+08 | 7,72E+08 | 26,94 | 0 | 323,31 | 98 |
| 1,01E+07 | 7,21E+06 | 9,32E+06 | 8,33E+06 | 68,505 | 0 | 12,339 | 17 |
| 9,11E+07 | 1,30E+08 | 1,09E+08 | 1,36E+08 | 19,965 | 0,000148 | 3,4034 | 25 |
| 3,54E+08 | 1,66E+08 | 1,87E+08 | 1,86E+08 | 21,782 | 0 | 323,31 | 86 |
| 9,32E+07 | 9,21E+07 | 9,45E+07 | 8,18E+07 | 88,975 | 0 | 46,824 | 62 |
| 2,02E+07 | 1,77E+07 | 1,85E+07 | 2,24E+07 | 67,026 | 0 | 6,719 | 21 |
| 4,46E+07 | 4,27E+07 | 3,76E+07 | 4,30E+07 | 60,705 | 0 | 48,523 | 36 |
| 1,05E+09 | 9,31E+08 | 9,22E+08 | 9,64E+08 | 40,703 | 0 | 323,31 | 128 |
| 7,99E+07 | 7,07E+07 | 6,42E+07 | 4,60E+07 | 20,594 | 0 | 17,425 | 16 |
| 3,34E+07 | 1,18E+07 | 1,29E+07 | 1,51E+07 | 58,512 | 0 | 16,942 | 18 |
| 4,45E+07 | 4,83E+07 | 4,86E+07 | 5,25E+07 | 98,144 | 0 | 29,537 | 46 |
| 5,32E+07 | 5,69E+07 | 4,15E+07 | 4,90E+07 | 71,601 | 0 | 29,977 | 51 |
| 4,63E+06 | 3,77E+06 | 5,65E+06 | 5,22E+06 | 35,207 | 0,000158 | 5,151 | 5 |
| 2,27E+08 | 2,41E+08 | 2,32E+08 | 2,46E+08 | 73,189 | 0 | 177,72 | 165 |
| 8,58E+06 | 4,60E+06 | 5,87E+06 | 2,96E+06 | 58,379 | 0,000155 | 4,5832 | 9 |
| 7,76E+06 | 8,99E+06 | 9,12E+06 | 7,19E+06 | 236,87 | 0,000157 | 4,9332 | 17 |
| 6,80E+08 | 6,94E+08 | 7,17E+08 | 7,62E+08 | 111,6 | 0 | 257,81 | 202 |
| 3,40E+07 | 2,68E+07 | 3,29E+07 | 2,91E+07 | 76,278 | 0 | 14,322 | 43 |
| 2,07E+07 | 4,25E+06 | 3,19E+06 | 4,25E+06 | 118,92 | 0,007678 | 1,5302 | 4 |
| 7,11E+07 | 9,63E+07 | 8,71E+07 | 1,13E+08 | 15,283 | 0 | 11,467 | 28 |
| 1,33E+07 | 1,41E+07 | 1,78E+07 | 1,32E+07 | 79,957 | 0 | 6,2773 | 21 |
| 5,98E+06 | 7,99E+06 | 9,07E+06 | 9,20E+06 | 37,157 | 0,000154 | 4,3351 | 14 |
| 1,94E+07 | 1,53E+07 | 2,44E+07 | 2,61E+07 | 54,163 | 0 | 15,502 | 28 |
| 4,82E+06 | 5,32E+06 | 5,46E+06 | 7,95E+06 | 111,54 | 0 | 5,3605 | 6 |
| 4,93E+08 | 4,85E+08 | 5,02E+08 | 4,64E+08 | 24,74 | 0 | 323,31 | 179 |
| 4,50E+06 | 7,63E+06 | 8,86E+06 | 9,30E+06 | 60,717 | 0,000434 | 2,8919 | 9 |
| 4,61E+06 | 1,30E+07 | 2,46E+07 | 2,08E+07 | 248,98 | 0 | 7,8748 | 13 |
| 1,93E+07 | 1,75E+07 | 1,70E+07 | 1,45E+07 | 62,491 | 0 | 39,162 | 17 |
| 2,19E+09 | 1,90E+09 | 1,98E+09 | 2,00E+09 | 44,714 | 0 | 323,31 | 364 |
| 2,32E+07 | 2,01E+07 | 2,58E+07 | 3,23E+07 | 45,824 | 0 | 8,6118 | 21 |
| 5,02E+07 | 5,09E+07 | 4,50E+07 | 3,96E+07 | 23,715 | 0 | 14,297 | 27 |
| 6,87E+08 | 7,09E+08 | 7,52E+08 | 7,40E+08 | 111,97 | 0 | 323,31 | 290 |
| 6,16E+07 | 6,13E+07 | 7,32E+07 | 7,13E+07 | 70,545 | 0 | 16,606 | 50 |
| 6,77E+06 | 4,90E+06 | 1,81E+07 | 5,55E+06 | 31,599 | 0 | 10,554 | 12 |
| 1,07E+08 | 1,01E+08 | 1,07E+08 | 1,10E+08 | 69,07 | 0 | 55,247 | 74 |
| 2,13E+07 | 2,12E+07 | 1,66E+07 | 2,24E+07 | 51,177 | 0 | 13,232 | 25 |
| 4,48E+09 | 3,83E+09 | 4,07E+09 | 4,58E+09 | 28,68 | 0 | 120,72 | 288 |
| 4,45E+07 | 5,55E+07 | 6,29E+07 | 5,29E+07 | 43,813 | 0 | 39,555 | 47 |
| 9,60E+06 | 1,60E+07 | 1,03E+07 | 1,51E+07 | 33,048 | 0 | 10,253 | 32 |
| 1,29E+08 | 1,42E+08 | 1,47E+08 | 1,55E+08 | 11,871 | 0 | 93,565 | 48 |
| 2,90E+08 | 3,09E+08 | 2,94E+08 | 2,88E+08 | 83,415 | 0 | 94,088 | 178 |
| 2,07E+07 | 1,80E+07 | 2,09E+07 | 2,53E+07 | 57,534 | 0 | 6,8064 | 24 |
| 5,03E+06 | 5,59E+06 | 4,29E+06 | 6,01E+06 | 20,026 | 0,001548 | 2,3079 | 6 |
| 3,96E+06 | 3,00E+06 | 2,81E+06 | 2,51E+06 | 40,748 | 0,000146 | 3,1334 | 9 |
| 2,46E+08 | 2,09E+08 | 2,13E+08 | 2,20E+08 | 65,225 | 0 | 128,23 | 109 |
| 9,95E+08 | 1,03E+09 | 1,02E+09 | 8,38E+08 | 22,965 | 0 | 157,29 | 184 |
| 3,10E+09 | 2,70E+09 | 2,98E+09 | 3,24E+09 | 23,464 | 0 | 169,11 | 254 |
| 3,00E+06 | 2,58E+06 | 3,66E+06 | 4,11E+06 | 37,485 | 0,000855 | 2,6069 | 6 |
| 8,05E+07 | 8,14E+07 | 9,05E+07 | 8,06E+07 | 111,17 | 0 | 51,938 | 95 |

| | | | | | | | |
|---|---|---|---|---|---|---|---|
| 4,65E+06 | 4,75E+06 | 9,99E+06 | 9,99E+06 | 56,701 | 0,008978 | 1,4541 | 6 |
| 3,15E+08 | 3,93E+08 | 3,98E+08 | 3,73E+08 | 14,585 | 0 | 65,433 | 84 |
| 5,81E+07 | 5,16E+07 | 4,98E+07 | 4,41E+07 | 86,769 | 0 | 28,963 | 55 |
| 3,72E+06 | 5,71E+06 | 2,17E+06 | 2,32E+06 | 45,281 | 0 | 6,5404 | 9 |
| 1,49E+08 | 1,36E+08 | 1,40E+08 | 1,41E+08 | 82,367 | 0 | 72,666 | 106 |
| 6,71E+06 | 1,10E+07 | 9,59E+06 | 1,08E+07 | 53,497 | 0,008458 | 1,4747 | 7 |
| 6,18E+07 | 8,31E+07 | 4,97E+07 | 6,36E+07 | 79,959 | 0 | 107,25 | 62 |
| 4,40E+06 | 3,48E+06 | 3,47E+06 | 2,80E+06 | 24,103 | 0,00237 | 2,0969 | 4 |
| 3,16E+07 | 3,60E+07 | 3,16E+07 | 3,37E+07 | 45,97 | 0 | 239,44 | 28 |
| 5,23E+07 | 5,25E+07 | 6,27E+07 | 5,07E+07 | 18,418 | 0,000148 | 3,3757 | 22 |
| 4,21E+06 | 2,60E+06 | 4,00E+06 | 1,75E+06 | 25,652 | 0,000152 | 4,0749 | 3 |
| 1,04E+06 | 1,95E+06 | 8,10E+06 | 2,26E+06 | 19,977 | 0,000992 | 2,489 | 3 |
| 1,17E+08 | 1,82E+08 | 3,42E+07 | 1,78E+08 | 17,04 | 0 | 75,182 | 71 |
| 5,12E+07 | 5,41E+07 | 5,80E+07 | 6,42E+07 | 69,399 | 0 | 58,522 | 42 |
